# Combinatorial approach with mass spectrometry and lectin microarray dissected glycoproteomic features of virion-derived spike protein of SARS-CoV-2

**DOI:** 10.1101/2021.04.10.439300

**Authors:** Takahiro Hiono, Azusa Tomioka, Hiroyuki Kaji, Michihito Sasaki, Yasuko Orba, Hirofumi Sawa, Atsushi Kuno

## Abstract

The COVID-19 pandemic caused by the novel coronavirus, SARS-CoV-2, has a global impact on public health. Since glycosylation of the viral envelope glycoproteins is known to be deeply associated with their immunogenicity, intensive studies on the glycans of its major glycoprotein, S protein, have been conducted. Nevertheless, the detailed site-specific glycan compositions of virion-associated S protein have not yet been clarified. Here, we conducted intensive glycoproteomic analyses of SARS-CoV-2 S protein using a combinatorial approach with two different technologies: mass spectrometry (MS) and lectin microarray. Using our unique MS1-based glycoproteomic technique, Glyco-RIDGE, in addition to MS2-based Byonic search, we identified 1,759 site-specific glycan compositions. The most frequent was HexNAc:Hex:Fuc:NeuAc:NeuGc = 6:6:1:0:0, suggesting a tri-antennary *N*-glycan terminating with LacNAc and having bisecting GlcNAc and a core fucose, which was found in 20 of 22 glycosylated sites. The subsequent lectin microarray analysis emphasized intensive outer arm fucosylation of glycans, which efficiently complemented the glycoproteomic features. The present results illustrate the high-resolution glycoproteomic features of SARS-CoV-2 S protein and significantly contribute to vaccine design, as well as the understanding of viral protein synthesis.

## Introduction

The COVID-19 pandemic caused by the novel coronavirus, SARS-CoV-2, has a global impact on public health^1^. As a countermeasure against the COVID-19 pandemic, great efforts have been made to establish vaccines targeting the spike (S) protein of SARS-CoV-2^2^. The coronavirus S protein is a glycoprotein required for virus attachment to host cells and is mainly targeted by the host immune system. Since glycosylation of the viral envelope glycoproteins is known to be strongly associated with their immunogenicity^3^, intensive studies on the glycans of SARS-CoV-2 S protein have been conducted^4^. Watanabe et al. first demonstrated a site-specific glycoform for all 22 *N*-glycosylation sites of the S protein^5^. Moreover, Shajahan et al. identified two *O*-glycosylation sites in the receptor-binding domain of the S protein^6^. Based on the revealed glycoform, the molecular dynamics of the glycosylated S proteins were simulated *in silico*^7^. The results demonstrated that two *N*-glycans at N165 and N234 modulated the conformational dynamics of the receptor-binding domain of S proteins. These results were demonstrated with recombinant S protein using a mammalian cell-protein expression system. Subsequently, Yao et al. conducted site-specific glycomic analyses using the S protein derived from intact virions, although detailed glycan structures were not provided^8^.

To date, analyses of the glycoforms of viral proteins have been mainly conducted using mass spectrometry (MS). MS-based approaches have been previously employed for glycomic and glycoproteomic analyses of viruses in many different taxonomic groups, including human immunodeficiency virus^9,10^, influenza virus^11–13^, and SARS coronavirus^14^. MS-based physical analyses are beneficial in that they provide detailed site-specific glycan compositions. We previously established a liquid chromatography (LC)/MS-based glycoproteomic approach^15,16^. In general, analyses of site-specific glycoforms of glycopeptides require tandem MS-based approaches. Tandem MS analyses of glycopeptides often showed low sensitivity because of the difficulty of obtaining fragment ions to identify peptide sequences, in addition to the poor ionization efficiency of glycopeptides. On the other hand, our approach enables MS1-based site-specific glycoproteomic analyses using accurate masses and LC-retention times of glycopeptides^15,17^ This method, called Glyco-RIDGE, is MS2-independent and therefore has an advantage in sensitivity. Using this unique approach, we successfully identified lists of core peptides for poly-*N*-acetyllactosamine using human promyelocytic leukemia HL60 cells^17^ and those for Lewis x antigens using mouse kidneys^15^.

Lectin microarray (LMA)-based glycan profiling is an alternative biochemical method for analyzing glycan attached to viruses without the liberation of the glycan moiety^18^. Our previous studies on glycan profiling of surface glycoproteins of hepatitis B and influenza A viruses demonstrated that this method provides highly sensitive and simple platforms for the structural interpretation of viral glycans^19,20^. As an interaction-based assay, the LMA provides information not only on the molecular structures of glycans but also on the accessibility of endogenous glycan-binding molecules to them^21^. However, this method does not provide detailed glycan composition or site-specific information on glycans, that is, glycan micro-heterogeneity in a glycoprotein. Accordingly, the combinatorial use of MS-and LMA-based approaches in the glycoproteomic analysis of the viral glycoprotein should compensate for each method’s limitations to synergistically dissect the detailed glycoforms of viral glycoproteins. In this study, we analyzed the glycoform of S proteins derived from intact SARS-CoV-2 viruses by combining MS1-based glycoproteomic analysis and LMA. These two approaches cooperatively illustrated the composition, structure, and conformation of the glycan at high resolution, which will enhance our in-depth exploration of the real world, i.e., “meta-heterogeneity,” of SARS-CoV-2 glyco-particles^22^.

## Materials and methods

### Viruses and cells

The SARS-CoV-2, 2019-nCoV/Japan/TY/WK-521/2020 strain was provided by the National Institute of Infectious Diseases. VeroE6/TMPRSS2 cells were established in our laboratory as described previously^23^. The cells were maintained in Dulbecco’s modified Eagle’s medium supplemented with 10% fetal bovine serum. The virus was inoculated onto the cells, and the infectious cell culture supernatant was collected and cleared by low-speed centrifugation. For the preparation of the sample subjected to MS analyses, virus particles in the supernatant were concentrated through a 20% sucrose cushion at 133,900× g for 2 h, washed with PBS, and resuspended^14^. For the samples subjected to LMA analyses, viruses were inactivated with 0.1% β-propiolactone at 4 °C for 1 d.

### Antibodies

Rabbit monoclonal anti-SARS-CoV-2 spike S1 antibody, 007 (Cat# 40150-R007), and mouse monoclonal anti-SARS-CoV-2 spike neutralizing antibody, 43 (Cat# 40591-V05H1), were purchased from Sino Biological (Beijing, China). These antibodies were biotinylated using the Biotin Labeling Kit-NH2 (Dojindo, Kumamoto, Japan).

### Site-specific glycan analysis by LC-MS

SARS-CoV-2 particles collected from the culture medium of Vero/TMPRSS2 cells by ultracentrifugation were precipitated with acetone (final concentration of 75%) at -80 °C overnight. The precipitate was recovered and dissolved in 100 mM Tris-HCl (pH9.0) containing 12 mM sodium deoxycholate and 12 mM sodium lauroylsarcosinate. Proteins were S-reduced with dithiothreitol and alkylated with iodoacetamide. After 5 x dilution, the proteins were digested with chymotrypsin, trypsin + Lys-C endopeptidase, or α-lytic protease. An aliquot of each digest was analyzed using an LC/MS system with a nanoflow LC, Ultimate 3000 (Thermo Fisher Scientific, Waltham, MA, USA), and Orbitrap Fusion Tribrid mass spectrometer (Thermo Fisher Scientific). Another aliquot of each digest was subjected to hydrophilic interaction chromatography (HILIC) on an Amide-80 column (TOSOH, Tokyo, Japan) to collect glycopeptides. An aliquot of the glycopeptide fraction was treated with peptide-*N*-glycanase F (PNGaseF, Takara, Shiga, Japan) in H_2_^18^O to remove *N*-glycans and to label glycosylated Asn with ^18^O as Asp (^18^O). The labeled peptides were identified by LC/MS analysis followed by a database search using Mascot (Matrix Science, London, UK)^24,25^. Another aliquot of glycopeptides was analyzed using LC/MS and the acquired MS2 spectra were analyzed by database search using Byonic. Simultaneously, the MS1 spectra were analyzed using the Glyco-RIDGE method as described previously^15,17^.

### Three-dimensional structural-based mapping of S protein

*N*-glycosylation sites were mapped on the S protein in a closed conformation (PDB ID: 6ZGE)^26^ using Discovery Studio Visualizer (BIOVIA, San Diego, CA, USA). Each monomer was depicted as a soft surface model and colored in light gray, light pink, or light blue.

### Immunoprecipitation

One hundred microliters of β-propiolactone-inactivated infectious cell culture supernatants were mixed with 200 ng biotinylated 007 antibody. After incubation at 37 °C for 1 h, the complexes were captured with Dynabeads MyOne Streptavidin T1 (Thermo Fisher Scientific) at 4 °C for 1 h. After washing, virus antigens were eluted from the magnetic beads and disrupted using 20 μL of 100 mM citric acid containing 1% Triton X-100. Biotinylated antibodies incorporated into the eluted samples were depleted with Dynabeads MyOne Streptavidin T1 at 4 °C for 1 h. Antibody-depleted samples were then pH adjusted with 5 μL of Tris-HCl buffer (pH 9.0). For two-step immunoprecipitation, antibody-depleted samples were neutralized with 5 μL of HEPES-NaOH buffer (pH 8.5). The 25 μL sample was then fluorescently labeled with 10 μg Cy3-succinimidyl-ester (GE Healthcare, Buckinghamshire, UK) for 1 h at room temperature in the dark. After adjusting the volume to 125 μL with probing buffer (25 mM Tris-HCl (pH 7.5), containing 137 mM NaCl, 2.7 mM KCl, 500 mM glycine, 1 mM CaCl_2_, 1 mM MgCl_2_, and 1% Triton X-100), the sample solutions were incubated for 2 h at room temperature to quench the excess fluorescent label. The sample solutions were mixed with 200 ng biotinylated 43 antibody and incubated at 4 °C overnight. The complexes were captured with Dynabeads MyOne Streptavidin T1 at 4 °C for 1 h. After washing, virus antigens were eluted from the magnetic beads and disrupted using 20 μL of 100 mM citric acid containing 1% Triton X-100. Biotinylated antibodies incorporated into the eluted samples were depleted with Dynabeads MyOne Streptavidin T1 at 4 °C for 1 h. Antibody-depleted samples were then pH adjusted with 5 μL of Tris-HCl buffer (pH 9.0). PBS was used as the negative control instead of the virus solution.

### LMA

An antibody-overlay LMA was performed as described previously, with some modifications^20,27^. Lectins printed on the array chips are listed in Supplementary Table S1. The glass slides were printed with 45 lectins with three replicates (LecChip Ver. 1.0; GlycoTechnica, Yokohama, Japan) were activated with probing buffer. Immunoprecipitated samples diluted with probing buffer were then applied to glass slides and incubated overnight at 20 °C. Subsequently, the slides were blocked with 0.3 mg/mL human IgG for 30 min at 20 °C. After removal of unbound samples and IgG by subsequent washing, 6.7 μg/mL biotinylated monoclonal antibody 43 was added to the slides and incubated for 1 h at 20 °C. Slides were washed three times and incubated with 1.7 μg/mL Streptavidin, Alexa Fluor 555 conjugate (Thermo Fisher Scientific) for 30 min at 20 °C. After washing, the slides were scanned with an evanescent-field excitation fluorescence imager (GlycoStation Reader 1200; GlycoTechnica). The obtained images were quantified using the GlycoStation™ToolsPro Suite ver. 1.5 (GlycoTechnica). The net intensity for each lectin was calculated as the mean value of three spots minus the background. For direct labeling methods, sample solutions were added to the array glass slide and incubated overnight at °C. After washing three times, the glass slides were scanned using an evanescent-field excitation fluorescence imager.

### Exoglycosidase digestion

Fluorescently labeled S protein was prepared as described above using two-step immunoprecipitation. The labeled proteins were subjected to digestion with Sialidase A (Agilent Technologies, Santa Clara, CA, USA), β1-4 galactosidase (Agilent Technologies), or double digestion with Sialidase A and β-galactosidase, according to the manufacturer’s instructions. Digestion was conducted at 37 °C for 12 h.

## Results

### Site-specific glycan composition analysis of SARS-CoV-2 S protein using the LC/MS-Glyco-RIDGE method

Protein digests obtained using chymotrypsin, trypsin+ Lys-endopeptidase, or α-Lytic protease were analyzed by LC/MS, and the data were searched using Mascot to identify the spike protein and confirm peptides with the *N*-glycosylation potential consensus sequence, N-! P-(S|T) (! P = any amino acid except proline). In tryptic and chymotryptic digests, no consensus sequence-containing peptide was identified by Mascot search even considering semi-digestion, suggesting high glycan-occupancy for all 22 potential sites (Supplementary Tables S2–4). Conversely, *N*-glycosylated sites were identified by IGOT-LC/MS analysis of amide-bound glycopeptides. Mascot search results are shown in Supplementary Tables S5–7. In the chymotryptic digest, 20 *N*-glycosylated sites were identified from 22 potential sites (Supplementary Table S5). In addition, 16 and 11 sites were identified in the tryptic digest and α-Lytic protease digest, respectively (Supplementary Tables S6-7). Therefore, all 22 potential sites were identified as previously glycosylated sites.

Next, using LC/MS2 analysis results of amide-bound glycopeptides of each digest, a Byonic search was carried out to identify glycopeptide forms using an MS/MS spectrum-based approach. The identified glycopeptides are listed in Supplementary Table S8. A total of 919 site-specific glycans were identified for all 22 potential sites (Supplementary Table S9). Finally, using the same LC/MS analysis data, Glyco-RIDGE analyses were performed to identify glycopeptide forms (site-specific glycan compositions). All matched results are shown in Supplementary Table S10. A total of 1,449 site-specific glycans were identified (Supplementary Table S11). Combining the Byonic and Glyco-RIDGE results, 1,759 site-specific glycan compositions and 463 glycan compositions were identified (Supplementary Tables S12-13). The frequencies of the compositions that emerged at the 22 sites are shown in Supplementary Table S14. Among them, the top compositions presented over 15 sites are listed in Table 1. The most frequently found composition was HexNAc:Hex:Fuc:NeuAc:NeuGc = 6:6:1:0:0, which appeared at 20 out of 22 glycosylated sites. The most plausible structure predicted from this composition is a tri-antennary glycan having a bisecting GlcNAc and a fucose, probably a core fucose. Please note that, in the following text, Figures, and Tables, “N:N:N:N:N” means the actual glycan composition of HexNAc:Hex:Fuc:NeuAc:NeuGc. To present glycan compositions indicating their structure intuitively, we used the number of saccharides on the trimannosyl core (Man(3)GlcNAc(2)) and showed the number in the order of Hex-HexNAc-Fuc-NeuAc-NeuGc, e.g., glycan id 20000 for 2:5:0:0:0, or 22100 for 4:5:1:0:0. The top 21 most frequent compositions contained three high-mannose compositions, (N0000), and complex type compositions, including one or more fucoses and no sialic acid (except one, 32010). The emerging frequency of glycan compositions shows that glycans on the S protein were relatively well processed, i.e., highly branched and fucosylated, moderately sialylated, and low extension (polyLacNAc). Fig. 1 lists the number and rate of compositions at each site. Except for three C-terminal sites, high-mannose compositions were found at each site. To estimate the abundance of the high-mannose compositions, extracted ion chromatograms of the high-mannose composition (20000, 30000, 40000) and the top three compositions of each site were obtained for the common core peptide (data not shown). According to the results, the content of high-mannose composition was classified into three categories: rare (or not detected), moderate, and abundant (Fig. 1, Supplementary Fig. S1). Eight sites were found to contain abundant high-mannose glycans: Asn-61, 122, 234, 603, 709, 717, 801, and 1074. These sites also contained a relatively higher content of hybrid compositions (#HexNAc=1, over 15%). Conversely, other sites have a tendency to show higher rates of branching, fucosylation, and sialylation.

**Table 1.** Top 21 frequent glycan compositions emerged in 22 potential sites of SARS CoV-2 Spike protein

| Ranking | Glycan composition | No. of HexNAc | No. of Hex | No. of Fuc | No. of NeuAc | No. of NeuGc | Frequency / 22 sites | Glycan id |
| --- | --- | --- | --- | --- | --- | --- | --- | --- |
| 1 | 6:6:1:0:0 | 6 | 6 | 1 | 0 | 0 | 20 | 34100 |
| 2 | 4:5:1:0:0 | 4 | 5 | 1 | 0 | 0 | 19 | 22100 |
| 3 | 5:6:1:0:0 | 5 | 6 | 1 | 0 | 0 | 19 | 33100 |
| 4 | 6:7:1:0:0 | 6 | 7 | 1 | 0 | 0 | 19 | 44100 |
| 5 | 4:5:2:0:0 | 4 | 5 | 2 | 0 | 0 | 18 | 22200 |
| 6 | 4:5:3:0:0 | 4 | 5 | 3 | 0 | 0 | 18 | 22300 |
| 7 | 5:5:1:0:0 | 5 | 5 | 1 | 0 | 0 | 18 | 23100 |
| 8 | 5:6:2:0:0 | 5 | 6 | 2 | 0 | 0 | 18 | 33200 |
| 9 | 4:5:0:0:0 | 4 | 5 | 0 | 0 | 0 | 17 | 22000 |
| 10 | 5:4:1:0:0 | 5 | 4 | 1 | 0 | 0 | 17 | 13100 |
| 11 | 5:6:3:0:0 | 5 | 6 | 3 | 0 | 0 | 17 | 33300 |
| 12 | 6:7:4:0:0 | 6 | 7 | 4 | 0 | 0 | 17 | 44400 |
| 13 | 2:5:0:0:0 | 2 | 5 | 0 | 0 | 0 | 16 | 20000 |
| 14 | 3:5:1:0:0 | 3 | 5 | 1 | 0 | 0 | 16 | 21100 |
| 15 | 6:7:2:0:0 | 6 | 7 | 2 | 0 | 0 | 16 | 44200 |
| 16 | 6:7:3:0:0 | 6 | 7 | 3 | 0 | 0 | 16 | 44300 |
| 17 | 2:6:0:0:0 | 2 | 6 | 0 | 0 | 0 | 15 | 30000 |
| 18 | 2:7:0:0:0 | 2 | 7 | 0 | 0 | 0 | 15 | 40000 |
| 19 | 3:6:1:0:0 | 3 | 6 | 1 | 0 | 0 | 15 | 31100 |
| 20 | 4:6:0:1:0 | 4 | 6 | 0 | 1 | 0 | 15 | 32010 |
| 21 | 5:5:2:0:0 | 5 | 5 | 2 | 0 | 0 | 15 | 23200 |

**Fig. 1.**
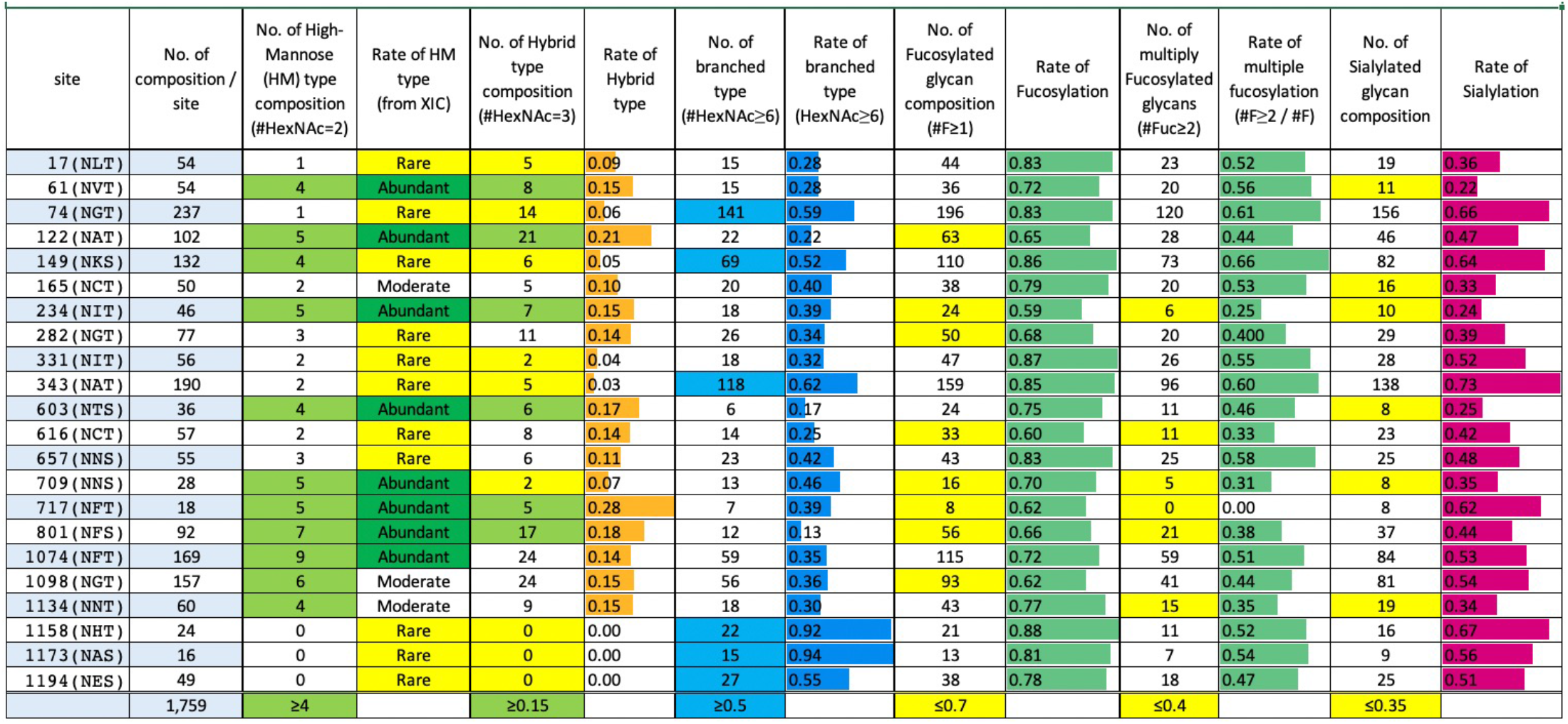
Rates of glycan types and terminal modifications on each glycosylation site. Rates of glycan types such as high-mannose type, hybrid type, and branched type are estimated based on the number of glycan compositions assigned on each site. The cells in which the numbers or rate are larger than criteria presented in the lowest cells, are colored. The rate of HM type is categorized from the extracted ion chromatograms (XIC) as described in the main text. Numbers and rate of terminal modifications such as fucosylation and sialylation are obtained by the same way for type analyses. The cells less than criteria are colored.

### Frequency of glycan compositions

Glycan modification susceptibility at each site was estimated by counting the compositions containing the focused motif or their rate. For example, when the fucosylation rates between sites X and Y are compared, the ratio of the number of fucosylated compositions for total number of compositions was compared. On the other hand, since high-mannose type glycans are limited in number, i.e., there are only five compositions for high-mannose glycans (20000-60000), we compared the rate of high-mannose type glycans, by the count of the high-mannose type compositions and the relative intensities of major high-mannose glycans to those of the top three glycopeptides of each site using extracted ion chromatography of the same core glycopeptides. For Asn-61 (NVT), peak intensities of 20000, 30000, and 40000 of glycopeptide positions 59–65 were compared with those of 22200, 21100, and 31100 of the same core glycopeptides. Because intensity of 30000 was significantly high compared to the others, the high-mannose rate of the site was classified as “abundant.” Conversely, for Asn-74 (NGT), the intensities of 20000, 30000, 40000, 23300, 13300, and 22300 glycopeptides (core positions 66–79) were compared. Slight peaks were observed for high-mannose compositions; thus, the rate is “rare (or not detected).” By comparison, eight sites were found to be high-mannose abundant, 61, 122, 234, 603, 709, 717, 801, and 1074 (Fig. 1).

### Glycan stem distribution

Next, we moved our attention to the glycan modification pathway from high-mannose to highly modified complex-type glycans in Vero/TMPRSS2 cells. Glycan modifications can be divided into two categories: extension or excision of the glycan stem, such as the addition of GlcNAc, Gal, GalNAc, and end-capping (leaves) with fucose and sialic acid, where polysialylation is not considered. From the identified compositions, it is clear that fucosylation actively occurs in Vero/TMPRSS2 cells; therefore, the formation of a glycan stem will be illustrated on #Hex-#HexNAc matrix considering the biosynthetic pathway. The *N*-glycan precursor is attached to the Asn side-chain en bloc as Glc(3)Man(9)GlcNAc(2), so the starting point is Hex(12)HexNAc(2) = 90000 (9-0 in Fig 2A). After successive removal of Glc3, mannoses were excised from the high-mannose glycans to form 20000 (2-0 in Fig 2A). Next, extension of glycan started with the addition of GlcNAc to form hybrid glycan 21000 (2-1 in Fig 2A), followed by Gal addition to 31000 (3-1 in Fig 2A). After removing 2 Man from glycan 3-1, successive addition of GlcNAc and Gal forms bi-antennary glycan 22000 (2-2 in Fig 2A). Further addition of 1 or 2 units of LacNAc (Gal-GlcNAc) formed tri-antennary 33000 (3-3 in Fig 2A) and tetra-antennary 44000 (4-4 in Fig 2A) glycans. Simultaneous addition of HexNAc (such as bisecting GlcNAc or GalNAc to form LacdiNAc) and removal of Hex (Gal or Man) will occur to form complex glycan stems. Fig. 2B shows the frequencies of each composition identified at 22 sites of the S protein. The most frequently observed compositions were 2-1, 2-2, 1-3, 3-3, 3-4, and 4-4, which are presumed to be bi-antennary to tetra-antennary glycans. Glycans 1-3 were presumed to have a LacdiNAc motif. From the frequency mapping, the glycan modification pathway of Vero/TMPRSS2 cells was elucidated. Mass data revealed glycan compositions, and composition mapping provided information on partial glycan motifs such as outer fucosylation, bisecting GlcNAc, LacdiNAc, and less polylactosamine formation. These motifs were suggested by LMA analysis.

**Fig. 2.**
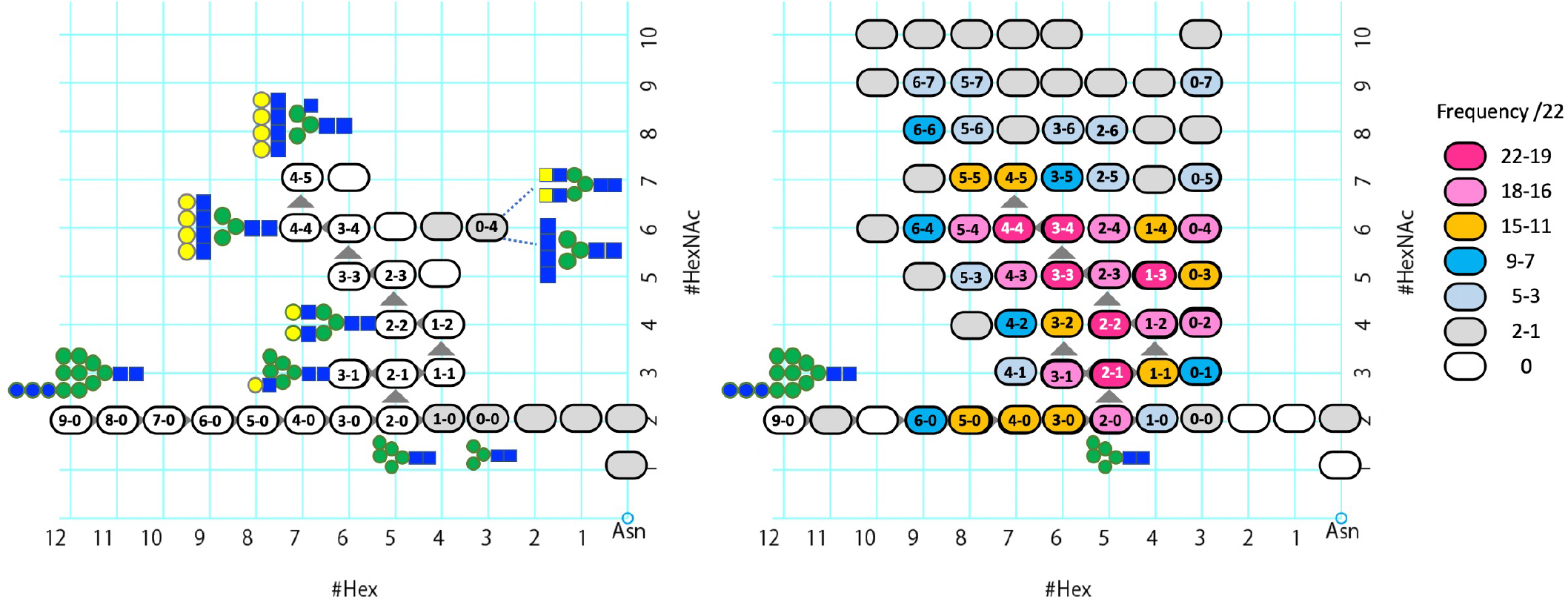
Illustration of formation of glycan stem (#Hex-#HexNAc matrix) considering the biosynthetic pathway. Extension or excision of glycan stem can be illustrated on a matrix of the numbers of Hex (Man or Gal) and HexNAc (GlcNAc or GalNAc). Frequencies of glycan stems (combination of Hex-HexNAc) in 22 potential glycosylation sites are plotted on the matrix.

### LMA analyses of SARS-CoV-2 S protein

Glycoforms of S protein were further analyzed using LMA analysis (Fig 3). First, we tried the “antibody-overlay method,” which we employed in our previous studies (Fig 3A)^20,27^. Immunoprecipitated S proteins were subjected to LMA glass slides and detected with a biotinylated anti-S monoclonal antibody and Alexa555-conjugated streptavidin. As a result, considerable signals in LacNAc-specific (ECA and RCA120) and mannose-specific lectins (NPA, GNA, and HHL) were obtained (Fig 4A), suggesting that both complex and high-mannose type glycans were displayed on the S protein. Signals in sialic acid-specific lectins (MAL-I, SNA, SSA, TJA-I, MAH) provided only low intensity, suggesting a low abundance of sialic acid-terminated glycans. Interestingly, the results were not necessarily consistent with those obtained by MS-based analyses. We then established direct labeling methods coupled with two-step antigen purification with immunoprecipitation (Fig 3B). For this purpose, we labeled immunoprecipitated viral antigens with Cy3 and further purified S proteins from fluorescence-labeled proteins by a second immunoprecipitation with another monoclonal antibody. The purified and Cy3-labeled S protein was then directly applied to LMA glass slides and scanned without overlaying detection antibodies. The results were clearly different from those obtained with the antibody-overlay LMA analyses. Notably, sialic acid-specific lectins showed high signals in this analysis (Fig 4B). In order to further investigate the glycan structures, especially for the outer non-reducing end of glycans, the purified and Cy3-labeled S proteins were digested with exoglycosidases, such as sialidase A and/or β1-4 galactosidase, and the digested antigens were subsequently subjected to LMA analyses (black bars in Fig 5). Signals in sialic acid-specific lectins were partially diminished by digestion with sialidase (Fig 5A). Interestingly, after digestion with β-galactosidase, signals in both sialic acid- and LacNAc-specific lectins partly decreased (Fig 5B). Moreover, double digestion with sialidase and β-galactosidase resulted in nearly complete disappearance of the signals in MAL-I, SNA, SSA, and TJA-I (Fig 5C). These results confirmed that complex-type glycans on S proteins were terminated with either sialic acid or exposed galactose. It should also be noted that signals in the AOL and AAL increased after β-galactosidase digestion. This may indicate that some of the S proteins were modified with Lewis-type fucose. Removing the terminal galactose interacted with fucose, an increase in the mobility of the fucose facilitated enhanced accessibility of AOL and AAL to the glycan. Along with low signals in UEA-I, which is an H-antigen specific lectin, most of the outer fucose structure can be assigned as the Lewis type rather than the H type. The simultaneous acquisition of the *O*-linked glycan information with *N*-glycan suggested the presence of disialyl T-, sialyl T-, and T-antigens indicated by signals in jacalin, ACA, MPA, and MAH. This was confirmed by the enhanced signals of HPA, a Tn-antigen-specific lectin, after β-galactosidase digestion. These *O*-glycan signals derived from the interaction-based assay were relatively weak compared to those of *N*-glycan signals, reflecting the difference in density and accessibility between *N*-and *O*-glycans against the surrounding molecules.

**Fig. 3.**
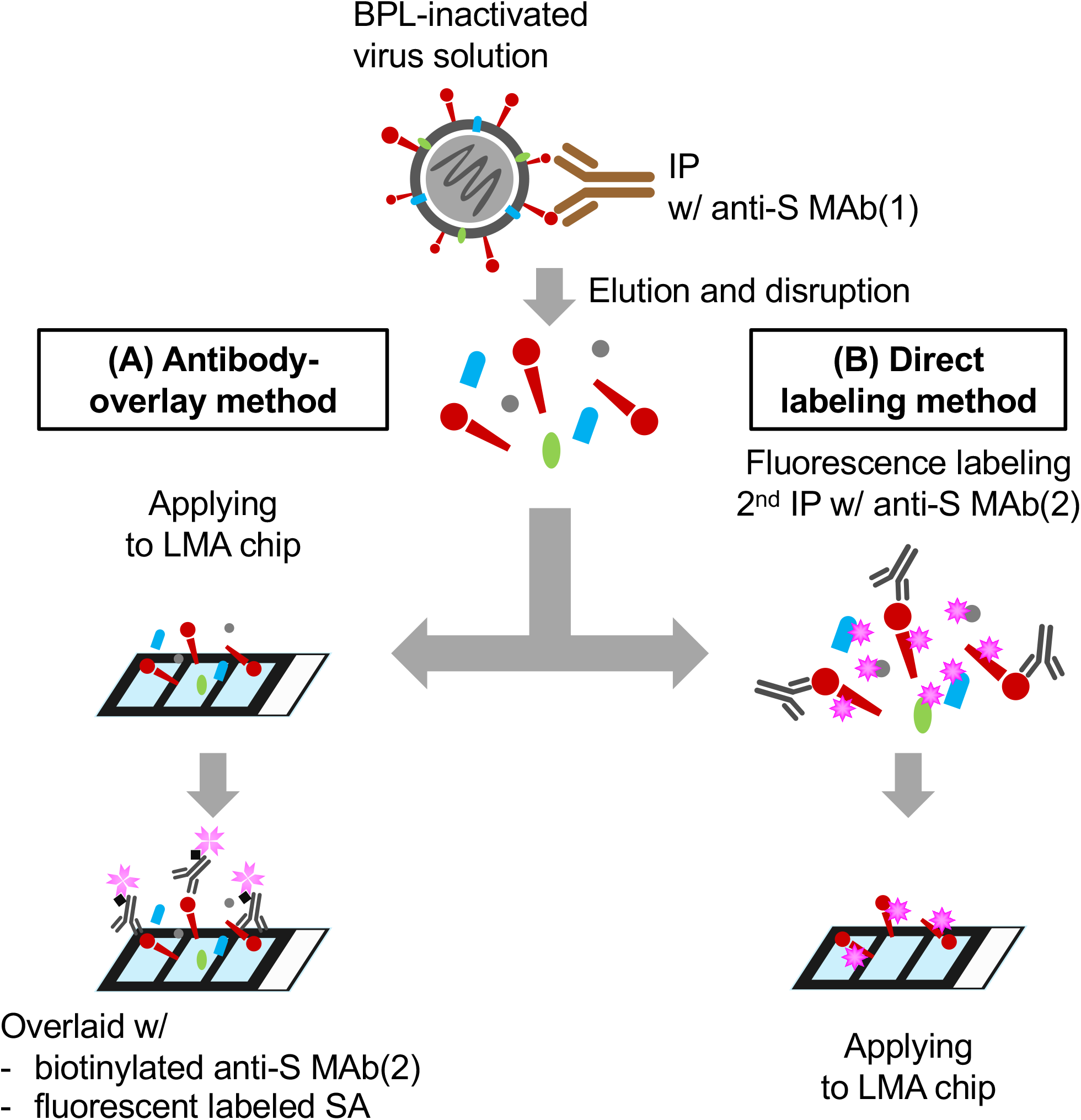
Schematic diagrams for LMA analyses. The samples for antibody-overlay method (A) and direct labeling method (B) were prepared from the same batch of the inactivated virus solution. In the antibody-overlay method, immunoprecipitated viral antigens were detected by overlaying an anti-S specific monoclonal antibody and fluorescent labeled streptavidin. In the direct labeling method, immunoprecipitated viral antigens were directly labeled with fluorophores and S proteins were further purified by second-round immunoprecipitation.

**Fig. 4.**
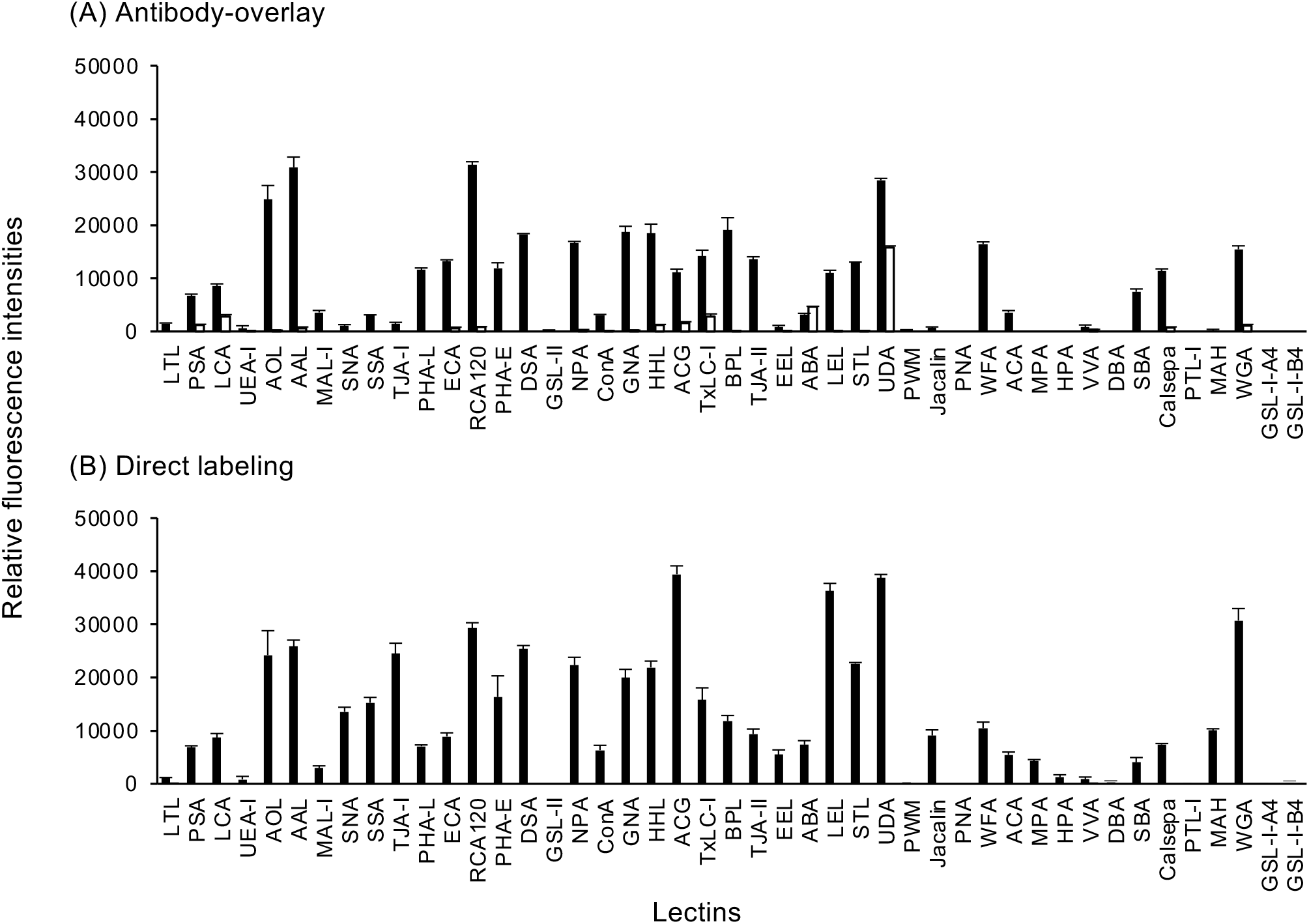
Glycan profiles of S protein. Glycoforms of S protein were analyzed with LMA analyses using 45 lectins. S proteins were enriched from inactivated viral solution (closed bars) or PBS (open bars), and signals were detected with either antibody-overlay (A) or direct labeling (B) methods. Data are represented as mean signals of three spots ± standard deviations (SD).

**Fig. 5.**
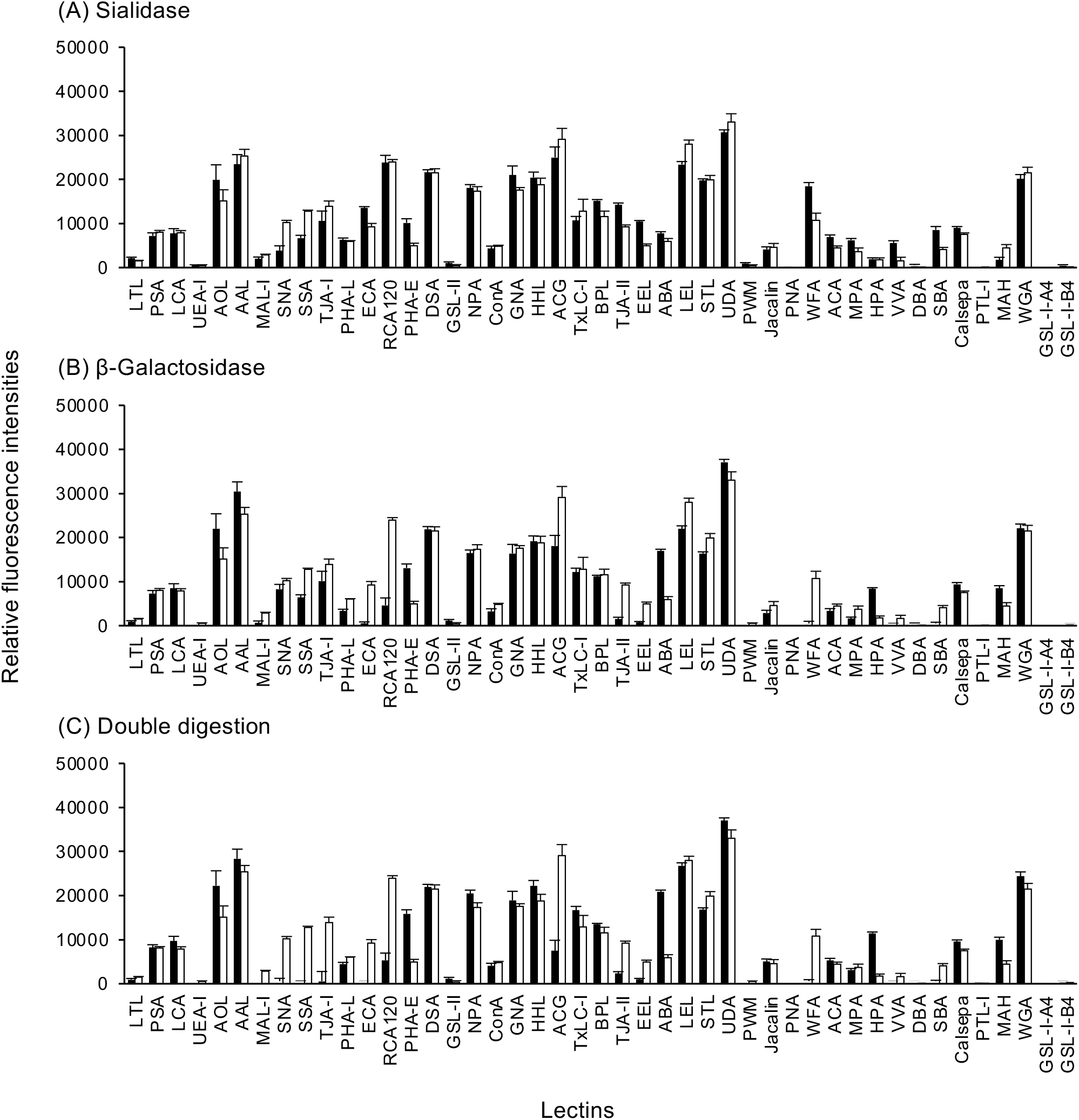
Effects of exoglycosidase digestion on glycan profiles of S protein. Glycoforms of S protein were analyzed with LMA analyses after the digestion with sialidase (A), β-galactosidase (B), and double digestion with sialidase and β-galactosidase (C). S protein prepared with the direct labeling method was digested with exoglycosidases and subjected to analyses. Data are represented as mean signals of three spots ± standard deviations (SD). Closed bars indicate signals in digested samples, and open bars indicate those in undigested samples.

### Glycosylation sites in S proteins were evolutionarily conserved among subgenus Sarbecovirus

To explore the biological significance of *N*-linked glycan, putative *N*-glycosylation sites of S proteins were compared among five viruses belonging to the subgenus *Sarbecovirus*. Therefore, the amino acid sequences of S proteins from five SARS-CoV-2 (2019/nCoV/Japan/TY/WK-521/2020, WHU-01, hCoV-19/England/MILK-9E05B3/2020 (VOC-202012/01), hCoV-19/South Africa/KRISP-K004312/2020 (501.V2 variant), hCoV-19/Japan/IC-0561/2021 (Lineage P.1)), two bat-derived coronaviruses (RaTG13, Rc-o319), a pangolin-derived coronavirus (PCoV-GX-P4L), and SARS coronavirus Tor2 strain were aligned (Supplementary Fig. S2). Although slight variations in glycosylation sites were observed in the N-terminal region, the N-!P-(S|T) sequon motifs were well conserved among viruses belonging to the subgenus *Sarbecovirus*. On the other hand, the lineage P.1 strain, hCoV-19/Japan/IC-0561/2021, acquired two additional N-!P-(S|T) sequences during evolution. Since stepwise aquations of *N*-glycans lead to antigenic changes in the virus^28^, any further acquisition of potential glycosylation in the SARS-CoV-2 S protein during evolution should be monitored.

## Discussion

In the present study, we utilized two approaches, MS and LMA, for glycoproteomic analyses of the SARS-CoV-2 S protein. MS-based analyses reveal site-specific glycans with accurate glycan composition (Supplementary Table S9, 11–12). This approach is especially useful for differentiating complex or high-mannose types or for analyzing the site occupancy of glycans at each site. In contrast, the glycoforms obtained in the present study and the other study were based on cell line-derived virions^8^. Considering that glycoforms of viral glycoproteins are host cell-dependent, it is essential to dissect glycoforms of human-derived virions to utilize glycomic data to develop vaccines or diagnostic methods^20^ For this purpose, the current MS-based approach does not have sufficient sensitivity to analyze human-derived virions. LMA is potentially sensitive enough to analyze human-derived virions. In addition, as an interaction-based approach, LMA is powerful for determining both the terminal structures and the accessibility of glycans. The present results demonstrated that the glycan profile obtained with the direct labeling method was consistent with the results obtained with the MS-based analysis (Fig. 4B). This suggests that the detailed glycoform of human-derived virions may be speculated by refining glycoproteomic data of cell line-derived virions obtained with the MS-based approach with the comparative data of glycan profiles of the cell line- and human-derived virions obtained by LMA.

For in-depth analysis, we further assigned the site-specific glycan compositions using two mass spectrometric approaches. One is the MS2-based approach using the Byonic database search engine, and the other is the MS1-based Glyco-RIDGE approach. The results identified by the two MS approaches are different from each other, and are increased in number when combined (Supplementary Table S14); however, it is therefore difficult to estimate the relative quantity between the identified members, especially when they were identified by different approaches. Similarly, when the compositions were identified from different core peptide sequences, the abundance ratio between the members was unclear. Thus, we compared glycan compositions between sites by counting the compositions containing the focused motif or their rate (Fig. 1).

Our results with MS-based glycoproteomics are in good agreement with the results of Yao et al. for the analysis of intact viral particles and of Watanabe et al. for the recombinant protein^5,8^. The major reason for the retention of high-mannose glycans at specific sites is thought to be the low accessibility of glycan-modifying enzymes (mannosidases, GlcNAc transferases, sialyl transferases, etc.) to the glycan. The fact that the rate of hybrid-type glycans (#HexNAc = #GlcNAc = 1) for all compositions is higher at the high-mannose-abundant sites supports this observation. In addition, these sites also tend to have low rates of branching (#HN ≥ 4) and sialylation, e.g., at Asn-61, 122, 234, 603, 709, and 801. This low accessibility appears to be caused by steric hindrance of the lipid bilayer (ER or Golgi membrane) around the site or by burial of the glycan in the protein cleft or the interface between subunits. In fact, sites 61, 122, and 234 were located in the cleft of the protein of the 3D model of the S protein trimer (Supplementary Fig. S1). In contrast, sites 709, 801, and 1047 were located on the surface of the protein. These sites are located on the bottom surface of the S protein, which is distant from the head domain. On the other hand, the C-terminus Asn-1158, 1173, 1194 were exclusively decorated with complex-type glycans. Since maturation of glycan and core-protein should proceed simultaneously, the higher frequency of sites 709, 801, and 1047 may reflect unknown intracellular events (e.g., host protein interactions or conformational changes) of S protein.

It is important to note that, at these sites, the high-mannose type is major but not exclusive, as indicated in soluble recombinant protein-based analyses by Watanabe et al^5^. On the eight sites, we also found many branched, highly fucosylated, or sialylated glycan compositions, as well as the results of Yao et al., who analyzed parental Vero cell-derived viral particles^8^. The reason for this difference is unclear; however, the sensitivity of the detection (depth/coverage of the glycome of each site) and molecular state (viral particle or over-expressed protein) may influence the distribution. In contrast, at the low high-mannose sites, the rate of HexNAc ≥ 3 (suggesting branching, including addition of bisecting GlcNAc) is relatively high, especially at Asn-1158, 1173, and 1194 (Fig. 1). Similarly, Asn-17, 149, 331, 657, 1158, and 1173 showed high fucosylation rates (> 80%). As described above, although there is a bias in the glycan composition for each site, there is no strict exclusion, and many complex-type glycans have been found at all sites. This may be partly because S proteins are not in exactly the same three-dimensional structural environment (symmetry) in the trimer of S proteins.

It is of interest that the glycoform of SARS-CoV-2 S protein was quite different from the previously reported glycoform of the SARS virus S protein^14^. Most of the complex-type *N*-glycans attached to the SARS virus S protein were represented in the agalactosyl form, suggesting incomplete maturation. In contrast, complex-type *N*-glycans of the SARS-CoV-2 S protein were terminated with either galactose or sialic acid. The difference in the glycoforms of S proteins of SARS virus and SARS-CoV-2 should arise from either a differential impact on the host glycosylation machinery or a differential route of the S protein synthesis pathway. It should also be noted that a murine coronavirus, mouse hepatitis virus, buds from the intermediate compartment between the endoplasmic reticulum and Golgi complex^29^. Virion formation of coronavirus during the early to mid-stage of *N*-glycan maturation is closely related to the glycoform of coronavirus S proteins. On the other hand, the Golgi localization of mannosyl (alpha-1,3-)-glycoprotein beta-1,2-*N*-acetylglucosaminyltransferase (MGAT1)^30^, which is the initiating enzyme for the synthesis of complex-type *N*-glycan, contradicts the fact that coronavirus S proteins are decorated with complex-type *N*-glycans. How the coronavirus S protein encounters MGAT1 is one of the key questions in understanding their post-translational modifications. Perhaps, analyzing the factors associated with the differential glycoforms of SARS and SARS-CoV-2 S proteins may unveil the widely unknown late-stage of the coronavirus lifecycle.

The glycoprotein of the viral protein is strongly associated with its immunogenicity^3^. Our previous study demonstrated that virions propagated in different cell lines showed different glycan profiles^20^. Accordingly, optimizing the glycoform of vaccine antigens by selecting the appropriate host cell line is one of the candidate strategies for developing better vaccines. Our approach, a combination of MS-based and lectin interaction-based glycoproteomics, provided a highly accurate glycan profile of SARS-CoV-2 S protein using a relatively lower amount of viral antigen, resulting in the exploration of meta-heterogeneity, to describe a higher level of glycan regulation: the variation in glycosylation across multiple sites of SARS-CoV-2 S protein. The platform also enables the quick provision of glycoform data of vaccine antigens. In summary, our new concept for glycoproteomic analyses of viral proteins should significantly contribute to establishing effective countermeasures against COVID-19, as well as future viral pandemics.

## Acknowledgements

We thank Ms. Misugi Nagai, Ms. Kaori Ohki, and Ms. Masako Sukegawa for their technical help. We thank the National Institute of Infectious Diseases for providing SARS-CoV-2, 2019-nCoV/Japan/TY/WK-521/2020 strain. We would like to thank all researchers who kindly deposited and shared genomic data on GISAID.

## Declarations

### Ethics approval

The present work does not contain any studies with human participants or animals performed by any of the authors.

### Authors’ contributions

Conceptualization: TH, HK, and AK; methodology: TH and HK; formal analysis: TH, AT, and HK; investigation: TH, HK, MS, and YO; resources: MS, YO, and HS; data curation: TH, AT, HK, and AK; writing original draft: TH and HK; writing review and editing; TH, HK, MS, YO, HS, and AK; visualization: TH and HK; supervision: HK, HS, and AK; project administration: HS and AK; funding acquisition: TH, HK, and HS.

---

**Supplementary Fig. S1.**
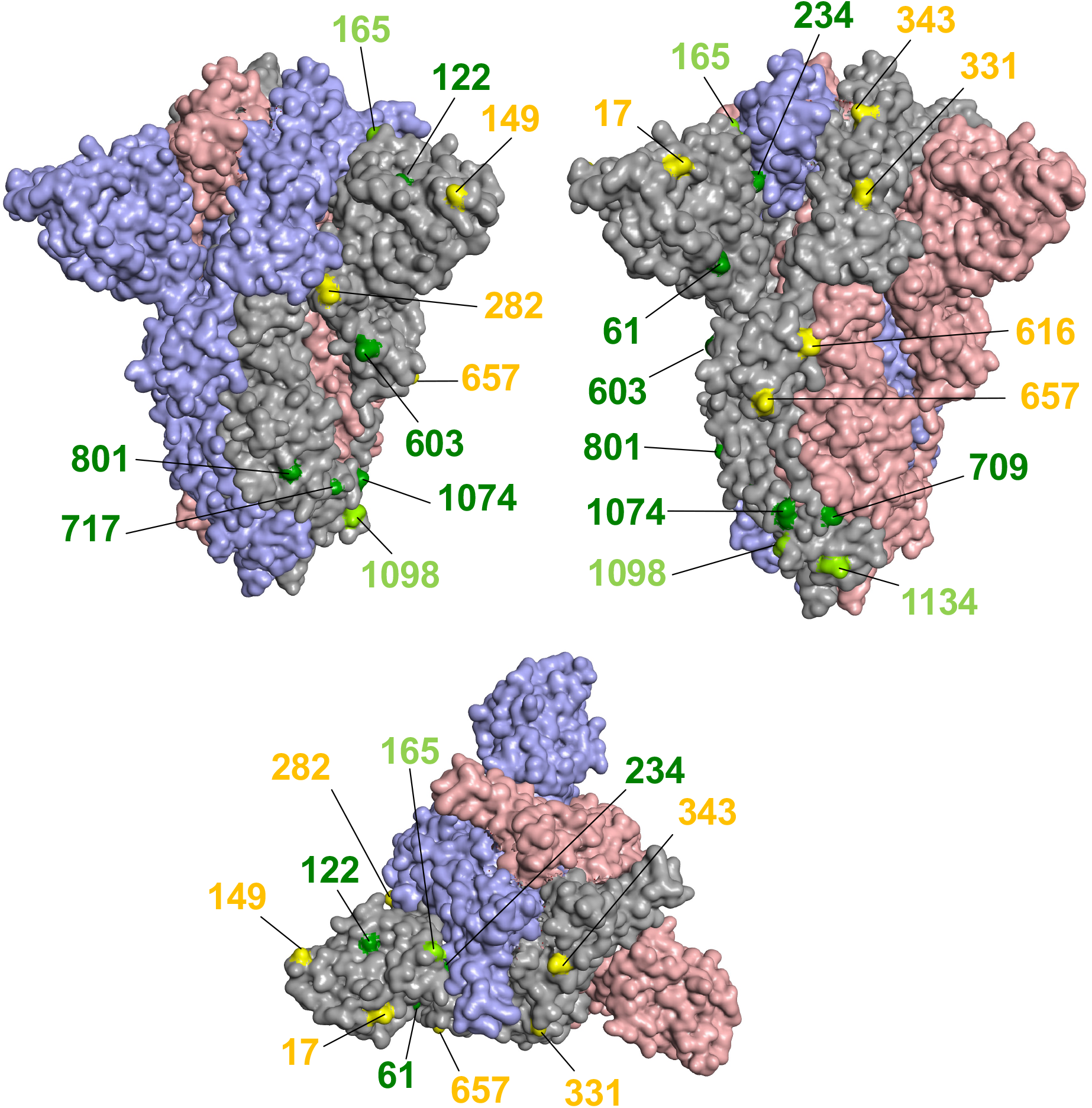
Three-dimensional structural-based mappings of *N*-linked glycosylation sites on S proteins. *N*-linked glycosylation sites of S protein were mapped on the 3D model of S protein in the closed conformation (PDB ID: 6ZGE). Each monomer is depicted in light gray, light pink, or light blue. *N*-linked glycosylation sites on the light gray monomer are colored yellow (rare), light green (moderate), or green (abundant) based on the frequency of high-mannose type glycans. The original structure missed structural information on amino acids 1–13, 71–75, and 1147–1273. Accordingly, sites 74, 1158, 1173, and 1194, all of which were categorized as rare (yellow), are not presented on the structure.

**Supplementary Fig. S2.**
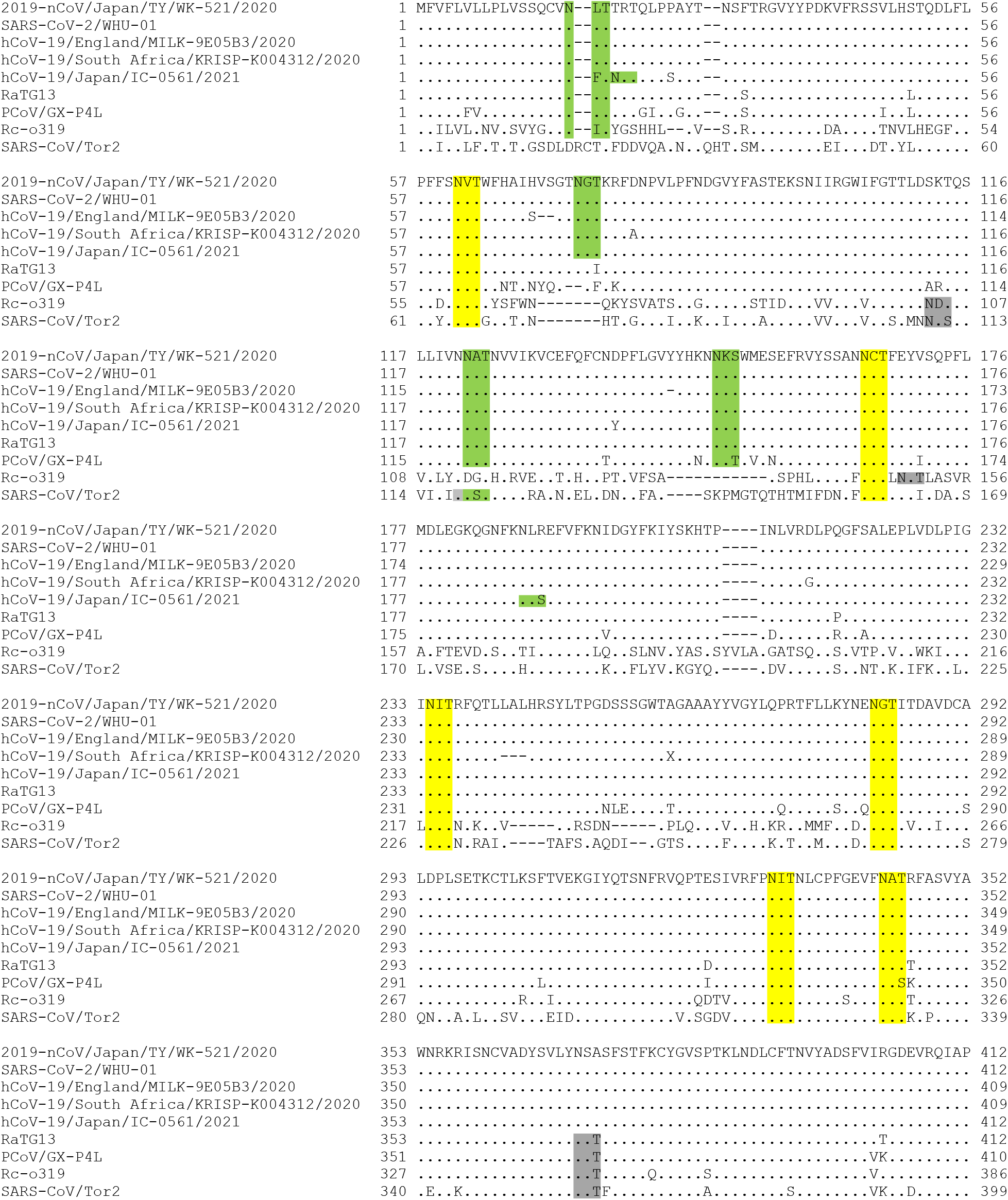
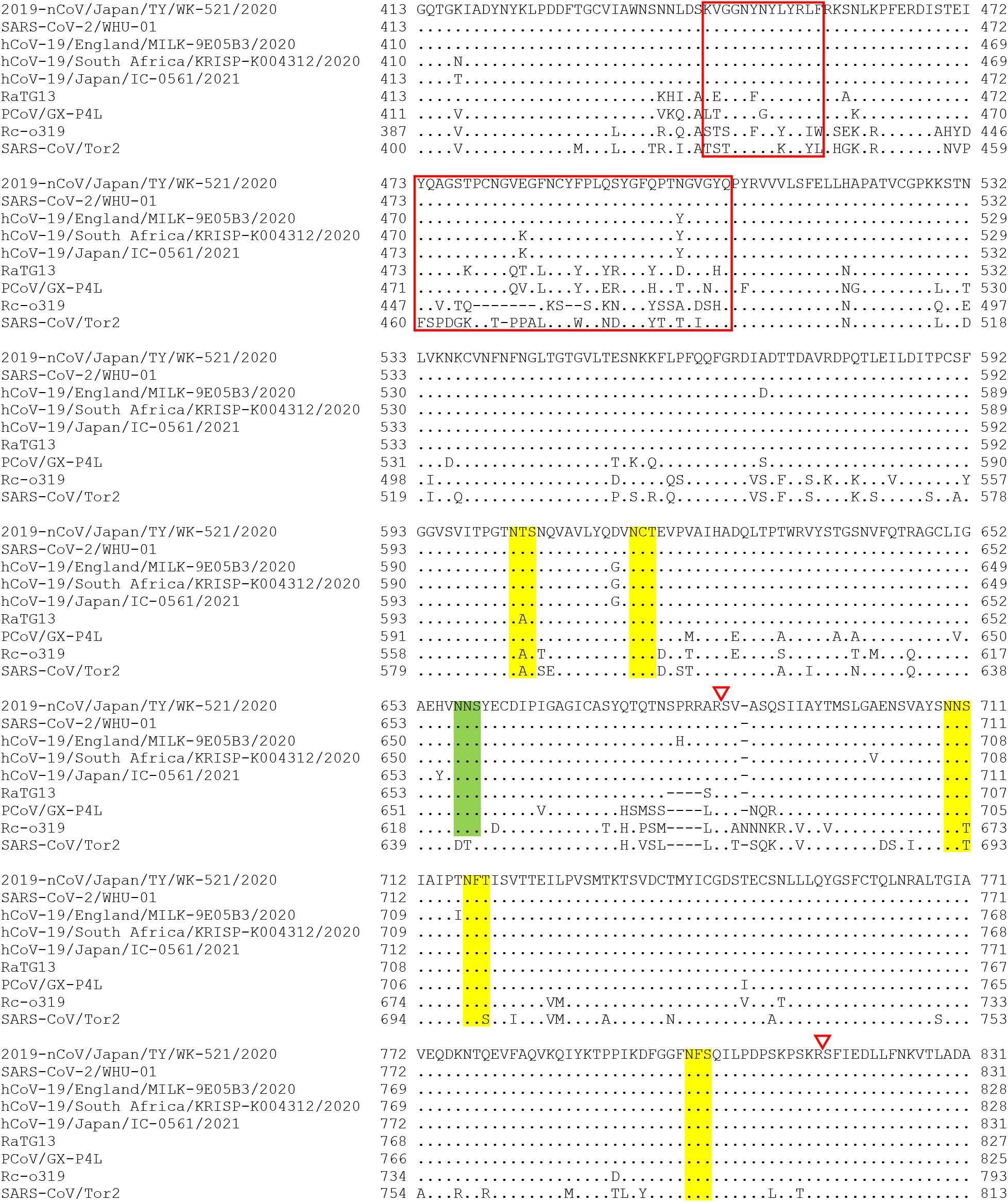
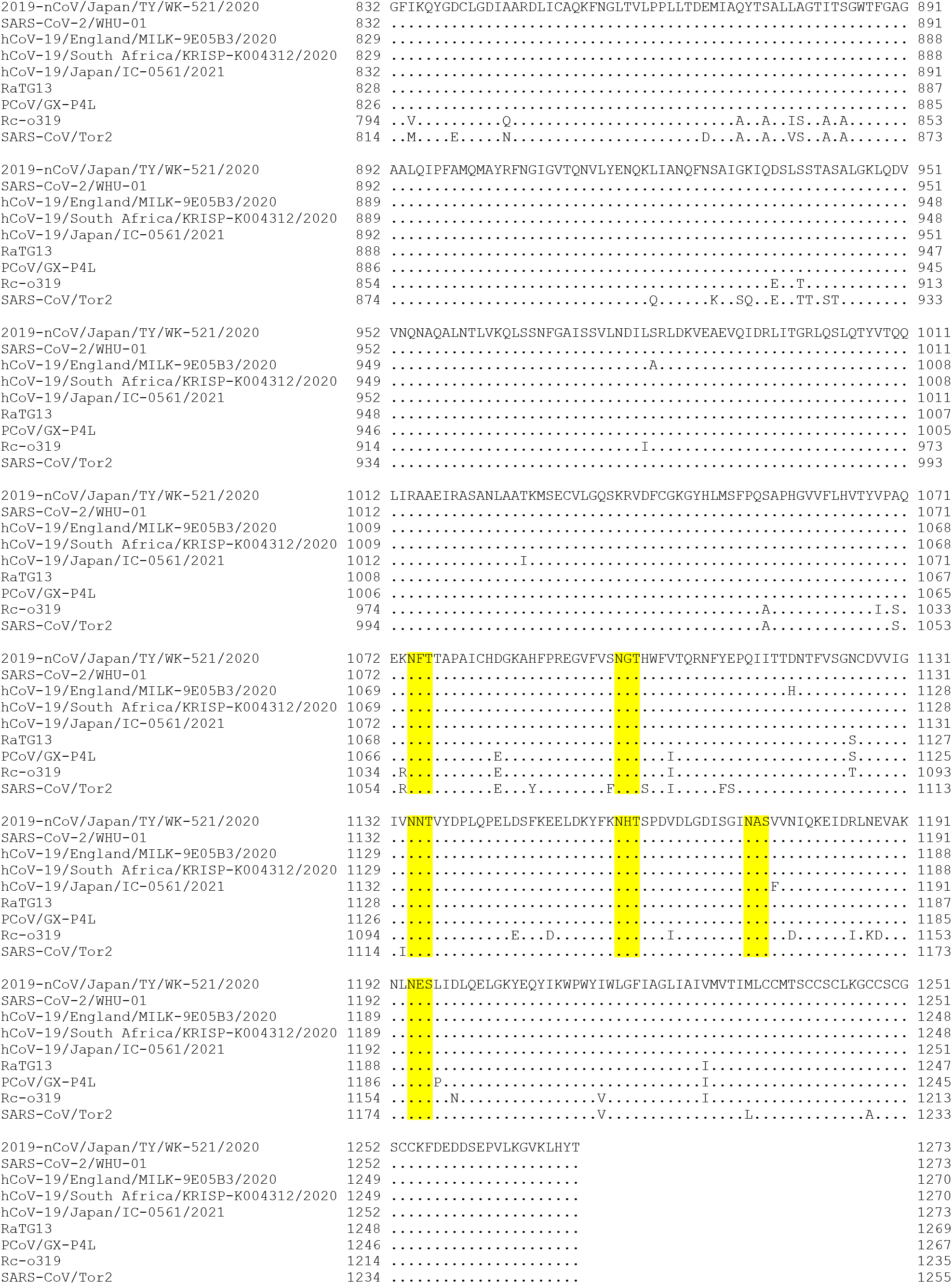
Comparison of potential *N*-glycosylation sites among the S protein of viruses belonging to subgenus *Sarbecovirus*. Amino acid sequences of S proteins from seven viruses belonging to subgenus *Sarbecovirus* were aligned with MUSCLE and N-!P-(S|T) sequons are highlighted. Sequons conserved across all the seven viruses are colored yellow. Sequons observed in SARS-CoV-2 but not conserved across all the members are colored green. Sequons not observed in SARS-CoV-2 are colored gray. ACE2 binding sites are indicated in red boxes. S1/S2 and S2’ proteolytic sites are shown with open red arrowheads.

