## Supplementary Tables for "Combinatorial approach with mass spectrometry and lectin microarray dissected glycoproteomic features of virion-derived spike protein of SARS-CoV-2"

Supplementary Table S1. List of 45 lectins that used in lectin microarray analyses

| No. | Name | Origin | Reported glycan-binding specificity[a] |
|---|---|---|---|
| 1 | LTL | Lotus tetragonolobus | Fucα1-3GlcNAc, Lewis x, Sialyl Lewis x |
| 2 | PSA | Pisum sativum | Fucα1-6GlcNAc , α-Man |
| 3 | LCA | Lens culinaris | Fucα1-6GlcNAc , α-Man, α-Glc |
| 4 | UEA-I | Ulex europaeus | Fucα1-2Galβ1-4GlcNAc |
| 5 | AOL | Aspergillus oryzae | Terminal αFuc, Lewis x, Sialyl Lewis x |
| 6 | AAL | Aleuria auranti | Terminal αFuc, Lewis x, Sialyl Lewis x |
| 7 | MAL-I | Maackia amurensis | Siaα2-3Gal |
| 8 | SNA | Sambucus nigra | Siaα2-6Gal/GalNAc |
| 9 | SSA | Sambucus sieboldiana | Siaα2-6Gal/GalNAc |
| 10 | TJA-I | Trichosanthes japonica | Siaα2-6Galβ1-4GlcNAc |
| 11 | PHA-L | Phaseolus vulgaris | Tri/tetra-antennary complex-type *N*-glycan |
| 12 | ECA | Erythrina cristagalli | Lac/LacNAc |
| 13 | RCA120 | Ricinus communis | Lac/LacNAc |
| 14 | PHA-E | Phaseolus vulgaris | Bi-antennary complex-type N-glycan with outer Gal and bisecting GlcNAc |
| 15 | DSA | Datura stramonium | Chitin, poly LacNAc, LacNAc |
| 16 | GSL-II | Griffonia simplicifolia | Agalactosylated tri/tetra antennary *N*-glycan |
| 17 | NPA | Narcissus pseudonarcissus | High-Mannose including Manα1-6Man |
| 18 | ConA | Canavalia ensiformis | High-Mannose including Manα1-6(Manα1-3)Man |
| 19 | GNA | Galanthus nivalis | High-Mannose including Manα1-3Man |
| 20 | HHL | Hippeastrum hybrid | High-Mannose including Manα1-3Man or Manα1-6Man |
| 21 | ACG | Agrocybe cylindracea | Siaα2-3Galβ1-4GlcNAc |
| 22 | TxLC-I | Tulipa gesneriana | Manα1-3(Manα1-6)Man, bi/tri-antennary complex-type *N*-glycan, GalNAc |
| 23 | BPL | Bauhinia purpurea | Galβ1-3GalNAc, tri/tetra-antennary complex-type N-glycan with outer Gal |
| 24 | TJA-II | ITanthes japonica | Fucα1-2Galβ1/GalNAcβ1, tri/tetra-antennary complex-type N-glycan with outer Gal |
| 25 | EEL | Euonymus europaeus | Galα1-3[Fuc α1-2 Gal], Gal α1-3 Gal |
| 26 | ABA | Agaricus bisporus | Galβ1-3GalNAcα-Thr/Ser (T) and sialyl-T |
| 27 | LEL | Lycopersicon esculentum | Chitin, poly LacNAc |
| 28 | STL | Solanum tuberosum | Chitin, poly LacNAc |
| 29 | UDA | Urtica dioica | Chitin, poly LacNAc |
| 30 | PWM | Phytolacca Americana | Chitin, poly LacNAc |
| 31 | Jacalin | Artocarpus integrifolia | Galβ1-3GalNAcα-Thr/Ser (T) and GalNAcα-Thr/Ser (Tn) |
| 32 | PNA | Arachis hypogaea | Galβ1-3GalNAcα-Thr/Ser (T) |
| 33 | WFA | Wisteria floribunda | GalNAcβ1-4GlcNAc (LacdiNAc), terminal GalNAc |
| 34 | ACA | Amaranthus caudatus | Galβ1-3GalNAcα-Thr/Ser (T) |
| 35 | MPA | Maclura pomifera | Galβ1-3GalNAcα-Thr/Ser (T) and GalNAcα-Thr/Ser (Tn) |
| 36 | HPA | Helix pomatia | Terminal GalNAc |
| 37 | VVA | Vicia villosa | α- or β-linked terminal GalNAc and GalNAcα-Thr/Ser (Tn) |
| 38 | DBA | Dolichos biflorus | GalNAcα-Thr/Ser (Tn) and GalNAcα1-3GalNAc |
| 39 | SBA | Glycine max | Terminal GalNAc |
| 40 | Calsepa | Calystegia sepium | Man, Maltose |
| 41 | PTL-I | Psophocarpus tetragonolobus | α-GalNAc, Gal |
| 42 | MAH | Maackia amurensis | Siaα2-3Galβ1-3(Siaα2-6)GalNAc (disialyl-T) |
| 43 | WGA | Triticum aestivum | Chitin, multivalent Sia |
| 44 | GSL-I-A4 | Griffonia simplicifolia | α-GalNAc, GalNAcα-Thr/Ser (Tn) |
| 45 | GSL-I-B4 | Griffonia simplicifolia | α-Gal |

[a]Specificitiy data of lectins were obtained form LfDB (Lectin frontier Database; http://riodb.ibase.aist.go.jp/rcmg/glycodb/LectinSerch).

Supplementary Table S2. Identification of SARS CoV-2 spike protein-derived peptides of chymotrypsin digest

| Enzyme for database search | No. of identified peptides / protein accession | Protein accession | Query No. | Rank | m/z of precursor ion | Calculated mass value of peptide | Charge (z) of precursor ion | Mass value of assigned peptide | Mass difference | Peptide position from | Peptide position to | No. of missed cleavage | Peptide score of Mascot | Expectation value | Amino acid before peptide | Peptide sequence | Amino acid after peptide | Type and position of variable modifications (1: ammonia-loss (N-term C), 2:pyroGlu (N-term Q), 3: oxidation (M)) | Type of variable modifications | No. of potential N-glycosylation site in peptide |
|---|---|---|---|---|---|---|---|---|---|---|---|---|---|---|---|---|---|---|---|---|
| chymotrypsin (semi) | 100 | 1::P0DTC2 | 7593 | 1 | 681.8544 | 1361.6943 | 2 | 1361.6939 | 0.0003 | 19 | 30 | 1 | 61.87 | 0.00015 | L | TTRTQLPPAYTN | S | | | 0 |
| chymotrypsin (semi) | 100 | 1::P0DTC2 | 4693 | 1 | 602.8037 | 1203.5929 | 2 | 1203.5924 | 0.0004 | 49 | 58 | 2 | 49.11 | 0.0027 | L | HSTQDLFLPF | F | | | 0 |
| chymotrypsin (semi) | 100 | 1::P0DTC2 | 7375 | 1 | 675.3222 | 1348.6298 | 2 | 1348.6299 | -0.0001 | 80 | 91 | 1 | 42.03 | 0.012 | F | DNPVLPFNDGVY | F | | | 0 |
| chymotrypsin (semi) | 100 | 1::P0DTC2 | 5816 | 1 | 422.5667 | 1264.6783 | 3 | 1264.6775 | 0.0008 | 92 | 102 | 1 | 59.79 | 0.00018 | Y | FASTEKSNIIR | G | | | 0 |
| chymotrypsin (semi) | 100 | 1::P0DTC2 | 10080 | 1 | 503.6009 | 1507.7808 | 3 | 1507.7783 | 0.0024 | 92 | 104 | 1 | 88.65 | 3.00E-07 | Y | FASTEKSNIIRGW | I | | | 0 |
| chymotrypsin (semi) | 100 | 1::P0DTC2 | 7575 | 1 | 454.5773 | 1360.7101 | 3 | 1360.7099 | 0.0002 | 93 | 104 | 0 | 89.23 | 2.50E-07 | F | ASTEKSNIIRGW | I | | | 0 |
| chymotrypsin (semi) | 100 | 1::P0DTC2 | 1635 | 1 | 487.2825 | 972.5505 | 2 | 972.5505 | 0 | 97 | 104 | 0 | 40.06 | 0.012 | E | KSNIIRGW | I | | | 0 |
| chymotrypsin (semi) | 100 | 1::P0DTC2 | 442 | 1 | 423.2347 | 844.4549 | 2 | 844.4555 | -0.0007 | 98 | 104 | 0 | 38.6 | 0.02 | K | SNIIRGW | I | | | 0 |
| chymotrypsin (semi) | 100 | 1::P0DTC2 | 7528 | 1 | 680.3053 | 1358.596 | 2 | 1358.5965 | -0.0005 | 134 | 144 | 3 | 60.63 | 8.50E-05 | F | QFCNDPFLGVY | Y | | | 0 |
| chymotrypsin (semi) | 100 | 1::P0DTC2 | 8200 | 1 | 699.8396 | 1397.6646 | 2 | 1397.6649 | -0.0003 | 175 | 186 | 3 | 57.71 | 0.00033 | P | FLMDLEGKQGNF | K | | | 0 |
| chymotrypsin (semi) | 100 | 1::P0DTC2 | 5517 | 1 | 626.305 | 1250.5955 | 2 | 1250.5965 | -0.001 | 176 | 186 | 2 | 51.06 | 0.0014 | F | LMDLEGKQGNF | K | | | 0 |
| chymotrypsin (semi) | 100 | 1::P0DTC2 | 3743 | 1 | 569.7638 | 1137.5131 | 2 | 1137.5125 | 0.0006 | 177 | 186 | 1 | 63.7 | 6.90E-05 | L | MDLEGKQGNF | K | | | 0 |
| chymotrypsin (semi) | 100 | 1::P0DTC2 | 524 | 1 | 428.7138 | 855.4131 | 2 | 855.4127 | 0.0004 | 195 | 201 | 1 | 36.53 | 0.029 | F | KNIDGYF | K | | | 0 |
| chymotrypsin (semi) | 100 | 1::P0DTC2 | 4014 | 1 | 579.8166 | 1157.6186 | 2 | 1157.6193 | -0.0007 | 211 | 220 | 1 | 63.76 | 7.30E-05 | I | NLVRDLPQGF | S | | | 0 |
| chymotrypsin (semi) | 100 | 1::P0DTC2 | 1180 | 1 | 466.2533 | 930.492 | 2 | 930.4923 | -0.0004 | 213 | 220 | 0 | 51.34 | 0.0011 | L | VRDLPQGF | S | | | 0 |
| chymotrypsin (semi) | 100 | 1::P0DTC2 | 15257 | 1 | 608.6738 | 1822.9995 | 3 | 1823.0013 | -0.0019 | 401 | 417 | 0 | 56.43 | 0.00035 | F | VIRGDEVRQIAPGQTGK | I | | | 0 |
| chymotrypsin (semi) | 100 | 1::P0DTC2 | 21173 | 1 | 762.7446 | 2285.2118 | 3 | 2285.2128 | -0.001 | 401 | 421 | 0 | 54.25 | 0.00086 | F | VIRGDEVRQIAPGQTGKIADY | N | | | 0 |
| chymotrypsin (semi) | 100 | 1::P0DTC2 | 5206 | 1 | 617.3271 | 1232.6397 | 2 | 1232.6401 | -0.0004 | 410 | 421 | 0 | 51.62 | 0.0014 | Q | IAPGQTGKIADY | N | | | 0 |
| chymotrypsin (semi) | 100 | 1::P0DTC2 | 14919 | 1 | 899.9269 | 1797.8392 | 2 | 1797.8396 | -0.0004 | 422 | 436 | 2 | 65.25 | 6.00E-05 | Y | NYKLPDDFTGCVIAW | N | | | 0 |
| chymotrypsin (semi) | 100 | 1::P0DTC2 | 10282 | 1 | 761.3738 | 1520.733 | 2 | 1520.7334 | -0.0004 | 424 | 436 | 1 | 38.84 | 0.028 | Y | KLPDDFTGCVIAW | N | | | 0 |
| chymotrypsin (semi) | 100 | 1::P0DTC2 | 7899 | 1 | 691.3201 | 1380.6257 | 2 | 1380.627 | -0.0012 | 437 | 449 | 1 | 77.17 | 3.20E-06 | W | NSNNLDSKVGGNY | N | | | 0 |
| chymotrypsin (semi) | 100 | 1::P0DTC2 | 3580 | 1 | 563.2751 | 1124.5356 | 2 | 1124.5349 | 0.0007 | 465 | 473 | 0 | 41.89 | 0.012 | F | ERDISTEIY | Q | | | 0 |
| chymotrypsin (semi) | 100 | 1::P0DTC2 | 4481 | 1 | 596.2605 | 1190.5065 | 2 | 1190.5066 | -0.0001 | 487 | 495 | 2 | 44.9 | 0.0029 | F | NCYFPLQSY | G | | | 0 |
| chymotrypsin (semi) | 100 | 1::P0DTC2 | 7278 | 1 | 672.8344 | 1343.6542 | 2 | 1343.6544 | -0.0001 | 514 | 525 | 3 | 67.72 | 3.30E-05 | L | SFELLHAPATVC | G | | | 0 |
| chymotrypsin (semi) | 100 | 1::P0DTC2 | 16608 | 1 | 643.0017 | 1925.9832 | 3 | 1925.9847 | -0.0015 | 542 | 559 | 3 | 105.83 | 7.20E-09 | F | NFNGLTGTGVLTESNKKF | L | | | 0 |
| chymotrypsin (semi) | 100 | 1::P0DTC2 | 7904 | 1 | 461.2489 | 1380.7248 | 3 | 1380.7249 | -0.0001 | 547 | 559 | 1 | 84.94 | 5.40E-07 | L | TGTGVLTESNKKF | L | | | 0 |
| chymotrypsin (semi) | 100 | 1::P0DTC2 | 27956 | 1 | 1141.5561 | 3421.6465 | 3 | 3421.646 | 0.0004 | 563 | 592 | 3 | 38.21 | 0.041 | F | QQFGRDIADTTDAVRDPQTLEILDITPCSF | G | | | 0 |
| chymotrypsin (semi) | 100 | 1::P0DTC2 | 15528 | 1 | 615.3093 | 1842.9061 | 3 | 1842.9072 | -0.001 | 566 | 582 | 0 | 91.83 | 1.80E-07 | F | GRDIADTTDAVRDPQTL | E | | | 0 |
| chymotrypsin (semi) | 100 | 1::P0DTC2 | 25070 | 1 | 929.1274 | 2784.3605 | 3 | 2784.36 | 0.0005 | 566 | 590 | 2 | 53.44 | 0.0014 | F | GRDIADTTDAVRDPQTLEILDITPC | S | | | 0 |
| chymotrypsin (semi) | 100 | 1::P0DTC2 | 4536 | 1 | 597.7893 | 1193.5639 | 2 | 1193.5638 | 0.0001 | 583 | 592 | 1 | 38.45 | 0.033 | L | EILDITPCSF | G | | | 0 |
| chymotrypsin (semi) | 100 | 1::P0DTC2 | 2389 | 1 | 519.777 | 1037.5395 | 2 | 1037.5393 | 0.0002 | 686 | 695 | 0 | 44.98 | 0.0058 | R | SVASQSIIAY | T | | | 0 |
| chymotrypsin (semi) | 100 | 1::P0DTC2 | 8072 | 1 | 695.8784 | 1389.7422 | 2 | 1389.7425 | -0.0004 | 719 | 731 | 0 | 39.12 | 0.027 | F | TISVTTEILPVSM | T | | | 0 |
| chymotrypsin (semi) | 100 | 1::P0DTC2 | 4707 | 1 | 603.262 | 1204.5095 | 2 | 1204.5104 | -0.0009 | 732 | 741 | 0 | 34.5 | 0.02 | M | TKTSVDCTMY | I | | | 0 |
| chymotrypsin (semi) | 100 | 1::P0DTC2 | 5586 | 1 | 628.2504 | 1254.4863 | 2 | 1254.4857 | 0.0006 | 742 | 752 | 0 | 29.92 | 0.03 | Y | ICGDSTECSNL | L | | | 0 |
| chymotrypsin (semi) | 100 | 1::P0DTC2 | 14475 | 1 | 886.8951 | 1771.7757 | 2 | 1771.7757 | 0.0001 | 742 | 756 | 3 | 36.71 | 0.022 | Y | ICGDSTECSNLLLQY | G | | | 0 |
| chymotrypsin (semi) | 100 | 1::P0DTC2 | 19357 | 1 | 711.7025 | 2132.0858 | 3 | 2132.0862 | -0.0004 | 764 | 782 | 1 | 97.75 | 5.00E-08 | L | NRALTGIAVEQDKNTQEVF | A | | | 0 |
| chymotrypsin (semi) | 100 | 1::P0DTC2 | 17826 | 1 | 673.6876 | 2018.0411 | 3 | 2018.0433 | -0.0021 | 765 | 782 | 1 | 113.39 | 1.30E-09 | N | RALTGIAVEQDKNTQEVF | A | | | 0 |
| chymotrypsin (semi) | 100 | 1::P0DTC2 | 15805 | 1 | 931.9778 | 1861.9409 | 2 | 1861.9422 | -0.0012 | 766 | 782 | 1 | 60.69 | 0.00026 | R | ALTGIAVEQDKNTQEVF | A | | | 0 |
| chymotrypsin (semi) | 100 | 1::P0DTC2 | 12922 | 1 | 839.9178 | 1677.821 | 2 | 1677.821 | 0 | 768 | 782 | 0 | 76.23 | 6.20E-06 | L | TGIAVEQDKNTQEVF | A | | | 0 |
| chymotrypsin (semi) | 100 | 1::P0DTC2 | 11205 | 1 | 789.3936 | 1576.7726 | 2 | 1576.7733 | -0.0007 | 769 | 782 | 0 | 46.74 | 0.0052 | T | GIAVEQDKNTQEVF | A | | | 0 |
| chymotrypsin (semi) | 100 | 1::P0DTC2 | 1324 | 1 | 473.2739 | 944.5332 | 2 | 944.5331 | 0.0001 | 790 | 797 | 0 | 54.8 | 0.00061 | Y | KTPPIKDF | G | | | 0 |
| chymotrypsin (semi) | 100 | 1::P0DTC2 | 4735 | 1 | 603.8287 | 1205.6429 | 2 | 1205.6445 | -0.0015 | 790 | 800 | 1 | 63.47 | 7.20E-05 | Y | KTPPIKDFGGF | N | | | 0 |
| chymotrypsin (semi) | 100 | 1::P0DTC2 | 4146 | 1 | 584.8142 | 1167.6138 | 2 | 1167.6135 | 0.0002 | 803 | 813 | 0 | 36.88 | 0.028 | F | SQILPDPSKPS | K | | | 0 |
| chymotrypsin (semi) | 100 | 1::P0DTC2 | 9191 | 1 | 484.9437 | 1451.8092 | 3 | 1451.8096 | -0.0004 | 803 | 815 | 0 | 79.79 | 1.30E-06 | F | SQILPDPSKPSKR | S | | | 0 |
| chymotrypsin (semi) | 100 | 1::P0DTC2 | 5854 | 1 | 634.3559 | 1266.6972 | 2 | 1266.6972 | 0 | 814 | 823 | 3 | 50.65 | 0.0011 | S | KRSFIEDLLF | N | | | 0 |
| chymotrypsin (semi) | 100 | 1::P0DTC2 | 3766 | 1 | 570.3084 | 1138.6023 | 2 | 1138.6022 | 0 | 815 | 823 | 3 | 51.84 | 0.0011 | K | RSFIEDLLF | N | | | 0 |
| chymotrypsin (semi) | 100 | 1::P0DTC2 | 6565 | 1 | 652.8765 | 1303.7385 | 2 | 1303.7387 | -0.0002 | 818 | 828 | 3 | 55.23 | 0.00028 | F | IEDLLFNKVTL | A | | | 0 |
| chymotrypsin (semi) | 100 | 1::P0DTC2 | 14369 | 1 | 883.4722 | 1764.9299 | 2 | 1764.9298 | 0.0001 | 818 | 833 | 4 | 77.61 | 3.90E-06 | F | IEDLLFNKVTLADAGF | I | | | 0 |
| chymotrypsin (semi) | 100 | 1::P0DTC2 | 6400 | 1 | 648.3532 | 1294.6918 | 2 | 1294.6921 | -0.0004 | 822 | 833 | 3 | 99.46 | 2.00E-08 | L | LFNKVTLADAGF | I | | | 0 |
| chymotrypsin (semi) | 100 | 1::P0DTC2 | 2368 | 1 | 518.277 | 1034.5393 | 2 | 1034.5397 | -0.0003 | 824 | 833 | 1 | 79.49 | 2.10E-06 | F | NKVTLADAGF | I | | | 0 |
| chymotrypsin (semi) | 100 | 1::P0DTC2 | 11047 | 1 | 784.4276 | 1566.8406 | 2 | 1566.8406 | 0 | 824 | 837 | 2 | 102.86 | 8.60E-09 | F | NKVTLADAGFIKQY | G | | | 0 |
| chymotrypsin (semi) | 100 | 1::P0DTC2 | 3587 | 1 | 563.2997 | 1124.5848 | 2 | 1124.5866 | -0.0018 | 828 | 837 | 2 | 69.09 | 2.00E-05 | T | LADAGFIKQY | G | | | 0 |
| chymotrypsin (semi) | 100 | 1::P0DTC2 | 2090 | 1 | 506.7583 | 1011.502 | 2 | 1011.5025 | -0.0005 | 829 | 837 | 1 | 60.13 | 0.00015 | L | ADAGFIKQY | G | | | 0 |
| chymotrypsin (semi) | 100 | 1::P0DTC2 | 20945 | 1 | 757.0353 | 2268.0842 | 3 | 2268.0845 | -0.0003 | 829 | 849 | 3 | 76.54 | 5.70E-06 | L | ADAGFIKQYGDCLGDIAARDL | I | | | 0 |
| chymotrypsin (semi) | 100 | 1::P0DTC2 | 24725 | 1 | 914.4382 | 2740.2927 | 3 | 2740.2949 | -0.0022 | 829 | 853 | 4 | 52.83 | 0.0013 | L | ADAGFIKQYGDCLGDIAARDLICAQ | K | | | 0 |
| chymotrypsin (semi) | 100 | 1::P0DTC2 | 15057 | 1 | 603.3045 | 1806.8917 | 3 | 1806.8934 | -0.0017 | 834 | 849 | 2 | 47.85 | 0.004 | F | IKQYGDCLGDIAARDL | I | | | 0 |
| chymotrypsin (semi) | 100 | 1::P0DTC2 | 21102 | 1 | 760.7083 | 2279.103 | 3 | 2279.1038 | -0.0009 | 834 | 853 | 3 | 66.37 | 6.00E-05 | F | IKQYGDCLGDIAARDLICAQ | K | | | 0 |
| chymotrypsin (semi) | 100 | 1::P0DTC2 | 5998 | 1 | 638.3026 | 1274.5905 | 2 | 1274.5925 | -0.002 | 838 | 849 | 1 | 62.84 | 8.30E-05 | Y | GDCLGDIAARDL | I | | | 0 |
| chymotrypsin (semi) | 100 | 1::P0DTC2 | 8028 | 1 | 694.8456 | 1387.6766 | 2 | 1387.6766 | 0.0001 | 838 | 850 | 2 | 45.43 | 0.0057 | Y | GDCLGDIAARDLI | C | | | 0 |
| chymotrypsin (semi) | 100 | 1::P0DTC2 | 10729 | 1 | 774.8613 | 1547.708 | 2 | 1547.7072 | 0.0008 | 838 | 851 | 2 | 45.62 | 0.0043 | Y | GDCLGDIAARDLIC | A | | | 0 |
| chymotrypsin (semi) | 100 | 1::P0DTC2 | 14039 | 1 | 874.4078 | 1746.801 | 2 | 1746.8029 | -0.0019 | 838 | 853 | 2 | 93.66 | 7.40E-08 | Y | GDCLGDIAARDLICAQ | K | | | 0 |
| chymotrypsin (semi) | 100 | 1::P0DTC2 | 17926 | 1 | 1011.9908 | 2021.967 | 2 | 2021.9663 | 0.0008 | 838 | 855 | 2 | 69.75 | 2.70E-05 | Y | GDCLGDIAARDLICAQKF | N | | | 0 |
| chymotrypsin (semi) | 100 | 1::P0DTC2 | 17389 | 1 | 994.5235 | 1987.0325 | 2 | 1987.0336 | -0.0011 | 856 | 873 | 3 | 55.58 | 0.00074 | F | NGLTVLPPLLTDEMIAQY | T | | | 0 |
| chymotrypsin (semi) | 100 | 1::P0DTC2 | 5103 | 1 | 614.8278 | 1227.6411 | 2 | 1227.6421 | -0.001 | 859 | 869 | 2 | 54.47 | 0.00057 | L | TVLPPLLTDEM | I | | | 0 |
| chymotrypsin (semi) | 100 | 1::P0DTC2 | 13335 | 1 | 852.4497 | 1702.8848 | 2 | 1702.8851 | -0.0003 | 859 | 873 | 2 | 45.79 | 0.0064 | L | TVLPPLLTDEMIAQY | T | | | 0 |
| chymotrypsin (semi) | 100 | 1::P0DTC2 | 6030 | 1 | 639.3405 | 1276.6663 | 2 | 1276.6663 | 0 | 874 | 886 | 2 | 73.53 | 9.10E-06 | Y | TSALLAGTITSGW | T | | | 0 |
| chymotrypsin (semi) | 100 | 1::P0DTC2 | 4504 | 1 | 596.8212 | 1191.6278 | 2 | 1191.6288 | -0.001 | 887 | 898 | 2 | 49.16 | 0.0025 | W | TFGAGAALQIPF | A | | | 0 |
| chymotrypsin (semi) | 100 | 1::P0DTC2 | 3314 | 1 | 553.2912 | 1104.5678 | 2 | 1104.5676 | 0.0002 | 905 | 914 | 1 | 47.37 | 0.0037 | Y | RFNGIGVTQN | V | | | 0 |
| chymotrypsin (semi) | 100 | 1::P0DTC2 | 4320 | 1 | 589.3131 | 1176.6117 | 2 | 1176.6139 | -0.0022 | 907 | 917 | 1 | 45.25 | 0.0061 | F | NGIGVTQNVLY | E | | | 0 |
| chymotrypsin (semi) | 100 | 1::P0DTC2 | 3838 | 1 | 573.3114 | 1144.6083 | 2 | 1144.6088 | -0.0005 | 928 | 938 | 0 | 48.26 | 0.003 | F | NSAIGKIQDSL | S | | | 0 |
| chymotrypsin (semi) | 100 | 1::P0DTC2 | 9787 | 1 | 746.3865 | 1490.7585 | 2 | 1490.7576 | 0.0009 | 928 | 942 | 1 | 63.53 | 9.60E-05 | F | NSAIGKIQDSLSSTA | S | | | 0 |
| chymotrypsin (semi) | 100 | 1::P0DTC2 | 11219 | 1 | 789.9019 | 1577.7892 | 2 | 1577.7897 | -0.0004 | 928 | 943 | 1 | 60.1 | 0.00023 | F | NSAIGKIQDSLSSTAS | A | | | 0 |
| chymotrypsin (semi) | 100 | 1::P0DTC2 | 14315 | 1 | 881.9628 | 1761.9111 | 2 | 1761.9108 | 0.0003 | 928 | 945 | 1 | 91.87 | 1.50E-07 | F | NSAIGKIQDSLSSTASAL | G | | | 0 |
| chymotrypsin (semi) | 100 | 1::P0DTC2 | 26539 | 1 | 1019.8654 | 3056.5744 | 3 | 3056.5738 | 0.0006 | 928 | 957 | 3 | 65.17 | 9.70E-05 | F | NSAIGKIQDSLSSTASALGKLQDVVNQNAQ | A | | | 0 |
| chymotrypsin (semi) | 100 | 1::P0DTC2 | 27326 | 1 | 1081.2381 | 3240.6925 | 3 | 3240.695 | -0.0025 | 928 | 959 | 3 | 43.27 | 0.014 | F | NSAIGKIQDSLSSTASALGKLQDVVNQNAQAL | N | | | 0 |
| chymotrypsin (semi) | 100 | 1::P0DTC2 | 27753 | 1 | 1119.2527 | 3354.7363 | 3 | 3354.7379 | -0.0016 | 928 | 960 | 4 | 57.37 | 0.00056 | F | NSAIGKIQDSLSSTASALGKLQDVVNQNAQALN | T | | | 0 |
| chymotrypsin (semi) | 100 | 1::P0DTC2 | 12422 | 1 | 824.9414 | 1647.8682 | 2 | 1647.8679 | 0.0003 | 929 | 945 | 1 | 73.96 | 9.40E-06 | N | SAIGKIQDSLSSTASAL | G | | | 0 |
| chymotrypsin (semi) | 100 | 1::P0DTC2 | 12987 | 1 | 841.4652 | 1680.9158 | 2 | 1680.9159 | -0.0001 | 944 | 959 | 2 | 68.35 | 2.50E-05 | S | ALGKLQDVVNQNAQAL | N | | | 0 |

| | | | | | | | | | | | | | | | | | | | | |
|---|---|---|---|---|---|---|---|---|---|---|---|---|---|---|---|---|---|---|---|---|
| chymotrypsin (semi) | 100 | 1::P0DTC2 | 6733 | 1 | 657.3439 | 1312.6732 | 2 | 1312.6735 | -0.0003 | 946 | 957 | 1 | 86.31 | 4.50E-07 | L | GKLQDVVNQNAQ | A | | | 0 |
| chymotrypsin (semi) | 100 | 1::P0DTC2 | 9892 | 1 | 749.4037 | 1496.7929 | 2 | 1496.7947 | -0.0018 | 946 | 959 | 1 | 88.52 | 2.70E-07 | L | GKLQDVVNQNAQAL | N | | | 0 |
| chymotrypsin (semi) | 100 | 1::P0DTC2 | 11809 | 1 | 806.4259 | 1610.8372 | 2 | 1610.8376 | -0.0004 | 946 | 960 | 2 | 87.59 | 4.00E-07 | L | GKLQDVVNQNAQALN | T | | | 0 |
| chymotrypsin (semi) | 100 | 1::P0DTC2 | 3256 | 1 | 551.311 | 1100.6075 | 2 | 1100.6077 | -0.0002 | 971 | 981 | 1 | 42.95 | 0.0076 | F | GAISSVLNDIL | S | | | 0 |
| chymotrypsin (semi) | 100 | 1::P0DTC2 | 11133 | 1 | 786.9324 | 1571.8502 | 2 | 1571.8519 | -0.0016 | 971 | 985 | 3 | 79.95 | 2.00E-06 | F | GAISSVLNDILSRLD | K | | | 0 |
| chymotrypsin (semi) | 100 | 1::P0DTC2 | 16991 | 1 | 653.0123 | 1956.0152 | 3 | 1956.0164 | -0.0012 | 978 | 994 | 2 | 97.05 | 4.90E-08 | L | NDILSRLDKVEAEVQID | R | | | 0 |
| chymotrypsin (semi) | 100 | 1::P0DTC2 | 5966 | 1 | 637.3409 | 1272.6672 | 2 | 1272.6674 | -0.0001 | 982 | 992 | 1 | 74.02 | 7.60E-06 | L | SRLDKVEAEVQ | I | | | 0 |
| chymotrypsin (semi) | 100 | 1::P0DTC2 | 9954 | 1 | 751.3966 | 1500.7787 | 2 | 1500.7784 | 0.0003 | 982 | 994 | 1 | 53.64 | 0.00095 | L | SRLDKVEAEVQID | R | | | 0 |
| chymotrypsin (semi) | 100 | 1::P0DTC2 | 14449 | 1 | 590.9946 | 1769.9619 | 3 | 1769.9635 | -0.0016 | 982 | 996 | 1 | 59.71 | 0.00021 | L | SRLDKVEAEVQIDRL | I | | | 0 |
| chymotrypsin (semi) | 100 | 1::P0DTC2 | 6141 | 1 | 428.9177 | 1283.7312 | 3 | 1283.7309 | 0.0003 | 1013 | 1024 | 0 | 50.87 | 0.00094 | L | IRAAEIRASANL | A | | | 0 |
| chymotrypsin (semi) | 100 | 1::P0DTC2 | 3364 | 1 | 555.2699 | 1108.5252 | 2 | 1108.5257 | -0.0004 | 1025 | 1034 | 0 | 54.9 | 0.00061 | L | AATKMSECVL | G | | | 0 |
| chymotrypsin (semi) | 100 | 1::P0DTC2 | 3578 | 1 | 563.2676 | 1124.5207 | 2 | 1124.5206 | 0.0001 | 1025 | 1034 | 0 | 46.3 | 0.0044 | L | AATKMSECVL | G | 0.0000300000.0 | Oxidation (M) | 0 |
| chymotrypsin (semi) | 100 | 1::P0DTC2 | 10342 | 1 | 508.9205 | 1523.7396 | 3 | 1523.7402 | -0.0007 | 1030 | 1042 | 1 | 64.82 | 7.70E-05 | M | SECVLGQSKRVDF | C | | | 0 |
| chymotrypsin (semi) | 100 | 1::P0DTC2 | 5918 | 1 | 636.8221 | 1271.6296 | 2 | 1271.6299 | -0.0003 | 1051 | 1062 | 0 | 44.29 | 0.0075 | M | SFPQSAPHGVVF | L | | | 0 |
| chymotrypsin (semi) | 100 | 1::P0DTC2 | 644 | 1 | 435.7106 | 869.4067 | 2 | 869.4065 | 0.0001 | 1076 | 1083 | 0 | 75.27 | 2.50E-06 | F | TTAPAICH | D | | | 0 |
| chymotrypsin (semi) | 100 | 1::P0DTC2 | 20292 | 1 | 1106.0402 | 2210.0658 | 2 | 2210.0691 | -0.0033 | 1076 | 1095 | 0 | 78.83 | 3.40E-06 | F | TTAPAICHDGKAHFPREGVF | V | | | 0 |
| chymotrypsin (semi) | 100 | 1::P0DTC2 | 7531 | 1 | 680.343 | 1358.6714 | 2 | 1358.6731 | -0.0017 | 1084 | 1095 | 0 | 42.11 | 0.011 | H | DGKAHFPREGVF | V | | | 0 |
| chymotrypsin (semi) | 100 | 1::P0DTC2 | 984 | 1 | 456.2405 | 910.4665 | 2 | 910.4661 | 0.0004 | 1103 | 1109 | 1 | 41.34 | 0.0073 | W | FVTQRNF | Y | | | 0 |
| chymotrypsin (semi) | 100 | 1::P0DTC2 | 21672 | 1 | 1167.5727 | 2333.1308 | 2 | 2333.1328 | -0.002 | 1103 | 1121 | 3 | 72.39 | 1.60E-05 | W | FVTQRNFYEPQIITTDNTF | V | | | 0 |
| chymotrypsin (semi) | 100 | 1::P0DTC2 | 20024 | 1 | 1094.0384 | 2186.0622 | 2 | 2186.0644 | -0.0022 | 1104 | 1121 | 2 | 44.92 | 0.0087 | F | VTQRNFYEPQIITTDNTF | V | | | 0 |
| chymotrypsin (semi) | 100 | 1::P0DTC2 | 15204 | 1 | 606.9876 | 1817.9409 | 3 | 1817.9411 | -0.0002 | 1255 | 1270 | 2 | 61.55 | 0.00018 | C | KFDEDDSEPVLKGVKL | H | | | 0 |
| chymotrypsin (semi) | 100 | 1::P0DTC2 | 10654 | 1 | 515.2664 | 1542.7774 | 3 | 1542.7777 | -0.0003 | 1257 | 1270 | 1 | 57.12 | 0.00045 | F | DEDDSEPVLKGVKL | H | | | 0 |
| chymotrypsin | 69 | 1::P0DTC2 | 3876 | 1 | 574.3085 | 1146.6024 | 2 | 1146.6033 | -0.0009 | 19 | 28 | 0 | 35.27 | 0.0051 | L | TTRTQLPPAY | T | | | 0 |
| chymotrypsin | 69 | 1::P0DTC2 | 7281 | 1 | 672.8511 | 1343.6876 | 2 | 1343.6874 | 0.0002 | 33 | 43 | 1 | 34.11 | 0.0077 | F | TRGVYYPDKVF | R | | | 0 |
| chymotrypsin | 69 | 1::P0DTC2 | 4693 | 1 | 602.8037 | 1203.5929 | 2 | 1203.5924 | 0.0004 | 49 | 58 | 2 | 49.11 | 0.00027 | L | HSTQDLFLPF | F | | | 0 |
| chymotrypsin | 69 | 1::P0DTC2 | 99 | 1 | 401.2108 | 800.4069 | 2 | 800.4069 | 0.0001 | 80 | 86 | 0 | 32.3 | 0.0056 | F | DNPVLPF | N | | | 0 |
| chymotrypsin | 69 | 1::P0DTC2 | 7375 | 1 | 675.3222 | 1348.6298 | 2 | 1348.6299 | -0.0001 | 80 | 91 | 1 | 42.03 | 0.00099 | F | DNPVLPFNDGVY | F | | | 0 |
| chymotrypsin | 69 | 1::P0DTC2 | 10080 | 1 | 503.6009 | 1507.7808 | 3 | 1507.7783 | 0.0024 | 92 | 104 | 1 | 88.65 | 3.20E-08 | Y | FASTEKSNIIRGW | I | | | 0 |
| chymotrypsin | 69 | 1::P0DTC2 | 7575 | 1 | 454.5773 | 1360.7101 | 3 | 1360.7099 | 0.0002 | 93 | 104 | 0 | 89.23 | 2.50E-08 | F | ASTEKSNIIRGW | I | | | 0 |
| chymotrypsin | 69 | 1::P0DTC2 | 7528 | 1 | 680.3053 | 1358.596 | 2 | 1358.5965 | -0.0005 | 134 | 144 | 3 | 60.63 | 9.70E-06 | F | QFCNDPFLGVY | Y | | | 0 |
| chymotrypsin | 69 | 1::P0DTC2 | 5518 | 1 | 626.3062 | 1250.5977 | 2 | 1250.5965 | 0.0012 | 176 | 186 | 2 | 51.66 | 0.00014 | F | LMDLEGKQGNF | K | | | 0 |
| chymotrypsin | 69 | 1::P0DTC2 | 3743 | 1 | 569.7638 | 1137.5131 | 2 | 1137.5125 | 0.0006 | 177 | 186 | 1 | 63.7 | 8.10E-06 | L | MDLEGKQGNF | K | | | 0 |
| chymotrypsin | 69 | 1::P0DTC2 | 1399 | 1 | 478.2475 | 954.4805 | 2 | 954.4811 | -0.0006 | 193 | 200 | 1 | 33.59 | 0.007 | F | VFKNIDGY | F | | | 0 |
| chymotrypsin | 69 | 1::P0DTC2 | 3260 | 1 | 551.7816 | 1101.5487 | 2 | 1101.5495 | -0.0008 | 193 | 201 | 2 | 35.21 | 0.0053 | F | VFKNIDGYF | K | | | 0 |
| chymotrypsin | 69 | 1::P0DTC2 | 524 | 1 | 428.7138 | 855.4131 | 2 | 855.4127 | 0.0004 | 195 | 201 | 1 | 36.53 | 0.0028 | F | KNIDGYF | K | | | 0 |
| chymotrypsin | 69 | 1::P0DTC2 | 1180 | 1 | 466.2533 | 930.492 | 2 | 930.4923 | -0.0004 | 213 | 220 | 0 | 51.34 | 0.00011 | L | VRDLPQGF | S | | | 0 |
| chymotrypsin | 69 | 1::P0DTC2 | 21173 | 1 | 762.7446 | 2285.2118 | 3 | 2285.2128 | -0.001 | 401 | 421 | 0 | 54.25 | 9.50E-05 | F | VIRGDEVRQIAPGQTGKIADY | N | | | 0 |
| chymotrypsin | 69 | 1::P0DTC2 | 14919 | 1 | 899.9269 | 1797.8392 | 2 | 1797.8396 | -0.0004 | 422 | 436 | 2 | 65.25 | 6.10E-06 | Y | NYKLPDDFTGCVIAW | N | | | 0 |
| chymotrypsin | 69 | 1::P0DTC2 | 10282 | 1 | 761.3738 | 1520.733 | 2 | 1520.7334 | -0.0004 | 424 | 436 | 1 | 38.84 | 0.0035 | Y | KLPDDFTGCVIAW | N | | | 0 |
| chymotrypsin | 69 | 1::P0DTC2 | 7899 | 1 | 691.3201 | 1380.6257 | 2 | 1380.627 | -0.0012 | 437 | 449 | 1 | 77.17 | 3.10E-07 | W | NSNNLDSKVGGNY | N | | | 0 |
| chymotrypsin | 69 | 1::P0DTC2 | 3580 | 1 | 563.2751 | 1124.5356 | 2 | 1124.5349 | 0.0007 | 465 | 473 | 0 | 41.89 | 0.0012 | F | ERDISTEIY | Q | | | 0 |
| chymotrypsin | 69 | 1::P0DTC2 | 4481 | 1 | 596.2605 | 1190.5065 | 2 | 1190.5066 | -0.0001 | 487 | 495 | 2 | 44.9 | 0.00023 | F | NCYFPLQSY | G | | | 0 |
| chymotrypsin | 69 | 1::P0DTC2 | 3127 | 1 | 546.7861 | 1091.5576 | 2 | 1091.5611 | -0.0036 | 542 | 552 | 2 | 33.75 | 0.0092 | F | NFNGLTGTGVL | T | | | 0 |
| chymotrypsin | 69 | 1::P0DTC2 | 16608 | 1 | 643.0017 | 1925.9832 | 3 | 1925.9847 | -0.0015 | 542 | 559 | 3 | 105.83 | 8.20E-10 | F | NFNGLTGTGVLTESNKKF | L | | | 0 |
| chymotrypsin | 69 | 1::P0DTC2 | 7904 | 1 | 461.2489 | 1380.7248 | 3 | 1380.7249 | -0.0001 | 547 | 559 | 1 | 84.94 | 5.20E-08 | L | TGTGVLTESNKKF | L | | | 0 |
| chymotrypsin | 69 | 1::P0DTC2 | 27956 | 1 | 1141.5561 | 3421.6465 | 3 | 3421.646 | 0.0004 | 563 | 592 | 3 | 38.21 | 0.0047 | F | QQFGRDIADTTDAVRDPQTLEILDITPCSF | G | | | 0 |
| chymotrypsin | 69 | 1::P0DTC2 | 15528 | 1 | 615.3093 | 1842.9061 | 3 | 1842.9072 | -0.001 | 566 | 582 | 0 | 91.83 | 1.60E-08 | F | GRDIADTTDAVRDPQTL | E | | | 0 |
| chymotrypsin | 69 | 1::P0DTC2 | 4536 | 1 | 597.7893 | 1193.5639 | 2 | 1193.5638 | 0.0001 | 583 | 592 | 1 | 38.45 | 0.0027 | L | EILDITPCSF | G | | | 0 |
| chymotrypsin | 69 | 1::P0DTC2 | 5362 | 1 | 621.7871 | 1241.5595 | 2 | 1241.5598 | -0.0002 | 696 | 707 | 1 | 32.62 | 0.0074 | Y | TMSLGAENSVAY | S | | | 0 |
| chymotrypsin | 69 | 1::P0DTC2 | 23650 | 1 | 1289.1279 | 2576.2412 | 2 | 2576.2424 | -0.0011 | 719 | 741 | 0 | 38.3 | 0.0047 | F | TISVTTEILPVSMTKTSVDCTMY | I | | | 0 |
| chymotrypsin | 69 | 1::P0DTC2 | 5586 | 1 | 628.2504 | 1254.4863 | 2 | 1254.4857 | 0.0006 | 742 | 752 | 0 | 29.92 | 0.0023 | Y | ICGDSTECSNL | L | | | 0 |
| chymotrypsin | 69 | 1::P0DTC2 | 14475 | 1 | 886.8951 | 1771.7757 | 2 | 1771.7757 | 0.0001 | 742 | 756 | 3 | 36.71 | 0.0021 | Y | ICGDSTECSNLLLQY | G | | | 0 |
| chymotrypsin | 69 | 1::P0DTC2 | 18440 | 1 | 1032.4551 | 2062.8956 | 2 | 2062.8976 | -0.0019 | 742 | 759 | 4 | 31.25 | 0.009 | Y | ICGDSTECSNLLLQYGSF | C | | | 0 |
| chymotrypsin | 69 | 1::P0DTC2 | 19357 | 1 | 711.7025 | 2132.0858 | 3 | 2132.0862 | -0.0004 | 764 | 782 | 1 | 97.75 | 5.00E-09 | L | NRALTGIAVEQDKNTQEVF | A | | | 0 |
| chymotrypsin | 69 | 1::P0DTC2 | 12922 | 1 | 839.9178 | 1677.821 | 2 | 1677.821 | 0 | 768 | 782 | 0 | 76.23 | 5.80E-07 | L | TGIAVEQDKNTQEVF | A | | | 0 |
| chymotrypsin | 69 | 1::P0DTC2 | 1324 | 1 | 473.2739 | 944.5332 | 2 | 944.5331 | 0.0001 | 790 | 797 | 0 | 54.8 | 7.10E-05 | Y | KTPPIKDF | G | | | 0 |
| chymotrypsin | 69 | 1::P0DTC2 | 4735 | 1 | 603.8287 | 1205.6429 | 2 | 1205.6445 | -0.0015 | 790 | 800 | 1 | 63.47 | 6.70E-06 | Y | KTPPIKDFGGF | N | | | 0 |
| chymotrypsin | 69 | 1::P0DTC2 | 6565 | 1 | 652.8765 | 1303.7385 | 2 | 1303.7387 | -0.0002 | 818 | 828 | 3 | 55.23 | 3.60E-05 | F | IEDLLFNKVTL | A | | | 0 |
| chymotrypsin | 69 | 1::P0DTC2 | 14369 | 1 | 883.4722 | 1764.9299 | 2 | 1764.9298 | 0.0001 | 818 | 833 | 4 | 77.61 | 4.60E-07 | F | IEDLLFNKVTLADAGF | I | | | 0 |
| chymotrypsin | 69 | 1::P0DTC2 | 6400 | 1 | 648.3532 | 1294.6918 | 2 | 1294.6921 | -0.0004 | 822 | 833 | 3 | 99.46 | 2.20E-09 | L | LFNKVTLADAGF | I | | | 0 |
| chymotrypsin | 69 | 1::P0DTC2 | 2368 | 1 | 518.277 | 1034.5393 | 2 | 1034.5397 | -0.0003 | 824 | 833 | 1 | 79.49 | 3.00E-07 | F | NKVTLADAGF | I | | | 0 |
| chymotrypsin | 69 | 1::P0DTC2 | 11047 | 1 | 784.4276 | 1566.8406 | 2 | 1566.8406 | 0 | 824 | 837 | 2 | 102.86 | 9.10E-10 | F | NKVTLADAGFIKQY | G | | | 0 |
| chymotrypsin | 69 | 1::P0DTC2 | 2090 | 1 | 506.7583 | 1011.502 | 2 | 1011.5025 | -0.0005 | 829 | 837 | 1 | 60.13 | 1.40E-05 | L | ADAGFIKQY | G | | | 0 |
| chymotrypsin | 69 | 1::P0DTC2 | 20945 | 1 | 757.0353 | 2268.0842 | 3 | 2268.0845 | -0.0003 | 829 | 849 | 3 | 76.54 | 5.20E-07 | L | ADAGFIKQYGDCLGDIAARDL | I | | | 0 |
| chymotrypsin | 69 | 1::P0DTC2 | 15057 | 1 | 603.3045 | 1806.8917 | 3 | 1806.8934 | -0.0017 | 834 | 849 | 2 | 47.85 | 0.00042 | F | IKQYGDCLGDIAARDL | I | | | 0 |
| chymotrypsin | 69 | 1::P0DTC2 | 5998 | 1 | 638.3026 | 1274.5905 | 2 | 1274.5925 | -0.002 | 838 | 849 | 1 | 62.84 | 9.20E-06 | Y | GDCLGDIAARDL | I | | | 0 |
| chymotrypsin | 69 | 1::P0DTC2 | 17926 | 1 | 1011.9908 | 2021.967 | 2 | 2021.9663 | 0.0008 | 838 | 855 | 2 | 69.75 | 2.60E-06 | Y | GDCLGDIAARDLICAQKF | N | | | 0 |
| chymotrypsin | 69 | 1::P0DTC2 | 17389 | 1 | 994.5235 | 1987.0325 | 2 | 1987.0336 | -0.0011 | 856 | 873 | 3 | 55.58 | 6.70E-05 | F | NGLTVLPPLLTDEMIAQY | T | | | 0 |
| chymotrypsin | 69 | 1::P0DTC2 | 13335 | 1 | 852.4497 | 1702.8848 | 2 | 1702.8851 | -0.0003 | 859 | 873 | 2 | 45.79 | 0.00056 | L | TVLPPLLTDEMIAQY | T | | | 0 |
| chymotrypsin | 69 | 1::P0DTC2 | 6030 | 1 | 639.3405 | 1276.6663 | 2 | 1276.6663 | 0 | 874 | 886 | 2 | 73.53 | 8.30E-07 | Y | TSALLAGTITSGW | T | | | 0 |
| chymotrypsin | 69 | 1::P0DTC2 | 5 | 1 | 707.3716 | 706.3644 | 1 | 706.365 | -0.0006 | 887 | 894 | 1 | 29.5 | 0.0067 | W | TFGAGAAL | Q | | | 0 |
| chymotrypsin | 69 | 1::P0DTC2 | 4504 | 1 | 596.8212 | 1191.6278 | 2 | 1191.6288 | -0.001 | 887 | 898 | 2 | 49.16 | 0.00021 | W | TFGAGAALQIPF | A | | | 0 |
| chymotrypsin | 69 | 1::P0DTC2 | 1307 | 1 | 472.7632 | 943.5119 | 2 | 943.5127 | -0.0008 | 889 | 898 | 1 | 29.72 | 0.017 | F | GAGAALQIPF | A | | | 0 |
| chymotrypsin | 69 | 1::P0DTC2 | 4320 | 1 | 589.3131 | 1176.6117 | 2 | 1176.6139 | -0.0022 | 907 | 917 | 1 | 45.25 | 0.00065 | F | NGIGVTQNVLY | E | | | 0 |
| chymotrypsin | 69 | 1::P0DTC2 | 4696 | 1 | 602.8186 | 1203.6227 | 2 | 1203.6248 | -0.0021 | 918 | 927 | 1 | 32.98 | 0.0086 | Y | ENQKLIANQF | N | | | 0 |
| chymotrypsin | 69 | 1::P0DTC2 | 3838 | 1 | 573.3114 | 1144.6083 | 2 | 1144.6088 | -0.0005 | 928 | 938 | 0 | 48.26 | 0.00026 | F | NSAIGKIQDSL | S | | | 0 |
| chymotrypsin | 69 | 1::P0DTC2 | 14315 | 1 | 881.9628 | 1761.9111 | 2 | 1761.9108 | 0.0003 | 928 | 945 | 1 | 91.87 | 1.50E-08 | F | NSAIGKIQDSLSSTASAL | G | | | 0 |
| chymotrypsin | 69 | 1::P0DTC2 | 27326 | 1 | 1081.2381 | 3240.6925 | 3 | 3240.695 | -0.0025 | 928 | 959 | 3 | 43.27 | 0.0015 | F | NSAIGKIQDSLSSTASALGKLQDVVNQNAQAL | N | | | 0 |
| chymotrypsin | 69 | 1::P0DTC2 | 9892 | 1 | 749.4037 | 1496.7929 | 2 | 1496.7947 | -0.0018 | 946 | 959 | 1 | 88.52 | 2.60E-08 | L | GKLQDVVNQNAQAL | N | | | 0 |
| chymotrypsin | 69 | 1::P0DTC2 | 202 | 1 | 408.253 | 814.4914 | 2 | 814.4912 | 0.0002 | 960 | 966 | 1 | 32.39 | 0.0081 | L | NTLVKQL | S | | | 0 |
| chymotrypsin | 69 | 1::P0DTC2 | 3256 | 1 | 551.311 | 1100.6075 | 2 | 1100.6077 | -0.0002 | 971 | 981 | 1 | 42.95 | 0.00086 | F | GAISSVLNDIL | S | | | 0 |
| chymotrypsin | 69 | 1::P0DTC2 | 14449 | 1 | 590.9946 | 1769.9619 | 3 | 1769.9635 | -0.0016 | 982 | 996 | 1 | 59.71 | 2.20E-05 | L | SRLDKVEAEVQIDRL | I | | | 0 |
| chymotrypsin | 69 | 1::P0DTC2 | 6141 | 1 | 428.9177 | 1283.7312 | 3 | 1283.7309 | 0.0003 | 1013 | 1024 | 0 | 50.87 | 0.00012 | L | IRAAEIRASANL | A | | | 0 |
| chymotrypsin | 69 | 1::P0DTC2 | 3364 | 1 | 555.2699 | 1108.5252 | 2 | 1108.5257 | -0.0004 | 1025 | 1034 | 0 | 54.9 | 6.20E-05 | L | AATKMSECVL | G | | | 0 |

| | | | | | | | | | | | | | | | | | | | | |
|---|---|---|---|---|---|---|---|---|---|---|---|---|---|---|---|---|---|---|---|---|
| chymotrypsin | 69 | 1::P0DTC2 | 3578 | 1 | 563.2676 | 1124.5207 | 2 | 1124.5206 | 0.0001 | 1025 | 1034 | 0 | 46.3 | 0.00042 | L | AATKMSECVL | G | 0.0000300000.0 | Oxidation (M) | 0 |
| chymotrypsin | 69 | 1::P0DTC2 | 20292 | 1 | 1106.0402 | 2210.0658 | 2 | 2210.0691 | -0.0033 | 1076 | 1095 | 0 | 78.83 | 3.10E-07 | F | TTAPAICHDGKAHFPREGVF | V | | | 0 |
| chymotrypsin | 69 | 1::P0DTC2 | 984 | 1 | 456.2405 | 910.4665 | 2 | 910.4661 | 0.0004 | 1103 | 1109 | 1 | 41.34 | 0.00079 | W | FVTQRNF | Y | | | 0 |
| chymotrypsin | 69 | 1::P0DTC2 | 21672 | 1 | 1167.5727 | 2333.1308 | 2 | 2333.1328 | -0.002 | 1103 | 1121 | 3 | 72.39 | 1.50E-06 | W | FVTQRNFYEPQIITTDNTF | V | | | 0 |
| chymotrypsin | 69 | 1::P0DTC2 | 20024 | 1 | 1094.0384 | 2186.0622 | 2 | 2186.0644 | -0.0022 | 1104 | 1121 | 2 | 44.92 | 0.00099 | F | VTQRNFYEPQIITTDNTF | V | | | 0 |
| chymotrypsin | 69 | 1::P0DTC2 | 4047 | 1 | 580.7766 | 1159.5387 | 2 | 1159.5397 | -0.0011 | 1139 | 1148 | 2 | 29.23 | 0.019 | Y | DPLQPELDSF | K | | | 0 |
| chymotrypsin | 69 | 1::P0DTC2 | 10654 | 1 | 515.2664 | 1542.7774 | 3 | 1542.7777 | -0.0003 | 1257 | 1270 | 1 | 57.12 | 5.40E-05 | F | DEDDSEPVLKGVKL | H | | | 0 |

* Results shown in blue are overlapped with those for semi-search.

Supplementary Table S3. Identification of SARS CoV-2 spike protein-derived peptides of tryptic digest

| Enzyme for database search | No. of identified peptides / protein accession | Protein accession | Query No. | Rank | m/z of precursor ion | Calculated mass value of peptide | Charge (z) of precursor ion | Mass value of assigned peptide | Mass difference | Peptide position from | Peptide position to | No. of missed cleavage | Peptide score of Mascot | Expectation value | Amino acid before peptide | Peptide sequence | Amino acid after peptide | Type and position of variable modifications (1: ammonia-loss (N-term C), 2:pyroGlu (N-term Q), 3: oxidation (M)) | Type of variable modifications | No. of potential N-glycosylation site in peptide |
|---|---|---|---|---|---|---|---|---|---|---|---|---|---|---|---|---|---|---|---|---|
| trypsin (semi) | 63 | 1::P0DTC2 | 9522 | 1 | 748.3814 | 1494.7481 | 2 | 1494.7467 | 0.0015 | 22 | 34 | 0 | 67.44 | 3.10E-05 | R | TQLPPAYTNSFTR | G | | | 0 |
| trypsin (semi) | 63 | 1::P0DTC2 | 374 | 1 | 421.2082 | 840.4018 | 2 | 840.4018 | 0.0001 | 35 | 41 | 0 | 43.45 | 0.0037 | R | GVYYPDK | V | | | 0 |
| trypsin (semi) | 63 | 1::P0DTC2 | 20198 | 1 | 772.7143 | 2315.1211 | 3 | 2315.1223 | -0.0011 | 78 | 97 | 1 | 111.56 | 1.00E-09 | K | RFDNPVLPFNDGVYFASTEK | S | | | 0 |
| trypsin (semi) | 63 | 1::P0DTC2 | 18924 | 1 | 720.6806 | 2159.0199 | 3 | 2159.0211 | -0.0012 | 79 | 97 | 0 | 46.81 | 0.0024 | R | FDNPVLPFNDGVYFASTEK | S | | | 0 |
| trypsin (semi) | 63 | 1::P0DTC2 | 14314 | 1 | 892.4496 | 1782.8847 | 2 | 1782.8829 | 0.0018 | 82 | 97 | 0 | 79.09 | 1.70E-06 | N | PVLPFNDGVYFASTEK | S | | | 0 |
| trypsin (semi) | 63 | 1::P0DTC2 | 4989 | 1 | 612.8164 | 1223.6183 | 2 | 1223.6187 | -0.0003 | 103 | 113 | 0 | 76.46 | 2.80E-06 | R | GWIFGTTLDSK | T | | | 0 |
| trypsin (semi) | 63 | 1::P0DTC2 | 20249 | 1 | 775.0149 | 2322.0227 | 3 | 2322.0238 | -0.0011 | 130 | 147 | 0 | 148.12 | 7.90E-14 | K | VCEFQFCNDPFLGVYYHK | N | | | 0 |
| trypsin (semi) | 63 | 1::P0DTC2 | 2810 | 1 | 536.2317 | 1070.4488 | 2 | 1070.4491 | -0.0003 | 151 | 158 | 0 | 63.52 | 1.20E-05 | K | SWMESEFR | V | | | 0 |
| trypsin (semi) | 63 | 1::P0DTC2 | 1423 | 1 | 475.2819 | 948.5492 | 2 | 948.5505 | -0.0013 | 207 | 214 | 0 | 47.41 | 0.00093 | K | HTPINLVR | D | | | 0 |
| trypsin (semi) | 63 | 1::P0DTC2 | 3183 | 1 | 549.8246 | 1097.6346 | 2 | 1097.6346 | 0.0001 | 238 | 246 | 0 | 90.07 | 4.80E-08 | R | FQTLLALHR | S | | | 0 |
| trypsin (semi) | 63 | 1::P0DTC2 | 1450 | 1 | 476.2903 | 950.5659 | 2 | 950.5661 | -0.0002 | 239 | 246 | 0 | 36.02 | 0.0059 | F | QTLLALHR | S | | | 0 |
| trypsin (semi) | 63 | 1::P0DTC2 | 226 | 1 | 412.2614 | 822.5083 | 2 | 822.5076 | 0.0008 | 240 | 246 | 0 | 34.64 | 0.0051 | Q | TLLALHR | S | | | 0 |
| trypsin (semi) | 63 | 1::P0DTC2 | 22831 | 1 | 1419.6712 | 2837.3278 | 2 | 2837.3297 | -0.0018 | 247 | 273 | 0 | 149.09 | 1.30E-13 | R | SYLTPGDSSSGWTAGAAAYYVGYLQPR | T | | | 0 |
| trypsin (semi) | 63 | 1::P0DTC2 | 6133 | 1 | 650.8378 | 1299.661 | 2 | 1299.6611 | -0.0002 | 263 | 273 | 0 | 67.11 | 2.50E-05 | A | AAYYVGYLQPR | T | | | 0 |
| trypsin (semi) | 63 | 1::P0DTC2 | 5060 | 1 | 615.3203 | 1228.6261 | 2 | 1228.624 | 0.002 | 264 | 273 | 0 | 40.22 | 0.011 | A | AYYVGYLQPR | T | | | 0 |
| trypsin (semi) | 63 | 1::P0DTC2 | 1928 | 1 | 498.2684 | 994.5223 | 2 | 994.5236 | -0.0013 | 266 | 273 | 0 | 53.9 | 0.00048 | Y | YVGYLQPR | T | | | 0 |
| trypsin (semi) | 63 | 1::P0DTC2 | 3006 | 1 | 543.272 | 1084.5295 | 2 | 1084.5302 | -0.0006 | 311 | 319 | 0 | 54.21 | 0.00036 | K | GIYQTSNFR | V | | | 0 |
| trypsin (semi) | 63 | 1::P0DTC2 | 2322 | 1 | 514.7901 | 1027.5657 | 2 | 1027.5662 | -0.0005 | 320 | 328 | 0 | 42.54 | 0.0051 | R | VQPTESIVR | F | | | 0 |
| trypsin (semi) | 63 | 1::P0DTC2 | 3398 | 1 | 557.2765 | 1112.5385 | 2 | 1112.5403 | -0.0018 | 347 | 355 | 0 | 49.37 | 0.0011 | R | FASVYAWNR | K | | | 0 |
| trypsin (semi) | 63 | 1::P0DTC2 | 21572 | 1 | 843.7408 | 2528.2005 | 3 | 2528.2006 | -0.0001 | 357 | 378 | 1 | 111.49 | 8.70E-10 | K | RISNCVADYSVLYNSASFSTFK | C | | | 0 |
| trypsin (semi) | 63 | 1::P0DTC2 | 20617 | 1 | 1187.0566 | 2372.0986 | 2 | 2372.0995 | -0.0008 | 358 | 378 | 0 | 117.04 | 1.90E-10 | R | ISNCVADYSVLYNSASFSTFK | C | | | 0 |
| trypsin (semi) | 63 | 1::P0DTC2 | 8724 | 1 | 725.8646 | 1449.7147 | 2 | 1449.714 | 0.0007 | 366 | 378 | 0 | 85.96 | 3.30E-07 | Y | SVLYNSASFSTFK | C | | | 0 |
| trypsin (semi) | 63 | 1::P0DTC2 | 690 | 1 | 437.7189 | 873.4232 | 2 | 873.4232 | 0 | 371 | 378 | 0 | 62.07 | 6.00E-05 | N | SASFSTFK | C | | | 0 |
| trypsin (semi) | 63 | 1::P0DTC2 | 1017 | 1 | 456.2182 | 910.4218 | 2 | 910.4219 | 0 | 379 | 386 | 0 | 55.09 | 0.00025 | K | CYGVSPTK | L | | | 0 |
| trypsin (semi) | 63 | 1::P0DTC2 | 17736 | 1 | 1024.0007 | 2045.9868 | 2 | 2045.9881 | -0.0012 | 387 | 403 | 0 | 135.45 | 3.70E-12 | K | LNDLCFTNVYADSFVIR | G | | | 0 |
| trypsin (semi) | 63 | 1::P0DTC2 | 801 | 1 | 443.7189 | 885.4233 | 2 | 885.4232 | 0.0001 | 418 | 424 | 0 | 43.48 | 0.0034 | K | IADYNYK | L | | | 0 |
| trypsin (semi) | 63 | 1::P0DTC2 | 19810 | 1 | 1133.5257 | 2265.0368 | 2 | 2265.0372 | -0.0004 | 425 | 444 | 0 | 82.06 | 5.50E-07 | K | LPDDFTGCVIAWNSNNLDSK | V | | | 0 |
| trypsin (semi) | 63 | 1::P0DTC2 | 4904 | 1 | 609.7982 | 1217.5819 | 2 | 1217.5829 | -0.001 | 445 | 454 | 0 | 92.57 | 6.70E-08 | K | VGGNYNYLYR | L | | | 0 |
| trypsin (semi) | 63 | 1::P0DTC2 | 17625 | 1 | 1019.063 | 2036.1114 | 2 | 2036.1129 | -0.0014 | 510 | 528 | 0 | 131.79 | 4.50E-12 | R | VVVLSFELLHAPATVCGPK | K | | | 0 |
| trypsin (semi) | 63 | 1::P0DTC2 | 19009 | 1 | 1086.5221 | 2171.0296 | 2 | 2171.0317 | -0.0021 | 538 | 557 | 0 | 109.03 | 1.40E-09 | K | CVNFNFNGLTGTGVLTESNK | K | | | 0 |
| trypsin (semi) | 63 | 1::P0DTC2 | 19952 | 1 | 1142.0562 | 2282.0978 | 2 | 2282.1002 | -0.0023 | 538 | 558 | 1 | 69.28 | 1.60E-05 | K | CVNFNFNGLTGTGVLTESNKK | F | 1.0000000000000000000000.0 | Ammonia-loss (N-term C) | 0 |
| trypsin (semi) | 63 | 1::P0DTC2 | 5820 | 1 | 638.8404 | 1275.6662 | 2 | 1275.6671 | -0.0008 | 545 | 557 | 0 | 70.37 | 1.30E-05 | N | GLTGTGVLTESNK | K | | | 0 |
| trypsin (semi) | 63 | 1::P0DTC2 | 3751 | 1 | 570.3035 | 1138.5924 | 2 | 1138.5924 | 0 | 559 | 567 | 0 | 51.42 | 0.00068 | K | FLPFQQFGR | D | | | 0 |
| trypsin (semi) | 63 | 1::P0DTC2 | 741 | 1 | 440.2275 | 878.4404 | 2 | 878.4399 | 0.0005 | 561 | 567 | 0 | 45.81 | 0.0025 | L | PFQQFGR | D | | | 0 |
| trypsin (semi) | 63 | 1::P0DTC2 | 21818 | 1 | 1286.625 | 2571.2354 | 2 | 2571.2374 | -0.002 | 568 | 590 | 1 | 109.51 | 1.50E-09 | R | DIADTTDAVRDPQTLEILDITPC | S | | | 0 |
| trypsin (semi) | 63 | 1::P0DTC2 | 7129 | 1 | 679.838 | 1357.6614 | 2 | 1357.6626 | -0.0013 | 635 | 646 | 0 | 103.47 | 5.30E-09 | R | VYSTGSNVFQTR | A | | | 0 |
| trypsin (semi) | 63 | 1::P0DTC2 | 3149 | 1 | 548.7728 | 1095.5311 | 2 | 1095.5309 | 0.0002 | 637 | 646 | 0 | 75.16 | 4.50E-06 | Y | STGSNVFQTR | A | | | 0 |
| trypsin (semi) | 63 | 1::P0DTC2 | 5432 | 1 | 626.7988 | 1251.5831 | 2 | 1251.5844 | -0.0013 | 672 | 682 | 0 | 54.4 | 0.00041 | C | ASYQTQTNSPR | R | | | 0 |
| trypsin (semi) | 63 | 1::P0DTC2 | 11524 | 1 | 810.4492 | 1618.8839 | 2 | 1618.8852 | -0.0013 | 719 | 733 | 0 | 85.71 | 2.50E-07 | F | TISVTTEILPVSMTK | T | | | 0 |
| trypsin (semi) | 63 | 1::P0DTC2 | 3812 | 1 | 572.8138 | 1143.613 | 2 | 1143.6135 | -0.0005 | 766 | 776 | 0 | 69.4 | 1.40E-05 | R | ALTGIAVEQDK | N | | | 0 |
| trypsin (semi) | 63 | 1::P0DTC2 | 4046 | 1 | 582.306 | 1162.5975 | 2 | 1162.5982 | -0.0007 | 777 | 786 | 0 | 64.33 | 5.40E-05 | K | NTQEVFAQVK | Q | | | 0 |
| trypsin (semi) | 63 | 1::P0DTC2 | 7464 | 1 | 691.3773 | 1380.7401 | 2 | 1380.7401 | 0 | 815 | 825 | 1 | 65.34 | 3.00E-05 | K | RSFIEDLLFNK | V | | | 0 |
| trypsin (semi) | 63 | 1::P0DTC2 | 5009 | 1 | 613.3266 | 1224.6387 | 2 | 1224.639 | -0.0004 | 816 | 825 | 0 | 67.65 | 1.90E-05 | R | SFIEDLLFNK | V | | | 0 |
| trypsin (semi) | 63 | 1::P0DTC2 | 2389 | 1 | 517.7975 | 1033.5804 | 2 | 1033.5808 | -0.0004 | 826 | 835 | 0 | 74.07 | 3.50E-06 | K | VTLADAGFIK | Q | | | 0 |
| trypsin (semi) | 63 | 1::P0DTC2 | 6743 | 1 | 669.8085 | 1337.6025 | 2 | 1337.6034 | -0.0009 | 836 | 847 | 0 | 89.75 | 7.70E-08 | K | QYGDCLGDIAAR | D | | | 0 |
| trypsin (semi) | 63 | 1::P0DTC2 | 6508 | 1 | 661.2953 | 1320.576 | 2 | 1320.5769 | -0.0008 | 836 | 847 | 0 | 61.77 | 3.20E-05 | K | QYGDCLGDIAAR | D | 2.000000000000.0 | Gln->pyro-Glu (N-term Q) | 0 |
| trypsin (semi) | 63 | 1::P0DTC2 | 650 | 1 | 435.7017 | 869.3888 | 2 | 869.3887 | 0.0001 | 899 | 905 | 0 | 38.03 | 0.006 | F | AMQMAYR | F | | | 0 |
| trypsin (semi) | 63 | 1::P0DTC2 | 14902 | 1 | 912.4672 | 1822.9198 | 2 | 1822.9214 | -0.0016 | 906 | 921 | 0 | 108.44 | 1.90E-09 | R | FNGIGVTQNVLYENQK | L | | | 0 |
| trypsin (semi) | 63 | 1::P0DTC2 | 5803 | 1 | 638.3567 | 1274.6989 | 2 | 1274.6983 | 0.0006 | 922 | 933 | 0 | 113.48 | 4.70E-10 | K | LIANQFNSAIGK | I | | | 0 |
| trypsin (semi) | 63 | 1::P0DTC2 | 7408 | 1 | 689.3645 | 1376.7144 | 2 | 1376.7147 | -0.0003 | 934 | 947 | 0 | 129.97 | 1.70E-11 | K | IQDSLSSTASALGK | L | | | 0 |
| trypsin (semi) | 63 | 1::P0DTC2 | 15517 | 1 | 934.5153 | 1867.0161 | 2 | 1867.0163 | -0.0002 | 948 | 964 | 0 | 135.27 | 2.50E-12 | K | LQDVVNQNAQALNTLVK | Q | | | 0 |
| trypsin (semi) | 63 | 1::P0DTC2 | 17428 | 1 | 1011.0366 | 2020.0586 | 2 | 2020.0589 | -0.0003 | 965 | 983 | 0 | 164.29 | 4.40E-15 | K | QLSSNFGAISSVLNDILSR | L | | | 0 |
| trypsin (semi) | 63 | 1::P0DTC2 | 6907 | 1 | 672.8773 | 1343.7401 | 2 | 1343.7409 | -0.0008 | 971 | 983 | 0 | 94.17 | 4.70E-08 | F | GAISSVLNDILSR | L | | | 0 |
| trypsin (semi) | 63 | 1::P0DTC2 | 8003 | 1 | 707.8803 | 1413.7459 | 2 | 1413.7463 | -0.0004 | 984 | 995 | 1 | 69.96 | 1.20E-05 | R | LDKVEAEVQIDR | L | | | 0 |
| trypsin (semi) | 63 | 1::P0DTC2 | 2663 | 1 | 529.7761 | 1057.5376 | 2 | 1057.5404 | -0.0028 | 987 | 995 | 0 | 70.97 | 9.10E-06 | K | VEAEVQIDR | L | | | 0 |
| trypsin (semi) | 63 | 1::P0DTC2 | 12792 | 1 | 845.9778 | 1689.941 | 2 | 1689.9414 | -0.0004 | 1001 | 1014 | 0 | 143.5 | 3.30E-13 | R | LQSLQTYVTQQLIR | A | | | 0 |
| trypsin (semi) | 63 | 1::P0DTC2 | 3921 | 1 | 577.7623 | 1153.5101 | 2 | 1153.5107 | -0.0006 | 1029 | 1038 | 0 | 75.06 | 1.90E-06 | K | MSECVLGQSK | R | 0.3000000000.0 | Oxidation (M) | 0 |
| trypsin (semi) | 63 | 1::P0DTC2 | 3733 | 1 | 569.7649 | 1137.5153 | 2 | 1137.5158 | -0.0006 | 1029 | 1038 | 0 | 67.73 | 1.30E-05 | K | MSECVLGQSK | R | | | 0 |
| trypsin (semi) | 63 | 1::P0DTC2 | 23776 | 1 | 628.722 | 3138.5735 | 5 | 3138.575 | -0.0015 | 1046 | 1073 | 0 | 46.6 | 0.0028 | K | GYHLMSFPQSAPHGVVFLHVTYVPAQEK | N | | | 0 |
| trypsin (semi) | 63 | 1::P0DTC2 | 21612 | 1 | 846.7769 | 2537.3089 | 3 | 2537.3067 | 0.0022 | 1051 | 1073 | 0 | 114.23 | 4.30E-10 | M | SFPQSAPHGVVFLHVTYVPAQEK | N | | | 0 |
| trypsin (semi) | 63 | 1::P0DTC2 | 5944 | 1 | 642.8511 | 1283.6876 | 2 | 1283.6874 | 0.0002 | 1063 | 1073 | 0 | 78.65 | 1.40E-06 | F | LHVTYVPAQEK | N | | | 0 |
| trypsin (semi) | 63 | 1::P0DTC2 | 4395 | 1 | 593.8252 | 1185.6359 | 2 | 1185.6353 | 0.0006 | 1182 | 1191 | 1 | 48.5 | 0.0017 | K | EIDRLNEVAK | N | | | 0 |
| trypsin (semi) | 63 | 1::P0DTC2 | 6066 | 1 | 647.2956 | 1292.5767 | 2 | 1292.5772 | -0.0005 | 1256 | 1266 | 0 | 57.92 | 0.00012 | K | FDEDDSEPVLK | G | | | 0 |
| trypsin | 48 | 1::P0DTC2 | 9522 | 1 | 748.3814 | 1494.7481 | 2 | 1494.7467 | 0.0015 | 22 | 34 | 0 | 67.44 | 1.20E-06 | R | TQLPPAYTNSFTR | G | | | 0 |
| trypsin | 48 | 1::P0DTC2 | 374 | 1 | 421.2082 | 840.4018 | 2 | 840.4018 | 0.0001 | 35 | 41 | 0 | 43.45 | 0.00017 | R | GVYYPDK | V | | | 0 |
| trypsin | 48 | 1::P0DTC2 | 20198 | 1 | 772.7143 | 2315.1211 | 3 | 2315.1223 | -0.0011 | 78 | 97 | 1 | 111.56 | 8.10E-11 | K | RFDNPVLPFNDGVYFASTEK | S | | | 0 |
| trypsin | 48 | 1::P0DTC2 | 18924 | 1 | 720.6806 | 2159.0199 | 3 | 2159.0211 | -0.0012 | 79 | 97 | 0 | 46.81 | 0.00016 | R | FDNPVLPFNDGVYFASTEK | S | | | 0 |
| trypsin | 48 | 1::P0DTC2 | 4989 | 1 | 612.8164 | 1223.6183 | 2 | 1223.6187 | -0.0003 | 103 | 113 | 0 | 76.46 | 1.50E-07 | R | GWIFGTTLDSK | T | | | 0 |
| trypsin | 48 | 1::P0DTC2 | 20249 | 1 | 775.0149 | 2322.0227 | 3 | 2322.0238 | -0.0011 | 130 | 147 | 0 | 148.12 | 3.90E-15 | K | VCEFQFCNDPFLGVYYHK | N | | | 0 |
| trypsin | 48 | 1::P0DTC2 | 2810 | 1 | 536.2317 | 1070.4488 | 2 | 1070.4491 | -0.0003 | 151 | 158 | 0 | 63.52 | 7.80E-07 | K | SWMESEFR | V | | | 0 |
| trypsin | 48 | 1::P0DTC2 | 536 | 1 | 428.7147 | 855.4149 | 2 | 855.4127 | 0.0022 | 196 | 202 | 0 | 26.24 | 0.009 | K | NIDGYFK | I | | | 0 |
| trypsin | 48 | 1::P0DTC2 | 1423 | 1 | 475.2819 | 948.5492 | 2 | 948.5505 | -0.0013 | 207 | 214 | 0 | 47.41 | 0.00011 | K | HTPINLVR | D | | | 0 |
| trypsin | 48 | 1::P0DTC2 | 3183 | 1 | 549.8246 | 1097.6346 | 2 | 1097.6346 | 0.0001 | 238 | 246 | 0 | 90.07 | 3.20E-09 | R | FQTLLALHR | S | | | 0 |
| trypsin | 48 | 1::P0DTC2 | 22831 | 1 | 1419.6712 | 2837.3278 | 2 | 2837.3297 | -0.0018 | 247 | 273 | 0 | 149.09 | 5.80E-15 | R | SYLTPGDSSSGWTAGAAAYYVGYLQPR | T | | | 0 |
| trypsin | 48 | 1::P0DTC2 | 3006 | 1 | 543.272 | 1084.5295 | 2 | 1084.5302 | -0.0006 | 311 | 319 | 0 | 54.21 | 1.80E-05 | K | GIYQTSNFR | V | | | 0 |
| trypsin | 48 | 1::P0DTC2 | 2322 | 1 | 514.7901 | 1027.5657 | 2 | 1027.5662 | -0.0005 | 320 | 328 | 0 | 42.54 | 0.00038 | R | VQPTESIVR | F | | | 0 |
| trypsin | 48 | 1::P0DTC2 | 3398 | 1 | 557.2765 | 1112.5385 | 2 | 1112.5403 | -0.0018 | 347 | 355 | 0 | 49.37 | 6.10E-05 | R | FASVYAWNR | K | | | 0 |
| trypsin | 48 | 1::P0DTC2 | 21572 | 1 | 843.7408 | 2528.2005 | 3 | 2528.2006 | -0.0001 | 357 | 378 | 1 | 111.49 | 4.60E-11 | K | RISNCVADYSVLYNSASFSTFK | C | | | 0 |
| trypsin | 48 | 1::P0DTC2 | 20617 | 1 | 1187.0566 | 2372.0986 | 2 | 2372.0995 | -0.0008 | 358 | 378 | 0 | 117.04 | 8.00E-12 | R | ISNCVADYSVLYNSASFSTFK | C | | | 0 |
| trypsin | 48 | 1::P0DTC2 | 1017 | 1 | 456.2182 | 910.4218 | 2 | 910.4219 | 0 | 379 | 386 | 0 | 55.09 | 1.30E-05 | K | CYGVSPTK | L | | | 0 |

| | | | | | | | | | | | | | | | | | | | | |
|---|---|---|---|---|---|---|---|---|---|---|---|---|---|---|---|---|---|---|---|---|
| trypsin | 48 | 1::P0DTC2 | 17736 | 1 | 1024.0007 | 2045.9868 | 2 | 2045.9881 | -0.0012 | 387 | 403 | 0 | 135.45 | 2.20E-13 | K | LNDLCFTNVYADSFVIR | G | | | 0 |
| trypsin | 48 | 1::P0DTC2 | 801 | 1 | 443.7189 | 885.4233 | 2 | 885.4232 | 0.0001 | 418 | 424 | 0 | 43.48 | 0.00011 | K | IADYNYK | L | | | 0 |
| trypsin | 48 | 1::P0DTC2 | 19810 | 1 | 1133.5257 | 2265.0368 | 2 | 2265.0372 | -0.0004 | 425 | 444 | 0 | 82.06 | 3.00E-08 | K | LPDDFTGCVIAWNSNNLDSK | V | | | 0 |
| trypsin | 48 | 1::P0DTC2 | 4904 | 1 | 609.7982 | 1217.5819 | 2 | 1217.5829 | -0.001 | 445 | 454 | 0 | 92.57 | 7.20E-09 | K | VGGNYNYLYR | L | | | 0 |
| trypsin | 48 | 1::P0DTC2 | 1876 | 1 | 495.7717 | 989.5288 | 2 | 989.5294 | -0.0006 | 459 | 466 | 1 | 31.65 | 0.0063 | K | SNLKPFER | D | | | 0 |
| trypsin | 48 | 1::P0DTC2 | 17625 | 1 | 1019.063 | 2036.1114 | 2 | 2036.1129 | -0.0014 | 510 | 528 | 0 | 131.79 | 2.90E-13 | R | VVVLSFELLHAPATVCGPK | K | | | 0 |
| trypsin | 48 | 1::P0DTC2 | 19009 | 1 | 1086.5221 | 2171.0296 | 2 | 2171.0317 | -0.0021 | 538 | 557 | 0 | 109.03 | 7.40E-11 | K | CVNFNFNGLTGTGVLTESNK | K | | | 0 |
| trypsin | 48 | 1::P0DTC2 | 19952 | 1 | 1142.0562 | 2282.0978 | 2 | 2282.1002 | -0.0023 | 538 | 558 | 1 | 69.28 | 7.80E-07 | K | CVNFNFNGLTGTGVLTESNKK | F | 1.00000000000000000000.0 | Ammonia-loss (N-term C) | 0 |
| trypsin | 48 | 1::P0DTC2 | 3751 | 1 | 570.3035 | 1138.5924 | 2 | 1138.5924 | 0 | 559 | 567 | 0 | 51.42 | 4.10E-05 | K | FLPFQQFGR | D | | | 0 |
| trypsin | 48 | 1::P0DTC2 | 2886 | 1 | 538.7646 | 1075.5146 | 2 | 1075.5146 | 0.0001 | 568 | 577 | 0 | 33.37 | 0.0032 | R | DIADTTDAVR | D | | | 0 |
| trypsin | 48 | 1::P0DTC2 | 7129 | 1 | 679.838 | 1357.6614 | 2 | 1357.6626 | -0.0013 | 635 | 646 | 0 | 103.47 | 2.30E-10 | R | VYSTGSNVFQTR | A | | | 0 |
| trypsin | 48 | 1::P0DTC2 | 3812 | 1 | 572.8138 | 1143.613 | 2 | 1143.6135 | -0.0005 | 766 | 776 | 0 | 69.4 | 1.00E-06 | R | ALTGIAVEQDK | N | | | 0 |
| trypsin | 48 | 1::P0DTC2 | 4046 | 1 | 582.306 | 1162.5975 | 2 | 1162.5982 | -0.0007 | 777 | 786 | 0 | 64.33 | 3.50E-06 | K | NTQEVFAQVK | Q | | | 0 |
| trypsin | 48 | 1::P0DTC2 | 7464 | 1 | 691.3773 | 1380.7401 | 2 | 1380.7401 | 0 | 815 | 825 | 1 | 65.34 | 2.90E-06 | K | RSFIEDLLFNK | V | | | 0 |
| trypsin | 48 | 1::P0DTC2 | 5009 | 1 | 613.3266 | 1224.6387 | 2 | 1224.639 | -0.0004 | 816 | 825 | 0 | 67.65 | 1.70E-06 | R | SFIEDLLFNK | V | | | 0 |
| trypsin | 48 | 1::P0DTC2 | 2389 | 1 | 517.7975 | 1033.5804 | 2 | 1033.5808 | -0.0004 | 826 | 835 | 0 | 74.07 | 3.10E-07 | K | VTLADAGFIK | Q | | | 0 |
| trypsin | 48 | 1::P0DTC2 | 6743 | 1 | 669.8085 | 1337.6025 | 2 | 1337.6034 | -0.0009 | 836 | 847 | 0 | 89.75 | 4.40E-09 | K | QYGDCLGDIAAR | D | | | 0 |
| trypsin | 48 | 1::P0DTC2 | 6508 | 1 | 661.2953 | 1320.576 | 2 | 1320.5769 | -0.0008 | 836 | 847 | 0 | 61.77 | 1.20E-06 | K | QYGDCLGDIAAR | D | 2.000000000000.0 | Gln->pyro-Glu (N-term Q) | 0 |
| trypsin | 48 | 1::P0DTC2 | 14902 | 1 | 912.4672 | 1822.9198 | 2 | 1822.9214 | -0.0016 | 906 | 921 | 0 | 108.44 | 1.20E-10 | R | FNGIGVTQNVLYENQK | L | | | 0 |
| trypsin | 48 | 1::P0DTC2 | 5803 | 1 | 638.3567 | 1274.6989 | 2 | 1274.6983 | 0.0006 | 922 | 933 | 0 | 113.48 | 3.60E-11 | K | LIANQFNSAIGK | I | | | 0 |
| trypsin | 48 | 1::P0DTC2 | 7408 | 1 | 689.3645 | 1376.7144 | 2 | 1376.7147 | -0.0003 | 934 | 947 | 0 | 129.97 | 1.10E-12 | K | IQDSLSSTASALGK | L | | | 0 |
| trypsin | 48 | 1::P0DTC2 | 15517 | 1 | 934.5153 | 1867.0161 | 2 | 1867.0163 | -0.0002 | 948 | 964 | 0 | 135.27 | 1.60E-13 | K | LQDVVNQNAQALNTLVK | Q | | | 0 |
| trypsin | 48 | 1::P0DTC2 | 17428 | 1 | 1011.0366 | 2020.0586 | 2 | 2020.0589 | -0.0003 | 965 | 983 | 0 | 164.29 | 2.90E-16 | K | QLSSNFGAISSVLNDILSR | L | | | 0 |
| trypsin | 48 | 1::P0DTC2 | 8003 | 1 | 707.8803 | 1413.7459 | 2 | 1413.7463 | -0.0004 | 984 | 995 | 1 | 69.96 | 7.50E-07 | R | LDKVEAEVQIDR | L | | | 0 |
| trypsin | 48 | 1::P0DTC2 | 2663 | 1 | 529.7761 | 1057.5376 | 2 | 1057.5404 | -0.0028 | 987 | 995 | 0 | 70.97 | 5.60E-07 | K | VEAEVQIDR | L | | | 0 |
| trypsin | 48 | 1::P0DTC2 | 12792 | 1 | 845.9778 | 1689.941 | 2 | 1689.9414 | -0.0004 | 1001 | 1014 | 0 | 143.5 | 2.10E-14 | R | LQSLQTYVTQQLIR | A | | | 0 |
| trypsin | 48 | 1::P0DTC2 | 3921 | 1 | 577.7623 | 1153.5101 | 2 | 1153.5107 | -0.0006 | 1029 | 1038 | 0 | 75.06 | 9.00E-08 | K | MSECVLGQSK | R | 0.3000000000.0 | Oxidation (M) | 0 |
| trypsin | 48 | 1::P0DTC2 | 3733 | 1 | 569.7649 | 1137.5153 | 2 | 1137.5158 | -0.0006 | 1029 | 1038 | 0 | 67.73 | 7.00E-07 | K | MSECVLGQSK | R | | | 0 |
| trypsin | 48 | 1::P0DTC2 | 23776 | 1 | 628.722 | 3138.5735 | 5 | 3138.575 | -0.0015 | 1046 | 1073 | 0 | 46.6 | 0.00014 | K | GYHLMSFPQSAPHGVVFLHVTYVPAQEK | N | | | 0 |
| trypsin | 48 | 1::P0DTC2 | 4395 | 1 | 593.8252 | 1185.6359 | 2 | 1185.6353 | 0.0006 | 1182 | 1191 | 1 | 48.5 | 0.00012 | K | EIDRLNEVAK | N | | | 0 |
| trypsin | 48 | 1::P0DTC2 | 6066 | 1 | 647.2956 | 1292.5767 | 2 | 1292.5772 | -0.0005 | 1256 | 1266 | 0 | 57.92 | 4.10E-06 | K | FDEDDSEPVLK | G | | | 0 |

* Results shown in blue are overlapped with those for semi-search.

Supplementary Table S4. Identification of SARS CoV-2 spike protein-derived peptides of alpha-Lytic protease digest

| Enzyme for database search | No. of identified peptides / protein accession | Protein accession | Query No. | Rank | m/z of precursor ion | Calculated mass value of peptide | Charge (z) of precursor ion | Mass value of assigned peptide | Mass difference | Peptide position from | Peptide position to | No. of missed cleavage | Peptide score of Mascot | Expectation value | Amino acid before peptide | Peptide sequence | Amino acid after peptide | Type and position of variable modifications (1: ammonia-loss (N-term C), 2:pyroGlu (N-term Q), 3: oxidation (M)) | Type of variable modifications | No. of potential N-glycosylation site in peptide |
|---|---|---|---|---|---|---|---|---|---|---|---|---|---|---|---|---|---|---|---|---|
| a-Lytic (semi) | 44 | 1::P0DTC2 | 5877 | 1 | 631.3302 | 1260.6458 | 2 | 1260.6462 | -0.0004 | 20 | 30 | 4 | 51.95 | 0.002 | T | TRTQLPPAYTN | S | | | 0 |
| a-Lytic (semi) | 44 | 1::P0DTC2 | 2696 | 1 | 523.7845 | 1045.5544 | 2 | 1045.5556 | -0.0012 | 21 | 29 | 2 | 58.53 | 0.00042 | T | RTQLPPAYT | N | | | 0 |
| a-Lytic (semi) | 44 | 1::P0DTC2 | 6112 | 1 | 425.2239 | 1272.6498 | 3 | 1272.6503 | -0.0004 | 36 | 45 | 2 | 58.35 | 0.00045 | G | VYYPDKVFRS | S | | | 0 |
| a-Lytic (semi) | 44 | 1::P0DTC2 | 6185 | 1 | 639.3538 | 1276.6931 | 2 | 1276.6928 | 0.0003 | 99 | 109 | 1 | 60.36 | 0.00018 | S | NIIRGWIFGTT | L | | | 0 |
| a-Lytic (semi) | 44 | 1::P0DTC2 | 14189 | 1 | 937.434 | 1872.8535 | 2 | 1872.8539 | -0.0004 | 128 | 142 | 1 | 66.37 | 5.00E-05 | V | IKVCEFQFCNDPFLG | V | | | 0 |
| a-Lytic (semi) | 44 | 1::P0DTC2 | 10063 | 1 | 767.3108 | 1532.607 | 2 | 1532.6065 | 0.0006 | 131 | 142 | 0 | 73.08 | 1.90E-06 | V | CEFQFCNDPFLG | V | | | 0 |
| a-Lytic (semi) | 44 | 1::P0DTC2 | 5555 | 1 | 623.333 | 1244.6515 | 2 | 1244.6513 | 0.0002 | 211 | 221 | 1 | 59.04 | 0.00037 | I | NLVRDLPQGFS | A | | | 0 |
| a-Lytic (semi) | 44 | 1::P0DTC2 | 6802 | 1 | 658.8516 | 1315.6886 | 2 | 1315.6884 | 0.0002 | 211 | 222 | 2 | 65.05 | 8.30E-05 | I | NLVRDLPQGFSA | L | | | 0 |
| a-Lytic (semi) | 44 | 1::P0DTC2 | 3679 | 1 | 560.7447 | 1119.4748 | 2 | 1119.4754 | -0.0006 | 290 | 299 | 2 | 82.45 | 6.00E-07 | V | DCALDPLSET | K | | | 0 |
| a-Lytic (semi) | 44 | 1::P0DTC2 | 4264 | 1 | 580.2765 | 1158.5384 | 2 | 1158.538 | 0.0004 | 376 | 385 | 3 | 53.04 | 0.0011 | S | TFKCYGVSPT | K | | | 0 |
| a-Lytic (semi) | 44 | 1::P0DTC2 | 2421 | 1 | 514.2584 | 1026.5023 | 2 | 1026.5022 | 0.0001 | 394 | 402 | 4 | 54.52 | 0.00094 | T | NVYADSFVI | R | | | 0 |
| a-Lytic (semi) | 44 | 1::P0DTC2 | 6717 | 1 | 437.5772 | 1309.7097 | 3 | 1309.7102 | -0.0005 | 402 | 413 | 2 | 47.55 | 0.0032 | V | IRGDEVRQIAPG | Q | | | 0 |
| a-Lytic (semi) | 44 | 1::P0DTC2 | 7898 | 1 | 695.8122 | 1389.6099 | 2 | 1389.6089 | 0.001 | 420 | 430 | 0 | 49.63 | 0.0016 | A | DYNYKLPDDFT | G | | | 0 |
| a-Lytic (semi) | 44 | 1::P0DTC2 | 8750 | 1 | 724.3229 | 1446.6312 | 2 | 1446.6303 | 0.0008 | 420 | 431 | 1 | 61.02 | 0.0001 | A | DYNYKLPDDFTG | C | | | 0 |
| a-Lytic (semi) | 44 | 1::P0DTC2 | 4475 | 1 | 588.7858 | 1175.5569 | 2 | 1175.5571 | -0.0001 | 436 | 445 | 2 | 47.98 | 0.0045 | A | WNSNNLDSKV | G | | | 0 |
| a-Lytic (semi) | 44 | 1::P0DTC2 | 13532 | 1 | 904.9941 | 1807.9736 | 2 | 1807.9733 | 0.0003 | 556 | 570 | 0 | 58.14 | 0.00035 | S | NKKFLPFQQFGRDIA | D | | | 0 |
| a-Lytic (semi) | 44 | 1::P0DTC2 | 8628 | 1 | 719.8779 | 1437.7413 | 2 | 1437.7405 | 0.0008 | 559 | 570 | 0 | 55.38 | 0.00088 | K | FLPFQQFGRDIA | D | | | 0 |
| a-Lytic (semi) | 44 | 1::P0DTC2 | 12906 | 1 | 878.439 | 1754.8634 | 2 | 1754.8628 | 0.0006 | 559 | 573 | 2 | 71.47 | 2.20E-05 | K | FLPFQQFGRDIADTT | D | | | 0 |
| a-Lytic (semi) | 44 | 1::P0DTC2 | 7599 | 1 | 685.3698 | 1368.7251 | 2 | 1368.7249 | 0.0002 | 574 | 585 | 3 | 57.74 | 0.00033 | T | DAVRDPQTLEIL | D | | | 0 |
| a-Lytic (semi) | 44 | 1::P0DTC2 | 12181 | 1 | 849.9491 | 1697.8837 | 2 | 1697.8836 | 0.0001 | 574 | 588 | 3 | 66.57 | 6.00E-05 | T | DAVRDPQTLEILDIT | P | | | 0 |
| a-Lytic (semi) | 44 | 1::P0DTC2 | 2944 | 1 | 533.7799 | 1065.5453 | 2 | 1065.5455 | -0.0001 | 623 | 632 | 3 | 57.34 | 0.0004 | V | AIHADQLTPT | W | | | 0 |
| a-Lytic (semi) | 44 | 1::P0DTC2 | 2065 | 1 | 498.2612 | 994.5078 | 2 | 994.5084 | -0.0006 | 624 | 632 | 2 | 49.88 | 0.002 | A | IHADQLTPT | W | | | 0 |
| a-Lytic (semi) | 44 | 1::P0DTC2 | 12334 | 1 | 855.4824 | 1708.9503 | 2 | 1708.9512 | -0.0009 | 814 | 827 | 2 | 80.27 | 1.40E-06 | S | KRSFIEDLLFNKVT | L | | | 0 |
| a-Lytic (semi) | 44 | 1::P0DTC2 | 10699 | 1 | 527.9592 | 1580.8557 | 3 | 1580.8562 | -0.0005 | 815 | 827 | 2 | 51.33 | 0.0015 | K | RSFIEDLLFNKVT | L | | | 0 |
| a-Lytic (semi) | 44 | 1::P0DTC2 | 5444 | 1 | 619.3452 | 1236.6758 | 2 | 1236.6754 | 0.0004 | 817 | 826 | 0 | 56.44 | 0.00044 | S | FIEDLLFNKV | T | | | 0 |
| a-Lytic (semi) | 44 | 1::P0DTC2 | 7096 | 1 | 669.8691 | 1337.7237 | 2 | 1337.7231 | 0.0006 | 817 | 827 | 1 | 69.3 | 2.00E-05 | S | FIEDLLFNKVT | L | | | 0 |
| a-Lytic (semi) | 44 | 1::P0DTC2 | 11354 | 1 | 814.3929 | 1626.7713 | 2 | 1626.7712 | 0.0001 | 828 | 842 | 2 | 76.32 | 6.80E-06 | T | LADAGFIKQYGDCLG | D | | | 0 |
| a-Lytic (semi) | 44 | 1::P0DTC2 | 8687 | 1 | 722.3326 | 1442.6506 | 2 | 1442.65 | 0.0006 | 830 | 842 | 1 | 50.81 | 0.0016 | A | DAGFIKQYGDCLG | D | | | 0 |
| a-Lytic (semi) | 44 | 1::P0DTC2 | 12735 | 1 | 871.9066 | 1741.7987 | 2 | 1741.7981 | 0.0005 | 830 | 845 | 1 | 94.34 | 8.10E-08 | A | DAGFIKQYGDCLGDIA | A | | | 0 |
| a-Lytic (semi) | 44 | 1::P0DTC2 | 3376 | 1 | 549.2907 | 1096.5669 | 2 | 1096.5665 | 0.0003 | 903 | 912 | 2 | 50.68 | 0.0015 | M | AYRFNGIGVT | Q | | | 0 |
| a-Lytic (semi) | 44 | 1::P0DTC2 | 5617 | 1 | 624.8328 | 1247.651 | 2 | 1247.651 | 0 | 913 | 922 | 1 | 48.73 | 0.0038 | T | QNVLYENQKL | I | | | 0 |
| a-Lytic (semi) | 44 | 1::P0DTC2 | 8543 | 1 | 716.8936 | 1431.7726 | 2 | 1431.7721 | 0.0005 | 913 | 924 | 1 | 54.49 | 0.00079 | T | QNVLYENQKLIA | N | | | 0 |
| a-Lytic (semi) | 44 | 1::P0DTC2 | 8270 | 1 | 708.3802 | 1414.7459 | 2 | 1414.7456 | 0.0003 | 913 | 924 | 1 | 52.84 | 0.0012 | T | QNVLYENQKLIA | N | 2.000000000000.0 | Gln->pyro-Glu (N-term Q) | 0 |
| a-Lytic (semi) | 44 | 1::P0DTC2 | 2492 | 1 | 516.2901 | 1030.5656 | 2 | 1030.5659 | -0.0002 | 930 | 939 | 2 | 51.95 | 0.0017 | S | AIGKIQDSLS | S | | | 0 |
| a-Lytic (semi) | 44 | 1::P0DTC2 | 1425 | 1 | 471.7846 | 941.5546 | 2 | 941.5546 | 0.0001 | 944 | 952 | 2 | 43.12 | 0.0062 | S | ALGKLQDVV | N | | | 0 |
| a-Lytic (semi) | 44 | 1::P0DTC2 | 3187 | 1 | 543.7803 | 1085.5461 | 2 | 1085.5465 | -0.0004 | 953 | 962 | 3 | 48.83 | 0.0029 | V | NQNAQALNTL | V | | | 0 |
| a-Lytic (semi) | 44 | 1::P0DTC2 | 6329 | 1 | 643.851 | 1285.6875 | 2 | 1285.6878 | -0.0003 | 988 | 998 | 2 | 51.12 | 0.0017 | V | EAEVQIDRLIT | G | | | 0 |
| a-Lytic (semi) | 44 | 1::P0DTC2 | 7169 | 1 | 672.3604 | 1342.7063 | 2 | 1342.7092 | -0.003 | 988 | 999 | 3 | 77.48 | 4.50E-06 | V | EAEVQIDRLITG | R | | | 0 |
| a-Lytic (semi) | 44 | 1::P0DTC2 | 3783 | 1 | 563.2879 | 1124.5612 | 2 | 1124.5615 | -0.0003 | 1051 | 1061 | 4 | 52.17 | 0.0016 | M | SFPQSAPHGVV | F | | | 0 |
| a-Lytic (semi) | 44 | 1::P0DTC2 | 8155 | 1 | 704.865 | 1407.7153 | 2 | 1407.7146 | 0.0007 | 1106 | 1116 | 0 | 49.48 | 0.0036 | T | QRNFYEPQIIT | T | | | 0 |
| a-Lytic (semi) | 44 | 1::P0DTC2 | 9091 | 1 | 735.8994 | 1469.7842 | 2 | 1469.7838 | 0.0004 | 1178 | 1189 | 0 | 47.99 | 0.0038 | V | NIQKEIDRLNEV | A | | | 0 |
| a-Lytic (semi) | 44 | 1::P0DTC2 | 10197 | 1 | 771.4181 | 1540.8216 | 2 | 1540.8209 | 0.0008 | 1178 | 1190 | 1 | 63.21 | 0.00012 | V | NIQKEIDRLNEVA | K | | | 0 |
| a-Lytic (semi) | 44 | 1::P0DTC2 | 10763 | 1 | 794.3006 | 1586.5865 | 2 | 1586.5865 | 0 | 1252 | 1264 | 2 | 50.2 | 0.00014 | G | SCCKFDEDDSEPV | L | | | 0 |
| a-Lytic (semi) | 44 | 1::P0DTC2 | 9571 | 1 | 750.7844 | 1499.5542 | 2 | 1499.5545 | -0.0002 | 1253 | 1264 | 1 | 32.58 | 0.007 | S | CCKFDEDDSEPV | L | | | 0 |
| a-Lytic | 30 | 1::P0DTC2 | 4069 | 1 | 574.3089 | 1146.6033 | 2 | 1146.6033 | 0 | 20 | 29 | 3 | 44.36 | 0.0019 | T | TRTQLPPAYT | N | | | 0 |
| a-Lytic | 30 | 1::P0DTC2 | 2696 | 1 | 523.7845 | 1045.5544 | 2 | 1045.5556 | -0.0012 | 21 | 29 | 2 | 58.53 | 6.20E-05 | T | RTQLPPAYT | N | | | 0 |
| a-Lytic | 30 | 1::P0DTC2 | 6185 | 1 | 639.3538 | 1276.6931 | 2 | 1276.6928 | 0.0003 | 99 | 109 | 1 | 60.36 | 2.40E-05 | S | NIIRGWIFGTT | L | | | 0 |
| a-Lytic | 30 | 1::P0DTC2 | 1191 | 1 | 460.2351 | 918.4556 | 2 | 918.4559 | -0.0003 | 214 | 221 | 0 | 42.35 | 0.0022 | V | RDLPQGFS | A | | | 0 |
| a-Lytic | 30 | 1::P0DTC2 | 3679 | 1 | 560.7447 | 1119.4748 | 2 | 1119.4754 | -0.0006 | 290 | 299 | 2 | 82.45 | 8.90E-08 | V | DCALDPLSET | K | | | 0 |
| a-Lytic | 30 | 1::P0DTC2 | 4264 | 1 | 580.2765 | 1158.5384 | 2 | 1158.538 | 0.0004 | 376 | 385 | 3 | 53.04 | 0.00018 | S | TFKCYGVSPT | K | | | 0 |
| a-Lytic | 30 | 1::P0DTC2 | 7898 | 1 | 695.8122 | 1389.6099 | 2 | 1389.6089 | 0.001 | 420 | 430 | 0 | 49.63 | 0.00022 | A | DYNYKLPDDFT | G | | | 0 |
| a-Lytic | 30 | 1::P0DTC2 | 4475 | 1 | 588.7858 | 1175.5569 | 2 | 1175.5571 | -0.0001 | 436 | 445 | 2 | 47.98 | 0.00072 | A | WNSNNLDSKV | G | | | 0 |
| a-Lytic | 30 | 1::P0DTC2 | 12863 | 1 | 584.3055 | 1749.8946 | 3 | 1749.8951 | -0.0005 | 446 | 459 | 0 | 42.63 | 0.0032 | V | GGNYNYLYRLFRKS | N | | | 0 |
| a-Lytic | 30 | 1::P0DTC2 | 6853 | 1 | 660.3513 | 1318.6881 | 2 | 1318.6881 | 0 | 460 | 470 | 1 | 41.47 | 0.0032 | S | NLKPFERDIST | E | | | 0 |
| a-Lytic | 30 | 1::P0DTC2 | 13532 | 1 | 904.9941 | 1807.9736 | 2 | 1807.9733 | 0.0003 | 556 | 570 | 0 | 58.14 | 5.80E-05 | S | NKKFLPFQQFGRDIA | D | | | 0 |
| a-Lytic | 30 | 1::P0DTC2 | 12181 | 1 | 849.9491 | 1697.8837 | 2 | 1697.8836 | 0.0001 | 574 | 588 | 3 | 66.57 | 9.70E-06 | T | DAVRDPQTLEILDIT | P | | | 0 |
| a-Lytic | 30 | 1::P0DTC2 | 2944 | 1 | 533.7799 | 1065.5453 | 2 | 1065.5455 | -0.0001 | 623 | 632 | 3 | 57.34 | 6.10E-05 | V | AIHADQLTPT | W | | | 0 |
| a-Lytic | 30 | 1::P0DTC2 | 2065 | 1 | 498.2612 | 994.5078 | 2 | 994.5084 | -0.0006 | 624 | 632 | 2 | 49.88 | 0.00031 | A | IHADQLTPT | W | | | 0 |
| a-Lytic | 30 | 1::P0DTC2 | 3025 | 1 | 537.7749 | 1073.5353 | 2 | 1073.5353 | 0 | 769 | 778 | 2 | 43.44 | 0.0022 | T | GIAVEQDKNT | Q | | | 0 |
| a-Lytic | 30 | 1::P0DTC2 | 3132 | 1 | 541.297 | 1080.5795 | 2 | 1080.5815 | -0.002 | 804 | 813 | 1 | 40.56 | 0.0019 | S | QILPDPSKPS | K | | | 0 |
| a-Lytic | 30 | 1::P0DTC2 | 12334 | 1 | 855.4824 | 1708.9503 | 2 | 1708.9512 | -0.0009 | 814 | 827 | 2 | 80.27 | 2.00E-07 | S | KRSFIEDLLFNKVT | L | | | 0 |
| a-Lytic | 30 | 1::P0DTC2 | 5444 | 1 | 619.3452 | 1236.6758 | 2 | 1236.6754 | 0.0004 | 817 | 826 | 0 | 56.44 | 6.40E-05 | S | FIEDLLFNKV | T | | | 0 |
| a-Lytic | 30 | 1::P0DTC2 | 7096 | 1 | 669.8691 | 1337.7237 | 2 | 1337.7231 | 0.0006 | 817 | 827 | 1 | 69.3 | 3.60E-06 | S | FIEDLLFNKVT | L | | | 0 |
| a-Lytic | 30 | 1::P0DTC2 | 12735 | 1 | 871.9066 | 1741.7987 | 2 | 1741.7981 | 0.0005 | 830 | 845 | 1 | 94.34 | 1.20E-08 | A | DAGFIKQYGDCLGDIA | A | | | 0 |
| a-Lytic | 30 | 1::P0DTC2 | 8543 | 1 | 716.8936 | 1431.7726 | 2 | 1431.7721 | 0.0005 | 913 | 924 | 1 | 54.49 | 0.00012 | T | QNVLYENQKLIA | N | | | 0 |
| a-Lytic | 30 | 1::P0DTC2 | 8270 | 1 | 708.3802 | 1414.7459 | 2 | 1414.7456 | 0.0003 | 913 | 924 | 1 | 52.84 | 0.00021 | T | QNVLYENQKLIA | N | 2.000000000000.0 | Gln->pyro-Glu (N-term Q) | 0 |
| a-Lytic | 30 | 1::P0DTC2 | 2492 | 1 | 516.2901 | 1030.5656 | 2 | 1030.5659 | -0.0002 | 930 | 939 | 2 | 51.95 | 0.00028 | S | AIGKIQDSLS | S | | | 0 |
| a-Lytic | 30 | 1::P0DTC2 | 1597 | 1 | 480.7717 | 959.5289 | 2 | 959.5287 | 0.0001 | 931 | 939 | 1 | 43.37 | 0.0022 | A | IGKIQDSLS | S | | | 0 |
| a-Lytic | 30 | 1::P0DTC2 | 1425 | 1 | 471.7846 | 941.5546 | 2 | 941.5546 | 0.0001 | 944 | 952 | 2 | 43.12 | 0.001 | S | ALGKLQDVV | N | | | 0 |
| a-Lytic | 30 | 1::P0DTC2 | 6329 | 1 | 643.851 | 1285.6875 | 2 | 1285.6878 | -0.0003 | 988 | 998 | 2 | 51.12 | 0.00028 | V | EAEVQIDRLIT | G | | | 0 |
| a-Lytic | 30 | 1::P0DTC2 | 8155 | 1 | 704.865 | 1407.7153 | 2 | 1407.7146 | 0.0007 | 1106 | 1116 | 0 | 49.48 | 0.00062 | T | QRNFYEPQIIT | T | | | 0 |
| a-Lytic | 30 | 1::P0DTC2 | 9091 | 1 | 735.8994 | 1469.7842 | 2 | 1469.7838 | 0.0004 | 1178 | 1189 | 0 | 47.99 | 0.00059 | V | NIQKEIDRLNEV | A | | | 0 |

| | | | | | | | | | | | | | | | | | | |
|---|---|---|---|---|---|---|---|---|---|---|---|---|---|---|---|---|---|---|
| a-Lytic | 30 | 1::P0DTC2 | 10197 | 1 | 771.4181 | 1540.8216 | 2 | 1540.8209 | 0.0008 | 1178 | 1190 | 1 | 63.21 | 1.90E-05 | V | NIQKEIDRLNEVA | K | 0 |
| a-Lytic | 30 | 1::P0DTC2 | 9571 | 1 | 750.7844 | 1499.5542 | 2 | 1499.5545 | -0.0002 | 1253 | 1264 | 1 | 32.58 | 0.00075 | S | CCKFDEDDSEPV | L | 0 |

* Results shown in blue are overlapped with those for semi-search.

Supplementary Table S5. Identification of SARS CoV-2 spike protein-derived N-glycosylated peptides of chymotrypsin digest

| Enzyme for database search | No. of identified peptides / protein accession | Protein accession | Query No. | Rank | m/z of precursor ion | Calculated mass value of peptide | Charge (z) of precursor ion | Mass value of assigned peptide | Mass difference | Peptide position from | Peptide position to | No. of missed cleavage | Peptide score of Mascot | Expectation value | Amino acid before peptide | Peptide sequence | Amino acid after peptide | Type and position of variable modifications (1: ammonia-loss (N-term C), 2: Delta:H(1)N(-1)18O(1) (N), 3: pyroGlu (N-term Q), 4: oxidation (M)) | Type of variable modifications | No. of deglycosylated site | No. of potential N-glycosylation site | Assigned deglycosylated site (sequon) | Potential site (sequon) | No. of potential N-glycosylation site in peptide |
|---|---|---|---|---|---|---|---|---|---|---|---|---|---|---|---|---|---|---|---|---|---|---|---|---|
| chymotrypsin (semi) | 91 | 1::P0DTC2 | 3889 | 1 | 874.4316 | 1746.8487 | 2 | 1746.8497 | -0.001 | 14 | 28 | 1 | 61.01 | 0.00034 | S | QCVNLTTRTQLPPAY | T | 3.000200000000000.0 | Delta:H(1)N(-1)18O(1) (N); Gln->pyro-Glu (N-term Q) | 1 | 1 | 17(NLT)/ | 17(NLT)/ | 1 |
| chymotrypsin (semi) | 91 | 1::P0DTC2 | 6439 | 1 | 1099.0271 | 2196.0396 | 2 | 2196.0407 | -0.0011 | 14 | 32 | 2 | 60.31 | 0.00046 | S | QCVNLTTRTQLPPAYTNSF | T | 3.0002000000000000000.0 | Delta:H(1)N(-1)18O(1) (N); Gln->pyro-Glu (N-term Q) | 1 | 1 | 17(NLT)/ | 17(NLT)/ | 1 |
| chymotrypsin (semi) | 91 | 1::P0DTC2 | 3427 | 1 | 419.4705 | 1673.8529 | 4 | 1673.8524 | 0.0005 | 65 | 79 | 1 | 92.76 | 0.0000002 | W | FHAIHVSGTNGTKRF | D | 0.000000000200000.0 | Delta:H(1)N(-1)18O(1) (N) | 1 | 1 | 74(NGT)/ | 74(NGT)/ | 1 |
| chymotrypsin (semi) | 91 | 1::P0DTC2 | 2681 | 1 | 509.935 | 1526.7831 | 3 | 1526.784 | -0.0009 | 66 | 79 | 0 | 101.54 | 0.000000023 | F | HAIHVSGTNGTKRF | D | 0.00000000200000.0 | Delta:H(1)N(-1)18O(1) (N) | 1 | 1 | 74(NGT)/ | 74(NGT)/ | 1 |
| chymotrypsin (semi) | 91 | 1::P0DTC2 | 7021 | 1 | 770.7347 | 2309.1824 | 3 | 2309.1803 | 0.0021 | 66 | 86 | 1 | 67.61 | 0.000094 | F | HAIHVSGTNGTKRFDNPVLPF | N | 0.000000002000000000000.0 | Delta:H(1)N(-1)18O(1) (N) | 1 | 1 | 74(NGT)/ | 74(NGT)/ | 1 |
| chymotrypsin (semi) | 91 | 1::P0DTC2 | 9432 | 1 | 953.4749 | 2857.4029 | 3 | 2857.4034 | -0.0004 | 66 | 91 | 2 | 56.03 | 0.0018 | F | HAIHVSGTNGTKRFDNPVLPFNDGVY | F | 0.00000000200000000000000000.0 | Delta:H(1)N(-1)18O(1) (N) | 1 | 1 | 74(NGT)/ | 74(NGT)/ | 1 |
| chymotrypsin (semi) | 91 | 1::P0DTC2 | 2140 | 1 | 464.2505 | 1389.7297 | 3 | 1389.7251 | 0.0046 | 67 | 79 | 0 | 71.45 | 0.000022 | H | AIHVSGTNGTKRF | D | 0.0000000200000.0 | Delta:H(1)N(-1)18O(1) (N) | 1 | 1 | 74(NGT)/ | 74(NGT)/ | 1 |
| chymotrypsin (semi) | 91 | 1::P0DTC2 | 1904 | 1 | 440.57 | 1318.6881 | 3 | 1318.688 | 0.0002 | 68 | 79 | 0 | 61.33 | 0.00022 | A | IHVSGTNGTKRF | D | 0.000000200000.0 | Delta:H(1)N(-1)18O(1) (N) | 1 | 1 | 74(NGT)/ | 74(NGT)/ | 1 |
| chymotrypsin (semi) | 91 | 1::P0DTC2 | 991 | 1 | 535.2795 | 1068.5445 | 2 | 1068.545 | -0.0004 | 70 | 79 | 0 | 68.39 | 0.000033 | H | VSGTNGTKRF | D | 0.0000200000.0 | Delta:H(1)N(-1)18O(1) (N) | 1 | 1 | 74(NGT)/ | 74(NGT)/ | 1 |
| chymotrypsin (semi) | 91 | 1::P0DTC2 | 607 | 1 | 485.7455 | 969.4765 | 2 | 969.4766 | -0.0001 | 71 | 79 | 0 | 55.71 | 0.00059 | V | SGTNGTKRF | D | 0.000200000.0 | Delta:H(1)N(-1)18O(1) (N) | 1 | 1 | 74(NGT)/ | 74(NGT)/ | 1 |
| chymotrypsin (semi) | 91 | 1::P0DTC2 | 5479 | 1 | 1018.5522 | 2035.0898 | 2 | 2035.091 | -0.0011 | 116 | 133 | 2 | 85.53 | 0.00000095 | Q | SLLIVNNATNVVIKVCEF | Q | 0.000000200000000000.0 | Delta:H(1)N(-1)18O(1) (N) | 1 | 1 | 122(NAT)/ | 122(NAT)/ | 1 |
| chymotrypsin (semi) | 91 | 1::P0DTC2 | 4426 | 1 | 918.4949 | 1834.9752 | 2 | 1834.9749 | 0.0003 | 118 | 133 | 1 | 109.08 | 4.1E-09 | L | LIVNNATNVVIKVCEF | Q | 0.0000200000000000.0 | Delta:H(1)N(-1)18O(1) (N) | 1 | 1 | 122(NAT)/ | 122(NAT)/ | 1 |
| chymotrypsin (semi) | 91 | 1::P0DTC2 | 1461 | 1 | 594.3475 | 1186.6804 | 2 | 1186.6807 | -0.0003 | 119 | 129 | 0 | 81.89 | 0.00000079 | L | IVNNATNVVIK | V | 0.00020000000.0 | Delta:H(1)N(-1)18O(1) (N) | 1 | 1 | 122(NAT)/ | 122(NAT)/ | 1 |
| chymotrypsin (semi) | 91 | 1::P0DTC2 | 3700 | 1 | 861.9517 | 1721.8889 | 2 | 1721.8908 | -0.0019 | 119 | 133 | 0 | 100.49 | 0.000000037 | L | IVNNATNVVIKVCEF | Q | 0.000200000000000.0 | Delta:H(1)N(-1)18O(1) (N) | 1 | 1 | 122(NAT)/ | 122(NAT)/ | 1 |
| chymotrypsin (semi) | 91 | 1::P0DTC2 | 3552 | 1 | 851.8747 | 1701.7349 | 2 | 1701.7343 | 0.0007 | 145 | 157 | 2 | 61.72 | 0.00014 | Y | YHKNNKSWMESEF | R | 0.0000200000000.0 | Delta:H(1)N(-1)18O(1) (N) | 1 | 1 | 149(NKS)/ | 149(NKS)/ | 1 |
| chymotrypsin (semi) | 91 | 1::P0DTC2 | 3632 | 1 | 573.5834 | 1717.7284 | 3 | 1717.7292 | -0.0008 | 145 | 157 | 2 | 56.79 | 0.00032 | Y | YHKNNKSWMESEF | R | 0.0000200040000.0 | Delta:H(1)N(-1)18O(1) (N); Oxidation (M) | 1 | 1 | 149(NKS)/ | 149(NKS)/ | 1 |
| chymotrypsin (semi) | 91 | 1::P0DTC2 | 2742 | 1 | 513.8975 | 1538.6705 | 3 | 1538.6709 | -0.0004 | 146 | 157 | 1 | 38.24 | 0.031 | Y | HKNNKSWMESEF | R | 0.000200000000.0 | Delta:H(1)N(-1)18O(1) (N) | 1 | 1 | 149(NKS)/ | 149(NKS)/ | 1 |
| chymotrypsin (semi) | 91 | 1::P0DTC2 | 1477 | 1 | 598.2264 | 1194.4383 | 2 | 1194.4385 | -0.0002 | 161 | 170 | 1 | 27.65 | 0.047 | Y | SSANNCTFEY | V | 0.0000200000.0 | Delta:H(1)N(-1)18O(1) (N) | 1 | 2 | 165(NCT)/ | 164(NNC)/165(NCT)/ | 1 |
| chymotrypsin (semi) | 91 | 1::P0DTC2 | 3394 | 1 | 834.4717 | 1666.9289 | 2 | 1666.9279 | 0.001 | 221 | 236 | 2 | 61.93 | 0.00011 | F | SALEPLVDLPIGINIT | R | 0.0000000000000200.0 | Delta:H(1)N(-1)18O(1) (N) | 1 | 1 | 234(NIT)/ | 234(NIT)/ | 1 |
| chymotrypsin (semi) | 91 | 1::P0DTC2 | 4337 | 1 | 912.5222 | 1823.0299 | 2 | 1823.029 | 0.0009 | 221 | 237 | 2 | 42.51 | 0.008 | F | SALEPLVDLPIGINITR | F | 0.00000000000002000.0 | Delta:H(1)N(-1)18O(1) (N) | 1 | 1 | 234(NIT)/ | 234(NIT)/ | 1 |
| chymotrypsin (semi) | 91 | 1::P0DTC2 | 5156 | 1 | 986.055 | 1970.0955 | 2 | 1970.0974 | -0.0019 | 221 | 238 | 2 | 105.42 | 5.9E-09 | F | SALEPLVDLPIGINITRF | Q | 0.000000000000020000.0 | Delta:H(1)N(-1)18O(1) (N) | 1 | 1 | 234(NIT)/ | 234(NIT)/ | 1 |
| chymotrypsin (semi) | 91 | 1::P0DTC2 | 5805 | 1 | 1050.0854 | 2098.1562 | 2 | 2098.156 | 0.0003 | 221 | 239 | 3 | 49.38 | 0.0028 | F | SALEPLVDLPIGINITRFQ | T | 0.0000000000000200000.0 | Delta:H(1)N(-1)18O(1) (N) | 1 | 1 | 234(NIT)/ | 234(NIT)/ | 1 |
| chymotrypsin (semi) | 91 | 1::P0DTC2 | 4701 | 1 | 942.5401 | 1883.0656 | 2 | 1883.0654 | 0.0002 | 222 | 238 | 2 | 62.49 | 0.000081 | S | ALEPLVDLPIGINITRF | Q | 0.00000000000020000.0 | Delta:H(1)N(-1)18O(1) (N) | 1 | 1 | 234(NIT)/ | 234(NIT)/ | 1 |
| chymotrypsin (semi) | 91 | 1::P0DTC2 | 2031 | 1 | 680.8905 | 1359.7664 | 2 | 1359.7648 | 0.0016 | 227 | 238 | 0 | 58.87 | 0.00019 | L | VDLPIGINITRF | Q | 0.000000020000.0 | Delta:H(1)N(-1)18O(1) (N) | 1 | 1 | 234(NIT)/ | 234(NIT)/ | 1 |
| chymotrypsin (semi) | 91 | 1::P0DTC2 | 4362 | 1 | 914.9299 | 1827.8452 | 2 | 1827.8446 | 0.0006 | 276 | 291 | 3 | 41.37 | 0.026 | F | LLKYNENGTITDAVDC | A | 0.0000002000000000.0 | Delta:H(1)N(-1)18O(1) (N) | 1 | 1 | 282(NGT)/ | 282(NGT)/ | 1 |
| chymotrypsin (semi) | 91 | 1::P0DTC2 | 9207 | 1 | 928.4565 | 2782.3477 | 3 | 2782.3468 | 0.0009 | 276 | 300 | 5 | 79.96 | 0.0000069 | F | LLKYNENGTITDAVDCALDPLSETK | C | 0.0000002000000000000000000.0 | Delta:H(1)N(-1)18O(1) (N) | 1 | 1 | 282(NGT)/ | 282(NGT)/ | 1 |
| chymotrypsin (semi) | 91 | 1::P0DTC2 | 9696 | 1 | 981.7996 | 2942.377 | 3 | 2942.3775 | -0.0005 | 276 | 301 | 5 | 84.99 | 0.0000018 | F | LLKYNENGTITDAVDCALDPLSETKC | T | 0.00000020000000000000000000.0 | Delta:H(1)N(-1)18O(1) (N) | 1 | 1 | 282(NGT)/ | 282(NGT)/ | 1 |
| chymotrypsin (semi) | 91 | 1::P0DTC2 | 10421 | 1 | 1053.1767 | 3156.5083 | 3 | 3156.5092 | -0.001 | 276 | 303 | 5 | 87.99 | 0.000001 | F | LLKYNENGTITDAVDCALDPLSETKCTL | K | 0.0000002000000000000000000000.0 | Delta:H(1)N(-1)18O(1) (N) | 1 | 1 | 282(NGT)/ | 282(NGT)/ | 1 |
| chymotrypsin (semi) | 91 | 1::P0DTC2 | 10972 | 1 | 822.1583 | 3284.6041 | 4 | 3284.6042 | -0.0001 | 276 | 304 | 6 | 70.98 | 0.0000058 | F | LLKYNENGTITDAVDCALDPLSETKCTLK | S | 0.00000020000000000000000000000.0 | Delta:H(1)N(-1)18O(1) (N) | 1 | 1 | 282(NGT)/ | 282(NGT)/ | 1 |
| chymotrypsin (semi) | 91 | 1::P0DTC2 | 11858 | 1 | 880.6837 | 3518.7056 | 4 | 3518.7046 | 0.0009 | 276 | 306 | 6 | 52.49 | 0.0042 | F | LLKYNENGTITDAVDCALDPLSETKCTLKSI | T | 0.0000002000000000000000000000000.0 | Delta:H(1)N(-1)18O(1) (N) | 1 | 1 | 282(NGT)/ | 282(NGT)/ | 1 |
| chymotrypsin (semi) | 91 | 1::P0DTC2 | 8781 | 1 | 890.7615 | 2669.2628 | 3 | 2669.2628 | 0 | 277 | 300 | 4 | 73.36 | 0.000027 | L | LKYNENGTITDAVDCALDPLSETK | C | 0.000002000000000000000000.0 | Delta:H(1)N(-1)18O(1) (N) | 1 | 1 | 282(NGT)/ | 282(NGT)/ | 1 |
| chymotrypsin (semi) | 91 | 1::P0DTC2 | 9362 | 1 | 944.1049 | 2829.293 | 3 | 2829.2934 | -0.0005 | 277 | 301 | 4 | 102.78 | 0.000000023 | L | LKYNENGTITDAVDCALDPLSETKC | T | 0.0000020000000000000000000.0 | Delta:H(1)N(-1)18O(1) (N) | 1 | 1 | 282(NGT)/ | 282(NGT)/ | 1 |
| chymotrypsin (semi) | 91 | 1::P0DTC2 | 10004 | 1 | 1015.4822 | 3043.4248 | 3 | 3043.4252 | -0.0004 | 277 | 303 | 4 | 85.68 | 0.0000016 | L | LKYNENGTITDAVDCALDPLSETKCTL | K | 0.000002000000000000000000000.0 | Delta:H(1)N(-1)18O(1) (N) | 1 | 1 | 282(NGT)/ | 282(NGT)/ | 1 |
| chymotrypsin (semi) | 91 | 1::P0DTC2 | 11515 | 1 | 852.4124 | 3405.6207 | 4 | 3405.6206 | 0.0001 | 277 | 306 | 5 | 53.75 | 0.003 | L | LKYNENGTITDAVDCALDPLSETKCTLKSF | T | 0.000002000000000000000000000000.0 | Delta:H(1)N(-1)18O(1) (N) | 1 | 1 | 282(NGT)/ | 282(NGT)/ | 1 |
| chymotrypsin (semi) | 91 | 1::P0DTC2 | 8287 | 1 | 853.0668 | 2556.1785 | 3 | 2556.1787 | -0.0002 | 278 | 300 | 3 | 63.9 | 0.00019 | L | KYNENGTITDAVDCALDPLSETK | C | 0.00002000000000000000000.0 | Delta:H(1)N(-1)18O(1) (N) | 1 | 1 | 282(NGT)/ | 282(NGT)/ | 1 |
| chymotrypsin (semi) | 91 | 1::P0DTC2 | 8966 | 1 | 906.4097 | 2716.2073 | 3 | 2716.2094 | -0.002 | 278 | 301 | 3 | 71.19 | 0.000023 | L | KYNENGTITDAVDCALDPLSETKC | T | 0.000020000000000000000000.0 | Delta:H(1)N(-1)18O(1) (N) | 1 | 1 | 282(NGT)/ | 282(NGT)/ | 1 |
| chymotrypsin (semi) | 91 | 1::P0DTC2 | 9657 | 1 | 977.7878 | 2930.3415 | 3 | 2930.3411 | 0.0003 | 278 | 303 | 3 | 79.82 | 0.0000047 | L | KYNENGTITDAVDCALDPLSETKCTL | K | 0.00002000000000000000000000.0 | Delta:H(1)N(-1)18O(1) (N) | 1 | 1 | 282(NGT)/ | 282(NGT)/ | 1 |
| chymotrypsin (semi) | 91 | 1::P0DTC2 | 11011 | 1 | 824.1413 | 3292.5362 | 4 | 3292.5365 | -0.0003 | 278 | 306 | 4 | 59.48 | 0.00066 | L | KYNENGTITDAVDCALDPLSETKCTLKSF | T | 0.00002000000000000000000000000.0 | Delta:H(1)N(-1)18O(1) (N) | 1 | 1 | 282(NGT)/ | 282(NGT)/ | 1 |
| chymotrypsin (semi) | 91 | 1::P0DTC2 | 7603 | 1 | 1213.532 | 2425.0494 | 2 | 2425.0511 | -0.0016 | 280 | 301 | 2 | 70.61 | 0.000019 | Y | NENGTITDAVDCALDPLSETKC | T | 0.0020000000000000000000.0 | Delta:H(1)N(-1)18O(1) (N) | 1 | 1 | 282(NGT)/ | 282(NGT)/ | 1 |
| chymotrypsin (semi) | 91 | 1::P0DTC2 | 8653 | 1 | 880.7351 | 2639.1834 | 3 | 2639.1828 | 0.0006 | 280 | 303 | 2 | 60.45 | 0.0003 | Y | NENGTITDAVDCALDPLSETKCTL | K | 0.002000000000000000000000.0 | Delta:H(1)N(-1)18O(1) (N) | 1 | 1 | 282(NGT)/ | 282(NGT)/ | 1 |
| chymotrypsin (semi) | 91 | 1::P0DTC2 | 5243 | 1 | 663.0335 | 1986.0786 | 3 | 1986.0784 | 0.0002 | 319 | 335 | 0 | 93.46 | 0.00000013 | F | RVQPTESIVRFPNITNL | C | 0.00000000000020000.0 | Delta:H(1)N(-1)18O(1) (N) | 1 | 2 | 331(NIT)/ | 331(NIT)/334(NLC)/ | 1 |
| chymotrypsin (semi) | 91 | 1::P0DTC2 | 11545 | 1 | 854.6863 | 3414.7162 | 4 | 3414.7167 | -0.0004 | 319 | 347 | 3 | 63.16 | 0.00038 | F | RVQPTESIVRFPNITNLCPFGEVFNATRF | A | 0.00000000000020000000000020000.0 | 2 Delta:H(1)N(-1)18O(1) (N) | 2 | 3 | 331(NIT)/343(NAT)/ | 331(NIT)/334(NLC)/343(NAT)/ | 2 |
| chymotrypsin (semi) | 91 | 1::P0DTC2 | 4368 | 1 | 915.9955 | 1829.9764 | 2 | 1829.9773 | -0.0009 | 320 | 335 | 0 | 39.74 | 0.034 | R | VQPTESIVRFPNITNL | C | 0.0000000000020000.0 | Delta:H(1)N(-1)18O(1) (N) | 1 | 2 | 331(NIT)/ | 331(NIT)/334(NLC)/ | 1 |
| chymotrypsin (semi) | 91 | 1::P0DTC2 | 2183 | 1 | 703.3824 | 1404.7502 | 2 | 1404.7499 | 0.0004 | 324 | 335 | 0 | 57.06 | 0.00051 | T | ESIVRFPNITNL | C | 0.000000020000.0 | Delta:H(1)N(-1)18O(1) (N) | 1 | 2 | 331(NIT)/ | 331(NIT)/334(NLC)/ | 1 |
| chymotrypsin (semi) | 91 | 1::P0DTC2 | 4261 | 1 | 905.4578 | 1808.9011 | 2 | 1808.9017 | -0.0006 | 324 | 338 | 1 | 62.56 | 0.00026 | T | ESIVRFPNITNLCPF | G | 0.000000020000000.0 | Delta:H(1)N(-1)18O(1) (N) | 1 | 2 | 331(NIT)/ | 331(NIT)/334(NLC)/ | 1 |
| chymotrypsin (semi) | 91 | 1::P0DTC2 | 3031 | 1 | 797.4209 | 1592.8273 | 2 | 1592.8271 | 0.0003 | 326 | 338 | 1 | 47.77 | 0.0063 | S | IVRFPNITNLCPF | G | 0.0000020000000.0 | Delta:H(1)N(-1)18O(1) (N) | 1 | 2 | 331(NIT)/ | 331(NIT)/334(NLC)/ | 1 |
| chymotrypsin (semi) | 91 | 1::P0DTC2 | 5430 | 1 | 1013.521 | 2025.0274 | 2 | 2025.028 | -0.0005 | 326 | 342 | 2 | 67.6 | 0.000086 | S | IVRFPNITNLCPFGEVF | N | 0.00000200000000000.0 | Delta:H(1)N(-1)18O(1) (N) | 1 | 2 | 331(NIT)/ | 331(NIT)/334(NLC)/ | 1 |
| chymotrypsin (semi) | 91 | 1::P0DTC2 | 4278 | 1 | 907.444 | 1812.8734 | 2 | 1812.8755 | -0.0021 | 328 | 342 | 2 | 57.63 | 0.00073 | V | RFPNITNLCPFGEVF | N | 0.000200000000000.0 | Delta:H(1)N(-1)18O(1) (N) | 1 | 2 | 331(NIT)/ | 331(NIT)/334(NLC)/ | 1 |
| chymotrypsin (semi) | 91 | 1::P0DTC2 | 2352 | 1 | 724.3313 | 1446.648 | 2 | 1446.6488 | -0.0008 | 336 | 347 | 2 | 80.02 | 0.0000023 | L | CPFGEVFNATRF | A | 0.000000020000.0 | Delta:H(1)N(-1)18O(1) (N) | 1 | 1 | 343(NAT)/ | 343(NAT)/ | 1 |
| chymotrypsin (semi) | 91 | 1::P0DTC2 | 2270 | 1 | 715.8185 | 1429.6224 | 2 | 1429.6223 | 0.0002 | 336 | 347 | 2 | 76.55 | 0.0000043 | L | CPFGEVFNATRF | A | 1.000000020000.0 | Ammonia-loss (N-term C); Delta:H(1)N(-1)18O(1) (N) | 1 | 1 | 343(NAT)/ | 343(NAT)/ | 1 |
| chymotrypsin (semi) | 91 | 1::P0DTC2 | 889 | 1 | 522.2556 | 1042.4966 | 2 | 1042.497 | -0.0003 | 339 | 347 | 1 | 52.34 | 0.0012 | F | GEVFNATRF | A | 0.000020000.0 | Delta:H(1)N(-1)18O(1) (N) | 1 | 1 | 343(NAT)/ | 343(NAT)/ | 1 |
| chymotrypsin (semi) | 91 | 1::P0DTC2 | 3276 | 1 | 824.3823 | 1646.75 | 2 | 1646.7496 | 0.0004 | 612 | 625 | 1 | 80.9 | 0.0000024 | L | YQDVNCTEVPVAIH | A | 0.00002000000000.0 | Delta:H(1)N(-1)18O(1) (N) | 1 | 1 | 616(NCT)/ | 616(NCT)/ | 1 |
| chymotrypsin (semi) | 91 | 1::P0DTC2 | 8300 | 1 | 1280.5983 | 2559.182 | 2 | 2559.1838 | -0.0017 | 612 | 633 | 2 | 53.38 | 0.0022 | L | YQDVNCTEVPVAIHADQLTPTW | R | 0.0000200000000000000000.0 | Delta:H(1)N(-1)18O(1) (N) | 1 | 1 | 616(NCT)/ | 616(NCT)/ | 1 |
| chymotrypsin (semi) | 91 | 1::P0DTC2 | 9793 | 1 | 993.4798 | 2977.4175 | 3 | 2977.4166 | 0.0009 | 612 | 636 | 3 | 98.99 | 0.000000084 | L | YQDVNCTEVPVAIHADQLTPTWRVY | S | 0.0000200000000000000000000.0 | Delta:H(1)N(-1)18O(1) (N) | 1 | 1 | 616(NCT)/ | 616(NCT)/ | 1 |
| chymotrypsin (semi) | 91 | 1::P0DTC2 | 2459 | 1 | 742.8504 | 1483.6862 | 2 | 1483.6863 | -0.0001 | 613 | 625 | 0 | 89.75 | 0.0000031 | Y | QDVNCTEVPVAIH | A | 0.0002000000000.0 | Delta:H(1)N(-1)18O(1) (N) | 1 | 1 | 616(NCT)/ | 616(NCT)/ | 1 |
| chymotrypsin (semi) | 91 | 1::P0DTC2 | 4846 | 1 | 956.4539 | 1910.8932 | 2 | 1910.893 | 0.0002 | 613 | 629 | 0 | 46.68 | 0.0085 | Y | QDVNCTEVPVAIHADQL | T | 0.00020000000000000.0 | Delta:H(1)N(-1)18O(1) (N) | 1 | 1 | 616(NCT)/ | 616(NCT)/ | 1 |
| chymotrypsin (semi) | 91 | 1::P0DTC2 | 7443 | 1 | 1199.0671 | 2396.1196 | 2 | 2396.1205 | -0.0008 | 613 | 633 | 1 | 62.19 | 0.0003 | Y | QDVNCTEVPVAIHADQLTPTW | R | 0.000200000000000000000.0 | Delta:H(1)N(-1)18O(1) (N) | 1 | 1 | 616(NCT)/ | 616(NCT)/ | 1 |
| chymotrypsin (semi) | 91 | 1::P0DTC2 | 7351 | 1 | 1190.5537 | 2379.0928 | 2 | 2379.0939 | -0.0011 | 613 | 633 | 1 | 52.48 | 0.0023 | Y | QDVNCTEVPVAIHADQLTPTW | R | 3.000200000000000000000.0 | Delta:H(1)N(-1)18O(1) (N); Gln->pyro-Glu (N-term Q) | 1 | 1 | 616(NCT)/ | 616(NCT)/ | 1 |
| chymotrypsin (semi) | 91 | 1::P0DTC2 | 8248 | 1 | 851.7478 | 2552.2215 | 3 | 2552.2216 | -0.0001 | 613 | 634 | 2 | 86.79 | 0.0000013 | Y | QDVNCTEVPVAIHADQLTPTWR | V | 0.0002000000000000000000.0 | Delta:H(1)N(-1)18O(1) (N) | 1 | 1 | 616(NCT)/ | 616(NCT)/ | 1 |
| chymotrypsin (semi) | 91 | 1::P0DTC2 | 9314 | 1 | 939.1251 | 2814.3534 | 3 | 2814.3533 | 0.0001 | 613 | 636 | 2 | 83.06 | 0.0000033 | Y | QDVNCTEVPVAIHADQLTPTWRVY | S | 0.000200000000000000000000.0 | Delta:H(1)N(-1)18O(1) (N) | 1 | 1 | 616(NCT)/ | 616(NCT)/ | 1 |
| chymotrypsin (semi) | 91 | 1::P0DTC2 | 10176 | 1 | 1026.8025 | 3077.3857 | 3 | 3077.3891 | -0.0034 | 644 | 671 | 2 | 58.63 | 0.00051 | F | QTRAGCLIGAEHVNNSYECDIPIGAGIC | A | 0.0000000000000020000000000000.0 | Delta:H(1)N(-1)18O(1) (N) | 1 | 1 | 657(NNS)/ | 657(NNS)/ | 1 |
| chymotrypsin (semi) | 91 | 1::P0DTC2 | 11345 | 1 | 1128.1723 | 3381.4951 | 3 | 3381.495 | 0.0001 | 644 | 674 | 2 | 48.98 | 0.0034 | F | QTRAGCLIGAEHVNNSYECDIPIGAGICASY | Q | 3.0000000000000020000000000000000.0 | Delta:H(1)N(-1)18O(1) (N); Gln->pyro-Glu (N-term Q) | 1 | 1 | 657(NNS)/ | 657(NNS)/ | 1 |
| chymotrypsin (semi) | 91 | 1::P0DTC2 | 9926 | 1 | 1005.4448 | 3013.3126 | 3 | 3013.3142 | -0.0016 | 647 | 674 | 2 | 53.68 | 0.00095 | R | AGCLIGAEHVNNSYECDIPIGAGICASY | Q | 0.0000000000020000000000000000.0 | Delta:H(1)N(-1)18O(1) (N) | 1 | 1 | 657(NNS)/ | 657(NNS)/ | 1 |
| chymotrypsin (semi) | 91 | 1::P0DTC2 | 9010 | 1 | 909.4153 | 2725.2242 | 3 | 2725.2249 | -0.0008 | 650 | 674 | 2 | 65.33 | 0.0001 | C | LIGAEHVNNSYECDIPIGAGICASY | Q | 0.0000000020000000000000000.0 | Delta:H(1)N(-1)18O(1) (N) | 1 | 1 | 657(NNS)/ | 657(NNS)/ | 1 |
| chymotrypsin (semi) | 91 | 1::P0DTC2 | 8527 | 1 | 1307.0768 | 2612.139 | 2 | 2612.1409 | -0.0018 | 651 | 674 | 1 | 61.19 | 0.00016 | L | IGAEHVNNSYECDIPIGAGICASY | Q | 0.000000200000000000000000.0 | Delta:H(1)N(-1)18O(1) (N) | 1 | 1 | 657(NNS)/ | 657(NNS)/ | 1 |
| chymotrypsin (semi) | 91 | 1::P0DTC2 | 1452 | 1 | 592.2843 | 1182.554 | 2 | 1182.554 | 0 | 708 | 718 | 0 | 46.66 | 0.0059 | Y | SNNSIAIPTNF | T | 0.02000000020.0 | 2 Delta:H(1)N(-1)18O(1) (N) | 2 | 2 | 709(NNS)/717(NFT)/ | 709(NNS)/717(NFT)/ | 2 |
| chymotrypsin (semi) | 91 | 1::P0DTC2 | 3506 | 1 | 847.4195 | 1692.8245 | 2 | 1692.8245 | 0 | 798 | 813 | 2 | 90.9 | 0.00000035 | F | GGFNFSQILPDPSKPS | K | 0.0002000000000000.0 | Delta:H(1)N(-1)18O(1) (N) | 1 | 1 | 801(NFS)/ | 801(NFS)/ | 1 |
| chymotrypsin (semi) | 91 | 1::P0DTC2 | 2281 | 1 | 716.8637 | 1431.7128 | 2 | 1431.7132 | -0.0004 | 801 | 813 | 1 | 50.95 | 0.0028 | F | NFSQILPDPSKPS | K | 0.2000000000000.0 | Delta:H(1)N(-1)18O(1) (N) | 1 | 1 | 801(NFS)/ | 801(NFS)/ | 1 |
| chymotrypsin (semi) | 91 | 1::P0DTC2 | 3620 | 1 | 572.9768 | 1715.9085 | 3 | 1715.9092 | -0.0007 | 801 | 815 | 1 | 89.81 | 0.0000032 | F | NFSQILPDPSKPSKR | S | 0.200000000000000.0 | Delta:H(1)N(-1)18O(1) (N) | 1 | 1 | 801(NFS)/ | 801(NFS)/ | 1 |
| chymotrypsin (semi) | 91 | 1::P0DTC2 | 2793 | 1 | 774.9008 | 1547.7871 | 2 | 1547.787 | 0.0001 | 1063 | 1075 | 2 | 66.71 | 0.00008 | F | LHVTYVPAQEKNF | T | 0.0000000000020.0 | Delta:H(1)N(-1)18O(1) (N) | 1 | 1 | 1074(NFT)/ | 1074(NFT)/ | 1 |
| chymotrypsin (semi) | 91 | 1::P0DTC2 | 7474 | 1 | 800.7347 | 2399.1823 | 3 | 2399.183 | -0.0006 | 1063 | 1083 | 3 | 91.83 | 0.00000041 | F | LHVTYVPAQEKNFTTAPAICH | D | 0.000000000002000000000.0 | Delta:H(1)N(-1)18O(1) (N) | 1 | 1 | 1074(NFT)/ | 1074(NFT)/ | 1 |
| chymotrypsin (semi) | 91 | 1::P0DTC2 | 12279 | 1 | 748.9763 | 3739.8451 | 5 | 3739.8456 | -0.0004 | 1063 | 1095 | 3 | 56.92 | 0.0017 | F | LHVTYVPAQEKNFTTAPAICHDGKAHFPRE( | V | 0.000000000002000000000000000000000.0 | Delta:H(1)N(-1)18O(1) (N) | 1 | 1 | 1074(NFT)/ | 1074(NFT)/ | 1 |
| chymotrypsin (semi) | 91 | 1::P0DTC2 | 6891 | 1 | 763.0402 | 2286.0988 | 3 | 2286.0989 | -0.0001 | 1064 | 1083 | 2 | 95.77 | 0.00000015 | L | HVTYVPAQEKNFTTAPAICH | D | 0.00000000002000000000.0 | Delta:H(1)N(-1)18O(1) (N) | 1 | 1 | 1074(NFT)/ | 1074(NFT)/ | 1 |
| chymotrypsin (semi) | 91 | 1::P0DTC2 | 453 | 1 | 468.2394 | 934.4643 | 2 | 934.4646 | -0.0002 | 1068 | 1075 | 0 | 45.06 | 0.0081 | Y | VPAQEKNF | T | 0.00000020.0 | Delta:H(1)N(-1)18O(1) (N) | 1 | 1 | 1074(NFT)/ | 1074(NFT)/ | 1 |
| chymotrypsin (semi) | 91 | 1::P0DTC2 | 2097 | 1 | 688.8507 | 1375.6869 | 2 | 1375.6869 | 0 | 1068 | 1080 | 1 | 51.06 | 0.0024 | Y | VPAQEKNFTTAPA | I | 0.0000002000000.0 | Delta:H(1)N(-1)18O(1) (N) | 1 | 1 | 1074(NFT)/ | 1074(NFT)/ | 1 |
| chymotrypsin (semi) | 91 | 1::P0DTC2 | 3289 | 1 | 825.4084 | 1648.8022 | 2 | 1648.8017 | 0.0006 | 1068 | 1082 | 1 | 61.26 | 0.00031 | Y | VPAQEKNFTTAPAIC | H | 0.000000200000000.0 | Delta:H(1)N(-1)18O(1) (N) | 1 | 1 | 1074(NFT)/ | 1074(NFT)/ | 1 |
| chymotrypsin (semi) | 91 | 1::P0DTC2 | 4103 | 1 | 893.9375 | 1785.8604 | 2 | 1785.8606 | -0.0002 | 1068 | 1083 | 1 | 82.81 | 0.0000022 | Y | VPAQEKNFTTAPAICH | D | 0.0000002000000000.0 | Delta:H(1)N(-1)18O(1) (N) | 1 | 1 | 1074(NFT)/ | 1074(NFT)/ | 1 |
| chymotrypsin (semi) | 91 | 1::P0DTC2 | 5727 | 1 | 696.3421 | 2086.0044 | 3 | 2086.0039 | 0.0005 | 1068 | 1086 | 1 | 61.88 | 0.00032 | Y | VPAQEKNFTTAPAICHDGK | A | 0.0000002000000000000.0 | Delta:H(1)N(-1)18O(1) (N) | 1 | 1 | 1074(NFT)/ | 1074(NFT)/ | 1 |
| chymotrypsin (semi) | 91 | 1::P0DTC2 | 8895 | 1 | 674.5876 | 2694.3213 | 4 | 2694.3223 | -0.001 | 1068 | 1091 | 1 | 68.93 | 0.00000087 | Y | VPAQEKNFTTAPAICHDGKAHFPR | E | 0.000000200000000000000000.0 | Delta:H(1)N(-1)18O(1) (N) | 1 | 1 | 1074(NFT)/ | 1074(NFT)/ | 1 |
| chymotrypsin (semi) | 91 | 1::P0DTC2 | 10322 | 1 | 782.6379 | 3126.5227 | 4 | 3126.5231 | -0.0005 | 1068 | 1095 | 1 | 71.43 | 0.0000053 | Y | VPAQEKNFTTAPAICHDGKAHFPREGVF | V | 0.0000002000000000000000000000.0 | Delta:H(1)N(-1)18O(1) (N) | 1 | 1 | 1074(NFT)/ | 1074(NFT)/ | 1 |
| chymotrypsin (semi) | 91 | 1::P0DTC2 | 1599 | 1 | 618.2827 | 1234.5509 | 2 | 1234.5505 | 0.0005 | 1092 | 1102 | 1 | 62.05 | 0.00012 | R | EGVFVSNGTHW | F | 0.00000020000.0 | Delta:H(1)N(-1)18O(1) (N) | 1 | 1 | 1098(NGT)/ | 1098(NGT)/ | 1 |
| chymotrypsin (semi) | 91 | 1::P0DTC2 | 2113 | 1 | 691.8169 | 1381.6192 | 2 | 1381.6189 | 0.0003 | 1092 | 1103 | 2 | 40.74 | 0.02 | R | EGVFVSNGTHWF | V | 0.000000200000.0 | Delta:H(1)N(-1)18O(1) (N) | 1 | 1 | 1098(NGT)/ | 1098(NGT)/ | 1 |
| chymotrypsin (semi) | 91 | 1::P0DTC2 | 904 | 1 | 525.2503 | 1048.4861 | 2 | 1048.4864 | -0.0004 | 1094 | 1102 | 1 | 40.24 | 0.022 | G | VFVSNGTHW | F | 0.000020000.0 | Delta:H(1)N(-1)18O(1) (N) | 1 | 1 | 1098(NGT)/ | 1098(NGT)/ | 1 |
| chymotrypsin (semi) | 91 | 1::P0DTC2 | 502 | 1 | 475.7161 | 949.4176 | 2 | 949.418 | -0.0004 | 1096 | 1103 | 1 | 35.85 | 0.047 | F | VSNGTHWF | V | 0.00200000.0 | Delta:H(1)N(-1)18O(1) (N) | 1 | 1 | 1098(NGT)/ | 1098(NGT)/ | 1 |
| chymotrypsin (semi) | 91 | 1::P0DTC2 | 12491 | 1 | 968.9731 | 3871.8633 | 4 | 3871.86 | 0.0033 | 1122 | 1155 | 5 | 59.47 | 0.00083 | F | VSGNCDVVIGIVNNTVYDPLQPELDSFKEEI | F | 0.0000000000002000000000000000000000.0 | Delta:H(1)N(-1)18O(1) (N) | 1 | 1 | 1134(NNT)/ | 1134(NNT)/ | 1 |
| chymotrypsin (semi) | 91 | 1::P0DTC2 | 6917 | 1 | 764.3773 | 2290.11 | 3 | 2290.1101 | -0.0001 | 1156 | 1177 | 2 | 68.19 | 0.000085 | Y | FKNHTSPDVDLGDISGINASVV | N | 0.0020000000000000020000.0 | 2 Delta:H(1)N(-1)18O(1) (N) | 2 | 2 | 1158(NHT)/1173(NAS)/ | 1158(NHT)/1173(NAS)/ | 2 |
| chymotrypsin (semi) | 91 | 1::P0DTC2 | 11460 | 1 | 850.939 | 3399.7267 | 4 | 3399.7294 | -0.0026 | 1156 | 1186 | 2 | 47.5 | 0.013 | Y | FKNHTSPDVDLGDISGINASVVNIQKEIDRL | N | 0.0020000000000000020000000000000.0 | 2 Delta:H(1)N(-1)18O(1) (N) | 2 | 2 | 1158(NHT)/1173(NAS)/ | 1158(NHT)/1173(NAS)/ | 2 |
| chymotrypsin (semi) | 91 | 1::P0DTC2 | 9615 | 1 | 730.6351 | 2918.5112 | 4 | 2918.5122 | -0.001 | 1182 | 1206 | 5 | 44.74 | 0.02 | K | EIDRLNEVAKNLNESLIDLQELGKY | E | 0.0000000000002000000000000.0 | Delta:H(1)N(-1)18O(1) (N) | 1 | 1 | 1194(NES)/ | 1194(NES)/ | 1 |
| chymotrypsin (semi) | 91 | 1::P0DTC2 | 6935 | 1 | 765.0648 | 2292.1727 | 3 | 2292.1734 | -0.0007 | 1187 | 1206 | 4 | 57.65 | 0.00097 | L | NEVAKNLNESLIDLQELGKY | E | 0.00000002000000000000.0 | Delta:H(1)N(-1)18O(1) (N) | 1 | 1 | 1194(NES)/ | 1194(NES)/ | 1 |
| chymotrypsin (semi) | 91 | 1::P0DTC2 | 6340 | 1 | 727.051 | 2178.1311 | 3 | 2178.1305 | 0.0005 | 1188 | 1206 | 4 | 60.62 | 0.0004 | N | EVAKNLNESLIDLQELGKY | E | 0.0000002000000000000.0 | Delta:H(1)N(-1)18O(1) (N) | 1 | 1 | 1194(NES)/ | 1194(NES)/ | 1 |
| chymotrypsin (semi) | 91 | 1::P0DTC2 | 3913 | 1 | 876.4511 | 1750.8876 | 2 | 1750.8875 | 0.0002 | 1192 | 1206 | 4 | 99.54 | 0.000000048 | K | NLNESLIDLQELGKY | E | 0.002000000000000.0 | Delta:H(1)N(-1)18O(1) (N) | 1 | 1 | 1194(NES)/ | 1194(NES)/ | 1 |
| chymotrypsin | 49 | 1::P0DTC2 | 3427 | 1 | 419.4705 | 1673.8529 | 4 | 1673.8524 | 0.0005 | 65 | 79 | 1 | 92.76 | 2.10E-08 | W | FHAIHVSGTNGTKRF | D | 0.000000000200000.0 | Delta:H(1)N(-1)18O(1) (N) | 1 | 1 | 74(NGT)/ | 74(NGT)/ | 1 |
| chymotrypsin | 49 | 1::P0DTC2 | 2681 | 1 | 509.935 | 1526.7831 | 3 | 1526.784 | -0.0009 | 66 | 79 | 0 | 101.54 | 2.70E-09 | F | HAIHVSGTNGTKRF | D | 0.00000000200000.0 | Delta:H(1)N(-1)18O(1) (N) | 1 | 1 | 74(NGT)/ | 74(NGT)/ | 1 |
| chymotrypsin | 49 | 1::P0DTC2 | 7021 | 1 | 770.7347 | 2309.1824 | 3 | 2309.1803 | 0.0021 | 66 | 86 | 1 | 67.61 | 1.00E-05 | F | HAIHVSGTNGTKRFDNPVLPF | N | 0.000000002000000000000.0 | Delta:H(1)N(-1)18O(1) (N) | 1 | 1 | 74(NGT)/ | 74(NGT)/ | 1 |
| chymotrypsin | 49 | 1::P0DTC2 | 9432 | 1 | 953.4749 | 2857.4029 | 3 | 2857.4034 | -0.0004 | 66 | 91 | 2 | 56.03 | 0.00019 | F | HAIHVSGTNGTKRFDNPVLPFNDGVY | F | 0.00000000200000000000000000.0 | Delta:H(1)N(-1)18O(1) (N) | 1 | 1 | 74(NGT)/ | 74(NGT)/ | 1 |
| chymotrypsin | 49 | 1::P0DTC2 | 4426 | 1 | 918.4949 | 1834.9752 | 2 | 1834.9749 | 0.0003 | 118 | 133 | 1 | 109.08 | 3.90E-10 | L | LIVNNATNVVIKVCEF | Q | 0.0000200000000000.0 | Delta:H(1)N(-1)18O(1) (N) | 1 | 1 | 122(NAT)/ | 122(NAT)/ | 1 |
| chymotrypsin | 49 | 1::P0DTC2 | 3700 | 1 | 861.9517 | 1721.8889 | 2 | 1721.8908 | -0.0019 | 119 | 133 | 0 | 100.49 | 3.50E-09 | L | IVNNATNVVIKVCEF | Q | 0.000200000000000.0 | Delta:H(1)N(-1)18O(1) (N) | 1 | 1 | 122(NAT)/ | 122(NAT)/ | 1 |
| chymotrypsin | 49 | 1::P0DTC2 | 3552 | 1 | 851.8747 | 1701.7349 | 2 | 1701.7343 | 0.0007 | 145 | 157 | 2 | 61.72 | 1.40E-05 | Y | YHKNNKSWMESEF | R | 0.0000200000000.0 | Delta:H(1)N(-1)18O(1) (N) | 1 | 1 | 149(NKS)/ | 149(NKS)/ | 1 |
| chymotrypsin | 49 | 1::P0DTC2 | 3632 | 1 | 573.5834 | 1717.7284 | 3 | 1717.7292 | -0.0008 | 145 | 157 | 2 | 56.79 | 2.70E-05 | Y | YHKNNKSWMESEF | R | 0.0000200040000.0 | Delta:H(1)N(-1)18O(1) (N); Oxidation (M) | 1 | 1 | 149(NKS)/ | 149(NKS)/ | 1 |
| chymotrypsin | 49 | 1::P0DTC2 | 2742 | 1 | 513.8975 | 1538.6705 | 3 | 1538.6709 | -0.0004 | 146 | 157 | 1 | 38.24 | 0.0036 | Y | HKNNKSWMESEF | R | 0.000200000000.0 | Delta:H(1)N(-1)18O(1) (N) | 1 | 1 | 149(NKS)/ | 149(NKS)/ | 1 |
| chymotrypsin | 49 | 1::P0DTC2 | 2805 | 1 | 519.229 | 1554.6652 | 3 | 1554.6659 | -0.0007 | 146 | 157 | 1 | 26.96 | 0.035 | Y | HKNNKSWMESEF | R | 0.000200040000.0 | Delta:H(1)N(-1)18O(1) (N); Oxidation (M) | 1 | 1 | 149(NKS)/ | 149(NKS)/ | 1 |
| chymotrypsin | 49 | 1::P0DTC2 | 1477 | 1 | 598.2264 | 1194.4383 | 2 | 1194.4385 | -0.0002 | 161 | 170 | 1 | 27.65 | 0.0051 | Y | SSANNCTFEY | V | 0.0000200000.0 | Delta:H(1)N(-1)18O(1) (N) | 1 | 2 | 165(NCT)/ | 164(NNC)/165(NCT)/ | 1 |
| chymotrypsin | 49 | 1::P0DTC2 | 3932 | 1 | 877.3663 | 1752.718 | 2 | 1752.7187 | -0.0007 | 161 | 175 | 2 | 31.65 | 0.0054 | Y | SSANNCTFEYVSQPF | L | 0.000020000000000.0 | Delta:H(1)N(-1)18O(1) (N) | 1 | 2 | 165(NCT)/ | 164(NNC)/165(NCT)/ | 1 |
| chymotrypsin | 49 | 1::P0DTC2 | 5156 | 1 | 986.055 | 1970.0955 | 2 | 1970.0974 | -0.0019 | 221 | 238 | 2 | 105.42 | 5.90E-10 | F | SALEPLVDLPIGINITRF | Q | 0.000000000000020000.0 | Delta:H(1)N(-1)18O(1) (N) | 1 | 1 | 234(NIT)/ | 234(NIT)/ | 1 |
| chymotrypsin | 49 | 1::P0DTC2 | 2031 | 1 | 680.8905 | 1359.7664 | 2 | 1359.7648 | 0.0016 | 227 | 238 | 0 | 58.87 | 1.80E-05 | L | VDLPIGINITRF | Q | 0.000000020000.0 | Delta:H(1)N(-1)18O(1) (N) | 1 | 1 | 234(NIT)/ | 234(NIT)/ | 1 |
| chymotrypsin | 49 | 1::P0DTC2 | 10421 | 1 | 1053.1767 | 3156.5083 | 3 | 3156.5092 | -0.001 | 276 | 303 | 5 | 87.99 | 1.00E-07 | F | LLKYNENGTITDAVDCALDPLSETKCTL | K | 0.0000002000000000000000000000.0 | Delta:H(1)N(-1)18O(1) (N) | 1 | 1 | 282(NGT)/ | 282(NGT)/ | 1 |
| chymotrypsin | 49 | 1::P0DTC2 | 11858 | 1 | 880.6837 | 3518.7056 | 4 | 3518.7046 | 0.0009 | 276 | 306 | 6 | 52.49 | 0.00041 | F | LLKYNENGTITDAVDCALDPLSETKCTLKSI | T | 0.0000002000000000000000000000000.0 | Delta:H(1)N(-1)18O(1) (N) | 1 | 1 | 282(NGT)/ | 282(NGT)/ | 1 |
| chymotrypsin | 49 | 1::P0DTC2 | 10004 | 1 | 1015.4822 | 3043.4248 | 3 | 3043.4252 | -0.0004 | 277 | 303 | 4 | 85.68 | 1.50E-07 | L | LKYNENGTITDAVDCALDPLSETKCTL | K | 0.000002000000000000000000000.0 | Delta:H(1)N(-1)18O(1) (N) | 1 | 1 | 282(NGT)/ | 282(NGT)/ | 1 |
| chymotrypsin | 49 | 1::P0DTC2 | 11515 | 1 | 852.4124 | 3405.6207 | 4 | 3405.6206 | 0.0001 | 277 | 306 | 5 | 53.75 | 0.00025 | L | LKYNENGTITDAVDCALDPLSETKCTLKSF | T | 0.000002000000000000000000000000.0 | Delta:H(1)N(-1)18O(1) (N) | 1 | 1 | 282(NGT)/ | 282(NGT)/ | 1 |
| chymotrypsin | 49 | 1::P0DTC2 | 9657 | 1 | 977.7878 | 2930.3415 | 3 | 2930.3411 | 0.0003 | 278 | 303 | 3 | 79.82 | 4.40E-07 | L | KYNENGTITDAVDCALDPLSETKCTL | K | 0.00002000000000000000000000.0 | Delta:H(1)N(-1)18O(1) (N) | 1 | 1 | 282(NGT)/ | 282(NGT)/ | 1 |
| chymotrypsin | 49 | 1::P0DTC2 | 11011 | 1 | 824.1413 | 3292.5362 | 4 | 3292.5365 | -0.0003 | 278 | 306 | 4 | 59.48 | 6.20E-05 | L | KYNENGTITDAVDCALDPLSETKCTLKSF | T | 0.00002000000000000000000000000.0 | Delta:H(1)N(-1)18O(1) (N) | 1 | 1 | 282(NGT)/ | 282(NGT)/ | 1 |

| | | | | | | | | | | | | | | | | | | | | | | | | |
|---|---|---|---|---|---|---|---|---|---|---|---|---|---|---|---|---|---|---|---|---|---|---|---|---|
| chymotrypsin | 49 | 1::P0DTC2 | 8653 | 1 | 880.7351 | 2639.1834 | 3 | 2639.1828 | 0.0006 | 280 | 303 | 2 | 60.44 | 2.90E-05 | Y | NENGTITDAVDCALDPLSETKCTL | K | 0.0020000000000000000000000.0 | Delta:H(1)N(-1)18O(1) (N) | 1 | 1 | 282(NGT)/ | 282(NGT)/ | 1 |
| chymotrypsin | 49 | 1::P0DTC2 | 9873 | 1 | 1001.4663 | 3001.3771 | 3 | 3001.3782 | -0.0012 | 280 | 306 | 3 | 37.86 | 0.0077 | Y | NENGTITDAVDCALDPLSETKCTLKSF | T | 0.0020000000000000000000000000.0 | Delta:H(1)N(-1)18O(1) (N) | 1 | 1 | 282(NGT)/ | 282(NGT)/ | 1 |
| chymotrypsin | 49 | 1::P0DTC2 | 5243 | 1 | 663.0335 | 1986.0786 | 3 | 1986.0784 | 0.0002 | 319 | 335 | 0 | 93.46 | 1.60E-08 | F | RVQPTESIVRFPNITNL | C | 0.00000000000020000.0 | Delta:H(1)N(-1)18O(1) (N) | 1 | 2 | 331(NIT)/ | 331(NIT)/334(NLC)/ | 1 |
| chymotrypsin | 49 | 1::P0DTC2 | 9338 | 1 | 941.8169 | 2822.4288 | 3 | 2822.4312 | -0.0023 | 319 | 342 | 2 | 37.07 | 0.014 | F | RVQPTESIVRFPNITNLCPFGEVF | N | 0.00000000000002000000000000.0 | Delta:H(1)N(-1)18O(1) (N) | 1 | 2 | 331(NIT)/ | 331(NIT)/334(NLC)/ | 1 |
| chymotrypsin | 49 | 1::P0DTC2 | 11545 | 1 | 854.6863 | 3414.7162 | 4 | 3414.7167 | -0.0004 | 319 | 347 | 3 | 63.16 | 3.80E-05 | F | RVQPTESIVRFPNITNLCPFGEVFNATRF | A | 0.0000000000000200000000000020000.0 | 2 Delta:H(1)N(-1)18O(1) (N) | 2 | 3 | 331(NIT)/343(NAT)/ | 331(NIT)/334(NLC)/343(NAT)/ | 2 |
| chymotrypsin | 49 | 1::P0DTC2 | 2352 | 1 | 724.3313 | 1446.648 | 2 | 1446.6488 | -0.0008 | 336 | 347 | 2 | 80.02 | 2.00E-07 | L | CPFGEVFNATRF | A | 0.000000020000.0 | Delta:H(1)N(-1)18O(1) (N) | 1 | 1 | 343(NAT)/ | 343(NAT)/ | 1 |
| chymotrypsin | 49 | 1::P0DTC2 | 2270 | 1 | 715.8185 | 1429.6224 | 2 | 1429.6223 | 0.0002 | 336 | 347 | 2 | 76.55 | 3.20E-07 | L | CPFGEVFNATRF | A | 1.000000020000.0 | Ammonia-loss (N-term C); Delta:H(1)N(-1)18O(1) (N) | 1 | 1 | 343(NAT)/ | 343(NAT)/ | 1 |
| chymotrypsin | 49 | 1::P0DTC2 | 889 | 1 | 522.2556 | 1042.4966 | 2 | 1042.497 | -0.0003 | 339 | 347 | 1 | 52.34 | 0.00016 | F | GEVFNATRF | A | 0.000020000.0 | Delta:H(1)N(-1)18O(1) (N) | 1 | 1 | 343(NAT)/ | 343(NAT)/ | 1 |
| chymotrypsin | 49 | 1::P0DTC2 | 8300 | 1 | 1280.5983 | 2559.182 | 2 | 2559.1838 | -0.0017 | 612 | 633 | 2 | 53.38 | 0.00021 | L | YQDVNCTEVPVAIHADQLTPTW | R | 0.0000200000000000000000000.0 | Delta:H(1)N(-1)18O(1) (N) | 1 | 1 | 616(NCT)/ | 616(NCT)/ | 1 |
| chymotrypsin | 49 | 1::P0DTC2 | 9793 | 1 | 993.4798 | 2977.4175 | 3 | 2977.4166 | 0.0009 | 612 | 636 | 3 | 98.99 | 7.90E-09 | L | YQDVNCTEVPVAIHADQLTPTWRVY | S | 0.0000200000000000000000000000.0 | Delta:H(1)N(-1)18O(1) (N) | 1 | 1 | 616(NCT)/ | 616(NCT)/ | 1 |
| chymotrypsin | 49 | 1::P0DTC2 | 4846 | 1 | 956.4539 | 1910.8932 | 2 | 1910.893 | 0.0002 | 613 | 629 | 0 | 46.68 | 0.0008 | Y | QDVNCTEVPVAIHADQL | T | 0.00020000000000000.0 | Delta:H(1)N(-1)18O(1) (N) | 1 | 1 | 616(NCT)/ | 616(NCT)/ | 1 |
| chymotrypsin | 49 | 1::P0DTC2 | 7443 | 1 | 1199.0671 | 2396.1196 | 2 | 2396.1205 | -0.0008 | 613 | 633 | 1 | 62.19 | 2.80E-05 | Y | QDVNCTEVPVAIHADQLTPTW | R | 0.000200000000000000000000.0 | Delta:H(1)N(-1)18O(1) (N) | 1 | 1 | 616(NCT)/ | 616(NCT)/ | 1 |
| chymotrypsin | 49 | 1::P0DTC2 | 7351 | 1 | 1190.5537 | 2379.0928 | 2 | 2379.0939 | -0.0011 | 613 | 633 | 1 | 52.48 | 0.00022 | Y | QDVNCTEVPVAIHADQLTPTW | R | 3.000200000000000000000.0 | Delta:H(1)N(-1)18O(1) (N); Gln->pyro-Glu (N-term Q) | 1 | 1 | 616(NCT)/ | 616(NCT)/ | 1 |
| chymotrypsin | 49 | 1::P0DTC2 | 9314 | 1 | 939.1251 | 2814.3534 | 3 | 2814.3533 | 0.0001 | 613 | 636 | 2 | 83.06 | 3.00E-07 | Y | QDVNCTEVPVAIHADQLTPTWRVY | S | 0.00020000000000000000000000.0 | Delta:H(1)N(-1)18O(1) (N) | 1 | 1 | 616(NCT)/ | 616(NCT)/ | 1 |
| chymotrypsin | 49 | 1::P0DTC2 | 11345 | 1 | 1128.1723 | 3381.4951 | 3 | 3381.495 | 0.0001 | 644 | 674 | 2 | 48.98 | 0.00032 | F | QTRAGCLIGAEHVNNSYECDIPIGAGICASY | Q | 3.0000000000000020000000000000000.0 | Delta:H(1)N(-1)18O(1) (N); Gln->pyro-Glu (N-term Q) | 1 | 1 | 657(NNS)/ | 657(NNS)/ | 1 |
| chymotrypsin | 49 | 1::P0DTC2 | 11444 | 1 | 1133.8469 | 3398.5189 | 3 | 3398.5215 | -0.0027 | 644 | 674 | 2 | 32.34 | 0.016 | F | QTRAGCLIGAEHVNNSYECDIPIGAGICASY | Q | 0.0000000000000020000000000000000.0 | Delta:H(1)N(-1)18O(1) (N) | 1 | 1 | 657(NNS)/ | 657(NNS)/ | 1 |
| chymotrypsin | 49 | 1::P0DTC2 | 8527 | 1 | 1307.0768 | 2612.139 | 2 | 2612.1409 | -0.0018 | 651 | 674 | 1 | 61.19 | 1.40E-05 | L | IGAEHVNNSYECDIPIGAGICASY | Q | 0.0000000200000000000000000.0 | Delta:H(1)N(-1)18O(1) (N) | 1 | 1 | 657(NNS)/ | 657(NNS)/ | 1 |
| chymotrypsin | 49 | 1::P0DTC2 | 1452 | 1 | 592.2843 | 1182.554 | 2 | 1182.554 | 0 | 708 | 718 | 0 | 46.66 | 0.00061 | Y | SNNSIAIPTNF | T | 0.02000000020.0 | 2 Delta:H(1)N(-1)18O(1) (N) | 2 | 2 | 709(NNS)/717(NFT)/ | 709(NNS)/717(NFT)/ | 2 |
| chymotrypsin | 49 | 1::P0DTC2 | 2793 | 1 | 774.9008 | 1547.7871 | 2 | 1547.787 | 0.0001 | 1063 | 1075 | 2 | 66.71 | 8.10E-06 | F | LHVTYVPAQEKNF | T | 0.0000000000020.0 | Delta:H(1)N(-1)18O(1) (N) | 1 | 1 | 1074(NFT)/ | 1074(NFT)/ | 1 |
| chymotrypsin | 49 | 1::P0DTC2 | 12279 | 1 | 748.9763 | 3739.8451 | 5 | 3739.8456 | -0.0004 | 1063 | 1095 | 3 | 56.92 | 0.00015 | F | LHVTYVPAQEKNFTTAPAICHDGKAHFPRE( | V | 0.000000000000200000000000000000000.0 | Delta:H(1)N(-1)18O(1) (N) | 1 | 1 | 1074(NFT)/ | 1074(NFT)/ | 1 |
| chymotrypsin | 49 | 1::P0DTC2 | 2300 | 1 | 718.3587 | 1434.7029 | 2 | 1434.7029 | 0 | 1064 | 1075 | 1 | 30.26 | 0.032 | L | HVTYVPAQEKNF | T | 0.000000000020.0 | Delta:H(1)N(-1)18O(1) (N) | 1 | 1 | 1074(NFT)/ | 1074(NFT)/ | 1 |
| chymotrypsin | 49 | 1::P0DTC2 | 453 | 1 | 468.2394 | 934.4643 | 2 | 934.4646 | -0.0002 | 1068 | 1075 | 0 | 45.06 | 0.00078 | Y | VPAQEKNF | T | 0.00000020.0 | Delta:H(1)N(-1)18O(1) (N) | 1 | 1 | 1074(NFT)/ | 1074(NFT)/ | 1 |
| chymotrypsin | 49 | 1::P0DTC2 | 10322 | 1 | 782.6379 | 3126.5227 | 4 | 3126.5231 | -0.0005 | 1068 | 1095 | 1 | 71.43 | 5.40E-06 | Y | VPAQEKNFTTAPAICHDGKAHFPREGVF | V | 0.0000000200000000000000000000.0 | Delta:H(1)N(-1)18O(1) (N) | 1 | 1 | 1074(NFT)/ | 1074(NFT)/ | 1 |
| chymotrypsin | 49 | 1::P0DTC2 | 10363 | 1 | 629.3023 | 3141.4752 | 5 | 3141.4766 | -0.0014 | 1076 | 1103 | 2 | 34.77 | 0.02 | F | TTAPAICHDGKAHFPREGVFVSNGTHWF | V | 0.0000000000000000000000200000.0 | Delta:H(1)N(-1)18O(1) (N) | 1 | 1 | 1098(NGT)/ | 1098(NGT)/ | 1 |
| chymotrypsin | 49 | 1::P0DTC2 | 62 | 1 | 402.1822 | 802.3498 | 2 | 802.3496 | 0.0002 | 1096 | 1102 | 0 | 30.97 | 0.0048 | F | VSNGTHW | F | 0.0020000.0 | Delta:H(1)N(-1)18O(1) (N) | 1 | 1 | 1098(NGT)/ | 1098(NGT)/ | 1 |
| chymotrypsin | 49 | 1::P0DTC2 | 502 | 1 | 475.7161 | 949.4176 | 2 | 949.418 | -0.0004 | 1096 | 1103 | 1 | 35.85 | 0.0038 | F | VSNGTHWF | V | 0.00200000.0 | Delta:H(1)N(-1)18O(1) (N) | 1 | 1 | 1098(NGT)/ | 1098(NGT)/ | 1 |
| chymotrypsin | 49 | 1::P0DTC2 | 12491 | 1 | 968.9731 | 3871.8633 | 4 | 3871.86 | 0.0033 | 1122 | 1155 | 5 | 59.47 | 8.00E-05 | F | VSGNCDVVIGIVNNTVYDPLQPELDSFKEEI | F | 0.0000000000002000000000000000000000.0 | Delta:H(1)N(-1)18O(1) (N) | 1 | 1 | 1134(NNT)/ | 1134(NNT)/ | 1 |
| chymotrypsin | 49 | 1::P0DTC2 | 11460 | 1 | 850.939 | 3399.7267 | 4 | 3399.7294 | -0.0026 | 1156 | 1186 | 2 | 47.5 | 0.0012 | Y | FKNHTSPDVDLGDISGINASVVNIQKEIDRL | N | 0.0020000000000000020000000000000.0 | 2 Delta:H(1)N(-1)18O(1) (N) | 2 | 2 | 1158(NHT)/1173(NAS)/ | 1158(NHT)/1173(NAS)/ | 2 |
| chymotrypsin | 49 | 1::P0DTC2 | 6935 | 1 | 765.0648 | 2292.1727 | 3 | 2292.1734 | -0.0007 | 1187 | 1206 | 4 | 57.65 | 0.0001 | L | NEVAKNLNESLIDLQELGKY | E | 0.00000002000000000000.0 | Delta:H(1)N(-1)18O(1) (N) | 1 | 1 | 1194(NES)/ | 1194(NES)/ | 1 |

* Results shown in blue are overlapped with those for semi-search.

Supplementary Table S6. Identification of SARS CoV-2 spike protein-derived N-glycosylated peptides of tryptic digest

| Enzyme for database search | No. of identified peptides / protein accession | Protein accession | Query No. | Rank | m/z of precursor ion | Calculated mass value of peptide | Charge (z) of precursor ion | Mass value of assigned peptide | Mass difference | Peptide position from | Peptide position to | No. of missed cleavage | Peptide score of Mascot | Expectation value | Amino acid before peptide | Peptide sequence | Amino acid after peptide | Type and position of variable modifications (1: ammonia-loss (N-term C), 2: Delta:H(1)N(-1)18O(1) (N), 3: pyroGlu (N-term Q), 4: oxidation (M)) | Type of variable modifications | No. of deglycosylated site | No. of potential N-glycosylation site | Assigned deglycosylated site (sequon) | Potential site (sequon) | No. of potential N-glycosylation site in peptide |
|---|---|---|---|---|---|---|---|---|---|---|---|---|---|---|---|---|---|---|---|---|---|---|---|---|
| trypsin (semi) | 32 | 1::P0DTC2 | 523 | 1 | 489.2339 | 976.4531 | 2 | 976.4534 | -0.0003 | 14 | 21 | 0 | 62.67 | 0.000069 | S | QCVNLTTR | T | 3.00020000.0 | Delta:H(1)N(-1)18O(1) (N); Gln->pyro-Glu (N-term Q) | 1 | 1 | 17(NLT)/ | 17(NLT)/ | 1 |
| trypsin (semi) | 32 | 1::P0DTC2 | 1411 | 1 | 408.8789 | 1223.6148 | 3 | 1223.6145 | 0.0003 | 66 | 77 | 0 | 57.92 | 0.00031 | F | HAIHVSGTNGTK | R | 0.000000002000.0 | Delta:H(1)N(-1)18O(1) (N) | 1 | 1 | 74(NGT)/ | 74(NGT)/ | 1 |
| trypsin (semi) | 32 | 1::P0DTC2 | 2905 | 1 | 865.5004 | 1728.9863 | 2 | 1728.9871 | -0.0008 | 114 | 129 | 0 | 133.37 | 2.6E-12 | K | TQSLLIVNNATNVVIK | V | 0.0000000020000000.0 | Delta:H(1)N(-1)18O(1) (N) | 1 | 1 | 122(NAT)/ | 122(NAT)/ | 1 |
| trypsin (semi) | 32 | 1::P0DTC2 | 2346 | 1 | 750.948 | 1499.8815 | 2 | 1499.8809 | 0.0007 | 116 | 129 | 0 | 46.31 | 0.00093 | Q | SLLIVNNATNVVIK | V | 0.00000020000000.0 | Delta:H(1)N(-1)18O(1) (N) | 1 | 1 | 122(NAT)/ | 122(NAT)/ | 1 |
| trypsin (semi) | 32 | 1::P0DTC2 | 2048 | 1 | 707.4314 | 1412.8483 | 2 | 1412.8488 | -0.0006 | 117 | 129 | 0 | 63.3 | 0.000013 | S | LLIVNNATNVVIK | V | 0.0000020000000.0 | Delta:H(1)N(-1)18O(1) (N) | 1 | 1 | 122(NAT)/ | 122(NAT)/ | 1 |
| trypsin (semi) | 32 | 1::P0DTC2 | 1626 | 1 | 650.8896 | 1299.7647 | 2 | 1299.7648 | -0.0001 | 118 | 129 | 0 | 62.87 | 0.000024 | L | LIVNNATNVVIK | V | 0.000020000000.0 | Delta:H(1)N(-1)18O(1) (N) | 1 | 1 | 122(NAT)/ | 122(NAT)/ | 1 |
| trypsin (semi) | 32 | 1::P0DTC2 | 5652 | 1 | 1401.6275 | 2801.2404 | 2 | 2801.245 | -0.0046 | 159 | 182 | 0 | 128.08 | 2.2E-11 | R | VYSSANNCTFEYVSQPFLMDLEGK | Q | 0.000000200000000000000000.0 | Delta:H(1)N(-1)18O(1) (N) | 1 | 2 | 165(NCT)/ | 164(NNC)/165(NCT)/ | 1 |
| trypsin (semi) | 32 | 1::P0DTC2 | 5691 | 1 | 1409.6263 | 2817.238 | 2 | 2817.2399 | -0.0019 | 159 | 182 | 0 | 111.98 | 7.4E-10 | R | VYSSANNCTFEYVSQPFLMDLEGK | Q | 0.000000200000000000400000.0 | Delta:H(1)N(-1)18O(1) (N); Oxidation (M) | 1 | 2 | 165(NCT)/ | 164(NNC)/165(NCT)/ | 1 |
| trypsin (semi) | 32 | 1::P0DTC2 | 4980 | 1 | 1241.1773 | 2480.34 | 2 | 2480.3412 | -0.0012 | 215 | 237 | 0 | 85.44 | 0.00000029 | R | DLPQGFSALEPLVDLPIGINITR | F | 0.00000000000000000002000.0 | Delta:H(1)N(-1)18O(1) (N) | 1 | 1 | 234(NIT)/ | 234(NIT)/ | 1 |
| trypsin (semi) | 32 | 1::P0DTC2 | 3415 | 1 | 912.5216 | 1823.0286 | 2 | 1823.029 | -0.0004 | 221 | 237 | 0 | 96.78 | 0.000000013 | F | SALEPLVDLPIGINITR | F | 0.00000000000002000.0 | Delta:H(1)N(-1)18O(1) (N) | 1 | 1 | 234(NIT)/ | 234(NIT)/ | 1 |
| trypsin (semi) | 32 | 1::P0DTC2 | 2964 | 1 | 869.0059 | 1735.9972 | 2 | 1735.997 | 0.0002 | 222 | 237 | 0 | 57.74 | 0.000067 | S | ALEPLVDLPIGINITR | F | 0.0000000000002000.0 | Delta:H(1)N(-1)18O(1) (N) | 1 | 1 | 234(NIT)/ | 234(NIT)/ | 1 |
| trypsin (semi) | 32 | 1::P0DTC2 | 4833 | 1 | 1215.0474 | 2428.0802 | 2 | 2428.0838 | -0.0035 | 279 | 300 | 0 | 151.04 | 1.3E-13 | K | YNENGTITDAVDCALDPLSETK | C | 0.0002000000000000000000.0 | Delta:H(1)N(-1)18O(1) (N) | 1 | 1 | 282(NGT)/ | 282(NGT)/ | 1 |
| trypsin (semi) | 32 | 1::P0DTC2 | 4204 | 1 | 1076.4954 | 2150.9762 | 2 | 2150.9775 | -0.0013 | 281 | 300 | 0 | 49.8 | 0.0018 | N | ENGTITDAVDCALDPLSETK | C | 0.02000000000000000000.0 | Delta:H(1)N(-1)18O(1) (N) | 1 | 1 | 282(NGT)/ | 282(NGT)/ | 1 |
| trypsin (semi) | 32 | 1::P0DTC2 | 4113 | 1 | 1052.0025 | 2101.9904 | 2 | 2101.9915 | -0.001 | 329 | 346 | 0 | 117.11 | 4.8E-10 | R | FPNITNLCPFGEVFNATR | F | 0.002000000000002000.0 | 2 Delta:H(1)N(-1)18O(1) (N) | 2 | 3 | 331(NIT)/343(NAT)/ | 331(NIT)/334(NLC)/343(NAT)/ | 2 |
| trypsin (semi) | 32 | 1::P0DTC2 | 2046 | 1 | 707.3395 | 1412.6644 | 2 | 1412.6644 | 0 | 335 | 346 | 0 | 67.37 | 0.000033 | N | LCPFGEVFNATR | F | 0.000000002000.0 | Delta:H(1)N(-1)18O(1) (N) | 1 | 1 | 343(NAT)/ | 343(NAT)/ | 1 |
| trypsin (semi) | 32 | 1::P0DTC2 | 5902 | 1 | 976.82 | 2927.4382 | 3 | 2927.4374 | 0.0009 | 610 | 634 | 0 | 125.34 | 8.5E-11 | A | VLYQDVNCTEVPVAIHADQLTPTWR | V | 0.0000002000000000000000000.0 | Delta:H(1)N(-1)18O(1) (N) | 1 | 1 | 616(NCT)/ | 616(NCT)/ | 1 |
| trypsin (semi) | 32 | 1::P0DTC2 | 1521 | 1 | 629.2921 | 1256.5696 | 2 | 1256.5705 | -0.0009 | 647 | 658 | 0 | 46.67 | 0.0026 | R | AGCLIGAEHVNN | S | 0.000000000002.0 | Delta:H(1)N(-1)18O(1) (N) | 1 | 1 | 657(NNS)/ | 657(NNS)/ | 1 |
| trypsin (semi) | 32 | 1::P0DTC2 | 3242 | 1 | 886.4242 | 1770.8338 | 2 | 1770.8351 | -0.0013 | 796 | 811 | 0 | 130.77 | 1.7E-11 | K | DFGGFNFSQILPDPSK | P | 0.0000020000000000.0 | Delta:H(1)N(-1)18O(1) (N) | 1 | 1 | 801(NFS)/ | 801(NFS)/ | 1 |
| trypsin (semi) | 32 | 1::P0DTC2 | 3827 | 1 | 978.4675 | 1954.9204 | 2 | 1954.9199 | 0.0006 | 796 | 813 | 1 | 50.9 | 0.0018 | K | DFGGFNFSQILPDPSKPS | K | 0.000002000000000000.0 | Delta:H(1)N(-1)18O(1) (N) | 1 | 1 | 801(NFS)/ | 801(NFS)/ | 1 |
| trypsin (semi) | 32 | 1::P0DTC2 | 4068 | 1 | 695.3451 | 2083.0134 | 3 | 2083.0148 | -0.0014 | 796 | 814 | 1 | 107.83 | 4.7E-09 | K | DFGGFNFSQILPDPSKPSK | R | 0.0000020000000000000.0 | Delta:H(1)N(-1)18O(1) (N) | 1 | 1 | 801(NFS)/ | 801(NFS)/ | 1 |
| trypsin (semi) | 32 | 1::P0DTC2 | 4372 | 1 | 747.3791 | 2239.1156 | 3 | 2239.1159 | -0.0003 | 796 | 815 | 2 | 73.27 | 0.000013 | K | DFGGFNFSQILPDPSKPSKR | S | 0.00000200000000000000.0 | Delta:H(1)N(-1)18O(1) (N) | 1 | 1 | 801(NFS)/ | 801(NFS)/ | 1 |
| trypsin (semi) | 32 | 1::P0DTC2 | 2733 | 1 | 828.9114 | 1655.8083 | 2 | 1655.8081 | 0.0002 | 797 | 811 | 0 | 70.9 | 0.000019 | D | FGGFNFSQILPDPSK | P | 0.000020000000000.0 | Delta:H(1)N(-1)18O(1) (N) | 1 | 1 | 801(NFS)/ | 801(NFS)/ | 1 |
| trypsin (semi) | 32 | 1::P0DTC2 | 2360 | 1 | 755.3771 | 1508.7396 | 2 | 1508.7397 | -0.0001 | 798 | 811 | 0 | 39.03 | 0.031 | F | GGFNFSQILPDPSK | P | 0.00020000000000.0 | Delta:H(1)N(-1)18O(1) (N) | 1 | 1 | 801(NFS)/ | 801(NFS)/ | 1 |
| trypsin (semi) | 32 | 1::P0DTC2 | 2143 | 1 | 478.8902 | 1433.6488 | 3 | 1433.6495 | -0.0007 | 1074 | 1086 | 0 | 81.84 | 0.00000078 | K | NFTTAPAICHDGK | A | 0.2000000000000.0 | Delta:H(1)N(-1)18O(1) (N) | 1 | 1 | 1074(NFT)/ | 1074(NFT)/ | 1 |
| trypsin (semi) | 32 | 1::P0DTC2 | 1449 | 1 | 618.2824 | 1234.5502 | 2 | 1234.5505 | -0.0003 | 1092 | 1102 | 0 | 54.79 | 0.00036 | R | EGVFVSNGTHW | F | 0.00000020000.0 | Delta:H(1)N(-1)18O(1) (N) | 1 | 1 | 1098(NGT)/ | 1098(NGT)/ | 1 |
| trypsin (semi) | 32 | 1::P0DTC2 | 3518 | 1 | 622.972 | 1865.8941 | 3 | 1865.8947 | -0.0006 | 1092 | 1107 | 0 | 123.04 | 1.2E-10 | R | EGVFVSNGTHWFVTQR | N | 0.0000002000000000.0 | Delta:H(1)N(-1)18O(1) (N) | 1 | 1 | 1098(NGT)/ | 1098(NGT)/ | 1 |
| trypsin (semi) | 32 | 1::P0DTC2 | 2966 | 1 | 579.9574 | 1736.8503 | 3 | 1736.8521 | -0.0018 | 1093 | 1107 | 0 | 67.37 | 0.000049 | E | GVFVSNGTHWFVTQR | N | 0.000002000000000.0 | Delta:H(1)N(-1)18O(1) (N) | 1 | 1 | 1098(NGT)/ | 1098(NGT)/ | 1 |
| trypsin (semi) | 32 | 1::P0DTC2 | 5023 | 1 | 833.7495 | 2498.2265 | 3 | 2498.2272 | -0.0007 | 1158 | 1181 | 0 | 117.05 | 5.5E-10 | K | NHTSPDVDLGDISGINASVVNIQK | E | 0.200000000000000200000000.0 | 2 Delta:H(1)N(-1)18O(1) (N) | 2 | 2 | 1158(NHT)/1173(NAS)/ | 1158(NHT)/1173(NAS)/ | 2 |
| trypsin (semi) | 32 | 1::P0DTC2 | 4387 | 1 | 1123.0757 | 2244.1368 | 2 | 2244.1371 | -0.0003 | 1160 | 1181 | 0 | 80.33 | 0.0000023 | H | TSPDVDLGDISGINASVVNIQK | E | 0.0000000000000200000000.0 | Delta:H(1)N(-1)18O(1) (N) | 1 | 1 | 1173(NAS)/ | 1173(NAS)/ | 1 |
| trypsin (semi) | 32 | 1::P0DTC2 | 2240 | 1 | 730.8949 | 1459.7752 | 2 | 1459.7768 | -0.0016 | 1168 | 1181 | 0 | 84.87 | 0.00000062 | G | DISGINASVVNIQK | E | 0.00000200000000.0 | Delta:H(1)N(-1)18O(1) (N) | 1 | 1 | 1173(NAS)/ | 1173(NAS)/ | 1 |
| trypsin (semi) | 32 | 1::P0DTC2 | 2558 | 1 | 794.9192 | 1587.8239 | 2 | 1587.8241 | -0.0003 | 1192 | 1205 | 0 | 103.84 | 8.5E-09 | K | NLNESLIDLQELGK | Y | 0.00200000000000.0 | Delta:H(1)N(-1)18O(1) (N) | 1 | 1 | 1194(NES)/ | 1194(NES)/ | 1 |
| trypsin (semi) | 32 | 1::P0DTC2 | 2280 | 1 | 737.8978 | 1473.781 | 2 | 1473.7812 | -0.0002 | 1193 | 1205 | 0 | 75.49 | 0.0000049 | N | LNESLIDLQELGK | Y | 0.0200000000000.0 | Delta:H(1)N(-1)18O(1) (N) | 1 | 1 | 1194(NES)/ | 1194(NES)/ | 1 |
| trypsin | 14 | 1::P0DTC2 | 2905 | 1 | 865.5004 | 1728.9863 | 2 | 1728.9871 | -0.0008 | 114 | 129 | 0 | 133.37 | 1.90E-13 | K | TQSLLIVNNATNVVIK | V | 0.0000000020000000.0 | Delta:H(1)N(-1)18O(1) (N) | 1 | 1 | 122(NAT)/ | 122(NAT)/ | 1 |
| trypsin | 14 | 1::P0DTC2 | 5652 | 1 | 1401.6275 | 2801.2404 | 2 | 2801.245 | -0.0046 | 159 | 182 | 0 | 128.08 | 1.10E-12 | R | VYSSANNCTFEYVSQPFLMDLEGK | Q | 0.000000200000000000000000.0 | Delta:H(1)N(-1)18O(1) (N) | 1 | 2 | 165(NCT)/ | 164(NNC)/165(NCT)/ | 1 |
| trypsin | 14 | 1::P0DTC2 | 5691 | 1 | 1409.6263 | 2817.238 | 2 | 2817.2399 | -0.0019 | 159 | 182 | 0 | 111.98 | 3.90E-11 | R | VYSSANNCTFEYVSQPFLMDLEGK | Q | 0.000000200000000000400000.0 | Delta:H(1)N(-1)18O(1) (N); Oxidation (M) | 1 | 2 | 165(NCT)/ | 164(NNC)/165(NCT)/ | 1 |
| trypsin | 14 | 1::P0DTC2 | 4980 | 1 | 1241.1773 | 2480.34 | 2 | 2480.3412 | -0.0012 | 215 | 237 | 0 | 85.44 | 2.60E-08 | R | DLPQGFSALEPLVDLPIGINITR | F | 0.00000000000000000002000.0 | Delta:H(1)N(-1)18O(1) (N) | 1 | 1 | 234(NIT)/ | 234(NIT)/ | 1 |
| trypsin | 14 | 1::P0DTC2 | 4833 | 1 | 1215.0474 | 2428.0802 | 2 | 2428.0838 | -0.0035 | 279 | 300 | 0 | 151.04 | 6.20E-15 | K | YNENGTITDAVDCALDPLSETK | C | 0.0002000000000000000000.0 | Delta:H(1)N(-1)18O(1) (N) | 1 | 1 | 282(NGT)/ | 282(NGT)/ | 1 |
| trypsin | 14 | 1::P0DTC2 | 4113 | 1 | 1052.0025 | 2101.9904 | 2 | 2101.9915 | -0.001 | 329 | 346 | 0 | 117.11 | 2.30E-11 | R | FPNITNLCPFGEVFNATR | F | 0.002000000000002000.0 | 2 Delta:H(1)N(-1)18O(1) (N) | 2 | 3 | 331(NIT)/343(NAT)/ | 331(NIT)/334(NLC)/343(NAT)/ | 2 |
| trypsin | 14 | 1::P0DTC2 | 3242 | 1 | 886.4242 | 1770.8338 | 2 | 1770.8351 | -0.0013 | 796 | 811 | 0 | 130.77 | 1.20E-12 | K | DFGGFNFSQILPDPSK | P | 0.0000020000000000.0 | Delta:H(1)N(-1)18O(1) (N) | 1 | 1 | 801(NFS)/ | 801(NFS)/ | 1 |
| trypsin | 14 | 1::P0DTC2 | 2143 | 1 | 478.8902 | 1433.6488 | 3 | 1433.6495 | -0.0007 | 1074 | 1086 | 0 | 81.84 | 3.80E-08 | K | NFTTAPAICHDGK | A | 0.2000000000000.0 | Delta:H(1)N(-1)18O(1) (N) | 1 | 1 | 1074(NFT)/ | 1074(NFT)/ | 1 |
| trypsin | 14 | 1::P0DTC2 | 3518 | 1 | 622.972 | 1865.8941 | 3 | 1865.8947 | -0.0006 | 1092 | 1107 | 0 | 123.04 | 7.40E-12 | R | EGVFVSNGTHWFVTQR | N | 0.0000002000000000.0 | Delta:H(1)N(-1)18O(1) (N) | 1 | 1 | 1098(NGT)/ | 1098(NGT)/ | 1 |
| trypsin | 14 | 1::P0DTC2 | 5023 | 1 | 833.7495 | 2498.2265 | 3 | 2498.2272 | -0.0007 | 1158 | 1181 | 0 | 117.05 | 3.20E-11 | K | NHTSPDVDLGDISGINASVVNIQK | E | 0.200000000000000200000000.0 | 2 Delta:H(1)N(-1)18O(1) (N) | 2 | 2 | 1158(NHT)/1173(NAS)/ | 1158(NHT)/1173(NAS)/ | 2 |
| trypsin | 14 | 1::P0DTC2 | 2558 | 1 | 794.9192 | 1587.8239 | 2 | 1587.8241 | -0.0003 | 1192 | 1205 | 0 | 103.84 | 5.60E-10 | K | NLNESLIDLQELGK | Y | 0.00200000000000.0 | Delta:H(1)N(-1)18O(1) (N) | 1 | 1 | 1194(NES)/ | 1194(NES)/ | 1 |
| trypsin | 14 | 1::P0DTC2 | 4486 | 1 | 770.0634 | 2307.1683 | 3 | 2307.1673 | 0.0009 | 791 | 811 | 1 | 28.63 | 0.019 | K | TPPIKDFGGFNFSQILPDPSK | P | 0.000000000020000000000.0 | Delta:H(1)N(-1)18O(1) (N) | 1 | 1 | 801(NFS)/ | 801(NFS)/ | 1 |
| trypsin | 14 | 1::P0DTC2 | 4068 | 1 | 695.3451 | 2083.0134 | 3 | 2083.0148 | -0.0014 | 796 | 814 | 1 | 107.83 | 2.60E-10 | K | DFGGFNFSQILPDPSKPSK | R | 0.0000020000000000000.0 | Delta:H(1)N(-1)18O(1) (N) | 1 | 1 | 801(NFS)/ | 801(NFS)/ | 1 |
| trypsin | 14 | 1::P0DTC2 | 4372 | 1 | 747.3791 | 2239.1156 | 3 | 2239.1159 | -0.0003 | 796 | 815 | 2 | 73.27 | 7.50E-07 | K | DFGGFNFSQILPDPSKPSKR | S | 0.00000200000000000000.0 | Delta:H(1)N(-1)18O(1) (N) | 1 | 1 | 801(NFS)/ | 801(NFS)/ | 1 |

* Results shown in blue are overlapped with those for semi-search.

Supplementary Table S7. Identification of SARS CoV-2 spike protein-derived N-glycosylated peptides of alpha-Lytic protease digest

| Enzyme for database search | No. of identified peptides / protein accession | Protein accession | Query No. | Rank | m/z of precursor ion | Calculated mass value of peptide | Charge (z) of precursor ion | Mass value of assigned peptide | Mass difference | Peptide position from | Peptide position to | No. of missed cleavage | Peptide score of Mascot | Expectation value | Amino acid before peptide | Peptide sequence | Amino acid after peptide | Type and position of variable modifications (1: ammonia-loss (N-term C), 2: Delta:H(1)N(-1)18O(1) (N), 3: pyroGlu (N-term Q), 4: oxidation (M)) | Type of variable modifications | No. of deglycosy lated site | No. of potential N-glycosylation site | Assigned deglycosylated site (sequon) | Potential site (sequon) | No. of potential N-glycosylation site in peptide |
|---|---|---|---|---|---|---|---|---|---|---|---|---|---|---|---|---|---|---|---|---|---|---|---|---|
| a-Lytic (semi) | 26 | 1::P0DTC2 | 1727 | 1 | 715.858 | 1429.7014 | 2 | 1429.7016 | -0.0001 | 52 | 63 | 2 | 48.05 | 0.0067 | T | QDLFLPFFSNVT | W | 0.000000000200.0 | Delta:H(1)N(-1)18O(1) (N) | 1 | 1 | 61(NVT)/ | 61(NVT)/ | 1 |
| a-Lytic (semi) | 26 | 1::P0DTC2 | 3290 | 1 | 840.4071 | 2518.1994 | 3 | 2518.2015 | -0.0021 | 71 | 93 | 5 | 44.41 | 0.019 | V | SGTNGTKRFDNPVLPFNDGVYFA | S | 0.00020000000000000000000.0 | Delta:H(1)N(-1)18O(1) (N) | 1 | 1 | 74(NGT)/ | 74(NGT)/ | 1 |
| a-Lytic (semi) | 26 | 1::P0DTC2 | 2778 | 1 | 684.3411 | 2050.0014 | 3 | 2050.0006 | 0.0008 | 72 | 90 | 3 | 76.03 | 0.000015 | S | GTNGTKRFDNPVLPFNDGV | Y | 0.0020000000000000000.0 | Delta:H(1)N(-1)18O(1) (N) | 1 | 1 | 74(NGT)/ | 74(NGT)/ | 1 |
| a-Lytic (semi) | 26 | 1::P0DTC2 | 785 | 1 | 560.7287 | 1119.4429 | 2 | 1119.4429 | 0 | 163 | 171 | 2 | 41.63 | 0.0069 | S | ANNCTFEYV | S | 0.002000000.0 | Delta:H(1)N(-1)18O(1) (N) | 1 | 2 | 165(NCT)/ | 164(NNC)/165(NCT)/ | 1 |
| a-Lytic (semi) | 26 | 1::P0DTC2 | 1695 | 1 | 708.3692 | 1414.7238 | 2 | 1414.723 | 0.0008 | 275 | 286 | 1 | 53.53 | 0.0018 | T | FLLKYNENGTIT | D | 0.000000020000.0 | Delta:H(1)N(-1)18O(1) (N) | 1 | 1 | 282(NGT)/ | 282(NGT)/ | 1 |
| a-Lytic (semi) | 26 | 1::P0DTC2 | 1248 | 1 | 634.8348 | 1267.655 | 2 | 1267.6546 | 0.0004 | 276 | 286 | 1 | 67.59 | 0.000062 | F | LLKYNENGTIT | D | 0.00000020000.0 | Delta:H(1)N(-1)18O(1) (N) | 1 | 1 | 282(NGT)/ | 282(NGT)/ | 1 |
| a-Lytic (semi) | 26 | 1::P0DTC2 | 934 | 1 | 578.2925 | 1154.5705 | 2 | 1154.5705 | 0 | 277 | 286 | 1 | 43.41 | 0.017 | L | LKYNENGTIT | D | 0.0000020000.0 | Delta:H(1)N(-1)18O(1) (N) | 1 | 1 | 282(NGT)/ | 282(NGT)/ | 1 |
| a-Lytic (semi) | 26 | 1::P0DTC2 | 516 | 1 | 521.7503 | 1041.4861 | 2 | 1041.4864 | -0.0003 | 278 | 286 | 1 | 51.37 | 0.0022 | L | KYNENGTIT | D | 0.000020000.0 | Delta:H(1)N(-1)18O(1) (N) | 1 | 1 | 282(NGT)/ | 282(NGT)/ | 1 |
| a-Lytic (semi) | 26 | 1::P0DTC2 | 1566 | 1 | 686.3095 | 1370.6044 | 2 | 1370.6063 | -0.0019 | 334 | 345 | 2 | 55.69 | 0.0008 | T | NLCPFGEVFNAT | R | 0.000000000200.0 | Delta:H(1)N(-1)18O(1) (N) | 1 | 2 | 343(NAT)/ | 334(NLC)/343(NAT)/ | 1 |
| a-Lytic (semi) | 26 | 1::P0DTC2 | 774 | 1 | 557.7521 | 1113.4896 | 2 | 1113.4898 | -0.0003 | 610 | 618 | 2 | 41.81 | 0.018 | A | VLYQDVNCT | E | 0.000000200.0 | Delta:H(1)N(-1)18O(1) (N) | 1 | 1 | 616(NCT)/ | 616(NCT)/ | 1 |
| a-Lytic (semi) | 26 | 1::P0DTC2 | 2437 | 1 | 896.3904 | 1790.7663 | 2 | 1790.7667 | -0.0004 | 653 | 668 | 3 | 108.65 | 0.000000003 | G | AEHVNNSYECDIPIGA | G | 0.0000200000000000.0 | Delta:H(1)N(-1)18O(1) (N) | 1 | 1 | 657(NNS)/ | 657(NNS)/ | 1 |
| a-Lytic (semi) | 26 | 1::P0DTC2 | 3057 | 1 | 767.3999 | 2299.1779 | 3 | 2299.1775 | 0.0004 | 784 | 803 | 2 | 49.8 | 0.0057 | A | QVKQIYKTPPIKDFGGFNFS | Q | 3.00000000000000000200.0 | Delta:H(1)N(-1)18O(1) (N); Gln->pyro-Glu (N-term Q) | 1 | 1 | 801(NFS)/ | 801(NFS)/ | 1 |
| a-Lytic (semi) | 26 | 1::P0DTC2 | 4406 | 1 | 841.4445 | 3361.749 | 4 | 3361.7484 | 0.0006 | 784 | 813 | 4 | 80.49 | 0.0000034 | A | QVKQIYKTPPIKDFGGFNFSQILPDPSKPS | K | 3.000000000000000002000000000000.0 | Delta:H(1)N(-1)18O(1) (N); Gln->pyro-Glu (N-term Q) | 1 | 1 | 801(NFS)/ | 801(NFS)/ | 1 |
| a-Lytic (semi) | 26 | 1::P0DTC2 | 4432 | 1 | 845.701 | 3378.7748 | 4 | 3378.775 | -0.0002 | 784 | 813 | 4 | 44.08 | 0.014 | A | QVKQIYKTPPIKDFGGFNFSQILPDPSKPS | K | 0.000000000000000002000000000000.0 | Delta:H(1)N(-1)18O(1) (N) | 1 | 1 | 801(NFS)/ | 801(NFS)/ | 1 |
| a-Lytic (semi) | 26 | 1::P0DTC2 | 2817 | 1 | 697.3663 | 2089.077 | 3 | 2089.077 | -0.0001 | 786 | 803 | 1 | 75.96 | 0.000013 | V | KQIYKTPPIKDFGGFNFS | Q | 0.000000000000000200.0 | Delta:H(1)N(-1)18O(1) (N) | 1 | 1 | 801(NFS)/ | 801(NFS)/ | 1 |
| a-Lytic (semi) | 26 | 1::P0DTC2 | 4068 | 1 | 788.9184 | 3151.6444 | 4 | 3151.648 | -0.0036 | 786 | 813 | 3 | 57.58 | 0.0007 | V | KQIYKTPPIKDFGGFNFSQILPDPSKPS | K | 0.0000000000000002000000000000.0 | Delta:H(1)N(-1)18O(1) (N) | 1 | 1 | 801(NFS)/ | 801(NFS)/ | 1 |
| a-Lytic (semi) | 26 | 1::P0DTC2 | 3848 | 1 | 724.8811 | 2895.4954 | 4 | 2895.4944 | 0.0009 | 788 | 813 | 3 | 45.69 | 0.014 | Q | IYKTPPIKDFGGFNFSQILPDPSKPS | K | 0.00000000000002000000000000.0 | Delta:H(1)N(-1)18O(1) (N) | 1 | 1 | 801(NFS)/ | 801(NFS)/ | 1 |
| a-Lytic (semi) | 26 | 1::P0DTC2 | 3714 | 1 | 696.61 | 2782.4111 | 4 | 2782.4104 | 0.0007 | 789 | 813 | 3 | 46.94 | 0.011 | I | YKTPPIKDFGGFNFSQILPDPSKPS | K | 0.0000000000002000000000000.0 | Delta:H(1)N(-1)18O(1) (N) | 1 | 1 | 801(NFS)/ | 801(NFS)/ | 1 |
| a-Lytic (semi) | 26 | 1::P0DTC2 | 3428 | 1 | 874.123 | 2619.3473 | 3 | 2619.3471 | 0.0002 | 790 | 813 | 3 | 46.93 | 0.011 | Y | KTPPIKDFGGFNFSQILPDPSKPS | K | 0.000000000002000000000000.0 | Delta:H(1)N(-1)18O(1) (N) | 1 | 1 | 801(NFS)/ | 801(NFS)/ | 1 |
| a-Lytic (semi) | 26 | 1::P0DTC2 | 2144 | 1 | 818.9092 | 1635.8039 | 2 | 1635.803 | 0.0008 | 799 | 813 | 2 | 46.24 | 0.013 | G | GFNFSQILPDPSKPS | K | 0.002000000000000.0 | Delta:H(1)N(-1)18O(1) (N) | 1 | 1 | 801(NFS)/ | 801(NFS)/ | 1 |
| a-Lytic (semi) | 26 | 1::P0DTC2 | 1736 | 1 | 716.8638 | 1431.7131 | 2 | 1431.7132 | 0 | 801 | 813 | 2 | 57.3 | 0.00083 | F | NFSQILPDPSKPS | K | 0.2000000000000.0 | Delta:H(1)N(-1)18O(1) (N) | 1 | 1 | 801(NFS)/ | 801(NFS)/ | 1 |
| a-Lytic (semi) | 26 | 1::P0DTC2 | 1534 | 1 | 680.8637 | 1359.7129 | 2 | 1359.7131 | -0.0002 | 1190 | 1201 | 2 | 49.07 | 0.0053 | V | AKNLNESLIDLQ | E | 0.000020000000.0 | Delta:H(1)N(-1)18O(1) (N) | 1 | 1 | 1194(NES)/ | 1194(NES)/ | 1 |
| a-Lytic (semi) | 26 | 1::P0DTC2 | 2176 | 1 | 830.4377 | 1658.8608 | 2 | 1658.8612 | -0.0004 | 1190 | 1204 | 2 | 92.16 | 0.00000027 | V | AKNLNESLIDLQELG | K | 0.000020000000000.0 | Delta:H(1)N(-1)18O(1) (N) | 1 | 1 | 1194(NES)/ | 1194(NES)/ | 1 |
| a-Lytic (semi) | 26 | 1::P0DTC2 | 239 | 1 | 467.2604 | 932.5062 | 2 | 932.5064 | -0.0002 | 1191 | 1198 | 1 | 55.22 | 0.0011 | A | KNLNESLI | D | 0.00020000.0 | Delta:H(1)N(-1)18O(1) (N) | 1 | 1 | 1194(NES)/ | 1194(NES)/ | 1 |
| a-Lytic (semi) | 26 | 1::P0DTC2 | 1316 | 1 | 645.3457 | 1288.6768 | 2 | 1288.676 | 0.0008 | 1191 | 1201 | 1 | 50.65 | 0.0031 | A | KNLNESLIDLQ | E | 0.00020000000.0 | Delta:H(1)N(-1)18O(1) (N) | 1 | 1 | 1194(NES)/ | 1194(NES)/ | 1 |
| a-Lytic (semi) | 26 | 1::P0DTC2 | 2051 | 1 | 794.9195 | 1587.8244 | 2 | 1587.8241 | 0.0002 | 1191 | 1204 | 1 | 81.47 | 0.0000033 | A | KNLNESLIDLQELG | K | 0.00020000000000.0 | Delta:H(1)N(-1)18O(1) (N) | 1 | 1 | 1194(NES)/ | 1194(NES)/ | 1 |
| a-Lytic | 15 | 1::P0DTC2 | 1954 | 1 | 766.382 | 1530.7495 | 2 | 1530.7492 | 0.0003 | 51 | 63 | 3 | 43.54 | 0.0032 | S | TQDLFLPFFSNVT | W | 0.0000000000200.0 | Delta:H(1)N(-1)18O(1) (N) | 1 | 1 | 61(NVT)/ | 61(NVT)/ | 1 |
| a-Lytic | 15 | 1::P0DTC2 | 1727 | 1 | 715.858 | 1429.7014 | 2 | 1429.7016 | -0.0001 | 52 | 63 | 2 | 48.05 | 0.001 | T | QDLFLPFFSNVT | W | 0.000000000200.0 | Delta:H(1)N(-1)18O(1) (N) | 1 | 1 | 61(NVT)/ | 61(NVT)/ | 1 |
| a-Lytic | 15 | 1::P0DTC2 | 3290 | 1 | 840.4071 | 2518.1994 | 3 | 2518.2015 | -0.0021 | 71 | 93 | 5 | 44.41 | 0.0028 | V | SGTNGTKRFDNPVLPFNDGVYFA | S | 0.00020000000000000000000.0 | Delta:H(1)N(-1)18O(1) (N) | 1 | 1 | 74(NGT)/ | 74(NGT)/ | 1 |
| a-Lytic | 15 | 1::P0DTC2 | 2778 | 1 | 684.3411 | 2050.0014 | 3 | 2050.0006 | 0.0008 | 72 | 90 | 3 | 76.03 | 2.10E-06 | S | GTNGTKRFDNPVLPFNDGV | Y | 0.0020000000000000000.0 | Delta:H(1)N(-1)18O(1) (N) | 1 | 1 | 74(NGT)/ | 74(NGT)/ | 1 |
| a-Lytic | 15 | 1::P0DTC2 | 785 | 1 | 560.7287 | 1119.4429 | 2 | 1119.4429 | 0 | 163 | 171 | 2 | 41.63 | 0.001 | S | ANNCTFEYV | S | 0.002000000.0 | Delta:H(1)N(-1)18O(1) (N) | 1 | 2 | 165(NCT)/ | 164(NNC)/165(NCT)/ | 1 |
| a-Lytic | 15 | 1::P0DTC2 | 1695 | 1 | 708.3692 | 1414.7238 | 2 | 1414.723 | 0.0008 | 275 | 286 | 1 | 53.53 | 0.00026 | T | FLLKYNENGTIT | D | 0.000000020000.0 | Delta:H(1)N(-1)18O(1) (N) | 1 | 1 | 282(NGT)/ | 282(NGT)/ | 1 |
| a-Lytic | 15 | 1::P0DTC2 | 1566 | 1 | 686.3095 | 1370.6044 | 2 | 1370.6063 | -0.0019 | 334 | 345 | 2 | 55.69 | 0.00012 | T | NLCPFGEVFNAT | R | 0.000000000200.0 | Delta:H(1)N(-1)18O(1) (N) | 1 | 2 | 343(NAT)/ | 334(NLC)/343(NAT)/ | 1 |
| a-Lytic | 15 | 1::P0DTC2 | 403 | 1 | 496.2327 | 990.4509 | 2 | 990.4504 | 0.0005 | 600 | 609 | 4 | 37.57 | 0.0086 | T | PGTNTSNQVA | V | 0.0002000000.0 | Delta:H(1)N(-1)18O(1) (N) | 1 | 1 | 603(NTS)/ | 603(NTS)/ | 1 |
| a-Lytic | 15 | 1::P0DTC2 | 774 | 1 | 557.7521 | 1113.4896 | 2 | 1113.4898 | -0.0003 | 610 | 618 | 2 | 41.81 | 0.0026 | A | VLYQDVNCT | E | 0.000000200.0 | Delta:H(1)N(-1)18O(1) (N) | 1 | 1 | 616(NCT)/ | 616(NCT)/ | 1 |
| a-Lytic | 15 | 1::P0DTC2 | 3057 | 1 | 767.3999 | 2299.1779 | 3 | 2299.1775 | 0.0004 | 784 | 803 | 2 | 49.8 | 0.00081 | A | QVKQIYKTPPIKDFGGFNFS | Q | 3.00000000000000000200.0 | Delta:H(1)N(-1)18O(1) (N); Gln->pyro-Glu (N-term Q) | 1 | 1 | 801(NFS)/ | 801(NFS)/ | 1 |
| a-Lytic | 15 | 1::P0DTC2 | 4406 | 1 | 841.4445 | 3361.749 | 4 | 3361.7484 | 0.0006 | 784 | 813 | 4 | 80.49 | 4.60E-07 | A | QVKQIYKTPPIKDFGGFNFSQILPDPSKPS | K | 3.000000000000000002000000000000.0 | Delta:H(1)N(-1)18O(1) (N); Gln->pyro-Glu (N-term Q) | 1 | 1 | 801(NFS)/ | 801(NFS)/ | 1 |
| a-Lytic | 15 | 1::P0DTC2 | 4432 | 1 | 845.701 | 3378.7748 | 4 | 3378.775 | -0.0002 | 784 | 813 | 4 | 44.08 | 0.0018 | A | QVKQIYKTPPIKDFGGFNFSQILPDPSKPS | K | 0.000000000000000002000000000000.0 | Delta:H(1)N(-1)18O(1) (N) | 1 | 1 | 801(NFS)/ | 801(NFS)/ | 1 |
| a-Lytic | 15 | 1::P0DTC2 | 2817 | 1 | 697.3663 | 2089.077 | 3 | 2089.077 | -0.0001 | 786 | 803 | 1 | 75.96 | 2.00E-06 | V | KQIYKTPPIKDFGGFNFS | Q | 0.000000000000000200.0 | Delta:H(1)N(-1)18O(1) (N) | 1 | 1 | 801(NFS)/ | 801(NFS)/ | 1 |
| a-Lytic | 15 | 1::P0DTC2 | 4068 | 1 | 788.9184 | 3151.6444 | 4 | 3151.648 | -0.0036 | 786 | 813 | 3 | 57.58 | 9.80E-05 | V | KQIYKTPPIKDFGGFNFSQILPDPSKPS | K | 0.0000000000000002000000000000.0 | Delta:H(1)N(-1)18O(1) (N) | 1 | 1 | 801(NFS)/ | 801(NFS)/ | 1 |
| a-Lytic | 15 | 1::P0DTC2 | 325 | 1 | 482.724 | 963.4334 | 2 | 963.4337 | -0.0003 | 1098 | 1105 | 2 | 31.06 | 0.042 | S | NGTHWFVT | Q | 0.20000000.0 | Delta:H(1)N(-1)18O(1) (N) | 1 | 1 | 1098(NGT)/ | 1098(NGT)/ | 1 |

* Results shown in blue are overlapped with those for semi-search.

Supplementary Table S8. Identification of SARS CoV-2 S protein-derived glycopeptides by Byonic search

| Enzyme for database search | Confidence | Identifying Node Type | PSM Ambiguity | Peptide position from | Peptide position to | Peptide sequence | Glycosylation site | Glycan Composition | No. of HexNAc | No. of Hex | No. of Fuc | No. of NeuAc | No. of NeuGc | Annotated Sequence | Modifications | Master Protein Accessions | # Missed Cleavages | Charge | Search Engine Rank | m/z [Da] | MH+ [Da] | Theo. MH+ [Da] | DeltaM [ppm] | Deltam/z [Da] | Intensity | Activation Type | MS Order | Isolation Interference [%] | Ion Inject Time [ms] | RT [min] | First Scan | Last Scan | Master Scan(s) | \|Log Prob\| | Byonic Score | Delta Byonic Score | Delta Mod Score | PEP 2D | q-Value 2D | FDR 2D | Peptide Group FDR 2D | PEP 1D | q-Value 1D | FDR 1D | Peptide Group FDR 1D | Glycan Composition |
|---|---|---|---|---|---|---|---|---|---|---|---|---|---|---|---|---|---|---|---|---|---|---|---|---|---|---|---|---|---|---|---|---|---|---|---|---|---|---|---|---|---|---|---|---|---|---|
| chymotrypsin | High | PMI-Byonic | Unambiguous | 11 | 24 | VSSQCVNLTTRTQL | 17 | HexNAc(4)Hex(3)NeuAc(1) | 4 | 3 | | 1 | | [L].VSSQcVnLTTRTQL.[P] | C5(Carbamidomethyl); N7(HexNAc(4)Hex(3)NeuAc(1)) | P0DTC2 | 1 | 2 | 1 | 1598.7 | 3196.4 | 3196.39 | 2.14 | 0.00342 | 79706.5938 | HCD (High Energy Collision Dissociation) | MS2 | 22.3752 | 200 | 112.538 | 31336 | 31336 | 31334 | 1.85 | 13.1 | 9.8 | 9.8 | 0.01409 | 0.00726 | 0.00725 | 0.00778 | 0.74461 | 0.22955 | 0.22937 | 0.28277 | HexNAc(4)Hex(3)NeuAc(1) |
| chymotrypsin | High | PMI-Byonic | Unambiguous | 59 | 65 | FSNVTWF | 61 | HexNAc(2)Hex(5) | 2 | 5 | | | | [F].FSnVTWF.[H] | N3(HexNAc(2)Hex(5)) | P0DTC2 | 2 | 2 | 1 | 1058.93 | 2116.85 | 2116.85 | 0.14 | 0.00015 | 230329.047 | HCD (High Energy Collision Dissociation) | MS2 | 51.3729 | 170.144 | 73.2588 | 19243 | 19243 | 19220 | 5.17 | 396 | 262.9 | 262.9 | 0.0000068 | 0.0000021 | 0 | 0 | 0.00139 | 0.00044 | 0 | 0 | HexNAc(2)Hex(5) |
| chymotrypsin | High | PMI-Byonic | Unambiguous | 59 | 65 | FSNVTWF | 61 | HexNAc(2)Hex(6) | 2 | 6 | | | | [F].FSnVTWF.[H] | N3(HexNAc(2)Hex(6)) | P0DTC2 | 2 | 2 | 1 | 1139.95 | 2278.9 | 2278.9 | 0.31 | 0.00035 | 3294839.25 | HCD (High Energy Collision Dissociation) | MS2 | 10.465 | 53.637 | 73.1481 | 19206 | 19206 | 19192 | 5.78 | 505.1 | 366 | 366 | 0.0000017 | 0.00000055 | 0 | 0 | 0.00034 | 0.00011 | 0 | 0 | HexNAc(2)Hex(6) |
| chymotrypsin | High | PMI-Byonic | Unambiguous | 59 | 65 | FSNVTWF | 61 | HexNAc(2)Hex(7) | 2 | 7 | | | | [F].FSnVTWF.[H] | N3(HexNAc(2)Hex(7)) | P0DTC2 | 2 | 2 | 1 | 1220.98 | 2440.95 | 2440.95 | -0.15 | -0.0002 | 1108232.25 | HCD (High Energy Collision Dissociation) | MS2 | 30.2011 | 200 | 72.3309 | 18930 | 18930 | 18924 | 5.63 | 485.1 | 349.8 | 349.8 | 0.0000024 | 0.00000075 | 0 | 0 | 0.00048 | 0.00015 | 0 | 0 | HexNAc(2)Hex(7) |
| chymotrypsin | High | PMI-Byonic | Unambiguous | 59 | 65 | FSNVTWF | 61 | HexNAc(2)Hex(8) | 2 | 8 | | | | [F].FSnVTWF.[H] | N3(HexNAc(2)Hex(8)) | P0DTC2 | 2 | 2 | 1 | 1302 | 2603 | 2603.01 | -1.68 | -0.0022 | 484424.219 | HCD (High Energy Collision Dissociation) | MS2 | 33.2685 | 148.233 | 72.1024 | 18858 | 18858 | 18845 | 4.61 | 425.9 | 296.6 | 296.6 | 0.000025 | 0.0000051 | 0 | 0 | 0.00506 | 0.00263 | 0.00225 | 0.00339 | HexNAc(2)Hex(8) |
| chymotrypsin | High | PMI-Byonic | Unambiguous | 59 | 65 | FSNVTWF | 61 | HexNAc(3)Hex(4)Fuc(1) | 3 | 4 | 1 | | | [F].FSnVTWF.[H] | N3(HexNAc(3)Hex(4)Fuc(1)) | P0DTC2 | 2 | 2 | 1 | 1152.47 | 2303.92 | 2303.93 | -3.68 | -0.0042 | 219117.516 | HCD (High Energy Collision Dissociation) | MS2 | 20.6773 | 200 | 73.3271 | 19265 | 19265 | 19247 | 3.77 | 336.6 | 185.6 | 185.6 | 0.00017 | 0.000021 | 0 | 0 | 0.03338 | 0.01701 | 0.0165 | 0.02041 | HexNAc(3)Hex(4)Fuc(1) |
| chymotrypsin | High | PMI-Byonic | Unambiguous | 59 | 65 | FSNVTWF | 61 | HexNAc(3)Hex(5)Fuc(1) | 3 | 5 | 1 | | | [F].FSnVTWF.[H] | N3(HexNAc(3)Hex(5)Fuc(1)) | P0DTC2 | 2 | 2 | 1 | 1233.5 | 2465.98 | 2465.99 | -1 | -0.0012 | 2489586.5 | HCD (High Energy Collision Dissociation) | MS2 | 11.8005 | 32.944 | 72.7501 | 19063 | 19063 | 19048 | 4.85 | 403.8 | 274.6 | 274.6 | 0.000014 | 0.0000034 | 0 | 0 | 0.00292 | 0.00263 | 0.00252 | 0.00391 | HexNAc(3)Hex(5)Fuc(1) |
| chymotrypsin | High | PMI-Byonic | Unambiguous | 59 | 65 | FSNVTWF | 61 | HexNAc(3)Hex(5) | 3 | 5 | | | | [F].FSnVTWF.[H] | N3(HexNAc(3)Hex(5)) | P0DTC2 | 2 | 2 | 1 | 1160.47 | 2319.93 | 2319.93 | -0.2 | -0.0002 | 206834.563 | HCD (High Energy Collision Dissociation) | MS2 | 48.731 | 192.712 | 73.1836 | 19218 | 19218 | 19192 | 5.1 | 394.7 | 223.5 | 223.5 | 0.000008 | 0.0000024 | 0 | 0 | 0.00163 | 0.00049 | 0 | 0 | HexNAc(3)Hex(5) |
| chymotrypsin | High | PMI-Byonic | Unambiguous | 59 | 65 | FSNVTWF | 61 | HexNAc(3)Hex(6)Fuc(1) | 3 | 6 | 1 | | | [F].FSnVTWF.[H] | N3(HexNAc(3)Hex(6)Fuc(1)) | P0DTC2 | 2 | 2 | 1 | 1314.52 | 2628.04 | 2628.04 | -0.6 | -0.0008 | 1548230 | HCD (High Energy Collision Dissociation) | MS2 | 2.02464 | 70.555 | 72.1863 | 18885 | 18885 | 18874 | 4.83 | 411.3 | 281.3 | 281.3 | 0.000015 | 0.0000036 | 0 | 0 | 0.00304 | 0.00263 | 0.00248 | 0.00392 | HexNAc(3)Hex(6)Fuc(1) |
| chymotrypsin | High | PMI-Byonic | Unambiguous | 59 | 65 | FSNVTWF | 61 | HexNAc(3)Hex(6)NeuGc(1) | 3 | 6 | | | 1 | [F].FSnVTWF.[H] | N3(HexNAc(3)Hex(6)NeuGc(1)) | P0DTC2 | 2 | 2 | 1 | 1395.04 | 2789.07 | 2789.07 | -0.37 | -0.0005 | 153476.813 | HCD (High Energy Collision Dissociation) | MS2 | 33.481 | 144.444 | 77.282 | 20579 | 20579 | 20558 | 5.33 | 378.9 | 251.3 | 251.3 | 0.0000046 | 0.0000016 | 0 | 0 | 0.00095 | 0.00033 | 0 | 0 | HexNAc(3)Hex(6)NeuGc(1) |
| chymotrypsin | High | PMI-Byonic | Unambiguous | 59 | 65 | FSNVTWF | 61 | HexNAc(3)Hex(6) | 3 | 6 | | | | [F].FSnVTWF.[H] | N3(HexNAc(3)Hex(6)) | P0DTC2 | 2 | 2 | 1 | 1241.49 | 2481.97 | 2481.98 | -2.19 | -0.0027 | 1003532.63 | HCD (High Energy Collision Dissociation) | MS2 | 16.4848 | 30.271 | 72.758 | 19067 | 19067 | 19048 | 4.37 | 381.7 | 262.9 | 262.9 | 0.000043 | 0.0000083 | 0 | 0 | 0.00871 | 0.01033 | 0.00992 | 0.00893 | HexNAc(3)Hex(6) |
| chymotrypsin | High | PMI-Byonic | Unambiguous | 59 | 65 | FSNVTWF | 61 | HexNAc(4)Hex(4)Fuc(1) | 4 | 4 | 1 | | | [F].FSnVTWF.[H] | N3(HexNAc(4)Hex(4)Fuc(1)) | P0DTC2 | 2 | 2 | 1 | 1254.01 | 2507.01 | 2507.01 | 0.6 | 0.00075 | 254520.016 | HCD (High Energy Collision Dissociation) | MS2 | 48.3508 | 176.781 | 72.8465 | 19098 | 19098 | 19079 | 4.74 | 275.8 | 175 | 175 | 0.000018 | 0.000004 | 0 | 0 | 0.00374 | 0.00263 | 0.0024 | 0.00364 | HexNAc(4)Hex(4)Fuc(1) |
| chymotrypsin | High | PMI-Byonic | Unambiguous | 59 | 65 | FSNVTWF | 61 | HexNAc(4)Hex(5)Fuc(2) | 4 | 5 | 2 | | | [F].FSnVTWF.[H] | N3(HexNAc(4)Hex(5)Fuc(2)) | P0DTC2 | 2 | 3 | 1 | 939.046 | 2815.12 | 2815.12 | -0.15 | -0.0001 | 204971.25 | HCD (High Energy Collision Dissociation) | MS2 | 26.8118 | 200 | 71.5143 | 18664 | 18664 | 18637 | 4.77 | 383.9 | 256 | 256 | 0.000017 | 0.0000038 | 0 | 0 | 0.00343 | 0.00263 | 0.00243 | 0 | HexNAc(4)Hex(5)Fuc(2) |
| chymotrypsin | High | PMI-Byonic | Unambiguous | 59 | 65 | FSNVTWF | 61 | HexNAc(4)Hex(5)Fuc(3) | 4 | 5 | 3 | | | [F].FSnVTWF.[H] | N3(HexNAc(4)Hex(5)Fuc(3)) | P0DTC2 | 2 | 3 | 1 | 987.731 | 2961.18 | 2961.18 | -1.12 | -0.0011 | 185613.547 | HCD (High Energy Collision Dissociation) | MS2 | 13.7627 | 117.268 | 70.9104 | 18464 | 18464 | 18444 | 4.52 | 385.7 | 254.8 | 254.8 | 0.00003 | 0.0000061 | 0 | 0 | 0.00619 | 0.00448 | 0.0043 | 0.00656 | HexNAc(4)Hex(5)Fuc(3) |
| [illegible] | [illegible] | [illegible] | [illegible] | [illegible] | [illegible] | [illegible] | [illegible] | [illegible] | [illegible] | [illegible] | [illegible] | [illegible] | [illegible] | [illegible] | [illegible] | [illegible] | [illegible] | [illegible] | [illegible] | [illegible] | [illegible] | [illegible] | [illegible] | [illegible] | [illegible] | [illegible] | [illegible] | [illegible] | [illegible] | [illegible] | [illegible] | [illegible] | [illegible] | [illegible] | [illegible] | [illegible] | [illegible] | [illegible] | [illegible] | [illegible] | [illegible] | [illegible] | [illegible] | [illegible] | [illegible] | [illegible] |

| | | | | | | | | |
|---|---|---|---|---|---|---|---|---|
| chymotrypsin | High | PMI-Byonic | Unambiguous | 593 | 612 | GGVSVITPGTNTSNQVAVLY | 603 | HexNAc(4)Hex(5) |
| chymotrypsin | High | PMI-Byonic | Unambiguous | 593 | 612 | GGVSVITPGTNTSNQVAVLY | 603 | HexNAc(5)Hex(4)Fuc(1) |
| chymotrypsin | High | PMI-Byonic | Unambiguous | 593 | 612 | GGVSVITPGTNTSNQVAVLY | 603 | HexNAc(5)Hex(5)Fuc(1) |
| chymotrypsin | High | PMI-Byonic | Unambiguous | 593 | 612 | GGVSVITPGTNTSNQVAVLY | 603 | HexNAc(5)Hex(6)Fuc(2)[illegible] |
| chymotrypsin | High | PMI-Byonic | Unambiguous | 593 | 612 | GGVSVITPGTNTSNQVAVLY | 603 | HexNAc(5)Hex(6)Fuc(3) |
| chymotrypsin | High | PMI-Byonic | Unambiguous | 593 | 611 | GGVSVITPGTNTSNQVAVL | 603 | HexNAc(5)Hex(6)NeuAc(1) |
| chymotrypsin | High | PMI-Byonic | Unambiguous | 593 | 611 | GGVSVITPGTNTSNQVAVL | 603 | HexNAc(6)Hex(7)Fuc(1) |
| chymotrypsin | High | PMI-Byonic | Unambiguous | 613 | 633 | QDVNCTEVPVAIHADQLTPTW | 616 | HexNAc(3)Hex(5)Fuc(1) |
| chymotrypsin | High | PMI-Byonic | Unambiguous | 613 | 633 | QDVNCTEVPVAIHADQLTPTW | 616 | HexNAc(3)Hex(5) |
| chymotrypsin | High | PMI-Byonic | Unambiguous | 613 | 633 | QDVNCTEVPVAIHADQLTPTW | 616 | HexNAc(4)Hex(5)Fuc(1)[illegible] |
| chymotrypsin | High | PMI-Byonic | Unambiguous | 612 | 633 | YQDVNCTEVPVAIHADQLTPTW | 616 | HexNAc(4)Hex(5)Fuc(1) |
| chymotrypsin | High | PMI-Byonic | Unambiguous | 612 | 633 | YQDVNCTEVPVAIHADQLTPTW | 616 | HexNAc(4)Hex(5)Fuc(2) |
| chymotrypsin | High | PMI-Byonic | Unambiguous | 612 | 633 | YQDVNCTEVPVAIHADQLTPTW | 616 | HexNAc(4)Hex(5) |
| chymotrypsin | High | PMI-Byonic | Unambiguous | 613 | 633 | QDVNCTEVPVAIHADQLTPTW | 616 | HexNAc(4)Hex(6) |
| chymotrypsin | High | PMI-Byonic | Unambiguous | 613 | 633 | QDVNCTEVPVAIHADQLTPTW | 616 | HexNAc(5)Hex(4)Fuc(1) |
| chymotrypsin | High | PMI-Byonic | Unambiguous | 612 | 633 | YQDVNCTEVPVAIHADQLTPTW | 616 | HexNAc(5)Hex(4)NeuGc(1) |
| chymotrypsin | High | PMI-Byonic | Unambiguous | 612 | 633 | YQDVNCTEVPVAIHADQLTPTW | 616 | HexNAc(5)Hex(5)Fuc(1) |
| chymotrypsin | High | PMI-Byonic | Unambiguous | 613 | 633 | QDVNCTEVPVAIHADQLTPTW | 616 | HexNAc(5)Hex(5) |
| chymotrypsin | High | PMI-Byonic | Unambiguous | 613 | 633 | QDVNCTEVPVAIHADQLTPTW | 616 | HexNAc(5)Hex(6)Fuc(1)[illegible] |
| chymotrypsin | High | PMI-Byonic | Unambiguous | 612 | 633 | YQDVNCTEVPVAIHADQLTPTW | 616 | HexNAc(5)Hex(6)Fuc(1) |
| chymotrypsin | High | PMI-Byonic | Unambiguous | 613 | 633 | QDVNCTEVPVAIHADQLTPTW | 616 | HexNAc(5)Hex(6)NeuAc(1) |
| chymotrypsin | High | PMI-Byonic | Unambiguous | 612 | 633 | YQDVNCTEVPVAIHADQLTPTW | 616 | HexNAc(5)Hex(6) |
| chymotrypsin | High | PMI-Byonic | Unambiguous | 613 | 633 | QDVNCTEVPVAIHADQLTPTW | 616 | HexNAc(6)Hex(6)Fuc(2) |
| chymotrypsin | High | PMI-Byonic | Unambiguous | 612 | 633 | YQDVNCTEVPVAIHADQLTPTW | 616 | HexNAc(6)Hex(7)Fuc(1) |
| [illegible] | [illegible] | [illegible] | [illegible] | [illegible] | [illegible] | [illegible] | [illegible] | [illegible] |

| | | | | | | | | | | | | | | | |
|---|---|---|---|---|---|---|---|---|---|---|---|---|---|---|---|
| chymotrypsin(semi) | High | PMI-Byonic | Unambiguous | 59 | 65 | FSNVTWF | 61 | HexNAc(4)Hex(6)NeuAc(1) | 4 | 6 | | 1 | | [F].FSnVTWF.[H] | [illegible] |
| chymotrypsin(semi) | High | PMI-Byonic | Unambiguous | 59 | 65 | FSNVTWF | 61 | HexNAc(4)Hex(7)Fuc(2) | 4 | 7 | 2 | | | [F].FSnVTWF.[H] | [illegible] |
| chymotrypsin(semi) | High | PMI-Byonic | Unambiguous | 59 | 65 | FSNVTWF | 61 | HexNAc(5)Hex(4)Fuc(2) | 5 | 4 | 2 | | | [F].FSnVTWF.[H] | [illegible] |
| chymotrypsin(semi) | High | PMI-Byonic | Unambiguous | 59 | 64 | FSNVTW | 61 | HexNAc(5)Hex(5)Fuc(2) | 5 | 5 | 2 | | | [F].FSnVTW.[F] | [illegible] |
| chymotrypsin(semi) | High | PMI-Byonic | Unambiguous | 59 | 65 | FSNVTWF | 61 | HexNAc(5)Hex(5)Fuc(3) | 5 | 5 | 3 | | | [F].FSnVTWF.[H] | [illegible] |
| chymotrypsin(semi) | High | PMI-Byonic | Unambiguous | 59 | 65 | FSNVTWF | 61 | HexNAc(5)Hex(5) | 5 | 5 | | | | [F].FSnVTWF.[H] | [illegible] |
| chymotrypsin(semi) | High | PMI-Byonic | Unambiguous | 59 | 65 | FSNVTWF | 61 | HexNAc(5)Hex(6)Fuc(1) | 5 | 6 | 1 | | | [F].FSnVTWF.[H] | [illegible] |
| chymotrypsin(semi) | High | PMI-Byonic | Unambiguous | 59 | 65 | FSNVTWF | 61 | HexNAc(5)Hex(6)Fuc(2) | 5 | 6 | 2 | | | [F].FSnVTWF.[H] | [illegible] |
| chymotrypsin(semi) | High | PMI-Byonic | Unambiguous | 59 | 65 | FSNVTWF | 61 | HexNAc(5)Hex(6)Fuc(3) | 5 | 6 | 3 | | | [F].FSnVTWF.[H] | [illegible] |
| chymotrypsin(semi) | High | PMI-Byonic | Unambiguous | 59 | 65 | FSNVTWF | 61 | HexNAc(5)Hex(6) | 5 | 6 | | | | [F].FSnVTWF.[H] | [illegible] |
| chymotrypsin(semi) | High | PMI-Byonic | Unambiguous | 59 | 64 | FSNVTW | 61 | HexNAc(6)Hex(4) | 6 | 4 | | | | [F].FSnVTW.[F] | [illegible] |
| chymotrypsin(semi) | High | PMI-Byonic | Unambiguous | 59 | 65 | FSNVTWF | 61 | HexNAc(6)Hex(6)Fuc(1) | 6 | 6 | 1 | | | [F].FSnVTWF.[H] | [illegible] |
| chymotrypsin(semi) | High | PMI-Byonic | Unambiguous | 59 | 65 | FSNVTWF | 61 | HexNAc(6)Hex(6)Fuc(2) | 6 | 6 | 2 | | | [F].FSnVTWF.[H] | [illegible] |
| chymotrypsin(semi) | High | PMI-Byonic | Unambiguous | 59 | 64 | FSNVTW | 61 | HexNAc(6)Hex(6)Fuc(3) | 6 | 6 | 3 | | | [F].FSnVTW.[F] | [illegible] |
| chymotrypsin(semi) | High | PMI-Byonic | Unambiguous | 59 | 65 | FSNVTWF | 61 | HexNAc(6)Hex(6) | 6 | 6 | | | | [F].FSnVTWF.[H] | [illegible] |
| chymotrypsin(semi) | High | PMI-Byonic | Unambiguous | 59 | 77 | FSNVTWFHAIHVSGTNGTK | 61 | HexNAc(6)Hex(8)Fuc(1)NeuAc(1) | 6 | 8 | 1 | 1 | | [F].FSnVTWFHAIHVSGTNGTK.[R] | [illegible] |
| chymotrypsin(semi) | High | PMI-Byonic | Unambiguous | 59 | 66 | FSNVTWFH | 61 | HexNAc(8)Hex(5) | 8 | 5 | | | | [F].FSnVTWFH.[A] | [illegible] |
| chymotrypsin(semi) | High | PMI-Byonic | Unambiguous | 59 | 65 | FSNVTWF | 61 | HexNAc(9)Hex(3)Fuc(1) | 9 | 3 | 1 | | | [F].FSnVTWF.[H] | [illegible] |
| chymotrypsin(semi) | High | PMI-Byonic | Unambiguous | 66 | 79 | HAIHVSGTNGTKRF | 74 | HexNAc(2)Hex(8) | 2 | 8 | | | | [F].HAIHVSGTnGTKRF.[D] | [illegible] |
| chymotrypsin(semi) | High | PMI-Byonic | Unambiguous | 66 | 79 | HAIHVSGTNGTKRF | 74 | HexNAc(4)Hex(3)Fuc(1) | 4 | 3 | 1 | | | [F].HAIHVSGTnGTKRF.[D] | [illegible] |
| chymotrypsin(semi) | High | PMI-Byonic | Unambiguous | 66 | 79 | HAIHVSGTNGTKRF | 74 | HexNAc(4)Hex(3)Fuc(2) | 4 | 3 | 2 | | | [F].HAIHVSGTnGTKRF.[D] | [illegible] |
| chymotrypsin(semi) | High | PMI-Byonic | Unambiguous | 66 | 79 | HAIHVSGTNGTKRF | 74 | HexNAc(4)Hex(4)Fuc(1)NeuGc(1) | 4 | 4 | 1 | | 1 | [F].HAIHVSGTnGTKRF.[D] | [illegible] |
| chymotrypsin(semi) | High | PMI-Byonic | Unambiguous | 66 | 79 | HAIHVSGTNGTKRF | 74 | HexNAc(4)Hex(4)Fuc(1) | 4 | 4 | 1 | | | [F].HAIHVSGTnGTKRF.[D] | [illegible] |
| chymotrypsin(semi) | High | PMI-Byonic | Unambiguous | 66 | 79 | HAIHVSGTNGTKRF | 74 | HexNAc(4)Hex(4)Fuc(2) | 4 | 4 | 2 | | | [F].HAIHVSGTnGTKRF.[D] | [illegible] |
| chymotrypsin(semi) | High | PMI-Byonic | Unambiguous | 66 | 91 | HAIHVSGTNGTKRFDNPVLPFNDGVY | 74 | HexNAc(4)Hex(5)Fuc(1)NeuAc(1) | 4 | 5 | 1 | 1 | | [illegible] | [illegible] |
| chymotrypsin(semi) | High | PMI-Byonic | Unambiguous | 66 | 79 | HAIHVSGTNGTKRF | 74 | HexNAc(4)Hex(5)Fuc(1)NeuGc(1) | 4 | 5 | 1 | | 1 | [F].HAIHVSGTnGTKRF.[D] | [illegible] |
| chymotrypsin(semi) | High | PMI-Byonic | Unambiguous | 66 | 79 | HAIHVSGTNGTKRF | 74 | HexNAc(4)Hex(5)Fuc(1)NeuGc(2) | 4 | 5 | 1 | | 2 | [F].HAIHVSGTnGTKRF.[D] | [illegible] |
| chymotrypsin(semi) | High | PMI-Byonic | Unambiguous | 66 | 79 | HAIHVSGTNGTKRF | 74 | HexNAc(4)Hex(5)Fuc(1) | 4 | 5 | 1 | | | [F].HAIHVSGTnGTKRF.[D] | [illegible] |
| chymotrypsin(semi) | High | PMI-Byonic | Unambiguous | 66 | 79 | HAIHVSGTNGTKRF | 74 | HexNAc(4)Hex(5)Fuc(2)NeuGc(1) | 4 | 5 | 2 | | 1 | [F].HAIHVSGTnGTKRF.[D] | [illegible] |
| chymotrypsin(semi) | High | PMI-Byonic | Unambiguous | 66 | 79 | HAIHVSGTNGTKRF | 74 | HexNAc(4)Hex(5)Fuc(2) | 4 | 5 | 2 | | | [F].HAIHVSGTnGTKRF.[D] | [illegible] |
| [illegible] | | | | | | | | | | | | | | | |

| | | | | | | | | | | | | |
|---|---|---|---|---|---|---|---|---|---|---|---|---|
| chymotrypsin(semi) | High | PMI-Byonic | Unambiguous | 319 | 338 | RVQPTESIVRFPNITNLCPF | 331 | HexNAc(4)Hex(5)Fuc(1)NeuAc(1) | 4 | 5 | 1 | 1 |
| chymotrypsin(semi) | High | PMI-Byonic | Unambiguous | 319 | 342 | RVQPTESIVRFPNITNLCPFGEVF | 331 | HexNAc(4)Hex(5)Fuc(1)NeuGc(1) | 4 | 5 | 1 | |
| chymotrypsin(semi) | High | PMI-Byonic | Unambiguous | 319 | 342 | RVQPTESIVRFPNITNLCPFGEVF | 331 | HexNAc(4)Hex(5)Fuc(1) | 4 | 5 | 1 | |
| chymotrypsin(semi) | High | PMI-Byonic | Unambiguous | 319 | 342 | RVQPTESIVRFPNITNLCPFGEVF | 331 | HexNAc(4)Hex(5)Fuc(2) | 4 | 5 | 2 | |
| chymotrypsin(semi) | High | PMI-Byonic | Unambiguous | 319 | 342 | RVQPTESIVRFPNITNLCPFGEVF | 331 | HexNAc(4)Hex(5)Fuc(3) | 4 | 5 | 3 | |
| chymotrypsin(semi) | High | PMI-Byonic | Unambiguous | 319 | 342 | RVQPTESIVRFPNITNLCPFGEVF | 331 | HexNAc(4)Hex(6)Fuc(1)NeuAc(1) | 4 | 6 | 1 | 1 |
| chymotrypsin(semi) | High | PMI-Byonic | Unambiguous | 319 | 342 | RVQPTESIVRFPNITNLCPFGEVF | 331 | HexNAc(4)Hex(6)Fuc(2) | 4 | 6 | 2 | |
| chymotrypsin(semi) | High | PMI-Byonic | Unambiguous | 319 | 342 | RVQPTESIVRFPNITNLCPFGEVF | 331 | HexNAc(5)Hex(4)Fuc(1) | 5 | 4 | 1 | |
| chymotrypsin(semi) | High | PMI-Byonic | Unambiguous | 319 | 342 | RVQPTESIVRFPNITNLCPFGEVF | 331 | HexNAc(5)Hex(5)Fuc(1)NeuAc(1) | 5 | 5 | 1 | 1 |
| chymotrypsin(semi) | High | PMI-Byonic | Unambiguous | 319 | 342 | RVQPTESIVRFPNITNLCPFGEVF | 331 | HexNAc(5)Hex(5)Fuc(1) | 5 | 5 | 1 | |
| chymotrypsin(semi) | High | PMI-Byonic | Unambiguous | 319 | 338 | RVQPTESIVRFPNITNLCPF | 331 | HexNAc(5)Hex(5)Fuc(3) | 5 | 5 | 3 | |
| chymotrypsin(semi) | High | PMI-Byonic | Unambiguous | 319 | 338 | RVQPTESIVRFPNITNLCPF | 331 | HexNAc(5)Hex(6)Fuc(1)NeuAc(1) | 5 | 6 | 1 | 1 |
| chymotrypsin(semi) | High | PMI-Byonic | Unambiguous | 319 | 342 | RVQPTESIVRFPNITNLCPFGEVF | 331 | HexNAc(5)Hex(6)Fuc(1)NeuGc(1) | 5 | 6 | 1 | |
| chymotrypsin(semi) | High | PMI-Byonic | Unambiguous | 319 | 342 | RVQPTESIVRFPNITNLCPFGEVF | 331 | HexNAc(5)Hex(6)Fuc(1) | 5 | 6 | 1 | |
| chymotrypsin(semi) | High | PMI-Byonic | Unambiguous | 319 | 338 | RVQPTESIVRFPNITNLCPF | 331 | HexNAc(5)Hex(6)Fuc(2)NeuAc(1) | 5 | 6 | 2 | 1 |
| chymotrypsin(semi) | High | PMI-Byonic | Unambiguous | 319 | 338 | RVQPTESIVRFPNITNLCPF | 331 | HexNAc(5)Hex(6)Fuc(2) | 5 | 6 | 2 | |
| chymotrypsin(semi) | High | PMI-Byonic | Unambiguous | 319 | 342 | RVQPTESIVRFPNITNLCPFGEVF | 331 | HexNAc(5)Hex(6)Fuc(3) | 5 | 6 | 3 | |
| chymotrypsin(semi) | High | PMI-Byonic | Unambiguous | 319 | 342 | RVQPTESIVRFPNITNLCPFGEVF | 331 | HexNAc(6)Hex(6)Fuc(1) | 6 | 6 | 1 | |
| chymotrypsin(semi) | High | PMI-Byonic | Unambiguous | 319 | 338 | RVQPTESIVRFPNITNLCPF | 331 | HexNAc(6)Hex(6)Fuc(2) | 6 | 6 | 2 | |
| chymotrypsin(semi) | High | PMI-Byonic | Unambiguous | 319 | 342 | RVQPTESIVRFPNITNLCPFGEVF | 331 | HexNAc(6)Hex(7)Fuc(1)NeuAc(1) | 6 | 7 | 1 | 1 |
| chymotrypsin(semi) | High | PMI-Byonic | Unambiguous | 319 | 342 | RVQPTESIVRFPNITNLCPFGEVF | 331 | HexNAc(6)Hex(7)Fuc(4) | 6 | 7 | 4 | |
| chymotrypsin(semi) | High | PMI-Byonic | Unambiguous | 319 | 338 | RVQPTESIVRFPNITNLCPF | 331 | HexNAc(6)Hex(7)NeuAc(1) | 6 | 7 | | 1 |
| chymotrypsin(semi) | High | PMI-Byonic | Unambiguous | 319 | 342 | RVQPTESIVRFPNITNLCPFGEVF | 331 | HexNAc(8)Hex(9) | 8 | 9 | | |
| chymotrypsin(semi) | High | PMI-Byonic | Unambiguous | 339 | 347 | GEVFNATRF | 343 | HexNAc(2)Hex(5) | 2 | 5 | | |

| | | | | | | | | |
|---|---|---|---|---|---|---|---|---|
| chymotrypsin(semi) | High | PMI-Byonic | Unambiguous | 1068 | 1075 | VPAQEKNF | 1074 | HexNAc(4)Hex(4)Fuc(1) |
| chymotrypsin(semi) | High | PMI-Byonic | Unambiguous | 1068 | 1075 | VPAQEKNF | 1074 | HexNAc(4)Hex(4) |
| chymotrypsin(semi) | High | PMI-Byonic | Unambiguous | 1064 | 1083 | HVTYVPAQEKNFTTAPAICH | 1074 | [illegible] |
| chymotrypsin(semi) | High | PMI-Byonic | Unambiguous | 1068 | 1086 | VPAQEKNFTTAPAICHDGK | 1074 | [illegible] |
| chymotrypsin(semi) | High | PMI-Byonic | Unambiguous | 1064 | 1083 | HVTYVPAQEKNFTTAPAICH | 1074 | [illegible] |
| chymotrypsin(semi) | High | PMI-Byonic | Unambiguous | 1068 | 1075 | VPAQEKNF | 1074 | [illegible] |
| chymotrypsin(semi) | High | PMI-Byonic | Unambiguous | 1063 | 1083 | LHVTYVPAQEKNFTTAPAICH | 1074 | HexNAc(4)Hex(5)Fuc(1) |
| chymotrypsin(semi) | High | PMI-Byonic | Unambiguous | 1068 | 1075 | VPAQEKNF | 1074 | [illegible] |
| chymotrypsin(semi) | High | PMI-Byonic | Unambiguous | 1064 | 1075 | HVTYVPAQEKNF | 1074 | HexNAc(4)Hex(5)Fuc(2) |
| chymotrypsin(semi) | High | PMI-Byonic | Unambiguous | 1068 | 1083 | VPAQEKNFTTAPAICH | 1074 | [illegible] |
| chymotrypsin(semi) | High | PMI-Byonic | Unambiguous | 1068 | 1083 | VPAQEKNFTTAPAICH | 1074 | [illegible] |
| chymotrypsin(semi) | High | PMI-Byonic | Unambiguous | 1063 | 1075 | LHVTYVPAQEKNF | 1074 | HexNAc(4)Hex(5)Fuc(3) |
| chymotrypsin(semi) | High | PMI-Byonic | Unambiguous | 1068 | 1080 | VPAQEKNFTTAPA | 1074 | [illegible] |
| chymotrypsin(semi) | High | PMI-Byonic | Unambiguous | 1063 | 1082 | LHVTYVPAQEKNFTTAPAIC | 1074 | HexNAc(4)Hex(5)NeuAc(2) |
| chymotrypsin(semi) | High | PMI-Byonic | Unambiguous | 1068 | 1075 | VPAQEKNF | 1074 | HexNAc(4)Hex(5)NeuGc(1) |
| chymotrypsin(semi) | High | PMI-Byonic | Unambiguous | 1063 | 1075 | LHVTYVPAQEKNF | 1074 | HexNAc(4)Hex(5) |
| chymotrypsin(semi) | High | PMI-Byonic | Unambiguous | 1068 | 1075 | VPAQEKNF | 1074 | HexNAc(4)Hex(6)Fuc(1) |
| chymotrypsin(semi) | High | PMI-Byonic | Unambiguous | 1068 | 1075 | VPAQEKNF | 1074 | HexNAc(4)Hex(6)Fuc(2) |
| chymotrypsin(semi) | High | PMI-Byonic | Unambiguous | 1068 | 1075 | VPAQEKNF | 1074 | HexNAc(4)Hex(6)Fuc(3) |
| chymotrypsin(semi) | High | PMI-Byonic | Unambiguous | 1068 | 1095 | VPAQEKNFTTAPAICHDGKAHFPREGVF | 1074 | HexNAc(4)Hex(6)NeuAc(1) |
| chymotrypsin(semi) | High | PMI-Byonic | Unambiguous | 1068 | 1075 | VPAQEKNF | 1074 | HexNAc(4)Hex(6) |
| chymotrypsin(semi) | High | PMI-Byonic | Unambiguous | 1066 | 1095 | TYVPAQEKNFTTAPAICHDGKAHFPREGVF | 1074 | HexNAc(4)Hex(7)NeuAc(1) |
| chymotrypsin(semi) | High | PMI-Byonic | Unambiguous | 1068 | 1075 | VPAQEKNF | 1074 | HexNAc(4)Hex(7) |
| chymotrypsin(semi) | High | PMI-Byonic | Unambiguous | 1068 | 1075 | VPAQEKNF | 1074 | HexNAc(5)Hex(3)Fuc(1) |
| chymotrypsin(semi) | High | PMI-Byonic | Unambiguous | 1064 | 1075 | HVTYVPAQEKNF | 1074 | HexNAc(5)Hex(4)Fuc(1) |

[illegible]

| | | | | | | | | | | | | | | | | | | | | |
|---|---|---|---|---|---|---|---|---|---|---|---|---|---|---|---|---|---|---|---|---|
| Trypsin | High | PMI-Byonic | Unambiguous | 159 | 182 | VYSSANNCTFEYVSQPFLMDLEGK | 165 | HexNAc(5)Hex(6) | 5 | 6 | | | | [R].VYSSANncTFEYVSQPFLMDLEGK.[Q] | N7(HexNAc(5)Hex(6)); C8(Carbamidomethyl) | P0DTC2 | 0 | 3 | 1 | [illegible] |
| Trypsin | High | PMI-Byonic | Unambiguous | 159 | 182 | VYSSANNCTFEYVSQPFLMDLEGK | 165 | HexNAc(6)Hex(6)Fuc(1) | 6 | 6 | 1 | | | [R].VYSSANncTFEYVSQPFLMDLEGK.[Q] | N7(HexNAc(6)Hex(6)Fuc(1)); C8(Carbamidomethyl) | P0DTC2 | 0 | 4 | 1 | [illegible] |
| Trypsin | High | PMI-Byonic | Unambiguous | 159 | 182 | VYSSANNCTFEYVSQPFLMDLEGK | 165 | HexNAc(6)Hex(7)Fuc(1)NeuAc(1) | 6 | 7 | 1 | 1 | | [R].VYSSANncTFEYVSQPFLMDLEGK.[Q] | N7(HexNAc(6)Hex(7)Fuc(1)NeuAc(1)); C8(Carbamidomethyl) | P0DTC2 | 0 | 3 | 1 | [illegible] |
| Trypsin | High | PMI-Byonic | Unambiguous | 159 | 182 | VYSSANNCTFEYVSQPFLMDLEGK | 165 | HexNAc(6)Hex(7)Fuc(1) | 6 | 7 | 1 | | | [R].VYSSANncTFEYVSQPFLMDLEGK.[Q] | N7(HexNAc(6)Hex(7)Fuc(1)); C8(Carbamidomethyl) | P0DTC2 | 0 | 3 | 1 | [illegible] |
| Trypsin | High | PMI-Byonic | Unambiguous | 159 | 182 | VYSSANNCTFEYVSQPFLMDLEGK | 165 | HexNAc(6)Hex(7)Fuc(4) | 6 | 7 | 4 | | | [R].VYSSANncTFEYVSQPFLMDLEGK.[Q] | N7(HexNAc(6)Hex(7)Fuc(4)); C8(Carbamidomethyl) | P0DTC2 | 0 | 4 | 1 | [illegible] |
| Trypsin | High | PMI-Byonic | Unambiguous | 159 | 182 | VYSSANNCTFEYVSQPFLMDLEGK | 165 | HexNAc(8)Hex(9) | 8 | 9 | | | | [R].VYSSANncTFEYVSQPFLMDLEGK.[Q] | N7(HexNAc(8)Hex(9)); C8(Carbamidomethyl) | P0DTC2 | 0 | 4 | 1 | [illegible] |
| Trypsin | High | PMI-Byonic | Unambiguous | 215 | 237 | DLPQGFSALEPLVDLPIGINITR | 234 | HexNAc(2)Hex(5) | 2 | 5 | | | | [R].DLPQGFSALEPLVDLPIGInITR.[F] | N20(HexNAc(2)Hex(5)) | P0DTC2 | 0 | 3 | 1 | [illegible] |
| Trypsin | High | PMI-Byonic | Unambiguous | 215 | 237 | DLPQGFSALEPLVDLPIGINITR | 234 | HexNAc(2)Hex(6) | 2 | 6 | | | | [R].DLPQGFSALEPLVDLPIGInITR.[F] | N20(HexNAc(2)Hex(6)) | P0DTC2 | 0 | 3 | 1 | [illegible] |
| Trypsin | High | PMI-Byonic | Unambiguous | 215 | 237 | DLPQGFSALEPLVDLPIGINITR | 234 | HexNAc(2)Hex(7) | 2 | 7 | | | | [R].DLPQGFSALEPLVDLPIGInITR.[F] | N20(HexNAc(2)Hex(7)) | P0DTC2 | 0 | 3 | 1 | [illegible] |
| Trypsin | High | PMI-Byonic | Unambiguous | 215 | 237 | DLPQGFSALEPLVDLPIGINITR | 234 | HexNAc(2)Hex(8) | 2 | 8 | | | | [R].DLPQGFSALEPLVDLPIGInITR.[F] | N20(HexNAc(2)Hex(8)) | P0DTC2 | 0 | 3 | 1 | [illegible] |
| Trypsin | High | PMI-Byonic | Unambiguous | 215 | 237 | DLPQGFSALEPLVDLPIGINITR | 234 | HexNAc(2)Hex(9) | 2 | 9 | | | | [R].DLPQGFSALEPLVDLPIGInITR.[F] | N20(HexNAc(2)Hex(9)) | P0DTC2 | 0 | 3 | 1 | [illegible] |
| Trypsin | High | PMI-Byonic | Unambiguous | 215 | 237 | DLPQGFSALEPLVDLPIGINITR | 234 | HexNAc(3)Hex(5)Fuc(1) | 3 | 5 | 1 | | | [R].DLPQGFSALEPLVDLPIGInITR.[F] | N20(HexNAc(3)Hex(5)Fuc(1)) | P0DTC2 | 0 | 3 | 1 | [illegible] |
| Trypsin | High | PMI-Byonic | Unambiguous | 215 | 237 | DLPQGFSALEPLVDLPIGINITR | 234 | HexNAc(3)Hex(5) | 3 | 5 | | | | [R].DLPQGFSALEPLVDLPIGInITR.[F] | N20(HexNAc(3)Hex(5)) | P0DTC2 | 0 | 3 | 1 | [illegible] |
| Trypsin | High | PMI-Byonic | Unambiguous | 215 | 237 | DLPQGFSALEPLVDLPIGINITR | 234 | HexNAc(3)Hex(6) | 3 | 6 | | | | [R].DLPQGFSALEPLVDLPIGInITR.[F] | N20(HexNAc(3)Hex(6)) | P0DTC2 | 0 | 3 | 1 | [illegible] |
| Trypsin | High | PMI-Byonic | Unambiguous | 215 | 237 | DLPQGFSALEPLVDLPIGINITR | 234 | HexNAc(4)Hex(3)Fuc(1) | 4 | 3 | 1 | | | [R].DLPQGFSALEPLVDLPIGInITR.[F] | N20(HexNAc(4)Hex(3)Fuc(1)) | P0DTC2 | 0 | 3 | 1 | [illegible] |
| Trypsin | High | PMI-Byonic | Unambiguous | 215 | 237 | DLPQGFSALEPLVDLPIGINITR | 234 | HexNAc(4)Hex(4) | 4 | 4 | | | | [R].DLPQGFSALEPLVDLPIGInITR.[F] | N20(HexNAc(4)Hex(4)) | P0DTC2 | 0 | 3 | 1 | [illegible] |
| Trypsin | High | PMI-Byonic | Unambiguous | 215 | 237 | DLPQGFSALEPLVDLPIGINITR | 234 | HexNAc(4)Hex(5)Fuc(1) | 4 | 5 | 1 | | | [R].DLPQGFSALEPLVDLPIGInITR.[F] | N20(HexNAc(4)Hex(5)Fuc(1)) | P0DTC2 | 0 | 3 | 1 | [illegible] |
| Trypsin | High | PMI-Byonic | Unambiguous | 215 | 237 | DLPQGFSALEPLVDLPIGINITR | 234 | HexNAc(4)Hex(5) | 4 | 5 | | | | [R].DLPQGFSALEPLVDLPIGInITR.[F] | N20(HexNAc(4)Hex(5)) | P0DTC2 | 0 | 3 | 1 | [illegible] |
| Trypsin | High | PMI-Byonic | Unambiguous | 215 | 237 | DLPQGFSALEPLVDLPIGINITR | 234 | HexNAc(5)Hex(3)Fuc(1) | 5 | 3 | 1 | | | [R].DLPQGFSALEPLVDLPIGInITR.[F] | N20(HexNAc(5)Hex(3)Fuc(1)) | P0DTC2 | 0 | 3 | 1 | [illegible] |
| Trypsin | High | PMI-Byonic | Unambiguous | 215 | 237 | DLPQGFSALEPLVDLPIGINITR | 234 | HexNAc(5)Hex(4)Fuc(1) | 5 | 4 | 1 | | | [R].DLPQGFSALEPLVDLPIGInITR.[F] | N20(HexNAc(5)Hex(4)Fuc(1)) | P0DTC2 | 0 | 3 | 1 | [illegible] |
| Trypsin | High | PMI-Byonic | Unambiguous | 215 | 237 | DLPQGFSALEPLVDLPIGINITR | 234 | HexNAc(6)Hex(7) | 6 | 7 | | | | [R].DLPQGFSALEPLVDLPIGInITR.[F] | N20(HexNAc(6)Hex(7)) | P0DTC2 | 0 | 4 | 1 | [illegible] |
| Trypsin | High | PMI-Byonic | Unambiguous | 279 | 300 | YNENGTITDAVDCALDPLSETK | 282 | HexNAc(2)Hex(5) | 2 | 5 | | | | [K].YNEnGTITDAVDcALDPLSETK.[C] | N4(HexNAc(2)Hex(5)); C13(Carbamidomethyl) | P0DTC2 | 0 | 3 | 1 | [illegible] |
| Trypsin | High | PMI-Byonic | Unambiguous | 279 | 300 | YNENGTITDAVDCALDPLSETK | 282 | HexNAc(2)Hex(6) | 2 | 6 | | | | [K].YNEnGTITDAVDcALDPLSETK.[C] | N4(HexNAc(2)Hex(6)); C13(Carbamidomethyl) | P0DTC2 | 0 | 3 | 1 | [illegible] |
| Trypsin | High | PMI-Byonic | Unambiguous | 279 | 300 | YNENGTITDAVDCALDPLSETK | 282 | HexNAc(3)Hex(3) | 3 | 3 | | | | [K].YNEnGTITDAVDcALDPLSETK.[C] | N4(HexNAc(3)Hex(3)); C13(Carbamidomethyl) | P0DTC2 | 0 | 3 | 1 | [illegible] |
| Trypsin | High | PMI-Byonic | Unambiguous | 279 | 300 | YNENGTITDAVDCALDPLSETK | 282 | HexNAc(3)Hex(5)Fuc(1) | 3 | 5 | 1 | | | [K].YNEnGTITDAVDcALDPLSETK.[C] | N4(HexNAc(3)Hex(5)Fuc(1)); C13(Carbamidomethyl) | P0DTC2 | 0 | 3 | 1 | [illegible] |
| Trypsin | High | PMI-Byonic | Unambiguous | 279 | 300 | YNENGTITDAVDCALDPLSETK | 282 | HexNAc(3)Hex(5)NeuGc(1) | 3 | 5 | | | 1 | [K].YNEnGTITDAVDcALDPLSETK.[C] | N4(HexNAc(3)Hex(5)NeuGc(1)); C13(Carbamidomethyl) | P0DTC2 | 0 | 3 | 1 | [illegible] |
| Trypsin | High | PMI-Byonic | Unambiguous | 279 | 300 | YNENGTITDAVDCALDPLSETK | 282 | HexNAc(3)Hex(5) | 3 | 5 | | | | [K].YNEnGTITDAVDcALDPLSETK.[C] | N4(HexNAc(3)Hex(5)); C13(Carbamidomethyl) | P0DTC2 | 0 | 3 | 1 | [illegible] |
| Trypsin | High | PMI-Byonic | Unambiguous | 279 | 300 | YNENGTITDAVDCALDPLSETK | 282 | HexNAc(4)Hex(3)Fuc(1) | 4 | 3 | 1 | | | [K].YNEnGTITDAVDcALDPLSETK.[C] | N4(HexNAc(4)Hex(3)Fuc(1)); C13(Carbamidomethyl) | P0DTC2 | 0 | 3 | 1 | [illegible] |
| Trypsin | High | PMI-Byonic | Unambiguous | 279 | 300 | YNENGTITDAVDCALDPLSETK | 282 | HexNAc(4)Hex(5)Fuc(1)NeuAc(1) | 4 | 5 | 1 | 1 | | [K].YNEnGTITDAVDcALDPLSETK.[C] | N4(HexNAc(4)Hex(5)Fuc(1)NeuAc(1)); C13(Carbamidomethyl) | P0DTC2 | 0 | 3 | 1 | [illegible] |
| Trypsin | High | PMI-Byonic | Unambiguous | 279 | 300 | YNENGTITDAVDCALDPLSETK | 282 | HexNAc(4)Hex(5)Fuc(1) | 4 | 5 | 1 | | | [K].YNEnGTITDAVDcALDPLSETK.[C] | N4(HexNAc(4)Hex(5)Fuc(1)); C13(Carbamidomethyl) | P0DTC2 | 0 | 3 | 1 | [illegible] |
| Trypsin | High | PMI-Byonic | Unambiguous | 279 | 300 | YNENGTITDAVDCALDPLSETK | 282 | HexNAc(4)Hex(5)Fuc(2) | 4 | 5 | 2 | | | [K].YNEnGTITDAVDcALDPLSETK.[C] | N4(HexNAc(4)Hex(5)Fuc(2)); C13(Carbamidomethyl) | P0DTC2 | 0 | 3 | 1 | [illegible] |
| Trypsin | High | PMI-Byonic | Unambiguous | 279 | 300 | YNENGTITDAVDCALDPLSETK | 282 | HexNAc(4)Hex(5)Fuc(3)NeuAc(1) | 4 | 5 | 3 | 1 | | [K].YNEnGTITDAVDcALDPLSETK.[C] | N4(HexNAc(4)Hex(5)Fuc(3)NeuAc(1)); C13(Carbamidomethyl) | P0DTC2 | 0 | 3 | 1 | [illegible] |
| Trypsin | High | PMI-Byonic | Unambiguous | 279 | 300 | YNENGTITDAVDCALDPLSETK | 282 | HexNAc(4)Hex(5) | 4 | 5 | | | | [K].YNEnGTITDAVDcALDPLSETK.[C] | N4(HexNAc(4)Hex(5)); C13(Carbamidomethyl) | P0DTC2 | 0 | 3 | 1 | [illegible] |
| Trypsin | High | PMI-Byonic | Unambiguous | 279 | 300 | YNENGTITDAVDCALDPLSETK | 282 | HexNAc(4)Hex(6)NeuAc(1) | 4 | 6 | | 1 | | [K].YNEnGTITDAVDcALDPLSETK.[C] | N4(HexNAc(4)Hex(6)NeuAc(1)); C13(Carbamidomethyl) | P0DTC2 | 0 | 3 | 1 | [illegible] |
| Trypsin | High | PMI-Byonic | Unambiguous | 279 | 300 | YNENGTITDAVDCALDPLSETK | 282 | HexNAc(4)Hex(6) | 4 | 6 | | | | [K].YNEnGTITDAVDcALDPLSETK.[C] | N4(HexNAc(4)Hex(6)); C13(Carbamidomethyl) | P0DTC2 | 0 | 3 | 1 | [illegible] |
| Trypsin | High | PMI-Byonic | Unambiguous | 279 | 300 | YNENGTITDAVDCALDPLSETK | 282 | HexNAc(5)Hex(5)Fuc(1)NeuAc(1) | 5 | 5 | 1 | 1 | | [K].YNEnGTITDAVDcALDPLSETK.[C] | N4(HexNAc(5)Hex(5)Fuc(1)NeuAc(1)); C13(Carbamidomethyl) | P0DTC2 | 0 | 3 | 1 | [illegible] |
| Trypsin | High | PMI-Byonic | Unambiguous | 279 | 300 | YNENGTITDAVDCALDPLSETK | 282 | HexNAc(5)Hex(5)Fuc(1) | 5 | 5 | 1 | | | [K].YNEnGTITDAVDcALDPLSETK.[C] | N4(HexNAc(5)Hex(5)Fuc(1)); C13(Carbamidomethyl) | P0DTC2 | 0 | 3 | 1 | [illegible] |
| Trypsin | High | PMI-Byonic | Unambiguous | 279 | 300 | YNENGTITDAVDCALDPLSETK | 282 | HexNAc(5)Hex(5)NeuAc(1) | 5 | 5 | | 1 | | [K].YNEnGTITDAVDcALDPLSETK.[C] | N4(HexNAc(5)Hex(5)NeuAc(1)); C13(Carbamidomethyl) | P0DTC2 | 0 | 3 | 1 | [illegible] |
| Trypsin | High | PMI-Byonic | Unambiguous | 279 | 300 | YNENGTITDAVDCALDPLSETK | 282 | HexNAc(5)Hex(5) | 5 | 5 | | | | [K].YNEnGTITDAVDcALDPLSETK.[C] | N4(HexNAc(5)Hex(5)); C13(Carbamidomethyl) | P0DTC2 | 0 | 3 | 1 | [illegible] |
| Trypsin | High | PMI-Byonic | Unambiguous | 279 | 300 | YNENGTITDAVDCALDPLSETK | 282 | HexNAc(5)Hex(6)Fuc(1)NeuAc(1) | 5 | 6 | 1 | 1 | | [K].YNEnGTITDAVDcALDPLSETK.[C] | N4(HexNAc(5)Hex(6)Fuc(1)NeuAc(1)); C13(Carbamidomethyl) | P0DTC2 | 0 | 4 | 1 | [illegible] |
| Trypsin | High | PMI-Byonic | Unambiguous | 279 | 300 | YNENGTITDAVDCALDPLSETK | 282 | HexNAc(5)Hex(6)Fuc(1)NeuGc(1) | 5 | 6 | 1 | | 1 | [K].YNEnGTITDAVDcALDPLSETK.[C] | N4(HexNAc(5)Hex(6)Fuc(1)NeuGc(1)); C13(Carbamidomethyl) | P0DTC2 | 0 | 3 | 1 | [illegible] |
| Trypsin | High | PMI-Byonic | Unambiguous | 279 | 300 | YNENGTITDAVDCALDPLSETK | 282 | HexNAc(5)Hex(6)Fuc(1) | 5 | 6 | 1 | | | [K].YNEnGTITDAVDcALDPLSETK.[C] | N4(HexNAc(5)Hex(6)Fuc(1)); C13(Carbamidomethyl) | P0DTC2 | 0 | 3 | 1 | [illegible] |
| Trypsin | High | PMI-Byonic | Unambiguous | 279 | 300 | YNENGTITDAVDCALDPLSETK | 282 | HexNAc(5)Hex(6)Fuc(3) | 5 | 6 | 3 | | | [K].YNEnGTITDAVDcALDPLSETK.[C] | N4(HexNAc(5)Hex(6)Fuc(3)); C13(Carbamidomethyl) | P0DTC2 | 0 | 3 | 1 | [illegible] |
| Trypsin | High | PMI-Byonic | Unambiguous | 279 | 300 | YNENGTITDAVDCALDPLSETK | 282 | HexNAc(5)Hex(6)NeuAc(1) | 5 | 6 | | 1 | | [K].YNEnGTITDAVDcALDPLSETK.[C] | N4(HexNAc(5)Hex(6)NeuAc(1)); C13(Carbamidomethyl) | P0DTC2 | 0 | 3 | 1 | [illegible] |
| Trypsin | High | PMI-Byonic | Unambiguous | 279 | 300 | YNENGTITDAVDCALDPLSETK | 282 | HexNAc(5)Hex(6) | 5 | 6 | | | | [K].YNEnGTITDAVDcALDPLSETK.[C] | N4(HexNAc(5)Hex(6)); C13(Carbamidomethyl) | P0DTC2 | 0 | 3 | 1 | [illegible] |
| Trypsin | High | PMI-Byonic | Unambiguous | 279 | 300 | YNENGTITDAVDCALDPLSETK | 282 | HexNAc(5)Hex(7)Fuc(1) | 5 | 7 | 1 | | | [K].YNEnGTITDAVDcALDPLSETK.[C] | N4(HexNAc(5)Hex(7)Fuc(1)); C13(Carbamidomethyl) | P0DTC2 | 0 | 3 | 1 | [illegible] |
| Trypsin | High | PMI-Byonic | Unambiguous | 279 | 300 | YNENGTITDAVDCALDPLSETK | 282 | HexNAc(5)Hex(7) | 5 | 7 | | | | [K].YNEnGTITDAVDcALDPLSETK.[C] | N4(HexNAc(5)Hex(7)); C13(Carbamidomethyl) | P0DTC2 | 0 | 3 | 1 | [illegible] |
| Trypsin | High | PMI-Byonic | Unambiguous | 279 | 300 | YNENGTITDAVDCALDPLSETK | 282 | HexNAc(6)Hex(3)Fuc(1) | 6 | 3 | 1 | | | [K].YNEnGTITDAVDcALDPLSETK.[C] | N4(HexNAc(6)Hex(3)Fuc(1)); C13(Carbamidomethyl) | P0DTC2 | 0 | 3 | 1 | [illegible] |
| Trypsin | High | PMI-Byonic | Unambiguous | 279 | 300 | YNENGTITDAVDCALDPLSETK | 282 | HexNAc(6)Hex(4)Fuc(1) | 6 | 4 | 1 | | | [K].YNEnGTITDAVDcALDPLSETK.[C] | N4(HexNAc(6)Hex(4)Fuc(1)); C13(Carbamidomethyl) | P0DTC2 | 0 | 3 | 1 | [illegible] |
| Trypsin | High | PMI-Byonic | Unambiguous | 279 | 300 | YNENGTITDAVDCALDPLSETK | 282 | HexNAc(6)Hex(5)Fuc(1) | 6 | 5 | 1 | | | [K].YNEnGTITDAVDcALDPLSETK.[C] | N4(HexNAc(6)Hex(5)Fuc(1)); C13(Carbamidomethyl) | P0DTC2 | 0 | 4 | 1 | [illegible] |
| Trypsin | High | PMI-Byonic | Unambiguous | 279 | 300 | YNENGTITDAVDCALDPLSETK | 282 | HexNAc(6)Hex(6)Fuc(1) | 6 | 6 | 1 | | | [K].YNEnGTITDAVDcALDPLSETK.[C] | N4(HexNAc(6)Hex(6)Fuc(1)); C13(Carbamidomethyl) | P0DTC2 | 0 | 3 | 1 | [illegible] |
| Trypsin | High | PMI-Byonic | Unambiguous | 279 | 300 | YNENGTITDAVDCALDPLSETK | 282 | HexNAc(6)Hex(6)Fuc(2) | 6 | 6 | 2 | | | [K].YNEnGTITDAVDcALDPLSETK.[C] | N4(HexNAc(6)Hex(6)Fuc(2)); C13(Carbamidomethyl) | P0DTC2 | 0 | 3 | 1 | [illegible] |
| Trypsin | High | PMI-Byonic | Unambiguous | 279 | 300 | YNENGTITDAVDCALDPLSETK | 282 | HexNAc(6)Hex(6)Fuc(3) | 6 | 6 | 3 | | | [K].YNEnGTITDAVDcALDPLSETK.[C] | N4(HexNAc(6)Hex(6)Fuc(3)); C13(Carbamidomethyl) | P0DTC2 | 0 | 3 | 1 | [illegible] |
| Trypsin | High | PMI-Byonic | Unambiguous | 279 | 300 | YNENGTITDAVDCALDPLSETK | 282 | HexNAc(6)Hex(6)NeuAc(1) | 6 | 6 | | 1 | | [K].YNEnGTITDAVDcALDPLSETK.[C] | N4(HexNAc(6)Hex(6)NeuAc(1)); C13(Carbamidomethyl) | P0DTC2 | 0 | 3 | 1 | [illegible] |
| Trypsin | High | PMI-Byonic | Unambiguous | 279 | 300 | YNENGTITDAVDCALDPLSETK | 282 | HexNAc(6)Hex(6) | 6 | 6 | | | | [K].YNEnGTITDAVDcALDPLSETK.[C] | N4(HexNAc(6)Hex(6)); C13(Carbamidomethyl) | P0DTC2 | 0 | 3 | 1 | [illegible] |
| Trypsin | High | PMI-Byonic | Unambiguous | 279 | 300 | YNENGTITDAVDCALDPLSETK | 282 | HexNAc(6)Hex(7)Fuc(1) | 6 | 7 | 1 | | | [K].YNEnGTITDAVDcALDPLSETK.[C] | N4(HexNAc(6)Hex(7)Fuc(1)); C13(Carbamidomethyl) | P0DTC2 | 0 | 3 | 1 | [illegible] |
| Trypsin | High | PMI-Byonic | Unambiguous | 279 | 300 | YNENGTITDAVDCALDPLSETK | 282 | HexNAc(6)Hex(7)Fuc(5) | 6 | 7 | 5 | | | [K].YNEnGTITDAVDcALDPLSETK.[C] | N4(HexNAc(6)Hex(7)Fuc(5)); C13(Carbamidomethyl) | P0DTC2 | 0 | 4 | 1 | [illegible] |
| Trypsin | High | PMI-Byonic | Unambiguous | 279 | 300 | YNENGTITDAVDCALDPLSETK | 282 | HexNAc(6)Hex(7)NeuAc(1) | 6 | 7 | | 1 | | [K].YNEnGTITDAVDcALDPLSETK.[C] | N4(HexNAc(6)Hex(7)NeuAc(1)); C13(Carbamidomethyl) | P0DTC2 | 0 | 3 | 1 | [illegible] |
| Trypsin | High | PMI-Byonic | Unambiguous | 279 | 300 | YNENGTITDAVDCALDPLSETK | 282 | HexNAc(6)Hex(7) | 6 | 7 | | | | [K].YNEnGTITDAVDcALDPLSETK.[C] | N4(HexNAc(6)Hex(7)); C13(Carbamidomethyl) | P0DTC2 | 0 | 3 | 1 | [illegible] |
| Trypsin | High | PMI-Byonic | Unambiguous | 279 | 300 | YNENGTITDAVDCALDPLSETK | 282 | HexNAc(6)Hex(8)Fuc(1)NeuAc(1) | 6 | 8 | 1 | 1 | | [K].YNEnGTITDAVDcALDPLSETK.[C] | N4(HexNAc(6)Hex(8)Fuc(1)NeuAc(1)); C13(Carbamidomethyl) | P0DTC2 | 0 | 4 | 1 | [illegible] |
| Trypsin | High | PMI-Byonic | Unambiguous | 279 | 300 | YNENGTITDAVDCALDPLSETK | 282 | HexNAc(6)Hex(8)NeuAc(1) | 6 | 8 | | 1 | | [K].YNEnGTITDAVDcALDPLSETK.[C] | N4(HexNAc(6)Hex(8)NeuAc(1)); C13(Carbamidomethyl) | P0DTC2 | 0 | 4 | 1 | [illegible] |
| Trypsin | High | PMI-Byonic | Unambiguous | 279 | 300 | YNENGTITDAVDCALDPLSETK | 282 | HexNAc(7)Hex(6)Fuc(1) | 7 | 6 | 1 | | | [K].YNEnGTITDAVDcALDPLSETK.[C] | N4(HexNAc(7)Hex(6)Fuc(1)); C13(Carbamidomethyl) | P0DTC2 | 0 | 3 | 1 | [illegible] |
| Trypsin | High | PMI-Byonic | Unambiguous | 647 | 682 | AGCLIGAEHVNNSYECDIPIGAGICASYQTQTNSPR | 657 | HexNAc(3)Hex(5)Fuc(1) | 3 | 5 | 1 | | | [R].AGcLIGAEHVnNSYEcDIPIGAGIcASYQTQTNSPR.[R] | C3(Carbamidomethyl); N11(HexNAc(3)Hex(5)Fuc(1)); C16(Carbamidomethyl); C25(Carbamidomethyl) | P0DTC2 | 0 | 4 | 1 | [illegible] |
| Trypsin | High | PMI-Byonic | Unambiguous | 647 | 682 | AGCLIGAEHVNNSYECDIPIGAGICASYQTQTNSPR | 657 | HexNAc(3)Hex(5)NeuGc(1) | 3 | 5 | | | 1 | [R].AGcLIGAEHVnNSYEcDIPIGAGIcASYQTQTNSPR.[R] | C3(Carbamidomethyl); N11(HexNAc(3)Hex(5)NeuGc(1)); C16(Carbamidomethyl); C25(Carbamidomethyl) | P0DTC2 | 0 | 4 | 1 | [illegible] |
| Trypsin | High | PMI-Byonic | Unambiguous | 647 | 682 | AGCLIGAEHVNNSYECDIPIGAGICASYQTQTNSPR | 657 | HexNAc(4)Hex(5)Fuc(1)NeuAc(1) | 4 | 5 | 1 | 1 | | [R].AGcLIGAEHVnNSYEcDIPIGAGIcASYQTQTNSPR.[R] | C3(Carbamidomethyl); N11(HexNAc(4)Hex(5)Fuc(1)NeuAc(1)); C16(Carbamidomethyl); C25(Carbamidomethyl) | P0DTC2 | 0 | 4 | 1 | [illegible] |
| Trypsin | High | PMI-Byonic | Unambiguous | 647 | 682 | AGCLIGAEHVNNSYECDIPIGAGICASYQTQTNSPR | 657 | HexNAc(4)Hex(5)Fuc(1) | 4 | 5 | 1 | | | [R].AGcLIGAEHVnNSYEcDIPIGAGIcASYQTQTNSPR.[R] | C3(Carbamidomethyl); N11(HexNAc(4)Hex(5)Fuc(1)); C16(Carbamidomethyl); C25(Carbamidomethyl) | P0DTC2 | 0 | 4 | 1 | [illegible] |
| Trypsin | High | PMI-Byonic | Unambiguous | 647 | 682 | AGCLIGAEHVNNSYECDIPIGAGICASYQTQTNSPR | 657 | HexNAc(4)Hex(5)Fuc(3) | 4 | 5 | 3 | | | [R].AGcLIGAEHVnNSYEcDIPIGAGIcASYQTQTNSPR.[R] | C3(Carbamidomethyl); N11(HexNAc(4)Hex(5)Fuc(3)); C16(Carbamidomethyl); C25(Carbamidomethyl) | P0DTC2 | 0 | 4 | 1 | [illegible] |
| Trypsin | High | PMI-Byonic | Unambiguous | 647 | 682 | AGCLIGAEHVNNSYECDIPIGAGICASYQTQTNSPR | 657 | HexNAc(4)Hex(5)NeuAc(1) | 4 | 5 | | 1 | | [R].AGcLIGAEHVnNSYEcDIPIGAGIcASYQTQTNSPR.[R] | C3(Carbamidomethyl); N11(HexNAc(4)Hex(5)NeuAc(1)); C16(Carbamidomethyl); C25(Carbamidomethyl) | P0DTC2 | 0 | 4 | 1 | [illegible] |
| Trypsin | High | PMI-Byonic | Unambiguous | 647 | 682 | AGCLIGAEHVNNSYECDIPIGAGICASYQTQTNSPR | 657 | HexNAc(5)Hex(5)Fuc(1)NeuAc(1) | 5 | 5 | 1 | 1 | | [R].AGcLIGAEHVnNSYEcDIPIGAGIcASYQTQTNSPR.[R] | C3(Carbamidomethyl); N11(HexNAc(5)Hex(5)Fuc(1)NeuAc(1)); C16(Carbamidomethyl); C25(Carbamidomethyl) | P0DTC2 | 0 | 4 | 1 | [illegible] |
| Trypsin | High | PMI-Byonic | Unambiguous | 647 | 682 | AGCLIGAEHVNNSYECDIPIGAGICASYQTQTNSPR | 657 | HexNAc(5)Hex(5)Fuc(1) | 5 | 5 | 1 | | | [R].AGcLIGAEHVnNSYEcDIPIGAGIcASYQTQTNSPR.[R] | C3(Carbamidomethyl); N11(HexNAc(5)Hex(5)Fuc(1)); C16(Carbamidomethyl); C25(Carbamidomethyl) | P0DTC2 | 0 | 4 | 1 | [illegible] |
| Trypsin | High | PMI-Byonic | Unambiguous | 647 | 682 | AGCLIGAEHVNNSYECDIPIGAGICASYQTQTNSPR | 657 | HexNAc(5)Hex(5)Fuc(2) | 5 | 5 | 2 | | | [R].AGcLIGAEHVnNSYEcDIPIGAGIcASYQTQTNSPR.[R] | C3(Carbamidomethyl); N11(HexNAc(5)Hex(5)Fuc(2)); C16(Carbamidomethyl); C25(Carbamidomethyl) | P0DTC2 | 0 | 4 | 1 | [illegible] |
| Trypsin | High | PMI-Byonic | Unambiguous | 647 | 682 | AGCLIGAEHVNNSYECDIPIGAGICASYQTQTNSPR | 657 | HexNAc(5)Hex(6)Fuc(1) | 5 | 6 | 1 | | | [R].AGcLIGAEHVnNSYEcDIPIGAGIcASYQTQTNSPR.[R] | C3(Carbamidomethyl); N11(HexNAc(5)Hex(6)Fuc(1)); C16(Carbamidomethyl); C25(Carbamidomethyl) | P0DTC2 | 0 | 4 | 1 | [illegible] |
| Trypsin | High | PMI-Byonic | Unambiguous | 647 | 682 | AGCLIGAEHVNNSYECDIPIGAGICASYQTQTNSPR | 657 | HexNAc(5)Hex(6)Fuc(2)NeuAc(1) | 5 | 6 | 2 | 1 | | [R].AGcLIGAEHVnNSYEcDIPIGAGIcASYQTQTNSPR.[R] | C3(Carbamidomethyl); N11(HexNAc(5)Hex(6)Fuc(2)NeuAc(1)); C16(Carbamidomethyl); C25(Carbamidomethyl) | P0DTC2 | 0 | 4 | 1 | [illegible] |
| Trypsin | High | PMI-Byonic | Unambiguous | 647 | 682 | AGCLIGAEHVNNSYECDIPIGAGICASYQTQTNSPR | 657 | HexNAc(5)Hex(6)Fuc(2)NeuGc(1) | 5 | 6 | 2 | | 1 | [R].AGcLIGAEHVnNSYEcDIPIGAGIcASYQTQTNSPR.[R] | C3(Carbamidomethyl); N11(HexNAc(5)Hex(6)Fuc(2)NeuGc(1)); C16(Carbamidomethyl); C25(Carbamidomethyl) | P0DTC2 | 0 | 4 | 1 | [illegible] |
| Trypsin | High | PMI-Byonic | Unambiguous | 647 | 682 | AGCLIGAEHVNNSYECDIPIGAGICASYQTQTNSPR | 657 | HexNAc(5)Hex(6)Fuc(2) | 5 | 6 | 2 | | | [R].AGcLIGAEHVnNSYEcDIPIGAGIcASYQTQTNSPR.[R] | C3(Carbamidomethyl); N11(HexNAc(5)Hex(6)Fuc(2)); C16(Carbamidomethyl); C25(Carbamidomethyl) | P0DTC2 | 0 | 4 | 1 | [illegible] |
| Trypsin | High | PMI-Byonic | Unambiguous | 647 | 682 | AGCLIGAEHVNNSYECDIPIGAGICASYQTQTNSPR | 657 | HexNAc(5)Hex(6)NeuAc(1) | 5 | 6 | | 1 | | [R].AGcLIGAEHVnNSYEcDIPIGAGIcASYQTQTNSPR.[R] | C3(Carbamidomethyl); N11(HexNAc(5)Hex(6)NeuAc(1)); C16(Carbamidomethyl); C25(Carbamidomethyl) | P0DTC2 | 0 | 4 | 1 | [illegible] |
| Trypsin | High | PMI-Byonic | Unambiguous | 647 | 682 | AGCLIGAEHVNNSYECDIPIGAGICASYQTQTNSPR | 657 | HexNAc(6)Hex(4)Fuc(1) | 6 | 4 | 1 | | | [R].AGcLIGAEHVnNSYEcDIPIGAGIcASYQTQTNSPR.[R] | C3(Carbamidomethyl); N11(HexNAc(6)Hex(4)Fuc(1)); C16(Carbamidomethyl); C25(Carbamidomethyl) | P0DTC2 | 0 | 4 | 1 | [illegible] |
| Trypsin | High | PMI-Byonic | Unambiguous | 647 | 682 | AGCLIGAEHVNNSYECDIPIGAGICASYQTQTNSPR | 657 | HexNAc(6)Hex(6)Fuc(1) | 6 | 6 | 1 | | | [R].AGcLIGAEHVnNSYEcDIPIGAGIcASYQTQTNSPR.[R] | C3(Carbamidomethyl); N11(HexNAc(6)Hex(6)Fuc(1)); C16(Carbamidomethyl); C25(Carbamidomethyl) | P0DTC2 | 0 | 4 | 1 | [illegible] |
| Trypsin | High | PMI-Byonic | Unambiguous | 647 | 682 | AGCLIGAEHVNNSYECDIPIGAGICASYQTQTNSPR | 657 | HexNAc(6)Hex(6)Fuc(3) | 6 | 6 | 3 | | | [R].AGcLIGAEHVnNSYEcDIPIGAGIcASYQTQTNSPR.[R] | C3(Carbamidomethyl); N11(HexNAc(6)Hex(6)Fuc(3)); C16(Carbamidomethyl); C25(Carbamidomethyl) | P0DTC2 | 0 | 4 | 1 | [illegible] |
| Trypsin | High | PMI-Byonic | Unambiguous | 647 | 682 | AGCLIGAEHVNNSYECDIPIGAGICASYQTQTNSPR | 657 | HexNAc(6)Hex(7)Fuc(1)NeuAc(1) | 6 | 7 | 1 | 1 | | [R].AGcLIGAEHVnNSYEcDIPIGAGIcASYQTQTNSPR.[R] | C3(Carbamidomethyl); N11(HexNAc(6)Hex(7)Fuc(1)NeuAc(1)); C16(Carbamidomethyl); C25(Carbamidomethyl) | P0DTC2 | 0 | 4 | 1 | [illegible] |
| Trypsin | High | PMI-Byonic | Unambiguous | 647 | 682 | AGCLIGAEHVNNSYECDIPIGAGICASYQTQTNSPR | 657 | HexNAc(6)Hex(7)NeuAc(1) | 6 | 7 | | 1 | | [R].AGcLIGAEHVnNSYEcDIPIGAGIcASYQTQTNSPR.[R] | C3(Carbamidomethyl); N11(HexNAc(6)Hex(7)NeuAc(1)); C16(Carbamidomethyl); C25(Carbamidomethyl) | P0DTC2 | 0 | 4 | 1 | [illegible] |
| Trypsin | High | PMI-Byonic | Unambiguous | 796 | 811 | DFGGFNFSQILPDPSK | 801 | HexNAc(2)Hex(4) | 2 | 4 | | | | [K].DFGGFnFSQILPDPSK.[P] | N6(HexNAc(2)Hex(4)) | P0DTC2 | 0 | 2 | 1 | [illegible] |
| Trypsin | High | PMI-Byonic | Unambiguous | 796 | 811 | DFGGFNFSQILPDPSK | 801 | HexNAc(2)Hex(5) | 2 | 5 | | | | [K].DFGGFnFSQILPDPSK.[P] | N6(HexNAc(2)Hex(5)) | P0DTC2 | 0 | 3 | 1 | [illegible] |
| Trypsin | High | PMI-Byonic | Unambiguous | 796 | 811 | DFGGFNFSQILPDPSK | 801 | HexNAc(2)Hex(6) | 2 | 6 | | | | [K].DFGGFnFSQILPDPSK.[P] | N6(HexNAc(2)Hex(6)) | P0DTC2 | 0 | 3 | 1 | [illegible] |
| Trypsin | High | PMI-Byonic | Unambiguous | 796 | 811 | DFGGFNFSQILPDPSK | 801 | HexNAc(2)Hex(7) | 2 | 7 | | | | [K].DFGGFnFSQILPDPSK.[P] | N6(HexNAc(2)Hex(7)) | P0DTC2 | 0 | 3 | 1 | [illegible] |
| Trypsin | High | PMI-Byonic | Unambiguous | 796 | 811 | DFGGFNFSQILPDPSK | 801 | HexNAc(2)Hex(8) | 2 | 8 | | | | [K].DFGGFnFSQILPDPSK.[P] | N6(HexNAc(2)Hex(8)) | P0DTC2 | 0 | 3 | 1 | [illegible] |
| Trypsin | High | PMI-Byonic | Unambiguous | 796 | 811 | DFGGFNFSQILPDPSK | 801 | HexNAc(2)Hex(9) | 2 | 9 | | | | [K].DFGGFnFSQILPDPSK.[P] | N6(HexNAc(2)Hex(9)) | P0DTC2 | 0 | 3 | 1 | [illegible] |
| Trypsin | High | PMI-Byonic | Unambiguous | 796 | 811 | DFGGFNFSQILPDPSK | 801 | HexNAc(3)Hex(5)Fuc(1) | 3 | 5 | 1 | | | [K].DFGGFnFSQILPDPSK.[P] | N6(HexNAc(3)Hex(5)Fuc(1)) | P0DTC2 | 0 | 3 | 1 | [illegible] |
| Trypsin | High | PMI-Byonic | Unambiguous | 796 | 811 | DFGGFNFSQILPDPSK | 801 | HexNAc(3)Hex(5)NeuAc(1) | 3 | 5 | | 1 | | [K].DFGGFnFSQILPDPSK.[P] | N6(HexNAc(3)Hex(5)NeuAc(1)) | P0DTC2 | 0 | 3 | 1 | [illegible] |
| Trypsin | High | PMI-Byonic | Unambiguous | 796 | 811 | DFGGFNFSQILPDPSK | 801 | HexNAc(3)Hex(5) | 3 | 5 | | | | [K].DFGGFnFSQILPDPSK.[P] | N6(HexNAc(3)Hex(5)) | P0DTC2 | 0 | 3 | 1 | [illegible] |
| Trypsin | High | PMI-Byonic | Unambiguous | 796 | 811 | DFGGFNFSQILPDPSK | 801 | HexNAc(3)Hex(6)Fuc(1) | 3 | 6 | 1 | | | [K].DFGGFnFSQILPDPSK.[P] | N6(HexNAc(3)Hex(6)Fuc(1)) | P0DTC2 | 0 | 3 | 1 | [illegible] |
| Trypsin | High | PMI-Byonic | Unambiguous | 796 | 811 | DFGGFNFSQILPDPSK | 801 | HexNAc(3)Hex(6)NeuAc(1) | 3 | 6 | | 1 | | [K].DFGGFnFSQILPDPSK.[P] | N6(HexNAc(3)Hex(6)NeuAc(1)) | P0DTC2 | 0 | 3 | 1 | [illegible] |
| Trypsin | High | PMI-Byonic | Unambiguous | 796 | 811 | DFGGFNFSQILPDPSK | 801 | HexNAc(3)Hex(6)NeuGc(1) | 3 | 6 | | | 1 | [K].DFGGFnFSQILPDPSK.[P] | N6(HexNAc(3)Hex(6)NeuGc(1)) | P0DTC2 | 0 | 3 | 1 | [illegible] |
| Trypsin | High | PMI-Byonic | Unambiguous | 796 | 811 | DFGGFNFSQILPDPSK | 801 | HexNAc(3)Hex(6) | 3 | 6 | | | | [K].DFGGFnFSQILPDPSK.[P] | N6(HexNAc(3)Hex(6)) | P0DTC2 | 0 | 3 | 1 | [illegible] |
| Trypsin | High | PMI-Byonic | Unambiguous | 796 | 811 | DFGGFNFSQILPDPSK | 801 | HexNAc(4)Hex(3)Fuc(1) | 4 | 3 | 1 | | | [K].DFGGFnFSQILPDPSK.[P] | N6(HexNAc(4)Hex(3)Fuc(1)) | P0DTC2 | 0 | 3 | 1 | [illegible] |
| Trypsin | High | PMI-Byonic | Unambiguous | 796 | 811 | DFGGFNFSQILPDPSK | 801 | HexNAc(4)Hex(3) | 4 | 3 | | | | [K].DFGGFnFSQILPDPSK.[P] | N6(HexNAc(4)Hex(3)) | P0DTC2 | 0 | 3 | 1 | [illegible] |
| Trypsin | High | PMI-Byonic | Unambiguous | 796 | 811 | DFGGFNFSQILPDPSK | 801 | HexNAc(4)Hex(4)Fuc(1) | 4 | 4 | 1 | | | [K].DFGGFnFSQILPDPSK.[P] | N6(HexNAc(4)Hex(4)Fuc(1)) | P0DTC2 | 0 | 3 | 1 | [illegible] |
| Trypsin | High | PMI-Byonic | Unambiguous | 796 | 811 | DFGGFNFSQILPDPSK | 801 | HexNAc(4)Hex(4)Fuc(2) | 4 | 4 | 2 | | | [K].DFGGFnFSQILPDPSK.[P] | N6(HexNAc(4)Hex(4)Fuc(2)) | P0DTC2 | 0 | 3 | 1 | [illegible] |
| Trypsin | High | PMI-Byonic | Unambiguous | 796 | 811 | DFGGFNFSQILPDPSK | 801 | HexNAc(4)Hex(5)Fuc(1)NeuAc(1) | 4 | 5 | 1 | 1 | | [K].DFGGFnFSQILPDPSK.[P] | N6(HexNAc(4)Hex(5)Fuc(1)NeuAc(1)) | P0DTC2 | 0 | 3 | 1 | [illegible] |
| Trypsin | High | PMI-Byonic | Unambiguous | 796 | 811 | DFGGFNFSQILPDPSK | 801 | HexNAc(4)Hex(5)Fuc(1)NeuGc(1) | 4 | 5 | 1 | | 1 | [K].DFGGFnFSQILPDPSK.[P] | N6(HexNAc(4)Hex(5)Fuc(1)NeuGc(1)) | P0DTC2 | 0 | 3 | 1 | [illegible] |
| Trypsin | High | PMI-Byonic | Unambiguous | 796 | 811 | DFGGFNFSQILPDPSK | 801 | HexNAc(4)Hex(5)Fuc(1) | 4 | 5 | 1 | | | [K].DFGGFnFSQILPDPSK.[P] | N6(HexNAc(4)Hex(5)Fuc(1)) | P0DTC2 | 0 | 3 | 1 | [illegible] |
| Trypsin | High | PMI-Byonic | Unambiguous | 796 | 811 | DFGGFNFSQILPDPSK | 801 | HexNAc(4)Hex(5)Fuc(2) | 4 | 5 | 2 | | | [K].DFGGFnFSQILPDPSK.[P] | N6(HexNAc(4)Hex(5)Fuc(2)) | P0DTC2 | 0 | 3 | 1 | [illegible] |
| Trypsin | High | PMI-Byonic | Unambiguous | 796 | 811 | DFGGFNFSQILPDPSK | 801 | HexNAc(4)Hex(5)Fuc(3) | 4 | 5 | 3 | | | [K].DFGGFnFSQILPDPSK.[P] | N6(HexNAc(4)Hex(5)Fuc(3)) | P0DTC2 | 0 | 3 | 1 | [illegible] |
| Trypsin | High | PMI-Byonic | Unambiguous | 796 | 811 | DFGGFNFSQILPDPSK | 801 | HexNAc(4)Hex(5)NeuGc(1) | 4 | 5 | | | 1 | [K].DFGGFnFSQILPDPSK.[P] | N6(HexNAc(4)Hex(5)NeuGc(1)) | P0DTC2 | 0 | 3 | 1 | [illegible] |
| Trypsin | High | PMI-Byonic | Unambiguous | 796 | 811 | DFGGFNFSQILPDPSK | 801 | HexNAc(4)Hex(5) | 4 | 5 | | | | [K].DFGGFnFSQILPDPSK.[P] | N6(HexNAc(4)Hex(5)) | P0DTC2 | 0 | 3 | 1 | [illegible] |
| Trypsin | High | PMI-Byonic | Unambiguous | 796 | 811 | DFGGFNFSQILPDPSK | 801 | HexNAc(4)Hex(6)Fuc(1) | 4 | 6 | 1 | | | [K].DFGGFnFSQILPDPSK.[P] | N6(HexNAc(4)Hex(6)Fuc(1)) | P0DTC2 | 0 | 3 | 1 | [illegible] |
| Trypsin | High | PMI-Byonic | Unambiguous | 796 | 811 | DFGGFNFSQILPDPSK | 801 | HexNAc(4)Hex(6)Fuc(2) | 4 | 6 | 2 | | | [K].DFGGFnFSQILPDPSK.[P] | N6(HexNAc(4)Hex(6)Fuc(2)) | P0DTC2 | 0 | 3 | 1 | [illegible] |
| Trypsin | High | PMI-Byonic | Unambiguous | 796 | 811 | DFGGFNFSQILPDPSK | 801 | HexNAc(4)Hex(6) | 4 | 6 | | | | [K].DFGGFnFSQILPDPSK.[P] | N6(HexNAc(4)Hex(6)) | P0DTC2 | 0 | 3 | 1 | [illegible] |
| Trypsin | High | PMI-Byonic | Unambiguous | 796 | 811 | DFGGFNFSQILPDPSK | 801 | HexNAc(5)Hex(3)Fuc(1) | 5 | 3 | 1 | | | [K].DFGGFnFSQILPDPSK.[P] | N6(HexNAc(5)Hex(3)Fuc(1)) | P0DTC2 | 0 | 3 | 1 | [illegible] |
| Trypsin | High | PMI-Byonic | Unambiguous | 796 | 811 | DFGGFNFSQILPDPSK | 801 | HexNAc(5)Hex(4)Fuc(1) | 5 | 4 | 1 | | | [K].DFGGFnFSQILPDPSK.[P] | N6(HexNAc(5)Hex(4)Fuc(1)) | P0DTC2 | 0 | 3 | 1 | [illegible] |
| Trypsin | High | PMI-Byonic | Unambiguous | 796 | 811 | DFGGFNFSQILPDPSK | 801 | HexNAc(5)Hex(4) | 5 | 4 | | | | [K].DFGGFnFSQILPDPSK.[P] | N6(HexNAc(5)Hex(4)) | P0DTC2 | 0 | 3 | 1 | [illegible] |
| Trypsin | High | PMI-Byonic | Unambiguous | 796 | 811 | DFGGFNFSQILPDPSK | 801 | HexNAc(5)Hex(5)Fuc(1) | 5 | 5 | 1 | | | [K].DFGGFnFSQILPDPSK.[P] | N6(HexNAc(5)Hex(5)Fuc(1)) | P0DTC2 | 0 | 3 | 1 | [illegible] |
| Trypsin | High | PMI-Byonic | Unambiguous | 796 | 811 | DFGGFNFSQILPDPSK | 801 | HexNAc(5)Hex(5)Fuc(2) | 5 | 5 | 2 | | | [K].DFGGFnFSQILPDPSK.[P] | N6(HexNAc(5)Hex(5)Fuc(2)) | P0DTC2 | 0 | 3 | 1 | [illegible] |
| Trypsin | High | PMI-Byonic | Unambiguous | 796 | 811 | DFGGFNFSQILPDPSK | 801 | HexNAc(5)Hex(5)Fuc(3) | 5 | 5 | 3 | | | [K].DFGGFnFSQILPDPSK.[P] | N6(HexNAc(5)Hex(5)Fuc(3)) | P0DTC2 | 0 | 3 | 1 | [illegible] |
| Trypsin | High | PMI-Byonic | Unambiguous | 796 | 811 | DFGGFNFSQILPDPSK | 801 | HexNAc(5)Hex(5)NeuAc(1)NeuGc(1) | 5 | 5 | | 1 | 1 | [K].DFGGFnFSQILPDPSK.[P] | N6(HexNAc(5)Hex(5)NeuAc(1)NeuGc(1)) | P0DTC2 | 0 | 3 | 1 | [illegible] |
| Trypsin | High | PMI-Byonic | Unambiguous | 796 | 811 | DFGGFNFSQILPDPSK | 801 | HexNAc(5)Hex(5) | 5 | 5 | | | | [K].DFGGFnFSQILPDPSK.[P] | N6(HexNAc(5)Hex(5)) | P0DTC2 | 0 | 3 | 1 | [illegible] |
| Trypsin | High | PMI-Byonic | Unambiguous | 796 | 811 | DFGGFNFSQILPDPSK | 801 | HexNAc(5)Hex(6)Fuc(1)NeuAc(1) | 5 | 6 | 1 | 1 | | [K].DFGGFnFSQILPDPSK.[P] | N6(HexNAc(5)Hex(6)Fuc(1)NeuAc(1)) | P0DTC2 | 0 | 3 | 1 | [illegible] |
| Trypsin | High | PMI-Byonic | Unambiguous | 796 | 811 | DFGGFNFSQILPDPSK | 801 | HexNAc(5)Hex(6)Fuc(1)NeuGc(1) | 5 | 6 | 1 | | 1 | [K].DFGGFnFSQILPDPSK.[P] | N6(HexNAc(5)Hex(6)Fuc(1)NeuGc(1)) | P0DTC2 | 0 | 3 | 1 | [illegible] |
| Trypsin | High | PMI-Byonic | Unambiguous | 796 | 811 | DFGGFNFSQILPDPSK | 801 | HexNAc(5)Hex(6)Fuc(1) | 5 | 6 | 1 | | | [K].DFGGFnFSQILPDPSK.[P] | N6(HexNAc(5)Hex(6)Fuc(1)) | P0DTC2 | 0 | 3 | 1 | [illegible] |
| Trypsin | High | PMI-Byonic | Unambiguous | 796 | 811 | DFGGFNFSQILPDPSK | 801 | HexNAc(5)Hex(6)Fuc(2)NeuAc(1) | 5 | 6 | 2 | 1 | | [K].DFGGFnFSQILPDPSK.[P] | N6(HexNAc(5)Hex(6)Fuc(2)NeuAc(1)) | P0DTC2 | 0 | 3 | 1 | [illegible] |
| Trypsin | High | PMI-Byonic | Unambiguous | 796 | 811 | DFGGFNFSQILPDPSK | 801 | HexNAc(5)Hex(6)Fuc(2) | 5 | 6 | 2 | | | [K].DFGGFnFSQILPDPSK.[P] | N6(HexNAc(5)Hex(6)Fuc(2)) | P0DTC2 | 0 | 3 | 1 | [illegible] |
| Trypsin | High | PMI-Byonic | Unambiguous | 796 | 811 | DFGGFNFSQILPDPSK | 801 | HexNAc(5)Hex(6)Fuc(3) | 5 | 6 | 3 | | | [K].DFGGFnFSQILPDPSK.[P] | N6(HexNAc(5)Hex(6)Fuc(3)) | P0DTC2 | 0 | 3 | 1 | [illegible] |
| Trypsin | High | PMI-Byonic | Unambiguous | 796 | 811 | DFGGFNFSQILPDPSK | 801 | HexNAc(5)Hex(6)NeuAc(1) | 5 | 6 | | 1 | | [K].DFGGFnFSQILPDPSK.[P] | N6(HexNAc(5)Hex(6)NeuAc(1)) | P0DTC2 | 0 | 3 | 1 | [illegible] |
| Trypsin | High | PMI-Byonic | Unambiguous | 796 | 811 | DFGGFNFSQILPDPSK | 801 | HexNAc(5)Hex(6) | 5 | 6 | | | | [K].DFGGFnFSQILPDPSK.[P] | N6(HexNAc(5)Hex(6)) | P0DTC2 | 0 | 3 | 1 | [illegible] |
| Trypsin | High | PMI-Byonic | Unambiguous | 796 | 811 | DFGGFNFSQILPDPSK | 801 | HexNAc(6)Hex(3)Fuc(1) | 6 | 3 | 1 | | | [K].DFGGFnFSQILPDPSK.[P] | N6(HexNAc(6)Hex(3)Fuc(1)) | P0DTC2 | 0 | 3 | 1 | [illegible] |
| Trypsin | High | PMI-Byonic | Unambiguous | 796 | 811 | DFGGFNFSQILPDPSK | 801 | HexNAc(6)Hex(4)Fuc(1) | 6 | 4 | 1 | | | [K].DFGGFnFSQILPDPSK.[P] | N6(HexNAc(6)Hex(4)Fuc(1)) | P0DTC2 | 0 | 3 | 1 | [illegible] |
| Trypsin | High | PMI-Byonic | Unambiguous | 796 | 811 | DFGGFNFSQILPDPSK | 801 | HexNAc(6)Hex(6)Fuc(1)NeuAc(1) | 6 | 6 | 1 | 1 | | [K].DFGGFnFSQILPDPSK.[P] | N6(HexNAc(6)Hex(6)Fuc(1)NeuAc(1)) | P0DTC2 | 0 | 3 | 1 | [illegible] |
| Trypsin | High | PMI-Byonic | Unambiguous | 796 | 811 | DFGGFNFSQILPDPSK | 801 | HexNAc(6)Hex(6)Fuc(1) | 6 | 6 | 1 | | | [K].DFGGFnFSQILPDPSK.[P] | N6(HexNAc(6)Hex(6)Fuc(1)) | P0DTC2 | 0 | 3 | 1 | [illegible] |
| Trypsin | High | PMI-Byonic | Unambiguous | 796 | 811 | DFGGFNFSQILPDPSK | 801 | HexNAc(6)Hex(6)NeuAc(1) | 6 | 6 | | 1 | | [K].DFGGFnFSQILPDPSK.[P] | N6(HexNAc(6)Hex(6)NeuAc(1)) | P0DTC2 | 0 | 3 | 1 | [illegible] |
| Trypsin | High | PMI-Byonic | Unambiguous | 796 | 811 | DFGGFNFSQILPDPSK | 801 | HexNAc(6)Hex(6) | 6 | 6 | | | | [K].DFGGFnFSQILPDPSK.[P] | N6(HexNAc(6)Hex(6)) | P0DTC2 | 0 | 3 | 1 | [illegible] |
| Trypsin | High | PMI-Byonic | Unambiguous | 1074 | 1086 | NFTTAPAICHDGK | 1074 | HexNAc(2)Hex(3) | 2 | 3 | | | | [K].nFTTAPAIcHDGK.[A] | N1(HexNAc(2)Hex(3)); C9(Carbamidomethyl) | P0DTC2 | 0 | 3 | 1 | [illegible] |
| Trypsin | High | PMI-Byonic | Unambiguous | 1074 | 1086 | NFTTAPAICHDGK | 1074 | HexNAc(2)Hex(4) | 2 | 4 | | | | [K].nFTTAPAIcHDGK.[A] | N1(HexNAc(2)Hex(4)); C9(Carbamidomethyl) | P0DTC2 | 0 | 3 | 1 | [illegible] |
| Trypsin | High | PMI-Byonic | Unambiguous | 1074 | 1086 | NFTTAPAICHDGK | 1074 | HexNAc(2)Hex(5) | 2 | 5 | | | | [K].nFTTAPAIcHDGK.[A] | N1(HexNAc(2)Hex(5)); C9(Carbamidomethyl) | P0DTC2 | 0 | 3 | 1 | [illegible] |
| Trypsin | High | PMI-Byonic | Unambiguous | 1074 | 1086 | NFTTAPAICHDGK | 1074 | HexNAc(2)Hex(6) | 2 | 6 | | | | [K].nFTTAPAIcHDGK.[A] | N1(HexNAc(2)Hex(6)); C9(Carbamidomethyl) | P0DTC2 | 0 | 3 | 1 | [illegible] |
| Trypsin | High | PMI-Byonic | Unambiguous | 1074 | 1086 | NFTTAPAICHDGK | 1074 | HexNAc(2)Hex(7) | 2 | 7 | | | | [K].nFTTAPAIcHDGK.[A] | N1(HexNAc(2)Hex(7)); C9(Carbamidomethyl) | P0DTC2 | 0 | 3 | 1 | [illegible] |
| Trypsin | High | PMI-Byonic | Unambiguous | 1074 | 1086 | NFTTAPAICHDGK | 1074 | HexNAc(2)Hex(8) | 2 | 8 | | | | [K].nFTTAPAIcHDGK.[A] | N1(HexNAc(2)Hex(8)); C9(Carbamidomethyl) | P0DTC2 | 0 | 3 | 1 | [illegible] |
| Trypsin | High | PMI-Byonic | Unambiguous | 1074 | 1086 | NFTTAPAICHDGK | 1074 | HexNAc(2)Hex(9) | 2 | 9 | | | | [K].nFTTAPAIcHDGK.[A] | N1(HexNAc(2)Hex(9)); C9(Carbamidomethyl) | P0DTC2 | 0 | 3 | 1 | [illegible] |
| Trypsin | High | PMI-Byonic | Unambiguous | 1074 | 1086 | NFTTAPAICHDGK | 1074 | HexNAc(3)Hex(3) | 3 | 3 | | | | [K].nFTTAPAIcHDGK.[A] | N1(HexNAc(3)Hex(3)); C9(Carbamidomethyl) | P0DTC2 | 0 | 3 | 1 | [illegible] |
| Trypsin | High | PMI-Byonic | Unambiguous | 1074 | 1086 | NFTTAPAICHDGK | 1074 | HexNAc(3)Hex(4)Fuc(1) | 3 | 4 | 1 | | | [K].nFTTAPAIcHDGK.[A] | N1(HexNAc(3)Hex(4)Fuc(1)); C9(Carbamidomethyl) | P0DTC2 | 0 | 3 | 1 | [illegible] |
| Trypsin | High | PMI-Byonic | Unambiguous | 1074 | 1086 | NFTTAPAICHDGK | 1074 | HexNAc(3)Hex(4)Fuc(2) | 3 | 4 | 2 | | | [K].nFTTAPAIcHDGK.[A] | N1(HexNAc(3)Hex(4)Fuc(2)); C9(Carbamidomethyl) | P0DTC2 | 0 | 3 | 1 | [illegible] |
| Trypsin | High | PMI-Byonic | Unambiguous | 1074 | 1086 | NFTTAPAICHDGK | 1074 | HexNAc(3)Hex(4) | 3 | 4 | | | | [K].nFTTAPAIcHDGK.[A] | N1(HexNAc(3)Hex(4)); C9(Carbamidomethyl) | P0DTC2 | 0 | 3 | 1 | [illegible] |
| Trypsin | High | PMI-Byonic | Unambiguous | 1074 | 1086 | NFTTAPAICHDGK | 1074 | HexNAc(3)Hex(5)Fuc(1) | 3 | 5 | 1 | | | [K].nFTTAPAIcHDGK.[A] | N1(HexNAc(3)Hex(5)Fuc(1)); C9(Carbamidomethyl) | P0DTC2 | 0 | 3 | 1 | [illegible] |
| Trypsin | High | PMI-Byonic | Unambiguous | 1074 | 1086 | NFTTAPAICHDGK | 1074 | HexNAc(3)Hex(5)NeuAc(1) | 3 | 5 | | 1 | | [K].nFTTAPAIcHDGK.[A] | N1(HexNAc(3)Hex(5)NeuAc(1)); C9(Carbamidomethyl) | P0DTC2 | 0 | 3 | 1 | [illegible] |
| Trypsin | High | PMI-Byonic | Unambiguous | 1074 | 1086 | NFTTAPAICHDGK | 1074 | HexNAc(3)Hex(5)NeuGc(1) | 3 | 5 | | | 1 | [K].nFTTAPAIcHDGK.[A] | N1(HexNAc(3)Hex(5)NeuGc(1)); C9(Carbamidomethyl) | P0DTC2 | 0 | 3 | 1 | [illegible] |
| Trypsin | High | PMI-Byonic | Unambiguous | 1074 | 1086 | NFTTAPAICHDGK | 1074 | HexNAc(3)Hex(5) | 3 | 5 | | | | [K].nFTTAPAIcHDGK.[A] | N1(HexNAc(3)Hex(5)); C9(Carbamidomethyl) | P0DTC2 | 0 | 3 | 1 | [illegible] |
| Trypsin | High | PMI-Byonic | Unambiguous | 1074 | 1086 | NFTTAPAICHDGK | 1074 | HexNAc(3)Hex(6)Fuc(1)NeuGc(1) | 3 | 6 | 1 | | 1 | [K].nFTTAPAIcHDGK.[A] | N1(HexNAc(3)Hex(6)Fuc(1)NeuGc(1)); C9(Carbamidomethyl) | P0DTC2 | 0 | 3 | 1 | [illegible] |
| Trypsin | High | PMI-Byonic | Unambiguous | 1074 | 1086 | NFTTAPAICHDGK | 1074 | HexNAc(3)Hex(6)Fuc(1) | 3 | 6 | 1 | | | [K].nFTTAPAIcHDGK.[A] | N1(HexNAc(3)Hex(6)Fuc(1)); C9(Carbamidomethyl) | P0DTC2 | 0 | 3 | 1 | [illegible] |
| Trypsin | High | PMI-Byonic | Unambiguous | 1074 | 1086 | NFTTAPAICHDGK | 1074 | HexNAc(3)Hex(6)NeuAc(1) | 3 | 6 | | 1 | | [K].nFTTAPAIcHDGK.[A] | N1(HexNAc(3)Hex(6)NeuAc(1)); C9(Carbamidomethyl) | P0DTC2 | 0 | 3 | 1 | [illegible] |
| Trypsin | High | PMI-Byonic | Unambiguous | 1074 | 1086 | NFTTAPAICHDGK | 1074 | HexNAc(3)Hex(6)NeuGc(1) | 3 | 6 | | | 1 | [K].nFTTAPAIcHDGK.[A] | N1(HexNAc(3)Hex(6)NeuGc(1)); C9(Carbamidomethyl) | P0DTC2 | 0 | 3 | 1 | [illegible] |
| Trypsin | High | PMI-Byonic | Unambiguous | 1074 | 1086 | NFTTAPAICHDGK | 1074 | HexNAc(3)Hex(6) | 3 | 6 | | | | [K].nFTTAPAIcHDGK.[A] | N1(HexNAc(3)Hex(6)); C9(Carbamidomethyl) | P0DTC2 | 0 | 3 | 1 | [illegible] |
| Trypsin | High | PMI-Byonic | Unambiguous | 1074 | 1086 | NFTTAPAICHDGK | 1074 | HexNAc(4)Hex(3)Fuc(1) | 4 | 3 | 1 | | | [K].nFTTAPAIcHDGK.[A] | N1(HexNAc(4)Hex(3)Fuc(1)); C9(Carbamidomethyl) | P0DTC2 | 0 | 3 | 1 | [illegible] |
| Trypsin | High | PMI-Byonic | Unambiguous | 1074 | 1086 | NFTTAPAICHDGK | 1074 | HexNAc(4)Hex(3) | 4 | 3 | | | | [K].nFTTAPAIcHDGK.[A] | N1(HexNAc(4)Hex(3)); C9(Carbamidomethyl) | P0DTC2 | 0 | 3 | 1 | [illegible] |
| Trypsin | High | PMI-Byonic | Unambiguous | 1074 | 1086 | NFTTAPAICHDGK | 1074 | HexNAc(4)Hex(4)Fuc(1) | 4 | 4 | 1 | | | [K].nFTTAPAIcHDGK.[A] | N1(HexNAc(4)Hex(4)Fuc(1)); C9(Carbamidomethyl) | P0DTC2 | 0 | 3 | 1 | [illegible] |
| Trypsin | High | PMI-Byonic | Unambiguous | 1074 | 1086 | NFTTAPAICHDGK | 1074 | HexNAc(4)Hex(4) | 4 | 4 | | | | [K].nFTTAPAIcHDGK.[A] | N1(HexNAc(4)Hex(4)); C9(Carbamidomethyl) | P0DTC2 | 0 | 3 | 1 | [illegible] |
| Trypsin | High | PMI-Byonic | Unambiguous | 1074 | 1086 | NFTTAPAICHDGK | 1074 | HexNAc(4)Hex(5)Fuc(1)NeuGc(1) | 4 | 5 | 1 | | 1 | [K].nFTTAPAIcHDGK.[A] | N1(HexNAc(4)Hex(5)Fuc(1)NeuGc(1)); C9(Carbamidomethyl) | P0DTC2 | 0 | 3 | 1 | [illegible] |
| Trypsin | High | PMI-Byonic | Unambiguous | 1074 | 1086 | NFTTAPAICHDGK | 1074 | HexNAc(4)Hex(5)Fuc(1) | 4 | 5 | 1 | | | [K].nFTTAPAIcHDGK.[A] | N1(HexNAc(4)Hex(5)Fuc(1)); C9(Carbamidomethyl) | P0DTC2 | 0 | 3 | 1 | [illegible] |
| Trypsin | High | PMI-Byonic | Unambiguous | 1074 | 1086 | NFTTAPAICHDGK | 1074 | HexNAc(4)Hex(5)Fuc(2)NeuGc(1) | 4 | 5 | 2 | | 1 | [K].nFTTAPAIcHDGK.[A] | N1(HexNAc(4)Hex(5)Fuc(2)NeuGc(1)); C9(Carbamidomethyl) | P0DTC2 | 0 | 3 | 1 | [illegible] |
| Trypsin | High | PMI-Byonic | Unambiguous | 1074 | 1086 | NFTTAPAICHDGK | 1074 | HexNAc(4)Hex(5)Fuc(2) | 4 | 5 | 2 | | | [K].nFTTAPAIcHDGK.[A] | N1(HexNAc(4)Hex(5)Fuc(2)); C9(Carbamidomethyl) | P0DTC2 | 0 | 3 | 1 | [illegible] |
| Trypsin | High | PMI-Byonic | Unambiguous | 1074 | 1086 | NFTTAPAICHDGK | 1074 | HexNAc(4)Hex(5)Fuc(3) | 4 | 5 | 3 | | | [K].nFTTAPAIcHDGK.[A] | N1(HexNAc(4)Hex(5)Fuc(3)); C9(Carbamidomethyl) | P0DTC2 | 0 | 3 | 1 | [illegible] |
| Trypsin | High | PMI-Byonic | Unambiguous | 1074 | 1086 | NFTTAPAICHDGK | 1074 | HexNAc(4)Hex(5)NeuGc(1) | 4 | 5 | | | 1 | [K].nFTTAPAIcHDGK.[A] | N1(HexNAc(4)Hex(5)NeuGc(1)); C9(Carbamidomethyl) | P0DTC2 | 0 | 3 | 1 | [illegible] |
| Trypsin | High | PMI-Byonic | Unambiguous | 1074 | 1086 | NFTTAPAICHDGK | 1074 | HexNAc(4)Hex(5) | 4 | 5 | | | | [K].nFTTAPAIcHDGK.[A] | N1(HexNAc(4)Hex(5)); C9(Carbamidomethyl) | P0DTC2 | 0 | 3 | 1 | [illegible] |
| Trypsin | High | PMI-Byonic | Unambiguous | 1074 | 1086 | NFTTAPAICHDGK | 1074 | HexNAc(4)Hex(6)Fuc(1) | 4 | 6 | 1 | | | [K].nFTTAPAIcHDGK.[A] | N1(HexNAc(4)Hex(6)Fuc(1)); C9(Carbamidomethyl) | P0DTC2 | 0 | 3 | 1 | [illegible] |
| Trypsin | High | PMI-Byonic | Unambiguous | 1074 | 1086 | NFTTAPAICHDGK | 1074 | HexNAc(4)Hex(6)Fuc(2) | 4 | 6 | 2 | | | [K].nFTTAPAIcHDGK.[A] | N1(HexNAc(4)Hex(6)Fuc(2)); C9(Carbamidomethyl) | P0DTC2 | 0 | 3 | 1 | [illegible] |
| Trypsin | High | PMI-Byonic | Unambiguous | 1074 | 1086 | NFTTAPAICHDGK | 1074 | HexNAc(4)Hex(6)Fuc(3) | 4 | 6 | 3 | | | [K].nFTTAPAIcHDGK.[A] | N1(HexNAc(4)Hex(6)Fuc(3)); C9(Carbamidomethyl) | P0DTC2 | 0 | 3 | 1 | [illegible] |
| Trypsin | High | PMI-Byonic | Unambiguous | 1074 | 1086 | NFTTAPAICHDGK | 1074 | HexNAc(4)Hex(6) | 4 | 6 | | | | [K].nFTTAPAIcHDGK.[A] | N1(HexNAc(4)Hex(6)); C9(Carbamidomethyl) | P0DTC2 | 0 | 3 | 1 | [illegible] |
| Trypsin | High | PMI-Byonic | Unambiguous | 1074 | 1086 | NFTTAPAICHDGK | 1074 | HexNAc(4)Hex(7)Fuc(2) | 4 | 7 | 2 | | | [K].nFTTAPAIcHDGK.[A] | N1(HexNAc(4)Hex(7)Fuc(2)); C9(Carbamidomethyl) | P0DTC2 | 0 | 3 | 1 | [illegible] |
| Trypsin | High | PMI-Byonic | Unambiguous | 1074 | 1086 | NFTTAPAICHDGK | 1074 | HexNAc(4)Hex(7) | 4 | 7 | | | | [K].nFTTAPAIcHDGK.[A] | N1(HexNAc(4)Hex(7)); C9(Carbamidomethyl) | P0DTC2 | 0 | 3 | 1 | [illegible] |
| Trypsin | High | PMI-Byonic | Unambiguous | 1074 | 1086 | NFTTAPAICHDGK | 1074 | HexNAc(5)Hex(3)Fuc(1) | 5 | 3 | 1 | | | [K].nFTTAPAIcHDGK.[A] | N1(HexNAc(5)Hex(3)Fuc(1)); C9(Carbamidomethyl) | P0DTC2 | 0 | 3 | 1 | [illegible] |
| Trypsin | High | PMI-Byonic | Unambiguous | 1074 | 1086 | NFTTAPAICHDGK | 1074 | HexNAc(5)Hex(3) | 5 | 3 | | | | [K].nFTTAPAIcHDGK.[A] | N1(HexNAc(5)Hex(3)); C9(Carbamidomethyl) | P0DTC2 | 0 | 3 | 1 | [illegible] |
| Trypsin | High | PMI-Byonic | Unambiguous | 1074 | 1086 | NFTTAPAICHDGK | 1074 | HexNAc(5)Hex(4)Fuc(1) | 5 | 4 | 1 | | | [K].nFTTAPAIcHDGK.[A] | N1(HexNAc(5)Hex(4)Fuc(1)); C9(Carbamidomethyl) | P0DTC2 | 0 | 3 | 1 | [illegible] |
| Trypsin | High | PMI-Byonic | Unambiguous | 1074 | 1086 | NFTTAPAICHDGK | 1074 | HexNAc(5)Hex(4)Fuc(2) | 5 | 4 | 2 | | | [K].nFTTAPAIcHDGK.[A] | N1(HexNAc(5)Hex(4)Fuc(2)); C9(Carbamidomethyl) | P0DTC2 | 0 | 3 | 1 | [illegible] |
| Trypsin | High | PMI-Byonic | Unambiguous | 1074 | 1086 | NFTTAPAICHDGK | 1074 | HexNAc(5)Hex(4) | 5 | 4 | | | | [K].nFTTAPAIcHDGK.[A] | N1(HexNAc(5)Hex(4)); C9(Carbamidomethyl) | P0DTC2 | 0 | 3 | 1 | [illegible] |
| Trypsin | High | PMI-Byonic | Unambiguous | 1074 | 1086 | NFTTAPAICHDGK | 1074 | HexNAc(5)Hex(5)Fuc(1) | 5 | 5 | 1 | | | [K].nFTTAPAIcHDGK.[A] | N1(HexNAc(5)Hex(5)Fuc(1)); C9(Carbamidomethyl) | P0DTC2 | 0 | 3 | 1 | [illegible] |
| Trypsin | High | PMI-Byonic | Unambiguous | 1074 | 1086 | NFTTAPAICHDGK | 1074 | HexNAc(5)Hex(5)Fuc(2) | 5 | 5 | 2 | | | [K].nFTTAPAIcHDGK.[A] | N1(HexNAc(5)Hex(5)Fuc(2)); C9(Carbamidomethyl) | P0DTC2 | 0 | 3 | 1 | [illegible] |
| Trypsin | High | PMI-Byonic | Unambiguous | 1074 | 1086 | NFTTAPAICHDGK | 1074 | HexNAc(5)Hex(5)Fuc(3) | 5 | 5 | 3 | | | [K].nFTTAPAIcHDGK.[A] | N1(HexNAc(5)Hex(5)Fuc(3)); C9(Carbamidomethyl) | P0DTC2 | 0 | 3 | 1 | [illegible] |
| Trypsin | High | PMI-Byonic | Unambiguous | 1074 | 1086 | NFTTAPAICHDGK | 1074 | HexNAc(5)Hex(5) | 5 | 5 | | | | [K].nFTTAPAIcHDGK.[A] | N1(HexNAc(5)Hex(5)); C9(Carbamidomethyl) | P0DTC2 | 0 | 3 | 1 | [illegible] |
| Trypsin | High | PMI-Byonic | Unambiguous | 1074 | 1086 | NFTTAPAICHDGK | 1074 | HexNAc(5)Hex(6)Fuc(1)NeuAc(1) | 5 | 6 | 1 | 1 | | [K].nFTTAPAIcHDGK.[A] | N1(HexNAc(5)Hex(6)Fuc(1)NeuAc(1)); C9(Carbamidomethyl) | P0DTC2 | 0 | 3 | 1 | [illegible] |
| Trypsin | High | PMI-Byonic | Unambiguous | 1074 | 1086 | NFTTAPAICHDGK | 1074 | HexNAc(5)Hex(6)Fuc(1)NeuGc(1) | 5 | 6 | 1 | | 1 | [K].nFTTAPAIcHDGK.[A] | N1(HexNAc(5)Hex(6)Fuc(1)NeuGc(1)); C9(Carbamidomethyl) | P0DTC2 | 0 | 3 | 1 | [illegible] |
| Trypsin | High | PMI-Byonic | Unambiguous | 1074 | 1086 | NFTTAPAICHDGK | 1074 | HexNAc(5)Hex(6)Fuc(1) | 5 | 6 | 1 | | | [K].nFTTAPAIcHDGK.[A] | N1(HexNAc(5)Hex(6)Fuc(1)); C9(Carbamidomethyl) | P0DTC2 | 0 | 3 | 1 | [illegible] |
| Trypsin | High | PMI-Byonic | Unambiguous | 1074 | 1086 | NFTTAPAICHDGK | 1074 | HexNAc(5)Hex(6)Fuc(2)NeuAc(1) | 5 | 6 | 2 | 1 | | [K].nFTTAPAIcHDGK.[A] | N1(HexNAc(5)Hex(6)Fuc(2)NeuAc(1)); C9(Carbamidomethyl) | P0DTC2 | 0 | 3 | 1 | [illegible] |
| Trypsin | High | PMI-Byonic | Unambiguous | 1074 | 1086 | NFTTAPAICHDGK | 1074 | HexNAc(5)Hex(6)Fuc(2)NeuGc(1) | 5 | 6 | 2 | | 1 | [K].nFTTAPAIcHDGK.[A] | N1(HexNAc(5)Hex(6)Fuc(2)NeuGc(1)); C9(Carbamidomethyl) | P0DTC2 | 0 | 3 | 1 | [illegible] |
| Trypsin | High | PMI-Byonic | Unambiguous | 1074 | 1086 | NFTTAPAICHDGK | 1074 | HexNAc(5)Hex(6)Fuc(2) | 5 | 6 | 2 | | | [K].nFTTAPAIcHDGK.[A] | N1(HexNAc(5)Hex(6)Fuc(2)); C9(Carbamidomethyl) | P0DTC2 | 0 | 3 | 1 | [illegible] |
| Trypsin | High | PMI-Byonic | Unambiguous | 1074 | 1086 | NFTTAPAICHDGK | 1074 | HexNAc(5)Hex(6)Fuc(3) | 5 | 6 | 3 | | | [K].nFTTAPAIcHDGK.[A] | N1(HexNAc(5)Hex(6)Fuc(3)); C9(Carbamidomethyl) | P0DTC2 | 0 | 3 | 1 | [illegible] |
| Trypsin | High | PMI-Byonic | Unambiguous | 1074 | 1086 | NFTTAPAICHDGK | 1074 | HexNAc(5)Hex(6)NeuGc(1) | 5 | 6 | | | 1 | [K].nFTTAPAIcHDGK.[A] | N1(HexNAc(5)Hex(6)NeuGc(1)); C9(Carbamidomethyl) | P0DTC2 | 0 | 3 | 1 | [illegible] |
| Trypsin | High | PMI-Byonic | Unambiguous | 1074 | 1086 | NFTTAPAICHDGK | 1074 | HexNAc(5)Hex(6) | 5 | 6 | | | | [K].nFTTAPAIcHDGK.[A] | N1(HexNAc(5)Hex(6)); C9(Carbamidomethyl) | P0DTC2 | 0 | 3 | 1 | [illegible] |
| Trypsin | High | PMI-Byonic | Unambiguous | 1074 | 1086 | NFTTAPAICHDGK | 1074 | HexNAc(6)Hex(3)Fuc(1) | 6 | 3 | 1 | | | [K].nFTTAPAIcHDGK.[A] | N1(HexNAc(6)Hex(3)Fuc(1)); C9(Carbamidomethyl) | P0DTC2 | 0 | 3 | 1 | [illegible] |
| Trypsin | High | PMI-Byonic | Unambiguous | 1074 | 1086 | NFTTAPAICHDGK | 1074 | HexNAc(6)Hex(4)Fuc(1) | 6 | 4 | 1 | | | [K].nFTTAPAIcHDGK.[A] | N1(HexNAc(6)Hex(4)Fuc(1)); C9(Carbamidomethyl) | P0DTC2 | 0 | 3 | 1 | [illegible] |
| Trypsin | High | PMI-Byonic | Unambiguous | 1074 | 1086 | NFTTAPAICHDGK | 1074 | HexNAc(6)Hex(5)Fuc(1) | 6 | 5 | 1 | | | [K].nFTTAPAIcHDGK.[A] | N1(HexNAc(6)Hex(5)Fuc(1)); C9(Carbamidomethyl) | P0DTC2 | 0 | 3 | 1 | [illegible] |
| Trypsin | High | PMI-Byonic | Unambiguous | 1074 | 1086 | NFTTAPAICHDGK | 1074 | HexNAc(6)Hex(5)Fuc(2) | 6 | 5 | 2 | | | [K].nFTTAPAIcHDGK.[A] | N1(HexNAc(6)Hex(5)Fuc(2)); C9(Carbamidomethyl) | P0DTC2 | 0 | 3 | 1 | [illegible] |
| Trypsin | High | PMI-Byonic | Unambiguous | 1074 | 1086 | NFTTAPAICHDGK | 1074 | HexNAc(6)Hex(6)Fuc(1) | 6 | 6 | 1 | | | [K].nFTTAPAIcHDGK.[A] | N1(HexNAc(6)Hex(6)Fuc(1)); C9(Carbamidomethyl) | P0DTC2 | 0 | 4 | 1 | [illegible] |
| Trypsin | High | PMI-Byonic | Unambiguous | 1074 | 1086 | NFTTAPAICHDGK | 1074 | HexNAc(6)Hex(6)Fuc(2) | 6 | 6 | 2 | | | [K].nFTTAPAIcHDGK.[A] | N1(HexNAc(6)Hex(6)Fuc(2)); C9(Carbamidomethyl) | P0DTC2 | 0 | 4 | 1 | [illegible] |
| Trypsin | High | PMI-Byonic | Unambiguous | 1074 | 1086 | NFTTAPAICHDGK | 1074 | HexNAc(6)Hex(6)Fuc(3) | 6 | 6 | 3 | | | [K].nFTTAPAIcHDGK.[A] | N1(HexNAc(6)Hex(6)Fuc(3)); C9(Carbamidomethyl) | P0DTC2 | 0 | 3 | 1 | [illegible] |
| Trypsin | High | PMI-Byonic | Unambiguous | 1074 | 1086 | NFTTAPAICHDGK | 1074 | HexNAc(6)Hex(6) | 6 | 6 | | | | [K].nFTTAPAIcHDGK.[A] | N1(HexNAc(6)Hex(6)); C9(Carbamidomethyl) | P0DTC2 | 0 | 3 | 1 | [illegible] |
| Trypsin | High | PMI-Byonic | Unambiguous | 1074 | 1086 | NFTTAPAICHDGK | 1074 | HexNAc(6)Hex(7)Fuc(1)NeuAc(1) | 6 | 7 | 1 | 1 | | [K].nFTTAPAIcHDGK.[A] | N1(HexNAc(6)Hex(7)Fuc(1)NeuAc(1)); C9(Carbamidomethyl) | P0DTC2 | 0 | 4 | 1 | [illegible] |
| Trypsin | High | PMI-Byonic | Unambiguous | 1074 | 1086 | NFTTAPAICHDGK | 1074 | HexNAc(6)Hex(7)Fuc(1)NeuGc(1) | 6 | 7 | 1 | | 1 | [K].nFTTAPAIcHDGK.[A] | N1(HexNAc(6)Hex(7)Fuc(1)NeuGc(1)); C9(Carbamidomethyl) | P0DTC2 | 0 | 3 | 1 | [illegible] |
| Trypsin | High | PMI-Byonic | Unambiguous | 1074 | 1086 | NFTTAPAICHDGK | 1074 | HexNAc(6)Hex(7)Fuc(1) | 6 | 7 | 1 | | | [K].nFTTAPAIcHDGK.[A] | N1(HexNAc(6)Hex(7)Fuc(1)); C9(Carbamidomethyl) | P0DTC2 | 0 | 3 | 1 | [illegible] |
| Trypsin | High | PMI-Byonic | Unambiguous | 1074 | 1086 | NFTTAPAICHDGK | 1074 | HexNAc(6)Hex(7)Fuc(4) | 6 | 7 | 4 | | | [K].nFTTAPAIcHDGK.[A] | N1(HexNAc(6)Hex(7)Fuc(4)); C9(Carbamidomethyl) | P0DTC2 | 0 | 3 | 1 | [illegible] |
| Trypsin | High | PMI-Byonic | Unambiguous | 1074 | 1086 | NFTTAPAICHDGK | 1074 | HexNAc(6)Hex(7)Fuc(5) | 6 | 7 | 5 | | | [K].nFTTAPAIcHDGK.[A] | N1(HexNAc(6)Hex(7)Fuc(5)); C9(Carbamidomethyl) | P0DTC2 | 0 | 3 | 1 | [illegible] |
| Trypsin | High | PMI-Byonic | Unambiguous | 1074 | 1086 | NFTTAPAICHDGK | 1074 | HexNAc(6)Hex(7)NeuAc(1) | 6 | 7 | | 1 | | [K].nFTTAPAIcHDGK.[A] | N1(HexNAc(6)Hex(7)NeuAc(1)); C9(Carbamidomethyl) | P0DTC2 | 0 | 3 | 1 | [illegible] |
| Trypsin | High | PMI-Byonic | Unambiguous | 1074 | 1086 | NFTTAPAICHDGK | 1074 | HexNAc(6)Hex(7)NeuGc(1) | 6 | 7 | | | 1 | [K].nFTTAPAIcHDGK.[A] | N1(HexNAc(6)Hex(7)NeuGc(1)); C9(Carbamidomethyl) | P0DTC2 | 0 | 3 | 1 | [illegible] |
| Trypsin | High | PMI-Byonic | Unambiguous | 1074 | 1086 | NFTTAPAICHDGK | 1074 | HexNAc(6)Hex(7) | 6 | 7 | | | | [K].nFTTAPAIcHDGK.[A] | N1(HexNAc(6)Hex(7)); C9(Carbamidomethyl) | P0DTC2 | 0 | 3 | 1 | [illegible] |
| Trypsin | High | PMI-Byonic | Unambiguous | 1074 | 1086 | NFTTAPAICHDGK | 1074 | HexNAc(7)Hex(3)Fuc(1) | 7 | 3 | 1 | | | [K].nFTTAPAIcHDGK.[A] | N1(HexNAc(7)Hex(3)Fuc(1)); C9(Carbamidomethyl) | P0DTC2 | 0 | 3 | 1 | [illegible] |
| Trypsin | High | PMI-Byonic | Unambiguous | 1074 | 1086 | NFTTAPAICHDGK | 1074 | HexNAc(7)Hex(6)Fuc(1) | 7 | 6 | 1 | | | [K].nFTTAPAIcHDGK.[A] | N1(HexNAc(7)Hex(6)Fuc(1)); C9(Carbamidomethyl) | P0DTC2 | 0 | 3 | 1 | [illegible] |
| Trypsin | High | PMI-Byonic | Unambiguous | 1074 | 1086 | NFTTAPAICHDGK | 1074 | HexNAc(7)Hex(7)Fuc(1) | 7 | 7 | 1 | | | [K].nFTTAPAIcHDGK.[A] | N1(HexNAc(7)Hex(7)Fuc(1)); C9(Carbamidomethyl) | P0DTC2 | 0 | 3 | 1 | [illegible] |
| Trypsin | High | PMI-Byonic | Unambiguous | 1092 | 1107 | EGVFVSNGTHWFVTQR | 1098 | HexNAc(2)Hex(5) | 2 | 5 | | | | [R].EGVFVSnGTHWFVTQR.[N] | N7(HexNAc(2)Hex(5)) | P0DTC2 | 0 | 3 | 1 | [illegible] |
| Trypsin | High | PMI-Byonic | Unambiguous | 1092 | 1107 | EGVFVSNGTHWFVTQR | 1098 | HexNAc(2)Hex(6) | 2 | 6 | | | | [R].EGVFVSnGTHWFVTQR.[N] | N7(HexNAc(2)Hex(6)) | P0DTC2 | 0 | 3 | 1 | [illegible] |
| Trypsin | High | PMI-Byonic | Unambiguous | 1092 | 1107 | EGVFVSNGTHWFVTQR | 1098 | HexNAc(2)Hex(7) | 2 | 7 | | | | [R].EGVFVSnGTHWFVTQR.[N] | N7(HexNAc(2)Hex(7)) | P0DTC2 | 0 | 3 | 1 | [illegible] |
| Trypsin | High | PMI-Byonic | Unambiguous | 1092 | 1107 | EGVFVSNGTHWFVTQR | 1098 | HexNAc(2)Hex(8) | 2 | 8 | | | | [R].EGVFVSnGTHWFVTQR.[N] | N7(HexNAc(2)Hex(8)) | P0DTC2 | 0 | 3 | 1 | [illegible] |
| Trypsin | High | PMI-Byonic | Unambiguous | 1092 | 1107 | EGVFVSNGTHWFVTQR | 1098 | HexNAc(2)Hex(9) | 2 | 9 | | | | [R].EGVFVSnGTHWFVTQR.[N] | N7(HexNAc(2)Hex(9)) | P0DTC2 | 0 | 3 | 1 | [illegible] |
| Trypsin | High | PMI-Byonic | Unambiguous | 1092 | 1107 | EGVFVSNGTHWFVTQR | 1098 | HexNAc(3)Hex(3) | 3 | 3 | | | | [R].EGVFVSnGTHWFVTQR.[N] | N7(HexNAc(3)Hex(3)) | P0DTC2 | 0 | 3 | 1 | [illegible] |
| Trypsin | High | PMI-Byonic | Unambiguous | 1092 | 1107 | EGVFVSNGTHWFVTQR | 1098 | HexNAc(3)Hex(4)NeuGc(1) | 3 | 4 | | | 1 | [R].EGVFVSnGTHWFVTQR.[N] | N7(HexNAc(3)Hex(4)NeuGc(1)) | P0DTC2 | 0 | 3 | 1 | [illegible] |
| Trypsin | High | PMI-Byonic | Unambiguous | 1092 | 1107 | EGVFVSNGTHWFVTQR | 1098 | HexNAc(3)Hex(4) | 3 | 4 | | | | [R].EGVFVSnGTHWFVTQR.[N] | N7(HexNAc(3)Hex(4)) | P0DTC2 | 0 | 3 | 1 | [illegible] |
| Trypsin | High | PMI-Byonic | Unambiguous | 1092 | 1107 | EGVFVSNGTHWFVTQR | 1098 | HexNAc(3)Hex(5)Fuc(1) | 3 | 5 | 1 | | | [R].EGVFVSnGTHWFVTQR.[N] | N7(HexNAc(3)Hex(5)Fuc(1)) | P0DTC2 | 0 | 3 | 1 | [illegible] |
| Trypsin | High | PMI-Byonic | Unambiguous | 1092 | 1107 | EGVFVSNGTHWFVTQR | 1098 | HexNAc(3)Hex(5)NeuAc(1) | 3 | 5 | | 1 | | [R].EGVFVSnGTHWFVTQR.[N] | N7(HexNAc(3)Hex(5)NeuAc(1)) | P0DTC2 | 0 | 3 | 1 | [illegible] |
| Trypsin | High | PMI-Byonic | Unambiguous | 1092 | 1107 | EGVFVSNGTHWFVTQR | 1098 | HexNAc(3)Hex(5)NeuGc(1) | 3 | 5 | | | 1 | [R].EGVFVSnGTHWFVTQR.[N] | N7(HexNAc(3)Hex(5)NeuGc(1)) | P0DTC2 | 0 | 3 | 1 | [illegible] |
| Trypsin | High | PMI-Byonic | Unambiguous | 1092 | 1107 | EGVFVSNGTHWFVTQR | 1098 | HexNAc(3)Hex(5) | 3 | 5 | | | | [R].EGVFVSnGTHWFVTQR.[N] | N7(HexNAc(3)Hex(5)) | P0DTC2 | 0 | 3 | 1 | [illegible] |
| Trypsin | High | PMI-Byonic | Unambiguous | 1092 | 1107 | EGVFVSNGTHWFVTQR | 1098 | HexNAc(3)Hex(6)Fuc(1)NeuGc(1) | 3 | 6 | 1 | | 1 | [R].EGVFVSnGTHWFVTQR.[N] | N7(HexNAc(3)Hex(6)Fuc(1)NeuGc(1)) | P0DTC2 | 0 | 3 | 1 | [illegible] |
| Trypsin | High | PMI-Byonic | Unambiguous | 1092 | 1107 | EGVFVSNGTHWFVTQR | 1098 | HexNAc(3)Hex(6)Fuc(1) | 3 | 6 | 1 | | | [R].EGVFVSnGTHWFVTQR.[N] | N7(HexNAc(3)Hex(6)Fuc(1)) | P0DTC2 | 0 | 3 | 1 | [illegible] |
| Trypsin | High | PMI-Byonic | Unambiguous | 1092 | 1107 | EGVFVSNGTHWFVTQR | 1098 | HexNAc(3)Hex(6)NeuAc(1) | 3 | 6 | | 1 | | [R].EGVFVSnGTHWFVTQR.[N] | N7(HexNAc(3)Hex(6)NeuAc(1)) | P0DTC2 | 0 | 3 | 1 | [illegible] |
| Trypsin | High | PMI-Byonic | Unambiguous | 1092 | 1107 | EGVFVSNGTHWFVTQR | 1098 | HexNAc(3)Hex(6)NeuGc(1) | 3 | 6 | | | 1 | [R].EGVFVSnGTHWFVTQR.[N] | N7(HexNAc(3)Hex(6)NeuGc(1)) | P0DTC2 | 0 | 3 | 1 | [illegible] |
| Trypsin | High | PMI-Byonic | Unambiguous | 1092 | 1107 | EGVFVSNGTHWFVTQR | 1098 | HexNAc(3)Hex(6) | 3 | 6 | | | | [R].EGVFVSnGTHWFVTQR.[N] | N7(HexNAc(3)Hex(6)) | P0DTC2 | 0 | 3 | 1 | [illegible] |
| Trypsin | High | PMI-Byonic | Unambiguous | 1092 | 1107 | EGVFVSNGTHWFVTQR | 1098 | HexNAc(4)Hex(3) | 4 | 3 | | | | [R].EGVFVSnGTHWFVTQR.[N] | N7(HexNAc(4)Hex(3)) | P0DTC2 | 0 | 3 | 1 | [illegible] |
| Trypsin | High | PMI-Byonic | Unambiguous | 1092 | 1107 | EGVFVSNGTHWFVTQR | 1098 | HexNAc(4)Hex(4)Fuc(1) | 4 | 4 | 1 | | | [R].EGVFVSnGTHWFVTQR.[N] | N7(HexNAc(4)Hex(4)Fuc(1)) | P0DTC2 | 0 | 3 | 1 | [illegible] |
| Trypsin | High | PMI-Byonic | Unambiguous | 1092 | 1107 | EGVFVSNGTHWFVTQR | 1098 | HexNAc(4)Hex(4) | 4 | 4 | | | | [R].EGVFVSnGTHWFVTQR.[N] | N7(HexNAc(4)Hex(4)) | P0DTC2 | 0 | 3 | 1 | [illegible] |
| Trypsin | High | PMI-Byonic | Unambiguous | 1092 | 1107 | EGVFVSNGTHWFVTQR | 1098 | HexNAc(4)Hex(5)Fuc(1)NeuAc(1) | 4 | 5 | 1 | 1 | | [R].EGVFVSnGTHWFVTQR.[N] | N7(HexNAc(4)Hex(5)Fuc(1)NeuAc(1)) | P0DTC2 | 0 | 3 | 1 | [illegible] |
| Trypsin | High | PMI-Byonic | Unambiguous | 1092 | 1107 | EGVFVSNGTHWFVTQR | 1098 | HexNAc(4)Hex(5)Fuc(1)NeuGc(1) | 4 | 5 | 1 | | 1 | [R].EGVFVSnGTHWFVTQR.[N] | N7(HexNAc(4)Hex(5)Fuc(1)NeuGc(1)) | P0DTC2 | 0 | 3 | 1 | [illegible] |
| Trypsin | High | PMI-Byonic | Unambiguous | 1092 | 1107 | EGVFVSNGTHWFVTQR | 1098 | HexNAc(4)Hex(5)Fuc(1) | 4 | 5 | 1 | | | [R].EGVFVSnGTHWFVTQR.[N] | N7(HexNAc(4)Hex(5)Fuc(1)) | P0DTC2 | 0 | 3 | 1 | [illegible] |
| Trypsin | High | PMI-Byonic | Unambiguous | 1092 | 1107 | EGVFVSNGTHWFVTQR | 1098 | HexNAc(4)Hex(5)Fuc(2) | 4 | 5 | 2 | | | [R].EGVFVSnGTHWFVTQR.[N] | N7(HexNAc(4)Hex(5)Fuc(2)) | P0DTC2 | 0 | 3 | 1 | [illegible] |
| Trypsin | High | PMI-Byonic | Unambiguous | 1092 | 1107 | EGVFVSNGTHWFVTQR | 1098 | HexNAc(4)Hex(5)Fuc(3)NeuAc(1)NeuGc(1) | 4 | 5 | 3 | 1 | 1 | [R].EGVFVSnGTHWFVTQR.[N] | N7(HexNAc(4)Hex(5)Fuc(3)NeuAc(1)NeuGc(1)) | P0DTC2 | 0 | 4 | 1 | [illegible] |
| Trypsin | High | PMI-Byonic | Unambiguous | 1092 | 1107 | EGVFVSNGTHWFVTQR | 1098 | HexNAc(4)Hex(5)Fuc(3)NeuAc(1) | 4 | 5 | 3 | 1 | | [R].EGVFVSnGTHWFVTQR.[N] | N7(HexNAc(4)Hex(5)Fuc(3)NeuAc(1)) | P0DTC2 | 0 | 3 | 1 | [illegible] |
| Trypsin | High | PMI-Byonic | Unambiguous | 1092 | 1107 | EGVFVSNGTHWFVTQR | 1098 | HexNAc(4)Hex(5)Fuc(3)NeuAc(2) | 4 | 5 | 3 | 2 | | [R].EGVFVSnGTHWFVTQR.[N] | N7(HexNAc(4)Hex(5)Fuc(3)NeuAc(2)) | P0DTC2 | 0 | 3 | 1 | [illegible] |
| Trypsin | High | PMI-Byonic | Unambiguous | 1092 | 1107 | EGVFVSNGTHWFVTQR | 1098 | HexNAc(4)Hex(5)NeuGc(1) | 4 | 5 | | | 1 | [R].EGVFVSnGTHWFVTQR.[N] | N7(HexNAc(4)Hex(5)NeuGc(1)) | P0DTC2 | 0 | 3 | 1 | [illegible] |
| Trypsin | High | PMI-Byonic | Unambiguous | 1092 | 1107 | EGVFVSNGTHWFVTQR | 1098 | HexNAc(4)Hex(5) | 4 | 5 | | | | [R].EGVFVSnGTHWFVTQR.[N] | N7(HexNAc(4)Hex(5)) | P0DTC2 | 0 | 3 | 1 | [illegible] |
| Trypsin | High | PMI-Byonic | Unambiguous | 1092 | 1107 | EGVFVSNGTHWFVTQR | 1098 | HexNAc(4)Hex(6)Fuc(1) | 4 | 6 | 1 | | | [R].EGVFVSnGTHWFVTQR.[N] | N7(HexNAc(4)Hex(6)Fuc(1)) | P0DTC2 | 0 | 3 | 1 | [illegible] |
| Trypsin | High | PMI-Byonic | Unambiguous | 1092 | 1107 | EGVFVSNGTHWFVTQR | 1098 | HexNAc(4)Hex(6)Fuc(2) | 4 | 6 | 2 | | | [R].EGVFVSnGTHWFVTQR.[N] | N7(HexNAc(4)Hex(6)Fuc(2)) | P0DTC2 | 0 | 3 | 1 | [illegible] |
| Trypsin | High | PMI-Byonic | Unambiguous | 1092 | 1107 | EGVFVSNGTHWFVTQR | 1098 | HexNAc(4)Hex(6)NeuAc(1) | 4 | 6 | | 1 | | [R].EGVFVSnGTHWFVTQR.[N] | N7(HexNAc(4)Hex(6)NeuAc(1)) | P0DTC2 | 0 | 3 | 1 | [illegible] |
| Trypsin | High | PMI-Byonic | Unambiguous | 1092 | 1107 | EGVFVSNGTHWFVTQR | 1098 | HexNAc(4)Hex(6)NeuGc(1) | 4 | 6 | | | 1 | [R].EGVFVSnGTHWFVTQR.[N] | N7(HexNAc(4)Hex(6)NeuGc(1)) | P0DTC2 | 0 | 3 | 1 | [illegible] |
| Trypsin | High | PMI-Byonic | Unambiguous | 1092 | 1107 | EGVFVSNGTHWFVTQR | 1098 | HexNAc(4)Hex(6) | 4 | 6 | | | | [R].EGVFVSnGTHWFVTQR.[N] | N7(HexNAc(4)Hex(6)) | P0DTC2 | 0 | 3 | 1 | [illegible] |
| Trypsin | High | PMI-Byonic | Unambiguous | 1092 | 1107 | EGVFVSNGTHWFVTQR | 1098 | HexNAc(4)Hex(7)Fuc(1) | 4 | 7 | 1 | | | [R].EGVFVSnGTHWFVTQR.[N] | N7(HexNAc(4)Hex(7)Fuc(1)) | P0DTC2 | 0 | 3 | 1 | [illegible] |
| Trypsin | High | PMI-Byonic | Unambiguous | 1092 | 1107 | EGVFVSNGTHWFVTQR | 1098 | HexNAc(4)Hex(7)NeuAc(1) | 4 | 7 | | 1 | | [R].EGVFVSnGTHWFVTQR.[N] | N7(HexNAc(4)Hex(7)NeuAc(1)) | P0DTC2 | 0 | 3 | 1 | [illegible] |
| Trypsin | High | PMI-Byonic | Unambiguous | 1092 | 1107 | EGVFVSNGTHWFVTQR | 1098 | HexNAc(4)Hex(7) | 4 | 7 | | | | [R].EGVFVSnGTHWFVTQR.[N] | N7(HexNAc(4)Hex(7)) | P0DTC2 | 0 | 3 | 1 | [illegible] |
| Trypsin | High | PMI-Byonic | Unambiguous | 1092 | 1107 | EGVFVSNGTHWFVTQR | 1098 | HexNAc(5)Hex(3)Fuc(1) | 5 | 3 | 1 | | | [R].EGVFVSnGTHWFVTQR.[N] | N7(HexNAc(5)Hex(3)Fuc(1)) | P0DTC2 | 0 | 3 | 1 | [illegible] |
| Trypsin | High | PMI-Byonic | Unambiguous | 1092 | 1107 | EGVFVSNGTHWFVTQR | 1098 | HexNAc(5)Hex(3) | 5 | 3 | | | | [R].EGVFVSnGTHWFVTQR.[N] | N7(HexNAc(5)Hex(3)) | P0DTC2 | 0 | 3 | 1 | [illegible] |
| Trypsin | High | PMI-Byonic | Unambiguous | 1092 | 1107 | EGVFVSNGTHWFVTQR | 1098 | HexNAc(5)Hex(4)Fuc(1) | 5 | 4 | 1 | | | [R].EGVFVSnGTHWFVTQR.[N] | N7(HexNAc(5)Hex(4)Fuc(1)) | P0DTC2 | 0 | 3 | 1 | [illegible] |
| Trypsin | High | PMI-Byonic | Unambiguous | 1092 | 1107 | EGVFVSNGTHWFVTQR | 1098 | HexNAc(5)Hex(4) | 5 | 4 | | | | [R].EGVFVSnGTHWFVTQR.[N] | N7(HexNAc(5)Hex(4)) | P0DTC2 | 0 | 3 | 1 | [illegible] |

| | | | | | | | | | | | | | | | | | | | | |
|---|---|---|---|---|---|---|---|---|---|---|---|---|---|---|---|---|---|---|---|---|
| Trypsin | High | PMI-Byonic | Unambiguous | 1092 | 1107 | EGVFVSNGTHWFVTQR | 1098 | HexNAc(5)Hex(5)Fuc(1)NeuGc(1) | 5 | 5 | 1 | | 1 | [R].EGVFVSnGTHWFVTQR.[N] | N7(HexNAc(5)Hex(5)Fuc(1)NeuGc(1)) | P0DTC2 | 0 | 3 | 1 | [illegible] |
| Trypsin | High | PMI-Byonic | Unambiguous | 1092 | 1107 | EGVFVSNGTHWFVTQR | 1098 | HexNAc(5)Hex(5)Fuc(1) | 5 | 5 | 1 | | | [R].EGVFVSnGTHWFVTQR.[N] | N7(HexNAc(5)Hex(5)Fuc(1)) | P0DTC2 | 0 | 3 | 1 | [illegible] |
| Trypsin | High | PMI-Byonic | Unambiguous | 1092 | 1107 | EGVFVSNGTHWFVTQR | 1098 | HexNAc(5)Hex(5)Fuc(2) | 5 | 5 | 2 | | | [R].EGVFVSnGTHWFVTQR.[N] | N7(HexNAc(5)Hex(5)Fuc(2)) | P0DTC2 | 0 | 3 | 1 | [illegible] |
| Trypsin | High | PMI-Byonic | Unambiguous | 1092 | 1107 | EGVFVSNGTHWFVTQR | 1098 | HexNAc(5)Hex(5)NeuGc(1) | 5 | 5 | | | 1 | [R].EGVFVSnGTHWFVTQR.[N] | N7(HexNAc(5)Hex(5)NeuGc(1)) | P0DTC2 | 0 | 3 | 1 | [illegible] |
| Trypsin | High | PMI-Byonic | Unambiguous | 1092 | 1107 | EGVFVSNGTHWFVTQR | 1098 | HexNAc(5)Hex(5) | 5 | 5 | | | | [R].EGVFVSnGTHWFVTQR.[N] | N7(HexNAc(5)Hex(5)) | P0DTC2 | 0 | 3 | 1 | [illegible] |
| Trypsin | High | PMI-Byonic | Unambiguous | 1092 | 1107 | EGVFVSNGTHWFVTQR | 1098 | HexNAc(5)Hex(6)Fuc(1)NeuAc(1) | 5 | 6 | 1 | 1 | | [R].EGVFVSnGTHWFVTQR.[N] | N7(HexNAc(5)Hex(6)Fuc(1)NeuAc(1)) | P0DTC2 | 0 | 3 | 1 | [illegible] |
| Trypsin | High | PMI-Byonic | Unambiguous | 1092 | 1107 | EGVFVSNGTHWFVTQR | 1098 | HexNAc(5)Hex(6)Fuc(1)NeuGc(1) | 5 | 6 | 1 | | 1 | [R].EGVFVSnGTHWFVTQR.[N] | N7(HexNAc(5)Hex(6)Fuc(1)NeuGc(1)) | P0DTC2 | 0 | 3 | 1 | [illegible] |
| Trypsin | High | PMI-Byonic | Unambiguous | 1092 | 1107 | EGVFVSNGTHWFVTQR | 1098 | HexNAc(5)Hex(6)Fuc(1)NeuGc(2) | 5 | 6 | 1 | | 2 | [R].EGVFVSnGTHWFVTQR.[N] | N7(HexNAc(5)Hex(6)Fuc(1)NeuGc(2)) | P0DTC2 | 0 | 3 | 1 | [illegible] |
| Trypsin | High | PMI-Byonic | Unambiguous | 1092 | 1107 | EGVFVSNGTHWFVTQR | 1098 | HexNAc(5)Hex(6)Fuc(2)NeuGc(1) | 5 | 6 | 2 | | 1 | [R].EGVFVSnGTHWFVTQR.[N] | N7(HexNAc(5)Hex(6)Fuc(2)NeuGc(1)) | P0DTC2 | 0 | 3 | 1 | [illegible] |
| Trypsin | High | PMI-Byonic | Unambiguous | 1092 | 1107 | EGVFVSNGTHWFVTQR | 1098 | HexNAc(5)Hex(6)Fuc(2) | 5 | 6 | 2 | | | [R].EGVFVSnGTHWFVTQR.[N] | N7(HexNAc(5)Hex(6)Fuc(2)) | P0DTC2 | 0 | 3 | 1 | [illegible] |
| Trypsin | High | PMI-Byonic | Unambiguous | 1092 | 1107 | EGVFVSNGTHWFVTQR | 1098 | HexNAc(5)Hex(6)Fuc(3) | 5 | 6 | 3 | | | [R].EGVFVSnGTHWFVTQR.[N] | N7(HexNAc(5)Hex(6)Fuc(3)) | P0DTC2 | 0 | 3 | 1 | [illegible] |
| Trypsin | High | PMI-Byonic | Unambiguous | 1092 | 1107 | EGVFVSNGTHWFVTQR | 1098 | HexNAc(5)Hex(6)NeuAc(1)NeuGc(1) | 5 | 6 | | 1 | 1 | [R].EGVFVSnGTHWFVTQR.[N] | N7(HexNAc(5)Hex(6)NeuAc(1)NeuGc(1)) | P0DTC2 | 0 | 3 | 1 | [illegible] |
| Trypsin | High | PMI-Byonic | Unambiguous | 1092 | 1107 | EGVFVSNGTHWFVTQR | 1098 | HexNAc(5)Hex(6)NeuAc(1) | 5 | 6 | | 1 | | [R].EGVFVSnGTHWFVTQR.[N] | N7(HexNAc(5)Hex(6)NeuAc(1)) | P0DTC2 | 0 | 3 | 1 | [illegible] |
| Trypsin | High | PMI-Byonic | Unambiguous | 1092 | 1107 | EGVFVSNGTHWFVTQR | 1098 | HexNAc(5)Hex(6)NeuGc(1) | 5 | 6 | | | 1 | [R].EGVFVSnGTHWFVTQR.[N] | N7(HexNAc(5)Hex(6)NeuGc(1)) | P0DTC2 | 0 | 3 | 1 | [illegible] |
| Trypsin | High | PMI-Byonic | Unambiguous | 1092 | 1107 | EGVFVSNGTHWFVTQR | 1098 | HexNAc(5)Hex(6)NeuGc(2) | 5 | 6 | | | 2 | [R].EGVFVSnGTHWFVTQR.[N] | N7(HexNAc(5)Hex(6)NeuGc(2)) | P0DTC2 | 0 | 3 | 1 | [illegible] |
| Trypsin | High | PMI-Byonic | Unambiguous | 1092 | 1107 | EGVFVSNGTHWFVTQR | 1098 | HexNAc(5)Hex(6) | 5 | 6 | | | | [R].EGVFVSnGTHWFVTQR.[N] | N7(HexNAc(5)Hex(6)) | P0DTC2 | 0 | 3 | 1 | [illegible] |
| Trypsin | High | PMI-Byonic | Unambiguous | 1092 | 1107 | EGVFVSNGTHWFVTQR | 1098 | HexNAc(5)Hex(7) | 5 | 7 | | | | [R].EGVFVSnGTHWFVTQR.[N] | N7(HexNAc(5)Hex(7)) | P0DTC2 | 0 | 3 | 1 | [illegible] |
| Trypsin | High | PMI-Byonic | Unambiguous | 1092 | 1107 | EGVFVSNGTHWFVTQR | 1098 | HexNAc(6)Hex(3)Fuc(1) | 6 | 3 | 1 | | | [R].EGVFVSnGTHWFVTQR.[N] | N7(HexNAc(6)Hex(3)Fuc(1)) | P0DTC2 | 0 | 3 | 1 | [illegible] |
| Trypsin | High | PMI-Byonic | Unambiguous | 1092 | 1107 | EGVFVSNGTHWFVTQR | 1098 | HexNAc(6)Hex(3) | 6 | 3 | | | | [R].EGVFVSnGTHWFVTQR.[N] | N7(HexNAc(6)Hex(3)) | P0DTC2 | 0 | 3 | 1 | [illegible] |
| Trypsin | High | PMI-Byonic | Unambiguous | 1092 | 1107 | EGVFVSNGTHWFVTQR | 1098 | HexNAc(6)Hex(4)Fuc(1) | 6 | 4 | 1 | | | [R].EGVFVSnGTHWFVTQR.[N] | N7(HexNAc(6)Hex(4)Fuc(1)) | P0DTC2 | 0 | 3 | 1 | [illegible] |
| Trypsin | High | PMI-Byonic | Unambiguous | 1092 | 1107 | EGVFVSNGTHWFVTQR | 1098 | HexNAc(6)Hex(4) | 6 | 4 | | | | [R].EGVFVSnGTHWFVTQR.[N] | N7(HexNAc(6)Hex(4)) | P0DTC2 | 0 | 3 | 1 | [illegible] |
| Trypsin | High | PMI-Byonic | Unambiguous | 1092 | 1107 | EGVFVSNGTHWFVTQR | 1098 | HexNAc(6)Hex(5)Fuc(1) | 6 | 5 | 1 | | | [R].EGVFVSnGTHWFVTQR.[N] | N7(HexNAc(6)Hex(5)Fuc(1)) | P0DTC2 | 0 | 3 | 1 | [illegible] |
| Trypsin | High | PMI-Byonic | Unambiguous | 1092 | 1107 | EGVFVSNGTHWFVTQR | 1098 | HexNAc(6)Hex(5) | 6 | 5 | | | | [R].EGVFVSnGTHWFVTQR.[N] | N7(HexNAc(6)Hex(5)) | P0DTC2 | 0 | 3 | 1 | [illegible] |
| Trypsin | High | PMI-Byonic | Unambiguous | 1092 | 1107 | EGVFVSNGTHWFVTQR | 1098 | HexNAc(6)Hex(6)Fuc(1)NeuGc(2) | 6 | 6 | 1 | | 2 | [R].EGVFVSnGTHWFVTQR.[N] | N7(HexNAc(6)Hex(6)Fuc(1)NeuGc(2)) | P0DTC2 | 0 | 3 | 1 | [illegible] |
| Trypsin | High | PMI-Byonic | Unambiguous | 1092 | 1107 | EGVFVSNGTHWFVTQR | 1098 | HexNAc(6)Hex(6)Fuc(1) | 6 | 6 | 1 | | | [R].EGVFVSnGTHWFVTQR.[N] | N7(HexNAc(6)Hex(6)Fuc(1)) | P0DTC2 | 0 | 3 | 1 | [illegible] |
| Trypsin | High | PMI-Byonic | Unambiguous | 1092 | 1107 | EGVFVSNGTHWFVTQR | 1098 | HexNAc(6)Hex(6)NeuAc(1)NeuGc(1) | 6 | 6 | | 1 | 1 | [R].EGVFVSnGTHWFVTQR.[N] | N7(HexNAc(6)Hex(6)NeuAc(1)NeuGc(1)) | P0DTC2 | 0 | 3 | 1 | [illegible] |
| Trypsin | High | PMI-Byonic | Unambiguous | 1092 | 1107 | EGVFVSNGTHWFVTQR | 1098 | HexNAc(6)Hex(6) | 6 | 6 | | | | [R].EGVFVSnGTHWFVTQR.[N] | N7(HexNAc(6)Hex(6)) | P0DTC2 | 0 | 3 | 1 | [illegible] |
| Trypsin | High | PMI-Byonic | Unambiguous | 1092 | 1107 | EGVFVSNGTHWFVTQR | 1098 | HexNAc(6)Hex(7)Fuc(1)NeuAc(1) | 6 | 7 | 1 | 1 | | [R].EGVFVSnGTHWFVTQR.[N] | N7(HexNAc(6)Hex(7)Fuc(1)NeuAc(1)) | P0DTC2 | 0 | 3 | 1 | [illegible] |
| Trypsin | High | PMI-Byonic | Unambiguous | 1092 | 1107 | EGVFVSNGTHWFVTQR | 1098 | HexNAc(6)Hex(7)Fuc(1)NeuGc(1) | 6 | 7 | 1 | | 1 | [R].EGVFVSnGTHWFVTQR.[N] | N7(HexNAc(6)Hex(7)Fuc(1)NeuGc(1)) | P0DTC2 | 0 | 3 | 1 | [illegible] |
| Trypsin | High | PMI-Byonic | Unambiguous | 1092 | 1107 | EGVFVSNGTHWFVTQR | 1098 | HexNAc(6)Hex(7)Fuc(1) | 6 | 7 | 1 | | | [R].EGVFVSnGTHWFVTQR.[N] | N7(HexNAc(6)Hex(7)Fuc(1)) | P0DTC2 | 0 | 3 | 1 | [illegible] |
| Trypsin | High | PMI-Byonic | Unambiguous | 1092 | 1107 | EGVFVSNGTHWFVTQR | 1098 | HexNAc(6)Hex(7)NeuAc(1)NeuGc(1) | 6 | 7 | | 1 | 1 | [R].EGVFVSnGTHWFVTQR.[N] | N7(HexNAc(6)Hex(7)NeuAc(1)NeuGc(1)) | P0DTC2 | 0 | 4 | 1 | [illegible] |
| Trypsin | High | PMI-Byonic | Unambiguous | 1092 | 1107 | EGVFVSNGTHWFVTQR | 1098 | HexNAc(6)Hex(7)NeuAc(1) | 6 | 7 | | 1 | | [R].EGVFVSnGTHWFVTQR.[N] | N7(HexNAc(6)Hex(7)NeuAc(1)) | P0DTC2 | 0 | 3 | 1 | [illegible] |
| Trypsin | High | PMI-Byonic | Unambiguous | 1092 | 1107 | EGVFVSNGTHWFVTQR | 1098 | HexNAc(6)Hex(7)NeuGc(2) | 6 | 7 | | | 2 | [R].EGVFVSnGTHWFVTQR.[N] | N7(HexNAc(6)Hex(7)NeuGc(2)) | P0DTC2 | 0 | 4 | 1 | [illegible] |
| Trypsin | High | PMI-Byonic | Unambiguous | 1092 | 1107 | EGVFVSNGTHWFVTQR | 1098 | HexNAc(6)Hex(7) | 6 | 7 | | | | [R].EGVFVSnGTHWFVTQR.[N] | N7(HexNAc(6)Hex(7)) | P0DTC2 | 0 | 4 | 1 | [illegible] |
| Trypsin | High | PMI-Byonic | Unambiguous | 1092 | 1107 | EGVFVSNGTHWFVTQR | 1098 | HexNAc(7)Hex(6) | 7 | 6 | | | | [R].EGVFVSnGTHWFVTQR.[N] | N7(HexNAc(7)Hex(6)) | P0DTC2 | 0 | 3 | 1 | [illegible] |
| Trypsin | High | PMI-Byonic | Unambiguous | 1092 | 1107 | EGVFVSNGTHWFVTQR | 1098 | HexNAc(7)Hex(7) | 7 | 7 | | | | [R].EGVFVSnGTHWFVTQR.[N] | N7(HexNAc(7)Hex(7)) | P0DTC2 | 0 | 3 | 1 | [illegible] |
| Trypsin | High | PMI-Byonic | Unambiguous | 1108 | 1154 | NFYEPQIITTDNTFVSGNCDVVIGIVNNTVYDPLQPELDSFKEELDK | 1134 | HexNAc(3)Hex(5) | 3 | 5 | | | | [R].NFYEPQIITTDNTFVSGNcDVVIGIVnNTVYDPLQPELDSFKEELDK.[Y] | C19(Carbamidomethyl); N27(HexNAc(3)Hex(5)) | P0DTC2 | 1 | 4 | 1 | [illegible] |
| Trypsin | High | PMI-Byonic | Unambiguous | 1192 | 1205 | NLNESLIDLQELGK | 1194 | HexNAc(4)Hex(3)Fuc(1) | 4 | 3 | 1 | | | [K].NLnESLIDLQELGK.[Y] | N3(HexNAc(4)Hex(3)Fuc(1)) | P0DTC2 | 0 | 3 | 1 | [illegible] |
| Trypsin | High | PMI-Byonic | Unambiguous | 1192 | 1205 | NLNESLIDLQELGK | 1194 | HexNAc(4)Hex(5)Fuc(1) | 4 | 5 | 1 | | | [K].NLnESLIDLQELGK.[Y] | N3(HexNAc(4)Hex(5)Fuc(1)) | P0DTC2 | 0 | 3 | 1 | [illegible] |
| Trypsin | High | PMI-Byonic | Unambiguous | 1192 | 1205 | NLNESLIDLQELGK | 1194 | HexNAc(4)Hex(5)Fuc(3) | 4 | 5 | 3 | | | [K].NLnESLIDLQELGK.[Y] | N3(HexNAc(4)Hex(5)Fuc(3)) | P0DTC2 | 0 | 3 | 1 | [illegible] |
| Trypsin | High | PMI-Byonic | Unambiguous | 1192 | 1205 | NLNESLIDLQELGK | 1194 | HexNAc(4)Hex(5) | 4 | 5 | | | | [K].NLnESLIDLQELGK.[Y] | N3(HexNAc(4)Hex(5)) | P0DTC2 | 0 | 3 | 1 | [illegible] |
| Trypsin | High | PMI-Byonic | Unambiguous | 1192 | 1205 | NLNESLIDLQELGK | 1194 | HexNAc(5)Hex(3)Fuc(1) | 5 | 3 | 1 | | | [K].NLnESLIDLQELGK.[Y] | N3(HexNAc(5)Hex(3)Fuc(1)) | P0DTC2 | 0 | 3 | 1 | [illegible] |
| Trypsin | High | PMI-Byonic | Unambiguous | 1192 | 1205 | NLNESLIDLQELGK | 1194 | HexNAc(5)Hex(4)Fuc(1) | 5 | 4 | 1 | | | [K].NLnESLIDLQELGK.[Y] | N3(HexNAc(5)Hex(4)Fuc(1)) | P0DTC2 | 0 | 3 | 1 | [illegible] |
| Trypsin | High | PMI-Byonic | Unambiguous | 1192 | 1205 | NLNESLIDLQELGK | 1194 | HexNAc(5)Hex(5)Fuc(1) | 5 | 5 | 1 | | | [K].NLnESLIDLQELGK.[Y] | N3(HexNAc(5)Hex(5)Fuc(1)) | P0DTC2 | 0 | 3 | 1 | [illegible] |
| Trypsin | High | PMI-Byonic | Unambiguous | 1192 | 1205 | NLNESLIDLQELGK | 1194 | HexNAc(5)Hex(6)Fuc(1)NeuGc(1) | 5 | 6 | 1 | | 1 | [K].NLnESLIDLQELGK.[Y] | N3(HexNAc(5)Hex(6)Fuc(1)NeuGc(1)) | P0DTC2 | 0 | 3 | 1 | [illegible] |
| Trypsin | High | PMI-Byonic | Unambiguous | 1192 | 1205 | NLNESLIDLQELGK | 1194 | HexNAc(5)Hex(6)Fuc(1) | 5 | 6 | 1 | | | [K].NLnESLIDLQELGK.[Y] | N3(HexNAc(5)Hex(6)Fuc(1)) | P0DTC2 | 0 | 3 | 1 | [illegible] |
| Trypsin | High | PMI-Byonic | Unambiguous | 1192 | 1205 | NLNESLIDLQELGK | 1194 | HexNAc(5)Hex(6)Fuc(2)NeuAc(1) | 5 | 6 | 2 | 1 | | [K].NLnESLIDLQELGK.[Y] | N3(HexNAc(5)Hex(6)Fuc(2)NeuAc(1)) | P0DTC2 | 0 | 3 | 1 | [illegible] |
| Trypsin | High | PMI-Byonic | Unambiguous | 1192 | 1205 | NLNESLIDLQELGK | 1194 | HexNAc(5)Hex(6)Fuc(2) | 5 | 6 | 2 | | | [K].NLnESLIDLQELGK.[Y] | N3(HexNAc(5)Hex(6)Fuc(2)) | P0DTC2 | 0 | 3 | 1 | [illegible] |
| Trypsin | High | PMI-Byonic | Unambiguous | 1192 | 1205 | NLNESLIDLQELGK | 1194 | HexNAc(5)Hex(6)NeuAc(1)NeuGc(1) | 5 | 6 | | 1 | 1 | [K].NLnESLIDLQELGK.[Y] | N3(HexNAc(5)Hex(6)NeuAc(1)NeuGc(1)) | P0DTC2 | 0 | 3 | 1 | [illegible] |
| Trypsin | High | PMI-Byonic | Unambiguous | 1192 | 1205 | NLNESLIDLQELGK | 1194 | HexNAc(5)Hex(6) | 5 | 6 | | | | [K].NLnESLIDLQELGK.[Y] | N3(HexNAc(5)Hex(6)) | P0DTC2 | 0 | 3 | 1 | [illegible] |
| Trypsin | High | PMI-Byonic | Unambiguous | 1192 | 1205 | NLNESLIDLQELGK | 1194 | HexNAc(5)Hex(7)Fuc(1)NeuAc(1) | 5 | 7 | 1 | 1 | | [K].NLnESLIDLQELGK.[Y] | N3(HexNAc(5)Hex(7)Fuc(1)NeuAc(1)) | P0DTC2 | 0 | 3 | 1 | [illegible] |
| Trypsin | High | PMI-Byonic | Unambiguous | 1192 | 1205 | NLNESLIDLQELGK | 1194 | HexNAc(6)Hex(3)Fuc(1) | 6 | 3 | 1 | | | [K].NLnESLIDLQELGK.[Y] | N3(HexNAc(6)Hex(3)Fuc(1)) | P0DTC2 | 0 | 3 | 1 | [illegible] |
| Trypsin | High | PMI-Byonic | Unambiguous | 1192 | 1205 | NLNESLIDLQELGK | 1194 | HexNAc(6)Hex(4)Fuc(1) | 6 | 4 | 1 | | | [K].NLnESLIDLQELGK.[Y] | N3(HexNAc(6)Hex(4)Fuc(1)) | P0DTC2 | 0 | 3 | 1 | [illegible] |
| Trypsin | High | PMI-Byonic | Unambiguous | 1192 | 1205 | NLNESLIDLQELGK | 1194 | HexNAc(6)Hex(5)Fuc(1) | 6 | 5 | 1 | | | [K].NLnESLIDLQELGK.[Y] | N3(HexNAc(6)Hex(5)Fuc(1)) | P0DTC2 | 0 | 3 | 1 | [illegible] |
| Trypsin | High | PMI-Byonic | Unambiguous | 1192 | 1205 | NLNESLIDLQELGK | 1194 | HexNAc(6)Hex(7)Fuc(1)NeuAc(1) | 6 | 7 | 1 | 1 | | [K].NLnESLIDLQELGK.[Y] | N3(HexNAc(6)Hex(7)Fuc(1)NeuAc(1)) | P0DTC2 | 0 | 3 | 1 | [illegible] |
| Trypsin | High | PMI-Byonic | Unambiguous | 1192 | 1205 | NLNESLIDLQELGK | 1194 | HexNAc(6)Hex(7)Fuc(1)NeuGc(1) | 6 | 7 | 1 | | 1 | [K].NLnESLIDLQELGK.[Y] | N3(HexNAc(6)Hex(7)Fuc(1)NeuGc(1)) | P0DTC2 | 0 | 3 | 1 | [illegible] |
| Trypsin | High | PMI-Byonic | Unambiguous | 1192 | 1205 | NLNESLIDLQELGK | 1194 | HexNAc(6)Hex(7)Fuc(1) | 6 | 7 | 1 | | | [K].NLnESLIDLQELGK.[Y] | N3(HexNAc(6)Hex(7)Fuc(1)) | P0DTC2 | 0 | 3 | 1 | [illegible] |
| Trypsin | High | PMI-Byonic | Unambiguous | 1192 | 1205 | NLNESLIDLQELGK | 1194 | HexNAc(6)Hex(7)Fuc(4)NeuGc(1) | 6 | 7 | 4 | | 1 | [K].NLnESLIDLQELGK.[Y] | N3(HexNAc(6)Hex(7)Fuc(4)NeuGc(1)) | P0DTC2 | 0 | 3 | 1 | [illegible] |
| Trypsin | High | PMI-Byonic | Unambiguous | 1192 | 1205 | NLNESLIDLQELGK | 1194 | HexNAc(6)Hex(7)Fuc(4) | 6 | 7 | 4 | | | [K].NLnESLIDLQELGK.[Y] | N3(HexNAc(6)Hex(7)Fuc(4)) | P0DTC2 | 0 | 3 | 1 | [illegible] |
| Trypsin | High | PMI-Byonic | Unambiguous | 1192 | 1205 | NLNESLIDLQELGK | 1194 | HexNAc(6)Hex(7)NeuAc(1)NeuGc(1) | 6 | 7 | | 1 | 1 | [K].NLnESLIDLQELGK.[Y] | N3(HexNAc(6)Hex(7)NeuAc(1)NeuGc(1)) | P0DTC2 | 0 | 3 | 1 | [illegible] |
| Trypsin | High | PMI-Byonic | Unambiguous | 1192 | 1205 | NLNESLIDLQELGK | 1194 | HexNAc(6)Hex(7)NeuAc(1) | 6 | 7 | | 1 | | [K].NLnESLIDLQELGK.[Y] | N3(HexNAc(6)Hex(7)NeuAc(1)) | P0DTC2 | 0 | 3 | 1 | [illegible] |
| Trypsin | High | PMI-Byonic | Unambiguous | 1192 | 1205 | NLNESLIDLQELGK | 1194 | HexNAc(6)Hex(7) | 6 | 7 | | | | [K].NLnESLIDLQELGK.[Y] | N3(HexNAc(6)Hex(7)) | P0DTC2 | 0 | 3 | 1 | [illegible] |
| Trypsin(semi) | High | PMI-Byonic | Unambiguous | 14 | 21 | QCVNLTTR | 17 | HexNAc(2)Hex(7) | 2 | 7 | | | | [S].qcVnLTTR.[T] | Q1(Gln->pyro-Glu); C2(Carbamidomethyl); N4(HexNAc(2)Hex(7)) | P0DTC2 | 0 | 2 | 1 | [illegible] |
| Trypsin(semi) | High | PMI-Byonic | Unambiguous | 14 | 21 | QCVNLTTR | 17 | HexNAc(3)Hex(6)Fuc(1) | 3 | 6 | 1 | | | [S].qcVnLTTR.[T] | Q1(Gln->pyro-Glu); C2(Carbamidomethyl); N4(HexNAc(3)Hex(6)Fuc(1)) | P0DTC2 | 0 | 2 | 1 | [illegible] |
| Trypsin(semi) | High | PMI-Byonic | Unambiguous | 14 | 21 | QCVNLTTR | 17 | HexNAc(4)Hex(3)Fuc(1) | 4 | 3 | 1 | | | [S].qcVnLTTR.[T] | Q1(Gln->pyro-Glu); C2(Carbamidomethyl); N4(HexNAc(4)Hex(3)Fuc(1)) | P0DTC2 | 0 | 2 | 1 | [illegible] |
| Trypsin(semi) | High | PMI-Byonic | Unambiguous | 14 | 21 | QCVNLTTR | 17 | HexNAc(4)Hex(4)Fuc(2) | 4 | 4 | 2 | | | [S].qcVnLTTR.[T] | Q1(Gln->pyro-Glu); C2(Carbamidomethyl); N4(HexNAc(4)Hex(4)Fuc(2)) | P0DTC2 | 0 | 2 | 1 | [illegible] |
| Trypsin(semi) | High | PMI-Byonic | Unambiguous | 14 | 21 | QCVNLTTR | 17 | HexNAc(4)Hex(5)Fuc(3) | 4 | 5 | 3 | | | [S].qcVnLTTR.[T] | Q1(Gln->pyro-Glu); C2(Carbamidomethyl); N4(HexNAc(4)Hex(5)Fuc(3)) | P0DTC2 | 0 | 3 | 1 | [illegible] |
| Trypsin(semi) | High | PMI-Byonic | Unambiguous | 14 | 21 | QCVNLTTR | 17 | HexNAc(5)Hex(5)Fuc(2) | 5 | 5 | 2 | | | [S].qcVnLTTR.[T] | Q1(Gln->pyro-Glu); C2(Carbamidomethyl); N4(HexNAc(5)Hex(5)Fuc(2)) | P0DTC2 | 0 | 3 | 1 | [illegible] |
| Trypsin(semi) | High | PMI-Byonic | Unambiguous | 66 | 77 | HAIHVSGTNGTK | 74 | HexNAc(5)Hex(4)Fuc(2)NeuGc(1) | 5 | 4 | 2 | | 1 | [F].HAIHVSGTnGTK.[R] | N9(HexNAc(5)Hex(4)Fuc(2)NeuGc(1)) | P0DTC2 | 0 | 4 | 1 | [illegible] |
| Trypsin(semi) | High | PMI-Byonic | Unambiguous | 66 | 77 | HAIHVSGTNGTK | 74 | HexNAc(5)Hex(5)Fuc(1) | 5 | 5 | 1 | | | [F].HAIHVSGTnGTK.[R] | N9(HexNAc(5)Hex(5)Fuc(1)) | P0DTC2 | 0 | 4 | 1 | [illegible] |
| Trypsin(semi) | High | PMI-Byonic | Unambiguous | 66 | 77 | HAIHVSGTNGTK | 74 | HexNAc(5)Hex(5)Fuc(3) | 5 | 5 | 3 | | | [F].HAIHVSGTnGTK.[R] | N9(HexNAc(5)Hex(5)Fuc(3)) | P0DTC2 | 0 | 4 | 1 | [illegible] |
| Trypsin(semi) | High | PMI-Byonic | Unambiguous | 114 | 129 | TQSLLIVNNATNVVIK | 122 | HexNAc(2)Hex(5) | 2 | 5 | | | | [K].TQSLLIVNnATNVVIK.[V] | N9(HexNAc(2)Hex(5)) | P0DTC2 | 0 | 2 | 1 | [illegible] |
| Trypsin(semi) | High | PMI-Byonic | Unambiguous | 114 | 129 | TQSLLIVNNATNVVIK | 122 | HexNAc(2)Hex(6) | 2 | 6 | | | | [K].TQSLLIVNnATNVVIK.[V] | N9(HexNAc(2)Hex(6)) | P0DTC2 | 0 | 3 | 1 | [illegible] |
| Trypsin(semi) | High | PMI-Byonic | Unambiguous | 114 | 129 | TQSLLIVNNATNVVIK | 122 | HexNAc(2)Hex(7) | 2 | 7 | | | | [K].TQSLLIVNnATNVVIK.[V] | N9(HexNAc(2)Hex(7)) | P0DTC2 | 0 | 3 | 1 | [illegible] |
| Trypsin(semi) | High | PMI-Byonic | Unambiguous | 114 | 129 | TQSLLIVNNATNVVIK | 122 | HexNAc(2)Hex(8) | 2 | 8 | | | | [K].TQSLLIVNnATNVVIK.[V] | N9(HexNAc(2)Hex(8)) | P0DTC2 | 0 | 3 | 1 | [illegible] |
| Trypsin(semi) | High | PMI-Byonic | Unambiguous | 114 | 129 | TQSLLIVNNATNVVIK | 122 | HexNAc(3)Hex(3)Fuc(1) | 3 | 3 | 1 | | | [K].TQSLLIVNnATNVVIK.[V] | N9(HexNAc(3)Hex(3)Fuc(1)) | P0DTC2 | 0 | 3 | 1 | [illegible] |
| Trypsin(semi) | High | PMI-Byonic | Unambiguous | 114 | 129 | TQSLLIVNNATNVVIK | 122 | HexNAc(3)Hex(3) | 3 | 3 | | | | [K].TQSLLIVNnATNVVIK.[V] | N9(HexNAc(3)Hex(3)) | P0DTC2 | 0 | 3 | 1 | [illegible] |
| Trypsin(semi) | High | PMI-Byonic | Unambiguous | 114 | 129 | TQSLLIVNNATNVVIK | 122 | HexNAc(3)Hex(4)Fuc(1) | 3 | 4 | 1 | | | [K].TQSLLIVNnATNVVIK.[V] | N9(HexNAc(3)Hex(4)Fuc(1)) | P0DTC2 | 0 | 3 | 1 | [illegible] |
| Trypsin(semi) | High | PMI-Byonic | Unambiguous | 114 | 129 | TQSLLIVNNATNVVIK | 122 | HexNAc(3)Hex(4)Fuc(2) | 3 | 4 | 2 | | | [K].TQSLLIVNnATNVVIK.[V] | N9(HexNAc(3)Hex(4)Fuc(2)) | P0DTC2 | 0 | 3 | 1 | [illegible] |
| Trypsin(semi) | High | PMI-Byonic | Unambiguous | 114 | 129 | TQSLLIVNNATNVVIK | 122 | HexNAc(3)Hex(4)NeuGc(1) | 3 | 4 | | | 1 | [K].TQSLLIVNnATNVVIK.[V] | N9(HexNAc(3)Hex(4)NeuGc(1)) | P0DTC2 | 0 | 3 | 1 | [illegible] |
| Trypsin(semi) | High | PMI-Byonic | Unambiguous | 114 | 129 | TQSLLIVNNATNVVIK | 122 | HexNAc(3)Hex(4) | 3 | 4 | | | | [K].TQSLLIVNnATNVVIK.[V] | N9(HexNAc(3)Hex(4)) | P0DTC2 | 0 | 3 | 1 | [illegible] |
| Trypsin(semi) | High | PMI-Byonic | Unambiguous | 114 | 129 | TQSLLIVNNATNVVIK | 122 | HexNAc(3)Hex(5)Fuc(1) | 3 | 5 | 1 | | | [K].TQSLLIVNnATNVVIK.[V] | N9(HexNAc(3)Hex(5)Fuc(1)) | P0DTC2 | 0 | 3 | 1 | [illegible] |
| Trypsin(semi) | High | PMI-Byonic | Unambiguous | 114 | 129 | TQSLLIVNNATNVVIK | 122 | HexNAc(3)Hex(5)NeuAc(1) | 3 | 5 | | 1 | | [K].TQSLLIVNnATNVVIK.[V] | N9(HexNAc(3)Hex(5)NeuAc(1)) | P0DTC2 | 0 | 3 | 1 | [illegible] |
| Trypsin(semi) | High | PMI-Byonic | Unambiguous | 114 | 129 | TQSLLIVNNATNVVIK | 122 | HexNAc(3)Hex(5)NeuGc(1) | 3 | 5 | | | 1 | [K].TQSLLIVNnATNVVIK.[V] | N9(HexNAc(3)Hex(5)NeuGc(1)) | P0DTC2 | 0 | 3 | 1 | [illegible] |
| Trypsin(semi) | High | PMI-Byonic | Unambiguous | 114 | 129 | TQSLLIVNNATNVVIK | 122 | HexNAc(3)Hex(5) | 3 | 5 | | | | [K].TQSLLIVNnATNVVIK.[V] | N9(HexNAc(3)Hex(5)) | P0DTC2 | 0 | 3 | 1 | [illegible] |
| Trypsin(semi) | High | PMI-Byonic | Unambiguous | 114 | 129 | TQSLLIVNNATNVVIK | 122 | HexNAc(3)Hex(6)Fuc(1) | 3 | 6 | 1 | | | [K].TQSLLIVNnATNVVIK.[V] | N9(HexNAc(3)Hex(6)Fuc(1)) | P0DTC2 | 0 | 3 | 1 | [illegible] |
| Trypsin(semi) | High | PMI-Byonic | Unambiguous | 114 | 129 | TQSLLIVNNATNVVIK | 122 | HexNAc(3)Hex(6)NeuAc(1) | 3 | 6 | | 1 | | [K].TQSLLIVNnATNVVIK.[V] | N9(HexNAc(3)Hex(6)NeuAc(1)) | P0DTC2 | 0 | 3 | 1 | [illegible] |
| Trypsin(semi) | High | PMI-Byonic | Unambiguous | 114 | 129 | TQSLLIVNNATNVVIK | 122 | HexNAc(3)Hex(6)NeuGc(1) | 3 | 6 | | | 1 | [K].TQSLLIVNnATNVVIK.[V] | N9(HexNAc(3)Hex(6)NeuGc(1)) | P0DTC2 | 0 | 3 | 1 | [illegible] |
| Trypsin(semi) | High | PMI-Byonic | Unambiguous | 114 | 129 | TQSLLIVNNATNVVIK | 122 | HexNAc(3)Hex(6) | 3 | 6 | | | | [K].TQSLLIVNnATNVVIK.[V] | N9(HexNAc(3)Hex(6)) | P0DTC2 | 0 | 3 | 1 | [illegible] |
| Trypsin(semi) | High | PMI-Byonic | Unambiguous | 114 | 129 | TQSLLIVNNATNVVIK | 122 | HexNAc(4)Hex(3)Fuc(1) | 4 | 3 | 1 | | | [K].TQSLLIVNnATNVVIK.[V] | N9(HexNAc(4)Hex(3)Fuc(1)) | P0DTC2 | 0 | 3 | 1 | [illegible] |
| Trypsin(semi) | High | PMI-Byonic | Unambiguous | 114 | 129 | TQSLLIVNNATNVVIK | 122 | HexNAc(4)Hex(3) | 4 | 3 | | | | [K].TQSLLIVNnATNVVIK.[V] | N9(HexNAc(4)Hex(3)) | P0DTC2 | 0 | 3 | 1 | [illegible] |
| Trypsin(semi) | High | PMI-Byonic | Unambiguous | 114 | 129 | TQSLLIVNNATNVVIK | 122 | HexNAc(4)Hex(4)Fuc(1) | 4 | 4 | 1 | | | [K].TQSLLIVNnATNVVIK.[V] | N9(HexNAc(4)Hex(4)Fuc(1)) | P0DTC2 | 0 | 3 | 1 | [illegible] |
| Trypsin(semi) | High | PMI-Byonic | Unambiguous | 114 | 129 | TQSLLIVNNATNVVIK | 122 | HexNAc(4)Hex(4)Fuc(2) | 4 | 4 | 2 | | | [K].TQSLLIVNnATNVVIK.[V] | N9(HexNAc(4)Hex(4)Fuc(2)) | P0DTC2 | 0 | 3 | 1 | [illegible] |
| Trypsin(semi) | High | PMI-Byonic | Unambiguous | 114 | 129 | TQSLLIVNNATNVVIK | 122 | HexNAc(4)Hex(4) | 4 | 4 | | | | [K].TQSLLIVNnATNVVIK.[V] | N9(HexNAc(4)Hex(4)) | P0DTC2 | 0 | 3 | 1 | [illegible] |
| Trypsin(semi) | High | PMI-Byonic | Unambiguous | 114 | 129 | TQSLLIVNNATNVVIK | 122 | HexNAc(4)Hex(5)Fuc(1)NeuAc(1) | 4 | 5 | 1 | 1 | | [K].TQSLLIVNnATNVVIK.[V] | N9(HexNAc(4)Hex(5)Fuc(1)NeuAc(1)) | P0DTC2 | 0 | 3 | 1 | [illegible] |
| Trypsin(semi) | High | PMI-Byonic | Unambiguous | 114 | 129 | TQSLLIVNNATNVVIK | 122 | HexNAc(4)Hex(5)Fuc(1)NeuGc(1) | 4 | 5 | 1 | | 1 | [K].TQSLLIVNnATNVVIK.[V] | N9(HexNAc(4)Hex(5)Fuc(1)NeuGc(1)) | P0DTC2 | 0 | 3 | 1 | [illegible] |
| Trypsin(semi) | High | PMI-Byonic | Unambiguous | 114 | 129 | TQSLLIVNNATNVVIK | 122 | HexNAc(4)Hex(5)Fuc(1) | 4 | 5 | 1 | | | [K].TQSLLIVNnATNVVIK.[V] | N9(HexNAc(4)Hex(5)Fuc(1)) | P0DTC2 | 0 | 3 | 1 | [illegible] |
| Trypsin(semi) | High | PMI-Byonic | Unambiguous | 114 | 129 | TQSLLIVNNATNVVIK | 122 | HexNAc(4)Hex(5)Fuc(2)NeuGc(1) | 4 | 5 | 2 | | 1 | [K].TQSLLIVNnATNVVIK.[V] | N9(HexNAc(4)Hex(5)Fuc(2)NeuGc(1)) | P0DTC2 | 0 | 3 | 1 | [illegible] |
| Trypsin(semi) | High | PMI-Byonic | Unambiguous | 114 | 129 | TQSLLIVNNATNVVIK | 122 | HexNAc(4)Hex(5)Fuc(2) | 4 | 5 | 2 | | | [K].TQSLLIVNnATNVVIK.[V] | N9(HexNAc(4)Hex(5)Fuc(2)) | P0DTC2 | 0 | 3 | 1 | [illegible] |
| Trypsin(semi) | High | PMI-Byonic | Unambiguous | 114 | 129 | TQSLLIVNNATNVVIK | 122 | HexNAc(4)Hex(5)Fuc(3) | 4 | 5 | 3 | | | [K].TQSLLIVNnATNVVIK.[V] | N9(HexNAc(4)Hex(5)Fuc(3)) | P0DTC2 | 0 | 3 | 1 | [illegible] |
| Trypsin(semi) | High | PMI-Byonic | Unambiguous | 114 | 129 | TQSLLIVNNATNVVIK | 122 | HexNAc(4)Hex(5)NeuAc(1) | 4 | 5 | | 1 | | [K].TQSLLIVNnATNVVIK.[V] | N9(HexNAc(4)Hex(5)NeuAc(1)) | P0DTC2 | 0 | 3 | 1 | [illegible] |
| Trypsin(semi) | High | PMI-Byonic | Unambiguous | 114 | 129 | TQSLLIVNNATNVVIK | 122 | HexNAc(4)Hex(5)NeuGc(1) | 4 | 5 | | | 1 | [K].TQSLLIVNnATNVVIK.[V] | N9(HexNAc(4)Hex(5)NeuGc(1)) | P0DTC2 | 0 | 3 | 1 | [illegible] |
| Trypsin(semi) | High | PMI-Byonic | Unambiguous | 114 | 129 | TQSLLIVNNATNVVIK | 122 | HexNAc(4)Hex(5) | 4 | 5 | | | | [K].TQSLLIVNnATNVVIK.[V] | N9(HexNAc(4)Hex(5)) | P0DTC2 | 0 | 3 | 1 | [illegible] |
| Trypsin(semi) | High | PMI-Byonic | Unambiguous | 114 | 129 | TQSLLIVNNATNVVIK | 122 | HexNAc(4)Hex(6)Fuc(2) | 4 | 6 | 2 | | | [K].TQSLLIVNnATNVVIK.[V] | N9(HexNAc(4)Hex(6)Fuc(2)) | P0DTC2 | 0 | 3 | 1 | [illegible] |
| Trypsin(semi) | High | PMI-Byonic | Unambiguous | 114 | 129 | TQSLLIVNNATNVVIK | 122 | HexNAc(4)Hex(6)NeuAc(1) | 4 | 6 | | 1 | | [K].TQSLLIVNnATNVVIK.[V] | N9(HexNAc(4)Hex(6)NeuAc(1)) | P0DTC2 | 0 | 3 | 1 | [illegible] |
| Trypsin(semi) | High | PMI-Byonic | Unambiguous | 114 | 129 | TQSLLIVNNATNVVIK | 122 | HexNAc(4)Hex(6) | 4 | 6 | | | | [K].TQSLLIVNnATNVVIK.[V] | N9(HexNAc(4)Hex(6)) | P0DTC2 | 0 | 3 | 1 | [illegible] |
| Trypsin(semi) | High | PMI-Byonic | Unambiguous | 114 | 129 | TQSLLIVNNATNVVIK | 122 | HexNAc(4)Hex(7)Fuc(1) | 4 | 7 | 1 | | | [K].TQSLLIVNnATNVVIK.[V] | N9(HexNAc(4)Hex(7)Fuc(1)) | P0DTC2 | 0 | 3 | 1 | [illegible] |
| Trypsin(semi) | High | PMI-Byonic | Unambiguous | 114 | 129 | TQSLLIVNNATNVVIK | 122 | HexNAc(5)Hex(3)Fuc(1) | 5 | 3 | 1 | | | [K].TQSLLIVNnATNVVIK.[V] | N9(HexNAc(5)Hex(3)Fuc(1)) | P0DTC2 | 0 | 3 | 1 | [illegible] |
| Trypsin(semi) | High | PMI-Byonic | Unambiguous | 114 | 129 | TQSLLIVNNATNVVIK | 122 | HexNAc(5)Hex(4)Fuc(1) | 5 | 4 | 1 | | | [K].TQSLLIVNnATNVVIK.[V] | N9(HexNAc(5)Hex(4)Fuc(1)) | P0DTC2 | 0 | 3 | 1 | [illegible] |
| Trypsin(semi) | High | PMI-Byonic | Unambiguous | 114 | 129 | TQSLLIVNNATNVVIK | 122 | HexNAc(5)Hex(4)Fuc(2) | 5 | 4 | 2 | | | [K].TQSLLIVNnATNVVIK.[V] | N9(HexNAc(5)Hex(4)Fuc(2)) | P0DTC2 | 0 | 3 | 1 | [illegible] |
| Trypsin(semi) | High | PMI-Byonic | Unambiguous | 114 | 129 | TQSLLIVNNATNVVIK | 122 | HexNAc(5)Hex(4) | 5 | 4 | | | | [K].TQSLLIVNnATNVVIK.[V] | N9(HexNAc(5)Hex(4)) | P0DTC2 | 0 | 3 | 1 | [illegible] |
| Trypsin(semi) | High | PMI-Byonic | Unambiguous | 114 | 129 | TQSLLIVNNATNVVIK | 122 | HexNAc(5)Hex(5)Fuc(1) | 5 | 5 | 1 | | | [K].TQSLLIVNnATNVVIK.[V] | N9(HexNAc(5)Hex(5)Fuc(1)) | P0DTC2 | 0 | 3 | 1 | [illegible] |
| Trypsin(semi) | High | PMI-Byonic | Unambiguous | 114 | 129 | TQSLLIVNNATNVVIK | 122 | HexNAc(5)Hex(5)Fuc(2) | 5 | 5 | 2 | | | [K].TQSLLIVNnATNVVIK.[V] | N9(HexNAc(5)Hex(5)Fuc(2)) | P0DTC2 | 0 | 3 | 1 | [illegible] |
| Trypsin(semi) | High | PMI-Byonic | Unambiguous | 117 | 129 | LLIVNNATNVVIK | 122 | HexNAc(5)Hex(5)Fuc(3) | 5 | 5 | 3 | | | [S].LLIVNnATNVVIK.[V] | N6(HexNAc(5)Hex(5)Fuc(3)) | P0DTC2 | 0 | 3 | 1 | [illegible] |
| Trypsin(semi) | High | PMI-Byonic | Unambiguous | 114 | 129 | TQSLLIVNNATNVVIK | 122 | HexNAc(5)Hex(5) | 5 | 5 | | | | [K].TQSLLIVNnATNVVIK.[V] | N9(HexNAc(5)Hex(5)) | P0DTC2 | 0 | 3 | 1 | [illegible] |
| Trypsin(semi) | High | PMI-Byonic | Unambiguous | 114 | 129 | TQSLLIVNNATNVVIK | 122 | HexNAc(5)Hex(6)Fuc(1) | 5 | 6 | 1 | | | [K].TQSLLIVNnATNVVIK.[V] | N9(HexNAc(5)Hex(6)Fuc(1)) | P0DTC2 | 0 | 3 | 1 | [illegible] |
| Trypsin(semi) | High | PMI-Byonic | Unambiguous | 114 | 129 | TQSLLIVNNATNVVIK | 122 | HexNAc(5)Hex(6)Fuc(3) | 5 | 6 | 3 | | | [K].TQSLLIVNnATNVVIK.[V] | N9(HexNAc(5)Hex(6)Fuc(3)) | P0DTC2 | 0 | 3 | 1 | [illegible] |
| Trypsin(semi) | High | PMI-Byonic | Unambiguous | 114 | 129 | TQSLLIVNNATNVVIK | 122 | HexNAc(5)Hex(6)NeuAc(1) | 5 | 6 | | 1 | | [K].TQSLLIVNnATNVVIK.[V] | N9(HexNAc(5)Hex(6)NeuAc(1)) | P0DTC2 | 0 | 3 | 1 | [illegible] |
| Trypsin(semi) | High | PMI-Byonic | Unambiguous | 114 | 129 | TQSLLIVNNATNVVIK | 122 | HexNAc(5)Hex(6)NeuGc(1) | 5 | 6 | | | 1 | [K].TQSLLIVNnATNVVIK.[V] | N9(HexNAc(5)Hex(6)NeuGc(1)) | P0DTC2 | 0 | 3 | 1 | [illegible] |
| Trypsin(semi) | High | PMI-Byonic | Unambiguous | 114 | 129 | TQSLLIVNNATNVVIK | 122 | HexNAc(5)Hex(6) | 5 | 6 | | | | [K].TQSLLIVNnATNVVIK.[V] | N9(HexNAc(5)Hex(6)) | P0DTC2 | 0 | 3 | 1 | [illegible] |
| Trypsin(semi) | High | PMI-Byonic | Unambiguous | 117 | 129 | LLIVNNATNVVIK | 122 | HexNAc(6)Hex(5) | 6 | 5 | | | | [S].LLIVNnATNVVIK.[V] | N6(HexNAc(6)Hex(5)) | P0DTC2 | 0 | 3 | 1 | [illegible] |
| Trypsin(semi) | High | PMI-Byonic | Unambiguous | 114 | 129 | TQSLLIVNNATNVVIK | 122 | HexNAc(6)Hex(6)Fuc(1)NeuAc(1) | 6 | 6 | 1 | 1 | | [K].TQSLLIVNnATNVVIK.[V] | N9(HexNAc(6)Hex(6)Fuc(1)NeuAc(1)) | P0DTC2 | 0 | 3 | 1 | [illegible] |
| Trypsin(semi) | High | PMI-Byonic | Unambiguous | 114 | 129 | TQSLLIVNNATNVVIK | 122 | HexNAc(6)Hex(6)Fuc(1) | 6 | 6 | 1 | | | [K].TQSLLIVNnATNVVIK.[V] | N9(HexNAc(6)Hex(6)Fuc(1)) | P0DTC2 | 0 | 3 | 1 | [illegible] |
| Trypsin(semi) | High | PMI-Byonic | Unambiguous | 114 | 129 | TQSLLIVNNATNVVIK | 122 | HexNAc(6)Hex(6)Fuc(2) | 6 | 6 | 2 | | | [K].TQSLLIVNnATNVVIK.[V] | N9(HexNAc(6)Hex(6)Fuc(2)) | P0DTC2 | 0 | 3 | 1 | [illegible] |
| Trypsin(semi) | High | PMI-Byonic | Unambiguous | 114 | 129 | TQSLLIVNNATNVVIK | 122 | HexNAc(6)Hex(6)NeuAc(1)NeuGc(1) | 6 | 6 | | 1 | 1 | [K].TQSLLIVNnATNVVIK.[V] | N9(HexNAc(6)Hex(6)NeuAc(1)NeuGc(1)) | P0DTC2 | 0 | 3 | 1 | [illegible] |
| Trypsin(semi) | High | PMI-Byonic | Unambiguous | 114 | 129 | TQSLLIVNNATNVVIK | 122 | HexNAc(6)Hex(6) | 6 | 6 | | | | [K].TQSLLIVNnATNVVIK.[V] | N9(HexNAc(6)Hex(6)) | P0DTC2 | 0 | 3 | 1 | [illegible] |
| Trypsin(semi) | High | PMI-Byonic | Unambiguous | 114 | 129 | TQSLLIVNNATNVVIK | 122 | HexNAc(6)Hex(7)Fuc(1)NeuAc(1) | 6 | 7 | 1 | 1 | | [K].TQSLLIVNnATNVVIK.[V] | N9(HexNAc(6)Hex(7)Fuc(1)NeuAc(1)) | P0DTC2 | 0 | 3 | 1 | [illegible] |
| Trypsin(semi) | High | PMI-Byonic | Unambiguous | 114 | 129 | TQSLLIVNNATNVVIK | 122 | HexNAc(6)Hex(7)Fuc(1) | 6 | 7 | 1 | | | [K].TQSLLIVNnATNVVIK.[V] | N9(HexNAc(6)Hex(7)Fuc(1)) | P0DTC2 | 0 | 3 | 1 | [illegible] |
| Trypsin(semi) | High | PMI-Byonic | Unambiguous | 114 | 129 | TQSLLIVNNATNVVIK | 122 | HexNAc(6)Hex(7)NeuAc(1) | 6 | 7 | | 1 | | [K].TQSLLIVNnATNVVIK.[V] | N9(HexNAc(6)Hex(7)NeuAc(1)) | P0DTC2 | 0 | 3 | 1 | [illegible] |
| Trypsin(semi) | High | PMI-Byonic | Unambiguous | 114 | 129 | TQSLLIVNNATNVVIK | 122 | HexNAc(7)Hex(7)Fuc(1) | 7 | 7 | 1 | | | [K].TQSLLIVNnATNVVIK.[V] | N9(HexNAc(7)Hex(7)Fuc(1)) | P0DTC2 | 0 | 3 | 1 | [illegible] |
| Trypsin(semi) | High | PMI-Byonic | Unambiguous | 159 | 182 | VYSSANNCTFEYVSQPFLMDLEGK | 165 | HexNAc(2)Hex(5) | 2 | 5 | | | | [R].VYSSANncTFEYVSQPFLMDLEGK.[Q] | N7(HexNAc(2)Hex(5)); C8(Carbamidomethyl) | P0DTC2 | 0 | 3 | 1 | [illegible] |
| Trypsin(semi) | High | PMI-Byonic | Unambiguous | 159 | 182 | VYSSANNCTFEYVSQPFLMDLEGK | 165 | HexNAc(2)Hex(6) | 2 | 6 | | | | [R].VYSSANncTFEYVSQPFLMDLEGK.[Q] | N7(HexNAc(2)Hex(6)); C8(Carbamidomethyl) | P0DTC2 | 0 | 3 | 1 | [illegible] |
| Trypsin(semi) | High | PMI-Byonic | Unambiguous | 159 | 182 | VYSSANNCTFEYVSQPFLMDLEGK | 165 | HexNAc(3)Hex(4)Fuc(1) | 3 | 4 | 1 | | | [R].VYSSANncTFEYVSQPFLMDLEGK.[Q] | N7(HexNAc(3)Hex(4)Fuc(1)); C8(Carbamidomethyl) | P0DTC2 | 0 | 3 | 1 | [illegible] |
| Trypsin(semi) | High | PMI-Byonic | Unambiguous | 159 | 182 | VYSSANNCTFEYVSQPFLMDLEGK | 165 | HexNAc(3)Hex(5)Fuc(1) | 3 | 5 | 1 | | | [R].VYSSANncTFEYVSQPFLMDLEGK.[Q] | N7(HexNAc(3)Hex(5)Fuc(1)); C8(Carbamidomethyl) | P0DTC2 | 0 | 3 | 1 | [illegible] |
| Trypsin(semi) | High | PMI-Byonic | Unambiguous | 159 | 182 | VYSSANNCTFEYVSQPFLMDLEGK | 165 | HexNAc(3)Hex(6)Fuc(1) | 3 | 6 | 1 | | | [R].VYSSANncTFEYVSQPFLMDLEGK.[Q] | N7(HexNAc(3)Hex(6)Fuc(1)); C8(Carbamidomethyl) | P0DTC2 | 0 | 3 | 1 | [illegible] |
| Trypsin(semi) | High | PMI-Byonic | Unambiguous | 159 | 182 | VYSSANNCTFEYVSQPFLMDLEGK | 165 | HexNAc(3)Hex(6)NeuGc(1) | 3 | 6 | | | 1 | [R].VYSSANncTFEYVSQPFLMDLEGK.[Q] | N7(HexNAc(3)Hex(6)NeuGc(1)); C8(Carbamidomethyl) | P0DTC2 | 0 | 3 | 1 | [illegible] |
| Trypsin(semi) | High | PMI-Byonic | Unambiguous | 159 | 182 | VYSSANNCTFEYVSQPFLMDLEGK | 165 | HexNAc(4)Hex(5)Fuc(1)NeuAc(1) | 4 | 5 | 1 | 1 | | [R].VYSSANncTFEYVSQPFLMDLEGK.[Q] | N7(HexNAc(4)Hex(5)Fuc(1)NeuAc(1)); C8(Carbamidomethyl) | P0DTC2 | 0 | 3 | 1 | [illegible] |
| Trypsin(semi) | High | PMI-Byonic | Unambiguous | 159 | 182 | VYSSANNCTFEYVSQPFLMDLEGK | 165 | HexNAc(4)Hex(5)Fuc(1)NeuGc(1) | 4 | 5 | 1 | | 1 | [R].VYSSANncTFEYVSQPFLMDLEGK.[Q] | N7(HexNAc(4)Hex(5)Fuc(1)NeuGc(1)); C8(Carbamidomethyl) | P0DTC2 | 0 | 3 | 1 | [illegible] |
| Trypsin(semi) | High | PMI-Byonic | Unambiguous | 159 | 182 | VYSSANNCTFEYVSQPFLMDLEGK | 165 | HexNAc(4)Hex(5)Fuc(1) | 4 | 5 | 1 | | | [R].VYSSANncTFEYVSQPFLMDLEGK.[Q] | N7(HexNAc(4)Hex(5)Fuc(1)); C8(Carbamidomethyl) | P0DTC2 | 0 | 3 | 1 | [illegible] |
| Trypsin(semi) | High | PMI-Byonic | Unambiguous | 159 | 182 | VYSSANNCTFEYVSQPFLMDLEGK | 165 | HexNAc(4)Hex(5)Fuc(3) | 4 | 5 | 3 | | | [R].VYSSANncTFEYVSQPFLMDLEGK.[Q] | N7(HexNAc(4)Hex(5)Fuc(3)); C8(Carbamidomethyl) | P0DTC2 | 0 | 4 | 1 | [illegible] |
| Trypsin(semi) | High | PMI-Byonic | Unambiguous | 159 | 182 | VYSSANNCTFEYVSQPFLMDLEGK | 165 | HexNAc(4)Hex(5)NeuAc(1) | 4 | 5 | | 1 | | [R].VYSSANncTFEYVSQPFLMDLEGK.[Q] | N7(HexNAc(4)Hex(5)NeuAc(1)); C8(Carbamidomethyl) | P0DTC2 | 0 | 3 | 1 | [illegible] |
| Trypsin(semi) | High | PMI-Byonic | Unambiguous | 159 | 182 | VYSSANNCTFEYVSQPFLMDLEGK | 165 | HexNAc(4)Hex(5) | 4 | 5 | | | | [R].VYSSANncTFEYVSQPFLMDLEGK.[Q] | N7(HexNAc(4)Hex(5)); C8(Carbamidomethyl) | P0DTC2 | 0 | 3 | 1 | [illegible] |
| Trypsin(semi) | High | PMI-Byonic | Unambiguous | 159 | 182 | VYSSANNCTFEYVSQPFLMDLEGK | 165 | HexNAc(4)Hex(6)Fuc(1)NeuAc(1) | 4 | 6 | 1 | 1 | | [R].VYSSANncTFEYVSQPFLMDLEGK.[Q] | N7(HexNAc(4)Hex(6)Fuc(1)NeuAc(1)); C8(Carbamidomethyl) | P0DTC2 | 0 | 3 | 1 | [illegible] |
| Trypsin(semi) | High | PMI-Byonic | Unambiguous | 159 | 182 | VYSSANNCTFEYVSQPFLMDLEGK | 165 | HexNAc(4)Hex(6)NeuAc(1) | 4 | 6 | | 1 | | [R].VYSSANncTFEYVSQPFLMDLEGK.[Q] | N7(HexNAc(4)Hex(6)NeuAc(1)); C8(Carbamidomethyl) | P0DTC2 | 0 | 4 | 1 | [illegible] |
| Trypsin(semi) | High | PMI-Byonic | Unambiguous | 159 | 182 | VYSSANNCTFEYVSQPFLMDLEGK | 165 | HexNAc(5)Hex(5)Fuc(1)NeuAc(1) | 5 | 5 | 1 | 1 | | [R].VYSSANncTFEYVSQPFLMDLEGK.[Q] | N7(HexNAc(5)Hex(5)Fuc(1)NeuAc(1)); C8(Carbamidomethyl) | P0DTC2 | 0 | 3 | 1 | [illegible] |
| Trypsin(semi) | High | PMI-Byonic | Unambiguous | 159 | 182 | VYSSANNCTFEYVSQPFLMDLEGK | 165 | HexNAc(5)Hex(5)Fuc(1) | 5 | 5 | 1 | | | [R].VYSSANncTFEYVSQPFLMDLEGK.[Q] | N7(HexNAc(5)Hex(5)Fuc(1)); C8(Carbamidomethyl) | P0DTC2 | 0 | 3 | 1 | [illegible] |
| Trypsin(semi) | High | PMI-Byonic | Unambiguous | 159 | 182 | VYSSANNCTFEYVSQPFLMDLEGK | 165 | HexNAc(5)Hex(6)Fuc(2) | 5 | 6 | 2 | | | [R].VYSSANncTFEYVSQPFLMDLEGK.[Q] | N7(HexNAc(5)Hex(6)Fuc(2)); C8(Carbamidomethyl) | P0DTC2 | 0 | 3 | 1 | [illegible] |
| Trypsin(semi) | High | PMI-Byonic | Unambiguous | 159 | 182 | VYSSANNCTFEYVSQPFLMDLEGK | 165 | HexNAc(5)Hex(6)Fuc(3) | 5 | 6 | 3 | | | [R].VYSSANncTFEYVSQPFLMDLEGK.[Q] | N7(HexNAc(5)Hex(6)Fuc(3)); C8(Carbamidomethyl) | P0DTC2 | 0 | 4 | 1 | [illegible] |
| Trypsin(semi) | High | PMI-Byonic | Unambiguous | 159 | 182 | VYSSANNCTFEYVSQPFLMDLEGK | 165 | HexNAc(5)Hex(6) | 5 | 6 | | | | [R].VYSSANncTFEYVSQPFLMDLEGK.[Q] | N7(HexNAc(5)Hex(6)); C8(Carbamidomethyl) | P0DTC2 | 0 | 3 | 1 | [illegible] |
| Trypsin(semi) | High | PMI-Byonic | Unambiguous | 163 | 182 | ANNCTFEYVSQPFLMDLEGK | 165 | HexNAc(6)Hex(3)Fuc(1)NeuAc(1) | 6 | 3 | 1 | 1 | | [S].ANncTFEYVSQPFLMDLEGK.[Q] | N3(HexNAc(6)Hex(3)Fuc(1)NeuAc(1)); C4(Carbamidomethyl) | P0DTC2 | 0 | 3 | 1 | [illegible] |
| Trypsin(semi) | High | PMI-Byonic | Unambiguous | 159 | 182 | VYSSANNCTFEYVSQPFLMDLEGK | 165 | HexNAc(6)Hex(6)Fuc(1) | 6 | 6 | 1 | | | [R].VYSSANncTFEYVSQPFLMDLEGK.[Q] | N7(HexNAc(6)Hex(6)Fuc(1)); C8(Carbamidomethyl) | P0DTC2 | 0 | 4 | 1 | [illegible] |
| Trypsin(semi) | High | PMI-Byonic | Unambiguous | 159 | 182 | VYSSANNCTFEYVSQPFLMDLEGK | 165 | HexNAc(6)Hex(7)Fuc(1)NeuAc(1) | 6 | 7 | 1 | 1 | | [R].VYSSANncTFEYVSQPFLMDLEGK.[Q] | N7(HexNAc(6)Hex(7)Fuc(1)NeuAc(1)); C8(Carbamidomethyl) | P0DTC2 | 0 | 4 | 1 | [illegible] |
| Trypsin(semi) | High | PMI-Byonic | Unambiguous | 163 | 182 | ANNCTFEYVSQPFLMDLEGK | 165 | HexNAc(6)Hex(7)Fuc(1)NeuAc(2) | 6 | 7 | 1 | 2 | | [S].ANncTFEYVSQPFLMDLEGK.[Q] | N3(HexNAc(6)Hex(7)Fuc(1)NeuAc(2)); C4(Carbamidomethyl) | P0DTC2 | 0 | 3 | 1 | [illegible] |
| Trypsin(semi) | High | PMI-Byonic | Unambiguous | 159 | 182 | VYSSANNCTFEYVSQPFLMDLEGK | 165 | HexNAc(6)Hex(7)Fuc(1) | 6 | 7 | 1 | | | [R].VYSSANncTFEYVSQPFLMDLEGK.[Q] | N7(HexNAc(6)Hex(7)Fuc(1)); C8(Carbamidomethyl) | P0DTC2 | 0 | 3 | 1 | [illegible] |
| Trypsin(semi) | High | PMI-Byonic | Unambiguous | 159 | 182 | VYSSANNCTFEYVSQPFLMDLEGK | 165 | HexNAc(6)Hex(7)Fuc(4) | 6 | 7 | 4 | | | [R].VYSSANncTFEYVSQPFLMDLEGK.[Q] | N7(HexNAc(6)Hex(7)Fuc(4)); C8(Carbamidomethyl) | P0DTC2 | 0 | 4 | 1 | [illegible] |
| Trypsin(semi) | High | PMI-Byonic | Unambiguous | 163 | 182 | ANNCTFEYVSQPFLMDLEGK | 165 | HexNAc(6)Hex(7)Fuc(5) | 6 | 7 | 5 | | | [S].ANncTFEYVSQPFLMDLEGK.[Q] | N3(HexNAc(6)Hex(7)Fuc(5)); C4(Carbamidomethyl) | P0DTC2 | 0 | 3 | 1 | [illegible] |
| Trypsin(semi) | High | PMI-Byonic | Unambiguous | 159 | 182 | VYSSANNCTFEYVSQPFLMDLEGK | 165 | HexNAc(8)Hex(9) | 8 | 9 | | | | [R].VYSSANncTFEYVSQPFLMDLEGK.[Q] | N7(HexNAc(8)Hex(9)); C8(Carbamidomethyl) | P0DTC2 | 0 | 4 | 1 | [illegible] |
| Trypsin(semi) | High | PMI-Byonic | Unambiguous | 215 | 237 | DLPQGFSALEPLVDLPIGINITR | 234 | HexNAc(2)Hex(5) | 2 | 5 | | | | [R].DLPQGFSALEPLVDLPIGInITR.[F] | N20(HexNAc(2)Hex(5)) | P0DTC2 | 0 | 3 | 1 | [illegible] |
| Trypsin(semi) | High | PMI-Byonic | Unambiguous | 215 | 237 | DLPQGFSALEPLVDLPIGINITR | 234 | HexNAc(2)Hex(6) | 2 | 6 | | | | [R].DLPQGFSALEPLVDLPIGInITR.[F] | N20(HexNAc(2)Hex(6)) | P0DTC2 | 0 | 3 | 1 | [illegible] |
| Trypsin(semi) | High | PMI-Byonic | Unambiguous | 215 | 237 | DLPQGFSALEPLVDLPIGINITR | 234 | HexNAc(2)Hex(7) | 2 | 7 | | | | [R].DLPQGFSALEPLVDLPIGInITR.[F] | N20(HexNAc(2)Hex(7)) | P0DTC2 | 0 | 3 | 1 | [illegible] |
| Trypsin(semi) | High | PMI-Byonic | Unambiguous | 215 | 237 | DLPQGFSALEPLVDLPIGINITR | 234 | HexNAc(2)Hex(8) | 2 | 8 | | | | [R].DLPQGFSALEPLVDLPIGInITR.[F] | N20(HexNAc(2)Hex(8)) | P0DTC2 | 0 | 3 | 1 | [illegible] |
| Trypsin(semi) | High | PMI-Byonic | Unambiguous | 215 | 237 | DLPQGFSALEPLVDLPIGINITR | 234 | HexNAc(2)Hex(9) | 2 | 9 | | | | [R].DLPQGFSALEPLVDLPIGInITR.[F] | N20(HexNAc(2)Hex(9)) | P0DTC2 | 0 | 3 | 1 | [illegible] |
| Trypsin(semi) | High | PMI-Byonic | Unambiguous | 214 | 237 | RDLPQGFSALEPLVDLPIGINITR | 234 | HexNAc(3)Hex(4)NeuGc(1) | 3 | 4 | | | 1 | [V].RDLPQGFSALEPLVDLPIGInITR.[F] | N21(HexNAc(3)Hex(4)NeuGc(1)) | P0DTC2 | 1 | 3 | 1 | [illegible] |
| Trypsin(semi) | High | PMI-Byonic | Unambiguous | 215 | 237 | DLPQGFSALEPLVDLPIGINITR | 234 | HexNAc(3)Hex(5)Fuc(1) | 3 | 5 | 1 | | | [R].DLPQGFSALEPLVDLPIGInITR.[F] | N20(HexNAc(3)Hex(5)Fuc(1)) | P0DTC2 | 0 | 3 | 1 | [illegible] |
| Trypsin(semi) | High | PMI-Byonic | Unambiguous | 215 | 237 | DLPQGFSALEPLVDLPIGINITR | 234 | HexNAc(3)Hex(5) | 3 | 5 | | | | [R].DLPQGFSALEPLVDLPIGInITR.[F] | N20(HexNAc(3)Hex(5)) | P0DTC2 | 0 | 3 | 1 | [illegible] |
| Trypsin(semi) | High | PMI-Byonic | Unambiguous | 215 | 237 | DLPQGFSALEPLVDLPIGINITR | 234 | HexNAc(3)Hex(6) | 3 | 6 | | | | [R].DLPQGFSALEPLVDLPIGInITR.[F] | N20(HexNAc(3)Hex(6)) | P0DTC2 | 0 | 3 | 1 | [illegible] |
| Trypsin(semi) | High | PMI-Byonic | Unambiguous | 215 | 237 | DLPQGFSALEPLVDLPIGINITR | 234 | HexNAc(4)Hex(3)Fuc(1) | 4 | 3 | 1 | | | [R].DLPQGFSALEPLVDLPIGInITR.[F] | N20(HexNAc(4)Hex(3)Fuc(1)) | P0DTC2 | 0 | 3 | 1 | [illegible] |
| Trypsin(semi) | High | PMI-Byonic | Unambiguous | 215 | 237 | DLPQGFSALEPLVDLPIGINITR | 234 | HexNAc(4)Hex(4) | 4 | 4 | | | | [R].DLPQGFSALEPLVDLPIGInITR.[F] | N20(HexNAc(4)Hex(4)) | P0DTC2 | 0 | 3 | 1 | [illegible] |
| Trypsin(semi) | High | PMI-Byonic | Unambiguous | 215 | 237 | DLPQGFSALEPLVDLPIGINITR | 234 | HexNAc(5)Hex(3)Fuc(1) | 5 | 3 | 1 | | | [R].DLPQGFSALEPLVDLPIGInITR.[F] | N20(HexNAc(5)Hex(3)Fuc(1)) | P0DTC2 | 0 | 3 | 1 | [illegible] |
| Trypsin(semi) | High | PMI-Byonic | Unambiguous | 215 | 237 | DLPQGFSALEPLVDLPIGINITR | 234 | HexNAc(5)Hex(4)Fuc(1) | 5 | 4 | 1 | | | [R].DLPQGFSALEPLVDLPIGInITR.[F] | N20(HexNAc(5)Hex(4)Fuc(1)) | P0DTC2 | 0 | 3 | 1 | [illegible] |
| Trypsin(semi) | High | PMI-Byonic | Unambiguous | 217 | 237 | PQGFSALEPLVDLPIGINITR | 234 | HexNAc(6)Hex(3)Fuc(1) | 6 | 3 | 1 | | | [L].PQGFSALEPLVDLPIGInITR.[F] | N18(HexNAc(6)Hex(3)Fuc(1)) | P0DTC2 | 0 | 3 | 1 | [illegible] |
| Trypsin(semi) | High | PMI-Byonic | Unambiguous | 217 | 237 | PQGFSALEPLVDLPIGINITR | 234 | HexNAc(6)Hex(3)Fuc(2) | 6 | 3 | 2 | | | [L].PQGFSALEPLVDLPIGInITR.[F] | N18(HexNAc(6)Hex(3)Fuc(2)) | P0DTC2 | 0 | 3 | 1 | [illegible] |
| Trypsin(semi) | High | PMI-Byonic | Unambiguous | 215 | 237 | DLPQGFSALEPLVDLPIGINITR | 234 | HexNAc(6)Hex(7) | 6 | 7 | | | | [R].DLPQGFSALEPLVDLPIGInITR.[F] | N20(HexNAc(6)Hex(7)) | P0DTC2 | 0 | 4 | 1 | [illegible] |
| Trypsin(semi) | High | PMI-Byonic | Unambiguous | 225 | 237 | PLVDLPIGINITR | 234 | HexNAc(8)Hex(6) | 8 | 6 | | | | [E].PLVDLPIGInITR.[F] | N10(HexNAc(8)Hex(6)) | P0DTC2 | 0 | 3 | 1 | [illegible] |
| Trypsin(semi) | High | PMI-Byonic | Unambiguous | 279 | 300 | YNENGTITDAVDCALDPLSETK | 282 | HexNAc(2)Hex(5) | 2 | 5 | | | | [K].YNEnGTITDAVDcALDPLSETK.[C] | N4(HexNAc(2)Hex(5)); C13(Carbamidomethyl) | P0DTC2 | 0 | 3 | 1 | [illegible] |
| Trypsin(semi) | High | PMI-Byonic | Unambiguous | 279 | 300 | YNENGTITDAVDCALDPLSETK | 282 | HexNAc(2)Hex(6) | 2 | 6 | | | | [K].YNEnGTITDAVDcALDPLSETK.[C] | N4(HexNAc(2)Hex(6)); C13(Carbamidomethyl) | P0DTC2 | 0 | 3 | 1 | [illegible] |
| Trypsin(semi) | High | PMI-Byonic | Unambiguous | 279 | 300 | YNENGTITDAVDCALDPLSETK | 282 | HexNAc(3)Hex(3) | 3 | 3 | | | | [K].YNEnGTITDAVDcALDPLSETK.[C] | N4(HexNAc(3)Hex(3)); C13(Carbamidomethyl) | P0DTC2 | 0 | 3 | 1 | [illegible] |
| Trypsin(semi) | High | PMI-Byonic | Unambiguous | 279 | 300 | YNENGTITDAVDCALDPLSETK | 282 | HexNAc(3)Hex(5)Fuc(1) | 3 | 5 | 1 | | | [K].YNEnGTITDAVDcALDPLSETK.[C] | N4(HexNAc(3)Hex(5)Fuc(1)); C13(Carbamidomethyl) | P0DTC2 | 0 | 3 | 1 | [illegible] |
| Trypsin(semi) | High | PMI-Byonic | Unambiguous | 279 | 300 | YNENGTITDAVDCALDPLSETK | 282 | HexNAc(3)Hex(5)NeuGc(1) | 3 | 5 | | | 1 | [K].YNEnGTITDAVDcALDPLSETK.[C] | N4(HexNAc(3)Hex(5)NeuGc(1)); C13(Carbamidomethyl) | P0DTC2 | 0 | 3 | 1 | [illegible] |
| Trypsin(semi) | High | PMI-Byonic | Unambiguous | 279 | 300 | YNENGTITDAVDCALDPLSETK | 282 | HexNAc(3)Hex(5) | 3 | 5 | | | | [K].YNEnGTITDAVDcALDPLSETK.[C] | N4(HexNAc(3)Hex(5)); C13(Carbamidomethyl) | P0DTC2 | 0 | 3 | 1 | [illegible] |
| Trypsin(semi) | High | PMI-Byonic | Unambiguous | 279 | 300 | YNENGTITDAVDCALDPLSETK | 282 | HexNAc(4)Hex(3)Fuc(1) | 4 | 3 | 1 | | | [K].YNEnGTITDAVDcALDPLSETK.[C] | N4(HexNAc(4)Hex(3)Fuc(1)); C13(Carbamidomethyl) | P0DTC2 | 0 | 3 | 1 | [illegible] |
| Trypsin(semi) | High | PMI-Byonic | Unambiguous | 279 | 300 | YNENGTITDAVDCALDPLSETK | 282 | HexNAc(4)Hex(5)Fuc(1)NeuAc(1) | 4 | 5 | 1 | 1 | | [K].YNEnGTITDAVDcALDPLSETK.[C] | N4(HexNAc(4)Hex(5)Fuc(1)NeuAc(1)); C13(Carbamidomethyl) | P0DTC2 | 0 | 3 | 1 | [illegible] |
| Trypsin(semi) | High | PMI-Byonic | Unambiguous | 279 | 300 | YNENGTITDAVDCALDPLSETK | 282 | HexNAc(4)Hex(5)Fuc(1) | 4 | 5 | 1 | | | [K].YNEnGTITDAVDcALDPLSETK.[C] | N4(HexNAc(4)Hex(5)Fuc(1)); C13(Carbamidomethyl) | P0DTC2 | 0 | 3 | 1 | [illegible] |
| Trypsin(semi) | High | PMI-Byonic | Unambiguous | 279 | 300 | YNENGTITDAVDCALDPLSETK | 282 | HexNAc(4)Hex(5)Fuc(2) | 4 | 5 | 2 | | | [K].YNEnGTITDAVDcALDPLSETK.[C] | N4(HexNAc(4)Hex(5)Fuc(2)); C13(Carbamidomethyl) | P0DTC2 | 0 | 3 | 1 | [illegible] |
| Trypsin(semi) | High | PMI-Byonic | Unambiguous | 279 | 300 | YNENGTITDAVDCALDPLSETK | 282 | HexNAc(4)Hex(5)Fuc(3)NeuAc(1) | 4 | 5 | 3 | 1 | | [K].YNEnGTITDAVDcALDPLSETK.[C] | N4(HexNAc(4)Hex(5)Fuc(3)NeuAc(1)); C13(Carbamidomethyl) | P0DTC2 | 0 | 3 | 1 | [illegible] |
| Trypsin(semi) | High | PMI-Byonic | Unambiguous | 279 | 300 | YNENGTITDAVDCALDPLSETK | 282 | HexNAc(4)Hex(5) | 4 | 5 | | | | [K].YNEnGTITDAVDcALDPLSETK.[C] | N4(HexNAc(4)Hex(5)); C13(Carbamidomethyl) | P0DTC2 | 0 | 3 | 1 | [illegible] |
| Trypsin(semi) | High | PMI-Byonic | Unambiguous | 279 | 300 | YNENGTITDAVDCALDPLSETK | 282 | HexNAc(4)Hex(6)NeuAc(1) | 4 | 6 | | 1 | | [K].YNEnGTITDAVDcALDPLSETK.[C] | N4(HexNAc(4)Hex(6)NeuAc(1)); C13(Carbamidomethyl) | P0DTC2 | 0 | 3 | 1 | [illegible] |
| Trypsin(semi) | High | PMI-Byonic | Unambiguous | 279 | 300 | YNENGTITDAVDCALDPLSETK | 282 | HexNAc(4)Hex(6) | 4 | 6 | | | | [K].YNEnGTITDAVDcALDPLSETK.[C] | N4(HexNAc(4)Hex(6)); C13(Carbamidomethyl) | P0DTC2 | 0 | 3 | 1 | [illegible] |
| Trypsin(semi) | High | PMI-Byonic | Unambiguous | 279 | 300 | YNENGTITDAVDCALDPLSETK | 282 | HexNAc(5)Hex(5)Fuc(1)NeuAc(1) | 5 | 5 | 1 | 1 | | [K].YNEnGTITDAVDcALDPLSETK.[C] | N4(HexNAc(5)Hex(5)Fuc(1)NeuAc(1)); C13(Carbamidomethyl) | P0DTC2 | 0 | 3 | 1 | [illegible] |
| Trypsin(semi) | High | PMI-Byonic | Unambiguous | 279 | 300 | YNENGTITDAVDCALDPLSETK | 282 | HexNAc(5)Hex(5)Fuc(1) | 5 | 5 | 1 | | | [K].YNEnGTITDAVDcALDPLSETK.[C] | N4(HexNAc(5)Hex(5)Fuc(1)); C13(Carbamidomethyl) | P0DTC2 | 0 | 3 | 1 | [illegible] |
| Trypsin(semi) | High | PMI-Byonic | Unambiguous | 279 | 300 | YNENGTITDAVDCALDPLSETK | 282 | HexNAc(5)Hex(5)NeuAc(1) | 5 | 5 | | 1 | | [K].YNEnGTITDAVDcALDPLSETK.[C] | N4(HexNAc(5)Hex(5)NeuAc(1)); C13(Carbamidomethyl) | P0DTC2 | 0 | 3 | 1 | [illegible] |
| Trypsin(semi) | High | PMI-Byonic | Unambiguous | 279 | 300 | YNENGTITDAVDCALDPLSETK | 282 | HexNAc(5)Hex(5) | 5 | 5 | | | | [K].YNEnGTITDAVDcALDPLSETK.[C] | N4(HexNAc(5)Hex(5)); C13(Carbamidomethyl) | P0DTC2 | 0 | 3 | 1 | [illegible] |
| Trypsin(semi) | High | PMI-Byonic | Unambiguous | 279 | 300 | YNENGTITDAVDCALDPLSETK | 282 | HexNAc(5)Hex(6)Fuc(1)NeuAc(1) | 5 | 6 | 1 | 1 | | [K].YNEnGTITDAVDcALDPLSETK.[C] | N4(HexNAc(5)Hex(6)Fuc(1)NeuAc(1)); C13(Carbamidomethyl) | P0DTC2 | 0 | 4 | 1 | [illegible] |
| Trypsin(semi) | High | PMI-Byonic | Unambiguous | 279 | 300 | YNENGTITDAVDCALDPLSETK | 282 | HexNAc(5)Hex(6)Fuc(1)NeuGc(1) | 5 | 6 | 1 | | 1 | [K].YNEnGTITDAVDcALDPLSETK.[C] | N4(HexNAc(5)Hex(6)Fuc(1)NeuGc(1)); C13(Carbamidomethyl) | P0DTC2 | 0 | 3 | 1 | [illegible] |
| Trypsin(semi) | High | PMI-Byonic | Unambiguous | 279 | 300 | YNENGTITDAVDCALDPLSETK | 282 | HexNAc(5)Hex(6)Fuc(1) | 5 | 6 | 1 | | | [K].YNEnGTITDAVDcALDPLSETK.[C] | N4(HexNAc(5)Hex(6)Fuc(1)); C13(Carbamidomethyl) | P0DTC2 | 0 | 3 | 1 | [illegible] |
| Trypsin(semi) | High | PMI-Byonic | Unambiguous | 279 | 300 | YNENGTITDAVDCALDPLSETK | 282 | HexNAc(5)Hex(6)Fuc(3) | 5 | 6 | 3 | | | [K].YNEnGTITDAVDcALDPLSETK.[C] | N4(HexNAc(5)Hex(6)Fuc(3)); C13(Carbamidomethyl) | P0DTC2 | 0 | 3 | 1 | [illegible] |
| Trypsin(semi) | High | PMI-Byonic | Unambiguous | 279 | 300 | YNENGTITDAVDCALDPLSETK | 282 | HexNAc(5)Hex(6)NeuAc(1) | 5 | 6 | | 1 | | [K].YNEnGTITDAVDcALDPLSETK.[C] | N4(HexNAc(5)Hex(6)NeuAc(1)); C13(Carbamidomethyl) | P0DTC2 | 0 | 3 | 1 | [illegible] |
| Trypsin(semi) | High | PMI-Byonic | Unambiguous | 279 | 300 | YNENGTITDAVDCALDPLSETK | 282 | HexNAc(5)Hex(6) | 5 | 6 | | | | [K].YNEnGTITDAVDcALDPLSETK.[C] | N4(HexNAc(5)Hex(6)); C13(Carbamidomethyl) | P0DTC2 | 0 | 3 | 1 | [illegible] |
| Trypsin(semi) | High | PMI-Byonic | Unambiguous | 279 | 300 | YNENGTITDAVDCALDPLSETK | 282 | HexNAc(5)Hex(7)Fuc(1) | 5 | 7 | 1 | | | [K].YNEnGTITDAVDcALDPLSETK.[C] | N4(HexNAc(5)Hex(7)Fuc(1)); C13(Carbamidomethyl) | P0DTC2 | 0 | 3 | 1 | [illegible] |
| Trypsin(semi) | High | PMI-Byonic | Unambiguous | 279 | 300 | YNENGTITDAVDCALDPLSETK | 282 | HexNAc(5)Hex(7) | 5 | 7 | | | | [K].YNEnGTITDAVDcALDPLSETK.[C] | N4(HexNAc(5)Hex(7)); C13(Carbamidomethyl) | P0DTC2 | 0 | 3 | 1 | [illegible] |
| Trypsin(semi) | High | PMI-Byonic | Unambiguous | 279 | 300 | YNENGTITDAVDCALDPLSETK | 282 | HexNAc(6)Hex(3)Fuc(1) | 6 | 3 | 1 | | | [K].YNEnGTITDAVDcALDPLSETK.[C] | N4(HexNAc(6)Hex(3)Fuc(1)); C13(Carbamidomethyl) | P0DTC2 | 0 | 3 | 1 | [illegible] |
| Trypsin(semi) | High | PMI-Byonic | Unambiguous | 279 | 300 | YNENGTITDAVDCALDPLSETK | 282 | HexNAc(6)Hex(4)Fuc(1) | 6 | 4 | 1 | | | [K].YNEnGTITDAVDcALDPLSETK.[C] | N4(HexNAc(6)Hex(4)Fuc(1)); C13(Carbamidomethyl) | P0DTC2 | 0 | 3 | 1 | [illegible] |
| Trypsin(semi) | High | PMI-Byonic | Unambiguous | 279 | 300 | YNENGTITDAVDCALDPLSETK | 282 | HexNAc(6)Hex(5)Fuc(1) | 6 | 5 | 1 | | | [K].YNEnGTITDAVDcALDPLSETK.[C] | N4(HexNAc(6)Hex(5)Fuc(1)); C13(Carbamidomethyl) | P0DTC2 | 0 | 4 | 1 | [illegible] |
| Trypsin(semi) | High | PMI-Byonic | Unambiguous | 279 | 300 | YNENGTITDAVDCALDPLSETK | 282 | HexNAc(6)Hex(6)Fuc(1) | 6 | 6 | 1 | | | [K].YNEnGTITDAVDcALDPLSETK.[C] | N4(HexNAc(6)Hex(6)Fuc(1)); C13(Carbamidomethyl) | P0DTC2 | 0 | 3 | 1 | [illegible] |
| Trypsin(semi) | High | PMI-Byonic | Unambiguous | 279 | 300 | YNENGTITDAVDCALDPLSETK | 282 | HexNAc(6)Hex(6)Fuc(2) | 6 | 6 | 2 | | | [K].YNEnGTITDAVDcALDPLSETK.[C] | N4(HexNAc(6)Hex(6)Fuc(2)); C13(Carbamidomethyl) | P0DTC2 | 0 | 3 | 1 | [illegible] |
| Trypsin(semi) | High | PMI-Byonic | Unambiguous | 279 | 300 | YNENGTITDAVDCALDPLSETK | 282 | HexNAc(6)Hex(6)Fuc(3) | 6 | 6 | 3 | | | [K].YNEnGTITDAVDcALDPLSETK.[C] | N4(HexNAc(6)Hex(6)Fuc(3)); C13(Carbamidomethyl) | P0DTC2 | 0 | 3 | 1 | [illegible] |
| Trypsin(semi) | High | PMI-Byonic | Unambiguous | 279 | 300 | YNENGTITDAVDCALDPLSETK | 282 | HexNAc(6)Hex(6)NeuAc(1) | 6 | 6 | | 1 | | [K].YNEnGTITDAVDcALDPLSETK.[C] | N4(HexNAc(6)Hex(6)NeuAc(1)); C13(Carbamidomethyl) | P0DTC2 | 0 | 3 | 1 | [illegible] |
| Trypsin(semi) | High | PMI-Byonic | Unambiguous | 279 | 300 | YNENGTITDAVDCALDPLSETK | 282 | HexNAc(6)Hex(6) | 6 | 6 | | | | [K].YNEnGTITDAVDcALDPLSETK.[C] | N4(HexNAc(6)Hex(6)); C13(Carbamidomethyl) | P0DTC2 | 0 | 3 | 1 | [illegible] |
| Trypsin(semi) | High | PMI-Byonic | Unambiguous | 279 | 300 | YNENGTITDAVDCALDPLSETK | 282 | HexNAc(6)Hex(7)Fuc(1) | 6 | 7 | 1 | | | [K].YNEnGTITDAVDcALDPLSETK.[C] | N4(HexNAc(6)Hex(7)Fuc(1)); C13(Carbamidomethyl) | P0DTC2 | 0 | 3 | 1 | [illegible] |
| Trypsin(semi) | High | PMI-Byonic | Unambiguous | 279 | 300 | YNENGTITDAVDCALDPLSETK | 282 | HexNAc(6)Hex(7)Fuc(5) | 6 | 7 | 5 | | | [K].YNEnGTITDAVDcALDPLSETK.[C] | N4(HexNAc(6)Hex(7)Fuc(5)); C13(Carbamidomethyl) | P0DTC2 | 0 | 4 | 1 | [illegible] |
| Trypsin(semi) | High | PMI-Byonic | Unambiguous | 279 | 300 | YNENGTITDAVDCALDPLSETK | 282 | HexNAc(6)Hex(7)NeuAc(1) | 6 | 7 | | 1 | | [K].YNEnGTITDAVDcALDPLSETK.[C] | N4(HexNAc(6)Hex(7)NeuAc(1)); C13(Carbamidomethyl) | P0DTC2 | 0 | 3 | 1 | [illegible] |
| Trypsin(semi) | High | PMI-Byonic | Unambiguous | 279 | 300 | YNENGTITDAVDCALDPLSETK | 282 | HexNAc(6)Hex(7) | 6 | 7 | | | | [K].YNEnGTITDAVDcALDPLSETK.[C] | N4(HexNAc(6)Hex(7)); C13(Carbamidomethyl) | P0DTC2 | 0 | 3 | 1 | [illegible] |
| Trypsin(semi) | High | PMI-Byonic | Unambiguous | 279 | 300 | YNENGTITDAVDCALDPLSETK | 282 | HexNAc(6)Hex(8)Fuc(1)NeuAc(1) | 6 | 8 | 1 | 1 | | [K].YNEnGTITDAVDcALDPLSETK.[C] | N4(HexNAc(6)Hex(8)Fuc(1)NeuAc(1)); C13(Carbamidomethyl) | P0DTC2 | 0 | 4 | 1 | [illegible] |
| Trypsin(semi) | High | PMI-Byonic | Unambiguous | 279 | 300 | YNENGTITDAVDCALDPLSETK | 282 | HexNAc(6)Hex(8)NeuAc(1) | 6 | 8 | | 1 | | [K].YNEnGTITDAVDcALDPLSETK.[C] | N4(HexNAc(6)Hex(8)NeuAc(1)); C13(Carbamidomethyl) | P0DTC2 | 0 | 4 | 1 | [illegible] |
| Trypsin(semi) | High | PMI-Byonic | Unambiguous | 279 | 300 | YNENGTITDAVDCALDPLSETK | 282 | HexNAc(7)Hex(6)Fuc(1) | 7 | 6 | 1 | | | [K].YNEnGTITDAVDcALDPLSETK.[C] | N4(HexNAc(7)Hex(6)Fuc(1)); C13(Carbamidomethyl) | P0DTC2 | 0 | 3 | 1 | [illegible] |
| Trypsin(semi) | High | PMI-Byonic | Unambiguous | 280 | 300 | NENGTITDAVDCALDPLSETK | 282 | HexNAc(9)Hex(3) | 9 | 3 | | | | [Y].NEnGTITDAVDcALDPLSETK.[C] | N3(HexNAc(9)Hex(3)); C12(Carbamidomethyl) | P0DTC2 | 0 | 3 | 1 | [illegible] |
| Trypsin(semi) | High | PMI-Byonic | Unambiguous | 645 | 682 | TRAGCLIGAEHVNNSYECDIPIGAGICASYQTQTNSPR | 657 | HexNAc(2)Hex(9) | 2 | 9 | | | | [Q].TRAGcLIGAEHVnNSYEcDIPIGAGIcASYQTQTNSPR.[R] | C5(Carbamidomethyl); N13(HexNAc(2)Hex(9)); C18(Carbamidomethyl); C27(Carbamidomethyl) | P0DTC2 | 1 | 4 | 1 | [illegible] |
| Trypsin(semi) | High | PMI-Byonic | Unambiguous | 656 | 682 | VNNSYECDIPIGAGICASYQTQTNSPR | 657 | HexNAc(3)Hex(3)Fuc(2) | 3 | 3 | 2 | | | [H].VnNSYEcDIPIGAGIcASYQTQTNSPR.[R] | N2(HexNAc(3)Hex(3)Fuc(2)); C7(Carbamidomethyl); C16(Carbamidomethyl) | P0DTC2 | 0 | 3 | 1 | [illegible] |
| Trypsin(semi) | High | PMI-Byonic | Unambiguous | 647 | 682 | AGCLIGAEHVNNSYECDIPIGAGICASYQTQTNSPR | 657 | HexNAc(3)Hex(5)Fuc(1) | 3 | 5 | 1 | | | [R].AGcLIGAEHVnNSYEcDIPIGAGIcASYQTQTNSPR.[R] | C3(Carbamidomethyl); N11(HexNAc(3)Hex(5)Fuc(1)); C16(Carbamidomethyl); C25(Carbamidomethyl) | P0DTC2 | 0 | 4 | 1 | [illegible] |
| Trypsin(semi) | High | PMI-Byonic | Unambiguous | 647 | 682 | AGCLIGAEHVNNSYECDIPIGAGICASYQTQTNSPR | 657 | HexNAc(3)Hex(5)NeuGc(1) | 3 | 5 | | | 1 | [R].AGcLIGAEHVnNSYEcDIPIGAGIcASYQTQTNSPR.[R] | C3(Carbamidomethyl); N11(HexNAc(3)Hex(5)NeuGc(1)); C16(Carbamidomethyl); C25(Carbamidomethyl) | P0DTC2 | 0 | 4 | 1 | [illegible] |
| Trypsin(semi) | High | PMI-Byonic | Unambiguous | 650 | 682 | LIGAEHVNNSYECDIPIGAGICASYQTQTNSPR | 657 | HexNAc(3)Hex(6)Fuc(1)NeuAc(1) | 3 | 6 | 1 | 1 | | [C].LIGAEHVnNSYEcDIPIGAGIcASYQTQTNSPR.[R] | N8(HexNAc(3)Hex(6)Fuc(1)NeuAc(1)); C13(Carbamidomethyl); C22(Carbamidomethyl) | P0DTC2 | 0 | 4 | 1 | [illegible] |
| Trypsin(semi) | High | PMI-Byonic | Unambiguous | 647 | 682 | AGCLIGAEHVNNSYECDIPIGAGICASYQTQTNSPR | 657 | HexNAc(4)Hex(5)Fuc(1)NeuAc(1) | 4 | 5 | 1 | 1 | | [R].AGcLIGAEHVnNSYEcDIPIGAGIcASYQTQTNSPR.[R] | C3(Carbamidomethyl); N11(HexNAc(4)Hex(5)Fuc(1)NeuAc(1)); C16(Carbamidomethyl); C25(Carbamidomethyl) | P0DTC2 | 0 | 4 | 1 | [illegible] |
| Trypsin(semi) | High | PMI-Byonic | Unambiguous | 647 | 682 | AGCLIGAEHVNNSYECDIPIGAGICASYQTQTNSPR | 657 | HexNAc(4)Hex(5)Fuc(1) | 4 | 5 | 1 | | | [R].AGcLIGAEHVnNSYEcDIPIGAGIcASYQTQTNSPR.[R] | C3(Carbamidomethyl); N11(HexNAc(4)Hex(5)Fuc(1)); C16(Carbamidomethyl); C25(Carbamidomethyl) | P0DTC2 | 0 | 4 | 1 | [illegible] |
| Trypsin(semi) | High | PMI-Byonic | Unambiguous | 647 | 682 | AGCLIGAEHVNNSYECDIPIGAGICASYQTQTNSPR | 657 | HexNAc(4)Hex(5)Fuc(3) | 4 | 5 | 3 | | | [R].AGcLIGAEHVnNSYEcDIPIGAGIcASYQTQTNSPR.[R] | C3(Carbamidomethyl); N11(HexNAc(4)Hex(5)Fuc(3)); C16(Carbamidomethyl); C25(Carbamidomethyl) | P0DTC2 | 0 | 4 | 1 | [illegible] |
| Trypsin(semi) | High | PMI-Byonic | Unambiguous | 647 | 682 | AGCLIGAEHVNNSYECDIPIGAGICASYQTQTNSPR | 657 | HexNAc(4)Hex(5)NeuAc(1) | 4 | 5 | | 1 | | [R].AGcLIGAEHVnNSYEcDIPIGAGIcASYQTQTNSPR.[R] | C3(Carbamidomethyl); N11(HexNAc(4)Hex(5)NeuAc(1)); C16(Carbamidomethyl); C25(Carbamidomethyl) | P0DTC2 | 0 | 4 | 1 | [illegible] |
| Trypsin(semi) | High | PMI-Byonic | Unambiguous | 650 | 682 | LIGAEHVNNSYECDIPIGAGICASYQTQTNSPR | 657 | HexNAc(4)Hex(6)NeuAc(1) | 4 | 6 | | 1 | | [C].LIGAEHVnNSYEcDIPIGAGIcASYQTQTNSPR.[R] | N8(HexNAc(4)Hex(6)NeuAc(1)); C13(Carbamidomethyl); C22(Carbamidomethyl) | P0DTC2 | 0 | 4 | 1 | [illegible] |
| Trypsin(semi) | High | PMI-Byonic | Unambiguous | 650 | 682 | LIGAEHVNNSYECDIPIGAGICASYQTQTNSPR | 657 | HexNAc(5)Hex(4)Fuc(2)NeuAc(1) | 5 | 4 | 2 | 1 | | [C].LIGAEHVnNSYEcDIPIGAGIcASYQTQTNSPR.[R] | N8(HexNAc(5)Hex(4)Fuc(2)NeuAc(1)); C13(Carbamidomethyl); C22(Carbamidomethyl) | P0DTC2 | 0 | 4 | 1 | [illegible] |
| Trypsin(semi) | High | PMI-Byonic | Unambiguous | 647 | 682 | AGCLIGAEHVNNSYECDIPIGAGICASYQTQTNSPR | 657 | HexNAc(5)Hex(5)Fuc(1)NeuAc(1) | 5 | 5 | 1 | 1 | | [R].AGcLIGAEHVnNSYEcDIPIGAGIcASYQTQTNSPR.[R] | C3(Carbamidomethyl); N11(HexNAc(5)Hex(5)Fuc(1)NeuAc(1)); C16(Carbamidomethyl); C25(Carbamidomethyl) | P0DTC2 | 0 | 4 | 1 | [illegible] |
| Trypsin(semi) | High | PMI-Byonic | Unambiguous | 647 | 682 | AGCLIGAEHVNNSYECDIPIGAGICASYQTQTNSPR | 657 | HexNAc(5)Hex(5)Fuc(1) | 5 | 5 | 1 | | | [R].AGcLIGAEHVnNSYEcDIPIGAGIcASYQTQTNSPR.[R] | C3(Carbamidomethyl); N11(HexNAc(5)Hex(5)Fuc(1)); C16(Carbamidomethyl); C25(Carbamidomethyl) | P0DTC2 | 0 | 4 | 1 | [illegible] |
| Trypsin(semi) | High | PMI-Byonic | Unambiguous | 647 | 682 | AGCLIGAEHVNNSYECDIPIGAGICASYQTQTNSPR | 657 | HexNAc(5)Hex(5)Fuc(2) | 5 | 5 | 2 | | | [R].AGcLIGAEHVnNSYEcDIPIGAGIcASYQTQTNSPR.[R] | C3(Carbamidomethyl); N11(HexNAc(5)Hex(5)Fuc(2)); C16(Carbamidomethyl); C25(Carbamidomethyl) | P0DTC2 | 0 | 4 | 1 | [illegible] |
| Trypsin(semi) | High | PMI-Byonic | Unambiguous | 647 | 682 | AGCLIGAEHVNNSYECDIPIGAGICASYQTQTNSPR | 657 | HexNAc(5)Hex(6)Fuc(1) | 5 | 6 | 1 | | | [R].AGcLIGAEHVnNSYEcDIPIGAGIcASYQTQTNSPR.[R] | C3(Carbamidomethyl); N11(HexNAc(5)Hex(6)Fuc(1)); C16(Carbamidomethyl); C25(Carbamidomethyl) | P0DTC2 | 0 | 4 | 1 | [illegible] |
| Trypsin(semi) | High | PMI-Byonic | Unambiguous | 647 | 682 | AGCLIGAEHVNNSYECDIPIGAGICASYQTQTNSPR | 657 | HexNAc(5)Hex(6)Fuc(2)NeuAc(1) | 5 | 6 | 2 | 1 | | [R].AGcLIGAEHVnNSYEcDIPIGAGIcASYQTQTNSPR.[R] | C3(Carbamidomethyl); N11(HexNAc(5)Hex(6)Fuc(2)NeuAc(1)); C16(Carbamidomethyl); C25(Carbamidomethyl) | P0DTC2 | 0 | 4 | 1 | [illegible] |
| Trypsin(semi) | High | PMI-Byonic | Unambiguous | 647 | 682 | AGCLIGAEHVNNSYECDIPIGAGICASYQTQTNSPR | 657 | HexNAc(5)Hex(6)Fuc(2)NeuGc(1) | 5 | 6 | 2 | | 1 | [R].AGcLIGAEHVnNSYEcDIPIGAGIcASYQTQTNSPR.[R] | C3(Carbamidomethyl); N11(HexNAc(5)Hex(6)Fuc(2)NeuGc(1)); C16(Carbamidomethyl); C25(Carbamidomethyl) | P0DTC2 | 0 | 4 | 1 | [illegible] |
| Trypsin(semi) | High | PMI-Byonic | Unambiguous | 647 | 682 | AGCLIGAEHVNNSYECDIPIGAGICASYQTQTNSPR | 657 | HexNAc(5)Hex(6)Fuc(2) | 5 | 6 | 2 | | | [R].AGcLIGAEHVnNSYEcDIPIGAGIcASYQTQTNSPR.[R] | C3(Carbamidomethyl); N11(HexNAc(5)Hex(6)Fuc(2)); C16(Carbamidomethyl); C25(Carbamidomethyl) | P0DTC2 | 0 | 4 | 1 | [illegible] |
| Trypsin(semi) | High | PMI-Byonic | Unambiguous | 647 | 682 | AGCLIGAEHVNNSYECDIPIGAGICASYQTQTNSPR | 657 | HexNAc(5)Hex(6)NeuAc(1) | 5 | 6 | | 1 | | [R].AGcLIGAEHVnNSYEcDIPIGAGIcASYQTQTNSPR.[R] | C3(Carbamidomethyl); N11(HexNAc(5)Hex(6)NeuAc(1)); C16(Carbamidomethyl); C25(Carbamidomethyl) | P0DTC2 | 0 | 4 | 1 | [illegible] |
| Trypsin(semi) | High | PMI-Byonic | Unambiguous | 646 | 682 | RAGCLIGAEHVNNSYECDIPIGAGICASYQTQTNSPR | 657 | HexNAc(6)Hex(3)Fuc(1)NeuAc(2) | 6 | 3 | 1 | 2 | | [T].RAGcLIGAEHVnNSYEcDIPIGAGIcASYQTQTNSPR.[R] | C4(Carbamidomethyl); N12(HexNAc(6)Hex(3)Fuc(1)NeuAc(2)); C17(Carbamidomethyl); C26(Carbamidomethyl) | P0DTC2 | 1 | 4 | 1 | [illegible] |
| Trypsin(semi) | High | PMI-Byonic | Unambiguous | 650 | 682 | LIGAEHVNNSYECDIPIGAGICASYQTQTNSPR | 657 | HexNAc(6)Hex(3)Fuc(2)NeuGc(1) | 6 | 3 | 2 | | 1 | [C].LIGAEHVnNSYEcDIPIGAGIcASYQTQTNSPR.[R] | N8(HexNAc(6)Hex(3)Fuc(2)NeuGc(1)); C13(Carbamidomethyl); C22(Carbamidomethyl) | P0DTC2 | 0 | 4 | 1 | [illegible] |
| Trypsin(semi) | High | PMI-Byonic | Unambiguous | 647 | 682 | AGCLIGAEHVNNSYECDIPIGAGICASYQTQTNSPR | 657 | HexNAc(6)Hex(4)Fuc(1) | 6 | 4 | 1 | | | [R].AGcLIGAEHVnNSYEcDIPIGAGIcASYQTQTNSPR.[R] | C3(Carbamidomethyl); N11(HexNAc(6)Hex(4)Fuc(1)); C16(Carbamidomethyl); C25(Carbamidomethyl) | P0DTC2 | 0 | 4 | 1 | [illegible] |
| Trypsin(semi) | High | PMI-Byonic | Unambiguous | 650 | 682 | LIGAEHVNNSYECDIPIGAGICASYQTQTNSPR | 657 | HexNAc(6)Hex(5)Fuc(1)NeuAc(1)NeuGc(1) | 6 | 5 | 1 | 1 | 1 | [C].LIGAEHVnNSYEcDIPIGAGIcASYQTQTNSPR.[R] | N8(HexNAc(6)Hex(5)Fuc(1)NeuAc(1)NeuGc(1)); C13(Carbamidomethyl); C22(Carbamidomethyl) | P0DTC2 | 0 | 4 | 1 | [illegible] |
| Trypsin(semi) | High | PMI-Byonic | Unambiguous | 647 | 682 | AGCLIGAEHVNNSYECDIPIGAGICASYQTQTNSPR | 657 | HexNAc(6)Hex(6)Fuc(1) | 6 | 6 | 1 | | | [R].AGcLIGAEHVnNSYEcDIPIGAGIcASYQTQTNSPR.[R] | C3(Carbamidomethyl); N11(HexNAc(6)Hex(6)Fuc(1)); C16(Carbamidomethyl); C25(Carbamidomethyl) | P0DTC2 | 0 | 4 | 1 | [illegible] |
| Trypsin(semi) | High | PMI-Byonic | Unambiguous | 647 | 682 | AGCLIGAEHVNNSYECDIPIGAGICASYQTQTNSPR | 657 | HexNAc(6)Hex(6)Fuc(3) | 6 | 6 | 3 | | | [R].AGcLIGAEHVnNSYEcDIPIGAGIcASYQTQTNSPR.[R] | C3(Carbamidomethyl); N11(HexNAc(6)Hex(6)Fuc(3)); C16(Carbamidomethyl); C25(Carbamidomethyl) | P0DTC2 | 0 | 4 | 1 | [illegible] |
| Trypsin(semi) | High | PMI-Byonic | Unambiguous | 644 | 682 | QTRAGCLIGAEHVNNSYECDIPIGAGICASYQTQTNSPR | 657 | HexNAc(6)Hex(6)NeuAc(2) | 6 | 6 | | 2 | | [F].qTRAGcLIGAEHVnNSYEcDIPIGAGIcASYQTQTNSPR.[R] | Q1(Gln->pyro-Glu); C6(Carbamidomethyl); N14(HexNAc(6)Hex(6)NeuAc(2)); C19(Carbamidomethyl); C28(Carbamidomethyl) | P0DTC2 | 1 | 4 | 1 | [illegible] |
| Trypsin(semi) | High | PMI-Byonic | Unambiguous | 650 | 682 | LIGAEHVNNSYECDIPIGAGICASYQTQTNSPR | 657 | HexNAc(6)Hex(7)Fuc(1)NeuAc(1) | 6 | 7 | 1 | 1 | | [C].LIGAEHVnNSYEcDIPIGAGIcASYQTQTNSPR.[R] | N8(HexNAc(6)Hex(7)Fuc(1)NeuAc(1)); C13(Carbamidomethyl); C22(Carbamidomethyl) | P0DTC2 | 0 | 4 | 1 | [illegible] |
| Trypsin(semi) | High | PMI-Byonic | Unambiguous | 650 | 682 | LIGAEHVNNSYECDIPIGAGICASYQTQTNSPR | 657 | HexNAc(6)Hex(7)Fuc(4)NeuAc(1) | 6 | 7 | 4 | 1 | | [C].LIGAEHVnNSYEcDIPIGAGIcASYQTQTNSPR.[R] | N8(HexNAc(6)Hex(7)Fuc(4)NeuAc(1)); C13(Carbamidomethyl); C22(Carbamidomethyl) | P0DTC2 | 0 | 4 | 1 | [illegible] |
| Trypsin(semi) | High | PMI-Byonic | Unambiguous | 647 | 682 | AGCLIGAEHVNNSYECDIPIGAGICASYQTQTNSPR | 657 | HexNAc(6)Hex(7)NeuAc(1) | 6 | 7 | | 1 | | [R].AGcLIGAEHVnNSYEcDIPIGAGIcASYQTQTNSPR.[R] | C3(Carbamidomethyl); N11(HexNAc(6)Hex(7)NeuAc(1)); C16(Carbamidomethyl); C25(Carbamidomethyl) | P0DTC2 | 0 | 4 | 1 | [illegible] |

| | | | | | | | | | | | | | | |
|---|---|---|---|---|---|---|---|---|---|---|---|---|---|---|
| Trypsin(semi) | High | PMI-Byonic | Unambiguous | 696 | 733 | TMSLGAENSVAYSNNSIAIPTNFTISVTTEILPVSMTK | 717 | HexNAc(6)Hex(5)Fuc(1)NeuAc(1)NeuGc(2) | 6 | 5 | 1 | 1 | 2 | [illegible] |
| Trypsin(semi) | High | PMI-Byonic | Unambiguous | 697 | 733 | MSLGAENSVAYSNNSIAIPTNFTISVTTEILPVSMTK | 717 | HexNAc(6)Hex(6)NeuAc(1) | 6 | 6 | | 1 | | [illegible] |
| Trypsin(semi) | High | PMI-Byonic | Unambiguous | 711 | 733 | SIAIPTNFTISVTTEILPVSMTK | 717 | HexNAc(8)Hex(9)Fuc(1)NeuAc(2)NeuGc(1) | 8 | 9 | 1 | 2 | 1 | [N].SIAIPTnFTISVTTEILPVSMTK.[T] |
| Trypsin(semi) | High | PMI-Byonic | Unambiguous | 796 | 811 | DFGGFNFSQILPDPSK | 801 | HexNAc(2)Hex(4) | 2 | 4 | | | | [K].DFGGFnFSQILPDPSK.[P] |
| Trypsin(semi) | High | PMI-Byonic | Unambiguous | 796 | 811 | DFGGFNFSQILPDPSK | 801 | HexNAc(2)Hex(5) | 2 | 5 | | | | [K].DFGGFnFSQILPDPSK.[P] |
| Trypsin(semi) | High | PMI-Byonic | Unambiguous | 796 | 811 | DFGGFNFSQILPDPSK | 801 | HexNAc(2)Hex(6) | 2 | 6 | | | | [K].DFGGFnFSQILPDPSK.[P] |
| Trypsin(semi) | High | PMI-Byonic | Unambiguous | 796 | 811 | DFGGFNFSQILPDPSK | 801 | HexNAc(2)Hex(7) | 2 | 7 | | | | [K].DFGGFnFSQILPDPSK.[P] |
| Trypsin(semi) | High | PMI-Byonic | Unambiguous | 796 | 811 | DFGGFNFSQILPDPSK | 801 | HexNAc(2)Hex(8) | 2 | 8 | | | | [K].DFGGFnFSQILPDPSK.[P] |
| Trypsin(semi) | High | PMI-Byonic | Unambiguous | 796 | 811 | DFGGFNFSQILPDPSK | 801 | HexNAc(2)Hex(9) | 2 | 9 | | | | [K].DFGGFnFSQILPDPSK.[P] |
| Trypsin(semi) | High | PMI-Byonic | Unambiguous | 796 | 811 | DFGGFNFSQILPDPSK | 801 | HexNAc(3)Hex(5)Fuc(1) | 3 | 5 | 1 | | | [K].DFGGFnFSQILPDPSK.[P] |
| Trypsin(semi) | High | PMI-Byonic | Unambiguous | 796 | 811 | DFGGFNFSQILPDPSK | 801 | HexNAc(3)Hex(5)NeuAc(1) | 3 | 5 | | 1 | | [K].DFGGFnFSQILPDPSK.[P] |
| Trypsin(semi) | High | PMI-Byonic | Unambiguous | 796 | 811 | DFGGFNFSQILPDPSK | 801 | HexNAc(3)Hex(5) | 3 | 5 | | | | [K].DFGGFnFSQILPDPSK.[P] |
| Trypsin(semi) | High | PMI-Byonic | Unambiguous | 796 | 811 | DFGGFNFSQILPDPSK | 801 | HexNAc(3)Hex(6)Fuc(1) | 3 | 6 | 1 | | | [K].DFGGFnFSQILPDPSK.[P] |
| Trypsin(semi) | High | PMI-Byonic | Unambiguous | 796 | 811 | DFGGFNFSQILPDPSK | 801 | HexNAc(3)Hex(6)NeuAc(1) | 3 | 6 | | 1 | | [K].DFGGFnFSQILPDPSK.[P] |
| Trypsin(semi) | High | PMI-Byonic | Unambiguous | 796 | 811 | DFGGFNFSQILPDPSK | 801 | HexNAc(3)Hex(6)NeuGc(1) | 3 | 6 | | | 1 | [K].DFGGFnFSQILPDPSK.[P] |
| Trypsin(semi) | High | PMI-Byonic | Unambiguous | 796 | 811 | DFGGFNFSQILPDPSK | 801 | HexNAc(3)Hex(6) | 3 | 6 | | | | [K].DFGGFnFSQILPDPSK.[P] |
| Trypsin(semi) | High | PMI-Byonic | Unambiguous | 796 | 811 | DFGGFNFSQILPDPSK | 801 | HexNAc(4)Hex(3)Fuc(1) | 4 | 3 | 1 | | | [K].DFGGFnFSQILPDPSK.[P] |
| Trypsin(semi) | High | PMI-Byonic | Unambiguous | 796 | 811 | DFGGFNFSQILPDPSK | 801 | HexNAc(4)Hex(3) | 4 | 3 | | | | [K].DFGGFnFSQILPDPSK.[P] |
| Trypsin(semi) | High | PMI-Byonic | Unambiguous | 796 | 811 | DFGGFNFSQILPDPSK | 801 | HexNAc(4)Hex(4)Fuc(1) | 4 | 4 | 1 | | | [K].DFGGFnFSQILPDPSK.[P] |
| Trypsin(semi) | High | PMI-Byonic | Unambiguous | 796 | 811 | DFGGFNFSQILPDPSK | 801 | HexNAc(4)Hex(4)Fuc(2) | 4 | 4 | 2 | | | [K].DFGGFnFSQILPDPSK.[P] |
| Trypsin(semi) | High | PMI-Byonic | Unambiguous | 796 | 811 | DFGGFNFSQILPDPSK | 801 | HexNAc(4)Hex(5)Fuc(1)NeuAc(1) | 4 | 5 | 1 | 1 | | [K].DFGGFnFSQILPDPSK.[P] |
| Trypsin(semi) | High | PMI-Byonic | Unambiguous | 796 | 811 | DFGGFNFSQILPDPSK | 801 | HexNAc(4)Hex(5)Fuc(1)NeuGc(1) | 4 | 5 | 1 | | 1 | [K].DFGGFnFSQILPDPSK.[P] |
| Trypsin(semi) | High | PMI-Byonic | Unambiguous | 796 | 811 | DFGGFNFSQILPDPSK | 801 | HexNAc(4)Hex(5)Fuc(1) | 4 | 5 | 1 | | | [K].DFGGFnFSQILPDPSK.[P] |
| Trypsin(semi) | High | PMI-Byonic | Unambiguous | 796 | 811 | DFGGFNFSQILPDPSK | 801 | HexNAc(4)Hex(5)Fuc(2) | 4 | 5 | 2 | | | [K].DFGGFnFSQILPDPSK.[P] |
| Trypsin(semi) | High | PMI-Byonic | Unambiguous | 796 | 811 | DFGGFNFSQILPDPSK | 801 | HexNAc(4)Hex(5)Fuc(3) | 4 | 5 | 3 | | | [K].DFGGFnFSQILPDPSK.[P] |
| Trypsin(semi) | High | PMI-Byonic | Unambiguous | 796 | 811 | DFGGFNFSQILPDPSK | 801 | HexNAc(4)Hex(5)NeuGc(1) | 4 | 5 | | | 1 | [K].DFGGFnFSQILPDPSK.[P] |
| Trypsin(semi) | High | PMI-Byonic | Unambiguous | 796 | 811 | DFGGFNFSQILPDPSK | 801 | HexNAc(4)Hex(5) | 4 | 5 | | | | [K].DFGGFnFSQILPDPSK.[P] |
| Trypsin(semi) | High | PMI-Byonic | Unambiguous | 796 | 811 | DFGGFNFSQILPDPSK | 801 | HexNAc(4)Hex(6)Fuc(1) | 4 | 6 | 1 | | | [K].DFGGFnFSQILPDPSK.[P] |
| [illegible] | | | | | | | | | | | | | | |

| | | | | | | | | | |
|---|---|---|---|---|---|---|---|---|---|
| a-Lytic | High | PMI-Byonic | Unambiguous | 52 | 63 | QDLFLPPFFSNVT | 61 | HexNAc(4)Hex(6)Fuc(2) | [illegible] |
| a-Lytic | High | PMI-Byonic | Unambiguous | 52 | 63 | QDLFLPPFFSNVT | 61 | HexNAc(4)Hex(7)Fuc(1) | [illegible] |
| a-Lytic | High | PMI-Byonic | Unambiguous | 52 | 63 | QDLFLPPFFSNVT | 61 | HexNAc(4)Hex(7)Fuc(2) | [illegible] |
| a-Lytic | High | PMI-Byonic | Unambiguous | 52 | 63 | QDLFLPPFFSNVT | 61 | HexNAc(5)Hex(5)Fuc(1) | [illegible] |
| a-Lytic | High | PMI-Byonic | Unambiguous | 52 | 63 | QDLFLPPFFSNVT | 61 | HexNAc(5)Hex(5)Fuc(2) | [illegible] |
| a-Lytic | High | PMI-Byonic | Unambiguous | 52 | 63 | QDLFLPPFFSNVT | 61 | HexNAc(5)Hex(5)Fuc(3) | [illegible] |
| a-Lytic | High | PMI-Byonic | Unambiguous | 52 | 63 | QDLFLPPFFSNVT | 61 | HexNAc(5)Hex(6)Fuc(1) | [illegible] |
| a-Lytic | High | PMI-Byonic | Unambiguous | 52 | 63 | QDLFLPPFFSNVT | 61 | HexNAc(5)Hex(6)Fuc(3) | [illegible] |
| a-Lytic | High | PMI-Byonic | Unambiguous | 52 | 63 | QDLFLPPFFSNVT | 61 | HexNAc(5)Hex(6) | [illegible] |
| a-Lytic | High | PMI-Byonic | Unambiguous | 52 | 63 | QDLFLPPFFSNVT | 61 | HexNAc(6)Hex(6)Fuc(1)Neu[illegible] | [illegible] |
| a-Lytic | High | PMI-Byonic | Unambiguous | 52 | 63 | QDLFLPPFFSNVT | 61 | HexNAc(6)Hex(6)Fuc(2) | [illegible] |
| a-Lytic | High | PMI-Byonic | Unambiguous | 52 | 63 | QDLFLPPFFSNVT | 61 | HexNAc(6)Hex(6)Fuc(3) | [illegible] |
| a-Lytic | High | PMI-Byonic | Unambiguous | 52 | 63 | QDLFLPPFFSNVT | 61 | HexNAc(8)Hex(3) | [illegible] |
| a-Lytic | High | PMI-Byonic | Unambiguous | 52 | 63 | QDLFLPPFFSNVT | 61 | HexNAc(9)Hex(3)Fuc(1) | [illegible] |
| a-Lytic | High | PMI-Byonic | Unambiguous | 72 | 90 | GTNGTKRFDNPVLPFNDGV | 74 | HexNAc(5)Hex(5)Fuc(1)Neu[illegible] | [illegible] |
| a-Lytic | High | PMI-Byonic | Unambiguous | 72 | 90 | GTNGTKRFDNPVLPFNDGV | 74 | HexNAc(6)Hex(7)Fuc(5) | [illegible] |
| a-Lytic | High | PMI-Byonic | Unambiguous | 163 | 171 | ANNCTFEYV | 165 | HexNAc(4)Hex(5)Fuc(3) | [illegible] |
| a-Lytic | High | PMI-Byonic | Unambiguous | 163 | 171 | ANNCTFEYV | 165 | HexNAc(5)Hex(5)Fuc(3) | [illegible] |
| a-Lytic | High | PMI-Byonic | Unambiguous | 228 | 236 | DLPIGINIT | 234 | HexNAc(2)Hex(6) | [illegible] |
| a-Lytic | High | PMI-Byonic | Unambiguous | 228 | 236 | DLPIGINIT | 234 | HexNAc(2)Hex(7) | [illegible] |
| a-Lytic | High | PMI-Byonic | Unambiguous | 228 | 236 | DLPIGINIT | 234 | HexNAc(2)Hex(8) | [illegible] |
| a-Lytic | High | PMI-Byonic | Unambiguous | 228 | 236 | DLPIGINIT | 234 | HexNAc(2)Hex(9) | [illegible] |
| a-Lytic | High | PMI-Byonic | Unambiguous | 334 | 345 | NLCPFGEVFNAT | 343 | HexNAc(2)Hex(6) | [illegible] |
| a-Lytic | High | PMI-Byonic | Unambiguous | 334 | 345 | NLCPFGEVFNAT | 343 | HexNAc(4)Hex(5)Fuc(1)Neu[illegible] | [illegible] |
| a-Lytic | High | PMI-Byonic | Unambiguous | 334 | 345 | NLCPFGEVFNAT | 343 | HexNAc(4)Hex(5)Fuc(1) | [illegible] |
| a-Lytic | High | PMI-Byonic | Unambiguous | 334 | 345 | NLCPFGEVFNAT | 343 | HexNAc(4)Hex(5)Fuc(2)Neu[illegible] | [illegible] |
| a-Lytic | High | PMI-Byonic | Unambiguous | 334 | 345 | NLCPFGEVFNAT | 343 | HexNAc(4)Hex(5)Fuc(2) | [illegible] |
| a-Lytic | High | PMI-Byonic | Unambiguous | 334 | 345 | NLCPFGEVFNAT | 343 | HexNAc(4)Hex(5)Fuc(3) | [illegible] |
| a-Lytic | High | PMI-Byonic | Unambiguous | 334 | 345 | NLCPFGEVFNAT | 343 | HexNAc(4)Hex(7)Fuc(2) | [illegible] |
| a-Lytic | High | PMI-Byonic | Unambiguous | 334 | 345 | NLCPFGEVFNAT | 343 | HexNAc(5)Hex(4)Fuc(1) | [illegible] |
| a-Lytic | High | PMI-Byonic | Unambiguous | 334 | 345 | NLCPFGEVFNAT | 343 | HexNAc(5)Hex(5)Fuc(1) | [illegible] |
| a-Lytic | High | PMI-Byonic | Unambiguous | 334 | 345 | NLCPFGEVFNAT | 343 | HexNAc(5)Hex(5)Fuc(2) | [illegible] |
| a-Lytic | High | PMI-Byonic | Unambiguous | 334 | 345 | NLCPFGEVFNAT | 343 | HexNAc(5)Hex(5)Fuc(3) | [illegible] |
| a-Lytic | High | PMI-Byonic | Unambiguous | 334 | 345 | NLCPFGEVFNAT | 343 | HexNAc(5)Hex(6)Fuc(1)Neu[illegible] | [illegible] |
| a-Lytic | High | PMI-Byonic | Unambiguous | 334 | 345 | NLCPFGEVFNAT | 343 | HexNAc(6)Hex(6)Fuc(2) | [illegible] |
| a-Lytic | High | PMI-Byonic | Unambiguous | 334 | 345 | NLCPFGEVFNAT | 343 | HexNAc(6)Hex(7)Fuc(4) | [illegible] |
| [illegible] | | | | | | | | | |

| | | | | | | | | | | | | | | | | | | | | | | | | | | | | | | | | | | | | | | | | | | | | | | |
|---|---|---|---|---|---|---|---|---|---|---|---|---|---|---|---|---|---|---|---|---|---|---|---|---|---|---|---|---|---|---|---|---|---|---|---|---|---|---|---|---|---|---|---|---|---|---|
| a-Lytic (semi) | High | PMI-Byonic | Unambiguous | 707 | 716 | YSNNSIAIPT | 709 | HexNAc(3)Hex(6) | 3 | 6 | | | | [A].YSnNSIAIPT.[N] | N3(HexNAc(3)Hex(6)) | P0DTC2 | 3 | 2 | 1 | 1331.05005 | 2661.09282 | 2661.09182 | 0.37 | 0.0005 | 317230.281 | HCD (High Energy Collision Dissociation) | MS2 | 9.215857 | 200 | 56.619 | 11806 | 11806 | 11783 | 3.4 | 212.6 | 127.4 | 127.4 | 0.00039436 | 0.000055243 | 0 | 0 | 0.07125709 | 0.02023121 | 0.0201729 | 0.0291262 | HexNAc(3)Hex(6) |
| a-Lytic (semi) | High | PMI-Byonic | Unambiguous | 707 | 716 | YSNNSIAIPT | 709 | HexNAc(4)Hex(5)Fuc(1) | 4 | 5 | 1 | | | [A].YSnNSIAIPT.[N] | N3(HexNAc(4)Hex(5)Fuc(1)) | P0DTC2 | 3 | 2 | 1 | 1424.58972 | 2848.17217 | 2848.17628 | -1.44 | -0.00206 | 238050.641 | HCD (High Energy Collision Dissociation) | MS2 | 46.91282 | 200 | 56.5497 | 11788 | 11788 | 11770 | 4.43 | 235.6 | 142.4 | 142.4 | 0.000037128 | 0.000007699 | 0 | 0 | 0.00715727 | 0.00148915 | 0 | 0 | HexNAc(4)Hex(5)Fuc(1) |
| a-Lytic (semi) | High | PMI-Byonic | Unambiguous | 707 | 716 | YSNNSIAIPT | 709 | HexNAc(4)Hex(5) | 4 | 5 | | | | [A].YSnNSIAIPT.[N] | N3(HexNAc(4)Hex(5)) | P0DTC2 | 3 | 2 | 1 | 1351.56226 | 2702.11724 | 2702.11837 | -0.42 | -0.00057 | 312605.156 | HCD (High Energy Collision Dissociation) | MS2 | 29.94578 | 200 | 56.5691 | 11793 | 11793 | 11770 | 3.29 | 194.3 | 93.9 | 93.9 | 0.00051002 | 0.000086067 | 0 | 0 | 0.09053083 | 0.02168021 | 0.02116402 | 0.03097344 | HexNAc(4)Hex(5) |
| a-Lytic (semi) | High | PMI-Byonic | Unambiguous | 707 | 716 | YSNNSIAIPT | 709 | HexNAc(5)Hex(6)Fuc(1) | 5 | 6 | 1 | | | [A].YSnNSIAIPT.[N] | N3(HexNAc(5)Hex(6)Fuc(1)) | P0DTC2 | 3 | 3 | 1 | 1071.77393 | 3213.30722 | 3213.30848 | -0.39 | -0.00042 | 356774.531 | HCD (High Energy Collision Dissociation) | MS2 | 4.940231 | 130.728 | 55.8691 | 11620 | 11620 | 11605 | 3.23 | 181.3 | 116.5 | 116.5 | 0.00058613 | 0.000097183 | 0 | 0 | 0.10279539 | 0.02331606 | 0.02307692 | 0.03404254 | HexNAc(5)Hex(6)Fuc(1) |
| a-Lytic (semi) | High | PMI-Byonic | Unambiguous | 707 | 716 | YSNNSIAIPT | 709 | HexNAc(6)Hex(5)Fuc(1) | 6 | 5 | 1 | | | [A].YSnNSIAIPT.[N] | N3(HexNAc(6)Hex(5)Fuc(1)) | P0DTC2 | 3 | 3 | 1 | 1085.44946 | 3254.33384 | 3254.33503 | -0.37 | -0.0004 | 261907.297 | HCD (High Energy Collision Dissociation) | MS2 | 15.50362 | 168.593 | 56.5387 | 11785 | 11785 | 11770 | 3.11 | 157.3 | 91.1 | 91.1 | 0.00077616 | 0.00013823 | 0 | 0 | 0.13205254 | 0.02612826 | 0.02606634 | 0.03952568 | HexNAc(6)Hex(5)Fuc(1) |
| a-Lytic (semi) | High | PMI-Byonic | Unambiguous | 717 | 721 | NFTIS | 717 | HexNAc(2)Hex(5) | 2 | 5 | | | | [T].nFTIS.[V] | N1(HexNAc(2)Hex(5)) | P0DTC2 | 1 | 2 | 1 | 899.36206 | 1797.71684 | 1797.71582 | 0.57 | 0.00051 | 694486.625 | HCD (High Energy Collision Dissociation) | MS2 | 10.3996 | 98.483 | 49.0461 | 9915 | 9915 | 9900 | 3.42 | 428.2 | 103.5 | 103.5 | 0.00038297 | 0.000054254 | 0 | 0 | 0.06931508 | 0.01754385 | 0.01734104 | 0.02439023 | HexNAc(2)Hex(5) |
| a-Lytic (semi) | High | PMI-Byonic | Unambiguous | 717 | 721 | NFTIS | 717 | HexNAc(2)Hex(6) | 2 | 6 | | | | [T].nFTIS.[V] | N1(HexNAc(2)Hex(6)) | P0DTC2 | 1 | 2 | 1 | 980.38788 | 1959.76848 | 1959.76864 | -0.08 | -0.00008 | 785133.313 | HCD (High Energy Collision Dissociation) | MS2 | 5.196741 | 181.32 | 48.3379 | 9719 | 9719 | 9701 | 3.82 | 459.4 | 110.7 | 110.7 | 0.00015202 | 0.000022413 | 0 | 0 | 0.02867779 | 0.00706714 | 0.00666666 | 0.01117318 | HexNAc(2)Hex(6) |
| a-Lytic (semi) | High | PMI-Byonic | Unambiguous | 717 | 721 | NFTIS | 717 | HexNAc(2)Hex(7) | 2 | 7 | | | | [T].nFTIS.[V] | N1(HexNAc(2)Hex(7)) | P0DTC2 | 1 | 2 | 1 | 1061.41431 | 2121.82134 | 2121.82146 | -0.06 | -0.00006 | 467063.063 | HCD (High Energy Collision Dissociation) | MS2 | 0 | 115.3 | 47.7826 | 9558 | 9558 | 9540 | 3.82 | 458.8 | 111.7 | 111.7 | 0.00015202 | 0.00002154 | 0 | 0 | 0.02867779 | 0.00706714 | 0.00671141 | 0.01123595 | HexNAc(2)Hex(7) |
| a-Lytic (semi) | High | PMI-Byonic | Unambiguous | 717 | 721 | NFTIS | 717 | HexNAc(2)Hex(8) | 2 | 8 | | | | [T].nFTIS.[V] | N1(HexNAc(2)Hex(8)) | P0DTC2 | 1 | 2 | 1 | 1142.44055 | 2283.87383 | 2283.87429 | -0.2 | -0.00023 | 233912.578 | HCD (High Energy Collision Dissociation) | MS2 | 0 | 200 | 47.7019 | 9538 | 9538 | 9516 | 3.82 | 450.2 | 116.9 | 116.9 | 0.00015202 | 0.0000211 | 0 | 0 | 0.02867779 | 0.00706714 | 0.006734 | 0.01129943 | HexNAc(2)Hex(8) |
| a-Lytic (semi) | High | PMI-Byonic | Unambiguous | 717 | 721 | NFTIS | 717 | HexNAc(2)Hex(9) | 2 | 9 | | | | [T].nFTIS.[V] | N1(HexNAc(2)Hex(9)) | P0DTC2 | 1 | 2 | 1 | 1223.46802 | 2445.92876 | 2445.92711 | 0.67 | 0.00082 | 355510.188 | HCD (High Energy Collision Dissociation) | MS2 | 0 | 200 | 47.7174 | 9542 | 9542 | 9529 | 3.16 | 357.3 | 48.3 | 48.3 | 0.00068773 | 0.00011532 | 0 | 0 | 0.11867245 | 0.02331606 | 0.02227722 | 0.03278687 | HexNAc(2)Hex(9) |
| a-Lytic (semi) | High | PMI-Byonic | Unambiguous | 717 | 721 | NFTIS | 717 | HexNAc(3)Hex(5)Fuc(1) | 3 | 5 | 1 | | | [T].nFTIS.[V] | N1(HexNAc(3)Hex(5)Fuc(1)) | P0DTC2 | 1 | 2 | 1 | 1073.92896 | 2146.85063 | 2146.8531 | -1.15 | -0.00123 | 477471.406 | HCD (High Energy Collision Dissociation) | MS2 | 0 | 148.01 | 48.58 | 9788 | 9788 | 9766 | 3.1 | 335.1 | 50.7 | 50.7 | 0.00079389 | 0.0001414 | 0 | 0 | 0.13468705 | 0.02612826 | 0.02588235 | 0.03906248 | HexNAc(3)Hex(5)Fuc(1) |
| a-Lytic (semi) | High | PMI-Byonic | Unambiguous | 717 | 721 | NFTIS | 717 | HexNAc(3)Hex(5) | 3 | 5 | | | | [T].nFTIS.[V] | N1(HexNAc(3)Hex(5)) | P0DTC2 | 1 | 2 | 1 | 1000.90106 | 2000.79485 | 2000.79519 | -0.17 | -0.00017 | 1091030.13 | HCD (High Energy Collision Dissociation) | MS2 | 46.73877 | 157.745 | 49.1867 | 9954 | 9954 | 9939 | 3.51 | 360.8 | 79.1 | 79.1 | 0.00030667 | 0.000039765 | 0 | 0 | 0.05622729 | 0.01273885 | 0.01219512 | 0.01538461 | HexNAc(3)Hex(5) |
| a-Lytic (semi) | High | PMI-Byonic | Unambiguous | 717 | 721 | NFTIS | 717 | HexNAc(3)Hex(6)Fuc(1) | 3 | 6 | 1 | | | [T].nFTIS.[V] | N1(HexNAc(3)Hex(6)Fuc(1)) | P0DTC2 | 1 | 2 | 1 | 1154.95654 | 2308.90581 | 2308.90592 | -0.05 | -0.00006 | 1323961.25 | HCD (High Energy Collision Dissociation) | MS2 | 2.63344 | 70.294 | 47.863 | 9580 | 9580 | 9566 | 3.58 | 372.2 | 85.2 | 85.2 | 0.00026199 | 0.000032041 | 0 | 0 | 0.04842654 | 0.01273885 | 0.01257861 | 0.01587301 | HexNAc(3)Hex(6)Fuc(1) |
| a-Lytic (semi) | High | PMI-Byonic | Unambiguous | 717 | 721 | NFTIS | 717 | HexNAc(3)Hex(6)NeuGc(1) | 3 | 6 | | | 1 | [T].nFTIS.[V] | N1(HexNAc(3)Hex(6)NeuGc(1)) | P0DTC2 | 1 | 2 | 1 | 1235.47253 | 2469.93779 | 2469.93834 | -0.22 | -0.00028 | 256662.063 | HCD (High Energy Collision Dissociation) | MS2 | 11.06873 | 194.055 | 51.8761 | 10633 | 10633 | 10618 | 3.47 | 339.1 | 70.4 | 70.4 | 0.00034227 | 0.000044035 | 0 | 0 | 0.06235193 | 0.01273885 | 0.01201201 | 0.01507537 | HexNAc(3)Hex(6)NeuGc(1) |
| a-Lytic (semi) | High | PMI-Byonic | Unambiguous | 717 | 721 | NFTIS | 717 | HexNAc(3)Hex(6) | 3 | 6 | | | | [T].nFTIS.[V] | N1(HexNAc(3)Hex(6)) | P0DTC2 | 1 | 2 | 1 | 1081.92749 | 2162.8477 | 2162.84801 | -0.14 | -0.00015 | 519258.563 | HCD (High Energy Collision Dissociation) | MS2 | 0 | 200 | 48.3877 | 9733 | 9733 | 9714 | 3.48 | 396.5 | 98.6 | 98.6 | 0.00033096 | 0.000043134 | 0 | 0 | 0.06041475 | 0.01273885 | 0.01204819 | 0.01515151 | HexNAc(3)Hex(6) |
| a-Lytic (semi) | High | PMI-Byonic | Unambiguous | 717 | 721 | NFTIS | 717 | HexNAc(4)Hex(5) | 4 | 5 | | | | [T].nFTIS.[V] | N1(HexNAc(4)Hex(5)) | P0DTC2 | 1 | 2 | 1 | 1102.4425 | 2203.87773 | 2203.87456 | 1.44 | 0.00159 | 217172.406 | HCD (High Energy Collision Dissociation) | MS2 | 14.5854 | 193.898 | 48.7083 | 9822 | 9822 | 9807 | 3.07 | 308.5 | 66.7 | 66.7 | 0.00085825 | 0.00014645 | 0 | 0 | 0.14412325 | 0.02803738 | 0.02790697 | 0.04230768 | HexNAc(4)Hex(5) |
| a-Lytic (semi) | High | PMI-Byonic | Unambiguous | 717 | 723 | NFTISVT | 717 | HexNAc(7)Hex(8)Fuc(1)NeuAc(1)NeuGc(1) | 7 | 8 | 1 | 1 | 1 | [T].nFTISVT.[T] | N1(HexNAc(7)Hex(8)Fuc(1)NeuAc(1)NeuGc(1)) | P0DTC2 | 3 | 3 | 1 | 1414.88428 | 4242.63828 | 4243.6309 | -233.91 | -0.33087 | 338462.781 | HCD (High Energy Collision Dissociation) | MS2 | 0 | 182.092 | 47.9227 | 9598 | 9598 | 9579 | 1.57 | 276.3 | 75.6 | 75.6 | 0.02676632 | 0.00578035 | 0.00575816 | 0.00632911 | 0.84119489 | 0.14188032 | 0.14139691 | 0.19280201 | HexNAc(7)Hex(8)Fuc(1)NeuAc(1)NeuGc(1) |
| a-Lytic (semi) | High | PMI-Byonic | Unambiguous | 717 | 723 | NFTISVT | 717 | HexNAc(7)Hex(8)Fuc(1)NeuAc(1) | 7 | 8 | 1 | 1 | | [T].nFTISVT.[T] | N1(HexNAc(7)Hex(8)Fuc(1)NeuAc(1)) | P0DTC2 | 3 | 3 | 1 | 1312.85583 | 3936.55295 | 3936.54057 | 3.15 | 0.00413 | 362706.438 | HCD (High Energy Collision Dissociation) | MS2 | 0 | 200 | 48.6331 | 9803 | 9803 | 9779 | 3.15 | 242.9 | 78.4 | 78.4 | 0.00070621 | 0.00011975 | 0 | 0 | 0.12150147 | 0.02331606 | 0.02211302 | 0.03252031 | HexNAc(7)Hex(8)Fuc(1)NeuAc(1) |
| a-Lytic (semi) | High | PMI-Byonic | Unambiguous | 786 | 813 | KQIYKTPPIKDFGGFNFSQILPDPSKPS | 801 | HexNAc(2)Hex(4) | 2 | 4 | | | | [V].KQIYKTPPIKDFGGFnFSQILPDPSKPS.[K] | N16(HexNAc(2)Hex(4)) | P0DTC2 | 2 | 5 | 1 | 841.81384 | 4205.04011 | 4204.03704 | 238.6 | 0.20061 | 177391.516 | HCD (High Energy Collision Dissociation) | MS2 | 9.485111 | 200 | 81.4766 | 18082 | 18082 | 18063 | 5.59 | 194 | 194 | 194 | 2.5673E-06 | 4.7636E-07 | 0 | 0 | 0.00049819 | 0.000092456 | 0 | 0 | HexNAc(2)Hex(4) |
| a-Lytic (semi) | High | PMI-Byonic | Unambiguous | 786 | 803 | KQIYKTPPIKDFGGFNFS | 801 | HexNAc(2)Hex(5) | 2 | 5 | | | | [V].KQIYKTPPIKDFGGFnFS.[Q] | N16(HexNAc(2)Hex(5)) | P0DTC2 | 0 | 4 | 1 | 826.63538 | 3303.51967 | 3303.5189 | 0.23 | 0.00019 | 230510.422 | HCD (High Energy Collision Dissociation) | MS2 | 46.01514 | 174.433 | 70.699 | 15335 | 15335 | 15317 | 5.57 | 476.4 | 473 | 473 | 0.000002667 | 4.9966E-07 | 0 | 0 | 0.00051753 | 0.000096978 | 0 | 0 | HexNAc(2)Hex(5) |
| a-Lytic (semi) | High | PMI-Byonic | Unambiguous | 786 | 803 | KQIYKTPPIKDFGGFNFS | 801 | HexNAc(2)Hex(6) | 2 | 6 | | | | [V].KQIYKTPPIKDFGGFnFS.[Q] | N16(HexNAc(2)Hex(6)) | P0DTC2 | 0 | 4 | 1 | 867.14825 | 3465.57119 | 3465.57172 | -0.15 | -0.00013 | 1602109.63 | HCD (High Energy Collision Dissociation) | MS2 | 34.30973 | 34.566 | 70.5911 | 15305 | 15305 | 15291 | 4.93 | 478.7 | 263.9 | 263.9 | 0.000011617 | 3.1098E-06 | 0 | 0 | 0.00225047 | 0.00060301 | 0 | 0 | HexNAc(2)Hex(6) |
| a-Lytic (semi) | High | PMI-Byonic | Unambiguous | 786 | 803 | KQIYKTPPIKDFGGFNFS | 801 | HexNAc(2)Hex(7) | 2 | 7 | | | | [V].KQIYKTPPIKDFGGFnFS.[Q] | N16(HexNAc(2)Hex(7)) | P0DTC2 | 0 | 4 | 1 | 907.66132 | 3627.62343 | 3627.62455 | -0.31 | -0.00028 | 516290.938 | HCD (High Energy Collision Dissociation) | MS2 | 2.268208 | 95.633 | 70.3946 | 15249 | 15249 | 15235 | 5.12 | 458.6 | 292.9 | 292.9 | 7.5279E-06 | 2.1179E-06 | 0 | 0 | 0.00145941 | 0.00041082 | 0 | 0 | HexNAc(2)Hex(7) |
| a-Lytic (semi) | High | PMI-Byonic | Unambiguous | 786 | 803 | KQIYKTPPIKDFGGFNFS | 801 | HexNAc(2)Hex(8) | 2 | 8 | | | | [V].KQIYKTPPIKDFGGFnFS.[Q] | N16(HexNAc(2)Hex(8)) | P0DTC2 | 0 | 4 | 1 | 948.17548 | 3789.68007 | 3789.67737 | 0.71 | 0.00068 | 277017.813 | HCD (High Energy Collision Dissociation) | MS2 | 0 | 200 | 70.3628 | 15240 | 15240 | 15224 | 4.8 | 482.2 | 255.8 | 255.8 | 0.000015862 | 4.1802E-06 | 0 | 0 | 0.0030702 | 0.00081021 | 0 | 0 | HexNAc(2)Hex(8) |
| a-Lytic (semi) | High | PMI-Byonic | Unambiguous | 785 | 813 | VKQIYKTPPIKDFGGFNFSQILPDPSKPS | 801 | HexNAc(3)Hex(4)Fuc(1)NeuAc(1) | 3 | 4 | 1 | 1 | | [Q].VKQIYKTPPIKDFGGFnFSQILPDPSKPS.[K] | N17(HexNAc(3)Hex(4)Fuc(1)NeuAc(1)) | P0DTC2 | 3 | 5 | 1 | 989.67694 | 4944.3556 | 4943.33815 | 205.82 | 0.20349 | 289734.219 | HCD (High Energy Collision Dissociation) | MS2 | 0 | 200 | 81.9971 | 18226 | 18226 | 18204 | 2.46 | 0.9 | 0.9 | 0.9 | 0.00348746 | 0.00034146 | 0 | 0 | 0.404537 | 0.05918366 | 0.05906312 | 0.08580855 | HexNAc(3)Hex(4)Fuc(1)NeuAc(1) |
| a-Lytic (semi) | High | PMI-Byonic | Unambiguous | 786 | 813 | KQIYKTPPIKDFGGFNFSQILPDPSKPS | 801 | HexNAc(3)Hex(4)NeuGc(1) | 3 | 4 | | | 1 | [V].KQIYKTPPIKDFGGFnFSQILPDPSKPS.[K] | N16(HexNAc(3)Hex(4)NeuGc(1)) | P0DTC2 | 2 | 5 | 1 | 943.85052 | 4715.22352 | 4714.20674 | 215.68 | 0.20336 | 1052157.75 | HCD (High Energy Collision Dissociation) | MS2 | 62.15588 | 17.158 | 80.8603 | 17913 | 17913 | 17895 | 6.03 | 356.7 | 306 | 0.8 | 9.2896E-07 | 1.404E-07 | 0 | 0 | 0.00018032 | 0.000027257 | 0 | 0 | HexNAc(3)Hex(4)NeuGc(1) |
| a-Lytic (semi) | High | PMI-Byonic | Unambiguous | 786 | 803 | KQIYKTPPIKDFGGFNFS | 801 | HexNAc(3)Hex(5)Fuc(1) | 3 | 5 | 1 | | | [V].KQIYKTPPIKDFGGFnFS.[Q] | N16(HexNAc(3)Hex(5)Fuc(1)) | P0DTC2 | 0 | 4 | 1 | 913.91998 | 3652.6581 | 3652.65618 | 0.53 | 0.00048 | 451859.656 | HCD (High Energy Collision Dissociation) | MS2 | 0 | 101.415 | 70.5591 | 15296 | 15296 | 15278 | 5.15 | 465.6 | 322.9 | 322.9 | 7.0523E-06 | 1.8362E-06 | 0 | 0 | 0.00136735 | 0.00035622 | 0 | 0 | HexNAc(3)Hex(5)Fuc(1) |
| a-Lytic (semi) | High | PMI-Byonic | Unambiguous | 786 | 813 | KQIYKTPPIKDFGGFNFSQILPDPSKPS | 801 | HexNAc(3)Hex(5)NeuAc(1) | 3 | 5 | | 1 | | [V].KQIYKTPPIKDFGGFnFSQILPDPSKPS.[K] | N16(HexNAc(3)Hex(5)NeuAc(1)) | P0DTC2 | 2 | 5 | 1 | 972.85822 | 4860.26197 | 4860.26465 | -0.55 | -0.00054 | 89707.7109 | HCD (High Energy Collision Dissociation) | MS2 | 0 | 200 | 83.7307 | 18678 | 18678 | 18663 | 7 | 360.4 | 309.8 | 309.8 | 1.0103E-07 | 6.218E-08 | 0 | 0 | 0.000019615 | 0.000012072 | 0 | 0 | HexNAc(3)Hex(5)NeuAc(1) |
| a-Lytic (semi) | High | PMI-Byonic | Unambiguous | 786 | 813 | KQIYKTPPIKDFGGFNFSQILPDPSKPS | 801 | HexNAc(3)Hex(5)NeuGc(1) | 3 | 5 | | | 1 | [V].KQIYKTPPIKDFGGFnFSQILPDPSKPS.[K] | N16(HexNAc(3)Hex(5)NeuGc(1)) | P0DTC2 | 2 | 5 | 1 | 976.0578 | 4876.25989 | 4876.25956 | 0.07 | 0.00007 | 210124.063 | HCD (High Energy Collision Dissociation) | MS2 | 0 | 200 | 83.4081 | 18603 | 18603 | 18589 | 7.23 | 381.5 | 296.4 | 44.5 | 5.8737E-08 | 3.7458E-08 | 0 | 0 | 0.000011404 | 7.2723E-06 | 0 | 0 | HexNAc(3)Hex(5)NeuGc(1) |
| a-Lytic (semi) | High | PMI-Byonic | Unambiguous | 786 | 803 | KQIYKTPPIKDFGGFNFS | 801 | HexNAc(3)Hex(5) | 3 | 5 | | | | [V].KQIYKTPPIKDFGGFnFS.[Q] | N16(HexNAc(3)Hex(5)) | P0DTC2 | 0 | 4 | 1 | 877.40479 | 3506.59731 | 3506.59827 | -0.27 | -0.00024 | 443783.563 | HCD (High Energy Collision Dissociation) | MS2 | 0 | 200 | 70.7324 | 15344 | 15344 | 15330 | 4.93 | 485.3 | 264.9 | 264.9 | 0.000011617 | 3.1568E-06 | 0 | 0 | 0.00225047 | 0.00061211 | 0 | 0 | HexNAc(3)Hex(5) |
| a-Lytic (semi) | High | PMI-Byonic | Unambiguous | 786 | 803 | KQIYKTPPIKDFGGFNFS | 801 | HexNAc(3)Hex(6)Fuc(1) | 3 | 6 | 1 | | | [V].KQIYKTPPIKDFGGFnFS.[Q] | N16(HexNAc(3)Hex(6)Fuc(1)) | P0DTC2 | 0 | 4 | 1 | 954.43262 | 3814.70864 | 3814.70901 | -0.1 | -0.00009 | 4669879 | HCD (High Energy Collision Dissociation) | MS2 | 52.83842 | 16.562 | 70.5013 | 15279 | 15279 | 15261 | 5.04 | 351.2 | 184.8 | 184.8 | 9.1651E-06 | 2.7368E-06 | 0 | 0 | 0.00177626 | 0.00053076 | 0 | 0 | HexNAc(3)Hex(6)Fuc(1) |
| a-Lytic (semi) | High | PMI-Byonic | Unambiguous | 786 | 813 | KQIYKTPPIKDFGGFNFSQILPDPSKPS | 801 | HexNAc(3)Hex(6)NeuAc(1) | 3 | 6 | | 1 | | [V].KQIYKTPPIKDFGGFnFSQILPDPSKPS.[K] | N16(HexNAc(3)Hex(6)NeuAc(1)) | P0DTC2 | 2 | 5 | 1 | 1005.47028 | 5023.32227 | 5022.31747 | 200.07 | 0.20096 | 245053.875 | HCD (High Energy Collision Dissociation) | MS2 | 42.45863 | 195.159 | 83.524 | 18631 | 18631 | 18613 | 7.1 | 283.3 | 231.4 | 231.4 | 8.0086E-08 | 5.4682E-08 | 0 | 0 | 0.000015548 | 0.000010616 | 0 | 0 | HexNAc(3)Hex(6)NeuAc(1) |
| a-Lytic (semi) | High | PMI-Byonic | Unambiguous | 786 | 813 | KQIYKTPPIKDFGGFNFSQILPDPSKPS | 801 | HexNAc(3)Hex(6)NeuGc(1) | 3 | 6 | | | 1 | [V].KQIYKTPPIKDFGGFnFSQILPDPSKPS.[K] | N16(HexNAc(3)Hex(6)NeuGc(1)) | P0DTC2 | 2 | 5 | 1 | 1008.46899 | 5038.31586 | 5038.31239 | 0.69 | 0.0007 | 116512.125 | HCD (High Energy Collision Dissociation) | MS2 | 0 | 160.83 | 83.3158 | 18579 | 18579 | 18566 | 7.17 | 457.3 | 361.6 | 361.6 | 6.7805E-08 | 4.347E-08 | 0 | 0 | 0.000013164 | 8.4396E-06 | 0 | 0 | HexNAc(3)Hex(6)NeuGc(1) |
| a-Lytic (semi) | High | PMI-Byonic | Unambiguous | 786 | 803 | KQIYKTPPIKDFGGFNFS | 801 | HexNAc(3)Hex(6) | 3 | 6 | | | | [V].KQIYKTPPIKDFGGFnFS.[Q] | N16(HexNAc(3)Hex(6)) | P0DTC2 | 0 | 4 | 1 | 917.91864 | 3668.65273 | 3668.6511 | 0.45 | 0.00041 | 271849.125 | HCD (High Energy Collision Dissociation) | MS2 | 0 | 149.805 | 70.5148 | 15283 | 15283 | 15261 | 4.74 | 483.6 | 287.3 | 287.3 | 0.000018171 | 4.3134E-06 | 0 | 0 | 0.00351562 | 0.00083597 | 0 | 0 | HexNAc(3)Hex(6) |
| a-Lytic (semi) | High | PMI-Byonic | Unambiguous | 786 | 813 | KQIYKTPPIKDFGGFNFSQILPDPSKPS | 801 | HexNAc(4)Hex(3) | 4 | 3 | | | | [V].KQIYKTPPIKDFGGFnFSQILPDPSKPS.[K] | N16(HexNAc(4)Hex(3)) | P0DTC2 | 2 | 5 | 1 | 890.63483 | 4449.14503 | 4448.14296 | 225.28 | 0.20041 | 177143.719 | HCD (High Energy Collision Dissociation) | MS2 | 0 | 200 | 81.0954 | 17984 | 17984 | 17961 | 5.53 | 183.1 | 176.8 | 176.8 | 0.000002947 | 7.0591E-07 | 0 | 0 | 0.00057182 | 0.000137 | 0 | 0 | HexNAc(4)Hex(3) |
| a-Lytic (semi) | High | PMI-Byonic | Unambiguous | 786 | 813 | KQIYKTPPIKDFGGFNFSQILPDPSKPS | 801 | HexNAc(4)Hex(4)Fuc(1) | 4 | 4 | 1 | | | [V].KQIYKTPPIKDFGGFnFSQILPDPSKPS.[K] | N16(HexNAc(4)Hex(4)Fuc(1)) | P0DTC2 | 2 | 5 | 1 | 952.25775 | 4757.25965 | 4756.25369 | 211.5 | 0.20119 | 472529.188 | HCD (High Energy Collision Dissociation) | MS2 | 9.479612 | 200 | 80.4401 | 17769 | 17769 | 17752 | 6.72 | 250.7 | 197.8 | 197.8 | 1.9035E-07 | 9.5278E-08 | 0 | 0 | 0.000036954 | 0.000018498 | 0 | 0 | HexNAc(4)Hex(4)Fuc(1) |
| a-Lytic (semi) | High | PMI-Byonic | Unambiguous | 786 | 813 | KQIYKTPPIKDFGGFNFSQILPDPSKPS | 801 | HexNAc(4)Hex(5)Fuc(1)NeuAc(1) | 4 | 5 | 1 | 1 | | [V].KQIYKTPPIKDFGGFnFSQILPDPSKPS.[K] | N16(HexNAc(4)Hex(5)Fuc(1)NeuAc(1)) | P0DTC2 | 2 | 4 | 1 | 1303.35962 | 5210.41665 | 5209.40193 | 194.79 | 0.25368 | 121103.773 | HCD (High Energy Collision Dissociation) | MS2 | 12.35386 | 82.657 | 79.7967 | 17577 | 17577 | 17562 | 6.71 | 290 | 218 | 15.4 | 1.9622E-07 | 9.991E-08 | 0 | 0 | 0.000038095 | 0.000019397 | 0 | 0 | HexNAc(4)Hex(5)Fuc(1)NeuAc(1) |
| a-Lytic (semi) | High | PMI-Byonic | Unambiguous | 786 | 813 | KQIYKTPPIKDFGGFNFSQILPDPSKPS | 801 | HexNAc(4)Hex(5)Fuc(1)NeuGc(1) | 4 | 5 | 1 | | 1 | [V].KQIYKTPPIKDFGGFnFSQILPDPSKPS.[K] | N16(HexNAc(4)Hex(5)Fuc(1)NeuGc(1)) | P0DTC2 | 2 | 5 | 1 | 1046.08704 | 5226.40607 | 5225.39685 | 193.14 | 0.20185 | 437755.938 | HCD (High Energy Collision Dissociation) | MS2 | 30.93658 | 183.792 | 82.39 | 18337 | 18337 | 18323 | 6.96 | 402.4 | 328.7 | 26.3 | 1.0941E-07 | 6.8568E-08 | 0 | 0 | 0.000021242 | 0.000013312 | 0 | 0 | HexNAc(4)Hex(5)Fuc(1)NeuGc(1) |
| a-Lytic (semi) | High | PMI-Byonic | Unambiguous | 786 | 813 | KQIYKTPPIKDFGGFNFSQILPDPSKPS | 801 | HexNAc(4)Hex(5)Fuc(1) | 4 | 5 | 1 | | | [V].KQIYKTPPIKDFGGFnFSQILPDPSKPS.[K] | N16(HexNAc(4)Hex(5)Fuc(1)) | P0DTC2 | 2 | 5 | 1 | 984.46606 | 4918.30122 | 4918.30651 | -1.08 | -0.00106 | 359460.656 | HCD (High Energy Collision Dissociation) | MS2 | 30.57154 | 84.048 | 79.935 | 17618 | 17618 | 17601 | 6.93 | 416.2 | 317.1 | 31.6 | 1.1634E-07 | 7.2569E-08 | 0 | 0 | 0.000022587 | 0.000014089 | 0 | 0 | HexNAc(4)Hex(5)Fuc(1) |
| a-Lytic (semi) | High | PMI-Byonic | Unambiguous | 786 | 813 | KQIYKTPPIKDFGGFNFSQILPDPSKPS | 801 | HexNAc(4)Hex(5)Fuc(2)NeuGc(1) | 4 | 5 | 2 | | 1 | [V].KQIYKTPPIKDFGGFnFSQILPDPSKPS.[K] | N16(HexNAc(4)Hex(5)Fuc(2)NeuGc(1)) | P0DTC2 | 2 | 5 | 1 | 1075.29382 | 5372.44001 | 5371.45475 | 183.42 | 0.19705 | 205432.313 | HCD (High Energy Collision Dissociation) | MS2 | 0 | 138.31 | 82.3634 | 18329 | 18329 | 18310 | 4.34 | 173.3 | 161.5 | 33.2 | 0.000045229 | 8.9385E-06 | 0 | 0 | 0.00870547 | 0.0017549 | 0 | 0 | HexNAc(4)Hex(5)Fuc(2)NeuGc(1) |
| a-Lytic (semi) | High | PMI-Byonic | Unambiguous | 786 | 813 | KQIYKTPPIKDFGGFNFSQILPDPSKPS | 801 | HexNAc(4)Hex(5)Fuc(2) | 4 | 5 | 2 | | | [V].KQIYKTPPIKDFGGFnFSQILPDPSKPS.[K] | N16(HexNAc(4)Hex(5)Fuc(2)) | P0DTC2 | 2 | 5 | 1 | 1013.67834 | 5064.36262 | 5064.36442 | -0.36 | -0.00036 | 299526.906 | HCD (High Energy Collision Dissociation) | MS2 | 63.05309 | 82.735 | 79.7923 | 17576 | 17576 | 17562 | 7.13 | 398 | 310.7 | 13 | 7.4755E-08 | 4.7865E-08 | 0 | 0 | 0.000014513 | 9.2927E-06 | 0 | 0 | HexNAc(4)Hex(5)Fuc(2) |
| a-Lytic (semi) | High | PMI-Byonic | Unambiguous | 786 | 813 | KQIYKTPPIKDFGGFNFSQILPDPSKPS | 801 | HexNAc(4)Hex(5)Fuc(3) | 4 | 5 | 3 | | | [V].KQIYKTPPIKDFGGFnFSQILPDPSKPS.[K] | N16(HexNAc(4)Hex(5)Fuc(3)) | P0DTC2 | 2 | 4 | 1 | 1303.6106 | 5211.42055 | 5210.42233 | 191.58 | 0.24956 | 664325.688 | HCD (High Energy Collision Dissociation) | MS2 | 0 | 115.641 | 79.9939 | 17637 | 17637 | 17616 | 6.99 | 303.1 | 228.2 | 228.2 | 1.0214E-07 | 6.4394E-08 | 0 | 0 | 0.000019831 | 0.000012502 | 0 | 0 | HexNAc(4)Hex(5)Fuc(3) |
| a-Lytic (semi) | High | PMI-Byonic | Unambiguous | 786 | 813 | KQIYKTPPIKDFGGFNFSQILPDPSKPS | 801 | HexNAc(4)Hex(5)NeuGc(1) | 4 | 5 | | | 1 | [V].KQIYKTPPIKDFGGFnFSQILPDPSKPS.[K] | N16(HexNAc(4)Hex(5)NeuGc(1)) | P0DTC2 | 2 | 5 | 1 | 1016.67352 | 5079.33851 | 5079.33894 | -0.08 | -0.00009 | 118240.734 | HCD (High Energy Collision Dissociation) | MS2 | 30.16124 | 173.851 | 83.3568 | 18590 | 18590 | 18577 | 7.1 | 291 | 256.7 | 43.7 | 7.9047E-08 | 5.1507E-08 | 0 | 0 | 0.000015347 | 9.9998E-06 | 0 | 0 | HexNAc(4)Hex(5)NeuGc(1) |
| a-Lytic (semi) | High | PMI-Byonic | Unambiguous | 786 | 813 | KQIYKTPPIKDFGGFNFSQILPDPSKPS | 801 | HexNAc(4)Hex(5) | 4 | 5 | | | | [V].KQIYKTPPIKDFGGFnFSQILPDPSKPS.[K] | N16(HexNAc(4)Hex(5)) | P0DTC2 | 2 | 5 | 1 | 955.25671 | 4772.25446 | 4772.2486 | 1.23 | 0.00117 | 150871.281 | HCD (High Energy Collision Dissociation) | MS2 | 19.8466 | 145.343 | 80.4671 | 17777 | 17777 | 17752 | 7.05 | 333.4 | 307.7 | 307.7 | 8.9892E-08 | 5.7881E-08 | 0 | 0 | 0.000017452 | 0.000011237 | 0 | 0 | HexNAc(4)Hex(5) |
| a-Lytic (semi) | High | PMI-Byonic | Unambiguous | 786 | 813 | KQIYKTPPIKDFGGFNFSQILPDPSKPS | 801 | HexNAc(4)Hex(6)Fuc(1) | 4 | 6 | 1 | | | [V].KQIYKTPPIKDFGGFnFSQILPDPSKPS.[K] | N16(HexNAc(4)Hex(6)Fuc(1)) | P0DTC2 | 2 | 5 | 1 | 1017.08026 | 5081.3722 | 5080.35934 | 199.37 | 0.20257 | 433511.063 | HCD (High Energy Collision Dissociation) | MS2 | 20.76618 | 187.092 | 80.7843 | 17887 | 17887 | 17859 | 6.74 | 369.4 | 355.9 | 355.9 | 1.8199E-07 | 9.0597E-08 | 0 | 0 | 0.000035332 | 0.000017589 | 0 | 0 | HexNAc(4)Hex(6)Fuc(1) |
| a-Lytic (semi) | High | PMI-Byonic | Unambiguous | 786 | 813 | KQIYKTPPIKDFGGFNFSQILPDPSKPS | 801 | HexNAc(4)Hex(6)Fuc(2) | 4 | 6 | 2 | | | [V].KQIYKTPPIKDFGGFnFSQILPDPSKPS.[K] | N16(HexNAc(4)Hex(6)Fuc(2)) | P0DTC2 | 2 | 5 | 1 | 1046.28967 | 5227.41926 | 5226.41725 | 191.72 | 0.2004 | 401105.469 | HCD (High Energy Collision Dissociation) | MS2 | 72.46484 | 147.814 | 80.8966 | 17925 | 17925 | 17895 | 5.55 | 137.5 | 137.5 | 137.5 | 2.8314E-06 | 5.9266E-07 | 0 | 0 | 0.00054942 | 0.00011502 | 0 | 0 | HexNAc(4)Hex(6)Fuc(2) |
| a-Lytic (semi) | High | PMI-Byonic | Unambiguous | 786 | 813 | KQIYKTPPIKDFGGFNFSQILPDPSKPS | 801 | HexNAc(4)Hex(6)NeuAc(1) | 4 | 6 | | 1 | | [V].KQIYKTPPIKDFGGFnFSQILPDPSKPS.[K] | N16(HexNAc(4)Hex(6)NeuAc(1)) | P0DTC2 | 2 | 5 | 1 | 1045.88635 | 5225.40266 | 5225.39685 | 1.11 | 0.00116 | 128788.906 | HCD (High Energy Collision Dissociation) | MS2 | 40.77452 | 200 | 83.2265 | 18559 | 18559 | 18540 | 6.71 | 243.6 | 243.6 | 26.5 | 1.9468E-07 | 9.6856E-08 | 0 | 0 | 0.000037795 | 0.000018804 | 0 | 0 | HexNAc(4)Hex(6)NeuAc(1) |
| a-Lytic (semi) | High | PMI-Byonic | Unambiguous | 786 | 813 | KQIYKTPPIKDFGGFNFSQILPDPSKPS | 801 | HexNAc(5)Hex(3)Fuc(1)NeuAc(1) | 5 | 3 | 1 | 1 | | [V].KQIYKTPPIKDFGGFnFSQILPDPSKPS.[K] | N16(HexNAc(5)Hex(3)Fuc(1)NeuAc(1)) | P0DTC2 | 2 | 4 | 1 | 1273.10107 | 5089.38247 | 5088.37566 | 197.86 | 0.2517 | 77553.6875 | HCD (High Energy Collision Dissociation) | MS2 | 11.69677 | 200 | 85.8931 | 19134 | 19134 | 19120 | 3.67 | 0.5 | 0.5 | 0.5 | 0.00021471 | 0.000029413 | 0 | 0 | 0.04003386 | 0.00990099 | 0.00955414 | 0.0108108 | HexNAc(5)Hex(3)Fuc(1)NeuAc(1) |
| a-Lytic (semi) | High | PMI-Byonic | Unambiguous | 786 | 813 | KQIYKTPPIKDFGGFNFSQILPDPSKPS | 801 | HexNAc(5)Hex(3)Fuc(1) | 5 | 3 | 1 | | | [V].KQIYKTPPIKDFGGFnFSQILPDPSKPS.[K] | N16(HexNAc(5)Hex(3)Fuc(1)) | P0DTC2 | 2 | 4 | 1 | 1200.32446 | 4798.27602 | 4797.28024 | 207.57 | 0.24895 | 269885.75 | HCD (High Energy Collision Dissociation) | MS2 | 0 | 200 | 80.1878 | 17696 | 17696 | 17674 | 6.73 | 294.2 | 178.8 | 178.8 | 1.8719E-07 | 9.372E-08 | 0 | 0 | 0.000036341 | 0.000018195 | 0 | 0 | HexNAc(5)Hex(3)Fuc(1) |
| a-Lytic (semi) | High | PMI-Byonic | Unambiguous | 786 | 813 | KQIYKTPPIKDFGGFNFSQILPDPSKPS | 801 | HexNAc(5)Hex(4)Fuc(1) | 5 | 4 | 1 | | | [V].KQIYKTPPIKDFGGFnFSQILPDPSKPS.[K] | N16(HexNAc(5)Hex(4)Fuc(1)) | P0DTC2 | 2 | 5 | 1 | 992.67194 | 4959.33057 | 4959.33306 | -0.5 | -0.0005 | 428256.188 | HCD (High Energy Collision Dissociation) | MS2 | 12.51172 | 107.341 | 79.9799 | 17632 | 17632 | 17616 | 6.86 | 385.7 | 250 | 250 | 1.3776E-07 | 8.0921E-08 | 0 | 0 | 0.000026745 | 0.00001571 | 0 | 0 | HexNAc(5)Hex(4)Fuc(1) |
| a-Lytic (semi) | High | PMI-Byonic | Unambiguous | 786 | 813 | KQIYKTPPIKDFGGFNFSQILPDPSKPS | 801 | HexNAc(5)Hex(4)Fuc(2) | 5 | 4 | 2 | | | [V].KQIYKTPPIKDFGGFnFSQILPDPSKPS.[K] | N16(HexNAc(5)Hex(4)Fuc(2)) | P0DTC2 | 2 | 5 | 1 | 1022.08386 | 5106.39021 | 5105.39097 | 195.72 | 0.19985 | 394890.531 | HCD (High Energy Collision Dissociation) | MS2 | 0 | 164.429 | 79.9421 | 17621 | 17621 | 17601 | 6.81 | 354.4 | 252.3 | 252.3 | 1.5434E-07 | 8.4604E-08 | 0 | 0 | 0.000029964 | 0.000016425 | 0 | 0 | HexNAc(5)Hex(4)Fuc(2) |
| a-Lytic (semi) | High | PMI-Byonic | Unambiguous | 786 | 813 | KQIYKTPPIKDFGGFNFSQILPDPSKPS | 801 | HexNAc(5)Hex(4) | 5 | 4 | | | | [V].KQIYKTPPIKDFGGFnFSQILPDPSKPS.[K] | N16(HexNAc(5)Hex(4)) | P0DTC2 | 2 | 4 | 1 | 1204.32739 | 4814.28774 | 4813.27515 | 210.37 | 0.25315 | 182207.281 | HCD (High Energy Collision Dissociation) | MS2 | 18.19911 | 200 | 80.6062 | 17821 | 17821 | 17795 | 6.54 | 411.3 | 285.2 | 285.2 | 2.8617E-07 | 1.1223E-07 | 0 | 0 | 0.000055556 | 0.000021788 | 0 | 0 | HexNAc(5)Hex(4) |
| a-Lytic (semi) | High | PMI-Byonic | Unambiguous | 786 | 813 | KQIYKTPPIKDFGGFNFSQILPDPSKPS | 801 | HexNAc(5)Hex(5)Fuc(1) | 5 | 5 | 1 | | | [V].KQIYKTPPIKDFGGFnFSQILPDPSKPS.[K] | N16(HexNAc(5)Hex(5)Fuc(1)) | P0DTC2 | 2 | 5 | 1 | 1025.08655 | 5121.40363 | 5121.38589 | 3.47 | 0.00355 | 365465.563 | HCD (High Energy Collision Dissociation) | MS2 | 95.56812 | 41.614 | 80.6261 | 17827 | 17827 | 17812 | 6.02 | 304.3 | 304.3 | 304.3 | 9.5758E-07 | 1.5115E-07 | 0 | 0 | 0.00018588 | 0.000029344 | 0 | 0 | HexNAc(5)Hex(5)Fuc(1) |
| a-Lytic (semi) | High | PMI-Byonic | Unambiguous | 786 | 813 | KQIYKTPPIKDFGGFNFSQILPDPSKPS | 801 | HexNAc(5)Hex(5)Fuc(2) | 5 | 5 | 2 | | | [V].KQIYKTPPIKDFGGFnFSQILPDPSKPS.[K] | N16(HexNAc(5)Hex(5)Fuc(2)) | P0DTC2 | 2 | 4 | 1 | 1317.86755 | 5268.44838 | 5267.4438 | 190.72 | 0.25115 | 283868.875 | HCD (High Energy Collision Dissociation) | MS2 | 21.90827 | 83.575 | 80.5969 | 17818 | 17818 | 17795 | 5.68 | 216.4 | 154.1 | 154.1 | 2.0997E-06 | 3.2787E-07 | 0 | 0 | 0.00040749 | 0.00006364 | 0 | 0 | HexNAc(5)Hex(5)Fuc(2) |
| a-Lytic (semi) | High | PMI-Byonic | Unambiguous | 786 | 813 | KQIYKTPPIKDFGGFNFSQILPDPSKPS | 801 | HexNAc(5)Hex(5)Fuc(3) | 5 | 5 | 3 | | | [V].KQIYKTPPIKDFGGFnFSQILPDPSKPS.[K] | N16(HexNAc(5)Hex(5)Fuc(3)) | P0DTC2 | 2 | 4 | 1 | 1354.37903 | 5414.49428 | 5413.5017 | 183.35 | 0.24814 | 353850.125 | HCD (High Energy Collision Dissociation) | MS2 | 0 | 200 | 79.7486 | 17564 | 17564 | 17549 | 6.59 | 315.4 | 188.9 | 188.9 | 2.5818E-07 | 1.0231E-07 | 0 | 0 | 0.000050123 | 0.000019862 | 0 | 0 | HexNAc(5)Hex(5)Fuc(3) |
| a-Lytic (semi) | High | PMI-Byonic | Unambiguous | 786 | 813 | KQIYKTPPIKDFGGFNFSQILPDPSKPS | 801 | HexNAc(5)Hex(5)NeuAc(1) | 5 | 5 | | 1 | | [V].KQIYKTPPIKDFGGFnFSQILPDPSKPS.[K] | N16(HexNAc(5)Hex(5)NeuAc(1)) | P0DTC2 | 2 | 5 | 1 | 1054.28821 | 5267.41193 | 5266.42339 | 187.71 | 0.19771 | 98182.5781 | HCD (High Energy Collision Dissociation) | MS2 | 0 | 141.213 | 80.4007 | 17757 | 17757 | 17740 | 5.88 | 296.5 | 231.4 | 33.9 | 1.3227E-06 | 1.9172E-07 | 0 | 0 | 0.00025673 | 0.000037219 | 0 | 0 | HexNAc(5)Hex(5)NeuAc(1) |
| a-Lytic (semi) | High | PMI-Byonic | Unambiguous | 786 | 813 | KQIYKTPPIKDFGGFNFSQILPDPSKPS | 801 | HexNAc(5)Hex(5) | 5 | 5 | | | | [V].KQIYKTPPIKDFGGFnFSQILPDPSKPS.[K] | N16(HexNAc(5)Hex(5)) | P0DTC2 | 2 | 5 | 1 | 995.87225 | 4975.33216 | 4975.32798 | 0.84 | 0.00084 | 437098.688 | HCD (High Energy Collision Dissociation) | MS2 | 21.43268 | 28.573 | 80.6717 | 17843 | 17843 | 17825 | 6.99 | 253.7 | 217.6 | 217.6 | 1.0286E-07 | 6.5434E-08 | 0 | 0 | 0.00001997 | 0.000012704 | 0 | 0 | HexNAc(5)Hex(5) |
| a-Lytic (semi) | High | PMI-Byonic | Unambiguous | 786 | 813 | KQIYKTPPIKDFGGFNFSQILPDPSKPS | 801 | HexNAc(5)Hex(6)Fuc(1)NeuGc(1) | 5 | 6 | 1 | | 1 | [V].KQIYKTPPIKDFGGFnFSQILPDPSKPS.[K] | N16(HexNAc(5)Hex(6)Fuc(1)NeuGc(1)) | P0DTC2 | 2 | 5 | 1 | 1119.11365 | 5591.53913 | 5590.52904 | 180.68 | 0.20202 | 156347.984 | HCD (High Energy Collision Dissociation) | MS2 | 28.42833 | 69.275 | 82.5325 | 18377 | 18377 | 18362 | 4.88 | 67.3 | 67.3 | 67.3 | 0.000013242 | 3.6981E-06 | 0 | 0 | 0.00256446 | 0.00071691 | 0 | 0 | HexNAc(5)Hex(6)Fuc(1)NeuGc(1) |
| a-Lytic (semi) | High | PMI-Byonic | Unambiguous | 786 | 813 | KQIYKTPPIKDFGGFNFSQILPDPSKPS | 801 | HexNAc(5)Hex(6)Fuc(1) | 5 | 6 | 1 | | | [V].KQIYKTPPIKDFGGFnFSQILPDPSKPS.[K] | N16(HexNAc(5)Hex(6)Fuc(1)) | P0DTC2 | 2 | 5 | 1 | 1057.49255 | 5283.43366 | 5283.43871 | -0.96 | -0.00101 | 151058.141 | HCD (High Energy Collision Dissociation) | MS2 | 34.68391 | 200 | 79.716 | 17555 | 17555 | 17536 | 6.86 | 299.5 | 240.1 | 29.2 | 1.3775E-07 | 7.9806E-08 | 0 | 0 | 0.000026744 | 0.000015494 | 0 | 0 | HexNAc(5)Hex(6)Fuc(1) |
| a-Lytic (semi) | High | PMI-Byonic | Unambiguous | 786 | 813 | KQIYKTPPIKDFGGFNFSQILPDPSKPS | 801 | HexNAc(5)Hex(6)Fuc(2) | 5 | 6 | 2 | | | [V].KQIYKTPPIKDFGGFnFSQILPDPSKPS.[K] | N16(HexNAc(5)Hex(6)Fuc(2)) | P0DTC2 | 2 | 5 | 1 | 1086.90613 | 5430.50153 | 5429.49662 | 185.08 | 0.20098 | 196732.531 | HCD (High Energy Collision Dissociation) | MS2 | 29.28654 | 200 | 79.656 | 17541 | 17541 | 17523 | 7.1 | 282 | 226.2 | 226.2 | 8.0086E-08 | 5.265E-08 | 0 | 0 | 0.000015548 | 0.000010222 | 0 | 0 | HexNAc(5)Hex(6)Fuc(2) |
| a-Lytic (semi) | High | PMI-Byonic | Unambiguous | 786 | 813 | KQIYKTPPIKDFGGFNFSQILPDPSKPS | 801 | HexNAc(5)Hex(6)NeuAc(1) | 5 | 6 | | 1 | | [V].KQIYKTPPIKDFGGFnFSQILPDPSKPS.[K] | N16(HexNAc(5)Hex(6)NeuAc(1)) | P0DTC2 | 2 | 5 | 1 | 1086.69995 | 5429.47065 | 5428.47622 | 183.19 | 0.19889 | 179884.453 | HCD (High Energy Collision Dissociation) | MS2 | 8.508872 | 158.761 | 82.6536 | 18410 | 18410 | 18388 | 4.63 | 149.7 | 104.6 | 36 | 0.000023427 | 5.2619E-06 | 0 | 0 | 0.0045279 | 0.00101925 | 0 | 0 | HexNAc(5)Hex(6)NeuAc(1) |
| a-Lytic (semi) | High | PMI-Byonic | Unambiguous | 786 | 813 | KQIYKTPPIKDFGGFNFSQILPDPSKPS | 801 | HexNAc(6)Hex(5)Fuc(1) | 6 | 5 | 1 | | | [V].KQIYKTPPIKDFGGFnFSQILPDPSKPS.[K] | N16(HexNAc(6)Hex(5)Fuc(1)) | P0DTC2 | 2 | 5 | 1 | 1065.90051 | 5325.47346 | 5324.46526 | 189.35 | 0.20164 | 193525 | HCD (High Energy Collision Dissociation) | MS2 | 39.15903 | 146.627 | 80.3342 | 17738 | 17738 | 17714 | 4.52 | 89.8 | 89.8 | 89.8 | 0.000030138 | 6.0517E-06 | 0 | 0 | 0.00581747 | 0.00117167 | 0 | 0 | HexNAc(6)Hex(5)Fuc(1) |
| a-Lytic (semi) | High | PMI-Byonic | Unambiguous | 786 | 813 | KQIYKTPPIKDFGGFNFSQILPDPSKPS | 801 | HexNAc(6)Hex(6)Fuc(1) | 6 | 6 | 1 | | | [V].KQIYKTPPIKDFGGFnFSQILPDPSKPS.[K] | N16(HexNAc(6)Hex(6)Fuc(1)) | P0DTC2 | 2 | 5 | 1 | 1098.11597 | 5486.55073 | 5486.51808 | 5.95 | 0.00653 | 182176.156 | HCD (High Energy Collision Dissociation) | MS2 | 19.5441 | 57.997 | 79.9453 | 17622 | 17622 | 17601 | 3.78 | 149.3 | 149.3 | 149.3 | 0.00016539 | 0.000025558 | 0 | 0 | 0.03112175 | 0.00990099 | 0.00977198 | 0 | HexNAc(6)Hex(6)Fuc(1) |
| a-Lytic (semi) | High | PMI-Byonic | Unambiguous | 786 | 813 | KQIYKTPPIKDFGGFNFSQILPDPSKPS | 801 | HexNAc(6)Hex(6)Fuc(3) | 6 | 6 | 3 | | | [V].KQIYKTPPIKDFGGFnFSQILPDPSKPS.[K] | N16(HexNAc(6)Hex(6)Fuc(3)) | P0DTC2 | 2 | 5 | 1 | 1156.73193 | 5779.63056 | 5778.6339 | 172.47 | 0.19933 | 333497.438 | HCD (High Energy Collision Dissociation) | MS2 | 0 | 103.964 | 79.9078 | 17610 | 17610 | 17588 | 5.31 | 170.1 | 153.7 | 153.7 | 4.8692E-06 | 1.3771E-06 | 0 | 0 | 0.00094446 | 0.0002672 | 0 | 0 | HexNAc(6)Hex(6)Fuc(3) |
| a-Lytic (semi) | High | PMI-Byonic | Unambiguous | 786 | 813 | KQIYKTPPIKDFGGFNFSQILPDPSKPS | 801 | HexNAc(6)Hex(6) | 6 | 6 | | | | [V].KQIYKTPPIKDFGGFnFSQILPDPSKPS.[K] | N16(HexNAc(6)Hex(6)) | P0DTC2 | 2 | 5 | 1 | 1069.09753 | 5341.45856 | 5340.46017 | 186.95 | 0.19968 | 497463 | HCD (High Energy Collision Dissociation) | MS2 | 10.27753 | 90.069 | 80.5414 | 17801 | 17801 | 17780 | 5.31 | 203.2 | 152.8 | 152.8 | 4.8692E-06 | 1.4036E-06 | 0 | 0 | 0.00094446 | 0.00027233 | 0 | 0 | HexNAc(6)Hex(6) |
| a-Lytic (semi) | High | PMI-Byonic | Unambiguous | 1071 | 1078 | QEKNFTTA | 1074 | HexNAc(2)Hex(5) | 2 | 5 | | | | [A].QEKnFTTA.[P] | N4(HexNAc(2)Hex(5)) | P0DTC2 | 2 | 2 | 1 | 1077.94299 | 2154.87871 | 2154.88065 | -0.9 | -0.00097 | 515752.25 | HCD (High Energy Collision Dissociation) | MS2 | 0 | 179.644 | 33.4271 | 5953 | 5953 | 5939 | 5.3 | 311.2 | 183.4 | 183.4 | 5.0374E-06 | 1.5073E-06 | 0 | 0 | 0.00097706 | 0.00029244 | 0 | 0 | HexNAc(2)Hex(5) |
| a-Lytic (semi) | High | PMI-Byonic | Unambiguous | 1071 | 1078 | QEKNFTTA | 1074 | HexNAc(2)Hex(6) | 2 | 6 | | | | [A].QEKnFTTA.[P] | N4(HexNAc(2)Hex(6)) | P0DTC2 | 2 | 2 | 1 | 1158.9707 | 2316.93413 | 2316.93347 | 0.28 | 0.00033 | 263108.313 | HCD (High Energy Collision Dissociation) | MS2 | 0 | 200 | 32.8943 | 5811 | 5811 | 5792 | 5.71 | 417.9 | 233.1 | 233.1 | 1.9486E-06 | 2.6576E-07 | 0 | 0 | 0.00037818 | 0.000051588 | 0 | 0 | HexNAc(2)Hex(6) |
| a-Lytic (semi) | High | PMI-Byonic | Unambiguous | 1071 | 1080 | QEKNFTTAPA | 1074 | HexNAc(2)Hex(7) | 2 | 7 | | | | [A].QEKnFTTAPA.[I] | N4(HexNAc(2)Hex(7)) | P0DTC2 | 2 | 2 | 1 | 1324.04211 | 2647.07695 | 2647.07617 | 0.29 | 0.00039 | 321597.656 | HCD (High Energy Collision Dissociation) | MS2 | 6.714626 | 200 | 37.3414 | 6987 | 6987 | 6970 | 3.54 | 111.6 | 59.6 | 59.6 | 0.00028833 | 0.000036489 | 0 | 0 | 0.05304038 | 0.01273885 | 0.01234568 | 0.01546391 | HexNAc(2)Hex(7) |
| a-Lytic (semi) | High | PMI-Byonic | Unambiguous | 1071 | 1080 | QEKNFTTAPA | 1074 | HexNAc(2)Hex(8) | 2 | 8 | | | | [A].QEKnFTTAPA.[I] | N4(HexNAc(2)Hex(8)) | P0DTC2 | 2 | 2 | 1 | 1405.06873 | 2809.13017 | 2809.129 | 0.42 | 0.00059 | 212497.891 | HCD (High Energy Collision Dissociation) | MS2 | 0 | 200 | 36.9271 | 6885 | 6885 | 6866 | 3.94 | 200.2 | 127.2 | 127.2 | 0.00011352 | 0.000019448 | 0 | 0 | 0.02156945 | 0.00706714 | 0.00682594 | 0.01156069 | HexNAc(2)Hex(8) |
| a-Lytic (semi) | High | PMI-Byonic | Unambiguous | 1071 | 1080 | QEKNFTTAPA | 1074 | HexNAc(3)Hex(5)Fuc(1) | 3 | 5 | 1 | | | [A].QEKnFTTAPA.[I] | N4(HexNAc(3)Hex(5)Fuc(1)) | P0DTC2 | 2 | 2 | 1 | 1336.55847 | 2672.10967 | 2672.10781 | 0.7 | 0.00093 | 243440.984 | HCD (High Energy Collision Dissociation) | MS2 | 0 | 200 | 38.2392 | 7210 | 7210 | 7187 | 3.62 | 218 | 101.8 | 101.8 | 0.00024041 | 0.000031313 | 0 | 0 | 0.04461461 | 0.01273885 | 0.01261829 | 0.01595744 | HexNAc(3)Hex(5)Fuc(1) |
| a-Lytic (semi) | High | PMI-Byonic | Unambiguous | 1071 | 1080 | QEKNFTTAPA | 1074 | HexNAc(3)Hex(6)Fuc(1) | 3 | 6 | 1 | | | [A].QEKnFTTAPA.[I] | N4(HexNAc(3)Hex(6)Fuc(1)) | P0DTC2 | 2 | 2 | 1 | 1417.58154 | 2834.15581 | 2834.16063 | -1.7 | -0.00241 | 190736.047 | HCD (High Energy Collision Dissociation) | MS2 | 0 | 200 | 37.1527 | 6942 | 6942 | 6924 | 2.86 | 149.4 | 57 | 57 | 0.0013662 | 0.00017388 | 0 | 0 | 0.21057055 | 0.03846153 | 0.03828828 | 0.0516605 | HexNAc(3)Hex(6)Fuc(1) |
| a-Lytic (semi) | High | PMI-Byonic | Unambiguous | 1098 | 1105 | NGTHWFVT | 1098 | HexNAc(2)Hex(5) | 2 | 5 | | | | [S].nGTHWFVT.[Q] | N1(HexNAc(2)Hex(5)) | P0DTC2 | 2 | 2 | 1 | 1089.44214 | 2177.877 | 2177.8755 | 0.69 | 0.00075 | 150786.891 | HCD (High Energy Collision Dissociation) | MS2 | 0 | 200 | 59.8588 | 12564 | 12564 | 12541 | 5.37 | 255.7 | 118.3 | 118.3 | 4.2334E-06 | 1.2208E-06 | 0 | 0 | 0.00082123 | 0.00023687 | 0 | 0 | HexNAc(2)Hex(5) |
| a-Lytic (semi) | High | PMI-Byonic | Unambiguous | 1098 | 1105 | NGTHWFVT | 1098 | HexNAc(2)Hex(6) | 2 | 6 | | | | [S].nGTHWFVT.[Q] | N1(HexNAc(2)Hex(6)) | P0DTC2 | 2 | 2 | 1 | 1170.46692 | 2339.92656 | 2339.92833 | -0.75 | -0.00088 | 198633.188 | HCD (High Energy Collision Dissociation) | MS2 | 9.61342 | 200 | 59.1527 | 12404 | 12404 | 12393 | 5.3 | 318.6 | 178.6 | 178.6 | 5.0374E-06 | 0.000001533 | 0 | 0 | 0.00097706 | 0.00029744 | 0 | 0 | HexNAc(2)Hex(6) |
| a-Lytic (semi) | High | PMI-Byonic | Unambiguous | 1098 | 1105 | NGTHWFVT | 1098 | HexNAc(2)Hex(7) | 2 | 7 | | | | [S].nGTHWFVT.[Q] | N1(HexNAc(2)Hex(7)) | P0DTC2 | 2 | 2 | 1 | 1251.49158 | 2501.97588 | 2501.98115 | -2.11 | -0.00264 | 136961.438 | HCD (High Energy Collision Dissociation) | MS2 | 0 | 200 | 58.6508 | 12295 | 12295 | 12278 | 5.08 | 327.2 | 179.9 | 179.9 | 0.00000833 | 2.4071E-06 | 0 | 0 | 0.00161468 | 0.00046687 | 0 | 0 | HexNAc(2)Hex(7) |
| a-Lytic (semi) | High | PMI-Byonic | Unambiguous | 1098 | 1105 | NGTHWFVT | 1098 | HexNAc(2)Hex(8) | 2 | 8 | | | | [S].nGTHWFVT.[Q] | N1(HexNAc(2)Hex(8)) | P0DTC2 | 2 | 2 | 1 | 1332.51953 | 2664.03179 | 2664.03397 | -0.82 | -0.00109 | 145310.813 | HCD (High Energy Collision Dissociation) | MS2 | 0 | 200 | 58.5156 | 12263 | 12263 | 12245 | 5.5 | 337.4 | 193.1 | 193.1 | 3.1468E-06 | 8.3617E-07 | 0 | 0 | 0.00061057 | 0.00016227 | 0 | 0 | HexNAc(2)Hex(8) |
| a-Lytic (semi) | High | PMI-Byonic | Unambiguous | 1098 | 1105 | NGTHWFVT | 1098 | HexNAc(2)Hex(9) | 2 | 9 | | | | [S].nGTHWFVT.[Q] | N1(HexNAc(2)Hex(9)) | P0DTC2 | 2 | 2 | 1 | 1413.54614 | 2826.08501 | 2826.0868 | -0.63 | -0.00089 | 136778.938 | HCD (High Energy Collision Dissociation) | MS2 | 0 | 200 | 58.4156 | 12239 | 12239 | 12221 | 3.25 | 215.2 | 52.5 | 52.5 | 0.00056054 | 0.000092062 | 0 | 0 | 0.09870876 | 0.02168021 | 0.02072538 | 0.0301724 | HexNAc(2)Hex(9) |
| a-Lytic (semi) | High | PMI-Byonic | Unambiguous | 1098 | 1105 | NGTHWFVT | 1098 | HexNAc(3)Hex(5)Fuc(1) | 3 | 5 | 1 | | | [S].nGTHWFVT.[Q] | N1(HexNAc(3)Hex(5)Fuc(1)) | P0DTC2 | 2 | 2 | 1 | 1264.01074 | 2527.01421 | 2527.01279 | 0.56 | 0.00071 | 145661.781 | HCD (High Energy Collision Dissociation) | MS2 | 0 | 200 | 58.887 | 12347 | 12347 | 12329 | 5.6 | 295.8 | 162 | 162 | 2.5272E-06 | 4.5363E-07 | 0 | 0 | 0.00049042 | 0.000088046 | 0 | 0 | HexNAc(3)Hex(5)Fuc(1) |
| a-Lytic (semi) | High | PMI-Byonic | Unambiguous | 1098 | 1105 | NGTHWFVT | 1098 | HexNAc(3)Hex(5)NeuGc(1) | 3 | 5 | | | 1 | [S].nGTHWFVT.[Q] | N1(HexNAc(3)Hex(5)NeuGc(1)) | P0DTC2 | 2 | 3 | 1 | 896.68689 | 2688.04612 | 2688.04521 | 0.34 | 0.0003 | 187519.844 | HCD (High Energy Collision Dissociation) | MS2 | 14.33206 | 200 | 63.5783 | 13596 | 13596 | 13574 | 4.84 | 294.8 | 97.6 | 97.6 | 0.000014598 | 3.9548E-06 | 0 | 0 | 0.00282617 | 0.00076659 | 0 | 0 | HexNAc(3)Hex(5)NeuGc(1) |
| a-Lytic (semi) | High | PMI-Byonic | Unambiguous | 1098 | 1105 | NGTHWFVT | 1098 | HexNAc(3)Hex(5) | 3 | 5 | | | | [S].nGTHWFVT.[Q] | N1(HexNAc(3)Hex(5)) | P0DTC2 | 2 | 2 | 1 | 1190.98108 | 2380.95488 | 2380.95488 | 0 | 0 | 186576.75 | HCD (High Energy Collision Dissociation) | MS2 | 0 | 175.582 | 59.2499 | 12427 | 12427 | 12413 | 5.75 | 292.3 | 140.9 | 140.9 | 1.7777E-06 | 2.2542E-07 | 0 | 0 | 0.00034503 | 0.00004376 | 0 | 0 | HexNAc(3)Hex(5) |
| a-Lytic (semi) | High | PMI-Byonic | Unambiguous | 1098 | 1105 | NGTHWFVT | 1098 | HexNAc(3)Hex(6)Fuc(1) | 3 | 6 | 1 | | | [S].nGTHWFVT.[Q] | N1(HexNAc(3)Hex(6)Fuc(1)) | P0DTC2 | 2 | 3 | 1 | 897.02783 | 2689.06894 | 2689.06561 | 1.24 | 0.00111 | 222229.563 | HCD (High Energy Collision Dissociation) | MS2 | 0 | 200 | 58.5618 | 12273 | 12273 | 12258 | 5.2 | 364.1 | 202.6 | 202.6 | 6.2913E-06 | 1.6582E-06 | 0 | 0 | 0.00121996 | 0.0003217 | 0 | 0 | HexNAc(3)Hex(6)Fuc(1) |
| a-Lytic (semi) | High | PMI-Byonic | Unambiguous | 1098 | 1105 | NGTHWFVT | 1098 | HexNAc(3)Hex(6)NeuAc(1) | 3 | 6 | | 1 | | [S].nGTHWFVT.[Q] | N1(HexNAc(3)Hex(6)NeuAc(1)) | P0DTC2 | 2 | 3 | 1 | 945.37305 | 2834.10459 | 2834.10312 | 0.52 | 0.00049 | 137569.203 | HCD (High Energy Collision Dissociation) | MS2 | 0 | 200 | 63.433 | 13558 | 13558 | 13535 | 1.25 | 221.5 | 25.6 | 25.6 | 0.05643664 | 0.00754717 | 0.00747663 | 0.009375 | 0.91939198 | 0.16366609 | 0.16313211 | 0.21951214 | HexNAc(3)Hex(6)NeuAc(1) |
| a-Lytic (semi) | High | PMI-Byonic | Unambiguous | 1098 | 1105 | NGTHWFVT | 1098 | HexNAc(3)Hex(6)NeuGc(1) | 3 | 6 | | | 1 | [S].nGTHWFVT.[Q] | N1(HexNAc(3)Hex(6)NeuGc(1)) | P0DTC2 | 2 | 3 | 1 | 950.70465 | 2850.0994 | 2850.09803 | 0.48 | 0.00046 | 351925.188 | HCD (High Energy Collision Dissociation) | MS2 | 59.23533 | 200 | 63.2174 | 13498 | 13498 | 13479 | 4.97 | 322.6 | 155.8 | 155.8 | 0.000010838 | 3.0178E-06 | 0 | 0 | 0.00209983 | 0.0005852 | 0 | 0 | HexNAc(3)Hex(6)NeuGc(1) |
| a-Lytic (semi) | High | PMI-Byonic | Unambiguous | 1098 | 1105 | NGTHWFVT | 1098 | HexNAc(3)Hex(6) | 3 | 6 | | | | [S].nGTHWFVT.[Q] | N1(HexNAc(3)Hex(6)) | P0DTC2 | 2 | 2 | 1 | 1272.00696 | 2543.00664 | 2543.0077 | -0.42 | -0.00053 | 386225.406 | HCD (High Energy Collision Dissociation) | MS2 | 55.90098 | 175.355 | 58.9146 | 12353 | 12353 | 12342 | 5.53 | 319.1 | 184.5 | 184.5 | 2.9293E-06 | 6.6153E-07 | 0 | 0 | 0.00056839 | 0.00012839 | 0 | 0 | HexNAc(3)Hex(6) |
| a-Lytic (semi) | High | PMI-Byonic | Unambiguous | 1098 | 1105 | NGTHWFVT | 1098 | HexNAc(4)Hex(5)Fuc(1) | 4 | 5 | 1 | | | [S].nGTHWFVT.[Q] | N1(HexNAc(4)Hex(5)Fuc(1)) | P0DTC2 | 2 | 2 | 1 | 1365.55066 | 2730.09404 | 2730.09216 | 0.69 | 0.00094 | 287710.875 | HCD (High Energy Collision Dissociation) | MS2 | 0 | 200 | 58.558 | 12272 | 12272 | 12258 | 5.64 | 279.5 | 148.7 | 148.7 | 2.2832E-06 | 4.0865E-07 | 0 | 0 | 0.00044309 | 0.000079318 | 0 | 0 | HexNAc(4)Hex(5)Fuc(1) |
| a-Lytic (semi) | High | PMI-Byonic | Unambiguous | 1098 | 1105 | NGTHWFVT | 1098 | HexNAc(4)Hex(5)Fuc(2) | 4 | 5 | 2 | | | [S].nGTHWFVT.[Q] | N1(HexNAc(4)Hex(5)Fuc(2)) | P0DTC2 | 2 | 2 | 1 | 1438.58105 | 2876.15483 | 2876.15007 | 1.66 | 0.00238 | 140725.922 | HCD (High Energy Collision Dissociation) | MS2 | 0 | 200 | 58.2621 | 12205 | 12205 | 12186 | 4.99 | 272.1 | 121.1 | 121.1 | 0.000010234 | 2.9736E-06 | 0 | 0 | 0.00198304 | 0.00057664 | 0 | 0 | HexNAc(4)Hex(5)Fuc(2) |
| a-Lytic (semi) | High | PMI-Byonic | Unambiguous | 1098 | 1105 | NGTHWFVT | 1098 | HexNAc(4)Hex(5)NeuGc(1) | 4 | 5 | | | 1 | [S].nGTHWFVT.[Q] | N1(HexNAc(4)Hex(5)NeuGc(1)) | P0DTC2 | 2 | 3 | 1 | 964.37842 | 2891.1207 | 2891.12458 | -1.34 | -0.00129 | 310140.781 | HCD (High Energy Collision Dissociation) | MS2 | 0 | 200 | 63.2463 | 13505 | 13505 | 13479 | 4.35 | 246.6 | 84 | 84 | 0.000045127 | 8.7904E-06 | 0 | 0 | 0.00868597 | 0.00172664 | 0 | 0 | HexNAc(4)Hex(5)NeuGc(1) |
| a-Lytic (semi) | High | PMI-Byonic | Unambiguous | 1098 | 1105 | NGTHWFVT | 1098 | HexNAc(4)Hex(5) | 4 | 5 | | | | [S].nGTHWFVT.[Q] | N1(HexNAc(4)Hex(5)) | P0DTC2 | 2 | 2 | 1 | 1292.52197 | 2584.03667 | 2584.03425 | 0.94 | 0.00121 | 225724.891 | HCD (High Energy Collision Dissociation) | MS2 | 22.17236 | 200 | 58.9636 | 12364 | 12364 | 12351 | 5.37 | 262.1 | 125.8 | 125.8 | 4.2334E-06 | 1.1965E-06 | 0 | 0 | 0.00082123 | 0.00023216 | 0 | 0 | HexNAc(4)Hex(5) |
| a-Lytic (semi) | High | PMI-Byonic | Unambiguous | 1098 | 1105 | NGTHWFVT | 1098 | HexNAc(5)Hex(6)Fuc(1)NeuAc(1) | 5 | 6 | 1 | 1 | | [S].nGTHWFVT.[Q] | N1(HexNAc(5)Hex(6)Fuc(1)NeuAc(1)) | P0DTC2 | 2 | 3 | 1 | 1129.44434 | 3386.31845 | 3386.31977 | -0.39 | -0.00044 | 161321.516 | HCD (High Energy Collision Dissociation) | MS2 | 0 | 156.397 | 62.219 | 13197 | 13197 | 13182 | 1.8 | 124 | 10.6 | 10.6 | 0.01571039 | 0.00196078 | 0.00194932 | 0.00321543 | 0.75527269 | 0.13111886 | 0.13089003 | 0.17819144 | HexNAc(5)Hex(6)Fuc(1)NeuAc(1) |
| a-Lytic (semi) | High | PMI-Byonic | Unambiguous | 1098 | 1105 | NGTHWFVT | 1098 | HexNAc(5)Hex(6)Fuc(1)NeuGc(1) | 5 | 6 | 1 | | 1 | [S].nGTHWFVT.[Q] | N1(HexNAc(5)Hex(6)Fuc(1)NeuGc(1)) | P0DTC2 | 2 | 3 | 1 | 1134.77686 | 3402.31601 | 3402.31469 | 0.39 | 0.00044 | 208474.875 | HCD (High Energy Collision Dissociation) | MS2 | 26.30257 | 200 | 61.8385 | 13096 | 13096 | 13078 | 5.06 | 264.7 | 127.9 | 127.9 | 8.7403E-06 | 0.000002625 | 0 | 0 | 0.00169407 | 0.00050909 | 0 | 0 | HexNAc(5)Hex(6)Fuc(1)NeuGc(1) |
| a-Lytic (semi) | High | PMI-Byonic | Unambiguous | 1098 | 1105 | NGTHWFVT | 1098 | HexNAc(5)Hex(6)Fuc(1) | 5 | 6 | 1 | | | [S].nGTHWFVT.[Q] | N1(HexNAc(5)Hex(6)Fuc(1)) | P0DTC2 | 2 | 3 | 1 | 1032.41333 | 3095.22544 | 3095.22435 | 0.35 | 0.00036 | 267979.844 | HCD (High Energy Collision Dissociation) | MS2 | 27.63937 | 165.524 | 57.88 | 12115 | 12115 | 12097 | 5.34 | 292.3 | 199.2 | 199.2 | 4.5466E-06 | 1.2728E-06 | 0 | 0 | 0.00088194 | 0.00024696 | 0 | 0 | HexNAc(5)Hex(6)Fuc(1) |
| a-Lytic (semi) | High | PMI-Byonic | Unambiguous | 1098 | 1105 | NGTHWFVT | 1098 | HexNAc(5)Hex(6)Fuc(2) | 5 | 6 | 2 | | | [S].nGTHWFVT.[Q] | N1(HexNAc(5)Hex(6)Fuc(2)) | P0DTC2 | 2 | 3 | 1 | 1081.09973 | 3241.28464 | 3241.28226 | 0.73 | 0.00079 | 241930.906 | HCD (High Energy Collision Dissociation) | MS2 | 32.33216 | 200 | 57.54 | 12033 | 12033 | 12022 | 5.21 | 287.7 | 193.7 | 193.7 | 6.2094E-06 | 1.6251E-06 | 0 | 0 | 0.00120411 | 0.00031529 | 0 | 0 | HexNAc(5)Hex(6)Fuc(2) |
| a-Lytic (semi) | High | PMI-Byonic | Unambiguous | 1098 | 1105 | NGTHWFVT | 1098 | HexNAc(5)Hex(6)Fuc(3) | 5 | 6 | 3 | | | [S].nGTHWFVT.[Q] | N1(HexNAc(5)Hex(6)Fuc(3)) | P0DTC2 | 2 | 3 | 1 | 1129.78503 | 3387.34055 | 3387.34017 | 0.11 | 0.00013 | 383158.781 | HCD (High Energy Collision Dissociation) | MS2 | 2.412698 | 200 | 57.2893 | 11974 | 11974 | 11959 | 5.33 | 269.7 | 160.4 | 160.4 | 4.7151E-06 | 1.3503E-06 | 0 | 0 | 0.00091459 | 0.00026199 | 0 | 0 | HexNAc(5)Hex(6)Fuc(3) |
| a-Lytic (semi) | High | PMI-Byonic | Unambiguous | 1098 | 1105 | NGTHWFVT | 1098 | HexNAc(5)Hex(6)NeuGc(1) | 5 | 6 | | | 1 | [S].nGTHWFVT.[Q] | N1(HexNAc(5)Hex(6)NeuGc(1)) | P0DTC2 | 2 | 3 | 1 | 1086.09143 | 3256.25974 | 3256.25678 | 0.91 | 0.00099 | 168626.953 | HCD (High Energy Collision Dissociation) | MS2 | 25.28661 | 181.244 | 62.1339 | 13175 | 13175 | 13156 | 5.15 | 297.7 | 175.5 | 175.5 | 7.1273E-06 | 1.8722E-06 | 0 | 0 | 0.00138186 | 0.0003632 | 0 | 0 | HexNAc(5)Hex(6)NeuGc(1) |
| a-Lytic (semi) | High | PMI-Byonic | Unambiguous | 1098 | 1105 | NGTHWFVT | 1098 | HexNAc(5)Hex(6)NeuGc(2) | 5 | 6 | | | 2 | [S].nGTHWFVT.[Q] | N1(HexNAc(5)Hex(6)NeuGc(2)) | P0DTC2 | 2 | 3 | 1 | 1188.45557 | 3563.35215 | 3563.34711 | 1.41 | 0.00168 | 112685.211 | HCD (High Energy Collision Dissociation) | MS2 | 8.15163 | 200 | 66.0762 | 14222 | 14222 | 14203 | 2.94 | 168.9 | 35.7 | 35.7 | 0.00115193 | 0.00016835 | 0 | 0 | 0.18385733 | 0.02988505 | 0.02941176 | 0.04089218 | HexNAc(5)Hex(6)NeuGc(2) |
| a-Lytic (semi) | High | PMI-Byonic | Unambiguous | 1098 | 1105 | NGTHWFVT | 1098 | HexNAc(5)Hex(6) | 5 | 6 | | | | [S].nGTHWFVT.[Q] | N1(HexNAc(5)Hex(6)) | P0DTC2 | 2 | 2 | 1 | 1475.08728 | 2949.16728 | 2949.16645 | 0.28 | 0.00042 | 180737.203 | HCD (High Energy Collision Dissociation) | MS2 | 0 | 200 | 58.1433 | 12178 | 12178 | 12162 | 5.51 | 265.3 | 110.4 | 110.4 | 3.0995E-06 | 8.1477E-07 | 0 | 0 | 0.00060141 | 0.00015812 | 0 | 0 | HexNAc(5)Hex(6) |
| a-Lytic (semi) | High | PMI-Byonic | Unambiguous | 1098 | 1105 | NGTHWFVT | 1098 | HexNAc(6)Hex(5) | 6 | 5 | | | | [S].nGTHWFVT.[Q] | N1(HexNAc(6)Hex(5)) | P0DTC2 | 2 | 3 | 1 | 997.40411 | 2990.19779 | 2990.19299 | 1.6 | 0.0016 | 132518.875 | HCD (High Energy Collision Dissociation) | MS2 | 14.1648 | 200 | 58.4194 | 12240 | 12240 | 12221 | 2.42 | 141 | 53.9 | 53.9 | 0.00377413 | 0.00041172 | 0 | 0 | 0.42396853 | 0.07142856 | 0.07128711 | 0.10126579 | HexNAc(6)Hex(5) |
| a-Lytic (semi) | High | PMI-Byonic | Unambiguous | 1098 | 1105 | NGTHWFVT | 1098 | HexNAc(6)Hex(6)Fuc(1)NeuAc(1)NeuGc(2) | 6 | 6 | 1 | 1 | 2 | [S].nGTHWFVT.[Q] | N1(HexNAc(6)Hex(6)Fuc(1)NeuAc(1)NeuGc(2)) | P0DTC2 | 2 | 3 | 1 | 1402.1991 | 4204.58274 | 4203.57981 | 238.59 | 0.33431 | 121311.039 | HCD (High Energy Collision Dissociation) | MS2 | 20.82641 | 178.188 | 69.1111 | 14944 | 14944 | 14929 | 1.8 | 120.2 | 19.5 | 19.5 | 0.01571039 | 0.00196078 | 0.00194552 | 0.00320513 | 0.75527269 | 0.13111886 | 0.130662 | 0.17771879 | HexNAc(6)Hex(6)Fuc(1)NeuAc(1)NeuGc(2) |
| a-Lytic (semi) | High | PMI-Byonic | Unambiguous | 1098 | 1105 | NGTHWFVT | 1098 | HexNAc(6)Hex(6) | 6 | 6 | | | | [S].nGTHWFVT.[Q] | N1(HexNAc(6)Hex(6)) | P0DTC2 | 2 | 3 | 1 | 1051.42017 | 3152.24594 | 3152.24582 | 0.04 | 0.00004 | 204259.922 | HCD (High Energy Collision Dissociation) | MS2 | 0 | 189.993 | 58.4496 | 12247 | 12247 | 12234 | 3.35 | 185.9 | 59.9 | 59.9 | 0.00045156 | 0.0000727 | 0 | 0 | 0.08088846 | 0.02023121 | 0.01917808 | 0.02790696 | HexNAc(6)Hex(6) |
| a-Lytic (semi) | High | PMI-Byonic | Unambiguous | 1098 | 1105 | NGTHWFVT | 1098 | HexNAc(6)Hex(7)Fuc(1)NeuAc(1) | 6 | 7 | 1 | 1 | | [S].nGTHWFVT.[Q] | N1(HexNAc(6)Hex(7)Fuc(1)NeuAc(1)) | P0DTC2 | 2 | 3 | 1 | 1251.1571 | 3751.45676 | 3751.45197 | 1.28 | 0.0016 | 58304.6445 | HCD (High Energy Collision Dissociation) | MS2 | 0 | 126.81 | 61.4729 | 13002 | 13002 | 12989 | 2.94 | 146.4 | 44.2 | 44.2 | 0.00115193 | 0.00016371 | 0 | 0 | 0.18385733 | 0.02988505 | 0.02954545 | 0.04104476 | HexNAc(6)Hex(7)Fuc(1)NeuAc(1) |
| a-Lytic (semi) | High | PMI-Byonic | Unambiguous | 1098 | 1105 | NGTHWFVT | 1098 | HexNAc(6)Hex(7)Fuc(1)NeuGc(1) | 6 | 7 | 1 | | 1 | [S].nGTHWFVT.[Q] | N1(HexNAc(6)Hex(7)Fuc(1)NeuGc(1)) | P0DTC2 | 2 | 3 | 1 | 1256.4856 | 3767.44223 | 3767.44688 | -1.23 | -0.00155 | 406470.969 | HCD (High Energy Collision Dissociation) | MS2 | 12.62921 | 114.345 | 61.2259 | 12939 | 12939 | 12924 | 5.06 | 297.8 | 140.7 | 140.7 | 8.6216E-06 | 2.5884E-06 | 0 | 0 | 0.0016711 | 0.00050199 | 0 | 0 | HexNAc(6)Hex(7)Fuc(1)NeuGc(1) |
| [illegible] | [illegible] | [illegible] | [illegible] | [illegible] | [illegible] | [illegible] | [illegible] | [illegible] | [illegible] | [illegible] | [illegible] | [illegible] | [illegible] | [illegible] | [illegible] | [illegible] | 2 | 3 | 1 | 1154.12573 | 3460.36264 | 3460.35655 | 1.76 | 0.00203 | 333866.781 | HCD (High Energy Collision Dissociation) | MS2 | 0 | 120.009 | 57.1865 | 11948 | 11948 | 11933 | 4.76 | 296.6 | 169.8 | 169.8 | 0.000017223 | 4.2448E-06 | 0 | 0 | 0.00333277 | 0.0008227 | 0 | 0 | HexNAc(6)Hex(7)Fuc(1) |
| [illegible] | [illegible] | [illegible] | [illegible] | [illegible] | [illegible] | [illegible] | [illegible] | [illegible] | [illegible] | [illegible] | [illegible] | [illegible] | [illegible] | [illegible] | [illegible] | [illegible] | 2 | 3 | 1 | 1202.47119 | 3605.39902 | 3605.39406 | 1.38 | 0.00165 | 331560.063 | HCD (High Energy Collision Dissociation) | MS2 | 12.72466 | 165.141 | 61.7725 | 13079 | 13079 | 13065 | 3.31 | 223.2 | 102.5 | 102.5 | 0.00049262 | 0.000081503 | 0 | 0 | 0.0876826 | 0.02168021 | 0.02139037 | 0.03153152 | HexNAc(6)Hex(7)NeuAc(1) |
| [illegible] | [illegible] | [illegible] | [illegible] | [illegible] | [illegible] | [illegible] | [illegible] | [illegible] | [illegible] | [illegible] | [illegible] | [illegible] | [illegible] | [illegible] | [illegible] | [illegible] | 2 | 3 | 1 | 1207.80188 | 3621.39109 | 3621.38897 | 0.58 | 0.0007 | 1053591.13 | HCD (High Energy Collision Dissociation) | MS2 | 65.67776 | 60.383 | 61.5674 | 13027 | 13027 | 13013 | 4.64 | 230.2 | 109.9 | 109.9 | 0.000022859 | 5.0925E-06 | 0 | 0 | 0.00441858 | 0.00098652 | 0 | 0 | HexNAc(6)Hex(7)NeuGc(1) |
| [illegible] | [illegible] | [illegible] | [illegible] | [illegible] | [illegible] | [illegible] | [illegible] | [illegible] | [illegible] | [illegible] | [illegible] | [illegible] | [illegible] | [illegible] | [illegible] | [illegible] | 2 | 3 | 1 | 1310.16675 | 3928.48569 | 3928.4793 | 1.63 | 0.00213 | 157260.391 | HCD (High Energy Collision Dissociation) | MS2 | 37.98656 | 200 | 65.1363 | 13988 | 13988 | 13974 | 2.42 | 174.3 | 41 | 41 | 0.00377413 | 0.00041887 | 0 | 0 | 0.42396853 | 0.07142856 | 0.07114623 | 0.10094634 | HexNAc(6)Hex(7)NeuGc(2) |
| [illegible] | [illegible] | [illegible] | [illegible] | [illegible] | [illegible] | [illegible] | [illegible] | [illegible] | [illegible] | [illegible] | [illegible] | [illegible] | [illegible] | [illegible] | [illegible] | [illegible] | 2 | 3 | 1 | 1105.43835 | 3314.30051 | 3314.29864 | 0.56 | 0.00062 | 144681.109 | HCD (High Energy Collision Dissociation) | MS2 | 4.635627 | 200 | 57.393 | 11998 | 11998 | 11985 | 5.21 | 292.2 | 188.9 | 188.9 | 6.2094E-06 | 1.5921E-06 | 0 | 0 | 0.00120411 | 0.00030889 | 0 | 0 | HexNAc(6)Hex(7) |
| [illegible] | [illegible] | [illegible] | [illegible] | [illegible] | [illegible] | [illegible] | [illegible] | [illegible] | [illegible] | [illegible] | [illegible] | [illegible] | [illegible] | [illegible] | [illegible] | [illegible] | 2 | 3 | 1 | 1256.48755 | 3767.44809 | 3767.44688 | 0.32 | 0.0004 | 200389.063 | HCD (High Energy Collision Dissociation) | MS2 | 17.81931 | 164.351 | 61.7926 | 13084 | 13084 | 13065 | 3.12 | 177.2 | 39.5 | 39.5 | 0.00076184 | 0.00013512 | 0 | 0 | 0.12991224 | 0.02457002 | 0.02386634 | 0 | HexNAc(6)Hex(8)NeuAc(1) |
| [illegible] | [illegible] | [illegible] | [illegible] | [illegible] | [illegible] | [illegible] | [illegible] | [illegible] | [illegible] | [illegible] | [illegible] | [illegible] | [illegible] | [illegible] | [illegible] | [illegible] | 2 | 3 | 1 | 1213.47717 | 3638.41697 | 3638.40429 | 3.48 | 0.00423 | 206058.609 | HCD (High Energy Collision Dissociation) | MS2 | 2.73648 | 192.448 | 61.7208 | 13066 | 13066 | 13052 | 2.06 | 190.8 | 93.3 | 93.3 | 0.00879703 | 0.00059487 | 0 | 0 | 0.63225407 | 0.10223046 | 0.10185183 | 0.13793099 | HexNAc(6)Hex(9) |
| [illegible] | [illegible] | [illegible] | [illegible] | [illegible] | [illegible] | [illegible] | [illegible] | [illegible] | [illegible] | [illegible] | [illegible] | [illegible] | [illegible] | [illegible] | [illegible] | [illegible] | [illegible] | [illegible] | [illegible] | [illegible] | [illegible] | [illegible] | [illegible] | [illegible] | [illegible] | [illegible] | [illegible] | [illegible] | [illegible] | [illegible] | [illegible] | [illegible] | [illegible] | [illegible] | [illegible] | [illegible] | [illegible] | [illegible] | [illegible] | [illegible] | [illegible] | [illegible] | [illegible] | [illegible] | [illegible] | [illegible] |

Supplementary Table S9. Non-redundant site-specific glycan compositions of SARS CoV-2 Spike protein identified by Byonic

| No. | Glycosylation site | No. of assigned composition / site | Glycan Composition | No. of HexNAc | No. of Hex | No. of Fuc | No. of NeuAc | No. of NeuGc | Glycan composition |
|---|---|---|---|---|---|---|---|---|---|
| 1 | 17 | 21 | HexNAc(2)Hex(7) | 2 | 7 | 0 | 0 | 0 | 2:7:0:0:0 |
| 2 | 17 | 21 | HexNAc(3)Hex(6)Fuc(1) | 3 | 6 | 1 | 0 | 0 | 3:6:1:0:0 |
| 3 | 17 | 21 | HexNAc(4)Hex(3)Fuc(1) | 4 | 3 | 1 | 0 | 0 | 4:3:1:0:0 |
| 4 | 17 | 21 | HexNAc(4)Hex(3)NeuAc(1) | 4 | 3 | 0 | 1 | 0 | 4:3:0:1:0 |
| 5 | 17 | 21 | HexNAc(4)Hex(4)Fuc(2) | 4 | 4 | 2 | 0 | 0 | 4:4:2:0:0 |
| 6 | 17 | 21 | HexNAc(4)Hex(5)Fuc(1) | 4 | 5 | 1 | 0 | 0 | 4:5:1:0:0 |
| 7 | 17 | 21 | HexNAc(4)Hex(5)Fuc(2) | 4 | 5 | 2 | 0 | 0 | 4:5:2:0:0 |
| 8 | 17 | 21 | HexNAc(4)Hex(5)Fuc(3) | 4 | 5 | 3 | 0 | 0 | 4:5:3:0:0 |
| 9 | 17 | 21 | HexNAc(5)Hex(4)Fuc(1) | 5 | 4 | 1 | 0 | 0 | 5:4:1:0:0 |
| 10 | 17 | 21 | HexNAc(5)Hex(4)Fuc(2) | 5 | 4 | 2 | 0 | 0 | 5:4:2:0:0 |
| 11 | 17 | 21 | HexNAc(5)Hex(5)Fuc(1) | 5 | 5 | 1 | 0 | 0 | 5:5:1:0:0 |
| 12 | 17 | 21 | HexNAc(5)Hex(5)Fuc(2) | 5 | 5 | 2 | 0 | 0 | 5:5:2:0:0 |
| 13 | 17 | 21 | HexNAc(5)Hex(5)Fuc(3) | 5 | 5 | 3 | 0 | 0 | 5:5:3:0:0 |
| 14 | 17 | 21 | HexNAc(5)Hex(6)Fuc(2) | 5 | 6 | 2 | 0 | 0 | 5:6:2:0:0 |
| 15 | 17 | 21 | HexNAc(5)Hex(6)Fuc(3) | 5 | 6 | 3 | 0 | 0 | 5:6:3:0:0 |
| 16 | 17 | 21 | HexNAc(5)Hex(6)NeuAc(3) | 5 | 6 | 0 | 3 | 0 | 5:6:0:3:0 |
| 17 | 17 | 21 | HexNAc(6)Hex(6)Fuc(3) | 6 | 6 | 3 | 0 | 0 | 6:6:3:0:0 |
| 18 | 17 | 21 | HexNAc(6)Hex(7)Fuc(4) | 6 | 7 | 4 | 0 | 0 | 6:7:4:0:0 |
| 19 | 17 | 21 | HexNAc(6)Hex(7)Fuc(5) | 6 | 7 | 5 | 0 | 0 | 6:7:5:0:0 |
| 20 | 17 | 21 | HexNAc(7)Hex(8)Fuc(1)NeuAc(1) | 7 | 8 | 1 | 1 | 0 | 7:8:1:1:0 |
| 21 | 17 | 21 | HexNAc(8)Hex(7) | 8 | 7 | 0 | 0 | 0 | 8:7:0:0:0 |
| 22 | 61 | 40 | HexNAc(2)Hex(5) | 2 | 5 | 0 | 0 | 0 | 2:5:0:0:0 |
| 23 | 61 | 40 | HexNAc(2)Hex(6) | 2 | 6 | 0 | 0 | 0 | 2:6:0:0:0 |
| 24 | 61 | 40 | HexNAc(2)Hex(7) | 2 | 7 | 0 | 0 | 0 | 2:7:0:0:0 |
| 25 | 61 | 40 | HexNAc(2)Hex(8) | 2 | 8 | 0 | 0 | 0 | 2:8:0:0:0 |
| 26 | 61 | 40 | HexNAc(3)Hex(4)Fuc(1) | 3 | 4 | 1 | 0 | 0 | 3:4:1:0:0 |
| 27 | 61 | 40 | HexNAc(3)Hex(5)Fuc(1) | 3 | 5 | 1 | 0 | 0 | 3:5:1:0:0 |
| 28 | 61 | 40 | HexNAc(3)Hex(5) | 3 | 5 | 0 | 0 | 0 | 3:5:0:0:0 |
| 29 | 61 | 40 | HexNAc(3)Hex(6)Fuc(1) | 3 | 6 | 1 | 0 | 0 | 3:6:1:0:0 |
| 30 | 61 | 40 | HexNAc(3)Hex(6)NeuGc(1) | 3 | 6 | 0 | 0 | 1 | 3:6:0:0:1 |
| 31 | 61 | 40 | HexNAc(3)Hex(6) | 3 | 6 | 0 | 0 | 0 | 3:6:0:0:0 |
| 32 | 61 | 40 | HexNAc(4)Hex(3)Fuc(3) | 4 | 3 | 3 | 0 | 0 | 4:3:3:0:0 |
| 33 | 61 | 40 | HexNAc(4)Hex(4)Fuc(1) | 4 | 4 | 1 | 0 | 0 | 4:4:1:0:0 |
| 34 | 61 | 40 | HexNAc(4)Hex(5)Fuc(1) | 4 | 5 | 1 | 0 | 0 | 4:5:1:0:0 |
| 35 | 61 | 40 | HexNAc(4)Hex(5)Fuc(2) | 4 | 5 | 2 | 0 | 0 | 4:5:2:0:0 |
| 36 | 61 | 40 | HexNAc(4)Hex(5)Fuc(3) | 4 | 5 | 3 | 0 | 0 | 4:5:3:0:0 |
| 37 | 61 | 40 | HexNAc(4)Hex(5) | 4 | 5 | 0 | 0 | 0 | 4:5:0:0:0 |
| 38 | 61 | 40 | HexNAc(4)Hex(6)Fuc(1) | 4 | 6 | 1 | 0 | 0 | 4:6:1:0:0 |
| 39 | 61 | 40 | HexNAc(4)Hex(6)Fuc(2) | 4 | 6 | 2 | 0 | 0 | 4:6:2:0:0 |
| 40 | 61 | 40 | HexNAc(4)Hex(6)NeuAc(1) | 4 | 6 | 0 | 1 | 0 | 4:6:0:1:0 |
| 41 | 61 | 40 | HexNAc(4)Hex(7)Fuc(1) | 4 | 7 | 1 | 0 | 0 | 4:7:1:0:0 |
| 42 | 61 | 40 | HexNAc(4)Hex(7)Fuc(2) | 4 | 7 | 2 | 0 | 0 | 4:7:2:0:0 |
| 43 | 61 | 40 | HexNAc(5)Hex(4)Fuc(2) | 5 | 4 | 2 | 0 | 0 | 5:4:2:0:0 |
| 44 | 61 | 40 | HexNAc(5)Hex(5)Fuc(1) | 5 | 5 | 1 | 0 | 0 | 5:5:1:0:0 |
| 45 | 61 | 40 | HexNAc(5)Hex(5)Fuc(2) | 5 | 5 | 2 | 0 | 0 | 5:5:2:0:0 |
| 46 | 61 | 40 | HexNAc(5)Hex(5)Fuc(3) | 5 | 5 | 3 | 0 | 0 | 5:5:3:0:0 |
| 47 | 61 | 40 | HexNAc(5)Hex(5) | 5 | 5 | 0 | 0 | 0 | 5:5:0:0:0 |
| 48 | 61 | 40 | HexNAc(5)Hex(6)Fuc(1) | 5 | 6 | 1 | 0 | 0 | 5:6:1:0:0 |
| 49 | 61 | 40 | HexNAc(5)Hex(6)Fuc(2) | 5 | 6 | 2 | 0 | 0 | 5:6:2:0:0 |
| 50 | 61 | 40 | HexNAc(5)Hex(6)Fuc(3) | 5 | 6 | 3 | 0 | 0 | 5:6:3:0:0 |
| 51 | 61 | 40 | HexNAc(5)Hex(6) | 5 | 6 | 0 | 0 | 0 | 5:6:0:0:0 |
| 52 | 61 | 40 | HexNAc(6)Hex(4) | 6 | 4 | 0 | 0 | 0 | 6:4:0:0:0 |
| 53 | 61 | 40 | HexNAc(6)Hex(6)Fuc(1)NeuAc(1) | 6 | 6 | 1 | 1 | 0 | 6:6:1:1:0 |
| 54 | 61 | 40 | HexNAc(6)Hex(6)Fuc(1) | 6 | 6 | 1 | 0 | 0 | 6:6:1:0:0 |
| 55 | 61 | 40 | HexNAc(6)Hex(6)Fuc(2) | 6 | 6 | 2 | 0 | 0 | 6:6:2:0:0 |
| 56 | 61 | 40 | HexNAc(6)Hex(6)Fuc(3) | 6 | 6 | 3 | 0 | 0 | 6:6:3:0:0 |
| 57 | 61 | 40 | HexNAc(6)Hex(6) | 6 | 6 | 0 | 0 | 0 | 6:6:0:0:0 |
| 58 | 61 | 40 | HexNAc(6)Hex(8)Fuc(1)NeuAc(1) | 6 | 8 | 1 | 1 | 0 | 6:8:1:1:0 |
| 59 | 61 | 40 | HexNAc(8)Hex(3) | 8 | 3 | 0 | 0 | 0 | 8:3:0:0:0 |
| 60 | 61 | 40 | HexNAc(8)Hex(5) | 8 | 5 | 0 | 0 | 0 | 8:5:0:0:0 |
| 61 | 61 | 40 | HexNAc(9)Hex(3)Fuc(1) | 9 | 3 | 1 | 0 | 0 | 9:3:1:0:0 |
| 62 | 74 | 64 | HexNAc(2)Hex(8) | 2 | 8 | 0 | 0 | 0 | 2:8:0:0:0 |
| 63 | 74 | 64 | HexNAc(4)Hex(3)Fuc(1) | 4 | 3 | 1 | 0 | 0 | 4:3:1:0:0 |
| 64 | 74 | 64 | HexNAc(4)Hex(3)Fuc(2) | 4 | 3 | 2 | 0 | 0 | 4:3:2:0:0 |
| 65 | 74 | 64 | HexNAc(4)Hex(4)Fuc(1)NeuGc(1) | 4 | 4 | 1 | 0 | 1 | 4:4:1:0:1 |
| 66 | 74 | 64 | HexNAc(4)Hex(4)Fuc(1) | 4 | 4 | 1 | 0 | 0 | 4:4:1:0:0 |
| 67 | 74 | 64 | HexNAc(4)Hex(4)Fuc(2) | 4 | 4 | 2 | 0 | 0 | 4:4:2:0:0 |
| 68 | 74 | 64 | HexNAc(4)Hex(5)Fuc(1)NeuAc(1) | 4 | 5 | 1 | 1 | 0 | 4:5:1:1:0 |
| 69 | 74 | 64 | HexNAc(4)Hex(5)Fuc(1)NeuGc(1) | 4 | 5 | 1 | 0 | 1 | 4:5:1:0:1 |
| 70 | 74 | 64 | HexNAc(4)Hex(5)Fuc(1)NeuGc(2) | 4 | 5 | 1 | 0 | 2 | 4:5:1:0:2 |
| 71 | 74 | 64 | HexNAc(4)Hex(5)Fuc(1) | 4 | 5 | 1 | 0 | 0 | 4:5:1:0:0 |
| 72 | 74 | 64 | HexNAc(4)Hex(5)Fuc(2)NeuGc(1) | 4 | 5 | 2 | 0 | 1 | 4:5:2:0:1 |
| 73 | 74 | 64 | HexNAc(4)Hex(5)Fuc(2) | 4 | 5 | 2 | 0 | 0 | 4:5:2:0:0 |

| | | | | | | | | | |
|---|---|---|---|---|---|---|---|---|---|
| 74 | 74 | 64 | HexNAc(4)Hex(5)Fuc(3)NeuAc(2) | 4 | 5 | 3 | 2 | 0 | 4:5:3:2:0 |
| 75 | 74 | 64 | HexNAc(4)Hex(5)Fuc(3) | 4 | 5 | 3 | 0 | 0 | 4:5:3:0:0 |
| 76 | 74 | 64 | HexNAc(4)Hex(6)Fuc(1)NeuAc(1) | 4 | 6 | 1 | 1 | 0 | 4:6:1:1:0 |
| 77 | 74 | 64 | HexNAc(4)Hex(6)Fuc(3) | 4 | 6 | 3 | 0 | 0 | 4:6:3:0:0 |
| 78 | 74 | 64 | HexNAc(4)Hex(7)NeuAc(1) | 4 | 7 | 0 | 1 | 0 | 4:7:0:1:0 |
| 79 | 74 | 64 | HexNAc(5)Hex(3)Fuc(1) | 5 | 3 | 1 | 0 | 0 | 5:3:1:0:0 |
| 80 | 74 | 64 | HexNAc(5)Hex(4)Fuc(1)NeuGc(1) | 5 | 4 | 1 | 0 | 1 | 5:4:1:0:1 |
| 81 | 74 | 64 | HexNAc(5)Hex(4)Fuc(1) | 5 | 4 | 1 | 0 | 0 | 5:4:1:0:0 |
| 82 | 74 | 64 | HexNAc(5)Hex(4)Fuc(2)NeuAc(1) | 5 | 4 | 2 | 1 | 0 | 5:4:2:1:0 |
| 83 | 74 | 64 | HexNAc(5)Hex(4)Fuc(2)NeuGc(1) | 5 | 4 | 2 | 0 | 1 | 5:4:2:0:1 |
| 84 | 74 | 64 | HexNAc(5)Hex(4)Fuc(2) | 5 | 4 | 2 | 0 | 0 | 5:4:2:0:0 |
| 85 | 74 | 64 | HexNAc(5)Hex(5)Fuc(1)NeuAc(1) | 5 | 5 | 1 | 1 | 0 | 5:5:1:1:0 |
| 86 | 74 | 64 | HexNAc(5)Hex(5)Fuc(1)NeuGc(1) | 5 | 5 | 1 | 0 | 1 | 5:5:1:0:1 |
| 87 | 74 | 64 | HexNAc(5)Hex(5)Fuc(1)NeuGc(2) | 5 | 5 | 1 | 0 | 2 | 5:5:1:0:2 |
| 88 | 74 | 64 | HexNAc(5)Hex(5)Fuc(1) | 5 | 5 | 1 | 0 | 0 | 5:5:1:0:0 |
| 89 | 74 | 64 | HexNAc(5)Hex(5)Fuc(2) | 5 | 5 | 2 | 0 | 0 | 5:5:2:0:0 |
| 90 | 74 | 64 | HexNAc(5)Hex(5)Fuc(3) | 5 | 5 | 3 | 0 | 0 | 5:5:3:0:0 |
| 91 | 74 | 64 | HexNAc(5)Hex(6)Fuc(1)NeuAc(1) | 5 | 6 | 1 | 1 | 0 | 5:6:1:1:0 |
| 92 | 74 | 64 | HexNAc(5)Hex(6)Fuc(1) | 5 | 6 | 1 | 0 | 0 | 5:6:1:0:0 |
| 93 | 74 | 64 | HexNAc(5)Hex(6)Fuc(2)NeuAc(1) | 5 | 6 | 2 | 1 | 0 | 5:6:2:1:0 |
| 94 | 74 | 64 | HexNAc(5)Hex(6)Fuc(2)NeuGc(1) | 5 | 6 | 2 | 0 | 1 | 5:6:2:0:1 |
| 95 | 74 | 64 | HexNAc(5)Hex(6)Fuc(2) | 5 | 6 | 2 | 0 | 0 | 5:6:2:0:0 |
| 96 | 74 | 64 | HexNAc(5)Hex(6)Fuc(3)NeuGc(1) | 5 | 6 | 3 | 0 | 1 | 5:6:3:0:1 |
| 97 | 74 | 64 | HexNAc(5)Hex(6)Fuc(3) | 5 | 6 | 3 | 0 | 0 | 5:6:3:0:0 |
| 98 | 74 | 64 | HexNAc(5)Hex(7)Fuc(1)NeuAc(1)NeuGc(1) | 5 | 7 | 1 | 1 | 1 | 5:7:1:1:1 |
| 99 | 74 | 64 | HexNAc(6)Hex(3)Fuc(1)NeuGc(1) | 6 | 3 | 1 | 0 | 1 | 6:3:1:0:1 |
| 100 | 74 | 64 | HexNAc(6)Hex(3)Fuc(1) | 6 | 3 | 1 | 0 | 0 | 6:3:1:0:0 |
| 101 | 74 | 64 | HexNAc(6)Hex(3)Fuc(2) | 6 | 3 | 2 | 0 | 0 | 6:3:2:0:0 |
| 102 | 74 | 64 | HexNAc(6)Hex(3)Fuc(3) | 6 | 3 | 3 | 0 | 0 | 6:3:3:0:0 |
| 103 | 74 | 64 | HexNAc(6)Hex(4)Fuc(1) | 6 | 4 | 1 | 0 | 0 | 6:4:1:0:0 |
| 104 | 74 | 64 | HexNAc(6)Hex(4)Fuc(2) | 6 | 4 | 2 | 0 | 0 | 6:4:2:0:0 |
| 105 | 74 | 64 | HexNAc(6)Hex(5)Fuc(1)NeuAc(1)NeuGc(1) | 6 | 5 | 1 | 1 | 1 | 6:5:1:1:1 |
| 106 | 74 | 64 | HexNAc(6)Hex(5)Fuc(1)NeuAc(1) | 6 | 5 | 1 | 1 | 0 | 6:5:1:1:0 |
| 107 | 74 | 64 | HexNAc(6)Hex(5)Fuc(1) | 6 | 5 | 1 | 0 | 0 | 6:5:1:0:0 |
| 108 | 74 | 64 | HexNAc(6)Hex(6)Fuc(1)NeuAc(1)NeuGc(1) | 6 | 6 | 1 | 1 | 1 | 6:6:1:1:1 |
| 109 | 74 | 64 | HexNAc(6)Hex(6)Fuc(1) | 6 | 6 | 1 | 0 | 0 | 6:6:1:0:0 |
| 110 | 74 | 64 | HexNAc(6)Hex(6)Fuc(2) | 6 | 6 | 2 | 0 | 0 | 6:6:2:0:0 |
| 111 | 74 | 64 | HexNAc(6)Hex(6)Fuc(3) | 6 | 6 | 3 | 0 | 0 | 6:6:3:0:0 |
| 112 | 74 | 64 | HexNAc(6)Hex(6)NeuAc(1)NeuGc(1) | 6 | 6 | 0 | 1 | 1 | 6:6:0:1:1 |
| 113 | 74 | 64 | HexNAc(6)Hex(7)Fuc(1)NeuAc(1) | 6 | 7 | 1 | 1 | 0 | 6:7:1:1:0 |
| 114 | 74 | 64 | HexNAc(6)Hex(7)Fuc(1)NeuGc(1) | 6 | 7 | 1 | 0 | 1 | 6:7:1:0:1 |
| 115 | 74 | 64 | HexNAc(6)Hex(7)Fuc(1) | 6 | 7 | 1 | 0 | 0 | 6:7:1:0:0 |
| 116 | 74 | 64 | HexNAc(6)Hex(7)Fuc(4)NeuGc(1) | 6 | 7 | 4 | 0 | 1 | 6:7:4:0:1 |
| 117 | 74 | 64 | HexNAc(6)Hex(7)Fuc(4) | 6 | 7 | 4 | 0 | 0 | 6:7:4:0:0 |
| 118 | 74 | 64 | HexNAc(6)Hex(7)Fuc(5) | 6 | 7 | 5 | 0 | 0 | 6:7:5:0:0 |
| 119 | 74 | 64 | HexNAc(6)Hex(7)NeuAc(1) | 6 | 7 | 0 | 1 | 0 | 6:7:0:1:0 |
| 120 | 74 | 64 | HexNAc(7)Hex(3)Fuc(1) | 7 | 3 | 1 | 0 | 0 | 7:3:1:0:0 |
| 121 | 74 | 64 | HexNAc(7)Hex(4)Fuc(2) | 7 | 4 | 2 | 0 | 0 | 7:4:2:0:0 |
| 122 | 74 | 64 | HexNAc(8)Hex(8) | 8 | 8 | 0 | 0 | 0 | 8:8:0:0:0 |
| 123 | 74 | 64 | HexNAc(8)Hex(9)Fuc(1) | 8 | 9 | 1 | 0 | 0 | 8:9:1:0:0 |
| 124 | 74 | 64 | HexNAc(9)Hex(9)Fuc(1) | 9 | 9 | 1 | 0 | 0 | 9:9:1:0:0 |
| 125 | 74 | 64 | HexNAc(9)Hex(10) | 9 | 10 | 0 | 0 | 0 | 9:10:0:0:0 |
| 126 | 122 | 72 | HexNAc(2)Hex(5) | 2 | 5 | 0 | 0 | 0 | 2:5:0:0:0 |
| 127 | 122 | 72 | HexNAc(2)Hex(6) | 2 | 6 | 0 | 0 | 0 | 2:6:0:0:0 |
| 128 | 122 | 72 | HexNAc(2)Hex(7) | 2 | 7 | 0 | 0 | 0 | 2:7:0:0:0 |
| 129 | 122 | 72 | HexNAc(2)Hex(8) | 2 | 8 | 0 | 0 | 0 | 2:8:0:0:0 |
| 130 | 122 | 72 | HexNAc(3)Hex(3)Fuc(1) | 3 | 3 | 1 | 0 | 0 | 3:3:1:0:0 |
| 131 | 122 | 72 | HexNAc(3)Hex(3) | 3 | 3 | 0 | 0 | 0 | 3:3:0:0:0 |
| 132 | 122 | 72 | HexNAc(3)Hex(4)Fuc(1) | 3 | 4 | 1 | 0 | 0 | 3:4:1:0:0 |
| 133 | 122 | 72 | HexNAc(3)Hex(4)Fuc(2) | 3 | 4 | 2 | 0 | 0 | 3:4:2:0:0 |
| 134 | 122 | 72 | HexNAc(3)Hex(4)NeuGc(1) | 3 | 4 | 0 | 0 | 1 | 3:4:0:0:1 |
| 135 | 122 | 72 | HexNAc(3)Hex(4) | 3 | 4 | 0 | 0 | 0 | 3:4:0:0:0 |
| 136 | 122 | 72 | HexNAc(3)Hex(5)Fuc(1) | 3 | 5 | 1 | 0 | 0 | 3:5:1:0:0 |
| 137 | 122 | 72 | HexNAc(3)Hex(5)NeuAc(1) | 3 | 5 | 0 | 1 | 0 | 3:5:0:1:0 |
| 138 | 122 | 72 | HexNAc(3)Hex(5)NeuGc(1) | 3 | 5 | 0 | 0 | 1 | 3:5:0:0:1 |
| 139 | 122 | 72 | HexNAc(3)Hex(5) | 3 | 5 | 0 | 0 | 0 | 3:5:0:0:0 |
| 140 | 122 | 72 | HexNAc(3)Hex(6)Fuc(1)NeuAc(1) | 3 | 6 | 1 | 1 | 0 | 3:6:1:1:0 |
| 141 | 122 | 72 | HexNAc(3)Hex(6)Fuc(1) | 3 | 6 | 1 | 0 | 0 | 3:6:1:0:0 |
| 142 | 122 | 72 | HexNAc(3)Hex(6)NeuAc(1) | 3 | 6 | 0 | 1 | 0 | 3:6:0:1:0 |
| 143 | 122 | 72 | HexNAc(3)Hex(6)NeuGc(1) | 3 | 6 | 0 | 0 | 1 | 3:6:0:0:1 |
| 144 | 122 | 72 | HexNAc(3)Hex(6) | 3 | 6 | 0 | 0 | 0 | 3:6:0:0:0 |
| 145 | 122 | 72 | HexNAc(4)Hex(3)Fuc(1) | 4 | 3 | 1 | 0 | 0 | 4:3:1:0:0 |
| 146 | 122 | 72 | HexNAc(4)Hex(3) | 4 | 3 | 0 | 0 | 0 | 4:3:0:0:0 |
| 147 | 122 | 72 | HexNAc(4)Hex(4)Fuc(1) | 4 | 4 | 1 | 0 | 0 | 4:4:1:0:0 |
| 148 | 122 | 72 | HexNAc(4)Hex(4)Fuc(2) | 4 | 4 | 2 | 0 | 0 | 4:4:2:0:0 |
| 149 | 122 | 72 | HexNAc(4)Hex(4) | 4 | 4 | 0 | 0 | 0 | 4:4:0:0:0 |
| 150 | 122 | 72 | HexNAc(4)Hex(5)Fuc(1)NeuAc(1) | 4 | 5 | 1 | 1 | 0 | 4:5:1:1:0 |
| 151 | 122 | 72 | HexNAc(4)Hex(5)Fuc(1)NeuGc(1) | 4 | 5 | 1 | 0 | 1 | 4:5:1:0:1 |

| | | | | | | | | | |
|---|---|---|---|---|---|---|---|---|---|
| 152 | 122 | 72 | HexNAc(4)Hex(5)Fuc(1) | 4 | 5 | 1 | 0 | 0 | 4:5:1:0:0 |
| 153 | 122 | 72 | HexNAc(4)Hex(5)Fuc(2)NeuGc(1) | 4 | 5 | 2 | 0 | 1 | 4:5:2:0:1 |
| 154 | 122 | 72 | HexNAc(4)Hex(5)Fuc(2) | 4 | 5 | 2 | 0 | 0 | 4:5:2:0:0 |
| 155 | 122 | 72 | HexNAc(4)Hex(5)Fuc(3) | 4 | 5 | 3 | 0 | 0 | 4:5:3:0:0 |
| 156 | 122 | 72 | HexNAc(4)Hex(5)NeuAc(1)NeuGc(1) | 4 | 5 | 0 | 1 | 1 | 4:5:0:1:1 |
| 157 | 122 | 72 | HexNAc(4)Hex(5)NeuAc(1) | 4 | 5 | 0 | 1 | 0 | 4:5:0:1:0 |
| 158 | 122 | 72 | HexNAc(4)Hex(5)NeuGc(1) | 4 | 5 | 0 | 0 | 1 | 4:5:0:0:1 |
| 159 | 122 | 72 | HexNAc(4)Hex(5) | 4 | 5 | 0 | 0 | 0 | 4:5:0:0:0 |
| 160 | 122 | 72 | HexNAc(4)Hex(6)Fuc(1) | 4 | 6 | 1 | 0 | 0 | 4:6:1:0:0 |
| 161 | 122 | 72 | HexNAc(4)Hex(6)Fuc(2) | 4 | 6 | 2 | 0 | 0 | 4:6:2:0:0 |
| 162 | 122 | 72 | HexNAc(4)Hex(6)NeuAc(1) | 4 | 6 | 0 | 1 | 0 | 4:6:0:1:0 |
| 163 | 122 | 72 | HexNAc(4)Hex(6) | 4 | 6 | 0 | 0 | 0 | 4:6:0:0:0 |
| 164 | 122 | 72 | HexNAc(4)Hex(7)Fuc(1) | 4 | 7 | 1 | 0 | 0 | 4:7:1:0:0 |
| 165 | 122 | 72 | HexNAc(4)Hex(7)Fuc(2) | 4 | 7 | 2 | 0 | 0 | 4:7:2:0:0 |
| 166 | 122 | 72 | HexNAc(5)Hex(3)Fuc(1) | 5 | 3 | 1 | 0 | 0 | 5:3:1:0:0 |
| 167 | 122 | 72 | HexNAc(5)Hex(4)Fuc(1) | 5 | 4 | 1 | 0 | 0 | 5:4:1:0:0 |
| 168 | 122 | 72 | HexNAc(5)Hex(4)Fuc(2)NeuAc(1) | 5 | 4 | 2 | 1 | 0 | 5:4:2:1:0 |
| 169 | 122 | 72 | HexNAc(5)Hex(4)Fuc(2) | 5 | 4 | 2 | 0 | 0 | 5:4:2:0:0 |
| 170 | 122 | 72 | HexNAc(5)Hex(4)NeuGc(1) | 5 | 4 | 0 | 0 | 1 | 5:4:0:0:1 |
| 171 | 122 | 72 | HexNAc(5)Hex(4) | 5 | 4 | 0 | 0 | 0 | 5:4:0:0:0 |
| 172 | 122 | 72 | HexNAc(5)Hex(5)Fuc(1) | 5 | 5 | 1 | 0 | 0 | 5:5:1:0:0 |
| 173 | 122 | 72 | HexNAc(5)Hex(5)Fuc(2) | 5 | 5 | 2 | 0 | 0 | 5:5:2:0:0 |
| 174 | 122 | 72 | HexNAc(5)Hex(5)Fuc(3) | 5 | 5 | 3 | 0 | 0 | 5:5:3:0:0 |
| 175 | 122 | 72 | HexNAc(5)Hex(5)NeuAc(1) | 5 | 5 | 0 | 1 | 0 | 5:5:0:1:0 |
| 176 | 122 | 72 | HexNAc(5)Hex(5) | 5 | 5 | 0 | 0 | 0 | 5:5:0:0:0 |
| 177 | 122 | 72 | HexNAc(5)Hex(6)Fuc(1)NeuAc(1) | 5 | 6 | 1 | 1 | 0 | 5:6:1:1:0 |
| 178 | 122 | 72 | HexNAc(5)Hex(6)Fuc(1) | 5 | 6 | 1 | 0 | 0 | 5:6:1:0:0 |
| 179 | 122 | 72 | HexNAc(5)Hex(6)Fuc(2) | 5 | 6 | 2 | 0 | 0 | 5:6:2:0:0 |
| 180 | 122 | 72 | HexNAc(5)Hex(6)Fuc(3) | 5 | 6 | 3 | 0 | 0 | 5:6:3:0:0 |
| 181 | 122 | 72 | HexNAc(5)Hex(6)NeuAc(1) | 5 | 6 | 0 | 1 | 0 | 5:6:0:1:0 |
| 182 | 122 | 72 | HexNAc(5)Hex(6)NeuGc(1) | 5 | 6 | 0 | 0 | 1 | 5:6:0:0:1 |
| 183 | 122 | 72 | HexNAc(5)Hex(6) | 5 | 6 | 0 | 0 | 0 | 5:6:0:0:0 |
| 184 | 122 | 72 | HexNAc(5)Hex(7)Fuc(1)NeuAc(1) | 5 | 7 | 1 | 1 | 0 | 5:7:1:1:0 |
| 185 | 122 | 72 | HexNAc(6)Hex(3) | 6 | 3 | 0 | 0 | 0 | 6:3:0:0:0 |
| 186 | 122 | 72 | HexNAc(6)Hex(5) | 6 | 5 | 0 | 0 | 0 | 6:5:0:0:0 |
| 187 | 122 | 72 | HexNAc(6)Hex(6)Fuc(1)NeuAc(1) | 6 | 6 | 1 | 1 | 0 | 6:6:1:1:0 |
| 188 | 122 | 72 | HexNAc(6)Hex(6)Fuc(1) | 6 | 6 | 1 | 0 | 0 | 6:6:1:0:0 |
| 189 | 122 | 72 | HexNAc(6)Hex(6)Fuc(2) | 6 | 6 | 2 | 0 | 0 | 6:6:2:0:0 |
| 190 | 122 | 72 | HexNAc(6)Hex(6)NeuAc(1)NeuGc(1) | 6 | 6 | 0 | 1 | 1 | 6:6:0:1:1 |
| 191 | 122 | 72 | HexNAc(6)Hex(6)NeuAc(1) | 6 | 6 | 0 | 1 | 0 | 6:6:0:1:0 |
| 192 | 122 | 72 | HexNAc(6)Hex(6) | 6 | 6 | 0 | 0 | 0 | 6:6:0:0:0 |
| 193 | 122 | 72 | HexNAc(6)Hex(7)Fuc(1)NeuAc(1) | 6 | 7 | 1 | 1 | 0 | 6:7:1:1:0 |
| 194 | 122 | 72 | HexNAc(6)Hex(7)Fuc(1) | 6 | 7 | 1 | 0 | 0 | 6:7:1:0:0 |
| 195 | 122 | 72 | HexNAc(6)Hex(7)Fuc(4) | 6 | 7 | 4 | 0 | 0 | 6:7:4:0:0 |
| 196 | 122 | 72 | HexNAc(6)Hex(7)NeuAc(1) | 6 | 7 | 0 | 1 | 0 | 6:7:0:1:0 |
| 197 | 122 | 72 | HexNAc(7)Hex(7)Fuc(1) | 7 | 7 | 1 | 0 | 0 | 7:7:1:0:0 |
| 198 | 149 | 32 | HexNAc(2)Hex(5) | 2 | 5 | 0 | 0 | 0 | 2:5:0:0:0 |
| 199 | 149 | 32 | HexNAc(4)Hex(4)Fuc(1)NeuAc(1) | 4 | 4 | 1 | 1 | 0 | 4:4:1:1:0 |
| 200 | 149 | 32 | HexNAc(4)Hex(4)Fuc(2) | 4 | 4 | 2 | 0 | 0 | 4:4:2:0:0 |
| 201 | 149 | 32 | HexNAc(4)Hex(5)Fuc(1) | 4 | 5 | 1 | 0 | 0 | 4:5:1:0:0 |
| 202 | 149 | 32 | HexNAc(4)Hex(5)Fuc(3)NeuAc(2) | 4 | 5 | 3 | 2 | 0 | 4:5:3:2:0 |
| 203 | 149 | 32 | HexNAc(4)Hex(5)Fuc(3) | 4 | 5 | 3 | 0 | 0 | 4:5:3:0:0 |
| 204 | 149 | 32 | HexNAc(4)Hex(5) | 4 | 5 | 0 | 0 | 0 | 4:5:0:0:0 |
| 205 | 149 | 32 | HexNAc(4)Hex(6)Fuc(1)NeuAc(1) | 4 | 6 | 1 | 1 | 0 | 4:6:1:1:0 |
| 206 | 149 | 32 | HexNAc(4)Hex(6)Fuc(2) | 4 | 6 | 2 | 0 | 0 | 4:6:2:0:0 |
| 207 | 149 | 32 | HexNAc(4)Hex(6)NeuAc(1) | 4 | 6 | 0 | 1 | 0 | 4:6:0:1:0 |
| 208 | 149 | 32 | HexNAc(5)Hex(3)Fuc(1) | 5 | 3 | 1 | 0 | 0 | 5:3:1:0:0 |
| 209 | 149 | 32 | HexNAc(5)Hex(4)Fuc(2)NeuGc(1) | 5 | 4 | 2 | 0 | 1 | 5:4:2:0:1 |
| 210 | 149 | 32 | HexNAc(5)Hex(4)Fuc(2) | 5 | 4 | 2 | 0 | 0 | 5:4:2:0:0 |
| 211 | 149 | 32 | HexNAc(5)Hex(5)Fuc(1)NeuAc(1) | 5 | 5 | 1 | 1 | 0 | 5:5:1:1:0 |
| 212 | 149 | 32 | HexNAc(5)Hex(5)Fuc(1) | 5 | 5 | 1 | 0 | 0 | 5:5:1:0:0 |
| 213 | 149 | 32 | HexNAc(5)Hex(5)Fuc(2) | 5 | 5 | 2 | 0 | 0 | 5:5:2:0:0 |
| 214 | 149 | 32 | HexNAc(5)Hex(5)Fuc(3) | 5 | 5 | 3 | 0 | 0 | 5:5:3:0:0 |
| 215 | 149 | 32 | HexNAc(5)Hex(5)NeuAc(1)NeuGc(1) | 5 | 5 | 0 | 1 | 1 | 5:5:0:1:1 |
| 216 | 149 | 32 | HexNAc(5)Hex(6)Fuc(1)NeuAc(1)NeuGc(1) | 5 | 6 | 1 | 1 | 1 | 5:6:1:1:1 |
| 217 | 149 | 32 | HexNAc(5)Hex(6)Fuc(1) | 5 | 6 | 1 | 0 | 0 | 5:6:1:0:0 |
| 218 | 149 | 32 | HexNAc(5)Hex(6)Fuc(2)NeuAc(1) | 5 | 6 | 2 | 1 | 0 | 5:6:2:1:0 |
| 219 | 149 | 32 | HexNAc(5)Hex(6)NeuAc(3) | 5 | 6 | 0 | 3 | 0 | 5:6:0:3:0 |
| 220 | 149 | 32 | HexNAc(5)Hex(7)Fuc(1)NeuAc(1)NeuGc(1) | 5 | 7 | 1 | 1 | 1 | 5:7:1:1:1 |
| 221 | 149 | 32 | HexNAc(6)Hex(3)Fuc(1) | 6 | 3 | 1 | 0 | 0 | 6:3:1:0:0 |
| 222 | 149 | 32 | HexNAc(6)Hex(6)Fuc(1)NeuAc(1)NeuGc(1) | 6 | 6 | 1 | 1 | 1 | 6:6:1:1:1 |
| 223 | 149 | 32 | HexNAc(6)Hex(6)Fuc(1) | 6 | 6 | 1 | 0 | 0 | 6:6:1:0:0 |
| 224 | 149 | 32 | HexNAc(6)Hex(6)Fuc(2) | 6 | 6 | 2 | 0 | 0 | 6:6:2:0:0 |
| 225 | 149 | 32 | HexNAc(6)Hex(7)Fuc(1)NeuAc(1) | 6 | 7 | 1 | 1 | 0 | 6:7:1:1:0 |
| 226 | 149 | 32 | HexNAc(6)Hex(7)Fuc(1) | 6 | 7 | 1 | 0 | 0 | 6:7:1:0:0 |
| 227 | 149 | 32 | HexNAc(6)Hex(7)Fuc(4)NeuAc(1) | 6 | 7 | 4 | 1 | 0 | 6:7:4:1:0 |
| 228 | 149 | 32 | HexNAc(6)Hex(7)Fuc(5) | 6 | 7 | 5 | 0 | 0 | 6:7:5:0:0 |
| 229 | 149 | 32 | HexNAc(6)Hex(7)NeuAc(1)NeuGc(1) | 6 | 7 | 0 | 1 | 1 | 6:7:0:1:1 |

| | | | | | | | | | |
|---|---|---|---|---|---|---|---|---|---|
| 230 | 165 | 34 | HexNAc(2)Hex(5) | 2 | 5 | 0 | 0 | 0 | 2:5:0:0:0 |
| 231 | 165 | 34 | HexNAc(2)Hex(6) | 2 | 6 | 0 | 0 | 0 | 2:6:0:0:0 |
| 232 | 165 | 34 | HexNAc(3)Hex(4)Fuc(1) | 3 | 4 | 1 | 0 | 0 | 3:4:1:0:0 |
| 233 | 165 | 34 | HexNAc(3)Hex(5)Fuc(1) | 3 | 5 | 1 | 0 | 0 | 3:5:1:0:0 |
| 234 | 165 | 34 | HexNAc(3)Hex(6)Fuc(1) | 3 | 6 | 1 | 0 | 0 | 3:6:1:0:0 |
| 235 | 165 | 34 | HexNAc(3)Hex(6)NeuGc(1) | 3 | 6 | 0 | 0 | 1 | 3:6:0:0:1 |
| 236 | 165 | 34 | HexNAc(4)Hex(5)Fuc(1)NeuAc(1) | 4 | 5 | 1 | 1 | 0 | 4:5:1:1:0 |
| 237 | 165 | 34 | HexNAc(4)Hex(5)Fuc(1)NeuGc(1) | 4 | 5 | 1 | 0 | 1 | 4:5:1:0:1 |
| 238 | 165 | 34 | HexNAc(4)Hex(5)Fuc(1) | 4 | 5 | 1 | 0 | 0 | 4:5:1:0:0 |
| 239 | 165 | 34 | HexNAc(4)Hex(5)Fuc(2) | 4 | 5 | 2 | 0 | 0 | 4:5:2:0:0 |
| 240 | 165 | 34 | HexNAc(4)Hex(5)Fuc(3) | 4 | 5 | 3 | 0 | 0 | 4:5:3:0:0 |
| 241 | 165 | 34 | HexNAc(4)Hex(5)NeuAc(1) | 4 | 5 | 0 | 1 | 0 | 4:5:0:1:0 |
| 242 | 165 | 34 | HexNAc(4)Hex(5) | 4 | 5 | 0 | 0 | 0 | 4:5:0:0:0 |
| 243 | 165 | 34 | HexNAc(4)Hex(6)Fuc(1)NeuAc(1) | 4 | 6 | 1 | 1 | 0 | 4:6:1:1:0 |
| 244 | 165 | 34 | HexNAc(4)Hex(6)NeuAc(1) | 4 | 6 | 0 | 1 | 0 | 4:6:0:1:0 |
| 245 | 165 | 34 | HexNAc(5)Hex(5)Fuc(1)NeuAc(1) | 5 | 5 | 1 | 1 | 0 | 5:5:1:1:0 |
| 246 | 165 | 34 | HexNAc(5)Hex(5)Fuc(1) | 5 | 5 | 1 | 0 | 0 | 5:5:1:0:0 |
| 247 | 165 | 34 | HexNAc(5)Hex(5)Fuc(2) | 5 | 5 | 2 | 0 | 0 | 5:5:2:0:0 |
| 248 | 165 | 34 | HexNAc(5)Hex(5)Fuc(3) | 5 | 5 | 3 | 0 | 0 | 5:5:3:0:0 |
| 249 | 165 | 34 | HexNAc(5)Hex(5) | 5 | 5 | 0 | 0 | 0 | 5:5:0:0:0 |
| 250 | 165 | 34 | HexNAc(5)Hex(6)Fuc(1)NeuAc(1) | 5 | 6 | 1 | 1 | 0 | 5:6:1:1:0 |
| 251 | 165 | 34 | HexNAc(5)Hex(6)Fuc(2)NeuAc(1) | 5 | 6 | 2 | 1 | 0 | 5:6:2:1:0 |
| 252 | 165 | 34 | HexNAc(5)Hex(6)Fuc(2) | 5 | 6 | 2 | 0 | 0 | 5:6:2:0:0 |
| 253 | 165 | 34 | HexNAc(5)Hex(6)Fuc(3) | 5 | 6 | 3 | 0 | 0 | 5:6:3:0:0 |
| 254 | 165 | 34 | HexNAc(5)Hex(6) | 5 | 6 | 0 | 0 | 0 | 5:6:0:0:0 |
| 255 | 165 | 34 | HexNAc(6)Hex(3)Fuc(1)NeuAc(1) | 6 | 3 | 1 | 1 | 0 | 6:3:1:1:0 |
| 256 | 165 | 34 | HexNAc(6)Hex(6)Fuc(1)NeuAc(1) | 6 | 6 | 1 | 1 | 0 | 6:6:1:1:0 |
| 257 | 165 | 34 | HexNAc(6)Hex(6)Fuc(1) | 6 | 6 | 1 | 0 | 0 | 6:6:1:0:0 |
| 258 | 165 | 34 | HexNAc(6)Hex(7)Fuc(1)NeuAc(1) | 6 | 7 | 1 | 1 | 0 | 6:7:1:1:0 |
| 259 | 165 | 34 | HexNAc(6)Hex(7)Fuc(1)NeuAc(2) | 6 | 7 | 1 | 2 | 0 | 6:7:1:2:0 |
| 260 | 165 | 34 | HexNAc(6)Hex(7)Fuc(1) | 6 | 7 | 1 | 0 | 0 | 6:7:1:0:0 |
| 261 | 165 | 34 | HexNAc(6)Hex(7)Fuc(4) | 6 | 7 | 4 | 0 | 0 | 6:7:4:0:0 |
| 262 | 165 | 34 | HexNAc(6)Hex(7)Fuc(5) | 6 | 7 | 5 | 0 | 0 | 6:7:5:0:0 |
| 263 | 165 | 34 | HexNAc(8)Hex(9) | 8 | 9 | 0 | 0 | 0 | 8:9:0:0:0 |
| 264 | 234 | 38 | HexNAc(2)Hex(5) | 2 | 5 | 0 | 0 | 0 | 2:5:0:0:0 |
| 265 | 234 | 38 | HexNAc(2)Hex(6) | 2 | 6 | 0 | 0 | 0 | 2:6:0:0:0 |
| 266 | 234 | 38 | HexNAc(2)Hex(7) | 2 | 7 | 0 | 0 | 0 | 2:7:0:0:0 |
| 267 | 234 | 38 | HexNAc(2)Hex(8) | 2 | 8 | 0 | 0 | 0 | 2:8:0:0:0 |
| 268 | 234 | 38 | HexNAc(2)Hex(9) | 2 | 9 | 0 | 0 | 0 | 2:9:0:0:0 |
| 269 | 234 | 38 | HexNAc(3)Hex(4)Fuc(1) | 3 | 4 | 1 | 0 | 0 | 3:4:1:0:0 |
| 270 | 234 | 38 | HexNAc(3)Hex(4)NeuGc(1) | 3 | 4 | 0 | 0 | 1 | 3:4:0:0:1 |
| 271 | 234 | 38 | HexNAc(3)Hex(5)Fuc(1) | 3 | 5 | 1 | 0 | 0 | 3:5:1:0:0 |
| 272 | 234 | 38 | HexNAc(3)Hex(5)NeuGc(1) | 3 | 5 | 0 | 0 | 1 | 3:5:0:0:1 |
| 273 | 234 | 38 | HexNAc(3)Hex(5) | 3 | 5 | 0 | 0 | 0 | 3:5:0:0:0 |
| 274 | 234 | 38 | HexNAc(3)Hex(6) | 3 | 6 | 0 | 0 | 0 | 3:6:0:0:0 |
| 275 | 234 | 38 | HexNAc(4)Hex(3)Fuc(1) | 4 | 3 | 1 | 0 | 0 | 4:3:1:0:0 |
| 276 | 234 | 38 | HexNAc(4)Hex(4) | 4 | 4 | 0 | 0 | 0 | 4:4:0:0:0 |
| 277 | 234 | 38 | HexNAc(4)Hex(5)Fuc(1) | 4 | 5 | 1 | 0 | 0 | 4:5:1:0:0 |
| 278 | 234 | 38 | HexNAc(4)Hex(5)Fuc(2)NeuAc(1) | 4 | 5 | 2 | 1 | 0 | 4:5:2:1:0 |
| 279 | 234 | 38 | HexNAc(4)Hex(5)Fuc(3)NeuAc(1) | 4 | 5 | 3 | 1 | 0 | 4:5:3:1:0 |
| 280 | 234 | 38 | HexNAc(4)Hex(5)Fuc(3) | 4 | 5 | 3 | 0 | 0 | 4:5:3:0:0 |
| 281 | 234 | 38 | HexNAc(4)Hex(5) | 4 | 5 | 0 | 0 | 0 | 4:5:0:0:0 |
| 282 | 234 | 38 | HexNAc(5)Hex(3)Fuc(1) | 5 | 3 | 1 | 0 | 0 | 5:3:1:0:0 |
| 283 | 234 | 38 | HexNAc(5)Hex(4)Fuc(1) | 5 | 4 | 1 | 0 | 0 | 5:4:1:0:0 |
| 284 | 234 | 38 | HexNAc(5)Hex(5)Fuc(1) | 5 | 5 | 1 | 0 | 0 | 5:5:1:0:0 |
| 285 | 234 | 38 | HexNAc(5)Hex(6)Fuc(1) | 5 | 6 | 1 | 0 | 0 | 5:6:1:0:0 |
| 286 | 234 | 38 | HexNAc(5)Hex(7) | 5 | 7 | 0 | 0 | 0 | 5:7:0:0:0 |
| 287 | 234 | 38 | HexNAc(6)Hex(3)Fuc(1)NeuAc(1) | 6 | 3 | 1 | 1 | 0 | 6:3:1:1:0 |
| 288 | 234 | 38 | HexNAc(6)Hex(3)Fuc(1) | 6 | 3 | 1 | 0 | 0 | 6:3:1:0:0 |
| 289 | 234 | 38 | HexNAc(6)Hex(3)Fuc(2) | 6 | 3 | 2 | 0 | 0 | 6:3:2:0:0 |
| 290 | 234 | 38 | HexNAc(6)Hex(3)Fuc(3) | 6 | 3 | 3 | 0 | 0 | 6:3:3:0:0 |
| 291 | 234 | 38 | HexNAc(6)Hex(3) | 6 | 3 | 0 | 0 | 0 | 6:3:0:0:0 |
| 292 | 234 | 38 | HexNAc(6)Hex(4)Fuc(1) | 6 | 4 | 1 | 0 | 0 | 6:4:1:0:0 |
| 293 | 234 | 38 | HexNAc(6)Hex(4) | 6 | 4 | 0 | 0 | 0 | 6:4:0:0:0 |
| 294 | 234 | 38 | HexNAc(6)Hex(5) | 6 | 5 | 0 | 0 | 0 | 6:5:0:0:0 |
| 295 | 234 | 38 | HexNAc(6)Hex(6)Fuc(1)NeuAc(2) | 6 | 6 | 1 | 2 | 0 | 6:6:1:2:0 |
| 296 | 234 | 38 | HexNAc(6)Hex(6)Fuc(1) | 6 | 6 | 1 | 0 | 0 | 6:6:1:0:0 |
| 297 | 234 | 38 | HexNAc(6)Hex(7) | 6 | 7 | 0 | 0 | 0 | 6:7:0:0:0 |
| 298 | 234 | 38 | HexNAc(7)Hex(3) | 7 | 3 | 0 | 0 | 0 | 7:3:0:0:0 |
| 299 | 234 | 38 | HexNAc(7)Hex(6)Fuc(1)NeuAc(3)NeuGc(1) | 7 | 6 | 1 | 3 | 1 | 7:6:1:3:1 |
| 300 | 234 | 38 | HexNAc(7)Hex(7)Fuc(1)NeuAc(4) | 7 | 7 | 1 | 4 | 0 | 7:7:1:4:0 |
| 301 | 234 | 38 | HexNAc(8)Hex(6) | 8 | 6 | 0 | 0 | 0 | 8:6:0:0:0 |
| 302 | 282 | 54 | HexNAc(2)Hex(5) | 2 | 5 | 0 | 0 | 0 | 2:5:0:0:0 |
| 303 | 282 | 54 | HexNAc(2)Hex(6) | 2 | 6 | 0 | 0 | 0 | 2:6:0:0:0 |
| 304 | 282 | 54 | HexNAc(2)Hex(7) | 2 | 7 | 0 | 0 | 0 | 2:7:0:0:0 |
| 305 | 282 | 54 | HexNAc(3)Hex(3) | 3 | 3 | 0 | 0 | 0 | 3:3:0:0:0 |
| 306 | 282 | 54 | HexNAc(3)Hex(5)Fuc(1) | 3 | 5 | 1 | 0 | 0 | 3:5:1:0:0 |
| 307 | 282 | 54 | HexNAc(3)Hex(5)NeuGc(1) | 3 | 5 | 0 | 0 | 1 | 3:5:0:0:1 |

| | | | | | | | | | |
|---|---|---|---|---|---|---|---|---|---|
| 308 | 282 | 54 | HexNAc(3)Hex(5) | 3 | 5 | 0 | 0 | 0 | 3:5:0:0:0 |
| 309 | 282 | 54 | HexNAc(3)Hex(6) | 3 | 6 | 0 | 0 | 0 | 3:6:0:0:0 |
| 310 | 282 | 54 | HexNAc(4)Hex(3)Fuc(1) | 4 | 3 | 1 | 0 | 0 | 4:3:1:0:0 |
| 311 | 282 | 54 | HexNAc(4)Hex(4)NeuGc(1) | 4 | 4 | 0 | 0 | 1 | 4:4:0:0:1 |
| 312 | 282 | 54 | HexNAc(4)Hex(5)Fuc(1)NeuAc(1) | 4 | 5 | 1 | 1 | 0 | 4:5:1:1:0 |
| 313 | 282 | 54 | HexNAc(4)Hex(5)Fuc(1) | 4 | 5 | 1 | 0 | 0 | 4:5:1:0:0 |
| 314 | 282 | 54 | HexNAc(4)Hex(5)Fuc(2) | 4 | 5 | 2 | 0 | 0 | 4:5:2:0:0 |
| 315 | 282 | 54 | HexNAc(4)Hex(5)Fuc(3)NeuAc(1) | 4 | 5 | 3 | 1 | 0 | 4:5:3:1:0 |
| 316 | 282 | 54 | HexNAc(4)Hex(5)Fuc(3) | 4 | 5 | 3 | 0 | 0 | 4:5:3:0:0 |
| 317 | 282 | 54 | HexNAc(4)Hex(5) | 4 | 5 | 0 | 0 | 0 | 4:5:0:0:0 |
| 318 | 282 | 54 | HexNAc(4)Hex(6)NeuAc(1) | 4 | 6 | 0 | 1 | 0 | 4:6:0:1:0 |
| 319 | 282 | 54 | HexNAc(4)Hex(6) | 4 | 6 | 0 | 0 | 0 | 4:6:0:0:0 |
| 320 | 282 | 54 | HexNAc(5)Hex(4)Fuc(1)NeuAc(1)NeuGc(1) | 5 | 4 | 1 | 1 | 1 | 5:4:1:1:1 |
| 321 | 282 | 54 | HexNAc(5)Hex(4)Fuc(1) | 5 | 4 | 1 | 0 | 0 | 5:4:1:0:0 |
| 322 | 282 | 54 | HexNAc(5)Hex(5)Fuc(1)NeuAc(1) | 5 | 5 | 1 | 1 | 0 | 5:5:1:1:0 |
| 323 | 282 | 54 | HexNAc(5)Hex(5)Fuc(1) | 5 | 5 | 1 | 0 | 0 | 5:5:1:0:0 |
| 324 | 282 | 54 | HexNAc(5)Hex(5)NeuAc(1) | 5 | 5 | 0 | 1 | 0 | 5:5:0:1:0 |
| 325 | 282 | 54 | HexNAc(5)Hex(5) | 5 | 5 | 0 | 0 | 0 | 5:5:0:0:0 |
| 326 | 282 | 54 | HexNAc(5)Hex(6)Fuc(1)NeuAc(1) | 5 | 6 | 1 | 1 | 0 | 5:6:1:1:0 |
| 327 | 282 | 54 | HexNAc(5)Hex(6)Fuc(1)NeuGc(1) | 5 | 6 | 1 | 0 | 1 | 5:6:1:0:1 |
| 328 | 282 | 54 | HexNAc(5)Hex(6)Fuc(1) | 5 | 6 | 1 | 0 | 0 | 5:6:1:0:0 |
| 329 | 282 | 54 | HexNAc(5)Hex(6)Fuc(2) | 5 | 6 | 2 | 0 | 0 | 5:6:2:0:0 |
| 330 | 282 | 54 | HexNAc(5)Hex(6)Fuc(3) | 5 | 6 | 3 | 0 | 0 | 5:6:3:0:0 |
| 331 | 282 | 54 | HexNAc(5)Hex(6)NeuAc(1) | 5 | 6 | 0 | 1 | 0 | 5:6:0:1:0 |
| 332 | 282 | 54 | HexNAc(5)Hex(6) | 5 | 6 | 0 | 0 | 0 | 5:6:0:0:0 |
| 333 | 282 | 54 | HexNAc(5)Hex(7)Fuc(1) | 5 | 7 | 1 | 0 | 0 | 5:7:1:0:0 |
| 334 | 282 | 54 | HexNAc(5)Hex(7) | 5 | 7 | 0 | 0 | 0 | 5:7:0:0:0 |
| 335 | 282 | 54 | HexNAc(6)Hex(3)Fuc(1) | 6 | 3 | 1 | 0 | 0 | 6:3:1:0:0 |
| 336 | 282 | 54 | HexNAc(6)Hex(4)Fuc(1) | 6 | 4 | 1 | 0 | 0 | 6:4:1:0:0 |
| 337 | 282 | 54 | HexNAc(6)Hex(5)Fuc(1) | 6 | 5 | 1 | 0 | 0 | 6:5:1:0:0 |
| 338 | 282 | 54 | HexNAc(6)Hex(6)Fuc(1) | 6 | 6 | 1 | 0 | 0 | 6:6:1:0:0 |
| 339 | 282 | 54 | HexNAc(6)Hex(6)Fuc(2) | 6 | 6 | 2 | 0 | 0 | 6:6:2:0:0 |
| 340 | 282 | 54 | HexNAc(6)Hex(6)Fuc(3) | 6 | 6 | 3 | 0 | 0 | 6:6:3:0:0 |
| 341 | 282 | 54 | HexNAc(6)Hex(6)NeuAc(1) | 6 | 6 | 0 | 1 | 0 | 6:6:0:1:0 |
| 342 | 282 | 54 | HexNAc(6)Hex(6) | 6 | 6 | 0 | 0 | 0 | 6:6:0:0:0 |
| 343 | 282 | 54 | HexNAc(6)Hex(7)Fuc(1)NeuAc(1) | 6 | 7 | 1 | 1 | 0 | 6:7:1:1:0 |
| 344 | 282 | 54 | HexNAc(6)Hex(7)Fuc(1)NeuGc(1) | 6 | 7 | 1 | 0 | 1 | 6:7:1:0:1 |
| 345 | 282 | 54 | HexNAc(6)Hex(7)Fuc(1) | 6 | 7 | 1 | 0 | 0 | 6:7:1:0:0 |
| 346 | 282 | 54 | HexNAc(6)Hex(7)Fuc(4) | 6 | 7 | 4 | 0 | 0 | 6:7:4:0:0 |
| 347 | 282 | 54 | HexNAc(6)Hex(7)Fuc(5) | 6 | 7 | 5 | 0 | 0 | 6:7:5:0:0 |
| 348 | 282 | 54 | HexNAc(6)Hex(7)NeuAc(1) | 6 | 7 | 0 | 1 | 0 | 6:7:0:1:0 |
| 349 | 282 | 54 | HexNAc(6)Hex(7)NeuAc(3)NeuGc(1) | 6 | 7 | 0 | 3 | 1 | 6:7:0:3:1 |
| 350 | 282 | 54 | HexNAc(6)Hex(7) | 6 | 7 | 0 | 0 | 0 | 6:7:0:0:0 |
| 351 | 282 | 54 | HexNAc(6)Hex(8)Fuc(1)NeuAc(1) | 6 | 8 | 1 | 1 | 0 | 6:8:1:1:0 |
| 352 | 282 | 54 | HexNAc(6)Hex(8)NeuAc(1) | 6 | 8 | 0 | 1 | 0 | 6:8:0:1:0 |
| 353 | 282 | 54 | HexNAc(7)Hex(6)Fuc(1) | 7 | 6 | 1 | 0 | 0 | 7:6:1:0:0 |
| 354 | 282 | 54 | HexNAc(7)Hex(7)Fuc(1) | 7 | 7 | 1 | 0 | 0 | 7:7:1:0:0 |
| 355 | 282 | 54 | HexNAc(9)Hex(3) | 9 | 3 | 0 | 0 | 0 | 9:3:0:0:0 |
| 356 | 331 | 26 | HexNAc(2)Hex(5) | 2 | 5 | 0 | 0 | 0 | 2:5:0:0:0 |
| 357 | 331 | 26 | HexNAc(2)Hex(7) | 2 | 7 | 0 | 0 | 0 | 2:7:0:0:0 |
| 358 | 331 | 26 | HexNAc(3)Hex(4)Fuc(1)NeuGc(1) | 3 | 4 | 1 | 0 | 1 | 3:4:1:0:1 |
| 359 | 331 | 26 | HexNAc(4)Hex(5)Fuc(1)NeuAc(1) | 4 | 5 | 1 | 1 | 0 | 4:5:1:1:0 |
| 360 | 331 | 26 | HexNAc(4)Hex(5)Fuc(1)NeuGc(1) | 4 | 5 | 1 | 0 | 1 | 4:5:1:0:1 |
| 361 | 331 | 26 | HexNAc(4)Hex(5)Fuc(1) | 4 | 5 | 1 | 0 | 0 | 4:5:1:0:0 |
| 362 | 331 | 26 | HexNAc(4)Hex(5)Fuc(2) | 4 | 5 | 2 | 0 | 0 | 4:5:2:0:0 |
| 363 | 331 | 26 | HexNAc(4)Hex(5)Fuc(3) | 4 | 5 | 3 | 0 | 0 | 4:5:3:0:0 |
| 364 | 331 | 26 | HexNAc(4)Hex(6)Fuc(1)NeuAc(1) | 4 | 6 | 1 | 1 | 0 | 4:6:1:1:0 |
| 365 | 331 | 26 | HexNAc(4)Hex(6)Fuc(2) | 4 | 6 | 2 | 0 | 0 | 4:6:2:0:0 |
| 366 | 331 | 26 | HexNAc(5)Hex(4)Fuc(1) | 5 | 4 | 1 | 0 | 0 | 5:4:1:0:0 |
| 367 | 331 | 26 | HexNAc(5)Hex(5)Fuc(1)NeuAc(1) | 5 | 5 | 1 | 1 | 0 | 5:5:1:1:0 |
| 368 | 331 | 26 | HexNAc(5)Hex(5)Fuc(1) | 5 | 5 | 1 | 0 | 0 | 5:5:1:0:0 |
| 369 | 331 | 26 | HexNAc(5)Hex(5)Fuc(3) | 5 | 5 | 3 | 0 | 0 | 5:5:3:0:0 |
| 370 | 331 | 26 | HexNAc(5)Hex(6)Fuc(1)NeuAc(1) | 5 | 6 | 1 | 1 | 0 | 5:6:1:1:0 |
| 371 | 331 | 26 | HexNAc(5)Hex(6)Fuc(1)NeuGc(1) | 5 | 6 | 1 | 0 | 1 | 5:6:1:0:1 |
| 372 | 331 | 26 | HexNAc(5)Hex(6)Fuc(1) | 5 | 6 | 1 | 0 | 0 | 5:6:1:0:0 |
| 373 | 331 | 26 | HexNAc(5)Hex(6)Fuc(2)NeuAc(1) | 5 | 6 | 2 | 1 | 0 | 5:6:2:1:0 |
| 374 | 331 | 26 | HexNAc(5)Hex(6)Fuc(2) | 5 | 6 | 2 | 0 | 0 | 5:6:2:0:0 |
| 375 | 331 | 26 | HexNAc(5)Hex(6)Fuc(3) | 5 | 6 | 3 | 0 | 0 | 5:6:3:0:0 |
| 376 | 331 | 26 | HexNAc(6)Hex(6)Fuc(1) | 6 | 6 | 1 | 0 | 0 | 6:6:1:0:0 |
| 377 | 331 | 26 | HexNAc(6)Hex(6)Fuc(2) | 6 | 6 | 2 | 0 | 0 | 6:6:2:0:0 |
| 378 | 331 | 26 | HexNAc(6)Hex(7)Fuc(1)NeuAc(1) | 6 | 7 | 1 | 1 | 0 | 6:7:1:1:0 |
| 379 | 331 | 26 | HexNAc(6)Hex(7)Fuc(4) | 6 | 7 | 4 | 0 | 0 | 6:7:4:0:0 |
| 380 | 331 | 26 | HexNAc(6)Hex(7)NeuAc(1) | 6 | 7 | 0 | 1 | 0 | 6:7:0:1:0 |
| 381 | 331 | 26 | HexNAc(8)Hex(9) | 8 | 9 | 0 | 0 | 0 | 8:9:0:0:0 |
| 382 | 343 | 55 | HexNAc(2)Hex(5) | 2 | 5 | 0 | 0 | 0 | 2:5:0:0:0 |
| 383 | 343 | 55 | HexNAc(2)Hex(6) | 2 | 6 | 0 | 0 | 0 | 2:6:0:0:0 |
| 384 | 343 | 55 | HexNAc(3)Hex(6)NeuAc(1) | 3 | 6 | 0 | 1 | 0 | 3:6:0:1:0 |
| 385 | 343 | 55 | HexNAc(4)Hex(4)Fuc(1) | 4 | 4 | 1 | 0 | 0 | 4:4:1:0:0 |

| | | | | | | | | | |
|---|---|---|---|---|---|---|---|---|---|
| 386 | 343 | 55 | HexNAc(4)Hex(5)Fuc(1)NeuAc(1) | 4 | 5 | 1 | 1 | 0 | 4:5:1:1:0 |
| 387 | 343 | 55 | HexNAc(4)Hex(5)Fuc(1)NeuGc(1) | 4 | 5 | 1 | 0 | 1 | 4:5:1:0:1 |
| 388 | 343 | 55 | HexNAc(4)Hex(5)Fuc(1) | 4 | 5 | 1 | 0 | 0 | 4:5:1:0:0 |
| 389 | 343 | 55 | HexNAc(4)Hex(5)Fuc(2)NeuAc(1) | 4 | 5 | 2 | 1 | 0 | 4:5:2:1:0 |
| 390 | 343 | 55 | HexNAc(4)Hex(5)Fuc(2)NeuGc(1) | 4 | 5 | 2 | 0 | 1 | 4:5:2:0:1 |
| 391 | 343 | 55 | HexNAc(4)Hex(5)Fuc(2) | 4 | 5 | 2 | 0 | 0 | 4:5:2:0:0 |
| 392 | 343 | 55 | HexNAc(4)Hex(5)Fuc(3)NeuAc(1) | 4 | 5 | 3 | 1 | 0 | 4:5:3:1:0 |
| 393 | 343 | 55 | HexNAc(4)Hex(5)Fuc(3)NeuAc(2) | 4 | 5 | 3 | 2 | 0 | 4:5:3:2:0 |
| 394 | 343 | 55 | HexNAc(4)Hex(5)Fuc(3) | 4 | 5 | 3 | 0 | 0 | 4:5:3:0:0 |
| 395 | 343 | 55 | HexNAc(4)Hex(5) | 4 | 5 | 0 | 0 | 0 | 4:5:0:0:0 |
| 396 | 343 | 55 | HexNAc(4)Hex(7)Fuc(2) | 4 | 7 | 2 | 0 | 0 | 4:7:2:0:0 |
| 397 | 343 | 55 | HexNAc(5)Hex(3)Fuc(1) | 5 | 3 | 1 | 0 | 0 | 5:3:1:0:0 |
| 398 | 343 | 55 | HexNAc(5)Hex(4)Fuc(1) | 5 | 4 | 1 | 0 | 0 | 5:4:1:0:0 |
| 399 | 343 | 55 | HexNAc(5)Hex(4)Fuc(2) | 5 | 4 | 2 | 0 | 0 | 5:4:2:0:0 |
| 400 | 343 | 55 | HexNAc(5)Hex(4) | 5 | 4 | 0 | 0 | 0 | 5:4:0:0:0 |
| 401 | 343 | 55 | HexNAc(5)Hex(5)Fuc(1)NeuGc(1) | 5 | 5 | 1 | 0 | 1 | 5:5:1:0:1 |
| 402 | 343 | 55 | HexNAc(5)Hex(5)Fuc(1) | 5 | 5 | 1 | 0 | 0 | 5:5:1:0:0 |
| 403 | 343 | 55 | HexNAc(5)Hex(5)Fuc(2) | 5 | 5 | 2 | 0 | 0 | 5:5:2:0:0 |
| 404 | 343 | 55 | HexNAc(5)Hex(5)Fuc(3) | 5 | 5 | 3 | 0 | 0 | 5:5:3:0:0 |
| 405 | 343 | 55 | HexNAc(5)Hex(5) | 5 | 5 | 0 | 0 | 0 | 5:5:0:0:0 |
| 406 | 343 | 55 | HexNAc(5)Hex(6)Fuc(1)NeuAc(1)NeuGc(1) | 5 | 6 | 1 | 1 | 1 | 5:6:1:1:1 |
| 407 | 343 | 55 | HexNAc(5)Hex(6)Fuc(1)NeuAc(1) | 5 | 6 | 1 | 1 | 0 | 5:6:1:1:0 |
| 408 | 343 | 55 | HexNAc(5)Hex(6)Fuc(1)NeuAc(2)NeuGc(1) | 5 | 6 | 1 | 2 | 1 | 5:6:1:2:1 |
| 409 | 343 | 55 | HexNAc(5)Hex(6)Fuc(1)NeuGc(1) | 5 | 6 | 1 | 0 | 1 | 5:6:1:0:1 |
| 410 | 343 | 55 | HexNAc(5)Hex(6)Fuc(2)NeuAc(1) | 5 | 6 | 2 | 1 | 0 | 5:6:2:1:0 |
| 411 | 343 | 55 | HexNAc(5)Hex(6)Fuc(2)NeuGc(1) | 5 | 6 | 2 | 0 | 1 | 5:6:2:0:1 |
| 412 | 343 | 55 | HexNAc(5)Hex(6)Fuc(2) | 5 | 6 | 2 | 0 | 0 | 5:6:2:0:0 |
| 413 | 343 | 55 | HexNAc(5)Hex(6)Fuc(3)NeuGc(1) | 5 | 6 | 3 | 0 | 1 | 5:6:3:0:1 |
| 414 | 343 | 55 | HexNAc(5)Hex(6)Fuc(3) | 5 | 6 | 3 | 0 | 0 | 5:6:3:0:0 |
| 415 | 343 | 55 | HexNAc(5)Hex(6) | 5 | 6 | 0 | 0 | 0 | 5:6:0:0:0 |
| 416 | 343 | 55 | HexNAc(5)Hex(7)Fuc(1)NeuAc(2) | 5 | 7 | 1 | 2 | 0 | 5:7:1:2:0 |
| 417 | 343 | 55 | HexNAc(6)Hex(3)Fuc(1) | 6 | 3 | 1 | 0 | 0 | 6:3:1:0:0 |
| 418 | 343 | 55 | HexNAc(6)Hex(5)Fuc(1) | 6 | 5 | 1 | 0 | 0 | 6:5:1:0:0 |
| 419 | 343 | 55 | HexNAc(6)Hex(5)Fuc(2) | 6 | 5 | 2 | 0 | 0 | 6:5:2:0:0 |
| 420 | 343 | 55 | HexNAc(6)Hex(6)Fuc(1) | 6 | 6 | 1 | 0 | 0 | 6:6:1:0:0 |
| 421 | 343 | 55 | HexNAc(6)Hex(6)Fuc(2) | 6 | 6 | 2 | 0 | 0 | 6:6:2:0:0 |
| 422 | 343 | 55 | HexNAc(6)Hex(7)Fuc(1) | 6 | 7 | 1 | 0 | 0 | 6:7:1:0:0 |
| 423 | 343 | 55 | HexNAc(6)Hex(7)Fuc(4)NeuGc(1) | 6 | 7 | 4 | 0 | 1 | 6:7:4:0:1 |
| 424 | 343 | 55 | HexNAc(6)Hex(7)Fuc(4) | 6 | 7 | 4 | 0 | 0 | 6:7:4:0:0 |
| 425 | 343 | 55 | HexNAc(6)Hex(7)Fuc(5) | 6 | 7 | 5 | 0 | 0 | 6:7:5:0:0 |
| 426 | 343 | 55 | HexNAc(6)Hex(7)NeuAc(1)NeuGc(1) | 6 | 7 | 0 | 1 | 1 | 6:7:0:1:1 |
| 427 | 343 | 55 | HexNAc(6)Hex(8)Fuc(1)NeuAc(1) | 6 | 8 | 1 | 1 | 0 | 6:8:1:1:0 |
| 428 | 343 | 55 | HexNAc(6)Hex(9)Fuc(1)NeuAc(2) | 6 | 9 | 1 | 2 | 0 | 6:9:1:2:0 |
| 429 | 343 | 55 | HexNAc(7)Hex(6)Fuc(1)NeuAc(3)NeuGc(1) | 7 | 6 | 1 | 3 | 1 | 7:6:1:3:1 |
| 430 | 343 | 55 | HexNAc(7)Hex(7)Fuc(1)NeuAc(1)NeuGc(2) | 7 | 7 | 1 | 1 | 2 | 7:7:1:1:2 |
| 431 | 343 | 55 | HexNAc(7)Hex(8)Fuc(1)NeuAc(1)NeuGc(2) | 7 | 8 | 1 | 1 | 2 | 7:8:1:1:2 |
| 432 | 343 | 55 | HexNAc(7)Hex(8)Fuc(1)NeuAc(2) | 7 | 8 | 1 | 2 | 0 | 7:8:1:2:0 |
| 433 | 343 | 55 | HexNAc(8)Hex(5) | 8 | 5 | 0 | 0 | 0 | 8:5:0:0:0 |
| 434 | 343 | 55 | HexNAc(8)Hex(9)Fuc(1)NeuAc(1)NeuGc(2) | 8 | 9 | 1 | 1 | 2 | 8:9:1:1:2 |
| 435 | 343 | 55 | HexNAc(8)Hex(9)Fuc(1)NeuAc(2)NeuGc(1) | 8 | 9 | 1 | 2 | 1 | 8:9:1:2:1 |
| 436 | 343 | 55 | HexNAc(10)Hex(10)Fuc(1) | 10 | 10 | 1 | 0 | 0 | 10:10:1:0:0 |
| 437 | 603 | 30 | HexNAc(2)Hex(5) | 2 | 5 | 0 | 0 | 0 | 2:5:0:0:0 |
| 438 | 603 | 30 | HexNAc(2)Hex(6) | 2 | 6 | 0 | 0 | 0 | 2:6:0:0:0 |
| 439 | 603 | 30 | HexNAc(2)Hex(7) | 2 | 7 | 0 | 0 | 0 | 2:7:0:0:0 |
| 440 | 603 | 30 | HexNAc(2)Hex(8) | 2 | 8 | 0 | 0 | 0 | 2:8:0:0:0 |
| 441 | 603 | 30 | HexNAc(3)Hex(4)NeuAc(1) | 3 | 4 | 0 | 1 | 0 | 3:4:0:1:0 |
| 442 | 603 | 30 | HexNAc(3)Hex(5)Fuc(1) | 3 | 5 | 1 | 0 | 0 | 3:5:1:0:0 |
| 443 | 603 | 30 | HexNAc(3)Hex(5) | 3 | 5 | 0 | 0 | 0 | 3:5:0:0:0 |
| 444 | 603 | 30 | HexNAc(3)Hex(6)Fuc(1) | 3 | 6 | 1 | 0 | 0 | 3:6:1:0:0 |
| 445 | 603 | 30 | HexNAc(3)Hex(6) | 3 | 6 | 0 | 0 | 0 | 3:6:0:0:0 |
| 446 | 603 | 30 | HexNAc(4)Hex(3)Fuc(1) | 4 | 3 | 1 | 0 | 0 | 4:3:1:0:0 |
| 447 | 603 | 30 | HexNAc(4)Hex(3)NeuAc(1) | 4 | 3 | 0 | 1 | 0 | 4:3:0:1:0 |
| 448 | 603 | 30 | HexNAc(4)Hex(4)Fuc(1) | 4 | 4 | 1 | 0 | 0 | 4:4:1:0:0 |
| 449 | 603 | 30 | HexNAc(4)Hex(4) | 4 | 4 | 0 | 0 | 0 | 4:4:0:0:0 |
| 450 | 603 | 30 | HexNAc(4)Hex(5)Fuc(1)NeuAc(1) | 4 | 5 | 1 | 1 | 0 | 4:5:1:1:0 |
| 451 | 603 | 30 | HexNAc(4)Hex(5)Fuc(1) | 4 | 5 | 1 | 0 | 0 | 4:5:1:0:0 |
| 452 | 603 | 30 | HexNAc(4)Hex(5)Fuc(2) | 4 | 5 | 2 | 0 | 0 | 4:5:2:0:0 |
| 453 | 603 | 30 | HexNAc(4)Hex(5)Fuc(3)NeuAc(1) | 4 | 5 | 3 | 1 | 0 | 4:5:3:1:0 |
| 454 | 603 | 30 | HexNAc(4)Hex(5) | 4 | 5 | 0 | 0 | 0 | 4:5:0:0:0 |
| 455 | 603 | 30 | HexNAc(5)Hex(4)Fuc(1) | 5 | 4 | 1 | 0 | 0 | 5:4:1:0:0 |
| 456 | 603 | 30 | HexNAc(5)Hex(5)Fuc(1)NeuAc(1) | 5 | 5 | 1 | 1 | 0 | 5:5:1:1:0 |
| 457 | 603 | 30 | HexNAc(5)Hex(5)Fuc(1) | 5 | 5 | 1 | 0 | 0 | 5:5:1:0:0 |
| 458 | 603 | 30 | HexNAc(5)Hex(6)Fuc(1)NeuAc(1) | 5 | 6 | 1 | 1 | 0 | 5:6:1:1:0 |
| 459 | 603 | 30 | HexNAc(5)Hex(6)Fuc(2)NeuAc(1) | 5 | 6 | 2 | 1 | 0 | 5:6:2:1:0 |
| 460 | 603 | 30 | HexNAc(5)Hex(6)Fuc(2) | 5 | 6 | 2 | 0 | 0 | 5:6:2:0:0 |
| 461 | 603 | 30 | HexNAc(5)Hex(6)Fuc(3) | 5 | 6 | 3 | 0 | 0 | 5:6:3:0:0 |
| 462 | 603 | 30 | HexNAc(5)Hex(6)NeuAc(1) | 5 | 6 | 0 | 1 | 0 | 5:6:0:1:0 |
| 463 | 603 | 30 | HexNAc(5)Hex(7)Fuc(1) | 5 | 7 | 1 | 0 | 0 | 5:7:1:0:0 |

| | | | | | | | | | |
|---|---|---|---|---|---|---|---|---|---|
| 464 | 603 | 30 | HexNAc(6)Hex(7)Fuc(1) | 6 | 7 | 1 | 0 | 0 | 6:7:1:0:0 |
| 465 | 603 | 30 | HexNAc(6)Hex(7)Fuc(4) | 6 | 7 | 4 | 0 | 0 | 6:7:4:0:0 |
| 466 | 603 | 30 | HexNAc(7)Hex(3) | 7 | 3 | 0 | 0 | 0 | 7:3:0:0:0 |
| 467 | 616 | 43 | HexNAc(2)Hex(4) | 2 | 4 | 0 | 0 | 0 | 2:4:0:0:0 |
| 468 | 616 | 43 | HexNAc(2)Hex(5) | 2 | 5 | 0 | 0 | 0 | 2:5:0:0:0 |
| 469 | 616 | 43 | HexNAc(3)Hex(4)Fuc(1)NeuAc(1) | 3 | 4 | 1 | 1 | 0 | 3:4:1:1:0 |
| 470 | 616 | 43 | HexNAc(3)Hex(5)Fuc(1) | 3 | 5 | 1 | 0 | 0 | 3:5:1:0:0 |
| 471 | 616 | 43 | HexNAc(3)Hex(5) | 3 | 5 | 0 | 0 | 0 | 3:5:0:0:0 |
| 472 | 616 | 43 | HexNAc(3)Hex(6)Fuc(1) | 3 | 6 | 1 | 0 | 0 | 3:6:1:0:0 |
| 473 | 616 | 43 | HexNAc(3)Hex(6) | 3 | 6 | 0 | 0 | 0 | 3:6:0:0:0 |
| 474 | 616 | 43 | HexNAc(4)Hex(4)Fuc(1) | 4 | 4 | 1 | 0 | 0 | 4:4:1:0:0 |
| 475 | 616 | 43 | HexNAc(4)Hex(4)NeuGc(1) | 4 | 4 | 0 | 0 | 1 | 4:4:0:0:1 |
| 476 | 616 | 43 | HexNAc(4)Hex(5)Fuc(1)NeuAc(1) | 4 | 5 | 1 | 1 | 0 | 4:5:1:1:0 |
| 477 | 616 | 43 | HexNAc(4)Hex(5)Fuc(1) | 4 | 5 | 1 | 0 | 0 | 4:5:1:0:0 |
| 478 | 616 | 43 | HexNAc(4)Hex(5)Fuc(2) | 4 | 5 | 2 | 0 | 0 | 4:5:2:0:0 |
| 479 | 616 | 43 | HexNAc(4)Hex(5)Fuc(3)NeuAc(1) | 4 | 5 | 3 | 1 | 0 | 4:5:3:1:0 |
| 480 | 616 | 43 | HexNAc(4)Hex(5) | 4 | 5 | 0 | 0 | 0 | 4:5:0:0:0 |
| 481 | 616 | 43 | HexNAc(4)Hex(6)Fuc(1)NeuAc(1) | 4 | 6 | 1 | 1 | 0 | 4:6:1:1:0 |
| 482 | 616 | 43 | HexNAc(4)Hex(6)Fuc(1) | 4 | 6 | 1 | 0 | 0 | 4:6:1:0:0 |
| 483 | 616 | 43 | HexNAc(4)Hex(6) | 4 | 6 | 0 | 0 | 0 | 4:6:0:0:0 |
| 484 | 616 | 43 | HexNAc(5)Hex(4)Fuc(1) | 5 | 4 | 1 | 0 | 0 | 5:4:1:0:0 |
| 485 | 616 | 43 | HexNAc(5)Hex(4)NeuAc(1)NeuGc(1) | 5 | 4 | 0 | 1 | 1 | 5:4:0:1:1 |
| 486 | 616 | 43 | HexNAc(5)Hex(4)NeuGc(1) | 5 | 4 | 0 | 0 | 1 | 5:4:0:0:1 |
| 487 | 616 | 43 | HexNAc(5)Hex(4) | 5 | 4 | 0 | 0 | 0 | 5:4:0:0:0 |
| 488 | 616 | 43 | HexNAc(5)Hex(5)Fuc(1) | 5 | 5 | 1 | 0 | 0 | 5:5:1:0:0 |
| 489 | 616 | 43 | HexNAc(5)Hex(5)Fuc(3) | 5 | 5 | 3 | 0 | 0 | 5:5:3:0:0 |
| 490 | 616 | 43 | HexNAc(5)Hex(5)NeuAc(1)NeuGc(1) | 5 | 5 | 0 | 1 | 1 | 5:5:0:1:1 |
| 491 | 616 | 43 | HexNAc(5)Hex(5)NeuAc(1) | 5 | 5 | 0 | 1 | 0 | 5:5:0:1:0 |
| 492 | 616 | 43 | HexNAc(5)Hex(5) | 5 | 5 | 0 | 0 | 0 | 5:5:0:0:0 |
| 493 | 616 | 43 | HexNAc(5)Hex(6)Fuc(1)NeuAc(1) | 5 | 6 | 1 | 1 | 0 | 5:6:1:1:0 |
| 494 | 616 | 43 | HexNAc(5)Hex(6)Fuc(1)NeuGc(1) | 5 | 6 | 1 | 0 | 1 | 5:6:1:0:1 |
| 495 | 616 | 43 | HexNAc(5)Hex(6)Fuc(1) | 5 | 6 | 1 | 0 | 0 | 5:6:1:0:0 |
| 496 | 616 | 43 | HexNAc(5)Hex(6)NeuAc(1)NeuGc(2) | 5 | 6 | 0 | 1 | 2 | 5:6:0:1:2 |
| 497 | 616 | 43 | HexNAc(5)Hex(6)NeuAc(1) | 5 | 6 | 0 | 1 | 0 | 5:6:0:1:0 |
| 498 | 616 | 43 | HexNAc(5)Hex(6) | 5 | 6 | 0 | 0 | 0 | 5:6:0:0:0 |
| 499 | 616 | 43 | HexNAc(5)Hex(7) | 5 | 7 | 0 | 0 | 0 | 5:7:0:0:0 |
| 500 | 616 | 43 | HexNAc(6)Hex(3)Fuc(1) | 6 | 3 | 1 | 0 | 0 | 6:3:1:0:0 |
| 501 | 616 | 43 | HexNAc(6)Hex(5)Fuc(1)NeuAc(1)NeuGc(1) | 6 | 5 | 1 | 1 | 1 | 6:5:1:1:1 |
| 502 | 616 | 43 | HexNAc(6)Hex(6)Fuc(1) | 6 | 6 | 1 | 0 | 0 | 6:6:1:0:0 |
| 503 | 616 | 43 | HexNAc(6)Hex(6)Fuc(2) | 6 | 6 | 2 | 0 | 0 | 6:6:2:0:0 |
| 504 | 616 | 43 | HexNAc(6)Hex(6)NeuAc(1) | 6 | 6 | 0 | 1 | 0 | 6:6:0:1:0 |
| 505 | 616 | 43 | HexNAc(6)Hex(7)Fuc(1)NeuAc(1) | 6 | 7 | 1 | 1 | 0 | 6:7:1:1:0 |
| 506 | 616 | 43 | HexNAc(6)Hex(7)Fuc(1) | 6 | 7 | 1 | 0 | 0 | 6:7:1:0:0 |
| 507 | 616 | 43 | HexNAc(6)Hex(7)NeuAc(1) | 6 | 7 | 0 | 1 | 0 | 6:7:0:1:0 |
| 508 | 616 | 43 | HexNAc(6)Hex(7) | 6 | 7 | 0 | 0 | 0 | 6:7:0:0:0 |
| 509 | 616 | 43 | HexNAc(7)Hex(7)Fuc(1) | 7 | 7 | 1 | 0 | 0 | 7:7:1:0:0 |
| 510 | 657 | 43 | HexNAc(2)Hex(6) | 2 | 6 | 0 | 0 | 0 | 2:6:0:0:0 |
| 511 | 657 | 43 | HexNAc(2)Hex(7) | 2 | 7 | 0 | 0 | 0 | 2:7:0:0:0 |
| 512 | 657 | 43 | HexNAc(2)Hex(9) | 2 | 9 | 0 | 0 | 0 | 2:9:0:0:0 |
| 513 | 657 | 43 | HexNAc(3)Hex(3)Fuc(2) | 3 | 3 | 2 | 0 | 0 | 3:3:2:0:0 |
| 514 | 657 | 43 | HexNAc(3)Hex(5)Fuc(1) | 3 | 5 | 1 | 0 | 0 | 3:5:1:0:0 |
| 515 | 657 | 43 | HexNAc(3)Hex(5)NeuGc(1) | 3 | 5 | 0 | 0 | 1 | 3:5:0:0:1 |
| 516 | 657 | 43 | HexNAc(3)Hex(6)Fuc(1)NeuAc(1) | 3 | 6 | 1 | 1 | 0 | 3:6:1:1:0 |
| 517 | 657 | 43 | HexNAc(3)Hex(6)Fuc(1) | 3 | 6 | 1 | 0 | 0 | 3:6:1:0:0 |
| 518 | 657 | 43 | HexNAc(4)Hex(5)Fuc(1)NeuAc(1) | 4 | 5 | 1 | 1 | 0 | 4:5:1:1:0 |
| 519 | 657 | 43 | HexNAc(4)Hex(5)Fuc(1)NeuAc(2) | 4 | 5 | 1 | 2 | 0 | 4:5:1:2:0 |
| 520 | 657 | 43 | HexNAc(4)Hex(5)Fuc(1) | 4 | 5 | 1 | 0 | 0 | 4:5:1:0:0 |
| 521 | 657 | 43 | HexNAc(4)Hex(5)Fuc(3) | 4 | 5 | 3 | 0 | 0 | 4:5:3:0:0 |
| 522 | 657 | 43 | HexNAc(4)Hex(5)NeuAc(1) | 4 | 5 | 0 | 1 | 0 | 4:5:0:1:0 |
| 523 | 657 | 43 | HexNAc(4)Hex(6)Fuc(2) | 4 | 6 | 2 | 0 | 0 | 4:6:2:0:0 |
| 524 | 657 | 43 | HexNAc(4)Hex(6)NeuAc(1) | 4 | 6 | 0 | 1 | 0 | 4:6:0:1:0 |
| 525 | 657 | 43 | HexNAc(5)Hex(4)Fuc(1)NeuAc(2) | 5 | 4 | 1 | 2 | 0 | 5:4:1:2:0 |
| 526 | 657 | 43 | HexNAc(5)Hex(4)Fuc(2)NeuAc(1) | 5 | 4 | 2 | 1 | 0 | 5:4:2:1:0 |
| 527 | 657 | 43 | HexNAc(5)Hex(5)Fuc(1)NeuAc(1) | 5 | 5 | 1 | 1 | 0 | 5:5:1:1:0 |
| 528 | 657 | 43 | HexNAc(5)Hex(5)Fuc(1) | 5 | 5 | 1 | 0 | 0 | 5:5:1:0:0 |
| 529 | 657 | 43 | HexNAc(5)Hex(5)Fuc(2) | 5 | 5 | 2 | 0 | 0 | 5:5:2:0:0 |
| 530 | 657 | 43 | HexNAc(5)Hex(5)Fuc(3) | 5 | 5 | 3 | 0 | 0 | 5:5:3:0:0 |
| 531 | 657 | 43 | HexNAc(5)Hex(5)NeuAc(1) | 5 | 5 | 0 | 1 | 0 | 5:5:0:1:0 |
| 532 | 657 | 43 | HexNAc(5)Hex(5)NeuGc(1) | 5 | 5 | 0 | 0 | 1 | 5:5:0:0:1 |
| 533 | 657 | 43 | HexNAc(5)Hex(6)Fuc(1) | 5 | 6 | 1 | 0 | 0 | 5:6:1:0:0 |
| 534 | 657 | 43 | HexNAc(5)Hex(6)Fuc(2)NeuAc(1) | 5 | 6 | 2 | 1 | 0 | 5:6:2:1:0 |
| 535 | 657 | 43 | HexNAc(5)Hex(6)Fuc(2)NeuGc(1) | 5 | 6 | 2 | 0 | 1 | 5:6:2:0:1 |
| 536 | 657 | 43 | HexNAc(5)Hex(6)Fuc(2) | 5 | 6 | 2 | 0 | 0 | 5:6:2:0:0 |
| 537 | 657 | 43 | HexNAc(5)Hex(6)NeuAc(1) | 5 | 6 | 0 | 1 | 0 | 5:6:0:1:0 |
| 538 | 657 | 43 | HexNAc(5)Hex(6) | 5 | 6 | 0 | 0 | 0 | 5:6:0:0:0 |
| 539 | 657 | 43 | HexNAc(6)Hex(3)Fuc(1)NeuAc(2) | 6 | 3 | 1 | 2 | 0 | 6:3:1:2:0 |
| 540 | 657 | 43 | HexNAc(6)Hex(3)Fuc(2)NeuGc(1) | 6 | 3 | 2 | 0 | 1 | 6:3:2:0:1 |
| 541 | 657 | 43 | HexNAc(6)Hex(4)Fuc(1) | 6 | 4 | 1 | 0 | 0 | 6:4:1:0:0 |

| | | | | | | | | | |
|---|---|---|---|---|---|---|---|---|---|
| 542 | 657 | 43 | HexNAc(6)Hex(5)Fuc(1)NeuAc(1)NeuGc(1) | 6 | 5 | 1 | 1 | 1 | 6:5:1:1:1 |
| 543 | 657 | 43 | HexNAc(6)Hex(6)Fuc(1) | 6 | 6 | 1 | 0 | 0 | 6:6:1:0:0 |
| 544 | 657 | 43 | HexNAc(6)Hex(6)Fuc(2) | 6 | 6 | 2 | 0 | 0 | 6:6:2:0:0 |
| 545 | 657 | 43 | HexNAc(6)Hex(6)Fuc(3) | 6 | 6 | 3 | 0 | 0 | 6:6:3:0:0 |
| 546 | 657 | 43 | HexNAc(6)Hex(6)NeuAc(2) | 6 | 6 | 0 | 2 | 0 | 6:6:0:2:0 |
| 547 | 657 | 43 | HexNAc(6)Hex(7)Fuc(1)NeuAc(1) | 6 | 7 | 1 | 1 | 0 | 6:7:1:1:0 |
| 548 | 657 | 43 | HexNAc(6)Hex(7)Fuc(1) | 6 | 7 | 1 | 0 | 0 | 6:7:1:0:0 |
| 549 | 657 | 43 | HexNAc(6)Hex(7)Fuc(4)NeuAc(1) | 6 | 7 | 4 | 1 | 0 | 6:7:4:1:0 |
| 550 | 657 | 43 | HexNAc(6)Hex(7)Fuc(4) | 6 | 7 | 4 | 0 | 0 | 6:7:4:0:0 |
| 551 | 657 | 43 | HexNAc(6)Hex(7)NeuAc(1) | 6 | 7 | 0 | 1 | 0 | 6:7:0:1:0 |
| 552 | 657 | 43 | HexNAc(7)Hex(8)Fuc(1) | 7 | 8 | 1 | 0 | 0 | 7:8:1:0:0 |
| 553 | 709 | 17 | HexNAc(2)Hex(4) | 2 | 4 | 0 | 0 | 0 | 2:4:0:0:0 |
| 554 | 709 | 17 | HexNAc(2)Hex(5) | 2 | 5 | 0 | 0 | 0 | 2:5:0:0:0 |
| 555 | 709 | 17 | HexNAc(2)Hex(6) | 2 | 6 | 0 | 0 | 0 | 2:6:0:0:0 |
| 556 | 709 | 17 | HexNAc(2)Hex(7) | 2 | 7 | 0 | 0 | 0 | 2:7:0:0:0 |
| 557 | 709 | 17 | HexNAc(2)Hex(8) | 2 | 8 | 0 | 0 | 0 | 2:8:0:0:0 |
| 558 | 709 | 17 | HexNAc(3)Hex(5)Fuc(1) | 3 | 5 | 1 | 0 | 0 | 3:5:1:0:0 |
| 559 | 709 | 17 | HexNAc(3)Hex(6) | 3 | 6 | 0 | 0 | 0 | 3:6:0:0:0 |
| 560 | 709 | 17 | HexNAc(4)Hex(3)NeuAc(1) | 4 | 3 | 0 | 1 | 0 | 4:3:0:1:0 |
| 561 | 709 | 17 | HexNAc(4)Hex(5)Fuc(1) | 4 | 5 | 1 | 0 | 0 | 4:5:1:0:0 |
| 562 | 709 | 17 | HexNAc(4)Hex(5) | 4 | 5 | 0 | 0 | 0 | 4:5:0:0:0 |
| 563 | 709 | 17 | HexNAc(5)Hex(6)Fuc(1) | 5 | 6 | 1 | 0 | 0 | 5:6:1:0:0 |
| 564 | 709 | 17 | HexNAc(5)Hex(6)NeuAc(2)NeuGc(1) | 5 | 6 | 0 | 2 | 1 | 5:6:0:2:1 |
| 565 | 709 | 17 | HexNAc(6)Hex(5)Fuc(1) | 6 | 5 | 1 | 0 | 0 | 6:5:1:0:0 |
| 566 | 709 | 17 | HexNAc(6)Hex(7)Fuc(4)NeuAc(1) | 6 | 7 | 4 | 1 | 0 | 6:7:4:1:0 |
| 567 | 709 | 17 | HexNAc(6)Hex(7)NeuAc(2) | 6 | 7 | 0 | 2 | 0 | 6:7:0:2:0 |
| 568 | 709 | 17 | HexNAc(6)Hex(9)Fuc(1)NeuAc(1)NeuGc(1) | 6 | 9 | 1 | 1 | 1 | 6:9:1:1:1 |
| 569 | 709 | 17 | HexNAc(6)Hex(10)Fuc(1)NeuAc(1) | 6 | 10 | 1 | 1 | 0 | 6:10:1:1:0 |
| 570 | 717 | 18 | HexNAc(2)Hex(5) | 2 | 5 | 0 | 0 | 0 | 2:5:0:0:0 |
| 571 | 717 | 18 | HexNAc(2)Hex(6) | 2 | 6 | 0 | 0 | 0 | 2:6:0:0:0 |
| 572 | 717 | 18 | HexNAc(2)Hex(7) | 2 | 7 | 0 | 0 | 0 | 2:7:0:0:0 |
| 573 | 717 | 18 | HexNAc(2)Hex(8) | 2 | 8 | 0 | 0 | 0 | 2:8:0:0:0 |
| 574 | 717 | 18 | HexNAc(2)Hex(9) | 2 | 9 | 0 | 0 | 0 | 2:9:0:0:0 |
| 575 | 717 | 18 | HexNAc(3)Hex(5)Fuc(1) | 3 | 5 | 1 | 0 | 0 | 3:5:1:0:0 |
| 576 | 717 | 18 | HexNAc(3)Hex(5) | 3 | 5 | 0 | 0 | 0 | 3:5:0:0:0 |
| 577 | 717 | 18 | HexNAc(3)Hex(6)Fuc(1) | 3 | 6 | 1 | 0 | 0 | 3:6:1:0:0 |
| 578 | 717 | 18 | HexNAc(3)Hex(6)NeuGc(1) | 3 | 6 | 0 | 0 | 1 | 3:6:0:0:1 |
| 579 | 717 | 18 | HexNAc(3)Hex(6) | 3 | 6 | 0 | 0 | 0 | 3:6:0:0:0 |
| 580 | 717 | 18 | HexNAc(4)Hex(5) | 4 | 5 | 0 | 0 | 0 | 4:5:0:0:0 |
| 581 | 717 | 18 | HexNAc(6)Hex(5)Fuc(1)NeuAc(1)NeuGc(2) | 6 | 5 | 1 | 1 | 2 | 6:5:1:1:2 |
| 582 | 717 | 18 | HexNAc(6)Hex(6)NeuAc(1) | 6 | 6 | 0 | 1 | 0 | 6:6:0:1:0 |
| 583 | 717 | 18 | HexNAc(6)Hex(7)Fuc(1)NeuAc(2) | 6 | 7 | 1 | 2 | 0 | 6:7:1:2:0 |
| 584 | 717 | 18 | HexNAc(6)Hex(8)Fuc(1)NeuAc(1) | 6 | 8 | 1 | 1 | 0 | 6:8:1:1:0 |
| 585 | 717 | 18 | HexNAc(7)Hex(8)Fuc(1)NeuAc(1)NeuGc(1) | 7 | 8 | 1 | 1 | 1 | 7:8:1:1:1 |
| 586 | 717 | 18 | HexNAc(7)Hex(8)Fuc(1)NeuAc(1) | 7 | 8 | 1 | 1 | 0 | 7:8:1:1:0 |
| 587 | 717 | 18 | HexNAc(8)Hex(9)Fuc(1)NeuAc(2)NeuGc(1) | 8 | 9 | 1 | 2 | 1 | 8:9:1:2:1 |
| 588 | 801 | 70 | HexNAc(2)Hex(4) | 2 | 4 | 0 | 0 | 0 | 2:4:0:0:0 |
| 589 | 801 | 70 | HexNAc(2)Hex(5)Fuc(1) | 2 | 5 | 1 | 0 | 0 | 2:5:1:0:0 |
| 590 | 801 | 70 | HexNAc(2)Hex(5) | 2 | 5 | 0 | 0 | 0 | 2:5:0:0:0 |
| 591 | 801 | 70 | HexNAc(2)Hex(6) | 2 | 6 | 0 | 0 | 0 | 2:6:0:0:0 |
| 592 | 801 | 70 | HexNAc(2)Hex(7) | 2 | 7 | 0 | 0 | 0 | 2:7:0:0:0 |
| 593 | 801 | 70 | HexNAc(2)Hex(8) | 2 | 8 | 0 | 0 | 0 | 2:8:0:0:0 |
| 594 | 801 | 70 | HexNAc(2)Hex(9) | 2 | 9 | 0 | 0 | 0 | 2:9:0:0:0 |
| 595 | 801 | 70 | HexNAc(3)Hex(4)Fuc(1)NeuAc(1) | 3 | 4 | 1 | 1 | 0 | 3:4:1:1:0 |
| 596 | 801 | 70 | HexNAc(3)Hex(4)Fuc(1) | 3 | 4 | 1 | 0 | 0 | 3:4:1:0:0 |
| 597 | 801 | 70 | HexNAc(3)Hex(4)Fuc(2) | 3 | 4 | 2 | 0 | 0 | 3:4:2:0:0 |
| 598 | 801 | 70 | HexNAc(3)Hex(4)NeuGc(1) | 3 | 4 | 0 | 0 | 1 | 3:4:0:0:1 |
| 599 | 801 | 70 | HexNAc(3)Hex(5)Fuc(1) | 3 | 5 | 1 | 0 | 0 | 3:5:1:0:0 |
| 600 | 801 | 70 | HexNAc(3)Hex(5)NeuAc(1) | 3 | 5 | 0 | 1 | 0 | 3:5:0:1:0 |
| 601 | 801 | 70 | HexNAc(3)Hex(5)NeuGc(1) | 3 | 5 | 0 | 0 | 1 | 3:5:0:0:1 |
| 602 | 801 | 70 | HexNAc(3)Hex(5) | 3 | 5 | 0 | 0 | 0 | 3:5:0:0:0 |
| 603 | 801 | 70 | HexNAc(3)Hex(6)Fuc(1) | 3 | 6 | 1 | 0 | 0 | 3:6:1:0:0 |
| 604 | 801 | 70 | HexNAc(3)Hex(6)NeuAc(1) | 3 | 6 | 0 | 1 | 0 | 3:6:0:1:0 |
| 605 | 801 | 70 | HexNAc(3)Hex(6)NeuGc(1) | 3 | 6 | 0 | 0 | 1 | 3:6:0:0:1 |
| 606 | 801 | 70 | HexNAc(3)Hex(6) | 3 | 6 | 0 | 0 | 0 | 3:6:0:0:0 |
| 607 | 801 | 70 | HexNAc(4)Hex(3)Fuc(1) | 4 | 3 | 1 | 0 | 0 | 4:3:1:0:0 |
| 608 | 801 | 70 | HexNAc(4)Hex(3) | 4 | 3 | 0 | 0 | 0 | 4:3:0:0:0 |
| 609 | 801 | 70 | HexNAc(4)Hex(4)Fuc(1) | 4 | 4 | 1 | 0 | 0 | 4:4:1:0:0 |
| 610 | 801 | 70 | HexNAc(4)Hex(4)Fuc(2) | 4 | 4 | 2 | 0 | 0 | 4:4:2:0:0 |
| 611 | 801 | 70 | HexNAc(4)Hex(5)Fuc(1)NeuAc(1) | 4 | 5 | 1 | 1 | 0 | 4:5:1:1:0 |
| 612 | 801 | 70 | HexNAc(4)Hex(5)Fuc(1)NeuGc(1) | 4 | 5 | 1 | 0 | 1 | 4:5:1:0:1 |
| 613 | 801 | 70 | HexNAc(4)Hex(5)Fuc(1) | 4 | 5 | 1 | 0 | 0 | 4:5:1:0:0 |
| 614 | 801 | 70 | HexNAc(4)Hex(5)Fuc(2)NeuGc(1) | 4 | 5 | 2 | 0 | 1 | 4:5:2:0:1 |
| 615 | 801 | 70 | HexNAc(4)Hex(5)Fuc(2) | 4 | 5 | 2 | 0 | 0 | 4:5:2:0:0 |
| 616 | 801 | 70 | HexNAc(4)Hex(5)Fuc(3) | 4 | 5 | 3 | 0 | 0 | 4:5:3:0:0 |
| 617 | 801 | 70 | HexNAc(4)Hex(5)NeuAc(1) | 4 | 5 | 0 | 1 | 0 | 4:5:0:1:0 |
| 618 | 801 | 70 | HexNAc(4)Hex(5)NeuGc(1) | 4 | 5 | 0 | 0 | 1 | 4:5:0:0:1 |
| 619 | 801 | 70 | HexNAc(4)Hex(5) | 4 | 5 | 0 | 0 | 0 | 4:5:0:0:0 |

| | | | | | | | | | |
|---|---|---|---|---|---|---|---|---|---|
| 620 | 801 | 70 | HexNAc(4)Hex(6)Fuc(1) | 4 | 6 | 1 | 0 | 0 | 4:6:1:0:0 |
| 621 | 801 | 70 | HexNAc(4)Hex(6)Fuc(2) | 4 | 6 | 2 | 0 | 0 | 4:6:2:0:0 |
| 622 | 801 | 70 | HexNAc(4)Hex(6)Fuc(3) | 4 | 6 | 3 | 0 | 0 | 4:6:3:0:0 |
| 623 | 801 | 70 | HexNAc(4)Hex(6)NeuAc(1) | 4 | 6 | 0 | 1 | 0 | 4:6:0:1:0 |
| 624 | 801 | 70 | HexNAc(4)Hex(6) | 4 | 6 | 0 | 0 | 0 | 4:6:0:0:0 |
| 625 | 801 | 70 | HexNAc(4)Hex(7) | 4 | 7 | 0 | 0 | 0 | 4:7:0:0:0 |
| 626 | 801 | 70 | HexNAc(5)Hex(3)Fuc(1)NeuAc(1) | 5 | 3 | 1 | 1 | 0 | 5:3:1:1:0 |
| 627 | 801 | 70 | HexNAc(5)Hex(3)Fuc(1) | 5 | 3 | 1 | 0 | 0 | 5:3:1:0:0 |
| 628 | 801 | 70 | HexNAc(5)Hex(3) | 5 | 3 | 0 | 0 | 0 | 5:3:0:0:0 |
| 629 | 801 | 70 | HexNAc(5)Hex(4)Fuc(1) | 5 | 4 | 1 | 0 | 0 | 5:4:1:0:0 |
| 630 | 801 | 70 | HexNAc(5)Hex(4)Fuc(2) | 5 | 4 | 2 | 0 | 0 | 5:4:2:0:0 |
| 631 | 801 | 70 | HexNAc(5)Hex(4) | 5 | 4 | 0 | 0 | 0 | 5:4:0:0:0 |
| 632 | 801 | 70 | HexNAc(5)Hex(5)Fuc(1) | 5 | 5 | 1 | 0 | 0 | 5:5:1:0:0 |
| 633 | 801 | 70 | HexNAc(5)Hex(5)Fuc(2) | 5 | 5 | 2 | 0 | 0 | 5:5:2:0:0 |
| 634 | 801 | 70 | HexNAc(5)Hex(5)Fuc(3) | 5 | 5 | 3 | 0 | 0 | 5:5:3:0:0 |
| 635 | 801 | 70 | HexNAc(5)Hex(5)NeuAc(1)NeuGc(1) | 5 | 5 | 0 | 1 | 1 | 5:5:0:1:1 |
| 636 | 801 | 70 | HexNAc(5)Hex(5)NeuAc(1) | 5 | 5 | 0 | 1 | 0 | 5:5:0:1:0 |
| 637 | 801 | 70 | HexNAc(5)Hex(5) | 5 | 5 | 0 | 0 | 0 | 5:5:0:0:0 |
| 638 | 801 | 70 | HexNAc(5)Hex(6)Fuc(1)NeuAc(1) | 5 | 6 | 1 | 1 | 0 | 5:6:1:1:0 |
| 639 | 801 | 70 | HexNAc(5)Hex(6)Fuc(1)NeuGc(1) | 5 | 6 | 1 | 0 | 1 | 5:6:1:0:1 |
| 640 | 801 | 70 | HexNAc(5)Hex(6)Fuc(1) | 5 | 6 | 1 | 0 | 0 | 5:6:1:0:0 |
| 641 | 801 | 70 | HexNAc(5)Hex(6)Fuc(2)NeuAc(1) | 5 | 6 | 2 | 1 | 0 | 5:6:2:1:0 |
| 642 | 801 | 70 | HexNAc(5)Hex(6)Fuc(2) | 5 | 6 | 2 | 0 | 0 | 5:6:2:0:0 |
| 643 | 801 | 70 | HexNAc(5)Hex(6)Fuc(3) | 5 | 6 | 3 | 0 | 0 | 5:6:3:0:0 |
| 644 | 801 | 70 | HexNAc(5)Hex(6)NeuAc(1) | 5 | 6 | 0 | 1 | 0 | 5:6:0:1:0 |
| 645 | 801 | 70 | HexNAc(5)Hex(6) | 5 | 6 | 0 | 0 | 0 | 5:6:0:0:0 |
| 646 | 801 | 70 | HexNAc(5)Hex(7)Fuc(1) | 5 | 7 | 1 | 0 | 0 | 5:7:1:0:0 |
| 647 | 801 | 70 | HexNAc(6)Hex(3)Fuc(1) | 6 | 3 | 1 | 0 | 0 | 6:3:1:0:0 |
| 648 | 801 | 70 | HexNAc(6)Hex(4)Fuc(1) | 6 | 4 | 1 | 0 | 0 | 6:4:1:0:0 |
| 649 | 801 | 70 | HexNAc(6)Hex(5)Fuc(1) | 6 | 5 | 1 | 0 | 0 | 6:5:1:0:0 |
| 650 | 801 | 70 | HexNAc(6)Hex(6)Fuc(1)NeuAc(1) | 6 | 6 | 1 | 1 | 0 | 6:6:1:1:0 |
| 651 | 801 | 70 | HexNAc(6)Hex(6)Fuc(1) | 6 | 6 | 1 | 0 | 0 | 6:6:1:0:0 |
| 652 | 801 | 70 | HexNAc(6)Hex(6)Fuc(3) | 6 | 6 | 3 | 0 | 0 | 6:6:3:0:0 |
| 653 | 801 | 70 | HexNAc(6)Hex(6)NeuAc(1) | 6 | 6 | 0 | 1 | 0 | 6:6:0:1:0 |
| 654 | 801 | 70 | HexNAc(6)Hex(6) | 6 | 6 | 0 | 0 | 0 | 6:6:0:0:0 |
| 655 | 801 | 70 | HexNAc(6)Hex(7)Fuc(1)NeuAc(3)NeuGc(1) | 6 | 7 | 1 | 3 | 1 | 6:7:1:3:1 |
| 656 | 801 | 70 | HexNAc(6)Hex(9)Fuc(1)NeuAc(1)NeuGc(1) | 6 | 9 | 1 | 1 | 1 | 6:9:1:1:1 |
| 657 | 801 | 70 | HexNAc(7)Hex(3) | 7 | 3 | 0 | 0 | 0 | 7:3:0:0:0 |
| 658 | 1074 | 98 | HexNAc(2)Hex(3) | 2 | 3 | 0 | 0 | 0 | 2:3:0:0:0 |
| 659 | 1074 | 98 | HexNAc(2)Hex(4) | 2 | 4 | 0 | 0 | 0 | 2:4:0:0:0 |
| 660 | 1074 | 98 | HexNAc(2)Hex(5) | 2 | 5 | 0 | 0 | 0 | 2:5:0:0:0 |
| 661 | 1074 | 98 | HexNAc(2)Hex(6) | 2 | 6 | 0 | 0 | 0 | 2:6:0:0:0 |
| 662 | 1074 | 98 | HexNAc(2)Hex(7) | 2 | 7 | 0 | 0 | 0 | 2:7:0:0:0 |
| 663 | 1074 | 98 | HexNAc(2)Hex(8) | 2 | 8 | 0 | 0 | 0 | 2:8:0:0:0 |
| 664 | 1074 | 98 | HexNAc(2)Hex(9) | 2 | 9 | 0 | 0 | 0 | 2:9:0:0:0 |
| 665 | 1074 | 98 | HexNAc(2)Hex(11) | 2 | 11 | 0 | 0 | 0 | 2:11:0:0:0 |
| 666 | 1074 | 98 | HexNAc(2)Fuc(1) | 2 | 0 | 1 | 0 | 0 | 2:0:1:0:0 |
| 667 | 1074 | 98 | HexNAc(3)Hex(3)Fuc(1) | 3 | 3 | 1 | 0 | 0 | 3:3:1:0:0 |
| 668 | 1074 | 98 | HexNAc(3)Hex(3) | 3 | 3 | 0 | 0 | 0 | 3:3:0:0:0 |
| 669 | 1074 | 98 | HexNAc(3)Hex(4)Fuc(1)NeuAc(1) | 3 | 4 | 1 | 1 | 0 | 3:4:1:1:0 |
| 670 | 1074 | 98 | HexNAc(3)Hex(4)Fuc(1)NeuGc(1) | 3 | 4 | 1 | 0 | 1 | 3:4:1:0:1 |
| 671 | 1074 | 98 | HexNAc(3)Hex(4)Fuc(1) | 3 | 4 | 1 | 0 | 0 | 3:4:1:0:0 |
| 672 | 1074 | 98 | HexNAc(3)Hex(4)Fuc(2) | 3 | 4 | 2 | 0 | 0 | 3:4:2:0:0 |
| 673 | 1074 | 98 | HexNAc(3)Hex(4) | 3 | 4 | 0 | 0 | 0 | 3:4:0:0:0 |
| 674 | 1074 | 98 | HexNAc(3)Hex(5)Fuc(1) | 3 | 5 | 1 | 0 | 0 | 3:5:1:0:0 |
| 675 | 1074 | 98 | HexNAc(3)Hex(5)NeuAc(1) | 3 | 5 | 0 | 1 | 0 | 3:5:0:1:0 |
| 676 | 1074 | 98 | HexNAc(3)Hex(5)NeuGc(1) | 3 | 5 | 0 | 0 | 1 | 3:5:0:0:1 |
| 677 | 1074 | 98 | HexNAc(3)Hex(5) | 3 | 5 | 0 | 0 | 0 | 3:5:0:0:0 |
| 678 | 1074 | 98 | HexNAc(3)Hex(6)Fuc(1)NeuGc(1) | 3 | 6 | 1 | 0 | 1 | 3:6:1:0:1 |
| 679 | 1074 | 98 | HexNAc(3)Hex(6)Fuc(1) | 3 | 6 | 1 | 0 | 0 | 3:6:1:0:0 |
| 680 | 1074 | 98 | HexNAc(3)Hex(6)NeuAc(1) | 3 | 6 | 0 | 1 | 0 | 3:6:0:1:0 |
| 681 | 1074 | 98 | HexNAc(3)Hex(6)NeuGc(1) | 3 | 6 | 0 | 0 | 1 | 3:6:0:0:1 |
| 682 | 1074 | 98 | HexNAc(3)Hex(6) | 3 | 6 | 0 | 0 | 0 | 3:6:0:0:0 |
| 683 | 1074 | 98 | HexNAc(4)Hex(3)Fuc(1) | 4 | 3 | 1 | 0 | 0 | 4:3:1:0:0 |
| 684 | 1074 | 98 | HexNAc(4)Hex(3) | 4 | 3 | 0 | 0 | 0 | 4:3:0:0:0 |
| 685 | 1074 | 98 | HexNAc(4)Hex(4)Fuc(1) | 4 | 4 | 1 | 0 | 0 | 4:4:1:0:0 |
| 686 | 1074 | 98 | HexNAc(4)Hex(4) | 4 | 4 | 0 | 0 | 0 | 4:4:0:0:0 |
| 687 | 1074 | 98 | HexNAc(4)Hex(5)Fuc(1)NeuAc(1)NeuGc(1) | 4 | 5 | 1 | 1 | 1 | 4:5:1:1:1 |
| 688 | 1074 | 98 | HexNAc(4)Hex(5)Fuc(1)NeuAc(1) | 4 | 5 | 1 | 1 | 0 | 4:5:1:1:0 |
| 689 | 1074 | 98 | HexNAc(4)Hex(5)Fuc(1)NeuAc(2) | 4 | 5 | 1 | 2 | 0 | 4:5:1:2:0 |
| 690 | 1074 | 98 | HexNAc(4)Hex(5)Fuc(1)NeuGc(1) | 4 | 5 | 1 | 0 | 1 | 4:5:1:0:1 |
| 691 | 1074 | 98 | HexNAc(4)Hex(5)Fuc(1) | 4 | 5 | 1 | 0 | 0 | 4:5:1:0:0 |
| 692 | 1074 | 98 | HexNAc(4)Hex(5)Fuc(2)NeuGc(1) | 4 | 5 | 2 | 0 | 1 | 4:5:2:0:1 |
| 693 | 1074 | 98 | HexNAc(4)Hex(5)Fuc(2) | 4 | 5 | 2 | 0 | 0 | 4:5:2:0:0 |
| 694 | 1074 | 98 | HexNAc(4)Hex(5)Fuc(3)NeuAc(1) | 4 | 5 | 3 | 1 | 0 | 4:5:3:1:0 |
| 695 | 1074 | 98 | HexNAc(4)Hex(5)Fuc(3)NeuAc(2) | 4 | 5 | 3 | 2 | 0 | 4:5:3:2:0 |
| 696 | 1074 | 98 | HexNAc(4)Hex(5)Fuc(3) | 4 | 5 | 3 | 0 | 0 | 4:5:3:0:0 |
| 697 | 1074 | 98 | HexNAc(4)Hex(5)NeuAc(1)NeuGc(1) | 4 | 5 | 0 | 1 | 1 | 4:5:0:1:1 |

| | | | | | | | | | |
|---|---|---|---|---|---|---|---|---|---|
| 698 | 1074 | 98 | HexNAc(4)Hex(5)NeuAc(2) | 4 | 5 | 0 | 2 | 0 | 4:5:0:2:0 |
| 699 | 1074 | 98 | HexNAc(4)Hex(5)NeuGc(1) | 4 | 5 | 0 | 0 | 1 | 4:5:0:0:1 |
| 700 | 1074 | 98 | HexNAc(4)Hex(5) | 4 | 5 | 0 | 0 | 0 | 4:5:0:0:0 |
| 701 | 1074 | 98 | HexNAc(4)Hex(6)Fuc(1) | 4 | 6 | 1 | 0 | 0 | 4:6:1:0:0 |
| 702 | 1074 | 98 | HexNAc(4)Hex(6)Fuc(2) | 4 | 6 | 2 | 0 | 0 | 4:6:2:0:0 |
| 703 | 1074 | 98 | HexNAc(4)Hex(6)Fuc(3) | 4 | 6 | 3 | 0 | 0 | 4:6:3:0:0 |
| 704 | 1074 | 98 | HexNAc(4)Hex(6)NeuAc(1) | 4 | 6 | 0 | 1 | 0 | 4:6:0:1:0 |
| 705 | 1074 | 98 | HexNAc(4)Hex(6) | 4 | 6 | 0 | 0 | 0 | 4:6:0:0:0 |
| 706 | 1074 | 98 | HexNAc(4)Hex(7)Fuc(2) | 4 | 7 | 2 | 0 | 0 | 4:7:2:0:0 |
| 707 | 1074 | 98 | HexNAc(4)Hex(7)NeuAc(1) | 4 | 7 | 0 | 1 | 0 | 4:7:0:1:0 |
| 708 | 1074 | 98 | HexNAc(4)Hex(7) | 4 | 7 | 0 | 0 | 0 | 4:7:0:0:0 |
| 709 | 1074 | 98 | HexNAc(5)Hex(3)Fuc(1) | 5 | 3 | 1 | 0 | 0 | 5:3:1:0:0 |
| 710 | 1074 | 98 | HexNAc(5)Hex(3) | 5 | 3 | 0 | 0 | 0 | 5:3:0:0:0 |
| 711 | 1074 | 98 | HexNAc(5)Hex(4)Fuc(1) | 5 | 4 | 1 | 0 | 0 | 5:4:1:0:0 |
| 712 | 1074 | 98 | HexNAc(5)Hex(4)Fuc(2) | 5 | 4 | 2 | 0 | 0 | 5:4:2:0:0 |
| 713 | 1074 | 98 | HexNAc(5)Hex(4)NeuAc(1) | 5 | 4 | 0 | 1 | 0 | 5:4:0:1:0 |
| 714 | 1074 | 98 | HexNAc(5)Hex(4)NeuGc(2) | 5 | 4 | 0 | 0 | 2 | 5:4:0:0:2 |
| 715 | 1074 | 98 | HexNAc(5)Hex(4) | 5 | 4 | 0 | 0 | 0 | 5:4:0:0:0 |
| 716 | 1074 | 98 | HexNAc(5)Hex(5)Fuc(1)NeuAc(1) | 5 | 5 | 1 | 1 | 0 | 5:5:1:1:0 |
| 717 | 1074 | 98 | HexNAc(5)Hex(5)Fuc(1) | 5 | 5 | 1 | 0 | 0 | 5:5:1:0:0 |
| 718 | 1074 | 98 | HexNAc(5)Hex(5)Fuc(2) | 5 | 5 | 2 | 0 | 0 | 5:5:2:0:0 |
| 719 | 1074 | 98 | HexNAc(5)Hex(5)Fuc(3) | 5 | 5 | 3 | 0 | 0 | 5:5:3:0:0 |
| 720 | 1074 | 98 | HexNAc(5)Hex(5) | 5 | 5 | 0 | 0 | 0 | 5:5:0:0:0 |
| 721 | 1074 | 98 | HexNAc(5)Hex(6)Fuc(1)NeuAc(1) | 5 | 6 | 1 | 1 | 0 | 5:6:1:1:0 |
| 722 | 1074 | 98 | HexNAc(5)Hex(6)Fuc(1)NeuGc(1) | 5 | 6 | 1 | 0 | 1 | 5:6:1:0:1 |
| 723 | 1074 | 98 | HexNAc(5)Hex(6)Fuc(1)NeuGc(2) | 5 | 6 | 1 | 0 | 2 | 5:6:1:0:2 |
| 724 | 1074 | 98 | HexNAc(5)Hex(6)Fuc(1) | 5 | 6 | 1 | 0 | 0 | 5:6:1:0:0 |
| 725 | 1074 | 98 | HexNAc(5)Hex(6)Fuc(2)NeuAc(1) | 5 | 6 | 2 | 1 | 0 | 5:6:2:1:0 |
| 726 | 1074 | 98 | HexNAc(5)Hex(6)Fuc(2)NeuGc(1) | 5 | 6 | 2 | 0 | 1 | 5:6:2:0:1 |
| 727 | 1074 | 98 | HexNAc(5)Hex(6)Fuc(2) | 5 | 6 | 2 | 0 | 0 | 5:6:2:0:0 |
| 728 | 1074 | 98 | HexNAc(5)Hex(6)Fuc(3) | 5 | 6 | 3 | 0 | 0 | 5:6:3:0:0 |
| 729 | 1074 | 98 | HexNAc(5)Hex(6)NeuAc(1) | 5 | 6 | 0 | 1 | 0 | 5:6:0:1:0 |
| 730 | 1074 | 98 | HexNAc(5)Hex(6)NeuGc(1) | 5 | 6 | 0 | 0 | 1 | 5:6:0:0:1 |
| 731 | 1074 | 98 | HexNAc(5)Hex(6) | 5 | 6 | 0 | 0 | 0 | 5:6:0:0:0 |
| 732 | 1074 | 98 | HexNAc(5)Hex(7)Fuc(1) | 5 | 7 | 1 | 0 | 0 | 5:7:1:0:0 |
| 733 | 1074 | 98 | HexNAc(6)Hex(3)Fuc(1) | 6 | 3 | 1 | 0 | 0 | 6:3:1:0:0 |
| 734 | 1074 | 98 | HexNAc(6)Hex(4)Fuc(1) | 6 | 4 | 1 | 0 | 0 | 6:4:1:0:0 |
| 735 | 1074 | 98 | HexNAc(6)Hex(5)Fuc(1) | 6 | 5 | 1 | 0 | 0 | 6:5:1:0:0 |
| 736 | 1074 | 98 | HexNAc(6)Hex(5)Fuc(2) | 6 | 5 | 2 | 0 | 0 | 6:5:2:0:0 |
| 737 | 1074 | 98 | HexNAc(6)Hex(6)Fuc(1) | 6 | 6 | 1 | 0 | 0 | 6:6:1:0:0 |
| 738 | 1074 | 98 | HexNAc(6)Hex(6)Fuc(2) | 6 | 6 | 2 | 0 | 0 | 6:6:2:0:0 |
| 739 | 1074 | 98 | HexNAc(6)Hex(6)Fuc(3) | 6 | 6 | 3 | 0 | 0 | 6:6:3:0:0 |
| 740 | 1074 | 98 | HexNAc(6)Hex(6)NeuAc(1) | 6 | 6 | 0 | 1 | 0 | 6:6:0:1:0 |
| 741 | 1074 | 98 | HexNAc(6)Hex(6) | 6 | 6 | 0 | 0 | 0 | 6:6:0:0:0 |
| 742 | 1074 | 98 | HexNAc(6)Hex(7)Fuc(1)NeuAc(1) | 6 | 7 | 1 | 1 | 0 | 6:7:1:1:0 |
| 743 | 1074 | 98 | HexNAc(6)Hex(7)Fuc(1)NeuGc(1) | 6 | 7 | 1 | 0 | 1 | 6:7:1:0:1 |
| 744 | 1074 | 98 | HexNAc(6)Hex(7)Fuc(1) | 6 | 7 | 1 | 0 | 0 | 6:7:1:0:0 |
| 745 | 1074 | 98 | HexNAc(6)Hex(7)Fuc(4)NeuGc(1) | 6 | 7 | 4 | 0 | 1 | 6:7:4:0:1 |
| 746 | 1074 | 98 | HexNAc(6)Hex(7)Fuc(4) | 6 | 7 | 4 | 0 | 0 | 6:7:4:0:0 |
| 747 | 1074 | 98 | HexNAc(6)Hex(7)Fuc(5) | 6 | 7 | 5 | 0 | 0 | 6:7:5:0:0 |
| 748 | 1074 | 98 | HexNAc(6)Hex(7)NeuAc(1) | 6 | 7 | 0 | 1 | 0 | 6:7:0:1:0 |
| 749 | 1074 | 98 | HexNAc(6)Hex(7)NeuGc(1) | 6 | 7 | 0 | 0 | 1 | 6:7:0:0:1 |
| 750 | 1074 | 98 | HexNAc(6)Hex(7) | 6 | 7 | 0 | 0 | 0 | 6:7:0:0:0 |
| 751 | 1074 | 98 | HexNAc(6)Hex(8)Fuc(1)NeuAc(1) | 6 | 8 | 1 | 1 | 0 | 6:8:1:1:0 |
| 752 | 1074 | 98 | HexNAc(7)Hex(3)Fuc(1) | 7 | 3 | 1 | 0 | 0 | 7:3:1:0:0 |
| 753 | 1074 | 98 | HexNAc(7)Hex(6)Fuc(1) | 7 | 6 | 1 | 0 | 0 | 7:6:1:0:0 |
| 754 | 1074 | 98 | HexNAc(7)Hex(7)Fuc(1) | 7 | 7 | 1 | 0 | 0 | 7:7:1:0:0 |
| 755 | 1074 | 98 | HexNAc(7)Hex(7) | 7 | 7 | 0 | 0 | 0 | 7:7:0:0:0 |
| 756 | 1098 | 91 | HexNAc(2)Hex(5) | 2 | 5 | 0 | 0 | 0 | 2:5:0:0:0 |
| 757 | 1098 | 91 | HexNAc(2)Hex(6) | 2 | 6 | 0 | 0 | 0 | 2:6:0:0:0 |
| 758 | 1098 | 91 | HexNAc(2)Hex(7) | 2 | 7 | 0 | 0 | 0 | 2:7:0:0:0 |
| 759 | 1098 | 91 | HexNAc(2)Hex(8) | 2 | 8 | 0 | 0 | 0 | 2:8:0:0:0 |
| 760 | 1098 | 91 | HexNAc(2)Hex(9) | 2 | 9 | 0 | 0 | 0 | 2:9:0:0:0 |
| 761 | 1098 | 91 | HexNAc(3)Hex(3) | 3 | 3 | 0 | 0 | 0 | 3:3:0:0:0 |
| 762 | 1098 | 91 | HexNAc(3)Hex(4)NeuGc(1) | 3 | 4 | 0 | 0 | 1 | 3:4:0:0:1 |
| 763 | 1098 | 91 | HexNAc(3)Hex(4) | 3 | 4 | 0 | 0 | 0 | 3:4:0:0:0 |
| 764 | 1098 | 91 | HexNAc(3)Hex(5)Fuc(1) | 3 | 5 | 1 | 0 | 0 | 3:5:1:0:0 |
| 765 | 1098 | 91 | HexNAc(3)Hex(5)NeuAc(1) | 3 | 5 | 0 | 1 | 0 | 3:5:0:1:0 |
| 766 | 1098 | 91 | HexNAc(3)Hex(5)NeuGc(1) | 3 | 5 | 0 | 0 | 1 | 3:5:0:0:1 |
| 767 | 1098 | 91 | HexNAc(3)Hex(5) | 3 | 5 | 0 | 0 | 0 | 3:5:0:0:0 |
| 768 | 1098 | 91 | HexNAc(3)Hex(6)Fuc(1)NeuGc(1) | 3 | 6 | 1 | 0 | 1 | 3:6:1:0:1 |
| 769 | 1098 | 91 | HexNAc(3)Hex(6)Fuc(1) | 3 | 6 | 1 | 0 | 0 | 3:6:1:0:0 |
| 770 | 1098 | 91 | HexNAc(3)Hex(6)NeuAc(1) | 3 | 6 | 0 | 1 | 0 | 3:6:0:1:0 |
| 771 | 1098 | 91 | HexNAc(3)Hex(6)NeuGc(1) | 3 | 6 | 0 | 0 | 1 | 3:6:0:0:1 |
| 772 | 1098 | 91 | HexNAc(3)Hex(6) | 3 | 6 | 0 | 0 | 0 | 3:6:0:0:0 |
| 773 | 1098 | 91 | HexNAc(4)Hex(3) | 4 | 3 | 0 | 0 | 0 | 4:3:0:0:0 |
| 774 | 1098 | 91 | HexNAc(4)Hex(4)Fuc(1) | 4 | 4 | 1 | 0 | 0 | 4:4:1:0:0 |
| 775 | 1098 | 91 | HexNAc(4)Hex(4) | 4 | 4 | 0 | 0 | 0 | 4:4:0:0:0 |

| | | | | | | | | | |
|---|---|---|---|---|---|---|---|---|---|
| 776 | 1098 | 91 | HexNAc(4)Hex(5)Fuc(1)NeuAc(1) | 4 | 5 | 1 | 1 | 0 | 4:5:1:1:0 |
| 777 | 1098 | 91 | HexNAc(4)Hex(5)Fuc(1)NeuGc(1) | 4 | 5 | 1 | 0 | 1 | 4:5:1:0:1 |
| 778 | 1098 | 91 | HexNAc(4)Hex(5)Fuc(1) | 4 | 5 | 1 | 0 | 0 | 4:5:1:0:0 |
| 779 | 1098 | 91 | HexNAc(4)Hex(5)Fuc(2) | 4 | 5 | 2 | 0 | 0 | 4:5:2:0:0 |
| 780 | 1098 | 91 | HexNAc(4)Hex(5)Fuc(3)NeuAc(1)NeuGc(1) | 4 | 5 | 3 | 1 | 1 | 4:5:3:1:1 |
| 781 | 1098 | 91 | HexNAc(4)Hex(5)Fuc(3)NeuAc(1) | 4 | 5 | 3 | 1 | 0 | 4:5:3:1:0 |
| 782 | 1098 | 91 | HexNAc(4)Hex(5)Fuc(3)NeuAc(2) | 4 | 5 | 3 | 2 | 0 | 4:5:3:2:0 |
| 783 | 1098 | 91 | HexNAc(4)Hex(5)Fuc(3) | 4 | 5 | 3 | 0 | 0 | 4:5:3:0:0 |
| 784 | 1098 | 91 | HexNAc(4)Hex(5)NeuAc(1) | 4 | 5 | 0 | 1 | 0 | 4:5:0:1:0 |
| 785 | 1098 | 91 | HexNAc(4)Hex(5)NeuGc(1) | 4 | 5 | 0 | 0 | 1 | 4:5:0:0:1 |
| 786 | 1098 | 91 | HexNAc(4)Hex(5)NeuGc(2) | 4 | 5 | 0 | 0 | 2 | 4:5:0:0:2 |
| 787 | 1098 | 91 | HexNAc(4)Hex(5) | 4 | 5 | 0 | 0 | 0 | 4:5:0:0:0 |
| 788 | 1098 | 91 | HexNAc(4)Hex(6)Fuc(1) | 4 | 6 | 1 | 0 | 0 | 4:6:1:0:0 |
| 789 | 1098 | 91 | HexNAc(4)Hex(6)Fuc(2) | 4 | 6 | 2 | 0 | 0 | 4:6:2:0:0 |
| 790 | 1098 | 91 | HexNAc(4)Hex(6)NeuAc(1) | 4 | 6 | 0 | 1 | 0 | 4:6:0:1:0 |
| 791 | 1098 | 91 | HexNAc(4)Hex(6)NeuGc(1) | 4 | 6 | 0 | 0 | 1 | 4:6:0:0:1 |
| 792 | 1098 | 91 | HexNAc(4)Hex(6) | 4 | 6 | 0 | 0 | 0 | 4:6:0:0:0 |
| 793 | 1098 | 91 | HexNAc(4)Hex(7)Fuc(1) | 4 | 7 | 1 | 0 | 0 | 4:7:1:0:0 |
| 794 | 1098 | 91 | HexNAc(4)Hex(7)NeuAc(1) | 4 | 7 | 0 | 1 | 0 | 4:7:0:1:0 |
| 795 | 1098 | 91 | HexNAc(4)Hex(7) | 4 | 7 | 0 | 0 | 0 | 4:7:0:0:0 |
| 796 | 1098 | 91 | HexNAc(5)Hex(3)Fuc(1) | 5 | 3 | 1 | 0 | 0 | 5:3:1:0:0 |
| 797 | 1098 | 91 | HexNAc(5)Hex(3) | 5 | 3 | 0 | 0 | 0 | 5:3:0:0:0 |
| 798 | 1098 | 91 | HexNAc(5)Hex(4)Fuc(1) | 5 | 4 | 1 | 0 | 0 | 5:4:1:0:0 |
| 799 | 1098 | 91 | HexNAc(5)Hex(4) | 5 | 4 | 0 | 0 | 0 | 5:4:0:0:0 |
| 800 | 1098 | 91 | HexNAc(5)Hex(5)Fuc(1)NeuGc(1) | 5 | 5 | 1 | 0 | 1 | 5:5:1:0:1 |
| 801 | 1098 | 91 | HexNAc(5)Hex(5)Fuc(1) | 5 | 5 | 1 | 0 | 0 | 5:5:1:0:0 |
| 802 | 1098 | 91 | HexNAc(5)Hex(5)Fuc(2) | 5 | 5 | 2 | 0 | 0 | 5:5:2:0:0 |
| 803 | 1098 | 91 | HexNAc(5)Hex(5)NeuGc(1) | 5 | 5 | 0 | 0 | 1 | 5:5:0:0:1 |
| 804 | 1098 | 91 | HexNAc(5)Hex(5)NeuGc(2) | 5 | 5 | 0 | 0 | 2 | 5:5:0:0:2 |
| 805 | 1098 | 91 | HexNAc(5)Hex(5) | 5 | 5 | 0 | 0 | 0 | 5:5:0:0:0 |
| 806 | 1098 | 91 | HexNAc(5)Hex(6)Fuc(1)NeuAc(1) | 5 | 6 | 1 | 1 | 0 | 5:6:1:1:0 |
| 807 | 1098 | 91 | HexNAc(5)Hex(6)Fuc(1)NeuGc(1) | 5 | 6 | 1 | 0 | 1 | 5:6:1:0:1 |
| 808 | 1098 | 91 | HexNAc(5)Hex(6)Fuc(1)NeuGc(2) | 5 | 6 | 1 | 0 | 2 | 5:6:1:0:2 |
| 809 | 1098 | 91 | HexNAc(5)Hex(6)Fuc(1) | 5 | 6 | 1 | 0 | 0 | 5:6:1:0:0 |
| 810 | 1098 | 91 | HexNAc(5)Hex(6)Fuc(2)NeuGc(1) | 5 | 6 | 2 | 0 | 1 | 5:6:2:0:1 |
| 811 | 1098 | 91 | HexNAc(5)Hex(6)Fuc(2) | 5 | 6 | 2 | 0 | 0 | 5:6:2:0:0 |
| 812 | 1098 | 91 | HexNAc(5)Hex(6)Fuc(3) | 5 | 6 | 3 | 0 | 0 | 5:6:3:0:0 |
| 813 | 1098 | 91 | HexNAc(5)Hex(6)NeuAc(1)NeuGc(1) | 5 | 6 | 0 | 1 | 1 | 5:6:0:1:1 |
| 814 | 1098 | 91 | HexNAc(5)Hex(6)NeuAc(1) | 5 | 6 | 0 | 1 | 0 | 5:6:0:1:0 |
| 815 | 1098 | 91 | HexNAc(5)Hex(6)NeuGc(1) | 5 | 6 | 0 | 0 | 1 | 5:6:0:0:1 |
| 816 | 1098 | 91 | HexNAc(5)Hex(6)NeuGc(2) | 5 | 6 | 0 | 0 | 2 | 5:6:0:0:2 |
| 817 | 1098 | 91 | HexNAc(5)Hex(6) | 5 | 6 | 0 | 0 | 0 | 5:6:0:0:0 |
| 818 | 1098 | 91 | HexNAc(5)Hex(7) | 5 | 7 | 0 | 0 | 0 | 5:7:0:0:0 |
| 819 | 1098 | 91 | HexNAc(6)Hex(3)Fuc(1) | 6 | 3 | 1 | 0 | 0 | 6:3:1:0:0 |
| 820 | 1098 | 91 | HexNAc(6)Hex(3)Fuc(2) | 6 | 3 | 2 | 0 | 0 | 6:3:2:0:0 |
| 821 | 1098 | 91 | HexNAc(6)Hex(3) | 6 | 3 | 0 | 0 | 0 | 6:3:0:0:0 |
| 822 | 1098 | 91 | HexNAc(6)Hex(4)Fuc(1) | 6 | 4 | 1 | 0 | 0 | 6:4:1:0:0 |
| 823 | 1098 | 91 | HexNAc(6)Hex(4) | 6 | 4 | 0 | 0 | 0 | 6:4:0:0:0 |
| 824 | 1098 | 91 | HexNAc(6)Hex(5)Fuc(1) | 6 | 5 | 1 | 0 | 0 | 6:5:1:0:0 |
| 825 | 1098 | 91 | HexNAc(6)Hex(5) | 6 | 5 | 0 | 0 | 0 | 6:5:0:0:0 |
| 826 | 1098 | 91 | HexNAc(6)Hex(6)Fuc(1)NeuAc(1)NeuGc(2) | 6 | 6 | 1 | 1 | 2 | 6:6:1:1:2 |
| 827 | 1098 | 91 | HexNAc(6)Hex(6)Fuc(1)NeuGc(2) | 6 | 6 | 1 | 0 | 2 | 6:6:1:0:2 |
| 828 | 1098 | 91 | HexNAc(6)Hex(6)Fuc(1) | 6 | 6 | 1 | 0 | 0 | 6:6:1:0:0 |
| 829 | 1098 | 91 | HexNAc(6)Hex(6)NeuAc(1)NeuGc(1) | 6 | 6 | 0 | 1 | 1 | 6:6:0:1:1 |
| 830 | 1098 | 91 | HexNAc(6)Hex(6) | 6 | 6 | 0 | 0 | 0 | 6:6:0:0:0 |
| 831 | 1098 | 91 | HexNAc(6)Hex(7)Fuc(1)NeuAc(1) | 6 | 7 | 1 | 1 | 0 | 6:7:1:1:0 |
| 832 | 1098 | 91 | HexNAc(6)Hex(7)Fuc(1)NeuGc(1) | 6 | 7 | 1 | 0 | 1 | 6:7:1:0:1 |
| 833 | 1098 | 91 | HexNAc(6)Hex(7)Fuc(1) | 6 | 7 | 1 | 0 | 0 | 6:7:1:0:0 |
| 834 | 1098 | 91 | HexNAc(6)Hex(7)Fuc(4) | 6 | 7 | 4 | 0 | 0 | 6:7:4:0:0 |
| 835 | 1098 | 91 | HexNAc(6)Hex(7)NeuAc(1)NeuGc(1) | 6 | 7 | 0 | 1 | 1 | 6:7:0:1:1 |
| 836 | 1098 | 91 | HexNAc(6)Hex(7)NeuAc(1) | 6 | 7 | 0 | 1 | 0 | 6:7:0:1:0 |
| 837 | 1098 | 91 | HexNAc(6)Hex(7)NeuAc(2)NeuGc(1) | 6 | 7 | 0 | 2 | 1 | 6:7:0:2:1 |
| 838 | 1098 | 91 | HexNAc(6)Hex(7)NeuGc(1) | 6 | 7 | 0 | 0 | 1 | 6:7:0:0:1 |
| 839 | 1098 | 91 | HexNAc(6)Hex(7)NeuGc(2) | 6 | 7 | 0 | 0 | 2 | 6:7:0:0:2 |
| 840 | 1098 | 91 | HexNAc(6)Hex(7) | 6 | 7 | 0 | 0 | 0 | 6:7:0:0:0 |
| 841 | 1098 | 91 | HexNAc(6)Hex(8)Fuc(1)NeuAc(1) | 6 | 8 | 1 | 1 | 0 | 6:8:1:1:0 |
| 842 | 1098 | 91 | HexNAc(6)Hex(8)NeuAc(1) | 6 | 8 | 0 | 1 | 0 | 6:8:0:1:0 |
| 843 | 1098 | 91 | HexNAc(6)Hex(9) | 6 | 9 | 0 | 0 | 0 | 6:9:0:0:0 |
| 844 | 1098 | 91 | HexNAc(7)Hex(6) | 7 | 6 | 0 | 0 | 0 | 7:6:0:0:0 |
| 845 | 1098 | 91 | HexNAc(7)Hex(7)Fuc(1) | 7 | 7 | 1 | 0 | 0 | 7:7:1:0:0 |
| 846 | 1098 | 91 | HexNAc(7)Hex(7) | 7 | 7 | 0 | 0 | 0 | 7:7:0:0:0 |
| 847 | 1134 | 41 | HexNAc(2)Hex(5) | 2 | 5 | 0 | 0 | 0 | 2:5:0:0:0 |
| 848 | 1134 | 41 | HexNAc(2)Hex(6) | 2 | 6 | 0 | 0 | 0 | 2:6:0:0:0 |
| 849 | 1134 | 41 | HexNAc(2)Hex(7) | 2 | 7 | 0 | 0 | 0 | 2:7:0:0:0 |
| 850 | 1134 | 41 | HexNAc(2)Hex(8) | 2 | 8 | 0 | 0 | 0 | 2:8:0:0:0 |
| 851 | 1134 | 41 | HexNAc(3)Hex(3)Fuc(1) | 3 | 3 | 1 | 0 | 0 | 3:3:1:0:0 |
| 852 | 1134 | 41 | HexNAc(3)Hex(4)Fuc(1) | 3 | 4 | 1 | 0 | 0 | 3:4:1:0:0 |
| 853 | 1134 | 41 | HexNAc(3)Hex(5)Fuc(1) | 3 | 5 | 1 | 0 | 0 | 3:5:1:0:0 |

| | | | | | | | | | |
|---|---|---|---|---|---|---|---|---|---|
| 854 | 1134 | 41 | HexNAc(3)Hex(5)NeuGc(1) | 3 | 5 | 0 | 0 | 1 | 3:5:0:0:1 |
| 855 | 1134 | 41 | HexNAc(3)Hex(5) | 3 | 5 | 0 | 0 | 0 | 3:5:0:0:0 |
| 856 | 1134 | 41 | HexNAc(3)Hex(6)Fuc(1) | 3 | 6 | 1 | 0 | 0 | 3:6:1:0:0 |
| 857 | 1134 | 41 | HexNAc(3)Hex(6)NeuAc(1) | 3 | 6 | 0 | 1 | 0 | 3:6:0:1:0 |
| 858 | 1134 | 41 | HexNAc(3)Hex(6) | 3 | 6 | 0 | 0 | 0 | 3:6:0:0:0 |
| 859 | 1134 | 41 | HexNAc(4)Hex(3)Fuc(1) | 4 | 3 | 1 | 0 | 0 | 4:3:1:0:0 |
| 860 | 1134 | 41 | HexNAc(4)Hex(4)Fuc(1) | 4 | 4 | 1 | 0 | 0 | 4:4:1:0:0 |
| 861 | 1134 | 41 | HexNAc(4)Hex(5)Fuc(1) | 4 | 5 | 1 | 0 | 0 | 4:5:1:0:0 |
| 862 | 1134 | 41 | HexNAc(4)Hex(5)Fuc(2) | 4 | 5 | 2 | 0 | 0 | 4:5:2:0:0 |
| 863 | 1134 | 41 | HexNAc(4)Hex(6)Fuc(1) | 4 | 6 | 1 | 0 | 0 | 4:6:1:0:0 |
| 864 | 1134 | 41 | HexNAc(4)Hex(6)Fuc(2) | 4 | 6 | 2 | 0 | 0 | 4:6:2:0:0 |
| 865 | 1134 | 41 | HexNAc(4)Hex(6)NeuAc(1) | 4 | 6 | 0 | 1 | 0 | 4:6:0:1:0 |
| 866 | 1134 | 41 | HexNAc(5)Hex(3)Fuc(1)NeuAc(1) | 5 | 3 | 1 | 1 | 0 | 5:3:1:1:0 |
| 867 | 1134 | 41 | HexNAc(5)Hex(3) | 5 | 3 | 0 | 0 | 0 | 5:3:0:0:0 |
| 868 | 1134 | 41 | HexNAc(5)Hex(4)Fuc(1) | 5 | 4 | 1 | 0 | 0 | 5:4:1:0:0 |
| 869 | 1134 | 41 | HexNAc(5)Hex(5)Fuc(1)NeuAc(1) | 5 | 5 | 1 | 1 | 0 | 5:5:1:1:0 |
| 870 | 1134 | 41 | HexNAc(5)Hex(6)Fuc(1)NeuAc(1) | 5 | 6 | 1 | 1 | 0 | 5:6:1:1:0 |
| 871 | 1134 | 41 | HexNAc(5)Hex(6)Fuc(1)NeuGc(1) | 5 | 6 | 1 | 0 | 1 | 5:6:1:0:1 |
| 872 | 1134 | 41 | HexNAc(5)Hex(6)Fuc(1)NeuGc(2) | 5 | 6 | 1 | 0 | 2 | 5:6:1:0:2 |
| 873 | 1134 | 41 | HexNAc(5)Hex(6)Fuc(1) | 5 | 6 | 1 | 0 | 0 | 5:6:1:0:0 |
| 874 | 1134 | 41 | HexNAc(5)Hex(6)Fuc(3)NeuGc(1) | 5 | 6 | 3 | 0 | 1 | 5:6:3:0:1 |
| 875 | 1134 | 41 | HexNAc(5)Hex(6)NeuAc(1) | 5 | 6 | 0 | 1 | 0 | 5:6:0:1:0 |
| 876 | 1134 | 41 | HexNAc(5)Hex(6) | 5 | 6 | 0 | 0 | 0 | 5:6:0:0:0 |
| 877 | 1134 | 41 | HexNAc(5)Hex(8)Fuc(1) | 5 | 8 | 1 | 0 | 0 | 5:8:1:0:0 |
| 878 | 1134 | 41 | HexNAc(6)Hex(4)Fuc(1) | 6 | 4 | 1 | 0 | 0 | 6:4:1:0:0 |
| 879 | 1134 | 41 | HexNAc(6)Hex(5)Fuc(1) | 6 | 5 | 1 | 0 | 0 | 6:5:1:0:0 |
| 880 | 1134 | 41 | HexNAc(6)Hex(5)Fuc(2) | 6 | 5 | 2 | 0 | 0 | 6:5:2:0:0 |
| 881 | 1134 | 41 | HexNAc(6)Hex(6)Fuc(1) | 6 | 6 | 1 | 0 | 0 | 6:6:1:0:0 |
| 882 | 1134 | 41 | HexNAc(6)Hex(6)NeuAc(1) | 6 | 6 | 0 | 1 | 0 | 6:6:0:1:0 |
| 883 | 1134 | 41 | HexNAc(6)Hex(7)Fuc(1) | 6 | 7 | 1 | 0 | 0 | 6:7:1:0:0 |
| 884 | 1134 | 41 | HexNAc(6)Hex(7) | 6 | 7 | 0 | 0 | 0 | 6:7:0:0:0 |
| 885 | 1134 | 41 | HexNAc(7)Hex(8)Fuc(1)NeuAc(2)NeuGc(1) | 7 | 8 | 1 | 2 | 1 | 7:8:1:2:1 |
| 886 | 1134 | 41 | HexNAc(8)Hex(5) | 8 | 5 | 0 | 0 | 0 | 8:5:0:0:0 |
| 887 | 1134 | 41 | HexNAc(9)Hex(3)Fuc(1) | 9 | 3 | 1 | 0 | 0 | 9:3:1:0:0 |
| 888 | 1158 | 1 | HexNAc(6)Hex(7)Fuc(1) | 6 | 7 | 1 | 0 | 0 | 6:7:1:0:0 |
| 889 | 1173 | 6 | HexNAc(5)Hex(6)Fuc(1) | 5 | 6 | 1 | 0 | 0 | 5:6:1:0:0 |
| 890 | 1173 | 6 | HexNAc(6)Hex(3)Fuc(1) | 6 | 3 | 1 | 0 | 0 | 6:3:1:0:0 |
| 891 | 1173 | 6 | HexNAc(6)Hex(7)Fuc(1)NeuGc(1) | 6 | 7 | 1 | 0 | 1 | 6:7:1:0:1 |
| 892 | 1173 | 6 | HexNAc(6)Hex(7)Fuc(1) | 6 | 7 | 1 | 0 | 0 | 6:7:1:0:0 |
| 893 | 1173 | 6 | HexNAc(6)Hex(7)Fuc(4) | 6 | 7 | 4 | 0 | 0 | 6:7:4:0:0 |
| 894 | 1173 | 6 | HexNAc(6)Hex(7)Fuc(5) | 6 | 7 | 5 | 0 | 0 | 6:7:5:0:0 |
| 895 | 1194 | 25 | HexNAc(4)Hex(3)Fuc(1) | 4 | 3 | 1 | 0 | 0 | 4:3:1:0:0 |
| 896 | 1194 | 25 | HexNAc(4)Hex(5)Fuc(1) | 4 | 5 | 1 | 0 | 0 | 4:5:1:0:0 |
| 897 | 1194 | 25 | HexNAc(4)Hex(5)Fuc(3) | 4 | 5 | 3 | 0 | 0 | 4:5:3:0:0 |
| 898 | 1194 | 25 | HexNAc(4)Hex(5) | 4 | 5 | 0 | 0 | 0 | 4:5:0:0:0 |
| 899 | 1194 | 25 | HexNAc(5)Hex(3)Fuc(1) | 5 | 3 | 1 | 0 | 0 | 5:3:1:0:0 |
| 900 | 1194 | 25 | HexNAc(5)Hex(4)Fuc(1) | 5 | 4 | 1 | 0 | 0 | 5:4:1:0:0 |
| 901 | 1194 | 25 | HexNAc(5)Hex(5)Fuc(1) | 5 | 5 | 1 | 0 | 0 | 5:5:1:0:0 |
| 902 | 1194 | 25 | HexNAc(5)Hex(6)Fuc(1)NeuGc(1) | 5 | 6 | 1 | 0 | 1 | 5:6:1:0:1 |
| 903 | 1194 | 25 | HexNAc(5)Hex(6)Fuc(1) | 5 | 6 | 1 | 0 | 0 | 5:6:1:0:0 |
| 904 | 1194 | 25 | HexNAc(5)Hex(6)Fuc(2)NeuAc(1) | 5 | 6 | 2 | 1 | 0 | 5:6:2:1:0 |
| 905 | 1194 | 25 | HexNAc(5)Hex(6)Fuc(2) | 5 | 6 | 2 | 0 | 0 | 5:6:2:0:0 |
| 906 | 1194 | 25 | HexNAc(5)Hex(6)NeuAc(1)NeuGc(1) | 5 | 6 | 0 | 1 | 1 | 5:6:0:1:1 |
| 907 | 1194 | 25 | HexNAc(5)Hex(6) | 5 | 6 | 0 | 0 | 0 | 5:6:0:0:0 |
| 908 | 1194 | 25 | HexNAc(5)Hex(7)Fuc(1)NeuAc(1) | 5 | 7 | 1 | 1 | 0 | 5:7:1:1:0 |
| 909 | 1194 | 25 | HexNAc(6)Hex(3)Fuc(1) | 6 | 3 | 1 | 0 | 0 | 6:3:1:0:0 |
| 910 | 1194 | 25 | HexNAc(6)Hex(4)Fuc(1) | 6 | 4 | 1 | 0 | 0 | 6:4:1:0:0 |
| 911 | 1194 | 25 | HexNAc(6)Hex(5)Fuc(1) | 6 | 5 | 1 | 0 | 0 | 6:5:1:0:0 |
| 912 | 1194 | 25 | HexNAc(6)Hex(7)Fuc(1)NeuAc(1) | 6 | 7 | 1 | 1 | 0 | 6:7:1:1:0 |
| 913 | 1194 | 25 | HexNAc(6)Hex(7)Fuc(1)NeuGc(1) | 6 | 7 | 1 | 0 | 1 | 6:7:1:0:1 |
| 914 | 1194 | 25 | HexNAc(6)Hex(7)Fuc(1) | 6 | 7 | 1 | 0 | 0 | 6:7:1:0:0 |
| 915 | 1194 | 25 | HexNAc(6)Hex(7)Fuc(4)NeuGc(1) | 6 | 7 | 4 | 0 | 1 | 6:7:4:0:1 |
| 916 | 1194 | 25 | HexNAc(6)Hex(7)Fuc(4) | 6 | 7 | 4 | 0 | 0 | 6:7:4:0:0 |
| 917 | 1194 | 25 | HexNAc(6)Hex(7)NeuAc(1)NeuGc(1) | 6 | 7 | 0 | 1 | 1 | 6:7:0:1:1 |
| 918 | 1194 | 25 | HexNAc(6)Hex(7)NeuAc(1) | 6 | 7 | 0 | 1 | 0 | 6:7:0:1:0 |
| 919 | 1194 | 25 | HexNAc(6)Hex(7) | 6 | 7 | 0 | 0 | 0 | 6:7:0:0:0 |

Supplementary Table S10. Assignment of glycopeptide signals to SARS CoV-2-derived prptides by Glyco-RIDGE method

| Enzyme | Cluster_no | No. of glycopeptides / cluster | Peak_no | Charge | Scan_no | Mass of glycopeptide signal | Retention time of glycopeptide peak | Signal intensity | Mass of glycopeptide signal obserbed | Core peptide no | Peptide position from | Peptide position to | Peptide sequence | Type and position of variable modifications (1: ammonia-loss (N-term C), 2: Delta:H(1)N(-1)18O(1) (N), 3: pyroGlu (N-term Q), 4: oxidation (M)) | Glycosylated site (sequon) | No. of glycosylated site | Calculated mass of IGOT-labeled peptide | Mass of core peptide | Signal intensity of core peptide | RT of core peptide | Rank | No. of Hex | No. of HexNAc | No. of Fuc | No. of NeuAc | No. of NeuGc | 0: usual, 1: unusual composition | Glycan point | Calculated mass of glycopeptide (Da) | delta (Da) | delta (ppm) | delta (RT) |
|---|---|---|---|---|---|---|---|---|---|---|---|---|---|---|---|---|---|---|---|---|---|---|---|---|---|---|---|---|---|---|---|---|
| Trypsin | 49 | 38 | 285 | 1 | 4040 | 3035.2274 | 44.4361 | 470,502 | 3034.219575 | 32 | 14 | 21 | QCVNLTTR | 3.00020000.0 | 17(NLT) | 1 | 976.4534 | 973.4651387 | 4303470 | 49.37 | 1 | 5 | 5 | 3 | 0 | 0 | 0 | 1 | 3034.220478 | -0.0009 | -0.29 | -4.94 |
| Trypsin | 49 | 38 | 367 | 1 | 4038 | 3238.306 | 44.3402 | 392,213 | 3237.298175 | 32 | 14 | 21 | QCVNLTTR | 3.00020000.0 | 17(NLT) | 1 | 976.4534 | 973.4651387 | 4303470 | 49.37 | 1 | 5 | 5 | 3 | 0 | 0 | 0 | 1 | 3237.299851 | -0.0017 | -0.51 | -5.03 |
| Trypsin | 49 | 38 | 371 | 1 | 4047 | 2889.1687 | 44.7547 | 389,754 | 2888.160875 | 32 | 14 | 21 | QCVNLTTR | 3.00020000.0 | 17(NLT) | 1 | 976.4534 | 973.4651387 | 4303470 | 49.37 | 1 | 5 | 4 | 2 | 0 | 0 | 0 | 1 | 2888.162569 | -0.0017 | -0.58 | -4.62 |
| Trypsin | 49 | 38 | 416 | 1 | 4051 | 2743.1118 | 44.9668 | 348,968 | 2742.103975 | 32 | 14 | 21 | QCVNLTTR | 3.00020000.0 | 17(NLT) | 1 | 976.4534 | 973.4651387 | 4303470 | 49.37 | 1 | 5 | 4 | 1 | 0 | 0 | 0 | 1 | 2742.10466 | -0.0007 | -0.24 | -4.41 |
| Trypsin | 49 | 38 | 432 | 1 | 4041 | 3076.2543 | 44.4774 | 336,646 | 3075.246475 | 32 | 14 | 21 | QCVNLTTR | 3.00020000.0 | 17(NLT) | 1 | 976.4534 | 973.4651387 | 4303470 | 49.37 | 1 | 4 | 5 | 3 | 0 | 0 | 0 | 0 | 3075.247027 | -0.0006 | -0.17 | -4.90 |
| Trypsin | 49 | 38 | 469 | 1 | 4052 | 2727.1185 | 45.0089 | 310,474 | 2726.110675 | 32 | 14 | 21 | QCVNLTTR | 3.00020000.0 | 17(NLT) | 1 | 976.4534 | 973.4651387 | 4303470 | 49.37 | 1 | 4 | 4 | 2 | 0 | 0 | 0 | 1 | 2726.109745 | 0.0009 | 0.34 | -4.36 |
| Trypsin | 49 | 38 | 558 | 1 | 4051 | 2686.0905 | 44.9668 | 257,762 | 2685.082675 | 32 | 14 | 21 | QCVNLTTR | 3.00020000.0 | 17(NLT) | 1 | 976.4534 | 973.4651387 | 4303470 | 49.37 | 1 | 5 | 3 | 2 | 0 | 0 | 0 | 0 | 2685.083196 | -0.0005 | -0.19 | -4.41 |
| Trypsin | 49 | 38 | 602 | 1 | 4047 | 3092.2469 | 44.7547 | 237,359 | 3091.239075 | 32 | 14 | 21 | QCVNLTTR | 3.00020000.0 | 17(NLT) | 1 | 976.4534 | 973.4651387 | 4303470 | 49.37 | 1 | 5 | 5 | 2 | 0 | 0 | 0 | 1 | 3091.241942 | -0.0029 | -0.92 | -4.62 |
| Trypsin | 49 | 38 | 647 | 1 | 4051 | 2702.0831 | 44.9668 | 216,331 | 2701.075275 | 32 | 14 | 21 | QCVNLTTR | 3.00020000.0 | 17(NLT) | 1 | 976.4534 | 973.4651387 | 4303470 | 49.37 | 1 | 6 | 3 | 1 | 0 | 0 | 0 | 1 | 2701.078111 | -0.0028 | -1.04 | -4.41 |
| Trypsin | 49 | 38 | 685 | 1 | 4056 | 2946.1914 | 45.2022 | 206,835 | 2945.183575 | 32 | 14 | 21 | QCVNLTTR | 3.00020000.0 | 17(NLT) | 1 | 976.4534 | 973.4651387 | 4303470 | 49.37 | 1 | 5 | 5 | 1 | 0 | 0 | 0 | 1 | 2945.184033 | -0.0005 | -0.15 | -4.17 |
| Trypsin | 49 | 38 | 699 | 1 | 4054 | 2784.1368 | 45.1105 | 202,127 | 2783.128975 | 32 | 14 | 21 | QCVNLTTR | 3.00020000.0 | 17(NLT) | 1 | 976.4534 | 973.4651387 | 4303470 | 49.37 | 1 | 4 | 5 | 1 | 0 | 0 | 0 | 1 | 2783.131209 | -0.0022 | -0.8 | -4.26 |
| Trypsin | 49 | 38 | 784 | 1 | 4060 | 2581.0601 | 45.3831 | 179,216 | 2580.052275 | 32 | 14 | 21 | QCVNLTTR | 3.00020000.0 | 17(NLT) | 1 | 976.4534 | 973.4651387 | 4303470 | 49.37 | 1 | 4 | 4 | 1 | 0 | 0 | 0 | 1 | 2580.051836 | 0.0004 | 0.17 | -3.99 |
| Trypsin | 49 | 38 | 806 | 1 | 4047 | 2930.1966 | 44.7547 | 175,063 | 2929.188775 | 32 | 14 | 21 | QCVNLTTR | 3.00020000.0 | 17(NLT) | 1 | 976.4534 | 973.4651387 | 4303470 | 49.37 | 1 | 4 | 5 | 2 | 0 | 0 | 0 | 1 | 2929.189118 | -0.0003 | -0.11 | -4.62 |
| Trypsin | 49 | 38 | 952 | 1 | 4064 | 2419.0085 | 45.564 | 148,944 | 2418.000675 | 32 | 14 | 21 | QCVNLTTR | 3.00020000.0 | 17(NLT) | 1 | 976.4534 | 973.4651387 | 4303470 | 49.37 | 1 | 3 | 4 | 1 | 0 | 0 | 0 | 1 | 2417.999012 | 0.0017 | 0.68 | -3.81 |
| Trypsin | 49 | 38 | 1390 | 1 | 4047 | 3457.3777 | 44.7547 | 98,322 | 3456.369875 | 32 | 14 | 21 | QCVNLTTR | 3.00020000.0 | 17(NLT) | 1 | 976.4534 | 973.4651387 | 4303470 | 49.37 | 1 | 6 | 6 | 2 | 0 | 0 | 0 | 1 | 3456.374139 | -0.0043 | -1.23 | -4.62 |
| Trypsin | 49 | 38 | 1482 | 1 | 4047 | 2848.1396 | 44.7547 | 91,941 | 2847.131775 | 32 | 14 | 21 | QCVNLTTR | 3.00020000.0 | 17(NLT) | 1 | 976.4534 | 973.4651387 | 4303470 | 49.37 | 1 | 6 | 3 | 2 | 0 | 0 | 0 | 0 | 2847.13602 | -0.0042 | -1.49 | -4.62 |
| Trypsin | 49 | 38 | 1524 | 1 | 4060 | 2622.0868 | 45.3831 | 89,872 | 2621.078975 | 32 | 14 | 21 | QCVNLTTR | 3.00020000.0 | 17(NLT) | 1 | 976.4534 | 973.4651387 | 4303470 | 49.37 | 1 | 3 | 5 | 1 | 0 | 0 | 0 | 1 | 2621.078385 | 0.0006 | 0.22 | -3.99 |
| Trypsin | 49 | 38 | 1613 | 1 | 4047 | 3108.238 | 44.7547 | 84,208 | 3107.230175 | 32 | 14 | 21 | QCVNLTTR | 3.00020000.0 | 17(NLT) | 1 | 976.4534 | 973.4651387 | 4303470 | 49.37 | 1 | 6 | 5 | 1 | 0 | 0 | 0 | 1 | 3107.236857 | -0.0067 | -2.15 | -4.62 |
| Trypsin | 49 | 38 | 1649 | 1 | 4141 | 3050.201 | 49.1615 | 82,100 | 3049.193175 | 32 | 14 | 21 | QCVNLTTR | 3.00020000.0 | 17(NLT) | 1 | 976.4534 | 973.4651387 | 4303470 | 49.37 | 1 | 5 | 4 | 1 | 0 | 1 | 0 | 1 | 3049.194991 | -0.0018 | -0.59 | -0.21 |
| Trypsin | 49 | 38 | 1649 | 1 | 4141 | 3050.201 | 49.1615 | 82,100 | 3049.193175 | 32 | 14 | 21 | QCVNLTTR | 3.00020000.0 | 17(NLT) | 1 | 976.4534 | 973.4651387 | 4303470 | 49.37 | 2 | 6 | 4 | 0 | 1 | 0 | 0 | 1 | 3049.194992 | -0.0018 | -0.59 | -0.21 |
| Trypsin | 49 | 38 | 1749 | 1 | 4057 | 2540.0311 | 45.2456 | 76,954 | 2539.023275 | 32 | 14 | 21 | QCVNLTTR | 3.00020000.0 | 17(NLT) | 1 | 976.4534 | 973.4651387 | 4303470 | 49.37 | 1 | 5 | 3 | 1 | 0 | 0 | 0 | 1 | 2539.025287 | -0.0020 | -0.79 | -4.13 |
| Trypsin | 49 | 38 | 1774 | 1 | 4055 | 2825.1694 | 45.1523 | 75,261 | 2824.161575 | 32 | 14 | 21 | QCVNLTTR | 3.00020000.0 | 17(NLT) | 1 | 976.4534 | 973.4651387 | 4303470 | 49.37 | 1 | 3 | 6 | 1 | 0 | 0 | 0 | 1 | 2824.157758 | 0.0038 | 1.35 | -4.22 |
| Trypsin | 49 | 38 | 2049 | 1 | 4134 | 3196.2553 | 48.8406 | 63,416 | 3195.247475 | 32 | 14 | 21 | QCVNLTTR | 3.00020000.0 | 17(NLT) | 1 | 976.4534 | 973.4651387 | 4303470 | 49.37 | 1 | 5 | 4 | 2 | 0 | 1 | 0 | 1 | 3195.2529 | -0.0054 | -1.69 | -0.53 |
| Trypsin | 49 | 38 | 2049 | 1 | 4134 | 3196.2553 | 48.8406 | 63,416 | 3195.247475 | 32 | 14 | 21 | QCVNLTTR | 3.00020000.0 | 17(NLT) | 1 | 976.4534 | 973.4651387 | 4303470 | 49.37 | 2 | 6 | 4 | 1 | 1 | 0 | 0 | 1 | 3195.252901 | -0.0054 | -1.69 | -0.53 |
| Trypsin | 49 | 38 | 2051 | 1 | 4053 | 3311.3243 | 45.0593 | 63,370 | 3310.316475 | 32 | 14 | 21 | QCVNLTTR | 3.00020000.0 | 17(NLT) | 1 | 976.4534 | 973.4651387 | 4303470 | 49.37 | 1 | 6 | 6 | 1 | 0 | 0 | 0 | 1 | 3310.31623 | 0.0002 | 0.07 | -4.31 |
| Trypsin | 49 | 38 | 2086 | 1 | 4040 | 3603.4383 | 44.4361 | 62,241 | 3602.430475 | 32 | 14 | 21 | QCVNLTTR | 3.00020000.0 | 17(NLT) | 1 | 976.4534 | 973.4651387 | 4303470 | 49.37 | 1 | 6 | 6 | 3 | 0 | 0 | 0 | 1 | 3602.432048 | -0.0016 | -0.43 | -4.94 |
| Trypsin | 49 | 38 | 2679 | 1 | 4138 | 3618.4127 | 49.0255 | 45,985 | 3617.404875 | 32 | 14 | 21 | QCVNLTTR | 3.00020000.0 | 17(NLT) | 1 | 976.4534 | 973.4651387 | 4303470 | 49.37 | 1 | 7 | 6 | 0 | 1 | 0 | 0 | 1 | 3617.406562 | -0.0017 | -0.46 | -0.35 |
| Trypsin | 49 | 38 | 2679 | 1 | 4138 | 3618.4127 | 49.0255 | 45,985 | 3617.404875 | 32 | 14 | 21 | QCVNLTTR | 3.00020000.0 | 17(NLT) | 1 | 976.4534 | 973.4651387 | 4303470 | 49.37 | 2 | 6 | 6 | 1 | 0 | 1 | 0 | 0 | 3617.406561 | -0.0017 | -0.46 | -0.35 |
| Trypsin | 49 | 38 | 2794 | 1 | 4035 | 3400.3524 | 44.2001 | 43,385 | 3399.344575 | 32 | 14 | 21 | QCVNLTTR | 3.00020000.0 | 17(NLT) | 1 | 976.4534 | 973.4651387 | 4303470 | 49.37 | 1 | 6 | 5 | 3 | 0 | 0 | 0 | 1 | 3399.352675 | -0.0081 | -2.38 | -5.17 |
| Trypsin | 49 | 38 | 3188 | 1 | 4130 | 3237.279 | 48.6478 | 36,448 | 3236.271175 | 32 | 14 | 21 | QCVNLTTR | 3.00020000.0 | 17(NLT) | 1 | 976.4534 | 973.4651387 | 4303470 | 49.37 | 1 | 4 | 5 | 2 | 0 | 1 | 0 | 1 | 3236.279449 | -0.0083 | -2.55 | -0.73 |
| Trypsin | 49 | 38 | 3188 | 1 | 4130 | 3237.279 | 48.6478 | 36,448 | 3236.271175 | 32 | 14 | 21 | QCVNLTTR | 3.00020000.0 | 17(NLT) | 1 | 976.4534 | 973.4651387 | 4303470 | 49.37 | 2 | 5 | 5 | 1 | 1 | 0 | 0 | 1 | 3236.27945 | -0.0083 | -2.55 | -0.73 |
| Trypsin | 49 | 38 | 3223 | 1 | 4143 | 2888.1485 | 49.2547 | 35,930 | 2887.140675 | 32 | 14 | 21 | QCVNLTTR | 3.00020000.0 | 17(NLT) | 1 | 976.4534 | 973.4651387 | 4303470 | 49.37 | 1 | 4 | 4 | 1 | 0 | 1 | 0 | 1 | 2887.142167 | -0.0015 | -0.51 | -0.12 |
| Trypsin | 49 | 38 | 3223 | 1 | 4143 | 2888.1485 | 49.2547 | 35,930 | 2887.140675 | 32 | 14 | 21 | QCVNLTTR | 3.00020000.0 | 17(NLT) | 1 | 976.4534 | 973.4651387 | 4303470 | 49.37 | 2 | 5 | 4 | 0 | 1 | 0 | 0 | 1 | 2887.142168 | -0.0015 | -0.51 | -0.12 |
| Trypsin | 49 | 38 | 3267 | 1 | 4058 | 2556.0259 | 45.288 | 35,070 | 2555.018075 | 32 | 14 | 21 | QCVNLTTR | 3.00020000.0 | 17(NLT) | 1 | 976.4534 | 973.4651387 | 4303470 | 49.37 | 1 | 6 | 3 | 0 | 0 | 0 | 0 | 1 | 2555.020202 | -0.0021 | -0.83 | -4.09 |
| Trypsin | 49 | 38 | 3271 | 1 | 4036 | 3473.3767 | 44.2457 | 35,029 | 3472.368875 | 32 | 14 | 21 | QCVNLTTR | 3.00020000.0 | 17(NLT) | 1 | 976.4534 | 973.4651387 | 4303470 | 49.37 | 1 | 7 | 6 | 1 | 0 | 0 | 0 | 1 | 3472.369054 | -0.0002 | -0.05 | -5.13 |
| Trypsin | 49 | 38 | 3349 | 1 | 4030 | 3749.4972 | 43.9588 | 33,856 | 3748.489375 | 32 | 14 | 21 | QCVNLTTR | 3.00020000.0 | 17(NLT) | 1 | 976.4534 | 973.4651387 | 4303470 | 49.37 | 1 | 6 | 6 | 4 | 0 | 0 | 0 | 0 | 3748.489957 | -0.0006 | -0.15 | -5.42 |
| Trypsin | 49 | 38 | 3511 | 1 | 4132 | 3415.3279 | 48.7441 | 31,701 | 3414.320075 | 32 | 14 | 21 | QCVNLTTR | 3.00020000.0 | 17(NLT) | 1 | 976.4534 | 973.4651387 | 4303470 | 49.37 | 1 | 6 | 5 | 1 | 0 | 1 | 0 | 1 | 3414.327188 | -0.0071 | -2.08 | -0.63 |
| Trypsin | 49 | 38 | 3511 | 1 | 4132 | 3415.3279 | 48.7441 | 31,701 | 3414.320075 | 32 | 14 | 21 | QCVNLTTR | 3.00020000.0 | 17(NLT) | 1 | 976.4534 | 973.4651387 | 4303470 | 49.37 | 2 | 7 | 5 | 0 | 1 | 0 | 0 | 1 | 3414.327189 | -0.0071 | -2.08 | -0.63 |
| Trypsin | 49 | 38 | 3975 | 1 | 4028 | 3546.4145 | 43.8618 | 26,447 | 3545.406675 | 32 | 14 | 21 | QCVNLTTR | 3.00020000.0 | 17(NLT) | 1 | 976.4534 | 973.4651387 | 4303470 | 49.37 | 1 | 6 | 5 | 4 | 0 | 0 | 0 | 0 | 3545.410584 | -0.0039 | -1.1 | -5.51 |
| Trypsin | 49 | 38 | 4359 | 1 | 4029 | 3587.4452 | 43.9034 | 22,651 | 3586.437375 | 32 | 14 | 21 | QCVNLTTR | 3.00020000.0 | 17(NLT) | 1 | 976.4534 | 973.4651387 | 4303470 | 49.37 | 1 | 5 | 6 | 4 | 0 | 0 | 0 | 0 | 3586.437133 | 0.0002 | 0.06 | -5.47 |
| Trypsin | 49 | 38 | 4455 | 1 | 4137 | 3091.2202 | 48.9745 | 21,777 | 3090.212375 | 32 | 14 | 21 | QCVNLTTR | 3.00020000.0 | 17(NLT) | 1 | 976.4534 | 973.4651387 | 4303470 | 49.37 | 1 | 4 | 5 | 1 | 0 | 1 | 0 | 1 | 3090.22154 | -0.0092 | -2.96 | -0.40 |
| Trypsin | 49 | 38 | 4455 | 1 | 4137 | 3091.2202 | 48.9745 | 21,777 | 3090.212375 | 32 | 14 | 21 | QCVNLTTR | 3.00020000.0 | 17(NLT) | 1 | 976.4534 | 973.4651387 | 4303470 | 49.37 | 2 | 5 | 5 | 0 | 1 | 0 | 0 | 1 | 3090.221541 | -0.0092 | -2.96 | -0.40 |
| Trypsin | 49 | 38 | 5008 | 1 | 4126 | 3561.3893 | 48.4645 | 17,656 | 3560.381475 | 32 | 14 | 21 | QCVNLTTR | 3.00020000.0 | 17(NLT) | 1 | 976.4534 | 973.4651387 | 4303470 | 49.37 | 1 | 6 | 5 | 2 | 0 | 1 | 0 | 1 | 3560.385097 | -0.0036 | -1.01 | -0.91 |
| Trypsin | 49 | 38 | 5008 | 1 | 4126 | 3561.3893 | 48.4645 | 17,656 | 3560.381475 | 32 | 14 | 21 | QCVNLTTR | 3.00020000.0 | 17(NLT) | 1 | 976.4534 | 973.4651387 | 4303470 | 49.37 | 2 | 7 | 5 | 1 | 1 | 0 | 0 | 1 | 3560.385098 | -0.0036 | -1.01 | -0.91 |
| Trypsin | 49 | 38 | 5426 | 1 | 4047 | 3133.2707 | 44.7547 | 14,645 | 3132.262875 | 32 | 14 | 21 | QCVNLTTR | 3.00020000.0 | 17(NLT) | 1 | 976.4534 | 973.4651387 | 4303470 | 49.37 | 1 | 4 | 6 | 2 | 0 | 0 | 0 | 1 | 3132.268491 | -0.0056 | -1.79 | -4.62 |
| Trypsin | 49 | 38 | 5777 | 1 | 4031 | 3660.4702 | 44.01 | 12,474 | 3659.462375 | 32 | 14 | 21 | QCVNLTTR | 3.00020000.0 | 17(NLT) | 1 | 976.4534 | 973.4651387 | 4303470 | 49.37 | 1 | 6 | 7 | 2 | 0 | 0 | 0 | 0 | 3659.453512 | 0.0089 | 2.42 | -5.36 |
| Chymotrypsin | 19 | 31 | 49 | 1 | 4627 | 2278.8999 | 72.9165 | 5,172,450 | 2277.892075 | 1001 | 59 | 65 | FSNVTWF | 0.0020000.0 | 61(NVT) | 1 |  | 899.41773 | 379236.06 | 82.81 | 1 | 6 | 2 | 0 | 0 | 0 | 0 | 1 | 2277.89342 | -0.0013 | -0.59 | -9.89 |
| Chymotrypsin | 19 | 31 | 95 | 1 | 4592 | 2815.1228 | 71.6958 | 3,112,780 | 2814.114975 | 1001 | 59 | 65 | FSNVTWF | 0.0020000.0 | 61(NVT) | 1 |  | 899.41773 | 379236.06 | 82.81 | 1 | 5 | 4 | 2 | 0 | 0 | 0 | 1 | 2814.11516 | -0.0002 | -0.06 | -11.12 |
| Chymotrypsin | 19 | 31 | 119 | 1 | 4625 | 2465.9844 | 72.8322 | 2,651,650 | 2464.976575 | 1001 | 59 | 65 | FSNVTWF | 0.0020000.0 | 61(NVT) | 1 |  | 899.41773 | 379236.06 | 82.81 | 1 | 5 | 3 | 1 | 0 | 0 | 0 | 1 | 2464.977878 | -0.0013 | -0.52 | -9.98 |
| Chymotrypsin | 19 | 31 | 168 | 1 | 4611 | 2628.036 | 72.3627 | 1,888,400 | 2627.028175 | 1001 | 59 | 65 | FSNVTWF | 0.0020000.0 | 61(NVT) | 1 |  | 899.41773 | 379236.06 | 82.81 | 1 | 6 | 3 | 1 | 0 | 0 | 0 | 1 | 2627.030702 | -0.0025 | -0.96 | -10.45 |
| Chymotrypsin | 19 | 31 | 226 | 1 | 4591 | 3018.2007 | 71.6519 | 1,443,960 | 3017.192875 | 1001 | 59 | 65 | FSNVTWF | 0.0020000.0 | 61(NVT) | 1 |  | 899.41773 | 379236.06 | 82.81 | 1 | 5 | 5 | 2 | 0 | 0 | 0 | 1 | 3017.194533 | -0.0017 | -0.54 | -11.16 |
| Chymotrypsin | 19 | 31 | 261 | 1 | 4565 | 3326.3118 | 70.5583 | 1,276,130 | 3325.303975 | 1001 | 59 | 65 | FSNVTWF | 0.0020000.0 | 61(NVT) | 1 |  | 899.41773 | 379236.06 | 82.81 | 1 | 6 | 5 | 3 | 0 | 0 | 0 | 1 | 3325.305266 | -0.0013 | -0.38 | -12.25 |
| Chymotrypsin | 19 | 31 | 269 | 1 | 4619 | 2440.9522 | 72.576 | 1,251,910 | 2439.944375 | 1001 | 59 | 65 | FSNVTWF | 0.0020000.0 | 61(NVT) | 1 |  | 899.41773 | 379236.06 | 82.81 | 1 | 7 | 2 | 0 | 0 | 0 | 0 | 1 | 2439.946244 | -0.0019 | -0.76 | -10.24 |
| Chymotrypsin | 19 | 31 | 283 | 1 | 4569 | 3529.393 | 70.722 | 1,215,500 | 3528.385175 | 1001 | 59 | 65 | FSNVTWF | 0.0020000.0 | 61(NVT) | 1 |  | 899.41773 | 379236.06 | 82.81 | 1 | 6 | 6 | 3 | 0 | 0 | 0 | 1 | 3528.384639 | 0.0005 | 0.15 | -12.09 |
| Chymotrypsin | 19 | 31 | 729 | 1 | 4555 | 3675.4409 | 70.1262 | 507,475 | 3674.433075 | 1001 | 59 | 65 | FSNVTWF | 0.0020000.0 | 61(NVT) | 1 |  | 899.41773 | 379236.06 | 82.81 | 1 | 6 | 6 | 4 | 0 | 0 | 0 | 0 | 3674.442548 | -0.0095 | -2.57 | -12.68 |
| Chymotrypsin | 19 | 31 | 787 | 1 | 4550 | 3472.3689 | 69.919 | 472,994 | 3471.361075 | 1001 | 59 | 65 | FSNVTWF | 0.0020000.0 | 61(NVT) | 1 |  | 899.41773 | 379236.06 | 82.81 | 1 | 5 | 6 | 4 | 0 | 0 | 0 | 0 | 3471.363175 | -0.0021 | -0.6 | -12.89 |
| Chymotrypsin | 19 | 31 | 832 | 1 | 4574 | 2961.1811 | 70.9407 | 448,384 | 2960.173275 | 1001 | 59 | 65 | FSNVTWF | 0.0020000.0 | 61(NVT) | 1 |  | 899.41773 | 379236.06 | 82.81 | 1 | 5 | 4 | 3 | 0 | 0 | 0 | 1 | 2960.173069 | 0.0002 | 0.06 | -11.87 |
| Chymotrypsin | 19 | 31 | 845 | 1 | 4606 | 2603.0047 | 72.221 | 444,475 | 2601.996875 | 1001 | 59 | 65 | FSNVTWF | 0.0020000.0 | 61(NVT) | 1 |  | 899.41773 | 379236.06 | 82.81 | 1 | 8 | 2 | 0 | 0 | 0 | 0 | 1 | 2601.999068 | -0.0022 | -0.84 | -10.59 |
| Chymotrypsin | 19 | 31 | 1040 | 1 | 4576 | 3164.2593 | 71.0184 | 364,002 | 3163.251475 | 1001 | 59 | 65 | FSNVTWF | 0.0020000.0 | 61(NVT) | 1 |  | 899.41773 | 379236.06 | 82.81 | 1 | 5 | 5 | 3 | 0 | 0 | 0 | 1 | 3163.252442 | -0.0010 | -0.3 | -11.79 |
| Chymotrypsin | 19 | 31 | 1204 | 1 | 4638 | 2116.8462 | 73.3583 | 318,060 | 2115.838375 | 1001 | 59 | 65 | FSNVTWF | 0.0020000.0 | 61(NVT) | 1 |  | 899.41773 | 379236.06 | 82.81 | 1 | 5 | 2 | 0 | 0 | 0 | 0 | 1 | 2115.840596 | -0.0022 | -1.04 | -9.45 |
| Chymotrypsin | 19 | 31 | 1382 | 1 | 4640 | 2303.9334 | 73.4449 | 278,424 | 2302.925575 | 1001 | 59 | 65 | FSNVTWF | 0.0020000.0 | 61(NVT) | 1 |  | 899.41773 | 379236.06 | 82.81 | 1 | 4 | 3 | 1 | 0 | 0 | 0 | 1 | 2302.925054 | 0.0005 | 0.22 | -9.37 |
| Chymotrypsin | 19 | 31 | 1393 | 1 | 4618 | 2523.0018 | 72.5462 | 277,482 | 2521.993975 | 1001 | 59 | 65 | FSNVTWF | 0.0020000.0 | 61(NVT) | 1 |  | 899.41773 | 379236.06 | 82.81 | 1 | 5 | 4 | 0 | 0 | 0 | 0 | 1 | 2521.999342 | -0.0054 | -2.12 | -10.26 |
| Chymotrypsin | 19 | 31 | 1699 | 1 | 4590 | 2888.1385 | 71.6073 | 229,577 | 2887.130675 | 1001 | 59 | 65 | FSNVTWF | 0.0020000.0 | 61(NVT) | 1 |  | 899.41773 | 379236.06 | 82.81 | 1 | 6 | 5 | 0 | 0 | 0 | 0 | 1 | 2887.131539 | -0.0009 | -0.29 | -11.20 |
| Chymotrypsin | 19 | 31 | 1963 | 1 | 4583 | 3383.3338 | 71.3077 | 198,484 | 3382.325975 | 1001 | 59 | 65 | FSNVTWF | 0.0020000.0 | 61(NVT) | 1 |  | 899.41773 | 379236.06 | 82.81 | 1 | 6 | 6 | 2 | 0 | 0 | 0 | 1 | 3382.32673 | -0.0008 | -0.22 | -11.50 |
| Chymotrypsin | 19 | 31 | 2097 | 1 | 4599 | 2612.0437 | 71.9825 | 186,451 | 2611.035875 | 1001 | 59 | 65 | FSNVTWF | 0.0020000.0 | 61(NVT) | 1 |  | 899.41773 | 379236.06 | 82.81 | 1 | 5 | 3 | 2 | 0 | 0 | 0 | 0 | 2611.035787 | 0.0001 | 0.03 | -10.83 |
| Chymotrypsin | 19 | 31 | 2109 | 1 | 4595 | 3139.2275 | 71.824 | 185,985 | 3138.219675 | 1001 | 59 | 65 | FSNVTWF | 0.0020000.0 | 61(NVT) | 1 |  | 899.41773 | 379236.06 | 82.81 | 1 | 7 | 4 | 2 | 0 | 0 | 0 | 1 | 3138.220808 | -0.0011 | -0.36 | -10.99 |
| Chymotrypsin | 19 | 31 | 2532 | 1 | 4576 | 3180.2433 | 71.0184 | 152,983 | 3179.235475 | 1001 | 59 | 65 | FSNVTWF | 0.0020000.0 | 61(NVT) | 1 |  | 899.41773 | 379236.06 | 82.81 | 1 | 6 | 5 | 2 | 0 | 0 | 0 | 1 | 3179.247357 | -0.0119 | -3.73 | -11.79 |
| Chymotrypsin | 19 | 31 | 2647 | 1 | 4598 | 3091.2217 | 71.9396 | 146,171 | 3090.213875 | 1001 | 59 | 65 | FSNVTWF | 0.0020000.0 | 61(NVT) | 1 |  | 899.41773 | 379236.06 | 82.81 | 1 | 6 | 6 | 0 | 0 | 0 | 0 | 1 | 3090.210912 | 0.0030 | 0.95 | -10.87 |
| Chymotrypsin | 19 | 31 | 2776 | 1 | 4591 | 2977.1703 | 71.6519 | 138,450 | 2976.162475 | 1001 | 59 | 65 | FSNVTWF | 0.0020000.0 | 61(NVT) | 1 |  | 899.41773 | 379236.06 | 82.81 | 1 | 6 | 4 | 2 | 0 | 0 | 0 | 1 | 2976.167984 | -0.0055 | -1.85 | -11.16 |
| Chymotrypsin | 19 | 31 | 2847 | 1 | 4620 | 2831.1167 | 72.6193 | 134,318 | 2830.108875 | 1001 | 59 | 65 | FSNVTWF | 0.0020000.0 | 61(NVT) | 1 |  | 899.41773 | 379236.06 | 82.81 | 1 | 6 | 4 | 1 | 0 | 0 | 0 | 1 | 2830.110075 | -0.0012 | -0.42 | -10.19 |
| Chymotrypsin | 19 | 31 | 2861 | 1 | 4618 | 2977.172 | 72.5462 | 133,684 | 2976.164175 | 1001 | 59 | 65 | FSNVTWF | 0.0020000.0 | 61(NVT) | 1 |  | 899.41773 | 379236.06 | 82.81 | 1 | 6 | 4 | 2 | 0 | 0 | 0 | 1 | 2976.167984 | -0.0038 | -1.27 | -10.26 |
| Chymotrypsin | 19 | 31 | 3109 | 1 | 4588 | 3342.3006 | 71.5222 | 121,700 | 3341.292775 | 1001 | 59 | 65 | FSNVTWF | 0.0020000.0 | 61(NVT) | 1 |  | 899.41773 | 379236.06 | 82.81 | 1 | 7 | 5 | 2 | 0 | 0 | 0 | 0 | 3341.300181 | -0.0074 | -2.21 | -11.29 |
| Chymotrypsin | 19 | 31 | 3379 | 1 | 4635 | 2319.9319 | 73.2328 | 111,223 | 2318.924075 | 1001 | 59 | 65 | FSNVTWF | 0.0020000.0 | 61(NVT) | 1 |  | 899.41773 | 379236.06 | 82.81 | 1 | 5 | 3 | 0 | 0 | 0 | 0 | 1 | 2318.919969 | 0.0041 | 1.77 | -9.58 |
| Chymotrypsin | 19 | 31 | 3387 | 1 | 4588 | 3237.2754 | 71.5222 | 110,840 | 3236.267575 | 1001 | 59 | 65 | FSNVTWF | 0.0020000.0 | 61(NVT) | 1 |  | 899.41773 | 379236.06 | 82.81 | 1 | 6 | 6 | 1 | 0 | 0 | 0 | 1 | 3236.268821 | -0.0012 | -0.38 | -11.29 |
| Chymotrypsin | 19 | 31 | 3400 | 1 | 4599 | 2993.1714 | 71.9825 | 110,479 | 2992.163575 | 1001 | 59 | 65 | FSNVTWF | 0.0020000.0 | 61(NVT) | 1 |  | 899.41773 | 379236.06 | 82.81 | 1 | 7 | 4 | 1 | 0 | 0 | 0 | 1 | 2992.162899 | 0.0007 | 0.22 | -10.83 |
| Chymotrypsin | 19 | 31 | 5362 | 1 | 4548 | 3837.5017 | 69.8341 | 62,394 | 3836.493875 | 1001 | 59 | 65 | FSNVTWF | 0.0020000.0 | 61(NVT) | 1 |  | 899.41773 | 379236.06 | 82.81 | 1 | 7 | 6 | 4 | 0 | 0 | 0 | 1 | 3836.495372 | -0.0015 | -0.39 | -12.98 |
| Chymotrypsin | 19 | 31 | 5362 | 1 | 4548 | 3837.5017 | 69.8341 | 62,394 | 3836.493875 | 1001 | 59 | 65 | FSNVTWF | 0.0020000.0 | 61(NVT) | 1 |  | 899.41773 | 379236.06 | 82.81 | 2 | 2 | 10 | 0 | 2 | 0 | 1 | -1 | 3836.507942 | -0.0141 | -3.66 | -12.98 |
| Chymotrypsin | 19 | 31 | 5826 | 1 | 4588 | 3180.2505 | 71.5222 | 54,643 | 3179.242675 | 1001 | 59 | 65 | FSNVTWF | 0.0020000.0 | 61(NVT) | 1 |  | 899.41773 | 379236.06 | 82.81 | 1 | 6 | 5 | 2 | 0 | 0 | 0 | 1 | 3179.247357 | -0.0047 | -1.47 | -11.29 |
| alpha-Lytic | 41 | 31 | 186 | 1 | 5326 | 3155.3336 | 95.7283 | 714,913 | 3154.325775 | 12 | 52 | 63 | QDLFLPFFSNVT | 0.000000000200.0 | 61(NVT) | 1 | 1429.7016 | 1426.713339 | 11168946 | 106.08 | 1 | 6 | 3 | 1 | 0 | 0 | 0 | 1 | 3154.326311 | -0.0005 | -0.16 | -10.35 |
| alpha-Lytic | 41 | 31 | 275 | 1 | 5315 | 3342.4171 | 95.2382 | 473,699 | 3341.409275 | 12 | 52 | 63 | QDLFLPFFSNVT | 0.000000000200.0 | 61(NVT) | 1 | 1429.7016 | 1426.713339 | 11168946 | 106.08 | 1 | 5 | 4 | 2 | 0 | 0 | 0 | 1 | 3341.410769 | -0.0015 | -0.44 | -10.84 |
| alpha-Lytic | 41 | 31 | 497 | 1 | 5292 | 3853.6082 | 94.1428 | 261,923 | 3852.600375 | 12 | 52 | 63 | QDLFLPFFSNVT | 0.000000000200.0 | 61(NVT) | 1 | 1429.7016 | 1426.713339 | 11168946 | 106.08 | 1 | 6 | 5 | 3 | 0 | 0 | 0 | 1 | 3852.600875 | -0.0005 | -0.12 | -11.94 |
| alpha-Lytic | 41 | 31 | 543 | 1 | 5293 | 4056.683 | 94.1879 | 244,179 | 4055.675175 | 12 | 52 | 63 | QDLFLPFFSNVT | 0.000000000200.0 | 61(NVT) | 1 | 1429.7016 | 1426.713339 | 11168946 | 106.08 | 1 | 6 | 6 | 3 | 0 | 0 | 0 | 1 | 4055.680248 | -0.0051 | -1.25 | -11.89 |
| alpha-Lytic | 41 | 31 | 573 | 1 | 5335 | 2993.2831 | 96.1392 | 231,671 | 2992.275275 | 12 | 52 | 63 | QDLFLPFFSNVT | 0.000000000200.0 | 61(NVT) | 1 | 1429.7016 | 1426.713339 | 11168946 | 106.08 | 1 | 5 | 3 | 1 | 0 | 0 | 0 | 1 | 2992.273487 | 0.0018 | 0.59 | -9.94 |
| alpha-Lytic | 41 | 31 | 600 | 1 | 5312 | 3545.4993 | 95.0789 | 222,574 | 3544.491475 | 12 | 52 | 63 | QDLFLPFFSNVT | 0.000000000200.0 | 61(NVT) | 1 | 1429.7016 | 1426.713339 | 11168946 | 106.08 | 1 | 5 | 5 | 2 | 0 | 0 | 0 | 1 | 3544.490142 | 0.0013 | 0.37 | -11.00 |
| alpha-Lytic | 41 | 31 | 1071 | 1 | 5276 | 4202.7473 | 93.493 | 124,112 | 4201.739475 | 12 | 52 | 63 | QDLFLPFFSNVT | 0.000000000200.0 | 61(NVT) | 1 | 1429.7016 | 1426.713339 | 11168946 | 106.08 | 1 | 6 | 6 | 4 | 0 | 0 | 0 | 0 | 4201.738157 | 0.0013 | 0.31 | -12.59 |
| alpha-Lytic | 41 | 31 | 1124 | 1 | 5298 | 3488.4761 | 94.4401 | 117,324 | 3487.468275 | 12 | 52 | 63 | QDLFLPFFSNVT | 0.000000000200.0 | 61(NVT) | 1 | 1429.7016 | 1426.713339 | 11168946 | 106.08 | 1 | 5 | 4 | 3 | 0 | 0 | 0 | 1 | 3487.468678 | -0.0004 | -0.11 | -11.64 |
| alpha-Lytic | 41 | 31 | 1217 | 1 | 5276 | 3999.6639 | 93.493 | 108,330 | 3998.656075 | 12 | 52 | 63 | QDLFLPFFSNVT | 0.000000000200.0 | 61(NVT) | 1 | 1429.7016 | 1426.713339 | 11168946 | 106.08 | 1 | 5 | 5 | 4 | 0 | 0 | 0 | 0 | 3998.658784 | -0.0027 | -0.67 | -12.59 |
| alpha-Lytic | 41 | 31 | 1814 | 1 | 5297 | 3691.5571 | 94.3968 | 67,256 | 3690.549275 | 12 | 52 | 63 | QDLFLPFFSNVT | 0.000000000200.0 | 61(NVT) | 1 | 1429.7016 | 1426.713339 | 11168946 | 106.08 | 1 | 5 | 5 | 3 | 0 | 0 | 0 | 1 | 3690.548051 | 0.0012 | 0.33 | -11.68 |
| alpha-Lytic | 41 | 31 | 1957 | 1 | 5326 | 3504.4731 | 95.7283 | 60,771 | 3503.465275 | 12 | 52 | 63 | QDLFLPFFSNVT | 0.000000000200.0 | 61(NVT) | 1 | 1429.7016 | 1426.713339 | 11168946 | 106.08 | 1 | 6 | 4 | 2 | 0 | 0 | 0 | 1 | 3503.463593 | 0.0017 | 0.48 | -10.35 |
| alpha-Lytic | 41 | 31 | 2105 | 1 | 5301 | 3910.63 | 94.5875 | 55,691 | 3909.622175 | 12 | 52 | 63 | QDLFLPFFSNVT | 0.000000000200.0 | 61(NVT) | 1 | 1429.7016 | 1426.713339 | 11168946 | 106.08 | 1 | 6 | 6 | 2 | 0 | 0 | 0 | 1 | 3909.622339 | -0.0002 | -0.04 | -11.49 |
| alpha-Lytic | 41 | 31 | 2189 | 1 | 5322 | 3196.3615 | 95.5522 | 52,721 | 3195.353675 | 12 | 52 | 63 | QDLFLPFFSNVT | 0.000000000200.0 | 61(NVT) | 1 | 1429.7016 | 1426.713339 | 11168946 | 106.08 | 1 | 5 | 4 | 1 | 0 | 0 | 0 | 1 | 3195.35286 | 0.0008 | 0.25 | -10.53 |
| alpha-Lytic | 41 | 31 | 2223 | 1 | 5321 | 3399.4471 | 95.5062 | 51,640 | 3398.439275 | 12 | 52 | 63 | QDLFLPFFSNVT | 0.000000000200.0 | 61(NVT) | 1 | 1429.7016 | 1426.713339 | 11168946 | 106.08 | 1 | 5 | 5 | 1 | 0 | 0 | 0 | 1 | 3398.432233 | 0.0070 | 2.07 | -10.57 |
| alpha-Lytic | 41 | 31 | 2693 | 1 | 5318 | 3666.5226 | 95.3701 | 37,685 | 3665.514775 | 12 | 52 | 63 | QDLFLPFFSNVT | 0.000000000200.0 | 61(NVT) | 1 | 1429.7016 | 1426.713339 | 11168946 | 106.08 | 1 | 7 | 4 | 2 | 0 | 0 | 0 | 1 | 3665.516417 | -0.0016 | -0.44 | -10.71 |
| alpha-Lytic | 41 | 31 | 2734 | 1 | 5318 | 3139.3385 | 95.3701 | 36,558 | 3138.330675 | 12 | 52 | 63 | QDLFLPFFSNVT | 0.000000000200.0 | 61(NVT) | 1 | 1429.7016 | 1426.713339 | 11168946 | 106.08 | 1 | 5 | 3 | 2 | 0 | 0 | 0 | 0 | 3138.331396 | -0.0007 | -0.22 | -10.71 |
| alpha-Lytic | 41 | 31 | 2791 | 1 | 5308 | 3764.5726 | 94.905 | 34,860 | 3763.564775 | 12 | 52 | 63 | QDLFLPFFSNVT | 0.000000000200.0 | 61(NVT) | 1 | 1429.7016 | 1426.713339 | 11168946 | 106.08 | 1 | 6 | 6 | 1 | 0 | 0 | 0 | 1 | 3763.56443 | 0.0003 | 0.09 | -11.18 |
| alpha-Lytic | 41 | 31 | 2900 | 1 | 5315 | 3618.5108 | 95.2382 | 32,110 | 3617.502975 | 12 | 52 | 63 | QDLFLPFFSNVT | 0.000000000200.0 | 61(NVT) | 1 | 1429.7016 | 1426.713339 | 11168946 | 106.08 | 1 | 6 | 6 | 0 | 0 | 0 | 0 | 1 | 3617.506521 | -0.0035 | -0.98 | -10.84 |
| alpha-Lytic | 41 | 31 | 2922 | 1 | 5322 | 3520.4652 | 95.5522 | 31,722 | 3519.457375 | 12 | 52 | 63 | QDLFLPFFSNVT | 0.000000000200.0 | 61(NVT) | 1 | 1429.7016 | 1426.713339 | 11168946 | 106.08 | 1 | 7 | 4 | 1 | 0 | 0 | 0 | 1 | 3519.458508 | -0.0011 | -0.32 | -10.53 |
| alpha-Lytic | 41 | 31 | 3166 | 1 | 5312 | 3504.4742 | 95.0789 | 26,470 | 3503.466375 | 12 | 52 | 63 | QDLFLPFFSNVT | 0.000000000200.0 | 61(NVT) | 1 | 1429.7016 | 1426.713339 | 11168946 | 106.08 | 1 | 6 | 4 | 2 | 0 | 0 | 0 | 1 | 3503.463593 | 0.0028 | 0.79 | -11.00 |
| alpha-Lytic | 41 | 31 | 3266 | 1 | 5308 | 3869.6023 | 94.905 | 24,134 | 3868.594475 | 12 | 52 | 63 | QDLFLPFFSNVT | 0.000000000200.0 | 61(NVT) | 1 | 1429.7016 | 1426.713339 | 11168946 | 106.08 | 1 | 7 | 5 | 2 | 0 | 0 | 0 | 0 | 3868.59579 | -0.0013 | -0.33 | -11.18 |
| alpha-Lytic | 41 | 31 | 3521 | 1 | 5275 | 4364.7905 | 93.4373 | 18,629 | 4363.782675 | 12 | 52 | 63 | QDLFLPFFSNVT | 0.000000000200.0 | 61(NVT) | 1 | 1429.7016 | 1426.713339 | 11168946 | 106.08 | 1 | 7 | 6 | 4 | 0 | 0 | 0 | 1 | 4363.790981 | -0.0083 | -1.9 | -12.64 |
| alpha-Lytic | 41 | 31 | 3521 | 1 | 5275 | 4364.7905 | 93.4373 | 18,629 | 4363.782675 | 12 | 52 | 63 | QDLFLPFFSNVT | 0.000000000200.0 | 61(NVT) | 1 | 1429.7016 | 1426.713339 | 11168946 | 106.08 | 2 | 2 | 10 | 0 | 2 | 0 | 1 | -1 | 4363.803551 | -0.0209 | -4.78 | -12.64 |
| alpha-Lytic | 41 | 31 | 3615 | 1 | 5258 | 4510.8479 | 92.8648 | 16,574 | 4509.840075 | 12 | 52 | 63 | QDLFLPFFSNVT | 0.000000000200.0 | 61(NVT) | 1 | 1429.7016 | 1426.713339 | 11168946 | 106.08 | 1 | 9 | 8 | 0 | 0 | 0 | 0 | 1 | 4509.823739 | 0.0163 | 3.62 | -13.22 |
| alpha-Lytic | 41 | 31 | 3615 | 1 | 5258 | 4510.8479 | 92.8648 | 16,574 | 4509.840075 | 12 | 52 | 63 | QDLFLPFFSNVT | 0.000000000200.0 | 61(NVT) | 1 | 1429.7016 | 1426.713339 | 11168946 | 106.08 | 2 | 2 | 10 | 1 | 2 | 0 | 1 | -1 | 4509.86146 | -0.0214 | -4.74 | -13.22 |
| alpha-Lytic | 41 | 31 | 3637 | 1 | 5569 | 3316.3648 | 100.108 | 16,071 | 3315.356975 | 12 | 52 | 63 | QDLFLPFFSNVT | 0.000000000200.0 | 61(NVT) | 1 | 1429.7016 | 1426.713339 | 11168946 | 106.08 | 1 | 6 | 3 | 0 | 0 | 1 | 0 | 1 | 3315.358733 | -0.0018 | -0.53 | -5.97 |
| alpha-Lytic | 41 | 31 | 3737 | 1 | 5501 | 3503.4503 | 99.4868 | 13,803 | 3502.442475 | 12 | 52 | 63 | QDLFLPFFSNVT | 0.000000000200.0 | 61(NVT) | 1 | 1429.7016 | 1426.713339 | 11168946 | 106.08 | 1 | 5 | 4 | 1 | 0 | 1 | 0 | 1 | 3502.443191 | -0.0007 | -0.2 | -6.59 |
| alpha-Lytic | 41 | 31 | 3737 | 1 | 5501 | 3503.4503 | 99.4868 | 13,803 | 3502.442475 | 12 | 52 | 63 | QDLFLPFFSNVT | 0.000000000200.0 | 61(NVT) | 1 | 1429.7016 | 1426.713339 | 11168946 | 106.08 | 2 | 6 | 4 | 0 | 1 | 0 | 0 | 1 | 3502.443192 | -0.0007 | -0.2 | -6.59 |
| alpha-Lytic | 41 | 31 | 3815 | 1 | 5304 | 4015.6552 | 94.728 | 12,156 | 4014.647375 | 12 | 52 | 63 | QDLFLPFFSNVT | 0.000000000200.0 | 61(NVT) | 1 | 1429.7016 | 1426.713339 | 11168946 | 106.08 | 1 | 7 | 5 | 3 | 0 | 0 | 0 | 0 | 4014.653699 | -0.0063 | -1.57 | -11.35 |
| alpha-Lytic | 41 | 31 | 3901 | 1 | 5617 | 3154.3165 | 100.524 | 10,348 | 3153.308675 | 12 | 52 | 63 | QDLFLPFFSNVT | 0.000000000200.0 | 61(NVT) | 1 | 1429.7016 | 1426.713339 | 11168946 | 106.08 | 1 | 5 | 3 | 0 | 0 | 1 | 0 | 1 | 3153.305909 | 0.0028 | 0.87 | -5.56 |
| alpha-Lytic | 41 | 31 | 3975 | 1 | 5305 | 3342.424 | 94.7712 | 8,768 | 3341.416175 | 12 | 52 | 63 | QDLFLPFFSNVT | 0.000000000200.0 | 61(NVT) | 1 | 1429.7016 | 1426.713339 | 11168946 | 106.08 | 1 | 5 | 4 | 2 | 0 | 0 | 0 | 1 | 3341.410769 | 0.0054 | 1.61 | -11.31 |
| alpha-Lytic | 41 | 31 | 4011 | 1 | 5452 | 4014.6434 | 98.7382 | 7,859 | 4013.635575 | 12 | 52 | 63 | QDLFLPFFSNVT | 0.000000000200.0 | 61(NVT) | 1 | 1429.7016 | 1426.713339 | 11168946 | 106.08 | 1 | 7 | 5 | 1 | 1 | 0 | 0 | 1 | 4013.633298 | 0.0023 | 0.56 | -7.34 |
| alpha-Lytic | 41 | 31 | 4011 | 1 | 5452 | 4014.6434 | 98.7382 | 7,859 | 4013.635575 | 12 | 52 | 63 | QDLFLPFFSNVT | 0.000000000200.0 | 61(NVT) | 1 | 1429.7016 | 1426.713339 | 11168946 | 106.08 | 2 | 6 | 5 | 2 | 0 | 1 | 0 | 1 | 4013.633297 | 0.0023 | 0.56 | -7.34 |
| alpha-Lytic | 41 | 31 | 4071 | 1 | 5543 | 3357.3926 | 99.8621 | 6,248 | 3356.384775 | 12 | 52 | 63 | QDLFLPFFSNVT | 0.000000000200.0 | 61(NVT) | 1 | 1429.7016 | 1426.713339 | 11168946 | 106.08 | 1 | 5 | 4 | 0 | 0 | 1 | 0 | 1 | 3356.385282 | -0.0005 | -0.15 | -6.22 |
| alpha-Lytic | 41 | 31 | 4102 | 1 | 5459 | 4217.7224 | 98.8157 | 5,463 | 4216.714575 | 12 | 52 | 63 | QDLFLPFFSNVT | 0.000000000200.0 | 61(NVT) | 1 | 1429.7016 | 1426.713339 | 11168946 | 106.08 | 1 | 7 | 6 | 1 | 1 | 0 | 0 | 1 | 4216.712671 | 0.0019 | 0.45 | -7.27 |
| alpha-Lytic | 41 | 31 | 4102 | 1 | 5459 | 4217.7224 | 98.8157 | 5,463 | 4216.714575 | 12 | 52 | 63 | QDLFLPFFSNVT | 0.000000000200.0 | 61(NVT) | 1 | 1429.7016 | 1426.713339 | 11168946 | 106.08 | 2 | 6 | 6 | 2 | 0 | 1 | 0 | 0 | 4216.71267 | 0.0019 | 0.45 | -7.27 |
| Chymotrypsin | 2 | 174 | 5 | 1 | 3791 | 3057.3491 | 37.9474 | 28,050,700 | 3056.341275 | 21 | 66 | 79 | HAIHVSGTNGTKRF | 0.00000000200000.0 | 74(NGT) | 1 | 1526.784 | 1523.795739 | 23511576 | 36.61 | 1 | 2 | 3 | 2 | 0 | 1 | 1 | -1 | 3056.345655 | -0.0044 | -1.43 | 1.34 |
| Chymotrypsin | 2 | 174 | 5 | 1 | 3791 | 3057.3491 | 37.9474 | 28,050,700 | 3056.341275 | 21 | 66 | 79 | HAIHVSGTNGTKRF | 0.00000000200000.0 | 74(NGT) | 1 | 1526.784 | 1523.795739 | 23511576 | 36.61 | 2 | 3 | 3 | 1 | 1 | 0 | 0 | 0 | 3056.345656 | -0.0044 | -1.43 | 1.34 |
| Chymotrypsin | 2 | 174 | 28 | 1 | 3788 | 3422.4816 | 37.8213 | 7,284,320 | 3421.473775 | 21 | 66 | 79 | HAIHVSGTNGTKRF | 0.00000000200000.0 | 74(NGT) | 1 | 1526.784 | 1523.795739 | 23511576 | 36.61 | 1 | 4 | 4 | 1 | 1 | 0 | 0 | 1 | 3421.477853 | -0.0041 | -1.19 | 1.21 |
| Chymotrypsin | 2 | 174 | 28 | 1 | 3788 | 3422.4816 | 37.8213 | 7,284,320 | 3421.473775 | 21 | 66 | 79 | HAIHVSGTNGTKRF | 0.00000000200000.0 | 74(NGT) | 1 | 1526.784 | 1523.795739 | 23511576 | 36.61 | 2 | 3 | 4 | 2 | 0 | 1 | 0 | 0 | 3421.477852 | -0.0041 | -1.19 | 1.21 |
| Chymotrypsin | 2 | 174 | 78 | 1 | 3658 | 4607.9393 | 32.2645 | 3,660,490 | 4606.931475 | 21 | 66 | 79 | HAIHVSGTNGTKRF | 0.00000000200000.0 | 74(NGT) | 1 | 1526.784 | 1523.795739 | 23511576 | 36.61 | 1 | 2 | 10 | 1 | 2 | 0 | 1 | -1 | 4606.94386 | -0.0124 | -2.68 | -4.34 |
| Chymotrypsin | 2 | 174 | 88 | 1 | 3787 | 3219.4006 | 37.7792 | 3,206,430 | 3218.392775 | 21 | 66 | 79 | HAIHVSGTNGTKRF | 0.00000000200000.0 | 74(NGT) | 1 | 1526.784 | 1523.795739 | 23511576 | 36.61 | 1 | 4 | 3 | 1 | 1 | 0 | 0 | 1 | 3218.39848 | -0.0057 | -1.77 | 1.17 |
| Chymotrypsin | 2 | 174 | 88 | 1 | 3787 | 3219.4006 | 37.7792 | 3,206,430 | 3218.392775 | 21 | 66 | 79 | HAIHVSGTNGTKRF | 0.00000000200000.0 | 74(NGT) | 1 | 1526.784 | 1523.795739 | 23511576 | 36.61 | 2 | 3 | 3 | 2 | 0 | 1 | 0 | 0 | 3218.398479 | -0.0057 | -1.77 | 1.17 |
| Chymotrypsin | 2 | 174 | 89 | 1 | 3838 | 3713.5765 | 39.9683 | 3,190,160 | 3712.568675 | 21 | 66 | 79 | HAIHVSGTNGTKRF | 0.00000000200000.0 | 74(NGT) | 1 | 1526.784 | 1523.795739 | 23511576 | 36.61 | 1 | 2 | 4 | 3 | 0 | 2 | 1 | -1 | 3712.573268 | -0.0046 | -1.23 | 3.36 |
| Chymotrypsin | 2 | 174 | 89 | 1 | 3838 | 3713.5765 | 39.9683 | 3,190,160 | 3712.568675 | 21 | 66 | 79 | HAIHVSGTNGTKRF | 0.00000000200000.0 | 74(NGT) | 1 | 1526.784 | 1523.795739 | 23511576 | 36.61 | 2 | 3 | 4 | 2 | 1 | 1 | 1 | -1 | 3712.573269 | -0.0046 | -1.23 | 3.36 |
| Chymotrypsin | 2 | 174 | 89 | 1 | 3838 | 3713.5765 | 39.9683 | 3,190,160 | 3712.568675 | 21 | 66 | 79 | HAIHVSGTNGTKRF | 0.00000000200000.0 | 74(NGT) | 1 | 1526.784 | 1523.795739 | 23511576 | 36.61 | 3 | 4 | 4 | 1 | 2 | 0 | 0 | 0 | 3712.57327 | -0.0046 | -1.23 | 3.36 |
| Chymotrypsin | 2 | 174 | 90 | 1 | 3672 | 3788.6374 | 32.8662 | 3,190,090 | 3787.629575 | 21 | 66 | 79 | HAIHVSGTNGTKRF | 0.00000000200000.0 | 74(NGT) | 1 | 1526.784 | 1523.795739 | 23511576 | 36.61 | 1 | 5 | 5 | 3 | 0 | 0 | 0 | 1 | 3787.630451 | -0.0009 | -0.23 | -3.74 |
| Chymotrypsin | 2 | 174 | 93 | 1 | 3670 | 3626.5844 | 32.7835 | 3,162,210 | 3625.576575 | 21 | 66 | 79 | HAIHVSGTNGTKRF | 0.00000000200000.0 | 74(NGT) | 1 | 1526.784 | 1523.795739 | 23511576 | 36.61 | 1 | 4 | 5 | 3 | 0 | 0 | 0 | 0 | 3625.577627 | -0.0011 | -0.29 | -3.83 |
| Chymotrypsin | 2 | 174 | 134 | 1 | 3654 | 4811.0187 | 32.094 | 2,347,830 | 4810.010875 | 21 | 66 | 79 | HAIHVSGTNGTKRF | 0.00000000200000.0 | 74(NGT) | 1 | 1526.784 | 1523.795739 | 23511576 | 36.61 |  |  |  |  |  |  |  |  |  |  |  |  |
| Chymotrypsin | 2 | 174 | 152 | 1 | 3669 | 3585.5593 | 32.7395 | 2,088,870 | 3584.551475 | 21 | 66 | 79 | HAIHVSGTNGTKRF | 0.00000000200000.0 | 74(NGT) | 1 | 1526.784 | 1523.795739 | 23511576 | 36.61 | 1 | 5 | 4 | 3 | 0 | 0 | 0 | 1 | 3584.551078 | 0.0004 | 0.11 | -3.87 |
| Chymotrypsin | 2 | 174 | 167 | 1 | 3670 | 4299.8264 | 32.7835 | 1,889,190 | 4298.818575 | 21 | 66 | 79 | HAIHVSGTNGTKRF | 0.00000000200000.0 | 74(NGT) | 1 | 1526.784 | 1523.795739 | 23511576 | 36.61 | 1 | 6 | 6 | 4 | 0 | 0 | 0 | 0 | 4298.820557 | -0.0020 | -0.46 | -3.83 |
| Chymotrypsin | 2 | 174 | 197 | 1 | 3843 | 3551.5229 | 40.1848 | 1,619,070 | 3550.515075 | 21 | 66 | 79 | HAIHVSGTNGTKRF | 0.00000000200000.0 | 74(NGT) | 1 | 1526.784 | 1523.795739 | 23511576 | 36.61 | 1 | 2 | 4 | 2 | 1 | 1 | 1 | -1 | 3550.520445 | -0.0054 | -1.51 | 3.58 |
| Chymotrypsin | 2 | 174 | 197 | 1 | 3843 | 3551.5229 | 40.1848 | 1,619,070 | 3550.515075 | 21 | 66 | 79 | HAIHVSGTNGTKRF | 0.00000000200000.0 | 74(NGT) | 1 | 1526.784 | 1523.795739 | 23511576 | 36.61 | 2 | 3 | 4 | 1 | 2 | 0 | 0 | 0 | 3550.520446 | -0.0054 | -1.51 | 3.58 |
| Chymotrypsin | 2 | 174 | 214 | 1 | 3841 | 3754.6025 | 40.0994 | 1,511,190 | 3753.594675 | 21 | 66 | 79 | HAIHVSGTNGTKRF | 0.00000000200000.0 | 74(NGT) | 1 | 1526.784 | 1523.795739 | 23511576 | 36.61 | 1 | 2 | 5 | 2 | 1 | 1 | 1 | -1 | 3753.599818 | -0.0051 | -1.37 | 3.49 |
| Chymotrypsin | 2 | 174 | 214 | 1 | 3841 | 3754.6025 | 40.0994 | 1,511,190 | 3753.594675 | 21 | 66 | 79 | HAIHVSGTNGTKRF | 0.00000000200000.0 | 74(NGT) | 1 | 1526.784 | 1523.795739 | 23511576 | 36.61 | 2 | 3 | 5 | 1 | 2 | 0 | 0 | 0 | 3753.599819 | -0.0051 | -1.37 | 3.49 |
| Chymotrypsin | 2 | 174 | 284 | 1 | 3664 | 4096.7487 | 32.5259 | 1,213,470 | 4095.740875 | 21 | 66 | 79 | HAIHVSGTNGTKRF | 0.00000000200000.0 | 74(NGT) | 1 | 1526.784 | 1523.795739 | 23511576 | 36.61 | 1 | 6 | 5 | 4 | 0 | 0 | 0 | 0 | 4095.741184 | -0.0003 | -0.07 | -4.08 |
| Chymotrypsin | 2 | 174 | 312 | 1 | 3785 | 3381.4541 | 37.6934 | 1,100,360 | 3380.446275 | 21 | 66 | 79 | HAIHVSGTNGTKRF | 0.00000000200000.0 | 74(NGT) | 1 | 1526.784 | 1523.795739 | 23511576 | 36.61 | 1 | 4 | 3 | 2 | 0 | 1 | 0 | 0 | 3380.451303 | -0.0050 | -1.48 | 1.08 |
| Chymotrypsin | 2 | 174 | 312 | 1 | 3785 | 3381.4541 | 37.6934 | 1,100,360 | 3380.446275 | 21 | 66 | 79 | HAIHVSGTNGTKRF | 0.00000000200000.0 | 74(NGT) | 1 | 1526.784 | 1523.795739 | 23511576 | 36.61 | 2 | 5 | 3 | 1 | 1 | 0 | 0 | 0 | 3380.451304 | -0.0050 | -1.48 | 1.08 |
| Chymotrypsin | 2 | 174 | 319 | 1 | 3793 | 3260.4265 | 38.0314 | 1,086,660 | 3259.418675 | 21 | 66 | 79 | HAIHVSGTNGTKRF | 0.00000000200000.0 | 74(NGT) | 1 | 1526.784 | 1523.795739 | 23511576 | 36.61 | 1 | 2 | 4 | 2 | 0 | 1 | 1 | -1 | 3259.425028 | -0.0064 | -1.94 | 1.42 |
| Chymotrypsin | 2 | 174 | 319 | 1 | 3793 | 3260.4265 | 38.0314 | 1,086,660 | 3259.418675 | 21 | 66 | 79 | HAIHVSGTNGTKRF | 0.00000000200000.0 | 74(NGT) | 1 | 1526.784 | 1523.795739 | 23511576 | 36.61 | 2 | 3 | 4 | 1 | 1 | 0 | 0 | 0 | 3259.425029 | -0.0064 | -1.94 | 1.42 |
| Chymotrypsin | 2 | 174 | 322 | 1 | 3666 | 4137.7753 | 32.6112 | 1,079,170 | 4136.767475 | 21 | 66 | 79 | HAIHVSGTNGTKRF | 0.00000000200000.0 | 74(NGT) | 1 | 1526.784 | 1523.795739 | 23511576 | 36.61 | 1 | 5 | 6 | 4 | 0 | 0 | 0 | 0 | 4136.767733 | -0.0003 | -0.06 | -4.00 |
| Chymotrypsin | 2 | 174 | 323 | 1 | 3660 | 4648.9663 | 32.3486 | 1,078,900 | 4647.958475 | 21 | 66 | 79 | HAIHVSGTNGTKRF | 0.00000000200000.0 | 74(NGT) | 1 | 1526.784 | 1523.795739 | 23511576 | 36.61 |  |  |  |  |  |  |  |  |  |  |  |  |
| Chymotrypsin | 2 | 174 | 442 | 1 | 3680 | 3277.4478 | 33.2082 | 802,343 | 3276.439975 | 21 | 66 | 79 | HAIHVSGTNGTKRF | 0.00000000200000.0 | 74(NGT) | 1 | 1526.784 | 1523.795739 | 23511576 | 36.61 | 1 | 4 | 4 | 2 | 0 | 0 | 0 | 1 | 3276.440345 | -0.0004 | -0.11 | -3.40 |
| Chymotrypsin | 2 | 174 | 464 | 1 | 3672 | 3667.6109 | 32.8662 | 770,130 | 3666.603075 | 21 | 66 | 79 | HAIHVSGTNGTKRF | 0.00000000200000.0 | 74(NGT) | 1 | 1526.784 | 1523.795739 | 23511576 | 36.61 | 1 | 3 | 6 | 3 | 0 | 0 | 0 | 0 | 3666.604176 | -0.0011 | -0.3 | -3.74 |
| Chymotrypsin | 2 | 174 | 510 | 1 | 3722 | 3787.6185 | 34.9983 | 701,158 | 3786.610675 | 21 | 66 | 79 | HAIHVSGTNGTKRF | 0.00000000200000.0 | 74(NGT) | 1 | 1526.784 | 1523.795739 | 23511576 | 36.61 | 1 | 5 | 5 | 1 | 1 | 0 | 0 | 1 | 3786.61005 | 0.0006 | 0.16 | -1.61 |
| Chymotrypsin | 2 | 174 | 510 | 1 | 3722 | 3787.6185 | 34.9983 | 701,158 | 3786.610675 | 21 | 66 | 79 | HAIHVSGTNGTKRF | 0.00000000200000.0 | 74(NGT) | 1 | 1526.784 | 1523.795739 | 23511576 | 36.61 | 2 | 4 | 5 | 2 | 0 | 1 | 0 | 0 | 3786.610049 | 0.0006 | 0.16 | -1.61 |
| Chymotrypsin | 2 | 174 | 541 | 1 | 3711 | 4768.9702 | 34.5332 | 659,494 | 4767.962375 | 21 | 66 | 79 | HAIHVSGTNGTKRF | 0.00000000200000.0 | 74(NGT) | 1 | 1526.784 | 1523.795739 | 23511576 | 36.61 | 1 | 7 | 6 | 4 | 0 | 1 | 0 | 1 | 4767.963712 | -0.0013 | -0.28 | -2.08 |
| Chymotrypsin | 2 | 174 | 541 | 1 | 3711 | 4768.9702 | 34.5332 | 659,494 | 4767.962375 | 21 | 66 | 79 | HAIHVSGTNGTKRF | 0.00000000200000.0 | 74(NGT) | 1 | 1526.784 | 1523.795739 | 23511576 | 36.61 | 2 | 8 | 6 | 3 | 1 | 0 | 0 | 0 | 4767.963713 | -0.0013 | -0.28 | -2.08 |
| Chymotrypsin | 2 | 174 | 541 | 1 | 3711 | 4768.9702 | 34.5332 | 659,494 | 4767.962375 | 21 | 66 | 79 | HAIHVSGTNGTKRF | 0.00000000200000.0 | 74(NGT) | 1 | 1526.784 | 1523.795739 | 23511576 | 36.61 | 3 | 2 | 10 | 0 | 2 | 1 | 1 | -1 | 4767.976282 | -0.0139 | -2.91 | -2.08 |
| Chymotrypsin | 2 | 174 | 616 | 1 | 3663 | 4502.9065 | 32.4813 | 585,892 | 4501.898675 | 21 | 66 | 79 | HAIHVSGTNGTKRF | 0.00000000200000.0 | 74(NGT) | 1 | 1526.784 | 1523.795739 | 23511576 | 36.61 | 1 | 6 | 7 | 4 | 0 | 0 | 0 | 0 | 4501.89993 | -0.0013 | -0.27 | -4.13 |
| Chymotrypsin | 2 | 174 | 640 | 1 | 3792 | 3463.5063 | 37.9887 | 567,755 | 3462.498475 | 21 | 66 | 79 | HAIHVSGTNGTKRF | 0.00000000200000.0 | 74(NGT) | 1 | 1526.784 | 1523.795739 | 23511576 | 36.61 | 1 | 3 | 5 | 1 | 1 | 0 | 0 | 1 | 3462.504402 | -0.0059 | -1.71 | 1.38 |
| Chymotrypsin | 2 | 174 | 640 | 1 | 3792 | 3463.5063 | 37.9887 | 567,755 | 3462.498475 | 21 | 66 | 79 | HAIHVSGTNGTKRF | 0.00000000200000.0 | 74(NGT) | 1 | 1526.784 | 1523.795739 | 23511576 | 36.61 | 2 | 2 | 5 | 2 | 0 | 1 | 1 | -1 | 3462.504401 | -0.0059 | -1.71 | 1.38 |
| Chymotrypsin | 2 | 174 | 672 | 1 | 3781 | 3543.5066 | 37.5237 | 538,365 | 3542.498775 | 21 | 66 | 79 | HAIHVSGTNGTKRF | 0.00000000200000.0 | 74(NGT) | 1 | 1526.784 | 1523.795739 | 23511576 | 36.61 | 1 | 6 | 3 | 1 | 1 | 0 | 0 | 1 | 3542.504128 | -0.0054 | -1.51 | 0.91 |
| Chymotrypsin | 2 | 174 | 672 | 1 | 3781 | 3543.5066 | 37.5237 | 538,365 | 3542.498775 | 21 | 66 | 79 | HAIHVSGTNGTKRF | 0.00000000200000.0 | 74(NGT) | 1 | 1526.784 | 1523.795739 | 23511576 | 36.61 | 2 | 5 | 3 | 2 | 0 | 1 | 0 | 0 | 3542.504127 | -0.0054 | -1.51 | 0.91 |
| Chymotrypsin | 2 | 174 | 712 | 1 | 3661 | 4461.8822 | 32.3932 | 514,503 | 4460.874375 | 21 | 66 | 79 | HAIHVSGTNGTKRF | 0.00000000200000.0 | 74(NGT) | 1 | 1526.784 | 1523.795739 | 23511576 | 36.61 | 1 | 7 | 6 | 4 | 0 | 0 | 0 | 1 | 4460.873381 | 0.0010 | 0.22 | -4.22 |
| Chymotrypsin | 2 | 174 | 712 | 1 | 3661 | 4461.8822 | 32.3932 | 514,503 | 4460.874375 | 21 | 66 | 79 | HAIHVSGTNGTKRF | 0.00000000200000.0 | 74(NGT) | 1 | 1526.784 | 1523.795739 | 23511576 | 36.61 | 2 | 2 | 10 | 0 | 2 | 0 | 1 | -1 | 4460.885951 | -0.0116 | -2.59 | -4.22 |
| Chymotrypsin | 2 | 174 | 752 | 1 | 3679 | 3480.5266 | 33.1661 | 492,622 | 3479.518775 | 21 | 66 | 79 | HAIHVSGTNGTKRF | 0.00000000200000.0 | 74(NGT) | 1 | 1526.784 | 1523.795739 | 23511576 | 36.61 | 1 | 4 | 5 | 2 | 0 | 0 | 0 | 1 | 3479.519718 | -0.0009 | -0.27 | -3.44 |
| Chymotrypsin | 2 | 174 | 770 | 1 | 3695 | 3496.521 | 33.8488 | 480,682 | 3495.513175 | 21 | 66 | 79 | HAIHVSGTNGTKRF | 0.00000000200000.0 | 74(NGT) | 1 | 1526.784 | 1523.795739 | 23511576 | 36.61 | 1 | 5 | 5 | 1 | 0 | 0 | 0 | 1 | 3495.514633 | -0.0015 | -0.41 | -2.76 |
| Chymotrypsin | 2 | 174 | 795 | 1 | 3676 | 3439.5005 | 33.036 | 465,233 | 3438.492675 | 21 | 66 | 79 | HAIHVSGTNGTKRF | 0.00000000200000.0 | 74(NGT) | 1 | 1526.784 | 1523.795739 | 23511576 | 36.61 | 1 | 5 | 4 | 2 | 0 | 0 | 0 | 1 | 3438.493169 | -0.0005 | -0.14 | -3.57 |
| Chymotrypsin | 2 | 174 | 824 | 1 | 3720 | 3949.6701 | 34.9149 | 452,102 | 3948.662275 | 21 | 66 | 79 | HAIHVSGTNGTKRF | 0.00000000200000.0 | 74(NGT) | 1 | 1526.784 | 1523.795739 | 23511576 | 36.61 | 1 | 6 | 5 | 1 | 1 | 0 | 0 | 1 | 3948.662874 | -0.0006 | -0.15 | -1.69 |
| Chymotrypsin | 2 | 174 | 824 | 1 | 3720 | 3949.6701 | 34.9149 | 452,102 | 3948.662275 | 21 | 66 | 79 | HAIHVSGTNGTKRF | 0.00000000200000.0 | 74(NGT) | 1 | 1526.784 | 1523.795739 | 23511576 | 36.61 | 2 | 5 | 5 | 2 | 0 | 1 | 0 | 0 | 3948.662873 | -0.0006 | -0.15 | -1.69 |
| Chymotrypsin | 2 | 174 | 887 | 1 | 3770 | 3057.3482 | 37.0538 | 422,683 | 3056.340375 | 21 | 66 | 79 | HAIHVSGTNGTKRF | 0.00000000200000.0 | 74(NGT) | 1 | 1526.784 | 1523.795739 | 23511576 | 36.61 | 1 | 2 | 3 | 2 | 0 | 1 | 1 | -1 | 3056.345655 | -0.0053 | -1.72 | 0.44 |
| Chymotrypsin | 2 | 174 | 887 | 1 | 3770 | 3057.3482 | 37.0538 | 422,683 | 3056.340375 | 21 | 66 | 79 | HAIHVSGTNGTKRF | 0.00000000200000.0 | 74(NGT) | 1 | 1526.784 | 1523.795739 | 23511576 | 36.61 | 2 | 3 | 3 | 1 | 1 | 0 | 0 | 0 | 3056.345656 | -0.0053 | -1.72 | 0.44 |
| Chymotrypsin | 2 | 174 | 951 | 1 | 3684 | 3642.5784 | 33.3755 | 400,144 | 3641.570575 | 21 | 66 | 79 | HAIHVSGTNGTKRF | 0.00000000200000.0 | 74(NGT) | 1 | 1526.784 | 1523.795739 | 23511576 | 36.61 | 1 | 5 | 5 | 2 | 0 | 0 | 0 | 1 | 3641.572542 | -0.0020 | -0.54 | -3.23 |
| Chymotrypsin | 2 | 174 | 974 | 1 | 3720 | 3746.5843 | 34.9149 | 392,858 | 3745.576475 | 21 | 66 | 79 | HAIHVSGTNGTKRF | 0.00000000200000.0 | 74(NGT) | 1 | 1526.784 | 1523.795739 | 23511576 | 36.61 | 1 | 5 | 4 | 2 | 0 | 1 | 0 | 1 | 3745.5835 | -0.0070 | -1.87 | -1.69 |
| Chymotrypsin | 2 | 174 | 974 | 1 | 3720 | 3746.5843 | 34.9149 | 392,858 | 3745.576475 | 21 | 66 | 79 | HAIHVSGTNGTKRF | 0.00000000200000.0 | 74(NGT) | 1 | 1526.784 | 1523.795739 | 23511576 | 36.61 | 2 | 6 | 4 | 1 | 1 | 0 | 0 | 1 | 3745.583501 | -0.0070 | -1.87 | -1.69 |
| Chymotrypsin | 2 | 174 | 1001 | 1 | 3787 | 3584.5325 | 37.7792 | 381,505 | 3583.524675 | 21 | 66 | 79 | HAIHVSGTNGTKRF | 0.00000000200000.0 | 74(NGT) | 1 | 1526.784 | 1523.795739 | 23511576 | 36.61 | 1 | 4 | 4 | 2 | 0 | 1 | 0 | 0 | 3583.530676 | -0.0060 | -1.67 | 1.17 |
| Chymotrypsin | 2 | 174 | 1001 | 1 | 3787 | 3584.5325 | 37.7792 | 381,505 | 3583.524675 | 21 | 66 | 79 | HAIHVSGTNGTKRF | 0.00000000200000.0 | 74(NGT) | 1 | 1526.784 | 1523.795739 | 23511576 | 36.61 | 2 | 5 | 4 | 1 | 1 | 0 | 0 | 0 | 3583.530677 | -0.0060 | -1.67 | 1.17 |
| Chymotrypsin | 2 | 174 | 1098 | 1 | 3666 | 4178.8035 | 32.6112 | 350,235 | 4177.795675 | 21 | 66 | 79 | HAIHVSGTNGTKRF | 0.00000000200000.0 | 74(NGT) | 1 | 1526.784 | 1523.795739 | 23511576 | 36.61 | 1 | 4 | 7 | 4 | 0 | 0 | 0 | 0 | 4177.794282 | 0.0014 | 0.33 | -4.00 |
| Chymotrypsin | 2 | 174 | 1223 | 1 | 3687 | 3334.4696 | 33.5058 | 312,779 | 3333.461775 | 21 | 66 | 79 | HAIHVSGTNGTKRF | 0.00000000200000.0 | 74(NGT) | 1 | 1526.784 | 1523.795739 | 23511576 | 36.61 | 1 | 4 | 5 | 1 | 0 | 0 | 0 | 1 | 3333.461809 | 0.0000 | -0.01 | -3.10 |
| Chymotrypsin | 2 | 174 | 1245 | 1 | 3658 | 4664.9707 | 32.2645 | 307,358 | 4663.962875 | 21 | 66 | 79 | HAIHVSGTNGTKRF | 0.00000000200000.0 | 74(NGT) | 1 | 1526.784 | 1523.795739 | 23511576 | 36.61 | 1 | 7 | 7 | 4 | 0 | 0 | 0 | 0 | 4663.952754 | 0.0101 | 2.17 | -4.34 |
| Chymotrypsin | 2 | 174 | 1343 | 1 | 3687 | 4153.77 | 33.5058 | 283,926 | 4152.762175 | 21 | 66 | 79 | HAIHVSGTNGTKRF | 0.00000000200000.0 | 74(NGT) | 1 | 1526.784 | 1523.795739 | 23511576 | 36.61 | 1 | 6 | 6 | 3 | 0 | 0 | 0 | 1 | 4152.762648 | -0.0005 | -0.11 | -3.10 |
| Chymotrypsin | 2 | 174 | 1373 | 1 | 3685 | 3375.495 | 33.4199 | 279,709 | 3374.487175 | 21 | 66 | 79 | HAIHVSGTNGTKRF | 0.00000000200000.0 | 74(NGT) | 1 | 1526.784 | 1523.795739 | 23511576 | 36.61 | 1 | 3 | 6 | 1 | 0 | 0 | 0 | 1 | 3374.488358 | -0.0012 | -0.35 | -3.19 |
| Chymotrypsin | 2 | 174 | 1396 | 1 | 3663 | 4689.9934 | 32.4813 | 276,809 | 4688.985575 | 21 | 66 | 79 | HAIHVSGTNGTKRF | 0.00000000200000.0 | 74(NGT) | 1 | 1526.784 | 1523.795739 | 23511576 | 36.61 |  |  |  |  |  |  |  |  |  |  |  |  |
| Chymotrypsin | 2 | 174 | 1425 | 1 | 3718 | 4257.779 | 34.8306 | 268,280 | 4256.771175 | 21 | 66 | 79 | HAIHVSGTNGTKRF | 0.00000000200000.0 | 74(NGT) | 1 | 1526.784 | 1523.795739 | 23511576 | 36.61 | 1 | 6 | 5 | 3 | 0 | 1 | 0 | 1 | 4256.773606 | -0.0024 | -0.57 | -1.78 |
| Chymotrypsin | 2 | 174 | 1425 | 1 | 3718 | 4257.779 | 34.8306 | 268,280 | 4256.771175 | 21 | 66 | 79 | HAIHVSGTNGTKRF | 0.00000000200000.0 | 74(NGT) | 1 | 1526.784 | 1523.795739 | 23511576 | 36.61 | 2 | 7 | 5 | 2 | 1 | 0 | 0 | 0 | 4256.773607 | -0.0024 | -0.57 | -1.78 |
| Chymotrypsin | 2 | 174 | 1453 | 1 | 3689 | 2969.3356 | 33.5921 | 264,304 | 2968.327775 | 21 | 66 | 79 | HAIHVSGTNGTKRF | 0.00000000200000.0 | 74(NGT) | 1 | 1526.784 | 1523.795739 | 23511576 | 36.61 | 1 | 3 | 4 | 1 | 0 | 0 | 0 | 1 | 2968.329612 | -0.0018 | -0.61 | -3.02 |
| Chymotrypsin | 2 | 174 | 1456 | 1 | 3665 | 4315.8232 | 32.5693 | 264,062 | 4314.815375 | 21 | 66 | 79 | HAIHVSGTNGTKRF | 0.00000000200000.0 | 74(NGT) | 1 | 1526.784 | 1523.795739 | 23511576 | 36.61 | 1 | 7 | 6 | 3 | 0 | 0 | 0 | 0 | 4314.815472 | -0.0001 | -0.02 | -4.04 |
| Chymotrypsin | 2 | 174 | 1461 | 1 | 3665 | 4543.9313 | 32.5693 | 263,629 | 4542.923475 | 21 | 66 | 79 | HAIHVSGTNGTKRF | 0.00000000200000.0 | 74(NGT) | 1 | 1526.784 | 1523.795739 | 23511576 | 36.61 | 1 | 5 | 8 | 4 | 0 | 0 | 0 | 0 | 4542.926479 | -0.0030 | -0.66 | -4.04 |
| Chymotrypsin | 2 | 174 | 1606 | 1 | 3676 | 4023.7065 | 33.036 | 241,714 | 4022.698675 | 21 | 66 | 79 | HAIHVSGTNGTKRF | 0.00000000200000.0 | 74(NGT) | 1 | 1526.784 | 1523.795739 | 23511576 | 36.61 | 1 | 7 | 6 | 1 | 0 | 0 | 0 | 0 | 4022.699654 | -0.0010 | -0.24 | -3.57 |
| Chymotrypsin | 2 | 174 | 1792 | 1 | 3833 | 3729.5722 | 39.7527 | 218,238 | 3728.564375 | 21 | 66 | 79 | HAIHVSGTNGTKRF | 0.00000000200000.0 | 74(NGT) | 1 | 1526.784 | 1523.795739 | 23511576 | 36.61 | 1 | 5 | 4 | 0 | 2 | 0 | 0 | 1 | 3728.568185 | -0.0038 | -1.02 | 3.14 |
| Chymotrypsin | 2 | 174 | 1792 | 1 | 3833 | 3729.5722 | 39.7527 | 218,238 | 3728.564375 | 21 | 66 | 79 | HAIHVSGTNGTKRF | 0.00000000200000.0 | 74(NGT) | 1 | 1526.784 | 1523.795739 | 23511576 | 36.61 | 2 | 3 | 4 | 2 | 0 | 2 | 1 | -1 | 3728.568183 | -0.0038 | -1.02 | 3.14 |
| Chymotrypsin | 2 | 174 | 1792 | 1 | 3833 | 3729.5722 | 39.7527 | 218,238 | 3728.564375 | 21 | 66 | 79 | HAIHVSGTNGTKRF | 0.00000000200000.0 | 74(NGT) | 1 | 1526.784 | 1523.795739 | 23511576 | 36.61 | 3 | 4 | 4 | 1 | 1 | 1 | 0 | 0 | 3728.568184 | -0.0038 | -1.02 | 3.14 |
| Chymotrypsin | 2 | 174 | 1796 | 1 | 3682 | 3521.5518 | 33.2916 | 217,780 | 3520.543975 | 21 | 66 | 79 | HAIHVSGTNGTKRF | 0.00000000200000.0 | 74(NGT) | 1 | 1526.784 | 1523.795739 | 23511576 | 36.61 | 1 | 3 | 6 | 2 | 0 | 0 | 0 | 0 | 3520.546267 | -0.0023 | -0.65 | -3.32 |
| Chymotrypsin | 2 | 174 | 1917 | 1 | 3788 | 3625.5557 | 37.8213 | 202,544 | 3624.547875 | 21 | 66 | 79 | HAIHVSGTNGTKRF | 0.00000000200000.0 | 74(NGT) | 1 | 1526.784 | 1523.795739 | 23511576 | 36.61 | 1 | 4 | 5 | 1 | 1 | 0 | 0 | 1 | 3624.557226 | -0.0094 | -2.57 | 1.21 |
| Chymotrypsin | 2 | 174 | 1917 | 1 | 3788 | 3625.5557 | 37.8213 | 202,544 | 3624.547875 | 21 | 66 | 79 | HAIHVSGTNGTKRF | 0.00000000200000.0 | 74(NGT) | 1 | 1526.784 | 1523.795739 | 23511576 | 36.61 | 2 | 3 | 5 | 2 | 0 | 1 | 0 | 0 | 3624.557225 | -0.0093 | -2.57 | 1.21 |
| Chymotrypsin | 2 | 174 | 1951 | 1 | 3898 | 4045.7059 | 42.5315 | 199,085 | 4044.698075 | 21 | 66 | 79 | HAIHVSGTNGTKRF | 0.00000000200000.0 | 74(NGT) | 1 | 1526.784 | 1523.795739 | 23511576 | 36.61 | 1 | 3 | 5 | 1 | 3 | 0 | 1 | -1 | 4044.695236 | 0.0028 | 0.7 | 5.92 |
| Chymotrypsin | 2 | 174 | 1951 | 1 | 3898 | 4045.7059 | 42.5315 | 199,085 | 4044.698075 | 21 | 66 | 79 | HAIHVSGTNGTKRF | 0.00000000200000.0 | 74(NGT) | 1 | 1526.784 | 1523.795739 | 23511576 | 36.61 | 2 | 2 | 5 | 2 | 2 | 1 | 1 | -1 | 4044.695235 | 0.0028 | 0.7 | 5.92 |
| Chymotrypsin | 2 | 174 | 2004 | 1 | 3687 | 3131.3895 | 33.5058 | 193,881 | 3130.381675 | 21 | 66 | 79 | HAIHVSGTNGTKRF | 0.00000000200000.0 | 74(NGT) | 1 | 1526.784 | 1523.795739 | 23511576 | 36.61 | 1 | 4 | 4 | 1 | 0 | 0 | 0 | 1 | 3130.382436 | -0.0008 | -0.24 | -3.10 |
| Chymotrypsin | 2 | 174 | 2015 | 1 | 3724 | 4460.8629 | 35.0816 | 192,782 | 4459.855075 | 21 | 66 | 79 | HAIHVSGTNGTKRF | 0.00000000200000.0 | 74(NGT) | 1 | 1526.784 | 1523.795739 | 23511576 | 36.61 | 1 | 7 | 6 | 2 | 1 | 0 | 0 | 0 | 4459.85298 | 0.0021 | 0.46 | -1.53 |
| Chymotrypsin | 2 | 174 | 2015 | 1 | 3724 | 4460.8629 | 35.0816 | 192,782 | 4459.855075 | 21 | 66 | 79 | HAIHVSGTNGTKRF | 0.00000000200000.0 | 74(NGT) | 1 | 1526.784 | 1523.795739 | 23511576 | 36.61 | 2 | 6 | 6 | 3 | 0 | 1 | 0 | 0 | 4459.852979 | 0.0021 | 0.47 | -1.53 |
| Chymotrypsin | 2 | 174 | 2031 | 1 | 3717 | 4663.9396 | 34.7882 | 191,184 | 4662.931775 | 21 | 66 | 79 | HAIHVSGTNGTKRF | 0.00000000200000.0 | 74(NGT) | 1 | 1526.784 | 1523.795739 | 23511576 | 36.61 | 1 | 6 | 7 | 3 | 0 | 1 | 0 | 0 | 4662.932352 | -0.0006 | -0.12 | -1.82 |
| Chymotrypsin | 2 | 174 | 2031 | 1 | 3717 | 4663.9396 | 34.7882 | 191,184 | 4662.931775 | 21 | 66 | 79 | HAIHVSGTNGTKRF | 0.00000000200000.0 | 74(NGT) | 1 | 1526.784 | 1523.795739 | 23511576 | 36.61 | 2 | 7 | 7 | 2 | 1 | 0 | 0 | 0 | 4662.932353 | -0.0006 | -0.12 | -1.82 |
| Chymotrypsin | 2 | 174 | 2077 | 1 | 3657 | 4852.0418 | 32.2234 | 187,976 | 4851.033975 | 21 | 66 | 79 | HAIHVSGTNGTKRF | 0.00000000200000.0 | 74(NGT) | 1 | 1526.784 | 1523.795739 | 23511576 | 36.61 | 1 | 8 | 10 | 0 | 0 | 0 | 0 | 1 | 4851.012061 | 0.0219 | 4.51 | -4.39 |
| Chymotrypsin | 2 | 174 | 2089 | 1 | 3712 | 4810.0001 | 34.5742 | 186,886 | 4808.992275 | 21 | 66 | 79 | HAIHVSGTNGTKRF | 0.00000000200000.0 | 74(NGT) | 1 | 1526.784 | 1523.795739 | 23511576 | 36.61 | 1 | 7 | 7 | 3 | 1 | 0 | 0 | 0 | 4808.990262 | 0.0020 | 0.41 | -2.03 |
| Chymotrypsin | 2 | 174 | 2089 | 1 | 3712 | 4810.0001 | 34.5742 | 186,886 | 4808.992275 | 21 | 66 | 79 | HAIHVSGTNGTKRF | 0.00000000200000.0 | 74(NGT) | 1 | 1526.784 | 1523.795739 | 23511576 | 36.61 | 2 | 6 | 7 | 4 | 0 | 1 | 0 | 0 | 4808.990261 | 0.0020 | 0.41 | -2.03 |
| Chymotrypsin | 2 | 174 | 2096 | 1 | 3685 | 3293.4439 | 33.4199 | 186,531 | 3292.436075 | 21 | 66 | 79 | HAIHVSGTNGTKRF | 0.00000000200000.0 | 74(NGT) | 1 | 1526.784 | 1523.795739 | 23511576 | 36.61 | 1 | 5 | 4 | 1 | 0 | 0 | 0 | 1 | 3292.43526 | 0.0008 | 0.24 | -3.19 |
| Chymotrypsin | 2 | 174 | 2188 | 1 | 3703 | 3861.6537 | 34.1926 | 179,008 | 3860.645875 | 21 | 66 | 79 | HAIHVSGTNGTKRF | 0.00000000200000.0 | 74(NGT) | 1 | 1526.784 | 1523.795739 | 23511576 | 36.61 | 1 | 6 | 6 | 1 | 0 | 0 | 0 | 1 | 3860.64683 | -0.0010 | -0.24 | -2.42 |
| Chymotrypsin | 2 | 174 | 2210 | 1 | 3667 | 4356.8498 | 32.6524 | 177,464 | 4355.841975 | 21 | 66 | 79 | HAIHVSGTNGTKRF | 0.00000000200000.0 | 74(NGT) | 1 | 1526.784 | 1523.795739 | 23511576 | 36.61 | 1 | 6 | 7 | 3 | 0 | 0 | 0 | 0 | 4355.842021 | 0.0000 | -0.01 | -3.96 |
| Chymotrypsin | 2 | 174 | 2214 | 1 | 3679 | 4372.8441 | 33.1661 | 177,254 | 4371.836275 | 21 | 66 | 79 | HAIHVSGTNGTKRF | 0.00000000200000.0 | 74(NGT) | 1 | 1526.784 | 1523.795739 | 23511576 | 36.61 | 1 | 7 | 7 | 2 | 0 | 0 | 0 | 0 | 4371.836936 | -0.0007 | -0.15 | -3.44 |
| Chymotrypsin | 2 | 174 | 2236 | 1 | 3707 | 4972.0544 | 34.3624 | 175,346 | 4971.046575 | 21 | 66 | 79 | HAIHVSGTNGTKRF | 0.00000000200000.0 | 74(NGT) | 1 | 1526.784 | 1523.795739 | 23511576 | 36.61 | 1 | 8 | 7 | 3 | 1 | 0 | 0 | 0 | 4971.043086 | 0.0035 | 0.7 | -2.25 |
| Chymotrypsin | 2 | 174 | 2236 | 1 | 3707 | 4972.0544 | 34.3624 | 175,346 | 4971.046575 | 21 | 66 | 79 | HAIHVSGTNGTKRF | 0.00000000200000.0 | 74(NGT) | 1 | 1526.784 | 1523.795739 | 23511576 | 36.61 | 2 | 7 | 7 | 4 | 0 | 1 | 0 | 0 | 4971.043085 | 0.0035 | 0.7 | -2.25 |
| Chymotrypsin | 2 | 174 | 2245 | 1 | 3676 | 3991.7202 | 33.036 | 174,094 | 3990.712375 | 21 | 66 | 79 | HAIHVSGTNGTKRF | 0.00000000200000.0 | 74(NGT) | 1 | 1526.784 | 1523.795739 | 23511576 | 36.61 | 1 | 5 | 6 | 3 | 0 | 0 | 0 | 0 | 3990.709824 | 0.0026 | 0.63 | -3.57 |
| Chymotrypsin | 2 | 174 | 2395 | 1 | 3663 | 4518.9032 | 32.4813 | 161,890 | 4517.895375 | 21 | 66 | 79 | HAIHVSGTNGTKRF | 0.00000000200000.0 | 74(NGT) | 1 | 1526.784 | 1523.795739 | 23511576 | 36.61 | 1 | 7 | 7 | 3 | 0 | 0 | 0 | 0 | 4517.894845 | 0.0005 | 0.11 | -4.13 |
| Chymotrypsin | 2 | 174 | 2414 | 1 | 3667 | 4032.7435 | 32.6524 | 160,525 | 4031.735675 | 21 | 66 | 79 | HAIHVSGTNGTKRF | 0.00000000200000.0 | 74(NGT) | 1 | 1526.784 | 1523.795739 | 23511576 | 36.61 | 1 | 4 | 7 | 3 | 0 | 0 | 0 | 0 | 4031.736373 | -0.0007 | -0.17 | -3.96 |
| Chymotrypsin | 2 | 174 | 2533 | 1 | 3700 | 4007.7096 | 34.0623 | 152,953 | 4006.701775 | 21 | 66 | 79 | HAIHVSGTNGTKRF | 0.00000000200000.0 | 74(NGT) | 1 | 1526.784 | 1523.795739 | 23511576 | 36.61 | 1 | 6 | 6 | 2 | 0 | 0 | 0 | 1 | 4006.704739 | -0.0030 | -0.73 | -2.55 |
| Chymotrypsin | 2 | 174 | 2646 | 1 | 3669 | 4169.7582 | 32.7395 | 146,196 | 4168.750375 | 21 | 66 | 79 | HAIHVSGTNGTKRF | 0.00000000200000.0 | 74(NGT) | 1 | 1526.784 | 1523.795739 | 23511576 | 36.61 | 1 | 7 | 6 | 2 | 0 | 0 | 0 | 0 | 4168.757563 | -0.0072 | -1.72 | -3.87 |
| Chymotrypsin | 2 | 174 | 2674 | 1 | 3682 | 4226.7864 | 33.2916 | 144,953 | 4225.778575 | 21 | 66 | 79 | HAIHVSGTNGTKRF | 0.00000000200000.0 | 74(NGT) | 1 | 1526.784 | 1523.795739 | 23511576 | 36.61 | 1 | 7 | 7 | 1 | 0 | 0 | 0 | 0 | 4225.779027 | -0.0005 | -0.1 | -3.32 |
| Chymotrypsin | 2 | 174 | 2681 | 1 | 3674 | 3950.6875 | 32.9519 | 144,569 | 3949.679675 | 21 | 66 | 79 | HAIHVSGTNGTKRF | 0.00000000200000.0 | 74(NGT) | 1 | 1526.784 | 1523.795739 | 23511576 | 36.61 | 1 | 6 | 5 | 3 | 0 | 0 | 0 | 0 | 3949.683275 | -0.0036 | -0.91 | -3.66 |
| Chymotrypsin | 2 | 174 | 2690 | 1 | 3698 | 3172.4162 | 33.9771 | 144,153 | 3171.408375 | 21 | 66 | 79 | HAIHVSGTNGTKRF | 0.00000000200000.0 | 74(NGT) | 1 | 1526.784 | 1523.795739 | 23511576 | 36.61 | 1 | 3 | 5 | 1 | 0 | 0 | 0 | 1 | 3171.408985 | -0.0006 | -0.19 | -2.63 |
| Chymotrypsin | 2 | 174 | 2734 | 1 | 3835 | 3770.5983 | 39.8388 | 140,944 | 3769.590475 | 21 | 66 | 79 | HAIHVSGTNGTKRF | 0.00000000200000.0 | 74(NGT) | 1 | 1526.784 | 1523.795739 | 23511576 | 36.61 | 1 | 4 | 5 | 0 | 2 | 0 | 0 | 1 | 3769.594734 | -0.0043 | -1.12 | 3.23 |
| Chymotrypsin | 2 | 174 | 2734 | 1 | 3835 | 3770.5983 | 39.8388 | 140,944 | 3769.590475 | 21 | 66 | 79 | HAIHVSGTNGTKRF | 0.00000000200000.0 | 74(NGT) | 1 | 1526.784 | 1523.795739 | 23511576 | 36.61 | 2 | 2 | 5 | 2 | 0 | 2 | 1 | -1 | 3769.594732 | -0.0043 | -1.12 | 3.23 |
| Chymotrypsin | 2 | 174 | 2734 | 1 | 3835 | 3770.5983 | 39.8388 | 140,944 | 3769.590475 | 21 | 66 | 79 | HAIHVSGTNGTKRF | 0.00000000200000.0 | 74(NGT) | 1 | 1526.784 | 1523.795739 | 23511576 | 36.61 | 3 | 3 | 5 | 1 | 1 | 1 | 1 | -1 | 3769.594733 | -0.0043 | -1.12 | 3.23 |
| Chymotrypsin | 2 | 174 | 2783 | 1 | 3727 | 3600.5354 | 35.2098 | 137,927 | 3599.527575 | 21 | 66 | 79 | HAIHVSGTNGTKRF | 0.00000000200000.0 | 74(NGT) | 1 | 1526.784 | 1523.795739 | 23511576 | 36.61 | 1 | 6 | 4 | 0 | 1 | 0 | 0 | 1 | 3599.525592 | 0.0020 | 0.55 | -1.40 |
| Chymotrypsin | 2 | 174 | 2783 | 1 | 3727 | 3600.5354 | 35.2098 | 137,927 | 3599.527575 | 21 | 66 | 79 | HAIHVSGTNGTKRF | 0.00000000200000.0 | 74(NGT) | 1 | 1526.784 | 1523.795739 | 23511576 | 36.61 | 2 | 5 | 4 | 1 | 0 | 1 | 0 | 1 | 3599.525591 | 0.0020 | 0.55 | -1.40 |
| Chymotrypsin | 2 | 174 | 2820 | 1 | 3663 | 4705.9887 | 32.4813 | 135,784 | 4704.980875 | 21 | 66 | 79 | HAIHVSGTNGTKRF | 0.00000000200000.0 | 74(NGT) | 1 | 1526.784 | 1523.795739 | 23511576 | 36.61 | 1 | 6 | 8 | 4 | 0 | 0 | 0 | 0 | 4704.979303 | 0.0016 | 0.33 | -4.13 |
| Chymotrypsin | 2 | 174 | 2825 | 1 | 3838 | 3567.5166 | 39.9683 | 135,349 | 3566.508775 | 21 | 66 | 79 | HAIHVSGTNGTKRF | 0.00000000200000.0 | 74(NGT) | 1 | 1526.784 | 1523.795739 | 23511576 | 36.61 | 1 | 2 | 4 | 2 | 0 | 2 | 1 | -1 | 3566.515359 | -0.0066 | -1.84 | 3.36 |
| Chymotrypsin | 2 | 174 | 2825 | 1 | 3838 | 3567.5166 | 39.9683 | 135,349 | 3566.508775 | 21 | 66 | 79 | HAIHVSGTNGTKRF | 0.00000000200000.0 | 74(NGT) | 1 | 1526.784 | 1523.795739 | 23511576 | 36.61 | 2 | 3 | 4 | 1 | 1 | 1 | 1 | -1 | 3566.51536 | -0.0066 | -1.84 | 3.36 |
| Chymotrypsin | 2 | 174 | 2825 | 1 | 3838 | 3567.5166 | 39.9683 | 135,349 | 3566.508775 | 21 | 66 | 79 | HAIHVSGTNGTKRF | 0.00000000200000.0 | 74(NGT) | 1 | 1526.784 | 1523.795739 | 23511576 | 36.61 | 3 | 4 | 4 | 0 | 2 | 0 | 0 | 0 | 3566.515361 | -0.0066 | -1.84 | 3.36 |
| Chymotrypsin | 2 | 174 | 2884 | 1 | 3673 | 4340.8568 | 32.9085 | 132,707 | 4339.848975 | 21 | 66 | 79 | HAIHVSGTNGTKRF | 0.00000000200000.0 | 74(NGT) | 1 | 1526.784 | 1523.795739 | 23511576 | 36.61 | 1 | 5 | 7 | 4 | 0 | 0 | 0 | 0 | 4339.847106 | 0.0019 | 0.43 | -3.70 |
| Chymotrypsin | 2 | 174 | 2888 | 1 | 3654 | 4957.0747 | 32.094 | 132,586 | 4956.066875 | 21 | 66 | 79 | HAIHVSGTNGTKRF | 0.00000000200000.0 | 74(NGT) | 1 | 1526.784 | 1523.795739 | 23511576 | 36.61 | 1 | 9 | 9 | 1 | 0 | 0 | 0 | 1 | 4956.043421 | 0.0235 | 4.73 | -4.51 |
| Chymotrypsin | 2 | 174 | 3019 | 1 | 3682 | 3115.3959 | 33.2916 | 126,060 | 3114.388075 | 21 | 66 | 79 | HAIHVSGTNGTKRF | 0.00000000200000.0 | 74(NGT) | 1 | 1526.784 | 1523.795739 | 23511576 | 36.61 | 1 | 3 | 4 | 2 | 0 | 0 | 0 | 1 | 3114.387521 | 0.0006 | 0.17 | -3.32 |
| Chymotrypsin | 2 | 174 | 3032 | 1 | 3667 | 3293.4433 | 32.6524 | 125,296 | 3292.435475 | 21 | 66 | 79 | HAIHVSGTNGTKRF | 0.00000000200000.0 | 74(NGT) | 1 | 1526.784 | 1523.795739 | 23511576 | 36.61 | 1 | 5 | 4 | 1 | 0 | 0 | 0 | 1 | 3292.43526 | 0.0002 | 0.06 | -3.96 |
| Chymotrypsin | 2 | 174 | 3153 | 1 | 3721 | 3828.6438 | 34.9566 | 119,668 | 3827.635975 | 21 | 66 | 79 | HAIHVSGTNGTKRF | 0.00000000200000.0 | 74(NGT) | 1 | 1526.784 | 1523.795739 | 23511576 | 36.61 | 1 | 3 | 6 | 2 | 0 | 1 | 0 | 1 | 3827.636598 | -0.0006 | -0.16 | -1.65 |
| Chymotrypsin | 2 | 174 | 3153 | 1 | 3721 | 3828.6438 | 34.9566 | 119,668 | 3827.635975 | 21 | 66 | 79 | HAIHVSGTNGTKRF | 0.00000000200000.0 | 74(NGT) | 1 | 1526.784 | 1523.795739 | 23511576 | 36.61 | 2 | 4 | 6 | 1 | 1 | 0 | 0 | 0 | 3827.636599 | -0.0006 | -0.16 | -1.65 |
| Chymotrypsin | 2 | 174 | 3250 | 1 | 3731 | 3438.4794 | 35.3805 | 115,770 | 3437.471575 | 21 | 66 | 79 | HAIHVSGTNGTKRF | 0.00000000200000.0 | 74(NGT) | 1 | 1526.784 | 1523.795739 | 23511576 | 36.61 | 1 | 4 | 4 | 1 | 0 | 1 | 0 | 1 | 3437.472767 | -0.0012 | -0.34 | -1.23 |
| Chymotrypsin | 2 | 174 | 3250 | 1 | 3731 | 3438.4794 | 35.3805 | 115,770 | 3437.471575 | 21 | 66 | 79 | HAIHVSGTNGTKRF | 0.00000000200000.0 | 74(NGT) | 1 | 1526.784 | 1523.795739 | 23511576 | 36.61 | 2 | 5 | 4 | 0 | 1 | 0 | 0 | 1 | 3437.472768 | -0.0012 | -0.34 | -1.23 |
| Chymotrypsin | 2 | 174 | 3302 | 1 | 3767 | 4930.0032 | 36.9242 | 113,783 | 4928.995375 | 21 | 66 | 79 | HAIHVSGTNGTKRF | 0.00000000200000.0 | 74(NGT) | 1 | 1526.784 | 1523.795739 | 23511576 | 36.61 | 1 | 9 | 6 | 1 | 2 | 0 | 0 | 1 | 4928.996136 | -0.0008 | -0.15 | 0.32 |
| Chymotrypsin | 2 | 174 | 3302 | 1 | 3767 | 4930.0032 | 36.9242 | 113,783 | 4928.995375 | 21 | 66 | 79 | HAIHVSGTNGTKRF | 0.00000000200000.0 | 74(NGT) | 1 | 1526.784 | 1523.795739 | 23511576 | 36.61 | 2 | 7 | 6 | 3 | 0 | 2 | 0 | 0 | 4928.996134 | -0.0008 | -0.15 | 0.32 |
| Chymotrypsin | 2 | 174 | 3302 | 1 | 3767 | 4930.0032 | 36.9242 | 113,783 | 4928.995375 | 21 | 66 | 79 | HAIHVSGTNGTKRF | 0.00000000200000.0 | 74(NGT) | 1 | 1526.784 | 1523.795739 | 23511576 | 36.61 | 3 | 8 | 6 | 2 | 1 | 1 | 0 | 0 | 4928.996135 | -0.0008 | -0.15 | 0.32 |
| Chymotrypsin | 2 | 174 | 3382 | 1 | 3677 | 3236.4206 | 33.0797 | 111,080 | 3235.412775 | 21 | 66 | 79 | HAIHVSGTNGTKRF | 0.00000000200000.0 | 74(NGT) | 1 | 1526.784 | 1523.795739 | 23511576 | 36.61 | 1 | 5 | 3 | 2 | 0 | 0 | 0 | 0 | 3235.413796 | -0.0010 | -0.31 | -3.53 |
| Chymotrypsin | 2 | 174 | 3444 | 1 | 3848 | 3389.466 | 40.3994 | 108,737 | 3388.458175 | 21 | 66 | 79 | HAIHVSGTNGTKRF | 0.00000000200000.0 | 74(NGT) | 1 | 1526.784 | 1523.795739 | 23511576 | 36.61 | 1 | 2 | 4 | 1 | 2 | 0 | 1 | -1 | 3388.467622 | -0.0094 | -2.78 | 3.79 |
| Chymotrypsin | 2 | 174 | 3463 | 1 | 3716 | 4476.8561 | 34.7441 | 108,072 | 4475.848275 | 21 | 66 | 79 | HAIHVSGTNGTKRF | 0.00000000200000.0 | 74(NGT) | 1 | 1526.784 | 1523.795739 | 23511576 | 36.61 | 1 | 8 | 6 | 1 | 1 | 0 | 0 | 1 | 4475.847895 | 0.0004 | 0.08 | -1.86 |
| Chymotrypsin | 2 | 174 | 3463 | 1 | 3716 | 4476.8561 | 34.7441 | 108,072 | 4475.848275 | 21 | 66 | 79 | HAIHVSGTNGTKRF | 0.00000000200000.0 | 74(NGT) | 1 | 1526.784 | 1523.795739 | 23511576 | 36.61 | 2 | 7 | 6 | 2 | 0 | 1 | 0 | 0 | 4475.847894 | 0.0004 | 0.08 | -1.86 |
| Chymotrypsin | 2 | 174 | 3536 | 1 | 3714 | 4752.9864 | 34.6605 | 105,853 | 4751.978575 | 21 | 66 | 79 | HAIHVSGTNGTKRF | 0.00000000200000.0 | 74(NGT) | 1 | 1526.784 | 1523.795739 | 23511576 | 36.61 | 1 | 7 | 6 | 4 | 1 | 0 | 0 | 0 | 4751.968798 | 0.0098 | 2.05 | -1.95 |
| Chymotrypsin | 2 | 174 | 3536 | 1 | 3714 | 4752.9864 | 34.6605 | 105,853 | 4751.978575 | 21 | 66 | 79 | HAIHVSGTNGTKRF | 0.00000000200000.0 | 74(NGT) | 1 | 1526.784 | 1523.795739 | 23511576 | 36.61 | 2 | 2 | 10 | 0 | 3 | 0 | 1 | -1 | 4751.981368 | -0.0028 | -0.58 | -1.95 |
| Chymotrypsin | 2 | 174 | 3614 | 1 | 3669 | 4445.8827 | 32.7395 | 103,205 | 4444.874875 | 21 | 66 | 79 | HAIHVSGTNGTKRF | 0.00000000200000.0 | 74(NGT) | 1 | 1526.784 | 1523.795739 | 23511576 | 36.61 | 1 | 8 | 8 | 0 | 0 | 0 | 0 | 1 | 4444.853315 | 0.0216 | 4.85 | -3.87 |
| Chymotrypsin | 2 | 174 | 3796 | 1 | 3780 | 3705.5616 | 37.4792 | 97,362 | 3704.553775 | 21 | 66 | 79 | HAIHVSGTNGTKRF | 0.00000000200000.0 | 74(NGT) | 1 | 1526.784 | 1523.795739 | 23511576 | 36.61 | 1 | 6 | 3 | 2 | 0 | 1 | 0 | 0 | 3704.556951 | -0.0032 | -0.85 | 0.87 |
| Chymotrypsin | 2 | 174 | 3796 | 1 | 3780 | 3705.5616 | 37.4792 | 97,362 | 3704.553775 | 21 | 66 | 79 | HAIHVSGTNGTKRF | 0.00000000200000.0 | 74(NGT) | 1 | 1526.784 | 1523.795739 | 23511576 | 36.61 | 2 | 7 | 3 | 1 | 1 | 0 | 1 | -1 | 3704.556952 | -0.0032 | -0.85 | 0.87 |
| Chymotrypsin | 2 | 174 | 3808 | 1 | 3796 | 3098.3748 | 38.1615 | 97,118 | 3097.366975 | 21 | 66 | 79 | HAIHVSGTNGTKRF | 0.00000000200000.0 | 74(NGT) | 1 | 1526.784 | 1523.795739 | 23511576 | 36.61 | 1 | 2 | 4 | 1 | 1 | 0 | 1 | -1 | 3097.372205 | -0.0052 | -1.68 | 1.55 |
| Chymotrypsin | 2 | 174 | 3819 | 1 | 3730 | 3803.6139 | 35.3364 | 96,958 | 3802.606075 | 21 | 66 | 79 | HAIHVSGTNGTKRF | 0.00000000200000.0 | 74(NGT) | 1 | 1526.784 | 1523.795739 | 23511576 | 36.61 | 1 | 6 | 5 | 0 | 1 | 0 | 0 | 1 | 3802.604965 | 0.0011 | 0.29 | -1.27 |
| Chymotrypsin | 2 | 174 | 3819 | 1 | 3730 | 3803.6139 | 35.3364 | 96,958 | 3802.606075 | 21 | 66 | 79 | HAIHVSGTNGTKRF | 0.00000000200000.0 | 74(NGT) | 1 | 1526.784 | 1523.795739 | 23511576 | 36.61 | 2 | 5 | 5 | 1 | 0 | 1 | 0 | 1 | 3802.604964 | 0.0011 | 0.29 | -1.27 |
| Chymotrypsin | 2 | 174 | 4288 | 1 | 3673 | 3829.6747 | 32.9085 | 84,165 | 3828.666875 | 21 | 66 | 79 | HAIHVSGTNGTKRF | 0.00000000200000.0 | 74(NGT) | 1 | 1526.784 | 1523.795739 | 23511576 | 36.61 | 1 | 4 | 6 | 3 | 0 | 0 | 0 | 0 | 3828.657 | 0.0099 | 2.57 | -3.70 |
| Chymotrypsin | 2 | 174 | 4356 | 1 | 3676 | 4064.7307 | 33.036 | 82,550 | 4063.722875 | 21 | 66 | 79 | HAIHVSGTNGTKRF | 0.00000000200000.0 | 74(NGT) | 1 | 1526.784 | 1523.795739 | 23511576 | 36.61 | 1 | 6 | 7 | 1 | 0 | 0 | 0 | 1 | 4063.726203 | -0.0033 | -0.81 | -3.57 |
| Chymotrypsin | 2 | 174 | 4357 | 1 | 3686 | 3658.5689 | 33.4641 | 82,547 | 3657.561075 | 21 | 66 | 79 | HAIHVSGTNGTKRF | 0.00000000200000.0 | 74(NGT) | 1 | 1526.784 | 1523.795739 | 23511576 | 36.61 | 1 | 6 | 5 | 1 | 0 | 0 | 0 | 1 | 3657.567457 | -0.0064 | -1.74 | -3.14 |
| Chymotrypsin | 2 | 174 | 4366 | 1 | 3728 | 4533.8824 | 35.2515 | 82,234 | 4532.874575 | 21 | 66 | 79 | HAIHVSGTNGTKRF | 0.00000000200000.0 | 74(NGT) | 1 | 1526.784 | 1523.795739 | 23511576 | 36.61 | 1 | 8 | 7 | 0 | 1 | 0 | 0 | 1 | 4532.869359 | 0.0052 | 1.15 | -1.36 |
| Chymotrypsin | 2 | 174 | 4366 | 1 | 3728 | 4533.8824 | 35.2515 | 82,234 | 4532.874575 | 21 | 66 | 79 | HAIHVSGTNGTKRF | 0.00000000200000.0 | 74(NGT) | 1 | 1526.784 | 1523.795739 | 23511576 | 36.61 | 2 | 7 | 7 | 1 | 0 | 1 | 0 | 0 | 4532.869358 | 0.0052 | 1.15 | -1.36 |
| Chymotrypsin | 2 | 174 | 4419 | 1 | 3737 | 4314.7929 | 35.6386 | 80,907 | 4313.785075 | 21 | 66 | 79 | HAIHVSGTNGTKRF | 0.00000000200000.0 | 74(NGT) | 1 | 1526.784 | 1523.795739 | 23511576 | 36.61 | 1 | 7 | 6 | 1 | 1 | 0 | 0 | 1 | 4313.795071 | -0.0100 | -2.31 | -0.97 |
| Chymotrypsin | 2 | 174 | 4419 | 1 | 3737 | 4314.7929 | 35.6386 | 80,907 | 4313.785075 | 21 | 66 | 79 | HAIHVSGTNGTKRF | 0.00000000200000.0 | 74(NGT) | 1 | 1526.784 | 1523.795739 | 23511576 | 36.61 | 2 | 6 | 6 | 2 | 0 | 1 | 0 | 1 | 4313.79507 | -0.0100 | -2.31 | -0.97 |
| Chymotrypsin | 2 | 174 | 4441 | 1 | 3743 | 3641.559 | 35.8951 | 80,409 | 3640.551175 | 21 | 66 | 79 | HAIHVSGTNGTKRF | 0.00000000200000.0 | 74(NGT) | 1 | 1526.784 | 1523.795739 | 23511576 | 36.61 | 1 | 4 | 5 | 1 | 0 | 1 | 0 | 1 | 3640.55214 | -0.0010 | -0.26 | -0.71 |
| Chymotrypsin | 2 | 174 | 4441 | 1 | 3743 | 3641.559 | 35.8951 | 80,409 | 3640.551175 | 21 | 66 | 79 | HAIHVSGTNGTKRF | 0.00000000200000.0 | 74(NGT) | 1 | 1526.784 | 1523.795739 | 23511576 | 36.61 | 2 | 5 | 5 | 0 | 1 | 0 | 0 | 1 | 3640.552141 | -0.0010 | -0.26 | -0.71 |
| Chymotrypsin | 2 | 174 | 4516 | 1 | 3670 | 4397.874 | 32.7835 | 78,680 | 4396.866175 | 21 | 66 | 79 | HAIHVSGTNGTKRF | 0.00000000200000.0 | 74(NGT) | 1 | 1526.784 | 1523.795739 | 23511576 | 36.61 | 1 | 5 | 8 | 3 | 0 | 0 | 0 | 0 | 4396.86857 | -0.0024 | -0.54 | -3.83 |
| Chymotrypsin | 2 | 174 | 4577 | 1 | 3723 | 4330.7916 | 35.04 | 77,349 | 4329.783775 | 21 | 66 | 79 | HAIHVSGTNGTKRF | 0.00000000200000.0 | 74(NGT) | 1 | 1526.784 | 1523.795739 | 23511576 | 36.61 | 1 | 7 | 6 | 1 | 0 | 1 | 0 | 1 | 4329.789985 | -0.0062 | -1.43 | -1.57 |
| Chymotrypsin | 2 | 174 | 4577 | 1 | 3723 | 4330.7916 | 35.04 | 77,349 | 4329.783775 | 21 | 66 | 79 | HAIHVSGTNGTKRF | 0.00000000200000.0 | 74(NGT) | 1 | 1526.784 | 1523.795739 | 23511576 | 36.61 | 2 | 8 | 6 | 0 | 1 | 0 | 0 | 1 | 4329.789986 | -0.0062 | -1.43 | -1.57 |
| Chymotrypsin | 2 | 174 | 4628 | 1 | 3730 | 3682.5851 | 35.3364 | 76,479 | 3681.577275 | 21 | 66 | 79 | HAIHVSGTNGTKRF | 0.00000000200000.0 | 74(NGT) | 1 | 1526.784 | 1523.795739 | 23511576 | 36.61 | 1 | 3 | 6 | 1 | 0 | 1 | 0 | 1 | 3681.578689 | -0.0014 | -0.38 | -1.27 |
| Chymotrypsin | 2 | 174 | 4628 | 1 | 3730 | 3682.5851 | 35.3364 | 76,479 | 3681.577275 | 21 | 66 | 79 | HAIHVSGTNGTKRF | 0.00000000200000.0 | 74(NGT) | 1 | 1526.784 | 1523.795739 | 23511576 | 36.61 | 2 | 4 | 6 | 0 | 1 | 0 | 0 | 1 | 3681.57869 | -0.0014 | -0.38 | -1.27 |
| Chymotrypsin | 2 | 174 | 4694 | 1 | 3810 | 3713.5754 | 38.7636 | 74,866 | 3712.567575 | 21 | 66 | 79 | HAIHVSGTNGTKRF | 0.00000000200000.0 | 74(NGT) | 1 | 1526.784 | 1523.795739 | 23511576 | 36.61 | 1 | 2 | 4 | 3 | 0 | 2 | 1 | -1 | 3712.573268 | -0.0057 | -1.53 | 2.15 |
| Chymotrypsin | 2 | 174 | 4694 | 1 | 3810 | 3713.5754 | 38.7636 | 74,866 | 3712.567575 | 21 | 66 | 79 | HAIHVSGTNGTKRF | 0.00000000200000.0 | 74(NGT) | 1 | 1526.784 | 1523.795739 | 23511576 | 36.61 | 2 | 3 | 4 | 2 | 1 | 1 | 1 | -1 | 3712.573269 | -0.0057 | -1.53 | 2.15 |
| Chymotrypsin | 2 | 174 | 4694 | 1 | 3810 | 3713.5754 | 38.7636 | 74,866 | 3712.567575 | 21 | 66 | 79 | HAIHVSGTNGTKRF | 0.00000000200000.0 | 74(NGT) | 1 | 1526.784 | 1523.795739 | 23511576 | 36.61 | 3 | 4 | 4 | 1 | 2 | 0 | 0 | 0 | 3712.57327 | -0.0057 | -1.53 | 2.15 |
| Chymotrypsin | 2 | 174 | 4731 | 1 | 3673 | 4210.7925 | 32.9085 | 74,082 | 4209.784675 | 21 | 66 | 79 | HAIHVSGTNGTKRF | 0.00000000200000.0 | 74(NGT) | 1 | 1526.784 | 1523.795739 | 23511576 | 36.61 | 1 | 6 | 7 | 2 | 0 | 0 | 0 | 0 | 4209.784112 | 0.0006 | 0.13 | -3.70 |
| Chymotrypsin | 2 | 174 | 4845 | 1 | 3786 | 3948.6542 | 37.7367 | 71,844 | 3947.646375 | 21 | 66 | 79 | HAIHVSGTNGTKRF | 0.00000000200000.0 | 74(NGT) | 1 | 1526.784 | 1523.795739 | 23511576 | 36.61 | 1 | 5 | 5 | 0 | 1 | 1 | 0 | 1 | 3947.642472 | 0.0039 | 0.98 | 1.13 |
| Chymotrypsin | 2 | 174 | 4845 | 1 | 3786 | 3948.6542 | 37.7367 | 71,844 | 3947.646375 | 21 | 66 | 79 | HAIHVSGTNGTKRF | 0.00000000200000.0 | 74(NGT) | 1 | 1526.784 | 1523.795739 | 23511576 | 36.61 | 2 | 4 | 5 | 1 | 0 | 2 | 0 | 1 | 3947.642471 | 0.0039 | 0.98 | 1.13 |
| Chymotrypsin | 2 | 174 | 5083 | 1 | 3673 | 3398.4735 | 32.9085 | 66,947 | 3397.465675 | 21 | 66 | 79 | HAIHVSGTNGTKRF | 0.00000000200000.0 | 74(NGT) | 1 | 1526.784 | 1523.795739 | 23511576 | 36.61 | 1 | 6 | 3 | 2 | 0 | 0 | 0 | 0 | 3397.46662 | -0.0009 | -0.27 | -3.70 |
| Chymotrypsin | 2 | 174 | 5111 | 1 | 3653 | 4973.0701 | 32.0502 | 66,506 | 4972.062275 | 21 | 66 | 79 | HAIHVSGTNGTKRF | 0.00000000200000.0 | 74(NGT) | 1 | 1526.784 | 1523.795739 | 23511576 | 36.61 | 1 | 10 | 9 | 0 | 0 | 0 | 0 | 1 | 4972.038336 | 0.0239 | 4.81 | -4.56 |
| Chymotrypsin | 2 | 174 | 5140 | 1 | 3649 | 5119.1253 | 31.8795 | 66,043 | 5118.117475 | 21 | 66 | 79 | HAIHVSGTNGTKRF | 0.00000000200000.0 | 74(NGT) | 1 | 1526.784 | 1523.795739 | 23511576 | 36.61 | 1 | 10 | 9 | 1 | 0 | 0 | 0 | 1 | 5118.096245 | 0.0212 | 4.14 | -4.73 |
| Chymotrypsin | 2 | 174 | 5157 | 1 | 3728 | 3730.5976 | 35.2515 | 65,770 | 3729.589775 | 21 | 66 | 79 | HAIHVSGTNGTKRF | 0.00000000200000.0 | 74(NGT) | 1 | 1526.784 | 1523.795739 | 23511576 | 36.61 | 1 | 5 | 4 | 2 | 1 | 0 | 0 | 1 | 3729.588586 | 0.0012 | 0.31 | -1.36 |
| Chymotrypsin | 2 | 174 | 5157 | 1 | 3728 | 3730.5976 | 35.2515 | 65,770 | 3729.589775 | 21 | 66 | 79 | HAIHVSGTNGTKRF | 0.00000000200000.0 | 74(NGT) | 1 | 1526.784 | 1523.795739 | 23511576 | 36.61 | 2 | 4 | 4 | 3 | 0 | 1 | 0 | 0 | 3729.588585 | 0.0012 | 0.31 | -1.36 |
| Chymotrypsin | 2 | 174 | 5195 | 1 | 3770 | 4637.8885 | 37.0538 | 65,186 | 4636.880675 | 21 | 66 | 79 | HAIHVSGTNGTKRF | 0.00000000200000.0 | 74(NGT) | 1 | 1526.784 | 1523.795739 | 23511576 | 36.61 | 1 | 8 | 6 | 0 | 1 | 1 | 0 | 0 | 4636.880317 | 0.0004 | 0.07 | 0.44 |
| Chymotrypsin | 2 | 174 | 5195 | 1 | 3770 | 4637.8885 | 37.0538 | 65,186 | 4636.880675 | 21 | 66 | 79 | HAIHVSGTNGTKRF | 0.00000000200000.0 | 74(NGT) | 1 | 1526.784 | 1523.795739 | 23511576 | 36.61 | 2 | 7 | 6 | 1 | 0 | 2 | 0 | 0 | 4636.880316 | 0.0004 | 0.07 | 0.44 |
| Chymotrypsin | 2 | 174 | 5273 | 1 | 3677 | 3861.6487 | 33.0797 | 64,074 | 3860.640875 | 21 | 66 | 79 | HAIHVSGTNGTKRF | 0.00000000200000.0 | 74(NGT) | 1 | 1526.784 | 1523.795739 | 23511576 | 36.61 | 1 | 6 | 6 | 1 | 0 | 0 | 0 | 1 | 3860.64683 | -0.0060 | -1.54 | -3.53 |
| Chymotrypsin | 2 | 174 | 5312 | 1 | 3652 | 5176.1527 | 32.0073 | 63,374 | 5175.144875 | 21 | 66 | 79 | HAIHVSGTNGTKRF | 0.00000000200000.0 | 74(NGT) | 1 | 1526.784 | 1523.795739 | 23511576 | 36.61 |  |  |  |  |  |  |  |  |  |  |  |  |
| Chymotrypsin | 2 | 174 | 5429 | 1 | 3672 | 3845.6603 | 32.8662 | 61,292 | 3844.652475 | 21 | 66 | 79 | HAIHVSGTNGTKRF | 0.00000000200000.0 | 74(NGT) | 1 | 1526.784 | 1523.795739 | 23511576 | 36.61 | 1 | 5 | 6 | 2 | 0 | 0 | 0 | 1 | 3844.651915 | 0.0006 | 0.14 | -3.74 |
| Chymotrypsin | 2 | 174 | 5481 | 1 | 3766 | 3422.4772 | 36.8797 | 60,396 | 3421.469375 | 21 | 66 | 79 | HAIHVSGTNGTKRF | 0.00000000200000.0 | 74(NGT) | 1 | 1526.784 | 1523.795739 | 23511576 | 36.61 | 1 | 4 | 4 | 1 | 1 | 0 | 0 | 1 | 3421.477853 | -0.0085 | -2.47 | 0.27 |
| Chymotrypsin | 2 | 174 | 5481 | 1 | 3766 | 3422.4772 | 36.8797 | 60,396 | 3421.469375 | 21 | 66 | 79 | HAIHVSGTNGTKRF | 0.00000000200000.0 | 74(NGT) | 1 | 1526.784 | 1523.795739 | 23511576 | 36.61 | 2 | 3 | 4 | 2 | 0 | 1 | 0 | 0 | 3421.477852 | -0.0085 | -2.47 | 0.27 |
| Chymotrypsin | 2 | 174 | 5494 | 1 | 3684 | 3537.5431 | 33.3755 | 60,239 | 3536.535275 | 21 | 66 | 79 | HAIHVSGTNGTKRF | 0.00000000200000.0 | 74(NGT) | 1 | 1526.784 | 1523.795739 | 23511576 | 36.61 | 1 | 4 | 6 | 1 | 0 | 0 | 0 | 1 | 3536.541182 | -0.0059 | -1.67 | -3.23 |
| Chymotrypsin | 2 | 174 | 5589 | 1 | 3678 | 3699.5986 | 33.1224 | 58,730 | 3698.590775 | 21 | 66 | 79 | HAIHVSGTNGTKRF | 0.00000000200000.0 | 74(NGT) | 1 | 1526.784 | 1523.795739 | 23511576 | 36.61 | 1 | 5 | 6 | 1 | 0 | 0 | 0 | 1 | 3698.594006 | -0.0032 | -0.87 | -3.49 |
| Chymotrypsin | 2 | 174 | 5697 | 1 | 3735 | 3277.434 | 35.5533 | 56,907 | 3276.426175 | 21 | 66 | 79 | HAIHVSGTNGTKRF | 0.00000000200000.0 | 74(NGT) | 1 | 1526.784 | 1523.795739 | 23511576 | 36.61 | 1 | 4 | 4 | 2 | 0 | 0 | 0 | 1 | 3276.440345 | -0.0142 | -4.32 | -1.06 |
| Chymotrypsin | 2 | 174 | 5746 | 1 | 3660 | 4753.9964 | 32.3486 | 55,826 | 4752.988575 | 21 | 66 | 79 | HAIHVSGTNGTKRF | 0.00000000200000.0 | 74(NGT) | 1 | 1526.784 | 1523.795739 | 23511576 | 36.61 | 1 | 2 | 10 | 2 | 2 | 0 | 1 | -1 | 4753.001769 | -0.0132 | -2.77 | -4.26 |
| Chymotrypsin | 2 | 174 | 5769 | 1 | 3720 | 4111.7181 | 34.9149 | 55,439 | 4110.710275 | 21 | 66 | 79 | HAIHVSGTNGTKRF | 0.00000000200000.0 | 74(NGT) | 1 | 1526.784 | 1523.795739 | 23511576 | 36.61 | 1 | 6 | 5 | 2 | 0 | 1 | 0 | 1 | 4110.715697 | -0.0054 | -1.31 | -1.69 |
| Chymotrypsin | 2 | 174 | 5769 | 1 | 3720 | 4111.7181 | 34.9149 | 55,439 | 4110.710275 | 21 | 66 | 79 | HAIHVSGTNGTKRF | 0.00000000200000.0 | 74(NGT) | 1 | 1526.784 | 1523.795739 | 23511576 | 36.61 | 2 | 7 | 5 | 1 | 1 | 0 | 0 | 1 | 4110.715698 | -0.0054 | -1.31 | -1.69 |
| Chymotrypsin | 2 | 174 | 5885 | 1 | 3781 | 3867.6139 | 37.5237 | 53,652 | 3866.606075 | 21 | 66 | 79 | HAIHVSGTNGTKRF | 0.00000000200000.0 | 74(NGT) | 1 | 1526.784 | 1523.795739 | 23511576 | 36.61 | 1 | 7 | 3 | 2 | 0 | 1 | 1 | -1 | 3866.609775 | -0.0037 | -0.95 | 0.91 |
| Chymotrypsin | 2 | 174 | 5885 | 1 | 3781 | 3867.6139 | 37.5237 | 53,652 | 3866.606075 | 21 | 66 | 79 | HAIHVSGTNGTKRF | 0.00000000200000.0 | 74(NGT) | 1 | 1526.784 | 1523.795739 | 23511576 | 36.61 | 2 | 8 | 3 | 1 | 1 | 0 | 1 | -1 | 3866.609776 | -0.0037 | -0.95 | 0.91 |
| Chymotrypsin | 2 | 174 | 5902 | 1 | 3714 | 4622.9121 | 34.6605 | 53,424 | 4621.904275 | 21 | 66 | 79 | HAIHVSGTNGTKRF | 0.00000000200000.0 | 74(NGT) | 1 | 1526.784 | 1523.795739 | 23511576 | 36.61 | 1 | 7 | 6 | 3 | 0 | 1 | 0 | 0 | 4621.905803 | -0.0015 | -0.33 | -1.95 |
| Chymotrypsin | 2 | 174 | 5902 | 1 | 3714 | 4622.9121 | 34.6605 | 53,424 | 4621.904275 | 21 | 66 | 79 | HAIHVSGTNGTKRF | 0.00000000200000.0 | 74(NGT) | 1 | 1526.784 | 1523.795739 | 23511576 | 36.61 | 2 | 8 | 6 | 2 | 1 | 0 | 0 | 0 | 4621.905804 | -0.0015 | -0.33 | -1.95 |
| Chymotrypsin | 2 | 174 | 5997 | 1 | 3727 | 4444.8607 | 35.2098 | 52,039 | 4443.852875 | 21 | 66 | 79 | HAIHVSGTNGTKRF | 0.00000000200000.0 | 74(NGT) | 1 | 1526.784 | 1523.795739 | 23511576 | 36.61 | 1 | 5 | 6 | 4 | 0 | 1 | 0 | 0 | 4443.858064 | -0.0052 | -1.16 | -1.40 |
| Chymotrypsin | 2 | 174 | 5997 | 1 | 3727 | 4444.8607 | 35.2098 | 52,039 | 4443.852875 | 21 | 66 | 79 | HAIHVSGTNGTKRF | 0.00000000200000.0 | 74(NGT) | 1 | 1526.784 | 1523.795739 | 23511576 | 36.61 | 2 | 6 | 6 | 3 | 1 | 0 | 0 | 0 | 4443.858065 | -0.0052 | -1.16 | -1.40 |
| Chymotrypsin | 2 | 174 | 6014 | 1 | 3745 | 3771.62 | 35.9834 | 51,783 | 3770.612175 | 21 | 66 | 79 | HAIHVSGTNGTKRF | 0.00000000200000.0 | 74(NGT) | 1 | 1526.784 | 1523.795739 | 23511576 | 36.61 | 1 | 4 | 5 | 2 | 1 | 0 | 0 | 1 | 3770.615135 | -0.0030 | -0.78 | -0.63 |

| | | | | | | | | | | | | | | | | | | | | | | | | | | | | | | | | |
|---|---|---|---|---|---|---|---|---|---|---|---|---|---|---|---|---|---|---|---|---|---|---|---|---|---|---|---|---|---|---|---|---|
| Chymotrypsin | 2 | 174 | 6014 | 1 | 3745 | 3771.62 | 35.9834 | 51,783 | 3770.612175 | 21 | 66 | 79 | HAIHVSGTNGTKRF | 0.00000000200000.0 | 74(NGT) | 1 | 1526.784 | 1523.795739 | 23511576 | 36.61 | 2 | 3 | 5 | 3 | 0 | 1 | 0 | 0 | 3770.615134 | -0.0030 | -0.78 | -0.63 |
| Chymotrypsin | 2 | 174 | 6131 | 1 | 3777 | 4621.8963 | 37.3532 | 50,212 | 4620.888475 | 21 | 66 | 79 | HAIHVSGTNGTKRF | 0.00000000200000.0 | 74(NGT) | 1 | 1526.784 | 1523.795739 | 23511576 | 36.61 | 1 | 8 | 6 | 0 | 2 | 0 | 0 | 0 | 4620.885403 | 0.0031 | 0.66 | 0.74 |
| Chymotrypsin | 2 | 174 | 6131 | 1 | 3777 | 4621.8963 | 37.3532 | 50,212 | 4620.888475 | 21 | 66 | 79 | HAIHVSGTNGTKRF | 0.00000000200000.0 | 74(NGT) | 1 | 1526.784 | 1523.795739 | 23511576 | 36.61 | 2 | 7 | 6 | 1 | 1 | 1 | 0 | 0 | 4620.885402 | 0.0031 | 0.66 | 0.74 |
| Chymotrypsin | 2 | 174 | 6131 | 1 | 3777 | 4621.8963 | 37.3532 | 50,212 | 4620.888475 | 21 | 66 | 79 | HAIHVSGTNGTKRF | 0.00000000200000.0 | 74(NGT) | 1 | 1526.784 | 1523.795739 | 23511576 | 36.61 | 3 | 6 | 6 | 2 | 0 | 2 | 0 | 0 | 4620.885401 | 0.0031 | 0.66 | 0.74 |
| Chymotrypsin | 2 | 174 | 6133 | 1 | 3767 | 4783.946 | 36.9242 | 50,206 | 4782.938175 | 21 | 66 | 79 | HAIHVSGTNGTKRF | 0.00000000200000.0 | 74(NGT) | 1 | 1526.784 | 1523.795739 | 23511576 | 36.61 | 1 | 7 | 6 | 2 | 0 | 2 | 0 | 0 | 4782.938225 | 0.0000 | -0.01 | 0.32 |
| Chymotrypsin | 2 | 174 | 6133 | 1 | 3767 | 4783.946 | 36.9242 | 50,206 | 4782.938175 | 21 | 66 | 79 | HAIHVSGTNGTKRF | 0.00000000200000.0 | 74(NGT) | 1 | 1526.784 | 1523.795739 | 23511576 | 36.61 | 2 | 8 | 6 | 1 | 1 | 1 | 0 | 0 | 4782.938226 | -0.0001 | -0.01 | 0.32 |
| Chymotrypsin | 2 | 174 | 6133 | 1 | 3767 | 4783.946 | 36.9242 | 50,206 | 4782.938175 | 21 | 66 | 79 | HAIHVSGTNGTKRF | 0.00000000200000.0 | 74(NGT) | 1 | 1526.784 | 1523.795739 | 23511576 | 36.61 | 3 | 9 | 6 | 0 | 2 | 0 | 0 | 0 | 4782.938227 | -0.0001 | -0.01 | 0.32 |
| Chymotrypsin | 2 | 174 | 6188 | 1 | 3707 | 4915.027 | 34.3624 | 49,370 | 4914.019175 | 21 | 66 | 79 | HAIHVSGTNGTKRF | 0.00000000200000.0 | 74(NGT) | 1 | 1526.784 | 1523.795739 | 23511576 | 36.61 | 1 | 8 | 6 | 4 | 1 | 0 | 0 | 0 | 4914.021622 | -0.0024 | -0.49 | -2.25 |
| Chymotrypsin | 2 | 174 | 6188 | 1 | 3707 | 4915.027 | 34.3624 | 49,370 | 4914.019175 | 21 | 66 | 79 | HAIHVSGTNGTKRF | 0.00000000200000.0 | 74(NGT) | 1 | 1526.784 | 1523.795739 | 23511576 | 36.61 | 2 | 2 | 10 | 1 | 2 | 1 | 1 | -1 | 4914.034191 | -0.0150 | -3.05 | -2.25 |
| Chymotrypsin | 2 | 174 | 6188 | 1 | 3707 | 4915.027 | 34.3624 | 49,370 | 4914.019175 | 21 | 66 | 79 | HAIHVSGTNGTKRF | 0.00000000200000.0 | 74(NGT) | 1 | 1526.784 | 1523.795739 | 23511576 | 36.61 | 3 | 3 | 10 | 0 | 3 | 0 | 0 | 0 | 4914.034192 | -0.0150 | -3.05 | -2.25 |
| Chymotrypsin | 2 | 174 | 6188 | 1 | 3707 | 4915.027 | 34.3624 | 49,370 | 4914.019175 | 21 | 66 | 79 | HAIHVSGTNGTKRF | 0.00000000200000.0 | 74(NGT) | 1 | 1526.784 | 1523.795739 | 23511576 | 36.61 | 4 | 9 | 8 | 0 | 0 | 1 | 0 | 0 | 4913.99647 | 0.0227 | 4.62 | -2.25 |
| Chymotrypsin | 2 | 174 | 6223 | 1 | 3648 | 5322.203 | 31.8378 | 48,860 | 5321.195175 | 21 | 66 | 79 | HAIHVSGTNGTKRF | 0.00000000200000.0 | 74(NGT) | 1 | 1526.784 | 1523.795739 | 23511576 | 36.61 | 1 | 10 | 10 | 1 | 0 | 0 | 0 | 1 | 5321.175618 | 0.0196 | 3.67 | -4.77 |
| Chymotrypsin | 2 | 174 | 6419 | 1 | 3781 | 4110.7018 | 37.5237 | 46,274 | 4109.693975 | 21 | 66 | 79 | HAIHVSGTNGTKRF | 0.00000000200000.0 | 74(NGT) | 1 | 1526.784 | 1523.795739 | 23511576 | 36.61 | 1 | 5 | 5 | 1 | 0 | 2 | 0 | 1 | 4109.695295 | -0.0013 | -0.32 | 0.91 |
| Chymotrypsin | 2 | 174 | 6419 | 1 | 3781 | 4110.7018 | 37.5237 | 46,274 | 4109.693975 | 21 | 66 | 79 | HAIHVSGTNGTKRF | 0.00000000200000.0 | 74(NGT) | 1 | 1526.784 | 1523.795739 | 23511576 | 36.61 | 2 | 6 | 5 | 0 | 1 | 1 | 0 | 1 | 4109.695296 | -0.0013 | -0.32 | 0.91 |
| Chymotrypsin | 2 | 174 | 6531 | 1 | 3675 | 3747.6098 | 32.9948 | 44,747 | 3746.601975 | 21 | 66 | 79 | HAIHVSGTNGTKRF | 0.00000000200000.0 | 74(NGT) | 1 | 1526.784 | 1523.795739 | 23511576 | 36.61 | 1 | 6 | 4 | 3 | 0 | 0 | 0 | 1 | 3746.603902 | -0.0019 | -0.51 | -3.61 |
| Chymotrypsin | 2 | 174 | 6547 | 1 | 3668 | 3804.6386 | 32.6951 | 44,547 | 3803.630775 | 21 | 66 | 79 | HAIHVSGTNGTKRF | 0.00000000200000.0 | 74(NGT) | 1 | 1526.784 | 1523.795739 | 23511576 | 36.61 | 1 | 6 | 5 | 2 | 0 | 0 | 0 | 1 | 3803.625366 | 0.0054 | 1.42 | -3.91 |
| Chymotrypsin | 2 | 174 | 6587 | 1 | 3753 | 3495.4877 | 36.3243 | 43,910 | 3494.479875 | 21 | 66 | 79 | HAIHVSGTNGTKRF | 0.00000000200000.0 | 74(NGT) | 1 | 1526.784 | 1523.795739 | 23511576 | 36.61 | 1 | 4 | 5 | 0 | 0 | 1 | 0 | 1 | 3494.494231 | -0.0144 | -4.1 | -0.28 |
| Chymotrypsin | 2 | 174 | 6598 | 1 | 3719 | 4606.9371 | 34.8717 | 43,816 | 4605.929275 | 21 | 66 | 79 | HAIHVSGTNGTKRF | 0.00000000200000.0 | 74(NGT) | 1 | 1526.784 | 1523.795739 | 23511576 | 36.61 | 1 | 7 | 6 | 3 | 1 | 0 | 0 | 0 | 4605.910889 | 0.0184 | 3.99 | -1.74 |
| Chymotrypsin | 2 | 174 | 6598 | 1 | 3719 | 4606.9371 | 34.8717 | 43,816 | 4605.929275 | 21 | 66 | 79 | HAIHVSGTNGTKRF | 0.00000000200000.0 | 74(NGT) | 1 | 1526.784 | 1523.795739 | 23511576 | 36.61 | 2 | 6 | 6 | 4 | 0 | 1 | 0 | 0 | 4605.910888 | 0.0184 | 3.99 | -1.74 |
| Chymotrypsin | 2 | 174 | 6635 | 1 | 3726 | 4371.8275 | 35.1674 | 43,283 | 4370.819675 | 21 | 66 | 79 | HAIHVSGTNGTKRF | 0.00000000200000.0 | 74(NGT) | 1 | 1526.784 | 1523.795739 | 23511576 | 36.61 | 1 | 7 | 7 | 0 | 1 | 0 | 0 | 0 | 4370.816535 | 0.0031 | 0.71 | -1.44 |
| Chymotrypsin | 2 | 174 | 6635 | 1 | 3726 | 4371.8275 | 35.1674 | 43,283 | 4370.819675 | 21 | 66 | 79 | HAIHVSGTNGTKRF | 0.00000000200000.0 | 74(NGT) | 1 | 1526.784 | 1523.795739 | 23511576 | 36.61 | 2 | 6 | 7 | 1 | 0 | 1 | 0 | 0 | 4370.816534 | 0.0031 | 0.71 | -1.44 |
| Chymotrypsin | 2 | 174 | 6663 | 1 | 3727 | 4152.7557 | 35.2098 | 42,928 | 4151.747875 | 21 | 66 | 79 | HAIHVSGTNGTKRF | 0.00000000200000.0 | 74(NGT) | 1 | 1526.784 | 1523.795739 | 23511576 | 36.61 | 1 | 6 | 6 | 1 | 1 | 0 | 0 | 1 | 4151.742247 | 0.0056 | 1.35 | -1.40 |
| Chymotrypsin | 2 | 174 | 6663 | 1 | 3727 | 4152.7557 | 35.2098 | 42,928 | 4151.747875 | 21 | 66 | 79 | HAIHVSGTNGTKRF | 0.00000000200000.0 | 74(NGT) | 1 | 1526.784 | 1523.795739 | 23511576 | 36.61 | 2 | 5 | 6 | 2 | 0 | 1 | 0 | 0 | 4151.742246 | 0.0056 | 1.35 | -1.40 |
| Chymotrypsin | 2 | 174 | 6692 | 1 | 3723 | 4241.777 | 35.04 | 42,565 | 4240.769175 | 21 | 66 | 79 | HAIHVSGTNGTKRF | 0.00000000200000.0 | 74(NGT) | 1 | 1526.784 | 1523.795739 | 23511576 | 36.61 | 1 | 6 | 5 | 3 | 1 | 0 | 0 | 1 | 4240.778692 | -0.0095 | -2.24 | -1.57 |
| Chymotrypsin | 2 | 174 | 6692 | 1 | 3723 | 4241.777 | 35.04 | 42,565 | 4240.769175 | 21 | 66 | 79 | HAIHVSGTNGTKRF | 0.00000000200000.0 | 74(NGT) | 1 | 1526.784 | 1523.795739 | 23511576 | 36.61 | 2 | 5 | 5 | 4 | 0 | 1 | 0 | 0 | 4240.778691 | -0.0095 | -2.24 | -1.57 |
| Chymotrypsin | 2 | 174 | 6768 | 1 | 3675 | 4251.8173 | 32.9948 | 41,486 | 4250.809475 | 21 | 66 | 79 | HAIHVSGTNGTKRF | 0.00000000200000.0 | 74(NGT) | 1 | 1526.784 | 1523.795739 | 23511576 | 36.61 | 1 | 5 | 8 | 2 | 0 | 0 | 0 | 0 | 4250.810661 | -0.0012 | -0.27 | -3.61 |
| Chymotrypsin | 2 | 174 | 6773 | 1 | 3712 | 4956.052 | 34.5742 | 41,370 | 4955.044175 | 21 | 66 | 79 | HAIHVSGTNGTKRF | 0.00000000200000.0 | 74(NGT) | 1 | 1526.784 | 1523.795739 | 23511576 | 36.61 | 1 | 7 | 7 | 4 | 1 | 0 | 0 | 0 | 4955.048171 | -0.0040 | -0.8 | -2.03 |
| Chymotrypsin | 2 | 174 | 6773 | 1 | 3712 | 4956.052 | 34.5742 | 41,370 | 4955.044175 | 21 | 66 | 79 | HAIHVSGTNGTKRF | 0.00000000200000.0 | 74(NGT) | 1 | 1526.784 | 1523.795739 | 23511576 | 36.61 | 2 | 8 | 9 | 0 | 0 | 1 | 0 | 0 | 4955.023019 | 0.0212 | 4.26 | -2.03 |
| Chymotrypsin | 2 | 174 | 6804 | 1 | 3855 | 3348.4313 | 40.7011 | 40,853 | 3347.423475 | 21 | 66 | 79 | HAIHVSGTNGTKRF | 0.00000000200000.0 | 74(NGT) | 1 | 1526.784 | 1523.795739 | 23511576 | 36.61 | | | | | | | | | | | | |
| Chymotrypsin | 2 | 174 | 6810 | 1 | 3687 | 3886.6891 | 33.5058 | 40,775 | 3885.681275 | 21 | 66 | 79 | HAIHVSGTNGTKRF | 0.00000000200000.0 | 74(NGT) | 1 | 1526.784 | 1523.795739 | 23511576 | 36.61 | 1 | 4 | 7 | 2 | 0 | 0 | 0 | 1 | 3885.678464 | 0.0028 | 0.72 | -3.10 |
| Chymotrypsin | 2 | 174 | 6815 | 1 | 3728 | 3771.6263 | 35.2515 | 40,735 | 3770.618475 | 21 | 66 | 79 | HAIHVSGTNGTKRF | 0.00000000200000.0 | 74(NGT) | 1 | 1526.784 | 1523.795739 | 23511576 | 36.61 | 1 | 4 | 5 | 2 | 1 | 0 | 0 | 1 | 3770.615135 | 0.0033 | 0.88 | -1.36 |
| Chymotrypsin | 2 | 174 | 6815 | 1 | 3728 | 3771.6263 | 35.2515 | 40,735 | 3770.618475 | 21 | 66 | 79 | HAIHVSGTNGTKRF | 0.00000000200000.0 | 74(NGT) | 1 | 1526.784 | 1523.795739 | 23511576 | 36.61 | 2 | 3 | 5 | 3 | 0 | 1 | 0 | 0 | 3770.615134 | 0.0033 | 0.88 | -1.36 |
| Chymotrypsin | 2 | 174 | 6844 | 1 | 3718 | 3892.6483 | 34.8306 | 40,336 | 3891.640475 | 21 | 66 | 79 | HAIHVSGTNGTKRF | 0.00000000200000.0 | 74(NGT) | 1 | 1526.784 | 1523.795739 | 23511576 | 36.61 | 1 | 5 | 4 | 3 | 0 | 1 | 0 | 0 | 3891.641409 | -0.0009 | -0.23 | -1.78 |
| Chymotrypsin | 2 | 174 | 6844 | 1 | 3718 | 3892.6483 | 34.8306 | 40,336 | 3891.640475 | 21 | 66 | 79 | HAIHVSGTNGTKRF | 0.00000000200000.0 | 74(NGT) | 1 | 1526.784 | 1523.795739 | 23511576 | 36.61 | 2 | 6 | 4 | 2 | 1 | 0 | 0 | 0 | 3891.64141 | -0.0009 | -0.24 | -1.78 |
| Chymotrypsin | 2 | 174 | 6916 | 1 | 3665 | 4559.9327 | 32.5693 | 39,402 | 4558.924875 | 21 | 66 | 79 | HAIHVSGTNGTKRF | 0.00000000200000.0 | 74(NGT) | 1 | 1526.784 | 1523.795739 | 23511576 | 36.61 | 1 | 6 | 8 | 3 | 0 | 0 | 0 | 0 | 4558.921394 | 0.0035 | 0.76 | -4.04 |
| Chymotrypsin | 2 | 174 | 6922 | 1 | 3689 | 4194.7949 | 33.5921 | 39,313 | 4193.787075 | 21 | 66 | 79 | HAIHVSGTNGTKRF | 0.00000000200000.0 | 74(NGT) | 1 | 1526.784 | 1523.795739 | 23511576 | 36.61 | 1 | 5 | 7 | 3 | 0 | 0 | 0 | 0 | 4193.789197 | -0.0021 | -0.5 | -3.02 |
| Chymotrypsin | 2 | 174 | 6986 | 1 | 3667 | 4153.7796 | 32.6524 | 38,242 | 4152.771775 | 21 | 66 | 79 | HAIHVSGTNGTKRF | 0.00000000200000.0 | 74(NGT) | 1 | 1526.784 | 1523.795739 | 23511576 | 36.61 | 1 | 6 | 6 | 3 | 0 | 0 | 0 | 1 | 4152.762648 | 0.0091 | 2.19 | -3.96 |
| Chymotrypsin | 2 | 174 | 7105 | 1 | 3733 | 4298.8131 | 35.4682 | 36,459 | 4297.805275 | 21 | 66 | 79 | HAIHVSGTNGTKRF | 0.00000000200000.0 | 74(NGT) | 1 | 1526.784 | 1523.795739 | 23511576 | 36.61 | 1 | 6 | 6 | 2 | 1 | 0 | 0 | 0 | 4297.800156 | 0.0051 | 1.19 | -1.14 |
| Chymotrypsin | 2 | 174 | 7105 | 1 | 3733 | 4298.8131 | 35.4682 | 36,459 | 4297.805275 | 21 | 66 | 79 | HAIHVSGTNGTKRF | 0.00000000200000.0 | 74(NGT) | 1 | 1526.784 | 1523.795739 | 23511576 | 36.61 | 2 | 5 | 6 | 3 | 0 | 1 | 0 | 0 | 4297.800155 | 0.0051 | 1.19 | -1.14 |
| Chymotrypsin | 2 | 174 | 7121 | 1 | 3719 | 4501.8825 | 34.8717 | 36,334 | 4500.874675 | 21 | 66 | 79 | HAIHVSGTNGTKRF | 0.00000000200000.0 | 74(NGT) | 1 | 1526.784 | 1523.795739 | 23511576 | 36.61 | 1 | 5 | 7 | 3 | 0 | 1 | 0 | 0 | 4500.879528 | -0.0049 | -1.07 | -1.74 |
| Chymotrypsin | 2 | 174 | 7121 | 1 | 3719 | 4501.8825 | 34.8717 | 36,334 | 4500.874675 | 21 | 66 | 79 | HAIHVSGTNGTKRF | 0.00000000200000.0 | 74(NGT) | 1 | 1526.784 | 1523.795739 | 23511576 | 36.61 | 2 | 6 | 7 | 2 | 1 | 0 | 0 | 0 | 4500.879529 | -0.0049 | -1.07 | -1.74 |
| Chymotrypsin | 2 | 174 | 7178 | 1 | 3778 | 4272.7575 | 37.3948 | 35,494 | 4271.749675 | 21 | 66 | 79 | HAIHVSGTNGTKRF | 0.00000000200000.0 | 74(NGT) | 1 | 1526.784 | 1523.795739 | 23511576 | 36.61 | 1 | 6 | 5 | 1 | 0 | 2 | 0 | 1 | 4271.748119 | 0.0016 | 0.36 | 0.79 |
| Chymotrypsin | 2 | 174 | 7178 | 1 | 3778 | 4272.7575 | 37.3948 | 35,494 | 4271.749675 | 21 | 66 | 79 | HAIHVSGTNGTKRF | 0.00000000200000.0 | 74(NGT) | 1 | 1526.784 | 1523.795739 | 23511576 | 36.61 | 2 | 7 | 5 | 0 | 1 | 1 | 0 | 0 | 4271.74812 | 0.0016 | 0.36 | 0.79 |
| Chymotrypsin | 2 | 174 | 7190 | 1 | 3714 | 4298.809 | 34.6605 | 35,441 | 4297.801175 | 21 | 66 | 79 | HAIHVSGTNGTKRF | 0.00000000200000.0 | 74(NGT) | 1 | 1526.784 | 1523.795739 | 23511576 | 36.61 | 1 | 6 | 6 | 2 | 1 | 0 | 0 | 0 | 4297.800156 | 0.0010 | 0.23 | -1.95 |
| Chymotrypsin | 2 | 174 | 7190 | 1 | 3714 | 4298.809 | 34.6605 | 35,441 | 4297.801175 | 21 | 66 | 79 | HAIHVSGTNGTKRF | 0.00000000200000.0 | 74(NGT) | 1 | 1526.784 | 1523.795739 | 23511576 | 36.61 | 2 | 5 | 6 | 3 | 0 | 1 | 0 | 0 | 4297.800155 | 0.0010 | 0.23 | -1.95 |
| Chymotrypsin | 2 | 174 | 7275 | 1 | 3688 | 3683.6076 | 33.5492 | 34,383 | 3682.599775 | 21 | 66 | 79 | HAIHVSGTNGTKRF | 0.00000000200000.0 | 74(NGT) | 1 | 1526.784 | 1523.795739 | 23511576 | 36.61 | 1 | 4 | 6 | 2 | 0 | 0 | 0 | 1 | 3682.599091 | 0.0007 | 0.18 | -3.06 |
| Chymotrypsin | 2 | 174 | 7282 | 1 | 3673 | 3886.6857 | 32.9085 | 34,287 | 3885.677875 | 21 | 66 | 79 | HAIHVSGTNGTKRF | 0.00000000200000.0 | 74(NGT) | 1 | 1526.784 | 1523.795739 | 23511576 | 36.61 | 1 | 4 | 7 | 2 | 0 | 0 | 0 | 1 | 3885.678464 | -0.0006 | -0.15 | -3.70 |
| Chymotrypsin | 2 | 174 | 7304 | 1 | 3650 | 5160.152 | 31.9215 | 34,087 | 5159.144175 | 21 | 66 | 79 | HAIHVSGTNGTKRF | 0.00000000200000.0 | 74(NGT) | 1 | 1526.784 | 1523.795739 | 23511576 | 36.61 | 1 | 9 | 10 | 1 | 0 | 0 | 0 | 0 | 5159.122794 | 0.0214 | 4.14 | -4.69 |
| Chymotrypsin | 2 | 174 | 7484 | 1 | 3784 | 3907.6176 | 37.6518 | 31,899 | 3906.609775 | 21 | 66 | 79 | HAIHVSGTNGTKRF | 0.00000000200000.0 | 74(NGT) | 1 | 1526.784 | 1523.795739 | 23511576 | 36.61 | 1 | 5 | 4 | 1 | 0 | 2 | 0 | 1 | 3906.615922 | -0.0061 | -1.57 | 1.04 |
| Chymotrypsin | 2 | 174 | 7484 | 1 | 3784 | 3907.6176 | 37.6518 | 31,899 | 3906.609775 | 21 | 66 | 79 | HAIHVSGTNGTKRF | 0.00000000200000.0 | 74(NGT) | 1 | 1526.784 | 1523.795739 | 23511576 | 36.61 | 2 | 6 | 4 | 0 | 1 | 1 | 0 | 0 | 3906.615923 | -0.0061 | -1.57 | 1.04 |
| Chymotrypsin | 2 | 174 | 7525 | 1 | 3680 | 3074.3673 | 33.2082 | 31,293 | 3073.359475 | 21 | 66 | 79 | HAIHVSGTNGTKRF | 0.00000000200000.0 | 74(NGT) | 1 | 1526.784 | 1523.795739 | 23511576 | 36.61 | 1 | 4 | 3 | 2 | 0 | 0 | 0 | 1 | 3073.360972 | -0.0015 | -0.48 | -3.40 |
| Chymotrypsin | 2 | 174 | 7539 | 1 | 3680 | 3804.6339 | 33.2082 | 31,111 | 3803.626075 | 21 | 66 | 79 | HAIHVSGTNGTKRF | 0.00000000200000.0 | 74(NGT) | 1 | 1526.784 | 1523.795739 | 23511576 | 36.61 | 1 | 6 | 5 | 2 | 0 | 0 | 0 | 1 | 3803.625366 | 0.0007 | 0.18 | -3.40 |
| Chymotrypsin | 2 | 174 | 7583 | 1 | 3735 | 3584.5348 | 35.5533 | 30,463 | 3583.526975 | 21 | 66 | 79 | HAIHVSGTNGTKRF | 0.00000000200000.0 | 74(NGT) | 1 | 1526.784 | 1523.795739 | 23511576 | 36.61 | 1 | 4 | 4 | 2 | 0 | 1 | 0 | 1 | 3583.530676 | -0.0037 | -1.03 | -1.06 |
| Chymotrypsin | 2 | 174 | 7583 | 1 | 3735 | 3584.5348 | 35.5533 | 30,463 | 3583.526975 | 21 | 66 | 79 | HAIHVSGTNGTKRF | 0.00000000200000.0 | 74(NGT) | 1 | 1526.784 | 1523.795739 | 23511576 | 36.61 | 2 | 5 | 4 | 1 | 1 | 0 | 0 | 1 | 3583.530677 | -0.0037 | -1.03 | -1.06 |
| Chymotrypsin | 2 | 174 | 7638 | 1 | 3751 | 4152.7526 | 36.2403 | 29,867 | 4151.744775 | 21 | 66 | 79 | HAIHVSGTNGTKRF | 0.00000000200000.0 | 74(NGT) | 1 | 1526.784 | 1523.795739 | 23511576 | 36.61 | 1 | 6 | 6 | 1 | 1 | 0 | 0 | 1 | 4151.742247 | 0.0025 | 0.6 | -0.37 |
| Chymotrypsin | 2 | 174 | 7638 | 1 | 3751 | 4152.7526 | 36.2403 | 29,867 | 4151.744775 | 21 | 66 | 79 | HAIHVSGTNGTKRF | 0.00000000200000.0 | 74(NGT) | 1 | 1526.784 | 1523.795739 | 23511576 | 36.61 | 2 | 5 | 6 | 2 | 0 | 1 | 0 | 0 | 4151.742246 | 0.0025 | 0.6 | -0.37 |
| Chymotrypsin | 2 | 174 | 7654 | 1 | 3742 | 4460.8693 | 35.8511 | 29,702 | 4459.861475 | 21 | 66 | 79 | HAIHVSGTNGTKRF | 0.00000000200000.0 | 74(NGT) | 1 | 1526.784 | 1523.795739 | 23511576 | 36.61 | 1 | 7 | 6 | 2 | 1 | 0 | 0 | 0 | 4459.85298 | 0.0085 | 1.9 | -0.76 |
| Chymotrypsin | 2 | 174 | 7654 | 1 | 3742 | 4460.8693 | 35.8511 | 29,702 | 4459.861475 | 21 | 66 | 79 | HAIHVSGTNGTKRF | 0.00000000200000.0 | 74(NGT) | 1 | 1526.784 | 1523.795739 | 23511576 | 36.61 | 2 | 6 | 6 | 3 | 0 | 1 | 0 | 0 | 4459.852979 | 0.0085 | 1.9 | -0.76 |
| Chymotrypsin | 2 | 174 | 7685 | 1 | 3835 | 4579.8576 | 39.8388 | 29,421 | 4578.849775 | 21 | 66 | 79 | HAIHVSGTNGTKRF | 0.00000000200000.0 | 74(NGT) | 1 | 1526.784 | 1523.795739 | 23511576 | 36.61 | 1 | 6 | 5 | 1 | 0 | 3 | 0 | 1 | 4578.83845 | 0.0113 | 2.47 | 3.23 |
| Chymotrypsin | 2 | 174 | 7685 | 1 | 3835 | 4579.8576 | 39.8388 | 29,421 | 4578.849775 | 21 | 66 | 79 | HAIHVSGTNGTKRF | 0.00000000200000.0 | 74(NGT) | 1 | 1526.784 | 1523.795739 | 23511576 | 36.61 | 2 | 7 | 5 | 0 | 1 | 2 | 0 | 0 | 4578.838451 | 0.0113 | 2.47 | 3.23 |
| Chymotrypsin | 2 | 174 | 7685 | 1 | 3835 | 4579.8576 | 39.8388 | 29,421 | 4578.849775 | 21 | 66 | 79 | HAIHVSGTNGTKRF | 0.00000000200000.0 | 74(NGT) | 1 | 1526.784 | 1523.795739 | 23511576 | 36.61 | 3 | 6 | 3 | 4 | 2 | 1 | 1 | -1 | 4578.863603 | -0.0138 | -3.01 | 3.23 |
| Chymotrypsin | 2 | 174 | 7685 | 1 | 3835 | 4579.8576 | 39.8388 | 29,421 | 4578.849775 | 21 | 66 | 79 | HAIHVSGTNGTKRF | 0.00000000200000.0 | 74(NGT) | 1 | 1526.784 | 1523.795739 | 23511576 | 36.61 | 4 | 7 | 3 | 3 | 3 | 0 | 1 | -1 | 4578.863604 | -0.0138 | -3.02 | 3.23 |
| Chymotrypsin | 2 | 174 | 7723 | 1 | 3718 | 4339.8248 | 34.8306 | 28,626 | 4338.816975 | 21 | 66 | 79 | HAIHVSGTNGTKRF | 0.00000000200000.0 | 74(NGT) | 1 | 1526.784 | 1523.795739 | 23511576 | 36.61 | 1 | 4 | 7 | 3 | 0 | 1 | 0 | 0 | 4338.826704 | -0.0097 | -2.24 | -1.78 |
| Chymotrypsin | 2 | 174 | 7723 | 1 | 3718 | 4339.8248 | 34.8306 | 28,626 | 4338.816975 | 21 | 66 | 79 | HAIHVSGTNGTKRF | 0.00000000200000.0 | 74(NGT) | 1 | 1526.784 | 1523.795739 | 23511576 | 36.61 | 2 | 5 | 7 | 2 | 1 | 0 | 0 | 0 | 4338.826705 | -0.0097 | -2.24 | -1.78 |
| Chymotrypsin | 2 | 174 | 7821 | 1 | 3827 | 5091.0319 | 39.4927 | 27,272 | 5090.024075 | 21 | 66 | 79 | HAIHVSGTNGTKRF | 0.00000000200000.0 | 74(NGT) | 1 | 1526.784 | 1523.795739 | 23511576 | 36.61 | 1 | 7 | 6 | 2 | 0 | 3 | 0 | 0 | 5090.028556 | -0.0045 | -0.88 | 2.88 |
| Chymotrypsin | 2 | 174 | 7821 | 1 | 3827 | 5091.0319 | 39.4927 | 27,272 | 5090.024075 | 21 | 66 | 79 | HAIHVSGTNGTKRF | 0.00000000200000.0 | 74(NGT) | 1 | 1526.784 | 1523.795739 | 23511576 | 36.61 | 2 | 8 | 6 | 1 | 1 | 2 | 0 | 0 | 5090.028557 | -0.0045 | -0.88 | 2.88 |
| Chymotrypsin | 2 | 174 | 7821 | 1 | 3827 | 5091.0319 | 39.4927 | 27,272 | 5090.024075 | 21 | 66 | 79 | HAIHVSGTNGTKRF | 0.00000000200000.0 | 74(NGT) | 1 | 1526.784 | 1523.795739 | 23511576 | 36.61 | 3 | 9 | 6 | 0 | 2 | 1 | 0 | 0 | 5090.028558 | -0.0045 | -0.88 | 2.88 |
| Chymotrypsin | 2 | 174 | 7835 | 1 | 3661 | 4340.8595 | 32.3932 | 27,021 | 4339.851675 | 21 | 66 | 79 | HAIHVSGTNGTKRF | 0.00000000200000.0 | 74(NGT) | 1 | 1526.784 | 1523.795739 | 23511576 | 36.61 | 1 | 5 | 7 | 4 | 0 | 0 | 0 | 0 | 4339.847106 | 0.0046 | 1.05 | -4.22 |
| Chymotrypsin | 2 | 174 | 7877 | 1 | 3729 | 4193.7736 | 35.2926 | 26,574 | 4192.765775 | 21 | 66 | 79 | HAIHVSGTNGTKRF | 0.00000000200000.0 | 74(NGT) | 1 | 1526.784 | 1523.795739 | 23511576 | 36.61 | 1 | 4 | 7 | 2 | 0 | 1 | 0 | 0 | 4192.768795 | -0.0030 | -0.72 | -1.32 |
| Chymotrypsin | 2 | 174 | 7877 | 1 | 3729 | 4193.7736 | 35.2926 | 26,574 | 4192.765775 | 21 | 66 | 79 | HAIHVSGTNGTKRF | 0.00000000200000.0 | 74(NGT) | 1 | 1526.784 | 1523.795739 | 23511576 | 36.61 | 2 | 5 | 7 | 1 | 1 | 0 | 0 | 0 | 4192.768796 | -0.0030 | -0.72 | -1.32 |
| Chymotrypsin | 2 | 174 | 7893 | 1 | 3775 | 4971.0421 | 37.2675 | 26,359 | 4970.034275 | 21 | 66 | 79 | HAIHVSGTNGTKRF | 0.00000000200000.0 | 74(NGT) | 1 | 1526.784 | 1523.795739 | 23511576 | 36.61 | 1 | 8 | 7 | 1 | 2 | 0 | 0 | 1 | 4970.022685 | 0.0116 | 2.33 | 0.66 |
| Chymotrypsin | 2 | 174 | 7893 | 1 | 3775 | 4971.0421 | 37.2675 | 26,359 | 4970.034275 | 21 | 66 | 79 | HAIHVSGTNGTKRF | 0.00000000200000.0 | 74(NGT) | 1 | 1526.784 | 1523.795739 | 23511576 | 36.61 | 2 | 7 | 7 | 2 | 1 | 1 | 0 | 0 | 4970.022684 | 0.0116 | 2.33 | 0.66 |
| Chymotrypsin | 2 | 174 | 7893 | 1 | 3775 | 4971.0421 | 37.2675 | 26,359 | 4970.034275 | 21 | 66 | 79 | HAIHVSGTNGTKRF | 0.00000000200000.0 | 74(NGT) | 1 | 1526.784 | 1523.795739 | 23511576 | 36.61 | 3 | 6 | 7 | 3 | 0 | 2 | 0 | 0 | 4970.022683 | 0.0116 | 2.33 | 0.66 |
| Chymotrypsin | 2 | 174 | 7940 | 1 | 3715 | 4403.8253 | 34.703 | 25,841 | 4402.817475 | 21 | 66 | 79 | HAIHVSGTNGTKRF | 0.00000000200000.0 | 74(NGT) | 1 | 1526.784 | 1523.795739 | 23511576 | 36.61 | 1 | 6 | 5 | 4 | 0 | 1 | 0 | 0 | 4402.831515 | -0.0140 | -3.18 | -1.91 |
| Chymotrypsin | 2 | 174 | 7940 | 1 | 3715 | 4403.8253 | 34.703 | 25,841 | 4402.817475 | 21 | 66 | 79 | HAIHVSGTNGTKRF | 0.00000000200000.0 | 74(NGT) | 1 | 1526.784 | 1523.795739 | 23511576 | 36.61 | 2 | 7 | 5 | 3 | 1 | 0 | 0 | 0 | 4402.831516 | -0.0140 | -3.18 | -1.91 |
| Chymotrypsin | 2 | 174 | 8093 | 1 | 3769 | 4914.0114 | 37.0103 | 24,117 | 4913.003575 | 21 | 66 | 79 | HAIHVSGTNGTKRF | 0.00000000200000.0 | 74(NGT) | 1 | 1526.784 | 1523.795739 | 23511576 | 36.61 | 1 | 8 | 6 | 2 | 2 | 0 | 0 | 0 | 4913.001221 | 0.0024 | 0.47 | 0.40 |
| Chymotrypsin | 2 | 174 | 8093 | 1 | 3769 | 4914.0114 | 37.0103 | 24,117 | 4913.003575 | 21 | 66 | 79 | HAIHVSGTNGTKRF | 0.00000000200000.0 | 74(NGT) | 1 | 1526.784 | 1523.795739 | 23511576 | 36.61 | 2 | 7 | 6 | 3 | 1 | 1 | 0 | 0 | 4913.00122 | 0.0024 | 0.47 | 0.40 |
| Chymotrypsin | 2 | 174 | 8093 | 1 | 3769 | 4914.0114 | 37.0103 | 24,117 | 4913.003575 | 21 | 66 | 79 | HAIHVSGTNGTKRF | 0.00000000200000.0 | 74(NGT) | 1 | 1526.784 | 1523.795739 | 23511576 | 36.61 | 3 | 6 | 6 | 4 | 0 | 2 | 0 | 0 | 4913.001219 | 0.0024 | 0.47 | 0.40 |
| Chymotrypsin | 2 | 174 | 8108 | 1 | 3726 | 4314.8068 | 35.1674 | 23,943 | 4313.798975 | 21 | 66 | 79 | HAIHVSGTNGTKRF | 0.00000000200000.0 | 74(NGT) | 1 | 1526.784 | 1523.795739 | 23511576 | 36.61 | 1 | 7 | 6 | 1 | 1 | 0 | 0 | 1 | 4313.795071 | 0.0039 | 0.9 | -1.44 |
| Chymotrypsin | 2 | 174 | 8108 | 1 | 3726 | 4314.8068 | 35.1674 | 23,943 | 4313.798975 | 21 | 66 | 79 | HAIHVSGTNGTKRF | 0.00000000200000.0 | 74(NGT) | 1 | 1526.784 | 1523.795739 | 23511576 | 36.61 | 2 | 6 | 6 | 2 | 0 | 1 | 0 | 0 | 4313.79507 | 0.0039 | 0.9 | -1.44 |
| Chymotrypsin | 2 | 174 | 8150 | 1 | 3681 | 4413.8691 | 33.2502 | 23,459 | 4412.861275 | 21 | 66 | 79 | HAIHVSGTNGTKRF | 0.00000000200000.0 | 74(NGT) | 1 | 1526.784 | 1523.795739 | 23511576 | 36.61 | 1 | 6 | 8 | 2 | 0 | 0 | 0 | 0 | 4412.863485 | -0.0022 | -0.5 | -3.36 |
| Chymotrypsin | 2 | 174 | 8368 | 1 | 3796 | 4110.7042 | 38.1615 | 20,644 | 4109.696375 | 21 | 66 | 79 | HAIHVSGTNGTKRF | 0.00000000200000.0 | 74(NGT) | 1 | 1526.784 | 1523.795739 | 23511576 | 36.61 | 1 | 6 | 5 | 0 | 1 | 1 | 0 | 1 | 4109.695296 | 0.0011 | 0.26 | 1.55 |

[illegible]

| | | | | | | | | | | | | | | | | | | | | | | | | | | | | | | | |
|---|---|---|---|---|---|---|---|---|---|---|---|---|---|---|---|---|---|---|---|---|---|---|---|---|---|---|---|---|---|---|---|
| Trypsin | 104 | 49 | 1916 | 1 | 3487 | 3282.3897 | 24.6075 | 69,139 | 3281.381875 | 150 | 66 | 77 | HAIHVSGTNGTK | 0.000000002000.0 | 74(NGT) | 1 | 1223.6145 | 1220.626239 | 377363.72 | 25.77 | 1 | 5 | 4 | 3 | 0 | 0 | 0 | 1 | 3281.381578 | 0.0003 | 0.09 | -1.16 |
| Trypsin | 104 | 49 | 2899 | 1 | 3502 | 2974.2791 | 24.9007 | 41,142 | 2973.271275 | 150 | 66 | 77 | HAIHVSGTNGTK | 0.000000002000.0 | 74(NGT) | 1 | 1223.6145 | 1220.626239 | 377363.72 | 25.77 | 1 | 4 | 4 | 2 | 0 | 0 | 0 | 1 | 2973.270845 | 0.0004 | 0.14 | -0.87 |
| Trypsin | 104 | 49 | 3222 | 1 | 3496 | 3364.4423 | 24.7829 | 35,956 | 3363.434475 | 150 | 66 | 77 | HAIHVSGTNGTK | 0.000000002000.0 | 74(NGT) | 1 | 1223.6145 | 1220.626239 | 377363.72 | 25.77 | 1 | 3 | 6 | 3 | 0 | 0 | 0 | 1 | 3363.434676 | -0.0002 | -0.05 | -0.99 |
| Trypsin | 104 | 49 | 3739 | 1 | 3607 | 3193.3501 | 26.4402 | 28,961 | 3192.342275 | 150 | 66 | 77 | HAIHVSGTNGTK | 0.000000002000.0 | 74(NGT) | 1 | 1223.6145 | 1220.626239 | 377363.72 | 25.77 | 1 | 5 | 5 | 1 | 0 | 0 | 0 | 1 | 3192.345133 | -0.0029 | -0.89 | 0.67 |
| Trypsin | 104 | 49 | 3758 | 1 | 3606 | 3484.4457 | 26.3904 | 28,813 | 3483.437875 | 150 | 66 | 77 | HAIHVSGTNGTK | 0.000000002000.0 | 74(NGT) | 1 | 1223.6145 | 1220.626239 | 377363.72 | 25.77 | 1 | 4 | 5 | 2 | 0 | 1 | 0 | 1 | 3483.440549 | -0.0027 | -0.76 | 0.62 |
| Trypsin | 104 | 49 | 3758 | 1 | 3606 | 3484.4457 | 26.3904 | 28,813 | 3483.437875 | 150 | 66 | 77 | HAIHVSGTNGTK | 0.000000002000.0 | 74(NGT) | 1 | 1223.6145 | 1220.626239 | 377363.72 | 25.77 | 2 | 5 | 5 | 1 | 1 | 0 | 0 | 1 | 3483.44055 | -0.0027 | -0.76 | 0.62 |
| Trypsin | 104 | 49 | 4460 | 1 | 3636 | 2770.1823 | 27.1208 | 21,737 | 2769.174475 | 150 | 66 | 77 | HAIHVSGTNGTK | 0.000000002000.0 | 74(NGT) | 1 | 1223.6145 | 1220.626239 | 377363.72 | 25.77 | 1 | 4 | 3 | 0 | 1 | 0 | 0 | 1 | 2769.171071 | 0.0034 | 1.22 | 1.35 |
| Trypsin | 104 | 49 | 4460 | 1 | 3636 | 2770.1823 | 27.1208 | 21,737 | 2769.174475 | 150 | 66 | 77 | HAIHVSGTNGTK | 0.000000002000.0 | 74(NGT) | 1 | 1223.6145 | 1220.626239 | 377363.72 | 25.77 | 2 | 3 | 3 | 1 | 0 | 1 | 0 | 0 | 2769.17107 | 0.0034 | 1.22 | 1.35 |
| Trypsin | 104 | 49 | 4691 | 1 | 3518 | 3177.3583 | 25.0728 | 19,937 | 3176.350475 | 150 | 66 | 77 | HAIHVSGTNGTK | 0.000000002000.0 | 74(NGT) | 1 | 1223.6145 | 1220.626239 | 377363.72 | 25.77 | 1 | 4 | 5 | 2 | 0 | 0 | 0 | 1 | 3176.350218 | 0.0003 | 0.08 | -0.70 |
| Trypsin | 104 | 49 | 4800 | 1 | 3558 | 2666.167 | 25.5717 | 19,170 | 2665.159175 | 150 | 66 | 77 | HAIHVSGTNGTK | 0.000000002000.0 | 74(NGT) | 1 | 1223.6145 | 1220.626239 | 377363.72 | 25.77 | 1 | 3 | 4 | 1 | 0 | 0 | 0 | 1 | 2665.160112 | -0.0009 | -0.35 | -0.20 |
| Trypsin | 104 | 49 | 4949 | 1 | 3561 | 3031.3 | 25.6002 | 18,125 | 3030.292175 | 150 | 66 | 77 | HAIHVSGTNGTK | 0.000000002000.0 | 74(NGT) | 1 | 1223.6145 | 1220.626239 | 377363.72 | 25.77 | 1 | 4 | 5 | 1 | 0 | 0 | 0 | 1 | 3030.292309 | -0.0001 | -0.04 | -0.17 |
| Trypsin | 104 | 49 | 5032 | 1 | 3543 | 3072.326 | 25.395 | 17,489 | 3071.318175 | 150 | 66 | 77 | HAIHVSGTNGTK | 0.000000002000.0 | 74(NGT) | 1 | 1223.6145 | 1220.626239 | 377363.72 | 25.77 | 1 | 3 | 6 | 1 | 0 | 0 | 0 | 1 | 3071.318858 | -0.0007 | -0.22 | -0.38 |
| Trypsin | 104 | 49 | 5113 | 1 | 3489 | 3136.3322 | 24.6437 | 16,852 | 3135.324375 | 150 | 66 | 77 | HAIHVSGTNGTK | 0.000000002000.0 | 74(NGT) | 1 | 1223.6145 | 1220.626239 | 377363.72 | 25.77 | 1 | 5 | 4 | 2 | 0 | 0 | 0 | 1 | 3135.323669 | 0.0007 | 0.22 | -1.13 |
| Trypsin | 104 | 49 | 5165 | 1 | 3573 | 3339.4116 | 25.7878 | 16,565 | 3338.403775 | 150 | 66 | 77 | HAIHVSGTNGTK | 0.000000002000.0 | 74(NGT) | 1 | 1223.6145 | 1220.626239 | 377363.72 | 25.77 | 1 | 5 | 5 | 2 | 0 | 0 | 0 | 1 | 3338.403042 | 0.0007 | 0.21 | 0.02 |
| Trypsin | 104 | 49 | 5473 | 1 | 3602 | 2723.1914 | 26.2899 | 14,358 | 2722.183575 | 150 | 66 | 77 | HAIHVSGTNGTK | 0.000000002000.0 | 74(NGT) | 1 | 1223.6145 | 1220.626239 | 377363.72 | 25.77 | 1 | 3 | 5 | 0 | 0 | 0 | 0 | 1 | 2722.181576 | 0.0020 | 0.73 | 0.52 |
| Trypsin | 104 | 49 | 5527 | 1 | 3539 | 3218.3831 | 25.3505 | 13,992 | 3217.375275 | 150 | 66 | 77 | HAIHVSGTNGTK | 0.000000002000.0 | 74(NGT) | 1 | 1223.6145 | 1220.626239 | 377363.72 | 25.77 | 1 | 3 | 6 | 2 | 0 | 0 | 0 | 1 | 3217.376767 | -0.0015 | -0.46 | -0.42 |
| Trypsin | 104 | 49 | 5624 | 1 | 3490 | 3793.5767 | 24.6761 | 13,362 | 3792.568875 | 150 | 66 | 77 | HAIHVSGTNGTK | 0.000000002000.0 | 74(NGT) | 1 | 1223.6145 | 1220.626239 | 377363.72 | 25.77 | 1 | 6 | 5 | 4 | 0 | 0 | 0 | 0 | 3792.571684 | -0.0028 | -0.74 | -1.09 |
| Trypsin | 104 | 49 | 5852 | 1 | 3519 | 3136.3279 | 25.0795 | 12,083 | 3135.320075 | 150 | 66 | 77 | HAIHVSGTNGTK | 0.000000002000.0 | 74(NGT) | 1 | 1223.6145 | 1220.626239 | 377363.72 | 25.77 | 1 | 5 | 4 | 2 | 0 | 0 | 0 | 1 | 3135.323669 | -0.0036 | -1.14 | -0.69 |
| Trypsin | 104 | 49 | 6047 | 1 | 3610 | 3646.4965 | 26.5334 | 10,832 | 3645.488675 | 150 | 66 | 77 | HAIHVSGTNGTK | 0.000000002000.0 | 74(NGT) | 1 | 1223.6145 | 1220.626239 | 377363.72 | 25.77 | 1 | 6 | 5 | 1 | 1 | 0 | 0 | 1 | 3645.493374 | -0.0047 | -1.28 | 0.76 |
| Trypsin | 104 | 49 | 6047 | 1 | 3610 | 3646.4965 | 26.5334 | 10,832 | 3645.488675 | 150 | 66 | 77 | HAIHVSGTNGTK | 0.000000002000.0 | 74(NGT) | 1 | 1223.6145 | 1220.626239 | 377363.72 | 25.77 | 2 | 5 | 5 | 2 | 0 | 1 | 0 | 0 | 3645.493373 | -0.0047 | -1.28 | 0.76 |
| Trypsin | 104 | 49 | 6103 | 1 | 3603 | 3443.4235 | 26.3192 | 10,551 | 3442.415675 | 150 | 66 | 77 | HAIHVSGTNGTK | 0.000000002000.0 | 74(NGT) | 1 | 1223.6145 | 1220.626239 | 377363.72 | 25.77 | 1 | 6 | 4 | 1 | 1 | 0 | 0 | 1 | 3442.414001 | 0.0017 | 0.48 | 0.55 |
| Trypsin | 104 | 49 | 6103 | 1 | 3603 | 3443.4235 | 26.3192 | 10,551 | 3442.415675 | 150 | 66 | 77 | HAIHVSGTNGTK | 0.000000002000.0 | 74(NGT) | 1 | 1223.6145 | 1220.626239 | 377363.72 | 25.77 | 2 | 5 | 4 | 2 | 0 | 1 | 0 | 1 | 3442.414 | 0.0017 | 0.48 | 0.55 |
| Trypsin | 104 | 49 | 6118 | 1 | 3492 | 2771.1927 | 24.6976 | 10,477 | 2770.184875 | 150 | 66 | 77 | HAIHVSGTNGTK | 0.000000002000.0 | 74(NGT) | 1 | 1223.6145 | 1220.626239 | 377363.72 | 25.77 | 1 | 4 | 3 | 2 | 0 | 0 | 0 | 1 | 2770.191472 | -0.0066 | -2.38 | -1.07 |
| Trypsin | 104 | 49 | 6175 | 1 | 3644 | 3558.4824 | 27.2509 | 10,124 | 3557.474575 | 150 | 66 | 77 | HAIHVSGTNGTK | 0.000000002000.0 | 74(NGT) | 1 | 1223.6145 | 1220.626239 | 377363.72 | 25.77 | 1 | 6 | 6 | 1 | 0 | 0 | 0 | 1 | 3557.47733 | -0.0028 | -0.77 | 1.48 |
| Trypsin | 104 | 49 | 6435 | 1 | 3532 | 3720.5375 | 25.2774 | 8,561 | 3719.529675 | 150 | 66 | 77 | HAIHVSGTNGTK | 0.000000002000.0 | 74(NGT) | 1 | 1223.6145 | 1220.626239 | 377363.72 | 25.77 | 1 | 7 | 6 | 1 | 0 | 0 | 0 | 1 | 3719.530154 | -0.0005 | -0.12 | -0.49 |
| Trypsin | 104 | 49 | 6472 | 1 | 3557 | 2990.273 | 25.5648 | 8,406 | 2989.265175 | 150 | 66 | 77 | HAIHVSGTNGTK | 0.000000002000.0 | 74(NGT) | 1 | 1223.6145 | 1220.626239 | 377363.72 | 25.77 | 1 | 5 | 4 | 1 | 0 | 0 | 0 | 1 | 2989.26576 | -0.0006 | -0.19 | -0.21 |
| Trypsin | 104 | 49 | 6559 | 1 | 3590 | 2520.1059 | 26.0059 | 7,775 | 2519.098075 | 150 | 66 | 77 | HAIHVSGTNGTK | 0.000000002000.0 | 74(NGT) | 1 | 1223.6145 | 1220.626239 | 377363.72 | 25.77 | 1 | 3 | 4 | 0 | 0 | 0 | 0 | 1 | 2519.102203 | -0.0041 | -1.63 | 0.23 |
| Trypsin | 104 | 49 | 6600 | 1 | 3505 | 2812.2259 | 24.9293 | 7,464 | 2811.218075 | 150 | 66 | 77 | HAIHVSGTNGTK | 0.000000002000.0 | 74(NGT) | 1 | 1223.6145 | 1220.626239 | 377363.72 | 25.77 | 1 | 3 | 4 | 2 | 0 | 0 | 0 | 1 | 2811.218021 | 0.0001 | 0.01 | -0.84 |
| Trypsin | 104 | 49 | 6603 | 1 | 3449 | 3834.607 | 24.2438 | 7,452 | 3833.599175 | 150 | 66 | 77 | HAIHVSGTNGTK | 0.000000002000.0 | 74(NGT) | 1 | 1223.6145 | 1220.626239 | 377363.72 | 25.77 | 1 | 5 | 6 | 4 | 0 | 0 | 0 | 0 | 3833.598233 | 0.0009 | 0.24 | -1.53 |
| Trypsin | 104 | 49 | 6747 | 1 | 3527 | 2828.2212 | 25.1963 | 6,394 | 2827.213375 | 150 | 66 | 77 | HAIHVSGTNGTK | 0.000000002000.0 | 74(NGT) | 1 | 1223.6145 | 1220.626239 | 377363.72 | 25.77 | 1 | 4 | 4 | 1 | 0 | 0 | 0 | 1 | 2827.212936 | 0.0004 | 0.15 | -0.57 |
| Trypsin | 104 | 49 | 6846 | 1 | 3581 | 2869.2419 | 25.9262 | 5,798 | 2868.234075 | 150 | 66 | 77 | HAIHVSGTNGTK | 0.000000002000.0 | 74(NGT) | 1 | 1223.6145 | 1220.626239 | 377363.72 | 25.77 | 1 | 3 | 5 | 1 | 0 | 0 | 0 | 1 | 2868.239485 | -0.0054 | -1.88 | 0.16 |
| Trypsin | 104 | 49 | 6902 | 1 | 3513 | 3526.4954 | 25.0204 | 5,429 | 3525.487575 | 150 | 66 | 77 | HAIHVSGTNGTK | 0.000000002000.0 | 74(NGT) | 1 | 1223.6145 | 1220.626239 | 377363.72 | 25.77 | 1 | 4 | 6 | 3 | 0 | 0 | 0 | 0 | 3525.4875 | 0.0001 | 0.02 | -0.75 |
| Trypsin | 104 | 49 | 6956 | 1 | 3607 | 3031.2964 | 26.4402 | 4,958 | 3030.288575 | 150 | 66 | 77 | HAIHVSGTNGTK | 0.000000002000.0 | 74(NGT) | 1 | 1223.6145 | 1220.626239 | 377363.72 | 25.77 | 1 | 4 | 5 | 1 | 0 | 0 | 0 | 1 | 3030.292309 | -0.0037 | -1.23 | 0.67 |
| [illegible] | [illegible] | [illegible] | [illegible] | [illegible] | [illegible] | [illegible] | [illegible] | [illegible] | [illegible] | [illegible] | [illegible] | [illegible] | [illegible] | [illegible] | [illegible] | [illegible] | [illegible] | [illegible] | [illegible] | [illegible] | [illegible] | [illegible] | [illegible] | [illegible] | [illegible] | [illegible] | [illegible] | [illegible] | [illegible] | [illegible] | [illegible] | [illegible] |

| | | | | | | | | | | | | | |
|---|---|---|---|---|---|---|---|---|---|---|---|---|---|
| Chymotrypsin | 53 | 78 | 6550 | 1 | 4184 | 4796.841 | 54.6598 | 44,506 | 4795.833175 | 98 | 145 | 157 | YHKNNKSWMESEF |
| Chymotrypsin | 53 | 78 | 6550 | 1 | 4184 | 4796.841 | 54.6598 | 44,506 | 4795.833175 | 98 | 145 | 157 | YHKNNKSWMESEF |
| Chymotrypsin | 53 | 78 | 6632 | 1 | 4098 | 4781.8693 | 51.0346 | 43,315 | 4780.861475 | 98 | 145 | 157 | YHKNNKSWMESEF |
| Chymotrypsin | 53 | 78 | 6632 | 1 | 4098 | 4781.8693 | 51.0346 | 43,315 | 4780.861475 | 98 | 145 | 157 | YHKNNKSWMESEF |
| Chymotrypsin | 53 | 78 | 6745 | 1 | 4016 | 4531.8239 | 47.5367 | 41,715 | 4530.816075 | 98 | 145 | 157 | YHKNNKSWMESEF |
| Chymotrypsin | 53 | 78 | 6822 | 1 | 4174 | 5104.9498 | 54.2318 | 40,689 | 5103.941975 | 98 | 145 | 157 | YHKNNKSWMESEF |
| Chymotrypsin | 53 | 78 | 6822 | 1 | 4174 | 5104.9498 | 54.2318 | 40,689 | 5103.941975 | 98 | 145 | 157 | YHKNNKSWMESEF |
| Chymotrypsin | 53 | 78 | 6822 | 1 | 4174 | 5104.9498 | 54.2318 | 40,689 | 5103.941975 | 98 | 145 | 157 | YHKNNKSWMESEF |
| Chymotrypsin | 53 | 78 | 6845 | 1 | 4096 | 4927.9249 | 50.951 | 40,321 | 4926.917075 | 98 | 145 | 157 | YHKNNKSWMESEF |
| Chymotrypsin | 53 | 78 | 6845 | 1 | 4096 | 4927.9249 | 50.951 | 40,321 | 4926.917075 | 98 | 145 | 157 | YHKNNKSWMESEF |
| Chymotrypsin | 53 | 78 | 6845 | 1 | 4096 | 4927.9249 | 50.951 | 40,321 | 4926.917075 | 98 | 145 | 157 | YHKNNKSWMESEF |
| Chymotrypsin | 53 | 78 | 6927 | 1 | 4184 | 4593.7677 | 54.6598 | 39,229 | 4592.759875 | 98 | 145 | 157 | YHKNNKSWMESEF |
| Chymotrypsin | 53 | 78 | 6927 | 1 | 4184 | 4593.7677 | 54.6598 | 39,229 | 4592.759875 | 98 | 145 | 157 | YHKNNKSWMESEF |
| Chymotrypsin | 53 | 78 | 6927 | 1 | 4184 | 4593.7677 | 54.6598 | 39,229 | 4592.759875 | 98 | 145 | 157 | YHKNNKSWMESEF |
| Chymotrypsin | 53 | 78 | 6983 | 1 | 4090 | 5000.9523 | 50.6924 | 38,316 | 4999.944475 | 98 | 145 | 157 | YHKNNKSWMESEF |
| Chymotrypsin | 53 | 78 | 6983 | 1 | 4090 | 5000.9523 | 50.6924 | 38,316 | 4999.944475 | 98 | 145 | 157 | YHKNNKSWMESEF |
| Chymotrypsin | 53 | 78 | 7205 | 1 | 4044 | 3712.506 | 48.7313 | 35,177 | 3711.498175 | 98 | 145 | 157 | YHKNNKSWMESEF |
| Chymotrypsin | 53 | 78 | 7351 | 1 | 4120 | 4416.7291 | 51.971 | 33,583 | 4415.721275 | 98 | 145 | 157 | YHKNNKSWMESEF |
| Chymotrypsin | 53 | 78 | 7351 | 1 | 4120 | 4416.7291 | 51.971 | 33,583 | 4415.721275 | 98 | 145 | 157 | YHKNNKSWMESEF |
| Chymotrypsin | 53 | 78 | 7462 | 1 | 4112 | 4327.7042 | 51.622 | 32,248 | 4326.696375 | 98 | 145 | 157 | YHKNNKSWMESEF |
| Chymotrypsin | 53 | 78 | 7462 | 1 | 4112 | 4327.7042 | 51.622 | 32,248 | 4326.696375 | 98 | 145 | 157 | YHKNNKSWMESEF |
| Chymotrypsin | 53 | 78 | 8024 | 1 | 4131 | 3921.5548 | 52.4426 | 24,861 | 3920.546975 | 98 | 145 | 157 | YHKNNKSWMESEF |
| Chymotrypsin | 53 | 78 | 8024 | 1 | 4131 | 3921.5548 | 52.4426 | 24,861 | 3920.546975 | 98 | 145 | 157 | YHKNNKSWMESEF |
| Chymotrypsin | 53 | 78 | 8189 | 1 | 4061 | 4822.902 | 49.4552 | 23,041 | 4821.894175 | 98 | 145 | 157 | YHKNNKSWMESEF |
| Chymotrypsin | 53 | 78 | 8189 | 1 | 4061 | 4822.902 | 49.4552 | 23,041 | 4821.894175 | 98 | 145 | 157 | YHKNNKSWMESEF |
| Chymotrypsin | 53 | 78 | 8195 | 1 | 4182 | 4942.9015 | 54.5763 | 22,953 | 4941.893675 | 98 | 145 | 157 | YHKNNKSWMESEF |
| Chymotrypsin | 53 | 78 | 8195 | 1 | 4182 | 4942.9015 | 54.5763 | 22,953 | 4941.893675 | 98 | 145 | 157 | YHKNNKSWMESEF |
| Chymotrypsin | 53 | 78 | 8195 | 1 | 4182 | 4942.9015 | 54.5763 | 22,953 | 4941.893675 | 98 | 145 | 157 | YHKNNKSWMESEF |
| Chymotrypsin | 53 | 78 | 8248 | 1 | 4161 | 4634.7799 | 53.6825 | 22,392 | 4633.772075 | 98 | 145 | 157 | YHKNNKSWMESEF |
| Chymotrypsin | 53 | 78 | 8248 | 1 | 4161 | 4634.7799 | 53.6825 | 22,392 | 4633.772075 | 98 | 145 | 157 | YHKNNKSWMESEF |
| Chymotrypsin | 53 | 78 | 8248 | 1 | 4161 | 4634.7799 | 53.6825 | 22,392 | 4633.772075 | 98 | 145 | 157 | YHKNNKSWMESEF |
| Chymotrypsin | 86 | 44 | 321 | 1 | 3983 | 4108.6309 | 46.1492 | 1,082,790 | 4107.623075 | 370 | 146 | 157 | HKNNKSWMESEF |
| Chymotrypsin | 86 | 44 | 343 | 1 | 3976 | 4619.822 | 45.859 | 1,005,300 | 4618.814175 | 370 | 146 | 157 | HKNNKSWMESEF |
| Chymotrypsin | 86 | 44 | 343 | 1 | 3976 | 4619.822 | 45.859 | 1,005,300 | 4618.814175 | 370 | 146 | 157 | HKNNKSWMESEF |
| Chymotrypsin | 86 | 44 | 380 | 1 | 3987 | 4311.7129 | 46.3079 | 923,563 | 4310.705075 | 370 | 146 | 157 | HKNNKSWMESEF |
| Chymotrypsin | 86 | 44 | 395 | 1 | 3999 | 3800.521 | 46.8152 | 872,109 | 3799.513175 | 370 | 146 | 157 | HKNNKSWMESEF |
| Chymotrypsin | 86 | 44 | 517 | 1 | 3998 | 3597.4408 | 46.7739 | 688,589 | 3596.432975 | 370 | 146 | 157 | HKNNKSWMESEF |
| Chymotrypsin | 86 | 44 | 1281 | 1 | 3979 | 4473.7697 | 45.9874 | 298,845 | 4472.761875 | 370 | 146 | 157 | HKNNKSWMESEF |
| Chymotrypsin | 86 | 44 | 1281 | 1 | 3979 | 4473.7697 | 45.9874 | 298,845 | 4472.761875 | 370 | 146 | 157 | HKNNKSWMESEF |
| Chymotrypsin | 86 | 44 | 1283 | 1 | 3972 | 4822.904 | 45.6906 | 298,548 | 4821.896175 | 370 | 146 | 157 | HKNNKSWMESEF |
| Chymotrypsin | 86 | 44 | 1423 | 1 | 4008 | 3451.3882 | 47.1937 | 268,990 | 3450.380375 | 370 | 146 | 157 | HKNNKSWMESEF |
| Chymotrypsin | 86 | 44 | 1426 | 1 | 3983 | 4149.6613 | 46.1492 | 267,706 | 4148.653475 | 370 | 146 | 157 | HKNNKSWMESEF |
| Chymotrypsin | 86 | 44 | 1603 | 1 | 4019 | 3508.4066 | 47.6659 | 242,294 | 3507.398775 | 370 | 146 | 157 | HKNNKSWMESEF |
| Chymotrypsin | 86 | 44 | 1615 | 1 | 4016 | 3289.3351 | 47.5367 | 240,958 | 3288.327275 | 370 | 146 | 157 | HKNNKSWMESEF |
| Chymotrypsin | 86 | 44 | 1888 | 1 | 4074 | 4269.6671 | 50.0071 | 206,300 | 4268.659275 | 370 | 146 | 157 | HKNNKSWMESEF |
| Chymotrypsin | 86 | 44 | 1888 | 1 | 4074 | 4269.6671 | 50.0071 | 206,300 | 4268.659275 | 370 | 146 | 157 | HKNNKSWMESEF |
| Chymotrypsin | 86 | 44 | 1940 | 1 | 4088 | 3758.4856 | 50.6098 | 200,350 | 3757.477775 | 370 | 146 | 157 | HKNNKSWMESEF |
| Chymotrypsin | 86 | 44 | 1940 | 1 | 4088 | 3758.4856 | 50.6098 | 200,350 | 3757.477775 | 370 | 146 | 157 | HKNNKSWMESEF |
| Chymotrypsin | 86 | 44 | 2075 | 1 | 4017 | 3305.3258 | 47.5783 | 188,086 | 3304.317975 | 370 | 146 | 157 | HKNNKSWMESEF |
| Chymotrypsin | 86 | 44 | 2239 | 1 | 4009 | 3492.4164 | 47.2356 | 175,094 | 3491.408575 | 370 | 146 | 157 | HKNNKSWMESEF |
| Chymotrypsin | 86 | 44 | 2320 | 1 | 3992 | 4165.6592 | 46.5242 | 167,622 | 4164.651375 | 370 | 146 | 157 | HKNNKSWMESEF |
| Chymotrypsin | 86 | 44 | 2325 | 1 | 4007 | 4019.5981 | 47.1519 | 166,966 | 4018.590275 | 370 | 146 | 157 | HKNNKSWMESEF |
| Chymotrypsin | 86 | 44 | 2433 | 1 | 3990 | 3962.5681 | 46.4389 | 159,624 | 3961.560275 | 370 | 146 | 157 | HKNNKSWMESEF |
| Chymotrypsin | 86 | 44 | 2520 | 1 | 3981 | 4327.7042 | 46.065 | 154,243 | 4326.696375 | 370 | 146 | 157 | HKNNKSWMESEF |
| Chymotrypsin | 86 | 44 | 2567 | 1 | 4029 | 2981.2247 | 48.0895 | 151,228 | 2980.216875 | 370 | 146 | 157 | HKNNKSWMESEF |
| Chymotrypsin | 86 | 44 | 2656 | 1 | 4017 | 3346.3516 | 47.5783 | 145,769 | 3345.343775 | 370 | 146 | 157 | HKNNKSWMESEF |
| Chymotrypsin | 86 | 44 | 2663 | 1 | 4089 | 3961.5628 | 50.6509 | 145,377 | 3960.554975 | 370 | 146 | 157 | HKNNKSWMESEF |
| Chymotrypsin | 86 | 44 | 2663 | 1 | 4089 | 3961.5628 | 50.6509 | 145,377 | 3960.554975 | 370 | 146 | 157 | HKNNKSWMESEF |
| Chymotrypsin | 86 | 44 | 2722 | 1 | 4014 | 3873.5425 | 47.4488 | 141,783 | 3872.534675 | 370 | 146 | 157 | HKNNKSWMESEF |
| Chymotrypsin | 86 | 44 | 2957 | 1 | 3977 | 4676.8576 | 45.9007 | 129,644 | 4675.849775 | 370 | 146 | 157 | HKNNKSWMESEF |
| Chymotrypsin | 86 | 44 | 3488 | 1 | 4027 | 3143.2758 | 48.0043 | 107,452 | 3142.267975 | 370 | 146 | 157 | HKNNKSWMESEF |
| Chymotrypsin | 86 | 44 | 3641 | 1 | 3991 | 4003.6012 | 46.4821 | 102,388 | 4002.593375 | 370 | 146 | 157 | HKNNKSWMESEF |
| Chymotrypsin | 86 | 44 | 3781 | 1 | 4015 | 3387.3844 | 47.4932 | 97,657 | 3386.376575 | 370 | 146 | 157 | HKNNKSWMESEF |
| Chymotrypsin | 86 | 44 | 3822 | 1 | 4021 | 3184.3014 | 47.7502 | 96,764 | 3183.293575 | 370 | 146 | 157 | HKNNKSWMESEF |
| Chymotrypsin | 86 | 44 | 4141 | 1 | 3976 | 4660.8532 | 45.859 | 88,256 | 4659.845375 | 370 | 146 | 157 | HKNNKSWMESEF |
| Chymotrypsin | 86 | 44 | 4146 | 1 | 4094 | 4180.6324 | 50.8629 | 88,025 | 4179.624575 | 370 | 146 | 157 | HKNNKSWMESEF |
| Chymotrypsin | 86 | 44 | 4146 | 1 | 4094 | 4180.6324 | 50.8629 | 88,025 | 4179.624575 | 370 | 146 | 157 | HKNNKSWMESEF |
| Chymotrypsin | 86 | 44 | 4273 | 1 | 4066 | 4634.7995 | 49.6671 | 84,541 | 4633.791675 | 370 | 146 | 157 | HKNNKSWMESEF |
| Chymotrypsin | 86 | 44 | 4273 | 1 | 4066 | 4634.7995 | 49.6671 | 84,541 | 4633.791675 | 370 | 146 | 157 | HKNNKSWMESEF |
| Chymotrypsin | 86 | 44 | 4557 | 1 | 4005 | 3670.4631 | 47.0678 | 77,628 | 3669.455275 | 370 | 146 | 157 | HKNNKSWMESEF |
| Chymotrypsin | 86 | 44 | 4822 | 1 | 4096 | 3612.4137 | 50.951 | 72,334 | 3611.405875 | 370 | 146 | 157 | HKNNKSWMESEF |
| Chymotrypsin | 86 | 44 | 4822 | 1 | 4096 | 3612.4137 | 50.951 | 72,334 | 3611.405875 | 370 | 146 | 157 | HKNNKSWMESEF |
| Chymotrypsin | 86 | 44 | 5617 | 1 | 4072 | 4488.7421 | 49.9241 | 58,274 | 4487.734275 | 370 | 146 | 157 | HKNNKSWMESEF |
| Chymotrypsin | 86 | 44 | 5617 | 1 | 4072 | 4488.7421 | 49.9241 | 58,274 | 4487.734275 | 370 | 146 | 157 | HKNNKSWMESEF |
| Chymotrypsin | 86 | 44 | 6725 | 1 | 3984 | 4181.6583 | 46.1925 | 41,939 | 4180.650475 | 370 | 146 | 157 | HKNNKSWMESEF |
| Chymotrypsin | 86 | 44 | 6799 | 1 | 4159 | 4430.6841 | 53.5959 | 40,905 | 4429.676275 | 370 | 146 | 157 | HKNNKSWMESEF |
| Chymotrypsin | 86 | 44 | 6799 | 1 | 4159 | 4430.6841 | 53.5959 | 40,905 | 4429.676275 | 370 | 146 | 157 | HKNNKSWMESEF |
| Chymotrypsin | 86 | 44 | 6799 | 1 | 4159 | 4430.6841 | 53.5959 | 40,905 | 4429.676275 | 370 | 146 | 157 | HKNNKSWMESEF |
| Chymotrypsin | 86 | 44 | 7130 | 1 | 4080 | 4123.6066 | 50.2642 | 36,164 | 4122.598775 | 370 | 146 | 157 | HKNNKSWMESEF |
| Chymotrypsin | 86 | 44 | 7130 | 1 | 4080 | 4123.6066 | 50.2642 | 36,164 | 4122.598775 | 370 | 146 | 157 | HKNNKSWMESEF |
| Chymotrypsin | 86 | 44 | 7652 | 1 | 4006 | 3711.4894 | 47.1096 | 29,720 | 3710.481575 | 370 | 146 | 157 | HKNNKSWMESEF |
| Chymotrypsin | 86 | 44 | 7699 | 1 | 4164 | 4284.6403 | 53.8091 | 29,288 | 4283.632475 | 370 | 146 | 157 | HKNNKSWMESEF |
| Chymotrypsin | 86 | 44 | 7699 | 1 | 4164 | 4284.6403 | 53.8091 | 29,288 | 4283.632475 | 370 | 146 | 157 | HKNNKSWMESEF |
| Chymotrypsin | 86 | 44 | 8237 | 1 | 4034 | 2778.1461 | 48.3028 | 22,490 | 2777.138275 | 370 | 146 | 157 | HKNNKSWMESEF |
| Chymotrypsin | 86 | 44 | 8495 | 1 | 4081 | 4456.7584 | 50.3078 | 18,791 | 4455.750575 | 370 | 146 | 157 | HKNNKSWMESEF |
| Chymotrypsin | 86 | 44 | 8495 | 1 | 4081 | 4456.7584 | 50.3078 | 18,791 | 4455.750575 | 370 | 146 | 157 | HKNNKSWMESEF |
| Chymotrypsin | 86 | 44 | 8551 | 1 | 4153 | 4795.8312 | 53.3384 | 18,014 | 4794.823375 | 370 | 146 | 157 | HKNNKSWMESEF |
| Chymotrypsin | 86 | 44 | 8551 | 1 | 4153 | 4795.8312 | 53.3384 | 18,014 | 4794.823375 | 370 | 146 | 157 | HKNNKSWMESEF |
| Chymotrypsin | 86 | 44 | 8551 | 1 | 4153 | 4795.8312 | 53.3384 | 18,014 | 4794.823375 | 370 | 146 | 157 | HKNNKSWMESEF |
| Chymotrypsin | 86 | 44 | 8637 | 1 | 4079 | 4253.6745 | 50.2207 | 17,022 | 4252.666675 | 370 | 146 | 157 | HKNNKSWMESEF |
| Chymotrypsin | 86 | 44 | 8637 | 1 | 4079 | 4253.6745 | 50.2207 | 17,022 | 4252.666675 | 370 | 146 | 157 | HKNNKSWMESEF |
| Chymotrypsin | 86 | 44 | 8918 | 1 | 4076 | 4545.7609 | 50.094 | 12,105 | 4544.753075 | 370 | 146 | 157 | HKNNKSWMESEF |
| Chymotrypsin | 86 | 44 | 8918 | 1 | 4076 | 4545.7609 | 50.094 | 12,105 | 4544.753075 | 370 | 146 | 157 | HKNNKSWMESEF |
| Chymotrypsin | 285 | 13 | 2988 | 1 | 3834 | 4490.7806 | 39.7963 | 127,789 | 4489.772775 | 150 | 145 | 157 | YHKNNKSWMESEF |
| Chymotrypsin | 285 | 13 | 3359 | 1 | 3841 | 3979.5792 | 40.0994 | 111,931 | 3978.571375 | 150 | 145 | 157 | YHKNNKSWMESEF |
| Chymotrypsin | 285 | 13 | 3494 | 1 | 3839 | 3776.4931 | 40.013 | 107,216 | 3775.485275 | 150 | 145 | 157 | YHKNNKSWMESEF |
| Chymotrypsin | 285 | 13 | 4299 | 1 | 3830 | 4287.6894 | 39.622 | 83,871 | 4286.681575 | 150 | 145 | 157 | YHKNNKSWMESEF |
| Chymotrypsin | 285 | 13 | 4631 | 1 | 3882 | 4797.8584 | 41.8517 | 76,390 | 4796.850575 | 150 | 145 | 157 | YHKNNKSWMESEF |
| Chymotrypsin | 285 | 13 | 4631 | 1 | 3882 | 4797.8584 | 41.8517 | 76,390 | 4796.850575 | 150 | 145 | 157 | YHKNNKSWMESEF |
| Chymotrypsin | 285 | 13 | 4684 | 1 | 3822 | 4798.8737 | 39.2785 | 75,281 | 4797.865875 | 150 | 145 | 157 | YHKNNKSWMESEF |
| Chymotrypsin | 285 | 13 | 4684 | 1 | 3822 | 4798.8737 | 39.2785 | 75,281 | 4797.865875 | 150 | 145 | 157 | YHKNNKSWMESEF |
| Chymotrypsin | 285 | 13 | 4806 | 1 | 3877 | 5000.9368 | 41.6372 | 72,727 | 4999.928975 | 150 | 145 | 157 | YHKNNKSWMESEF |
| Chymotrypsin | 285 | 13 | 4806 | 1 | 3877 | 5000.9368 | 41.6372 | 72,727 | 4999.928975 | 150 | 145 | 157 | YHKNNKSWMESEF |
| Chymotrypsin | 285 | 13 | 5664 | 1 | 3884 | 4854.882 | 41.9392 | 57,413 | 4853.874175 | 150 | 145 | 157 | YHKNNKSWMESEF |
| Chymotrypsin | 285 | 13 | 5664 | 1 | 3884 | 4854.882 | 41.9392 | 57,413 | 4853.874175 | 150 | 145 | 157 | YHKNNKSWMESEF |
| Chymotrypsin | 285 | 13 | 6375 | 1 | 3840 | 3817.5288 | 40.0555 | 46,806 | 3816.520975 | 150 | 145 | 157 | YHKNNKSWMESEF |
| Chymotrypsin | 285 | 13 | 7947 | 1 | 3825 | 4652.8107 | 39.4066 | 25,771 | 4651.802875 | 150 | 145 | 157 | YHKNNKSWMESEF |
| Chymotrypsin | 285 | 13 | 8559 | 1 | 3886 | 4651.8102 | 42.0224 | 17,944 | 4650.802375 | 150 | 145 | 157 | YHKNNKSWMESEF |
| Chymotrypsin | 285 | 13 | 8559 | 1 | 3886 | 4651.8102 | 42.0224 | 17,944 | 4650.802375 | 150 | 145 | 157 | YHKNNKSWMESEF |
| Chymotrypsin | 285 | 13 | 8693 | 1 | 3843 | 4344.715 | 40.1848 | 16,005 | 4343.707175 | 150 | 145 | 157 | YHKNNKSWMESEF |
| Chymotrypsin | 285 | 13 | 9109 | 1 | 3830 | 4506.7472 | 39.622 | 7,626 | 4505.739375 | 150 | 145 | 157 | YHKNNKSWMESEF |
| Chymotrypsin | 290 | 4 | 3113 | 1 | 4064 | 2916.1771 | 49.5822 | 121,562 | 2915.169275 | 98 | 145 | 157 | YHKNNKSWMESEF |
| Chymotrypsin | 290 | 4 | 4606 | 1 | 4052 | 3240.2806 | 49.0762 | 76,835 | 3239.272775 | 98 | 145 | 157 | YHKNNKSWMESEF |
| Chymotrypsin | 290 | 4 | 6246 | 1 | 4046 | 3402.3339 | 48.8182 | 48,546 | 3401.326075 | 98 | 145 | 157 | YHKNNKSWMESEF |
| Chymotrypsin | 290 | 4 | 6883 | 1 | 4059 | 3078.2303 | 49.371 | 39,839 | 3077.222475 | 98 | 145 | 157 | YHKNNKSWMESEF |
| Chymotrypsin | 311 | 3 | 4048 | 1 | 4033 | 2753.1112 | 48.2607 | 90,602 | 2752.103375 | 370 | 146 | 157 | HKNNKSWMESEF |
| Chymotrypsin | 311 | 3 | 5416 | 1 | 4019 | 3077.2225 | 47.6659 | 61,576 | 3076.214675 | 370 | 146 | 157 | HKNNKSWMESEF |
| Chymotrypsin | 311 | 3 | 6395 | 1 | 4026 | 2915.1652 | 47.9623 | 46,585 | 2914.157375 | 370 | 146 | 157 | HKNNKSWMESEF |
| alpha-Lytic | 81 | 10 | 566 | 1 | 4111 | 3949.6109 | 41.6158 | 234,936 | 3948.603075 | 1023 | 143 | 155 | VYYHKNNKSWMES |
| alpha-Lytic | 81 | 10 | 640 | 1 | 4110 | 3746.5289 | 41.5698 | 211,107 | 3745.521075 | 1023 | 143 | 155 | VYYHKNNKSWMES |
| alpha-Lytic | 81 | 10 | 1391 | 1 | 4112 | 3787.5578 | 41.6615 | 92,985 | 3786.549975 | 1023 | 143 | 155 | VYYHKNNKSWMES |
| alpha-Lytic | 81 | 10 | 2681 | 1 | 4116 | 3600.4779 | 41.8648 | 37,897 | 3599.470075 | 1023 | 143 | 155 | VYYHKNNKSWMES |
| alpha-Lytic | 81 | 10 | 2924 | 1 | 4122 | 3438.4201 | 42.1679 | 31,708 | 3437.412275 | 1023 | 143 | 155 | VYYHKNNKSWMES |
| alpha-Lytic | 81 | 10 | 3017 | 1 | 4121 | 3454.4127 | 42.1172 | 29,637 | 3453.404875 | 1023 | 143 | 155 | VYYHKNNKSWMES |
| alpha-Lytic | 81 | 10 | 3184 | 1 | 4119 | 3803.5541 | 42.0164 | 26,076 | 3802.546275 | 1023 | 143 | 155 | VYYHKNNKSWMES |
| alpha-Lytic | 81 | 10 | 3242 | 1 | 4124 | 4022.6325 | 42.269 | 24,743 | 4021.624675 | 1023 | 143 | 155 | VYYHKNNKSWMES |
| alpha-Lytic | 81 | 10 | 3416 | 1 | 4126 | 3657.4892 | 42.3524 | 20,711 | 3656.481375 | 1023 | 143 | 155 | VYYHKNNKSWMES |
| alpha-Lytic | 81 | 10 | 3574 | 1 | 4122 | 3397.3893 | 42.1679 | 17,460 | 3396.381475 | 1023 | 143 | 155 | VYYHKNNKSWMES |
| Chymotrypsin | 180 | 33 | 1008 | 1 | 4499 | 3811.493 | 67.7617 | 377,397 | 3810.485175 | 38 | 161 | 175 | SSANNCTFEYVSQPF |
| Chymotrypsin | 180 | 33 | 1176 | 1 | 4503 | 3665.4367 | 67.9176 | 324,880 | 3664.428875 | 38 | 161 | 175 | SSANNCTFEYVSQPF |
| Chymotrypsin | 180 | 33 | 1583 | 1 | 4497 | 4014.5874 | 67.678 | 244,698 | 4013.579575 | 38 | 161 | 175 | SSANNCTFEYVSQPF |
| Chymotrypsin | 180 | 33 | 1800 | 1 | 4474 | 4687.8113 | 66.6959 | 217,433 | 4686.803475 | 38 | 161 | 175 | SSANNCTFEYVSQPF |
| Chymotrypsin | 180 | 33 | 1800 | 1 | 4474 | 4687.8113 | 66.6959 | 217,433 | 4686.803475 | 38 | 161 | 175 | SSANNCTFEYVSQPF |
| Chymotrypsin | 180 | 33 | 1922 | 1 | 4470 | 4833.8683 | 66.5335 | 202,119 | 4832.860475 | 38 | 161 | 175 | SSANNCTFEYVSQPF |
| Chymotrypsin | 180 | 33 | 1922 | 1 | 4470 | 4833.8683 | 66.5335 | 202,119 | 4832.860475 | 38 | 161 | 175 | SSANNCTFEYVSQPF |
| Chymotrypsin | 180 | 33 | 1967 | 1 | 4514 | 3478.3518 | 68.3876 | 197,520 | 3477.343975 | 38 | 161 | 175 | SSANNCTFEYVSQPF |
| Chymotrypsin | 180 | 33 | 2129 | 1 | 4514 | 3722.4624 | 68.3876 | 184,451 | 3721.454575 | 38 | 161 | 175 | SSANNCTFEYVSQPF |
| Chymotrypsin | 180 | 33 | 2221 | 1 | 4511 | 3519.3786 | 68.2594 | 176,367 | 3518.370775 | 38 | 161 | 175 | SSANNCTFEYVSQPF |
| Chymotrypsin | 180 | 33 | 2853 | 1 | 4505 | 3868.5186 | 68.002 | 134,081 | 3867.510775 | 38 | 161 | 175 | SSANNCTFEYVSQPF |
| Chymotrypsin | 180 | 33 | 2876 | 1 | 4490 | 4176.6079 | 67.3761 | 132,880 | 4175.600075 | 38 | 161 | 175 | SSANNCTFEYVSQPF |
| Chymotrypsin | 180 | 33 | 2978 | 1 | 4486 | 4249.6351 | 67.2075 | 128,467 | 4248.627275 | 38 | 161 | 175 | SSANNCTFEYVSQPF |
| Chymotrypsin | 180 | 33 | 3233 | 1 | 4414 | 3253.3088 | 64.2176 | 116,578 | 3252.300975 | 38 | 161 | 175 | SSANNCTFEYVSQPF |
| Chymotrypsin | 180 | 33 | 3264 | 1 | 4484 | 4525.768 | 67.1215 | 115,364 | 4524.760175 | 38 | 161 | 175 | SSANNCTFEYVSQPF |
| Chymotrypsin | 180 | 33 | 3358 | 1 | 4520 | 3316.3015 | 68.6398 | 111,954 | 3315.293675 | 38 | 161 | 175 | SSANNCTFEYVSQPF |
| Chymotrypsin | 180 | 33 | 3367 | 1 | 4479 | 4598.7835 | 66.9086 | 111,578 | 4597.775675 | 38 | 161 | 175 | SSANNCTFEYVSQPF |
| Chymotrypsin | 180 | 33 | 3375 | 1 | 4493 | 4030.571 | 67.506 | 111,288 | 4029.563175 | 38 | 161 | 175 | SSANNCTFEYVSQPF |
| Chymotrypsin | 180 | 33 | 3603 | 1 | 4517 | 3373.3191 | 68.5146 | 103,638 | 3372.311275 | 38 | 161 | 175 | SSANNCTFEYVSQPF |
| Chymotrypsin | 180 | 33 | 3655 | 1 | 4518 | 3576.3999 | 68.5567 | 101,760 | 3575.392075 | 38 | 161 | 175 | SSANNCTFEYVSQPF |
| Chymotrypsin | 180 | 33 | 3802 | 1 | 4483 | 4322.6957 | 67.0789 | 97,280 | 4321.687875 | 38 | 161 | 175 | SSANNCTFEYVSQPF |
| Chymotrypsin | 180 | 33 | 3817 | 1 | 4479 | 4541.7462 | 66.9086 | 96,992 | 4540.738375 | 38 | 161 | 175 | SSANNCTFEYVSQPF |
| Chymotrypsin | 180 | 33 | 3954 | 1 | 4472 | 4890.9143 | 66.6111 | 93,062 | 4889.906475 | 38 | 161 | 175 | SSANNCTFEYVSQPF |
| Chymotrypsin | 180 | 33 | 4647 | 1 | 4487 | 4379.7081 | 67.2489 | 76,059 | 4378.700275 | 38 | 161 | 175 | SSANNCTFEYVSQPF |
| Chymotrypsin | 180 | 33 | 4713 | 1 | 4501 | 4087.5831 | 67.847 | 74,494 | 4086.575275 | 38 | 161 | 175 | SSANNCTFEYVSQPF |
| Chymotrypsin | 180 | 33 | 5437 | 1 | 4474 | 4744.8581 | 66.6959 | 61,091 | 4743.850275 | 38 | 161 | 175 | SSANNCTFEYVSQPF |
| Chymotrypsin | 180 | 33 | 5601 | 1 | 4482 | 4395.6888 | 67.0353 | 58,519 | 4394.680975 | 38 | 161 | 175 | SSANNCTFEYVSQPF |
| Chymotrypsin | 180 | 33 | 6560 | 1 | 4418 | 3091.2557 | 64.3742 | 44,274 | 3090.247875 | 38 | 161 | 175 | SSANNCTFEYVSQPF |
| Chymotrypsin | 180 | 33 | 6612 | 1 | 4486 | 4452.7302 | 67.2075 | 43,673 | 4451.722375 | 38 | 161 | 175 | SSANNCTFEYVSQPF |
| Chymotrypsin | 180 | 33 | 6631 | 1 | 4470 | 5036.9513 | 66.5335 | 43,320 | 5035.943475 | 38 | 161 | 175 | SSANNCTFEYVSQPF |
| Chymotrypsin | 180 | 33 | 6687 | 1 | 4519 | 3560.4067 | 68.598 | 42,653 | 3559.398875 | 38 | 161 | 175 | SSANNCTFEYVSQPF |
| Chymotrypsin | 180 | 33 | 7131 | 1 | 4565 | 4848.8485 | 70.5583 | 36,163 | 4847.840675 | 38 | 161 | 175 | SSANNCTFEYVSQPF |
| Chymotrypsin | 180 | 33 | 7131 | 1 | 4565 | 4848.8485 | 70.5583 | 36,163 | 4847.840675 | 38 | 161 | 175 | SSANNCTFEYVSQPF |
| Chymotrypsin | 180 | 33 | 7730 | 1 | 4413 | 3050.2205 | 64.1758 | 28,583 | 3049.212675 | 38 | 161 | 175 | SSANNCTFEYVSQPF |
| Chymotrypsin | 180 | 33 | 7775 | 1 | 4499 | 3852.535 | 67.7617 | 27,875 | 3851.527175 | 38 | 161 | 175 | SSANNCTFEYVSQPF |
| Chymotrypsin | 180 | 33 | 7907 | 1 | 4520 | 3332.3064 | 68.6398 | 26,182 | 3331.298575 | 38 | 161 | 175 | SSANNCTFEYVSQPF |
| Chymotrypsin | 295 | 6 | 3355 | 1 | 4028 | 3107.1555 | 48.0463 | 112,019 | 3106.147675 | 252 | 161 | 170 | SSANNCTFEY |
| Chymotrypsin | 295 | 6 | 3384 | 1 | 4024 | 3253.2103 | 47.8766 | 111,066 | 3252.202475 | 252 | 161 | 170 | SSANNCTFEY |
| Chymotrypsin | 295 | 6 | 4579 | 1 | 4026 | 3456.2915 | 47.9623 | 77,341 | 3455.283675 | 252 | 161 | 170 | SSANNCTFEY |
| Chymotrypsin | 295 | 6 | 5121 | 1 | 4037 | 3310.2334 | 48.4341 | 66,281 | 3309.225575 | 252 | 161 | 170 | SSANNCTFEY |
| Chymotrypsin | 295 | 6 | 5587 | 1 | 4052 | 3164.1745 | 49.0762 | 58,763 | 3163.166675 | 252 | 161 | 170 | SSANNCTFEY |
| Chymotrypsin | 295 | 6 | 7513 | 1 | 4052 | 3018.1142 | 49.0762 | 31,490 | 3017.106375 | 252 | 161 | 170 | SSANNCTFEY |
| Trypsin | 80 | 16 | 564 | 1 | 4799 | 4177.7395 | 80.6607 | 253,749 | 4176.731675 | 13 | 159 | 182 | VYSSANNCTFEYVSQPFLMDLEGK |
| Trypsin | 80 | 16 | 691 | 1 | 4788 | 4713.9591 | 80.1059 | 204,480 | 4712.951275 | 13 | 159 | 182 | VYSSANNCTFEYVSQPFLMDLEGK |
| Trypsin | 80 | 16 | 1158 | 1 | 4794 | 4526.8703 | 80.392 | 119,967 | 4525.862475 | 13 | 159 | 182 | VYSSANNCTFEYVSQPFLMDLEGK |
| Trypsin | 80 | 16 | 1199 | 1 | 4786 | 4860.0283 | 79.996 | 115,455 | 4859.020475 | 13 | 159 | 182 | VYSSANNCTFEYVSQPFLMDLEGK |
| Trypsin | 80 | 16 | 1400 | 1 | 4797 | 4364.8322 | 80.5512 | 97,632 | 4363.824375 | 13 | 159 | 182 | VYSSANNCTFEYVSQPFLMDLEGK |
| Trypsin | 80 | 16 | 1602 | 1 | 4792 | 4567.9061 | 80.307 | 84,797 | 4566.898275 | 13 | 159 | 182 | VYSSANNCTFEYVSQPFLMDLEGK |
| Trypsin | 80 | 16 | 1836 | 1 | 4805 | 4015.6856 | 80.9368 | 72,514 | 4014.677775 | 13 | 159 | 182 | VYSSANNCTFEYVSQPFLMDLEGK |
| Trypsin | 80 | 16 | 2298 | 1 | 4792 | 4770.9986 | 80.307 | 55,394 | 4769.990775 | 13 | 159 | 182 | VYSSANNCTFEYVSQPFLMDLEGK |
| Trypsin | 80 | 16 | 2334 | 1 | 4785 | 5063.109 | 79.9434 | 54,447 | 5062.101175 | 13 | 159 | 182 | VYSSANNCTFEYVSQPFLMDLEGK |
| Trypsin | 80 | 16 | 2770 | 1 | 4795 | 4421.8491 | 80.4452 | 43,898 | 4420.841275 | 13 | 159 | 182 | VYSSANNCTFEYVSQPFLMDLEGK |
| Trypsin | 80 | 16 | 3165 | 1 | 4798 | 4380.8181 | 80.6094 | 36,759 | 4379.810275 | 13 | 159 | 182 | VYSSANNCTFEYVSQPFLMDLEGK |
| Trypsin | 80 | 16 | 3171 | 1 | 4795 | 4624.9207 | 80.4452 | 36,686 | 4623.912875 | 13 | 159 | 182 | VYSSANNCTFEYVSQPFLMDLEGK |
| Trypsin | 80 | 16 | 3270 | 1 | 4871 | 4874.9966 | 84.2268 | 35,054 | 4873.988775 | 13 | 159 | 182 | VYSSANNCTFEYVSQPFLMDLEGK |
| Trypsin | 80 | 16 | 3270 | 1 | 4871 | 4874.9966 | 84.2268 | 35,054 | 4873.988775 | 13 | 159 | 182 | VYSSANNCTFEYVSQPFLMDLEGK |
| Trypsin | 80 | 16 | 3996 | 1 | 4801 | 4202.7605 | 80.7578 | 26,241 | 4201.752675 | 13 | 159 | 182 | VYSSANNCTFEYVSQPFLMDLEGK |
| Trypsin | 80 | 16 | 5099 | 1 | 4879 | 4687.9193 | 84.5756 | 16,908 | 4686.911475 | 13 | 159 | 182 | VYSSANNCTFEYVSQPFLMDLEGK |
| Trypsin | 80 | 16 | 5770 | 1 | 4876 | 4728.9195 | 84.4399 | 12,511 | 4727.911675 | 13 | 159 | 182 | VYSSANNCTFEYVSQPFLMDLEGK |
| Trypsin | 173 | 5 | 3235 | 1 | 4756 | 4542.884 | 78.5045 | 35,710 | 4541.876175 | 70 | 159 | 182 | VYSSANNCTFEYVSQPFLMDLEGK |
| Trypsin | 173 | 5 | 3258 | 1 | 4752 | 4729.9468 | 78.3334 | 35,250 | 4728.938975 | 70 | 159 | 182 | VYSSANNCTFEYVSQPFLMDLEGK |
| Trypsin | 173 | 5 | 3948 | 1 | 4761 | 4380.8181 | 78.7474 | 26,825 | 4379.810275 | 70 | 159 | 182 | VYSSANNCTFEYVSQPFLMDLEGK |
| Trypsin | 173 | 5 | 5095 | 1 | 4750 | 4933.0331 | 78.24 | 16,937 | 4932.025275 | 70 | 159 | 182 | VYSSANNCTFEYVSQPFLMDLEGK |
| Trypsin | 173 | 5 | 5535 | 1 | 4755 | 4583.8985 | 78.4627 | 13,918 | 4582.890675 | 70 | 159 | 182 | VYSSANNCTFEYVSQPFLMDLEGK |
| Chymotrypsin | 7 | 6 | 14 | 1 | 5118 | 3670.6991 | 92.9655 | 11,373,300 | 3669.691275 | 8 | 221 | 238 | SALEPLVDLPIGINITRF |
| Chymotrypsin | 7 | 6 | 66 | 1 | 5125 | 3508.6454 | 93.2607 | 4,164,350 | 3507.637575 | 8 | 221 | 238 | SALEPLVDLPIGINITRF |
| Chymotrypsin | 7 | 6 | 94 | 1 | 5115 | 3832.7468 | 92.8598 | 3,151,610 | 3831.738975 | 8 | 221 | 238 | SALEPLVDLPIGINITRF |
| Chymotrypsin | 7 | 6 | 779 | 1 | 5129 | 3346.5928 | 93.4192 | 475,819 | 3345.584975 | 8 | 221 | 238 | SALEPLVDLPIGINITRF |
| Chymotrypsin | 7 | 6 | 961 | 1 | 5125 | 3549.6749 | 93.2607 | 397,843 | 3548.667075 | 8 | 221 | 238 | SALEPLVDLPIGINITRF |
| Chymotrypsin | 7 | 6 | 7614 | 1 | 5219 | 3856.7574 | 97.1653 | 30,121 | 3855.749575 | 8 | 221 | 238 | SALEPLVDLPIGINITRF |
| Chymotrypsin | 100 | 3 | 408 | 1 | 4837 | 3060.3727 | 81.5915 | 852,910 | 3059.364875 | 241 | 227 | 238 | VDLPIGINITRF |
| Chymotrypsin | 100 | 3 | 1722 | 1 | 4841 | 2898.3158 | 81.7617 | 225,905 | 2897.307975 | 241 | 227 | 238 | VDLPIGINITRF |
| Chymotrypsin | 100 | 3 | 3685 | 1 | 4834 | 3222.4185 | 81.467 | 100,806 | 3221.410675 | 241 | 227 | 238 | VDLPIGINITRF |
| Chymotrypsin | 182 | 15 | 1027 | 1 | 5104 | 3736.7578 | 92.4577 | 369,179 | 3735.749975 | 8 | 221 | 238 | SALEPLVDLPIGINITRF |
| Chymotrypsin | 182 | 15 | 1480 | 1 | 5095 | 4101.8899 | 92.0769 | 260,655 | 4100.882075 | 8 | 221 | 238 | SALEPLVDLPIGINITRF |
| Chymotrypsin | 182 | 15 | 1552 | 1 | 5121 | 3590.6986 | 93.091 | 249,578 | 3589.690775 | 8 | 221 | 238 | SALEPLVDLPIGINITRF |
| Chymotrypsin | 182 | 15 | 1776 | 1 | 5111 | 3955.8222 | 92.6993 | 219,629 | 3954.814375 | 8 | 221 | 238 | SALEPLVDLPIGINITRF |
| Chymotrypsin | 182 | 15 | 1936 | 1 | 5098 | 4142.9007 | 92.2041 | 200,809 | 4141.892875 | 8 | 221 | 238 | SALEPLVDLPIGINITRF |
| Chymotrypsin | 182 | 15 | 1968 | 1 | 5102 | 3980.8725 | 92.3719 | 197,469 | 3979.864675 | 8 | 221 | 238 | SALEPLVDLPIGINITRF |
| Chymotrypsin | 182 | 15 | 2635 | 1 | 5086 | 4467.0183 | 91.7018 | 147,160 | 4466.010475 | 8 | 221 | 238 | SALEPLVDLPIGINITRF |
| Chymotrypsin | 182 | 15 | 2856 | 1 | 5105 | 3777.7859 | 92.5019 | 133,847 | 3776.778075 | 8 | 221 | 238 | SALEPLVDLPIGINITRF |
| Chymotrypsin | 182 | 15 | 3112 | 1 | 5097 | 3939.8382 | 92.1625 | 121,575 | 3938.830375 | 8 | 221 | 238 | SALEPLVDLPIGINITRF |
| Chymotrypsin | 182 | 15 | 3166 | 1 | 5115 | 3996.8441 | 92.8598 | 119,086 | 3995.836275 | 8 | 221 | 238 | SALEPLVDLPIGINITRF |
| Chymotrypsin | 182 | 15 | 3445 | 1 | 5104 | 4320.9723 | 92.4577 | 108,660 | 4319.964475 | 8 | 221 | 238 | SALEPLVDLPIGINITRF |
| Chymotrypsin | 182 | 15 | 3687 | 1 | 5089 | 4304.9626 | 91.8323 | 100,703 | 4303.954775 | 8 | 221 | 238 | SALEPLVDLPIGINITRF |
| Chymotrypsin | 182 | 15 | 5996 | 1 | 5099 | 3882.8133 | 92.2455 | 52,042 | 3881.805475 | 8 | 221 | 238 | SALEPLVDLPIGINITRF |
| Chymotrypsin | 182 | 15 | 6215 | 1 | 5009 | 4013.8725 | 88.5463 | 48,932 | 4012.864675 | 8 | 221 | 238 | SALEPLVDLPIGINITRF |
| Chymotrypsin | 182 | 15 | 6599 | 1 | 5215 | 3897.7804 | 96.9957 | 43,808 | 3896.772575 | 8 | 221 | 238 | SALEPLVDLPIGINITRF |
| Trypsin | 13 | 17 | 36 | 1 | 5064 | 4180.9404 | 93.5623 | 3,292,110 | 4179.932575 | 28 | 215 | 237 | DLPQGFSALEPLVDLPIGINITR |
| Trypsin | 13 | 17 | 73 | 1 | 5067 | 4018.8867 | 93.7099 | 1,661,670 | 4017.878875 | 28 | 215 | 237 | DLPQGFSALEPLVDLPIGINITR |
| Trypsin | 13 | 17 | 122 | 1 | 5070 | 3856.8336 | 93.8586 | 1,020,590 | 3855.825775 | 28 | 215 | 237 | DLPQGFSALEPLVDLPIGINITR |
| Trypsin | 13 | 17 | 185 | 1 | 5062 | 4342.9918 | 93.4498 | 689,790 | 4341.983975 | 28 | 215 | 237 | DLPQGFSALEPLVDLPIGINITR |
| Trypsin | 13 | 17 | 399 | 1 | 5075 | 3694.7801 | 94.1043 | 362,485 | 3693.772275 | 28 | 215 | 237 | DLPQGFSALEPLVDLPIGINITR |
| Trypsin | 13 | 17 | 631 | 1 | 5071 | 3897.8646 | 93.9041 | 227,986 | 3896.856775 | 28 | 215 | 237 | DLPQGFSALEPLVDLPIGINITR |
| Trypsin | 13 | 17 | 716 | 1 | 5068 | 4059.9181 | 93.7589 | 196,527 | 4058.910275 | 28 | 215 | 237 | DLPQGFSALEPLVDLPIGINITR |
| Trypsin | 13 | 17 | 1343 | 1 | 5065 | 4100.9447 | 93.6129 | 102,383 | 4099.936875 | 28 | 215 | 237 | DLPQGFSALEPLVDLPIGINITR |
| Trypsin | 13 | 17 | 1633 | 1 | 5065 | 4043.9223 | 93.6129 | 83,136 | 4042.914475 | 28 | 215 | 237 | DLPQGFSALEPLVDLPIGINITR |
| Trypsin | 13 | 17 | 1795 | 1 | 5069 | 3922.8944 | 93.8089 | 74,149 | 3921.886575 | 28 | 215 | 237 | DLPQGFSALEPLVDLPIGINITR |
| Trypsin | 13 | 17 | 2229 | 1 | 5056 | 4466.0661 | 93.1501 | 57,739 | 4465.058275 | 28 | 215 | 237 | DLPQGFSALEPLVDLPIGINITR |
| Trypsin | 13 | 17 | 2276 | 1 | 5063 | 4084.9409 | 93.5059 | 56,200 | 4083.933075 | 28 | 215 | 237 | DLPQGFSALEPLVDLPIGINITR |
| Trypsin | 13 | 17 | 2457 | 1 | 5050 | 4612.1239 | 92.8755 | 51,086 | 4611.116075 | 28 | 215 | 237 | DLPQGFSALEPLVDLPIGINITR |
| Trypsin | 13 | 17 | 2700 | 1 | 5070 | 3938.8791 | 93.8586 | 45,536 | 3937.871275 | 28 | 215 | 237 | DLPQGFSALEPLVDLPIGINITR |
| Trypsin | 13 | 17 | 3509 | 1 | 5048 | 4831.2062 | 92.7925 | 31,742 | 4830.198375 | 28 | 215 | 237 | DLPQGFSALEPLVDLPIGINITR |
| Trypsin | 13 | 17 | 4823 | 1 | 5054 | 4669.1594 | 93.0496 | 19,011 | 4668.151575 | 28 | 215 | 237 | DLPQGFSALEPLVDLPIGINITR |
| Trypsin | 13 | 17 | 5982 | 1 | 5041 | 4977.2614 | 92.4726 | 11,252 | 4976.253575 | 28 | 215 | 237 | DLPQGFSALEPLVDLPIGINITR |
| Chymotrypsin | 166 | 21 | 889 | 1 | 4810 | 5653.4299 | 80.4895 | 421,643 | 5652.422075 | 111 | 276 | 303 | LLKYNENGTITDAVDCALDPLSETKCTL |
| Chymotrypsin | 166 | 21 | 1155 | 1 | 4816 | 5288.2908 | 80.7471 | 331,403 | 5287.282975 | 111 | 276 | 303 | LLKYNENGTITDAVDCALDPLSETKCTL |
| Chymotrypsin | 166 | 21 | 1474 | 1 | 4823 | 4923.1698 | 81.0357 | 261,836 | 4922.161975 | 111 | 276 | 303 | LLKYNENGTITDAVDCALDPLSETKCTL |
| Chymotrypsin | 166 | 21 | 1573 | 1 | 4808 | 5799.487 | 80.4011 | 246,326 | 5798.479175 | 111 | 276 | 303 | LLKYNENGTITDAVDCALDPLSETKCTL |
| Chymotrypsin | 166 | 21 | 1969 | 1 | 4814 | 5434.3602 | 80.6621 | 197,430 | 5433.352375 | 111 | 276 | 303 | LLKYNENGTITDAVDCALDPLSETKCTL |
| Chymotrypsin | 166 | 21 | 2044 | 1 | 4820 | 5069.2119 | 80.915 | 189,813 | 5068.204075 | 111 | 276 | 303 | LLKYNENGTITDAVDCALDPLSETKCTL |
| Chymotrypsin | 166 | 21 | 2393 | 1 | 4806 | 5945.5314 | 80.335 | 161,946 | 5944.523575 | 111 | 276 | 303 | LLKYNENGTITDAVDCALDPLSETKCTL |
| Chymotrypsin | 166 | 21 | 2513 | 1 | 4825 | 4964.2075 | 81.1194 | 154,699 | 4963.199675 | 111 | 276 | 303 | LLKYNENGTITDAVDCALDPLSETKCTL |
| Chymotrypsin | 166 | 21 | 3084 | 1 | 4823 | 5126.2415 | 81.0357 | 123,067 | 5125.233675 | 111 | 276 | 303 | LLKYNENGTITDAVDCALDPLSETKCTL |
| Chymotrypsin | 166 | 21 | 3314 | 1 | 4805 | 6002.5879 | 80.2967 | 113,305 | 6001.580075 | 111 | 276 | 303 | LLKYNENGTITDAVDCALDPLSETKCTL |
| Chymotrypsin | 166 | 21 | 3449 | 1 | 4804 | 6091.5983 | 80.2538 | 108,594 | 6090.590475 | 111 | 276 | 303 | LLKYNENGTITDAVDCALDPLSETKCTL |
| Chymotrypsin | 166 | 21 | 3449 | 1 | 4804 | 6091.5983 | 80.2538 | 108,594 | 6090.590475 | 111 | 276 | 303 | LLKYNENGTITDAVDCALDPLSETKCTL |
| Chymotrypsin | 166 | 21 | 3752 | 1 | 4819 | 5272.3121 | 80.8724 | 98,756 | 5271.304275 | 111 | 276 | 303 | LLKYNENGTITDAVDCALDPLSETKCTL |
| Chymotrypsin | 166 | 21 | 4013 | 1 | 4814 | 5491.3954 | 80.6621 | 91,459 | 5490.387575 | 111 | 276 | 303 | LLKYNENGTITDAVDCALDPLSETKCTL |
| Chymotrypsin | 166 | 21 | 4799 | 1 | 4826 | 4777.0993 | 81.1614 | 72,872 | 4776.091475 | 111 | 276 | 303 | LLKYNENGTITDAVDCALDPLSETKCTL |
| Chymotrypsin | 166 | 21 | 4841 | 1 | 4811 | 5580.4049 | 80.5311 | 71,942 | 5579.397075 | 111 | 276 | 303 | LLKYNENGTITDAVDCALDPLSETKCTL |
| Chymotrypsin | 166 | 21 | 5235 | 1 | 4816 | 5637.4489 | 80.7471 | 64,667 | 5636.441075 | 111 | 276 | 303 | LLKYNENGTITDAVDCALDPLSETKCTL |
| Chymotrypsin | 166 | 21 | 5623 | 1 | 4829 | 4720.0777 | 81.2824 | 58,170 | 4719.069875 | 111 | 276 | 303 | LLKYNENGTITDAVDCALDPLSETKCTL |
| Chymotrypsin | 166 | 21 | 5858 | 1 | 4815 | 5329.3167 | 80.7054 | 54,211 | 5328.308875 | 111 | 276 | 303 | LLKYNENGTITDAVDCALDPLSETKCTL |
| Chymotrypsin | 166 | 21 | 5944 | 1 | 4803 | 6148.6398 | 80.212 | 52,758 | 6147.631975 | 111 | 276 | 303 | LLKYNENGTITDAVDCALDPLSETKCTL |
| Chymotrypsin | 166 | 21 | 7480 | 1 | 4877 | 5960.504 | 83.2403 | 31,946 | 5959.496175 | 111 | 276 | 303 | LLKYNENGTITDAVDCALDPLSETKCTL |
| Chymotrypsin | 166 | 21 | 7480 | 1 | 4877 | 5960.504 | 83.2403 | 31,946 | 5959.496175 | 111 | 276 | 303 | LLKYNENGTITDAVDCALDPLSETKCTL |
| Chymotrypsin | 166 | 21 | 7684 | 1 | 4876 | 5636.423 | 83.1962 | 29,428 | 5635.415175 | 111 | 276 | 303 | LLKYNENGTITDAVDCALDPLSETKCTL |
| Chymotrypsin | 166 | 21 | 7684 | 1 | 4876 | 5636.423 | 83.1962 | 29,428 | 5635.415175 | 111 | 276 | 303 | LLKYNENGTITDAVDCALDPLSETKCTL |
| Chymotrypsin | 265 | 9 | 2381 | 1 | 4877 | 5288.3063 | 83.2403 | 162,976 | 5287.298475 | 111 | 276 | 303 | LLKYNENGTITDAVDCALDPLSETKCTL |
| Chymotrypsin | 265 | 9 | 2469 | 1 | 4870 | 5653.4474 | 82.9393 | 157,227 | 5652.439575 | 111 | 276 | 303 | LLKYNENGTITDAVDCALDPLSETKCTL |
| Chymotrypsin | 265 | 9 | 3433 | 1 | 4868 | 5799.5036 | 82.8525 | 109,079 | 5798.495775 | 111 | 276 | 303 | LLKYNENGTITDAVDCALDPLSETKCTL |
| Chymotrypsin | 265 | 9 | 4169 | 1 | 4867 | 5945.5524 | 82.8103 | 87,217 | 5944.544575 | 111 | 276 | 303 | LLKYNENGTITDAVDCALDPLSETKCTL |
| Chymotrypsin | 265 | 9 | 4619 | 1 | 4885 | 4923.158 | 83.5739 | 76,625 | 4922.150175 | 111 | 276 | 303 | LLKYNENGTITDAVDCALDPLSETKCTL |
| Chymotrypsin | 265 | 9 | 5384 | 1 | 4864 | 6091.5998 | 82.6922 | 61,984 | 6090.591975 | 111 | 276 | 303 | LLKYNENGTITDAVDCALDPLSETKCTL |
| Chymotrypsin | 265 | 9 | 5384 | 1 | 4864 | 6091.5998 | 82.6922 | 61,984 | 6090.591975 | 111 | 276 | 303 | LLKYNENGTITDAVDCALDPLSETKCTL |
| Chymotrypsin | 265 | 9 | 6302 | 1 | 4876 | 5434.3622 | 83.1962 | 47,769 | 5433.354375 | 111 | 276 | 303 | LLKYNENGTITDAVDCALDPLSETKCTL |
| Chymotrypsin | 265 | 9 | 7521 | 1 | 4873 | 5580.4235 | 83.0666 | 31,351 | 5579.415675 | 111 | 276 | 303 | LLKYNENGTITDAVDCALDPLSETKCTL |
| Chymotrypsin | 265 | 9 | 8330 | 1 | 4880 | 5272.2916 | 83.359 | 21,131 | 5271.283775 | 111 | 276 | 303 | LLKYNENGTITDAVDCALDPLSETKCTL |
| Chymotrypsin | 307 | 6 | 3888 | 1 | 4778 | 5488.401 | 79.1719 | 94,951 | 5487.393175 | 231 | 276 | 304 | LLKYNENGTITDAVDCALDPLSETKCTLK |
| Chymotrypsin | 307 | 6 | 3888 | 1 | 4778 | 5488.401 | 79.1719 | 94,951 | 5487.393175 | 231 | 276 | 304 | LLKYNENGTITDAVDCALDPLSETKCTLK |
| Chymotrypsin | 307 | 6 | 4974 | 1 | 4779 | 5342.3615 | 79.2141 | 69,095 | 5341.353675 | 231 | 276 | 304 | LLKYNENGTITDAVDCALDPLSETKCTLK |
| Chymotrypsin | 307 | 6 | 4974 | 1 | 4779 | 5342.3615 | 79.2141 | 69,095 | 5341.353675 | 231 | 276 | 304 | LLKYNENGTITDAVDCALDPLSETKCTLK |
| Chymotrypsin | 307 | 6 | 6067 | 1 | 4782 | 5196.3024 | 79.3389 | 51,135 | 5195.294575 | 231 | 276 | 304 | LLKYNENGTITDAVDCALDPLSETKCTLK |
| Chymotrypsin | 307 | 6 | 6067 | 1 | 4782 | 5196.3024 | 79.3389 | 51,135 | 5195.294575 | 231 | 276 | 304 | LLKYNENGTITDAVDCALDPLSETKCTLK |
| Chymotrypsin | 307 | 6 | 6530 | 1 | 4796 | 4466.0309 | 79.9368 | 44,758 | 4465.023075 | 231 | 276 | 304 | LLKYNENGTITDAVDCALDPLSETKCTLK |
| Chymotrypsin | 307 | 6 | 6530 | 1 | 4796 | 4466.0309 | 79.9368 | 44,758 | 4465.023075 | 231 | 276 | 304 | LLKYNENGTITDAVDCALDPLSETKCTLK |
| Chymotrypsin | 307 | 6 | 7598 | 1 | 4785 | 4977.2223 | 79.468 | 30,290 | 4976.214475 | 231 | 276 | 304 | LLKYNENGTITDAVDCALDPLSETKCTLK |
| Chymotrypsin | 307 | 6 | 7598 | 1 | 4785 | 4977.2223 | 79.468 | 30,290 | 4976.214475 | 231 | 276 | 304 | LLKYNENGTITDAVDCALDPLSETKCTLK |
| Chymotrypsin | 307 | 6 | 8419 | 1 | 4788 | 4831.1642 | 79.5993 | 19,729 | 4830.156375 | 231 | 276 | 304 | LLKYNENGTITDAVDCALDPLSETKCTLK |
| Chymotrypsin | 307 | 6 | 8419 | 1 | 4788 | 4831.1642 | 79.5993 | 19,729 | 4830.156375 | 231 | 276 | 304 | LLKYNENGTITDAVDCALDPLSETKCTLK |
| Trypsin | 51 | 34 | 293 | 1 | 4637 | 4924.9996 | 72.7589 | 460,657 | 4923.991775 | 9 | 279 | 300 | YNENGTITDAVDCALDPLSETK |
| Trypsin | 51 | 34 | 337 | 1 | 4644 | 4559.8719 | 73.1226 | 413,357 | 4558.864075 | 9 | 279 | 300 | YNENGTITDAVDCALDPLSETK |
| Trypsin | 51 | 34 | 425 | 1 | 4634 | 5071.0549 | 72.6033 | 343,360 | 5070.047075 | 9 | 279 | 300 | YNENGTITDAVDCALDPLSETK |
| Trypsin | 51 | 34 | 498 | 1 | 4641 | 4705.9262 | 72.9672 | 290,525 | 4704.918375 | 9 | 279 | 300 | YNENGTITDAVDCALDPLSETK |
| Trypsin | 51 | 34 | 502 | 1 | 4651 | 4194.7408 | 73.4904 | 288,236 | 4193.732975 | 9 | 279 | 300 | YNENGTITDAVDCALDPLSETK |
| Trypsin | 51 | 34 | 623 | 1 | 4651 | 4397.8197 | 73.4904 | 231,090 | 4396.811875 | 9 | 279 | 300 | YNENGTITDAVDCALDPLSETK |
| Trypsin | 51 | 34 | 726 | 1 | 4648 | 4340.7982 | 73.3151 | 193,131 | 4339.790375 | 9 | 279 | 300 | YNENGTITDAVDCALDPLSETK |
| Trypsin | 51 | 34 | 745 | 1 | 4632 | 5217.1244 | 72.4991 | 187,871 | 5216.116575 | 9 | 279 | 300 | YNENGTITDAVDCALDPLSETK |
| Trypsin | 51 | 34 | 771 | 1 | 4655 | 4235.7638 | 73.6614 | 181,886 | 4234.755975 | 9 | 279 | 300 | YNENGTITDAVDCALDPLSETK |
| Trypsin | 51 | 34 | 1065 | 1 | 4645 | 4486.8515 | 73.1714 | 128,678 | 4485.843675 | 9 | 279 | 300 | YNENGTITDAVDCALDPLSETK |
| Trypsin | 51 | 34 | 1091 | 1 | 4630 | 5363.1778 | 72.4031 | 126,590 | 5362.169975 | 9 | 279 | 300 | YNENGTITDAVDCALDPLSETK |
| Trypsin | 51 | 34 | 1091 | 1 | 4630 | 5363.1778 | 72.4031 | 126,590 | 5362.169975 | 9 | 279 | 300 | YNENGTITDAVDCALDPLSETK |

| | | | | | | | | | | | | | | | | | | | | | | | | | | | | | | | | |
|---|---|---|---|---|---|---|---|---|---|---|---|---|---|---|---|---|---|---|---|---|---|---|---|---|---|---|---|---|---|---|---|---|
| Trypsin | 51 | 34 | 1107 | 1 | 4639 | 4851.9909 | 72.8674 | 124,678 | 4850.983075 | 9 | 279 | 300 | YNENGTITDAVDCALDPLSETK | 0.00020000000000000000000.0 | 282(NGT) | 1 | 2428.0838 | 2425.095539 | 14489701 | 79.38 | 1 | 6 | 5 | 3 | 0 | 0 | 0 | 1 | 4850.983075 | 0.0000 | 0 | -6.52 |
| Trypsin | 51 | 34 | 1376 | 1 | 4655 | 4276.7922 | 73.6614 | 99,311 | 4275.784375 | 9 | 279 | 300 | YNENGTITDAVDCALDPLSETK | 0.00020000000000000000000.0 | 282(NGT) | 1 | 2428.0838 | 2425.095539 | 14489701 | 79.38 | 1 | 3 | 6 | 1 | 0 | 0 | 0 | 1 | 4275.788158 | -0.0038 | -0.88 | -5.72 |
| Trypsin | 51 | 34 | 1394 | 1 | 4649 | 4543.8812 | 73.3574 | 98,020 | 4542.873375 | 9 | 279 | 300 | YNENGTITDAVDCALDPLSETK | 0.00020000000000000000000.0 | 282(NGT) | 1 | 2428.0838 | 2425.095539 | 14489701 | 79.38 | 1 | 5 | 5 | 2 | 0 | 0 | 0 | 1 | 4542.872342 | 0.0010 | 0.22 | -6.03 |
| Trypsin | 51 | 34 | 1429 | 1 | 4643 | 4909.0088 | 73.0741 | 95,302 | 4908.000975 | 9 | 279 | 300 | YNENGTITDAVDCALDPLSETK | 0.00020000000000000000000.0 | 282(NGT) | 1 | 2428.0838 | 2425.095539 | 14489701 | 79.38 | 1 | 6 | 6 | 2 | 0 | 0 | 0 | 1 | 4908.004539 | -0.0036 | -0.72 | -6.31 |
| Trypsin | 51 | 34 | 1466 | 1 | 4656 | 4048.6772 | 73.7193 | 92,526 | 4047.669375 | 9 | 279 | 300 | YNENGTITDAVDCALDPLSETK | 0.00020000000000000000000.0 | 282(NGT) | 1 | 2428.0838 | 2425.095539 | 14489701 | 79.38 | 1 | 5 | 4 | 0 | 0 | 0 | 0 | 1 | 4047.677151 | -0.0078 | -1.92 | -5.66 |
| Trypsin | 51 | 34 | 1706 | 1 | 4644 | 4600.9087 | 73.1226 | 78,739 | 4599.900875 | 9 | 279 | 300 | YNENGTITDAVDCALDPLSETK | 0.00020000000000000000000.0 | 282(NGT) | 1 | 2428.0838 | 2425.095539 | 14489701 | 79.38 | 1 | 5 | 6 | 1 | 0 | 0 | 0 | 1 | 4599.893806 | 0.0071 | 1.53 | -6.26 |
| Trypsin | 51 | 34 | 1771 | 1 | 4647 | 4413.8254 | 73.2665 | 75,617 | 4412.817575 | 9 | 279 | 300 | YNENGTITDAVDCALDPLSETK | 0.00020000000000000000000.0 | 282(NGT) | 1 | 2428.0838 | 2425.095539 | 14489701 | 79.38 | 1 | 6 | 5 | 0 | 0 | 0 | 0 | 1 | 4412.809348 | 0.0082 | 1.86 | -6.12 |
| Trypsin | 51 | 34 | 1977 | 1 | 4659 | 3991.6622 | 73.8634 | 66,247 | 3990.654375 | 9 | 279 | 300 | YNENGTITDAVDCALDPLSETK | 0.00020000000000000000000.0 | 282(NGT) | 1 | 2428.0838 | 2425.095539 | 14489701 | 79.38 | 1 | 5 | 3 | 1 | 0 | 0 | 0 | 1 | 3990.655687 | -0.0013 | -0.32 | -5.52 |
| Trypsin | 51 | 34 | 2055 | 1 | 4662 | 3870.6379 | 74.01 | 63,312 | 3869.630075 | 9 | 279 | 300 | YNENGTITDAVDCALDPLSETK | 0.00020000000000000000000.0 | 282(NGT) | 1 | 2428.0838 | 2425.095539 | 14489701 | 79.38 | 1 | 3 | 4 | 1 | 0 | 0 | 0 | 1 | 3869.629412 | 0.0007 | 0.17 | -5.37 |
| Trypsin | 51 | 34 | 2802 | 1 | 4645 | 4689.9323 | 73.1714 | 43,266 | 4688.924475 | 9 | 279 | 300 | YNENGTITDAVDCALDPLSETK | 0.00020000000000000000000.0 | 282(NGT) | 1 | 2428.0838 | 2425.095539 | 14489701 | 79.38 | 1 | 5 | 5 | 3 | 0 | 0 | 0 | 1 | 4688.930251 | -0.0058 | -1.23 | -6.21 |
| Trypsin | 51 | 34 | 2950 | 1 | 4665 | 3642.5299 | 74.1459 | 40,212 | 3641.522075 | 9 | 279 | 300 | YNENGTITDAVDCALDPLSETK | 0.00020000000000000000000.0 | 282(NGT) | 1 | 2428.0838 | 2425.095539 | 14489701 | 79.38 | 1 | 5 | 2 | 0 | 0 | 0 | 0 | 1 | 3641.518405 | 0.0037 | 1 | -5.24 |
| Trypsin | 51 | 34 | 3187 | 1 | 4654 | 4153.7124 | 73.6167 | 36,457 | 4152.704575 | 9 | 279 | 300 | YNENGTITDAVDCALDPLSETK | 0.00020000000000000000000.0 | 282(NGT) | 1 | 2428.0838 | 2425.095539 | 14489701 | 79.38 | 1 | 6 | 3 | 1 | 0 | 0 | 0 | 1 | 4152.708511 | -0.0039 | -0.94 | -5.77 |
| Trypsin | 51 | 34 | 3504 | 1 | 4664 | 3845.6001 | 74.0975 | 31,807 | 3844.592275 | 9 | 279 | 300 | YNENGTITDAVDCALDPLSETK | 0.00020000000000000000000.0 | 282(NGT) | 1 | 2428.0838 | 2425.095539 | 14489701 | 79.38 | 1 | 5 | 3 | 0 | 0 | 0 | 0 | 1 | 3844.597778 | -0.0055 | -1.43 | -5.29 |
| Trypsin | 51 | 34 | 3566 | 1 | 4661 | 3804.5815 | 73.9659 | 31,086 | 3803.573675 | 9 | 279 | 300 | YNENGTITDAVDCALDPLSETK | 0.00020000000000000000000.0 | 282(NGT) | 1 | 2428.0838 | 2425.095539 | 14489701 | 79.38 | 1 | 6 | 2 | 0 | 0 | 0 | 0 | 1 | 3803.571229 | 0.0024 | 0.64 | -5.42 |
| Trypsin | 51 | 34 | 3656 | 1 | 4640 | 4778.956 | 72.9166 | 29,958 | 4777.948175 | 9 | 279 | 300 | YNENGTITDAVDCALDPLSETK | 0.00020000000000000000000.0 | 282(NGT) | 1 | 2428.0838 | 2425.095539 | 14489701 | 79.38 | 1 | 7 | 6 | 0 | 0 | 0 | 0 | 1 | 4777.941545 | 0.0066 | 1.38 | -6.47 |
| Trypsin | 51 | 34 | 3824 | 1 | 4651 | 4381.8167 | 73.4904 | 28,037 | 4380.808875 | 9 | 279 | 300 | YNENGTITDAVDCALDPLSETK | 0.00020000000000000000000.0 | 282(NGT) | 1 | 2428.0838 | 2425.095539 | 14489701 | 79.38 | 1 | 4 | 5 | 2 | 0 | 0 | 0 | 1 | 4380.819518 | -0.0106 | -2.42 | -5.89 |
| Trypsin | 51 | 34 | 4271 | 1 | 4667 | 3724.575 | 74.24 | 23,391 | 3723.567175 | 9 | 279 | 300 | YNENGTITDAVDCALDPLSETK | 0.00020000000000000000000.0 | 282(NGT) | 1 | 2428.0838 | 2425.095539 | 14489701 | 79.38 | 1 | 3 | 4 | 0 | 0 | 0 | 0 | 1 | 3723.571503 | -0.0043 | -1.16 | -5.14 |
| Trypsin | 51 | 34 | 4335 | 1 | 4663 | 3927.6599 | 74.0543 | 22,842 | 3926.652075 | 9 | 279 | 300 | YNENGTITDAVDCALDPLSETK | 0.00020000000000000000000.0 | 282(NGT) | 1 | 2428.0838 | 2425.095539 | 14489701 | 79.38 | 1 | 3 | 5 | 0 | 0 | 0 | 0 | 1 | 3926.650876 | 0.0012 | 0.3 | -5.33 |
| Trypsin | 51 | 34 | 4408 | 1 | 4657 | 4032.6778 | 73.7621 | 22,212 | 4031.669975 | 9 | 279 | 300 | YNENGTITDAVDCALDPLSETK | 0.00020000000000000000000.0 | 282(NGT) | 1 | 2428.0838 | 2425.095539 | 14489701 | 79.38 | 1 | 4 | 4 | 1 | 0 | 0 | 0 | 1 | 4031.682236 | -0.0123 | -3.04 | -5.62 |
| Trypsin | 51 | 34 | 4422 | 1 | 4664 | 3829.6124 | 74.0975 | 22,111 | 3828.604575 | 9 | 279 | 300 | YNENGTITDAVDCALDPLSETK | 0.00020000000000000000000.0 | 282(NGT) | 1 | 2428.0838 | 2425.095539 | 14489701 | 79.38 | 1 | 4 | 3 | 1 | 0 | 0 | 0 | 1 | 3828.602863 | 0.0017 | 0.44 | -5.29 |
| Trypsin | 51 | 34 | 4519 | 1 | 4650 | 4438.8548 | 73.4469 | 21,254 | 4437.846975 | 9 | 279 | 300 | YNENGTITDAVDCALDPLSETK | 0.00020000000000000000000.0 | 282(NGT) | 1 | 2428.0838 | 2425.095539 | 14489701 | 79.38 | 1 | 4 | 6 | 1 | 0 | 0 | 0 | 1 | 4437.840982 | 0.0060 | 1.35 | -5.94 |
| Trypsin | 51 | 34 | 5214 | 1 | 4641 | 5055.063 | 72.9672 | 16,214 | 5054.055175 | 9 | 279 | 300 | YNENGTITDAVDCALDPLSETK | 0.00020000000000000000000.0 | 282(NGT) | 1 | 2428.0838 | 2425.095539 | 14489701 | 79.38 | 1 | 6 | 6 | 3 | 0 | 0 | 0 | 1 | 5054.062448 | -0.0073 | -1.43 | -6.42 |
| Trypsin | 51 | 34 | 5550 | 1 | 4725 | 4501.8226 | 77.0302 | 13,794 | 4500.814775 | 9 | 279 | 300 | YNENGTITDAVDCALDPLSETK | 0.00020000000000000000000.0 | 282(NGT) | 1 | 2428.0838 | 2425.095539 | 14489701 | 79.38 | 1 | 5 | 4 | 1 | 0 | 1 | 0 | 1 | 4500.825391 | -0.0106 | -2.35 | -2.35 |
| Trypsin | 51 | 34 | 5550 | 1 | 4725 | 4501.8226 | 77.0302 | 13,794 | 4500.814775 | 9 | 279 | 300 | YNENGTITDAVDCALDPLSETK | 0.00020000000000000000000.0 | 282(NGT) | 1 | 2428.0838 | 2425.095539 | 14489701 | 79.38 | 2 | 6 | 4 | 0 | 1 | 0 | 0 | 1 | 4500.825392 | -0.0106 | -2.35 | -2.35 |
| alpha-Lytic | 60 | 15 | 341 | 1 | 3977 | 3538.4128 | 35.0953 | 391,578 | 3537.404975 | 9 | 278 | 286 | KYNENGTIT | 0.000020000.0 | 282(NGT) | 1 | 1041.4864 | 1038.498139 | 23571974 | 37.86 | 1 | 7 | 6 | 1 | 0 | 0 | 0 | 1 | 3537.402054 | 0.0029 | 0.82 | -2.76 |
| alpha-Lytic | 60 | 15 | 420 | 1 | 3971 | 3684.472 | 34.8024 | 309,244 | 3683.464175 | 9 | 278 | 286 | KYNENGTIT | 0.000020000.0 | 282(NGT) | 1 | 1041.4864 | 1038.498139 | 23571974 | 37.86 | 1 | 7 | 6 | 2 | 0 | 0 | 0 | 0 | 3683.459963 | 0.0042 | 1.14 | -3.06 |
| alpha-Lytic | 60 | 15 | 461 | 1 | 3977 | 3173.2789 | 35.0953 | 275,120 | 3172.271075 | 9 | 278 | 286 | KYNENGTIT | 0.000020000.0 | 282(NGT) | 1 | 1041.4864 | 1038.498139 | 23571974 | 37.86 | 1 | 6 | 5 | 1 | 0 | 0 | 0 | 1 | 3172.269857 | 0.0012 | 0.38 | -2.76 |
| alpha-Lytic | 60 | 15 | 540 | 1 | 3990 | 2808.1486 | 35.7468 | 245,201 | 2807.140775 | 9 | 278 | 286 | KYNENGTIT | 0.000020000.0 | 282(NGT) | 1 | 1041.4864 | 1038.498139 | 23571974 | 37.86 | 1 | 5 | 4 | 1 | 0 | 0 | 0 | 1 | 2807.13766 | 0.0031 | 1.1 | -2.11 |
| alpha-Lytic | 60 | 15 | 565 | 1 | 3964 | 3830.5232 | 34.4767 | 235,077 | 3829.515375 | 9 | 278 | 286 | KYNENGTIT | 0.000020000.0 | 282(NGT) | 1 | 1041.4864 | 1038.498139 | 23571974 | 37.86 | 1 | 7 | 6 | 3 | 0 | 0 | 0 | 0 | 3829.517872 | -0.0025 | -0.65 | -3.38 |
| alpha-Lytic | 60 | 15 | 716 | 1 | 4003 | 3011.2236 | 36.3821 | 186,506 | 3010.215775 | 9 | 278 | 286 | KYNENGTIT | 0.000020000.0 | 282(NGT) | 1 | 1041.4864 | 1038.498139 | 23571974 | 37.86 | 1 | 5 | 5 | 1 | 0 | 0 | 0 | 1 | 3010.217033 | -0.0013 | -0.41 | -1.48 |
| alpha-Lytic | 60 | 15 | 841 | 1 | 3960 | 3976.5904 | 34.2839 | 156,298 | 3975.582575 | 9 | 278 | 286 | KYNENGTIT | 0.000020000.0 | 282(NGT) | 1 | 1041.4864 | 1038.498139 | 23571974 | 37.86 | 1 | 7 | 6 | 4 | 0 | 0 | 0 | 1 | 3975.575781 | 0.0068 | 1.7 | -3.58 |
| alpha-Lytic | 60 | 15 | 841 | 1 | 3960 | 3976.5904 | 34.2839 | 156,298 | 3975.582575 | 9 | 278 | 286 | KYNENGTIT | 0.000020000.0 | 282(NGT) | 1 | 1041.4864 | 1038.498139 | 23571974 | 37.86 | 2 | 2 | 10 | 0 | 2 | 0 | 1 | -1 | 3975.588351 | -0.0058 | -1.45 | -3.58 |
| alpha-Lytic | 60 | 15 | 893 | 1 | 3982 | 2954.2088 | 35.3591 | 148,158 | 2953.200975 | 9 | 278 | 286 | KYNENGTIT | 0.000020000.0 | 282(NGT) | 1 | 1041.4864 | 1038.498139 | 23571974 | 37.86 | 1 | 5 | 4 | 2 | 0 | 0 | 0 | 1 | 2953.195569 | 0.0054 | 1.83 | -2.50 |
| alpha-Lytic | 60 | 15 | 896 | 1 | 3971 | 3319.3348 | 34.8024 | 147,973 | 3318.326975 | 9 | 278 | 286 | KYNENGTIT | 0.000020000.0 | 282(NGT) | 1 | 1041.4864 | 1038.498139 | 23571974 | 37.86 | 1 | 6 | 5 | 2 | 0 | 0 | 0 | 1 | 3318.327766 | -0.0008 | -0.23 | -3.06 |
| alpha-Lytic | 60 | 15 | 908 | 1 | 4005 | 2849.172 | 36.4819 | 146,611 | 2848.164175 | 9 | 278 | 286 | KYNENGTIT | 0.000020000.0 | 282(NGT) | 1 | 1041.4864 | 1038.498139 | 23571974 | 37.86 | 1 | 4 | 5 | 1 | 0 | 0 | 0 | 1 | 2848.164209 | 0.0000 | -0.01 | -1.38 |
| alpha-Lytic | 60 | 15 | 1154 | 1 | 3976 | 3376.359 | 35.0495 | 113,865 | 3375.351175 | 9 | 278 | 286 | KYNENGTIT | 0.000020000.0 | 282(NGT) | 1 | 1041.4864 | 1038.498139 | 23571974 | 37.86 | 1 | 6 | 6 | 1 | 0 | 0 | 0 | 1 | 3375.34923 | 0.0019 | 0.57 | -2.81 |
| alpha-Lytic | 60 | 15 | 1542 | 1 | 3991 | 3157.2824 | 35.8043 | 84,038 | 3156.274575 | 9 | 278 | 286 | KYNENGTIT | 0.000020000.0 | 282(NGT) | 1 | 1041.4864 | 1038.498139 | 23571974 | 37.86 | 1 | 5 | 5 | 2 | 0 | 0 | 0 | 1 | 3156.274942 | -0.0004 | -0.11 | -2.06 |
| alpha-Lytic | 60 | 15 | 1799 | 1 | 3971 | 3100.2597 | 34.8024 | 67,907 | 3099.251875 | 9 | 278 | 286 | KYNENGTIT | 0.000020000.0 | 282(NGT) | 1 | 1041.4864 | 1038.498139 | 23571974 | 37.86 | 1 | 5 | 4 | 3 | 0 | 0 | 0 | 1 | 3099.253478 | -0.0016 | -0.51 | -3.06 |
| alpha-Lytic | 60 | 15 | 2114 | 1 | 3991 | 3173.2778 | 35.8043 | 55,235 | 3172.269975 | 9 | 278 | 286 | KYNENGTIT | 0.000020000.0 | 282(NGT) | 1 | 1041.4864 | 1038.498139 | 23571974 | 37.86 | 1 | 6 | 5 | 1 | 0 | 0 | 0 | 1 | 3172.269857 | 0.0001 | 0.03 | -2.06 |
| alpha-Lytic | 60 | 15 | 2165 | 1 | 3977 | 3214.3022 | 35.0953 | 53,460 | 3213.294375 | 9 | 278 | 286 | KYNENGTIT | 0.000020000.0 | 282(NGT) | 1 | 1041.4864 | 1038.498139 | 23571974 | 37.86 | 1 | 5 | 6 | 1 | 0 | 0 | 0 | 1 | 3213.296406 | -0.0020 | -0.63 | -2.76 |
| alpha-Lytic | 83 | 12 | 572 | 1 | 4320 | 3764.5752 | 51.9218 | 232,243 | 3763.567375 | 45 | 276 | 286 | LLKYNENGTIT | 0.00000020000.0 | 282(NGT) | 1 | 1267.6546 | 1264.666339 | 2528809 | 55.99 | 1 | 7 | 6 | 1 | 0 | 0 | 0 | 1 | 3763.570254 | -0.0029 | -0.76 | -4.07 |
| alpha-Lytic | 83 | 12 | 857 | 1 | 4317 | 3910.6329 | 51.7747 | 153,187 | 3909.625075 | 45 | 276 | 286 | LLKYNENGTIT | 0.00000020000.0 | 282(NGT) | 1 | 1267.6546 | 1264.666339 | 2528809 | 55.99 | 1 | 7 | 6 | 2 | 0 | 0 | 0 | 0 | 3909.628163 | -0.0031 | -0.78 | -4.22 |
| alpha-Lytic | 83 | 12 | 972 | 1 | 4314 | 4056.6954 | 51.6274 | 136,918 | 4055.687575 | 45 | 276 | 286 | LLKYNENGTIT | 0.00000020000.0 | 282(NGT) | 1 | 1267.6546 | 1264.666339 | 2528809 | 55.99 | 1 | 7 | 6 | 3 | 0 | 0 | 0 | 0 | 4055.686072 | 0.0015 | 0.37 | -4.37 |
| alpha-Lytic | 83 | 12 | 1127 | 1 | 4325 | 3399.4457 | 52.1641 | 116,819 | 3398.437875 | 45 | 276 | 286 | LLKYNENGTIT | 0.00000020000.0 | 282(NGT) | 1 | 1267.6546 | 1264.666339 | 2528809 | 55.99 | 1 | 6 | 5 | 1 | 0 | 0 | 0 | 1 | 3398.438057 | -0.0002 | -0.05 | -3.83 |
| alpha-Lytic | 83 | 12 | 1477 | 1 | 4312 | 4202.7513 | 51.5289 | 87,411 | 4201.743475 | 45 | 276 | 286 | LLKYNENGTIT | 0.00000020000.0 | 282(NGT) | 1 | 1267.6546 | 1264.666339 | 2528809 | 55.99 | 1 | 7 | 6 | 4 | 0 | 0 | 0 | 1 | 4201.743981 | -0.0005 | -0.12 | -4.47 |
| alpha-Lytic | 83 | 12 | 1477 | 1 | 4312 | 4202.7513 | 51.5289 | 87,411 | 4201.743475 | 45 | 276 | 286 | LLKYNENGTIT | 0.00000020000.0 | 282(NGT) | 1 | 1267.6546 | 1264.666339 | 2528809 | 55.99 | 2 | 2 | 10 | 0 | 2 | 0 | 1 | -1 | 4201.756551 | -0.0131 | -3.11 | -4.47 |
| alpha-Lytic | 83 | 12 | 1677 | 1 | 4309 | 4113.7159 | 51.3831 | 74,488 | 4112.708075 | 45 | 276 | 286 | LLKYNENGTIT | 0.00000020000.0 | 282(NGT) | 1 | 1267.6546 | 1264.666339 | 2528809 | 55.99 | 1 | 7 | 7 | 2 | 0 | 0 | 0 | 0 | 4112.707536 | 0.0005 | 0.13 | -4.61 |
| alpha-Lytic | 83 | 12 | 2239 | 1 | 4333 | 3180.3786 | 52.5613 | 51,311 | 3179.370775 | 45 | 276 | 286 | LLKYNENGTIT | 0.00000020000.0 | 282(NGT) | 1 | 1267.6546 | 1264.666339 | 2528809 | 55.99 | 1 | 5 | 4 | 2 | 0 | 0 | 0 | 1 | 3179.363769 | 0.0070 | 2.2 | -3.43 |
| alpha-Lytic | 83 | 12 | 2302 | 1 | 4319 | 3691.5685 | 51.8712 | 48,880 | 3690.560675 | 45 | 276 | 286 | LLKYNENGTIT | 0.00000020000.0 | 282(NGT) | 1 | 1267.6546 | 1264.666339 | 2528809 | 55.99 | 1 | 6 | 5 | 3 | 0 | 0 | 0 | 1 | 3690.553875 | 0.0068 | 1.84 | -4.12 |
| alpha-Lytic | 83 | 12 | 2382 | 1 | 4326 | 3326.4309 | 52.2119 | 46,019 | 3325.423075 | 45 | 276 | 286 | LLKYNENGTIT | 0.00000020000.0 | 282(NGT) | 1 | 1267.6546 | 1264.666339 | 2528809 | 55.99 | 1 | 5 | 4 | 3 | 0 | 0 | 0 | 1 | 3325.421678 | 0.0014 | 0.42 | -3.78 |
| alpha-Lytic | 83 | 12 | 2440 | 1 | 4324 | 3253.3879 | 52.1133 | 44,524 | 3252.380075 | 45 | 276 | 286 | LLKYNENGTIT | 0.00000020000.0 | 282(NGT) | 1 | 1267.6546 | 1264.666339 | 2528809 | 55.99 | 1 | 6 | 5 | 0 | 0 | 0 | 0 | 1 | 3252.380148 | -0.0001 | -0.02 | -3.88 |
| alpha-Lytic | 83 | 12 | 2462 | 1 | 4312 | 3967.6523 | 51.5289 | 43,650 | 3966.644475 | 45 | 276 | 286 | LLKYNENGTIT | 0.00000020000.0 | 282(NGT) | 1 | 1267.6546 | 1264.666339 | 2528809 | 55.99 | 1 | 7 | 7 | 1 | 0 | 0 | 0 | 1 | 3966.649627 | -0.0052 | -1.29 | -4.47 |
| alpha-Lytic | 83 | 12 | 2837 | 1 | 4305 | 4259.7707 | 51.1756 | 33,634 | 4258.762875 | 45 | 276 | 286 | LLKYNENGTIT | 0.00000020000.0 | 282(NGT) | 1 | 1267.6546 | 1264.666339 | 2528809 | 55.99 | 1 | 7 | 7 | 3 | 0 | 0 | 0 | 0 | 4258.765445 | -0.0026 | -0.6 | -4.82 |
| Chymotrypsin | 114 | 36 | 495 | 1 | 4986 | 4881.207 | 87.5865 | 721,026 | 4880.199175 | 9 | 319 | 342 | RVQPTESIVRFPNITNLCPFGEVF | 0.000000000000200000000000.0 | 331(NIT) | 1 | 2822.4312 | 2819.442939 | 76519264 | 93.02 | 1 | 5 | 4 | 3 | 0 | 0 | 0 | 1 | 4880.198278 | 0.0009 | 0.18 | -5.43 |
| Chymotrypsin | 114 | 36 | 849 | 1 | 4986 | 5084.2824 | 87.5865 | 440,524 | 5083.274575 | 9 | 319 | 342 | RVQPTESIVRFPNITNLCPFGEVF | 0.000000000000200000000000.0 | 331(NIT) | 1 | 2822.4312 | 2819.442939 | 76519264 | 93.02 | 1 | 5 | 5 | 3 | 0 | 0 | 0 | 1 | 5083.277651 | -0.0031 | -0.6 | -5.43 |
| Chymotrypsin | 114 | 36 | 1147 | 1 | 4990 | 4735.1495 | 87.7602 | 332,705 | 4734.141675 | 9 | 319 | 342 | RVQPTESIVRFPNITNLCPFGEVF | 0.000000000000200000000000.0 | 331(NIT) | 1 | 2822.4312 | 2819.442939 | 76519264 | 93.02 | 1 | 5 | 4 | 2 | 0 | 0 | 0 | 1 | 4734.140369 | 0.0013 | 0.27 | -5.26 |
| Chymotrypsin | 114 | 36 | 1251 | 1 | 4993 | 4792.1709 | 87.8855 | 306,119 | 4791.163075 | 9 | 319 | 342 | RVQPTESIVRFPNITNLCPFGEVF | 0.000000000000200000000000.0 | 331(NIT) | 1 | 2822.4312 | 2819.442939 | 76519264 | 93.02 | 1 | 5 | 5 | 1 | 0 | 0 | 0 | 1 | 4791.161833 | 0.0012 | 0.25 | -5.13 |
| Chymotrypsin | 114 | 36 | 1384 | 1 | 4991 | 4938.2279 | 87.802 | 278,330 | 4937.220075 | 9 | 319 | 342 | RVQPTESIVRFPNITNLCPFGEVF | 0.000000000000200000000000.0 | 331(NIT) | 1 | 2822.4312 | 2819.442939 | 76519264 | 93.02 | 1 | 5 | 5 | 2 | 0 | 0 | 0 | 1 | 4937.219742 | 0.0003 | 0.06 | -5.21 |
| Chymotrypsin | 114 | 36 | 1494 | 1 | 4974 | 5757.5234 | 87.0755 | 258,260 | 5756.515575 | 9 | 319 | 342 | RVQPTESIVRFPNITNLCPFGEVF | 0.000000000000200000000000.0 | 331(NIT) | 1 | 2822.4312 | 2819.442939 | 76519264 | 93.02 | 1 | 7 | 6 | 4 | 0 | 0 | 0 | 1 | 5756.520581 | -0.0050 | -0.86 | -5.94 |
| Chymotrypsin | 114 | 36 | 1494 | 1 | 4974 | 5757.5234 | 87.0755 | 258,260 | 5756.515575 | 9 | 319 | 342 | RVQPTESIVRFPNITNLCPFGEVF | 0.000000000000200000000000.0 | 331(NIT) | 1 | 2822.4312 | 2819.442939 | 76519264 | 93.02 | 2 | 2 | 10 | 0 | 2 | 0 | 1 | -1 | 5756.533151 | -0.0176 | -3.05 | -5.94 |
| Chymotrypsin | 114 | 36 | 1577 | 1 | 5077 | 4896.1823 | 91.3267 | 245,146 | 4895.174475 | 9 | 319 | 342 | RVQPTESIVRFPNITNLCPFGEVF | 0.000000000000200000000000.0 | 331(NIT) | 1 | 2822.4312 | 2819.442939 | 76519264 | 93.02 | 1 | 6 | 4 | 0 | 1 | 0 | 0 | 1 | 4895.172792 | 0.0017 | 0.34 | -1.69 |
| Chymotrypsin | 114 | 36 | 1577 | 1 | 5077 | 4896.1823 | 91.3267 | 245,146 | 4895.174475 | 9 | 319 | 342 | RVQPTESIVRFPNITNLCPFGEVF | 0.000000000000200000000000.0 | 331(NIT) | 1 | 2822.4312 | 2819.442939 | 76519264 | 93.02 | 2 | 5 | 4 | 1 | 0 | 1 | 0 | 1 | 4895.172791 | 0.0017 | 0.34 | -1.69 |
| Chymotrypsin | 114 | 36 | 1638 | 1 | 4977 | 5595.4751 | 87.2014 | 238,297 | 5594.467275 | 9 | 319 | 342 | RVQPTESIVRFPNITNLCPFGEVF | 0.000000000000200000000000.0 | 331(NIT) | 1 | 2822.4312 | 2819.442939 | 76519264 | 93.02 | 1 | 6 | 6 | 4 | 0 | 0 | 0 | 0 | 5594.467757 | -0.0005 | -0.08 | -5.81 |
| Chymotrypsin | 114 | 36 | 1685 | 1 | 4993 | 4589.0892 | 87.8855 | 231,757 | 4588.081375 | 9 | 319 | 342 | RVQPTESIVRFPNITNLCPFGEVF | 0.000000000000200000000000.0 | 331(NIT) | 1 | 2822.4312 | 2819.442939 | 76519264 | 93.02 | 1 | 5 | 4 | 1 | 0 | 0 | 0 | 1 | 4588.08246 | -0.0011 | -0.23 | -5.13 |
| Chymotrypsin | 114 | 36 | 1939 | 1 | 5073 | 5042.2511 | 91.1547 | 200,426 | 5041.243275 | 9 | 319 | 342 | RVQPTESIVRFPNITNLCPFGEVF | 0.000000000000200000000000.0 | 331(NIT) | 1 | 2822.4312 | 2819.442939 | 76519264 | 93.02 | 1 | 6 | 4 | 1 | 1 | 0 | 0 | 1 | 5041.230701 | 0.0126 | 2.49 | -1.86 |
| Chymotrypsin | 114 | 36 | 1939 | 1 | 5073 | 5042.2511 | 91.1547 | 200,426 | 5041.243275 | 9 | 319 | 342 | RVQPTESIVRFPNITNLCPFGEVF | 0.000000000000200000000000.0 | 331(NIT) | 1 | 2822.4312 | 2819.442939 | 76519264 | 93.02 | 2 | 5 | 4 | 2 | 0 | 1 | 0 | 1 | 5041.2307 | 0.0126 | 2.49 | -1.86 |
| Chymotrypsin | 114 | 36 | 2501 | 1 | 4985 | 5303.369 | 87.5423 | 155,240 | 5302.361175 | 9 | 319 | 342 | RVQPTESIVRFPNITNLCPFGEVF | 0.000000000000200000000000.0 | 331(NIT) | 1 | 2822.4312 | 2819.442939 | 76519264 | 93.02 | 1 | 6 | 6 | 2 | 0 | 0 | 0 | 1 | 5302.351939 | 0.0092 | 1.74 | -5.47 |
| Chymotrypsin | 114 | 36 | 2629 | 1 | 4985 | 4954.2369 | 87.5423 | 147,354 | 4953.229075 | 9 | 319 | 342 | RVQPTESIVRFPNITNLCPFGEVF | 0.000000000000200000000000.0 | 331(NIT) | 1 | 2822.4312 | 2819.442939 | 76519264 | 93.02 | 1 | 6 | 5 | 1 | 0 | 0 | 0 | 1 | 4953.214657 | 0.0144 | 2.91 | -5.47 |
| Chymotrypsin | 114 | 36 | 2645 | 1 | 4980 | 5449.4255 | 87.3293 | 146,198 | 5448.417675 | 9 | 319 | 342 | RVQPTESIVRFPNITNLCPFGEVF | 0.000000000000200000000000.0 | 331(NIT) | 1 | 2822.4312 | 2819.442939 | 76519264 | 93.02 | 1 | 6 | 6 | 3 | 0 | 0 | 0 | 1 | 5448.409848 | 0.0078 | 1.43 | -5.69 |
| Chymotrypsin | 114 | 36 | 2669 | 1 | 4975 | 5611.4795 | 87.1178 | 145,189 | 5610.471675 | 9 | 319 | 342 | RVQPTESIVRFPNITNLCPFGEVF | 0.000000000000200000000000.0 | 331(NIT) | 1 | 2822.4312 | 2819.442939 | 76519264 | 93.02 | 1 | 7 | 6 | 3 | 0 | 0 | 0 | 0 | 5610.462672 | 0.0090 | 1.6 | -5.90 |
| Chymotrypsin | 114 | 36 | 2682 | 1 | 4981 | 5246.3454 | 87.3714 | 144,541 | 5245.337575 | 9 | 319 | 342 | RVQPTESIVRFPNITNLCPFGEVF | 0.000000000000200000000000.0 | 331(NIT) | 1 | 2822.4312 | 2819.442939 | 76519264 | 93.02 | 1 | 6 | 5 | 3 | 0 | 0 | 0 | 1 | 5245.330475 | 0.0071 | 1.35 | -5.64 |
| Chymotrypsin | 114 | 36 | 2741 | 1 | 4971 | 5903.5899 | 86.9455 | 140,592 | 5902.582075 | 9 | 319 | 342 | RVQPTESIVRFPNITNLCPFGEVF | 0.000000000000200000000000.0 | 331(NIT) | 1 | 2822.4312 | 2819.442939 | 76519264 | 93.02 | 1 | 9 | 8 | 0 | 0 | 0 | 0 | 1 | 5902.553339 | 0.0287 | 4.86 | -6.07 |
| Chymotrypsin | 114 | 36 | 2741 | 1 | 4971 | 5903.5899 | 86.9455 | 140,592 | 5902.582075 | 9 | 319 | 342 | RVQPTESIVRFPNITNLCPFGEVF | 0.000000000000200000000000.0 | 331(NIT) | 1 | 2822.4312 | 2819.442939 | 76519264 | 93.02 | 2 | 2 | 10 | 1 | 2 | 0 | 1 | -1 | 5902.59106 | -0.0090 | -1.52 | -6.07 |
| Chymotrypsin | 114 | 36 | 2762 | 1 | 4980 | 5319.3542 | 87.3293 | 139,337 | 5318.346375 | 9 | 319 | 342 | RVQPTESIVRFPNITNLCPFGEVF | 0.000000000000200000000000.0 | 331(NIT) | 1 | 2822.4312 | 2819.442939 | 76519264 | 93.02 | 1 | 7 | 6 | 1 | 0 | 0 | 0 | 1 | 5318.346854 | -0.0005 | -0.09 | -5.69 |
| Chymotrypsin | 114 | 36 | 2839 | 1 | 4987 | 5157.2938 | 87.63 | 134,687 | 5156.285975 | 9 | 319 | 342 | RVQPTESIVRFPNITNLCPFGEVF | 0.000000000000200000000000.0 | 331(NIT) | 1 | 2822.4312 | 2819.442939 | 76519264 | 93.02 | 1 | 6 | 6 | 1 | 0 | 0 | 0 | 1 | 5156.29403 | -0.0081 | -1.56 | -5.39 |
| Chymotrypsin | 114 | 36 | 3702 | 1 | 4995 | 4630.1109 | 87.9646 | 100,156 | 4629.103075 | 9 | 319 | 342 | RVQPTESIVRFPNITNLCPFGEVF | 0.000000000000200000000000.0 | 331(NIT) | 1 | 2822.4312 | 2819.442939 | 76519264 | 93.02 | 1 | 4 | 5 | 1 | 0 | 0 | 0 | 1 | 4629.109009 | -0.0059 | -1.28 | -5.05 |
| Chymotrypsin | 114 | 36 | 3894 | 1 | 4977 | 5465.4103 | 87.2014 | 94,833 | 5464.402475 | 9 | 319 | 342 | RVQPTESIVRFPNITNLCPFGEVF | 0.000000000000200000000000.0 | 331(NIT) | 1 | 2822.4312 | 2819.442939 | 76519264 | 93.02 | 1 | 7 | 6 | 2 | 0 | 0 | 0 | 0 | 5464.404763 | -0.0023 | -0.41 | -5.81 |
| Chymotrypsin | 114 | 36 | 3940 | 1 | 4986 | 4922.2385 | 87.5865 | 93,465 | 4921.230675 | 9 | 319 | 342 | RVQPTESIVRFPNITNLCPFGEVF | 0.000000000000200000000000.0 | 331(NIT) | 1 | 2822.4312 | 2819.442939 | 76519264 | 93.02 | 1 | 4 | 5 | 3 | 0 | 0 | 0 | 0 | 4921.224827 | 0.0058 | 1.18 | -5.43 |
| Chymotrypsin | 114 | 36 | 3991 | 1 | 4985 | 5100.2697 | 87.5423 | 92,206 | 5099.261875 | 9 | 319 | 342 | RVQPTESIVRFPNITNLCPFGEVF | 0.000000000000200000000000.0 | 331(NIT) | 1 | 2822.4312 | 2819.442939 | 76519264 | 93.02 | 1 | 6 | 5 | 2 | 0 | 0 | 0 | 1 | 5099.272566 | -0.0107 | -2.09 | -5.47 |
| Chymotrypsin | 114 | 36 | 4079 | 1 | 5079 | 5099.2626 | 91.4109 | 89,584 | 5098.254775 | 9 | 319 | 342 | RVQPTESIVRFPNITNLCPFGEVF | 0.000000000000200000000000.0 | 331(NIT) | 1 | 2822.4312 | 2819.442939 | 76519264 | 93.02 | 1 | 6 | 5 | 0 | 1 | 0 | 0 | 1 | 5098.252165 | 0.0026 | 0.51 | -1.61 |
| Chymotrypsin | 114 | 36 | 4079 | 1 | 5079 | 5099.2626 | 91.4109 | 89,584 | 5098.254775 | 9 | 319 | 342 | RVQPTESIVRFPNITNLCPFGEVF | 0.000000000000200000000000.0 | 331(NIT) | 1 | 2822.4312 | 2819.442939 | 76519264 | 93.02 | 2 | 5 | 5 | 1 | 0 | 1 | 0 | 1 | 5098.252164 | 0.0026 | 0.51 | -1.61 |
| Chymotrypsin | 114 | 36 | 4618 | 1 | 5000 | 4532.0683 | 88.1604 | 76,630 | 4531.060475 | 9 | 319 | 342 | RVQPTESIVRFPNITNLCPFGEVF | 0.000000000000200000000000.0 | 331(NIT) | 1 | 2822.4312 | 2819.442939 | 76519264 | 93.02 | 1 | 5 | 3 | 2 | 0 | 0 | 0 | 0 | 4531.060996 | -0.0005 | -0.11 | -4.86 |
| Chymotrypsin | 114 | 36 | 5227 | 1 | 5003 | 4264.9812 | 88.2893 | 64,762 | 4263.973375 | 9 | 319 | 342 | RVQPTESIVRFPNITNLCPFGEVF | 0.000000000000200000000000.0 | 331(NIT) | 1 | 2822.4312 | 2819.442939 | 76519264 | 93.02 | 1 | 3 | 4 | 1 | 0 | 0 | 0 | 1 | 4263.976812 | -0.0034 | -0.8 | -4.73 |
| Chymotrypsin | 114 | 36 | 5332 | 1 | 5000 | 4468.0774 | 88.1604 | 62,931 | 4467.069575 | 9 | 319 | 342 | RVQPTESIVRFPNITNLCPFGEVF | 0.000000000000200000000000.0 | 331(NIT) | 1 | 2822.4312 | 2819.442939 | 76519264 | 93.02 | 1 | 3 | 5 | 1 | 0 | 0 | 0 | 1 | 4467.056185 | 0.0134 | 2.99 | -4.86 |
| Chymotrypsin | 114 | 36 | 5441 | 1 | 5074 | 5245.322 | 91.1976 | 61,045 | 5244.314175 | 9 | 319 | 342 | RVQPTESIVRFPNITNLCPFGEVF | 0.000000000000200000000000.0 | 331(NIT) | 1 | 2822.4312 | 2819.442939 | 76519264 | 93.02 | 1 | 6 | 5 | 1 | 1 | 0 | 0 | 1 | 5244.310074 | 0.0041 | 0.78 | -1.82 |
| Chymotrypsin | 114 | 36 | 5441 | 1 | 5074 | 5245.322 | 91.1976 | 61,045 | 5244.314175 | 9 | 319 | 342 | RVQPTESIVRFPNITNLCPFGEVF | 0.000000000000200000000000.0 | 331(NIT) | 1 | 2822.4312 | 2819.442939 | 76519264 | 93.02 | 2 | 5 | 5 | 2 | 0 | 1 | 0 | 0 | 5244.310073 | 0.0041 | 0.78 | -1.82 |
| Chymotrypsin | 114 | 36 | 5556 | 1 | 5081 | 4880.1893 | 91.4969 | 59,256 | 4879.181475 | 9 | 319 | 342 | RVQPTESIVRFPNITNLCPFGEVF | 0.000000000000200000000000.0 | 331(NIT) | 1 | 2822.4312 | 2819.442939 | 76519264 | 93.02 | 1 | 5 | 4 | 1 | 1 | 0 | 0 | 1 | 4879.177877 | 0.0036 | 0.73 | -1.52 |
| Chymotrypsin | 114 | 36 | 5556 | 1 | 5081 | 4880.1893 | 91.4969 | 59,256 | 4879.181475 | 9 | 319 | 342 | RVQPTESIVRFPNITNLCPFGEVF | 0.000000000000200000000000.0 | 331(NIT) | 1 | 2822.4312 | 2819.442939 | 76519264 | 93.02 | 2 | 4 | 4 | 2 | 0 | 1 | 0 | 1 | 4879.177876 | 0.0036 | 0.73 | -1.52 |
| Chymotrypsin | 114 | 36 | 5990 | 1 | 5061 | 5772.5031 | 90.6658 | 52,157 | 5771.495275 | 9 | 319 | 342 | RVQPTESIVRFPNITNLCPFGEVF | 0.000000000000200000000000.0 | 331(NIT) | 1 | 2822.4312 | 2819.442939 | 76519264 | 93.02 | 1 | 8 | 6 | 1 | 1 | 0 | 0 | 1 | 5771.495095 | 0.0002 | 0.03 | -2.35 |
| Chymotrypsin | 114 | 36 | 5990 | 1 | 5061 | 5772.5031 | 90.6658 | 52,157 | 5771.495275 | 9 | 319 | 342 | RVQPTESIVRFPNITNLCPFGEVF | 0.000000000000200000000000.0 | 331(NIT) | 1 | 2822.4312 | 2819.442939 | 76519264 | 93.02 | 2 | 7 | 6 | 2 | 0 | 1 | 0 | 0 | 5771.495094 | 0.0002 | 0.03 | -2.35 |
| Chymotrypsin | 114 | 36 | 6474 | 1 | 5079 | 5464.3832 | 91.4109 | 45,407 | 5463.375375 | 9 | 319 | 342 | RVQPTESIVRFPNITNLCPFGEVF | 0.000000000000200000000000.0 | 331(NIT) | 1 | 2822.4312 | 2819.442939 | 76519264 | 93.02 | 1 | 7 | 6 | 0 | 1 | 0 | 0 | 1 | 5463.384362 | -0.0090 | -1.64 | -1.61 |
| Chymotrypsin | 114 | 36 | 6474 | 1 | 5079 | 5464.3832 | 91.4109 | 45,407 | 5463.375375 | 9 | 319 | 342 | RVQPTESIVRFPNITNLCPFGEVF | 0.000000000000200000000000.0 | 331(NIT) | 1 | 2822.4312 | 2819.442939 | 76519264 | 93.02 | 2 | 6 | 6 | 1 | 0 | 1 | 0 | 0 | 5463.384361 | -0.0090 | -1.64 | -1.61 |
| Chymotrypsin | 114 | 36 | 6616 | 1 | 5073 | 5261.3239 | 91.1547 | 43,653 | 5260.316075 | 9 | 319 | 342 | RVQPTESIVRFPNITNLCPFGEVF | 0.000000000000200000000000.0 | 331(NIT) | 1 | 2822.4312 | 2819.442939 | 76519264 | 93.02 | 1 | 6 | 5 | 1 | 0 | 1 | 0 | 1 | 5260.304988 | 0.0111 | 2.1 | -1.86 |
| Chymotrypsin | 114 | 36 | 6616 | 1 | 5073 | 5261.3239 | 91.1547 | 43,653 | 5260.316075 | 9 | 319 | 342 | RVQPTESIVRFPNITNLCPFGEVF | 0.000000000000200000000000.0 | 331(NIT) | 1 | 2822.4312 | 2819.442939 | 76519264 | 93.02 | 2 | 7 | 5 | 0 | 1 | 0 | 0 | 0 | 5260.304989 | 0.0111 | 2.1 | -1.86 |
| Chymotrypsin | 114 | 36 | 6723 | 1 | 4999 | 4573.1041 | 88.1343 | 41,946 | 4572.096275 | 9 | 319 | 342 | RVQPTESIVRFPNITNLCPFGEVF | 0.000000000000200000000000.0 | 331(NIT) | 1 | 2822.4312 | 2819.442939 | 76519264 | 93.02 | 1 | 4 | 4 | 2 | 0 | 0 | 0 | 1 | 4572.087545 | 0.0087 | 1.9 | -4.88 |
| Chymotrypsin | 114 | 36 | 6735 | 1 | 5063 | 5553.4162 | 90.7504 | 41,841 | 5552.408375 | 9 | 319 | 342 | RVQPTESIVRFPNITNLCPFGEVF | 0.000000000000200000000000.0 | 331(NIT) | 1 | 2822.4312 | 2819.442939 | 76519264 | 93.02 | 1 | 6 | 5 | 3 | 0 | 1 | 0 | 1 | 5552.420806 | -0.0124 | -2.23 | -2.27 |
| Chymotrypsin | 114 | 36 | 6735 | 1 | 5063 | 5553.4162 | 90.7504 | 41,841 | 5552.408375 | 9 | 319 | 342 | RVQPTESIVRFPNITNLCPFGEVF | 0.000000000000200000000000.0 | 331(NIT) | 1 | 2822.4312 | 2819.442939 | 76519264 | 93.02 | 2 | 7 | 5 | 2 | 1 | 0 | 0 | 0 | 5552.420807 | -0.0124 | -2.23 | -2.27 |
| Chymotrypsin | 114 | 36 | 6796 | 1 | 5064 | 5626.4479 | 90.793 | 40,955 | 5625.440075 | 9 | 319 | 342 | RVQPTESIVRFPNITNLCPFGEVF | 0.000000000000200000000000.0 | 331(NIT) | 1 | 2822.4312 | 2819.442939 | 76519264 | 93.02 | 1 | 8 | 6 | 0 | 1 | 0 | 0 | 1 | 5625.437186 | 0.0029 | 0.51 | -2.22 |
| Chymotrypsin | 114 | 36 | 6796 | 1 | 5064 | 5626.4479 | 90.793 | 40,955 | 5625.440075 | 9 | 319 | 342 | RVQPTESIVRFPNITNLCPFGEVF | 0.000000000000200000000000.0 | 331(NIT) | 1 | 2822.4312 | 2819.442939 | 76519264 | 93.02 | 2 | 7 | 6 | 1 | 0 | 1 | 0 | 1 | 5625.437185 | 0.0029 | 0.51 | -2.22 |
| Chymotrypsin | 114 | 36 | 8022 | 1 | 5169 | 5568.3957 | 95.1067 | 24,893 | 5567.387875 | 9 | 319 | 342 | RVQPTESIVRFPNITNLCPFGEVF | 0.000000000000200000000000.0 | 331(NIT) | 1 | 2822.4312 | 2819.442939 | 76519264 | 93.02 | 1 | 6 | 5 | 1 | 0 | 2 | 0 | 1 | 5567.395319 | -0.0074 | -1.33 | 2.09 |
| Chymotrypsin | 114 | 36 | 8022 | 1 | 5169 | 5568.3957 | 95.1067 | 24,893 | 5567.387875 | 9 | 319 | 342 | RVQPTESIVRFPNITNLCPFGEVF | 0.000000000000200000000000.0 | 331(NIT) | 1 | 2822.4312 | 2819.442939 | 76519264 | 93.02 | 2 | 7 | 5 | 0 | 1 | 1 | 0 | 0 | 5567.39532 | -0.0074 | -1.33 | 2.09 |
| Chymotrypsin | 114 | 36 | 8793 | 1 | 5056 | 6064.632 | 90.4544 | 14,355 | 6063.624175 | 9 | 319 | 342 | RVQPTESIVRFPNITNLCPFGEVF | 0.000000000000200000000000.0 | 331(NIT) | 1 | 2822.4312 | 2819.442939 | 76519264 | 93.02 | 1 | 7 | 6 | 4 | 0 | 1 | 0 | 1 | 6063.610912 | 0.0133 | 2.18 | -2.56 |
| Chymotrypsin | 114 | 36 | 8793 | 1 | 5056 | 6064.632 | 90.4544 | 14,355 | 6063.624175 | 9 | 319 | 342 | RVQPTESIVRFPNITNLCPFGEVF | 0.000000000000200000000000.0 | 331(NIT) | 1 | 2822.4312 | 2819.442939 | 76519264 | 93.02 | 2 | 2 | 10 | 0 | 2 | 1 | 1 | -1 | 6063.623482 | 0.0007 | 0.11 | -2.56 |
| Chymotrypsin | 114 | 36 | 8793 | 1 | 5056 | 6064.632 | 90.4544 | 14,355 | 6063.624175 | 9 | 319 | 342 | RVQPTESIVRFPNITNLCPFGEVF | 0.000000000000200000000000.0 | 331(NIT) | 1 | 2822.4312 | 2819.442939 | 76519264 | 93.02 | 3 | 8 | 6 | 3 | 1 | 0 | 0 | 0 | 6063.610913 | 0.0133 | 2.18 | -2.56 |
| Chymotrypsin | 219 | 4 | 1472 | 1 | 4599 | 4044.8547 | 71.9825 | 262,179 | 4043.846875 | 25 | 319 | 335 | RVQPTESIVRFPNITNL | 0.00000000000020000.0 | 331(NIT) | 1 | 1986.0784 | 1983.090139 | 18904074 | 77.77 | 1 | 5 | 4 | 3 | 0 | 0 | 0 | 1 | 4043.845478 | 0.0014 | 0.34 | -5.79 |
| Chymotrypsin | 219 | 4 | 2488 | 1 | 4598 | 4247.9238 | 71.9396 | 156,012 | 4246.915975 | 25 | 319 | 335 | RVQPTESIVRFPNITNL | 0.00000000000020000.0 | 331(NIT) | 1 | 1986.0784 | 1983.090139 | 18904074 | 77.77 | 1 | 5 | 5 | 3 | 0 | 0 | 0 | 1 | 4246.924851 | -0.0089 | -2.08 | -5.83 |
| Chymotrypsin | 219 | 4 | 3320 | 1 | 4613 | 3955.8186 | 72.4061 | 113,083 | 3954.810775 | 25 | 319 | 335 | RVQPTESIVRFPNITNL | 0.00000000000020000.0 | 331(NIT) | 1 | 1986.0784 | 1983.090139 | 18904074 | 77.77 | 1 | 5 | 5 | 1 | 0 | 0 | 0 | 1 | 3954.809033 | 0.0017 | 0.44 | -5.36 |
| Chymotrypsin | 219 | 4 | 4482 | 1 | 4606 | 4101.88 | 72.221 | 79,459 | 4100.872175 | 25 | 319 | 335 | RVQPTESIVRFPNITNL | 0.00000000000020000.0 | 331(NIT) | 1 | 1986.0784 | 1983.090139 | 18904074 | 77.77 | 1 | 5 | 5 | 2 | 0 | 0 | 0 | 1 | 4100.866942 | 0.0052 | 1.27 | -5.55 |
| alpha-Lytic | 63 | 15 | 352 | 1 | 4268 | 2808.1714 | 49.3336 | 378,110 | 2807.163575 | 1004 | 328 | 333 | RFPNIT | - | 331(NIT) | 1 | | 746.4075 | - | 54.00 | 1 | 5 | 4 | 3 | 0 | 0 | 0 | 1 | 2807.162839 | 0.0007 | 0.26 | -4.67 |
| alpha-Lytic | 63 | 15 | 702 | 1 | 4268 | 3011.2492 | 49.3336 | 190,866 | 3010.241375 | 1004 | 328 | 333 | RFPNIT | - | 331(NIT) | 1 | | 746.4075 | - | 54.00 | 1 | 5 | 5 | 3 | 0 | 0 | 0 | 1 | 3010.242212 | -0.0008 | -0.27 | -4.67 |
| alpha-Lytic | 63 | 15 | 791 | 1 | 4275 | 2662.1131 | 49.6859 | 166,641 | 2661.105275 | 1004 | 328 | 333 | RFPNIT | - | 331(NIT) | 1 | | 746.4075 | - | 54.00 | 1 | 5 | 4 | 2 | 0 | 0 | 0 | 1 | 2661.10493 | 0.0003 | 0.12 | -4.31 |
| alpha-Lytic | 63 | 15 | 939 | 1 | 4284 | 2516.0538 | 50.1345 | 141,527 | 2515.045975 | 1004 | 328 | 333 | RFPNIT | - | 331(NIT) | 1 | | 746.4075 | - | 54.00 | 1 | 5 | 4 | 1 | 0 | 0 | 0 | 1 | 2515.047021 | -0.0010 | -0.41 | -3.87 |
| alpha-Lytic | 63 | 15 | 1011 | 1 | 4291 | 2719.1343 | 50.4832 | 131,673 | 2718.126475 | 1004 | 328 | 333 | RFPNIT | - | 331(NIT) | 1 | | 746.4075 | - | 54.00 | 1 | 5 | 5 | 1 | 0 | 0 | 0 | 1 | 2718.126394 | 0.0001 | 0.02 | -3.52 |
| alpha-Lytic | 63 | 15 | 1043 | 1 | 4279 | 2865.1963 | 49.8877 | 128,512 | 2864.188475 | 1004 | 328 | 333 | RFPNIT | - | 331(NIT) | 1 | | 746.4075 | - | 54.00 | 1 | 5 | 5 | 2 | 0 | 0 | 0 | 1 | 2864.184303 | 0.0042 | 1.45 | -4.11 |
| alpha-Lytic | 63 | 15 | 1233 | 1 | 4350 | 2823.1453 | 53.4027 | 107,068 | 2822.137475 | 1004 | 328 | 333 | RFPNIT | - | 331(NIT) | 1 | | 746.4075 | - | 54.00 | 1 | 6 | 4 | 0 | 1 | 0 | 0 | 1 | 2822.137353 | 0.0001 | 0.04 | -0.60 |
| alpha-Lytic | 63 | 15 | 1233 | 1 | 4350 | 2823.1453 | 53.4027 | 107,068 | 2822.137475 | 1004 | 328 | 333 | RFPNIT | - | 331(NIT) | 1 | | 746.4075 | - | 54.00 | 2 | 5 | 4 | 1 | 0 | 1 | 0 | 1 | 2822.137352 | 0.0001 | 0.04 | -0.60 |
| alpha-Lytic | 63 | 15 | 1394 | 1 | 4251 | 3319.3601 | 48.5069 | 92,668 | 3318.352275 | 1004 | 328 | 333 | RFPNIT | - | 331(NIT) | 1 | | 746.4075 | - | 54.00 | 1 | 6 | 5 | 4 | 0 | 0 | 0 | 0 | 3318.352945 | -0.0007 | -0.2 | -5.49 |
| alpha-Lytic | 63 | 15 | 1602 | 1 | 4343 | 2969.2056 | 53.0488 | 78,720 | 2968.197775 | 1004 | 328 | 333 | RFPNIT | - | 331(NIT) | 1 | | 746.4075 | - | 54.00 | 1 | 6 | 4 | 1 | 1 | 0 | 0 | 1 | 2968.195262 | 0.0025 | 0.84 | -0.95 |
| alpha-Lytic | 63 | 15 | 1602 | 1 | 4343 | 2969.2056 | 53.0488 | 78,720 | 2968.197775 | 1004 | 328 | 333 | RFPNIT | - | 331(NIT) | 1 | | 746.4075 | - | 54.00 | 2 | 5 | 4 | 2 | 0 | 1 | 0 | 1 | 2968.195261 | 0.0025 | 0.84 | -0.95 |
| alpha-Lytic | 63 | 15 | 2066 | 1 | 4253 | 3538.4363 | 48.6035 | 57,028 | 3537.428475 | 1004 | 328 | 333 | RFPNIT | - | 331(NIT) | 1 | | 746.4075 | - | 54.00 | 1 | 7 | 6 | 3 | 0 | 0 | 0 | 0 | 3537.427233 | 0.0012 | 0.35 | -5.40 |
| alpha-Lytic | 63 | 15 | 2417 | 1 | 4263 | 3173.302 | 49.0893 | 45,047 | 3172.294175 | 1004 | 328 | 333 | RFPNIT | - | 331(NIT) | 1 | | 746.4075 | - | 54.00 | 1 | 6 | 5 | 3 | 0 | 0 | 0 | 1 | 3172.295036 | -0.0009 | -0.27 | -4.91 |
| alpha-Lytic | 63 | 15 | 2632 | 1 | 4291 | 3084.2666 | 50.4832 | 38,868 | 3083.258775 | 1004 | 328 | 333 | RFPNIT | - | 331(NIT) | 1 | | 746.4075 | - | 54.00 | 1 | 6 | 6 | 1 | 0 | 0 | 0 | 1 | 3083.258591 | 0.0002 | 0.05 | -3.52 |
| alpha-Lytic | 63 | 15 | 2859 | 1 | 4352 | 2969.1983 | 53.5005 | 33,157 | 2968.190475 | 1004 | 328 | 333 | RFPNIT | - | 331(NIT) | 1 | | 746.4075 | - | 54.00 | 1 | 5 | 4 | 2 | 0 | 1 | 0 | 1 | 2968.195261 | -0.0048 | -1.61 | -0.50 |
| alpha-Lytic | 63 | 15 | 2859 | 1 | 4352 | 2969.1983 | 53.5005 | 33,157 | 2968.190475 | 1004 | 328 | 333 | RFPNIT | - | 331(NIT) | 1 | | 746.4075 | - | 54.00 | 2 | 6 | 4 | 1 | 1 | 0 | 0 | 1 | 2968.195262 | -0.0048 | -1.61 | -0.50 |
| alpha-Lytic | 63 | 15 | 3153 | 1 | 4280 | 2881.1852 | 49.9361 | 26,719 | 2880.177375 | 1004 | 328 | 333 | RFPNIT | - | 331(NIT) | 1 | | 746.4075 | - | 54.00 | 1 | 6 | 5 | 1 | 0 | 0 | 0 | 1 | 2880.179218 | -0.0018 | -0.63 | -4.06 |
| alpha-Lytic | 63 | 15 | 3386 | 1 | 4349 | 3172.2777 | 53.3563 | 21,303 | 3171.269875 | 1004 | 328 | 333 | RFPNIT | - | 331(NIT) | 1 | | 746.4075 | - | 54.00 | 1 | 6 | 5 | 1 | 1 | 0 | 0 | 1 | 3171.274635 | -0.0048 | -1.5 | -0.64 |
| alpha-Lytic | 63 | 15 | 3386 | 1 | 4349 | 3172.2777 | 53.3563 | 21,303 | 3171.269875 | 1004 | 328 | 333 | RFPNIT | - | 331(NIT) | 1 | | 746.4075 | - | 54.00 | 2 | 5 | 5 | 2 | 0 | 1 | 0 | 0 | 3171.274634 | -0.0048 | -1.5 | -0.64 |
| Chymotrypsin | 16 | 71 | 43 | 1 | 4177 | 3101.27 | 54.3618 | 5,278,720 | 3100.262175 | 63 | 339 | 347 | GEVFNATRF | 0.000020000.0 | 343(NAT) | 1 | 1042.497 | 1039.508739 | 5711414 | 62.15 | 1 | 5 | 4 | 3 | 0 | 0 | 0 | 1 | 3100.264078 | -0.0019 | -0.61 | -7.79 |
| Chymotrypsin | 16 | 71 | 72 | 1 | 4174 | 3304.3504 | 54.2318 | 4,050,230 | 3303.342575 | 63 | 339 | 347 | GEVFNATRF | 0.000020000.0 | 343(NAT) | 1 | 1042.497 | 1039.508739 | 5711414 | 62.15 | 1 | 5 | 5 | 3 | 0 | 0 | 0 | 1 | 3303.343451 | -0.0009 | -0.26 | -7.92 |
| Chymotrypsin | 16 | 71 | 158 | 1 | 4190 | 2809.1539 | 54.9062 | 1,993,320 | 2808.146075 | 63 | 339 | 347 | GEVFNATRF | 0.000020000.0 | 343(NAT) | 1 | 1042.497 | 1039.508739 | 5711414 | 62.15 | 1 | 5 | 4 | 1 | 0 | 0 | 0 | 1 | 2808.14826 | -0.0022 | -0.77 | -7.24 |
| Chymotrypsin | 16 | 71 | 172 | 1 | 4375 | 4258.641 | 62.5564 | 1,845,370 | 4257.633175 | 63 | 339 | 347 | GEVFNATRF | 0.000020000.0 | 343(NAT) | 1 | 1042.497 | 1039.508739 | 5711414 | 62.15 | 1 | 10 | 5 | 0 | 2 | 0 | 1 | -1 | 4257.624678 | 0.0085 | 1.99 | 0.41 |
| Chymotrypsin | 16 | 71 | 172 | 1 | 4375 | 4258.641 | 62.5564 | 1,845,370 | 4257.633175 | 63 | 339 | 347 | GEVFNATRF | 0.000020000.0 | 343(NAT) | 1 | 1042.497 | 1039.508739 | 5711414 | 62.15 | 2 | 9 | 5 | 1 | 1 | 1 | 0 | 0 | 4257.624677 | 0.0085 | 1.99 | 0.41 |
| Chymotrypsin | 16 | 71 | 172 | 1 | 4375 | 4258.641 | 62.5564 | 1,845,370 | 4257.633175 | 63 | 339 | 347 | GEVFNATRF | 0.000020000.0 | 343(NAT) | 1 | 1042.497 | 1039.508739 | 5711414 | 62.15 | 3 | 8 | 5 | 2 | 0 | 2 | 0 | 0 | 4257.624676 | 0.0085 | 1.99 | 0.41 |
| Chymotrypsin | 16 | 71 | 177 | 1 | 4380 | 4096.5901 | 62.7702 | 1,791,210 | 4095.582275 | 63 | 339 | 347 | GEVFNATRF | 0.000020000.0 | 343(NAT) | 1 | 1042.497 | 1039.508739 | 5711414 | 62.15 | 1 | 9 | 5 | 0 | 2 | 0 | 0 | 0 | 4095.571854 | 0.0104 | 2.54 | 0.62 |
| Chymotrypsin | 16 | 71 | 177 | 1 | 4380 | 4096.5901 | 62.7702 | 1,791,210 | 4095.582275 | 63 | 339 | 347 | GEVFNATRF | 0.000020000.0 | 343(NAT) | 1 | 1042.497 | 1039.508739 | 5711414 | 62.15 | 2 | 8 | 5 | 1 | 1 | 1 | 0 | 0 | 4095.571853 | 0.0104 | 2.54 | 0.62 |
| Chymotrypsin | 16 | 71 | 177 | 1 | 4380 | 4096.5901 | 62.7702 | 1,791,210 | 4095.582275 | 63 | 339 | 347 | GEVFNATRF | 0.000020000.0 | 343(NAT) | 1 | 1042.497 | 1039.508739 | 5711414 | 62.15 | 3 | 7 | 5 | 2 | 0 | 2 | 0 | 0 | 4095.571852 | 0.0104 | 2.54 | 0.62 |
| Chymotrypsin | 16 | 71 | 208 | 1 | 4388 | 3934.5367 | 63.1108 | 1,549,080 | 3933.528875 | 63 | 339 | 347 | GEVFNATRF | 0.000020000.0 | 343(NAT) | 1 | 1042.497 | 1039.508739 | 5711414 | 62.15 | 1 | 7 | 5 | 1 | 1 | 1 | 0 | 1 | 3933.519029 | 0.0098 | 2.5 | 0.96 |
| Chymotrypsin | 16 | 71 | 208 | 1 | 4388 | 3934.5367 | 63.1108 | 1,549,080 | 3933.528875 | 63 | 339 | 347 | GEVFNATRF | 0.000020000.0 | 343(NAT) | 1 | 1042.497 | 1039.508739 | 5711414 | 62.15 | 2 | 8 | 5 | 0 | 2 | 0 | 0 | 0 | 3933.51903 | 0.0098 | 2.5 | 0.96 |
| Chymotrypsin | 16 | 71 | 208 | 1 | 4388 | 3934.5367 | 63.1108 | 1,549,080 | 3933.528875 | 63 | 339 | 347 | GEVFNATRF | 0.000020000.0 | 343(NAT) | 1 | 1042.497 | 1039.508739 | 5711414 | 62.15 | 3 | 6 | 5 | 2 | 0 | 2 | 0 | 0 | 3933.519028 | 0.0098 | 2.5 | 0.96 |
| Chymotrypsin | 16 | 71 | 259 | 1 | 4192 | 3012.2342 | 54.994 | 1,289,860 | 3011.226375 | 63 | 339 | 347 | GEVFNATRF | 0.000020000.0 | 343(NAT) | 1 | 1042.497 | 1039.508739 | 5711414 | 62.15 | 1 | 5 | 5 | 1 | 0 | 0 | 0 | 1 | 3011.227633 | -0.0013 | -0.41 | -7.15 |
| Chymotrypsin | 16 | 71 | 263 | 1 | 4369 | 4420.6941 | 62.3022 | 1,266,680 | 4419.686275 | 63 | 339 | 347 | GEVFNATRF | 0.000020000.0 | 343(NAT) | 1 | 1042.497 | 1039.508739 | 5711414 | 62.15 | 1 | 2 | 9 | 0 | 0 | 4 | 1 | -1 | 4419.690068 | -0.0038 | -0.85 | 0.15 |
| Chymotrypsin | 16 | 71 | 263 | 1 | 4369 | 4420.6941 | 62.3022 | 1,266,680 | 4419.686275 | 63 | 339 | 347 | GEVFNATRF | 0.000020000.0 | 343(NAT) | 1 | 1042.497 | 1039.508739 | 5711414 | 62.15 | 2 | 10 | 5 | 1 | 1 | 1 | 1 | -1 | 4419.677501 | 0.0088 | 1.98 | 0.15 |
| Chymotrypsin | 16 | 71 | 263 | 1 | 4369 | 4420.6941 | 62.3022 | 1,266,680 | 4419.686275 | 63 | 339 | 347 | GEVFNATRF | 0.000020000.0 | 343(NAT) | 1 | 1042.497 | 1039.508739 | 5711414 | 62.15 | 3 | 9 | 5 | 2 | 0 | 2 | 0 | 0 | 4419.6775 | 0.0088 | 1.98 | 0.15 |
| Chymotrypsin | 16 | 71 | 277 | 1 | 4183 | 2955.2114 | 54.6183 | 1,237,980 | 2954.203575 | 63 | 339 | 347 | GEVFNATRF | 0.000020000.0 | 343(NAT) | 1 | 1042.497 | 1039.508739 | 5711414 | 62.15 | 1 | 5 | 4 | 2 | 0 | 0 | 0 | 1 | 2954.206169 | -0.0026 | -0.87 | -7.53 |
| Chymotrypsin | 16 | 71 | 324 | 1 | 4183 | 3158.2889 | 54.6183 | 1,075,560 | 3157.281075 | 63 | 339 | 347 | GEVFNATRF | 0.000020000.0 | 343(NAT) | 1 | 1042.497 | 1039.508739 | 5711414 | 62.15 | 1 | 5 | 5 | 2 | 0 | 0 | 0 | 1 | 3157.285542 | -0.0045 | -1.41 | -7.53 |
| Chymotrypsin | 16 | 71 | 445 | 1 | 4192 | 2752.1344 | 54.994 | 800,887 | 2751.126575 | 63 | 339 | 347 | GEVFNATRF | 0.000020000.0 | 343(NAT) | 1 | 1042.497 | 1039.508739 | 5711414 | 62.15 | 1 | 5 | 3 | 2 | 0 | 0 | 0 | 0 | 2751.126796 | -0.0002 | -0.08 | -7.15 |
| Chymotrypsin | 16 | 71 | 574 | 1 | 4166 | 3612.4602 | 53.8946 | 625,150 | 3611.452375 | 63 | 339 | 347 | GEVFNATRF | 0.000020000.0 | 343(NAT) | 1 | 1042.497 | 1039.508739 | 5711414 | 62.15 | 1 | 6 | 5 | 4 | 0 | 0 | 0 | 0 | 3611.454184 | -0.0018 | -0.5 | -8.25 |
| Chymotrypsin | 16 | 71 | 603 | 1 | 4272 | 3262.3025 | 58.4038 | 597,243 | 3261.294675 | 63 | 339 | 347 | GEVFNATRF | 0.000020000.0 | 343(NAT) | 1 | 1042.497 | 1039.508739 | 5711414 | 62.15 | 1 | 5 | 4 | 2 | 0 | 1 | 0 | 1 | 3261.2965 | -0.0018 | -0.55 | -3.74 |
| Chymotrypsin | 16 | 71 | 603 | 1 | 4272 | 3262.3025 | 58.4038 | 597,243 | 3261.294675 | 63 | 339 | 347 | GEVFNATRF | 0.000020000.0 | 343(NAT) | 1 | 1042.497 | 1039.508739 | 5711414 | 62.15 | 2 | 6 | 4 | 1 | 1 | 0 | 0 | 1 | 3261.296501 | -0.0018 | -0.55 | -3.74 |
| Chymotrypsin | 16 | 71 | 647 | 1 | 4395 | 3772.4809 | 63.4102 | 559,844 | 3771.473075 | 63 | 339 | 347 | GEVFNATRF | 0.000020000.0 | 343(NAT) | 1 | 1042.497 | 1039.508739 | 5711414 | 62.15 | 1 | 6 | 5 | 1 | 1 | 1 | 0 | 1 | 3771.466205 | 0.0069 | 1.82 | 1.26 |
| Chymotrypsin | 16 | 71 | 647 | 1 | 4395 | 3772.4809 | 63.4102 | 559,844 | 3771.473075 | 63 | 339 | 347 | GEVFNATRF | 0.000020000.0 | 343(NAT) | 1 | 1042.497 | 1039.508739 | 5711414 | 62.15 | 2 | 7 | 5 | 0 | 2 | 0 | 0 | 1 | 3771.466206 | 0.0069 | 1.82 | 1.26 |
| Chymotrypsin | 16 | 71 | 647 | 1 | 4395 | 3772.4809 | 63.4102 | 559,844 | 3771.473075 | 63 | 339 | 347 | GEVFNATRF | 0.000020000.0 | 343(NAT) | 1 | 1042.497 | 1039.508739 | 5711414 | 62.15 | 3 | 5 | 5 | 2 | 0 | 2 | 0 | 0 | 3771.466204 | 0.0069 | 1.82 | 1.26 |
| Chymotrypsin | 16 | 71 | 690 | 1 | 4203 | 2485.0528 | 55.4605 | 527,320 | 2484.044975 | 63 | 339 | 347 | GEVFNATRF | 0.000020000.0 | 343(NAT) | 1 | 1042.497 | 1039.508739 | 5711414 | 62.15 | 1 | 3 | 4 | 1 | 0 | 0 | 0 | 1 | 2484.042612 | 0.0024 | 0.95 | -6.69 |
| Chymotrypsin | 16 | 71 | 833 | 1 | 4386 | 4137.6126 | 63.0266 | 448,338 | 4136.604775 | 63 | 339 | 347 | GEVFNATRF | 0.000020000.0 | 343(NAT) | 1 | 1042.497 | 1039.508739 | 5711414 | 62.15 | 1 | 8 | 6 | 0 | 2 | 0 | 0 | 0 | 4136.598403 | 0.0064 | 1.54 | 0.88 |
| Chymotrypsin | 16 | 71 | 833 | 1 | 4386 | 4137.6126 | 63.0266 | 448,338 | 4136.604775 | 63 | 339 | 347 | GEVFNATRF | 0.000020000.0 | 343(NAT) | 1 | 1042.497 | 1039.508739 | 5711414 | 62.15 | 2 | 7 | 6 | 1 | 1 | 1 | 0 | 0 | 4136.598402 | 0.0064 | 1.54 | 0.88 |
| Chymotrypsin | 16 | 71 | 833 | 1 | 4386 | 4137.6126 | 63.0266 | 448,338 | 4136.604775 | 63 | 339 | 347 | GEVFNATRF | 0.000020000.0 | 343(NAT) | 1 | 1042.497 | 1039.508739 | 5711414 | 62.15 | 3 | 6 | 6 | 2 | 0 | 2 | 0 | 0 | 4136.598401 | 0.0064 | 1.54 | 0.88 |
| Chymotrypsin | 16 | 71 | 883 | 1 | 4166 | 3815.5378 | 53.8946 | 423,754 | 3814.529975 | 63 | 339 | 347 | GEVFNATRF | 0.000020000.0 | 343(NAT) | 1 | 1042.497 | 1039.508739 | 5711414 | 62.15 | 1 | 6 | 6 | 4 | 0 | 0 | 0 | 0 | 3814.533557 | -0.0036 | -0.93 | -8.25 |
| Chymotrypsin | 16 | 71 | 1024 | 1 | 4280 | 3116.2462 | 58.7494 | 370,360 | 3115.238375 | 63 | 339 | 347 | GEVFNATRF | 0.000020000.0 | 343(NAT) | 1 | 1042.497 | 1039.508739 | 5711414 | 62.15 | 1 | 5 | 4 | 1 | 0 | 1 | 0 | 1 | 3115.238591 | -0.0002 | -0.06 | -3.40 |
| Chymotrypsin | 16 | 71 | 1024 | 1 | 4280 | 3116.2462 | 58.7494 | 370,360 | 3115.238375 | 63 | 339 | 347 | GEVFNATRF | 0.000020000.0 | 343(NAT) | 1 | 1042.497 | 1039.508739 | 5711414 | 62.15 | 2 | 6 | 4 | 0 | 1 | 0 | 0 | 1 | 3115.238592 | -0.0002 | -0.06 | -3.40 |
| Chymotrypsin | 16 | 71 | 1048 | 1 | 4205 | 2768.1282 | 55.5481 | 362,016 | 2767.120375 | 63 | 339 | 347 | GEVFNATRF | 0.000020000.0 | 343(NAT) | 1 | 1042.497 | 1039.508739 | 5711414 | 62.15 | 1 | 6 | 3 | 1 | 0 | 0 | 0 | 1 | 2767.121711 | -0.0013 | -0.48 | -6.60 |
| Chymotrypsin | 16 | 71 | 1051 | 1 | 4188 | 2996.2418 | 54.8278 | 361,199 | 2995.233975 | 63 | 339 | 347 | GEVFNATRF | 0.000020000.0 | 343(NAT) | 1 | 1042.497 | 1039.508739 | 5711414 | 62.15 | 1 | 4 | 5 | 2 | 0 | 0 | 0 | 1 | 2995.232718 | 0.0013 | 0.41 | -7.32 |
| Chymotrypsin | 16 | 71 | 1129 | 1 | 4393 | 3975.5533 | 63.3238 | 340,668 | 3974.545475 | 63 | 339 | 347 | GEVFNATRF | 0.000020000.0 | 343(NAT) | 1 | 1042.497 | 1039.508739 | 5711414 | 62.15 | 1 | 6 | 6 | 1 | 1 | 1 | 0 | 1 | 3974.545578 | -0.0001 | -0.02 | 1.18 |
| Chymotrypsin | 16 | 71 | 1129 | 1 | 4393 | 3975.5533 | 63.3238 | 340,668 | 3974.545475 | 63 | 339 | 347 | GEVFNATRF | 0.000020000.0 | 343(NAT) | 1 | 1042.497 | 1039.508739 | 5711414 | 62.15 | 2 | 7 | 6 | 0 | 2 | 0 | 0 | 1 | 3974.545579 | -0.0001 | -0.02 | 1.18 |
| Chymotrypsin | 16 | 71 | 1129 | 1 | 4393 | 3975.5533 | 63.3238 | 340,668 | 3974.545475 | 63 | 339 | 347 | GEVFNATRF | 0.000020000.0 | 343(NAT) | 1 | 1042.497 | 1039.508739 | 5711414 | 62.15 | 3 | 5 | 6 | 2 | 0 | 2 | 0 | 0 | 3974.545577 | -0.0001 | -0.02 | 1.18 |
| Chymotrypsin | 16 | 71 | 1242 | 1 | 4196 | 2850.1808 | 55.1654 | 308,144 | 2849.172975 | 63 | 339 | 347 | GEVFNATRF | 0.000020000.0 | 343(NAT) | 1 | 1042.497 | 1039.508739 | 5711414 | 62.15 | 1 | 4 | 5 | 1 | 0 | 0 | 0 | 1 | 2849.174809 | -0.0018 | -0.64 | -6.98 |
| Chymotrypsin | 16 | 71 | 1364 | 1 | 4171 | 3466.4007 | 54.1076 | 281,575 | 3465.392875 | 63 | 339 | 347 | GEVFNATRF | 0.000020000.0 | 343(NAT) | 1 | 1042.497 | 1039.508739 | 5711414 | 62.15 | 1 | 6 | 5 | 3 | 0 | 0 | 0 | 1 | 3465.396275 | -0.0034 | -0.98 | -8.04 |
| Chymotrypsin | 16 | 71 | 1399 | 1 | 4186 | 2914.1862 | 54.7443 | 275,936 | 2913.178375 | 63 | 339 | 347 | GEVFNATRF | 0.000020000.0 | 343(NAT) | 1 | 1042.497 | 1039.508739 | 5711414 | 62.15 | 1 | 6 | 3 | 2 | 0 | 0 | 0 | 0 | 2913.17962 | -0.0012 | -0.42 | -7.40 |
| Chymotrypsin | 16 | 71 | 1448 | 1 | 4263 | 4277.7213 | 58.0177 | 264,563 | 4276.713475 | 63 | 339 | 347 | GEVFNATRF | 0.000020000.0 | 343(NAT) | 1 | 1042.497 | 1039.508739 | 5711414 | 62.15 | 1 | 6 | 9 | 1 | 1 | 0 | 0 | 0 | 4276.693366 | 0.0201 | 4.7 | -4.13 |
| Chymotrypsin | 16 | 71 | 1448 | 1 | 4263 | 4277.7213 | 58.0177 | 264,563 | 4276.713475 | 63 | 339 | 347 | GEVFNATRF | 0.000020000.0 | 343(NAT) | 1 | 1042.497 | 1039.508739 | 5711414 | 62.15 | 2 | 5 | 9 | 2 | 0 | 1 | 0 | 0 | 4276.693365 | 0.0201 | 4.7 | -4.13 |
| Chymotrypsin | 16 | 71 | 1529 | 1 | 4378 | 4283.681 | 62.6857 | 252,040 | 4282.673175 | 63 | 339 | 347 | GEVFNATRF | 0.000020000.0 | 343(NAT) | 1 | 1042.497 | 1039.508739 | 5711414 | 62.15 | 1 | 8 | 6 | 1 | 2 | 0 | 0 | 0 | 4282.656312 | 0.0169 | 3.93 | 0.54 |
| Chymotrypsin | 16 | 71 | 1529 | 1 | 4378 | 4283.681 | 62.6857 | 252,040 | 4282.673175 | 63 | 339 | 347 | GEVFNATRF | 0.000020000.0 | 343(NAT) | 1 | 1042.497 | 1039.508739 | 5711414 | 62.15 | 2 | 7 | 6 | 2 | 1 | 1 | 0 | 0 | 4282.656311 | 0.0169 | 3.93 | 0.54 |
| Chymotrypsin | 16 | 71 | 1529 | 1 | 4378 | 4283.681 | 62.6857 | 252,040 | 4282.673175 | 63 | 339 | 347 | GEVFNATRF | 0.000020000.0 | 343(NAT) | 1 | 1042.497 | 1039.508739 | 5711414 | 62.15 | 3 | 6 | 6 | 3 | 0 | 2 | 0 | 0 | 4282.65631 | 0.0169 | 3.93 | 0.54 |
| Chymotrypsin | 16 | 71 | 1596 | 1 | 4171 | 3669.4819 | 54.1076 | 243,031 | 3668.474075 | 63 | 339 | 347 | GEVFNATRF | 0.000020000.0 | 343(NAT) | 1 | 1042.497 | 1039.508739 | 5711414 | 62.15 | 1 | 6 | 6 | 3 | 0 | 0 | 0 | 1 | 3668.475648 | -0.0016 | -0.42 | -8.04 |
| Chymotrypsin | 16 | 71 | 1797 | 1 | 4403 | 3610.4237 | 63.7535 | 217,763 | 3609.415875 | 63 | 339 | 347 | GEVFNATRF | 0.000020000.0 | 343(NAT) | 1 | 1042.497 | 1039.508739 | 5711414 | 62.15 | 1 | 6 | 5 | 0 | 2 | 0 | 0 | 1 | 3609.413382 | 0.0025 | 0.69 | 1.61 |
| Chymotrypsin | 16 | 71 | 1797 | 1 | 4403 | 3610.4237 | 63.7535 | 217,763 | 3609.415875 | 63 | 339 | 347 | GEVFNATRF | 0.000020000.0 | 343(NAT) | 1 | 1042.497 | 1039.508739 | 5711414 | 62.15 | 2 | 5 | 5 | 1 | 1 | 1 | 0 | 1 | 3609.413381 | 0.0025 | 0.69 | 1.61 |
| Chymotrypsin | 16 | 71 | 1797 | 1 | 4403 | 3610.4237 | 63.7535 | 217,763 | 3609.415875 | 63 | 339 | 347 | GEVFNATRF | 0.000020000.0 | 343(NAT) | 1 | 1042.497 | 1039.508739 | 5711414 | 62.15 | 3 | 4 | 5 | 2 | 0 | 2 | 0 | 0 | 3609.41338 | 0.0025 | 0.69 | 1.61 |
| Chymotrypsin | 16 | 71 | 1799 | 1 | 4199 | 2647.1028 | 55.2919 | 217,452 | 2646.094975 | 63 | 339 | 347 | GEVFNATRF | 0.000020000.0 | 343(NAT) | 1 | 1042.497 | 1039.508739 | 5711414 | 62.15 | 1 | 4 | 4 | 1 | 0 | 0 | 0 | 1 | 2646.095436 | -0.0005 | -0.17 | -6.86 |
| Chymotrypsin | 16 | 71 | 1820 | 1 | 4203 | 2688.1294 | 55.4605 | 214,964 | 2687.121575 | 63 | 339 | 347 | GEVFNATRF | 0.000020000.0 | 343(NAT) | 1 | 1042.497 | 1039.508739 | 5711414 | 62.15 | 1 | 3 | 5 | 1 | 0 | 0 | 0 | 1 | 2687.121985 | -0.0004 | -0.15 | -6.69 |
| Chymotrypsin | 16 | 71 | 1869 | 1 | 4198 | 2590.0843 | 55.2502 | 208,951 | 2589.076475 | 63 | 339 | 347 | GEVFNATRF | 0.000020000.0 | 343(NAT) | 1 | 1042.497 | 1039.508739 | 5711414 | 62.15 | 1 | 4 | 3 | 2 | 0 | 0 | 0 | 1 | 2589.073972 | 0.0025 | 0.96 | -6.90 |
| Chymotrypsin | 16 | 71 | 2017 | 1 | 4379 | 4299.6683 | 62.7281 | 192,686 | 4298.660475 | 63 | 339 | 347 | GEVFNATRF | 0.000020000.0 | 343(NAT) | 1 | 1042.497 | 1039.508739 | 5711414 | 62.15 | 1 | 9 | 6 | 0 | 2 | 0 | 0 | 0 | 4298.651227 | 0.0092 | 2.15 | 0.58 |
| Chymotrypsin | 16 | 71 | 2017 | 1 | 4379 | 4299.6683 | 62.7281 | 192,686 | 4298.660475 | 63 | 339 | 347 | GEVFNATRF | 0.000020000.0 | 343(NAT) | 1 | 1042.497 | 1039.508739 | 5711414 | 62.15 | 2 | 8 | 6 | 1 | 1 | 1 | 0 | 0 | 4298.651226 | 0.0092 | 2.15 | 0.58 |
| Chymotrypsin | 16 | 71 | 2017 | 1 | 4379 | 4299.6683 | 62.7281 | 192,686 | 4298.660475 | 63 | 339 | 347 | GEVFNATRF | 0.000020000.0 | 343(NAT) | 1 | 1042.497 | 1039.508739 | 5711414 | 62.15 | 3 | 7 | 6 | 2 | 0 | 2 | 0 | 0 | 4298.651225 | 0.0093 | 2.15 | 0.58 |
| Chymotrypsin | 16 | 71 | 2037 | 1 | 4388 | 4121.6231 | 63.1108 | 190,668 | 4120.615275 | 63 | 339 | 347 | GEVFNATRF | 0.000020000.0 | 343(NAT) | 1 | 1042.497 | 1039.508739 | 5711414 | 62.15 | 1 | 7 | 6 | 1 | 2 | 0 | 0 | 1 | 4120.603488 | 0.0118 | 2.86 | 0.96 |
| Chymotrypsin | 16 | 71 | 2037 | 1 | 4388 | 4121.6231 | 63.1108 | 190,668 | 4120.615275 | 63 | 339 | 347 | GEVFNATRF | 0.000020000.0 | 343(NAT) | 1 | 1042.497 | 1039.508739 | 5711414 | 62.15 | 2 | 6 | 6 | 2 | 1 | 1 | 0 | 0 | 4120.603487 | 0.0118 | 2.86 | 0.96 |
| Chymotrypsin | 16 | 71 | 2037 | 1 | 4388 | 4121.6231 | 63.1108 | 190,668 | 4120.615275 | 63 | 339 | 347 | GEVFNATRF | 0.000020000.0 | 343(NAT) | 1 | 1042.497 | 1039.508739 | 5711414 | 62.15 | 3 | 5 | 6 | 3 | 0 | 2 | 0 | 0 | 4120.603486 | 0.0118 | 2.86 | 0.96 |
| Chymotrypsin | 16 | 71 | 2079 | 1 | 4182 | 3174.2869 | 54.5763 | 187,752 | 3173.279075 | 63 | 339 | 347 | GEVFNATRF | 0.000020000.0 | 343(NAT) | 1 | 1042.497 | 1039.508739 | 5711414 | 62.15 | 1 | 6 | 5 | 1 | 0 | 0 | 0 | 1 | 3173.280457 | -0.0014 | -0.43 | -7.57 |
| Chymotrypsin | 16 | 71 | 2161 | 1 | 4269 | 3465.3823 | 58.2744 | 181,647 | 3464.374475 | 63 | 339 | 347 | GEVFNATRF | 0.000020000.0 | 343(NAT) | 1 | 1042.497 | 1039.508739 | 5711414 | 62.15 | 1 | 6 | 5 | 1 | 1 | 0 | 0 | 1 | 3464.375874 | -0.0014 | -0.4 | -3.87 |
| Chymotrypsin | 16 | 71 | 2161 | 1 | 4269 | 3465.3823 | 58.2744 | 181,647 | 3464.374475 | 63 | 339 | 347 | GEVFNATRF | 0.000020000.0 | 343(NAT) | 1 | 1042.497 | 1039.508739 | 5711414 | 62.15 | 2 | 5 | 5 | 2 | 0 | 1 | 0 | 0 | 3464.375873 | -0.0014 | -0.4 | -3.87 |
| Chymotrypsin | 16 | 71 | 2274 | 1 | 4182 | 3523.4278 | 54.5763 | 171,441 | 3522.419975 | 63 | 339 | 347 | GEVFNATRF | 0.000020000.0 | 343(NAT) | 1 | 1042.497 | 1039.508739 | 5711414 | 62.15 | 1 | 6 | 6 | 2 | 0 | 0 | 0 | 1 | 3522.417739 | 0.0022 | 0.63 | -7.57 |
| Chymotrypsin | 16 | 71 | 2278 | 1 | 4349 | 5217.0139 | 61.4484 | 171,228 | 5216.006075 | 63 | 339 | 347 | GEVFNATRF | 0.000020000.0 | 343(NAT) | 1 | 1042.497 | 1039.508739 | 5711414 | 62.15 | 1 | 10 | 9 | 1 | 2 | 0 | 0 | 0 | 5216.000079 | 0.0060 | 1.14 | -0.70 |
| Chymotrypsin | 16 | 71 | 2278 | 1 | 4349 | 5217.0139 | 61.4484 | 171,228 | 5216.006075 | 63 | 339 | 347 | GEVFNATRF | 0.000020000.0 | 343(NAT) | 1 | 1042.497 | 1039.508739 | 5711414 | 62.15 | 2 | 9 | 9 | 2 | 1 | 1 | 0 | 0 | 5216.000078 | 0.0060 | 1.14 | -0.70 |
| Chymotrypsin | 16 | 71 | 2278 | 1 | 4349 | 5217.0139 | 61.4484 | 171,228 | 5216.006075 | 63 | 339 | 347 | GEVFNATRF | 0.000020000.0 | 343(NAT) | 1 | 1042.497 | 1039.508739 | 5711414 | 62.15 | 3 | 8 | 9 | 3 | 0 | 2 | 0 | 0 | 5216.000077 | 0.0060 | 1.14 | -0.70 |
| Chymotrypsin | 16 | 71 | 2330 | 1 | 4356 | 5054.9669 | 61.7476 | 166,462 | 5053.959075 | 63 | 339 | 347 | GEVFNATRF | 0.000020000.0 | 343(NAT) | 1 | 1042.497 | 1039.508739 | 5711414 | 62.15 | 1 | 9 | 9 | 1 | 2 | 0 | 0 | 0 | 5053.947255 | 0.0118 | 2.33 | -0.40 |
| Chymotrypsin | 16 | 71 | 2330 | 1 | 4356 | 5054.9669 | 61.7476 | 166,462 | 5053.959075 | 63 | 339 | 347 | GEVFNATRF | 0.000020000.0 | 343(NAT) | 1 | 1042.497 | 1039.508739 | 5711414 | 62.15 | 2 | 8 | 9 | 2 | 1 | 1 | 0 | 0 | 5053.947254 | 0.0118 | 2.33 | -0.40 |
| Chymotrypsin | 16 | 71 | 2330 | 1 | 4356 | 5054.9669 | 61.7476 | 166,462 | 5053.959075 | 63 | 339 | 347 | GEVFNATRF | 0.000020000.0 | 343(NAT) | 1 | 1042.497 | 1039.508739 | 5711414 | 62.15 | 3 | 7 | 9 | 3 | 0 | 2 | 0 | 0 | 5053.947253 | 0.0118 | 2.33 | -0.40 |
| Chymotrypsin | 16 | 71 | 2730 | 1 | 4177 | 3142.3003 | 54.3618 | 141,177 | 3141.292475 | 63 | 339 | 347 | GEVFNATRF | 0.000020000.0 | 343(NAT) | 1 | 1042.497 | 1039.508739 | 5711414 | 62.15 | 1 | 4 | 5 | 3 | 0 | 0 | 0 | 0 | 3141.290627 | 0.0018 | 0.58 | -7.79 |
| Chymotrypsin | 16 | 71 | 2832 | 1 | 4366 | 4582.7354 | 62.1721 | 135,140 | 4581.727575 | 63 | 339 | 347 | GEVFNATRF | 0.000020000.0 | 343(NAT) | 1 | 1042.497 | 1039.508739 | 5711414 | 62.15 | 1 | 10 | 5 | 2 | 0 | 2 | 1 | -1 | 4581.730324 | -0.0027 | -0.59 | 0.02 |
| Chymotrypsin | 16 | 71 | 2832 | 1 | 4366 | 4582.7354 | 62.1721 | 135,140 | 4581.727575 | 63 | 339 | 347 | GEVFNATRF | 0.000020000.0 | 343(NAT) | 1 | 1042.497 | 1039.508739 | 5711414 | 62.15 | 2 | 3 | 9 | 0 | 0 | 4 | 1 | -1 | 4581.742892 | -0.0153 | -3.34 | 0.02 |
| Chymotrypsin | 16 | 71 | 2951 | 1 | 4373 | 4486.7571 | 62.474 | 129,922 | 4485.749275 | 63 | 339 | 347 | GEVFNATRF | 0.000020000.0 | 343(NAT) | 1 | 1042.497 | 1039.508739 | 5711414 | 62.15 | 1 | 8 | 7 | 1 | 2 | 0 | 0 | 1 | 4485.735685 | 0.0136 | 3.02 | 0.33 |
| Chymotrypsin | 16 | 71 | 2951 | 1 | 4373 | 4486.7571 | 62.474 | 129,922 | 4485.749275 | 63 | 339 | 347 | GEVFNATRF | 0.000020000.0 | 343(NAT) | 1 | 1042.497 | 1039.508739 | 5711414 | 62.15 | 2 | 7 | 7 | 2 | 1 | 1 | 0 | 0 | 4485.735684 | 0.0136 | 3.02 | 0.33 |
| Chymotrypsin | 16 | 71 | 2951 | 1 | 4373 | 4486.7571 | 62.474 | 129,922 | 4485.749275 | 63 | 339 | 347 | GEVFNATRF | 0.000020000.0 | 343(NAT) | 1 | 1042.497 | 1039.508739 | 5711414 | 62.15 | 3 | 6 | 7 | 3 | 0 | 2 | 0 | 0 | 4485.735683 | 0.0136 | 3.03 | 0.33 |
| Chymotrypsin | 16 | 71 | 2993 | 1 | 4188 | 3377.3624 | 54.8278 | 127,360 | 3376.354575 | 63 | 339 | 347 | GEVFNATRF | 0.000020000.0 | 343(NAT) | 1 | 1042.497 | 1039.508739 | 5711414 | 62.15 | 1 | 6 | 6 | 1 | 0 | 0 | 0 | 1 | 3376.35983 | -0.0053 | -1.55 | -7.32 |
| Chymotrypsin | 16 | 71 | 3071 | 1 | 4202 | 2809.1614 | 55.4187 | 123,515 | 2808.153575 | 63 | 339 | 347 | GEVFNATRF | 0.000020000.0 | 343(NAT) | 1 | 1042.497 | 1039.508739 | 5711414 | 62.15 | 1 | 5 | 4 | 1 | 0 | 0 | 0 | 1 | 2808.14826 | 0.0053 | 1.89 | -6.73 |
| Chymotrypsin | 16 | 71 | 3198 | 1 | 4195 | 2955.2203 | 55.1217 | 117,972 | 2954.212475 | 63 | 339 | 347 | GEVFNATRF | 0.000020000.0 | 343(NAT) | 1 | 1042.497 | 1039.508739 | 5711414 | 62.15 | 1 | 5 | 4 | 2 | 0 | 0 | 0 | 1 | 2954.206169 | 0.0063 | 2.13 | -7.03 |
| Chymotrypsin | 16 | 71 | 3210 | 1 | 4374 | 4461.7284 | 62.513 | 117,396 | 4460.720575 | 63 | 339 | 347 | GEVFNATRF | 0.000020000.0 | 343(NAT) | 1 | 1042.497 | 1039.508739 | 5711414 | 62.15 | 1 | 9 | 6 | 1 | 1 | 1 | 0 | 1 | 4460.70405 | 0.0165 | 3.7 | 0.36 |
| Chymotrypsin | 16 | 71 | 3210 | 1 | 4374 | 4461.7284 | 62.513 | 117,396 | 4460.720575 | 63 | 339 | 347 | GEVFNATRF | 0.000020000.0 | 343(NAT) | 1 | 1042.497 | 1039.508739 | 5711414 | 62.15 | 2 | 10 | 6 | 0 | 2 | 0 | 0 | 0 | 4460.704051 | 0.0165 | 3.7 | 0.36 |
| Chymotrypsin | 16 | 71 | 3210 | 1 | 4374 | 4461.7284 | 62.513 | 117,396 | 4460.720575 | 63 | 339 | 347 | GEVFNATRF | 0.000020000.0 | 343(NAT) | 1 | 1042.497 | 1039.508739 | 5711414 | 62.15 | 3 | 8 | 6 | 2 | 0 | 2 | 0 | 0 | 4460.704049 | 0.0165 | 3.7 | 0.36 |
| Chymotrypsin | 16 | 71 | 3210 | 1 | 4374 | 4461.7284 | 62.513 | 117,396 | 4460.720575 | 63 | 339 | 347 | GEVFNATRF | 0.000020000.0 | 343(NAT) | 1 | 1042.497 | 1039.508739 | 5711414 | 62.15 | 4 | 2 | 8 | 2 | 3 | 1 | 1 | -1 | 4460.741771 | -0.0212 | -4.75 | 0.36 |
| Chymotrypsin | 16 | 71 | 3210 | 1 | 4374 | 4461.7284 | 62.513 | 117,396 | 4460.720575 | 63 | 339 | 347 | GEVFNATRF | 0.000020000.0 | 343(NAT) | 1 | 1042.497 | 1039.508739 | 5711414 | 62.15 | 5 | 3 | 8 | 1 | 4 | 0 | 1 | -1 | 4460.741772 | -0.0212 | -4.75 | 0.36 |
| Chymotrypsin | 16 | 71 | 3767 | 1 | 4379 | 4324.7094 | 62.7281 | 98,406 | 4323.701575 | 63 | 339 | 347 | GEVFNATRF | 0.000020000.0 | 343(NAT) | 1 | 1042.497 | 1039.508739 | 5711414 | 62.15 | 1 | 7 | 7 | 1 | 2 | 0 | 0 | 1 | 4323.682861 | 0.0187 | 4.32 | 0.58 |
| Chymotrypsin | 16 | 71 | 3767 | 1 | 4379 | 4324.7094 | 62.7281 | 98,406 | 4323.701575 | 63 | 339 | 347 | GEVFNATRF | 0.000020000.0 | 343(NAT) | 1 | 1042.497 | 1039.508739 | 5711414 | 62.15 | 2 | 6 | 7 | 2 | 1 | 1 | 0 | 0 | 4323.68286 | 0.0187 | 4.32 | 0.58 |
| Chymotrypsin | 16 | 71 | 3767 | 1 | 4379 | 4324.7094 | 62.7281 | 98,406 | 4323.701575 | 63 | 339 | 347 | GEVFNATRF | 0.000020000.0 | 343(NAT) | 1 | 1042.497 | 1039.508739 | 5711414 | 62.15 | 3 | 5 | 7 | 3 | 0 | 2 | 0 | 0 | 4323.682859 | 0.0187 | 4.32 | 0.58 |
| Chymotrypsin | 16 | 71 | 3797 | 1 | 4367 | 4689.8382 | 62.2157 | 97,325 | 4688.830375 | 63 | 339 | 347 | GEVFNATRF | 0.000020000.0 | 343(NAT) | 1 | 1042.497 | 1039.508739 | 5711414 | 62.15 | 1 | 8 | 8 | 1 | 2 | 0 | 0 | 0 | 4688.815058 | 0.0153 | 3.26 | 0.07 |
| Chymotrypsin | 16 | 71 | 3797 | 1 | 4367 | 4689.8382 | 62.2157 | 97,325 | 4688.830375 | 63 | 339 | 347 | GEVFNATRF | 0.000020000.0 | 343(NAT) | 1 | 1042.497 | 1039.508739 | 5711414 | 62.15 | 2 | 7 | 8 | 2 | 1 | 1 | 0 | 0 | 4688.815057 | 0.0153 | 3.26 | 0.07 |
| Chymotrypsin | 16 | 71 | 3797 | 1 | 4367 | 4689.8382 | 62.2157 | 97,325 | 4688.830375 | 63 | 339 | 347 | GEVFNATRF | 0.000020000.0 | 343(NAT) | 1 | 1042.497 | 1039.508739 | 5711414 | 62.15 | 3 | 6 | 8 | 3 | 0 | 2 | 0 | 0 | 4688.815056 | 0.0153 | 3.26 | 0.07 |
| Chymotrypsin | 16 | 71 | 3885 | 1 | 4362 | 4851.8954 | 62.0013 | 94,996 | 4850.887575 | 63 | 339 | 347 | GEVFNATRF | 0.000020000.0 | 343(NAT) | 1 | 1042.497 | 1039.508739 | 5711414 | 62.15 | 1 | 2 | 10 | 2 | 4 | 0 | 1 | -1 | 4850.905603 | -0.0180 | -3.71 | -0.15 |
| Chymotrypsin | 16 | 71 | 3885 | 1 | 4362 | 4851.8954 | 62.0013 | 94,996 | 4850.887575 | 63 | 339 | 347 | GEVFNATRF | 0.000020000.0 | 343(NAT) | 1 | 1042.497 | 1039.508739 | 5711414 | 62.15 | 2 | 9 | 8 | 1 | 2 | 0 | 0 | 0 | 4850.867882 | 0.0197 | 4.05 | -0.15 |
| Chymotrypsin | 16 | 71 | 3885 | 1 | 4362 | 4851.8954 | 62.0013 | 94,996 | 4850.887575 | 63 | 339 | 347 | GEVFNATRF | 0.000020000.0 | 343(NAT) | 1 | 1042.497 | 1039.508739 | 5711414 | 62.15 | 3 | 8 | 8 | 2 | 1 | 1 | 0 | 0 | 4850.867881 | 0.0197 | 4.05 | -0.15 |
| Chymotrypsin | 16 | 71 | 3885 | 1 | 4362 | 4851.8954 | 62.0013 | 94,996 | 4850.887575 | 63 | 339 | 347 | GEVFNATRF | 0.000020000.0 | 343(NAT) | 1 | 1042.497 | 1039.508739 | 5711414 | 62.15 | 4 | 7 | 8 | 3 | 0 | 2 | 0 | 0 | 4850.86788 | 0.0197 | 4.06 | -0.15 |
| Chymotrypsin | 16 | 71 | 4061 | 1 | 4362 | 4892.9161 | 62.0013 | 90,216 | 4891.908275 | 63 | 339 | 347 | GEVFNATRF | 0.000020000.0 | 343(NAT) | 1 | 1042.497 | 1039.508739 | 5711414 | 62.15 | 1 | 8 | 9 | 1 | 2 | 0 | 0 | 0 | 4891.894431 | 0.0138 | 2.83 | -0.15 |
| Chymotrypsin | 16 | 71 | 4061 | 1 | 4362 | 4892.9161 | 62.0013 | 90,216 | 4891.908275 | 63 | 339 | 347 | GEVFNATRF | 0.000020000.0 | 343(NAT) | 1 | 1042.497 | 1039.508739 | 5711414 | 62.15 | 2 | 7 | 9 | 2 | 1 | 1 | 0 | 0 | 4891.89443 | 0.0138 | 2.83 | -0.15 |
| Chymotrypsin | 16 | 71 | 4061 | 1 | 4362 | 4892.9161 | 62.0013 | 90,216 | 4891.908275 | 63 | 339 | 347 | GEVFNATRF | 0.000020000.0 | 343(NAT) | 1 | 1042.497 | 1039.508739 | 5711414 | 62.15 | 3 | 6 | 9 | 3 | 0 | 2 | 0 | 0 | 4891.894429 | 0.0138 | 2.83 | -0.15 |
| Chymotrypsin | 16 | 71 | 4245 | 1 | 4265 | 3773.4891 | 58.1019 | 85,241 | 3772.481275 | 63 | 339 | 347 | GEVFNATRF | 0.000020000.0 | 343(NAT) | 1 | 1042.497 | 1039.508739 | 5711414 | 62.15 | 1 | 6 | 5 | 3 | 0 | 1 | 0 | 1 | 3772.486606 | -0.0053 | -1.41 | -4.05 |
| Chymotrypsin | 16 | 71 | 4245 | 1 | 4265 | 3773.4891 | 58.1019 | 85,241 | 3772.481275 | 63 | 339 | 347 | GEVFNATRF | 0.000020000.0 | 343(NAT) | 1 | 1042.497 | 1039.508739 | 5711414 | 62.15 | 2 | 7 | 5 | 2 | 1 | 0 | 0 | 0 | 3772.486607 | -0.0053 | -1.41 | -4.05 |
| Chymotrypsin | 16 | 71 | 4280 | 1 | 4152 | 4180.6743 | 53.2993 | 84,395 | 4179.666475 | 63 | 339 | 347 | GEVFNATRF | 0.000020000.0 | 343(NAT) | 1 | 1042.497 | 1039.508739 | 5711414 | 62.15 | 1 | 7 | 7 | 4 | 0 | 0 | 0 | 0 | 4179.665754 | 0.0007 | 0.17 | -8.85 |
| Chymotrypsin | 16 | 71 | 4363 | 1 | 4376 | 4527.7789 | 62.6003 | 82,350 | 4526.771075 | 63 | 339 | 347 | GEVFNATRF | 0.000020000.0 | 343(NAT) | 1 | 1042.497 | 1039.508739 | 5711414 | 62.15 | 1 | 7 | 8 | 1 | 2 | 0 | 0 | 0 | 4526.762234 | 0.0088 | 1.95 | 0.45 |
| Chymotrypsin | 16 | 71 | 4363 | 1 | 4376 | 4527.7789 | 62.6003 | 82,350 | 4526.771075 | 63 | 339 | 347 | GEVFNATRF | 0.000020000.0 | 343(NAT) | 1 | 1042.497 | 1039.508739 | 5711414 | 62.15 | 2 | 6 | 8 | 2 | 1 | 1 | 0 | 0 | 4526.762233 | 0.0088 | 1.95 | 0.45 |
| Chymotrypsin | 16 | 71 | 4363 | 1 | 4376 | 4527.7789 | 62.6003 | 82,350 | 4526.771075 | 63 | 339 | 347 | GEVFNATRF | 0.000020000.0 | 343(NAT) | 1 | 1042.497 | 1039.508739 | 5711414 | 62.15 | 3 | 5 | 8 | 3 | 0 | 2 | 0 | 0 | 4526.762232 | 0.0088 | 1.95 | 0.45 |
| Chymotrypsin | 16 | 71 | 4384 | 1 | 4376 | 4730.848 | 62.6003 | 81,859 | 4729.840175 | 63 | 339 | 347 | GEVFNATRF | 0.000020000.0 | 343(NAT) | 1 | 1042.497 | 1039.508739 | 5711414 | 62.15 | 1 | 5 | 9 | 3 | 0 | 2 | 0 | 0 | 4729.841605 | -0.0014 | -0.3 | 0.45 |
| Chymotrypsin | 16 | 71 | 4384 | 1 | 4376 | 4730.848 | 62.6003 | 81,859 | 4729.840175 | 63 | 339 | 347 | GEVFNATRF | 0.000020000.0 | 343(NAT) | 1 | 1042.497 | 1039.508739 | 5711414 | 62.15 | 2 | 6 | 9 | 2 | 1 | 1 | 0 | 0 | 4729.841606 | -0.0014 | -0.3 | 0.45 |
| Chymotrypsin | 16 | 71 | 4384 | 1 | 4376 | 4730.848 | 62.6003 | 81,859 | 4729.840175 | 63 | 339 | 347 | GEVFNATRF | 0.000020000.0 | 343(NAT) | 1 | 1042.497 | 1039.508739 | 5711414 | 62.15 | 3 | 7 | 9 | 1 | 2 | 0 | 0 | 0 | 4729.841607 | -0.0014 | -0.3 | 0.45 |
| Chymotrypsin | 16 | 71 | 4648 | 1 | 4381 | 4381.7226 | 62.8147 | 76,052 | 4380.714775 | 63 | 339 | 347 | GEVFNATRF | 0.000020000.0 | 343(NAT) | 1 | 1042.497 | 1039.508739 | 5711414 | 62.15 | 1 | 7 | 8 | 0 | 2 | 0 | 0 | 0 | 4380.704325 | 0.0105 | 2.38 | 0.67 |
| Chymotrypsin | 16 | 71 | 4648 | 1 | 4381 | 4381.7226 | 62.8147 | 76,052 | 4380.714775 | 63 | 339 | 347 | GEVFNATRF | 0.000020000.0 | 343(NAT) | 1 | 1042.497 | 1039.508739 | 5711414 | 62.15 | 2 | 6 | 8 | 1 | 1 | 1 | 0 | 0 | 4380.704324 | 0.0105 | 2.38 | 0.67 |
| Chymotrypsin | 16 | 71 | 4648 | 1 | 4381 | 4381.7226 | 62.8147 | 76,052 | 4380.714775 | 63 | 339 | 347 | GEVFNATRF | 0.000020000.0 | 343(NAT) | 1 | 1042.497 | 1039.508739 | 5711414 | 62.15 | 3 | 5 | 8 | 2 | 0 | 2 | 0 | 0 | 4380.704323 | 0.0105 | 2.38 | 0.67 |
| Chymotrypsin | 16 | 71 | 4708 | 1 | 4277 | 3319.3257 | 58.6235 | 74,595 | 3318.317875 | 63 | 339 | 347 | GEVFNATRF | 0.000020000.0 | 343(NAT) | 1 | 1042.497 | 1039.508739 | 5711414 | 62.15 | 1 | 5 | 5 | 1 | 0 | 1 | 0 | 1 | 3318.317964 | -0.0001 | -0.02 | -3.52 |
| Chymotrypsin | 16 | 71 | 4708 | 1 | 4277 | 3319.3257 | 58.6235 | 74,595 | 3318.317875 | 63 | 339 | 347 | GEVFNATRF | 0.000020000.0 | 343(NAT) | 1 | 1042.497 | 1039.508739 | 5711414 | 62.15 | 2 | 6 | 5 | 0 | 1 | 0 | 0 | 1 | 3318.317965 | -0.0001 | -0.02 | -3.52 |
| Chymotrypsin | 16 | 71 | 4760 | 1 | 4388 | 4365.7309 | 63.1108 | 73,631 | 4364.723075 | 63 | 339 | 347 | GEVFNATRF | 0.000020000.0 | 343(NAT) | 1 | 1042.497 | 1039.508739 | 5711414 | 62.15 | 1 | 6 | 8 | 1 | 2 | 0 | 0 | 0 | 4364.70941 | 0.0137 | 3.13 | 0.96 |
| Chymotrypsin | 16 | 71 | 4760 | 1 | 4388 | 4365.7309 | 63.1108 | 73,631 | 4364.723075 | 63 | 339 | 347 | GEVFNATRF | 0.000020000.0 | 343(NAT) | 1 | 1042.497 | 1039.508739 | 5711414 | 62.15 | 2 | 5 | 8 | 2 | 1 | 1 | 0 | 0 | 4364.709409 | 0.0137 | 3.13 | 0.96 |
| Chymotrypsin | 16 | 71 | 4760 | 1 | 4388 | 4365.7309 | 63.1108 | 73,631 | 4364.723075 | 63 | 339 | 347 | GEVFNATRF | 0.000020000.0 | 343(NAT) | 1 | 1042.497 | 1039.508739 | 5711414 | 62.15 | 3 | 4 | 8 | 3 | 0 | 2 | 0 | 0 | 4364.709408 | 0.0137 | 3.13 | 0.96 |
| Chymotrypsin | 16 | 71 | 4983 | 1 | 4282 | 3684.4515 | 58.8341 | 68,994 | 3683.443675 | 63 | 339 | 347 | GEVFNATRF | 0.000020000.0 | 343(NAT) | 1 | 1042.497 | 1039.508739 | 5711414 | 62.15 | 1 | 7 | 6 | 0 | 1 | 0 | 0 | 1 | 3683.450162 | -0.0065 | -1.76 | -3.31 |
| Chymotrypsin | 16 | 71 | 4983 | 1 | 4282 | 3684.4515 | 58.8341 | 68,994 | 3683.443675 | 63 | 339 | 347 | GEVFNATRF | 0.000020000.0 | 343(NAT) | 1 | 1042.497 | 1039.508739 | 5711414 | 62.15 | 2 | 6 | 6 | 1 | 0 | 1 | 0 | 0 | 3683.450161 | -0.0065 | -1.76 | -3.31 |
| Chymotrypsin | 16 | 71 | 5177 | 1 | 4368 | 4632.8137 | 62.2594 | 65,457 | 4631.805875 | 63 | 339 | 347 | GEVFNATRF | 0.000020000.0 | 343(NAT) | 1 | 1042.497 | 1039.508739 | 5711414 | 62.15 | 1 | 8 | 7 | 2 | 2 | 0 | 0 | 0 | 4631.793594 | 0.0123 | 2.65 | 0.11 |
| Chymotrypsin | 16 | 71 | 5177 | 1 | 4368 | 4632.8137 | 62.2594 | 65,457 | 4631.805875 | 63 | 339 | 347 | GEVFNATRF | 0.000020000.0 | 343(NAT) | 1 | 1042.497 | 1039.508739 | 5711414 | 62.15 | 2 | 7 | 7 | 3 | 1 | 1 | 0 | 0 | 4631.793593 | 0.0123 | 2.65 | 0.11 |
| Chymotrypsin | 16 | 71 | 5177 | 1 | 4368 | 4632.8137 | 62.2594 | 65,457 | 4631.805875 | 63 | 339 | 347 | GEVFNATRF | 0.000020000.0 | 343(NAT) | 1 | 1042.497 | 1039.508739 | 5711414 | 62.15 | 3 | 6 | 7 | 4 | 0 | 2 | 0 | 0 | 4631.793592 | 0.0123 | 2.65 | 0.11 |
| Chymotrypsin | 16 | 71 | 5338 | 1 | 4375 | 4470.7481 | 62.5564 | 62,802 | 4469.740275 | 63 | 339 | 347 | GEVFNATRF | 0.000020000.0 | 343(NAT) | 1 | 1042.497 | 1039.508739 | 5711414 | 62.15 | 1 | 5 | 7 | 4 | 0 | 2 | 0 | 0 | 4469.740768 | -0.0005 | -0.11 | 0.41 |
| Chymotrypsin | 16 | 71 | 5338 | 1 | 4375 | 4470.7481 | 62.5564 | 62,802 | 4469.740275 | 63 | 339 | 347 | GEVFNATRF | 0.000020000.0 | 343(NAT) | 1 | 1042.497 | 1039.508739 | 5711414 | 62.15 | 2 | 6 | 7 | 3 | 1 | 1 | 0 | 0 | 4469.740769 | -0.0005 | -0.11 | 0.41 |
| Chymotrypsin | 16 | 71 | 5338 | 1 | 4375 | 4470.7481 | 62.5564 | 62,802 | 4469.740275 | 63 | 339 | 347 | GEVFNATRF | 0.000020000.0 | 343(NAT) | 1 | 1042.497 | 1039.508739 | 5711414 | 62.15 | 3 | 7 | 7 | 2 | 2 | 0 | 0 | 0 | 4469.74077 | -0.0005 | -0.11 | 0.41 |
| Chymotrypsin | 16 | 71 | 5537 | 1 | 4150 | 4326.7423 | 53.2082 | 59,591 | 4325.734475 | 63 | 339 | 347 | GEVFNATRF | 0.000020000.0 | 343(NAT) | 1 | 1042.497 | 1039.508739 | 5711414 | 62.15 | | | | | | | | | | | | |
| Chymotrypsin | 16 | 71 | 5641 | 1 | 4383 | 4178.6382 | 62.8992 | 57,855 | 4177.630375 | 63 | 339 | 347 | GEVFNATRF | 0.000020000.0 | 343(NAT) | 1 | 1042.497 | 1039.508739 | 5711414 | 62.15 | 1 | 7 | 7 | 0 | 2 | 0 | 0 | 0 | 4177.624952 | 0.0054 | 1.29 | 0.75 |
| Chymotrypsin | 16 | 71 | 5641 | 1 | 4383 | 4178.6382 | 62.8992 | 57,855 | 4177.630375 | 63 | 339 | 347 | GEVFNATRF | 0.000020000.0 | 343(NAT) | 1 | 1042.497 | 1039.508739 | 5711414 | 62.15 | 2 | 6 | 7 | 1 | 1 | 1 | 0 | 0 | 4177.624951 | 0.0054 | 1.29 | 0.75 |
| Chymotrypsin | 16 | 71 | 5641 | 1 | 4383 | 4178.6382 | 62.8992 | 57,855 | 4177.630375 | 63 | 339 | 347 | GEVFNATRF | 0.000020000.0 | 343(NAT) | 1 | 1042.497 | 1039.508739 | 5711414 | 62.15 | 3 | 5 | 7 | 2 | 0 | 2 | 0 | 0 | 4177.62495 | 0.0054 | 1.29 | 0.75 |
| Chymotrypsin | 16 | 71 | 5792 | 1 | 4279 | 3246.3058 | 58.7076 | 55,203 | 3245.297975 | 63 | 339 | 347 | GEVFNATRF | 0.000020000.0 | 343(NAT) | 1 | 1042.497 | 1039.508739 | 5711414 | 62.15 | 1 | 5 | 4 | 2 | 1 | 0 | 0 | 1 | 3245.301586 | -0.0036 | -1.11 | -3.44 |
| Chymotrypsin | 16 | 71 | 5792 | 1 | 4279 | 3246.3058 | 58.7076 | 55,203 | 3245.297975 | 63 | 339 | 347 | GEVFNATRF | 0.000020000.0 | 343(NAT) | 1 | 1042.497 | 1039.508739 | 5711414 | 62.15 | 2 | 4 | 4 | 3 | 0 | 1 | 0 | 0 | 3245.301585 | -0.0036 | -1.11 | -3.44 |
| Chymotrypsin | 16 | 71 | 6033 | 1 | 4367 | 4607.7905 | 62.2157 | 51,501 | 4606.782675 | 63 | 339 | 347 | GEVFNATRF | 0.000020000.0 | 343(NAT) | 1 | 1042.497 | 1039.508739 | 5711414 | 62.15 | 1 | 2 | 10 | 0 | 1 | 3 | 1 | -1 | 4606.774527 | 0.0081 | 1.76 | 0.07 |
| Chymotrypsin | 16 | 71 | 6033 | 1 | 4367 | 4607.7905 | 62.2157 | 51,501 | 4606.782675 | 63 | 339 | 347 | GEVFNATRF | 0.000020000.0 | 343(NAT) | 1 | 1042.497 | 1039.508739 | 5711414 | 62.15 | 2 | 2 | 8 | 3 | 3 | 1 | 1 | -1 | 4606.79968 | -0.0170 | -3.69 | 0.07 |
| Chymotrypsin | 16 | 71 | 6033 | 1 | 4367 | 4607.7905 | 62.2157 | 51,501 | 4606.782675 | 63 | 339 | 347 | GEVFNATRF | 0.000020000.0 | 343(NAT) | 1 | 1042.497 | 1039.508739 | 5711414 | 62.15 | 3 | 3 | 8 | 2 | 4 | 0 | 1 | -1 | 4606.799681 | -0.0170 | -3.69 | 0.07 |
| Chymotrypsin | 16 | 71 | 6033 | 1 | 4367 | 4607.7905 | 62.2157 | 51,501 | 4606.782675 | 63 | 339 | 347 | GEVFNATRF | 0.000020000.0 | 343(NAT) | 1 | 1042.497 | 1039.508739 | 5711414 | 62.15 | 4 | 10 | 6 | 1 | 2 | 0 | 0 | 0 | 4606.76196 | 0.0207 | 4.49 | 0.07 |
| Chymotrypsin | 16 | 71 | 6033 | 1 | 4367 | 4607.7905 | 62.2157 | 51,501 | 4606.782675 | 63 | 339 | 347 | GEVFNATRF | 0.000020000.0 | 343(NAT) | 1 | 1042.497 | 1039.508739 | 5711414 | 62.15 | 5 | 9 | 6 | 2 | 1 | 1 | 0 | 0 | 4606.761959 | 0.0207 | 4.49 | 0.07 |
| Chymotrypsin | 16 | 71 | 6033 | 1 | 4367 | 4607.7905 | 62.2157 | 51,501 | 4606.782675 | 63 | 339 | 347 | GEVFNATRF | 0.000020000.0 | 343(NAT) | 1 | 1042.497 | 1039.508739 | 5711414 | 62.15 | 6 | 8 | 6 | 3 | 0 | 2 | 0 | 0 | 4606.761958 | 0.0207 | 4.49 | 0.07 |
| Chymotrypsin | 16 | 71 | 6249 | 1 | 4344 | 5363.0881 | 61.2384 | 48,516 | 5362.080275 | 63 | 339 | 347 | GEVFNATRF | 0.000020000.0 | 343(NAT) | 1 | 1042.497 | 1039.508739 | 5711414 | 62.15 | 1 | 10 | 9 | 2 | 2 | 0 | 0 | 0 | 5362.057988 | 0.0223 | 4.15 | -0.91 |
| Chymotrypsin | 16 | 71 | 6249 | 1 | 4344 | 5363.0881 | 61.2384 | 48,516 | 5362.080275 | 63 | 339 | 347 | GEVFNATRF | 0.000020000.0 | 343(NAT) | 1 | 1042.497 | 1039.508739 | 5711414 | 62.15 | 2 | 9 | 9 | 3 | 1 | 1 | 0 | 0 | 5362.057987 | 0.0223 | 4.15 | -0.91 |
| Chymotrypsin | 16 | 71 | 6249 | 1 | 4344 | 5363.0881 | 61.2384 | 48,516 | 5362.080275 | 63 | 339 | 347 | GEVFNATRF | 0.000020000.0 | 343(NAT) | 1 | 1042.497 | 1039.508739 | 5711414 | 62.15 | 3 | 8 | 9 | 4 | 0 | 2 | 0 | 0 | 5362.057986 | 0.0223 | 4.15 | -0.91 |
| Chymotrypsin | 16 | 71 | 6262 | 1 | 4358 | 5095.987 | 61.8327 | 48,311 | 5094.979175 | 63 | 339 | 347 | GEVFNATRF | 0.000020000.0 | 343(NAT) | 1 | 1042.497 | 1039.508739 | 5711414 | 62.15 | 1 | 8 | 10 | 1 | 2 | 0 | 0 | 0 | 5094.973804 | 0.0054 | 1.05 | -0.32 |
| Chymotrypsin | 16 | 71 | 6262 | 1 | 4358 | 5095.987 | 61.8327 | 48,311 | 5094.979175 | 63 | 339 | 347 | GEVFNATRF | 0.000020000.0 | 343(NAT) | 1 | 1042.497 | 1039.508739 | 5711414 | 62.15 | 2 | 7 | 10 | 2 | 1 | 1 | 0 | 0 | 5094.973803 | 0.0054 | 1.05 | -0.32 |
| Chymotrypsin | 16 | 71 | 6262 | 1 | 4358 | 5095.987 | 61.8327 | 48,311 | 5094.979175 | 63 | 339 | 347 | GEVFNATRF | 0.000020000.0 | 343(NAT) | 1 | 1042.497 | 1039.508739 | 5711414 | 62.15 | 3 | 6 | 10 | 3 | 0 | 2 | 0 | 0 | 5094.973802 | 0.0054 | 1.05 | -0.32 |
| Chymotrypsin | 16 | 71 | 6391 | 1 | 4351 | 5201.0442 | 61.5339 | 46,657 | 5200.036375 | 63 | 339 | 347 | GEVFNATRF | 0.000020000.0 | 343(NAT) | 1 | 1042.497 | 1039.508739 | 5711414 | 62.15 | | | | | | | | | | | | |
| Chymotrypsin | 16 | 71 | 6401 | 1 | 4157 | 4034.6176 | 53.5109 | 46,540 | 4033.609775 | 63 | 339 | 347 | GEVFNATRF | 0.000020000.0 | 343(NAT) | 1 | 1042.497 | 1039.508739 | 5711414 | 62.15 | 1 | 7 | 7 | 3 | 0 | 0 | 0 | 0 | 4033.607845 | 0.0019 | 0.47 | -8.64 |
| Chymotrypsin | 16 | 71 | 6478 | 1 | 4392 | 4219.667 | 63.2822 | 45,314 | 4218.659175 | 63 | 339 | 347 | GEVFNATRF | 0.000020000.0 | 343(NAT) | 1 | 1042.497 | 1039.508739 | 5711414 | 62.15 | 1 | 6 | 8 | 0 | 2 | 0 | 0 | 0 | 4218.651501 | 0.0077 | 1.81 | 1.13 |
| Chymotrypsin | 16 | 71 | 6478 | 1 | 4392 | 4219.667 | 63.2822 | 45,314 | 4218.659175 | 63 | 339 | 347 | GEVFNATRF | 0.000020000.0 | 343(NAT) | 1 | 1042.497 | 1039.508739 | 5711414 | 62.15 | 2 | 5 | 8 | 1 | 1 | 1 | 0 | 0 | 4218.6515 | 0.0077 | 1.81 | 1.13 |
| Chymotrypsin | 16 | 71 | 6478 | 1 | 4392 | 4219.667 | 63.2822 | 45,314 | 4218.659175 | 63 | 339 | 347 | GEVFNATRF | 0.000020000.0 | 343(NAT) | 1 | 1042.497 | 1039.508739 | 5711414 | 62.15 | 3 | 4 | 8 | 2 | 0 | 2 | 0 | 0 | 4218.651499 | 0.0077 | 1.81 | 1.13 |
| Chymotrypsin | 16 | 71 | 6716 | 1 | 4374 | 4568.8037 | 62.513 | 42,111 | 4567.795875 | 63 | 339 | 347 | GEVFNATRF | 0.000020000.0 | 343(NAT) | 1 | 1042.497 | 1039.508739 | 5711414 | 62.15 | 1 | 6 | 9 | 1 | 2 | 0 | 0 | 0 | 4567.788783 | 0.0071 | 1.55 | 0.36 |
| Chymotrypsin | 16 | 71 | 6716 | 1 | 4374 | 4568.8037 | 62.513 | 42,111 | 4567.795875 | 63 | 339 | 347 | GEVFNATRF | 0.000020000.0 | 343(NAT) | 1 | 1042.497 | 1039.508739 | 5711414 | 62.15 | 2 | 5 | 9 | 2 | 1 | 1 | 0 | 0 | 4567.788782 | 0.0071 | 1.55 | 0.36 |
| Chymotrypsin | 16 | 71 | 6716 | 1 | 4374 | 4568.8037 | 62.513 | 42,111 | 4567.795875 | 63 | 339 | 347 | GEVFNATRF | 0.000020000.0 | 343(NAT) | 1 | 1042.497 | 1039.508739 | 5711414 | 62.15 | 3 | 4 | 9 | 3 | 0 | 2 | 0 | 0 | 4567.788781 | 0.0071 | 1.55 | 0.36 |
| Chymotrypsin | 16 | 71 | 7185 | 1 | 4267 | 3976.5758 | 58.1897 | 35,471 | 3975.567975 | 63 | 339 | 347 | GEVFNATRF | 0.000020000.0 | 343(NAT) | 1 | 1042.497 | 1039.508739 | 5711414 | 62.15 | 1 | 7 | 6 | 2 | 1 | 0 | 0 | 0 | 3975.56598 | 0.0020 | 0.5 | -3.96 |
| Chymotrypsin | 16 | 71 | 7185 | 1 | 4267 | 3976.5758 | 58.1897 | 35,471 | 3975.567975 | 63 | 339 | 347 | GEVFNATRF | 0.000020000.0 | 343(NAT) | 1 | 1042.497 | 1039.508739 | 5711414 | 62.15 | 2 | 6 | 6 | 3 | 0 | 1 | 0 | 0 | 3975.565979 | 0.0020 | 0.5 | -3.96 |
| Chymotrypsin | 16 | 71 | 8032 | 1 | 4163 | 3977.5949 | 53.7679 | 24,794 | 3976.587075 | 63 | 339 | 347 | GEVFNATRF | 0.000020000.0 | 343(NAT) | 1 | 1042.497 | 1039.508739 | 5711414 | 62.15 | 1 | 7 | 6 | 4 | 0 | 0 | 0 | 1 | 3976.586381 | 0.0007 | 0.17 | -8.38 |
| Chymotrypsin | 16 | 71 | 8032 | 1 | 4163 | 3977.5949 | 53.7679 | 24,794 | 3976.587075 | 63 | 339 | 347 | GEVFNATRF | 0.000020000.0 | 343(NAT) | 1 | 1042.497 | 1039.508739 | 5711414 | 62.15 | 2 | 2 | 10 | 0 | 2 | 0 | 1 | -1 | 3976.598951 | -0.0119 | -2.98 | -8.38 |
| Chymotrypsin | 21 | 39 | 54 | 1 | 3622 | 2580.0331 | 30.7174 | 5,121,020 | 2579.025275 | 1006 | 343 | 347 | NATRF | - | 343(NAT) | 1 | | 607.30779 | 3442227.5 | 39.85 | 1 | 5 | 5 | 1 | 0 | 0 | 0 | 1 | 2579.026684 | -0.0014 | -0.54 | -9.13 |
| Chymotrypsin | 21 | 39 | 77 | 1 | 3596 | 2376.9537 | 29.5861 | 3,706,630 | 2375.945875 | 1006 | 343 | 347 | NATRF | - | 343(NAT) | 1 | | 607.30779 | 3442227.5 | 39.85 | 1 | 5 | 4 | 1 | 0 | 0 | 0 | 1 | 2375.947311 | -0.0014 | -0.6 | -10.26 |
| Chymotrypsin | 21 | 39 | 139 | 1 | 3597 | 2726.091 | 29.6297 | 2,245,450 | 2725.083175 | 1006 | 343 | 347 | NATRF | - | 343(NAT) | 1 | | 607.30779 | 3442227.5 | 39.85 | 1 | 5 | 5 | 2 | 0 | 0 | 0 | 1 | 2725.084593 | -0.0014 | -0.52 | -10.22 |
| Chymotrypsin | 21 | 39 | 191 | 1 | 3570 | 2872.1493 | 28.4748 | 1,693,870 | 2871.141475 | 1006 | 343 | 347 | NATRF | - | 343(NAT) | 1 | | 607.30779 | 3442227.5 | 39.85 | 1 | 5 | 5 | 3 | 0 | 0 | 0 | 1 | 2871.142502 | -0.0010 | -0.35 | -11.37 |
| Chymotrypsin | 21 | 39 | 341 | 1 | 3626 | 2417.9804 | 30.8857 | 1,007,260 | 2416.972575 | 1006 | 343 | 347 | NATRF | - | 343(NAT) | 1 | | 607.30779 | 3442227.5 | 39.85 | 1 | 4 | 5 | 1 | 0 | 0 | 0 | 1 | 2416.97386 | -0.0013 | -0.53 | -8.96 |
| Chymotrypsin | 21 | 39 | 491 | 1 | 3618 | 2830.1024 | 30.5421 | 728,179 | 2829.094575 | 1006 | 343 | 347 | NATRF | - | 343(NAT) | 1 | | 607.30779 | 3442227.5 | 39.85 | 1 | 5 | 4 | 2 | 0 | 1 | 0 | 1 | 2829.095551 | -0.0010 | -0.34 | -9.30 |
| Chymotrypsin | 21 | 39 | 491 | 1 | 3618 | 2830.1024 | 30.5421 | 728,179 | 2829.094575 | 1006 | 343 | 347 | NATRF | - | 343(NAT) | 1 | | 607.30779 | 3442227.5 | 39.85 | 2 | 6 | 4 | 1 | 1 | 0 | 0 | 1 | 2829.095552 | -0.0010 | -0.34 | -9.30 |
| Chymotrypsin | 21 | 39 | 578 | 1 | 3653 | 2684.0438 | 32.0502 | 621,184 | 2683.035975 | 1006 | 343 | 347 | NATRF | - | 343(NAT) | 1 | | 607.30779 | 3442227.5 | 39.85 | 1 | 5 | 4 | 1 | 0 | 1 | 0 | 1 | 2683.037642 | -0.0017 | -0.62 | -7.80 |
| Chymotrypsin | 21 | 39 | 578 | 1 | 3653 | 2684.0438 | 32.0502 | 621,184 | 2683.035975 | 1006 | 343 | 347 | NATRF | - | 343(NAT) | 1 | | 607.30779 | 3442227.5 | 39.85 | 2 | 6 | 4 | 0 | 1 | 0 | 0 | 1 | 2683.037643 | -0.0017 | -0.62 | -7.80 |
| Chymotrypsin | 21 | 39 | 610 | 1 | 3608 | 2742.0861 | 30.1076 | 593,233 | 2741.078275 | 1006 | 343 | 347 | NATRF | - | 343(NAT) | 1 | | 607.30779 | 3442227.5 | 39.85 | 1 | 6 | 5 | 1 | 0 | 0 | 0 | 1 | 2741.079508 | -0.0012 | -0.44 | -9.74 |
| Chymotrypsin | 21 | 39 | 665 | 1 | 3556 | 3180.2592 | 27.8743 | 546,937 | 3179.251375 | 1006 | 343 | 347 | NATRF | - | 343(NAT) | 1 | | 607.30779 | 3442227.5 | 39.85 | 1 | 6 | 5 | 4 | 0 | 0 | 0 | 0 | 3179.253235 | -0.0019 | -0.58 | -11.97 |
| Chymotrypsin | 21 | 39 | 816 | 1 | 3598 | 2564.0377 | 29.6741 | 456,432 | 2563.029875 | 1006 | 343 | 347 | NATRF | - | 343(NAT) | 1 | | 607.30779 | 3442227.5 | 39.85 | 1 | 4 | 5 | 2 | 0 | 0 | 0 | 1 | 2563.031769 | -0.0019 | -0.73 | -10.17 |
| Chymotrypsin | 21 | 39 | 858 | 1 | 3598 | 2214.9014 | 29.6741 | 437,436 | 2213.893575 | 1006 | 343 | 347 | NATRF | - | 343(NAT) | 1 | | 607.30779 | 3442227.5 | 39.85 | 1 | 4 | 4 | 1 | 0 | 0 | 0 | 1 | 2213.894487 | -0.0009 | -0.41 | -10.17 |
| Chymotrypsin | 21 | 39 | 874 | 1 | 3578 | 2523.0127 | 28.8163 | 426,356 | 2522.004875 | 1006 | 343 | 347 | NATRF | - | 343(NAT) | 1 | | 607.30779 | 3442227.5 | 39.85 | 1 | 5 | 4 | 2 | 0 | 0 | 0 | 1 | 2522.00522 | -0.0003 | -0.13 | -11.03 |

| | | | | | | | | | | | | | | | | | | | | | | | | | | | | | | | | |
|---|---|---|---|---|---|---|---|---|---|---|---|---|---|---|---|---|---|---|---|---|---|---|---|---|---|---|---|---|---|---|---|---|
| Chymotrypsin | 21 | 39 | 1067 | 1 | 3599 | 2888.1432 | 29.7188 | 356,405 | 2887.135375 | 1006 | 343 | 347 | NATRF | - | 343(NAT) | 1 | | 607.30779 | 3442227.5 | 39.85 | 1 | 6 | 5 | 2 | 0 | 0 | 0 | 1 | 2887.137417 | -0.0020 | -0.7 | -10.13 |
| Chymotrypsin | 21 | 39 | 1498 | 1 | 3622 | 3033.1827 | 30.7174 | 257,489 | 3032.174875 | 1006 | 343 | 347 | NATRF | - | 343(NAT) | 1 | | 607.30779 | 3442227.5 | 39.85 | 1 | 6 | 5 | 1 | 1 | 0 | 0 | 1 | 3032.174925 | -0.0001 | -0.01 | -9.13 |
| Chymotrypsin | 21 | 39 | 1498 | 1 | 3622 | 3033.1827 | 30.7174 | 257,489 | 3032.174875 | 1006 | 343 | 347 | NATRF | - | 343(NAT) | 1 | | 607.30779 | 3442227.5 | 39.85 | 2 | 5 | 5 | 2 | 0 | 1 | 0 | 0 | 3032.174924 | 0.0000 | -0.01 | -9.13 |
| Chymotrypsin | 21 | 39 | 1697 | 1 | 3600 | 2052.8489 | 29.7623 | 230,353 | 2051.841075 | 1006 | 343 | 347 | NATRF | - | 343(NAT) | 1 | | 607.30779 | 3442227.5 | 39.85 | 1 | 3 | 4 | 1 | 0 | 0 | 0 | 1 | 2051.841663 | -0.0006 | -0.28 | -10.08 |
| Chymotrypsin | 21 | 39 | 1750 | 1 | 3627 | 2742.0853 | 30.9274 | 222,107 | 2741.077475 | 1006 | 343 | 347 | NATRF | - | 343(NAT) | 1 | | 607.30779 | 3442227.5 | 39.85 | 1 | 6 | 5 | 1 | 0 | 0 | 0 | 1 | 2741.079508 | -0.0020 | -0.74 | -8.92 |
| Chymotrypsin | 21 | 39 | 1880 | 1 | 3646 | 2887.1237 | 31.752 | 207,156 | 2886.115875 | 1006 | 343 | 347 | NATRF | - | 343(NAT) | 1 | | 607.30779 | 3442227.5 | 39.85 | 1 | 5 | 5 | 1 | 0 | 1 | 0 | 1 | 2886.117015 | -0.0011 | -0.39 | -8.10 |
| Chymotrypsin | 21 | 39 | 1880 | 1 | 3646 | 2887.1237 | 31.752 | 207,156 | 2886.115875 | 1006 | 343 | 347 | NATRF | - | 343(NAT) | 1 | | 607.30779 | 3442227.5 | 39.85 | 2 | 6 | 5 | 0 | 1 | 0 | 0 | 1 | 2886.117016 | -0.0011 | -0.39 | -8.10 |
| Chymotrypsin | 21 | 39 | 2032 | 1 | 3576 | 3034.2024 | 28.728 | 191,174 | 3033.194575 | 1006 | 343 | 347 | NATRF | - | 343(NAT) | 1 | | 607.30779 | 3442227.5 | 39.85 | 1 | 6 | 5 | 3 | 0 | 0 | 0 | 1 | 3033.195326 | -0.0008 | -0.24 | -11.12 |
| Chymotrypsin | 21 | 39 | 2585 | 1 | 3578 | 3107.2185 | 28.8163 | 149,758 | 3106.210675 | 1006 | 343 | 347 | NATRF | - | 343(NAT) | 1 | | 607.30779 | 3442227.5 | 39.85 | 1 | 7 | 6 | 1 | 0 | 0 | 0 | 1 | 3106.211705 | -0.0010 | -0.33 | -11.03 |
| Chymotrypsin | 21 | 39 | 4036 | 1 | 3542 | 3748.4718 | 27.2749 | 90,868 | 3747.463975 | 1006 | 343 | 347 | NATRF | - | 343(NAT) | 1 | | 607.30779 | 3442227.5 | 39.85 | 1 | 7 | 7 | 4 | 0 | 0 | 0 | 0 | 3747.464805 | -0.0008 | -0.22 | -12.57 |
| Chymotrypsin | 21 | 39 | 4227 | 1 | 3616 | 3341.2921 | 30.4556 | 85,678 | 3340.284275 | 1006 | 343 | 347 | NATRF | - | 343(NAT) | 1 | | 607.30779 | 3442227.5 | 39.85 | 1 | 6 | 5 | 3 | 0 | 1 | 0 | 1 | 3340.285657 | -0.0014 | -0.41 | -9.39 |
| Chymotrypsin | 21 | 39 | 4227 | 1 | 3616 | 3341.2921 | 30.4556 | 85,678 | 3340.284275 | 1006 | 343 | 347 | NATRF | - | 343(NAT) | 1 | | 607.30779 | 3442227.5 | 39.85 | 2 | 7 | 5 | 2 | 1 | 0 | 0 | 0 | 3340.285658 | -0.0014 | -0.41 | -9.39 |
| Chymotrypsin | 21 | 39 | 4260 | 1 | 3661 | 3049.1773 | 32.3932 | 84,958 | 3048.169475 | 1006 | 343 | 347 | NATRF | - | 343(NAT) | 1 | | 607.30779 | 3442227.5 | 39.85 | 1 | 6 | 5 | 1 | 0 | 1 | 0 | 1 | 3048.169839 | -0.0004 | -0.11 | -7.45 |
| Chymotrypsin | 21 | 39 | 4260 | 1 | 3661 | 3049.1773 | 32.3932 | 84,958 | 3048.169475 | 1006 | 343 | 347 | NATRF | - | 343(NAT) | 1 | | 607.30779 | 3442227.5 | 39.85 | 2 | 7 | 5 | 0 | 1 | 0 | 0 | 0 | 3048.16984 | -0.0004 | -0.11 | -7.45 |
| Chymotrypsin | 21 | 39 | 4447 | 1 | 3570 | 2360.9615 | 28.4748 | 80,342 | 2359.953675 | 1006 | 343 | 347 | NATRF | - | 343(NAT) | 1 | | 607.30779 | 3442227.5 | 39.85 | 1 | 4 | 4 | 2 | 0 | 0 | 0 | 1 | 2359.952396 | 0.0013 | 0.54 | -11.37 |
| Chymotrypsin | 21 | 39 | 4548 | 1 | 3567 | 3253.2744 | 28.3466 | 77,799 | 3252.266575 | 1006 | 343 | 347 | NATRF | - | 343(NAT) | 1 | | 607.30779 | 3442227.5 | 39.85 | 1 | 7 | 6 | 2 | 0 | 0 | 0 | 0 | 3252.269614 | -0.0030 | -0.93 | -11.50 |
| Chymotrypsin | 21 | 39 | 5097 | 1 | 3647 | 2668.052 | 31.7937 | 66,738 | 2667.044175 | 1006 | 343 | 347 | NATRF | - | 343(NAT) | 1 | | 607.30779 | 3442227.5 | 39.85 | 1 | 5 | 4 | 1 | 1 | 0 | 0 | 1 | 2667.042728 | 0.0014 | 0.54 | -8.05 |
| Chymotrypsin | 21 | 39 | 5097 | 1 | 3647 | 2668.052 | 31.7937 | 66,738 | 2667.044175 | 1006 | 343 | 347 | NATRF | - | 343(NAT) | 1 | | 607.30779 | 3442227.5 | 39.85 | 2 | 4 | 4 | 2 | 0 | 1 | 0 | 1 | 2667.042727 | 0.0014 | 0.54 | -8.05 |
| Chymotrypsin | 21 | 39 | 5500 | 1 | 3583 | 2945.1689 | 29.0292 | 60,144 | 2944.161075 | 1006 | 343 | 347 | NATRF | - | 343(NAT) | 1 | | 607.30779 | 3442227.5 | 39.85 | 1 | 6 | 6 | 1 | 0 | 0 | 0 | 1 | 2944.158881 | 0.0022 | 0.74 | -10.82 |
| Chymotrypsin | 21 | 39 | 5903 | 1 | 3605 | 3341.2877 | 29.9774 | 53,412 | 3340.279875 | 1006 | 343 | 347 | NATRF | - | 343(NAT) | 1 | | 607.30779 | 3442227.5 | 39.85 | 1 | 6 | 5 | 3 | 0 | 1 | 0 | 1 | 3340.285657 | -0.0058 | -1.73 | -9.87 |
| Chymotrypsin | 21 | 39 | 5903 | 1 | 3605 | 3341.2877 | 29.9774 | 53,412 | 3340.279875 | 1006 | 343 | 347 | NATRF | - | 343(NAT) | 1 | | 607.30779 | 3442227.5 | 39.85 | 2 | 7 | 5 | 2 | 1 | 0 | 0 | 0 | 3340.285658 | -0.0058 | -1.73 | -9.87 |
| Chymotrypsin | 21 | 39 | 5909 | 1 | 3541 | 3545.3902 | 27.2322 | 53,357 | 3544.382375 | 1006 | 343 | 347 | NATRF | - | 343(NAT) | 1 | | 607.30779 | 3442227.5 | 39.85 | 1 | 7 | 6 | 4 | 0 | 0 | 0 | 1 | 3544.385432 | -0.0031 | -0.86 | -12.61 |
| Chymotrypsin | 21 | 39 | 5909 | 1 | 3541 | 3545.3902 | 27.2322 | 53,357 | 3544.382375 | 1006 | 343 | 347 | NATRF | - | 343(NAT) | 1 | | 607.30779 | 3442227.5 | 39.85 | 2 | 2 | 10 | 0 | 2 | 0 | 1 | -1 | 3544.398002 | -0.0156 | -4.4 | -12.61 |
| Chymotrypsin | 21 | 39 | 6353 | 1 | 3621 | 3195.2351 | 30.6732 | 47,161 | 3194.227275 | 1006 | 343 | 347 | NATRF | - | 343(NAT) | 1 | | 607.30779 | 3442227.5 | 39.85 | 1 | 6 | 5 | 2 | 0 | 1 | 0 | 1 | 3194.227748 | -0.0005 | -0.14 | -9.17 |
| Chymotrypsin | 21 | 39 | 6353 | 1 | 3621 | 3195.2351 | 30.6732 | 47,161 | 3194.227275 | 1006 | 343 | 347 | NATRF | - | 343(NAT) | 1 | | 607.30779 | 3442227.5 | 39.85 | 2 | 7 | 5 | 1 | 1 | 0 | 0 | 1 | 3194.227749 | -0.0005 | -0.14 | -9.17 |
| Chymotrypsin | 21 | 39 | 6928 | 1 | 3645 | 3049.1739 | 31.7098 | 39,173 | 3048.166075 | 1006 | 343 | 347 | NATRF | - | 343(NAT) | 1 | | 607.30779 | 3442227.5 | 39.85 | 1 | 6 | 5 | 1 | 0 | 1 | 0 | 1 | 3048.169839 | -0.0038 | -1.23 | -8.14 |
| Chymotrypsin | 21 | 39 | 6928 | 1 | 3645 | 3049.1739 | 31.7098 | 39,173 | 3048.166075 | 1006 | 343 | 347 | NATRF | - | 343(NAT) | 1 | | 607.30779 | 3442227.5 | 39.85 | 2 | 7 | 5 | 0 | 1 | 0 | 0 | 0 | 3048.16984 | -0.0038 | -1.23 | -8.14 |
| Chymotrypsin | 21 | 39 | 7042 | 1 | 3693 | 2887.1252 | 33.766 | 37,361 | 2886.117375 | 1006 | 343 | 347 | NATRF | - | 343(NAT) | 1 | | 607.30779 | 3442227.5 | 39.85 | 1 | 6 | 5 | 0 | 1 | 0 | 0 | 1 | 2886.117016 | 0.0004 | 0.12 | -6.08 |
| Chymotrypsin | 21 | 39 | 7042 | 1 | 3693 | 2887.1252 | 33.766 | 37,361 | 2886.117375 | 1006 | 343 | 347 | NATRF | - | 343(NAT) | 1 | | 607.30779 | 3442227.5 | 39.85 | 2 | 5 | 5 | 1 | 0 | 1 | 0 | 1 | 2886.117015 | 0.0004 | 0.12 | -6.08 |
| Chymotrypsin | 21 | 39 | 7257 | 1 | 3633 | 3414.3116 | 31.1882 | 34,602 | 3413.303775 | 1006 | 343 | 347 | NATRF | - | 343(NAT) | 1 | | 607.30779 | 3442227.5 | 39.85 | 1 | 8 | 6 | 0 | 1 | 0 | 0 | 1 | 3413.302037 | 0.0017 | 0.5 | -8.66 |
| Chymotrypsin | 21 | 39 | 7257 | 1 | 3633 | 3414.3116 | 31.1882 | 34,602 | 3413.303775 | 1006 | 343 | 347 | NATRF | - | 343(NAT) | 1 | | 607.30779 | 3442227.5 | 39.85 | 2 | 7 | 6 | 1 | 0 | 1 | 0 | 1 | 3413.302036 | 0.0017 | 0.5 | -8.66 |
| Chymotrypsin | 21 | 39 | 7454 | 1 | 3549 | 3399.3347 | 27.5744 | 32,402 | 3398.326875 | 1006 | 343 | 347 | NATRF | - | 343(NAT) | 1 | | 607.30779 | 3442227.5 | 39.85 | 1 | 7 | 6 | 3 | 0 | 0 | 0 | 0 | 3398.327523 | -0.0006 | -0.19 | -12.27 |
| Chymotrypsin | 21 | 39 | 7606 | 1 | 3642 | 3195.2245 | 31.579 | 30,236 | 3194.216675 | 1006 | 343 | 347 | NATRF | - | 343(NAT) | 1 | | 607.30779 | 3442227.5 | 39.85 | 1 | 6 | 5 | 2 | 0 | 1 | 0 | 1 | 3194.227748 | -0.0111 | -3.46 | -8.27 |
| Chymotrypsin | 21 | 39 | 7606 | 1 | 3642 | 3195.2245 | 31.579 | 30,236 | 3194.216675 | 1006 | 343 | 347 | NATRF | - | 343(NAT) | 1 | | 607.30779 | 3442227.5 | 39.85 | 2 | 7 | 5 | 1 | 1 | 0 | 0 | 1 | 3194.227749 | -0.0111 | -3.46 | -8.27 |
| Chymotrypsin | 21 | 39 | 8014 | 1 | 3634 | 3049.1784 | 31.2312 | 24,952 | 3048.170575 | 1006 | 343 | 347 | NATRF | - | 343(NAT) | 1 | | 607.30779 | 3442227.5 | 39.85 | 1 | 6 | 5 | 1 | 0 | 1 | 0 | 1 | 3048.169839 | 0.0007 | 0.24 | -8.62 |
| Chymotrypsin | 21 | 39 | 8014 | 1 | 3634 | 3049.1784 | 31.2312 | 24,952 | 3048.170575 | 1006 | 343 | 347 | NATRF | - | 343(NAT) | 1 | | 607.30779 | 3442227.5 | 39.85 | 2 | 7 | 5 | 0 | 1 | 0 | 0 | 0 | 3048.16984 | 0.0007 | 0.24 | -8.62 |
| Chymotrypsin | 21 | 39 | 8062 | 1 | 3569 | 3018.2069 | 28.4311 | 24,447 | 3017.199075 | 1006 | 343 | 347 | NATRF | - | 343(NAT) | 1 | | 607.30779 | 3442227.5 | 39.85 | 1 | 5 | 5 | 4 | 0 | 0 | 0 | 0 | 3017.200411 | -0.0013 | -0.44 | -11.42 |
| Chymotrypsin | 21 | 39 | 8382 | 1 | 3563 | 3399.3373 | 28.1732 | 20,503 | 3398.329475 | 1006 | 343 | 347 | NATRF | - | 343(NAT) | 1 | | 607.30779 | 3442227.5 | 39.85 | 1 | 7 | 6 | 3 | 0 | 0 | 0 | 0 | 3398.327523 | 0.0020 | 0.57 | -11.67 |
| Chymotrypsin | 21 | 39 | 8514 | 1 | 3597 | 3852.479 | 29.6297 | 18,533 | 3851.471175 | 1006 | 343 | 347 | NATRF | - | 343(NAT) | 1 | | 607.30779 | 3442227.5 | 39.85 | 1 | 7 | 6 | 4 | 0 | 1 | 0 | 1 | 3851.475763 | -0.0046 | -1.19 | -10.22 |
| Chymotrypsin | 21 | 39 | 8514 | 1 | 3597 | 3852.479 | 29.6297 | 18,533 | 3851.471175 | 1006 | 343 | 347 | NATRF | - | 343(NAT) | 1 | | 607.30779 | 3442227.5 | 39.85 | 2 | 8 | 6 | 3 | 1 | 0 | 0 | 0 | 3851.475764 | -0.0046 | -1.19 | -10.22 |
| Chymotrypsin | 21 | 39 | 8514 | 1 | 3597 | 3852.479 | 29.6297 | 18,533 | 3851.471175 | 1006 | 343 | 347 | NATRF | - | 343(NAT) | 1 | | 607.30779 | 3442227.5 | 39.85 | 3 | 2 | 10 | 0 | 2 | 1 | 1 | -1 | 3851.488333 | -0.0172 | -4.45 | -10.22 |
| Chymotrypsin | 21 | 39 | 8587 | 1 | 3638 | 3017.1769 | 31.4036 | 17,618 | 3016.169075 | 1006 | 343 | 347 | NATRF | - | 343(NAT) | 1 | | 607.30779 | 3442227.5 | 39.85 | 1 | 4 | 5 | 3 | 0 | 1 | 0 | 0 | 3016.180009 | -0.0109 | -3.62 | -8.44 |
| Chymotrypsin | 21 | 39 | 8587 | 1 | 3638 | 3017.1769 | 31.4036 | 17,618 | 3016.169075 | 1006 | 343 | 347 | NATRF | - | 343(NAT) | 1 | | 607.30779 | 3442227.5 | 39.85 | 2 | 5 | 5 | 2 | 1 | 0 | 0 | 0 | 3016.18001 | -0.0109 | -3.62 | -8.44 |
| Chymotrypsin | 121 | 7 | 532 | 1 | 3638 | 3091.2245 | 31.4036 | 666,958 | 3090.216675 | 1006 | 343 | 347 | NATRF | - | 343(NAT) | 1 | | 607.30779 | 3442227.5 | 39.85 | 1 | 6 | 6 | 2 | 0 | 0 | 0 | 1 | 3090.21679 | -0.0001 | -0.03 | -8.44 |
| Chymotrypsin | 121 | 7 | 678 | 1 | 3644 | 2945.1655 | 31.6673 | 531,999 | 2944.157675 | 1006 | 343 | 347 | NATRF | - | 343(NAT) | 1 | | 607.30779 | 3442227.5 | 39.85 | 1 | 6 | 6 | 1 | 0 | 0 | 0 | 1 | 2944.158881 | -0.0012 | -0.4 | -8.18 |
| Chymotrypsin | 121 | 7 | 2803 | 1 | 3698 | 3252.2556 | 33.9771 | 137,014 | 3251.247775 | 1006 | 343 | 347 | NATRF | - | 343(NAT) | 1 | | 607.30779 | 3442227.5 | 39.85 | 1 | 7 | 6 | 0 | 1 | 0 | 0 | 1 | 3251.249213 | -0.0014 | -0.44 | -5.87 |
| Chymotrypsin | 121 | 7 | 2803 | 1 | 3698 | 3252.2556 | 33.9771 | 137,014 | 3251.247775 | 1006 | 343 | 347 | NATRF | - | 343(NAT) | 1 | | 607.30779 | 3442227.5 | 39.85 | 2 | 6 | 6 | 1 | 0 | 1 | 0 | 0 | 3251.249212 | -0.0014 | -0.44 | -5.87 |
| Chymotrypsin | 121 | 7 | 4414 | 1 | 3645 | 2783.1152 | 31.7098 | 81,054 | 2782.107375 | 1006 | 343 | 347 | NATRF | - | 343(NAT) | 1 | | 607.30779 | 3442227.5 | 39.85 | 1 | 5 | 6 | 1 | 0 | 0 | 0 | 1 | 2782.106057 | 0.0013 | 0.47 | -8.14 |
| Chymotrypsin | 121 | 7 | 5827 | 1 | 3638 | 2929.1749 | 31.4036 | 54,629 | 2928.167075 | 1006 | 343 | 347 | NATRF | - | 343(NAT) | 1 | | 607.30779 | 3442227.5 | 39.85 | 1 | 5 | 6 | 2 | 0 | 0 | 0 | 1 | 2928.163966 | 0.0031 | 1.06 | -8.44 |
| Chymotrypsin | 121 | 7 | 7128 | 1 | 3702 | 3236.2599 | 34.1484 | 36,246 | 3235.252075 | 1006 | 343 | 347 | NATRF | - | 343(NAT) | 1 | | 607.30779 | 3442227.5 | 39.85 | 1 | 6 | 6 | 1 | 1 | 0 | 0 | 1 | 3235.254298 | -0.0022 | -0.68 | -5.70 |
| Chymotrypsin | 121 | 7 | 7128 | 1 | 3702 | 3236.2599 | 34.1484 | 36,246 | 3235.252075 | 1006 | 343 | 347 | NATRF | - | 343(NAT) | 1 | | 607.30779 | 3442227.5 | 39.85 | 2 | 5 | 6 | 2 | 0 | 1 | 0 | 0 | 3235.254297 | -0.0022 | -0.68 | -5.70 |
| Chymotrypsin | 121 | 7 | 8305 | 1 | 3636 | 3237.2801 | 31.3175 | 21,610 | 3236.272275 | 1006 | 343 | 347 | NATRF | - | 343(NAT) | 1 | | 607.30779 | 3442227.5 | 39.85 | 1 | 6 | 6 | 3 | 0 | 0 | 0 | 1 | 3236.274699 | -0.0024 | -0.74 | -8.53 |
| Chymotrypsin | 206 | 8 | 1322 | 1 | 3557 | 4202.613 | 27.9172 | 288,269 | 4201.605175 | 1006 | 343 | 347 | NATRF | - | 343(NAT) | 1 | | 607.30779 | 3442227.5 | 39.85 | 1 | 10 | 9 | 1 | 0 | 0 | 0 | 1 | 4201.608296 | -0.0031 | -0.74 | -11.93 |
| Chymotrypsin | 206 | 8 | 2965 | 1 | 3556 | 4405.691 | 27.8743 | 129,033 | 4404.683175 | 1006 | 343 | 347 | NATRF | - | 343(NAT) | 1 | | 607.30779 | 3442227.5 | 39.85 | 1 | 10 | 10 | 1 | 0 | 0 | 0 | 1 | 4404.687669 | -0.0045 | -1.02 | -11.97 |
| Chymotrypsin | 206 | 8 | 3479 | 1 | 3580 | 3894.495 | 28.9017 | 107,673 | 3893.487175 | 1006 | 343 | 347 | NATRF | - | 343(NAT) | 1 | | 607.30779 | 3442227.5 | 39.85 | 1 | 9 | 9 | 0 | 0 | 0 | 0 | 0 | 3893.497563 | -0.0104 | -2.66 | -10.95 |
| Chymotrypsin | 206 | 8 | 5256 | 1 | 3560 | 4243.6361 | 28.0457 | 64,303 | 4242.628275 | 1006 | 343 | 347 | NATRF | - | 343(NAT) | 1 | | 607.30779 | 3442227.5 | 39.85 | 1 | 9 | 10 | 1 | 0 | 0 | 0 | 0 | 4242.634845 | -0.0066 | -1.54 | -11.80 |
| Chymotrypsin | 206 | 8 | 6901 | 1 | 3561 | 4056.555 | 28.0893 | 39,678 | 4055.547175 | 1006 | 343 | 347 | NATRF | - | 343(NAT) | 1 | | 607.30779 | 3442227.5 | 39.85 | 1 | 10 | 9 | 0 | 0 | 0 | 0 | 1 | 4055.550387 | -0.0032 | -0.79 | -11.76 |
| Chymotrypsin | 206 | 8 | 7149 | 1 | 3571 | 3732.4486 | 28.5174 | 35,929 | 3731.440775 | 1006 | 343 | 347 | NATRF | - | 343(NAT) | 1 | | 607.30779 | 3442227.5 | 39.85 | 1 | 8 | 9 | 0 | 0 | 0 | 0 | 0 | 3731.444739 | -0.0040 | -1.06 | -11.33 |
| Chymotrypsin | 206 | 8 | 7688 | 1 | 3566 | 4097.5787 | 28.3031 | 29,384 | 4096.570875 | 1006 | 343 | 347 | NATRF | - | 343(NAT) | 1 | | 607.30779 | 3442227.5 | 39.85 | 1 | 9 | 10 | 0 | 0 | 0 | 0 | 0 | 4096.576936 | -0.0061 | -1.47 | -11.54 |
| Chymotrypsin | 206 | 8 | 9236 | 1 | 3561 | 4259.63 | 28.0893 | 4,551 | 4258.622175 | 1006 | 343 | 347 | NATRF | - | 343(NAT) | 1 | | 607.30779 | 3442227.5 | 39.85 | 1 | 10 | 10 | 0 | 0 | 0 | 0 | 0 | 4258.62976 | -0.0076 | -1.78 | -11.76 |
| alpha-Lytic | 26 | 19 | 116 | 1 | 4876 | 3429.3818 | 79.3352 | 1,193,560 | 3428.373975 | 5 | 334 | 345 | NLCPFGEVFNAT | 0.000000000200.0 | 343(NAT) | 1 | 1370.6063 | 1367.618039 | 29579572 | 87.51 | 1 | 5 | 4 | 3 | 0 | 0 | 0 | 1 | 3428.373378 | 0.0006 | 0.17 | -8.17 |
| alpha-Lytic | 26 | 19 | 126 | 1 | 4874 | 3632.4568 | 79.2391 | 1,083,140 | 3631.448975 | 5 | 334 | 345 | NLCPFGEVFNAT | 0.000000000200.0 | 343(NAT) | 1 | 1370.6063 | 1367.618039 | 29579572 | 87.51 | 1 | 5 | 5 | 3 | 0 | 0 | 0 | 1 | 3631.452751 | -0.0038 | -1.03 | -8.27 |
| alpha-Lytic | 26 | 19 | 371 | 1 | 4887 | 3137.2636 | 79.8822 | 353,020 | 3136.255775 | 5 | 334 | 345 | NLCPFGEVFNAT | 0.000000000200.0 | 343(NAT) | 1 | 1370.6063 | 1367.618039 | 29579572 | 87.51 | 1 | 5 | 4 | 1 | 0 | 0 | 0 | 1 | 3136.25756 | -0.0018 | -0.56 | -7.63 |
| alpha-Lytic | 26 | 19 | 419 | 1 | 4887 | 3340.3436 | 79.8822 | 311,021 | 3339.335775 | 5 | 334 | 345 | NLCPFGEVFNAT | 0.000000000200.0 | 343(NAT) | 1 | 1370.6063 | 1367.618039 | 29579572 | 87.51 | 1 | 5 | 5 | 1 | 0 | 0 | 0 | 1 | 3339.336933 | -0.0012 | -0.34 | -7.63 |
| alpha-Lytic | 26 | 19 | 452 | 1 | 4881 | 3283.3223 | 79.5869 | 279,182 | 3282.314475 | 5 | 334 | 345 | NLCPFGEVFNAT | 0.000000000200.0 | 343(NAT) | 1 | 1370.6063 | 1367.618039 | 29579572 | 87.51 | 1 | 5 | 4 | 2 | 0 | 0 | 0 | 1 | 3282.315469 | -0.0010 | -0.3 | -7.92 |
| alpha-Lytic | 26 | 19 | 569 | 1 | 4882 | 3486.4046 | 79.638 | 233,612 | 3485.396775 | 5 | 334 | 345 | NLCPFGEVFNAT | 0.000000000200.0 | 343(NAT) | 1 | 1370.6063 | 1367.618039 | 29579572 | 87.51 | 1 | 5 | 5 | 2 | 0 | 0 | 0 | 1 | 3485.394842 | 0.0019 | 0.55 | -7.87 |
| alpha-Lytic | 26 | 19 | 826 | 1 | 4893 | 3080.2407 | 80.161 | 160,029 | 3079.232875 | 5 | 334 | 345 | NLCPFGEVFNAT | 0.000000000200.0 | 343(NAT) | 1 | 1370.6063 | 1367.618039 | 29579572 | 87.51 | 1 | 5 | 3 | 2 | 0 | 0 | 0 | 0 | 3079.236096 | -0.0032 | -1.04 | -7.35 |
| alpha-Lytic | 26 | 19 | 831 | 1 | 4859 | 3940.5692 | 78.4873 | 158,871 | 3939.561375 | 5 | 334 | 345 | NLCPFGEVFNAT | 0.000000000200.0 | 343(NAT) | 1 | 1370.6063 | 1367.618039 | 29579572 | 87.51 | 1 | 6 | 5 | 4 | 0 | 0 | 0 | 0 | 3939.563484 | -0.0021 | -0.53 | -9.02 |
| alpha-Lytic | 26 | 19 | 833 | 1 | 4956 | 3590.4093 | 83.156 | 158,239 | 3589.401475 | 5 | 334 | 345 | NLCPFGEVFNAT | 0.000000000200.0 | 343(NAT) | 1 | 1370.6063 | 1367.618039 | 29579572 | 87.51 | 1 | 5 | 4 | 2 | 0 | 1 | 0 | 1 | 3589.4058 | -0.0043 | -1.2 | -4.35 |
| alpha-Lytic | 26 | 19 | 833 | 1 | 4956 | 3590.4093 | 83.156 | 158,239 | 3589.401475 | 5 | 334 | 345 | NLCPFGEVFNAT | 0.000000000200.0 | 343(NAT) | 1 | 1370.6063 | 1367.618039 | 29579572 | 87.51 | 2 | 6 | 4 | 1 | 1 | 0 | 0 | 1 | 3589.405801 | -0.0043 | -1.2 | -4.35 |
| alpha-Lytic | 26 | 19 | 898 | 1 | 4859 | 4143.6482 | 78.4873 | 147,371 | 4142.640375 | 5 | 334 | 345 | NLCPFGEVFNAT | 0.000000000200.0 | 343(NAT) | 1 | 1370.6063 | 1367.618039 | 29579572 | 87.51 | 1 | 6 | 6 | 4 | 0 | 0 | 0 | 0 | 4142.642857 | -0.0025 | -0.59 | -9.02 |
| alpha-Lytic | 26 | 19 | 1722 | 1 | 4873 | 3794.51 | 79.1924 | 71,956 | 3793.502175 | 5 | 334 | 345 | NLCPFGEVFNAT | 0.000000000200.0 | 343(NAT) | 1 | 1370.6063 | 1367.618039 | 29579572 | 87.51 | 1 | 6 | 5 | 3 | 0 | 0 | 0 | 1 | 3793.505575 | -0.0034 | -0.89 | -8.32 |
| alpha-Lytic | 26 | 19 | 1796 | 1 | 4963 | 3444.3548 | 83.5078 | 68,008 | 3443.346975 | 5 | 334 | 345 | NLCPFGEVFNAT | 0.000000000200.0 | 343(NAT) | 1 | 1370.6063 | 1367.618039 | 29579572 | 87.51 | 1 | 5 | 4 | 1 | 0 | 1 | 0 | 1 | 3443.347891 | -0.0009 | -0.26 | -4.00 |
| alpha-Lytic | 26 | 19 | 1796 | 1 | 4963 | 3444.3548 | 83.5078 | 68,008 | 3443.346975 | 5 | 334 | 345 | NLCPFGEVFNAT | 0.000000000200.0 | 343(NAT) | 1 | 1370.6063 | 1367.618039 | 29579572 | 87.51 | 2 | 6 | 4 | 0 | 1 | 0 | 0 | 1 | 3443.347892 | -0.0009 | -0.26 | -4.00 |
| alpha-Lytic | 26 | 19 | 1893 | 1 | 4851 | 4451.7674 | 78.0675 | 63,545 | 4450.759575 | 5 | 334 | 345 | NLCPFGEVFNAT | 0.000000000200.0 | 343(NAT) | 1 | 1370.6063 | 1367.618039 | 29579572 | 87.51 | 1 | 2 | 10 | 1 | 2 | 0 | 1 | -1 | 4450.76616 | -0.0066 | -1.47 | -9.44 |
| alpha-Lytic | 26 | 19 | 2349 | 1 | 4885 | 3324.3453 | 79.7918 | 47,339 | 3323.337475 | 5 | 334 | 345 | NLCPFGEVFNAT | 0.000000000200.0 | 343(NAT) | 1 | 1370.6063 | 1367.618039 | 29579572 | 87.51 | 1 | 4 | 5 | 2 | 0 | 0 | 0 | 1 | 3323.342018 | -0.0045 | -1.36 | -7.72 |
| alpha-Lytic | 26 | 19 | 2433 | 1 | 4894 | 3178.2883 | 80.202 | 44,711 | 3177.280475 | 5 | 334 | 345 | NLCPFGEVFNAT | 0.000000000200.0 | 343(NAT) | 1 | 1370.6063 | 1367.618039 | 29579572 | 87.51 | 1 | 4 | 5 | 1 | 0 | 0 | 0 | 1 | 3177.284109 | -0.0036 | -1.14 | -7.31 |
| alpha-Lytic | 26 | 19 | 2513 | 1 | 4870 | 3851.5342 | 79.0403 | 42,299 | 3850.526375 | 5 | 334 | 345 | NLCPFGEVFNAT | 0.000000000200.0 | 343(NAT) | 1 | 1370.6063 | 1367.618039 | 29579572 | 87.51 | 1 | 6 | 6 | 2 | 0 | 0 | 0 | 1 | 3850.527039 | -0.0007 | -0.17 | -8.47 |
| alpha-Lytic | 26 | 19 | 2627 | 1 | 4853 | 4305.7046 | 78.1691 | 38,981 | 4304.696775 | 5 | 334 | 345 | NLCPFGEVFNAT | 0.000000000200.0 | 343(NAT) | 1 | 1370.6063 | 1367.618039 | 29579572 | 87.51 | 1 | 7 | 6 | 4 | 0 | 0 | 0 | 1 | 4304.695681 | 0.0011 | 0.25 | -9.34 |
| alpha-Lytic | 26 | 19 | 2627 | 1 | 4853 | 4305.7046 | 78.1691 | 38,981 | 4304.696775 | 5 | 334 | 345 | NLCPFGEVFNAT | 0.000000000200.0 | 343(NAT) | 1 | 1370.6063 | 1367.618039 | 29579572 | 87.51 | 2 | 2 | 10 | 0 | 2 | 0 | 1 | -1 | 4304.708251 | -0.0115 | -2.66 | -9.34 |
| alpha-Lytic | 26 | 19 | 2824 | 1 | 4786 | 4144.6736 | 74.957 | 33,979 | 4143.665775 | 5 | 334 | 345 | NLCPFGEVFNAT | 0.000000000200.0 | 343(NAT) | 1 | 1370.6063 | 1367.618039 | 29579572 | 87.51 | 1 | 0 | 10 | 3 | 0 | 1 | 1 | -1 | 4143.675827 | -0.0101 | -2.42 | -12.55 |
| alpha-Lytic | 26 | 19 | 3684 | 1 | 5032 | 3751.452 | 86.6885 | 14,695 | 3750.444175 | 5 | 334 | 345 | NLCPFGEVFNAT | 0.000000000200.0 | 343(NAT) | 1 | 1370.6063 | 1367.618039 | 29579572 | 87.51 | 1 | 5 | 4 | 1 | 0 | 2 | 0 | 1 | 3750.438222 | 0.0060 | 1.58 | -0.82 |
| alpha-Lytic | 26 | 19 | 3684 | 1 | 5032 | 3751.452 | 86.6885 | 14,695 | 3750.444175 | 5 | 334 | 345 | NLCPFGEVFNAT | 0.000000000200.0 | 343(NAT) | 1 | 1370.6063 | 1367.618039 | 29579572 | 87.51 | 2 | 6 | 4 | 0 | 1 | 1 | 0 | 0 | 3750.438223 | 0.0060 | 1.58 | -0.82 |
| Chymotrypsin | 112 | 26 | 482 | 1 | 4580 | 3355.5075 | 71.1781 | 742,156 | 3354.499675 | 1004 | 593 | 612 | GGVSVITPGTNTSNQVAVLY | 0.0000000000000002000000.0 | 603(NTS) | 1 | | 1976.02143 | 6159764 | 76.04 | 1 | 6 | 2 | 0 | 0 | 0 | 0 | 1 | 3354.49712 | 0.0026 | 0.76 | -4.87 |
| Chymotrypsin | 112 | 26 | 741 | 1 | 4586 | 3193.4504 | 71.4403 | 498,700 | 3192.442575 | 1004 | 593 | 612 | GGVSVITPGTNTSNQVAVLY | 0.0000000000000002000000.0 | 603(NTS) | 1 | | 1976.02143 | 6159764 | 76.04 | 1 | 5 | 2 | 0 | 0 | 0 | 0 | 1 | 3192.444296 | -0.0017 | -0.53 | -4.60 |
| Chymotrypsin | 112 | 26 | 768 | 1 | 4571 | 3745.6662 | 70.8087 | 481,516 | 3744.658375 | 1004 | 593 | 612 | GGVSVITPGTNTSNQVAVLY | 0.0000000000000002000000.0 | 603(NTS) | 1 | | 1976.02143 | 6159764 | 76.04 | 1 | 5 | 4 | 1 | 0 | 0 | 0 | 1 | 3744.660951 | -0.0026 | -0.68 | -5.24 |
| Chymotrypsin | 112 | 26 | 975 | 1 | 4574 | 3517.5557 | 70.9407 | 392,133 | 3516.547875 | 1004 | 593 | 612 | GGVSVITPGTNTSNQVAVLY | 0.0000000000000002000000.0 | 603(NTS) | 1 | | 1976.02143 | 6159764 | 76.04 | 1 | 7 | 2 | 0 | 0 | 0 | 0 | 1 | 3516.549944 | -0.0021 | -0.58 | -5.10 |
| Chymotrypsin | 112 | 26 | 1311 | 1 | 4550 | 4475.9231 | 69.919 | 291,370 | 4474.915275 | 1004 | 593 | 612 | GGVSVITPGTNTSNQVAVLY | 0.0000000000000002000000.0 | 603(NTS) | 1 | | 1976.02143 | 6159764 | 76.04 | 1 | 7 | 6 | 1 | 0 | 0 | 0 | 1 | 4474.925345 | -0.0101 | -2.25 | -6.13 |
| Chymotrypsin | 112 | 26 | 1352 | 1 | 4578 | 3542.5865 | 71.0927 | 283,023 | 3541.578675 | 1004 | 593 | 612 | GGVSVITPGTNTSNQVAVLY | 0.0000000000000002000000.0 | 603(NTS) | 1 | | 1976.02143 | 6159764 | 76.04 | 1 | 5 | 3 | 1 | 0 | 0 | 0 | 1 | 3541.581578 | -0.0029 | -0.81 | -4.95 |
| Chymotrypsin | 112 | 26 | 1431 | 1 | 4576 | 3558.5819 | 71.0184 | 267,339 | 3557.574075 | 1004 | 593 | 612 | GGVSVITPGTNTSNQVAVLY | 0.0000000000000002000000.0 | 603(NTS) | 1 | | 1976.02143 | 6159764 | 76.04 | 1 | 6 | 3 | 0 | 0 | 0 | 0 | 1 | 3557.576493 | -0.0024 | -0.67 | -5.03 |
| Chymotrypsin | 112 | 26 | 1505 | 1 | 4564 | 3891.728 | 70.5137 | 256,509 | 3890.720175 | 1004 | 593 | 612 | GGVSVITPGTNTSNQVAVLY | 0.0000000000000002000000.0 | 603(NTS) | 1 | | 1976.02143 | 6159764 | 76.04 | 1 | 5 | 4 | 2 | 0 | 0 | 0 | 1 | 3890.71886 | 0.0013 | 0.33 | -5.53 |
| Chymotrypsin | 112 | 26 | 1670 | 1 | 4561 | 4037.7868 | 70.3832 | 233,296 | 4036.778975 | 1004 | 593 | 612 | GGVSVITPGTNTSNQVAVLY | 0.0000000000000002000000.0 | 603(NTS) | 1 | | 1976.02143 | 6159764 | 76.04 | 1 | 5 | 4 | 3 | 0 | 0 | 0 | 1 | 4036.776769 | 0.0022 | 0.54 | -5.66 |
| Chymotrypsin | 112 | 26 | 1834 | 1 | 4573 | 3704.6491 | 70.8976 | 213,200 | 3703.641275 | 1004 | 593 | 612 | GGVSVITPGTNTSNQVAVLY | 0.0000000000000002000000.0 | 603(NTS) | 1 | | 1976.02143 | 6159764 | 76.04 | 1 | 6 | 3 | 1 | 0 | 0 | 0 | 1 | 3703.634402 | 0.0069 | 1.85 | -5.15 |
| Chymotrypsin | 112 | 26 | 1916 | 1 | 4582 | 3396.5284 | 71.2634 | 202,581 | 3395.520575 | 1004 | 593 | 612 | GGVSVITPGTNTSNQVAVLY | 0.0000000000000002000000.0 | 603(NTS) | 1 | | 1976.02143 | 6159764 | 76.04 | 1 | 5 | 3 | 0 | 0 | 0 | 0 | 1 | 3395.523669 | -0.0031 | -0.91 | -4.78 |
| Chymotrypsin | 112 | 26 | 2040 | 1 | 4556 | 4256.8572 | 70.1696 | 189,979 | 4255.849375 | 1004 | 593 | 612 | GGVSVITPGTNTSNQVAVLY | 0.0000000000000002000000.0 | 603(NTS) | 1 | | 1976.02143 | 6159764 | 76.04 | 1 | 6 | 5 | 2 | 0 | 0 | 0 | 1 | 4255.851057 | -0.0017 | -0.39 | -5.87 |
| Chymotrypsin | 112 | 26 | 2134 | 1 | 4571 | 3948.7467 | 70.8087 | 183,960 | 3947.738875 | 1004 | 593 | 612 | GGVSVITPGTNTSNQVAVLY | 0.0000000000000002000000.0 | 603(NTS) | 1 | | 1976.02143 | 6159764 | 76.04 | 1 | 5 | 5 | 1 | 0 | 0 | 0 | 1 | 3947.740324 | -0.0014 | -0.36 | -5.24 |
| Chymotrypsin | 112 | 26 | 2326 | 1 | 4548 | 4621.9955 | 69.8341 | 166,874 | 4620.987675 | 1004 | 593 | 612 | GGVSVITPGTNTSNQVAVLY | 0.0000000000000002000000.0 | 603(NTS) | 1 | | 1976.02143 | 6159764 | 76.04 | 1 | 7 | 6 | 2 | 0 | 0 | 0 | 0 | 4620.983254 | 0.0044 | 0.95 | -6.21 |
| Chymotrypsin | 112 | 26 | 2451 | 1 | 4498 | 3600.635 | 67.7198 | 158,284 | 3599.627175 | 1004 | 593 | 612 | GGVSVITPGTNTSNQVAVLY | 0.0000000000000002000000.0 | 603(NTS) | 1 | | 1976.02143 | 6159764 | 76.04 | | | | | | | | | | | | |
| Chymotrypsin | 112 | 26 | 2487 | 1 | 4545 | 4768.0484 | 69.7016 | 156,049 | 4767.040575 | 1004 | 593 | 612 | GGVSVITPGTNTSNQVAVLY | 0.0000000000000002000000.0 | 603(NTS) | 1 | | 1976.02143 | 6159764 | 76.04 | 1 | 7 | 6 | 3 | 0 | 0 | 0 | 0 | 4767.041163 | -0.0006 | -0.12 | -6.34 |
| Chymotrypsin | 112 | 26 | 2617 | 1 | 4576 | 3786.6998 | 71.0184 | 147,898 | 3785.691975 | 1004 | 593 | 612 | GGVSVITPGTNTSNQVAVLY | 0.0000000000000002000000.0 | 603(NTS) | 1 | | 1976.02143 | 6159764 | 76.04 | 1 | 4 | 5 | 1 | 0 | 0 | 0 | 1 | 3785.6875 | 0.0045 | 1.18 | -5.03 |
| Chymotrypsin | 112 | 26 | 3211 | 1 | 4570 | 3679.6135 | 70.7656 | 117,266 | 3678.605675 | 1004 | 593 | 612 | GGVSVITPGTNTSNQVAVLY | 0.0000000000000002000000.0 | 603(NTS) | 1 | | 1976.02143 | 6159764 | 76.04 | 1 | 8 | 2 | 0 | 0 | 0 | 0 | 1 | 3678.602768 | 0.0029 | 0.79 | -5.28 |
| Chymotrypsin | 112 | 26 | 3292 | 1 | 4574 | 3688.656 | 70.9407 | 114,168 | 3687.648175 | 1004 | 593 | 612 | GGVSVITPGTNTSNQVAVLY | 0.0000000000000002000000.0 | 603(NTS) | 1 | | 1976.02143 | 6159764 | 76.04 | 1 | 5 | 3 | 2 | 0 | 0 | 0 | 0 | 3687.639487 | 0.0087 | 2.35 | -5.10 |
| Chymotrypsin | 112 | 26 | 3298 | 1 | 4585 | 3421.5644 | 71.3959 | 113,917 | 3420.556575 | 1004 | 593 | 612 | GGVSVITPGTNTSNQVAVLY | 0.0000000000000002000000.0 | 603(NTS) | 1 | | 1976.02143 | 6159764 | 76.04 | 1 | 3 | 4 | 1 | 0 | 0 | 0 | 1 | 3420.555303 | 0.0013 | 0.37 | -4.65 |
| Chymotrypsin | 112 | 26 | 3594 | 1 | 4541 | 4914.1026 | 69.528 | 104,034 | 4913.094775 | 1004 | 593 | 612 | GGVSVITPGTNTSNQVAVLY | 0.0000000000000002000000.0 | 603(NTS) | 1 | | 1976.02143 | 6159764 | 76.04 | 1 | 7 | 6 | 4 | 0 | 0 | 0 | 1 | 4913.099072 | -0.0043 | -0.87 | -6.52 |
| Chymotrypsin | 112 | 26 | 3594 | 1 | 4541 | 4914.1026 | 69.528 | 104,034 | 4913.094775 | 1004 | 593 | 612 | GGVSVITPGTNTSNQVAVLY | 0.0000000000000002000000.0 | 603(NTS) | 1 | | 1976.02143 | 6159764 | 76.04 | 2 | 2 | 10 | 0 | 2 | 0 | 1 | -1 | 4913.111642 | -0.0169 | -3.43 | -6.52 |
| Chymotrypsin | 112 | 26 | 4032 | 1 | 4573 | 3599.6067 | 70.8976 | 90,975 | 3598.598875 | 1004 | 593 | 612 | GGVSVITPGTNTSNQVAVLY | 0.0000000000000002000000.0 | 603(NTS) | 1 | | 1976.02143 | 6159764 | 76.04 | 1 | 5 | 4 | 0 | 0 | 0 | 0 | 1 | 3598.603042 | -0.0042 | -1.15 | -5.15 |
| Chymotrypsin | 112 | 26 | 4812 | 1 | 4548 | 4548.9898 | 69.8341 | 72,540 | 4547.981975 | 1004 | 593 | 612 | GGVSVITPGTNTSNQVAVLY | 0.0000000000000002000000.0 | 603(NTS) | 1 | | 1976.02143 | 6159764 | 76.04 | 1 | 6 | 5 | 4 | 0 | 0 | 0 | 0 | 4547.966875 | 0.0151 | 3.32 | -6.21 |
| Chymotrypsin | 112 | 26 | 5101 | 1 | 4565 | 4313.8945 | 70.5583 | 66,682 | 4312.886675 | 1004 | 593 | 612 | GGVSVITPGTNTSNQVAVLY | 0.0000000000000002000000.0 | 603(NTS) | 1 | | 1976.02143 | 6159764 | 76.04 | 1 | 6 | 6 | 1 | 0 | 0 | 0 | 1 | 4312.872521 | 0.0142 | 3.28 | -5.49 |
| Chymotrypsin | 112 | 26 | 5609 | 1 | 4554 | 4402.9171 | 70.0915 | 58,348 | 4401.909275 | 1004 | 593 | 612 | GGVSVITPGTNTSNQVAVLY | 0.0000000000000002000000.0 | 603(NTS) | 1 | | 1976.02143 | 6159764 | 76.04 | 1 | 6 | 5 | 3 | 0 | 0 | 0 | 1 | 4401.908966 | 0.0003 | 0.07 | -5.95 |
| Chymotrypsin | 112 | 26 | 6454 | 1 | 4577 | 3583.6168 | 71.0629 | 45,749 | 3582.608975 | 1004 | 593 | 612 | GGVSVITPGTNTSNQVAVLY | 0.0000000000000002000000.0 | 603(NTS) | 1 | | 1976.02143 | 6159764 | 76.04 | 1 | 4 | 4 | 1 | 0 | 0 | 0 | 1 | 3582.608127 | 0.0008 | 0.23 | -4.98 |
| Chymotrypsin | 189 | 9 | 1123 | 1 | 4505 | 3192.4373 | 68.002 | 341,850 | 3191.429475 | 1003 | 593 | 611 | GGVSVITPGTNTSNQVAVL | 0.000000000000000200000.0 | 603(NTS) | 1 | | 1812.9581 | 4140285 | 73.30 | 1 | 6 | 2 | 0 | 0 | 0 | 0 | 1 | 3191.43379 | -0.0043 | -1.35 | -5.30 |
| Chymotrypsin | 189 | 9 | 1928 | 1 | 4496 | 3582.6046 | 67.6362 | 201,765 | 3581.596775 | 1003 | 593 | 611 | GGVSVITPGTNTSNQVAVL | 0.000000000000000200000.0 | 603(NTS) | 1 | | 1812.9581 | 4140285 | 73.30 | 1 | 5 | 4 | 1 | 0 | 0 | 0 | 1 | 3581.597621 | -0.0008 | -0.23 | -5.67 |
| Chymotrypsin | 189 | 9 | 2411 | 1 | 4500 | 3354.4933 | 67.8034 | 160,913 | 3353.485475 | 1003 | 593 | 611 | GGVSVITPGTNTSNQVAVL | 0.000000000000000200000.0 | 603(NTS) | 1 | | 1812.9581 | 4140285 | 73.30 | 1 | 7 | 2 | 0 | 0 | 0 | 0 | 1 | 3353.486614 | -0.0011 | -0.33 | -5.50 |
| Chymotrypsin | 189 | 9 | 2998 | 1 | 4507 | 3379.5271 | 68.0881 | 127,144 | 3378.519275 | 1003 | 593 | 611 | GGVSVITPGTNTSNQVAVL | 0.000000000000000200000.0 | 603(NTS) | 1 | | 1812.9581 | 4140285 | 73.30 | 1 | 5 | 3 | 1 | 0 | 0 | 0 | 1 | 3378.518248 | 0.0010 | 0.3 | -5.21 |
| Chymotrypsin | 189 | 9 | 4009 | 1 | 4501 | 3395.5235 | 67.847 | 91,543 | 3394.515675 | 1003 | 593 | 611 | GGVSVITPGTNTSNQVAVL | 0.000000000000000200000.0 | 603(NTS) | 1 | | 1812.9581 | 4140285 | 73.30 | 1 | 6 | 3 | 0 | 0 | 0 | 0 | 1 | 3394.513163 | 0.0025 | 0.74 | -5.46 |
| Chymotrypsin | 189 | 9 | 5313 | 1 | 4509 | 3233.4675 | 68.1743 | 63,359 | 3232.459675 | 1003 | 593 | 611 | GGVSVITPGTNTSNQVAVL | 0.000000000000000200000.0 | 603(NTS) | 1 | | 1812.9581 | 4140285 | 73.30 | 1 | 5 | 3 | 0 | 0 | 0 | 0 | 1 | 3232.460339 | -0.0007 | -0.2 | -5.13 |
| Chymotrypsin | 189 | 9 | 5775 | 1 | 4501 | 3525.5811 | 67.847 | 55,350 | 3524.573275 | 1003 | 593 | 611 | GGVSVITPGTNTSNQVAVL | 0.000000000000000200000.0 | 603(NTS) | 1 | | 1812.9581 | 4140285 | 73.30 | 1 | 5 | 3 | 2 | 0 | 0 | 0 | 0 | 3524.576157 | -0.0029 | -0.81 | -5.46 |
| Chymotrypsin | 189 | 9 | 5883 | 1 | 4513 | 3030.3896 | 68.3449 | 53,712 | 3029.381775 | 1003 | 593 | 611 | GGVSVITPGTNTSNQVAVL | 0.000000000000000200000.0 | 603(NTS) | 1 | | 1812.9581 | 4140285 | 73.30 | 1 | 5 | 2 | 0 | 0 | 0 | 0 | 1 | 3029.380966 | 0.0008 | 0.26 | -4.96 |
| Chymotrypsin | 189 | 9 | 6433 | 1 | 4497 | 3516.5456 | 67.678 | 45,978 | 3515.537775 | 1003 | 593 | 611 | GGVSVITPGTNTSNQVAVL | 0.000000000000000200000.0 | 603(NTS) | 1 | | 1812.9581 | 4140285 | 73.30 | 1 | 8 | 2 | 0 | 0 | 0 | 0 | 1 | 3515.539438 | -0.0017 | -0.47 | -5.62 |
| alpha-Lytic | 73 | 4 | 458 | 1 | 4033 | 2658.1034 | 37.8531 | 277,424 | 2657.095575 | 1005 | 597 | 605 | VITPGTNTS | - | 603(NTS) | 1 | | 888.45524 | - | 42.00 | 1 | 5 | 4 | 1 | 0 | 0 | 0 | 1 | 2657.094761 | 0.0008 | 0.3 | -4.15 |
| alpha-Lytic | 73 | 4 | 1047 | 1 | 4033 | 2455.0249 | 37.8531 | 127,726 | 2454.017075 | 1005 | 597 | 605 | VITPGTNTS | - | 603(NTS) | 1 | | 888.45524 | - | 42.00 | 1 | 5 | 3 | 1 | 0 | 0 | 0 | 1 | 2454.015388 | 0.0017 | 0.68 | -4.15 |
| alpha-Lytic | 73 | 4 | 1278 | 1 | 4020 | 2804.1617 | 37.2267 | 102,253 | 2803.153875 | 1005 | 597 | 605 | VITPGTNTS | - | 603(NTS) | 1 | | 888.45524 | - | 42.00 | 1 | 5 | 4 | 2 | 0 | 0 | 0 | 1 | 2803.15267 | 0.0012 | 0.42 | -4.77 |
| alpha-Lytic | 73 | 4 | 1316 | 1 | 4009 | 2950.2178 | 36.699 | 99,076 | 2949.209975 | 1005 | 597 | 605 | VITPGTNTS | - | 603(NTS) | 1 | | 888.45524 | - | 42.00 | 1 | 5 | 4 | 3 | 0 | 0 | 0 | 1 | 2949.210579 | -0.0006 | -0.2 | -5.30 |
| alpha-Lytic | 112 | 4 | 1157 | 1 | 4011 | 3388.3711 | 36.7956 | 113,602 | 3387.363275 | 1005 | 597 | 605 | VITPGTNTS | - | 603(NTS) | 1 | | 888.45524 | - | 42.00 | 1 | 7 | 6 | 1 | 0 | 0 | 0 | 1 | 3387.359155 | 0.0041 | 1.21 | -5.20 |
| alpha-Lytic | 112 | 4 | 2312 | 1 | 4004 | 3534.4336 | 36.432 | 48,688 | 3533.425775 | 1005 | 597 | 605 | VITPGTNTS | - | 603(NTS) | 1 | | 888.45524 | - | 42.00 | 1 | 7 | 6 | 2 | 0 | 0 | 0 | 0 | 3533.417064 | 0.0087 | 2.46 | -5.57 |
| alpha-Lytic | 112 | 4 | 2377 | 1 | 3999 | 3680.4843 | 36.1872 | 46,141 | 3679.476475 | 1005 | 597 | 605 | VITPGTNTS | - | 603(NTS) | 1 | | 888.45524 | - | 42.00 | 1 | 7 | 6 | 3 | 0 | 0 | 0 | 0 | 3679.474973 | 0.0015 | 0.4 | -5.81 |
| alpha-Lytic | 112 | 4 | 2665 | 1 | 3993 | 3826.5406 | 35.8991 | 38,358 | 3825.532775 | 1005 | 597 | 605 | VITPGTNTS | - | 603(NTS) | 1 | | 888.45524 | - | 42.00 | 1 | 7 | 6 | 4 | 0 | 0 | 0 | 1 | 3825.532882 | -0.0001 | -0.02 | -6.10 |
| alpha-Lytic | 112 | 4 | 2665 | 1 | 3993 | 3826.5406 | 35.8991 | 38,358 | 3825.532775 | 1005 | 597 | 605 | VITPGTNTS | - | 603(NTS) | 1 | | 888.45524 | - | 42.00 | 2 | 2 | 10 | 0 | 2 | 0 | 1 | -1 | 3825.545452 | -0.0127 | -3.31 | -6.10 |
| Chymotrypsin | 110 | 21 | 471 | 1 | 4633 | 4893.039 | 73.1477 | 757,923 | 4892.031175 | 48 | 613 | 633 | QDVNCTEVPVAIHADQLTPTW | 0.0002000000000000000000.0 | 616(NCT) | 1 | 2396.1205 | 2393.132239 | 8493031 | 79.12 | 1 | 7 | 6 | 1 | 0 | 0 | 0 | 1 | 4892.036154 | -0.0050 | -1.01 | -5.97 |
| Chymotrypsin | 110 | 21 | 515 | 1 | 4651 | 4162.7783 | 73.909 | 689,538 | 4161.770475 | 48 | 613 | 633 | QDVNCTEVPVAIHADQLTPTW | 0.0002000000000000000000.0 | 616(NCT) | 1 | 2396.1205 | 2393.132239 | 8493031 | 79.12 | 1 | 5 | 4 | 1 | 0 | 0 | 0 | 1 | 4161.77176 | -0.0013 | -0.3 | -5.21 |
| Chymotrypsin | 110 | 21 | 774 | 1 | 4642 | 4527.9115 | 73.5272 | 478,294 | 4526.903675 | 48 | 613 | 633 | QDVNCTEVPVAIHADQLTPTW | 0.0002000000000000000000.0 | 616(NCT) | 1 | 2396.1205 | 2393.132239 | 8493031 | 79.12 | 1 | 6 | 5 | 1 | 0 | 0 | 0 | 1 | 4526.903957 | -0.0003 | -0.06 | -5.59 |
| Chymotrypsin | 110 | 21 | 1077 | 1 | 4656 | 4016.7249 | 74.1216 | 353,888 | 4015.717075 | 48 | 613 | 633 | QDVNCTEVPVAIHADQLTPTW | 0.0002000000000000000000.0 | 616(NCT) | 1 | 2396.1205 | 2393.132239 | 8493031 | 79.12 | 1 | 5 | 4 | 0 | 0 | 0 | 0 | 1 | 4015.713851 | 0.0032 | 0.8 | -5.00 |
| Chymotrypsin | 110 | 21 | 1115 | 1 | 4648 | 4308.8368 | 73.7831 | 343,989 | 4307.828975 | 48 | 613 | 633 | QDVNCTEVPVAIHADQLTPTW | 0.0002000000000000000000.0 | 616(NCT) | 1 | 2396.1205 | 2393.132239 | 8493031 | 79.12 | 1 | 5 | 4 | 2 | 0 | 0 | 0 | 1 | 4307.829669 | -0.0007 | -0.16 | -5.34 |
| Chymotrypsin | 110 | 21 | 1455 | 1 | 4630 | 5039.0978 | 73.0203 | 264,063 | 5038.089975 | 48 | 613 | 633 | QDVNCTEVPVAIHADQLTPTW | 0.0002000000000000000000.0 | 616(NCT) | 1 | 2396.1205 | 2393.132239 | 8493031 | 79.12 | 1 | 7 | 6 | 2 | 0 | 0 | 0 | 0 | 5038.094063 | -0.0041 | -0.81 | -6.10 |
| Chymotrypsin | 110 | 21 | 1976 | 1 | 4643 | 4454.8974 | 73.5696 | 197,083 | 4453.889575 | 48 | 613 | 633 | QDVNCTEVPVAIHADQLTPTW | 0.0002000000000000000000.0 | 616(NCT) | 1 | 2396.1205 | 2393.132239 | 8493031 | 79.12 | 1 | 5 | 4 | 3 | 0 | 0 | 0 | 1 | 4453.887578 | 0.0020 | 0.44 | -5.55 |
| Chymotrypsin | 110 | 21 | 2183 | 1 | 4647 | 4381.8493 | 73.7412 | 179,193 | 4380.841475 | 48 | 613 | 633 | QDVNCTEVPVAIHADQLTPTW | 0.0002000000000000000000.0 | 616(NCT) | 1 | 2396.1205 | 2393.132239 | 8493031 | 79.12 | 1 | 6 | 5 | 0 | 0 | 0 | 0 | 1 | 4380.846048 | -0.0046 | -1.04 | -5.38 |
| Chymotrypsin | 110 | 21 | 2345 | 1 | 4628 | 5185.1531 | 72.9577 | 165,663 | 5184.145275 | 48 | 613 | 633 | QDVNCTEVPVAIHADQLTPTW | 0.0002000000000000000000.0 | 616(NCT) | 1 | 2396.1205 | 2393.132239 | 8493031 | 79.12 | 1 | 7 | 6 | 3 | 0 | 0 | 0 | 0 | 5184.151972 | -0.0067 | -1.29 | -6.16 |
| Chymotrypsin | 110 | 21 | 2517 | 1 | 4638 | 4673.9732 | 73.3583 | 154,346 | 4672.965375 | 48 | 613 | 633 | QDVNCTEVPVAIHADQLTPTW | 0.0002000000000000000000.0 | 616(NCT) | 1 | 2396.1205 | 2393.132239 | 8493031 | 79.12 | 1 | 6 | 5 | 2 | 0 | 0 | 0 | 1 | 4672.961866 | 0.0035 | 0.75 | -5.76 |
| Chymotrypsin | 110 | 21 | 2943 | 1 | 4664 | 3813.6445 | 74.446 | 130,215 | 3812.636675 | 48 | 613 | 633 | QDVNCTEVPVAIHADQLTPTW | 0.0002000000000000000000.0 | 616(NCT) | 1 | 2396.1205 | 2393.132239 | 8493031 | 79.12 | 1 | 5 | 3 | 0 | 0 | 0 | 0 | 1 | 3812.634478 | 0.0022 | 0.57 | -4.68 |
| Chymotrypsin | 110 | 21 | 3056 | 1 | 4658 | 3959.6872 | 74.1975 | 124,174 | 3958.679375 | 48 | 613 | 633 | QDVNCTEVPVAIHADQLTPTW | 0.0002000000000000000000.0 | 616(NCT) | 1 | 2396.1205 | 2393.132239 | 8493031 | 79.12 | 1 | 5 | 3 | 1 | 0 | 0 | 0 | 1 | 3958.692387 | -0.0130 | -3.28 | -4.92 |
| Chymotrypsin | 110 | 21 | 4044 | 1 | 4726 | 4469.8773 | 77.0005 | 90,618 | 4468.869475 | 48 | 613 | 633 | QDVNCTEVPVAIHADQLTPTW | 0.0002000000000000000000.0 | 616(NCT) | 1 | 2396.1205 | 2393.132239 | 8493031 | 79.12 | 1 | 6 | 4 | 0 | 1 | 0 | 0 | 1 | 4468.862092 | 0.0074 | 1.65 | -2.12 |
| Chymotrypsin | 110 | 21 | 4044 | 1 | 4726 | 4469.8773 | 77.0005 | 90,618 | 4468.869475 | 48 | 613 | 633 | QDVNCTEVPVAIHADQLTPTW | 0.0002000000000000000000.0 | 616(NCT) | 1 | 2396.1205 | 2393.132239 | 8493031 | 79.12 | 2 | 5 | 4 | 1 | 0 | 1 | 0 | 1 | 4468.862091 | 0.0074 | 1.65 | -2.12 |
| Chymotrypsin | 110 | 21 | 4545 | 1 | 4710 | 5200.1328 | 76.309 | 77,884 | 5199.124975 | 48 | 613 | 633 | QDVNCTEVPVAIHADQLTPTW | 0.0002000000000000000000.0 | 616(NCT) | 1 | 2396.1205 | 2393.132239 | 8493031 | 79.12 | 1 | 7 | 6 | 1 | 0 | 1 | 0 | 1 | 5199.126485 | -0.0015 | -0.29 | -2.81 |
| Chymotrypsin | 110 | 21 | 4545 | 1 | 4710 | 5200.1328 | 76.309 | 77,884 | 5199.124975 | 48 | 613 | 633 | QDVNCTEVPVAIHADQLTPTW | 0.0002000000000000000000.0 | 616(NCT) | 1 | 2396.1205 | 2393.132239 | 8493031 | 79.12 | 2 | 8 | 6 | 0 | 1 | 0 | 0 | 1 | 5199.126486 | -0.0015 | -0.29 | -2.81 |
| Chymotrypsin | 110 | 21 | 4917 | 1 | 4637 | 4820.0218 | 73.3159 | 70,297 | 4819.013975 | 48 | 613 | 633 | QDVNCTEVPVAIHADQLTPTW | 0.0002000000000000000000.0 | 616(NCT) | 1 | 2396.1205 | 2393.132239 | 8493031 | 79.12 | 1 | 6 | 5 | 3 | 0 | 0 | 0 | 1 | 4819.019775 | -0.0058 | -1.2 | -5.81 |
| Chymotrypsin | 110 | 21 | 5051 | 1 | 4630 | 4966.0755 | 73.0203 | 67,518 | 4965.067675 | 48 | 613 | 633 | QDVNCTEVPVAIHADQLTPTW | 0.0002000000000000000000.0 | 616(NCT) | 1 | 2396.1205 | 2393.132239 | 8493031 | 79.12 | 1 | 6 | 5 | 4 | 0 | 0 | 0 | 0 | 4965.077684 | -0.0100 | -2.01 | -6.10 |
| Chymotrypsin | 110 | 21 | 5667 | 1 | 4656 | 4219.8091 | 74.1216 | 57,368 | 4218.801275 | 48 | 613 | 633 | QDVNCTEVPVAIHADQLTPTW | 0.0002000000000000000000.0 | 616(NCT) | 1 | 2396.1205 | 2393.132239 | 8493031 | 79.12 | 1 | 5 | 5 | 0 | 0 | 0 | 0 | 1 | 4218.793224 | 0.0081 | 1.9 | -5.00 |
| Chymotrypsin | 110 | 21 | 5897 | 1 | 4648 | 4511.9168 | 73.7831 | 53,468 | 4510.908975 | 48 | 613 | 633 | QDVNCTEVPVAIHADQLTPTW | 0.0002000000000000000000.0 | 616(NCT) | 1 | 2396.1205 | 2393.132239 | 8493031 | 79.12 | 1 | 5 | 5 | 2 | 0 | 0 | 0 | 1 | 4510.909042 | -0.0001 | -0.01 | -5.34 |
| Chymotrypsin | 110 | 21 | 6709 | 1 | 4674 | 4893.0319 | 74.8727 | 42,219 | 4892.024075 | 48 | 613 | 633 | QDVNCTEVPVAIHADQLTPTW | 0.0002000000000000000000.0 | 616(NCT) | 1 | 2396.1205 | 2393.132239 | 8493031 | 79.12 | 1 | 7 | 6 | 1 | 0 | 0 | 0 | 1 | 4892.036154 | -0.0121 | -2.46 | -4.25 |
| Chymotrypsin | 110 | 21 | 6907 | 1 | 4668 | 3651.5998 | 74.6196 | 39,572 | 3650.591975 | 48 | 613 | 633 | QDVNCTEVPVAIHADQLTPTW | 0.0002000000000000000000.0 | 616(NCT) | 1 | 2396.1205 | 2393.132239 | 8493031 | 79.12 | 1 | 4 | 3 | 0 | 0 | 0 | 0 | 1 | 3650.581654 | 0.0103 | 2.82 | -4.50 |
| Chymotrypsin | 110 | 21 | 7006 | 1 | 4665 | 3610.5619 | 74.4901 | 37,915 | 3609.554075 | 48 | 613 | 633 | QDVNCTEVPVAIHADQLTPTW | 0.0002000000000000000000.0 | 616(NCT) | 1 | 2396.1205 | 2393.132239 | 8493031 | 79.12 | 1 | 5 | 2 | 0 | 0 | 0 | 0 | 1 | 3609.555105 | -0.0010 | -0.28 | -4.63 |
| Chymotrypsin | 235 | 4 | 1802 | 1 | 4679 | 4325.8311 | 75.0833 | 217,259 | 4324.823275 | 229 | 612 | 633 | YQDVNCTEVPVAIHADQLTPTW | 0.00002000000000000000000.0 | 616(NCT) | 1 | 2559.1838 | 2556.195539 | 1001733.9 | 79.94 | 1 | 5 | 4 | 1 | 0 | 0 | 0 | 1 | 4324.83506 | -0.0118 | -2.72 | -4.85 |
| Chymotrypsin | 235 | 4 | 3127 | 1 | 4671 | 4690.9684 | 74.7468 | 120,750 | 4689.960575 | 229 | 612 | 633 | YQDVNCTEVPVAIHADQLTPTW | 0.00002000000000000000000.0 | 616(NCT) | 1 | 2559.1838 | 2556.195539 | 1001733.9 | 79.94 | 1 | 6 | 5 | 1 | 0 | 0 | 0 | 1 | 4689.967257 | -0.0067 | -1.42 | -5.19 |
| Chymotrypsin | 235 | 4 | 3260 | 1 | 4663 | 5056.1065 | 74.4041 | 115,504 | 5055.098675 | 229 | 612 | 633 | YQDVNCTEVPVAIHADQLTPTW | 0.00002000000000000000000.0 | 616(NCT) | 1 | 2559.1838 | 2556.195539 | 1001733.9 | 79.94 | 1 | 7 | 6 | 1 | 0 | 0 | 0 | 1 | 5055.099454 | -0.0008 | -0.15 | -5.53 |
| Chymotrypsin | 235 | 4 | 5881 | 1 | 4675 | 4471.8922 | 74.9141 | 53,733 | 4470.884375 | 229 | 612 | 633 | YQDVNCTEVPVAIHADQLTPTW | 0.00002000000000000000000.0 | 616(NCT) | 1 | 2559.1838 | 2556.195539 | 1001733.9 | 79.94 | 1 | 5 | 4 | 2 | 0 | 0 | 0 | 1 | 4470.892969 | -0.0086 | -1.92 | -5.02 |
| Chymotrypsin | 251 | 6 | 2094 | 1 | 4647 | 5165.2272 | 73.7412 | 186,697 | 5164.219375 | 117 | 613 | 636 | QDVNCTEVPVAIHADQLTPTW | 0.000200000000000000000000.0 | 616(NCT) | 1 | 2814.3533 | 2811.365039 | 2689609 | 78.66 | 1 | 7 | 6 | 0 | 0 | 0 | 0 | 1 | 5164.211045 | 0.0083 | 1.61 | -4.92 |
| Chymotrypsin | 251 | 6 | 2215 | 1 | 4645 | 5311.2776 | 73.6559 | 177,026 | 5310.269775 | 117 | 613 | 636 | QDVNCTEVPVAIHADQLTPTW | 0.000200000000000000000000.0 | 616(NCT) | 1 | 2814.3533 | 2811.365039 | 2689609 | 78.66 | 1 | 7 | 6 | 1 | 0 | 0 | 0 | 1 | 5310.268954 | 0.0008 | 0.15 | -5.00 |
| Chymotrypsin | 251 | 6 | 3010 | 1 | 4652 | 4946.1384 | 73.9523 | 126,497 | 4945.130575 | 117 | 613 | 636 | QDVNCTEVPVAIHADQLTPTW | 0.000200000000000000000000.0 | 616(NCT) | 1 | 2814.3533 | 2811.365039 | 2689609 | 78.66 | 1 | 6 | 5 | 1 | 0 | 0 | 0 | 1 | 4945.136757 | -0.0062 | -1.25 | -4.70 |
| Chymotrypsin | 251 | 6 | 3785 | 1 | 4660 | 4581.0022 | 74.2802 | 97,538 | 4579.994375 | 117 | 613 | 636 | QDVNCTEVPVAIHADQLTPTW | 0.000200000000000000000000.0 | 616(NCT) | 1 | 2814.3533 | 2811.365039 | 2689609 | 78.66 | 1 | 5 | 4 | 1 | 0 | 0 | 0 | 1 | 4580.00456 | -0.0102 | -2.22 | -4.38 |
| Chymotrypsin | 251 | 6 | 4827 | 1 | 4643 | 5457.3298 | 73.5696 | 72,186 | 5456.321975 | 117 | 613 | 636 | QDVNCTEVPVAIHADQLTPTW | 0.000200000000000000000000.0 | 616(NCT) | 1 | 2814.3533 | 2811.365039 | 2689609 | 78.66 | 1 | 7 | 6 | 2 | 0 | 0 | 0 | 0 | 5456.326863 | -0.0049 | -0.89 | -5.09 |
| Chymotrypsin | 251 | 6 | 7592 | 1 | 4640 | 5603.3913 | 73.4449 | 30,336 | 5602.383475 | 117 | 613 | 636 | QDVNCTEVPVAIHADQLTPTW | 0.000200000000000000000000.0 | 616(NCT) | 1 | 2814.3533 | 2811.365039 | 2689609 | 78.66 | 1 | 7 | 6 | 3 | 0 | 0 | 0 | 0 | 5602.384772 | -0.0013 | -0.23 | -5.21 |
| alpha-Lytic | 54 | 11 | 288 | 1 | 4244 | 3610.4117 | 48.1773 | 455,276 | 3609.403875 | 32 | 610 | 618 | VLYQDVNCT | 0.000000200.0 | 616(NCT) | 1 | 1113.4898 | 1110.501539 | 2961110.8 | 55.24 | 1 | 7 | 6 | 1 | 0 | 0 | 0 | 1 | 3609.405454 | -0.0016 | -0.43 | -7.07 |
| alpha-Lytic | 54 | 11 | 297 | 1 | 4265 | 2880.1509 | 49.1818 | 440,525 | 2879.143075 | 32 | 610 | 618 | VLYQDVNCT | 0.000000200.0 | 616(NCT) | 1 | 1113.4898 | 1110.501539 | 2961110.8 | 55.24 | 1 | 5 | 4 | 1 | 0 | 0 | 0 | 1 | 2879.14106 | 0.0020 | 0.69 | -6.06 |
| alpha-Lytic | 54 | 11 | 475 | 1 | 4265 | 2734.0898 | 49.1818 | 269,995 | 2733.081975 | 32 | 610 | 618 | VLYQDVNCT | 0.000000200.0 | 616(NCT) | 1 | 1113.4898 | 1110.501539 | 2961110.8 | 55.24 | 1 | 5 | 4 | 0 | 0 | 0 | 0 | 1 | 2733.083151 | -0.0012 | -0.43 | -6.06 |
| alpha-Lytic | 54 | 11 | 561 | 1 | 4258 | 3245.2789 | 48.8497 | 237,312 | 3244.271075 | 32 | 610 | 618 | VLYQDVNCT | 0.000000200.0 | 616(NCT) | 1 | 1113.4898 | 1110.501539 | 2961110.8 | 55.24 | 1 | 6 | 5 | 1 | 0 | 0 | 0 | 1 | 3244.273257 | -0.0022 | -0.67 | -6.39 |
| alpha-Lytic | 54 | 11 | 787 | 1 | 4237 | 3756.468 | 47.8626 | 168,269 | 3755.460175 | 32 | 610 | 618 | VLYQDVNCT | 0.000000200.0 | 616(NCT) | 1 | 1113.4898 | 1110.501539 | 2961110.8 | 55.24 | 1 | 7 | 6 | 2 | 0 | 0 | 0 | 0 | 3755.463363 | -0.0032 | -0.84 | -7.38 |
| alpha-Lytic | 54 | 11 | 900 | 1 | 4268 | 3448.3641 | 49.3336 | 147,190 | 3447.356275 | 32 | 610 | 618 | VLYQDVNCT | 0.000000200.0 | 616(NCT) | 1 | 1113.4898 | 1110.501539 | 2961110.8 | 55.24 | 1 | 6 | 6 | 1 | 0 | 0 | 0 | 1 | 3447.35263 | 0.0036 | 1.05 | -5.91 |
| alpha-Lytic | 54 | 11 | 916 | 1 | 4257 | 3026.2118 | 48.7999 | 145,556 | 3025.203975 | 32 | 610 | 618 | VLYQDVNCT | 0.000000200.0 | 616(NCT) | 1 | 1113.4898 | 1110.501539 | 2961110.8 | 55.24 | 1 | 5 | 4 | 2 | 0 | 0 | 0 | 1 | 3025.198969 | 0.0050 | 1.65 | -6.44 |
| alpha-Lytic | 54 | 11 | 1037 | 1 | 4273 | 2531.0088 | 49.5869 | 129,063 | 2530.000975 | 32 | 610 | 618 | VLYQDVNCT | 0.000000200.0 | 616(NCT) | 1 | 1113.4898 | 1110.501539 | 2961110.8 | 55.24 | 1 | 5 | 3 | 0 | 0 | 0 | 0 | 1 | 2530.003778 | -0.0028 | -1.1 | -5.66 |
| alpha-Lytic | 54 | 11 | 1190 | 1 | 4233 | 3902.529 | 47.6653 | 110,296 | 3901.521175 | 32 | 610 | 618 | VLYQDVNCT | 0.000000200.0 | 616(NCT) | 1 | 1113.4898 | 1110.501539 | 2961110.8 | 55.24 | 1 | 7 | 6 | 3 | 0 | 0 | 0 | 0 | 3901.521272 | -0.0001 | -0.02 | -7.58 |
| alpha-Lytic | 54 | 11 | 1549 | 1 | 4271 | 3083.2339 | 49.4865 | 83,495 | 3082.226075 | 32 | 610 | 618 | VLYQDVNCT | 0.000000200.0 | 616(NCT) | 1 | 1113.4898 | 1110.501539 | 2961110.8 | 55.24 | 1 | 5 | 5 | 1 | 0 | 0 | 0 | 1 | 3082.220433 | 0.0056 | 1.83 | -5.76 |
| alpha-Lytic | 54 | 11 | 1894 | 1 | 4300 | 2822.1051 | 50.926 | 63,507 | 2821.097275 | 32 | 610 | 618 | VLYQDVNCT | 0.000000200.0 | 616(NCT) | 1 | 1113.4898 | 1110.501539 | 2961110.8 | 55.24 | 1 | 4 | 3 | 1 | 0 | 1 | 0 | 1 | 2821.099194 | -0.0019 | -0.68 | -4.32 |
| alpha-Lytic | 54 | 11 | 1894 | 1 | 4300 | 2822.1051 | 50.926 | 63,507 | 2821.097275 | 32 | 610 | 618 | VLYQDVNCT | 0.000000200.0 | 616(NCT) | 1 | 1113.4898 | 1110.501539 | 2961110.8 | 55.24 | 2 | 5 | 3 | 0 | 1 | 0 | 0 | 1 | 2821.099195 | -0.0019 | -0.68 | -4.32 |
| alpha-Lytic | 67 | 5 | 404 | 1 | 4117 | 3511.3451 | 41.9172 | 321,936 | 3510.337275 | 1006 | 611 | 618 | LYQDVNCT | - | 616(NCT) | 1 | | 1011.43313 | - | 46.00 | 1 | 7 | 6 | 1 | 0 | 0 | 0 | 1 | 3510.337045 | 0.0002 | 0.06 | -4.08 |
| alpha-Lytic | 67 | 5 | 500 | 1 | 4136 | 2781.0809 | 42.8605 | 259,708 | 2780.073075 | 1006 | 611 | 618 | LYQDVNCT | - | 616(NCT) | 1 | | 1011.43313 | - | 46.00 | 1 | 5 | 4 | 1 | 0 | 0 | 0 | 1 | 2780.072651 | 0.0004 | 0.15 | -3.14 |
| alpha-Lytic | 67 | 5 | 1413 | 1 | 4112 | 3657.4002 | 41.6615 | 91,022 | 3656.392375 | 1006 | 611 | 618 | LYQDVNCT | - | 616(NCT) | 1 | | 1011.43313 | - | 46.00 | 1 | 7 | 6 | 2 | 0 | 0 | 0 | 0 | 3656.394954 | -0.0026 | -0.7 | -4.34 |
| alpha-Lytic | 67 | 5 | 2505 | 1 | 4122 | 3146.2145 | 42.1679 | 42,473 | 3145.206675 | 1006 | 611 | 618 | LYQDVNCT | - | 616(NCT) | 1 | | 1011.43313 | - | 46.00 | 1 | 6 | 5 | 1 | 0 | 0 | 0 | 1 | 3145.204848 | 0.0018 | 0.58 | -3.83 |
| alpha-Lytic | 67 | 5 | 3525 | 1 | 4125 | 3292.2696 | 42.3109 | 18,468 | 3291.261775 | 1006 | 611 | 618 | LYQDVNCT | - | 616(NCT) | 1 | | 1011.43313 | - | 46.00 | 1 | 6 | 5 | 2 | 0 | 0 | 0 | 1 | 3291.262757 | -0.0010 | -0.29 | -3.69 |
| Chymotrypsin | 215 | 19 | 1411 | 1 | 4572 | 5368.2589 | 70.8534 | 272,585 | 5367.251075 | 73 | 644 | 674 | QTRAGCLIGAEHVNNSYECDIPIGAGICASY | 0.0000000000000020000000000000000.0 | 657(NNS) | 1 | 3398.5215 | 3395.533239 | 4677428.5 | 74.56 | 1 | 5 | 5 | 1 | 0 | 0 | 0 | 1 | 5367.252133 | -0.0011 | -0.19 | -3.70 |
| Chymotrypsin | 215 | 19 | 1773 | 1 | 4566 | 5457.3055 | 70.6012 | 220,058 | 5456.297675 | 73 | 644 | 674 | QTRAGCLIGAEHVNNSYECDIPIGAGICASY | 0.0000000000000020000000000000000.0 | 657(NNS) | 1 | 3398.5215 | 3395.533239 | 4677428.5 | 74.56 | 1 | 5 | 4 | 3 | 0 | 0 | 0 | 1 | 5456.288578 | 0.0091 | 1.66 | -3.96 |
| Chymotrypsin | 215 | 19 | 1795 | 1 | 4569 | 5514.299 | 70.722 | 217,797 | 5513.291175 | 73 | 644 | 674 | QTRAGCLIGAEHVNNSYECDIPIGAGICASY | 0.0000000000000020000000000000000.0 | 657(NNS) | 1 | 3398.5215 | 3395.533239 | 4677428.5 | 74.56 | 1 | 5 | 5 | 2 | 0 | 0 | 0 | 1 | 5513.310042 | -0.0189 | -3.42 | -3.83 |
| Chymotrypsin | 215 | 19 | 1991 | 1 | 4564 | 5879.4523 | 70.5137 | 195,013 | 5878.444475 | 73 | 644 | 674 | QTRAGCLIGAEHVNNSYECDIPIGAGICASY | 0.0000000000000020000000000000000.0 | 657(NNS) | 1 | 3398.5215 | 3395.533239 | 4677428.5 | 74.56 | 1 | 6 | 6 | 2 | 0 | 0 | 0 | 1 | 5878.442239 | 0.0022 | 0.38 | -4.04 |
| Chymotrypsin | 215 | 19 | 2023 | 1 | 4567 | 5660.3703 | 70.6445 | 192,271 | 5659.362475 | 73 | 644 | 674 | QTRAGCLIGAEHVNNSYECDIPIGAGICASY | 0.0000000000000020000000000000000.0 | 657(NNS) | 1 | 3398.5215 | 3395.533239 | 4677428.5 | 74.56 | 1 | 5 | 5 | 3 | 0 | 0 | 0 | 1 | 5659.367951 | -0.0055 | -0.96 | -3.91 |
| Chymotrypsin | 215 | 19 | 2554 | 1 | 4573 | 5165.177 | 70.8976 | 151,886 | 5164.169175 | 73 | 644 | 674 | QTRAGCLIGAEHVNNSYECDIPIGAGICASY | 0.0000000000000020000000000000000.0 | 657(NNS) | 1 | 3398.5215 | 3395.533239 | 4677428.5 | 74.56 | 1 | 5 | 4 | 1 | 0 | 0 | 0 | 1 | 5164.17276 | -0.0036 | -0.69 | -3.66 |
| Chymotrypsin | 215 | 19 | 2587 | 1 | 4570 | 5311.2304 | 70.7656 | 149,560 | 5310.222575 | 73 | 644 | 674 | QTRAGCLIGAEHVNNSYECDIPIGAGICASY | 0.0000000000000020000000000000000.0 | 657(NNS) | 1 | 3398.5215 | 3395.533239 | 4677428.5 | 74.56 | 1 | 5 | 4 | 2 | 0 | 0 | 0 | 1 | 5310.230669 | -0.0081 | -1.52 | -3.79 |
| Chymotrypsin | 215 | 19 | 2613 | 1 | 4553 | 6333.6232 | 70.0474 | 148,011 | 6332.615375 | 73 | 644 | 674 | QTRAGCLIGAEHVNNSYECDIPIGAGICASY | 0.0000000000000020000000000000000.0 | 657(NNS) | 1 | 3398.5215 | 3395.533239 | 4677428.5 | 74.56 | 1 | 7 | 6 | 4 | 0 | 0 | 0 | 1 | 6332.610881 | 0.0045 | 0.7 | -4.51 |
| Chymotrypsin | 215 | 19 | 2613 | 1 | 4553 | 6333.6232 | 70.0474 | 148,011 | 6332.615375 | 73 | 644 | 674 | QTRAGCLIGAEHVNNSYECDIPIGAGICASY | 0.0000000000000020000000000000000.0 | 657(NNS) | 1 | 3398.5215 | 3395.533239 | 4677428.5 | 74.56 | 2 | 2 | 10 | 0 | 2 | 0 | 1 | -1 | 6332.623451 | -0.0081 | -1.27 | -4.51 |
| Chymotrypsin | 215 | 19 | 2979 | 1 | 4558 | 6171.5739 | 70.2543 | 128,452 | 6170.566075 | 73 | 644 | 674 | QTRAGCLIGAEHVNNSYECDIPIGAGICASY | 0.0000000000000020000000000000000.0 | 657(NNS) | 1 | 3398.5215 | 3395.533239 | 4677428.5 | 74.56 | 1 | 6 | 6 | 4 | 0 | 0 | 0 | 0 | 6170.558057 | 0.0080 | 1.29 | -4.30 |
| Chymotrypsin | 215 | 19 | 3047 | 1 | 4561 | 6025.5089 | 70.3832 | 124,447 | 6024.501075 | 73 | 644 | 674 | QTRAGCLIGAEHVNNSYECDIPIGAGICASY | 0.0000000000000020000000000000000.0 | 657(NNS) | 1 | 3398.5215 | 3395.533239 | 4677428.5 | 74.56 | 1 | 6 | 6 | 3 | 0 | 0 | 0 | 1 | 6024.500148 | 0.0009 | 0.15 | -4.17 |
| Chymotrypsin | 215 | 19 | 3880 | 1 | 4556 | 6187.5529 | 70.1696 | 95,394 | 6186.545075 | 73 | 644 | 674 | QTRAGCLIGAEHVNNSYECDIPIGAGICASY | 0.0000000000000020000000000000000.0 | 657(NNS) | 1 | 3398.5215 | 3395.533239 | 4677428.5 | 74.56 | 1 | 7 | 6 | 3 | 0 | 0 | 0 | 0 | 6186.552972 | -0.0079 | -1.27 | -4.39 |
| Chymotrypsin | 215 | 19 | 4029 | 1 | 4560 | 5822.4327 | 70.3415 | 91,029 | 5821.424875 | 73 | 644 | 674 | QTRAGCLIGAEHVNNSYECDIPIGAGICASY | 0.0000000000000020000000000000000.0 | 657(NNS) | 1 | 3398.5215 | 3395.533239 | 4677428.5 | 74.56 | 1 | 6 | 5 | 3 | 0 | 0 | 0 | 1 | 5821.420775 | 0.0041 | 0.7 | -4.22 |
| Chymotrypsin | 215 | 19 | 4315 | 1 | 4555 | 6244.5866 | 70.1262 | 83,397 | 6243.578775 | 73 | 644 | 674 | QTRAGCLIGAEHVNNSYECDIPIGAGICASY | 0.0000000000000020000000000000000.0 | 657(NNS) | 1 | 3398.5215 | 3395.533239 | 4677428.5 | 74.56 | 1 | 7 | 7 | 2 | 0 | 0 | 0 | 0 | 6243.574436 | 0.0043 | 0.69 | -4.43 |
| Chymotrypsin | 215 | 19 | 4381 | 1 | 4575 | 5108.1605 | 70.984 | 81,941 | 5107.152675 | 73 | 644 | 674 | QTRAGCLIGAEHVNNSYECDIPIGAGICASY | 0.0000000000000020000000000000000.0 | 657(NNS) | 1 | 3398.5215 | 3395.533239 | 4677428.5 | 74.56 | 1 | 5 | 3 | 2 | 0 | 0 | 0 | 0 | 5107.151296 | 0.0014 | 0.27 | -3.57 |
| Chymotrypsin | 215 | 19 | 5191 | 1 | 4557 | 6041.4899 | 70.2108 | 65,245 | 6040.482075 | 73 | 644 | 674 | QTRAGCLIGAEHVNNSYECDIPIGAGICASY | 0.0000000000000020000000000000000.0 | 657(NNS) | 1 | 3398.5215 | 3395.533239 | 4677428.5 | 74.56 | 1 | 7 | 6 | 2 | 0 | 0 | 0 | 0 | 6040.495063 | -0.0130 | -2.15 | -4.35 |
| Chymotrypsin | 215 | 19 | 5344 | 1 | 4541 | 5061.1867 | 69.528 | 62,672 | 5060.178875 | 73 | 644 | 674 | QTRAGCLIGAEHVNNSYECDIPIGAGICASY | 0.0000000000000020000000000000000.0 | 657(NNS) | 1 | 3398.5215 | 3395.533239 | 4677428.5 | 74.56 | | | | | | | | | | | | |
| Chymotrypsin | 215 | 19 | 5381 | 1 | 4554 | 6390.6461 | 70.0915 | 61,993 | 6389.638275 | 73 | 644 | 674 | QTRAGCLIGAEHVNNSYECDIPIGAGICASY | 0.0000000000000020000000000000000.0 | 657(NNS) | 1 | 3398.5215 | 3395.533239 | 4677428.5 | 74.56 | 1 | 7 | 7 | 3 | 0 | 0 | 0 | 0 | 6389.632345 | 0.0059 | 0.92 | -4.47 |
| Chymotrypsin | 215 | 19 | 8381 | 1 | 4622 | 6348.5802 | 72.7051 | 20,511 | 6347.572375 | 73 | 644 | 674 | QTRAGCLIGAEHVNNSYECDIPIGAGICASY | 0.0000000000000020000000000000000.0 | 657(NNS) | 1 | 3398.5215 | 3395.533239 | 4677428.5 | 74.56 | 1 | 8 | 6 | 1 | 1 | 0 | 0 | 1 | 6347.585395 | -0.0130 | -2.05 | -1.85 |
| Chymotrypsin | 215 | 19 | 8381 | 1 | 4622 | 6348.5802 | 72.7051 | 20,511 | 6347.572375 | 73 | 644 | 674 | QTRAGCLIGAEHVNNSYECDIPIGAGICASY | 0.0000000000000020000000000000000.0 | 657(NNS) | 1 | 3398.5215 | 3395.533239 | 4677428.5 | 74.56 | 2 | 7 | 6 | 2 | 0 | 1 | 0 | 0 | 6347.585394 | -0.0130 | -2.05 | -1.85 |
| Chymotrypsin | 215 | 19 | 8554 | 1 | 4620 | 6494.6632 | 72.6193 | 17,995 | 6493.655375 | 73 | 644 | 674 | QTRAGCLIGAEHVNNSYECDIPIGAGICASY | 0.0000000000000020000000000000000.0 | 657(NNS) | 1 | 3398.5215 | 3395.533239 | 4677428.5 | 74.56 | 1 | 8 | 6 | 2 | 1 | 0 | 0 | 0 | 6493.643304 | 0.0121 | 1.85 | -1.94 |
| Chymotrypsin | 215 | 19 | 8554 | 1 | 4620 | 6494.6632 | 72.6193 | 17,995 | 6493.655375 | 73 | 644 | 674 | QTRAGCLIGAEHVNNSYECDIPIGAGICASY | 0.0000000000000020000000000000000.0 | 657(NNS) | 1 | 3398.5215 | 3395.533239 | 4677428.5 | 74.56 | 2 | 7 | 6 | 3 | 0 | 1 | 0 | 0 | 6493.643303 | 0.0121 | 1.85 | -1.94 |
| alpha-Lytic | 122 | 5 | 1887 | 1 | 4474 | 3760.5084 | 59.6188 | 63,810 | 3759.500575 | 21 | 653 | 668 | AEHVNNSYECDIPIGA | 0.0000200000000000.0 | 657(NNS) | 1 | 1790.7667 | 1787.778439 | 5874644.5 | 64.55 | 1 | 5 | 5 | 1 | 0 | 0 | 0 | 1 | 3759.497333 | 0.0032 | 0.86 | -4.93 |
| alpha-Lytic | 122 | 5 | 1920 | 1 | 4464 | 4052.6228 | 59.1478 | 62,629 | 4051.614975 | 21 | 653 | 668 | AEHVNNSYECDIPIGA | 0.0000200000000000.0 | 657(NNS) | 1 | 1790.7667 | 1787.778439 | 5874644.5 | 64.55 | 1 | 5 | 5 | 3 | 0 | 0 | 0 | 1 | 4051.613151 | 0.0018 | 0.45 | -5.40 |
| alpha-Lytic | 122 | 5 | 1997 | 1 | 4469 | 3906.5621 | 59.3848 | 59,166 | 3905.554275 | 21 | 653 | 668 | AEHVNNSYECDIPIGA | 0.0000200000000000.0 | 657(NNS) | 1 | 1790.7667 | 1787.778439 | 5874644.5 | 64.55 | 1 | 5 | 5 | 2 | 0 | 0 | 0 | 1 | 3905.555242 | -0.0010 | -0.24 | -5.17 |
| alpha-Lytic | 122 | 5 | 2187 | 1 | 4463 | 3849.5379 | 59.1043 | 52,750 | 3848.530075 | 21 | 653 | 668 | AEHVNNSYECDIPIGA | 0.0000200000000000.0 | 657(NNS) | 1 | 1790.7667 | 1787.778439 | 5874644.5 | 64.55 | 1 | 5 | 4 | 3 | 0 | 0 | 0 | 1 | 3848.533778 | -0.0037 | -0.96 | -5.45 |
| alpha-Lytic | 122 | 5 | 2774 | 1 | 4462 | 4271.6957 | 59.054 | 35,510 | 4270.687875 | 21 | 653 | 668 | AEHVNNSYECDIPIGA | 0.0000200000000000.0 | 657(NNS) | 1 | 1790.7667 | 1787.778439 | 5874644.5 | 64.55 | 1 | 6 | 6 | 2 | 0 | 0 | 0 | 1 | 4270.687439 | 0.0004 | 0.1 | -5.50 |
| alpha-Lytic | 12 | 5 | 56 | 1 | 4414 | 2458.0137 | 56.6331 | 2,342,680 | 2457.005875 | 1009 | 707 | 716 | YSNNSIAIPT | - | 709(NNS) | 1 | | 1078.52947 | - | 61.00 | 1 | 6 | 2 | 0 | 0 | 0 | 0 | 1 | 2457.00516 | 0.0007 | 0.29 | -4.37 |
| alpha-Lytic | 12 | 5 | 75 | 1 | 4408 | 2620.0638 | 56.3319 | 1,711,110 | 2619.055975 | 1009 | 707 | 716 | YSNNSIAIPT | - | 709(NNS) | 1 | | 1078.52947 | - | 61.00 | 1 | 7 | 2 | 0 | 0 | 0 | 0 | 1 | 2619.057984 | -0.0020 | -0.76 | -4.67 |
| alpha-Lytic | 12 | 5 | 86 | 1 | 4403 | 2782.117 | 56.0677 | 1,545,130 | 2781.109175 | 1009 | 707 | 716 | YSNNSIAIPT | - | 709(NNS) | 1 | | 1078.52947 | - | 61.00 | 1 | 8 | 2 | 0 | 0 | 0 | 0 | 1 | 2781.110808 | -0.0016 | -0.58 | -4.93 |
| alpha-Lytic | 12 | 5 | 147 | 1 | 4420 | 2295.961 | 56.9532 | 949,055 | 2294.953175 | 1009 | 707 | 716 | YSNNSIAIPT | - | 709(NNS) | 1 | | 1078.52947 | - | 61.00 | 1 | 5 | 2 | 0 | 0 | 0 | 0 | 1 | 2294.952336 | 0.0008 | 0.36 | -4.05 |
| alpha-Lytic | 12 | 5 | 1065 | 1 | 4412 | 2661.0928 | 56.5338 | 124,758 | 2660.084975 | 1009 | 707 | 716 | YSNNSIAIPT | - | 709(NNS) | 1 | | 1078.52947 | - | 61.00 | 1 | 6 | 3 | 0 | 0 | 0 | 0 | 1 | 2660.084533 | 0.0004 | 0.16 | -4.47 |
| alpha-Lytic | 69 | 16 | 445 | 1 | 4388 | 3578.4411 | 55.2969 | 284,513 | 3577.433275 | 1009 | 707 | 716 | YSNNSIAIPT | - | 709(NNS) | 1 | | 1078.52947 | - | 61.00 | 1 | 7 | 6 | 1 | 0 | 0 | 0 | 1 | 3577.433385 | -0.0001 | -0.03 | -5.70 |
| alpha-Lytic | 69 | 16 | 717 | 1 | 4412 | 2848.1747 | 56.5338 | 186,497 | 2847.166875 | 1009 | 707 | 716 | YSNNSIAIPT | - | 709(NNS) | 1 | | 1078.52947 | - | 61.00 | 1 | 5 | 4 | 1 | 0 | 0 | 0 | 1 | 2847.168991 | -0.0021 | -0.74 | -4.47 |
| alpha-Lytic | 69 | 16 | 1135 | 1 | 4412 | 2702.1187 | 56.5338 | 115,702 | 2701.110875 | 1009 | 707 | 716 | YSNNSIAIPT | - | 709(NNS) | 1 | | 1078.52947 | - | 61.00 | 1 | 5 | 4 | 0 | 0 | 0 | 0 | 1 | 2701.111082 | -0.0002 | -0.07 | -4.47 |
| alpha-Lytic | 69 | 16 | 1150 | 1 | 4381 | 3724.4963 | 54.9622 | 114,582 | 3723.488475 | 1009 | 707 | 716 | YSNNSIAIPT | - | 709(NNS) | 1 | | 1078.52947 | - | 61.00 | 1 | 7 | 6 | 2 | 0 | 0 | 0 | 0 | 3723.491294 | -0.0028 | -0.75 | -6.04 |
| alpha-Lytic | 69 | 16 | 1263 | 1 | 4400 | 3213.3091 | 55.916 | 103,711 | 3212.301275 | 1009 | 707 | 716 | YSNNSIAIPT | - | 709(NNS) | 1 | | 1078.52947 | - | 61.00 | 1 | 6 | 5 | 1 | 0 | 0 | 0 | 1 | 3212.301188 | 0.0001 | 0.02 | -5.08 |
| alpha-Lytic | 69 | 16 | 1472 | 1 | 4390 | 3416.3906 | 55.3939 | 87,543 | 3415.382775 | 1009 | 707 | 716 | YSNNSIAIPT | - | 709(NNS) | 1 | | 1078.52947 | - | 61.00 | 1 | 6 | 6 | 1 | 0 | 0 | 0 | 1 | 3415.380561 | 0.0022 | 0.64 | -5.61 |
| alpha-Lytic | 69 | 16 | 1668 | 1 | 4421 | 2889.2027 | 56.9971 | 75,381 | 2888.194875 | 1009 | 707 | 716 | YSNNSIAIPT | - | 709(NNS) | 1 | | 1078.52947 | - | 61.00 | 1 | 4 | 5 | 1 | 0 | 0 | 0 | 1 | 2888.19554 | -0.0007 | -0.23 | -4.00 |
| alpha-Lytic | 69 | 16 | 1832 | 1 | 4412 | 3254.3364 | 56.5338 | 66,368 | 3253.328575 | 1009 | 707 | 716 | YSNNSIAIPT | - | 709(NNS) | 1 | | 1078.52947 | - | 61.00 | 1 | 5 | 6 | 1 | 0 | 0 | 0 | 1 | 3253.327737 | 0.0008 | 0.25 | -4.47 |
| alpha-Lytic | 69 | 16 | 1879 | 1 | 4376 | 3870.5533 | 54.7073 | 64,082 | 3869.545475 | 1009 | 707 | 716 | YSNNSIAIPT | - | 709(NNS) | 1 | | 1078.52947 | - | 61.00 | 1 | 7 | 6 | 3 | 0 | 0 | 0 | 0 | 3869.549203 | -0.0037 | -0.96 | -6.29 |
| alpha-Lytic | 69 | 16 | 2280 | 1 | 4389 | 3619.4586 | 55.3468 | 49,666 | 3618.450775 | 1009 | 707 | 716 | YSNNSIAIPT | - | 709(NNS) | 1 | | 1078.52947 | - | 61.00 | 1 | 6 | 7 | 1 | 0 | 0 | 0 | 1 | 3618.459934 | -0.0092 | -2.53 | -5.65 |
| alpha-Lytic | 69 | 16 | 2288 | 1 | 4372 | 4016.6187 | 54.5138 | 49,244 | 4015.610875 | 1009 | 707 | 716 | YSNNSIAIPT | - | 709(NNS) | 1 | | 1078.52947 | - | 61.00 | 1 | 7 | 6 | 4 | 0 | 0 | 0 | 1 | 4015.607112 | 0.0038 | 0.93 | -6.49 |
| alpha-Lytic | 69 | 16 | 2288 | 1 | 4372 | 4016.6187 | 54.5138 | 49,244 | 4015.610875 | 1009 | 707 | 716 | YSNNSIAIPT | - | 709(NNS) | 1 | | 1078.52947 | - | 61.00 | 2 | 2 | 10 | 0 | 2 | 0 | 1 | -1 | 4015.619682 | -0.0088 | -2.19 | -6.49 |
| alpha-Lytic | 69 | 16 | 2469 | 1 | 4396 | 3254.3375 | 55.7021 | 43,401 | 3253.329675 | 1009 | 707 | 716 | YSNNSIAIPT | - | 709(NNS) | 1 | | 1078.52947 | - | 61.00 | 1 | 5 | 6 | 1 | 0 | 0 | 0 | 1 | 3253.327737 | 0.0019 | 0.59 | -5.30 |
| alpha-Lytic | 69 | 16 | 2503 | 1 | 4392 | 3359.3668 | 55.5051 | 42,501 | 3358.358975 | 1009 | 707 | 716 | YSNNSIAIPT | - | 709(NNS) | 1 | | 1078.52947 | - | 61.00 | 1 | 6 | 5 | 2 | 0 | 0 | 0 | 1 | 3358.359097 | -0.0001 | -0.03 | -5.49 |
| alpha-Lytic | 69 | 16 | 2624 | 1 | 4444 | 3885.5299 | 58.1864 | 39,017 | 3884.522075 | 1009 | 707 | 716 | YSNNSIAIPT | - | 709(NNS) | 1 | | 1078.52947 | - | 61.00 | 1 | 7 | 6 | 1 | 0 | 1 | 0 | 1 | 3884.523716 | -0.0016 | -0.42 | -2.81 |
| alpha-Lytic | 69 | 16 | 2624 | 1 | 4444 | 3885.5299 | 58.1864 | 39,017 | 3884.522075 | 1009 | 707 | 716 | YSNNSIAIPT | - | 709(NNS) | 1 | | 1078.52947 | - | 61.00 | 2 | 8 | 6 | 0 | 1 | 0 | 0 | 1 | 3884.523717 | -0.0016 | -0.42 | -2.81 |
| alpha-Lytic | 69 | 16 | 2721 | 1 | 4408 | 3416.3895 | 56.3319 | 36,828 | 3415.381675 | 1009 | 707 | 716 | YSNNSIAIPT | - | 709(NNS) | 1 | | 1078.52947 | - | 61.00 | 1 | 6 | 6 | 1 | 0 | 0 | 0 | 1 | 3415.380561 | 0.0011 | 0.32 | -4.67 |
| alpha-Lytic | 69 | 16 | 2927 | 1 | 4419 | 2743.1442 | 56.9025 | 31,673 | 2742.136375 | 1009 | 707 | 716 | YSNNSIAIPT | - | 709(NNS) | 1 | | 1078.52947 | - | 61.00 | 1 | 4 | 5 | 0 | 0 | 0 | 0 | 1 | 2742.137631 | -0.0013 | -0.45 | -4.10 |
| alpha-Lytic | 4 | 9 | 19 | 1 | 4238 | 2121.8211 | 47.9046 | 7,128,640 | 2120.813275 | 1010 | 717 | 721 | NFTIS | - | 717(NFT) | 1 | | 580.28566 | - | 53.00 | 1 | 7 | 2 | 0 | 0 | 0 | 0 | 1 | 2120.814174 | -0.0009 | -0.42 | -5.10 |
| alpha-Lytic | 4 | 9 | 36 | 1 | 4251 | 1959.7688 | 48.5069 | 4,348,810 | 1958.760975 | 1010 | 717 | 721 | NFTIS | - | 717(NFT) | 1 | | 580.28566 | - | 53.00 | 1 | 6 | 2 | 0 | 0 | 0 | 0 | 1 | 1958.76135 | -0.0004 | -0.19 | -4.49 |
| alpha-Lytic | 4 | 9 | 67 | 1 | 4251 | 2162.8472 | 48.5069 | 1,962,420 | 2161.839375 | 1010 | 717 | 721 | NFTIS | - | 717(NFT) | 1 | | 580.28566 | - | 53.00 | 1 | 6 | 3 | 0 | 0 | 0 | 0 | 1 | 2161.840723 | -0.0013 | -0.62 | -4.49 |
| alpha-Lytic | 4 | 9 | 110 | 1 | 4239 | 2308.9068 | 47.9505 | 1,214,120 | 2307.898975 | 1010 | 717 | 721 | NFTIS | - | 717(NFT) | 1 | | 580.28566 | - | 53.00 | 1 | 6 | 3 | 1 | 0 | 0 | 0 | 1 | 2307.898632 | 0.0003 | 0.14 | -5.05 |
| alpha-Lytic | 4 | 9 | 232 | 1 | 4266 | 2000.7948 | 49.2292 | 588,392 | 1999.786975 | 1010 | 717 | 721 | NFTIS | - | 717(NFT) | 1 | | 580.28566 | - | 53.00 | 1 | 5 | 3 | 0 | 0 | 0 | 0 | 1 | 1999.787899 | -0.0009 | -0.46 | -3.77 |
| alpha-Lytic | 4 | 9 | 346 | 1 | 4235 | 2283.8738 | 47.7664 | 387,594 | 2282.865975 | 1010 | 717 | 721 | NFTIS | - | 717(NFT) | 1 | | 580.28566 | - | 53.00 | 1 | 8 | 2 | 0 | 0 | 0 | 0 | 1 | 2282.866998 | -0.0010 | -0.44 | -5.23 |
| alpha-Lytic | 4 | 9 | 488 | 1 | 4254 | 2146.8558 | 48.6504 | 264,297 | 2145.847975 | 1010 | 717 | 721 | NFTIS | - | 717(NFT) | 1 | | 580.28566 | - | 53.00 | 1 | 5 | 3 | 1 | 0 | 0 | 0 | 1 | 2145.845808 | 0.0022 | 1 | -4.35 |
| alpha-Lytic | 4 | 9 | 1303 | 1 | 4328 | 2469.9375 | 52.3094 | 100,313 | 2468.929675 | 1010 | 717 | 721 | NFTIS | - | 717(NFT) | 1 | | 580.28566 | - | 53.00 | 1 | 6 | 3 | 0 | 0 | 1 | 0 | 1 | 2468.931054 | -0.0014 | -0.55 | -0.69 |
| alpha-Lytic | 4 | 9 | 1310 | 1 | 4319 | 2469.9383 | 51.8712 | 99,543 | 2468.930475 | 1010 | 717 | 721 | NFTIS | - | 717(NFT) | 1 | | 580.28566 | - | 53.00 | 1 | 6 | 3 | 0 | 0 | 1 | 0 | 1 | 2468.931054 | -0.0006 | -0.23 | -1.13 |
| Chymotrypsin | 42 | 29 | 121 | 1 | 4296 | 2970.2605 | 59.4065 | 2,623,220 | 2969.252675 | 86 | 801 | 813 | NFSQILPDPSKPS | 0.2000000000000.0 | 801(NFS) | 1 | 1431.7132 | 1428.724939 | 3756061 | 67.81 | 1 | 7 | 2 | 0 | 0 | 0 | 0 | 1 | 2969.253453 | -0.0008 | -0.26 | -8.40 |
| Chymotrypsin | 42 | 29 | 169 | 1 | 4308 | 2808.2079 | 59.8489 | 1,887,910 | 2807.200075 | 86 | 801 | 813 | NFSQILPDPSKPS | 0.2000000000000.0 | 801(NFS) | 1 | 1431.7132 | 1428.724939 | 3756061 | 67.81 | 1 | 6 | 2 | 0 | 0 | 0 | 0 | 1 | 2807.200629 | -0.0006 | -0.19 | -7.96 |
| Chymotrypsin | 42 | 29 | 245 | 1 | 4321 | 2646.1542 | 60.2996 | 1,363,720 | 2645.146375 | 86 | 801 | 813 | NFSQILPDPSKPS | 0.2000000000000.0 | 801(NFS) | 1 | 1431.7132 | 1428.724939 | 3756061 | 67.81 | 1 | 5 | 2 | 0 | 0 | 0 | 0 | 1 | 2645.147805 | -0.0014 | -0.54 | -7.51 |
| Chymotrypsin | 42 | 29 | 286 | 1 | 4296 | 3157.3455 | 59.4065 | 1,208,260 | 3156.337675 | 86 | 801 | 813 | NFSQILPDPSKPS | 0.2000000000000.0 | 801(NFS) | 1 | 1431.7132 | 1428.724939 | 3756061 | 67.81 | 1 | 6 | 3 | 1 | 0 | 0 | 0 | 1 | 3156.337911 | -0.0002 | -0.07 | -8.40 |
| Chymotrypsin | 42 | 29 | 396 | 1 | 4296 | 3132.3131 | 59.4065 | 869,136 | 3131.305275 | 86 | 801 | 813 | NFSQILPDPSKPS | 0.2000000000000.0 | 801(NFS) | 1 | 1431.7132 | 1428.724939 | 3756061 | 67.81 | 1 | 8 | 2 | 0 | 0 | 0 | 0 | 1 | 3131.306277 | -0.0010 | -0.31 | -8.40 |
| Chymotrypsin | 42 | 29 | 461 | 1 | 4306 | 2995.2926 | 59.7633 | 771,400 | 2994.284775 | 86 | 801 | 813 | NFSQILPDPSKPS | 0.2000000000000.0 | 801(NFS) | 1 | 1431.7132 | 1428.724939 | 3756061 | 67.81 | 1 | 5 | 3 | 1 | 0 | 0 | 0 | 1 | 2994.285087 | -0.0003 | -0.1 | -8.05 |
| Chymotrypsin | 42 | 29 | 693 | 1 | 4315 | 2849.2341 | 60.078 | 525,159 | 2848.226275 | 86 | 801 | 813 | NFSQILPDPSKPS | 0.2000000000000.0 | 801(NFS) | 1 | 1431.7132 | 1428.724939 | 3756061 | 67.81 | 1 | 5 | 3 | 0 | 0 | 0 | 0 | 1 | 2848.227178 | -0.0009 | -0.31 | -7.73 |
| Chymotrypsin | 42 | 29 | 819 | 1 | 4292 | 3198.3669 | 59.2355 | 453,614 | 3197.359075 | 86 | 801 | 813 | NFSQILPDPSKPS | 0.2000000000000.0 | 801(NFS) | 1 | 1431.7132 | 1428.724939 | 3756061 | 67.81 | 1 | 5 | 4 | 1 | 0 | 0 | 0 | 1 | 3197.36446 | -0.0054 | -1.68 | -8.58 |
| Chymotrypsin | 42 | 29 | 1028 | 1 | 4306 | 3011.2879 | 59.7633 | 369,053 | 3010.280075 | 86 | 801 | 813 | NFSQILPDPSKPS | 0.2000000000000.0 | 801(NFS) | 1 | 1431.7132 | 1428.724939 | 3756061 | 67.81 | 1 | 6 | 3 | 0 | 0 | 0 | 0 | 1 | 3010.280002 | 0.0001 | 0.02 | -8.05 |
| Chymotrypsin | 42 | 29 | 1202 | 1 | 4282 | 3344.4298 | 58.8341 | 318,513 | 3343.421975 | 86 | 801 | 813 | NFSQILPDPSKPS | 0.2000000000000.0 | 801(NFS) | 1 | 1431.7132 | 1428.724939 | 3756061 | 67.81 | 1 | 5 | 4 | 2 | 0 | 0 | 0 | 1 | 3343.422369 | -0.0004 | -0.11 | -8.98 |
| Chymotrypsin | 42 | 29 | 1203 | 1 | 4301 | 3052.3132 | 59.6181 | 318,369 | 3051.305375 | 86 | 801 | 813 | NFSQILPDPSKPS | 0.2000000000000.0 | 801(NFS) | 1 | 1431.7132 | 1428.724939 | 3756061 | 67.81 | 1 | 5 | 4 | 0 | 0 | 0 | 0 | 1 | 3051.306551 | -0.0012 | -0.38 | -8.19 |
| Chymotrypsin | 42 | 29 | 1607 | 1 | 4395 | 3318.3746 | 63.4102 | 241,689 | 3317.366775 | 86 | 801 | 813 | NFSQILPDPSKPS | 0.2000000000000.0 | 801(NFS) | 1 | 1431.7132 | 1428.724939 | 3756061 | 67.81 | 1 | 6 | 3 | 0 | 0 | 1 | 0 | 1 | 3317.370333 | -0.0036 | -1.07 | -4.40 |
| Chymotrypsin | 42 | 29 | 1635 | 1 | 4288 | 3401.4485 | 59.0768 | 238,676 | 3400.440675 | 86 | 801 | 813 | NFSQILPDPSKPS | 0.2000000000000.0 | 801(NFS) | 1 | 1431.7132 | 1428.724939 | 3756061 | 67.81 | 1 | 5 | 5 | 1 | 0 | 0 | 0 | 1 | 3400.443833 | -0.0032 | -0.92 | -8.73 |
| Chymotrypsin | 42 | 29 | 2178 | 1 | 4282 | 3547.5106 | 58.8341 | 179,773 | 3546.502775 | 86 | 801 | 813 | NFSQILPDPSKPS | 0.2000000000000.0 | 801(NFS) | 1 | 1431.7132 | 1428.724939 | 3756061 | 67.81 | 1 | 5 | 5 | 2 | 0 | 0 | 0 | 1 | 3546.501742 | 0.0010 | 0.29 | -8.98 |
| Chymotrypsin | 42 | 29 | 2341 | 1 | 4308 | 3093.3398 | 59.8489 | 165,935 | 3092.331975 | 86 | 801 | 813 | NFSQILPDPSKPS | 0.2000000000000.0 | 801(NFS) | 1 | 1431.7132 | 1428.724939 | 3756061 | 67.81 | 1 | 4 | 5 | 0 | 0 | 0 | 0 | 1 | 3092.3331 | -0.0011 | -0.36 | -7.96 |
| Chymotrypsin | 42 | 29 | 2477 | 1 | 4296 | 3239.4009 | 59.4065 | 156,681 | 3238.393075 | 86 | 801 | 813 | NFSQILPDPSKPS | 0.2000000000000.0 | 801(NFS) | 1 | 1431.7132 | 1428.724939 | 3756061 | 67.81 | 1 | 4 | 5 | 1 | 0 | 0 | 0 | 1 | 3238.391009 | 0.0021 | 0.63 | -8.40 |
| Chymotrypsin | 42 | 29 | 2772 | 1 | 4296 | 3255.3969 | 59.4065 | 138,677 | 3254.389075 | 86 | 801 | 813 | NFSQILPDPSKPS | 0.2000000000000.0 | 801(NFS) | 1 | 1431.7132 | 1428.724939 | 3756061 | 67.81 | 1 | 5 | 5 | 0 | 0 | 0 | 0 | 1 | 3254.385924 | 0.0032 | 0.96 | -8.40 |
| Chymotrypsin | 42 | 29 | 3544 | 1 | 4391 | 3359.4027 | 63.2382 | 105,644 | 3358.394875 | 86 | 801 | 813 | NFSQILPDPSKPS | 0.2000000000000.0 | 801(NFS) | 1 | 1431.7132 | 1428.724939 | 3756061 | 67.81 | 1 | 5 | 4 | 0 | 0 | 1 | 0 | 1 | 3358.396882 | -0.0020 | -0.59 | -4.57 |
| Chymotrypsin | 42 | 29 | 4016 | 1 | 4286 | 3766.593 | 58.9906 | 91,411 | 3765.585175 | 86 | 801 | 813 | NFSQILPDPSKPS | 0.2000000000000.0 | 801(NFS) | 1 | 1431.7132 | 1428.724939 | 3756061 | 67.81 | 1 | 6 | 6 | 1 | 0 | 0 | 0 | 1 | 3765.57603 | 0.0091 | 2.42 | -8.82 |
| Chymotrypsin | 42 | 29 | 4402 | 1 | 4322 | 2931.2834 | 60.3441 | 81,510 | 2930.275575 | 86 | 801 | 813 | NFSQILPDPSKPS | 0.2000000000000.0 | 801(NFS) | 1 | 1431.7132 | 1428.724939 | 3756061 | 67.81 | 1 | 3 | 5 | 0 | 0 | 0 | 0 | 1 | 2930.280276 | -0.0047 | -1.6 | -7.47 |
| Chymotrypsin | 42 | 29 | 4498 | 1 | 4275 | 3490.4765 | 58.5361 | 79,077 | 3489.468675 | 86 | 801 | 813 | NFSQILPDPSKPS | 0.2000000000000.0 | 801(NFS) | 1 | 1431.7132 | 1428.724939 | 3756061 | 67.81 | 1 | 5 | 4 | 3 | 0 | 0 | 0 | 1 | 3489.480278 | -0.0116 | -3.32 | -9.28 |
| Chymotrypsin | 42 | 29 | 4546 | 1 | 4294 | 3214.3648 | 59.3213 | 77,860 | 3213.356975 | 86 | 801 | 813 | NFSQILPDPSKPS | 0.2000000000000.0 | 801(NFS) | 1 | 1431.7132 | 1428.724939 | 3756061 | 67.81 | 1 | 6 | 4 | 0 | 0 | 0 | 0 | 1 | 3213.359375 | -0.0024 | -0.74 | -8.49 |
| Chymotrypsin | 42 | 29 | 4728 | 1 | 4272 | 3693.5661 | 58.4038 | 74,149 | 3692.558275 | 86 | 801 | 813 | NFSQILPDPSKPS | 0.2000000000000.0 | 801(NFS) | 1 | 1431.7132 | 1428.724939 | 3756061 | 67.81 | 1 | 5 | 5 | 3 | 0 | 0 | 0 | 1 | 3692.559651 | -0.0014 | -0.37 | -9.41 |
| Chymotrypsin | 42 | 29 | 4793 | 1 | 4371 | 3505.459 | 62.3869 | 73,005 | 3504.451175 | 86 | 801 | 813 | NFSQILPDPSKPS | 0.2000000000000.0 | 801(NFS) | 1 | 1431.7132 | 1428.724939 | 3756061 | 67.81 | 1 | 5 | 4 | 1 | 0 | 1 | 0 | 1 | 3504.454791 | -0.0036 | -1.03 | -5.42 |
| Chymotrypsin | 42 | 29 | 4793 | 1 | 4371 | 3505.459 | 62.3869 | 73,005 | 3504.451175 | 86 | 801 | 813 | NFSQILPDPSKPS | 0.2000000000000.0 | 801(NFS) | 1 | 1431.7132 | 1428.724939 | 3756061 | 67.81 | 2 | 6 | 4 | 0 | 1 | 0 | 0 | 1 | 3504.454792 | -0.0036 | -1.03 | -5.42 |
| Chymotrypsin | 42 | 29 | 6166 | 1 | 4278 | 3709.5589 | 58.6649 | 49,662 | 3708.551075 | 86 | 801 | 813 | NFSQILPDPSKPS | 0.2000000000000.0 | 801(NFS) | 1 | 1431.7132 | 1428.724939 | 3756061 | 67.81 | 1 | 6 | 5 | 2 | 0 | 0 | 0 | 1 | 3708.554566 | -0.0035 | -0.94 | -9.15 |
| Chymotrypsin | 42 | 29 | 6318 | 1 | 4297 | 3417.4486 | 59.448 | 47,531 | 3416.440775 | 86 | 801 | 813 | NFSQILPDPSKPS | 0.2000000000000.0 | 801(NFS) | 1 | 1431.7132 | 1428.724939 | 3756061 | 67.81 | 1 | 6 | 5 | 0 | 0 | 0 | 0 | 1 | 3416.438748 | 0.0020 | 0.59 | -8.36 |
| Chymotrypsin | 42 | 29 | 6363 | 1 | 4396 | 3302.3727 | 63.4536 | 47,037 | 3301.364875 | 86 | 801 | 813 | NFSQILPDPSKPS | 0.2000000000000.0 | 801(NFS) | 1 | 1431.7132 | 1428.724939 | 3756061 | 67.81 | 1 | 6 | 3 | 0 | 1 | 0 | 0 | 1 | 3301.375419 | -0.0105 | -3.19 | -4.36 |
| Chymotrypsin | 42 | 29 | 6363 | 1 | 4396 | 3302.3727 | 63.4536 | 47,037 | 3301.364875 | 86 | 801 | 813 | NFSQILPDPSKPS | 0.2000000000000.0 | 801(NFS) | 1 | 1431.7132 | 1428.724939 | 3756061 | 67.81 | 2 | 5 | 3 | 1 | 0 | 1 | 0 | 0 | 3301.375418 | -0.0105 | -3.19 | -4.36 |
| Chymotrypsin | 42 | 29 | 6575 | 1 | 4410 | 3140.3386 | 64.049 | 44,030 | 3139.330775 | 86 | 801 | 813 | NFSQILPDPSKPS | 0.2000000000000.0 | 801(NFS) | 1 | 1431.7132 | 1428.724939 | 3756061 | 67.81 | 1 | 5 | 3 | 0 | 1 | 0 | 0 | 1 | 3139.322595 | 0.0082 | 2.6 | -3.76 |
| Chymotrypsin | 42 | 29 | 6575 | 1 | 4410 | 3140.3386 | 64.049 | 44,030 | 3139.330775 | 86 | 801 | 813 | NFSQILPDPSKPS | 0.2000000000000.0 | 801(NFS) | 1 | 1431.7132 | 1428.724939 | 3756061 | 67.81 | 2 | 4 | 3 | 1 | 0 | 1 | 0 | 1 | 3139.322594 | 0.0082 | 2.6 | -3.76 |
| Chymotrypsin | 42 | 29 | 7698 | 1 | 4208 | 3126.3655 | 55.6734 | 29,297 | 3125.357675 | 86 | 801 | 813 | NFSQILPDPSKPS | 0.2000000000000.0 | 801(NFS) | 1 | 1431.7132 | 1428.724939 | 3756061 | 67.81 | | | | | | | | | | | | |
| Chymotrypsin | 48 | 27 | 166 | 1 | 4097 | 3254.4561 | 50.9928 | 1,898,070 | 3253.448275 | 124 | 801 | 815 | NFSQILPDPSKPSKR | 0.200000000000000.0 | 801(NFS) | 1 | 1715.9092 | 1712.920939 | 2437807.5 | 58.25 | 1 | 7 | 2 | 0 | 0 | 0 | 0 | 1 | 3253.449453 | -0.0012 | -0.36 | -7.26 |
| Chymotrypsin | 48 | 27 | 296 | 1 | 4104 | 3092.4034 | 51.2876 | 1,180,530 | 3091.395575 | 124 | 801 | 815 | NFSQILPDPSKPSKR | 0.200000000000000.0 | 801(NFS) | 1 | 1715.9092 | 1712.920939 | 2437807.5 | 58.25 | 1 | 6 | 2 | 0 | 0 | 0 | 0 | 1 | 3091.396629 | -0.0011 | -0.34 | -6.96 |
| Chymotrypsin | 48 | 27 | 402 | 1 | 4092 | 3441.5396 | 50.779 | 859,723 | 3440.531775 | 124 | 801 | 815 | NFSQILPDPSKPSKR | 0.200000000000000.0 | 801(NFS) | 1 | 1715.9092 | 1712.920939 | 2437807.5 | 58.25 | 1 | 6 | 3 | 1 | 0 | 0 | 0 | 1 | 3440.533911 | -0.0021 | -0.62 | -7.47 |
| Chymotrypsin | 48 | 27 | 604 | 1 | 4103 | 3279.4871 | 51.2464 | 596,486 | 3278.479275 | 124 | 801 | 815 | NFSQILPDPSKPSKR | 0.200000000000000.0 | 801(NFS) | 1 | 1715.9092 | 1712.920939 | 2437807.5 | 58.25 | 1 | 5 | 3 | 1 | 0 | 0 | 0 | 1 | 3278.481087 | -0.0018 | -0.55 | -7.00 |
| Chymotrypsin | 48 | 27 | 739 | 1 | 4111 | 2930.3509 | 51.5799 | 499,473 | 2929.343075 | 124 | 801 | 815 | NFSQILPDPSKPSKR | 0.200000000000000.0 | 801(NFS) | 1 | 1715.9092 | 1712.920939 | 2437807.5 | 58.25 | 1 | 5 | 2 | 0 | 0 | 0 | 0 | 1 | 2929.343805 | -0.0007 | -0.24 | -6.67 |
| Chymotrypsin | 48 | 27 | 985 | 1 | 4097 | 3295.4788 | 50.9928 | 388,518 | 3294.470975 | 124 | 801 | 815 | NFSQILPDPSKPSKR | 0.200000000000000.0 | 801(NFS) | 1 | 1715.9092 | 1712.920939 | 2437807.5 | 58.25 | 1 | 6 | 3 | 0 | 0 | 0 | 0 | 1 | 3294.476002 | -0.0050 | -1.52 | -7.26 |
| Chymotrypsin | 48 | 27 | 1013 | 1 | 4094 | 3416.5091 | 50.8629 | 376,294 | 3415.501275 | 124 | 801 | 815 | NFSQILPDPSKPSKR | 0.200000000000000.0 | 801(NFS) | 1 | 1715.9092 | 1712.920939 | 2437807.5 | 58.25 | 1 | 8 | 2 | 0 | 0 | 0 | 0 | 1 | 3415.502277 | -0.0010 | -0.29 | -7.39 |
| Chymotrypsin | 48 | 27 | 1137 | 1 | 4107 | 3133.4299 | 51.4128 | 335,204 | 3132.422075 | 124 | 801 | 815 | NFSQILPDPSKPSKR | 0.200000000000000.0 | 801(NFS) | 1 | 1715.9092 | 1712.920939 | 2437807.5 | 58.25 | 1 | 5 | 3 | 0 | 0 | 0 | 0 | 1 | 3132.423178 | -0.0011 | -0.35 | -6.84 |
| Chymotrypsin | 48 | 27 | 1312 | 1 | 4087 | 3482.5691 | 50.566 | 291,045 | 3481.561275 | 124 | 801 | 815 | NFSQILPDPSKPSKR | 0.200000000000000.0 | 801(NFS) | 1 | 1715.9092 | 1712.920939 | 2437807.5 | 58.25 | 1 | 5 | 4 | 1 | 0 | 0 | 0 | 1 | 3481.56046 | 0.0008 | 0.23 | -7.68 |
| Chymotrypsin | 48 | 27 | 1488 | 1 | 4174 | 3602.5664 | 54.2318 | 259,351 | 3601.558575 | 124 | 801 | 815 | NFSQILPDPSKPSKR | 0.200000000000000.0 | 801(NFS) | 1 | 1715.9092 | 1712.920939 | 2437807.5 | 58.25 | 1 | 6 | 3 | 0 | 0 | 1 | 0 | 1 | 3601.566333 | -0.0078 | -2.15 | -4.02 |
| Chymotrypsin | 48 | 27 | 1626 | 1 | 4087 | 3628.6273 | 50.566 | 239,353 | 3627.619475 | 124 | 801 | 815 | NFSQILPDPSKPSKR | 0.200000000000000.0 | 801(NFS) | 1 | 1715.9092 | 1712.920939 | 2437807.5 | 58.25 | 1 | 5 | 4 | 2 | 0 | 0 | 0 | 1 | 3627.618369 | 0.0011 | 0.3 | -7.68 |
| Chymotrypsin | 48 | 27 | 1693 | 1 | 4104 | 3336.5034 | 51.2876 | 230,735 | 3335.495575 | 124 | 801 | 815 | NFSQILPDPSKPSKR | 0.200000000000000.0 | 801(NFS) | 1 | 1715.9092 | 1712.920939 | 2437807.5 | 58.25 | 1 | 5 | 4 | 0 | 0 | 0 | 0 | 1 | 3335.502551 | -0.0070 | -2.09 | -6.96 |
| Chymotrypsin | 48 | 27 | 1813 | 1 | 4180 | 3440.5184 | 54.4918 | 215,859 | 3439.510575 | 124 | 801 | 815 | NFSQILPDPSKPSKR | 0.200000000000000.0 | 801(NFS) | 1 | 1715.9092 | 1712.920939 | 2437807.5 | 58.25 | 1 | 5 | 3 | 0 | 0 | 1 | 0 | 1 | 3439.513509 | -0.0029 | -0.85 | -3.76 |
| Chymotrypsin | 48 | 27 | 2495 | 1 | 4171 | 3643.5966 | 54.1076 | 155,479 | 3642.588775 | 124 | 801 | 815 | NFSQILPDPSKPSKR | 0.200000000000000.0 | 801(NFS) | 1 | 1715.9092 | 1712.920939 | 2437807.5 | 58.25 | 1 | 5 | 4 | 0 | 0 | 1 | 0 | 1 | 3642.592882 | -0.0041 | -1.12 | -4.14 |
| Chymotrypsin | 48 | 27 | 3454 | 1 | 4094 | 3685.6469 | 50.8629 | 108,461 | 3684.639075 | 124 | 801 | 815 | NFSQILPDPSKPSKR | 0.200000000000000.0 | 801(NFS) | 1 | 1715.9092 | 1712.920939 | 2437807.5 | 58.25 | 1 | 5 | 5 | 1 | 0 | 0 | 0 | 1 | 3684.639833 | -0.0008 | -0.2 | -7.39 |
| Chymotrypsin | 48 | 27 | 3684 | 1 | 4102 | 3539.5861 | 51.2046 | 100,810 | 3538.578275 | 124 | 801 | 815 | NFSQILPDPSKPSKR | 0.200000000000000.0 | 801(NFS) | 1 | 1715.9092 | 1712.920939 | 2437807.5 | 58.25 | 1 | 5 | 5 | 0 | 0 | 0 | 0 | 1 | 3538.581924 | -0.0036 | -1.03 | -7.04 |
| Chymotrypsin | 48 | 27 | 3842 | 1 | 4102 | 3377.5392 | 51.2046 | 96,306 | 3376.531375 | 124 | 801 | 815 | NFSQILPDPSKPSKR | 0.200000000000000.0 | 801(NFS) | 1 | 1715.9092 | 1712.920939 | 2437807.5 | 58.25 | 1 | 4 | 5 | 0 | 0 | 0 | 0 | 1 | 3376.5291 | 0.0023 | 0.67 | -7.04 |
| Chymotrypsin | 48 | 27 | 5395 | 1 | 4161 | 3789.6664 | 53.6825 | 61,855 | 3788.658575 | 124 | 801 | 815 | NFSQILPDPSKPSKR | 0.200000000000000.0 | 801(NFS) | 1 | 1715.9092 | 1712.920939 | 2437807.5 | 58.25 | 1 | 6 | 4 | 0 | 1 | 0 | 0 | 1 | 3788.650792 | 0.0078 | 2.05 | -4.57 |
| Chymotrypsin | 48 | 27 | 5395 | 1 | 4161 | 3789.6664 | 53.6825 | 61,855 | 3788.658575 | 124 | 801 | 815 | NFSQILPDPSKPSKR | 0.200000000000000.0 | 801(NFS) | 1 | 1715.9092 | 1712.920939 | 2437807.5 | 58.25 | 2 | 5 | 4 | 1 | 0 | 1 | 0 | 1 | 3788.650791 | 0.0078 | 2.05 | -4.57 |
| Chymotrypsin | 48 | 27 | 6088 | 1 | 4184 | 3424.5251 | 54.6598 | 50,815 | 3423.517275 | 124 | 801 | 815 | NFSQILPDPSKPSKR | 0.200000000000000.0 | 801(NFS) | 1 | 1715.9092 | 1712.920939 | 2437807.5 | 58.25 | 1 | 4 | 3 | 1 | 0 | 1 | 0 | 1 | 3423.518594 | -0.0013 | -0.38 | -3.59 |
| Chymotrypsin | 48 | 27 | 6088 | 1 | 4184 | 3424.5251 | 54.6598 | 50,815 | 3423.517275 | 124 | 801 | 815 | NFSQILPDPSKPSKR | 0.200000000000000.0 | 801(NFS) | 1 | 1715.9092 | 1712.920939 | 2437807.5 | 58.25 | 2 | 5 | 3 | 0 | 1 | 0 | 0 | 1 | 3423.518595 | -0.0013 | -0.38 | -3.59 |
| Chymotrypsin | 48 | 27 | 6292 | 1 | 4087 | 4050.768 | 50.566 | 47,912 | 4049.760175 | 124 | 801 | 815 | NFSQILPDPSKPSKR | 0.200000000000000.0 | 801(NFS) | 1 | 1715.9092 | 1712.920939 | 2437807.5 | 58.25 | 1 | 6 | 6 | 1 | 0 | 0 | 0 | 1 | 4049.77203 | -0.0119 | -2.92 | -7.68 |

| | | | | | | | | | | | | | | | | | | | | | | | | | | | | | | | | |
|---|---|---|---|---|---|---|---|---|---|---|---|---|---|---|---|---|---|---|---|---|---|---|---|---|---|---|---|---|---|---|---|---|
| Chymotrypsin | 48 | 27 | 6678 | 1 | 4085 | 3847.6962 | 50.4817 | 42,746 | 3846.688375 | 124 | 801 | 815 | NFSQILPDPSKPSKR | 0.200000000000000.0 | 801(NFS) | 1 | 1715.9092 | 1712.920939 | 2437807.5 | 58.25 | 1 | 6 | 5 | 1 | 0 | 0 | 0 | 1 | 3846.692657 | -0.0043 | -1.11 | -7.77 |
| Chymotrypsin | 48 | 27 | 6760 | 1 | 4181 | 3586.5853 | 54.5331 | 41,567 | 3585.577475 | 124 | 801 | 815 | NFSQILPDPSKPSKR | 0.200000000000000.0 | 801(NFS) | 1 | 1715.9092 | 1712.920939 | 2437807.5 | 58.25 | 1 | 6 | 3 | 0 | 1 | 0 | 0 | 1 | 3585.571419 | 0.0061 | 1.68 | -3.72 |
| Chymotrypsin | 48 | 27 | 6760 | 1 | 4181 | 3586.5853 | 54.5331 | 41,567 | 3585.577475 | 124 | 801 | 815 | NFSQILPDPSKPSKR | 0.200000000000000.0 | 801(NFS) | 1 | 1715.9092 | 1712.920939 | 2437807.5 | 58.25 | 2 | 5 | 3 | 1 | 0 | 1 | 0 | 0 | 3585.571418 | 0.0061 | 1.68 | -3.72 |
| Chymotrypsin | 48 | 27 | 6896 | 1 | 4093 | 3701.6383 | 50.8215 | 39,777 | 3700.630475 | 124 | 801 | 815 | NFSQILPDPSKPSKR | 0.200000000000000.0 | 801(NFS) | 1 | 1715.9092 | 1712.920939 | 2437807.5 | 58.25 | 1 | 6 | 5 | 0 | 0 | 0 | 0 | 1 | 3700.634748 | -0.0043 | -1.15 | -7.43 |
| Chymotrypsin | 48 | 27 | 7293 | 1 | 4097 | 3498.5592 | 50.9928 | 34,236 | 3497.551375 | 124 | 801 | 815 | NFSQILPDPSKPSKR | 0.200000000000000.0 | 801(NFS) | 1 | 1715.9092 | 1712.920939 | 2437807.5 | 58.25 | 1 | 6 | 4 | 0 | 0 | 0 | 0 | 1 | 3497.555375 | -0.0040 | -1.14 | -7.26 |
| Chymotrypsin | 48 | 27 | 7765 | 1 | 4098 | 3904.7123 | 51.0346 | 28,043 | 3903.704475 | 124 | 801 | 815 | NFSQILPDPSKPSKR | 0.200000000000000.0 | 801(NFS) | 1 | 1715.9092 | 1712.920939 | 2437807.5 | 58.25 | 1 | 6 | 6 | 0 | 0 | 0 | 0 | 1 | 3903.714121 | -0.0096 | -2.47 | -7.21 |
| Chymotrypsin | 48 | 27 | 8125 | 1 | 4107 | 2768.2987 | 51.4128 | 23,766 | 2767.290875 | 124 | 801 | 815 | NFSQILPDPSKPSKR | 0.200000000000000.0 | 801(NFS) | 1 | 1715.9092 | 1712.920939 | 2437807.5 | 58.25 | 1 | 4 | 2 | 0 | 0 | 0 | 0 | 1 | 2767.290981 | -0.0001 | -0.03 | -6.84 |
| Chymotrypsin | 48 | 27 | 8437 | 1 | 4088 | 3523.5946 | 50.6098 | 19,475 | 3522.586775 | 124 | 801 | 815 | NFSQILPDPSKPSKR | 0.200000000000000.0 | 801(NFS) | 1 | 1715.9092 | 1712.920939 | 2437807.5 | 58.25 | 1 | 4 | 5 | 1 | 0 | 0 | 0 | 1 | 3522.587009 | -0.0002 | -0.06 | -7.64 |
| Chymotrypsin | 117 | 30 | 506 | 1 | 4728 | 3231.3728 | 77.087 | 705,499 | 3230.364975 | 32 | 798 | 813 | GGFNFSQILPDPSKPS | 0.0002000000000000.0 | 801(NFS) | 1 | 1692.8245 | 1689.836239 | 13656227 | 81.08 | 1 | 7 | 2 | 0 | 0 | 0 | 0 | 1 | 3230.364753 | 0.0002 | 0.06 | -4.00 |
| Chymotrypsin | 117 | 30 | 594 | 1 | 4736 | 3069.315 | 77.4279 | 606,163 | 3068.307175 | 32 | 798 | 813 | GGFNFSQILPDPSKPS | 0.0002000000000000.0 | 801(NFS) | 1 | 1692.8245 | 1689.836239 | 13656227 | 81.08 | 1 | 6 | 2 | 0 | 0 | 0 | 0 | 1 | 3068.311929 | -0.0048 | -1.54 | -3.66 |
| Chymotrypsin | 117 | 30 | 1234 | 1 | 4728 | 3418.4579 | 77.087 | 309,818 | 3417.450075 | 32 | 798 | 813 | GGFNFSQILPDPSKPS | 0.0002000000000000.0 | 801(NFS) | 1 | 1692.8245 | 1689.836239 | 13656227 | 81.08 | 1 | 6 | 3 | 1 | 0 | 0 | 0 | 1 | 3417.449211 | 0.0009 | 0.25 | -4.00 |
| Chymotrypsin | 117 | 30 | 1783 | 1 | 4737 | 2907.2703 | 77.4695 | 218,871 | 2906.262475 | 32 | 798 | 813 | GGFNFSQILPDPSKPS | 0.0002000000000000.0 | 801(NFS) | 1 | 1692.8245 | 1689.836239 | 13656227 | 81.08 | 1 | 5 | 2 | 0 | 0 | 0 | 0 | 1 | 2906.259105 | 0.0034 | 1.15 | -3.61 |
| Chymotrypsin | 117 | 30 | 2425 | 1 | 4732 | 3272.3941 | 77.2564 | 160,040 | 3271.386275 | 32 | 798 | 813 | GGFNFSQILPDPSKPS | 0.0002000000000000.0 | 801(NFS) | 1 | 1692.8245 | 1689.836239 | 13656227 | 81.08 | 1 | 6 | 3 | 0 | 0 | 0 | 0 | 1 | 3271.391302 | -0.0050 | -1.53 | -3.83 |
| Chymotrypsin | 117 | 30 | 2436 | 1 | 4727 | 3393.4237 | 77.0431 | 159,296 | 3392.415875 | 32 | 798 | 813 | GGFNFSQILPDPSKPS | 0.0002000000000000.0 | 801(NFS) | 1 | 1692.8245 | 1689.836239 | 13656227 | 81.08 | 1 | 8 | 2 | 0 | 0 | 0 | 0 | 1 | 3392.417577 | -0.0017 | -0.5 | -4.04 |
| Chymotrypsin | 117 | 30 | 2464 | 1 | 4734 | 3256.4127 | 77.3423 | 157,482 | 3255.404875 | 32 | 798 | 813 | GGFNFSQILPDPSKPS | 0.0002000000000000.0 | 801(NFS) | 1 | 1692.8245 | 1689.836239 | 13656227 | 81.08 | 1 | 5 | 3 | 1 | 0 | 0 | 0 | 1 | 3255.396387 | 0.0085 | 2.6 | -3.74 |
| Chymotrypsin | 117 | 30 | 2636 | 1 | 4829 | 3579.4848 | 81.2824 | 147,126 | 3578.476975 | 32 | 798 | 813 | GGFNFSQILPDPSKPS | 0.0002000000000000.0 | 801(NFS) | 1 | 1692.8245 | 1689.836239 | 13656227 | 81.08 | 1 | 6 | 3 | 0 | 0 | 1 | 0 | 1 | 3578.481633 | -0.0047 | -1.3 | 0.20 |
| Chymotrypsin | 117 | 30 | 2644 | 1 | 4693 | 3605.5415 | 75.6548 | 146,462 | 3604.533675 | 32 | 798 | 813 | GGFNFSQILPDPSKPS | 0.0002000000000000.0 | 801(NFS) | 1 | 1692.8245 | 1689.836239 | 13656227 | 81.08 | 1 | 5 | 4 | 2 | 0 | 0 | 0 | 1 | 3604.533669 | 0.0000 | 0 | -5.43 |
| Chymotrypsin | 117 | 30 | 2873 | 1 | 4819 | 3620.5103 | 80.8724 | 132,917 | 3619.502475 | 32 | 798 | 813 | GGFNFSQILPDPSKPS | 0.0002000000000000.0 | 801(NFS) | 1 | 1692.8245 | 1689.836239 | 13656227 | 81.08 | 1 | 5 | 4 | 0 | 0 | 1 | 0 | 1 | 3619.508182 | -0.0057 | -1.57 | -0.21 |
| Chymotrypsin | 117 | 30 | 2938 | 1 | 4835 | 3417.4361 | 81.5088 | 130,611 | 3416.428275 | 32 | 798 | 813 | GGFNFSQILPDPSKPS | 0.0002000000000000.0 | 801(NFS) | 1 | 1692.8245 | 1689.836239 | 13656227 | 81.08 | 1 | 5 | 3 | 0 | 0 | 1 | 0 | 1 | 3416.428809 | -0.0005 | -0.15 | 0.42 |
| Chymotrypsin | 117 | 30 | 3680 | 1 | 4727 | 3313.4212 | 77.0431 | 100,925 | 3312.413375 | 32 | 798 | 813 | GGFNFSQILPDPSKPS | 0.0002000000000000.0 | 801(NFS) | 1 | 1692.8245 | 1689.836239 | 13656227 | 81.08 | 1 | 5 | 4 | 0 | 0 | 0 | 0 | 1 | 3312.417851 | -0.0045 | -1.35 | -4.04 |
| Chymotrypsin | 117 | 30 | 3809 | 1 | 4721 | 3459.4826 | 76.7836 | 97,114 | 3458.474775 | 32 | 798 | 813 | GGFNFSQILPDPSKPS | 0.0002000000000000.0 | 801(NFS) | 1 | 1692.8245 | 1689.836239 | 13656227 | 81.08 | 1 | 5 | 4 | 1 | 0 | 0 | 0 | 1 | 3458.47576 | -0.0010 | -0.28 | -4.30 |
| Chymotrypsin | 117 | 30 | 4072 | 1 | 4691 | 3808.6212 | 75.5866 | 89,892 | 3807.613375 | 32 | 798 | 813 | GGFNFSQILPDPSKPS | 0.0002000000000000.0 | 801(NFS) | 1 | 1692.8245 | 1689.836239 | 13656227 | 81.08 | 1 | 5 | 5 | 2 | 0 | 0 | 0 | 1 | 3807.613042 | 0.0003 | 0.08 | -5.50 |
| Chymotrypsin | 117 | 30 | 4252 | 1 | 4794 | 3766.5791 | 79.8522 | 85,101 | 3765.571275 | 32 | 798 | 813 | GGFNFSQILPDPSKPS | 0.0002000000000000.0 | 801(NFS) | 1 | 1692.8245 | 1689.836239 | 13656227 | 81.08 | 1 | 6 | 4 | 0 | 1 | 0 | 0 | 1 | 3765.566092 | 0.0052 | 1.37 | -1.23 |
| Chymotrypsin | 117 | 30 | 4252 | 1 | 4794 | 3766.5791 | 79.8522 | 85,101 | 3765.571275 | 32 | 798 | 813 | GGFNFSQILPDPSKPS | 0.0002000000000000.0 | 801(NFS) | 1 | 1692.8245 | 1689.836239 | 13656227 | 81.08 | 2 | 5 | 4 | 1 | 0 | 1 | 0 | 1 | 3765.566091 | 0.0052 | 1.37 | -1.23 |
| Chymotrypsin | 117 | 30 | 4443 | 1 | 4703 | 3662.5506 | 76.016 | 80,396 | 3661.542775 | 32 | 798 | 813 | GGFNFSQILPDPSKPS | 0.0002000000000000.0 | 801(NFS) | 1 | 1692.8245 | 1689.836239 | 13656227 | 81.08 | 1 | 5 | 5 | 1 | 0 | 0 | 0 | 1 | 3661.555133 | -0.0124 | -3.37 | -5.07 |
| Chymotrypsin | 117 | 30 | 4480 | 1 | 4702 | 3459.4885 | 75.9814 | 79,521 | 3458.480675 | 32 | 798 | 813 | GGFNFSQILPDPSKPS | 0.0002000000000000.0 | 801(NFS) | 1 | 1692.8245 | 1689.836239 | 13656227 | 81.08 | 1 | 5 | 4 | 1 | 0 | 0 | 0 | 1 | 3458.47576 | 0.0049 | 1.42 | -5.10 |
| Chymotrypsin | 117 | 30 | 4652 | 1 | 4741 | 3110.3486 | 77.6424 | 75,978 | 3109.340775 | 32 | 798 | 813 | GGFNFSQILPDPSKPS | 0.0002000000000000.0 | 801(NFS) | 1 | 1692.8245 | 1689.836239 | 13656227 | 81.08 | 1 | 5 | 3 | 0 | 0 | 0 | 0 | 1 | 3109.338478 | 0.0023 | 0.73 | -3.44 |
| Chymotrypsin | 117 | 30 | 4656 | 1 | 4831 | 3807.613 | 81.3473 | 75,914 | 3806.605175 | 32 | 798 | 813 | GGFNFSQILPDPSKPS | 0.0002000000000000.0 | 801(NFS) | 1 | 1692.8245 | 1689.836239 | 13656227 | 81.08 | 1 | 5 | 5 | 0 | 1 | 0 | 0 | 1 | 3806.592641 | 0.0125 | 3.29 | 0.26 |
| Chymotrypsin | 117 | 30 | 4656 | 1 | 4831 | 3807.613 | 81.3473 | 75,914 | 3806.605175 | 32 | 798 | 813 | GGFNFSQILPDPSKPS | 0.0002000000000000.0 | 801(NFS) | 1 | 1692.8245 | 1689.836239 | 13656227 | 81.08 | 2 | 4 | 5 | 1 | 0 | 1 | 0 | 1 | 3806.59264 | 0.0125 | 3.29 | 0.26 |
| Chymotrypsin | 117 | 30 | 4766 | 1 | 4864 | 4203.7514 | 82.6922 | 73,488 | 4202.743575 | 32 | 798 | 813 | GGFNFSQILPDPSKPS | 0.0002000000000000.0 | 801(NFS) | 1 | 1692.8245 | 1689.836239 | 13656227 | 81.08 | | | | | | | | | | | | |
| Chymotrypsin | 117 | 30 | 4783 | 1 | 4689 | 3751.5985 | 75.5015 | 73,205 | 3750.590675 | 32 | 798 | 813 | GGFNFSQILPDPSKPS | 0.0002000000000000.0 | 801(NFS) | 1 | 1692.8245 | 1689.836239 | 13656227 | 81.08 | 1 | 5 | 4 | 3 | 0 | 0 | 0 | 1 | 3750.591578 | -0.0009 | -0.24 | -5.58 |
| Chymotrypsin | 117 | 30 | 4823 | 1 | 4724 | 3516.5042 | 76.9143 | 72,309 | 3515.496375 | 32 | 798 | 813 | GGFNFSQILPDPSKPS | 0.0002000000000000.0 | 801(NFS) | 1 | 1692.8245 | 1689.836239 | 13656227 | 81.08 | 1 | 5 | 5 | 0 | 0 | 0 | 0 | 1 | 3515.497224 | -0.0008 | -0.24 | -4.17 |
| Chymotrypsin | 117 | 30 | 5086 | 1 | 4717 | 3605.5356 | 76.6131 | 66,898 | 3604.527775 | 32 | 798 | 813 | GGFNFSQILPDPSKPS | 0.0002000000000000.0 | 801(NFS) | 1 | 1692.8245 | 1689.836239 | 13656227 | 81.08 | 1 | 5 | 4 | 2 | 0 | 0 | 0 | 1 | 3604.533669 | -0.0059 | -1.63 | -4.47 |
| Chymotrypsin | 117 | 30 | 5294 | 1 | 4722 | 3662.5587 | 76.8272 | 63,747 | 3661.550875 | 32 | 798 | 813 | GGFNFSQILPDPSKPS | 0.0002000000000000.0 | 801(NFS) | 1 | 1692.8245 | 1689.836239 | 13656227 | 81.08 | 1 | 5 | 5 | 1 | 0 | 0 | 0 | 1 | 3661.555133 | -0.0043 | -1.16 | -4.26 |
| Chymotrypsin | 117 | 30 | 5498 | 1 | 4730 | 3354.4526 | 77.1712 | 60,164 | 3353.444775 | 32 | 798 | 813 | GGFNFSQILPDPSKPS | 0.0002000000000000.0 | 801(NFS) | 1 | 1692.8245 | 1689.836239 | 13656227 | 81.08 | 1 | 4 | 5 | 0 | 0 | 0 | 0 | 1 | 3353.4444 | 0.0004 | 0.11 | -3.91 |
| Chymotrypsin | 117 | 30 | 5795 | 1 | 4827 | 3563.4898 | 81.2032 | 55,176 | 3562.481975 | 32 | 798 | 813 | GGFNFSQILPDPSKPS | 0.0002000000000000.0 | 801(NFS) | 1 | 1692.8245 | 1689.836239 | 13656227 | 81.08 | 1 | 6 | 3 | 0 | 1 | 0 | 0 | 1 | 3562.486719 | -0.0047 | -1.33 | 0.12 |
| Chymotrypsin | 117 | 30 | 5795 | 1 | 4827 | 3563.4898 | 81.2032 | 55,176 | 3562.481975 | 32 | 798 | 813 | GGFNFSQILPDPSKPS | 0.0002000000000000.0 | 801(NFS) | 1 | 1692.8245 | 1689.836239 | 13656227 | 81.08 | 2 | 5 | 3 | 1 | 0 | 1 | 0 | 0 | 3562.486718 | -0.0047 | -1.33 | 0.12 |
| Chymotrypsin | 117 | 30 | 6269 | 1 | 4686 | 3954.6815 | 75.3721 | 48,238 | 3953.673675 | 32 | 798 | 813 | GGFNFSQILPDPSKPS | 0.0002000000000000.0 | 801(NFS) | 1 | 1692.8245 | 1689.836239 | 13656227 | 81.08 | 1 | 5 | 5 | 3 | 0 | 0 | 0 | 1 | 3953.670951 | 0.0027 | 0.68 | -5.71 |
| Chymotrypsin | 117 | 30 | 6642 | 1 | 4853 | 4568.8912 | 82.252 | 43,165 | 4567.883375 | 32 | 798 | 813 | GGFNFSQILPDPSKPS | 0.0002000000000000.0 | 801(NFS) | 1 | 1692.8245 | 1689.836239 | 13656227 | 81.08 | 1 | 0 | 7 | 2 | 4 | 0 | 1 | -1 | 4567.889336 | -0.0060 | -1.3 | 1.17 |
| Chymotrypsin | 117 | 30 | 6658 | 1 | 4715 | 3808.6179 | 76.5274 | 42,947 | 3807.610075 | 32 | 798 | 813 | GGFNFSQILPDPSKPS | 0.0002000000000000.0 | 801(NFS) | 1 | 1692.8245 | 1689.836239 | 13656227 | 81.08 | 1 | 5 | 5 | 2 | 0 | 0 | 0 | 1 | 3807.613042 | -0.0030 | -0.77 | -4.56 |
| Chymotrypsin | 117 | 30 | 6918 | 1 | 4876 | 4057.6816 | 83.1962 | 39,373 | 4056.673775 | 32 | 798 | 813 | GGFNFSQILPDPSKPS | 0.0002000000000000.0 | 801(NFS) | 1 | 1692.8245 | 1689.836239 | 13656227 | 81.08 | 1 | 5 | 4 | 1 | 1 | 1 | 0 | 1 | 4056.661508 | 0.0123 | 3.02 | 2.11 |
| Chymotrypsin | 117 | 30 | 6918 | 1 | 4876 | 4057.6816 | 83.1962 | 39,373 | 4056.673775 | 32 | 798 | 813 | GGFNFSQILPDPSKPS | 0.0002000000000000.0 | 801(NFS) | 1 | 1692.8245 | 1689.836239 | 13656227 | 81.08 | 2 | 6 | 4 | 0 | 2 | 0 | 0 | 0 | 4056.661509 | 0.0123 | 3.02 | 2.11 |
| Chymotrypsin | 117 | 30 | 6918 | 1 | 4876 | 4057.6816 | 83.1962 | 39,373 | 4056.673775 | 32 | 798 | 813 | GGFNFSQILPDPSKPS | 0.0002000000000000.0 | 801(NFS) | 1 | 1692.8245 | 1689.836239 | 13656227 | 81.08 | 3 | 4 | 4 | 2 | 0 | 2 | 0 | 0 | 4056.661507 | 0.0123 | 3.02 | 2.11 |
| Trypsin | 7 | 59 | 14 | 1 | 4846 | 3309.3804 | 82.9755 | 6,614,410 | 3308.372575 | 3 | 796 | 811 | DFGGFNFSQILPDPSK | 0.0000200000000000.0 | 801(NFS) | 1 | 1770.8351 | 1767.846839 | 102420190 | 87.03 | 1 | 7 | 2 | 0 | 0 | 0 | 0 | 1 | 3308.375353 | -0.0028 | -0.83 | -4.05 |
| Trypsin | 7 | 59 | 19 | 1 | 4850 | 3147.3283 | 83.1682 | 5,814,610 | 3146.320475 | 3 | 796 | 811 | DFGGFNFSQILPDPSK | 0.0000200000000000.0 | 801(NFS) | 1 | 1770.8351 | 1767.846839 | 102420190 | 87.03 | 1 | 6 | 2 | 0 | 0 | 0 | 0 | 1 | 3146.322529 | -0.0021 | -0.65 | -3.86 |
| Trypsin | 7 | 59 | 42 | 1 | 4853 | 2985.2745 | 83.325 | 2,977,020 | 2984.266675 | 3 | 796 | 811 | DFGGFNFSQILPDPSK | 0.0000200000000000.0 | 801(NFS) | 1 | 1770.8351 | 1767.846839 | 102420190 | 87.03 | 1 | 5 | 2 | 0 | 0 | 0 | 0 | 1 | 2984.269705 | -0.0030 | -1.01 | -3.70 |
| Trypsin | 7 | 59 | 63 | 1 | 4844 | 3496.4646 | 82.877 | 1,953,880 | 3495.456775 | 3 | 796 | 811 | DFGGFNFSQILPDPSK | 0.0000200000000000.0 | 801(NFS) | 1 | 1770.8351 | 1767.846839 | 102420190 | 87.03 | 1 | 6 | 3 | 1 | 0 | 0 | 0 | 1 | 3495.459811 | -0.0030 | -0.86 | -4.15 |
| Trypsin | 7 | 59 | 71 | 1 | 4849 | 3334.4149 | 83.1192 | 1,691,660 | 3333.407075 | 3 | 796 | 811 | DFGGFNFSQILPDPSK | 0.0000200000000000.0 | 801(NFS) | 1 | 1770.8351 | 1767.846839 | 102420190 | 87.03 | 1 | 5 | 3 | 1 | 0 | 0 | 0 | 1 | 3333.406987 | 0.0001 | 0.02 | -3.91 |
| Trypsin | 7 | 59 | 99 | 1 | 4843 | 3471.4347 | 82.8269 | 1,228,520 | 3470.426875 | 3 | 796 | 811 | DFGGFNFSQILPDPSK | 0.0000200000000000.0 | 801(NFS) | 1 | 1770.8351 | 1767.846839 | 102420190 | 87.03 | 1 | 8 | 2 | 0 | 0 | 0 | 0 | 1 | 3470.428177 | -0.0013 | -0.37 | -4.20 |
| Trypsin | 7 | 59 | 103 | 1 | 4847 | 3350.4081 | 83.0222 | 1,202,570 | 3349.400275 | 3 | 796 | 811 | DFGGFNFSQILPDPSK | 0.0000200000000000.0 | 801(NFS) | 1 | 1770.8351 | 1767.846839 | 102420190 | 87.03 | 1 | 6 | 3 | 0 | 0 | 0 | 0 | 1 | 3349.401902 | -0.0016 | -0.48 | -4.01 |
| Trypsin | 7 | 59 | 117 | 1 | 4853 | 3188.3535 | 83.325 | 1,045,410 | 3187.345675 | 3 | 796 | 811 | DFGGFNFSQILPDPSK | 0.0000200000000000.0 | 801(NFS) | 1 | 1770.8351 | 1767.846839 | 102420190 | 87.03 | 1 | 5 | 3 | 0 | 0 | 0 | 0 | 1 | 3187.349078 | -0.0034 | -1.06 | -3.70 |
| Trypsin | 7 | 59 | 181 | 1 | 4820 | 3683.5567 | 81.6545 | 696,043 | 3682.548875 | 3 | 796 | 811 | DFGGFNFSQILPDPSK | 0.0000200000000000.0 | 801(NFS) | 1 | 1770.8351 | 1767.846839 | 102420190 | 87.03 | 1 | 5 | 4 | 2 | 0 | 0 | 0 | 1 | 3682.544269 | 0.0046 | 1.25 | -5.37 |
| Trypsin | 7 | 59 | 201 | 1 | 4826 | 3578.5205 | 81.9701 | 634,772 | 3577.512675 | 3 | 796 | 811 | DFGGFNFSQILPDPSK | 0.0000200000000000.0 | 801(NFS) | 1 | 1770.8351 | 1767.846839 | 102420190 | 87.03 | 1 | 4 | 5 | 1 | 0 | 0 | 0 | 1 | 3577.512909 | -0.0002 | -0.06 | -5.06 |
| Trypsin | 7 | 59 | 227 | 1 | 4839 | 3537.4939 | 82.6166 | 581,456 | 3536.486075 | 3 | 796 | 811 | DFGGFNFSQILPDPSK | 0.0000200000000000.0 | 801(NFS) | 1 | 1770.8351 | 1767.846839 | 102420190 | 87.03 | 1 | 5 | 4 | 1 | 0 | 0 | 0 | 1 | 3536.48636 | -0.0003 | -0.08 | -4.41 |
| Trypsin | 7 | 59 | 246 | 1 | 4844 | 3391.4255 | 82.877 | 552,593 | 3390.417675 | 3 | 796 | 811 | DFGGFNFSQILPDPSK | 0.0000200000000000.0 | 801(NFS) | 1 | 1770.8351 | 1767.846839 | 102420190 | 87.03 | 1 | 5 | 4 | 0 | 0 | 0 | 0 | 1 | 3390.428451 | -0.0108 | -3.17 | -4.15 |
| Trypsin | 7 | 59 | 276 | 1 | 4822 | 3740.5748 | 81.756 | 486,319 | 3739.566975 | 3 | 796 | 811 | DFGGFNFSQILPDPSK | 0.0000200000000000.0 | 801(NFS) | 1 | 1770.8351 | 1767.846839 | 102420190 | 87.03 | 1 | 5 | 5 | 1 | 0 | 0 | 0 | 1 | 3739.565733 | 0.0012 | 0.33 | -5.27 |
| Trypsin | 7 | 59 | 370 | 1 | 4843 | 3594.5138 | 82.8269 | 390,074 | 3593.505975 | 3 | 796 | 811 | DFGGFNFSQILPDPSK | 0.0000200000000000.0 | 801(NFS) | 1 | 1770.8351 | 1767.846839 | 102420190 | 87.03 | 1 | 5 | 5 | 0 | 0 | 0 | 0 | 1 | 3593.507824 | -0.0018 | -0.51 | -4.20 |
| Trypsin | 7 | 59 | 393 | 1 | 4847 | 3432.4606 | 83.0222 | 368,589 | 3431.452775 | 3 | 796 | 811 | DFGGFNFSQILPDPSK | 0.0000200000000000.0 | 801(NFS) | 1 | 1770.8351 | 1767.846839 | 102420190 | 87.03 | 1 | 4 | 5 | 0 | 0 | 0 | 0 | 1 | 3431.455 | -0.0022 | -0.64 | -4.01 |
| Trypsin | 7 | 59 | 478 | 1 | 4814 | 3829.6106 | 81.3531 | 303,649 | 3828.602775 | 3 | 796 | 811 | DFGGFNFSQILPDPSK | 0.0000200000000000.0 | 801(NFS) | 1 | 1770.8351 | 1767.846839 | 102420190 | 87.03 | 1 | 5 | 4 | 3 | 0 | 0 | 0 | 1 | 3828.602178 | 0.0006 | 0.15 | -5.67 |
| Trypsin | 7 | 59 | 513 | 1 | 4819 | 3886.6318 | 81.6003 | 279,637 | 3885.623975 | 3 | 796 | 811 | DFGGFNFSQILPDPSK | 0.0000200000000000.0 | 801(NFS) | 1 | 1770.8351 | 1767.846839 | 102420190 | 87.03 | 1 | 5 | 5 | 2 | 0 | 0 | 0 | 1 | 3885.623642 | 0.0003 | 0.08 | -5.43 |
| Trypsin | 7 | 59 | 632 | 1 | 4938 | 3657.4955 | 87.3503 | 227,434 | 3656.487675 | 3 | 796 | 811 | DFGGFNFSQILPDPSK | 0.0000200000000000.0 | 801(NFS) | 1 | 1770.8351 | 1767.846839 | 102420190 | 87.03 | 1 | 6 | 3 | 0 | 0 | 1 | 0 | 1 | 3656.492233 | -0.0046 | -1.24 | 0.32 |
| Trypsin | 7 | 59 | 643 | 1 | 4834 | 3416.4665 | 82.3529 | 218,012 | 3415.458675 | 3 | 796 | 811 | DFGGFNFSQILPDPSK | 0.0000200000000000.0 | 801(NFS) | 1 | 1770.8351 | 1767.846839 | 102420190 | 87.03 | 1 | 3 | 5 | 1 | 0 | 0 | 0 | 1 | 3415.460085 | -0.0014 | -0.41 | -4.67 |
| Trypsin | 7 | 59 | 671 | 1 | 4814 | 4032.6877 | 81.3531 | 210,794 | 4031.679875 | 3 | 796 | 811 | DFGGFNFSQILPDPSK | 0.0000200000000000.0 | 801(NFS) | 1 | 1770.8351 | 1767.846839 | 102420190 | 87.03 | 1 | 5 | 5 | 3 | 0 | 0 | 0 | 1 | 4031.681551 | -0.0017 | -0.41 | -5.67 |
| Trypsin | 7 | 59 | 743 | 1 | 4943 | 3495.4453 | 87.6084 | 187,990 | 3494.437475 | 3 | 796 | 811 | DFGGFNFSQILPDPSK | 0.0000200000000000.0 | 801(NFS) | 1 | 1770.8351 | 1767.846839 | 102420190 | 87.03 | 1 | 5 | 3 | 0 | 0 | 1 | 0 | 1 | 3494.439409 | -0.0019 | -0.55 | 0.58 |
| Trypsin | 7 | 59 | 765 | 1 | 4845 | 3553.4916 | 82.9268 | 183,429 | 3552.483775 | 3 | 796 | 811 | DFGGFNFSQILPDPSK | 0.0000200000000000.0 | 801(NFS) | 1 | 1770.8351 | 1767.846839 | 102420190 | 87.03 | 1 | 6 | 4 | 0 | 0 | 0 | 0 | 1 | 3552.481275 | 0.0025 | 0.7 | -4.10 |
| Trypsin | 7 | 59 | 778 | 1 | 4910 | 3844.5816 | 86.0306 | 179,792 | 3843.573775 | 3 | 796 | 811 | DFGGFNFSQILPDPSK | 0.0000200000000000.0 | 801(NFS) | 1 | 1770.8351 | 1767.846839 | 102420190 | 87.03 | 1 | 5 | 4 | 1 | 0 | 1 | 0 | 1 | 3843.576691 | -0.0029 | -0.75 | -1.00 |
| Trypsin | 7 | 59 | 778 | 1 | 4910 | 3844.5816 | 86.0306 | 179,792 | 3843.573775 | 3 | 796 | 811 | DFGGFNFSQILPDPSK | 0.0000200000000000.0 | 801(NFS) | 1 | 1770.8351 | 1767.846839 | 102420190 | 87.03 | 2 | 6 | 4 | 0 | 1 | 0 | 0 | 1 | 3843.576692 | -0.0029 | -0.75 | -1.00 |
| Trypsin | 7 | 59 | 965 | 1 | 4815 | 3902.6242 | 81.4074 | 146,433 | 3901.616375 | 3 | 796 | 811 | DFGGFNFSQILPDPSK | 0.0000200000000000.0 | 801(NFS) | 1 | 1770.8351 | 1767.846839 | 102420190 | 87.03 | 1 | 6 | 5 | 1 | 0 | 0 | 0 | 1 | 3901.618557 | -0.0022 | -0.55 | -5.62 |
| Trypsin | 7 | 59 | 969 | 1 | 4851 | 3229.3715 | 83.2179 | 145,304 | 3228.363675 | 3 | 796 | 811 | DFGGFNFSQILPDPSK | 0.0000200000000000.0 | 801(NFS) | 1 | 1770.8351 | 1767.846839 | 102420190 | 87.03 | 1 | 4 | 4 | 0 | 0 | 0 | 0 | 1 | 3228.375627 | -0.0120 | -3.7 | -3.81 |
| Trypsin | 7 | 59 | 997 | 1 | 4836 | 3756.5606 | 82.4588 | 140,612 | 3755.552775 | 3 | 796 | 811 | DFGGFNFSQILPDPSK | 0.0000200000000000.0 | 801(NFS) | 1 | 1770.8351 | 1767.846839 | 102420190 | 87.03 | 1 | 6 | 5 | 0 | 0 | 0 | 0 | 1 | 3755.560648 | -0.0079 | -2.09 | -4.57 |
| Trypsin | 7 | 59 | 1085 | 1 | 4837 | 3213.3844 | 82.5037 | 126,691 | 3212.376575 | 3 | 796 | 811 | DFGGFNFSQILPDPSK | 0.0000200000000000.0 | 801(NFS) | 1 | 1770.8351 | 1767.846839 | 102420190 | 87.03 | 1 | 3 | 4 | 1 | 0 | 0 | 0 | 1 | 3212.380712 | -0.0041 | -1.28 | -4.52 |
| Trypsin | 7 | 59 | 1118 | 1 | 4826 | 3480.4731 | 81.9701 | 123,790 | 3479.465275 | 3 | 796 | 811 | DFGGFNFSQILPDPSK | 0.0000200000000000.0 | 801(NFS) | 1 | 1770.8351 | 1767.846839 | 102420190 | 87.03 | 1 | 5 | 3 | 2 | 0 | 0 | 0 | 0 | 3479.464896 | 0.0004 | 0.1 | -5.06 |
| Trypsin | 7 | 59 | 1140 | 1 | 4816 | 4105.7061 | 81.4569 | 121,878 | 4104.698275 | 3 | 796 | 811 | DFGGFNFSQILPDPSK | 0.0000200000000000.0 | 801(NFS) | 1 | 1770.8351 | 1767.846839 | 102420190 | 87.03 | 1 | 6 | 6 | 1 | 0 | 0 | 0 | 1 | 4104.69793 | 0.0003 | 0.08 | -5.57 |
| Trypsin | 7 | 59 | 1167 | 1 | 4858 | 3026.3031 | 83.5663 | 118,691 | 3025.295275 | 3 | 796 | 811 | DFGGFNFSQILPDPSK | 0.0000200000000000.0 | 801(NFS) | 1 | 1770.8351 | 1767.846839 | 102420190 | 87.03 | 1 | 4 | 3 | 0 | 0 | 0 | 0 | 1 | 3025.296254 | -0.0010 | -0.32 | -3.46 |
| Trypsin | 7 | 59 | 1280 | 1 | 4831 | 3375.4413 | 82.2056 | 106,886 | 3374.433475 | 3 | 796 | 811 | DFGGFNFSQILPDPSK | 0.0000200000000000.0 | 801(NFS) | 1 | 1770.8351 | 1767.846839 | 102420190 | 87.03 | 1 | 4 | 4 | 1 | 0 | 0 | 0 | 1 | 3374.433536 | -0.0001 | -0.01 | -4.82 |
| Trypsin | 7 | 59 | 1314 | 1 | 4935 | 3698.5217 | 87.2062 | 104,742 | 3697.513875 | 3 | 796 | 811 | DFGGFNFSQILPDPSK | 0.0000200000000000.0 | 801(NFS) | 1 | 1770.8351 | 1767.846839 | 102420190 | 87.03 | 1 | 5 | 4 | 0 | 0 | 1 | 0 | 1 | 3697.518782 | -0.0049 | -1.32 | 0.18 |
| Trypsin | 7 | 59 | 1403 | 1 | 4836 | 3959.6521 | 82.4588 | 97,348 | 3958.644275 | 3 | 796 | 811 | DFGGFNFSQILPDPSK | 0.0000200000000000.0 | 801(NFS) | 1 | 1770.8351 | 1767.846839 | 102420190 | 87.03 | 1 | 6 | 6 | 0 | 0 | 0 | 0 | 1 | 3958.640021 | 0.0043 | 1.07 | -4.57 |
| Trypsin | 7 | 59 | 1590 | 1 | 4854 | 3067.3202 | 83.3702 | 85,338 | 3066.312375 | 3 | 796 | 811 | DFGGFNFSQILPDPSK | 0.0000200000000000.0 | 801(NFS) | 1 | 1770.8351 | 1767.846839 | 102420190 | 87.03 | 1 | 3 | 4 | 0 | 0 | 0 | 0 | 1 | 3066.322803 | -0.0104 | -3.4 | -3.66 |
| Trypsin | 7 | 59 | 1605 | 1 | 4814 | 4048.6838 | 81.3531 | 84,682 | 4047.675975 | 3 | 796 | 811 | DFGGFNFSQILPDPSK | 0.0000200000000000.0 | 801(NFS) | 1 | 1770.8351 | 1767.846839 | 102420190 | 87.03 | 1 | 6 | 5 | 2 | 0 | 0 | 0 | 1 | 4047.676466 | -0.0005 | -0.12 | -5.67 |
| Trypsin | 7 | 59 | 1730 | 1 | 4797 | 4340.7912 | 80.5512 | 77,751 | 4339.783375 | 3 | 796 | 811 | DFGGFNFSQILPDPSK | 0.0000200000000000.0 | 801(NFS) | 1 | 1770.8351 | 1767.846839 | 102420190 | 87.03 | 1 | 6 | 5 | 4 | 0 | 0 | 0 | 0 | 4339.792284 | -0.0089 | -2.05 | -6.48 |
| Trypsin | 7 | 59 | 1809 | 1 | 4941 | 3641.5067 | 87.5004 | 73,651 | 3640.498875 | 3 | 796 | 811 | DFGGFNFSQILPDPSK | 0.0000200000000000.0 | 801(NFS) | 1 | 1770.8351 | 1767.846839 | 102420190 | 87.03 | 1 | 6 | 3 | 0 | 1 | 0 | 0 | 1 | 3640.497319 | 0.0016 | 0.42 | 0.47 |
| Trypsin | 7 | 59 | 1809 | 1 | 4941 | 3641.5067 | 87.5004 | 73,651 | 3640.498875 | 3 | 796 | 811 | DFGGFNFSQILPDPSK | 0.0000200000000000.0 | 801(NFS) | 1 | 1770.8351 | 1767.846839 | 102420190 | 87.03 | 2 | 5 | 3 | 1 | 0 | 1 | 0 | 0 | 3640.497318 | 0.0016 | 0.42 | 0.47 |
| Trypsin | 7 | 59 | 1878 | 1 | 4810 | 4194.743 | 81.1665 | 70,953 | 4193.735175 | 3 | 796 | 811 | DFGGFNFSQILPDPSK | 0.0000200000000000.0 | 801(NFS) | 1 | 1770.8351 | 1767.846839 | 102420190 | 87.03 | 1 | 6 | 5 | 3 | 0 | 0 | 0 | 1 | 4193.734375 | 0.0008 | 0.19 | -5.86 |
| Trypsin | 7 | 59 | 1931 | 1 | 4945 | 3479.4492 | 87.7217 | 68,712 | 3478.441375 | 3 | 796 | 811 | DFGGFNFSQILPDPSK | 0.0000200000000000.0 | 801(NFS) | 1 | 1770.8351 | 1767.846839 | 102420190 | 87.03 | 1 | 4 | 3 | 1 | 0 | 1 | 0 | 1 | 3478.444494 | -0.0031 | -0.89 | 0.69 |
| Trypsin | 7 | 59 | 1931 | 1 | 4945 | 3479.4492 | 87.7217 | 68,712 | 3478.441375 | 3 | 796 | 811 | DFGGFNFSQILPDPSK | 0.0000200000000000.0 | 801(NFS) | 1 | 1770.8351 | 1767.846839 | 102420190 | 87.03 | 2 | 5 | 3 | 0 | 1 | 0 | 0 | 1 | 3478.444495 | -0.0031 | -0.89 | 0.69 |
| Trypsin | 7 | 59 | 1935 | 1 | 4861 | 2823.2261 | 83.7142 | 68,420 | 2822.218275 | 3 | 796 | 811 | DFGGFNFSQILPDPSK | 0.0000200000000000.0 | 801(NFS) | 1 | 1770.8351 | 1767.846839 | 102420190 | 87.03 | 1 | 4 | 2 | 0 | 0 | 0 | 0 | 1 | 2822.216881 | 0.0014 | 0.49 | -3.31 |
| Trypsin | 7 | 59 | 1959 | 1 | 4814 | 4251.7716 | 81.3531 | 67,076 | 4250.763775 | 3 | 796 | 811 | DFGGFNFSQILPDPSK | 0.0000200000000000.0 | 801(NFS) | 1 | 1770.8351 | 1767.846839 | 102420190 | 87.03 | 1 | 6 | 6 | 2 | 0 | 0 | 0 | 1 | 4250.755839 | 0.0079 | 1.86 | -5.67 |
| Trypsin | 7 | 59 | 2102 | 1 | 4839 | 3658.5219 | 82.6166 | 61,762 | 3657.514075 | 3 | 796 | 811 | DFGGFNFSQILPDPSK | 0.0000200000000000.0 | 801(NFS) | 1 | 1770.8351 | 1767.846839 | 102420190 | 87.03 | 1 | 7 | 3 | 1 | 0 | 0 | 0 | 0 | 3657.512635 | 0.0014 | 0.39 | -4.41 |
| Trypsin | 7 | 59 | 2143 | 1 | 4826 | 3521.503 | 81.9701 | 60,408 | 3520.495175 | 3 | 796 | 811 | DFGGFNFSQILPDPSK | 0.0000200000000000.0 | 801(NFS) | 1 | 1770.8351 | 1767.846839 | 102420190 | 87.03 | 1 | 4 | 4 | 2 | 0 | 0 | 0 | 1 | 3520.491445 | 0.0037 | 1.05 | -5.06 |
| Trypsin | 7 | 59 | 2222 | 1 | 4822 | 3943.6684 | 81.756 | 57,939 | 3942.660575 | 3 | 796 | 811 | DFGGFNFSQILPDPSK | 0.0000200000000000.0 | 801(NFS) | 1 | 1770.8351 | 1767.846839 | 102420190 | 87.03 | 1 | 5 | 6 | 1 | 0 | 0 | 0 | 1 | 3942.645106 | 0.0155 | 3.92 | -5.27 |
| Trypsin | 7 | 59 | 2492 | 1 | 4824 | 3724.5787 | 81.8587 | 50,382 | 3723.570875 | 3 | 796 | 811 | DFGGFNFSQILPDPSK | 0.0000200000000000.0 | 801(NFS) | 1 | 1770.8351 | 1767.846839 | 102420190 | 87.03 | 1 | 4 | 5 | 2 | 0 | 0 | 0 | 1 | 3723.570818 | 0.0001 | 0.01 | -5.17 |
| Trypsin | 7 | 59 | 2494 | 1 | 4842 | 3633.4889 | 82.7696 | 50,316 | 3632.481075 | 3 | 796 | 811 | DFGGFNFSQILPDPSK | 0.0000200000000000.0 | 801(NFS) | 1 | 1770.8351 | 1767.846839 | 102420190 | 87.03 | 1 | 9 | 2 | 0 | 0 | 0 | 0 | 1 | 3632.481001 | 0.0001 | 0.02 | -4.26 |
| Trypsin | 7 | 59 | 2717 | 1 | 4811 | 4397.8205 | 81.219 | 45,100 | 4396.812675 | 3 | 796 | 811 | DFGGFNFSQILPDPSK | 0.0000200000000000.0 | 801(NFS) | 1 | 1770.8351 | 1767.846839 | 102420190 | 87.03 | 1 | 6 | 6 | 3 | 0 | 0 | 0 | 1 | 4396.813748 | -0.0011 | -0.24 | -5.81 |
| Trypsin | 7 | 59 | 2733 | 1 | 4901 | 4209.7155 | 85.5943 | 44,584 | 4208.707675 | 3 | 796 | 811 | DFGGFNFSQILPDPSK | 0.0000200000000000.0 | 801(NFS) | 1 | 1770.8351 | 1767.846839 | 102420190 | 87.03 | 1 | 6 | 5 | 1 | 0 | 1 | 0 | 1 | 4208.708888 | -0.0012 | -0.28 | -1.43 |
| Trypsin | 7 | 59 | 2733 | 1 | 4901 | 4209.7155 | 85.5943 | 44,584 | 4208.707675 | 3 | 796 | 811 | DFGGFNFSQILPDPSK | 0.0000200000000000.0 | 801(NFS) | 1 | 1770.8351 | 1767.846839 | 102420190 | 87.03 | 2 | 7 | 5 | 0 | 1 | 0 | 0 | 0 | 4208.708889 | -0.0012 | -0.28 | -1.43 |
| Trypsin | 7 | 59 | 2861 | 1 | 4904 | 3990.6416 | 85.7353 | 41,701 | 3989.633775 | 3 | 796 | 811 | DFGGFNFSQILPDPSK | 0.0000200000000000.0 | 801(NFS) | 1 | 1770.8351 | 1767.846839 | 102420190 | 87.03 | 1 | 5 | 4 | 2 | 0 | 1 | 0 | 1 | 3989.6346 | -0.0008 | -0.2 | -1.29 |
| Trypsin | 7 | 59 | 2861 | 1 | 4904 | 3990.6416 | 85.7353 | 41,701 | 3989.633775 | 3 | 796 | 811 | DFGGFNFSQILPDPSK | 0.0000200000000000.0 | 801(NFS) | 1 | 1770.8351 | 1767.846839 | 102420190 | 87.03 | 2 | 6 | 4 | 1 | 1 | 0 | 0 | 1 | 3989.634601 | -0.0008 | -0.2 | -1.29 |
| Trypsin | 7 | 59 | 3127 | 1 | 4912 | 3828.5881 | 86.1173 | 37,420 | 3827.580275 | 3 | 796 | 811 | DFGGFNFSQILPDPSK | 0.0000200000000000.0 | 801(NFS) | 1 | 1770.8351 | 1767.846839 | 102420190 | 87.03 | 1 | 4 | 4 | 2 | 0 | 1 | 0 | 1 | 3827.581776 | -0.0015 | -0.39 | -0.91 |
| Trypsin | 7 | 59 | 3127 | 1 | 4912 | 3828.5881 | 86.1173 | 37,420 | 3827.580275 | 3 | 796 | 811 | DFGGFNFSQILPDPSK | 0.0000200000000000.0 | 801(NFS) | 1 | 1770.8351 | 1767.846839 | 102420190 | 87.03 | 2 | 5 | 4 | 1 | 1 | 0 | 0 | 1 | 3827.581777 | -0.0015 | -0.39 | -0.91 |
| Trypsin | 7 | 59 | 3263 | 1 | 4838 | 3845.6184 | 82.5607 | 35,127 | 3844.610575 | 3 | 796 | 811 | DFGGFNFSQILPDPSK | 0.0000200000000000.0 | 801(NFS) | 1 | 1770.8351 | 1767.846839 | 102420190 | 87.03 | 1 | 6 | 4 | 2 | 0 | 0 | 0 | 1 | 3844.597093 | 0.0135 | 3.5 | -4.47 |
| Trypsin | 7 | 59 | 3297 | 1 | 4829 | 4251.7536 | 82.1157 | 34,739 | 4250.745775 | 3 | 796 | 811 | DFGGFNFSQILPDPSK | 0.0000200000000000.0 | 801(NFS) | 1 | 1770.8351 | 1767.846839 | 102420190 | 87.03 | 1 | 6 | 6 | 2 | 0 | 0 | 0 | 1 | 4250.755839 | -0.0101 | -2.36 | -4.91 |
| Trypsin | 7 | 59 | 3582 | 1 | 4836 | 4007.6587 | 82.4588 | 30,758 | 4006.650875 | 3 | 796 | 811 | DFGGFNFSQILPDPSK | 0.0000200000000000.0 | 801(NFS) | 1 | 1770.8351 | 1767.846839 | 102420190 | 87.03 | 1 | 7 | 4 | 2 | 0 | 0 | 0 | 1 | 4006.649917 | 0.0010 | 0.23 | -4.57 |
| Trypsin | 7 | 59 | 4626 | 1 | 4831 | 3619.5509 | 82.2056 | 20,401 | 3618.543075 | 3 | 796 | 811 | DFGGFNFSQILPDPSK | 0.0000200000000000.0 | 801(NFS) | 1 | 1770.8351 | 1767.846839 | 102420190 | 87.03 | 1 | 3 | 6 | 1 | 0 | 0 | 0 | 1 | 3618.539458 | 0.0036 | 0.99 | -4.82 |
| Trypsin | 7 | 59 | 5042 | 1 | 4838 | 3861.5973 | 82.5607 | 17,401 | 3860.589475 | 3 | 796 | 811 | DFGGFNFSQILPDPSK | 0.0000200000000000.0 | 801(NFS) | 1 | 1770.8351 | 1767.846839 | 102420190 | 87.03 | 1 | 7 | 4 | 1 | 0 | 0 | 0 | 1 | 3860.592008 | -0.0025 | -0.65 | -4.47 |
| Trypsin | 7 | 59 | 5102 | 1 | 4825 | 4194.7478 | 81.9124 | 16,896 | 4193.739975 | 3 | 796 | 811 | DFGGFNFSQILPDPSK | 0.0000200000000000.0 | 801(NFS) | 1 | 1770.8351 | 1767.846839 | 102420190 | 87.03 | 1 | 6 | 5 | 3 | 0 | 0 | 0 | 1 | 4193.734375 | 0.0056 | 1.33 | -5.12 |
| Trypsin | 7 | 59 | 5608 | 1 | 4906 | 4193.7162 | 85.8333 | 13,462 | 4192.708375 | 3 | 796 | 811 | DFGGFNFSQILPDPSK | 0.0000200000000000.0 | 801(NFS) | 1 | 1770.8351 | 1767.846839 | 102420190 | 87.03 | 1 | 6 | 5 | 1 | 1 | 0 | 0 | 1 | 4192.713974 | -0.0056 | -1.33 | -1.19 |
| Trypsin | 7 | 59 | 5608 | 1 | 4906 | 4193.7162 | 85.8333 | 13,462 | 4192.708375 | 3 | 796 | 811 | DFGGFNFSQILPDPSK | 0.0000200000000000.0 | 801(NFS) | 1 | 1770.8351 | 1767.846839 | 102420190 | 87.03 | 2 | 5 | 5 | 2 | 0 | 1 | 0 | 0 | 4192.713973 | -0.0056 | -1.33 | -1.19 |
| Trypsin | 7 | 59 | 5800 | 1 | 4901 | 4047.6639 | 85.5943 | 12,349 | 4046.656075 | 3 | 796 | 811 | DFGGFNFSQILPDPSK | 0.0000200000000000.0 | 801(NFS) | 1 | 1770.8351 | 1767.846839 | 102420190 | 87.03 | 1 | 6 | 5 | 0 | 1 | 0 | 0 | 1 | 4046.656065 | 0.0000 | 0 | -1.43 |
| Trypsin | 7 | 59 | 5800 | 1 | 4901 | 4047.6639 | 85.5943 | 12,349 | 4046.656075 | 3 | 796 | 811 | DFGGFNFSQILPDPSK | 0.0000200000000000.0 | 801(NFS) | 1 | 1770.8351 | 1767.846839 | 102420190 | 87.03 | 2 | 5 | 5 | 1 | 0 | 1 | 0 | 1 | 4046.656064 | 0.0000 | 0 | -1.43 |
| Trypsin | 7 | 59 | 6659 | 1 | 4841 | 3715.5417 | 82.7165 | 7,093 | 3714.533875 | 3 | 796 | 811 | DFGGFNFSQILPDPSK | 0.0000200000000000.0 | 801(NFS) | 1 | 1770.8351 | 1767.846839 | 102420190 | 87.03 | 1 | 7 | 4 | 0 | 0 | 0 | 0 | 1 | 3714.534099 | -0.0002 | -0.06 | -4.31 |
| Trypsin | 63 | 16 | 426 | 1 | 4745 | 3621.5606 | 77.9978 | 343,264 | 3620.552775 | 43 | 796 | 814 | DFGGFNFSQILPDPSKPSK | 0.00000200000000000000.0 | 801(NFS) | 1 | 2083.0148 | 2080.026539 | 3024780.3 | 81.56 | 1 | 7 | 2 | 0 | 0 | 0 | 0 | 1 | 3620.555053 | -0.0023 | -0.62 | -3.56 |
| Trypsin | 63 | 16 | 540 | 1 | 4748 | 3459.5075 | 78.1353 | 267,172 | 3458.499675 | 43 | 796 | 814 | DFGGFNFSQILPDPSKPSK | 0.00000200000000000000.0 | 801(NFS) | 1 | 2083.0148 | 2080.026539 | 3024780.3 | 81.56 | 1 | 6 | 2 | 0 | 0 | 0 | 0 | 1 | 3458.502229 | -0.0026 | -0.73 | -3.42 |
| Trypsin | 63 | 16 | 1069 | 1 | 4742 | 3808.6424 | 77.8546 | 128,487 | 3807.634575 | 43 | 796 | 814 | DFGGFNFSQILPDPSKPSK | 0.00000200000000000000.0 | 801(NFS) | 1 | 2083.0148 | 2080.026539 | 3024780.3 | 81.56 | 1 | 6 | 3 | 1 | 0 | 0 | 0 | 1 | 3807.639511 | -0.0049 | -1.29 | -3.70 |
| Trypsin | 63 | 16 | 1196 | 1 | 4752 | 3297.4566 | 78.3334 | 115,665 | 3296.448775 | 43 | 796 | 814 | DFGGFNFSQILPDPSKPSK | 0.00000200000000000000.0 | 801(NFS) | 1 | 2083.0148 | 2080.026539 | 3024780.3 | 81.56 | 1 | 5 | 2 | 0 | 0 | 0 | 0 | 1 | 3296.449405 | -0.0006 | -0.19 | -3.23 |
| Trypsin | 63 | 16 | 1374 | 1 | 4748 | 3646.5886 | 78.1353 | 99,412 | 3645.580775 | 43 | 796 | 814 | DFGGFNFSQILPDPSKPSK | 0.00000200000000000000.0 | 801(NFS) | 1 | 2083.0148 | 2080.026539 | 3024780.3 | 81.56 | 1 | 5 | 3 | 1 | 0 | 0 | 0 | 1 | 3645.586687 | -0.0059 | -1.62 | -3.42 |
| Trypsin | 63 | 16 | 1887 | 1 | 4742 | 3783.6101 | 77.8546 | 70,507 | 3782.602275 | 43 | 796 | 814 | DFGGFNFSQILPDPSKPSK | 0.00000200000000000000.0 | 801(NFS) | 1 | 2083.0148 | 2080.026539 | 3024780.3 | 81.56 | 1 | 8 | 2 | 0 | 0 | 0 | 0 | 1 | 3782.607877 | -0.0056 | -1.48 | -3.70 |
| Trypsin | 63 | 16 | 2181 | 1 | 4745 | 3662.5946 | 77.9978 | 59,177 | 3661.586775 | 43 | 796 | 814 | DFGGFNFSQILPDPSKPSK | 0.00000200000000000000.0 | 801(NFS) | 1 | 2083.0148 | 2080.026539 | 3024780.3 | 81.56 | 1 | 6 | 3 | 0 | 0 | 0 | 0 | 1 | 3661.581602 | 0.0052 | 1.41 | -3.56 |
| Trypsin | 63 | 16 | 2466 | 1 | 4750 | 3500.5367 | 78.24 | 50,869 | 3499.528875 | 43 | 796 | 814 | DFGGFNFSQILPDPSKPSK | 0.00000200000000000000.0 | 801(NFS) | 1 | 2083.0148 | 2080.026539 | 3024780.3 | 81.56 | 1 | 5 | 3 | 0 | 0 | 0 | 0 | 1 | 3499.528778 | 0.0001 | 0.02 | -3.32 |
| Trypsin | 63 | 16 | 3290 | 1 | 4739 | 3849.669 | 77.7182 | 34,796 | 3848.661175 | 43 | 796 | 814 | DFGGFNFSQILPDPSKPSK | 0.00000200000000000000.0 | 801(NFS) | 1 | 2083.0148 | 2080.026539 | 3024780.3 | 81.56 | 1 | 5 | 4 | 1 | 0 | 0 | 0 | 1 | 3848.66606 | -0.0049 | -1.26 | -3.84 |
| Trypsin | 63 | 16 | 3534 | 1 | 4741 | 3703.6179 | 77.804 | 31,558 | 3702.610075 | 43 | 796 | 814 | DFGGFNFSQILPDPSKPSK | 0.00000200000000000000.0 | 801(NFS) | 1 | 2083.0148 | 2080.026539 | 3024780.3 | 81.56 | 1 | 5 | 4 | 0 | 0 | 0 | 0 | 1 | 3702.608151 | 0.0019 | 0.51 | -3.76 |
| Trypsin | 63 | 16 | 4000 | 1 | 4739 | 3906.6869 | 77.7182 | 26,217 | 3905.679075 | 43 | 796 | 814 | DFGGFNFSQILPDPSKPSK | 0.00000200000000000000.0 | 801(NFS) | 1 | 2083.0148 | 2080.026539 | 3024780.3 | 81.56 | 1 | 5 | 5 | 0 | 0 | 0 | 0 | 1 | 3905.687524 | -0.0084 | -2.16 | -3.84 |
| Trypsin | 63 | 16 | 4214 | 1 | 4733 | 3995.7315 | 77.4041 | 23,889 | 3994.723675 | 43 | 796 | 814 | DFGGFNFSQILPDPSKPSK | 0.00000200000000000000.0 | 801(NFS) | 1 | 2083.0148 | 2080.026539 | 3024780.3 | 81.56 | 1 | 5 | 4 | 2 | 0 | 0 | 0 | 1 | 3994.723969 | -0.0003 | -0.07 | -4.16 |
| Trypsin | 63 | 16 | 4464 | 1 | 4745 | 3744.6438 | 77.9978 | 21,723 | 3743.635975 | 43 | 796 | 814 | DFGGFNFSQILPDPSKPSK | 0.00000200000000000000.0 | 801(NFS) | 1 | 2083.0148 | 2080.026539 | 3024780.3 | 81.56 | 1 | 4 | 5 | 0 | 0 | 0 | 0 | 1 | 3743.6347 | 0.0013 | 0.34 | -3.56 |
| Trypsin | 63 | 16 | 4986 | 1 | 4832 | 3807.6244 | 82.2602 | 17,815 | 3806.616575 | 43 | 796 | 814 | DFGGFNFSQILPDPSKPSK | 0.00000200000000000000.0 | 801(NFS) | 1 | 2083.0148 | 2080.026539 | 3024780.3 | 81.56 | 1 | 5 | 3 | 0 | 0 | 1 | 0 | 1 | 3806.619109 | -0.0025 | -0.66 | 0.70 |
| Trypsin | 63 | 16 | 5821 | 1 | 4737 | 4052.7517 | 77.6036 | 12,292 | 4051.743875 | 43 | 796 | 814 | DFGGFNFSQILPDPSKPSK | 0.00000200000000000000.0 | 801(NFS) | 1 | 2083.0148 | 2080.026539 | 3024780.3 | 81.56 | 1 | 5 | 5 | 1 | 0 | 0 | 0 | 1 | 4051.745433 | -0.0016 | -0.38 | -3.96 |
| Trypsin | 63 | 16 | 6259 | 1 | 4726 | 3890.7052 | 77.0809 | 9,572 | 3889.697375 | 43 | 796 | 814 | DFGGFNFSQILPDPSKPSK | 0.00000200000000000000.0 | 801(NFS) | 1 | 2083.0148 | 2080.026539 | 3024780.3 | 81.56 | 1 | 4 | 5 | 1 | 0 | 0 | 0 | 1 | 3889.692609 | 0.0048 | 1.22 | -4.48 |
| Trypsin | 102 | 4 | 891 | 1 | 4833 | 4541.9498 | 82.3114 | 159,159 | 4540.941975 | 365 | 791 | 811 | TPPIKDFGGFNFSQILPDPSK | 0.0000000000020000000000.0 | 801(NFS) | 1 | 2307.1673 | 2304.179039 | 79845.984 | 83.15 | 1 | 5 | 4 | 0 | 0 | 2 | 0 | 1 | 4540.941313 | 0.0007 | 0.14 | -0.83 |
| Trypsin | 102 | 4 | 5476 | 1 | 4906 | 5198.1735 | 85.8333 | 14,341 | 5197.165675 | 365 | 791 | 811 | TPPIKDFGGFNFSQILPDPSK | 0.0000000000020000000000.0 | 801(NFS) | 1 | 2307.1673 | 2304.179039 | 79845.984 | 83.15 | 1 | 6 | 5 | 0 | 1 | 2 | 0 | 1 | 5197.168927 | -0.0033 | -0.62 | 2.69 |
| Trypsin | 102 | 4 | 5476 | 1 | 4906 | 5198.1735 | 85.8333 | 14,341 | 5197.165675 | 365 | 791 | 811 | TPPIKDFGGFNFSQILPDPSK | 0.0000000000020000000000.0 | 801(NFS) | 1 | 2307.1673 | 2304.179039 | 79845.984 | 83.15 | 2 | 5 | 5 | 1 | 0 | 3 | 1 | -1 | 5197.168926 | -0.0033 | -0.62 | 2.69 |
| Trypsin | 102 | 4 | 6469 | 1 | 4888 | 4833.035 | 84.9797 | 8,414 | 4832.027175 | 365 | 791 | 811 | TPPIKDFGGFNFSQILPDPSK | 0.0000000000020000000000.0 | 801(NFS) | 1 | 2307.1673 | 2304.179039 | 79845.984 | 83.15 | 1 | 4 | 4 | 1 | 0 | 3 | 1 | -1 | 4832.036729 | -0.0096 | -1.97 | 1.83 |
| Trypsin | 102 | 4 | 6469 | 1 | 4888 | 4833.035 | 84.9797 | 8,414 | 4832.027175 | 365 | 791 | 811 | TPPIKDFGGFNFSQILPDPSK | 0.0000000000020000000000.0 | 801(NFS) | 1 | 2307.1673 | 2304.179039 | 79845.984 | 83.15 | 2 | 5 | 4 | 0 | 1 | 2 | 1 | -1 | 4832.03673 | -0.0096 | -1.97 | 1.83 |
| Trypsin | 102 | 4 | 6829 | 1 | 4905 | 4833.0392 | 85.7818 | 5,889 | 4832.031375 | 365 | 791 | 811 | TPPIKDFGGFNFSQILPDPSK | 0.0000000000020000000000.0 | 801(NFS) | 1 | 2307.1673 | 2304.179039 | 79845.984 | 83.15 | 1 | 4 | 4 | 1 | 0 | 3 | 1 | -1 | 4832.036729 | -0.0054 | -1.1 | 2.64 |
| Trypsin | 102 | 4 | 6829 | 1 | 4905 | 4833.0392 | 85.7818 | 5,889 | 4832.031375 | 365 | 791 | 811 | TPPIKDFGGFNFSQILPDPSK | 0.0000000000020000000000.0 | 801(NFS) | 1 | 2307.1673 | 2304.179039 | 79845.984 | 83.15 | 2 | 5 | 4 | 0 | 1 | 2 | 1 | -1 | 4832.03673 | -0.0054 | -1.1 | 2.64 |
| alpha-Lytic | 11 | 35 | 52 | 1 | 4695 | 3627.628 | 70.5445 | 2,631,870 | 3626.620175 | 14 | 786 | 803 | KQIYKTPPIKDFGGFNFS | 0.000000000000000200.0 | 801(NFS) | 1 | 2089.077 | 2086.088739 | 8296301.5 | 73.10 | 1 | 7 | 2 | 0 | 0 | 0 | 0 | 1 | 3626.617253 | 0.0029 | 0.8 | -2.56 |
| alpha-Lytic | 11 | 35 | 76 | 1 | 4698 | 3465.5724 | 70.6832 | 1,696,420 | 3464.564575 | 14 | 786 | 803 | KQIYKTPPIKDFGGFNFS | 0.000000000000000200.0 | 801(NFS) | 1 | 2089.077 | 2086.088739 | 8296301.5 | 73.10 | 1 | 6 | 2 | 0 | 0 | 0 | 0 | 1 | 3464.564429 | 0.0001 | 0.04 | -2.42 |
| alpha-Lytic | 11 | 35 | 133 | 1 | 4693 | 3814.7093 | 70.4482 | 1,049,650 | 3813.701475 | 14 | 786 | 803 | KQIYKTPPIKDFGGFNFS | 0.000000000000000200.0 | 801(NFS) | 1 | 2089.077 | 2086.088739 | 8296301.5 | 73.10 | 1 | 6 | 3 | 1 | 0 | 0 | 0 | 1 | 3813.701711 | -0.0002 | -0.06 | -2.65 |
| alpha-Lytic | 11 | 35 | 164 | 1 | 4702 | 3303.5186 | 70.8824 | 831,407 | 3302.510775 | 14 | 786 | 803 | KQIYKTPPIKDFGGFNFS | 0.000000000000000200.0 | 801(NFS) | 1 | 2089.077 | 2086.088739 | 8296301.5 | 73.10 | 1 | 5 | 2 | 0 | 0 | 0 | 0 | 1 | 3302.511605 | -0.0008 | -0.25 | -2.22 |
| alpha-Lytic | 11 | 35 | 185 | 1 | 4693 | 3789.6801 | 70.4482 | 719,566 | 3788.672275 | 14 | 786 | 803 | KQIYKTPPIKDFGGFNFS | 0.000000000000000200.0 | 801(NFS) | 1 | 2089.077 | 2086.088739 | 8296301.5 | 73.10 | 1 | 8 | 2 | 0 | 0 | 0 | 0 | 1 | 3788.670077 | 0.0022 | 0.58 | -2.65 |
| alpha-Lytic | 11 | 35 | 212 | 1 | 4698 | 3652.6586 | 70.6832 | 645,757 | 3651.650775 | 14 | 786 | 803 | KQIYKTPPIKDFGGFNFS | 0.000000000000000200.0 | 801(NFS) | 1 | 2089.077 | 2086.088739 | 8296301.5 | 73.10 | 1 | 5 | 3 | 1 | 0 | 0 | 0 | 1 | 3651.648887 | 0.0019 | 0.51 | -2.42 |
| alpha-Lytic | 11 | 35 | 315 | 1 | 4697 | 3668.6496 | 70.636 | 422,323 | 3667.641775 | 14 | 786 | 803 | KQIYKTPPIKDFGGFNFS | 0.000000000000000200.0 | 801(NFS) | 1 | 2089.077 | 2086.088739 | 8296301.5 | 73.10 | 1 | 6 | 3 | 0 | 0 | 0 | 0 | 1 | 3667.643802 | -0.0020 | -0.55 | -2.46 |
| alpha-Lytic | 11 | 35 | 338 | 1 | 4692 | 3855.7379 | 70.3941 | 396,267 | 3854.730075 | 14 | 786 | 803 | KQIYKTPPIKDFGGFNFS | 0.000000000000000200.0 | 801(NFS) | 1 | 2089.077 | 2086.088739 | 8296301.5 | 73.10 | 1 | 5 | 4 | 1 | 0 | 0 | 0 | 1 | 3854.72826 | 0.0018 | 0.47 | -2.71 |
| alpha-Lytic | 11 | 35 | 418 | 1 | 4701 | 3506.5976 | 70.8287 | 311,198 | 3505.589775 | 14 | 786 | 803 | KQIYKTPPIKDFGGFNFS | 0.000000000000000200.0 | 801(NFS) | 1 | 2089.077 | 2086.088739 | 8296301.5 | 73.10 | 1 | 5 | 3 | 0 | 0 | 0 | 0 | 1 | 3505.590978 | -0.0012 | -0.34 | -2.27 |
| alpha-Lytic | 11 | 35 | 491 | 1 | 4689 | 4001.7977 | 70.2528 | 263,656 | 4000.789875 | 14 | 786 | 803 | KQIYKTPPIKDFGGFNFS | 0.000000000000000200.0 | 801(NFS) | 1 | 2089.077 | 2086.088739 | 8296301.5 | 73.10 | 1 | 5 | 4 | 2 | 0 | 0 | 0 | 1 | 4000.786169 | 0.0037 | 0.92 | -2.85 |
| alpha-Lytic | 11 | 35 | 612 | 1 | 4693 | 3709.68 | 70.4482 | 218,651 | 3708.672175 | 14 | 786 | 803 | KQIYKTPPIKDFGGFNFS | 0.000000000000000200.0 | 801(NFS) | 1 | 2089.077 | 2086.088739 | 8296301.5 | 73.10 | 1 | 5 | 4 | 0 | 0 | 0 | 0 | 1 | 3708.670351 | 0.0018 | 0.49 | -2.65 |
| alpha-Lytic | 11 | 35 | 649 | 1 | 4757 | 3975.7429 | 73.5854 | 208,363 | 3974.735075 | 14 | 786 | 803 | KQIYKTPPIKDFGGFNFS | 0.000000000000000200.0 | 801(NFS) | 1 | 2089.077 | 2086.088739 | 8296301.5 | 73.10 | 1 | 6 | 3 | 0 | 0 | 1 | 0 | 1 | 3974.734133 | 0.0009 | 0.23 | 0.48 |
| alpha-Lytic | 11 | 35 | 657 | 1 | 4691 | 4058.8128 | 70.3482 | 205,197 | 4057.804975 | 14 | 786 | 803 | KQIYKTPPIKDFGGFNFS | 0.000000000000000200.0 | 801(NFS) | 1 | 2089.077 | 2086.088739 | 8296301.5 | 73.10 | 1 | 5 | 5 | 1 | 0 | 0 | 0 | 1 | 4057.807633 | -0.0027 | -0.65 | -2.75 |
| alpha-Lytic | 11 | 35 | 680 | 1 | 4694 | 3912.7616 | 70.5009 | 196,311 | 3911.753775 | 14 | 786 | 803 | KQIYKTPPIKDFGGFNFS | 0.000000000000000200.0 | 801(NFS) | 1 | 2089.077 | 2086.088739 | 8296301.5 | 73.10 | 1 | 5 | 5 | 0 | 0 | 0 | 0 | 1 | 3911.749724 | 0.0041 | 1.03 | -2.60 |
| alpha-Lytic | 11 | 35 | 724 | 1 | 4760 | 3813.6897 | 73.7401 | 184,039 | 3812.681875 | 14 | 786 | 803 | KQIYKTPPIKDFGGFNFS | 0.000000000000000200.0 | 801(NFS) | 1 | 2089.077 | 2086.088739 | 8296301.5 | 73.10 | 1 | 5 | 3 | 0 | 0 | 1 | 0 | 1 | 3812.681309 | 0.0006 | 0.14 | 0.64 |
| alpha-Lytic | 11 | 35 | 974 | 1 | 4688 | 4204.8724 | 70.1988 | 136,620 | 4203.864575 | 14 | 786 | 803 | KQIYKTPPIKDFGGFNFS | 0.000000000000000200.0 | 801(NFS) | 1 | 2089.077 | 2086.088739 | 8296301.5 | 73.10 | 1 | 5 | 5 | 2 | 0 | 0 | 0 | 1 | 4203.865542 | -0.0010 | -0.22 | -2.90 |
| alpha-Lytic | 11 | 35 | 1148 | 1 | 4753 | 4016.7715 | 73.3868 | 114,755 | 4015.763675 | 14 | 786 | 803 | KQIYKTPPIKDFGGFNFS | 0.000000000000000200.0 | 801(NFS) | 1 | 2089.077 | 2086.088739 | 8296301.5 | 73.10 | 1 | 5 | 4 | 0 | 0 | 1 | 0 | 1 | 4015.760682 | 0.0030 | 0.74 | 0.29 |
| alpha-Lytic | 11 | 35 | 1167 | 1 | 4696 | 3750.7069 | 70.5906 | 112,057 | 3749.699075 | 14 | 786 | 803 | KQIYKTPPIKDFGGFNFS | 0.000000000000000200.0 | 801(NFS) | 1 | 2089.077 | 2086.088739 | 8296301.5 | 73.10 | 1 | 4 | 4 | 0 | 0 | 0 | 0 | 1 | 3749.6969 | 0.0022 | 0.58 | -2.51 |
| alpha-Lytic | 11 | 35 | 1359 | 1 | 4692 | 3896.7645 | 70.3941 | 95,458 | 3895.756675 | 14 | 786 | 803 | KQIYKTPPIKDFGGFNFS | 0.000000000000000200.0 | 801(NFS) | 1 | 2089.077 | 2086.088739 | 8296301.5 | 73.10 | 1 | 4 | 5 | 1 | 0 | 0 | 0 | 1 | 3895.754809 | 0.0019 | 0.47 | -2.71 |
| alpha-Lytic | 11 | 35 | 1645 | 1 | 4696 | 3871.7418 | 70.5906 | 76,344 | 3870.733975 | 14 | 786 | 803 | KQIYKTPPIKDFGGFNFS | 0.000000000000000200.0 | 801(NFS) | 1 | 2089.077 | 2086.088739 | 8296301.5 | 73.10 | 1 | 6 | 4 | 0 | 0 | 0 | 0 | 1 | 3870.723175 | 0.0108 | 2.79 | -2.51 |
| alpha-Lytic | 11 | 35 | 1810 | 1 | 4685 | 4147.8563 | 70.0561 | 67,370 | 4146.848475 | 14 | 786 | 803 | KQIYKTPPIKDFGGFNFS | 0.000000000000000200.0 | 801(NFS) | 1 | 2089.077 | 2086.088739 | 8296301.5 | 73.10 | 1 | 5 | 4 | 3 | 0 | 0 | 0 | 1 | 4146.844078 | 0.0044 | 1.06 | -3.04 |
| alpha-Lytic | 11 | 35 | 2083 | 1 | 4683 | 4350.9264 | 69.9553 | 56,192 | 4349.918575 | 14 | 786 | 803 | KQIYKTPPIKDFGGFNFS | 0.000000000000000200.0 | 801(NFS) | 1 | 2089.077 | 2086.088739 | 8296301.5 | 73.10 | 1 | 5 | 5 | 3 | 0 | 0 | 0 | 1 | 4349.923451 | -0.0049 | -1.12 | -3.15 |
| alpha-Lytic | 11 | 35 | 2250 | 1 | 4758 | 3959.748 | 73.6401 | 50,885 | 3958.740175 | 14 | 786 | 803 | KQIYKTPPIKDFGGFNFS | 0.000000000000000200.0 | 801(NFS) | 1 | 2089.077 | 2086.088739 | 8296301.5 | 73.10 | 1 | 6 | 3 | 0 | 1 | 0 | 0 | 1 | 3958.739219 | 0.0010 | 0.24 | 0.54 |
| alpha-Lytic | 11 | 35 | 2250 | 1 | 4758 | 3959.748 | 73.6401 | 50,885 | 3958.740175 | 14 | 786 | 803 | KQIYKTPPIKDFGGFNFS | 0.000000000000000200.0 | 801(NFS) | 1 | 2089.077 | 2086.088739 | 8296301.5 | 73.10 | 2 | 5 | 3 | 1 | 0 | 1 | 0 | 0 | 3958.739218 | 0.0010 | 0.24 | 0.54 |
| alpha-Lytic | 11 | 35 | 2489 | 1 | 4690 | 4074.8104 | 70.2985 | 42,846 | 4073.802575 | 14 | 786 | 803 | KQIYKTPPIKDFGGFNFS | 0.000000000000000200.0 | 801(NFS) | 1 | 2089.077 | 2086.088739 | 8296301.5 | 73.10 | 1 | 6 | 5 | 0 | 0 | 0 | 0 | 1 | 4073.802548 | 0.0000 | 0 | -2.80 |
| alpha-Lytic | 11 | 35 | 2565 | 1 | 4687 | 4423.9503 | 70.1459 | 40,825 | 4422.942475 | 14 | 786 | 803 | KQIYKTPPIKDFGGFNFS | 0.000000000000000200.0 | 801(NFS) | 1 | 2089.077 | 2086.088739 | 8296301.5 | 73.10 | 1 | 6 | 6 | 1 | 0 | 0 | 0 | 1 | 4422.93983 | 0.0026 | 0.59 | -2.95 |
| alpha-Lytic | 11 | 35 | 2578 | 1 | 4689 | 4277.8937 | 70.2528 | 40,258 | 4276.885875 | 14 | 786 | 803 | KQIYKTPPIKDFGGFNFS | 0.000000000000000200.0 | 801(NFS) | 1 | 2089.077 | 2086.088739 | 8296301.5 | 73.10 | 1 | 6 | 6 | 0 | 0 | 0 | 0 | 1 | 4276.881921 | 0.0040 | 0.92 | -2.85 |
| alpha-Lytic | 11 | 35 | 2797 | 1 | 4685 | 4366.9293 | 70.0561 | 34,651 | 4365.921475 | 14 | 786 | 803 | KQIYKTPPIKDFGGFNFS | 0.000000000000000200.0 | 801(NFS) | 1 | 2089.077 | 2086.088739 | 8296301.5 | 73.10 | 1 | 6 | 5 | 2 | 0 | 0 | 0 | 1 | 4365.918366 | 0.0031 | 0.71 | -3.04 |
| alpha-Lytic | 11 | 35 | 2956 | 1 | 4703 | 3588.6498 | 70.9331 | 30,899 | 3587.641975 | 14 | 786 | 803 | KQIYKTPPIKDFGGFNFS | 0.000000000000000200.0 | 801(NFS) | 1 | 2089.077 | 2086.088739 | 8296301.5 | 73.10 | 1 | 3 | 5 | 0 | 0 | 0 | 0 | 1 | 3587.644076 | -0.0021 | -0.58 | -2.17 |
| alpha-Lytic | 11 | 35 | 2966 | 1 | 4762 | 3797.6979 | 73.8377 | 30,798 | 3796.690075 | 14 | 786 | 803 | KQIYKTPPIKDFGGFNFS | 0.000000000000000200.0 | 801(NFS) | 1 | 2089.077 | 2086.088739 | 8296301.5 | 73.10 | 1 | 5 | 3 | 0 | 1 | 0 | 0 | 1 | 3796.686395 | 0.0037 | 0.96 | 0.74 |
| alpha-Lytic | 11 | 35 | 2966 | 1 | 4762 | 3797.6979 | 73.8377 | 30,798 | 3796.690075 | 14 | 786 | 803 | KQIYKTPPIKDFGGFNFS | 0.000000000000000200.0 | 801(NFS) | 1 | 2089.077 | 2086.088739 | 8296301.5 | 73.10 | 2 | 4 | 3 | 1 | 0 | 1 | 0 | 1 | 3796.686394 | 0.0037 | 0.96 | 0.74 |
| alpha-Lytic | 11 | 35 | 3105 | 1 | 4687 | 4220.8649 | 70.1459 | 27,613 | 4219.857075 | 14 | 786 | 803 | KQIYKTPPIKDFGGFNFS | 0.000000000000000200.0 | 801(NFS) | 1 | 2089.077 | 2086.088739 | 8296301.5 | 73.10 | 1 | 6 | 5 | 1 | 0 | 0 | 0 | 1 | 4219.860457 | -0.0034 | -0.8 | -2.95 |
| alpha-Lytic | 11 | 35 | 3174 | 1 | 4692 | 3976.7619 | 70.3941 | 26,212 | 3975.754075 | 14 | 786 | 803 | KQIYKTPPIKDFGGFNFS | 0.000000000000000200.0 | 801(NFS) | 1 | 2089.077 | 2086.088739 | 8296301.5 | 73.10 | 1 | 7 | 3 | 1 | 0 | 0 | 0 | 0 | 3975.754535 | -0.0005 | -0.11 | -2.71 |
| alpha-Lytic | 11 | 35 | 3283 | 1 | 4683 | 4512.9835 | 69.9553 | 23,798 | 4511.975675 | 14 | 786 | 803 | KQIYKTPPIKDFGGFNFS | 0.000000000000000200.0 | 801(NFS) | 1 | 2089.077 | 2086.088739 | 8296301.5 | 73.10 | 1 | 6 | 5 | 3 | 0 | 0 | 0 | 1 | 4511.976275 | -0.0006 | -0.13 | -3.15 |
| alpha-Lytic | 11 | 35 | 3627 | 1 | 4755 | 4000.7782 | 73.4945 | 16,342 | 3999.770375 | 14 | 786 | 803 | KQIYKTPPIKDFGGFNFS | 0.000000000000000200.0 | 801(NFS) | 1 | 2089.077 | 2086.088739 | 8296301.5 | 73.10 | 1 | 5 | 4 | 0 | 1 | 0 | 0 | 1 | 3999.765768 | 0.0046 | 1.15 | 0.39 |
| alpha-Lytic | 11 | 35 | 3627 | 1 | 4755 | 4000.7782 | 73.4945 | 16,342 | 3999.770375 | 14 | 786 | 803 | KQIYKTPPIKDFGGFNFS | 0.000000000000000200.0 | 801(NFS) | 1 | 2089.077 | 2086.088739 | 8296301.5 | 73.10 | 2 | 4 | 4 | 1 | 0 | 1 | 0 | 1 | 3999.765767 | 0.0046 | 1.15 | 0.39 |
| alpha-Lytic | 11 | 35 | 3670 | 1 | 4681 | 4716.0685 | 69.8656 | 15,102 | 4715.060675 | 14 | 786 | 803 | KQIYKTPPIKDFGGFNFS | 0.000000000000000200.0 | 801(NFS) | 1 | 2089.077 | 2086.088739 | 8296301.5 | 73.10 | 1 | 6 | 6 | 3 | 0 | 0 | 0 | 1 | 4715.055648 | 0.0050 | 1.06 | -3.24 |
| alpha-Lytic | 11 | 35 | 3984 | 1 | 4704 | 3344.5465 | 70.9774 | 8,480 | 3343.538675 | 14 | 786 | 803 | KQIYKTPPIKDFGGFNFS | 0.000000000000000200.0 | 801(NFS) | 1 | 2089.077 | 2086.088739 | 8296301.5 | 73.10 | 1 | 4 | 3 | 0 | 0 | 0 | 0 | 1 | 3343.538154 | 0.0005 | 0.15 | -2.12 |
| alpha-Lytic | 20 | 31 | 88 | 1 | 4905 | 4690.1971 | 80.7185 | 1,513,530 | 4689.189275 | 4 | 786 | 813 | KQIYKTPPIKDFGGFNFSQILPDPSKPS | 0.000000000000000020000000000000.0 | 801(NFS) | 1 | 3151.648 | 3148.659739 | 51370068 | 82.77 | 1 | 7 | 2 | 0 | 0 | 0 | 0 | 1 | 4689.188253 | 0.0010 | 0.21 | -2.05 |
| alpha-Lytic | 20 | 31 | 127 | 1 | 4909 | 4528.1419 | 80.9149 | 1,074,590 | 4527.134075 | 4 | 786 | 813 | KQIYKTPPIKDFGGFNFSQILPDPSKPS | 0.000000000000000020000000000000.0 | 801(NFS) | 1 | 3151.648 | 3148.659739 | 51370068 | 82.77 | 1 | 6 | 2 | 0 | 0 | 0 | 0 | 1 | 4527.135429 | -0.0014 | -0.29 | -1.86 |
| alpha-Lytic | 20 | 31 | 227 | 1 | 4905 | 4877.2847 | 80.7185 | 603,765 | 4876.276875 | 4 | 786 | 813 | KQIYKTPPIKDFGGFNFSQILPDPSKPS | 0.000000000000000020000000000000.0 | 801(NFS) | 1 | 3151.648 | 3148.659739 | 51370068 | 82.77 | 1 | 6 | 3 | 1 | 0 | 0 | 0 | 1 | 4876.272711 | 0.0042 | 0.85 | -2.05 |
| alpha-Lytic | 20 | 31 | 327 | 1 | 4911 | 4366.0896 | 81.0152 | 406,964 | 4365.081775 | 4 | 786 | 813 | KQIYKTPPIKDFGGFNFSQILPDPSKPS | 0.000000000000000020000000000000.0 | 801(NFS) | 1 | 3151.648 | 3148.659739 | 51370068 | 82.77 | 1 | 5 | 2 | 0 | 0 | 0 | 0 | 1 | 4365.082605 | -0.0008 | -0.19 | -1.76 |
| alpha-Lytic | 20 | 31 | 350 | 1 | 4905 | 4852.2524 | 80.7185 | 380,599 | 4851.244575 | 4 | 786 | 813 | KQIYKTPPIKDFGGFNFSQILPDPSKPS | 0.000000000000000020000000000000.0 | 801(NFS) | 1 | 3151.648 | 3148.659739 | 51370068 | 82.77 | 1 | 8 | 2 | 0 | 0 | 0 | 0 | 1 | 4851.241077 | 0.0035 | 0.72 | -2.05 |
| alpha-Lytic | 20 | 31 | 379 | 1 | 4909 | 4715.2262 | 80.9149 | 346,449 | 4714.218375 | 4 | 786 | 813 | KQIYKTPPIKDFGGFNFSQILPDPSKPS | 0.000000000000000020000000000000.0 | 801(NFS) | 1 | 3151.648 | 3148.659739 | 51370068 | 82.77 | 1 | 5 | 3 | 1 | 0 | 0 | 0 | 1 | 4714.219887 | -0.0015 | -0.32 | -1.86 |
| alpha-Lytic | 20 | 31 | 480 | 1 | 4889 | 5064.3646 | 79.9794 | 267,912 | 5063.356775 | 4 | 786 | 813 | KQIYKTPPIKDFGGFNFSQILPDPSKPS | 0.000000000000000020000000000000.0 | 801(NFS) | 1 | 3151.648 | 3148.659739 | 51370068 | 82.77 | 1 | 5 | 4 | 2 | 0 | 0 | 0 | 1 | 5063.357169 | -0.0004 | -0.07 | -2.79 |
| alpha-Lytic | 20 | 31 | 513 | 1 | 4907 | 4731.2208 | 80.8114 | 256,620 | 4730.212975 | 4 | 786 | 813 | KQIYKTPPIKDFGGFNFSQILPDPSKPS | 0.000000000000000020000000000000.0 | 801(NFS) | 1 | 3151.648 | 3148.659739 | 51370068 | 82.77 | 1 | 6 | 3 | 0 | 0 | 0 | 0 | 1 | 4730.214802 | -0.0018 | -0.38 | -1.96 |
| alpha-Lytic | 20 | 31 | 675 | 1 | 4903 | 4918.3012 | 80.6212 | 198,254 | 4917.293375 | 4 | 786 | 813 | KQIYKTPPIKDFGGFNFSQILPDPSKPS | 0.000000000000000020000000000000.0 | 801(NFS) | 1 | 3151.648 | 3148.659739 | 51370068 | 82.77 | 1 | 5 | 4 | 1 | 0 | 0 | 0 | 1 | 4917.29926 | -0.0059 | -1.19 | -2.15 |
| alpha-Lytic | 20 | 31 | 752 | 1 | 4891 | 4959.3343 | 80.0723 | 175,822 | 4958.326475 | 4 | 786 | 813 | KQIYKTPPIKDFGGFNFSQILPDPSKPS | 0.000000000000000020000000000000.0 | 801(NFS) | 1 | 3151.648 | 3148.659739 | 51370068 | 82.77 | 1 | 4 | 5 | 1 | 0 | 0 | 0 | 1 | 4958.325809 | 0.0007 | 0.13 | -2.70 |
| alpha-Lytic | 20 | 31 | 809 | 1 | 4913 | 4569.1736 | 81.1108 | 162,859 | 4568.165775 | 4 | 786 | 813 | KQIYKTPPIKDFGGFNFSQILPDPSKPS | 0.000000000000000020000000000000.0 | 801(NFS) | 1 | 3151.648 | 3148.659739 | 51370068 | 82.77 | 1 | 5 | 3 | 0 | 0 | 1 | 0 | 1 | 4568.161978 | 0.0038 | 0.83 | -1.66 |
| alpha-Lytic | 20 | 31 | 824 | 1 | 4890 | 5121.3824 | 80.0254 | 160,479 | 5120.374575 | 4 | 786 | 813 | KQIYKTPPIKDFGGFNFSQILPDPSKPS | 0.000000000000000020000000000000.0 | 801(NFS) | 1 | 3151.648 | 3148.659739 | 51370068 | 82.77 | 1 | 5 | 5 | 1 | 0 | 0 | 0 | 1 | 5120.378633 | -0.0041 | -0.79 | -2.75 |
| alpha-Lytic | 20 | 31 | 838 | 1 | 4886 | 5210.4255 | 79.8405 | 157,064 | 5209.417675 | 4 | 786 | 813 | KQIYKTPPIKDFGGFNFSQILPDPSKPS | 0.000000000000000020000000000000.0 | 801(NFS) | 1 | 3151.648 | 3148.659739 | 51370068 | 82.77 | 1 | 5 | 4 | 3 | 0 | 0 | 0 | 1 | 5209.415078 | 0.0026 | 0.49 | -2.93 |
| alpha-Lytic | 20 | 31 | 966 | 1 | 4904 | 4772.251 | 80.6712 | 137,334 | 4771.243175 | 4 | 786 | 813 | KQIYKTPPIKDFGGFNFSQILPDPSKPS | 0.000000000000000020000000000000.0 | 801(NFS) | 1 | 3151.648 | 3148.659739 | 51370068 | 82.77 | 1 | 5 | 4 | 0 | 0 | 0 | 0 | 1 | 4771.241351 | 0.0018 | 0.38 | -2.10 |
| alpha-Lytic | 20 | 31 | 1045 | 1 | 4905 | 4975.3282 | 80.7185 | 128,058 | 4974.320375 | 4 | 786 | 813 | KQIYKTPPIKDFGGFNFSQILPDPSKPS | 0.000000000000000020000000000000.0 | 801(NFS) | 1 | 3151.648 | 3148.659739 | 51370068 | 82.77 | 1 | 5 | 5 | 0 | 0 | 0 | 0 | 1 | 4974.320724 | -0.0003 | -0.07 | -2.05 |
| alpha-Lytic | 20 | 31 | 1189 | 1 | 4966 | 4876.2569 | 83.6682 | 110,493 | 4875.249075 | 4 | 786 | 813 | KQIYKTPPIKDFGGFNFSQILPDPSKPS | 0.000000000000000020000000000000.0 | 801(NFS) | 1 | 3151.648 | 3148.659739 | 51370068 | 82.77 | 1 | 5 | 3 | 0 | 0 | 1 | 0 | 1 | 4875.252309 | -0.0032 | -0.66 | 0.90 |
| alpha-Lytic | 20 | 31 | 1227 | 1 | 4886 | 5413.5002 | 79.8405 | 107,686 | 5412.492375 | 4 | 786 | 813 | KQIYKTPPIKDFGGFNFSQILPDPSKPS | 0.000000000000000020000000000000.0 | 801(NFS) | 1 | 3151.648 | 3148.659739 | 51370068 | 82.77 | 1 | 5 | 5 | 3 | 0 | 0 | 0 | 1 | 5412.494451 | -0.0021 | -0.38 | -2.93 |
| alpha-Lytic | 20 | 31 | 1243 | 1 | 4944 | 5225.3901 | 82.5743 | 106,034 | 5224.382275 | 4 | 786 | 813 | KQIYKTPPIKDFGGFNFSQILPDPSKPS | 0.000000000000000020000000000000.0 | 801(NFS) | 1 | 3151.648 | 3148.659739 | 51370068 | 82.77 | 1 | 5 | 4 | 1 | 0 | 1 | 0 | 1 | 5224.389591 | -0.0073 | -1.4 | -0.20 |
| alpha-Lytic | 20 | 31 | 1243 | 1 | 4944 | 5225.3901 | 82.5743 | 106,034 | 5224.382275 | 4 | 786 | 813 | KQIYKTPPIKDFGGFNFSQILPDPSKPS | 0.000000000000000020000000000000.0 | 801(NFS) | 1 | 3151.648 | 3148.659739 | 51370068 | 82.77 | 2 | 6 | 4 | 0 | 1 | 0 | 0 | 1 | 5224.389592 | -0.0073 | -1.4 | -0.20 |
| alpha-Lytic | 20 | 31 | 1265 | 1 | 4904 | 4813.2776 | 80.6712 | 103,559 | 4812.269775 | 4 | 786 | 813 | KQIYKTPPIKDFGGFNFSQILPDPSKPS | 0.000000000000000020000000000000.0 | 801(NFS) | 1 | 3151.648 | 3148.659739 | 51370068 | 82.77 | 1 | 4 | 5 | 0 | 0 | 0 | 0 | 1 | 4812.2679 | 0.0019 | 0.38 | -2.10 |
| alpha-Lytic | 20 | 31 | 1375 | 1 | 4887 | 5267.4493 | 79.8822 | 94,105 | 5266.441475 | 4 | 786 | 813 | KQIYKTPPIKDFGGFNFSQILPDPSKPS | 0.000000000000000020000000000000.0 | 801(NFS) | 1 | 3151.648 | 3148.659739 | 51370068 | 82.77 | 1 | 5 | 5 | 2 | 0 | 0 | 0 | 1 | 5266.436542 | 0.0049 | 0.93 | -2.89 |
| alpha-Lytic | 20 | 31 | 1567 | 1 | 4898 | 4797.2822 | 80.3886 | 81,963 | 4796.274375 | 4 | 786 | 813 | KQIYKTPPIKDFGGFNFSQILPDPSKPS | 0.000000000000000020000000000000.0 | 801(NFS) | 1 | 3151.648 | 3148.659739 | 51370068 | 82.77 | 1 | 3 | 5 | 1 | 0 | 0 | 0 | 1 | 4796.272985 | 0.0014 | 0.28 | -2.38 |
| alpha-Lytic | 20 | 31 | 1633 | 1 | 4962 | 5038.3124 | 83.457 | 76,785 | 5037.304575 | 4 | 786 | 813 | KQIYKTPPIKDFGGFNFSQILPDPSKPS | 0.000000000000000020000000000000.0 | 801(NFS) | 1 | 3151.648 | 3148.659739 | 51370068 | 82.77 | 1 | 6 | 3 | 0 | 0 | 1 | 0 | 1 | 5037.305133 | -0.0006 | -0.11 | 0.69 |
| alpha-Lytic | 20 | 31 | 1655 | 1 | 4958 | 5079.3349 | 83.2608 | 75,951 | 5078.327075 | 4 | 786 | 813 | KQIYKTPPIKDFGGFNFSQILPDPSKPS | 0.000000000000000020000000000000.0 | 801(NFS) | 1 | 3151.648 | 3148.659739 | 51370068 | 82.77 | 1 | 5 | 4 | 0 | 0 | 1 | 0 | 1 | 5078.331682 | -0.0046 | -0.9 | 0.49 |
| alpha-Lytic | 20 | 31 | 2771 | 1 | 4886 | 5283.4396 | 79.8405 | 35,603 | 5282.431775 | 4 | 786 | 813 | KQIYKTPPIKDFGGFNFSQILPDPSKPS | 0.000000000000000020000000000000.0 | 801(NFS) | 1 | 3151.648 | 3148.659739 | 51370068 | 82.77 | 1 | 6 | 5 | 1 | 0 | 0 | 0 | 1 | 5282.431457 | 0.0003 | 0.06 | -2.93 |
| alpha-Lytic | 20 | 31 | 2980 | 1 | 4965 | 4860.2639 | 83.6176 | 30,342 | 4859.256075 | 4 | 786 | 813 | KQIYKTPPIKDFGGFNFSQILPDPSKPS | 0.000000000000000020000000000000.0 | 801(NFS) | 1 | 3151.648 | 3148.659739 | 51370068 | 82.77 | 1 | 4 | 3 | 1 | 0 | 1 | 0 | 1 | 4859.257394 | -0.0013 | -0.27 | 0.85 |
| alpha-Lytic | 20 | 31 | 2980 | 1 | 4965 | 4860.2639 | 83.6176 | 30,342 | 4859.256075 | 4 | 786 | 813 | KQIYKTPPIKDFGGFNFSQILPDPSKPS | 0.000000000000000020000000000000.0 | 801(NFS) | 1 | 3151.648 | 3148.659739 | 51370068 | 82.77 | 2 | 5 | 3 | 0 | 1 | 0 | 0 | 1 | 4859.257395 | -0.0013 | -0.27 | 0.85 |
| alpha-Lytic | 20 | 31 | 3037 | 1 | 4886 | 5429.5012 | 79.8405 | 29,125 | 5428.493375 | 4 | 786 | 813 | KQIYKTPPIKDFGGFNFSQILPDPSKPS | 0.000000000000000020000000000000.0 | 801(NFS) | 1 | 3151.648 | 3148.659739 | 51370068 | 82.77 | 1 | 6 | 5 | 2 | 0 | 0 | 0 | 1 | 5428.489166 | 0.0040 | 0.73 | -2.93 |
| alpha-Lytic | 20 | 31 | 3040 | 1 | 4897 | 4756.2474 | 80.3425 | 29,043 | 4755.239575 | 4 | 786 | 813 | KQIYKTPPIKDFGGFNFSQILPDPSKPS | 0.000000000000000020000000000000.0 | 801(NFS) | 1 | 3151.648 | 3148.659739 | 51370068 | 82.77 | 1 | 4 | 4 | 1 | 0 | 0 | 0 | 1 | 4755.246436 | -0.0069 | -1.44 | -2.43 |
| alpha-Lytic | 20 | 31 | 3307 | 1 | 4884 | 5575.5644 | 79.7437 | 23,212 | 5574.556575 | 4 | 786 | 813 | KQIYKTPPIKDFGGFNFSQILPDPSKPS | 0.000000000000000020000000000000.0 | 801(NFS) | 1 | 3151.648 | 3148.659739 | 51370068 | 82.77 | 1 | 6 | 5 | 3 | 0 | 0 | 0 | 1 | 5574.547275 | 0.0093 | 1.66 | -3.03 |
| alpha-Lytic | 20 | 31 | 3538 | 1 | 4948 | 5209.3943 | 82.7615 | 18,248 | 5208.386475 | 4 | 786 | 813 | KQIYKTPPIKDFGGFNFSQILPDPSKPS | 0.000000000000000020000000000000.0 | 801(NFS) | 1 | 3151.648 | 3148.659739 | 51370068 | 82.77 | 1 | 4 | 4 | 0 | 0 | 1 | 0 | 1 | 5208.394676 | -0.0082 | -1.57 | -0.01 |
| alpha-Lytic | 20 | 31 | 3538 | 1 | 4948 | 5209.3943 | 82.7615 | 18,248 | 5208.386475 | 4 | 786 | 813 | KQIYKTPPIKDFGGFNFSQILPDPSKPS | 0.000000000000000020000000000000.0 | 801(NFS) | 1 | 3151.648 | 3148.659739 | 51370068 | 82.77 | 2 | 5 | 4 | 1 | 1 | 0 | 0 | 1 | 5208.394677 | -0.0082 | -1.57 | -0.01 |
| alpha-Lytic | 20 | 31 | 3788 | 1 | 4884 | 5721.6252 | 79.7437 | 12,692 | 5720.617375 | 4 | 786 | 813 | KQIYKTPPIKDFGGFNFSQILPDPSKPS | 0.000000000000000020000000000000.0 | 801(NFS) | 1 | 3151.648 | 3148.659739 | 51370068 | 82.77 | 1 | 6 | 5 | 4 | 0 | 0 | 0 | 0 | 5720.605184 | 0.0122 | 2.13 | -3.03 |
| alpha-Lytic | 20 | 31 | 3991 | 1 | 4941 | 5590.5247 | 82.4345 | 8,338 | 5589.516875 | 4 | 786 | 813 | KQIYKTPPIKDFGGFNFSQILPDPSKPS | 0.000000000000000020000000000000.0 | 801(NFS) | 1 | 3151.648 | 3148.659739 | 51370068 | 82.77 | 1 | 6 | 5 | 1 | 0 | 1 | 0 | 1 | 5589.521788 | -0.0049 | -0.87 | -0.34 |
| alpha-Lytic | 20 | 31 | 3991 | 1 | 4941 | 5590.5247 | 82.4345 | 8,338 | 5589.516875 | 4 | 786 | 813 | KQIYKTPPIKDFGGFNFSQILPDPSKPS | 0.000000000000000020000000000000.0 | 801(NFS) | 1 | 3151.648 | 3148.659739 | 51370068 | 82.77 | 2 | 7 | 5 | 0 | 1 | 0 | 0 | 0 | 5589.521789 | -0.0049 | -0.87 | -0.34 |
| alpha-Lytic | 89 | 9 | 651 | 1 | 4924 | 4157.899 | 81.6405 | 208,102 | 4156.891175 | 29 | 790 | 813 | KTPPIKDFGGFNFSQILPDPSKPS | 0.0000000000002000000000.0 | 801(NFS) | 1 | 2619.3471 | 2616.358839 | 3300966.5 | 83.97 | 1 | 7 | 2 | 0 | 0 | 0 | 0 | 1 | 4156.887353 | 0.0038 | 0.91 | -2.32 |
| alpha-Lytic | 89 | 9 | 1280 | 1 | 4927 | 3995.8395 | 81.7851 | 102,145 | 3994.831675 | 29 | 790 | 813 | KTPPIKDFGGFNFSQILPDPSKPS | 0.0000000000002000000000.0 | 801(NFS) | 1 | 2619.3471 | 2616.358839 | 3300966.5 | 83.97 | 1 | 6 | 2 | 0 | 0 | 0 | 0 | 1 | 3994.834529 | -0.0029 | -0.71 | -2.18 |
| alpha-Lytic | 89 | 9 | 1833 | 1 | 4923 | 4344.9713 | 81.5958 | 66,363 | 4343.963475 | 29 | 790 | 813 | KTPPIKDFGGFNFSQILPDPSKPS | 0.0000000000002000000000.0 | 801(NFS) | 1 | 2619.3471 | 2616.358839 | 3300966.5 | 83.97 | 1 | 6 | 3 | 1 | 0 | 0 | 0 | 1 | 4343.971811 | -0.0083 | -1.91 | -2.37 |
| alpha-Lytic | 89 | 9 | 2164 | 1 | 4930 | 3833.7863 | 81.9212 | 53,468 | 3832.778475 | 29 | 790 | 813 | KTPPIKDFGGFNFSQILPDPSKPS | 0.0000000000002000000000.0 | 801(NFS) | 1 | 2619.3471 | 2616.358839 | 3300966.5 | 83.97 | 1 | 5 | 2 | 0 | 0 | 0 | 0 | 1 | 3832.781705 | -0.0032 | -0.84 | -2.04 |
| alpha-Lytic | 89 | 9 | 2589 | 1 | 4926 | 4182.9259 | 81.7372 | 39,967 | 4181.918075 | 29 | 790 | 813 | KTPPIKDFGGFNFSQILPDPSKPS | 0.0000000000002000000000.0 | 801(NFS) | 1 | 2619.3471 | 2616.358839 | 3300966.5 | 83.97 | 1 | 5 | 3 | 1 | 0 | 0 | 0 | 1 | 4181.918987 | -0.0009 | -0.21 | -2.23 |
| alpha-Lytic | 89 | 9 | 2858 | 1 | 4925 | 4198.9186 | 81.686 | 33,162 | 4197.910775 | 29 | 790 | 813 | KTPPIKDFGGFNFSQILPDPSKPS | 0.0000000000002000000000.0 | 801(NFS) | 1 | 2619.3471 | 2616.358839 | 3300966.5 | 83.97 | 1 | 6 | 3 | 0 | 0 | 0 | 0 | 1 | 4197.913902 | -0.0031 | -0.74 | -2.28 |
| alpha-Lytic | 89 | 9 | 3018 | 1 | 4921 | 4319.9479 | 81.5023 | 29,628 | 4318.940075 | 29 | 790 | 813 | KTPPIKDFGGFNFSQILPDPSKPS | 0.0000000000002000000000.0 | 801(NFS) | 1 | 2619.3471 | 2616.358839 | 3300966.5 | 83.97 | 1 | 8 | 2 | 0 | 0 | 0 | 0 | 1 | 4318.940177 | -0.0001 | -0.02 | -2.46 |
| alpha-Lytic | 89 | 9 | 3783 | 1 | 4985 | 4506.006 | 84.5626 | 12,753 | 4504.998175 | 29 | 790 | 813 | KTPPIKDFGGFNFSQILPDPSKPS | 0.0000000000002000000000.0 | 801(NFS) | 1 | 2619.3471 | 2616.358839 | 3300966.5 | 83.97 | 1 | 6 | 3 | 0 | 0 | 1 | 0 | 1 | 4505.004233 | -0.0061 | -1.34 | 0.60 |
| alpha-Lytic | 89 | 9 | 3853 | 1 | 4989 | 4343.9538 | 84.7488 | 11,332 | 4342.945975 | 29 | 790 | 813 | KTPPIKDFGGFNFSQILPDPSKPS | 0.0000000000002000000000.0 | 801(NFS) | 1 | 2619.3471 | 2616.358839 | 3300966.5 | 83.97 | 1 | 5 | 3 | 0 | 0 | 1 | 0 | 1 | 4342.951409 | -0.0054 | -1.25 | 0.78 |
| alpha-Lytic | 132 | 4 | 3049 | 1 | 5020 | 4900.2941 | 86.1417 | 28,917 | 4899.286275 | 75 | 784 | 813 | QVKQIYKTPPIKDFGGFNFSQILPDPSKPS | 3.000000000000000002000000000000.0 | 801(NFS) | 1 | 3361.7484 | 3358.760139 | 1409069.1 | 88.00 | 1 | 7 | 2 | 0 | 0 | 0 | 0 | 1 | 4899.288653 | -0.0024 | -0.48 | -1.86 |
| alpha-Lytic | 132 | 4 | 3201 | 1 | 5022 | 4738.2399 | 86.2316 | 25,655 | 4737.232075 | 75 | 784 | 813 | QVKQIYKTPPIKDFGGFNFSQILPDPSKPS | 3.000000000000000002000000000000.0 | 801(NFS) | 1 | 3361.7484 | 3358.760139 | 1409069.1 | 88.00 | 1 | 6 | 2 | 0 | 0 | 0 | 0 | 1 | 4737.235829 | -0.0038 | -0.79 | -1.77 |
| alpha-Lytic | 132 | 4 | 3539 | 1 | 4928 | 4593.21 | 81.8305 | 18,235 | 4592.202175 | 75 | 784 | 813 | QVKQIYKTPPIKDFGGFNFSQILPDPSKPS | 3.000000000000000002000000000000.0 | 801(NFS) | 1 | 3361.7484 | 3358.760139 | 1409069.1 | 88.00 | | | | | | | | | | | | |
| alpha-Lytic | 132 | 4 | 3883 | 1 | 5024 | 4576.1828 | 86.3185 | 10,736 | 4575.174975 | 75 | 784 | 813 | QVKQIYKTPPIKDFGGFNFSQILPDPSKPS | 3.000000000000000002000000000000.0 | 801(NFS) | 1 | 3361.7484 | 3358.760139 | 1409069.1 | 88.00 | 1 | 5 | 2 | 0 | 0 | 0 | 0 | 1 | 4575.183005 | -0.0080 | -1.75 | -1.68 |
| Chymotrypsin | 1 | 104 | 4 | 1 | 3711 | 2310.9571 | 34.5332 | 35,849,000 | 2309.949275 | 7 | 1068 | 1075 | VPAQEKNF | 0.0000020.0 | 1074(NFT) | 1 | 934.4646 | 931.4763387 | 90297392 | 39.48 | 1 | 6 | 2 | 0 | 0 | 0 | 0 | 1 | 2309.952029 | -0.0028 | -1.19 | -4.95 |
| Chymotrypsin | 1 | 104 | 9 | 1 | 3720 | 2148.9051 | 34.9149 | 15,705,300 | 2147.897275 | 7 | 1068 | 1075 | VPAQEKNF | 0.0000020.0 | 1074(NFT) | 1 | 934.4646 | 931.4763387 | 90297392 | 39.48 | 1 | 5 | 2 | 0 | 0 | 0 | 0 | 1 | 2147.899205 | -0.0019 | -0.89 | -4.57 |
| Chymotrypsin | 1 | 104 | 21 | 1 | 3703 | 2660.0952 | 34.1926 | 9,562,620 | 2659.087375 | 7 | 1068 | 1075 | VPAQEKNF | 0.0000020.0 | 1074(NFT) | 1 | 934.4646 | 931.4763387 | 90297392 | 39.48 | 1 | 6 | 3 | 1 | 0 | 0 | 0 | 1 | 2659.089311 | -0.0019 | -0.72 | -5.29 |
| Chymotrypsin | 1 | 104 | 25 | 1 | 3703 | 2473.0108 | 34.1926 | 7,775,230 | 2472.002975 | 7 | 1068 | 1075 | VPAQEKNF | 0.0000020.0 | 1074(NFT) | 1 | 934.4646 | 931.4763387 | 90297392 | 39.48 | 1 | 7 | 2 | 0 | 0 | 0 | 0 | 1 | 2472.004853 | -0.0019 | -0.75 | -5.29 |
| Chymotrypsin | 1 | 104 | 27 | 1 | 3714 | 2514.0377 | 34.6605 | 7,309,110 | 2513.029875 | 7 | 1068 | 1075 | VPAQEKNF | 0.0000020.0 | 1074(NFT) | 1 | 934.4646 | 931.4763387 | 90297392 | 39.48 | 1 | 6 | 3 | 0 | 0 | 0 | 0 | 1 | 2513.031402 | -0.0015 | -0.6 | -4.82 |
| Chymotrypsin | 1 | 104 | 44 | 1 | 3700 | 2635.0641 | 34.0623 | 5,257,770 | 2634.056275 | 7 | 1068 | 1075 | VPAQEKNF | 0.0000020.0 | 1074(NFT) | 1 | 934.4646 | 931.4763387 | 90297392 | 39.48 | 1 | 8 | 2 | 0 | 0 | 0 | 0 | 1 | 2634.057677 | -0.0014 | -0.53 | -5.42 |
| Chymotrypsin | 1 | 104 | 50 | 1 | 3711 | 2498.0421 | 34.5332 | 5,171,680 | 2497.034275 | 7 | 1068 | 1075 | VPAQEKNF | 0.0000020.0 | 1074(NFT) | 1 | 934.4646 | 931.4763387 | 90297392 | 39.48 | 1 | 5 | 3 | 1 | 0 | 0 | 0 | 1 | 2497.036487 | -0.0022 | -0.88 | -4.95 |
| Chymotrypsin | 1 | 104 | 76 | 1 | 3723 | 2351.9851 | 35.04 | 3,908,620 | 2350.977275 | 7 | 1068 | 1075 | VPAQEKNF | 0.0000020.0 | 1074(NFT) | 1 | 934.4646 | 931.4763387 | 90297392 | 39.48 | 1 | 5 | 3 | 0 | 0 | 0 | 0 | 1 | 2350.978578 | -0.0013 | -0.55 | -4.44 |
| Chymotrypsin | 1 | 104 | 97 | 1 | 3736 | 2701.1222 | 35.5966 | 3,002,100 | 2700.114375 | 7 | 1068 | 1075 | VPAQEKNF | 0.0000020.0 | 1074(NFT) | 1 | 934.4646 | 931.4763387 | 90297392 | 39.48 | 1 | 5 | 4 | 1 | 0 | 0 | 0 | 1 | 2700.11586 | -0.0015 | -0.54 | -3.89 |
| Chymotrypsin | 1 | 104 | 128 | 1 | 3728 | 2644.1009 | 35.2515 | 2,440,280 | 2643.093075 | 7 | 1068 | 1075 | VPAQEKNF | 0.0000020.0 | 1074(NFT) | 1 | 934.4646 | 931.4763387 | 90297392 | 39.48 | 1 | 5 | 3 | 2 | 0 | 0 | 0 | 0 | 2643.094396 | -0.0013 | -0.49 | -4.23 |
| Chymotrypsin | 1 | 104 | 132 | 1 | 3723 | 2555.0645 | 35.04 | 2,381,230 | 2554.056675 | 7 | 1068 | 1075 | VPAQEKNF | 0.0000020.0 | 1074(NFT) | 1 | 934.4646 | 931.4763387 | 90297392 | 39.48 | 1 | 5 | 4 | 0 | 0 | 0 | 0 | 1 | 2554.057951 | -0.0013 | -0.49 | -4.44 |
| Chymotrypsin | 1 | 104 | 147 | 1 | 3738 | 2498.0425 | 35.6796 | 2,129,400 | 2497.034675 | 7 | 1068 | 1075 | VPAQEKNF | 0.0000020.0 | 1074(NFT) | 1 | 934.4646 | 931.4763387 | 90297392 | 39.48 | 1 | 5 | 3 | 1 | 0 | 0 | 0 | 1 | 2497.036487 | -0.0018 | -0.72 | -3.80 |
| Chymotrypsin | 1 | 104 | 153 | 1 | 3756 | 2742.1482 | 36.4537 | 2,056,520 | 2741.140375 | 7 | 1068 | 1075 | VPAQEKNF | 0.0000020.0 | 1074(NFT) | 1 | 934.4646 | 931.4763387 | 90297392 | 39.48 | 1 | 4 | 5 | 1 | 0 | 0 | 0 | 1 | 2741.142409 | -0.0020 | -0.74 | -3.03 |
| Chymotrypsin | 1 | 104 | 155 | 1 | 3718 | 3431.3868 | 34.8306 | 2,043,200 | 3430.378975 | 7 | 1068 | 1075 | VPAQEKNF | 0.0000020.0 | 1074(NFT) | 1 | 934.4646 | 931.4763387 | 90297392 | 39.48 | 1 | 7 | 6 | 1 | 0 | 0 | 0 | 1 | 3430.380254 | -0.0013 | -0.37 | -4.65 |
| Chymotrypsin | 1 | 104 | 205 | 1 | 3743 | 2596.0913 | 35.8951 | 1,571,670 | 2595.083475 | 7 | 1068 | 1075 | VPAQEKNF | 0.0000020.0 | 1074(NFT) | 1 | 934.4646 | 931.4763387 | 90297392 | 39.48 | 1 | 4 | 5 | 0 | 0 | 0 | 0 | 1 | 2595.0845 | -0.0010 | -0.39 | -3.59 |
| Chymotrypsin | 1 | 104 | 232 | 1 | 3754 | 2904.2008 | 36.3689 | 1,430,960 | 2903.192975 | 7 | 1068 | 1075 | VPAQEKNF | 0.0000020.0 | 1074(NFT) | 1 | 934.4646 | 931.4763387 | 90297392 | 39.48 | 1 | 5 | 5 | 1 | 0 | 0 | 0 | 1 | 2903.195233 | -0.0023 | -0.77 | -3.11 |
| Chymotrypsin | 1 | 104 | 285 | 1 | 3701 | 2847.1805 | 34.1056 | 1,209,730 | 2846.172675 | 7 | 1068 | 1075 | VPAQEKNF | 0.0000020.0 | 1074(NFT) | 1 | 934.4646 | 931.4763387 | 90297392 | 39.48 | 1 | 5 | 4 | 2 | 0 | 0 | 0 | 1 | 2846.173769 | -0.0011 | -0.38 | -5.38 |
| Chymotrypsin | 1 | 104 | 326 | 1 | 3711 | 2993.236 | 34.5332 | 1,064,110 | 2992.228175 | 7 | 1068 | 1075 | VPAQEKNF | 0.0000020.0 | 1074(NFT) | 1 | 934.4646 | 931.4763387 | 90297392 | 39.48 | 1 | 5 | 4 | 3 | 0 | 0 | 0 | 1 | 2992.231678 | -0.0035 | -1.17 | -4.95 |
| Chymotrypsin | 1 | 104 | 329 | 1 | 3737 | 3066.2539 | 35.6386 | 1,050,970 | 3065.246075 | 7 | 1068 | 1075 | VPAQEKNF | 0.0000020.0 | 1074(NFT) | 1 | 934.4646 | 931.4763387 | 90297392 | 39.48 | 1 | 6 | 5 | 1 | 0 | 0 | 0 | 1 | 3065.248057 | -0.0020 | -0.64 | -3.84 |
| Chymotrypsin | 1 | 104 | 346 | 1 | 3741 | 2758.1442 | 35.8092 | 992,520 | 2757.136375 | 7 | 1068 | 1075 | VPAQEKNF | 0.0000020.0 | 1074(NFT) | 1 | 934.4646 | 931.4763387 | 90297392 | 39.48 | 1 | 5 | 5 | 0 | 0 | 0 | 0 | 1 | 2757.137324 | -0.0009 | -0.34 | -3.67 |
| Chymotrypsin | 1 | 104 | 350 | 1 | 3763 | 3269.335 | 36.7509 | 982,277 | 3268.327175 | 7 | 1068 | 1075 | VPAQEKNF | 0.0000020.0 | 1074(NFT) | 1 | 934.4646 | 931.4763387 | 90297392 | 39.48 | 1 | 6 | 6 | 1 | 0 | 0 | 0 | 1 | 3268.32743 | -0.0003 | -0.07 | -2.73 |
| Chymotrypsin | 1 | 104 | 384 | 1 | 3717 | 2717.1181 | 34.7882 | 919,431 | 2716.110275 | 7 | 1068 | 1075 | VPAQEKNF | 0.0000020.0 | 1074(NFT) | 1 | 934.4646 | 931.4763387 | 90297392 | 39.48 | 1 | 6 | 4 | 0 | 0 | 0 | 0 | 1 | 2716.110775 | -0.0005 | -0.18 | -4.69 |
| Chymotrypsin | 1 | 104 | 394 | 1 | 3711 | 2701.1217 | 34.5332 | 886,983 | 2700.113875 | 7 | 1068 | 1075 | VPAQEKNF | 0.0000020.0 | 1074(NFT) | 1 | 934.4646 | 931.4763387 | 90297392 | 39.48 | 1 | 5 | 4 | 1 | 0 | 0 | 0 | 1 | 2700.11586 | -0.0020 | -0.73 | -4.95 |
| Chymotrypsin | 1 | 104 | 417 | 1 | 3725 | 2847.1792 | 35.1233 | 839,215 | 2846.171375 | 7 | 1068 | 1075 | VPAQEKNF | 0.0000020.0 | 1074(NFT) | 1 | 934.4646 | 931.4763387 | 90297392 | 39.48 | 1 | 5 | 4 | 2 | 0 | 0 | 0 | 1 | 2846.173769 | -0.0024 | -0.84 | -4.36 |
| Chymotrypsin | 1 | 104 | 450 | 1 | 3712 | 3577.4454 | 34.5742 | 790,430 | 3576.437575 | 7 | 1068 | 1075 | VPAQEKNF | 0.0000020.0 | 1074(NFT) | 1 | 934.4646 | 931.4763387 | 90297392 | 39.48 | 1 | 7 | 6 | 2 | 0 | 0 | 0 | 0 | 3576.438163 | -0.0006 | -0.16 | -4.91 |
| Chymotrypsin | 1 | 104 | 484 | 1 | 3757 | 3415.3923 | 36.4954 | 739,009 | 3414.384475 | 7 | 1068 | 1075 | VPAQEKNF | 0.0000020.0 | 1074(NFT) | 1 | 934.4646 | 931.4763387 | 90297392 | 39.48 | 1 | 6 | 6 | 2 | 0 | 0 | 0 | 1 | 3414.385339 | -0.0009 | -0.25 | -2.99 |
| Chymotrypsin | 1 | 104 | 488 | 1 | 3725 | 2189.9332 | 35.1233 | 730,917 | 2188.925375 | 7 | 1068 | 1075 | VPAQEKNF | 0.0000020.0 | 1074(NFT) | 1 | 934.4646 | 931.4763387 | 90297392 | 39.48 | 1 | 4 | 3 | 0 | 0 | 0 | 0 | 1 | 2188.925754 | -0.0004 | -0.17 | -4.36 |
| Chymotrypsin | 1 | 104 | 492 | 1 | 3727 | 1986.8532 | 35.2098 | 727,779 | 1985.845375 | 7 | 1068 | 1075 | VPAQEKNF | 0.0000020.0 | 1074(NFT) | 1 | 934.4646 | 931.4763387 | 90297392 | 39.48 | 1 | 4 | 2 | 0 | 0 | 0 | 0 | 1 | 1985.846381 | -0.0010 | -0.5 | -4.27 |
| Chymotrypsin | 1 | 104 | 502 | 1 | 3745 | 2377.0166 | 35.9834 | 707,779 | 2376.008775 | 7 | 1068 | 1075 | VPAQEKNF | 0.0000020.0 | 1074(NFT) | 1 | 934.4646 | 931.4763387 | 90297392 | 39.48 | 1 | 3 | 4 | 1 | 0 | 0 | 0 | 1 | 2376.010212 | -0.0014 | -0.6 | -3.50 |
| Chymotrypsin | 1 | 104 | 575 | 1 | 3703 | 3723.5028 | 34.1926 | 623,756 | 3722.494975 | 7 | 1068 | 1075 | VPAQEKNF | 0.0000020.0 | 1074(NFT) | 1 | 934.4646 | 931.4763387 | 90297392 | 39.48 | 1 | 7 | 6 | 3 | 0 | 0 | 0 | 0 | 3722.496072 | -0.0011 | -0.29 | -5.29 |
| Chymotrypsin | 1 | 104 | 633 | 1 | 3699 | 2797.1172 | 34.0205 | 572,013 | 2796.109375 | 7 | 1068 | 1075 | VPAQEKNF | 0.0000020.0 | 1074(NFT) | 1 | 934.4646 | 931.4763387 | 90297392 | 39.48 | 1 | 9 | 2 | 0 | 0 | 0 | 0 | 1 | 2796.110501 | -0.0011 | -0.4 | -5.46 |
| Chymotrypsin | 1 | 104 | 660 | 1 | 3705 | 3285.3332 | 34.2792 | 551,038 | 3284.325375 | 7 | 1068 | 1075 | VPAQEKNF | 0.0000020.0 | 1074(NFT) | 1 | 934.4646 | 931.4763387 | 90297392 | 39.48 | 1 | 7 | 6 | 0 | 0 | 0 | 0 | 1 | 3284.322345 | 0.0030 | 0.92 | -5.20 |
| Chymotrypsin | 1 | 104 | 714 | 1 | 3736 | 2783.1754 | 35.5966 | 514,155 | 2782.167575 | 7 | 1068 | 1075 | VPAQEKNF | 0.0000020.0 | 1074(NFT) | 1 | 934.4646 | 931.4763387 | 90297392 | 39.48 | 1 | 3 | 6 | 1 | 0 | 0 | 0 | 1 | 2782.168958 | -0.0014 | -0.49 | -3.89 |
| Chymotrypsin | 1 | 104 | 743 | 1 | 3717 | 2335.9895 | 34.7882 | 497,596 | 2334.981675 | 7 | 1068 | 1075 | VPAQEKNF | 0.0000020.0 | 1074(NFT) | 1 | 934.4646 | 931.4763387 | 90297392 | 39.48 | 1 | 4 | 3 | 1 | 0 | 0 | 0 | 1 | 2334.983663 | -0.0020 | -0.85 | -4.69 |
| Chymotrypsin | 1 | 104 | 761 | 1 | 3725 | 2920.1967 | 35.1233 | 486,176 | 2919.188875 | 7 | 1068 | 1075 | VPAQEKNF | 0.0000020.0 | 1074(NFT) | 1 | 934.4646 | 931.4763387 | 90297392 | 39.48 | 1 | 6 | 5 | 0 | 0 | 0 | 0 | 1 | 2919.190148 | -0.0013 | -0.43 | -4.36 |
| Chymotrypsin | 1 | 104 | 875 | 1 | 3730 | 2027.8805 | 35.3364 | 426,338 | 2026.872675 | 7 | 1068 | 1075 | VPAQEKNF | 0.0000020.0 | 1074(NFT) | 1 | 934.4646 | 931.4763387 | 90297392 | 39.48 | 1 | 3 | 3 | 0 | 0 | 0 | 0 | 1 | 2026.87293 | -0.0003 | -0.12 | -4.15 |
| Chymotrypsin | 1 | 104 | 897 | 1 | 3779 | 2821.1287 | 37.4363 | 420,015 | 2820.120875 | 7 | 1068 | 1075 | VPAQEKNF | 0.0000020.0 | 1074(NFT) | 1 | 934.4646 | 931.4763387 | 90297392 | 39.48 | 1 | 6 | 3 | 0 | 0 | 1 | 0 | 1 | 2820.121733 | -0.0009 | -0.3 | -2.05 |
| Chymotrypsin | 1 | 104 | 909 | 1 | 3726 | 3634.4674 | 35.1674 | 413,155 | 3633.459575 | 7 | 1068 | 1075 | VPAQEKNF | 0.0000020.0 | 1074(NFT) | 1 | 934.4646 | 931.4763387 | 90297392 | 39.48 | 1 | 7 | 7 | 1 | 0 | 0 | 0 | 1 | 3633.459627 | -0.0001 | -0.01 | -4.32 |
| Chymotrypsin | 1 | 104 | 913 | 1 | 3697 | 3869.5602 | 33.9356 | 411,861 | 3868.552375 | 7 | 1068 | 1075 | VPAQEKNF | 0.0000020.0 | 1074(NFT) | 1 | 934.4646 | 931.4763387 | 90297392 | 39.48 | 1 | 7 | 6 | 4 | 0 | 0 | 0 | 1 | 3868.553981 | -0.0016 | -0.41 | -5.55 |
| Chymotrypsin | 1 | 104 | 913 | 1 | 3697 | 3869.5602 | 33.9356 | 411,861 | 3868.552375 | 7 | 1068 | 1075 | VPAQEKNF | 0.0000020.0 | 1074(NFT) | 1 | 934.4646 | 931.4763387 | 90297392 | 39.48 | 2 | 2 | 10 | 0 | 2 | 0 | 1 | -1 | 3868.566551 | -0.0142 | -3.66 | -5.55 |
| Chymotrypsin | 1 | 104 | 926 | 1 | 3743 | 2335.9903 | 35.8951 | 407,337 | 2334.982475 | 7 | 1068 | 1075 | VPAQEKNF | 0.0000020.0 | 1074(NFT) | 1 | 934.4646 | 931.4763387 | 90297392 | 39.48 | 1 | 4 | 3 | 1 | 0 | 0 | 0 | 1 | 2334.983663 | -0.0012 | -0.5 | -3.59 |
| Chymotrypsin | 1 | 104 | 987 | 1 | 3720 | 3780.5222 | 34.9149 | 387,185 | 3779.514375 | 7 | 1068 | 1075 | VPAQEKNF | 0.0000020.0 | 1074(NFT) | 1 | 934.4646 | 931.4763387 | 90297392 | 39.48 | 1 | 7 | 7 | 2 | 0 | 0 | 0 | 0 | 3779.517536 | -0.0032 | -0.83 | -4.57 |
| Chymotrypsin | 1 | 104 | 993 | 1 | 3718 | 2806.1514 | 34.8306 | 383,257 | 2805.143575 | 7 | 1068 | 1075 | VPAQEKNF | 0.0000020.0 | 1074(NFT) | 1 | 934.4646 | 931.4763387 | 90297392 | 39.48 | 1 | 6 | 3 | 2 | 0 | 0 | 0 | 0 | 2805.14722 | -0.0036 | -1.29 | -4.65 |
| Chymotrypsin | 1 | 104 | 1039 | 1 | 3742 | 2539.0701 | 35.8511 | 364,032 | 2538.062275 | 7 | 1068 | 1075 | VPAQEKNF | 0.0000020.0 | 1074(NFT) | 1 | 934.4646 | 931.4763387 | 90297392 | 39.48 | 1 | 4 | 4 | 1 | 0 | 0 | 0 | 1 | 2538.063036 | -0.0008 | -0.29 | -3.63 |
| Chymotrypsin | 1 | 104 | 1085 | 1 | 3703 | 3504.4299 | 34.1926 | 352,081 | 3503.422075 | 7 | 1068 | 1075 | VPAQEKNF | 0.0000020.0 | 1074(NFT) | 1 | 934.4646 | 931.4763387 | 90297392 | 39.48 | 1 | 6 | 5 | 4 | 0 | 0 | 0 | 0 | 3503.421784 | 0.0003 | 0.08 | -5.29 |
| Chymotrypsin | 1 | 104 | 1143 | 1 | 3734 | 2888.2057 | 35.5106 | 333,383 | 2887.197875 | 7 | 1068 | 1075 | VPAQEKNF | 0.0000020.0 | 1074(NFT) | 1 | 934.4646 | 931.4763387 | 90297392 | 39.48 | 1 | 4 | 5 | 2 | 0 | 0 | 0 | 1 | 2887.200318 | -0.0024 | -0.84 | -3.97 |
| Chymotrypsin | 1 | 104 | 1150 | 1 | 3731 | 3212.3121 | 35.3805 | 332,131 | 3211.304275 | 7 | 1068 | 1075 | VPAQEKNF | 0.0000020.0 | 1074(NFT) | 1 | 934.4646 | 931.4763387 | 90297392 | 39.48 | 1 | 6 | 5 | 2 | 0 | 0 | 0 | 1 | 3211.305966 | -0.0017 | -0.52 | -4.10 |
| Chymotrypsin | 1 | 104 | 1235 | 1 | 3746 | 2580.0975 | 36.0261 | 309,611 | 2579.089675 | 7 | 1068 | 1075 | VPAQEKNF | 0.0000020.0 | 1074(NFT) | 1 | 934.4646 | 931.4763387 | 90297392 | 39.48 | 1 | 3 | 5 | 1 | 0 | 0 | 0 | 1 | 2579.089585 | 0.0001 | 0.03 | -3.46 |
| Chymotrypsin | 1 | 104 | 1294 | 1 | 3697 | 3431.3877 | 33.9356 | 295,345 | 3430.379875 | 7 | 1068 | 1075 | VPAQEKNF | 0.0000020.0 | 1074(NFT) | 1 | 934.4646 | 931.4763387 | 90297392 | 39.48 | 1 | 7 | 6 | 1 | 0 | 0 | 0 | 1 | 3430.380254 | -0.0004 | -0.11 | -5.55 |
| Chymotrypsin | 1 | 104 | 1309 | 1 | 3725 | 2904.2039 | 35.1233 | 292,027 | 2903.196075 | 7 | 1068 | 1075 | VPAQEKNF | 0.0000020.0 | 1074(NFT) | 1 | 934.4646 | 931.4763387 | 90297392 | 39.48 | 1 | 5 | 5 | 1 | 0 | 0 | 0 | 1 | 2903.195233 | 0.0008 | 0.29 | -4.36 |
| Chymotrypsin | 1 | 104 | 1398 | 1 | 3728 | 2393.0113 | 35.2515 | 276,266 | 2392.003475 | 7 | 1068 | 1075 | VPAQEKNF | 0.0000020.0 | 1074(NFT) | 1 | 934.4646 | 931.4763387 | 90297392 | 39.48 | 1 | 4 | 4 | 0 | 0 | 0 | 0 | 1 | 2392.005127 | -0.0017 | -0.69 | -4.23 |
| Chymotrypsin | 1 | 104 | 1433 | 1 | 3744 | 3269.3335 | 35.9388 | 267,059 | 3268.325675 | 7 | 1068 | 1075 | VPAQEKNF | 0.0000020.0 | 1074(NFT) | 1 | 934.4646 | 931.4763387 | 90297392 | 39.48 | 1 | 6 | 6 | 1 | 0 | 0 | 0 | 1 | 3268.32743 | -0.0018 | -0.53 | -3.54 |
| Chymotrypsin | 1 | 104 | 1451 | 1 | 3750 | 3123.2754 | 36.1973 | 264,345 | 3122.267575 | 7 | 1068 | 1075 | VPAQEKNF | 0.0000020.0 | 1074(NFT) | 1 | 934.4646 | 931.4763387 | 90297392 | 39.48 | 1 | 6 | 6 | 0 | 0 | 0 | 0 | 1 | 3122.269521 | -0.0019 | -0.62 | -3.29 |
| Chymotrypsin | 1 | 104 | 1454 | 1 | 3706 | 2920.203 | 34.321 | 264,135 | 2919.195175 | 7 | 1068 | 1075 | VPAQEKNF | 0.0000020.0 | 1074(NFT) | 1 | 934.4646 | 931.4763387 | 90297392 | 39.48 | 1 | 6 | 5 | 0 | 0 | 0 | 0 | 1 | 2919.190148 | 0.0050 | 1.72 | -5.16 |
| Chymotrypsin | 1 | 104 | 1476 | 1 | 3716 | 2879.1676 | 34.7441 | 261,173 | 2878.159775 | 7 | 1068 | 1075 | VPAQEKNF | 0.0000020.0 | 1074(NFT) | 1 | 934.4646 | 931.4763387 | 90297392 | 39.48 | 1 | 7 | 4 | 0 | 0 | 0 | 0 | 1 | 2878.163599 | -0.0038 | -1.32 | -4.74 |
| Chymotrypsin | 1 | 104 | 1571 | 1 | 3738 | 3050.2592 | 35.6796 | 246,475 | 3049.251375 | 7 | 1068 | 1075 | VPAQEKNF | 0.0000020.0 | 1074(NFT) | 1 | 934.4646 | 931.4763387 | 90297392 | 39.48 | 1 | 5 | 5 | 2 | 0 | 0 | 0 | 1 | 3049.253142 | -0.0018 | -0.57 | -3.80 |
| Chymotrypsin | 1 | 104 | 1627 | 1 | 3735 | 2482.0457 | 35.5533 | 239,300 | 2481.037875 | 7 | 1068 | 1075 | VPAQEKNF | 0.0000020.0 | 1074(NFT) | 1 | 934.4646 | 931.4763387 | 90297392 | 39.48 | 1 | 4 | 3 | 2 | 0 | 0 | 0 | 1 | 2481.041572 | -0.0037 | -1.48 | -3.93 |
| Chymotrypsin | 1 | 104 | 1640 | 1 | 3719 | 3358.3706 | 34.8717 | 237,214 | 3357.362775 | 7 | 1068 | 1075 | VPAQEKNF | 0.0000020.0 | 1074(NFT) | 1 | 934.4646 | 931.4763387 | 90297392 | 39.48 | 1 | 6 | 5 | 3 | 0 | 0 | 0 | 1 | 3357.363875 | -0.0011 | -0.32 | -4.61 |
| Chymotrypsin | 1 | 104 | 1647 | 1 | 3691 | 4015.6183 | 33.6788 | 235,872 | 4014.610475 | 7 | 1068 | 1075 | VPAQEKNF | 0.0000020.0 | 1074(NFT) | 1 | 934.4646 | 931.4763387 | 90297392 | 39.48 | 1 | 2 | 10 | 1 | 2 | 0 | 1 | -1 | 4014.62446 | -0.0140 | -3.48 | -5.80 |
| Chymotrypsin | 1 | 104 | 1678 | 1 | 3785 | 2659.0799 | 37.6934 | 232,308 | 2658.072075 | 7 | 1068 | 1075 | VPAQEKNF | 0.0000020.0 | 1074(NFT) | 1 | 934.4646 | 931.4763387 | 90297392 | 39.48 | 1 | 5 | 3 | 0 | 0 | 1 | 0 | 1 | 2658.068909 | 0.0032 | 1.19 | -1.79 |
| Chymotrypsin | 1 | 104 | 1751 | 1 | 3692 | 3358.3718 | 33.7235 | 222,045 | 3357.363975 | 7 | 1068 | 1075 | VPAQEKNF | 0.0000020.0 | 1074(NFT) | 1 | 934.4646 | 931.4763387 | 90297392 | 39.48 | 1 | 6 | 5 | 3 | 0 | 0 | 0 | 1 | 3357.363875 | 0.0001 | 0.02 | -5.76 |
| Chymotrypsin | 1 | 104 | 1789 | 1 | 3765 | 3107.2815 | 36.8357 | 218,338 | 3106.274575 | 7 | 1068 | 1075 | VPAQEKNF | 0.0000020.0 | 1074(NFT) | 1 | 934.4646 | 931.4763387 | 90297392 | 39.48 | 1 | 5 | 6 | 1 | 0 | 0 | 0 | 1 | 3106.274606 | -0.0009 | -0.29 | -2.65 |
| Chymotrypsin | 1 | 104 | 1814 | 1 | 3718 | 3269.3338 | 34.8306 | 215,715 | 3268.325975 | 7 | 1068 | 1075 | VPAQEKNF | 0.0000020.0 | 1074(NFT) | 1 | 934.4646 | 931.4763387 | 90297392 | 39.48 | 1 | 6 | 6 | 1 | 0 | 0 | 0 | 1 | 3268.32743 | -0.0015 | -0.44 | -4.65 |
| Chymotrypsin | 1 | 104 | 1842 | 1 | 3746 | 2173.938 | 36.0261 | 212,016 | 2172.930175 | 7 | 1068 | 1075 | VPAQEKNF | 0.0000020.0 | 1074(NFT) | 1 | 934.4646 | 931.4763387 | 90297392 | 39.48 | 1 | 3 | 3 | 1 | 0 | 0 | 0 | 1 | 2172.930839 | -0.0007 | -0.3 | -3.46 |
| Chymotrypsin | 1 | 104 | 1921 | 1 | 3723 | 2742.1501 | 35.04 | 202,131 | 2741.142275 | 7 | 1068 | 1075 | VPAQEKNF | 0.0000020.0 | 1074(NFT) | 1 | 934.4646 | 931.4763387 | 90297392 | 39.48 | 1 | 4 | 5 | 1 | 0 | 0 | 0 | 1 | 2741.142409 | -0.0001 | -0.04 | -4.44 |
| Chymotrypsin | 1 | 104 | 2100 | 1 | 3720 | 3107.2811 | 34.9149 | 186,268 | 3106.273275 | 7 | 1068 | 1075 | VPAQEKNF | 0.0000020.0 | 1074(NFT) | 1 | 934.4646 | 931.4763387 | 90297392 | 39.48 | 1 | 5 | 6 | 1 | 0 | 0 | 0 | 1 | 3106.274606 | -0.0013 | -0.42 | -4.57 |
| Chymotrypsin | 1 | 104 | 2222 | 1 | 3735 | 2945.2275 | 35.5533 | 176,330 | 2944.219675 | 7 | 1068 | 1075 | VPAQEKNF | 0.0000020.0 | 1074(NFT) | 1 | 934.4646 | 931.4763387 | 90297392 | 39.48 | 1 | 4 | 6 | 1 | 0 | 0 | 0 | 1 | 2944.221782 | -0.0021 | -0.71 | -3.93 |
| Chymotrypsin | 1 | 104 | 2321 | 1 | 3736 | 3561.4508 | 35.5966 | 167,429 | 3560.442975 | 7 | 1068 | 1075 | VPAQEKNF | 0.0000020.0 | 1074(NFT) | 1 | 934.4646 | 931.4763387 | 90297392 | 39.48 | 1 | 6 | 6 | 3 | 0 | 0 | 0 | 1 | 3560.443248 | -0.0003 | -0.07 | -3.89 |
| Chymotrypsin | 1 | 104 | 2346 | 1 | 3777 | 3738.4757 | 37.3532 | 165,616 | 3737.467875 | 7 | 1068 | 1075 | VPAQEKNF | 0.0000020.0 | 1074(NFT) | 1 | 934.4646 | 931.4763387 | 90297392 | 39.48 | 1 | 7 | 6 | 1 | 0 | 1 | 0 | 1 | 3737.470585 | -0.0027 | -0.72 | -2.13 |
| Chymotrypsin | 1 | 104 | 2346 | 1 | 3777 | 3738.4757 | 37.3532 | 165,616 | 3737.467875 | 7 | 1068 | 1075 | VPAQEKNF | 0.0000020.0 | 1074(NFT) | 1 | 934.4646 | 931.4763387 | 90297392 | 39.48 | 2 | 8 | 6 | 0 | 1 | 0 | 0 | 1 | 3737.470586 | -0.0027 | -0.72 | -2.13 |
| Chymotrypsin | 1 | 104 | 2438 | 1 | 3691 | 3577.4425 | 33.6788 | 159,224 | 3576.434675 | 7 | 1068 | 1075 | VPAQEKNF | 0.0000020.0 | 1074(NFT) | 1 | 934.4646 | 931.4763387 | 90297392 | 39.48 | 1 | 7 | 6 | 2 | 0 | 0 | 0 | 0 | 3576.438163 | -0.0035 | -0.97 | -5.80 |
| Chymotrypsin | 1 | 104 | 2472 | 1 | 3702 | 2863.1817 | 34.1484 | 157,067 | 2862.173875 | 7 | 1068 | 1075 | VPAQEKNF | 0.0000020.0 | 1074(NFT) | 1 | 934.4646 | 931.4763387 | 90297392 | 39.48 | 1 | 6 | 4 | 0 | 0 | 0 | 0 | 1 | 2862.168684 | 0.0052 | 1.81 | -5.33 |

| | | | | | | | | | | | | | | | | | | | | | | | | | | | | | | | | |
|---|---|---|---|---|---|---|---|---|---|---|---|---|---|---|---|---|---|---|---|---|---|---|---|---|---|---|---|---|---|---|---|---|
| Chymotrypsin | 1 | 104 | 2528 | 1 | 3684 | 3723.5068 | 33.3755 | 153,374 | 3722.498975 | 7 | 1068 | 1075 | VPAQEKNF | 0.00000020.0 | 1074(NFT) | 1 | 934.4646 | 931.4763387 | 90297392 | 39.48 | 1 | 7 | 6 | 3 | 0 | 0 | 0 | 0 | 3722.496072 | 0.0029 | 0.77 | -6.11 |
| Chymotrypsin | 1 | 104 | 2651 | 1 | 3705 | 3050.2599 | 34.2792 | 146,062 | 3049.252075 | 7 | 1068 | 1075 | VPAQEKNF | 0.00000020.0 | 1074(NFT) | 1 | 934.4646 | 931.4763387 | 90297392 | 39.48 | 1 | 5 | 5 | 2 | 0 | 0 | 0 | 1 | 3049.253142 | -0.0011 | -0.34 | -5.20 |
| Chymotrypsin | 1 | 104 | 2724 | 1 | 3760 | 3253.341 | 36.6218 | 141,463 | 3252.333175 | 7 | 1068 | 1075 | VPAQEKNF | 0.00000020.0 | 1074(NFT) | 1 | 934.4646 | 931.4763387 | 90297392 | 39.48 | 1 | 5 | 6 | 2 | 0 | 0 | 0 | 1 | 3252.332515 | 0.0007 | 0.2 | -2.86 |
| Chymotrypsin | 1 | 104 | 3042 | 1 | 3728 | 3472.4128 | 35.2515 | 124,706 | 3471.404975 | 7 | 1068 | 1075 | VPAQEKNF | 0.00000020.0 | 1074(NFT) | 1 | 934.4646 | 931.4763387 | 90297392 | 39.48 | 1 | 6 | 7 | 1 | 0 | 0 | 0 | 1 | 3471.406803 | -0.0018 | -0.52 | -4.23 |
| Chymotrypsin | 1 | 104 | 3052 | 1 | 3738 | 2863.1795 | 35.6796 | 124,306 | 2862.171675 | 7 | 1068 | 1075 | VPAQEKNF | 0.00000020.0 | 1074(NFT) | 1 | 934.4646 | 931.4763387 | 90297392 | 39.48 | 1 | 6 | 4 | 1 | 0 | 0 | 0 | 1 | 2862.168684 | 0.0030 | 1.04 | -3.80 |
| Chymotrypsin | 1 | 104 | 3074 | 1 | 3799 | 2805.1283 | 38.2909 | 123,448 | 2804.120475 | 7 | 1068 | 1075 | VPAQEKNF | 0.00000020.0 | 1074(NFT) | 1 | 934.4646 | 931.4763387 | 90297392 | 39.48 | 1 | 6 | 3 | 0 | 1 | 0 | 0 | 1 | 2804.126819 | -0.0063 | -2.26 | -1.19 |
| Chymotrypsin | 1 | 104 | 3074 | 1 | 3799 | 2805.1283 | 38.2909 | 123,448 | 2804.120475 | 7 | 1068 | 1075 | VPAQEKNF | 0.00000020.0 | 1074(NFT) | 1 | 934.4646 | 931.4763387 | 90297392 | 39.48 | 2 | 5 | 3 | 1 | 0 | 1 | 0 | 0 | 2804.126818 | -0.0063 | -2.26 | -1.19 |
| Chymotrypsin | 1 | 104 | 3137 | 1 | 3761 | 2580.0936 | 36.6644 | 120,451 | 2579.085775 | 7 | 1068 | 1075 | VPAQEKNF | 0.00000020.0 | 1074(NFT) | 1 | 934.4646 | 931.4763387 | 90297392 | 39.48 | 1 | 3 | 5 | 1 | 0 | 0 | 0 | 1 | 2579.089585 | -0.0038 | -1.47 | -2.82 |
| Chymotrypsin | 1 | 104 | 3269 | 1 | 3788 | 2805.1339 | 37.8213 | 115,032 | 2804.126075 | 7 | 1068 | 1075 | VPAQEKNF | 0.00000020.0 | 1074(NFT) | 1 | 934.4646 | 931.4763387 | 90297392 | 39.48 | 1 | 6 | 3 | 0 | 1 | 0 | 0 | 1 | 2804.126819 | -0.0007 | -0.26 | -1.66 |
| Chymotrypsin | 1 | 104 | 3269 | 1 | 3788 | 2805.1339 | 37.8213 | 115,032 | 2804.126075 | 7 | 1068 | 1075 | VPAQEKNF | 0.00000020.0 | 1074(NFT) | 1 | 934.4646 | 931.4763387 | 90297392 | 39.48 | 2 | 5 | 3 | 1 | 0 | 1 | 0 | 0 | 2804.126818 | -0.0007 | -0.26 | -1.66 |
| Chymotrypsin | 1 | 104 | 3402 | 1 | 3706 | 3926.5843 | 34.321 | 110,439 | 3925.576475 | 7 | 1068 | 1075 | VPAQEKNF | 0.00000020.0 | 1074(NFT) | 1 | 934.4646 | 931.4763387 | 90297392 | 39.48 | 1 | 7 | 7 | 3 | 0 | 0 | 0 | 0 | 3925.575445 | 0.0010 | 0.26 | -5.16 |
| Chymotrypsin | 1 | 104 | 3509 | 1 | 3723 | 3415.393 | 35.04 | 106,822 | 3414.385175 | 7 | 1068 | 1075 | VPAQEKNF | 0.00000020.0 | 1074(NFT) | 1 | 934.4646 | 931.4763387 | 90297392 | 39.48 | 1 | 6 | 6 | 2 | 0 | 0 | 0 | 1 | 3414.385339 | -0.0002 | -0.04 | -4.44 |
| Chymotrypsin | 1 | 104 | 3607 | 1 | 3697 | 3066.2587 | 33.9356 | 103,614 | 3065.250875 | 7 | 1068 | 1075 | VPAQEKNF | 0.00000020.0 | 1074(NFT) | 1 | 934.4646 | 931.4763387 | 90297392 | 39.48 | 1 | 6 | 5 | 1 | 0 | 0 | 0 | 1 | 3065.248057 | 0.0028 | 0.91 | -5.55 |
| Chymotrypsin | 1 | 104 | 3778 | 1 | 3794 | 3008.2125 | 38.0744 | 97,795 | 3007.204675 | 7 | 1068 | 1075 | VPAQEKNF | 0.00000020.0 | 1074(NFT) | 1 | 934.4646 | 931.4763387 | 90297392 | 39.48 | 1 | 5 | 4 | 1 | 0 | 1 | 0 | 1 | 3007.206191 | -0.0015 | -0.5 | -1.41 |
| Chymotrypsin | 1 | 104 | 3778 | 1 | 3794 | 3008.2125 | 38.0744 | 97,795 | 3007.204675 | 7 | 1068 | 1075 | VPAQEKNF | 0.00000020.0 | 1074(NFT) | 1 | 934.4646 | 931.4763387 | 90297392 | 39.48 | 2 | 6 | 4 | 0 | 1 | 0 | 0 | 1 | 3007.206192 | -0.0015 | -0.5 | -1.41 |
| Chymotrypsin | 1 | 104 | 3903 | 1 | 3722 | 3618.4706 | 34.9983 | 94,556 | 3617.462775 | 7 | 1068 | 1075 | VPAQEKNF | 0.00000020.0 | 1074(NFT) | 1 | 934.4646 | 931.4763387 | 90297392 | 39.48 | 1 | 6 | 7 | 2 | 0 | 0 | 0 | 0 | 3617.464712 | -0.0019 | -0.53 | -4.48 |
| Chymotrypsin | 1 | 104 | 4047 | 1 | 3783 | 2830.173 | 37.6075 | 90,602 | 2829.165175 | 7 | 1068 | 1075 | VPAQEKNF | 0.00000020.0 | 1074(NFT) | 1 | 934.4646 | 931.4763387 | 90297392 | 39.48 | 1 | 4 | 4 | 1 | 1 | 0 | 0 | 1 | 2829.158453 | 0.0067 | 2.37 | -1.88 |
| Chymotrypsin | 1 | 104 | 4047 | 1 | 3783 | 2830.173 | 37.6075 | 90,602 | 2829.165175 | 7 | 1068 | 1075 | VPAQEKNF | 0.00000020.0 | 1074(NFT) | 1 | 934.4646 | 931.4763387 | 90297392 | 39.48 | 2 | 3 | 4 | 2 | 0 | 1 | 0 | 0 | 2829.158452 | 0.0067 | 2.37 | -1.88 |
| Chymotrypsin | 1 | 104 | 4235 | 1 | 3680 | 3869.5656 | 33.2082 | 85,513 | 3868.557775 | 7 | 1068 | 1075 | VPAQEKNF | 0.00000020.0 | 1074(NFT) | 1 | 934.4646 | 931.4763387 | 90297392 | 39.48 | 1 | 7 | 6 | 4 | 0 | 0 | 0 | 1 | 3868.553981 | 0.0038 | 0.98 | -6.27 |
| Chymotrypsin | 1 | 104 | 4235 | 1 | 3680 | 3869.5656 | 33.2082 | 85,513 | 3868.557775 | 7 | 1068 | 1075 | VPAQEKNF | 0.00000020.0 | 1074(NFT) | 1 | 934.4646 | 931.4763387 | 90297392 | 39.48 | 2 | 2 | 10 | 0 | 2 | 0 | 1 | -1 | 3868.566551 | -0.0088 | -2.26 | -6.27 |
| Chymotrypsin | 1 | 104 | 4362 | 1 | 3785 | 3008.2072 | 37.6934 | 82,457 | 3007.199375 | 7 | 1068 | 1075 | VPAQEKNF | 0.00000020.0 | 1074(NFT) | 1 | 934.4646 | 931.4763387 | 90297392 | 39.48 | 1 | 5 | 4 | 1 | 0 | 1 | 0 | 1 | 3007.206191 | -0.0068 | -2.26 | -1.79 |
| Chymotrypsin | 1 | 104 | 4362 | 1 | 3785 | 3008.2072 | 37.6934 | 82,457 | 3007.199375 | 7 | 1068 | 1075 | VPAQEKNF | 0.00000020.0 | 1074(NFT) | 1 | 934.4646 | 931.4763387 | 90297392 | 39.48 | 2 | 6 | 4 | 0 | 1 | 0 | 0 | 1 | 3007.206192 | -0.0068 | -2.26 | -1.79 |
| Chymotrypsin | 1 | 104 | 4596 | 1 | 3709 | 3123.2754 | 34.4489 | 77,053 | 3122.267575 | 7 | 1068 | 1075 | VPAQEKNF | 0.00000020.0 | 1074(NFT) | 1 | 934.4646 | 931.4763387 | 90297392 | 39.48 | 1 | 6 | 6 | 0 | 0 | 0 | 0 | 1 | 3122.269521 | -0.0019 | -0.62 | -5.03 |
| Chymotrypsin | 1 | 104 | 4778 | 1 | 3770 | 3884.5283 | 37.0538 | 73,261 | 3883.520475 | 7 | 1068 | 1075 | VPAQEKNF | 0.00000020.0 | 1074(NFT) | 1 | 934.4646 | 931.4763387 | 90297392 | 39.48 | 1 | 8 | 6 | 1 | 1 | 0 | 0 | 1 | 3883.528495 | -0.0080 | -2.06 | -2.43 |
| Chymotrypsin | 1 | 104 | 4778 | 1 | 3770 | 3884.5283 | 37.0538 | 73,261 | 3883.520475 | 7 | 1068 | 1075 | VPAQEKNF | 0.00000020.0 | 1074(NFT) | 1 | 934.4646 | 931.4763387 | 90297392 | 39.48 | 2 | 7 | 6 | 2 | 0 | 1 | 0 | 0 | 3883.528494 | -0.0080 | -2.06 | -2.43 |
| Chymotrypsin | 1 | 104 | 4871 | 1 | 3697 | 2822.1478 | 33.9356 | 71,293 | 2821.139975 | 7 | 1068 | 1075 | VPAQEKNF | 0.00000020.0 | 1074(NFT) | 1 | 934.4646 | 931.4763387 | 90297392 | 39.48 | 1 | 7 | 3 | 1 | 0 | 0 | 0 | 0 | 2821.142135 | -0.0022 | -0.76 | -5.55 |
| Chymotrypsin | 1 | 104 | 4924 | 1 | 3725 | 3155.2925 | 35.1233 | 70,147 | 3154.284675 | 7 | 1068 | 1075 | VPAQEKNF | 0.00000020.0 | 1074(NFT) | 1 | 934.4646 | 931.4763387 | 90297392 | 39.48 | 1 | 6 | 4 | 3 | 0 | 0 | 0 | 1 | 3154.284502 | 0.0002 | 0.05 | -4.36 |
| Chymotrypsin | 1 | 104 | 5074 | 1 | 3755 | 2961.2144 | 36.4125 | 67,099 | 2960.206575 | 7 | 1068 | 1075 | VPAQEKNF | 0.00000020.0 | 1074(NFT) | 1 | 934.4646 | 931.4763387 | 90297392 | 39.48 | 1 | 5 | 6 | 0 | 0 | 0 | 0 | 1 | 2960.216697 | -0.0101 | -3.41 | -3.07 |
| Chymotrypsin | 1 | 104 | 5100 | 1 | 3795 | 3373.3421 | 38.1172 | 66,703 | 3372.334275 | 7 | 1068 | 1075 | VPAQEKNF | 0.00000020.0 | 1074(NFT) | 1 | 934.4646 | 931.4763387 | 90297392 | 39.48 | 1 | 6 | 5 | 1 | 0 | 1 | 0 | 1 | 3372.338388 | -0.0041 | -1.21 | -1.37 |
| Chymotrypsin | 1 | 104 | 5100 | 1 | 3795 | 3373.3421 | 38.1172 | 66,703 | 3372.334275 | 7 | 1068 | 1075 | VPAQEKNF | 0.00000020.0 | 1074(NFT) | 1 | 934.4646 | 931.4763387 | 90297392 | 39.48 | 2 | 7 | 5 | 0 | 1 | 0 | 0 | 0 | 3372.338389 | -0.0041 | -1.21 | -1.37 |
| Chymotrypsin | 1 | 104 | 5521 | 1 | 3816 | 3576.4303 | 39.0186 | 59,807 | 3575.422475 | 7 | 1068 | 1075 | VPAQEKNF | 0.00000020.0 | 1074(NFT) | 1 | 934.4646 | 931.4763387 | 90297392 | 39.48 | 1 | 7 | 6 | 0 | 1 | 0 | 0 | 1 | 3575.417762 | 0.0047 | 1.31 | -0.46 |
| Chymotrypsin | 1 | 104 | 5521 | 1 | 3816 | 3576.4303 | 39.0186 | 59,807 | 3575.422475 | 7 | 1068 | 1075 | VPAQEKNF | 0.00000020.0 | 1074(NFT) | 1 | 934.4646 | 931.4763387 | 90297392 | 39.48 | 2 | 6 | 6 | 1 | 0 | 1 | 0 | 0 | 3575.417761 | 0.0047 | 1.31 | -0.46 |
| Chymotrypsin | 1 | 104 | 5675 | 1 | 3733 | 3009.2329 | 35.4682 | 57,317 | 3008.225075 | 7 | 1068 | 1075 | VPAQEKNF | 0.00000020.0 | 1074(NFT) | 1 | 934.4646 | 931.4763387 | 90297392 | 39.48 | 1 | 6 | 4 | 2 | 0 | 0 | 0 | 1 | 3008.226593 | -0.0015 | -0.5 | -4.01 |
| [illegible] | | | | | | | | | | | | | | | | | | | | | | | | | | | | | | | | |

| | | | | | | | | | | | | | | | | | | | | | | | | | | | | | | | | |
|---|---|---|---|---|---|---|---|---|---|---|---|---|---|---|---|---|---|---|---|---|---|---|---|---|---|---|---|---|---|---|---|---|
| Trypsin | 2 | 152 | 1879 | 1 | 3951 | 3485.4422 | 40.0664 | 70,928 | 3484.434375 | 14 | 1074 | 1086 | NFTTAPAICHDGK | 0.2000000000000.0 | 1074(NFT) | 1 | 1433.6495 | 1430.661239 | 11165932 | 45.93 | 1 | 3 | 7 | 1 | 0 | 0 | 0 | 1 | 3484.433231 | 0.0011 | 0.32 | -5.87 |
| Trypsin | 2 | 152 | 1928 | 1 | 3966 | 3282.3611 | 40.7927 | 68,793 | 3281.353275 | 14 | 1074 | 1086 | NFTTAPAICHDGK | 0.2000000000000.0 | 1074(NFT) | 1 | 1433.6495 | 1430.661239 | 11165932 | 45.93 | 1 | 3 | 6 | 1 | 0 | 0 | 0 | 1 | 3281.353858 | -0.0006 | -0.17 | -5.14 |
| Trypsin | 2 | 152 | 1930 | 1 | 3929 | 3914.5817 | 39.0403 | 68,715 | 3913.573875 | 14 | 1074 | 1086 | NFTTAPAICHDGK | 0.2000000000000.0 | 1074(NFT) | 1 | 1433.6495 | 1430.661239 | 11165932 | 45.93 | 1 | 6 | 6 | 2 | 0 | 0 | 0 | 1 | 3913.570239 | 0.0036 | 0.92 | -6.89 |
| Trypsin | 2 | 152 | 1939 | 1 | 3919 | 3752.4959 | 38.5433 | 68,189 | 3751.488075 | 14 | 1074 | 1086 | NFTTAPAICHDGK | 0.2000000000000.0 | 1074(NFT) | 1 | 1433.6495 | 1430.661239 | 11165932 | 45.93 | | | | | | | | | | | | |
| Trypsin | 2 | 152 | 1983 | 1 | 3902 | 4149.647 | 37.7549 | 65,920 | 4148.639175 | 14 | 1074 | 1086 | NFTTAPAICHDGK | 0.2000000000000.0 | 1074(NFT) | 1 | 1433.6495 | 1430.661239 | 11165932 | 45.93 | 1 | 8 | 7 | 0 | 0 | 0 | 0 | 1 | 4148.639442 | -0.0003 | -0.06 | -8.18 |
| Trypsin | 2 | 152 | 2018 | 1 | 4009 | 3361.3407 | 42.9004 | 64,807 | 3360.332875 | 14 | 1074 | 1086 | NFTTAPAICHDGK | 0.2000000000000.0 | 1074(NFT) | 1 | 1433.6495 | 1430.661239 | 11165932 | 45.93 | 1 | 5 | 4 | 0 | 0 | 1 | 0 | 1 | 3360.333182 | -0.0003 | -0.09 | -3.03 |
| Trypsin | 2 | 152 | 2035 | 1 | 4022 | 3142.2638 | 43.5734 | 64,115 | 3141.255975 | 14 | 1074 | 1086 | NFTTAPAICHDGK | 0.2000000000000.0 | 1074(NFT) | 1 | 1433.6495 | 1430.661239 | 11165932 | 45.93 | 1 | 4 | 3 | 1 | 0 | 1 | 0 | 1 | 3141.258894 | -0.0029 | -0.92 | -2.36 |
| Trypsin | 2 | 152 | 2035 | 1 | 4022 | 3142.2638 | 43.5734 | 64,115 | 3141.255975 | 14 | 1074 | 1086 | NFTTAPAICHDGK | 0.2000000000000.0 | 1074(NFT) | 1 | 1433.6495 | 1430.661239 | 11165932 | 45.93 | 2 | 5 | 3 | 0 | 1 | 0 | 0 | 1 | 3141.258895 | -0.0029 | -0.92 | -2.36 |
| Trypsin | 2 | 152 | 2119 | 1 | 4023 | 3288.3222 | 43.6148 | 60,990 | 3287.314375 | 14 | 1074 | 1086 | NFTTAPAICHDGK | 0.2000000000000.0 | 1074(NFT) | 1 | 1433.6495 | 1430.661239 | 11165932 | 45.93 | 1 | 4 | 3 | 2 | 0 | 1 | 0 | 0 | 3287.316803 | -0.0024 | -0.73 | -2.32 |
| Trypsin | 2 | 152 | 2119 | 1 | 4023 | 3288.3222 | 43.6148 | 60,990 | 3287.314375 | 14 | 1074 | 1086 | NFTTAPAICHDGK | 0.2000000000000.0 | 1074(NFT) | 1 | 1433.6495 | 1430.661239 | 11165932 | 45.93 | 2 | 5 | 3 | 1 | 1 | 0 | 0 | 0 | 3287.316804 | -0.0024 | -0.73 | -2.32 |
| Trypsin | 2 | 152 | 2148 | 1 | 3934 | 3524.4159 | 39.2675 | 60,305 | 3523.408075 | 14 | 1074 | 1086 | NFTTAPAICHDGK | 0.2000000000000.0 | 1074(NFT) | 1 | 1433.6495 | 1430.661239 | 11165932 | 45.93 | 1 | 7 | 4 | 1 | 0 | 0 | 0 | 1 | 3523.406408 | 0.0017 | 0.47 | -6.67 |
| Trypsin | 2 | 152 | 2237 | 1 | 4004 | 4018.587 | 42.6557 | 57,547 | 4017.579175 | 14 | 1074 | 1086 | NFTTAPAICHDGK | 0.2000000000000.0 | 1074(NFT) | 1 | 1433.6495 | 1430.661239 | 11165932 | 45.93 | 1 | 6 | 5 | 2 | 0 | 1 | 0 | 1 | 4017.581197 | -0.0020 | -0.5 | -3.28 |
| Trypsin | 2 | 152 | 2237 | 1 | 4004 | 4018.587 | 42.6557 | 57,547 | 4017.579175 | 14 | 1074 | 1086 | NFTTAPAICHDGK | 0.2000000000000.0 | 1074(NFT) | 1 | 1433.6495 | 1430.661239 | 11165932 | 45.93 | 2 | 7 | 5 | 1 | 1 | 0 | 0 | 1 | 4017.581198 | -0.0020 | -0.5 | -3.28 |
| Trypsin | 2 | 152 | 2250 | 1 | 3916 | 4060.6086 | 38.402 | 57,080 | 4059.600775 | 14 | 1074 | 1086 | NFTTAPAICHDGK | 0.2000000000000.0 | 1074(NFT) | 1 | 1433.6495 | 1430.661239 | 11165932 | 45.93 | | | | | | | | | | | | |
| Trypsin | 2 | 152 | 2300 | 1 | 3934 | 3460.4081 | 39.2675 | 55,377 | 3459.400275 | 14 | 1074 | 1086 | NFTTAPAICHDGK | 0.2000000000000.0 | 1074(NFT) | 1 | 1433.6495 | 1430.661239 | 11165932 | 45.93 | 1 | 5 | 6 | 0 | 0 | 0 | 0 | 1 | 3459.401597 | -0.0013 | -0.38 | -6.67 |
| Trypsin | 2 | 152 | 2385 | 1 | 3932 | 3670.4685 | 39.1839 | 52,998 | 3669.460675 | 14 | 1074 | 1086 | NFTTAPAICHDGK | 0.2000000000000.0 | 1074(NFT) | 1 | 1433.6495 | 1430.661239 | 11165932 | 45.93 | 1 | 7 | 4 | 2 | 0 | 0 | 0 | 1 | 3669.464317 | -0.0036 | -0.99 | -6.75 |
| Trypsin | 2 | 152 | 2398 | 1 | 4001 | 4221.6694 | 42.5066 | 52,550 | 4220.661575 | 14 | 1074 | 1086 | NFTTAPAICHDGK | 0.2000000000000.0 | 1074(NFT) | 1 | 1433.6495 | 1430.661239 | 11165932 | 45.93 | 1 | 7 | 6 | 1 | 1 | 0 | 0 | 1 | 4220.660571 | 0.0010 | 0.23 | -3.43 |
| Trypsin | 2 | 152 | 2398 | 1 | 4001 | 4221.6694 | 42.5066 | 52,550 | 4220.661575 | 14 | 1074 | 1086 | NFTTAPAICHDGK | 0.2000000000000.0 | 1074(NFT) | 1 | 1433.6495 | 1430.661239 | 11165932 | 45.93 | 2 | 6 | 6 | 2 | 0 | 1 | 0 | 0 | 4220.66057 | 0.0010 | 0.23 | -3.43 |
| Trypsin | 2 | 152 | 2488 | 1 | 3973 | 3143.287 | 41.1282 | 50,444 | 3142.279175 | 14 | 1074 | 1086 | NFTTAPAICHDGK | 0.2000000000000.0 | 1074(NFT) | 1 | 1433.6495 | 1430.661239 | 11165932 | 45.93 | 1 | 5 | 3 | 2 | 0 | 0 | 0 | 0 | 3142.279296 | -0.0001 | -0.03 | -4.81 |
| Trypsin | 2 | 152 | 2519 | 1 | 3956 | 3930.5665 | 40.3122 | 49,724 | 3929.558675 | 14 | 1074 | 1086 | NFTTAPAICHDGK | 0.2000000000000.0 | 1074(NFT) | 1 | 1433.6495 | 1430.661239 | 11165932 | 45.93 | 1 | 7 | 6 | 1 | 0 | 0 | 0 | 1 | 3929.565154 | -0.0065 | -1.64 | -5.62 |
| Trypsin | 2 | 152 | 2552 | 1 | 3961 | 2933.2254 | 40.5637 | 48,947 | 2932.217575 | 14 | 1074 | 1086 | NFTTAPAICHDGK | 0.2000000000000.0 | 1074(NFT) | 1 | 1433.6495 | 1430.661239 | 11165932 | 45.93 | 1 | 3 | 5 | 0 | 0 | 0 | 0 | 1 | 2932.216576 | 0.0010 | 0.34 | -5.37 |
| Trypsin | 2 | 152 | 2591 | 1 | 3918 | 4514.8042 | 38.4982 | 47,791 | 4513.796375 | 14 | 1074 | 1086 | NFTTAPAICHDGK | 0.2000000000000.0 | 1074(NFT) | 1 | 1433.6495 | 1430.661239 | 11165932 | 45.93 | 1 | 2 | 10 | 1 | 2 | 0 | 1 | -1 | 4513.80936 | -0.0130 | -2.87 | -7.44 |
| Trypsin | 2 | 152 | 2645 | 1 | 3951 | 3873.5515 | 40.0664 | 46,683 | 3872.543675 | 14 | 1074 | 1086 | NFTTAPAICHDGK | 0.2000000000000.0 | 1074(NFT) | 1 | 1433.6495 | 1430.661239 | 11165932 | 45.93 | 1 | 7 | 5 | 2 | 0 | 0 | 0 | 0 | 3872.54369 | 0.0000 | 0 | -5.87 |
| Trypsin | 2 | 152 | 2699 | 1 | 3929 | 3752.5239 | 39.0403 | 45,622 | 3751.516075 | 14 | 1074 | 1086 | NFTTAPAICHDGK | 0.2000000000000.0 | 1074(NFT) | 1 | 1433.6495 | 1430.661239 | 11165932 | 45.93 | 1 | 5 | 6 | 2 | 0 | 0 | 0 | 1 | 3751.517415 | -0.0013 | -0.35 | -6.89 |
| Trypsin | 2 | 152 | 2769 | 1 | 3933 | 3321.3373 | 39.2257 | 43,953 | 3320.329475 | 14 | 1074 | 1086 | NFTTAPAICHDGK | 0.2000000000000.0 | 1074(NFT) | 1 | 1433.6495 | 1430.661239 | 11165932 | 45.93 | 1 | 7 | 3 | 1 | 0 | 0 | 0 | 0 | 3320.327035 | 0.0024 | 0.73 | -6.71 |
| Trypsin | 2 | 152 | 2779 | 1 | 3981 | 3241.3363 | 41.5409 | 43,803 | 3240.328475 | 14 | 1074 | 1086 | NFTTAPAICHDGK | 0.2000000000000.0 | 1074(NFT) | 1 | 1433.6495 | 1430.661239 | 11165932 | 45.93 | 1 | 4 | 5 | 1 | 0 | 0 | 0 | 1 | 3240.327309 | 0.0012 | 0.35 | -4.39 |
| Trypsin | 2 | 152 | 2820 | 1 | 4014 | 3466.3703 | 43.1518 | 42,834 | 3465.362475 | 14 | 1074 | 1086 | NFTTAPAICHDGK | 0.2000000000000.0 | 1074(NFT) | 1 | 1433.6495 | 1430.661239 | 11165932 | 45.93 | 1 | 6 | 3 | 1 | 0 | 1 | 0 | 1 | 3465.364542 | -0.0021 | -0.59 | -2.78 |
| Trypsin | 2 | 152 | 2820 | 1 | 4014 | 3466.3703 | 43.1518 | 42,834 | 3465.362475 | 14 | 1074 | 1086 | NFTTAPAICHDGK | 0.2000000000000.0 | 1074(NFT) | 1 | 1433.6495 | 1430.661239 | 11165932 | 45.93 | 2 | 7 | 3 | 0 | 1 | 0 | 1 | -1 | 3465.364543 | -0.0021 | -0.59 | -2.78 |
| Trypsin | 2 | 152 | 2858 | 1 | 3997 | 4164.6357 | 42.317 | 41,717 | 4163.627875 | 14 | 1074 | 1086 | NFTTAPAICHDGK | 0.2000000000000.0 | 1074(NFT) | 1 | 1433.6495 | 1430.661239 | 11165932 | 45.93 | 1 | 6 | 5 | 3 | 0 | 1 | 0 | 1 | 4163.639106 | -0.0112 | -2.69 | -3.62 |
| Trypsin | 2 | 152 | 2858 | 1 | 3997 | 4164.6357 | 42.317 | 41,717 | 4163.627875 | 14 | 1074 | 1086 | NFTTAPAICHDGK | 0.2000000000000.0 | 1074(NFT) | 1 | 1433.6495 | 1430.661239 | 11165932 | 45.93 | 2 | 7 | 5 | 2 | 1 | 0 | 0 | 0 | 4163.639107 | -0.0112 | -2.69 | -3.62 |
| Trypsin | 2 | 152 | 3015 | 1 | 3955 | 3419.3822 | 40.2648 | 39,231 | 3418.374375 | 14 | 1074 | 1086 | NFTTAPAICHDGK | 0.2000000000000.0 | 1074(NFT) | 1 | 1433.6495 | 1430.661239 | 11165932 | 45.93 | 1 | 6 | 5 | 0 | 0 | 0 | 0 | 1 | 3418.375048 | -0.0007 | -0.19 | -5.67 |
| Trypsin | 2 | 152 | 3112 | 1 | 4012 | 3856.5337 | 43.0514 | 37,623 | 3855.525875 | 14 | 1074 | 1086 | NFTTAPAICHDGK | 0.2000000000000.0 | 1074(NFT) | 1 | 1433.6495 | 1430.661239 | 11165932 | 45.93 | 1 | 6 | 5 | 1 | 1 | 0 | 0 | 1 | 3855.528374 | -0.0025 | -0.64 | -2.88 |
| Trypsin | 2 | 152 | 3112 | 1 | 4012 | 3856.5337 | 43.0514 | 37,623 | 3855.525875 | 14 | 1074 | 1086 | NFTTAPAICHDGK | 0.2000000000000.0 | 1074(NFT) | 1 | 1433.6495 | 1430.661239 | 11165932 | 45.93 | 2 | 5 | 5 | 2 | 0 | 1 | 0 | 0 | 3855.528373 | -0.0025 | -0.64 | -2.88 |
| Trypsin | 2 | 152 | 3126 | 1 | 4016 | 3491.4041 | 43.2877 | 37,465 | 3490.396275 | 14 | 1074 | 1086 | NFTTAPAICHDGK | 0.2000000000000.0 | 1074(NFT) | 1 | 1433.6495 | 1430.661239 | 11165932 | 45.93 | 1 | 5 | 4 | 1 | 1 | 0 | 0 | 1 | 3490.396177 | 0.0001 | 0.02 | -2.65 |
| Trypsin | 2 | 152 | 3126 | 1 | 4016 | 3491.4041 | 43.2877 | 37,465 | 3490.396275 | 14 | 1074 | 1086 | NFTTAPAICHDGK | 0.2000000000000.0 | 1074(NFT) | 1 | 1433.6495 | 1430.661239 | 11165932 | 45.93 | 2 | 4 | 4 | 2 | 0 | 1 | 0 | 1 | 3490.396176 | 0.0001 | 0.02 | -2.65 |
| Trypsin | 2 | 152 | 3211 | 1 | 3989 | 4529.774 | 41.9449 | 36,109 | 4528.766175 | 14 | 1074 | 1086 | NFTTAPAICHDGK | 0.2000000000000.0 | 1074(NFT) | 1 | 1433.6495 | 1430.661239 | 11165932 | 45.93 | 1 | 7 | 6 | 3 | 0 | 1 | 0 | 0 | 4528.771303 | -0.0051 | -1.13 | -3.99 |
| Trypsin | 2 | 152 | 3211 | 1 | 3989 | 4529.774 | 41.9449 | 36,109 | 4528.766175 | 14 | 1074 | 1086 | NFTTAPAICHDGK | 0.2000000000000.0 | 1074(NFT) | 1 | 1433.6495 | 1430.661239 | 11165932 | 45.93 | 2 | 8 | 6 | 2 | 1 | 0 | 0 | 0 | 4528.771304 | -0.0051 | -1.13 | -3.99 |
| Trypsin | 2 | 152 | 3274 | 1 | 3997 | 4091.6066 | 42.317 | 34,975 | 4090.598775 | 14 | 1074 | 1086 | NFTTAPAICHDGK | 0.2000000000000.0 | 1074(NFT) | 1 | 1433.6495 | 1430.661239 | 11165932 | 45.93 | 1 | 7 | 6 | 0 | 0 | 1 | 0 | 1 | 4090.597576 | 0.0012 | 0.29 | -3.62 |
| Trypsin | 2 | 152 | 3336 | 1 | 3904 | 4222.6663 | 37.855 | 34,043 | 4221.658475 | 14 | 1074 | 1086 | NFTTAPAICHDGK | 0.2000000000000.0 | 1074(NFT) | 1 | 1433.6495 | 1430.661239 | 11165932 | 45.93 | | | | | | | | | | | | |
| Trypsin | 2 | 152 | 3343 | 1 | 3950 | 4060.6318 | 40.0233 | 33,975 | 4059.623975 | 14 | 1074 | 1086 | NFTTAPAICHDGK | 0.2000000000000.0 | 1074(NFT) | 1 | 1433.6495 | 1430.661239 | 11165932 | 45.93 | 1 | 6 | 6 | 3 | 0 | 0 | 0 | 1 | 4059.628148 | -0.0042 | -1.02 | -5.91 |
| Trypsin | 2 | 152 | 3371 | 1 | 3978 | 3403.3824 | 41.3915 | 33,575 | 3402.374575 | 14 | 1074 | 1086 | NFTTAPAICHDGK | 0.2000000000000.0 | 1074(NFT) | 1 | 1433.6495 | 1430.661239 | 11165932 | 45.93 | 1 | 5 | 5 | 1 | 0 | 0 | 0 | 1 | 3402.380133 | -0.0056 | -1.63 | -4.54 |
| Trypsin | 2 | 152 | 3406 | 1 | 3937 | 3727.4883 | 39.4087 | 33,160 | 3726.480475 | 14 | 1074 | 1086 | NFTTAPAICHDGK | 0.2000000000000.0 | 1074(NFT) | 1 | 1433.6495 | 1430.661239 | 11165932 | 45.93 | 1 | 7 | 5 | 1 | 0 | 0 | 0 | 1 | 3726.485781 | -0.0053 | -1.42 | -6.53 |
| Trypsin | 2 | 152 | 3412 | 1 | 3972 | 3054.2444 | 41.0823 | 33,038 | 3053.236575 | 14 | 1074 | 1086 | NFTTAPAICHDGK | 0.2000000000000.0 | 1074(NFT) | 1 | 1433.6495 | 1430.661239 | 11165932 | 45.93 | 1 | 5 | 4 | 0 | 0 | 0 | 0 | 1 | 3053.242851 | -0.0063 | -2.05 | -4.85 |
| Trypsin | 2 | 152 | 3475 | 1 | 3975 | 2851.1715 | 41.2324 | 32,196 | 2850.163675 | 14 | 1074 | 1086 | NFTTAPAICHDGK | 0.2000000000000.0 | 1074(NFT) | 1 | 1433.6495 | 1430.661239 | 11165932 | 45.93 | 1 | 5 | 3 | 0 | 0 | 0 | 0 | 1 | 2850.163478 | 0.0002 | 0.06 | -4.70 |
| Trypsin | 2 | 152 | 3667 | 1 | 3955 | 3727.4978 | 40.2648 | 29,845 | 3726.489975 | 14 | 1074 | 1086 | NFTTAPAICHDGK | 0.2000000000000.0 | 1074(NFT) | 1 | 1433.6495 | 1430.661239 | 11165932 | 45.93 | 1 | 7 | 5 | 1 | 0 | 0 | 0 | 1 | 3726.485781 | 0.0042 | 1.12 | -5.67 |
| Trypsin | 2 | 152 | 3736 | 1 | 3982 | 3200.3016 | 41.5986 | 28,993 | 3199.293775 | 14 | 1074 | 1086 | NFTTAPAICHDGK | 0.2000000000000.0 | 1074(NFT) | 1 | 1433.6495 | 1430.661239 | 11165932 | 45.93 | 1 | 5 | 4 | 1 | 0 | 0 | 0 | 1 | 3199.30076 | -0.0070 | -2.18 | -4.34 |
| Trypsin | 2 | 152 | 3761 | 1 | 3916 | 3914.5488 | 38.402 | 28,795 | 3913.540975 | 14 | 1074 | 1086 | NFTTAPAICHDGK | 0.2000000000000.0 | 1074(NFT) | 1 | 1433.6495 | 1430.661239 | 11165932 | 45.93 | | | | | | | | | | | | |
| Trypsin | 2 | 152 | 3804 | 1 | 3996 | 4440.7492 | 42.2728 | 28,232 | 4439.741375 | 14 | 1074 | 1086 | NFTTAPAICHDGK | 0.2000000000000.0 | 1074(NFT) | 1 | 1433.6495 | 1430.661239 | 11165932 | 45.93 | 1 | 8 | 7 | 0 | 1 | 0 | 0 | 1 | 4439.734859 | 0.0065 | 1.46 | -3.66 |
| Trypsin | 2 | 152 | 3804 | 1 | 3996 | 4440.7492 | 42.2728 | 28,232 | 4439.741375 | 14 | 1074 | 1086 | NFTTAPAICHDGK | 0.2000000000000.0 | 1074(NFT) | 1 | 1433.6495 | 1430.661239 | 11165932 | 45.93 | 2 | 7 | 7 | 1 | 0 | 1 | 0 | 0 | 4439.734858 | 0.0065 | 1.46 | -3.66 |
| Trypsin | 2 | 152 | 3808 | 1 | 4017 | 3345.3398 | 43.3405 | 28,204 | 3344.331975 | 14 | 1074 | 1086 | NFTTAPAICHDGK | 0.2000000000000.0 | 1074(NFT) | 1 | 1433.6495 | 1430.661239 | 11165932 | 45.93 | 1 | 4 | 4 | 1 | 0 | 1 | 0 | 1 | 3344.338267 | -0.0063 | -1.88 | -2.59 |
| Trypsin | 2 | 152 | 3808 | 1 | 4017 | 3345.3398 | 43.3405 | 28,204 | 3344.331975 | 14 | 1074 | 1086 | NFTTAPAICHDGK | 0.2000000000000.0 | 1074(NFT) | 1 | 1433.6495 | 1430.661239 | 11165932 | 45.93 | 2 | 5 | 4 | 0 | 1 | 0 | 0 | 1 | 3344.338268 | -0.0063 | -1.88 | -2.59 |
| Trypsin | 2 | 152 | 4038 | 1 | 4010 | 3710.4796 | 42.9471 | 25,720 | 3709.471775 | 14 | 1074 | 1086 | NFTTAPAICHDGK | 0.2000000000000.0 | 1074(NFT) | 1 | 1433.6495 | 1430.661239 | 11165932 | 45.93 | 1 | 6 | 5 | 0 | 1 | 0 | 0 | 1 | 3709.470465 | 0.0013 | 0.35 | -2.99 |
| Trypsin | 2 | 152 | 4038 | 1 | 4010 | 3710.4796 | 42.9471 | 25,720 | 3709.471775 | 14 | 1074 | 1086 | NFTTAPAICHDGK | 0.2000000000000.0 | 1074(NFT) | 1 | 1433.6495 | 1430.661239 | 11165932 | 45.93 | 2 | 5 | 5 | 1 | 0 | 1 | 0 | 1 | 3709.470464 | 0.0013 | 0.35 | -2.99 |
| Trypsin | 2 | 152 | 4117 | 1 | 3947 | 3647.4909 | 39.8786 | 24,787 | 3646.483075 | 14 | 1074 | 1086 | NFTTAPAICHDGK | 0.2000000000000.0 | 1074(NFT) | 1 | 1433.6495 | 1430.661239 | 11165932 | 45.93 | 1 | 4 | 7 | 1 | 0 | 0 | 0 | 1 | 3646.486055 | -0.0030 | -0.81 | -6.06 |
| Trypsin | 2 | 152 | 4160 | 1 | 3950 | 4076.618 | 40.0233 | 24,314 | 4075.610175 | 14 | 1074 | 1086 | NFTTAPAICHDGK | 0.2000000000000.0 | 1074(NFT) | 1 | 1433.6495 | 1430.661239 | 11165932 | 45.93 | 1 | 7 | 6 | 2 | 0 | 0 | 0 | 0 | 4075.623063 | -0.0129 | -3.16 | -5.91 |
| Trypsin | 2 | 152 | 4194 | 1 | 4020 | 4059.6185 | 43.4816 | 24,086 | 4058.610675 | 14 | 1074 | 1086 | NFTTAPAICHDGK | 0.2000000000000.0 | 1074(NFT) | 1 | 1433.6495 | 1430.661239 | 11165932 | 45.93 | 1 | 6 | 6 | 1 | 1 | 0 | 0 | 1 | 4058.607747 | 0.0029 | 0.72 | -2.45 |
| Trypsin | 2 | 152 | 4194 | 1 | 4020 | 4059.6185 | 43.4816 | 24,086 | 4058.610675 | 14 | 1074 | 1086 | NFTTAPAICHDGK | 0.2000000000000.0 | 1074(NFT) | 1 | 1433.6495 | 1430.661239 | 11165932 | 45.93 | 2 | 5 | 6 | 2 | 0 | 1 | 0 | 0 | 4058.607746 | 0.0029 | 0.72 | -2.45 |
| Trypsin | 2 | 152 | 4428 | 1 | 4074 | 4179.6156 | 46.0307 | 22,021 | 4178.607775 | 14 | 1074 | 1086 | NFTTAPAICHDGK | 0.2000000000000.0 | 1074(NFT) | 1 | 1433.6495 | 1430.661239 | 11165932 | 45.93 | 1 | 6 | 5 | 1 | 0 | 2 | 0 | 1 | 4178.613619 | -0.0058 | -1.39 | 0.10 |
| Trypsin | 2 | 152 | 4428 | 1 | 4074 | 4179.6156 | 46.0307 | 22,021 | 4178.607775 | 14 | 1074 | 1086 | NFTTAPAICHDGK | 0.2000000000000.0 | 1074(NFT) | 1 | 1433.6495 | 1430.661239 | 11165932 | 45.93 | 2 | 7 | 5 | 0 | 1 | 1 | 0 | 0 | 4178.61362 | -0.0058 | -1.39 | 0.10 |
| Trypsin | 2 | 152 | 4499 | 1 | 3965 | 3346.3667 | 40.748 | 21,391 | 3345.358875 | 14 | 1074 | 1086 | NFTTAPAICHDGK | 0.2000000000000.0 | 1074(NFT) | 1 | 1433.6495 | 1430.661239 | 11165932 | 45.93 | 1 | 5 | 4 | 2 | 0 | 0 | 0 | 1 | 3345.358669 | 0.0002 | 0.06 | -5.19 |
| Trypsin | 2 | 152 | 4663 | 1 | 3996 | 4367.721 | 42.2728 | 20,135 | 4366.713175 | 14 | 1074 | 1086 | NFTTAPAICHDGK | 0.2000000000000.0 | 1074(NFT) | 1 | 1433.6495 | 1430.661239 | 11165932 | 45.93 | 1 | 6 | 6 | 3 | 0 | 1 | 0 | 0 | 4366.718479 | -0.0053 | -1.21 | -3.66 |
| Trypsin | 2 | 152 | 4663 | 1 | 3996 | 4367.721 | 42.2728 | 20,135 | 4366.713175 | 14 | 1074 | 1086 | NFTTAPAICHDGK | 0.2000000000000.0 | 1074(NFT) | 1 | 1433.6495 | 1430.661239 | 11165932 | 45.93 | 2 | 7 | 6 | 2 | 1 | 0 | 0 | 0 | 4366.71848 | -0.0053 | -1.21 | -3.66 |
| Trypsin | 2 | 152 | 4729 | 1 | 3966 | 4351.7243 | 40.7927 | 19,647 | 4350.716475 | 14 | 1074 | 1086 | NFTTAPAICHDGK | 0.2000000000000.0 | 1074(NFT) | 1 | 1433.6495 | 1430.661239 | 11165932 | 45.93 | 1 | 5 | 6 | 4 | 0 | 1 | 0 | 0 | 4350.723564 | -0.0071 | -1.62 | -5.14 |
| Trypsin | 2 | 152 | 4729 | 1 | 3966 | 4351.7243 | 40.7927 | 19,647 | 4350.716475 | 14 | 1074 | 1086 | NFTTAPAICHDGK | 0.2000000000000.0 | 1074(NFT) | 1 | 1433.6495 | 1430.661239 | 11165932 | 45.93 | 2 | 6 | 6 | 3 | 1 | 0 | 0 | 0 | 4350.723565 | -0.0071 | -1.62 | -5.14 |
| Trypsin | 2 | 152 | 4733 | 1 | 4063 | 4544.7477 | 45.5216 | 19,619 | 4543.739875 | 14 | 1074 | 1086 | NFTTAPAICHDGK | 0.2000000000000.0 | 1074(NFT) | 1 | 1433.6495 | 1430.661239 | 11165932 | 45.93 | 1 | 7 | 6 | 1 | 0 | 2 | 0 | 0 | 4543.745816 | -0.0059 | -1.3 | -0.41 |
| Trypsin | 2 | 152 | 4733 | 1 | 4063 | 4544.7477 | 45.5216 | 19,619 | 4543.739875 | 14 | 1074 | 1086 | NFTTAPAICHDGK | 0.2000000000000.0 | 1074(NFT) | 1 | 1433.6495 | 1430.661239 | 11165932 | 45.93 | 2 | 8 | 6 | 0 | 1 | 1 | 0 | 0 | 4543.745817 | -0.0059 | -1.3 | -0.41 |
| Trypsin | 2 | 152 | 4747 | 1 | 3900 | 4368.7204 | 37.6575 | 19,490 | 4367.712575 | 14 | 1074 | 1086 | NFTTAPAICHDGK | 0.2000000000000.0 | 1074(NFT) | 1 | 1433.6495 | 1430.661239 | 11165932 | 45.93 | | | | | | | | | | | | |
| Trypsin | 2 | 152 | 4829 | 1 | 4025 | 2996.2024 | 43.7131 | 18,978 | 2995.194575 | 14 | 1074 | 1086 | NFTTAPAICHDGK | 0.2000000000000.0 | 1074(NFT) | 1 | 1433.6495 | 1430.661239 | 11165932 | 45.93 | 1 | 4 | 3 | 0 | 0 | 1 | 0 | 1 | 2995.200985 | -0.0064 | -2.14 | -2.22 |
| Trypsin | 2 | 152 | 4856 | 1 | 4011 | 3637.459 | 43.0002 | 18,800 | 3636.451175 | 14 | 1074 | 1086 | NFTTAPAICHDGK | 0.2000000000000.0 | 1074(NFT) | 1 | 1433.6495 | 1430.661239 | 11165932 | 45.93 | 1 | 5 | 4 | 2 | 1 | 0 | 0 | 1 | 3636.454086 | -0.0029 | -0.8 | -2.93 |
| Trypsin | 2 | 152 | 4856 | 1 | 4011 | 3637.459 | 43.0002 | 18,800 | 3636.451175 | 14 | 1074 | 1086 | NFTTAPAICHDGK | 0.2000000000000.0 | 1074(NFT) | 1 | 1433.6495 | 1430.661239 | 11165932 | 45.93 | 2 | 4 | 4 | 3 | 0 | 1 | 0 | 0 | 3636.454085 | -0.0029 | -0.8 | -2.93 |
| Trypsin | 2 | 152 | 5011 | 1 | 3987 | 2648.0958 | 41.854 | 17,635 | 2647.087975 | 14 | 1074 | 1086 | NFTTAPAICHDGK | 0.2000000000000.0 | 1074(NFT) | 1 | 1433.6495 | 1430.661239 | 11165932 | 45.93 | 1 | 5 | 2 | 0 | 0 | 0 | 0 | 1 | 2647.084105 | 0.0039 | 1.46 | -4.08 |
| Trypsin | 2 | 152 | 5049 | 1 | 3926 | 4295.6971 | 38.9042 | 17,296 | 4294.689275 | 14 | 1074 | 1086 | NFTTAPAICHDGK | 0.2000000000000.0 | 1074(NFT) | 1 | 1433.6495 | 1430.661239 | 11165932 | 45.93 | 1 | 8 | 7 | 1 | 0 | 0 | 0 | 1 | 4294.697351 | -0.0081 | -1.88 | -7.03 |
| Trypsin | 2 | 152 | 5098 | 1 | 3963 | 2892.2012 | 40.6536 | 16,921 | 2891.193375 | 14 | 1074 | 1086 | NFTTAPAICHDGK | 0.2000000000000.0 | 1074(NFT) | 1 | 1433.6495 | 1430.661239 | 11165932 | 45.93 | 1 | 4 | 4 | 0 | 0 | 0 | 0 | 1 | 2891.190027 | 0.0033 | 1.15 | -5.28 |
| Trypsin | 2 | 152 | 5124 | 1 | 3904 | 4263.6945 | 37.855 | 16,765 | 4262.686675 | 14 | 1074 | 1086 | NFTTAPAICHDGK | 0.2000000000000.0 | 1074(NFT) | 1 | 1433.6495 | 1430.661239 | 11165932 | 45.93 | 1 | 6 | 7 | 3 | 0 | 0 | 0 | 0 | 4262.707521 | -0.0208 | -4.89 | -8.08 |
| Trypsin | 2 | 152 | 5238 | 1 | 4011 | 3815.5096 | 43.0002 | 16,093 | 3814.501775 | 14 | 1074 | 1086 | NFTTAPAICHDGK | 0.2000000000000.0 | 1074(NFT) | 1 | 1433.6495 | 1430.661239 | 11165932 | 45.93 | 1 | 6 | 4 | 2 | 0 | 1 | 0 | 0 | 3814.501824 | 0.0000 | -0.01 | -2.93 |
| Trypsin | 2 | 152 | 5238 | 1 | 4011 | 3815.5096 | 43.0002 | 16,093 | 3814.501775 | 14 | 1074 | 1086 | NFTTAPAICHDGK | 0.2000000000000.0 | 1074(NFT) | 1 | 1433.6495 | 1430.661239 | 11165932 | 45.93 | 2 | 7 | 4 | 1 | 1 | 0 | 0 | 0 | 3814.501825 | 0.0000 | -0.01 | -2.93 |
| Trypsin | 2 | 152 | 5299 | 1 | 4017 | 3929.5517 | 43.3405 | 15,679 | 3928.543875 | 14 | 1074 | 1086 | NFTTAPAICHDGK | 0.2000000000000.0 | 1074(NFT) | 1 | 1433.6495 | 1430.661239 | 11165932 | 45.93 | 1 | 6 | 6 | 0 | 0 | 1 | 0 | 0 | 3928.544752 | -0.0009 | -0.22 | -2.59 |
| Trypsin | 2 | 152 | 5303 | 1 | 3973 | 3914.5781 | 41.1282 | 15,638 | 3913.570275 | 14 | 1074 | 1086 | NFTTAPAICHDGK | 0.2000000000000.0 | 1074(NFT) | 1 | 1433.6495 | 1430.661239 | 11165932 | 45.93 | 1 | 6 | 6 | 2 | 0 | 0 | 0 | 1 | 3913.570239 | 0.0000 | 0 | -4.81 |
| Trypsin | 2 | 152 | 5338 | 1 | 3904 | 3946.5728 | 37.855 | 15,400 | 3945.564975 | 14 | 1074 | 1086 | NFTTAPAICHDGK | 0.2000000000000.0 | 1074(NFT) | 1 | 1433.6495 | 1430.661239 | 11165932 | 45.93 | 1 | 8 | 6 | 0 | 0 | 0 | 0 | 0 | 3945.560069 | 0.0049 | 1.24 | -8.08 |
| Trypsin | 2 | 152 | 5345 | 1 | 3922 | 3159.279 | 38.7023 | 15,353 | 3158.271175 | 14 | 1074 | 1086 | NFTTAPAICHDGK | 0.2000000000000.0 | 1074(NFT) | 1 | 1433.6495 | 1430.661239 | 11165932 | 45.93 | 1 | 6 | 3 | 1 | 0 | 0 | 0 | 1 | 3158.274211 | -0.0030 | -0.96 | -7.23 |
| Trypsin | 2 | 152 | 5349 | 1 | 3922 | 4263.7079 | 38.7023 | 15,330 | 4262.700075 | 14 | 1074 | 1086 | NFTTAPAICHDGK | 0.2000000000000.0 | 1074(NFT) | 1 | 1433.6495 | 1430.661239 | 11165932 | 45.93 | 1 | 6 | 7 | 3 | 0 | 0 | 0 | 0 | 4262.707521 | -0.0074 | -1.74 | -7.23 |
| Trypsin | 2 | 152 | 5430 | 1 | 3910 | 4206.6647 | 38.1306 | 14,625 | 4205.656875 | 14 | 1074 | 1086 | NFTTAPAICHDGK | 0.2000000000000.0 | 1074(NFT) | 1 | 1433.6495 | 1430.661239 | 11165932 | 45.93 | | | | | | | | | | | | |
| Trypsin | 2 | 152 | 5613 | 1 | 4021 | 3913.5611 | 43.5241 | 13,437 | 3912.553275 | 14 | 1074 | 1086 | NFTTAPAICHDGK | 0.2000000000000.0 | 1074(NFT) | 1 | 1433.6495 | 1430.661239 | 11165932 | 45.93 | 1 | 6 | 6 | 0 | 1 | 0 | 0 | 1 | 3912.549838 | 0.0034 | 0.87 | -2.41 |
| Trypsin | 2 | 152 | 5613 | 1 | 4021 | 3913.5611 | 43.5241 | 13,437 | 3912.553275 | 14 | 1074 | 1086 | NFTTAPAICHDGK | 0.2000000000000.0 | 1074(NFT) | 1 | 1433.6495 | 1430.661239 | 11165932 | 45.93 | 2 | 5 | 6 | 1 | 0 | 1 | 0 | 0 | 3912.549837 | 0.0034 | 0.87 | -2.41 |
| Trypsin | 2 | 152 | 5890 | 1 | 4000 | 4148.6517 | 42.465 | 11,849 | 4147.643875 | 14 | 1074 | 1086 | NFTTAPAICHDGK | 0.2000000000000.0 | 1074(NFT) | 1 | 1433.6495 | 1430.661239 | 11165932 | 45.93 | 1 | 6 | 5 | 3 | 1 | 0 | 0 | 1 | 4147.644192 | -0.0003 | -0.07 | -3.47 |
| Trypsin | 2 | 152 | 5890 | 1 | 4000 | 4148.6517 | 42.465 | 11,849 | 4147.643875 | 14 | 1074 | 1086 | NFTTAPAICHDGK | 0.2000000000000.0 | 1074(NFT) | 1 | 1433.6495 | 1430.661239 | 11165932 | 45.93 | 2 | 5 | 5 | 4 | 0 | 1 | 0 | 0 | 4147.644191 | -0.0003 | -0.07 | -3.47 |
| Trypsin | 2 | 152 | 5956 | 1 | 3983 | 3013.2215 | 41.6523 | 11,412 | 3012.213675 | 14 | 1074 | 1086 | NFTTAPAICHDGK | 0.2000000000000.0 | 1074(NFT) | 1 | 1433.6495 | 1430.661239 | 11165932 | 45.93 | 1 | 6 | 3 | 0 | 0 | 0 | 0 | 1 | 3012.216302 | -0.0026 | -0.87 | -4.28 |
| Trypsin | 2 | 152 | 6019 | 1 | 3973 | 3282.36 | 41.1282 | 10,984 | 3281.352175 | 14 | 1074 | 1086 | NFTTAPAICHDGK | 0.2000000000000.0 | 1074(NFT) | 1 | 1433.6495 | 1430.661239 | 11165932 | 45.93 | 1 | 3 | 6 | 1 | 0 | 0 | 0 | 1 | 3281.353858 | -0.0017 | -0.51 | -4.81 |
| Trypsin | 2 | 152 | 6238 | 1 | 4008 | 4002.5947 | 42.8568 | 9,703 | 4001.586875 | 14 | 1074 | 1086 | NFTTAPAICHDGK | 0.2000000000000.0 | 1074(NFT) | 1 | 1433.6495 | 1430.661239 | 11165932 | 45.93 | 1 | 6 | 5 | 2 | 1 | 0 | 0 | 1 | 4001.586283 | 0.0006 | 0.14 | -3.08 |
| Trypsin | 2 | 152 | 6238 | 1 | 4008 | 4002.5947 | 42.8568 | 9,703 | 4001.586875 | 14 | 1074 | 1086 | NFTTAPAICHDGK | 0.2000000000000.0 | 1074(NFT) | 1 | 1433.6495 | 1430.661239 | 11165932 | 45.93 | 2 | 5 | 5 | 3 | 0 | 1 | 0 | 0 | 4001.586282 | 0.0006 | 0.14 | -3.08 |
| Trypsin | 2 | 152 | 6261 | 1 | 3971 | 3216.305 | 41.0356 | 9,567 | 3215.297175 | 14 | 1074 | 1086 | NFTTAPAICHDGK | 0.2000000000000.0 | 1074(NFT) | 1 | 1433.6495 | 1430.661239 | 11165932 | 45.93 | 1 | 6 | 4 | 0 | 0 | 0 | 0 | 1 | 3215.295675 | 0.0015 | 0.46 | -4.90 |
| Trypsin | 2 | 152 | 6293 | 1 | 3993 | 4513.7846 | 42.1338 | 9,427 | 4512.776775 | 14 | 1074 | 1086 | NFTTAPAICHDGK | 0.2000000000000.0 | 1074(NFT) | 1 | 1433.6495 | 1430.661239 | 11165932 | 45.93 | 1 | 7 | 6 | 3 | 1 | 0 | 0 | 0 | 4512.776389 | 0.0004 | 0.08 | -3.80 |
| Trypsin | 2 | 152 | 6293 | 1 | 3993 | 4513.7846 | 42.1338 | 9,427 | 4512.776775 | 14 | 1074 | 1086 | NFTTAPAICHDGK | 0.2000000000000.0 | 1074(NFT) | 1 | 1433.6495 | 1430.661239 | 11165932 | 45.93 | 2 | 6 | 6 | 4 | 0 | 1 | 0 | 0 | 4512.776388 | 0.0004 | 0.08 | -3.80 |
| Trypsin | 2 | 152 | 6332 | 1 | 4012 | 3669.4533 | 43.0514 | 9,224 | 3668.445475 | 14 | 1074 | 1086 | NFTTAPAICHDGK | 0.2000000000000.0 | 1074(NFT) | 1 | 1433.6495 | 1430.661239 | 11165932 | 45.93 | 1 | 7 | 4 | 0 | 1 | 0 | 0 | 1 | 3668.443916 | 0.0016 | 0.42 | -2.88 |
| Trypsin | 2 | 152 | 6332 | 1 | 4012 | 3669.4533 | 43.0514 | 9,224 | 3668.445475 | 14 | 1074 | 1086 | NFTTAPAICHDGK | 0.2000000000000.0 | 1074(NFT) | 1 | 1433.6495 | 1430.661239 | 11165932 | 45.93 | 2 | 6 | 4 | 1 | 0 | 1 | 0 | 1 | 3668.443915 | 0.0016 | 0.42 | -2.88 |
| Trypsin | 2 | 152 | 6334 | 1 | 3921 | 4441.7622 | 38.6488 | 9,221 | 4440.754375 | 14 | 1074 | 1086 | NFTTAPAICHDGK | 0.2000000000000.0 | 1074(NFT) | 1 | 1433.6495 | 1430.661239 | 11165932 | 45.93 | 1 | 8 | 7 | 2 | 0 | 0 | 0 | 0 | 4440.75526 | -0.0009 | -0.19 | -7.29 |
| Trypsin | 2 | 152 | 6346 | 1 | 3900 | 4514.7831 | 37.6575 | 9,151 | 4513.775275 | 14 | 1074 | 1086 | NFTTAPAICHDGK | 0.2000000000000.0 | 1074(NFT) | 1 | 1433.6495 | 1430.661239 | 11165932 | 45.93 | 1 | 9 | 8 | 0 | 0 | 0 | 0 | 1 | 4513.771639 | 0.0036 | 0.8 | -8.28 |
| alpha-Lytic | 36 | 5 | 174 | 1 | 4026 | 2485.0235 | 37.5282 | 765,840 | 2484.015675 | 1016 | 1071 | 1080 | QEKNFTTAPA | - | 1074(NFT) | 1 | | 1105.54036 | - | 42.00 | 1 | 6 | 2 | 0 | 0 | 0 | 0 | 1 | 2484.01605 | -0.0004 | -0.15 | -4.47 |
| [illegible] | [illegible] | [illegible] | [illegible] | [illegible] | [illegible] | [illegible] | [illegible] | [illegible] | [illegible] | [illegible] | [illegible] | [illegible] | [illegible] | [illegible] | [illegible] | [illegible] | [illegible] | [illegible] | [illegible] | [illegible] | [illegible] | [illegible] | [illegible] | [illegible] | [illegible] | [illegible] | [illegible] | [illegible] | [illegible] | [illegible] | [illegible] | [illegible] |

| | | | | | | | | | | | | | | | | | | | | | | | | | | | | | | | | |
|---|---|---|---|---|---|---|---|---|---|---|---|---|---|---|---|---|---|---|---|---|---|---|---|---|---|---|---|---|---|---|---|---|
| Chymotrypsin | 83 | 31 | 3601 | 1 | 4205 | 3242.2346 | 55.5481 | 103,753 | 3241.226775 | 42 | 1096 | 1103 | VSNGTHWF | 0.00200000.0 | 1098(NGT) | 1 | 949.418 | 946.4297387 | 9738917 | 60.37 | 1 | 6 | 5 | 0 | 0 | 1 | 0 | 1 | 3241.233879 | -0.0071 | -2.19 | -4.83 |
| Chymotrypsin | 83 | 31 | 3723 | 1 | 4110 | 2659.0594 | 51.5362 | 99,462 | 2658.051575 | 42 | 1096 | 1103 | VSNGTHWF | 0.00200000.0 | 1098(NGT) | 1 | 949.418 | 946.4297387 | 9738917 | 60.37 | 1 | 5 | 3 | 2 | 0 | 0 | 0 | 0 | 2658.047796 | 0.0038 | 1.42 | -8.84 |
| Chymotrypsin | 83 | 31 | 3761 | 1 | 4111 | 3138.2322 | 51.5799 | 98,535 | 3137.224375 | 42 | 1096 | 1103 | VSNGTHWF | 0.00200000.0 | 1098(NGT) | 1 | 949.418 | 946.4297387 | 9738917 | 60.37 | 1 | 6 | 6 | 0 | 0 | 0 | 0 | 1 | 3137.222921 | 0.0015 | 0.46 | -8.79 |
| Chymotrypsin | 83 | 31 | 4727 | 1 | 4070 | 3373.3282 | 49.8409 | 74,168 | 3372.320375 | 42 | 1096 | 1103 | VSNGTHWF | 0.00200000.0 | 1098(NGT) | 1 | 949.418 | 946.4297387 | 9738917 | 60.37 | 1 | 6 | 5 | 3 | 0 | 0 | 0 | 1 | 3372.317275 | 0.0031 | 0.91 | -10.53 |
| Chymotrypsin | 83 | 31 | 6115 | 1 | 4186 | 3591.3834 | 54.7443 | 50,508 | 3590.375575 | 42 | 1096 | 1103 | VSNGTHWF | 0.00200000.0 | 1098(NGT) | 1 | 949.418 | 946.4297387 | 9738917 | 60.37 | 1 | 7 | 6 | 0 | 1 | 0 | 0 | 1 | 3590.371162 | 0.0044 | 1.22 | -5.63 |
| Chymotrypsin | 83 | 31 | 6115 | 1 | 4186 | 3591.3834 | 54.7443 | 50,508 | 3590.375575 | 42 | 1096 | 1103 | VSNGTHWF | 0.00200000.0 | 1098(NGT) | 1 | 949.418 | 946.4297387 | 9738917 | 60.37 | 2 | 6 | 6 | 1 | 0 | 1 | 0 | 0 | 3590.371161 | 0.0044 | 1.22 | -5.63 |
| Chymotrypsin | 83 | 31 | 6274 | 1 | 4055 | 3884.5257 | 49.2017 | 48,193 | 3883.517875 | 42 | 1096 | 1103 | VSNGTHWF | 0.00200000.0 | 1098(NGT) | 1 | 949.418 | 946.4297387 | 9738917 | 60.37 | 1 | 7 | 6 | 4 | 0 | 0 | 0 | 1 | 3883.507381 | 0.0105 | 2.7 | -11.17 |
| Chymotrypsin | 83 | 31 | 6274 | 1 | 4055 | 3884.5257 | 49.2017 | 48,193 | 3883.517875 | 42 | 1096 | 1103 | VSNGTHWF | 0.00200000.0 | 1098(NGT) | 1 | 949.418 | 946.4297387 | 9738917 | 60.37 | 2 | 2 | 10 | 0 | 2 | 0 | 1 | -1 | 3883.519951 | -0.0021 | -0.53 | -11.17 |
| Trypsin | 18 | 116 | 53 | 1 | 4369 | 4216.7575 | 59.8897 | 2,213,270 | 4215.749675 | 17 | 1092 | 1107 | EGVFVSNGTHWFVTQR | 0.0000002000000000.0 | 1098(NGT) | 1 | 1865.8947 | 1862.906439 | 9669981 | 67.72 | 1 | 7 | 6 | 0 | 0 | 0 | 0 | 1 | 4215.752445 | -0.0028 | -0.65 | -7.83 |
| Trypsin | 18 | 116 | 68 | 1 | 4382 | 3591.5255 | 60.5509 | 1,902,650 | 3590.517675 | 17 | 1092 | 1107 | EGVFVSNGTHWFVTQR | 0.0000002000000000.0 | 1098(NGT) | 1 | 1865.8947 | 1862.906439 | 9669981 | 67.72 | 1 | 6 | 3 | 1 | 0 | 0 | 0 | 1 | 3590.519411 | -0.0017 | -0.48 | -7.17 |
| Trypsin | 18 | 116 | 74 | 1 | 4377 | 3851.6254 | 60.2882 | 1,643,340 | 3850.617575 | 17 | 1092 | 1107 | EGVFVSNGTHWFVTQR | 0.0000002000000000.0 | 1098(NGT) | 1 | 1865.8947 | 1862.906439 | 9669981 | 67.72 | 1 | 6 | 5 | 0 | 0 | 0 | 0 | 1 | 3850.620248 | -0.0027 | -0.69 | -7.43 |
| Trypsin | 18 | 116 | 82 | 1 | 4366 | 4362.8173 | 59.7429 | 1,508,800 | 4361.809475 | 17 | 1092 | 1107 | EGVFVSNGTHWFVTQR | 0.0000002000000000.0 | 1098(NGT) | 1 | 1865.8947 | 1862.906439 | 9669981 | 67.72 | 1 | 7 | 6 | 1 | 0 | 0 | 0 | 1 | 4361.810354 | -0.0009 | -0.2 | -7.98 |
| Trypsin | 18 | 116 | 90 | 1 | 4364 | 4508.874 | 59.6474 | 1,349,040 | 4507.866175 | 17 | 1092 | 1107 | EGVFVSNGTHWFVTQR | 0.0000002000000000.0 | 1098(NGT) | 1 | 1865.8947 | 1862.906439 | 9669981 | 67.72 | 1 | 7 | 6 | 2 | 0 | 0 | 0 | 0 | 4507.868263 | -0.0021 | -0.46 | -8.07 |
| Trypsin | 18 | 116 | 95 | 1 | 4386 | 3445.4677 | 60.7457 | 1,266,530 | 3444.459875 | 17 | 1092 | 1107 | EGVFVSNGTHWFVTQR | 0.0000002000000000.0 | 1098(NGT) | 1 | 1865.8947 | 1862.906439 | 9669981 | 67.72 | 1 | 6 | 3 | 0 | 0 | 0 | 0 | 1 | 3444.461502 | -0.0016 | -0.47 | -6.97 |
| Trypsin | 18 | 116 | 110 | 1 | 4374 | 3997.6839 | 60.146 | 1,114,250 | 3996.676075 | 17 | 1092 | 1107 | EGVFVSNGTHWFVTQR | 0.0000002000000000.0 | 1098(NGT) | 1 | 1865.8947 | 1862.906439 | 9669981 | 67.72 | 1 | 6 | 5 | 1 | 0 | 0 | 0 | 1 | 3996.678157 | -0.0021 | -0.52 | -7.57 |
| Trypsin | 18 | 116 | 124 | 1 | 4386 | 3429.4735 | 60.7457 | 998,586 | 3428.465675 | 17 | 1092 | 1107 | EGVFVSNGTHWFVTQR | 0.0000002000000000.0 | 1098(NGT) | 1 | 1865.8947 | 1862.906439 | 9669981 | 67.72 | 1 | 5 | 3 | 1 | 0 | 0 | 0 | 1 | 3428.466587 | -0.0009 | -0.26 | -6.97 |
| Trypsin | 18 | 116 | 130 | 1 | 4449 | 4523.8461 | 63.7991 | 975,458 | 4522.838275 | 17 | 1092 | 1107 | EGVFVSNGTHWFVTQR | 0.0000002000000000.0 | 1098(NGT) | 1 | 1865.8947 | 1862.906439 | 9669981 | 67.72 | 1 | 7 | 6 | 0 | 0 | 1 | 0 | 1 | 4522.842776 | -0.0045 | -0.99 | -3.92 |
| Trypsin | 18 | 116 | 143 | 1 | 4390 | 3283.4143 | 60.9585 | 880,810 | 3282.406475 | 17 | 1092 | 1107 | EGVFVSNGTHWFVTQR | 0.0000002000000000.0 | 1098(NGT) | 1 | 1865.8947 | 1862.906439 | 9669981 | 67.72 | 1 | 5 | 3 | 0 | 0 | 0 | 0 | 1 | 3282.408678 | -0.0022 | -0.67 | -6.76 |
| Trypsin | 18 | 116 | 148 | 1 | 4379 | 3753.5786 | 60.3951 | 824,079 | 3752.570775 | 17 | 1092 | 1107 | EGVFVSNGTHWFVTQR | 0.0000002000000000.0 | 1098(NGT) | 1 | 1865.8947 | 1862.906439 | 9669981 | 67.72 | 1 | 7 | 3 | 1 | 0 | 0 | 0 | 0 | 3752.572235 | -0.0015 | -0.38 | -7.33 |
| Trypsin | 18 | 116 | 149 | 1 | 4383 | 3607.5197 | 60.5957 | 814,909 | 3606.511875 | 17 | 1092 | 1107 | EGVFVSNGTHWFVTQR | 0.0000002000000000.0 | 1098(NGT) | 1 | 1865.8947 | 1862.906439 | 9669981 | 67.72 | 1 | 7 | 3 | 0 | 0 | 0 | 0 | 0 | 3606.514326 | -0.0025 | -0.67 | -7.12 |
| Trypsin | 18 | 116 | 157 | 1 | 4464 | 3752.5569 | 64.51 | 795,335 | 3751.549075 | 17 | 1092 | 1107 | EGVFVSNGTHWFVTQR | 0.0000002000000000.0 | 1098(NGT) | 1 | 1865.8947 | 1862.906439 | 9669981 | 67.72 | 1 | 6 | 3 | 0 | 0 | 1 | 0 | 1 | 3751.551833 | -0.0028 | -0.73 | -3.21 |
| Trypsin | 18 | 116 | 163 | 1 | 4380 | 3632.5517 | 60.4506 | 764,955 | 3631.543875 | 17 | 1092 | 1107 | EGVFVSNGTHWFVTQR | 0.0000002000000000.0 | 1098(NGT) | 1 | 1865.8947 | 1862.906439 | 9669981 | 67.72 | 1 | 5 | 4 | 1 | 0 | 0 | 0 | 1 | 3631.54596 | -0.0021 | -0.57 | -7.27 |
| Trypsin | 18 | 116 | 168 | 1 | 4384 | 3404.4411 | 60.6462 | 741,675 | 3403.433275 | 17 | 1092 | 1107 | EGVFVSNGTHWFVTQR | 0.0000002000000000.0 | 1098(NGT) | 1 | 1865.8947 | 1862.906439 | 9669981 | 67.72 | 1 | 7 | 2 | 0 | 0 | 0 | 0 | 1 | 3403.434953 | -0.0017 | -0.49 | -7.07 |
| Trypsin | 18 | 116 | 172 | 1 | 4455 | 4158.7148 | 64.0885 | 721,181 | 4157.706975 | 17 | 1092 | 1107 | EGVFVSNGTHWFVTQR | 0.0000002000000000.0 | 1098(NGT) | 1 | 1865.8947 | 1862.906439 | 9669981 | 67.72 | 1 | 6 | 5 | 0 | 0 | 1 | 0 | 1 | 4157.710579 | -0.0036 | -0.86 | -3.63 |
| Trypsin | 18 | 116 | 174 | 1 | 4443 | 4669.9046 | 63.5267 | 719,554 | 4668.896775 | 17 | 1092 | 1107 | EGVFVSNGTHWFVTQR | 0.0000002000000000.0 | 1098(NGT) | 1 | 1865.8947 | 1862.906439 | 9669981 | 67.72 | 1 | 7 | 6 | 1 | 0 | 1 | 0 | 1 | 4668.900685 | -0.0039 | -0.83 | -4.19 |
| Trypsin | 18 | 116 | 174 | 1 | 4443 | 4669.9046 | 63.5267 | 719,554 | 4668.896775 | 17 | 1092 | 1107 | EGVFVSNGTHWFVTQR | 0.0000002000000000.0 | 1098(NGT) | 1 | 1865.8947 | 1862.906439 | 9669981 | 67.72 | 2 | 8 | 6 | 0 | 1 | 0 | 0 | 1 | 4668.900686 | -0.0039 | -0.83 | -4.19 |
| Trypsin | 18 | 116 | 176 | 1 | 4370 | 4143.7408 | 59.9461 | 710,592 | 4142.732975 | 17 | 1092 | 1107 | EGVFVSNGTHWFVTQR | 0.0000002000000000.0 | 1098(NGT) | 1 | 1865.8947 | 1862.906439 | 9669981 | 67.72 | 1 | 6 | 5 | 2 | 0 | 0 | 0 | 1 | 4142.736066 | -0.0031 | -0.74 | -7.77 |
| Trypsin | 18 | 116 | 268 | 1 | 4362 | 4654.9291 | 59.545 | 499,899 | 4653.921275 | 17 | 1092 | 1107 | EGVFVSNGTHWFVTQR | 0.0000002000000000.0 | 1098(NGT) | 1 | 1865.8947 | 1862.906439 | 9669981 | 67.72 | 1 | 7 | 6 | 3 | 0 | 0 | 0 | 0 | 4653.926172 | -0.0049 | -1.05 | -8.18 |
| Trypsin | 18 | 116 | 269 | 1 | 4451 | 4304.7771 | 63.8999 | 494,772 | 4303.769275 | 17 | 1092 | 1107 | EGVFVSNGTHWFVTQR | 0.0000002000000000.0 | 1098(NGT) | 1 | 1865.8947 | 1862.906439 | 9669981 | 67.72 | 1 | 6 | 5 | 1 | 0 | 1 | 0 | 1 | 4303.768488 | 0.0008 | 0.18 | -3.82 |
| Trypsin | 18 | 116 | 269 | 1 | 4451 | 4304.7771 | 63.8999 | 494,772 | 4303.769275 | 17 | 1092 | 1107 | EGVFVSNGTHWFVTQR | 0.0000002000000000.0 | 1098(NGT) | 1 | 1865.8947 | 1862.906439 | 9669981 | 67.72 | 2 | 7 | 5 | 0 | 1 | 0 | 0 | 0 | 4303.768489 | 0.0008 | 0.18 | -3.82 |
| Trypsin | 18 | 116 | 287 | 1 | 4377 | 4054.7054 | 60.2882 | 468,899 | 4053.697575 | 17 | 1092 | 1107 | EGVFVSNGTHWFVTQR | 0.0000002000000000.0 | 1098(NGT) | 1 | 1865.8947 | 1862.906439 | 9669981 | 67.72 | 1 | 6 | 6 | 0 | 0 | 0 | 0 | 1 | 4053.699621 | -0.0020 | -0.5 | -7.43 |
| Trypsin | 18 | 116 | 306 | 1 | 4461 | 3793.5835 | 64.3794 | 442,675 | 3792.575675 | 17 | 1092 | 1107 | EGVFVSNGTHWFVTQR | 0.0000002000000000.0 | 1098(NGT) | 1 | 1865.8947 | 1862.906439 | 9669981 | 67.72 | 1 | 5 | 4 | 0 | 0 | 1 | 0 | 1 | 3792.578382 | -0.0027 | -0.71 | -3.34 |
| Trypsin | 18 | 116 | 307 | 1 | 4384 | 3486.494 | 60.6462 | 442,493 | 3485.486175 | 17 | 1092 | 1107 | EGVFVSNGTHWFVTQR | 0.0000002000000000.0 | 1098(NGT) | 1 | 1865.8947 | 1862.906439 | 9669981 | 67.72 | 1 | 5 | 4 | 0 | 0 | 0 | 0 | 1 | 3485.488051 | -0.0019 | -0.53 | -7.07 |
| Trypsin | 18 | 116 | 313 | 1 | 4393 | 3080.336 | 61.1097 | 438,128 | 3079.328175 | 17 | 1092 | 1107 | EGVFVSNGTHWFVTQR | 0.0000002000000000.0 | 1098(NGT) | 1 | 1865.8947 | 1862.906439 | 9669981 | 67.72 | 1 | 5 | 2 | 0 | 0 | 0 | 0 | 1 | 3079.329305 | -0.0011 | -0.36 | -6.61 |
| Trypsin | 18 | 116 | 350 | 1 | 4377 | 3778.6095 | 60.2882 | 403,250 | 3777.601675 | 17 | 1092 | 1107 | EGVFVSNGTHWFVTQR | 0.0000002000000000.0 | 1098(NGT) | 1 | 1865.8947 | 1862.906439 | 9669981 | 67.72 | 1 | 5 | 4 | 2 | 0 | 0 | 0 | 1 | 3777.603869 | -0.0022 | -0.58 | -7.43 |
| Trypsin | 18 | 116 | 359 | 1 | 4367 | 4289.8026 | 59.784 | 397,512 | 4288.794775 | 17 | 1092 | 1107 | EGVFVSNGTHWFVTQR | 0.0000002000000000.0 | 1098(NGT) | 1 | 1865.8947 | 1862.906439 | 9669981 | 67.72 | 1 | 6 | 5 | 3 | 0 | 0 | 0 | 1 | 4288.793975 | 0.0008 | 0.18 | -7.94 |
| Trypsin | 18 | 116 | 365 | 1 | 4388 | 3242.3884 | 60.8532 | 393,984 | 3241.380575 | 17 | 1092 | 1107 | EGVFVSNGTHWFVTQR | 0.0000002000000000.0 | 1098(NGT) | 1 | 1865.8947 | 1862.906439 | 9669981 | 67.72 | 1 | 6 | 2 | 0 | 0 | 0 | 0 | 1 | 3241.382129 | -0.0016 | -0.47 | -6.87 |
| Trypsin | 18 | 116 | 409 | 1 | 4469 | 3590.5045 | 64.7493 | 355,150 | 3589.496675 | 17 | 1092 | 1107 | EGVFVSNGTHWFVTQR | 0.0000002000000000.0 | 1098(NGT) | 1 | 1865.8947 | 1862.906439 | 9669981 | 67.72 | 1 | 5 | 3 | 0 | 0 | 1 | 0 | 1 | 3589.499009 | -0.0023 | -0.65 | -2.97 |
| Trypsin | 18 | 116 | 452 | 1 | 4382 | 3566.4961 | 60.5509 | 322,768 | 3565.488275 | 17 | 1092 | 1107 | EGVFVSNGTHWFVTQR | 0.0000002000000000.0 | 1098(NGT) | 1 | 1865.8947 | 1862.906439 | 9669981 | 67.72 | 1 | 8 | 2 | 0 | 0 | 0 | 0 | 1 | 3565.487777 | 0.0005 | 0.13 | -7.17 |
| Trypsin | 18 | 116 | 507 | 1 | 4394 | 3121.3615 | 61.1533 | 284,871 | 3120.353675 | 17 | 1092 | 1107 | EGVFVSNGTHWFVTQR | 0.0000002000000000.0 | 1098(NGT) | 1 | 1865.8947 | 1862.906439 | 9669981 | 67.72 | 1 | 4 | 3 | 0 | 0 | 0 | 0 | 1 | 3120.355854 | -0.0022 | -0.69 | -6.57 |
| Trypsin | 18 | 116 | 517 | 1 | 4384 | 3648.5541 | 60.6462 | 277,797 | 3647.546275 | 17 | 1092 | 1107 | EGVFVSNGTHWFVTQR | 0.0000002000000000.0 | 1098(NGT) | 1 | 1865.8947 | 1862.906439 | 9669981 | 67.72 | 1 | 6 | 4 | 0 | 0 | 0 | 0 | 1 | 3647.540875 | 0.0054 | 1.48 | -7.07 |
| Trypsin | 18 | 116 | 538 | 1 | 4374 | 4200.7625 | 60.146 | 267,274 | 4199.754675 | 17 | 1092 | 1107 | EGVFVSNGTHWFVTQR | 0.0000002000000000.0 | 1098(NGT) | 1 | 1865.8947 | 1862.906439 | 9669981 | 67.72 | 1 | 6 | 6 | 1 | 0 | 0 | 0 | 1 | 4199.75753 | -0.0029 | -0.67 | -7.57 |
| Trypsin | 18 | 116 | 553 | 1 | 4383 | 3689.5719 | 60.5957 | 260,409 | 3688.564075 | 17 | 1092 | 1107 | EGVFVSNGTHWFVTQR | 0.0000002000000000.0 | 1098(NGT) | 1 | 1865.8947 | 1862.906439 | 9669981 | 67.72 | 1 | 5 | 5 | 0 | 0 | 0 | 0 | 1 | 3688.567424 | -0.0033 | -0.9 | -7.12 |
| Trypsin | 18 | 116 | 557 | 1 | 4450 | 4507.8613 | 63.8506 | 258,277 | 4506.853475 | 17 | 1092 | 1107 | EGVFVSNGTHWFVTQR | 0.0000002000000000.0 | 1098(NGT) | 1 | 1865.8947 | 1862.906439 | 9669981 | 67.72 | 1 | 7 | 6 | 0 | 1 | 0 | 0 | 1 | 4506.847862 | 0.0056 | 1.24 | -3.87 |
| Trypsin | 18 | 116 | 557 | 1 | 4450 | 4507.8613 | 63.8506 | 258,277 | 4506.853475 | 17 | 1092 | 1107 | EGVFVSNGTHWFVTQR | 0.0000002000000000.0 | 1098(NGT) | 1 | 1865.8947 | 1862.906439 | 9669981 | 67.72 | 2 | 6 | 6 | 1 | 0 | 1 | 0 | 0 | 4506.847861 | 0.0056 | 1.24 | -3.87 |
| Trypsin | 18 | 116 | 568 | 1 | 4380 | 3794.6059 | 60.4506 | 252,587 | 3793.598075 | 17 | 1092 | 1107 | EGVFVSNGTHWFVTQR | 0.0000002000000000.0 | 1098(NGT) | 1 | 1865.8947 | 1862.906439 | 9669981 | 67.72 | 1 | 6 | 4 | 1 | 0 | 0 | 0 | 1 | 3793.598784 | -0.0007 | -0.18 | -7.27 |
| Trypsin | 18 | 116 | 581 | 1 | 4382 | 3575.5309 | 60.5509 | 246,826 | 3574.523075 | 17 | 1092 | 1107 | EGVFVSNGTHWFVTQR | 0.0000002000000000.0 | 1098(NGT) | 1 | 1865.8947 | 1862.906439 | 9669981 | 67.72 | 1 | 5 | 3 | 2 | 0 | 0 | 0 | 0 | 3574.524496 | -0.0014 | -0.39 | -7.17 |
| Trypsin | 18 | 116 | 610 | 1 | 4466 | 3736.5611 | 64.6114 | 235,708 | 3735.553275 | 17 | 1092 | 1107 | EGVFVSNGTHWFVTQR | 0.0000002000000000.0 | 1098(NGT) | 1 | 1865.8947 | 1862.906439 | 9669981 | 67.72 | 1 | 6 | 3 | 0 | 1 | 0 | 0 | 1 | 3735.556919 | -0.0036 | -0.97 | -3.11 |
| Trypsin | 18 | 116 | 610 | 1 | 4466 | 3736.5611 | 64.6114 | 235,708 | 3735.553275 | 17 | 1092 | 1107 | EGVFVSNGTHWFVTQR | 0.0000002000000000.0 | 1098(NGT) | 1 | 1865.8947 | 1862.906439 | 9669981 | 67.72 | 2 | 5 | 3 | 1 | 0 | 1 | 0 | 0 | 3735.556918 | -0.0036 | -0.97 | -3.11 |
| Trypsin | 18 | 116 | 622 | 1 | 4458 | 4142.7223 | 64.2422 | 231,149 | 4141.714475 | 17 | 1092 | 1107 | EGVFVSNGTHWFVTQR | 0.0000002000000000.0 | 1098(NGT) | 1 | 1865.8947 | 1862.906439 | 9669981 | 67.72 | 1 | 5 | 5 | 1 | 0 | 1 | 0 | 1 | 4141.715664 | -0.0012 | -0.28 | -3.48 |
| Trypsin | 18 | 116 | 622 | 1 | 4458 | 4142.7223 | 64.2422 | 231,149 | 4141.714475 | 17 | 1092 | 1107 | EGVFVSNGTHWFVTQR | 0.0000002000000000.0 | 1098(NGT) | 1 | 1865.8947 | 1862.906439 | 9669981 | 67.72 | 2 | 6 | 5 | 0 | 1 | 0 | 0 | 1 | 4141.715665 | -0.0012 | -0.28 | -3.48 |
| Trypsin | 18 | 116 | 694 | 1 | 4380 | 3892.6523 | 60.4506 | 203,241 | 3891.644475 | 17 | 1092 | 1107 | EGVFVSNGTHWFVTQR | 0.0000002000000000.0 | 1098(NGT) | 1 | 1865.8947 | 1862.906439 | 9669981 | 67.72 | 1 | 5 | 6 | 0 | 0 | 0 | 0 | 1 | 3891.646797 | -0.0023 | -0.59 | -7.27 |
| Trypsin | 18 | 116 | 786 | 1 | 4390 | 3324.4413 | 60.9585 | 178,997 | 3323.433475 | 17 | 1092 | 1107 | EGVFVSNGTHWFVTQR | 0.0000002000000000.0 | 1098(NGT) | 1 | 1865.8947 | 1862.906439 | 9669981 | 67.72 | 1 | 4 | 4 | 0 | 0 | 0 | 0 | 1 | 3323.435227 | -0.0018 | -0.52 | -6.76 |
| Trypsin | 18 | 116 | 818 | 1 | 4390 | 3267.4218 | 60.9585 | 172,392 | 3266.413975 | 17 | 1092 | 1107 | EGVFVSNGTHWFVTQR | 0.0000002000000000.0 | 1098(NGT) | 1 | 1865.8947 | 1862.906439 | 9669981 | 67.72 | 1 | 4 | 3 | 1 | 0 | 0 | 0 | 1 | 3266.413763 | 0.0002 | 0.06 | -6.76 |
| Trypsin | 18 | 116 | 833 | 1 | 4442 | 4815.9625 | 63.4847 | 169,300 | 4814.954675 | 17 | 1092 | 1107 | EGVFVSNGTHWFVTQR | 0.0000002000000000.0 | 1098(NGT) | 1 | 1865.8947 | 1862.906439 | 9669981 | 67.72 | 1 | 8 | 6 | 1 | 1 | 0 | 0 | 1 | 4814.958595 | -0.0039 | -0.81 | -4.24 |
| Trypsin | 18 | 116 | 833 | 1 | 4442 | 4815.9625 | 63.4847 | 169,300 | 4814.954675 | 17 | 1092 | 1107 | EGVFVSNGTHWFVTQR | 0.0000002000000000.0 | 1098(NGT) | 1 | 1865.8947 | 1862.906439 | 9669981 | 67.72 | 2 | 7 | 6 | 2 | 0 | 1 | 0 | 0 | 4814.958594 | -0.0039 | -0.81 | -4.24 |
| Trypsin | 18 | 116 | 834 | 1 | 4461 | 3914.6125 | 64.3794 | 169,274 | 3913.604675 | 17 | 1092 | 1107 | EGVFVSNGTHWFVTQR | 0.0000002000000000.0 | 1098(NGT) | 1 | 1865.8947 | 1862.906439 | 9669981 | 67.72 | 1 | 7 | 3 | 0 | 0 | 1 | 1 | -1 | 3913.604657 | 0.0000 | 0 | -3.34 |
| Trypsin | 18 | 116 | 846 | 1 | 4446 | 4653.9099 | 63.6567 | 167,178 | 4652.902075 | 17 | 1092 | 1107 | EGVFVSNGTHWFVTQR | 0.0000002000000000.0 | 1098(NGT) | 1 | 1865.8947 | 1862.906439 | 9669981 | 67.72 | 1 | 7 | 6 | 1 | 1 | 0 | 0 | 1 | 4652.905771 | -0.0037 | -0.79 | -4.06 |
| Trypsin | 18 | 116 | 846 | 1 | 4446 | 4653.9099 | 63.6567 | 167,178 | 4652.902075 | 17 | 1092 | 1107 | EGVFVSNGTHWFVTQR | 0.0000002000000000.0 | 1098(NGT) | 1 | 1865.8947 | 1862.906439 | 9669981 | 67.72 | 2 | 6 | 6 | 2 | 0 | 1 | 0 | 0 | 4652.90577 | -0.0037 | -0.79 | -4.06 |
| Trypsin | 18 | 116 | 852 | 1 | 4380 | 3728.5499 | 60.4506 | 166,339 | 3727.542075 | 17 | 1092 | 1107 | EGVFVSNGTHWFVTQR | 0.0000002000000000.0 | 1098(NGT) | 1 | 1865.8947 | 1862.906439 | 9669981 | 67.72 | 1 | 9 | 2 | 0 | 0 | 0 | 0 | 1 | 3727.540601 | 0.0015 | 0.39 | -7.27 |
| Trypsin | 18 | 116 | 859 | 1 | 4377 | 3940.6633 | 60.2882 | 164,482 | 3939.655475 | 17 | 1092 | 1107 | EGVFVSNGTHWFVTQR | 0.0000002000000000.0 | 1098(NGT) | 1 | 1865.8947 | 1862.906439 | 9669981 | 67.72 | 1 | 6 | 4 | 2 | 0 | 0 | 0 | 1 | 3939.656693 | -0.0012 | -0.3 | -7.43 |
| Trypsin | 18 | 116 | 898 | 1 | 4390 | 3365.4675 | 60.9585 | 157,669 | 3364.459675 | 17 | 1092 | 1107 | EGVFVSNGTHWFVTQR | 0.0000002000000000.0 | 1098(NGT) | 1 | 1865.8947 | 1862.906439 | 9669981 | 67.72 | 1 | 3 | 5 | 0 | 0 | 0 | 0 | 1 | 3364.461776 | -0.0021 | -0.62 | -6.76 |
| Trypsin | 18 | 116 | 910 | 1 | 4398 | 2959.3106 | 61.3464 | 155,152 | 2958.302775 | 17 | 1092 | 1107 | EGVFVSNGTHWFVTQR | 0.0000002000000000.0 | 1098(NGT) | 1 | 1865.8947 | 1862.906439 | 9669981 | 67.72 | 1 | 3 | 3 | 0 | 0 | 0 | 0 | 1 | 2958.30303 | -0.0003 | -0.08 | -6.37 |
| Trypsin | 18 | 116 | 915 | 1 | 4458 | 3939.6471 | 64.2422 | 153,740 | 3938.639275 | 17 | 1092 | 1107 | EGVFVSNGTHWFVTQR | 0.0000002000000000.0 | 1098(NGT) | 1 | 1865.8947 | 1862.906439 | 9669981 | 67.72 | 1 | 6 | 4 | 0 | 1 | 0 | 0 | 1 | 3938.636292 | 0.0030 | 0.75 | -3.48 |
| Trypsin | 18 | 116 | 915 | 1 | 4458 | 3939.6471 | 64.2422 | 153,740 | 3938.639275 | 17 | 1092 | 1107 | EGVFVSNGTHWFVTQR | 0.0000002000000000.0 | 1098(NGT) | 1 | 1865.8947 | 1862.906439 | 9669981 | 67.72 | 2 | 5 | 4 | 1 | 0 | 1 | 0 | 1 | 3938.636291 | 0.0030 | 0.75 | -3.48 |
| Trypsin | 18 | 116 | 937 | 1 | 4379 | 3835.6296 | 60.3951 | 151,563 | 3834.621775 | 17 | 1092 | 1107 | EGVFVSNGTHWFVTQR | 0.0000002000000000.0 | 1098(NGT) | 1 | 1865.8947 | 1862.906439 | 9669981 | 67.72 | 1 | 5 | 5 | 1 | 0 | 0 | 0 | 1 | 3834.625333 | -0.0036 | -0.92 | -7.33 |
| Trypsin | 18 | 116 | 998 | 1 | 4380 | 3810.6057 | 60.4506 | 140,499 | 3809.597875 | 17 | 1092 | 1107 | EGVFVSNGTHWFVTQR | 0.0000002000000000.0 | 1098(NGT) | 1 | 1865.8947 | 1862.906439 | 9669981 | 67.72 | 1 | 7 | 4 | 0 | 0 | 0 | 0 | 1 | 3809.593699 | 0.0042 | 1.09 | -7.27 |
| Trypsin | 18 | 116 | 1011 | 1 | 4453 | 4288.7826 | 64.001 | 138,147 | 4287.774775 | 17 | 1092 | 1107 | EGVFVSNGTHWFVTQR | 0.0000002000000000.0 | 1098(NGT) | 1 | 1865.8947 | 1862.906439 | 9669981 | 67.72 | 1 | 6 | 5 | 1 | 1 | 0 | 0 | 1 | 4287.773574 | 0.0012 | 0.28 | -3.72 |
| Trypsin | 18 | 116 | 1011 | 1 | 4453 | 4288.7826 | 64.001 | 138,147 | 4287.774775 | 17 | 1092 | 1107 | EGVFVSNGTHWFVTQR | 0.0000002000000000.0 | 1098(NGT) | 1 | 1865.8947 | 1862.906439 | 9669981 | 67.72 | 2 | 5 | 5 | 2 | 0 | 1 | 0 | 0 | 4287.773573 | 0.0012 | 0.28 | -3.72 |
| Trypsin | 18 | 116 | 1123 | 1 | 4393 | 3162.3893 | 61.1097 | 123,409 | 3161.381475 | 17 | 1092 | 1107 | EGVFVSNGTHWFVTQR | 0.0000002000000000.0 | 1098(NGT) | 1 | 1865.8947 | 1862.906439 | 9669981 | 67.72 | 1 | 3 | 4 | 0 | 0 | 0 | 0 | 1 | 3161.382403 | -0.0009 | -0.29 | -6.61 |
| Trypsin | 18 | 116 | 1132 | 1 | 4446 | 4450.8276 | 63.6567 | 122,246 | 4449.819775 | 17 | 1092 | 1107 | EGVFVSNGTHWFVTQR | 0.0000002000000000.0 | 1098(NGT) | 1 | 1865.8947 | 1862.906439 | 9669981 | 67.72 | 1 | 6 | 5 | 2 | 0 | 1 | 0 | 1 | 4449.826397 | -0.0066 | -1.48 | -4.06 |
| Trypsin | 18 | 116 | 1132 | 1 | 4446 | 4450.8276 | 63.6567 | 122,246 | 4449.819775 | 17 | 1092 | 1107 | EGVFVSNGTHWFVTQR | 0.0000002000000000.0 | 1098(NGT) | 1 | 1865.8947 | 1862.906439 | 9669981 | 67.72 | 2 | 7 | 5 | 1 | 1 | 0 | 0 | 1 | 4449.826398 | -0.0066 | -1.48 | -4.06 |
| Trypsin | 18 | 116 | 1149 | 1 | 4386 | 3527.5254 | 60.7457 | 121,337 | 3526.517575 | 17 | 1092 | 1107 | EGVFVSNGTHWFVTQR | 0.0000002000000000.0 | 1098(NGT) | 1 | 1865.8947 | 1862.906439 | 9669981 | 67.72 | 1 | 4 | 5 | 0 | 0 | 0 | 0 | 1 | 3526.5146 | 0.0030 | 0.84 | -6.97 |
| Trypsin | 18 | 116 | 1201 | 1 | 4386 | 3470.5056 | 60.7457 | 115,316 | 3469.497775 | 17 | 1092 | 1107 | EGVFVSNGTHWFVTQR | 0.0000002000000000.0 | 1098(NGT) | 1 | 1865.8947 | 1862.906439 | 9669981 | 67.72 | 1 | 4 | 4 | 1 | 0 | 0 | 0 | 1 | 3469.493136 | 0.0046 | 1.33 | -6.97 |
| Trypsin | 18 | 116 | 1328 | 1 | 4383 | 3673.5798 | 60.5957 | 103,788 | 3672.571975 | 17 | 1092 | 1107 | EGVFVSNGTHWFVTQR | 0.0000002000000000.0 | 1098(NGT) | 1 | 1865.8947 | 1862.906439 | 9669981 | 67.72 | 1 | 4 | 5 | 1 | 0 | 0 | 0 | 1 | 3672.572509 | -0.0005 | -0.14 | -7.12 |
| Trypsin | 18 | 116 | 1345 | 1 | 4464 | 3777.5866 | 64.51 | 102,289 | 3776.578775 | 17 | 1092 | 1107 | EGVFVSNGTHWFVTQR | 0.0000002000000000.0 | 1098(NGT) | 1 | 1865.8947 | 1862.906439 | 9669981 | 67.72 | 1 | 4 | 4 | 1 | 0 | 1 | 0 | 1 | 3776.583467 | -0.0047 | -1.24 | -3.21 |
| Trypsin | 18 | 116 | 1345 | 1 | 4464 | 3777.5866 | 64.51 | 102,289 | 3776.578775 | 17 | 1092 | 1107 | EGVFVSNGTHWFVTQR | 0.0000002000000000.0 | 1098(NGT) | 1 | 1865.8947 | 1862.906439 | 9669981 | 67.72 | 2 | 5 | 4 | 0 | 1 | 0 | 0 | 1 | 3776.583468 | -0.0047 | -1.24 | -3.21 |
| Trypsin | 18 | 116 | 1351 | 1 | 4528 | 4830.9418 | 67.5705 | 101,592 | 4829.933975 | 17 | 1092 | 1107 | EGVFVSNGTHWFVTQR | 0.0000002000000000.0 | 1098(NGT) | 1 | 1865.8947 | 1862.906439 | 9669981 | 67.72 | 1 | 7 | 6 | 0 | 0 | 2 | 0 | 1 | 4829.933107 | 0.0009 | 0.17 | -0.15 |
| Trypsin | 18 | 116 | 1386 | 1 | 4377 | 4038.7075 | 60.2882 | 98,547 | 4037.699675 | 17 | 1092 | 1107 | EGVFVSNGTHWFVTQR | 0.0000002000000000.0 | 1098(NGT) | 1 | 1865.8947 | 1862.906439 | 9669981 | 67.72 | 1 | 5 | 6 | 1 | 0 | 0 | 0 | 1 | 4037.704706 | -0.0050 | -1.24 | -7.43 |
| Trypsin | 18 | 116 | 1399 | 1 | 4380 | 3876.6599 | 60.4506 | 97,748 | 3875.652075 | 17 | 1092 | 1107 | EGVFVSNGTHWFVTQR | 0.0000002000000000.0 | 1098(NGT) | 1 | 1865.8947 | 1862.906439 | 9669981 | 67.72 | 1 | 4 | 6 | 1 | 0 | 0 | 0 | 1 | 3875.651882 | 0.0002 | 0.04 | -7.27 |
| Trypsin | 18 | 116 | 1563 | 1 | 4384 | 3511.5267 | 60.6462 | 87,160 | 3510.518875 | 17 | 1092 | 1107 | EGVFVSNGTHWFVTQR | 0.0000002000000000.0 | 1098(NGT) | 1 | 1865.8947 | 1862.906439 | 9669981 | 67.72 | 1 | 3 | 5 | 1 | 0 | 0 | 0 | 1 | 3510.519685 | -0.0008 | -0.23 | -7.07 |
| Trypsin | 18 | 116 | 1569 | 1 | 4370 | 4346.8198 | 59.9461 | 86,569 | 4345.811975 | 17 | 1092 | 1107 | EGVFVSNGTHWFVTQR | 0.0000002000000000.0 | 1098(NGT) | 1 | 1865.8947 | 1862.906439 | 9669981 | 67.72 | 1 | 6 | 6 | 2 | 0 | 0 | 0 | 1 | 4345.815439 | -0.0035 | -0.79 | -7.77 |
| Trypsin | 18 | 116 | 1578 | 1 | 4360 | 4800.9895 | 59.4504 | 86,092 | 4799.981675 | 17 | 1092 | 1107 | EGVFVSNGTHWFVTQR | 0.0000002000000000.0 | 1098(NGT) | 1 | 1865.8947 | 1862.906439 | 9669981 | 67.72 | 1 | 7 | 6 | 4 | 0 | 0 | 0 | 1 | 4799.984081 | -0.0024 | -0.5 | -8.27 |
| Trypsin | 18 | 116 | 1578 | 1 | 4360 | 4800.9895 | 59.4504 | 86,092 | 4799.981675 | 17 | 1092 | 1107 | EGVFVSNGTHWFVTQR | 0.0000002000000000.0 | 1098(NGT) | 1 | 1865.8947 | 1862.906439 | 9669981 | 67.72 | 2 | 2 | 10 | 0 | 2 | 0 | 1 | -1 | 4799.996651 | -0.0150 | -3.11 | -8.27 |
| Trypsin | 18 | 116 | 1592 | 1 | 4463 | 3955.638 | 64.4678 | 85,296 | 3954.630175 | 17 | 1092 | 1107 | EGVFVSNGTHWFVTQR | 0.0000002000000000.0 | 1098(NGT) | 1 | 1865.8947 | 1862.906439 | 9669981 | 67.72 | 1 | 6 | 4 | 0 | 0 | 1 | 0 | 1 | 3954.631206 | -0.0010 | -0.26 | -3.25 |
| Trypsin | 18 | 116 | 1638 | 1 | 4455 | 4361.7916 | 64.0885 | 82,583 | 4360.783775 | 17 | 1092 | 1107 | EGVFVSNGTHWFVTQR | 0.0000002000000000.0 | 1098(NGT) | 1 | 1865.8947 | 1862.906439 | 9669981 | 67.72 | 1 | 6 | 6 | 0 | 0 | 1 | 0 | 0 | 4360.789952 | -0.0062 | -1.41 | -3.63 |
| Trypsin | 18 | 116 | 1662 | 1 | 4536 | 4465.8121 | 67.9767 | 81,306 | 4464.804275 | 17 | 1092 | 1107 | EGVFVSNGTHWFVTQR | 0.0000002000000000.0 | 1098(NGT) | 1 | 1865.8947 | 1862.906439 | 9669981 | 67.72 | 1 | 6 | 5 | 0 | 0 | 2 | 0 | 1 | 4464.80091 | 0.0034 | 0.75 | 0.26 |
| Trypsin | 18 | 116 | 1662 | 1 | 4536 | 4465.8121 | 67.9767 | 81,306 | 4464.804275 | 17 | 1092 | 1107 | EGVFVSNGTHWFVTQR | 0.0000002000000000.0 | 1098(NGT) | 1 | 1865.8947 | 1862.906439 | 9669981 | 67.72 | 2 | 5 | 3 | 4 | 1 | 1 | 1 | -1 | 4464.826062 | -0.0218 | -4.87 | 0.26 |
| Trypsin | 18 | 116 | 1662 | 1 | 4536 | 4465.8121 | 67.9767 | 81,306 | 4464.804275 | 17 | 1092 | 1107 | EGVFVSNGTHWFVTQR | 0.0000002000000000.0 | 1098(NGT) | 1 | 1865.8947 | 1862.906439 | 9669981 | 67.72 | 3 | 6 | 3 | 3 | 2 | 0 | 1 | -1 | 4464.826063 | -0.0218 | -4.87 | 0.26 |
| Trypsin | 18 | 116 | 1684 | 1 | 4458 | 4101.6939 | 64.2422 | 80,038 | 4100.686075 | 17 | 1092 | 1107 | EGVFVSNGTHWFVTQR | 0.0000002000000000.0 | 1098(NGT) | 1 | 1865.8947 | 1862.906439 | 9669981 | 67.72 | 1 | 6 | 4 | 1 | 0 | 1 | 0 | 1 | 4100.689115 | -0.0030 | -0.74 | -3.48 |
| Trypsin | 18 | 116 | 1684 | 1 | 4458 | 4101.6939 | 64.2422 | 80,038 | 4100.686075 | 17 | 1092 | 1107 | EGVFVSNGTHWFVTQR | 0.0000002000000000.0 | 1098(NGT) | 1 | 1865.8947 | 1862.906439 | 9669981 | 67.72 | 2 | 7 | 4 | 0 | 1 | 0 | 0 | 1 | 4100.689116 | -0.0030 | -0.74 | -3.48 |
| Trypsin | 18 | 116 | 1689 | 1 | 4377 | 3972.6518 | 60.2882 | 79,954 | 3971.643975 | 17 | 1092 | 1107 | EGVFVSNGTHWFVTQR | 0.0000002000000000.0 | 1098(NGT) | 1 | 1865.8947 | 1862.906439 | 9669981 | 67.72 | 1 | 8 | 4 | 0 | 0 | 0 | 0 | 0 | 3971.646523 | -0.0025 | -0.64 | -7.43 |
| Trypsin | 18 | 116 | 1750 | 1 | 4376 | 3956.6601 | 60.2381 | 76,906 | 3955.652275 | 17 | 1092 | 1107 | EGVFVSNGTHWFVTQR | 0.0000002000000000.0 | 1098(NGT) | 1 | 1865.8947 | 1862.906439 | 9669981 | 67.72 | 1 | 7 | 4 | 1 | 0 | 0 | 0 | 1 | 3955.651608 | 0.0007 | 0.16 | -7.48 |
| Trypsin | 18 | 116 | 1827 | 1 | 4382 | 3714.5991 | 60.5509 | 72,983 | 3713.591275 | 17 | 1092 | 1107 | EGVFVSNGTHWFVTQR | 0.0000002000000000.0 | 1098(NGT) | 1 | 1865.8947 | 1862.906439 | 9669981 | 67.72 | 1 | 3 | 6 | 1 | 0 | 0 | 0 | 1 | 3713.599058 | -0.0078 | -2.09 | -7.17 |
| Trypsin | 18 | 116 | 1844 | 1 | 4471 | 3574.5029 | 64.8473 | 72,308 | 3573.495075 | 17 | 1092 | 1107 | EGVFVSNGTHWFVTQR | 0.0000002000000000.0 | 1098(NGT) | 1 | 1865.8947 | 1862.906439 | 9669981 | 67.72 | 1 | 4 | 3 | 1 | 0 | 1 | 0 | 1 | 3573.504094 | -0.0090 | -2.52 | -2.87 |
| Trypsin | 18 | 116 | 1844 | 1 | 4471 | 3574.5029 | 64.8473 | 72,308 | 3573.495075 | 17 | 1092 | 1107 | EGVFVSNGTHWFVTQR | 0.0000002000000000.0 | 1098(NGT) | 1 | 1865.8947 | 1862.906439 | 9669981 | 67.72 | 2 | 5 | 3 | 0 | 1 | 0 | 0 | 1 | 3573.504095 | -0.0090 | -2.52 | -2.87 |
| Trypsin | 18 | 116 | 1991 | 1 | 4363 | 4435.8581 | 59.5965 | 65,641 | 4434.850275 | 17 | 1092 | 1107 | EGVFVSNGTHWFVTQR | 0.0000002000000000.0 | 1098(NGT) | 1 | 1865.8947 | 1862.906439 | 9669981 | 67.72 | 1 | 6 | 5 | 4 | 0 | 0 | 0 | 0 | 4434.851884 | -0.0016 | -0.36 | -8.12 |
| Trypsin | 18 | 116 | 2070 | 1 | 4462 | 3898.6168 | 64.4226 | 62,835 | 3897.608975 | 17 | 1092 | 1107 | EGVFVSNGTHWFVTQR | 0.0000002000000000.0 | 1098(NGT) | 1 | 1865.8947 | 1862.906439 | 9669981 | 67.72 | 1 | 6 | 3 | 1 | 0 | 1 | 0 | 1 | 3897.609742 | -0.0008 | -0.19 | -3.30 |
| Trypsin | 18 | 116 | 2070 | 1 | 4462 | 3898.6168 | 64.4226 | 62,835 | 3897.608975 | 17 | 1092 | 1107 | EGVFVSNGTHWFVTQR | 0.0000002000000000.0 | 1098(NGT) | 1 | 1865.8947 | 1862.906439 | 9669981 | 67.72 | 2 | 7 | 3 | 0 | 1 | 0 | 1 | -1 | 3897.609743 | -0.0008 | -0.19 | -3.30 |
| Trypsin | 18 | 116 | 2154 | 1 | 4372 | 4102.7157 | 60.0423 | 60,117 | 4101.707875 | 17 | 1092 | 1107 | EGVFVSNGTHWFVTQR | 0.0000002000000000.0 | 1098(NGT) | 1 | 1865.8947 | 1862.906439 | 9669981 | 67.72 | 1 | 7 | 4 | 2 | 0 | 0 | 0 | 1 | 4101.709517 | -0.0016 | -0.4 | -7.68 |
| Trypsin | 18 | 116 | 2401 | 1 | 4344 | 3868.6292 | 58.703 | 52,466 | 3867.621375 | 17 | 1092 | 1107 | EGVFVSNGTHWFVTQR | 0.0000002000000000.0 | 1098(NGT) | 1 | 1865.8947 | 1862.906439 | 9669981 | 67.72 | | | | | | | | | | | | |
| Trypsin | 18 | 116 | 2537 | 1 | 4445 | 4596.8789 | 63.6123 | 49,483 | 4595.871075 | 17 | 1092 | 1107 | EGVFVSNGTHWFVTQR | 0.0000002000000000.0 | 1098(NGT) | 1 | 1865.8947 | 1862.906439 | 9669981 | 67.72 | 1 | 6 | 5 | 3 | 0 | 1 | 0 | 1 | 4595.884306 | -0.0132 | -2.87 | -4.11 |
| Trypsin | 18 | 116 | 2537 | 1 | 4445 | 4596.8789 | 63.6123 | 49,483 | 4595.871075 | 17 | 1092 | 1107 | EGVFVSNGTHWFVTQR | 0.0000002000000000.0 | 1098(NGT) | 1 | 1865.8947 | 1862.906439 | 9669981 | 67.72 | 2 | 7 | 5 | 2 | 1 | 0 | 0 | 0 | 4595.884307 | -0.0132 | -2.87 | -4.11 |
| Trypsin | 18 | 116 | 2596 | 1 | 4372 | 3924.6701 | 60.0423 | 47,717 | 3923.662275 | 17 | 1092 | 1107 | EGVFVSNGTHWFVTQR | 0.0000002000000000.0 | 1098(NGT) | 1 | 1865.8947 | 1862.906439 | 9669981 | 67.72 | 1 | 5 | 4 | 3 | 0 | 0 | 0 | 1 | 3923.661778 | 0.0005 | 0.12 | -7.68 |
| Trypsin | 18 | 116 | 2669 | 1 | 4389 | 3308.4499 | 60.903 | 46,189 | 3307.442075 | 17 | 1092 | 1107 | EGVFVSNGTHWFVTQR | 0.0000002000000000.0 | 1098(NGT) | 1 | 1865.8947 | 1862.906439 | 9669981 | 67.72 | 1 | 3 | 4 | 1 | 0 | 0 | 0 | 1 | 3307.440312 | 0.0018 | 0.53 | -6.82 |
| Trypsin | 18 | 116 | 2715 | 1 | 4335 | 3665.5517 | 58.2445 | 45,110 | 3664.543875 | 17 | 1092 | 1107 | EGVFVSNGTHWFVTQR | 0.0000002000000000.0 | 1098(NGT) | 1 | 1865.8947 | 1862.906439 | 9669981 | 67.72 | | | | | | | | | | | | |
| Trypsin | 18 | 116 | 2797 | 1 | 4457 | 4117.6864 | 64.2008 | 43,365 | 4116.678575 | 17 | 1092 | 1107 | EGVFVSNGTHWFVTQR | 0.0000002000000000.0 | 1098(NGT) | 1 | 1865.8947 | 1862.906439 | 9669981 | 67.72 | 1 | 7 | 4 | 0 | 0 | 1 | 0 | 0 | 4116.68403 | -0.0055 | -1.32 | -3.52 |
| Trypsin | 18 | 116 | 2816 | 1 | 4360 | 4727.9603 | 59.4504 | 42,908 | 4726.952475 | 17 | 1092 | 1107 | EGVFVSNGTHWFVTQR | 0.0000002000000000.0 | 1098(NGT) | 1 | 1865.8947 | 1862.906439 | 9669981 | 67.72 | 1 | 8 | 7 | 1 | 0 | 0 | 0 | 1 | 4726.942551 | 0.0099 | 2.09 | -8.27 |
| Trypsin | 18 | 116 | 2965 | 1 | 4362 | 4581.8854 | 59.545 | 40,092 | 4580.877575 | 17 | 1092 | 1107 | EGVFVSNGTHWFVTQR | 0.0000002000000000.0 | 1098(NGT) | 1 | 1865.8947 | 1862.906439 | 9669981 | 67.72 | 1 | 8 | 7 | 0 | 0 | 0 | 0 | 1 | 4580.884642 | -0.0071 | -1.54 | -8.18 |
| Trypsin | 18 | 116 | 3025 | 1 | 4375 | 3981.6932 | 60.188 | 38,947 | 3980.685375 | 17 | 1092 | 1107 | EGVFVSNGTHWFVTQR | 0.0000002000000000.0 | 1098(NGT) | 1 | 1865.8947 | 1862.906439 | 9669981 | 67.72 | 1 | 5 | 5 | 2 | 0 | 0 | 0 | 1 | 3980.683242 | 0.0021 | 0.53 | -7.53 |
| Trypsin | 18 | 116 | 3152 | 1 | 4531 | 4814.9503 | 67.7355 | 36,956 | 4813.942475 | 17 | 1092 | 1107 | EGVFVSNGTHWFVTQR | 0.0000002000000000.0 | 1098(NGT) | 1 | 1865.8947 | 1862.906439 | 9669981 | 67.72 | 1 | 7 | 6 | 0 | 1 | 1 | 0 | 1 | 4813.938193 | 0.0043 | 0.88 | 0.01 |
| Trypsin | 18 | 116 | 3152 | 1 | 4531 | 4814.9503 | 67.7355 | 36,956 | 4813.942475 | 17 | 1092 | 1107 | EGVFVSNGTHWFVTQR | 0.0000002000000000.0 | 1098(NGT) | 1 | 1865.8947 | 1862.906439 | 9669981 | 67.72 | 2 | 6 | 6 | 1 | 0 | 2 | 0 | 1 | 4813.938192 | 0.0043 | 0.88 | 0.01 |
| Trypsin | 18 | 116 | 3152 | 1 | 4531 | 4814.9503 | 67.7355 | 36,956 | 4813.942475 | 17 | 1092 | 1107 | EGVFVSNGTHWFVTQR | 0.0000002000000000.0 | 1098(NGT) | 1 | 1865.8947 | 1862.906439 | 9669981 | 67.72 | 3 | 6 | 4 | 4 | 2 | 0 | 0 | 0 | 4813.963345 | -0.0209 | -4.33 | 0.01 |
| Trypsin | 18 | 116 | 3221 | 1 | 4525 | 4976.9975 | 67.4253 | 35,957 | 4975.989675 | 17 | 1092 | 1107 | EGVFVSNGTHWFVTQR | 0.0000002000000000.0 | 1098(NGT) | 1 | 1865.8947 | 1862.906439 | 9669981 | 67.72 | 1 | 7 | 6 | 1 | 0 | 2 | 0 | 0 | 4975.991016 | -0.0013 | -0.26 | -0.30 |
| Trypsin | 18 | 116 | 3221 | 1 | 4525 | 4976.9975 | 67.4253 | 35,957 | 4975.989675 | 17 | 1092 | 1107 | EGVFVSNGTHWFVTQR | 0.0000002000000000.0 | 1098(NGT) | 1 | 1865.8947 | 1862.906439 | 9669981 | 67.72 | 2 | 8 | 6 | 0 | 1 | 1 | 0 | 0 | 4975.991017 | -0.0013 | -0.26 | -0.30 |
| Trypsin | 18 | 116 | 3293 | 1 | 4339 | 3690.5833 | 58.4704 | 34,788 | 3689.575475 | 17 | 1092 | 1107 | EGVFVSNGTHWFVTQR | 0.0000002000000000.0 | 1098(NGT) | 1 | 1865.8947 | 1862.906439 | 9669981 | 67.72 | | | | | | | | | | | | |
| Trypsin | 18 | 116 | 3414 | 1 | 4400 | 4727.9386 | 61.4502 | 32,975 | 4726.930775 | 17 | 1092 | 1107 | EGVFVSNGTHWFVTQR | 0.0000002000000000.0 | 1098(NGT) | 1 | 1865.8947 | 1862.906439 | 9669981 | 67.72 | 1 | 8 | 7 | 1 | 0 | 0 | 0 | 1 | 4726.942551 | -0.0118 | -2.49 | -6.27 |
| Trypsin | 18 | 116 | 3417 | 1 | 4472 | 3428.4587 | 64.9029 | 32,936 | 3427.450875 | 17 | 1092 | 1107 | EGVFVSNGTHWFVTQR | 0.0000002000000000.0 | 1098(NGT) | 1 | 1865.8947 | 1862.906439 | 9669981 | 67.72 | 1 | 4 | 3 | 0 | 0 | 1 | 0 | 1 | 3427.446185 | 0.0047 | 1.36 | -2.82 |
| Trypsin | 18 | 116 | 3421 | 1 | 4367 | 4492.8852 | 59.784 | 32,908 | 4491.877375 | 17 | 1092 | 1107 | EGVFVSNGTHWFVTQR | 0.0000002000000000.0 | 1098(NGT) | 1 | 1865.8947 | 1862.906439 | 9669981 | 67.72 | 1 | 6 | 6 | 3 | 0 | 0 | 0 | 1 | 4491.873348 | 0.0040 | 0.89 | -7.94 |
| Trypsin | 18 | 116 | 3486 | 1 | 4370 | 4159.7292 | 59.9461 | 32,023 | 4158.721375 | 17 | 1092 | 1107 | EGVFVSNGTHWFVTQR | 0.0000002000000000.0 | 1098(NGT) | 1 | 1865.8947 | 1862.906439 | 9669981 | 67.72 | 1 | 7 | 5 | 1 | 0 | 0 | 0 | 1 | 4158.730981 | -0.0096 | -2.3 | -7.77 |
| Trypsin | 18 | 116 | 3554 | 1 | 4443 | 4799.9706 | 63.5267 | 31,327 | 4798.962775 | 17 | 1092 | 1107 | EGVFVSNGTHWFVTQR | 0.0000002000000000.0 | 1098(NGT) | 1 | 1865.8947 | 1862.906439 | 9669981 | 67.72 | 1 | 6 | 6 | 3 | 0 | 1 | 0 | 0 | 4798.963679 | -0.0009 | -0.18 | -4.19 |
| Trypsin | 18 | 116 | 3554 | 1 | 4443 | 4799.9706 | 63.5267 | 31,327 | 4798.962775 | 17 | 1092 | 1107 | EGVFVSNGTHWFVTQR | 0.0000002000000000.0 | 1098(NGT) | 1 | 1865.8947 | 1862.906439 | 9669981 | 67.72 | 2 | 7 | 6 | 2 | 1 | 0 | 0 | 0 | 4798.96368 | -0.0009 | -0.18 | -4.19 |
| Trypsin | 18 | 116 | 3555 | 1 | 4460 | 3923.6543 | 64.3359 | 31,318 | 3922.646475 | 17 | 1092 | 1107 | EGVFVSNGTHWFVTQR | 0.0000002000000000.0 | 1098(NGT) | 1 | 1865.8947 | 1862.906439 | 9669981 | 67.72 | 1 | 5 | 4 | 1 | 1 | 0 | 0 | 1 | 3922.641377 | 0.0051 | 1.29 | -3.38 |
| Trypsin | 18 | 116 | 3555 | 1 | 4460 | 3923.6543 | 64.3359 | 31,318 | 3922.646475 | 17 | 1092 | 1107 | EGVFVSNGTHWFVTQR | 0.0000002000000000.0 | 1098(NGT) | 1 | 1865.8947 | 1862.906439 | 9669981 | 67.72 | 2 | 4 | 4 | 2 | 0 | 1 | 0 | 1 | 3922.641376 | 0.0051 | 1.29 | -3.38 |
| Trypsin | 18 | 116 | 3556 | 1 | 4449 | 4434.8452 | 63.7991 | 31,244 | 4433.837375 | 17 | 1092 | 1107 | EGVFVSNGTHWFVTQR | 0.0000002000000000.0 | 1098(NGT) | 1 | 1865.8947 | 1862.906439 | 9669981 | 67.72 | 1 | 6 | 5 | 2 | 1 | 0 | 0 | 1 | 4433.831483 | 0.0059 | 1.32 | -3.92 |
| Trypsin | 18 | 116 | 3556 | 1 | 4449 | 4434.8452 | 63.7991 | 31,244 | 4433.837375 | 17 | 1092 | 1107 | EGVFVSNGTHWFVTQR | 0.0000002000000000.0 | 1098(NGT) | 1 | 1865.8947 | 1862.906439 | 9669981 | 67.72 | 2 | 5 | 5 | 3 | 0 | 1 | 0 | 0 | 4433.831482 | 0.0059 | 1.32 | -3.92 |
| Trypsin | 18 | 116 | 3558 | 1 | 4374 | 4013.6784 | 60.146 | 31,204 | 4012.670575 | 17 | 1092 | 1107 | EGVFVSNGTHWFVTQR | 0.0000002000000000.0 | 1098(NGT) | 1 | 1865.8947 | 1862.906439 | 9669981 | 67.72 | 1 | 7 | 5 | 0 | 0 | 0 | 0 | 1 | 4012.673072 | -0.0025 | -0.62 | -7.57 |
| Trypsin | 18 | 116 | 3615 | 1 | 4366 | 4378.8087 | 59.7429 | 30,480 | 4377.800875 | 17 | 1092 | 1107 | EGVFVSNGTHWFVTQR | 0.0000002000000000.0 | 1098(NGT) | 1 | 1865.8947 | 1862.906439 | 9669981 | 67.72 | 1 | 8 | 6 | 0 | 0 | 0 | 0 | 0 | 4377.805269 | -0.0044 | -1 | -7.98 |
| Trypsin | 18 | 116 | 3630 | 1 | 4538 | 4449.8089 | 68.0797 | 30,307 | 4448.801075 | 17 | 1092 | 1107 | EGVFVSNGTHWFVTQR | 0.0000002000000000.0 | 1098(NGT) | 1 | 1865.8947 | 1862.906439 | 9669981 | 67.72 | 1 | 5 | 5 | 1 | 0 | 2 | 0 | 1 | 4448.805995 | -0.0049 | -1.1 | 0.36 |
| Trypsin | 18 | 116 | 3630 | 1 | 4538 | 4449.8089 | 68.0797 | 30,307 | 4448.801075 | 17 | 1092 | 1107 | EGVFVSNGTHWFVTQR | 0.0000002000000000.0 | 1098(NGT) | 1 | 1865.8947 | 1862.906439 | 9669981 | 67.72 | 2 | 6 | 5 | 0 | 1 | 1 | 0 | 1 | 4448.805996 | -0.0049 | -1.1 | 0.36 |
| Trypsin | 18 | 116 | 3675 | 1 | 4303 | 3811.6246 | 56.751 | 29,715 | 3810.616775 | 17 | 1092 | 1107 | EGVFVSNGTHWFVTQR | 0.0000002000000000.0 | 1098(NGT) | 1 | 1865.8947 | 1862.906439 | 9669981 | 67.72 | | | | | | | | | | | | |
| Trypsin | 18 | 116 | 3717 | 1 | 4369 | 4403.8575 | 59.8897 | 29,302 | 4402.849675 | 17 | 1092 | 1107 | EGVFVSNGTHWFVTQR | 0.0000002000000000.0 | 1098(NGT) | 1 | 1865.8947 | 1862.906439 | 9669981 | 67.72 | 1 | 6 | 7 | 1 | 0 | 0 | 0 | 1 | 4402.836903 | 0.0128 | 2.9 | -7.83 |
| Trypsin | 18 | 116 | 3779 | 1 | 4358 | 4874.0079 | 59.3497 | 28,548 | 4873.000075 | 17 | 1092 | 1107 | EGVFVSNGTHWFVTQR | 0.0000002000000000.0 | 1098(NGT) | 1 | 1865.8947 | 1862.906439 | 9669981 | 67.72 | 1 | 8 | 7 | 2 | 0 | 0 | 0 | 0 | 4873.00046 | -0.0004 | -0.07 | -8.37 |
| Trypsin | 18 | 116 | 3997 | 1 | 4466 | 3720.5687 | 64.6114 | 26,239 | 3719.560875 | 17 | 1092 | 1107 | EGVFVSNGTHWFVTQR | 0.0000002000000000.0 | 1098(NGT) | 1 | 1865.8947 | 1862.906439 | 9669981 | 67.72 | 1 | 4 | 3 | 2 | 0 | 1 | 0 | 0 | 3719.562003 | -0.0011 | -0.3 | -3.11 |
| Trypsin | 18 | 116 | 3997 | 1 | 4466 | 3720.5687 | 64.6114 | 26,239 | 3719.560875 | 17 | 1092 | 1107 | EGVFVSNGTHWFVTQR | 0.0000002000000000.0 | 1098(NGT) | 1 | 1865.8947 | 1862.906439 | 9669981 | 67.72 | 2 | 5 | 3 | 1 | 1 | 0 | 0 | 0 | 3719.562004 | -0.0011 | -0.3 | -3.11 |
| Trypsin | 18 | 116 | 4035 | 1 | 4374 | 4118.7246 | 60.146 | 25,747 | 4117.716775 | 17 | 1092 | 1107 | EGVFVSNGTHWFVTQR | 0.0000002000000000.0 | 1098(NGT) | 1 | 1865.8947 | 1862.906439 | 9669981 | 67.72 | 1 | 8 | 4 | 1 | 0 | 0 | 0 | 0 | 4117.704432 | 0.0123 | 2.99 | -7.57 |
| Trypsin | 18 | 116 | 4105 | 1 | 4440 | 4962.0205 | 63.3741 | 24,923 | 4961.012675 | 17 | 1092 | 1107 | EGVFVSNGTHWFVTQR | 0.0000002000000000.0 | 1098(NGT) | 1 | 1865.8947 | 1862.906439 | 9669981 | 67.72 | 1 | 7 | 6 | 3 | 0 | 1 | 0 | 0 | 4961.016503 | -0.0038 | -0.77 | -4.35 |
| Trypsin | 18 | 116 | 4105 | 1 | 4440 | 4962.0205 | 63.3741 | 24,923 | 4961.012675 | 17 | 1092 | 1107 | EGVFVSNGTHWFVTQR | 0.0000002000000000.0 | 1098(NGT) | 1 | 1865.8947 | 1862.906439 | 9669981 | 67.72 | 2 | 8 | 6 | 2 | 1 | 0 | 0 | 0 | 4961.016504 | -0.0038 | -0.77 | -4.35 |
| Trypsin | 18 | 116 | 4107 | 1 | 4373 | 4086.7203 | 60.0965 | 24,905 | 4085.712475 | 17 | 1092 | 1107 | EGVFVSNGTHWFVTQR | 0.0000002000000000.0 | 1098(NGT) | 1 | 1865.8947 | 1862.906439 | 9669981 | 67.72 | 1 | 6 | 4 | 3 | 0 | 0 | 0 | 1 | 4085.714602 | -0.0021 | -0.52 | -7.62 |
| Trypsin | 18 | 116 | 4121 | 1 | 4436 | 5035.0414 | 63.165 | 24,741 | 5034.033575 | 17 | 1092 | 1107 | EGVFVSNGTHWFVTQR | 0.0000002000000000.0 | 1098(NGT) | 1 | 1865.8947 | 1862.906439 | 9669981 | 67.72 | 1 | 9 | 7 | 0 | 1 | 0 | 0 | 0 | 5034.032883 | 0.0007 | 0.13 | -4.56 |
| Trypsin | 18 | 116 | 4121 | 1 | 4436 | 5035.0414 | 63.165 | 24,741 | 5034.033575 | 17 | 1092 | 1107 | EGVFVSNGTHWFVTQR | 0.0000002000000000.0 | 1098(NGT) | 1 | 1865.8947 | 1862.906439 | 9669981 | 67.72 | 2 | 8 | 7 | 1 | 0 | 1 | 0 | 0 | 5034.032882 | 0.0007 | 0.13 | -4.56 |
| Trypsin | 18 | 116 | 4267 | 1 | 4459 | 4085.7055 | 64.2919 | 23,406 | 4084.697675 | 17 | 1092 | 1107 | EGVFVSNGTHWFVTQR | 0.0000002000000000.0 | 1098(NGT) | 1 | 1865.8947 | 1862.906439 | 9669981 | 67.72 | 1 | 6 | 4 | 1 | 1 | 0 | 0 | 1 | 4084.694201 | 0.0035 | 0.85 | -3.43 |
| Trypsin | 18 | 116 | 4267 | 1 | 4459 | 4085.7055 | 64.2919 | 23,406 | 4084.697675 | 17 | 1092 | 1107 | EGVFVSNGTHWFVTQR | 0.0000002000000000.0 | 1098(NGT) | 1 | 1865.8947 | 1862.906439 | 9669981 | 67.72 | 2 | 5 | 4 | 2 | 0 | 1 | 0 | 1 | 4084.6942 | 0.0035 | 0.85 | -3.43 |
| Trypsin | 18 | 116 | 4268 | 1 | 4393 | 3105.3676 | 61.1097 | 23,404 | 3104.359775 | 17 | 1092 | 1107 | EGVFVSNGTHWFVTQR | 0.0000002000000000.0 | 1098(NGT) | 1 | 1865.8947 | 1862.906439 | 9669981 | 67.72 | 1 | 3 | 3 | 1 | 0 | 0 | 0 | 1 | 3104.360939 | -0.0012 | -0.37 | -6.61 |
| Trypsin | 18 | 116 | 4425 | 1 | 4366 | 4305.7932 | 59.7429 | 22,064 | 4304.785375 | 17 | 1092 | 1107 | EGVFVSNGTHWFVTQR | 0.0000002000000000.0 | 1098(NGT) | 1 | 1865.8947 | 1862.906439 | 9669981 | 67.72 | 1 | 7 | 5 | 2 | 0 | 0 | 0 | 0 | 4304.78889 | -0.0035 | -0.81 | -7.98 |
| Trypsin | 18 | 116 | 4511 | 1 | 4374 | 4184.7806 | 60.146 | 21,314 | 4183.772775 | 17 | 1092 | 1107 | EGVFVSNGTHWFVTQR | 0.0000002000000000.0 | 1098(NGT) | 1 | 1865.8947 | 1862.906439 | 9669981 | 67.72 | 1 | 5 | 6 | 2 | 0 | 0 | 0 | 1 | 4183.762615 | 0.0102 | 2.42 | -7.57 |
| Trypsin | 18 | 116 | 5483 | 1 | 4416 | 3794.5939 | 62.1803 | 14,286 | 3793.586075 | 17 | 1092 | 1107 | EGVFVSNGTHWFVTQR | 0.0000002000000000.0 | 1098(NGT) | 1 | 1865.8947 | 1862.906439 | 9669981 | 67.72 | 1 | 6 | 4 | 1 | 0 | 0 | 0 | 1 | 3793.598784 | -0.0127 | -3.35 | -5.54 |
| Trypsin | 18 | 116 | 6028 | 1 | 4452 | 4320.7496 | 63.9483 | 10,948 | 4319.741775 | 17 | 1092 | 1107 | EGVFVSNGTHWFVTQR | 0.0000002000000000.0 | 1098(NGT) | 1 | 1865.8947 | 1862.906439 | 9669981 | 67.72 | | | | | | | | | | | | |
| Trypsin | 18 | 116 | 6036 | 1 | 4408 | 4362.8038 | 61.8309 | 10,909 | 4361.795975 | 17 | 1092 | 1107 | EGVFVSNGTHWFVTQR | 0.0000002000000000.0 | 1098(NGT) | 1 | 1865.8947 | 1862.906439 | 9669981 | 67.72 | 1 | 7 | 6 | 1 | 0 | 0 | 0 | 1 | 4361.810354 | -0.0144 | -3.29 | -5.89 |
| Trypsin | 18 | 116 | 6075 | 1 | 4398 | 2918.2903 | 61.3464 | 10,679 | 2917.282475 | 17 | 1092 | 1107 | EGVFVSNGTHWFVTQR | 0.0000002000000000.0 | 1098(NGT) | 1 | 1865.8947 | 1862.906439 | 9669981 | 67.72 | 1 | 4 | 2 | 0 | 0 | 0 | 0 | 1 | 2917.276481 | 0.0060 | 2.05 | -6.37 |
| Trypsin | 18 | 116 | 6700 | 1 | 4450 | 4466.8227 | 63.8506 | 6,753 | 4465.814875 | 17 | 1092 | 1107 | EGVFVSNGTHWFVTQR | 0.0000002000000000.0 | 1098(NGT) | 1 | 1865.8947 | 1862.906439 | 9669981 | 67.72 | 1 | 7 | 5 | 1 | 0 | 1 | 0 | 0 | 4465.821312 | -0.0064 | -1.44 | -3.87 |
| Trypsin | 18 | 116 | 6700 | 1 | 4450 | 4466.8227 | 63.8506 | 6,753 | 4465.814875 | 17 | 1092 | 1107 | EGVFVSNGTHWFVTQR | 0.0000002000000000.0 | 1098(NGT) | 1 | 1865.8947 | 1862.906439 | 9669981 | 67.72 | 2 | 8 | 5 | 0 | 1 | 0 | 0 | 0 | 4465.821313 | -0.0064 | -1.44 | -3.87 |
| Trypsin | 18 | 116 | 7118 | 1 | 4332 | 3893.7008 | 58.1056 | 3,740 | 3892.692975 | 17 | 1092 | 1107 | EGVFVSNGTHWFVTQR | 0.0000002000000000.0 | 1098(NGT) | 1 | 1865.8947 | 1862.906439 | 9669981 | 67.72 | | | | | | | | | | | | |
| Trypsin | 18 | 116 | 7175 | 1 | 4551 | 4027.6613 | 68.6773 | 3,308 | 4026.653475 | 17 | 1092 | 1107 | EGVFVSNGTHWFVTQR | 0.0000002000000000.0 | 1098(NGT) | 1 | 1865.8947 | 1862.906439 | 9669981 | 67.72 | 1 | 6 | 3 | 0 | 2 | 0 | 1 | -1 | 4026.652336 | 0.0011 | 0.28 | 0.96 |
| Trypsin | 18 | 116 | 7175 | 1 | 4551 | 4027.6613 | 68.6773 | 3,308 | 4026.653475 | 17 | 1092 | 1107 | EGVFVSNGTHWFVTQR | 0.0000002000000000.0 | 1098(NGT) | 1 | 1865.8947 | 1862.906439 | 9669981 | 67.72 | 2 | 5 | 3 | 1 | 1 | 1 | 0 | 0 | 4026.652335 | 0.0011 | 0.28 | 0.96 |
| Trypsin | 18 | 116 | 7175 | 1 | 4551 | 4027.6613 | 68.6773 | 3,308 | 4026.653475 | 17 | 1092 | 1107 | EGVFVSNGTHWFVTQR | 0.0000002000000000.0 | 1098(NGT) | 1 | 1865.8947 | 1862.906439 | 9669981 | 67.72 | 3 | 4 | 3 | 2 | 0 | 2 | 1 | -1 | 4026.652334 | 0.0011 | 0.28 | 0.96 |
| Trypsin | 182 | 7 | 4004 | 1 | 4363 | 4525.9152 | 59.5965 | 26,171 | 4524.907375 | 231 | 1093 | 1107 | GVFVSNGTHWFVTQR | 0.000002000000000.0 | 1098(NGT) | 1 | 1736.8521 | 1733.863839 | 210731.66 | 65.93 | 1 | 1 | 10 | 0 | 1 | 1 | 1 | -1 | 4524.896141 | 0.0112 | 2.48 | -6.33 |
| Trypsin | 182 | 7 | 4090 | 1 | 4443 | 4686.9249 | 63.5267 | 25,101 | 4685.917075 | 231 | 1093 | 1107 | GVFVSNGTHWFVTQR | 0.000002000000000.0 | 1098(NGT) | 1 | 1736.8521 | 1733.863839 | 210731.66 | 65.93 | 1 | 8 | 6 | 1 | 1 | 0 | 0 | 1 | 4685.915995 | 0.0011 | 0.23 | -2.40 |
| Trypsin | 182 | 7 | 4090 | 1 | 4443 | 4686.9249 | 63.5267 | 25,101 | 4685.917075 | 231 | 1093 | 1107 | GVFVSNGTHWFVTQR | 0.000002000000000.0 | 1098(NGT) | 1 | 1736.8521 | 1733.863839 | 210731.66 | 65.93 | 2 | 7 | 6 | 2 | 0 | 1 | 0 | 0 | 4685.915994 | 0.0011 | 0.23 | -2.40 |
| Trypsin | 182 | 7 | 5063 | 1 | 4361 | 4671.9713 | 59.5024 | 17,147 | 4670.963475 | 231 | 1093 | 1107 | GVFVSNGTHWFVTQR | 0.000002000000000.0 | 1098(NGT) | 1 | 1736.8521 | 1733.863839 | 210731.66 | 65.93 | 1 | 7 | 6 | 4 | 0 | 0 | 0 | 1 | 4670.941481 | 0.0220 | 4.7 | -6.42 |
| Trypsin | 182 | 7 | 5063 | 1 | 4361 | 4671.9713 | 59.5024 | 17,147 | 4670.963475 | 231 | 1093 | 1107 | GVFVSNGTHWFVTQR | 0.000002000000000.0 | 1098(NGT) | 1 | 1736.8521 | 1733.863839 | 210731.66 | 65.93 | 2 | 2 | 10 | 0 | 2 | 0 | 1 | -1 | 4670.954051 | 0.0094 | 2.01 | -6.42 |
| Trypsin | 182 | 7 | 5109 | 1 | 4447 | 4540.8553 | 63.7083 | 16,870 | 4539.847475 | 231 | 1093 | 1107 | GVFVSNGTHWFVTQR | 0.000002000000000.0 | 1098(NGT) | 1 | 1736.8521 | 1733.863839 | 210731.66 | 65.93 | 1 | 7 | 6 | 1 | 0 | 1 | 0 | 1 | 4539.858085 | -0.0106 | -2.33 | -2.22 |
| Trypsin | 182 | 7 | 5109 | 1 | 4447 | 4540.8553 | 63.7083 | 16,870 | 4539.847475 | 231 | 1093 | 1107 | GVFVSNGTHWFVTQR | 0.000002000000000.0 | 1098(NGT) | 1 | 1736.8521 | 1733.863839 | 210731.66 | 65.93 | 2 | 8 | 6 | 0 | 1 | 0 | 0 | 1 | 4539.858086 | -0.0106 | -2.33 | -2.22 |
| Trypsin | 182 | 7 | 5448 | 1 | 4438 | 4889.9862 | 63.2661 | 14,495 | 4888.978375 | 231 | 1093 | 1107 | GVFVSNGTHWFVTQR | 0.000002000000000.0 | 1098(NGT) | 1 | 1736.8521 | 1733.863839 | 210731.66 | 65.93 | 1 | 8 | 7 | 1 | 1 | 0 | 0 | 1 | 4888.995368 | -0.0170 | -3.47 | -2.66 |
| Trypsin | 182 | 7 | 5448 | 1 | 4438 | 4889.9862 | 63.2661 | 14,495 | 4888.978375 | 231 | 1093 | 1107 | GVFVSNGTHWFVTQR | 0.000002000000000.0 | 1098(NGT) | 1 | 1736.8521 | 1733.863839 | 210731.66 | 65.93 | 2 | 7 | 7 | 2 | 0 | 1 | 0 | 0 | 4888.995367 | -0.0170 | -3.47 | -2.66 |
| Trypsin | 182 | 7 | 5992 | 1 | 4443 | 4832.9889 | 63.5267 | 11,151 | 4831.981075 | 231 | 1093 | 1107 | GVFVSNGTHWFVTQR | 0.000002000000000.0 | 1098(NGT) | 1 | 1736.8521 | 1733.863839 | 210731.66 | 65.93 | 1 | 8 | 6 | 2 | 1 | 0 | 0 | 1 | 4831.973904 | 0.0072 | 1.48 | -2.40 |
| Trypsin | 182 | 7 | 5992 | 1 | 4443 | 4832.9889 | 63.5267 | 11,151 | 4831.981075 | 231 | 1093 | 1107 | GVFVSNGTHWFVTQR | 0.000002000000000.0 | 1098(NGT) | 1 | 1736.8521 | 1733.863839 | 210731.66 | 65.93 | 2 | 7 | 6 | 3 | 0 | 1 | 0 | 0 | 4831.973903 | 0.0072 | 1.48 | -2.40 |
| Trypsin | 182 | 7 | 6476 | 1 | 4511 | 5197.07 | 66.7439 | 8,366 | 5196.062175 | 231 | 1093 | 1107 | GVFVSNGTHWFVTQR | 0.000002000000000.0 | 1098(NGT) | 1 | 1736.8521 | 1733.863839 | 210731.66 | 65.93 | 1 | 8 | 7 | 1 | 1 | 1 | 0 | 1 | 5196.085699 | -0.0235 | -4.52 | 0.82 |
| Trypsin | 182 | 7 | 6476 | 1 | 4511 | 5197.07 | 66.7439 | 8,366 | 5196.062175 | 231 | 1093 | 1107 | GVFVSNGTHWFVTQR | 0.000002000000000.0 | 1098(NGT) | 1 | 1736.8521 | 1733.863839 | 210731.66 | 65.93 | 2 | 7 | 7 | 2 | 0 | 2 | 0 | 0 | 5196.085698 | -0.0235 | -4.52 | 0.82 |
| Trypsin | 182 | 7 | 6476 | 1 | 4511 | 5197.07 | 66.7439 | 8,366 | 5196.062175 | 231 | 1093 | 1107 | GVFVSNGTHWFVTQR | 0.000002000000000.0 | 1098(NGT) | 1 | 1736.8521 | 1733.863839 | 210731.66 | 65.93 | 3 | 9 | 7 | 0 | 2 | 0 | 0 | 0 | 5196.0857 | -0.0235 | -4.52 | 0.82 |
| alpha-Lytic | 33 | 36 | 167 | 1 | 4433 | 3314.2998 | 57.5947 | 794,370 | 3313.291975 | 23 | 1098 | 1105 | NGTHWFVT | 0.20000000.0 | 1098(NGT) | 1 | 963.4337 | 960.4454387 | 5022417.5 | 69.02 | 1 | 7 | 6 | 0 | 0 | 0 | 0 | 1 | 3313.291445 | 0.0005 | 0.16 | -11.42 |
| alpha-Lytic | 33 | 36 | 211 | 1 | 4453 | 2689.0656 | 58.6333 | 654,141 | 2688.057775 | 23 | 1098 | 1105 | NGTHWFVT | 0.20000000.0 | 1098(NGT) | 1 | 963.4337 | 960.4454387 | 5022417.5 | 69.02 | 1 | 6 | 3 | 1 | 0 | 0 | 0 | 1 | 2688.058411 | -0.0006 | -0.23 | -10.38 |
| alpha-Lytic | 33 | 36 | 264 | 1 | 4428 | 3460.3593 | 57.3394 | 510,298 | 3459.351475 | 23 | 1098 | 1105 | NGTHWFVT | 0.20000000.0 | 1098(NGT) | 1 | 963.4337 | 960.4454387 | 5022417.5 | 69.02 | 1 | 7 | 6 | 1 | 0 | 0 | 0 | 1 | 3459.349354 | 0.0021 | 0.61 | -11.68 |
| alpha-Lytic | 33 | 36 | 280 | 1 | 4423 | 3606.4138 | 57.0931 | 464,477 | 3605.405975 | 23 | 1098 | 1105 | NGTHWFVT | 0.20000000.0 | 1098(NGT) | 1 | 963.4337 | 960.4454387 | 5022417.5 | 69.02 | 1 | 7 | 6 | 2 | 0 | 0 | 0 | 0 | 3605.407263 | -0.0013 | -0.35 | -11.92 |
| alpha-Lytic | 33 | 36 | 335 | 1 | 4445 | 2949.1665 | 58.2399 | 400,193 | 2948.158675 | 23 | 1098 | 1105 | NGTHWFVT | 0.20000000.0 | 1098(NGT) | 1 | 963.4337 | 960.4454387 | 5022417.5 | 69.02 | 1 | 6 | 5 | 0 | 0 | 0 | 0 | 1 | 2948.159248 | -0.0006 | -0.19 | -10.78 |
| alpha-Lytic | 33 | 36 | 392 | 1 | 4461 | 2543.0086 | 59.002 | 330,706 | 2542.000775 | 23 | 1098 | 1105 | NGTHWFVT | 0.20000000.0 | 1098(NGT) | 1 | 963.4337 | 960.4454387 | 5022417.5 | 69.02 | 1 | 6 | 3 | 0 | 0 | 0 | 0 | 1 | 2542.000502 | 0.0003 | 0.1 | -10.01 |
| alpha-Lytic | 33 | 36 | 410 | 1 | 4514 | 3621.3904 | 61.567 | 318,068 | 3620.382575 | 23 | 1098 | 1105 | NGTHWFVT | 0.20000000.0 | 1098(NGT) | 1 | 963.4337 | 960.4454387 | 5022417.5 | 69.02 | 1 | 7 | 6 | 0 | 0 | 1 | 0 | 1 | 3620.381776 | 0.0008 | 0.22 | -7.45 |
| alpha-Lytic | 33 | 36 | 424 | 1 | 4440 | 3095.2241 | 57.972 | 307,259 | 3094.216275 | 23 | 1098 | 1105 | NGTHWFVT | 0.20000000.0 | 1098(NGT) | 1 | 963.4337 | 960.4454387 | 5022417.5 | 69.02 | 1 | 6 | 5 | 1 | 0 | 0 | 0 | 1 | 3094.217157 | -0.0009 | -0.28 | -11.04 |
| alpha-Lytic | 33 | 36 | 455 | 1 | 4461 | 2527.0152 | 59.002 | 278,324 | 2526.007375 | 23 | 1098 | 1105 | NGTHWFVT | 0.20000000.0 | 1098(NGT) | 1 | 963.4337 | 960.4454387 | 5022417.5 | 69.02 | 1 | 5 | 3 | 1 | 0 | 0 | 0 | 1 | 2526.005587 | 0.0018 | 0.7 | -10.01 |
| alpha-Lytic | 33 | 36 | 467 | 1 | 4509 | 3767.4444 | 61.3161 | 273,378 | 3766.436575 | 23 | 1098 | 1105 | NGTHWFVT | 0.20000000.0 | 1098(NGT) | 1 | 963.4337 | 960.4454387 | 5022417.5 | 69.02 | 1 | 7 | 6 | 1 | 0 | 1 | 0 | 1 | 3766.439685 | -0.0031 | -0.82 | -7.70 |
| alpha-Lytic | 33 | 36 | 467 | 1 | 4509 | 3767.4444 | 61.3161 | 273,378 | 3766.436575 | 23 | 1098 | 1105 | NGTHWFVT | 0.20000000.0 | 1098(NGT) | 1 | 963.4337 | 960.4454387 | 5022417.5 | 69.02 | 2 | 8 | 6 | 0 | 1 | 0 | 0 | 1 | 3766.439686 | -0.0031 | -0.82 | -7.70 |
| alpha-Lytic | 33 | 36 | 527 | 1 | 4446 | 2851.1185 | 58.2876 | 249,810 | 2850.110675 | 23 | 1098 | 1105 | NGTHWFVT | 0.20000000.0 | 1098(NGT) | 1 | 963.4337 | 960.4454387 | 5022417.5 | 69.02 | 1 | 7 | 3 | 1 | 0 | 0 | 0 | 0 | 2850.111235 | -0.0006 | -0.19 | -10.73 |
| alpha-Lytic | 33 | 36 | 544 | 1 | 4454 | 2705.0606 | 58.6843 | 243,087 | 2704.052775 | 23 | 1098 | 1105 | NGTHWFVT | 0.20000000.0 | 1098(NGT) | 1 | 963.4337 | 960.4454387 | 5022417.5 | 69.02 | 1 | 7 | 3 | 0 | 0 | 0 | 0 | 0 | 2704.053326 | -0.0006 | -0.2 | -10.33 |
| alpha-Lytic | 33 | 36 | 594 | 1 | 4550 | 2850.1007 | 63.3041 | 224,048 | 2849.092875 | 23 | 1098 | 1105 | NGTHWFVT | 0.20000000.0 | 1098(NGT) | 1 | 963.4337 | 960.4454387 | 5022417.5 | 69.02 | 1 | 6 | 3 | 0 | 0 | 1 | 0 | 1 | 2849.090833 | 0.0020 | 0.71 | -5.71 |
| alpha-Lytic | 33 | 36 | 602 | 1 | 4456 | 2501.9795 | 58.768 | 222,052 | 2500.971675 | 23 | 1098 | 1105 | NGTHWFVT | 0.20000000.0 | 1098(NGT) | 1 | 963.4337 | 960.4454387 | 5022417.5 | 69.02 | 1 | 7 | 2 | 0 | 0 | 0 | 0 | 1 | 2500.973953 | -0.0023 | -0.91 | -10.25 |
| alpha-Lytic | 33 | 36 | 603 | 1 | 4475 | 2380.9544 | 59.6675 | 221,839 | 2379.946575 | 23 | 1098 | 1105 | NGTHWFVT | 0.20000000.0 | 1098(NGT) | 1 | 963.4337 | 960.4454387 | 5022417.5 | 69.02 | 1 | 5 | 3 | 0 | 0 | 0 | 0 | 1 | 2379.947678 | -0.0011 | -0.46 | -9.35 |
| alpha-Lytic | 33 | 36 | 644 | 1 | 4453 | 2730.0933 | 58.6333 | 210,079 | 2729.085475 | 23 | 1098 | 1105 | NGTHWFVT | 0.20000000.0 | 1098(NGT) | 1 | 963.4337 | 960.4454387 | 5022417.5 | 69.02 | 1 | 5 | 4 | 1 | 0 | 0 | 0 | 1 | 2729.08496 | 0.0005 | 0.18 | -10.38 |
| alpha-Lytic | 33 | 36 | 650 | 1 | 4434 | 3241.2839 | 57.6452 | 208,306 | 3240.276075 | 23 | 1098 | 1105 | NGTHWFVT | 0.20000000.0 | 1098(NGT) | 1 | 963.4337 | 960.4454387 | 5022417.5 | 69.02 | 1 | 6 | 5 | 2 | 0 | 0 | 0 | 1 | 3240.275066 | 0.0010 | 0.31 | -11.37 |
| alpha-Lytic | 33 | 36 | 729 | 1 | 4419 | 3752.4686 | 56.9025 | 183,057 | 3751.460775 | 23 | 1098 | 1105 | NGTHWFVT | 0.20000000.0 | 1098(NGT) | 1 | 963.4337 | 960.4454387 | 5022417.5 | 69.02 | 1 | 7 | 6 | 3 | 0 | 0 | 0 | 0 | 3751.465172 | -0.0044 | -1.17 | -12.11 |
| alpha-Lytic | 33 | 36 | 758 | 1 | 4555 | 2688.0462 | 63.5444 | 173,893 | 2687.038375 | 23 | 1098 | 1105 | NGTHWFVT | 0.20000000.0 | 1098(NGT) | 1 | 963.4337 | 960.4454387 | 5022417.5 | 69.02 | 1 | 5 | 3 | 0 | 0 | 1 | 0 | 1 | 2687.038009 | 0.0004 | 0.13 | -5.47 |
| alpha-Lytic | 33 | 36 | 845 | 1 | 4523 | 3402.3153 | 62.0193 | 155,897 | 3401.307475 | 23 | 1098 | 1105 | NGTHWFVT | 0.20000000.0 | 1098(NGT) | 1 | 963.4337 | 960.4454387 | 5022417.5 | 69.02 | 1 | 6 | 5 | 1 | 0 | 1 | 0 | 1 | 3401.307488 | 0.0000 | 0 | -7.00 |
| alpha-Lytic | 33 | 36 | 845 | 1 | 4523 | 3402.3153 | 62.0193 | 155,897 | 3401.307475 | 23 | 1098 | 1105 | NGTHWFVT | 0.20000000.0 | 1098(NGT) | 1 | 963.4337 | 960.4454387 | 5022417.5 | 69.02 | 2 | 7 | 5 | 0 | 1 | 0 | 0 | 0 | 3401.307489 | 0.0000 | 0 | -7.00 |
| alpha-Lytic | 33 | 36 | 879 | 1 | 4464 | 2584.0354 | 59.1478 | 150,896 | 2583.027575 | 23 | 1098 | 1105 | NGTHWFVT | 0.20000000.0 | 1098(NGT) | 1 | 963.4337 | 960.4454387 | 5022417.5 | 69.02 | 1 | 5 | 4 | 0 | 0 | 0 | 0 | 1 | 2583.027051 | 0.0005 | 0.2 | -9.87 |
| alpha-Lytic | 33 | 36 | 1080 | 1 | 4529 | 3256.2586 | 62.3168 | 122,754 | 3255.250775 | 23 | 1098 | 1105 | NGTHWFVT | 0.20000000.0 | 1098(NGT) | 1 | 963.4337 | 960.4454387 | 5022417.5 | 69.02 | 1 | 6 | 5 | 0 | 0 | 1 | 0 | 1 | 3255.249579 | 0.0012 | 0.36 | -6.70 |
| alpha-Lytic | 33 | 36 | 1144 | 1 | 4446 | 2876.1482 | 58.2876 | 115,090 | 2875.140375 | 23 | 1098 | 1105 | NGTHWFVT | 0.20000000.0 | 1098(NGT) | 1 | 963.4337 | 960.4454387 | 5022417.5 | 69.02 | 1 | 5 | 4 | 2 | 0 | 0 | 0 | 1 | 2875.142869 | -0.0025 | -0.86 | -10.73 |
| alpha-Lytic | 33 | 36 | 1163 | 1 | 4467 | 2339.9263 | 59.2934 | 112,382 | 2338.918475 | 23 | 1098 | 1105 | NGTHWFVT | 0.20000000.0 | 1098(NGT) | 1 | 963.4337 | 960.4454387 | 5022417.5 | 69.02 | 1 | 6 | 2 | 0 | 0 | 0 | 0 | 1 | 2338.921129 | -0.0027 | -1.13 | -9.72 |
| alpha-Lytic | 33 | 36 | 1195 | 1 | 4427 | 3387.3405 | 57.2844 | 109,953 | 3386.332675 | 23 | 1098 | 1105 | NGTHWFVT | 0.20000000.0 | 1098(NGT) | 1 | 963.4337 | 960.4454387 | 5022417.5 | 69.02 | 1 | 6 | 5 | 3 | 0 | 0 | 0 | 1 | 3386.332975 | -0.0003 | -0.08 | -11.73 |
| alpha-Lytic | 33 | 36 | 1372 | 1 | 4450 | 2664.0313 | 58.4965 | 94,218 | 2663.023475 | 23 | 1098 | 1105 | NGTHWFVT | 0.20000000.0 | 1098(NGT) | 1 | 963.4337 | 960.4454387 | 5022417.5 | 69.02 | 1 | 8 | 2 | 0 | 0 | 0 | 0 | 1 | 2663.026777 | -0.0033 | -1.23 | -10.52 |
| alpha-Lytic | 33 | 36 | 1560 | 1 | 4506 | 3913.5011 | 61.169 | 82,651 | 3912.493275 | 23 | 1098 | 1105 | NGTHWFVT | 0.20000000.0 | 1098(NGT) | 1 | 963.4337 | 960.4454387 | 5022417.5 | 69.02 | 1 | 8 | 6 | 1 | 1 | 0 | 0 | 1 | 3912.497595 | -0.0043 | -1.1 | -7.85 |
| alpha-Lytic | 33 | 36 | 1560 | 1 | 4506 | 3913.5011 | 61.169 | 82,651 | 3912.493275 | 23 | 1098 | 1105 | NGTHWFVT | 0.20000000.0 | 1098(NGT) | 1 | 963.4337 | 960.4454387 | 5022417.5 | 69.02 | 2 | 7 | 6 | 2 | 0 | 1 | 0 | 0 | 3912.497594 | -0.0043 | -1.1 | -7.85 |
| alpha-Lytic | 33 | 36 | 1600 | 1 | 4442 | 3152.2511 | 58.0806 | 78,971 | 3151.243275 | 23 | 1098 | 1105 | NGTHWFVT | 0.20000000.0 | 1098(NGT) | 1 | 963.4337 | 960.4454387 | 5022417.5 | 69.02 | 1 | 6 | 6 | 0 | 0 | 0 | 0 | 1 | 3151.238621 | 0.0047 | 1.47 | -10.94 |
| alpha-Lytic | 33 | 36 | 1609 | 1 | 4548 | 2891.1244 | 63.2125 | 78,332 | 2890.116575 | 23 | 1098 | 1105 | NGTHWFVT | 0.20000000.0 | 1098(NGT) | 1 | 963.4337 | 960.4454387 | 5022417.5 | 69.02 | 1 | 5 | 4 | 0 | 0 | 0 | 0 | 1 | 2890.117382 | -0.0008 | -0.27 | -5.80 |
| alpha-Lytic | 33 | 36 | 1700 | 1 | 4523 | 3605.3979 | 62.0193 | 73,333 | 3604.390075 | 23 | 1098 | 1105 | NGTHWFVT | 0.20000000.0 | 1098(NGT) | 1 | 963.4337 | 960.4454387 | 5022417.5 | 69.02 | 1 | 7 | 6 | 0 | 1 | 0 | 0 | 1 | 3604.386862 | 0.0032 | 0.89 | -7.00 |
| alpha-Lytic | 33 | 36 | 1700 | 1 | 4523 | 3605.3979 | 62.0193 | 73,333 | 3604.390075 | 23 | 1098 | 1105 | NGTHWFVT | 0.20000000.0 | 1098(NGT) | 1 | 963.4337 | 960.4454387 | 5022417.5 | 69.02 | 2 | 6 | 6 | 1 | 0 | 1 | 0 | 0 | 3604.386861 | 0.0032 | 0.89 | -7.00 |
| alpha-Lytic | 33 | 36 | 2516 | 1 | 4437 | 3298.3114 | 57.8095 | 42,154 | 3297.303575 | 23 | 1098 | 1105 | NGTHWFVT | 0.20000000.0 | 1098(NGT) | 1 | 963.4337 | 960.4454387 | 5022417.5 | 69.02 | 1 | 6 | 6 | 1 | 0 | 0 | 0 | 1 | 3297.29653 | 0.0070 | 2.13 | -11.21 |
| alpha-Lytic | 33 | 36 | 2613 | 1 | 4515 | 3548.3701 | 61.6204 | 39,365 | 3547.362275 | 23 | 1098 | 1105 | NGTHWFVT | 0.20000000.0 | 1098(NGT) | 1 | 963.4337 | 960.4454387 | 5022417.5 | 69.02 | 1 | 6 | 5 | 2 | 0 | 1 | 0 | 1 | 3547.365397 | -0.0031 | -0.88 | -7.40 |
| alpha-Lytic | 33 | 36 | 2613 | 1 | 4515 | 3548.3701 | 61.6204 | 39,365 | 3547.362275 | 23 | 1098 | 1105 | NGTHWFVT | 0.20000000.0 | 1098(NGT) | 1 | 963.4337 | 960.4454387 | 5022417.5 | 69.02 | 2 | 7 | 5 | 1 | 1 | 0 | 0 | 1 | 3547.365398 | -0.0031 | -0.88 | -7.40 |
| alpha-Lytic | 33 | 36 | 2746 | 1 | 4603 | 3563.3551 | 65.9065 | 36,257 | 3562.347275 | 23 | 1098 | 1105 | NGTHWFVT | 0.20000000.0 | 1098(NGT) | 1 | 963.4337 | 960.4454387 | 5022417.5 | 69.02 | 1 | 6 | 5 | 0 | 0 | 2 | 0 | 1 | 3562.33991 | 0.0074 | 2.06 | -3.11 |
| alpha-Lytic | 33 | 36 | 2746 | 1 | 4603 | 3563.3551 | 65.9065 | 36,257 | 3562.347275 | 23 | 1098 | 1105 | NGTHWFVT | 0.20000000.0 | 1098(NGT) | 1 | 963.4337 | 960.4454387 | 5022417.5 | 69.02 | 2 | 5 | 3 | 4 | 1 | 1 | 1 | -1 | 3562.365062 | -0.0178 | -4.99 | -3.11 |
| alpha-Lytic | 33 | 36 | 2746 | 1 | 4603 | 3563.3551 | 65.9065 | 36,257 | 3562.347275 | 23 | 1098 | 1105 | NGTHWFVT | 0.20000000.0 | 1098(NGT) | 1 | 963.4337 | 960.4454387 | 5022417.5 | 69.02 | 3 | 6 | 3 | 3 | 2 | 0 | 1 | -1 | 3562.365063 | -0.0178 | -4.99 | -3.11 |
| alpha-Lytic | 33 | 36 | 2804 | 1 | 4448 | 2990.1903 | 58.3957 | 34,522 | 2989.182475 | 23 | 1098 | 1105 | NGTHWFVT | 0.20000000.0 | 1098(NGT) | 1 | 963.4337 | 960.4454387 | 5022417.5 | 69.02 | 1 | 5 | 6 | 0 | 0 | 0 | 0 | 1 | 2989.185797 | -0.0033 | -1.11 | -10.62 |
| alpha-Lytic | 33 | 36 | 3378 | 1 | 4444 | 3136.252 | 58.1864 | 21,440 | 3135.244175 | 23 | 1098 | 1105 | NGTHWFVT | 0.20000000.0 | 1098(NGT) | 1 | 963.4337 | 960.4454387 | 5022417.5 | 69.02 | 1 | 5 | 6 | 1 | 0 | 0 | 0 | 1 | 3135.243706 | 0.0005 | 0.14 | -10.83 |
| alpha-Lytic | 33 | 36 | 3620 | 1 | 4458 | 2828.1425 | 58.8678 | 16,393 | 2827.134675 | 23 | 1098 | 1105 | NGTHWFVT | 0.20000000.0 | 1098(NGT) | 1 | 963.4337 | 960.4454387 | 5022417.5 | 69.02 | 1 | 4 | 6 | 0 | 0 | 0 | 0 | 1 | 2827.132973 | 0.0017 | 0.6 | -10.15 |
| Chymotrypsin | 198 | 34 | 1222 | 1 | 5195 | 6368.7819 | 96.1571 | 313,094 | 6367.774075 | 59 | 1122 | 1155 | VSGNCDVVIGIVNNTVYDPLQPELDSFKEELDKY | 0.00000000000002000000000000000000000.0 | 1134(NNT) | 1 | 3871.86 | 3868.871739 | 6584575 | 101.39 | 1 | 7 | 6 | 1 | 0 | 0 | 0 | 1 | 6367.775654 | -0.0016 | -0.24 | -5.24 |
| Chymotrypsin | 198 | 34 | 1366 | 1 | 5192 | 6514.8444 | 96.0433 | 281,012 | 6513.836575 | 59 | 1122 | 1155 | VSGNCDVVIGIVNNTVYDPLQPELDSFKEELDKY | 0.00000000000002000000000000000000000.0 | 1134(NNT) | 1 | 3871.86 | 3868.871739 | 6584575 | 101.39 | 1 | 7 | 6 | 2 | 0 | 0 | 0 | 0 | 6513.833563 | 0.0030 | 0.46 | -5.35 |
| Chymotrypsin | 198 | 34 | 1522 | 1 | 5206 | 5638.517 | 96.6108 | 252,696 | 5637.509175 | 59 | 1122 | 1155 | VSGNCDVVIGIVNNTVYDPLQPELDSFKEELDKY | 0.00000000000002000000000000000000000.0 | 1134(NNT) | 1 | 3871.86 | 3868.871739 | 6584575 | 101.39 | 1 | 5 | 4 | 1 | 0 | 0 | 0 | 1 | 5637.51126 | -0.0021 | -0.36 | -4.78 |
| Chymotrypsin | 198 | 34 | 1755 | 1 | 5201 | 6003.6481 | 96.411 | 221,851 | 6002.640275 | 59 | 1122 | 1155 | VSGNCDVVIGIVNNTVYDPLQPELDSFKEELDKY | 0.00000000000002000000000000000000000.0 | 1134(NNT) | 1 | 3871.86 | 3868.871739 | 6584575 | 101.39 | 1 | 6 | 5 | 1 | 0 | 0 | 0 | 1 | 6002.643457 | -0.0032 | -0.53 | -4.98 |
| Chymotrypsin | 198 | 34 | 2020 | 1 | 5199 | 6149.7204 | 96.325 | 192,471 | 6148.712575 | 59 | 1122 | 1155 | VSGNCDVVIGIVNNTVYDPLQPELDSFKEELDKY | 0.00000000000002000000000000000000000.0 | 1134(NNT) | 1 | 3871.86 | 3868.871739 | 6584575 | 101.39 | 1 | 6 | 5 | 2 | 0 | 0 | 0 | 1 | 6148.701366 | 0.0112 | 1.82 | -5.07 |
| Chymotrypsin | 198 | 34 | 2156 | 1 | 5212 | 5435.432 | 96.8688 | 182,035 | 5434.424175 | 59 | 1122 | 1155 | VSGNCDVVIGIVNNTVYDPLQPELDSFKEELDKY | 0.00000000000002000000000000000000000.0 | 1134(NNT) | 1 | 3871.86 | 3868.871739 | 6584575 | 101.39 | 1 | 5 | 3 | 1 | 0 | 0 | 0 | 1 | 5434.431887 | -0.0077 | -1.41 | -4.52 |
| Chymotrypsin | 198 | 34 | 2176 | 1 | 5192 | 6660.8869 | 96.0433 | 179,904 | 6659.879075 | 59 | 1122 | 1155 | VSGNCDVVIGIVNNTVYDPLQPELDSFKEELDKY | 0.00000000000002000000000000000000000.0 | 1134(NNT) | 1 | 3871.86 | 3868.871739 | 6584575 | 101.39 | 1 | 7 | 6 | 3 | 0 | 0 | 0 | 0 | 6659.891472 | -0.0124 | -1.86 | -5.35 |
| Chymotrypsin | 198 | 34 | 2633 | 1 | 5206 | 5841.6171 | 96.6108 | 147,215 | 5840.609275 | 59 | 1122 | 1155 | VSGNCDVVIGIVNNTVYDPLQPELDSFKEELDKY | 0.00000000000002000000000000000000000.0 | 1134(NNT) | 1 | 3871.86 | 3868.871739 | 6584575 | 101.39 | 1 | 5 | 5 | 1 | 0 | 0 | 0 | 1 | 5840.590633 | 0.0186 | 3.19 | -4.78 |
| Chymotrypsin | 198 | 34 | 3276 | 1 | 5213 | 5289.3847 | 96.9114 | 114,791 | 5288.376875 | 59 | 1122 | 1155 | VSGNCDVVIGIVNNTVYDPLQPELDSFKEELDKY | 0.00000000000002000000000000000000000.0 | 1134(NNT) | 1 | 3871.86 | 3868.871739 | 6584575 | 101.39 | 1 | 5 | 3 | 0 | 0 | 0 | 0 | 1 | 5288.373978 | 0.0029 | 0.54 | -4.48 |
| [illegible] | [illegible] | [illegible] | [illegible] | [illegible] | [illegible] | [illegible] | [illegible] | [illegible] | [illegible] | [illegible] | [illegible] | [illegible] | [illegible] | [illegible] | [illegible] | [illegible] | [illegible] | [illegible] | [illegible] | [illegible] | [illegible] | [illegible] | [illegible] | [illegible] | [illegible] | [illegible] | [illegible] | [illegible] | [illegible] | [illegible] | [illegible] | [illegible] |

| | | | | | | | | | | | | | | | | | | | | | | | | | | | | | | | | |
|---|---|---|---|---|---|---|---|---|---|---|---|---|---|---|---|---|---|---|---|---|---|---|---|---|---|---|---|---|---|---|---|---|
| alpha-Lytic | 40 | 10 | 560 | 1 | 4237 | 3304.3322 | 47.8626 | 237,593 | 3303.324375 | 1020 | 1171 | 1177 | GINASVV | - | 1173(NAS) | 1 | | 658.36496 | - | 54.00 | 1 | 7 | 6 | 2 | 0 | 0 | 0 | 0 | 3303.326784 | -0.0024 | -0.72 | -6.14 |
| alpha-Lytic | 40 | 10 | 751 | 1 | 4307 | 3465.3662 | 51.281 | 175,822 | 3464.358375 | 1020 | 1171 | 1177 | GINASVV | - | 1173(NAS) | 1 | | 658.36496 | - | 54.00 | 1 | 7 | 6 | 1 | 0 | 1 | 0 | 1 | 3464.359206 | -0.0008 | -0.23 | -2.72 |
| alpha-Lytic | 40 | 10 | 751 | 1 | 4307 | 3465.3662 | 51.281 | 175,822 | 3464.358375 | 1020 | 1171 | 1177 | GINASVV | - | 1173(NAS) | 1 | | 658.36496 | - | 54.00 | 2 | 8 | 6 | 0 | 1 | 0 | 0 | 1 | 3464.359207 | -0.0008 | -0.24 | -2.72 |
| alpha-Lytic | 40 | 10 | 767 | 1 | 4232 | 3450.394 | 47.6098 | 172,305 | 3449.386175 | 1020 | 1171 | 1177 | GINASVV | - | 1173(NAS) | 1 | | 658.36496 | - | 54.00 | 1 | 7 | 6 | 3 | 0 | 0 | 0 | 0 | 3449.384693 | 0.0015 | 0.42 | -6.39 |
| alpha-Lytic | 40 | 10 | 1281 | 1 | 4262 | 3595.4359 | 49.0412 | 101,828 | 3594.428075 | 1020 | 1171 | 1177 | GINASVV | - | 1173(NAS) | 1 | | 658.36496 | - | 54.00 | 1 | 7 | 6 | 2 | 1 | 0 | 0 | 0 | 3594.422201 | 0.0059 | 1.63 | -4.96 |
| alpha-Lytic | 40 | 10 | 1281 | 1 | 4262 | 3595.4359 | 49.0412 | 101,828 | 3594.428075 | 1020 | 1171 | 1177 | GINASVV | - | 1173(NAS) | 1 | | 658.36496 | - | 54.00 | 2 | 6 | 6 | 3 | 0 | 1 | 0 | 0 | 3594.4222 | 0.0059 | 1.63 | -4.96 |
| alpha-Lytic | 40 | 10 | 1526 | 1 | 4223 | 3742.5106 | 47.1549 | 84,828 | 3741.502775 | 1020 | 1171 | 1177 | GINASVV | - | 1173(NAS) | 1 | | 658.36496 | - | 54.00 | 1 | 2 | 10 | 1 | 2 | 0 | 1 | -1 | 3741.513081 | -0.0103 | -2.75 | -6.85 |
| alpha-Lytic | 40 | 10 | 1562 | 1 | 4227 | 3596.4518 | 47.3671 | 82,286 | 3595.443975 | 1020 | 1171 | 1177 | GINASVV | - | 1173(NAS) | 1 | | 658.36496 | - | 54.00 | 1 | 7 | 6 | 4 | 0 | 0 | 0 | 1 | 3595.442602 | 0.0014 | 0.38 | -6.63 |
| alpha-Lytic | 40 | 10 | 1562 | 1 | 4227 | 3596.4518 | 47.3671 | 82,286 | 3595.443975 | 1020 | 1171 | 1177 | GINASVV | - | 1173(NAS) | 1 | | 658.36496 | - | 54.00 | 2 | 2 | 10 | 0 | 2 | 0 | 1 | -1 | 3595.455172 | -0.0112 | -3.11 | -6.63 |
| alpha-Lytic | 40 | 10 | 2008 | 1 | 4365 | 3772.4586 | 54.163 | 58,969 | 3771.450775 | 1020 | 1171 | 1177 | GINASVV | - | 1173(NAS) | 1 | | 658.36496 | - | 54.00 | 1 | 8 | 6 | 0 | 1 | 1 | 0 | 0 | 3771.449538 | 0.0012 | 0.32 | 0.16 |
| alpha-Lytic | 40 | 10 | 2008 | 1 | 4365 | 3772.4586 | 54.163 | 58,969 | 3771.450775 | 1020 | 1171 | 1177 | GINASVV | - | 1173(NAS) | 1 | | 658.36496 | - | 54.00 | 2 | 7 | 6 | 1 | 0 | 2 | 0 | 0 | 3771.449537 | 0.0012 | 0.32 | 0.16 |
| alpha-Lytic | 40 | 10 | 3036 | 1 | 4371 | 3756.4574 | 54.4613 | 29,176 | 3755.449575 | 1020 | 1171 | 1177 | GINASVV | - | 1173(NAS) | 1 | | 658.36496 | - | 54.00 | 1 | 6 | 6 | 2 | 0 | 2 | 0 | 0 | 3755.454622 | -0.0050 | -1.34 | 0.46 |
| alpha-Lytic | 40 | 10 | 3036 | 1 | 4371 | 3756.4574 | 54.4613 | 29,176 | 3755.449575 | 1020 | 1171 | 1177 | GINASVV | - | 1173(NAS) | 1 | | 658.36496 | - | 54.00 | 2 | 7 | 6 | 1 | 1 | 1 | 0 | 0 | 3755.454623 | -0.0050 | -1.34 | 0.46 |
| alpha-Lytic | 40 | 10 | 3036 | 1 | 4371 | 3756.4574 | 54.4613 | 29,176 | 3755.449575 | 1020 | 1171 | 1177 | GINASVV | - | 1173(NAS) | 1 | | 658.36496 | - | 54.00 | 3 | 8 | 6 | 0 | 2 | 0 | 0 | 0 | 3755.454624 | -0.0050 | -1.34 | 0.46 |
| Trypsin | 44 | 34 | 254 | 1 | 4529 | 4084.7475 | 67.6277 | 526,933 | 4083.739675 | 34 | 1192 | 1205 | NLNESLIDLQELGK | 0.00200000000000.0 | 1194(NES) | 1 | 1587.8241 | 1584.835839 | 3736906.5 | 75.63 | 1 | 7 | 6 | 1 | 0 | 0 | 0 | 1 | 4083.739754 | -0.0001 | -0.01 | -8.00 |
| Trypsin | 44 | 34 | 410 | 1 | 4539 | 3719.6123 | 68.1234 | 355,095 | 3718.604475 | 34 | 1192 | 1205 | NLNESLIDLQELGK | 0.00200000000000.0 | 1194(NES) | 1 | 1587.8241 | 1584.835839 | 3736906.5 | 75.63 | 1 | 6 | 5 | 1 | 0 | 0 | 0 | 1 | 3718.607557 | -0.0031 | -0.82 | -7.51 |
| Trypsin | 44 | 34 | 971 | 1 | 4531 | 3922.6926 | 67.7355 | 144,623 | 3921.684775 | 34 | 1192 | 1205 | NLNESLIDLQELGK | 0.00200000000000.0 | 1194(NES) | 1 | 1587.8241 | 1584.835839 | 3736906.5 | 75.63 | 1 | 6 | 6 | 1 | 0 | 0 | 0 | 1 | 3921.68693 | -0.0022 | -0.54 | -7.89 |
| Trypsin | 44 | 34 | 1025 | 1 | 4526 | 4230.8053 | 67.4761 | 136,411 | 4229.797475 | 34 | 1192 | 1205 | NLNESLIDLQELGK | 0.00200000000000.0 | 1194(NES) | 1 | 1587.8241 | 1584.835839 | 3736906.5 | 75.63 | 1 | 7 | 6 | 2 | 0 | 0 | 0 | 0 | 4229.797663 | -0.0002 | -0.04 | -8.15 |
| Trypsin | 44 | 34 | 1150 | 1 | 4546 | 3354.4798 | 68.4486 | 120,867 | 3353.471975 | 34 | 1192 | 1205 | NLNESLIDLQELGK | 0.00200000000000.0 | 1194(NES) | 1 | 1587.8241 | 1584.835839 | 3736906.5 | 75.63 | 1 | 5 | 4 | 1 | 0 | 0 | 0 | 1 | 3353.47536 | -0.0034 | -1 | -7.18 |
| Trypsin | 44 | 34 | 1405 | 1 | 4613 | 4391.8358 | 71.5577 | 97,256 | 4390.827975 | 34 | 1192 | 1205 | NLNESLIDLQELGK | 0.00200000000000.0 | 1194(NES) | 1 | 1587.8241 | 1584.835839 | 3736906.5 | 75.63 | 1 | 7 | 6 | 1 | 0 | 1 | 0 | 1 | 4390.830085 | -0.0021 | -0.48 | -4.07 |
| Trypsin | 44 | 34 | 1405 | 1 | 4613 | 4391.8358 | 71.5577 | 97,256 | 4390.827975 | 34 | 1192 | 1205 | NLNESLIDLQELGK | 0.00200000000000.0 | 1194(NES) | 1 | 1587.8241 | 1584.835839 | 3736906.5 | 75.63 | 2 | 8 | 6 | 0 | 1 | 0 | 0 | 1 | 4390.830086 | -0.0021 | -0.48 | -4.07 |
| Trypsin | 44 | 34 | 1773 | 1 | 4550 | 3436.534 | 68.6351 | 75,424 | 3435.526175 | 34 | 1192 | 1205 | NLNESLIDLQELGK | 0.00200000000000.0 | 1194(NES) | 1 | 1587.8241 | 1584.835839 | 3736906.5 | 75.63 | 1 | 3 | 6 | 1 | 0 | 0 | 0 | 1 | 3435.528458 | -0.0023 | -0.66 | -6.99 |
| Trypsin | 44 | 34 | 1968 | 1 | 4541 | 3598.5879 | 68.2208 | 66,663 | 3597.580075 | 34 | 1192 | 1205 | NLNESLIDLQELGK | 0.00200000000000.0 | 1194(NES) | 1 | 1587.8241 | 1584.835839 | 3736906.5 | 75.63 | 1 | 4 | 6 | 1 | 0 | 0 | 0 | 1 | 3597.581282 | -0.0012 | -0.33 | -7.41 |
| Trypsin | 44 | 34 | 1989 | 1 | 4535 | 3865.6717 | 67.9334 | 65,713 | 3864.663875 | 34 | 1192 | 1205 | NLNESLIDLQELGK | 0.00200000000000.0 | 1194(NES) | 1 | 1587.8241 | 1584.835839 | 3736906.5 | 75.63 | 1 | 6 | 5 | 2 | 0 | 0 | 0 | 1 | 3864.665466 | -0.0016 | -0.41 | -7.70 |
| Trypsin | 44 | 34 | 2090 | 1 | 4524 | 4376.8546 | 67.3744 | 62,088 | 4375.846775 | 34 | 1192 | 1205 | NLNESLIDLQELGK | 0.00200000000000.0 | 1194(NES) | 1 | 1587.8241 | 1584.835839 | 3736906.5 | 75.63 | 1 | 7 | 6 | 3 | 0 | 0 | 0 | 0 | 4375.855572 | -0.0088 | -2.01 | -8.25 |
| Trypsin | 44 | 34 | 2243 | 1 | 4536 | 3760.6413 | 67.9767 | 57,349 | 3759.633475 | 34 | 1192 | 1205 | NLNESLIDLQELGK | 0.00200000000000.0 | 1194(NES) | 1 | 1587.8241 | 1584.835839 | 3736906.5 | 75.63 | 1 | 5 | 6 | 1 | 0 | 0 | 0 | 1 | 3759.634106 | -0.0006 | -0.16 | -7.65 |
| Trypsin | 44 | 34 | 2352 | 1 | 4546 | 3395.5085 | 68.4486 | 53,842 | 3394.500675 | 34 | 1192 | 1205 | NLNESLIDLQELGK | 0.00200000000000.0 | 1194(NES) | 1 | 1587.8241 | 1584.835839 | 3736906.5 | 75.63 | 1 | 4 | 5 | 1 | 0 | 0 | 0 | 1 | 3394.501909 | -0.0012 | -0.36 | -7.18 |
| Trypsin | 44 | 34 | 2491 | 1 | 4541 | 3557.5656 | 68.2208 | 50,384 | 3556.557775 | 34 | 1192 | 1205 | NLNESLIDLQELGK | 0.00200000000000.0 | 1194(NES) | 1 | 1587.8241 | 1584.835839 | 3736906.5 | 75.63 | 1 | 5 | 5 | 1 | 0 | 0 | 0 | 1 | 3556.554733 | 0.0030 | 0.85 | -7.41 |
| Trypsin | 44 | 34 | 2541 | 1 | 4546 | 3208.4194 | 68.4486 | 49,348 | 3207.411575 | 34 | 1192 | 1205 | NLNESLIDLQELGK | 0.00200000000000.0 | 1194(NES) | 1 | 1587.8241 | 1584.835839 | 3736906.5 | 75.63 | 1 | 5 | 4 | 0 | 0 | 0 | 0 | 1 | 3207.417451 | -0.0059 | -1.83 | -7.18 |
| Trypsin | 44 | 34 | 2609 | 1 | 4538 | 3573.5508 | 68.0797 | 47,530 | 3572.542975 | 34 | 1192 | 1205 | NLNESLIDLQELGK | 0.00200000000000.0 | 1194(NES) | 1 | 1587.8241 | 1584.835839 | 3736906.5 | 75.63 | 1 | 6 | 5 | 0 | 0 | 0 | 0 | 1 | 3572.549648 | -0.0067 | -1.86 | -7.55 |
| Trypsin | 44 | 34 | 2640 | 1 | 4555 | 3233.4568 | 68.857 | 46,869 | 3232.448975 | 34 | 1192 | 1205 | NLNESLIDLQELGK | 0.00200000000000.0 | 1194(NES) | 1 | 1587.8241 | 1584.835839 | 3736906.5 | 75.63 | 1 | 3 | 5 | 1 | 0 | 0 | 0 | 1 | 3232.449085 | -0.0001 | -0.03 | -6.77 |
| Trypsin | 44 | 34 | 2807 | 1 | 4528 | 3938.6916 | 67.5705 | 43,141 | 3937.683775 | 34 | 1192 | 1205 | NLNESLIDLQELGK | 0.00200000000000.0 | 1194(NES) | 1 | 1587.8241 | 1584.835839 | 3736906.5 | 75.63 | 1 | 7 | 6 | 0 | 0 | 0 | 0 | 1 | 3937.681845 | 0.0019 | 0.49 | -8.06 |
| Trypsin | 44 | 34 | 2877 | 1 | 4527 | 4157.7773 | 67.5179 | 41,500 | 4156.769475 | 34 | 1192 | 1205 | NLNESLIDLQELGK | 0.00200000000000.0 | 1194(NES) | 1 | 1587.8241 | 1584.835839 | 3736906.5 | 75.63 | 1 | 6 | 5 | 4 | 0 | 0 | 0 | 0 | 4156.781284 | -0.0118 | -2.84 | -8.11 |
| Trypsin | 44 | 34 | 2923 | 1 | 4624 | 4026.6976 | 72.0913 | 40,722 | 4025.689775 | 34 | 1192 | 1205 | NLNESLIDLQELGK | 0.00200000000000.0 | 1194(NES) | 1 | 1587.8241 | 1584.835839 | 3736906.5 | 75.63 | 1 | 6 | 5 | 1 | 0 | 1 | 0 | 1 | 4025.697888 | -0.0081 | -2.01 | -3.54 |
| Trypsin | 44 | 34 | 2923 | 1 | 4624 | 4026.6976 | 72.0913 | 40,722 | 4025.689775 | 34 | 1192 | 1205 | NLNESLIDLQELGK | 0.00200000000000.0 | 1194(NES) | 1 | 1587.8241 | 1584.835839 | 3736906.5 | 75.63 | 2 | 7 | 5 | 0 | 1 | 0 | 0 | 0 | 4025.697889 | -0.0081 | -2.01 | -3.54 |
| Trypsin | 44 | 34 | 2992 | 1 | 4537 | 3646.5937 | 68.0279 | 39,591 | 3645.585875 | 34 | 1192 | 1205 | NLNESLIDLQELGK | 0.00200000000000.0 | 1194(NES) | 1 | 1587.8241 | 1584.835839 | 3736906.5 | 75.63 | 1 | 5 | 4 | 3 | 0 | 0 | 0 | 1 | 3645.591178 | -0.0053 | -1.45 | -7.60 |
| Trypsin | 44 | 34 | 3014 | 1 | 4531 | 4011.7218 | 67.7355 | 39,242 | 4010.713975 | 34 | 1192 | 1205 | NLNESLIDLQELGK | 0.00200000000000.0 | 1194(NES) | 1 | 1587.8241 | 1584.835839 | 3736906.5 | 75.63 | 1 | 6 | 5 | 3 | 0 | 0 | 0 | 1 | 4010.723375 | -0.0094 | -2.34 | -7.89 |
| Trypsin | 44 | 34 | 3337 | 1 | 4611 | 4537.9028 | 71.4631 | 34,033 | 4536.894975 | 34 | 1192 | 1205 | NLNESLIDLQELGK | 0.00200000000000.0 | 1194(NES) | 1 | 1587.8241 | 1584.835839 | 3736906.5 | 75.63 | 1 | 8 | 6 | 1 | 1 | 0 | 0 | 1 | 4536.887995 | 0.0070 | 1.53 | -4.17 |
| Trypsin | 44 | 34 | 3337 | 1 | 4611 | 4537.9028 | 71.4631 | 34,033 | 4536.894975 | 34 | 1192 | 1205 | NLNESLIDLQELGK | 0.00200000000000.0 | 1194(NES) | 1 | 1587.8241 | 1584.835839 | 3736906.5 | 75.63 | 2 | 7 | 6 | 2 | 0 | 1 | 0 | 0 | 4536.887994 | 0.0070 | 1.53 | -4.17 |
| Trypsin | 44 | 34 | 3391 | 1 | 4583 | 3337.4575 | 70.1628 | 33,348 | 3336.449675 | 34 | 1192 | 1205 | NLNESLIDLQELGK | 0.00200000000000.0 | 1194(NES) | 1 | 1587.8241 | 1584.835839 | 3736906.5 | 75.63 | 1 | 4 | 4 | 0 | 1 | 0 | 0 | 1 | 3336.460044 | -0.0104 | -3.1 | -5.47 |
| Trypsin | 44 | 34 | 3391 | 1 | 4583 | 3337.4575 | 70.1628 | 33,348 | 3336.449675 | 34 | 1192 | 1205 | NLNESLIDLQELGK | 0.00200000000000.0 | 1194(NES) | 1 | 1587.8241 | 1584.835839 | 3736906.5 | 75.63 | 2 | 3 | 4 | 1 | 0 | 1 | 0 | 0 | 3336.460043 | -0.0104 | -3.1 | -5.47 |
| Trypsin | 44 | 34 | 3429 | 1 | 4520 | 4522.927 | 67.1921 | 32,753 | 4521.919175 | 34 | 1192 | 1205 | NLNESLIDLQELGK | 0.00200000000000.0 | 1194(NES) | 1 | 1587.8241 | 1584.835839 | 3736906.5 | 75.63 | 1 | 7 | 6 | 4 | 0 | 0 | 0 | 1 | 4521.913481 | 0.0057 | 1.25 | -8.44 |
| Trypsin | 44 | 34 | 3429 | 1 | 4520 | 4522.927 | 67.1921 | 32,753 | 4521.919175 | 34 | 1192 | 1205 | NLNESLIDLQELGK | 0.00200000000000.0 | 1194(NES) | 1 | 1587.8241 | 1584.835839 | 3736906.5 | 75.63 | 2 | 2 | 10 | 0 | 2 | 0 | 1 | -1 | 4521.926051 | -0.0069 | -1.52 | -8.44 |
| Trypsin | 44 | 34 | 3545 | 1 | 4518 | 4668.9873 | 67.0921 | 31,429 | 4667.979475 | 34 | 1192 | 1205 | NLNESLIDLQELGK | 0.00200000000000.0 | 1194(NES) | 1 | 1587.8241 | 1584.835839 | 3736906.5 | 75.63 | 1 | 2 | 10 | 1 | 2 | 0 | 1 | -1 | 4667.98396 | -0.0045 | -0.96 | -8.54 |
| Trypsin | 44 | 34 | 4096 | 1 | 4610 | 4683.958 | 71.411 | 25,037 | 4682.950175 | 34 | 1192 | 1205 | NLNESLIDLQELGK | 0.00200000000000.0 | 1194(NES) | 1 | 1587.8241 | 1584.835839 | 3736906.5 | 75.63 | 1 | 8 | 6 | 2 | 1 | 0 | 0 | 0 | 4682.945904 | 0.0043 | 0.91 | -4.22 |
| Trypsin | 44 | 34 | 4096 | 1 | 4610 | 4683.958 | 71.411 | 25,037 | 4682.950175 | 34 | 1192 | 1205 | NLNESLIDLQELGK | 0.00200000000000.0 | 1194(NES) | 1 | 1587.8241 | 1584.835839 | 3736906.5 | 75.63 | 2 | 7 | 6 | 3 | 0 | 1 | 0 | 0 | 4682.945903 | 0.0043 | 0.91 | -4.22 |
| Trypsin | 44 | 34 | 4324 | 1 | 4560 | 3030.3735 | 69.081 | 22,944 | 3029.365675 | 34 | 1192 | 1205 | NLNESLIDLQELGK | 0.00200000000000.0 | 1194(NES) | 1 | 1587.8241 | 1584.835839 | 3736906.5 | 75.63 | 1 | 3 | 4 | 1 | 0 | 0 | 0 | 1 | 3029.369712 | -0.0040 | -1.33 | -6.55 |
| Trypsin | 44 | 34 | 4390 | 1 | 4620 | 4172.7513 | 71.8975 | 22,403 | 4171.743475 | 34 | 1192 | 1205 | NLNESLIDLQELGK | 0.00200000000000.0 | 1194(NES) | 1 | 1587.8241 | 1584.835839 | 3736906.5 | 75.63 | 1 | 6 | 5 | 2 | 0 | 1 | 0 | 1 | 4171.755797 | -0.0123 | -2.95 | -3.73 |
| Trypsin | 44 | 34 | 4390 | 1 | 4620 | 4172.7513 | 71.8975 | 22,403 | 4171.743475 | 34 | 1192 | 1205 | NLNESLIDLQELGK | 0.00200000000000.0 | 1194(NES) | 1 | 1587.8241 | 1584.835839 | 3736906.5 | 75.63 | 2 | 7 | 5 | 1 | 1 | 0 | 0 | 1 | 4171.755798 | -0.0123 | -2.95 | -3.73 |
| Trypsin | 44 | 34 | 4549 | 1 | 4540 | 3500.5407 | 68.1798 | 21,037 | 3499.532875 | 34 | 1192 | 1205 | NLNESLIDLQELGK | 0.00200000000000.0 | 1194(NES) | 1 | 1587.8241 | 1584.835839 | 3736906.5 | 75.63 | 1 | 5 | 4 | 2 | 0 | 0 | 0 | 1 | 3499.533269 | -0.0004 | -0.11 | -7.45 |
| Trypsin | 44 | 34 | 4628 | 1 | 4617 | 4375.8398 | 71.751 | 20,397 | 4374.831975 | 34 | 1192 | 1205 | NLNESLIDLQELGK | 0.00200000000000.0 | 1194(NES) | 1 | 1587.8241 | 1584.835839 | 3736906.5 | 75.63 | 1 | 7 | 6 | 1 | 1 | 0 | 0 | 1 | 4374.835171 | -0.0032 | -0.73 | -3.88 |
| Trypsin | 44 | 34 | 4628 | 1 | 4617 | 4375.8398 | 71.751 | 20,397 | 4374.831975 | 34 | 1192 | 1205 | NLNESLIDLQELGK | 0.00200000000000.0 | 1194(NES) | 1 | 1587.8241 | 1584.835839 | 3736906.5 | 75.63 | 2 | 6 | 6 | 2 | 0 | 1 | 0 | 0 | 4374.83517 | -0.0032 | -0.73 | -3.88 |
| Trypsin | 44 | 34 | 4862 | 1 | 4691 | 4844.9839 | 75.34 | 18,763 | 4843.976075 | 34 | 1192 | 1205 | NLNESLIDLQELGK | 0.00200000000000.0 | 1194(NES) | 1 | 1587.8241 | 1584.835839 | 3736906.5 | 75.63 | 1 | 7 | 6 | 2 | 0 | 2 | 0 | 0 | 4843.978325 | -0.0022 | -0.46 | -0.29 |
| Trypsin | 44 | 34 | 4862 | 1 | 4691 | 4844.9839 | 75.34 | 18,763 | 4843.976075 | 34 | 1192 | 1205 | NLNESLIDLQELGK | 0.00200000000000.0 | 1194(NES) | 1 | 1587.8241 | 1584.835839 | 3736906.5 | 75.63 | 2 | 8 | 6 | 1 | 1 | 1 | 0 | 0 | 4843.978326 | -0.0023 | -0.46 | -0.29 |
| Trypsin | 44 | 34 | 4862 | 1 | 4691 | 4844.9839 | 75.34 | 18,763 | 4843.976075 | 34 | 1192 | 1205 | NLNESLIDLQELGK | 0.00200000000000.0 | 1194(NES) | 1 | 1587.8241 | 1584.835839 | 3736906.5 | 75.63 | 3 | 9 | 6 | 0 | 2 | 0 | 0 | 0 | 4843.978327 | -0.0023 | -0.46 | -0.29 |
| Trypsin | 44 | 34 | 6289 | 1 | 4512 | 4449.8833 | 66.7908 | 9,448 | 4448.875475 | 34 | 1192 | 1205 | NLNESLIDLQELGK | 0.00200000000000.0 | 1194(NES) | 1 | 1587.8241 | 1584.835839 | 3736906.5 | 75.63 | 1 | 8 | 7 | 1 | 0 | 0 | 0 | 1 | 4448.871951 | 0.0035 | 0.79 | -8.84 |
| Trypsin | 44 | 34 | 6918 | 1 | 4687 | 4991.0401 | 75.1487 | 5,330 | 4990.032275 | 34 | 1192 | 1205 | NLNESLIDLQELGK | 0.00200000000000.0 | 1194(NES) | 1 | 1587.8241 | 1584.835839 | 3736906.5 | 75.63 | 1 | 9 | 6 | 1 | 2 | 0 | 0 | 1 | 4990.036236 | -0.0040 | -0.79 | -0.48 |
| Trypsin | 44 | 34 | 6918 | 1 | 4687 | 4991.0401 | 75.1487 | 5,330 | 4990.032275 | 34 | 1192 | 1205 | NLNESLIDLQELGK | 0.00200000000000.0 | 1194(NES) | 1 | 1587.8241 | 1584.835839 | 3736906.5 | 75.63 | 2 | 7 | 6 | 3 | 0 | 2 | 0 | 0 | 4990.036234 | -0.0040 | -0.79 | -0.48 |
| Trypsin | 44 | 34 | 6918 | 1 | 4687 | 4991.0401 | 75.1487 | 5,330 | 4990.032275 | 34 | 1192 | 1205 | NLNESLIDLQELGK | 0.00200000000000.0 | 1194(NES) | 1 | 1587.8241 | 1584.835839 | 3736906.5 | 75.63 | 3 | 8 | 6 | 2 | 1 | 1 | 0 | 0 | 4990.036235 | -0.0040 | -0.79 | -0.48 |
| Trypsin | 44 | 34 | 6995 | 1 | 4550 | 3046.3762 | 68.6351 | 4,633 | 3045.368375 | 34 | 1192 | 1205 | NLNESLIDLQELGK | 0.00200000000000.0 | 1194(NES) | 1 | 1587.8241 | 1584.835839 | 3736906.5 | 75.63 | 1 | 4 | 4 | 0 | 0 | 0 | 0 | 1 | 3045.364627 | 0.0037 | 1.23 | -6.99 |
| alpha-Lytic | 88 | 5 | 618 | 1 | 4745 | 4084.7461 | 72.9789 | 216,670 | 4083.738275 | 68 | 1191 | 1204 | KNLNESLIDLQELG | 0.00020000000000.0 | 1194(NES) | 1 | 1587.8241 | 1584.835839 | 1668065.5 | 81.99 | 1 | 7 | 6 | 1 | 0 | 0 | 0 | 1 | 4083.739754 | -0.0015 | -0.36 | -9.01 |
| alpha-Lytic | 88 | 5 | 1141 | 1 | 4755 | 3719.6162 | 73.4945 | 115,287 | 3718.608375 | 68 | 1191 | 1204 | KNLNESLIDLQELG | 0.00020000000000.0 | 1194(NES) | 1 | 1587.8241 | 1584.835839 | 1668065.5 | 81.99 | 1 | 6 | 5 | 1 | 0 | 0 | 0 | 1 | 3718.607557 | 0.0008 | 0.22 | -8.50 |
| alpha-Lytic | 88 | 5 | 2537 | 1 | 4747 | 3922.6955 | 73.0897 | 41,692 | 3921.687675 | 68 | 1191 | 1204 | KNLNESLIDLQELG | 0.00020000000000.0 | 1194(NES) | 1 | 1587.8241 | 1584.835839 | 1668065.5 | 81.99 | 1 | 6 | 6 | 1 | 0 | 0 | 0 | 1 | 3921.68693 | 0.0007 | 0.19 | -8.90 |
| alpha-Lytic | 88 | 5 | 2788 | 1 | 4764 | 3354.4867 | 73.9282 | 34,955 | 3353.478875 | 68 | 1191 | 1204 | KNLNESLIDLQELG | 0.00020000000000.0 | 1194(NES) | 1 | 1587.8241 | 1584.835839 | 1668065.5 | 81.99 | 1 | 5 | 4 | 1 | 0 | 0 | 0 | 1 | 3353.47536 | 0.0035 | 1.04 | -8.06 |
| alpha-Lytic | 88 | 5 | 3148 | 1 | 4750 | 3865.678 | 73.2447 | 26,780 | 3864.670175 | 68 | 1191 | 1204 | KNLNESLIDLQELG | 0.00020000000000.0 | 1194(NES) | 1 | 1587.8241 | 1584.835839 | 1668065.5 | 81.99 | 1 | 6 | 5 | 2 | 0 | 0 | 0 | 1 | 3864.665466 | 0.0047 | 1.21 | -8.75 |
| alpha-Lytic | 98 | 5 | 873 | 1 | 4752 | 4155.7862 | 73.3372 | 151,673 | 4154.778375 | 100 | 1190 | 1204 | AKNLNESLIDLQELG | 0.000020000000000.0 | 1194(NES) | 1 | 1658.8612 | 1655.872939 | 784978 | 81.90 | 1 | 7 | 6 | 1 | 0 | 0 | 0 | 1 | 4154.776854 | 0.0015 | 0.36 | -8.57 |
| alpha-Lytic | 98 | 5 | 1836 | 1 | 4761 | 3790.6509 | 73.7884 | 66,163 | 3789.643075 | 100 | 1190 | 1204 | AKNLNESLIDLQELG | 0.000020000000000.0 | 1194(NES) | 1 | 1658.8612 | 1655.872939 | 784978 | 81.90 | 1 | 6 | 5 | 1 | 0 | 0 | 0 | 1 | 3789.644657 | -0.0016 | -0.41 | -8.12 |
| alpha-Lytic | 98 | 5 | 2845 | 1 | 4753 | 3993.7346 | 73.3868 | 33,481 | 3992.726775 | 100 | 1190 | 1204 | AKNLNESLIDLQELG | 0.000020000000000.0 | 1194(NES) | 1 | 1658.8612 | 1655.872939 | 784978 | 81.90 | 1 | 6 | 6 | 1 | 0 | 0 | 0 | 1 | 3992.72403 | 0.0027 | 0.68 | -8.52 |
| alpha-Lytic | 98 | 5 | 2977 | 1 | 4747 | 4301.8429 | 73.0897 | 30,382 | 4300.835075 | 100 | 1190 | 1204 | AKNLNESLIDLQELG | 0.000020000000000.0 | 1194(NES) | 1 | 1658.8612 | 1655.872939 | 784978 | 81.90 | 1 | 7 | 6 | 2 | 0 | 0 | 0 | 0 | 4300.834763 | 0.0003 | 0.07 | -8.81 |
| alpha-Lytic | 98 | 5 | 3235 | 1 | 4771 | 3425.5195 | 74.2494 | 24,855 | 3424.511675 | 100 | 1190 | 1204 | AKNLNESLIDLQELG | 0.000020000000000.0 | 1194(NES) | 1 | 1658.8612 | 1655.872939 | 784978 | 81.90 | 1 | 5 | 4 | 1 | 0 | 0 | 0 | 1 | 3424.51246 | -0.0008 | -0.22 | -7.65 |

Supplementary Table S11. Non-redundant site-specific glycans assigned by Glyco-RIDGE

| # | Glycosylation site | No. of composition / site | Glycan composition | No. of HexNAc | No. of Hex | No. of Fuc | No. of NeuAc | No. of NeuGc | Frequency / 22 sites |
|---|---|---|---|---|---|---|---|---|---|
| 1 | 17 | 46 | 3:5:1:0:0 | 3 | 5 | 1 | 0 | 0 | 14 |
| 2 | 17 | 46 | 3:5:2:0:0 | 3 | 5 | 2 | 0 | 0 | 13 |
| 3 | 17 | 46 | 3:6:0:0:0 | 3 | 6 | 0 | 0 | 0 | 11 |
| 4 | 17 | 46 | 3:6:1:0:0 | 3 | 6 | 1 | 0 | 0 | 13 |
| 5 | 17 | 46 | 3:6:2:0:0 | 3 | 6 | 2 | 0 | 0 | 6 |
| 6 | 17 | 46 | 4:3:1:0:0 | 4 | 3 | 1 | 0 | 0 | 14 |
| 7 | 17 | 46 | 4:4:1:0:0 | 4 | 4 | 1 | 0 | 0 | 12 |
| 8 | 17 | 46 | 4:4:1:0:1 | 4 | 4 | 1 | 0 | 1 | 7 |
| 9 | 17 | 46 | 4:4:2:0:0 | 4 | 4 | 2 | 0 | 0 | 8 |
| 10 | 17 | 46 | 4:5:0:1:0 | 4 | 5 | 0 | 1 | 0 | 7 |
| 11 | 17 | 46 | 4:5:1:0:0 | 4 | 5 | 1 | 0 | 0 | 19 |
| 12 | 17 | 46 | 4:5:1:0:1 | 4 | 5 | 1 | 0 | 1 | 14 |
| 13 | 17 | 46 | 4:5:2:0:0 | 4 | 5 | 2 | 0 | 0 | 18 |
| 14 | 17 | 46 | 4:5:2:0:1 | 4 | 5 | 2 | 0 | 1 | 9 |
| 15 | 17 | 46 | 4:5:3:0:0 | 4 | 5 | 3 | 0 | 0 | 17 |
| 16 | 17 | 46 | 4:6:0:1:0 | 4 | 6 | 0 | 1 | 0 | 14 |
| 17 | 17 | 46 | 4:6:1:1:0 | 4 | 6 | 1 | 1 | 0 | 9 |
| 18 | 17 | 46 | 5:3:1:0:0 | 5 | 3 | 1 | 0 | 0 | 11 |
| 19 | 17 | 46 | 5:4:1:0:0 | 5 | 4 | 1 | 0 | 0 | 16 |
| 20 | 17 | 46 | 5:4:1:0:1 | 5 | 4 | 1 | 0 | 1 | 4 |
| 21 | 17 | 46 | 5:4:2:0:0 | 5 | 4 | 2 | 0 | 0 | 8 |
| 22 | 17 | 46 | 5:4:2:0:1 | 5 | 4 | 2 | 0 | 1 | 3 |
| 23 | 17 | 46 | 5:4:3:0:0 | 5 | 4 | 3 | 0 | 0 | 7 |
| 24 | 17 | 46 | 5:5:0:1:0 | 5 | 5 | 0 | 1 | 0 | 4 |
| 25 | 17 | 46 | 5:5:1:0:0 | 5 | 5 | 1 | 0 | 0 | 18 |
| 26 | 17 | 46 | 5:5:1:1:0 | 5 | 5 | 1 | 1 | 0 | 3 |
| 27 | 17 | 46 | 5:5:2:0:0 | 5 | 5 | 2 | 0 | 0 | 15 |
| 28 | 17 | 46 | 5:5:3:0:0 | 5 | 5 | 3 | 0 | 0 | 12 |
| 29 | 17 | 46 | 5:6:1:0:0 | 5 | 6 | 1 | 0 | 0 | 16 |
| 30 | 17 | 46 | 5:6:1:0:1 | 5 | 6 | 1 | 0 | 1 | 8 |
| 31 | 17 | 46 | 5:6:2:0:1 | 5 | 6 | 2 | 0 | 1 | 9 |
| 32 | 17 | 46 | 5:6:3:0:0 | 5 | 6 | 3 | 0 | 0 | 17 |
| 33 | 17 | 46 | 5:6:4:0:0 | 5 | 6 | 4 | 0 | 0 | 14 |
| 34 | 17 | 46 | 5:7:0:1:0 | 5 | 7 | 0 | 1 | 0 | 8 |
| 35 | 17 | 46 | 5:7:1:1:0 | 5 | 7 | 1 | 1 | 0 | 9 |
| 36 | 17 | 46 | 6:3:1:0:0 | 6 | 3 | 1 | 0 | 0 | 9 |
| 37 | 17 | 46 | 6:4:2:0:0 | 6 | 4 | 2 | 0 | 0 | 2 |
| 38 | 17 | 46 | 6:5:4:0:0 | 6 | 5 | 4 | 0 | 0 | 3 |
| 39 | 17 | 46 | 6:6:1:0:0 | 6 | 6 | 1 | 0 | 0 | 19 |
| 40 | 17 | 46 | 6:6:1:0:1 | 6 | 6 | 1 | 0 | 1 | 7 |
| 41 | 17 | 46 | 6:6:2:0:0 | 6 | 6 | 2 | 0 | 0 | 12 |
| 42 | 17 | 46 | 6:6:3:0:0 | 6 | 6 | 3 | 0 | 0 | 14 |
| 43 | 17 | 46 | 6:6:4:0:0 | 6 | 6 | 4 | 0 | 0 | 10 |
| 44 | 17 | 46 | 6:7:0:1:0 | 6 | 7 | 0 | 1 | 0 | 7 |
| 45 | 17 | 46 | 6:7:1:0:0 | 6 | 7 | 1 | 0 | 0 | 17 |
| 46 | 17 | 46 | 7:6:2:0:0 | 7 | 6 | 2 | 0 | 0 | 3 |
| 47 | 61 | 42 | 2:5:0:0:0 | 2 | 5 | 0 | 0 | 0 | 13 |
| 48 | 61 | 42 | 2:6:0:0:0 | 2 | 6 | 0 | 0 | 0 | 13 |
| 49 | 61 | 42 | 2:7:0:0:0 | 2 | 7 | 0 | 0 | 0 | 11 |
| 50 | 61 | 42 | 2:8:0:0:0 | 2 | 8 | 0 | 0 | 0 | 10 |
| 51 | 61 | 42 | 3:4:1:0:0 | 3 | 4 | 1 | 0 | 0 | 8 |
| 52 | 61 | 42 | 3:5:0:0:0 | 3 | 5 | 0 | 0 | 0 | 11 |
| 53 | 61 | 42 | 3:5:0:0:1 | 3 | 5 | 0 | 0 | 1 | 6 |
| 54 | 61 | 42 | 3:5:1:0:0 | 3 | 5 | 1 | 0 | 0 | 14 |

| | | | | | | | | | |
|---|---|---|---|---|---|---|---|---|---|
| 55 | 61 | 42 | 3:5:2:0:0 | 3 | 5 | 2 | 0 | 0 | 13 |
| 56 | 61 | 42 | 3:6:0:0:1 | 3 | 6 | 0 | 0 | 1 | 8 |
| 57 | 61 | 42 | 3:6:1:0:0 | 3 | 6 | 1 | 0 | 0 | 13 |
| 58 | 61 | 42 | 4:5:0:0:0 | 4 | 5 | 0 | 0 | 0 | 14 |
| 59 | 61 | 42 | 4:5:0:0:1 | 4 | 5 | 0 | 0 | 1 | 7 |
| 60 | 61 | 42 | 4:5:1:0:0 | 4 | 5 | 1 | 0 | 0 | 19 |
| 61 | 61 | 42 | 4:5:1:0:1 | 4 | 5 | 1 | 0 | 1 | 14 |
| 62 | 61 | 42 | 4:5:2:0:0 | 4 | 5 | 2 | 0 | 0 | 18 |
| 63 | 61 | 42 | 4:5:3:0:0 | 4 | 5 | 3 | 0 | 0 | 17 |
| 64 | 61 | 42 | 4:6:0:1:0 | 4 | 6 | 0 | 1 | 0 | 14 |
| 65 | 61 | 42 | 4:6:1:0:0 | 4 | 6 | 1 | 0 | 0 | 3 |
| 66 | 61 | 42 | 4:6:2:0:0 | 4 | 6 | 2 | 0 | 0 | 5 |
| 67 | 61 | 42 | 4:7:1:0:0 | 4 | 7 | 1 | 0 | 0 | 5 |
| 68 | 61 | 42 | 4:7:2:0:0 | 4 | 7 | 2 | 0 | 0 | 5 |
| 69 | 61 | 42 | 5:5:1:0:0 | 5 | 5 | 1 | 0 | 0 | 18 |
| 70 | 61 | 42 | 5:5:2:0:0 | 5 | 5 | 2 | 0 | 0 | 15 |
| 71 | 61 | 42 | 5:5:3:0:0 | 5 | 5 | 3 | 0 | 0 | 12 |
| 72 | 61 | 42 | 5:6:0:0:0 | 5 | 6 | 0 | 0 | 0 | 9 |
| 73 | 61 | 42 | 5:6:2:0:0 | 5 | 6 | 2 | 0 | 0 | 16 |
| 74 | 61 | 42 | 5:6:2:0:1 | 5 | 6 | 2 | 0 | 1 | 9 |
| 75 | 61 | 42 | 5:6:3:0:0 | 5 | 6 | 3 | 0 | 0 | 17 |
| 76 | 61 | 42 | 5:6:4:0:0 | 5 | 6 | 4 | 0 | 0 | 14 |
| 77 | 61 | 42 | 5:7:1:1:0 | 5 | 7 | 1 | 1 | 0 | 9 |
| 78 | 61 | 42 | 5:7:2:0:0 | 5 | 7 | 2 | 0 | 0 | 3 |
| 79 | 61 | 42 | 5:7:3:0:0 | 5 | 7 | 3 | 0 | 0 | 1 |
| 80 | 61 | 42 | 6:6:0:0:0 | 6 | 6 | 0 | 0 | 0 | 6 |
| 81 | 61 | 42 | 6:6:1:0:0 | 6 | 6 | 1 | 0 | 0 | 19 |
| 82 | 61 | 42 | 6:6:2:0:0 | 6 | 6 | 2 | 0 | 0 | 12 |
| 83 | 61 | 42 | 6:6:2:0:1 | 6 | 6 | 2 | 0 | 1 | 8 |
| 84 | 61 | 42 | 6:6:3:0:0 | 6 | 6 | 3 | 0 | 0 | 14 |
| 85 | 61 | 42 | 6:6:4:0:0 | 6 | 6 | 4 | 0 | 0 | 10 |
| 86 | 61 | 42 | 6:7:1:1:0 | 6 | 7 | 1 | 1 | 0 | 8 |
| 87 | 61 | 42 | 6:7:4:0:0 | 6 | 7 | 4 | 0 | 0 | 16 |
| 88 | 61 | 42 | 8:9:0:0:0 | 8 | 9 | 0 | 0 | 0 | 5 |
| 89 | 74 | 228 | 10:10:1:0:0 | 10 | 10 | 1 | 0 | 0 | 2 |
| 90 | 74 | 228 | 10:3:0:3:0 | 10 | 3 | 0 | 3 | 0 | 1 |
| 91 | 74 | 228 | 10:8:0:0:0 | 10 | 8 | 0 | 0 | 0 | 1 |
| 92 | 74 | 228 | 10:9:0:0:0 | 10 | 9 | 0 | 0 | 0 | 2 |
| 93 | 74 | 228 | 10:9:1:0:0 | 10 | 9 | 1 | 0 | 0 | 2 |
| 94 | 74 | 228 | 3:3:1:0:1 | 3 | 3 | 1 | 0 | 1 | 2 |
| 95 | 74 | 228 | 3:3:1:1:0 | 3 | 3 | 1 | 1 | 0 | 1 |
| 96 | 74 | 228 | 3:3:2:0:1 | 3 | 3 | 2 | 0 | 1 | 2 |
| 97 | 74 | 228 | 3:4:0:1:0 | 3 | 4 | 0 | 1 | 0 | 2 |
| 98 | 74 | 228 | 3:4:1:0:0 | 3 | 4 | 1 | 0 | 0 | 8 |
| 99 | 74 | 228 | 3:4:1:1:0 | 3 | 4 | 1 | 1 | 0 | 2 |
| 100 | 74 | 228 | 3:4:2:0:0 | 3 | 4 | 2 | 0 | 0 | 5 |
| 101 | 74 | 228 | 3:4:2:0:1 | 3 | 4 | 2 | 0 | 1 | 4 |
| 102 | 74 | 228 | 3:5:1:1:0 | 3 | 5 | 1 | 1 | 0 | 4 |
| 103 | 74 | 228 | 3:5:2:0:0 | 3 | 5 | 2 | 0 | 0 | 13 |
| 104 | 74 | 228 | 3:5:2:0:1 | 3 | 5 | 2 | 0 | 1 | 1 |
| 105 | 74 | 228 | 3:6:1:1:0 | 3 | 6 | 1 | 1 | 0 | 1 |
| 106 | 74 | 228 | 3:6:2:0:0 | 3 | 6 | 2 | 0 | 0 | 6 |
| 107 | 74 | 228 | 3:6:2:0:1 | 3 | 6 | 2 | 0 | 1 | 1 |
| 108 | 74 | 228 | 4:3:0:0:0 | 4 | 3 | 0 | 0 | 0 | 6 |
| 109 | 74 | 228 | 4:3:1:0:0 | 4 | 3 | 1 | 0 | 0 | 14 |
| 110 | 74 | 228 | 4:3:1:1:0 | 4 | 3 | 1 | 1 | 0 | 1 |
| 111 | 74 | 228 | 4:3:2:0:0 | 4 | 3 | 2 | 0 | 0 | 1 |
| 112 | 74 | 228 | 4:3:2:0:1 | 4 | 3 | 2 | 0 | 1 | 2 |
| 113 | 74 | 228 | 4:4:0:2:0 | 4 | 4 | 0 | 2 | 0 | 1 |

| | | | | | | | | | |
|---|---|---|---|---|---|---|---|---|---|
| 114 | 74 | 228 | 4:4:1:0:0 | 4 | 4 | 1 | 0 | 0 | 12 |
| 115 | 74 | 228 | 4:4:1:0:1 | 4 | 4 | 1 | 0 | 1 | 7 |
| 116 | 74 | 228 | 4:4:1:1:0 | 4 | 4 | 1 | 1 | 0 | 2 |
| 117 | 74 | 228 | 4:4:1:1:1 | 4 | 4 | 1 | 1 | 1 | 1 |
| 118 | 74 | 228 | 4:4:1:2:0 | 4 | 4 | 1 | 2 | 0 | 1 |
| 119 | 74 | 228 | 4:4:2:0:0 | 4 | 4 | 2 | 0 | 0 | 8 |
| 120 | 74 | 228 | 4:4:2:0:1 | 4 | 4 | 2 | 0 | 1 | 8 |
| 121 | 74 | 228 | 4:4:3:0:1 | 4 | 4 | 3 | 0 | 1 | 4 |
| 122 | 74 | 228 | 4:5:0:0:0 | 4 | 5 | 0 | 0 | 0 | 14 |
| 123 | 74 | 228 | 4:5:0:1:0 | 4 | 5 | 0 | 1 | 0 | 7 |
| 124 | 74 | 228 | 4:5:0:2:0 | 4 | 5 | 0 | 2 | 0 | 1 |
| 125 | 74 | 228 | 4:5:1:0:0 | 4 | 5 | 1 | 0 | 0 | 19 |
| 126 | 74 | 228 | 4:5:1:0:1 | 4 | 5 | 1 | 0 | 1 | 14 |
| 127 | 74 | 228 | 4:5:1:0:2 | 4 | 5 | 1 | 0 | 2 | 3 |
| 128 | 74 | 228 | 4:5:1:1:0 | 4 | 5 | 1 | 1 | 0 | 8 |
| 129 | 74 | 228 | 4:5:2:0:0 | 4 | 5 | 2 | 0 | 0 | 18 |
| 130 | 74 | 228 | 4:5:2:0:1 | 4 | 5 | 2 | 0 | 1 | 9 |
| 131 | 74 | 228 | 4:5:2:1:0 | 4 | 5 | 2 | 1 | 0 | 4 |
| 132 | 74 | 228 | 4:5:3:0:0 | 4 | 5 | 3 | 0 | 0 | 17 |
| 133 | 74 | 228 | 4:5:3:0:1 | 4 | 5 | 3 | 0 | 1 | 1 |
| 134 | 74 | 228 | 4:6:0:1:0 | 4 | 6 | 0 | 1 | 0 | 14 |
| 135 | 74 | 228 | 4:6:0:1:1 | 4 | 6 | 0 | 1 | 1 | 3 |
| 136 | 74 | 228 | 4:6:1:1:0 | 4 | 6 | 1 | 1 | 0 | 9 |
| 137 | 74 | 228 | 4:6:2:0:0 | 4 | 6 | 2 | 0 | 0 | 5 |
| 138 | 74 | 228 | 4:6:2:1:0 | 4 | 6 | 2 | 1 | 0 | 1 |
| 139 | 74 | 228 | 4:6:3:0:0 | 4 | 6 | 3 | 0 | 0 | 3 |
| 140 | 74 | 228 | 5:3:0:0:0 | 5 | 3 | 0 | 0 | 0 | 6 |
| 141 | 74 | 228 | 5:3:1:0:0 | 5 | 3 | 1 | 0 | 0 | 11 |
| 142 | 74 | 228 | 5:3:1:1:0 | 5 | 3 | 1 | 1 | 0 | 1 |
| 143 | 74 | 228 | 5:3:2:0:0 | 5 | 3 | 2 | 0 | 0 | 1 |
| 144 | 74 | 228 | 5:3:2:0:1 | 5 | 3 | 2 | 0 | 1 | 2 |
| 145 | 74 | 228 | 5:3:3:0:1 | 5 | 3 | 3 | 0 | 1 | 1 |
| 146 | 74 | 228 | 5:4:0:0:1 | 5 | 4 | 0 | 0 | 1 | 1 |
| 147 | 74 | 228 | 5:4:0:2:0 | 5 | 4 | 0 | 2 | 0 | 1 |
| 148 | 74 | 228 | 5:4:1:0:0 | 5 | 4 | 1 | 0 | 0 | 16 |
| 149 | 74 | 228 | 5:4:1:0:1 | 5 | 4 | 1 | 0 | 1 | 4 |
| 150 | 74 | 228 | 5:4:1:0:2 | 5 | 4 | 1 | 0 | 2 | 1 |
| 151 | 74 | 228 | 5:4:1:1:0 | 5 | 4 | 1 | 1 | 0 | 2 |
| 152 | 74 | 228 | 5:4:2:0:0 | 5 | 4 | 2 | 0 | 0 | 8 |
| 153 | 74 | 228 | 5:4:2:0:1 | 5 | 4 | 2 | 0 | 1 | 3 |
| 154 | 74 | 228 | 5:4:2:1:0 | 5 | 4 | 2 | 1 | 0 | 1 |
| 155 | 74 | 228 | 5:4:3:0:0 | 5 | 4 | 3 | 0 | 0 | 7 |
| 156 | 74 | 228 | 5:5:0:1:0 | 5 | 5 | 0 | 1 | 0 | 4 |
| 157 | 74 | 228 | 5:5:0:1:1 | 5 | 5 | 0 | 1 | 1 | 1 |
| 158 | 74 | 228 | 5:5:1:0:0 | 5 | 5 | 1 | 0 | 0 | 18 |
| 159 | 74 | 228 | 5:5:1:0:1 | 5 | 5 | 1 | 0 | 1 | 7 |
| 160 | 74 | 228 | 5:5:1:0:2 | 5 | 5 | 1 | 0 | 2 | 2 |
| 161 | 74 | 228 | 5:5:1:1:0 | 5 | 5 | 1 | 1 | 0 | 3 |
| 162 | 74 | 228 | 5:5:2:0:0 | 5 | 5 | 2 | 0 | 0 | 15 |
| 163 | 74 | 228 | 5:5:2:0:1 | 5 | 5 | 2 | 0 | 1 | 7 |
| 164 | 74 | 228 | 5:5:3:0:0 | 5 | 5 | 3 | 0 | 0 | 12 |
| 165 | 74 | 228 | 5:5:4:0:0 | 5 | 5 | 4 | 0 | 0 | 2 |
| 166 | 74 | 228 | 5:5:4:0:1 | 5 | 5 | 4 | 0 | 1 | 3 |
| 167 | 74 | 228 | 5:6:0:1:0 | 5 | 6 | 0 | 1 | 0 | 7 |
| 168 | 74 | 228 | 5:6:0:1:1 | 5 | 6 | 0 | 1 | 1 | 2 |
| 169 | 74 | 228 | 5:6:1:0:0 | 5 | 6 | 1 | 0 | 0 | 16 |
| 170 | 74 | 228 | 5:6:1:0:2 | 5 | 6 | 1 | 0 | 2 | 4 |
| 171 | 74 | 228 | 5:6:1:0:3 | 5 | 6 | 1 | 0 | 3 | 1 |
| 172 | 74 | 228 | 5:6:1:1:0 | 5 | 6 | 1 | 1 | 0 | 7 |

| | | | | | | | | | |
|---|---|---|---|---|---|---|---|---|---|
| 173 | 74 | 228 | 5:6:2:0:0 | 5 | 6 | 2 | 0 | 0 | 16 |
| 174 | 74 | 228 | 5:6:2:0:1 | 5 | 6 | 2 | 0 | 1 | 9 |
| 175 | 74 | 228 | 5:6:3:0:0 | 5 | 6 | 3 | 0 | 0 | 17 |
| 176 | 74 | 228 | 5:6:3:0:1 | 5 | 6 | 3 | 0 | 1 | 7 |
| 177 | 74 | 228 | 5:6:3:1:0 | 5 | 6 | 3 | 1 | 0 | 3 |
| 178 | 74 | 228 | 5:6:4:0:0 | 5 | 6 | 4 | 0 | 0 | 14 |
| 179 | 74 | 228 | 5:6:4:0:1 | 5 | 6 | 4 | 0 | 1 | 1 |
| 180 | 74 | 228 | 5:7:0:1:1 | 5 | 7 | 0 | 1 | 1 | 4 |
| 181 | 74 | 228 | 5:7:0:1:2 | 5 | 7 | 0 | 1 | 2 | 1 |
| 182 | 74 | 228 | 5:7:1:1:0 | 5 | 7 | 1 | 1 | 0 | 9 |
| 183 | 74 | 228 | 5:7:2:1:0 | 5 | 7 | 2 | 1 | 0 | 7 |
| 184 | 74 | 228 | 5:7:3:1:0 | 5 | 7 | 3 | 1 | 0 | 1 |
| 185 | 74 | 228 | 6:3:1:0:0 | 6 | 3 | 1 | 0 | 0 | 9 |
| 186 | 74 | 228 | 6:3:1:0:1 | 6 | 3 | 1 | 0 | 1 | 1 |
| 187 | 74 | 228 | 6:3:2:0:0 | 6 | 3 | 2 | 0 | 0 | 1 |
| 188 | 74 | 228 | 6:3:2:0:1 | 6 | 3 | 2 | 0 | 1 | 1 |
| 189 | 74 | 228 | 6:3:3:0:0 | 6 | 3 | 3 | 0 | 0 | 1 |
| 190 | 74 | 228 | 6:4:0:1:0 | 6 | 4 | 0 | 1 | 0 | 1 |
| 191 | 74 | 228 | 6:4:1:0:0 | 6 | 4 | 1 | 0 | 0 | 8 |
| 192 | 74 | 228 | 6:4:1:1:0 | 6 | 4 | 1 | 1 | 0 | 1 |
| 193 | 74 | 228 | 6:4:2:0:0 | 6 | 4 | 2 | 0 | 0 | 2 |
| 194 | 74 | 228 | 6:4:3:0:0 | 6 | 4 | 3 | 0 | 0 | 1 |
| 195 | 74 | 228 | 6:5:1:0:0 | 6 | 5 | 1 | 0 | 0 | 10 |
| 196 | 74 | 228 | 6:5:2:0:0 | 6 | 5 | 2 | 0 | 0 | 4 |
| 197 | 74 | 228 | 6:5:2:0:1 | 6 | 5 | 2 | 0 | 1 | 4 |
| 198 | 74 | 228 | 6:5:2:0:2 | 6 | 5 | 2 | 0 | 2 | 3 |
| 199 | 74 | 228 | 6:5:3:0:0 | 6 | 5 | 3 | 0 | 0 | 2 |
| 200 | 74 | 228 | 6:5:3:0:1 | 6 | 5 | 3 | 0 | 1 | 1 |
| 201 | 74 | 228 | 6:5:3:0:2 | 6 | 5 | 3 | 0 | 2 | 2 |
| 202 | 74 | 228 | 6:5:4:0:0 | 6 | 5 | 4 | 0 | 0 | 3 |
| 203 | 74 | 228 | 6:5:4:0:1 | 6 | 5 | 4 | 0 | 1 | 3 |
| 204 | 74 | 228 | 6:6:1:0:0 | 6 | 6 | 1 | 0 | 0 | 19 |
| 205 | 74 | 228 | 6:6:1:0:1 | 6 | 6 | 1 | 0 | 1 | 7 |
| 206 | 74 | 228 | 6:6:1:0:2 | 6 | 6 | 1 | 0 | 2 | 2 |
| 207 | 74 | 228 | 6:6:1:1:0 | 6 | 6 | 1 | 1 | 0 | 4 |
| 208 | 74 | 228 | 6:6:1:1:1 | 6 | 6 | 1 | 1 | 1 | 3 |
| 209 | 74 | 228 | 6:6:2:0:0 | 6 | 6 | 2 | 0 | 0 | 12 |
| 210 | 74 | 228 | 6:6:2:0:1 | 6 | 6 | 2 | 0 | 1 | 8 |
| 211 | 74 | 228 | 6:6:2:0:2 | 6 | 6 | 2 | 0 | 2 | 4 |
| 212 | 74 | 228 | 6:6:2:1:0 | 6 | 6 | 2 | 1 | 0 | 1 |
| 213 | 74 | 228 | 6:6:2:1:1 | 6 | 6 | 2 | 1 | 1 | 2 |
| 214 | 74 | 228 | 6:6:3:0:0 | 6 | 6 | 3 | 0 | 0 | 14 |
| 215 | 74 | 228 | 6:6:3:0:1 | 6 | 6 | 3 | 0 | 1 | 7 |
| 216 | 74 | 228 | 6:6:3:0:2 | 6 | 6 | 3 | 0 | 2 | 3 |
| 217 | 74 | 228 | 6:6:3:1:0 | 6 | 6 | 3 | 1 | 0 | 3 |
| 218 | 74 | 228 | 6:6:4:0:0 | 6 | 6 | 4 | 0 | 0 | 10 |
| 219 | 74 | 228 | 6:6:4:0:1 | 6 | 6 | 4 | 0 | 1 | 3 |
| 220 | 74 | 228 | 6:6:4:0:2 | 6 | 6 | 4 | 0 | 2 | 1 |
| 221 | 74 | 228 | 6:7:0:1:0 | 6 | 7 | 0 | 1 | 0 | 7 |
| 222 | 74 | 228 | 6:7:0:1:1 | 6 | 7 | 0 | 1 | 1 | 2 |
| 223 | 74 | 228 | 6:7:0:2:0 | 6 | 7 | 0 | 2 | 0 | 3 |
| 224 | 74 | 228 | 6:7:1:0:0 | 6 | 7 | 1 | 0 | 0 | 17 |
| 225 | 74 | 228 | 6:7:1:0:1 | 6 | 7 | 1 | 0 | 1 | 13 |
| 226 | 74 | 228 | 6:7:1:0:2 | 6 | 7 | 1 | 0 | 2 | 5 |
| 227 | 74 | 228 | 6:7:1:0:3 | 6 | 7 | 1 | 0 | 3 | 1 |
| 228 | 74 | 228 | 6:7:1:1:0 | 6 | 7 | 1 | 1 | 0 | 8 |
| 229 | 74 | 228 | 6:7:1:1:1 | 6 | 7 | 1 | 1 | 1 | 4 |
| 230 | 74 | 228 | 6:7:1:2:0 | 6 | 7 | 1 | 2 | 0 | 2 |
| 231 | 74 | 228 | 6:7:2:0:0 | 6 | 7 | 2 | 0 | 0 | 16 |

| | | | | | | | | | |
|---|---|---|---|---|---|---|---|---|---|
| 232 | 74 | 228 | 6:7:2:0:1 | 6 | 7 | 2 | 0 | 1 | 9 |
| 233 | 74 | 228 | 6:7:2:0:2 | 6 | 7 | 2 | 0 | 2 | 5 |
| 234 | 74 | 228 | 6:7:2:0:3 | 6 | 7 | 2 | 0 | 3 | 1 |
| 235 | 74 | 228 | 6:7:2:1:0 | 6 | 7 | 2 | 1 | 0 | 7 |
| 236 | 74 | 228 | 6:7:2:1:1 | 6 | 7 | 2 | 1 | 1 | 3 |
| 237 | 74 | 228 | 6:7:3:0:0 | 6 | 7 | 3 | 0 | 0 | 16 |
| 238 | 74 | 228 | 6:7:3:0:1 | 6 | 7 | 3 | 0 | 1 | 8 |
| 239 | 74 | 228 | 6:7:3:0:2 | 6 | 7 | 3 | 0 | 2 | 4 |
| 240 | 74 | 228 | 6:7:3:1:0 | 6 | 7 | 3 | 1 | 0 | 3 |
| 241 | 74 | 228 | 6:7:3:1:1 | 6 | 7 | 3 | 1 | 1 | 1 |
| 242 | 74 | 228 | 6:7:4:0:0 | 6 | 7 | 4 | 0 | 0 | 16 |
| 243 | 74 | 228 | 6:7:4:0:1 | 6 | 7 | 4 | 0 | 1 | 5 |
| 244 | 74 | 228 | 6:7:4:1:0 | 6 | 7 | 4 | 1 | 0 | 2 |
| 245 | 74 | 228 | 6:8:0:1:0 | 6 | 8 | 0 | 1 | 0 | 13 |
| 246 | 74 | 228 | 6:8:0:1:1 | 6 | 8 | 0 | 1 | 1 | 5 |
| 247 | 74 | 228 | 6:8:0:1:2 | 6 | 8 | 0 | 1 | 2 | 1 |
| 248 | 74 | 228 | 6:8:0:2:0 | 6 | 8 | 0 | 2 | 0 | 4 |
| 249 | 74 | 228 | 6:8:1:1:0 | 6 | 8 | 1 | 1 | 0 | 9 |
| 250 | 74 | 228 | 6:8:1:1:1 | 6 | 8 | 1 | 1 | 1 | 5 |
| 251 | 74 | 228 | 6:8:1:1:2 | 6 | 8 | 1 | 1 | 2 | 1 |
| 252 | 74 | 228 | 6:8:1:2:0 | 6 | 8 | 1 | 2 | 0 | 3 |
| 253 | 74 | 228 | 6:8:2:1:0 | 6 | 8 | 2 | 1 | 0 | 8 |
| 254 | 74 | 228 | 6:8:2:1:1 | 6 | 8 | 2 | 1 | 1 | 4 |
| 255 | 74 | 228 | 6:8:2:2:0 | 6 | 8 | 2 | 2 | 0 | 1 |
| 256 | 74 | 228 | 6:8:3:1:0 | 6 | 8 | 3 | 1 | 0 | 5 |
| 257 | 74 | 228 | 6:8:4:1:0 | 6 | 8 | 4 | 1 | 0 | 1 |
| 258 | 74 | 228 | 6:9:0:2:0 | 6 | 9 | 0 | 2 | 0 | 5 |
| 259 | 74 | 228 | 6:9:0:2:1 | 6 | 9 | 0 | 2 | 1 | 1 |
| 260 | 74 | 228 | 6:9:1:2:0 | 6 | 9 | 1 | 2 | 0 | 4 |
| 261 | 74 | 228 | 7:3:1:0:0 | 7 | 3 | 1 | 0 | 0 | 2 |
| 262 | 74 | 228 | 7:4:1:0:0 | 7 | 4 | 1 | 0 | 0 | 2 |
| 263 | 74 | 228 | 7:4:2:0:0 | 7 | 4 | 2 | 0 | 0 | 1 |
| 264 | 74 | 228 | 7:4:2:0:1 | 7 | 4 | 2 | 0 | 1 | 1 |
| 265 | 74 | 228 | 7:4:3:0:0 | 7 | 4 | 3 | 0 | 0 | 1 |
| 266 | 74 | 228 | 7:4:3:0:1 | 7 | 4 | 3 | 0 | 1 | 1 |
| 267 | 74 | 228 | 7:4:4:0:0 | 7 | 4 | 4 | 0 | 0 | 1 |
| 268 | 74 | 228 | 7:5:1:1:0 | 7 | 5 | 1 | 1 | 0 | 1 |
| 269 | 74 | 228 | 7:5:2:1:0 | 7 | 5 | 2 | 1 | 0 | 1 |
| 270 | 74 | 228 | 7:5:3:0:0 | 7 | 5 | 3 | 0 | 0 | 1 |
| 271 | 74 | 228 | 7:5:3:0:1 | 7 | 5 | 3 | 0 | 1 | 1 |
| 272 | 74 | 228 | 7:5:4:0:0 | 7 | 5 | 4 | 0 | 0 | 1 |
| 273 | 74 | 228 | 7:6:1:0:0 | 7 | 6 | 1 | 0 | 0 | 4 |
| 274 | 74 | 228 | 7:6:1:0:1 | 7 | 6 | 1 | 0 | 1 | 1 |
| 275 | 74 | 228 | 7:6:2:0:0 | 7 | 6 | 2 | 0 | 0 | 3 |
| 276 | 74 | 228 | 7:6:2:0:2 | 7 | 6 | 2 | 0 | 2 | 1 |
| 277 | 74 | 228 | 7:6:2:1:0 | 7 | 6 | 2 | 1 | 0 | 1 |
| 278 | 74 | 228 | 7:6:3:0:0 | 7 | 6 | 3 | 0 | 0 | 2 |
| 279 | 74 | 228 | 7:6:3:0:1 | 7 | 6 | 3 | 0 | 1 | 2 |
| 280 | 74 | 228 | 7:6:3:0:2 | 7 | 6 | 3 | 0 | 2 | 2 |
| 281 | 74 | 228 | 7:6:4:0:0 | 7 | 6 | 4 | 0 | 0 | 2 |
| 282 | 74 | 228 | 7:6:4:0:1 | 7 | 6 | 4 | 0 | 1 | 2 |
| 283 | 74 | 228 | 7:7:0:1:0 | 7 | 7 | 0 | 1 | 0 | 1 |
| 284 | 74 | 228 | 7:7:1:0:0 | 7 | 7 | 1 | 0 | 0 | 6 |
| 285 | 74 | 228 | 7:7:1:0:1 | 7 | 7 | 1 | 0 | 1 | 3 |
| 286 | 74 | 228 | 7:7:1:1:1 | 7 | 7 | 1 | 1 | 1 | 1 |
| 287 | 74 | 228 | 7:7:2:0:0 | 7 | 7 | 2 | 0 | 0 | 6 |
| 288 | 74 | 228 | 7:7:2:1:0 | 7 | 7 | 2 | 1 | 0 | 2 |
| 289 | 74 | 228 | 7:7:2:1:1 | 7 | 7 | 2 | 1 | 1 | 2 |
| 290 | 74 | 228 | 7:7:3:0:0 | 7 | 7 | 3 | 0 | 0 | 8 |

| | | | | | | | | | |
|---|---|---|---|---|---|---|---|---|---|
| 291 | 74 | 228 | 7:7:3:1:0 | 7 | 7 | 3 | 1 | 0 | 2 |
| 292 | 74 | 228 | 7:7:4:0:0 | 7 | 7 | 4 | 0 | 0 | 4 |
| 293 | 74 | 228 | 7:7:4:0:1 | 7 | 7 | 4 | 0 | 1 | 1 |
| 294 | 74 | 228 | 7:7:4:1:0 | 7 | 7 | 4 | 1 | 0 | 1 |
| 295 | 74 | 228 | 7:8:0:1:0 | 7 | 8 | 0 | 1 | 0 | 3 |
| 296 | 74 | 228 | 7:8:0:2:0 | 7 | 8 | 0 | 2 | 0 | 1 |
| 297 | 74 | 228 | 7:8:1:2:0 | 7 | 8 | 1 | 2 | 0 | 2 |
| 298 | 74 | 228 | 7:8:3:1:0 | 7 | 8 | 3 | 1 | 0 | 1 |
| 299 | 74 | 228 | 8:3:1:0:0 | 8 | 3 | 1 | 0 | 0 | 1 |
| 300 | 74 | 228 | 8:3:1:1:0 | 8 | 3 | 1 | 1 | 0 | 1 |
| 301 | 74 | 228 | 8:5:1:0:0 | 8 | 5 | 1 | 0 | 0 | 1 |
| 302 | 74 | 228 | 8:5:1:0:1 | 8 | 5 | 1 | 0 | 1 | 1 |
| 303 | 74 | 228 | 8:5:2:0:0 | 8 | 5 | 2 | 0 | 0 | 1 |
| 304 | 74 | 228 | 8:5:3:0:0 | 8 | 5 | 3 | 0 | 0 | 1 |
| 305 | 74 | 228 | 8:5:4:0:0 | 8 | 5 | 4 | 0 | 0 | 1 |
| 306 | 74 | 228 | 8:6:0:1:0 | 8 | 6 | 0 | 1 | 0 | 1 |
| 307 | 74 | 228 | 8:6:2:0:0 | 8 | 6 | 2 | 0 | 0 | 1 |
| 308 | 74 | 228 | 8:6:3:0:0 | 8 | 6 | 3 | 0 | 0 | 1 |
| 309 | 74 | 228 | 8:6:4:0:0 | 8 | 6 | 4 | 0 | 0 | 1 |
| 310 | 74 | 228 | 8:8:0:0:0 | 8 | 8 | 0 | 0 | 0 | 1 |
| 311 | 74 | 228 | 8:9:0:0:1 | 8 | 9 | 0 | 0 | 1 | 1 |
| 312 | 74 | 228 | 9:10:0:0:0 | 9 | 10 | 0 | 0 | 0 | 2 |
| 313 | 74 | 228 | 9:10:1:0:0 | 9 | 10 | 1 | 0 | 0 | 2 |
| 314 | 74 | 228 | 9:8:0:0:1 | 9 | 8 | 0 | 0 | 1 | 1 |
| 315 | 74 | 228 | 9:9:1:0:0 | 9 | 9 | 1 | 0 | 0 | 1 |
| 316 | 122 | 86 | 2:5:0:0:0 | 2 | 5 | 0 | 0 | 0 | 13 |
| 317 | 122 | 86 | 2:6:0:0:0 | 2 | 6 | 0 | 0 | 0 | 13 |
| 318 | 122 | 86 | 2:7:0:0:0 | 2 | 7 | 0 | 0 | 0 | 11 |
| 319 | 122 | 86 | 2:8:0:0:0 | 2 | 8 | 0 | 0 | 0 | 10 |
| 320 | 122 | 86 | 2:9:0:0:0 | 2 | 9 | 0 | 0 | 0 | 5 |
| 321 | 122 | 86 | 3:3:0:0:0 | 3 | 3 | 0 | 0 | 0 | 3 |
| 322 | 122 | 86 | 3:3:1:0:0 | 3 | 3 | 1 | 0 | 0 | 4 |
| 323 | 122 | 86 | 3:4:0:0:0 | 3 | 4 | 0 | 0 | 0 | 5 |
| 324 | 122 | 86 | 3:4:0:0:1 | 3 | 4 | 0 | 0 | 1 | 3 |
| 325 | 122 | 86 | 3:4:1:0:0 | 3 | 4 | 1 | 0 | 0 | 8 |
| 326 | 122 | 86 | 3:4:1:0:1 | 3 | 4 | 1 | 0 | 1 | 5 |
| 327 | 122 | 86 | 3:4:2:0:0 | 3 | 4 | 2 | 0 | 0 | 5 |
| 328 | 122 | 86 | 3:4:2:0:1 | 3 | 4 | 2 | 0 | 1 | 4 |
| 329 | 122 | 86 | 3:5:0:0:0 | 3 | 5 | 0 | 0 | 0 | 11 |
| 330 | 122 | 86 | 3:5:0:0:1 | 3 | 5 | 0 | 0 | 1 | 6 |
| 331 | 122 | 86 | 3:5:0:1:0 | 3 | 5 | 0 | 1 | 0 | 5 |
| 332 | 122 | 86 | 3:5:1:0:0 | 3 | 5 | 1 | 0 | 0 | 14 |
| 333 | 122 | 86 | 3:5:1:0:1 | 3 | 5 | 1 | 0 | 1 | 4 |
| 334 | 122 | 86 | 3:5:1:1:0 | 3 | 5 | 1 | 1 | 0 | 4 |
| 335 | 122 | 86 | 3:5:2:0:0 | 3 | 5 | 2 | 0 | 0 | 13 |
| 336 | 122 | 86 | 3:6:0:0:0 | 3 | 6 | 0 | 0 | 0 | 11 |
| 337 | 122 | 86 | 3:6:0:0:1 | 3 | 6 | 0 | 0 | 1 | 8 |
| 338 | 122 | 86 | 3:6:0:1:0 | 3 | 6 | 0 | 1 | 0 | 4 |
| 339 | 122 | 86 | 3:6:1:0:0 | 3 | 6 | 1 | 0 | 0 | 13 |
| 340 | 122 | 86 | 3:7:1:0:0 | 3 | 7 | 1 | 0 | 0 | 4 |
| 341 | 122 | 86 | 4:3:0:0:0 | 4 | 3 | 0 | 0 | 0 | 6 |
| 342 | 122 | 86 | 4:3:1:0:0 | 4 | 3 | 1 | 0 | 0 | 14 |
| 343 | 122 | 86 | 4:4:0:0:0 | 4 | 4 | 0 | 0 | 0 | 6 |
| 344 | 122 | 86 | 4:4:1:0:0 | 4 | 4 | 1 | 0 | 0 | 12 |
| 345 | 122 | 86 | 4:4:1:0:1 | 4 | 4 | 1 | 0 | 1 | 7 |
| 346 | 122 | 86 | 4:4:2:0:0 | 4 | 4 | 2 | 0 | 0 | 8 |
| 347 | 122 | 86 | 4:4:2:0:1 | 4 | 4 | 2 | 0 | 1 | 8 |
| 348 | 122 | 86 | 4:5:0:0:0 | 4 | 5 | 0 | 0 | 0 | 14 |
| 349 | 122 | 86 | 4:5:0:0:1 | 4 | 5 | 0 | 0 | 1 | 7 |

| | | | | | | | | | |
|---|---|---|---|---|---|---|---|---|---|
| 350 | 122 | 86 | 4:5:0:1:0 | 4 | 5 | 0 | 1 | 0 | 7 |
| 351 | 122 | 86 | 4:5:1:0:0 | 4 | 5 | 1 | 0 | 0 | 19 |
| 352 | 122 | 86 | 4:5:1:0:1 | 4 | 5 | 1 | 0 | 1 | 14 |
| 353 | 122 | 86 | 4:5:1:1:0 | 4 | 5 | 1 | 1 | 0 | 8 |
| 354 | 122 | 86 | 4:5:2:0:0 | 4 | 5 | 2 | 0 | 0 | 18 |
| 355 | 122 | 86 | 4:5:2:0:1 | 4 | 5 | 2 | 0 | 1 | 9 |
| 356 | 122 | 86 | 4:5:3:0:0 | 4 | 5 | 3 | 0 | 0 | 17 |
| 357 | 122 | 86 | 4:6:0:1:0 | 4 | 6 | 0 | 1 | 0 | 14 |
| 358 | 122 | 86 | 4:6:1:1:0 | 4 | 6 | 1 | 1 | 0 | 9 |
| 359 | 122 | 86 | 4:7:1:0:0 | 4 | 7 | 1 | 0 | 0 | 5 |
| 360 | 122 | 86 | 4:7:2:0:0 | 4 | 7 | 2 | 0 | 0 | 5 |
| 361 | 122 | 86 | 5:3:0:0:0 | 5 | 3 | 0 | 0 | 0 | 6 |
| 362 | 122 | 86 | 5:3:1:0:0 | 5 | 3 | 1 | 0 | 0 | 11 |
| 363 | 122 | 86 | 5:3:1:0:1 | 5 | 3 | 1 | 0 | 1 | 1 |
| 364 | 122 | 86 | 5:4:0:0:0 | 5 | 4 | 0 | 0 | 0 | 5 |
| 365 | 122 | 86 | 5:4:0:1:0 | 5 | 4 | 0 | 1 | 0 | 1 |
| 366 | 122 | 86 | 5:4:1:0:0 | 5 | 4 | 1 | 0 | 0 | 16 |
| 367 | 122 | 86 | 5:4:1:0:1 | 5 | 4 | 1 | 0 | 1 | 4 |
| 368 | 122 | 86 | 5:4:2:0:0 | 5 | 4 | 2 | 0 | 0 | 8 |
| 369 | 122 | 86 | 5:4:3:0:0 | 5 | 4 | 3 | 0 | 0 | 7 |
| 370 | 122 | 86 | 5:5:0:0:0 | 5 | 5 | 0 | 0 | 0 | 6 |
| 371 | 122 | 86 | 5:5:0:1:0 | 5 | 5 | 0 | 1 | 0 | 4 |
| 372 | 122 | 86 | 5:5:1:0:0 | 5 | 5 | 1 | 0 | 0 | 18 |
| 373 | 122 | 86 | 5:5:1:0:1 | 5 | 5 | 1 | 0 | 1 | 7 |
| 374 | 122 | 86 | 5:5:2:0:0 | 5 | 5 | 2 | 0 | 0 | 15 |
| 375 | 122 | 86 | 5:5:3:0:0 | 5 | 5 | 3 | 0 | 0 | 12 |
| 376 | 122 | 86 | 5:6:0:0:0 | 5 | 6 | 0 | 0 | 0 | 9 |
| 377 | 122 | 86 | 5:6:0:0:1 | 5 | 6 | 0 | 0 | 1 | 2 |
| 378 | 122 | 86 | 5:6:0:1:0 | 5 | 6 | 0 | 1 | 0 | 7 |
| 379 | 122 | 86 | 5:6:1:0:1 | 5 | 6 | 1 | 0 | 1 | 8 |
| 380 | 122 | 86 | 5:6:2:0:0 | 5 | 6 | 2 | 0 | 0 | 16 |
| 381 | 122 | 86 | 5:6:3:0:0 | 5 | 6 | 3 | 0 | 0 | 17 |
| 382 | 122 | 86 | 5:6:3:0:1 | 5 | 6 | 3 | 0 | 1 | 7 |
| 383 | 122 | 86 | 5:6:4:0:0 | 5 | 6 | 4 | 0 | 0 | 14 |
| 384 | 122 | 86 | 5:7:0:1:0 | 5 | 7 | 0 | 1 | 0 | 8 |
| 385 | 122 | 86 | 5:7:2:1:0 | 5 | 7 | 2 | 1 | 0 | 7 |
| 386 | 122 | 86 | 6:6:0:0:0 | 6 | 6 | 0 | 0 | 0 | 6 |
| 387 | 122 | 86 | 6:6:1:0:0 | 6 | 6 | 1 | 0 | 0 | 19 |
| 388 | 122 | 86 | 6:6:2:0:0 | 6 | 6 | 2 | 0 | 0 | 12 |
| 389 | 122 | 86 | 6:6:2:0:1 | 6 | 6 | 2 | 0 | 1 | 8 |
| 390 | 122 | 86 | 6:6:3:0:0 | 6 | 6 | 3 | 0 | 0 | 14 |
| 391 | 122 | 86 | 6:6:4:0:0 | 6 | 6 | 4 | 0 | 0 | 10 |
| 392 | 122 | 86 | 6:7:1:1:0 | 6 | 7 | 1 | 1 | 0 | 8 |
| 393 | 122 | 86 | 6:7:2:0:1 | 6 | 7 | 2 | 0 | 1 | 9 |
| 394 | 122 | 86 | 6:7:2:0:2 | 6 | 7 | 2 | 0 | 2 | 5 |
| 395 | 122 | 86 | 6:7:4:0:0 | 6 | 7 | 4 | 0 | 0 | 16 |
| 396 | 122 | 86 | 6:8:1:1:0 | 6 | 8 | 1 | 1 | 0 | 9 |
| 397 | 122 | 86 | 6:8:1:1:1 | 6 | 8 | 1 | 1 | 1 | 5 |
| 398 | 122 | 86 | 6:9:0:2:0 | 6 | 9 | 0 | 2 | 0 | 5 |
| 399 | 122 | 86 | 7:7:1:0:0 | 7 | 7 | 1 | 0 | 0 | 6 |
| 400 | 122 | 86 | 7:7:3:0:0 | 7 | 7 | 3 | 0 | 0 | 8 |
| 401 | 149 | 123 | 2:5:0:0:0 | 2 | 5 | 0 | 0 | 0 | 13 |
| 402 | 149 | 123 | 2:6:0:0:0 | 2 | 6 | 0 | 0 | 0 | 13 |
| 403 | 149 | 123 | 2:7:0:0:0 | 2 | 7 | 0 | 0 | 0 | 11 |
| 404 | 149 | 123 | 2:8:0:0:0 | 2 | 8 | 0 | 0 | 0 | 10 |
| 405 | 149 | 123 | 3:3:1:0:0 | 3 | 3 | 1 | 0 | 0 | 4 |
| 406 | 149 | 123 | 3:4:2:0:0 | 3 | 4 | 2 | 0 | 0 | 5 |
| 407 | 149 | 123 | 3:5:1:0:0 | 3 | 5 | 1 | 0 | 0 | 14 |
| 408 | 149 | 123 | 3:5:2:0:0 | 3 | 5 | 2 | 0 | 0 | 13 |

| | | | | | | | | | |
|---|---|---|---|---|---|---|---|---|---|
| 409 | 149 | 123 | 3:6:1:0:0 | 3 | 6 | 1 | 0 | 0 | 13 |
| 410 | 149 | 123 | 3:6:2:0:0 | 3 | 6 | 2 | 0 | 0 | 6 |
| 411 | 149 | 123 | 4:3:1:0:0 | 4 | 3 | 1 | 0 | 0 | 14 |
| 412 | 149 | 123 | 4:4:1:0:0 | 4 | 4 | 1 | 0 | 0 | 12 |
| 413 | 149 | 123 | 4:4:2:0:0 | 4 | 4 | 2 | 0 | 0 | 8 |
| 414 | 149 | 123 | 4:5:1:0:0 | 4 | 5 | 1 | 0 | 0 | 19 |
| 415 | 149 | 123 | 4:5:1:0:1 | 4 | 5 | 1 | 0 | 1 | 14 |
| 416 | 149 | 123 | 4:5:1:0:2 | 4 | 5 | 1 | 0 | 2 | 3 |
| 417 | 149 | 123 | 4:5:2:0:0 | 4 | 5 | 2 | 0 | 0 | 18 |
| 418 | 149 | 123 | 4:5:2:0:1 | 4 | 5 | 2 | 0 | 1 | 9 |
| 419 | 149 | 123 | 4:5:3:0:0 | 4 | 5 | 3 | 0 | 0 | 17 |
| 420 | 149 | 123 | 4:5:4:0:2 | 4 | 5 | 4 | 0 | 2 | 1 |
| 421 | 149 | 123 | 4:6:0:1:0 | 4 | 6 | 0 | 1 | 0 | 14 |
| 422 | 149 | 123 | 4:6:0:1:1 | 4 | 6 | 0 | 1 | 1 | 3 |
| 423 | 149 | 123 | 4:6:1:1:0 | 4 | 6 | 1 | 1 | 0 | 9 |
| 424 | 149 | 123 | 4:6:3:0:1 | 4 | 6 | 3 | 0 | 1 | 1 |
| 425 | 149 | 123 | 4:6:3:1:1 | 4 | 6 | 3 | 1 | 1 | 1 |
| 426 | 149 | 123 | 4:7:2:1:0 | 4 | 7 | 2 | 1 | 0 | 1 |
| 427 | 149 | 123 | 4:7:2:2:0 | 4 | 7 | 2 | 2 | 0 | 1 |
| 428 | 149 | 123 | 5:3:1:0:0 | 5 | 3 | 1 | 0 | 0 | 11 |
| 429 | 149 | 123 | 5:3:2:0:1 | 5 | 3 | 2 | 0 | 1 | 2 |
| 430 | 149 | 123 | 5:4:1:0:0 | 5 | 4 | 1 | 0 | 0 | 16 |
| 431 | 149 | 123 | 5:4:1:1:0 | 5 | 4 | 1 | 1 | 0 | 2 |
| 432 | 149 | 123 | 5:4:2:0:0 | 5 | 4 | 2 | 0 | 0 | 8 |
| 433 | 149 | 123 | 5:4:2:0:1 | 5 | 4 | 2 | 0 | 1 | 3 |
| 434 | 149 | 123 | 5:4:3:0:0 | 5 | 4 | 3 | 0 | 0 | 7 |
| 435 | 149 | 123 | 5:5:1:0:0 | 5 | 5 | 1 | 0 | 0 | 18 |
| 436 | 149 | 123 | 5:5:1:1:0 | 5 | 5 | 1 | 1 | 0 | 3 |
| 437 | 149 | 123 | 5:5:2:0:0 | 5 | 5 | 2 | 0 | 0 | 15 |
| 438 | 149 | 123 | 5:5:2:0:1 | 5 | 5 | 2 | 0 | 1 | 7 |
| 439 | 149 | 123 | 5:5:3:0:0 | 5 | 5 | 3 | 0 | 0 | 12 |
| 440 | 149 | 123 | 5:5:4:0:1 | 5 | 5 | 4 | 0 | 1 | 3 |
| 441 | 149 | 123 | 5:6:1:0:0 | 5 | 6 | 1 | 0 | 0 | 16 |
| 442 | 149 | 123 | 5:6:1:0:2 | 5 | 6 | 1 | 0 | 2 | 4 |
| 443 | 149 | 123 | 5:6:1:1:0 | 5 | 6 | 1 | 1 | 0 | 7 |
| 444 | 149 | 123 | 5:6:2:0:0 | 5 | 6 | 2 | 0 | 0 | 16 |
| 445 | 149 | 123 | 5:6:2:0:1 | 5 | 6 | 2 | 0 | 1 | 9 |
| 446 | 149 | 123 | 5:6:2:0:2 | 5 | 6 | 2 | 0 | 2 | 2 |
| 447 | 149 | 123 | 5:6:3:0:0 | 5 | 6 | 3 | 0 | 0 | 17 |
| 448 | 149 | 123 | 5:6:3:0:1 | 5 | 6 | 3 | 0 | 1 | 7 |
| 449 | 149 | 123 | 5:6:3:1:0 | 5 | 6 | 3 | 1 | 0 | 3 |
| 450 | 149 | 123 | 5:6:4:0:0 | 5 | 6 | 4 | 0 | 0 | 14 |
| 451 | 149 | 123 | 5:7:0:1:1 | 5 | 7 | 0 | 1 | 1 | 4 |
| 452 | 149 | 123 | 5:7:1:1:0 | 5 | 7 | 1 | 1 | 0 | 9 |
| 453 | 149 | 123 | 5:7:1:1:1 | 5 | 7 | 1 | 1 | 1 | 2 |
| 454 | 149 | 123 | 5:7:2:1:0 | 5 | 7 | 2 | 1 | 0 | 7 |
| 455 | 149 | 123 | 5:8:0:2:0 | 5 | 8 | 0 | 2 | 0 | 2 |
| 456 | 149 | 123 | 6:3:1:0:0 | 6 | 3 | 1 | 0 | 0 | 9 |
| 457 | 149 | 123 | 6:4:1:0:0 | 6 | 4 | 1 | 0 | 0 | 8 |
| 458 | 149 | 123 | 6:5:1:0:0 | 6 | 5 | 1 | 0 | 0 | 10 |
| 459 | 149 | 123 | 6:5:2:0:1 | 6 | 5 | 2 | 0 | 1 | 4 |
| 460 | 149 | 123 | 6:5:2:0:2 | 6 | 5 | 2 | 0 | 2 | 3 |
| 461 | 149 | 123 | 6:5:3:0:0 | 6 | 5 | 3 | 0 | 0 | 2 |
| 462 | 149 | 123 | 6:5:4:0:0 | 6 | 5 | 4 | 0 | 0 | 3 |
| 463 | 149 | 123 | 6:5:4:0:1 | 6 | 5 | 4 | 0 | 1 | 3 |
| 464 | 149 | 123 | 6:6:1:0:0 | 6 | 6 | 1 | 0 | 0 | 19 |
| 465 | 149 | 123 | 6:6:1:0:1 | 6 | 6 | 1 | 0 | 1 | 7 |
| 466 | 149 | 123 | 6:6:1:1:0 | 6 | 6 | 1 | 1 | 0 | 4 |
| 467 | 149 | 123 | 6:6:1:1:1 | 6 | 6 | 1 | 1 | 1 | 3 |

| | | | | | | | | | |
|---|---|---|---|---|---|---|---|---|---|
| 468 | 149 | 123 | 6:6:2:0:0 | 6 | 6 | 2 | 0 | 0 | 12 |
| 469 | 149 | 123 | 6:6:2:0:1 | 6 | 6 | 2 | 0 | 1 | 8 |
| 470 | 149 | 123 | 6:6:2:0:2 | 6 | 6 | 2 | 0 | 2 | 4 |
| 471 | 149 | 123 | 6:6:3:0:0 | 6 | 6 | 3 | 0 | 0 | 14 |
| 472 | 149 | 123 | 6:6:3:0:1 | 6 | 6 | 3 | 0 | 1 | 7 |
| 473 | 149 | 123 | 6:6:3:0:2 | 6 | 6 | 3 | 0 | 2 | 3 |
| 474 | 149 | 123 | 6:6:3:1:0 | 6 | 6 | 3 | 1 | 0 | 3 |
| 475 | 149 | 123 | 6:6:4:0:0 | 6 | 6 | 4 | 0 | 0 | 10 |
| 476 | 149 | 123 | 6:6:4:0:1 | 6 | 6 | 4 | 0 | 1 | 3 |
| 477 | 149 | 123 | 6:7:0:1:0 | 6 | 7 | 0 | 1 | 0 | 7 |
| 478 | 149 | 123 | 6:7:0:2:0 | 6 | 7 | 0 | 2 | 0 | 3 |
| 479 | 149 | 123 | 6:7:1:0:0 | 6 | 7 | 1 | 0 | 0 | 17 |
| 480 | 149 | 123 | 6:7:1:0:1 | 6 | 7 | 1 | 0 | 1 | 13 |
| 481 | 149 | 123 | 6:7:1:1:0 | 6 | 7 | 1 | 1 | 0 | 8 |
| 482 | 149 | 123 | 6:7:1:1:1 | 6 | 7 | 1 | 1 | 1 | 4 |
| 483 | 149 | 123 | 6:7:2:0:0 | 6 | 7 | 2 | 0 | 0 | 16 |
| 484 | 149 | 123 | 6:7:2:0:1 | 6 | 7 | 2 | 0 | 1 | 9 |
| 485 | 149 | 123 | 6:7:2:0:2 | 6 | 7 | 2 | 0 | 2 | 5 |
| 486 | 149 | 123 | 6:7:2:1:0 | 6 | 7 | 2 | 1 | 0 | 7 |
| 487 | 149 | 123 | 6:7:2:1:1 | 6 | 7 | 2 | 1 | 1 | 3 |
| 488 | 149 | 123 | 6:7:3:0:0 | 6 | 7 | 3 | 0 | 0 | 16 |
| 489 | 149 | 123 | 6:7:3:0:1 | 6 | 7 | 3 | 0 | 1 | 8 |
| 490 | 149 | 123 | 6:7:3:0:2 | 6 | 7 | 3 | 0 | 2 | 4 |
| 491 | 149 | 123 | 6:7:3:1:0 | 6 | 7 | 3 | 1 | 0 | 3 |
| 492 | 149 | 123 | 6:7:4:0:0 | 6 | 7 | 4 | 0 | 0 | 16 |
| 493 | 149 | 123 | 6:7:4:0:1 | 6 | 7 | 4 | 0 | 1 | 5 |
| 494 | 149 | 123 | 6:7:4:1:0 | 6 | 7 | 4 | 1 | 0 | 2 |
| 495 | 149 | 123 | 6:8:0:1:0 | 6 | 8 | 0 | 1 | 0 | 13 |
| 496 | 149 | 123 | 6:8:0:2:0 | 6 | 8 | 0 | 2 | 0 | 4 |
| 497 | 149 | 123 | 6:8:1:1:0 | 6 | 8 | 1 | 1 | 0 | 9 |
| 498 | 149 | 123 | 6:8:1:1:1 | 6 | 8 | 1 | 1 | 1 | 5 |
| 499 | 149 | 123 | 6:8:1:2:0 | 6 | 8 | 1 | 2 | 0 | 3 |
| 500 | 149 | 123 | 6:8:2:1:0 | 6 | 8 | 2 | 1 | 0 | 8 |
| 501 | 149 | 123 | 6:8:2:1:1 | 6 | 8 | 2 | 1 | 1 | 4 |
| 502 | 149 | 123 | 6:8:3:1:0 | 6 | 8 | 3 | 1 | 0 | 5 |
| 503 | 149 | 123 | 6:9:0:2:0 | 6 | 9 | 0 | 2 | 0 | 5 |
| 504 | 149 | 123 | 6:9:1:2:0 | 6 | 9 | 1 | 2 | 0 | 4 |
| 505 | 149 | 123 | 7:5:4:0:1 | 7 | 5 | 4 | 0 | 1 | 1 |
| 506 | 149 | 123 | 7:6:3:0:1 | 7 | 6 | 3 | 0 | 1 | 2 |
| 507 | 149 | 123 | 7:6:3:1:0 | 7 | 6 | 3 | 1 | 0 | 1 |
| 508 | 149 | 123 | 7:6:4:0:0 | 7 | 6 | 4 | 0 | 0 | 2 |
| 509 | 149 | 123 | 7:6:4:0:1 | 7 | 6 | 4 | 0 | 1 | 2 |
| 510 | 149 | 123 | 7:7:1:0:1 | 7 | 7 | 1 | 0 | 1 | 3 |
| 511 | 149 | 123 | 7:7:2:0:0 | 7 | 7 | 2 | 0 | 0 | 6 |
| 512 | 149 | 123 | 7:7:2:1:0 | 7 | 7 | 2 | 1 | 0 | 2 |
| 513 | 149 | 123 | 7:7:3:0:0 | 7 | 7 | 3 | 0 | 0 | 8 |
| 514 | 149 | 123 | 7:7:3:0:1 | 7 | 7 | 3 | 0 | 1 | 1 |
| 515 | 149 | 123 | 7:7:3:1:0 | 7 | 7 | 3 | 1 | 0 | 2 |
| 516 | 149 | 123 | 7:7:4:0:0 | 7 | 7 | 4 | 0 | 0 | 4 |
| 517 | 149 | 123 | 7:8:0:1:0 | 7 | 8 | 0 | 1 | 0 | 3 |
| 518 | 149 | 123 | 7:8:2:1:0 | 7 | 8 | 2 | 1 | 0 | 1 |
| 519 | 149 | 123 | 8:8:0:0:1 | 8 | 8 | 0 | 0 | 1 | 1 |
| 520 | 149 | 123 | 8:9:0:0:0 | 8 | 9 | 0 | 0 | 0 | 5 |
| 521 | 149 | 123 | 9:8:0:0:0 | 9 | 8 | 0 | 0 | 0 | 2 |
| 522 | 149 | 123 | 9:9:0:0:0 | 9 | 9 | 0 | 0 | 0 | 3 |
| 523 | 165 | 39 | 2:5:0:0:0 | 2 | 5 | 0 | 0 | 0 | 13 |
| 524 | 165 | 39 | 2:6:0:0:0 | 2 | 6 | 0 | 0 | 0 | 13 |
| 525 | 165 | 39 | 3:4:1:0:0 | 3 | 4 | 1 | 0 | 0 | 8 |
| 526 | 165 | 39 | 3:5:1:0:0 | 3 | 5 | 1 | 0 | 0 | 14 |

| | | | | | | | | | |
|---|---|---|---|---|---|---|---|---|---|
| 527 | 165 | 39 | 3:6:0:0:0 | 3 | 6 | 0 | 0 | 0 | 11 |
| 528 | 165 | 39 | 3:6:0:0:1 | 3 | 6 | 0 | 0 | 1 | 8 |
| 529 | 165 | 39 | 3:6:1:0:0 | 3 | 6 | 1 | 0 | 0 | 13 |
| 530 | 165 | 39 | 4:5:0:0:0 | 4 | 5 | 0 | 0 | 0 | 14 |
| 531 | 165 | 39 | 4:5:0:0:1 | 4 | 5 | 0 | 0 | 1 | 7 |
| 532 | 165 | 39 | 4:5:1:0:0 | 4 | 5 | 1 | 0 | 0 | 19 |
| 533 | 165 | 39 | 4:5:1:0:1 | 4 | 5 | 1 | 0 | 1 | 14 |
| 534 | 165 | 39 | 4:5:2:0:0 | 4 | 5 | 2 | 0 | 0 | 18 |
| 535 | 165 | 39 | 4:5:3:0:0 | 4 | 5 | 3 | 0 | 0 | 17 |
| 536 | 165 | 39 | 4:6:0:1:0 | 4 | 6 | 0 | 1 | 0 | 14 |
| 537 | 165 | 39 | 5:4:1:0:0 | 5 | 4 | 1 | 0 | 0 | 16 |
| 538 | 165 | 39 | 5:4:3:0:0 | 5 | 4 | 3 | 0 | 0 | 7 |
| 539 | 165 | 39 | 5:5:0:0:0 | 5 | 5 | 0 | 0 | 0 | 6 |
| 540 | 165 | 39 | 5:5:1:0:0 | 5 | 5 | 1 | 0 | 0 | 18 |
| 541 | 165 | 39 | 5:5:2:0:0 | 5 | 5 | 2 | 0 | 0 | 15 |
| 542 | 165 | 39 | 5:5:3:0:0 | 5 | 5 | 3 | 0 | 0 | 12 |
| 543 | 165 | 39 | 5:6:2:0:0 | 5 | 6 | 2 | 0 | 0 | 16 |
| 544 | 165 | 39 | 5:6:3:0:0 | 5 | 6 | 3 | 0 | 0 | 17 |
| 545 | 165 | 39 | 5:6:4:0:0 | 5 | 6 | 4 | 0 | 0 | 14 |
| 546 | 165 | 39 | 6:6:1:0:0 | 6 | 6 | 1 | 0 | 0 | 19 |
| 547 | 165 | 39 | 6:6:3:0:0 | 6 | 6 | 3 | 0 | 0 | 14 |
| 548 | 165 | 39 | 6:6:4:0:0 | 6 | 6 | 4 | 0 | 0 | 10 |
| 549 | 165 | 39 | 6:7:1:0:0 | 6 | 7 | 1 | 0 | 0 | 17 |
| 550 | 165 | 39 | 6:7:2:0:0 | 6 | 7 | 2 | 0 | 0 | 16 |
| 551 | 165 | 39 | 6:7:3:0:0 | 6 | 7 | 3 | 0 | 0 | 16 |
| 552 | 165 | 39 | 6:7:3:0:1 | 6 | 7 | 3 | 0 | 1 | 8 |
| 553 | 165 | 39 | 6:7:4:0:0 | 6 | 7 | 4 | 0 | 0 | 16 |
| 554 | 165 | 39 | 6:8:2:1:0 | 6 | 8 | 2 | 1 | 0 | 8 |
| 555 | 165 | 39 | 7:7:1:0:0 | 7 | 7 | 1 | 0 | 0 | 6 |
| 556 | 165 | 39 | 7:7:2:0:0 | 7 | 7 | 2 | 0 | 0 | 6 |
| 557 | 165 | 39 | 7:7:3:0:0 | 7 | 7 | 3 | 0 | 0 | 8 |
| 558 | 165 | 39 | 7:7:4:0:0 | 7 | 7 | 4 | 0 | 0 | 4 |
| 559 | 165 | 39 | 8:9:0:0:0 | 8 | 9 | 0 | 0 | 0 | 5 |
| 560 | 165 | 39 | 9:9:0:0:0 | 9 | 9 | 0 | 0 | 0 | 3 |
| 561 | 234 | 28 | 2:5:0:0:0 | 2 | 5 | 0 | 0 | 0 | 13 |
| 562 | 234 | 28 | 2:6:0:0:0 | 2 | 6 | 0 | 0 | 0 | 13 |
| 563 | 234 | 28 | 2:7:0:0:0 | 2 | 7 | 0 | 0 | 0 | 11 |
| 564 | 234 | 28 | 2:8:0:0:0 | 2 | 8 | 0 | 0 | 0 | 10 |
| 565 | 234 | 28 | 2:9:0:0:0 | 2 | 9 | 0 | 0 | 0 | 5 |
| 566 | 234 | 28 | 3:5:0:0:0 | 3 | 5 | 0 | 0 | 0 | 11 |
| 567 | 234 | 28 | 3:5:1:0:0 | 3 | 5 | 1 | 0 | 0 | 14 |
| 568 | 234 | 28 | 3:6:0:0:0 | 3 | 6 | 0 | 0 | 0 | 11 |
| 569 | 234 | 28 | 3:6:0:0:1 | 3 | 6 | 0 | 0 | 1 | 8 |
| 570 | 234 | 28 | 4:3:1:0:0 | 4 | 3 | 1 | 0 | 0 | 14 |
| 571 | 234 | 28 | 4:4:0:0:0 | 4 | 4 | 0 | 0 | 0 | 6 |
| 572 | 234 | 28 | 4:4:1:0:0 | 4 | 4 | 1 | 0 | 0 | 12 |
| 573 | 234 | 28 | 4:5:0:0:0 | 4 | 5 | 0 | 0 | 0 | 14 |
| 574 | 234 | 28 | 4:5:0:0:1 | 4 | 5 | 0 | 0 | 1 | 7 |
| 575 | 234 | 28 | 4:5:1:0:0 | 4 | 5 | 1 | 0 | 0 | 19 |
| 576 | 234 | 28 | 4:5:2:0:0 | 4 | 5 | 2 | 0 | 0 | 18 |
| 577 | 234 | 28 | 5:4:1:0:0 | 5 | 4 | 1 | 0 | 0 | 16 |
| 578 | 234 | 28 | 5:5:1:0:0 | 5 | 5 | 1 | 0 | 0 | 18 |
| 579 | 234 | 28 | 5:6:0:0:0 | 5 | 6 | 0 | 0 | 0 | 9 |
| 580 | 234 | 28 | 5:6:1:0:0 | 5 | 6 | 1 | 0 | 0 | 16 |
| 581 | 234 | 28 | 6:4:1:0:0 | 6 | 4 | 1 | 0 | 0 | 8 |
| 582 | 234 | 28 | 6:5:0:0:0 | 6 | 5 | 0 | 0 | 0 | 3 |
| 583 | 234 | 28 | 6:5:1:0:0 | 6 | 5 | 1 | 0 | 0 | 10 |
| 584 | 234 | 28 | 6:6:0:0:0 | 6 | 6 | 0 | 0 | 0 | 6 |
| 585 | 234 | 28 | 6:6:1:0:0 | 6 | 6 | 1 | 0 | 0 | 19 |

| | | | | | | | | | |
|---|---|---|---|---|---|---|---|---|---|
| 586 | 234 | 28 | 6:7:0:0:0 | 6 | 7 | 0 | 0 | 0 | 6 |
| 587 | 234 | 28 | 6:7:1:0:0 | 6 | 7 | 1 | 0 | 0 | 17 |
| 588 | 282 | 53 | 2:5:0:0:0 | 2 | 5 | 0 | 0 | 0 | 13 |
| 589 | 282 | 53 | 2:6:0:0:0 | 2 | 6 | 0 | 0 | 0 | 13 |
| 590 | 282 | 53 | 3:3:1:0:1 | 3 | 3 | 1 | 0 | 1 | 2 |
| 591 | 282 | 53 | 3:3:2:0:1 | 3 | 3 | 2 | 0 | 1 | 2 |
| 592 | 282 | 53 | 3:4:0:1:0 | 3 | 4 | 0 | 1 | 0 | 2 |
| 593 | 282 | 53 | 3:4:1:0:0 | 3 | 4 | 1 | 0 | 0 | 8 |
| 594 | 282 | 53 | 3:4:1:1:0 | 3 | 4 | 1 | 1 | 0 | 2 |
| 595 | 282 | 53 | 3:5:0:0:0 | 3 | 5 | 0 | 0 | 0 | 11 |
| 596 | 282 | 53 | 3:5:1:0:0 | 3 | 5 | 1 | 0 | 0 | 14 |
| 597 | 282 | 53 | 3:6:1:0:0 | 3 | 6 | 1 | 0 | 0 | 13 |
| 598 | 282 | 53 | 4:3:0:0:0 | 4 | 3 | 0 | 0 | 0 | 6 |
| 599 | 282 | 53 | 4:3:1:0:0 | 4 | 3 | 1 | 0 | 0 | 14 |
| 600 | 282 | 53 | 4:4:1:0:0 | 4 | 4 | 1 | 0 | 0 | 12 |
| 601 | 282 | 53 | 4:4:1:0:1 | 4 | 4 | 1 | 0 | 1 | 7 |
| 602 | 282 | 53 | 4:4:2:0:1 | 4 | 4 | 2 | 0 | 1 | 8 |
| 603 | 282 | 53 | 4:4:3:0:1 | 4 | 4 | 3 | 0 | 1 | 4 |
| 604 | 282 | 53 | 4:5:0:0:0 | 4 | 5 | 0 | 0 | 0 | 14 |
| 605 | 282 | 53 | 4:5:0:1:0 | 4 | 5 | 0 | 1 | 0 | 7 |
| 606 | 282 | 53 | 4:5:1:0:0 | 4 | 5 | 1 | 0 | 0 | 19 |
| 607 | 282 | 53 | 4:5:1:0:1 | 4 | 5 | 1 | 0 | 1 | 14 |
| 608 | 282 | 53 | 4:5:1:1:0 | 4 | 5 | 1 | 1 | 0 | 8 |
| 609 | 282 | 53 | 4:5:2:0:0 | 4 | 5 | 2 | 0 | 0 | 18 |
| 610 | 282 | 53 | 4:5:2:1:0 | 4 | 5 | 2 | 1 | 0 | 4 |
| 611 | 282 | 53 | 4:5:3:0:0 | 4 | 5 | 3 | 0 | 0 | 17 |
| 612 | 282 | 53 | 4:6:0:1:0 | 4 | 6 | 0 | 1 | 0 | 14 |
| 613 | 282 | 53 | 5:3:0:0:0 | 5 | 3 | 0 | 0 | 0 | 6 |
| 614 | 282 | 53 | 5:4:1:0:0 | 5 | 4 | 1 | 0 | 0 | 16 |
| 615 | 282 | 53 | 5:4:2:0:0 | 5 | 4 | 2 | 0 | 0 | 8 |
| 616 | 282 | 53 | 5:5:1:0:0 | 5 | 5 | 1 | 0 | 0 | 18 |
| 617 | 282 | 53 | 5:5:2:0:0 | 5 | 5 | 2 | 0 | 0 | 15 |
| 618 | 282 | 53 | 5:5:3:0:0 | 5 | 5 | 3 | 0 | 0 | 12 |
| 619 | 282 | 53 | 5:6:0:0:0 | 5 | 6 | 0 | 0 | 0 | 9 |
| 620 | 282 | 53 | 5:6:1:0:0 | 5 | 6 | 1 | 0 | 0 | 16 |
| 621 | 282 | 53 | 5:6:2:0:0 | 5 | 6 | 2 | 0 | 0 | 16 |
| 622 | 282 | 53 | 5:6:3:0:0 | 5 | 6 | 3 | 0 | 0 | 17 |
| 623 | 282 | 53 | 6:3:1:0:0 | 6 | 3 | 1 | 0 | 0 | 9 |
| 624 | 282 | 53 | 6:4:1:0:0 | 6 | 4 | 1 | 0 | 0 | 8 |
| 625 | 282 | 53 | 6:5:1:0:0 | 6 | 5 | 1 | 0 | 0 | 10 |
| 626 | 282 | 53 | 6:5:1:0:1 | 6 | 5 | 1 | 0 | 1 | 2 |
| 627 | 282 | 53 | 6:6:0:1:0 | 6 | 6 | 0 | 1 | 0 | 2 |
| 628 | 282 | 53 | 6:6:1:0:0 | 6 | 6 | 1 | 0 | 0 | 19 |
| 629 | 282 | 53 | 6:6:2:0:0 | 6 | 6 | 2 | 0 | 0 | 12 |
| 630 | 282 | 53 | 6:6:3:0:0 | 6 | 6 | 3 | 0 | 0 | 14 |
| 631 | 282 | 53 | 6:7:0:0:0 | 6 | 7 | 0 | 0 | 0 | 6 |
| 632 | 282 | 53 | 6:7:1:0:0 | 6 | 7 | 1 | 0 | 0 | 17 |
| 633 | 282 | 53 | 6:7:1:0:1 | 6 | 7 | 1 | 0 | 1 | 13 |
| 634 | 282 | 53 | 6:7:2:0:0 | 6 | 7 | 2 | 0 | 0 | 16 |
| 635 | 282 | 53 | 6:7:3:0:0 | 6 | 7 | 3 | 0 | 0 | 16 |
| 636 | 282 | 53 | 6:7:4:0:0 | 6 | 7 | 4 | 0 | 0 | 16 |
| 637 | 282 | 53 | 6:8:0:1:0 | 6 | 8 | 0 | 1 | 0 | 13 |
| 638 | 282 | 53 | 7:7:1:0:0 | 7 | 7 | 1 | 0 | 0 | 6 |
| 639 | 282 | 53 | 7:7:2:0:0 | 7 | 7 | 2 | 0 | 0 | 6 |
| 640 | 282 | 53 | 7:7:3:0:0 | 7 | 7 | 3 | 0 | 0 | 8 |
| 641 | 331 | 49 | 3:5:2:0:0 | 3 | 5 | 2 | 0 | 0 | 13 |
| 642 | 331 | 49 | 4:3:1:0:0 | 4 | 3 | 1 | 0 | 0 | 14 |
| 643 | 331 | 49 | 4:4:2:0:0 | 4 | 4 | 2 | 0 | 0 | 8 |
| 644 | 331 | 49 | 4:4:2:0:1 | 4 | 4 | 2 | 0 | 1 | 8 |

| | | | | | | | | | |
|---|---|---|---|---|---|---|---|---|---|
| 645 | 331 | 49 | 4:5:1:0:0 | 4 | 5 | 1 | 0 | 0 | 19 |
| 646 | 331 | 49 | 4:5:1:0:1 | 4 | 5 | 1 | 0 | 1 | 14 |
| 647 | 331 | 49 | 4:5:1:1:0 | 4 | 5 | 1 | 1 | 0 | 8 |
| 648 | 331 | 49 | 4:5:2:0:0 | 4 | 5 | 2 | 0 | 0 | 18 |
| 649 | 331 | 49 | 4:5:2:0:1 | 4 | 5 | 2 | 0 | 1 | 9 |
| 650 | 331 | 49 | 4:5:3:0:0 | 4 | 5 | 3 | 0 | 0 | 17 |
| 651 | 331 | 49 | 4:6:0:1:0 | 4 | 6 | 0 | 1 | 0 | 14 |
| 652 | 331 | 49 | 4:6:1:1:0 | 4 | 6 | 1 | 1 | 0 | 9 |
| 653 | 331 | 49 | 5:3:1:0:0 | 5 | 3 | 1 | 0 | 0 | 11 |
| 654 | 331 | 49 | 5:4:1:0:0 | 5 | 4 | 1 | 0 | 0 | 16 |
| 655 | 331 | 49 | 5:4:3:0:0 | 5 | 4 | 3 | 0 | 0 | 7 |
| 656 | 331 | 49 | 5:5:1:0:0 | 5 | 5 | 1 | 0 | 0 | 18 |
| 657 | 331 | 49 | 5:5:1:0:1 | 5 | 5 | 1 | 0 | 1 | 7 |
| 658 | 331 | 49 | 5:5:2:0:0 | 5 | 5 | 2 | 0 | 0 | 15 |
| 659 | 331 | 49 | 5:5:2:0:1 | 5 | 5 | 2 | 0 | 1 | 7 |
| 660 | 331 | 49 | 5:5:3:0:0 | 5 | 5 | 3 | 0 | 0 | 12 |
| 661 | 331 | 49 | 5:6:0:1:0 | 5 | 6 | 0 | 1 | 0 | 7 |
| 662 | 331 | 49 | 5:6:1:0:0 | 5 | 6 | 1 | 0 | 0 | 16 |
| 663 | 331 | 49 | 5:6:1:0:1 | 5 | 6 | 1 | 0 | 1 | 8 |
| 664 | 331 | 49 | 5:6:1:0:2 | 5 | 6 | 1 | 0 | 2 | 4 |
| 665 | 331 | 49 | 5:6:1:1:0 | 5 | 6 | 1 | 1 | 0 | 7 |
| 666 | 331 | 49 | 5:6:2:0:0 | 5 | 6 | 2 | 0 | 0 | 16 |
| 667 | 331 | 49 | 5:6:3:0:0 | 5 | 6 | 3 | 0 | 0 | 17 |
| 668 | 331 | 49 | 5:6:3:0:1 | 5 | 6 | 3 | 0 | 1 | 7 |
| 669 | 331 | 49 | 5:6:4:0:0 | 5 | 6 | 4 | 0 | 0 | 14 |
| 670 | 331 | 49 | 5:7:0:1:0 | 5 | 7 | 0 | 1 | 0 | 8 |
| 671 | 331 | 49 | 5:7:0:1:1 | 5 | 7 | 0 | 1 | 1 | 4 |
| 672 | 331 | 49 | 5:7:2:1:0 | 5 | 7 | 2 | 1 | 0 | 7 |
| 673 | 331 | 49 | 6:6:1:0:0 | 6 | 6 | 1 | 0 | 0 | 19 |
| 674 | 331 | 49 | 6:6:1:0:1 | 6 | 6 | 1 | 0 | 1 | 7 |
| 675 | 331 | 49 | 6:6:2:0:0 | 6 | 6 | 2 | 0 | 0 | 12 |
| 676 | 331 | 49 | 6:6:3:0:0 | 6 | 6 | 3 | 0 | 0 | 14 |
| 677 | 331 | 49 | 6:6:4:0:0 | 6 | 6 | 4 | 0 | 0 | 10 |
| 678 | 331 | 49 | 6:7:0:1:0 | 6 | 7 | 0 | 1 | 0 | 7 |
| 679 | 331 | 49 | 6:7:1:0:0 | 6 | 7 | 1 | 0 | 0 | 17 |
| 680 | 331 | 49 | 6:7:1:0:1 | 6 | 7 | 1 | 0 | 1 | 13 |
| 681 | 331 | 49 | 6:7:2:0:0 | 6 | 7 | 2 | 0 | 0 | 16 |
| 682 | 331 | 49 | 6:7:2:0:1 | 6 | 7 | 2 | 0 | 1 | 9 |
| 683 | 331 | 49 | 6:7:3:0:0 | 6 | 7 | 3 | 0 | 0 | 16 |
| 684 | 331 | 49 | 6:7:4:0:0 | 6 | 7 | 4 | 0 | 0 | 16 |
| 685 | 331 | 49 | 6:7:4:0:1 | 6 | 7 | 4 | 0 | 1 | 5 |
| 686 | 331 | 49 | 6:8:0:1:0 | 6 | 8 | 0 | 1 | 0 | 13 |
| 687 | 331 | 49 | 6:8:1:1:0 | 6 | 8 | 1 | 1 | 0 | 9 |
| 688 | 331 | 49 | 6:8:3:1:0 | 6 | 8 | 3 | 1 | 0 | 5 |
| 689 | 331 | 49 | 8:9:0:0:0 | 8 | 9 | 0 | 0 | 0 | 5 |
| 690 | 343 | 167 | 10:10:0:0:0 | 10 | 10 | 0 | 0 | 0 | 1 |
| 691 | 343 | 167 | 10:10:1:0:0 | 10 | 10 | 1 | 0 | 0 | 2 |
| 692 | 343 | 167 | 10:6:3:0:2 | 10 | 6 | 3 | 0 | 2 | 1 |
| 693 | 343 | 167 | 10:7:2:1:1 | 10 | 7 | 2 | 1 | 1 | 1 |
| 694 | 343 | 167 | 10:8:1:2:0 | 10 | 8 | 1 | 2 | 0 | 1 |
| 695 | 343 | 167 | 10:9:0:0:0 | 10 | 9 | 0 | 0 | 0 | 2 |
| 696 | 343 | 167 | 10:9:1:0:0 | 10 | 9 | 1 | 0 | 0 | 2 |
| 697 | 343 | 167 | 3:4:2:0:0 | 3 | 4 | 2 | 0 | 0 | 5 |
| 698 | 343 | 167 | 3:5:2:0:0 | 3 | 5 | 2 | 0 | 0 | 13 |
| 699 | 343 | 167 | 3:6:1:0:0 | 3 | 6 | 1 | 0 | 0 | 13 |
| 700 | 343 | 167 | 3:6:2:0:0 | 3 | 6 | 2 | 0 | 0 | 6 |
| 701 | 343 | 167 | 4:3:1:0:0 | 4 | 3 | 1 | 0 | 0 | 14 |
| 702 | 343 | 167 | 4:4:1:0:0 | 4 | 4 | 1 | 0 | 0 | 12 |
| 703 | 343 | 167 | 4:4:2:0:0 | 4 | 4 | 2 | 0 | 0 | 8 |

| | | | | | | | | | |
|---|---|---|---|---|---|---|---|---|---|
| 704 | 343 | 167 | 4:4:2:0:1 | 4 | 4 | 2 | 0 | 1 | 8 |
| 705 | 343 | 167 | 4:4:3:0:1 | 4 | 4 | 3 | 0 | 1 | 4 |
| 706 | 343 | 167 | 4:5:1:0:0 | 4 | 5 | 1 | 0 | 0 | 19 |
| 707 | 343 | 167 | 4:5:1:0:1 | 4 | 5 | 1 | 0 | 1 | 14 |
| 708 | 343 | 167 | 4:5:1:0:2 | 4 | 5 | 1 | 0 | 2 | 3 |
| 709 | 343 | 167 | 4:5:1:1:0 | 4 | 5 | 1 | 1 | 0 | 8 |
| 710 | 343 | 167 | 4:5:2:0:0 | 4 | 5 | 2 | 0 | 0 | 18 |
| 711 | 343 | 167 | 4:5:2:0:1 | 4 | 5 | 2 | 0 | 1 | 9 |
| 712 | 343 | 167 | 4:5:2:1:0 | 4 | 5 | 2 | 1 | 0 | 4 |
| 713 | 343 | 167 | 4:5:3:0:0 | 4 | 5 | 3 | 0 | 0 | 17 |
| 714 | 343 | 167 | 4:6:0:1:0 | 4 | 6 | 0 | 1 | 0 | 14 |
| 715 | 343 | 167 | 4:6:0:1:1 | 4 | 6 | 0 | 1 | 1 | 3 |
| 716 | 343 | 167 | 4:6:1:1:0 | 4 | 6 | 1 | 1 | 0 | 9 |
| 717 | 343 | 167 | 5:3:1:0:0 | 5 | 3 | 1 | 0 | 0 | 11 |
| 718 | 343 | 167 | 5:4:1:0:0 | 5 | 4 | 1 | 0 | 0 | 16 |
| 719 | 343 | 167 | 5:4:2:0:0 | 5 | 4 | 2 | 0 | 0 | 8 |
| 720 | 343 | 167 | 5:4:2:0:2 | 5 | 4 | 2 | 0 | 2 | 1 |
| 721 | 343 | 167 | 5:4:3:0:0 | 5 | 4 | 3 | 0 | 0 | 7 |
| 722 | 343 | 167 | 5:4:3:0:1 | 5 | 4 | 3 | 0 | 1 | 1 |
| 723 | 343 | 167 | 5:5:1:0:0 | 5 | 5 | 1 | 0 | 0 | 18 |
| 724 | 343 | 167 | 5:5:1:0:1 | 5 | 5 | 1 | 0 | 1 | 7 |
| 725 | 343 | 167 | 5:5:1:1:1 | 5 | 5 | 1 | 1 | 1 | 1 |
| 726 | 343 | 167 | 5:5:2:0:0 | 5 | 5 | 2 | 0 | 0 | 15 |
| 727 | 343 | 167 | 5:5:2:0:1 | 5 | 5 | 2 | 0 | 1 | 7 |
| 728 | 343 | 167 | 5:5:2:0:2 | 5 | 5 | 2 | 0 | 2 | 1 |
| 729 | 343 | 167 | 5:5:2:1:0 | 5 | 5 | 2 | 1 | 0 | 1 |
| 730 | 343 | 167 | 5:5:3:0:0 | 5 | 5 | 3 | 0 | 0 | 12 |
| 731 | 343 | 167 | 5:5:4:0:0 | 5 | 5 | 4 | 0 | 0 | 2 |
| 732 | 343 | 167 | 5:6:0:1:0 | 5 | 6 | 0 | 1 | 0 | 7 |
| 733 | 343 | 167 | 5:6:0:2:0 | 5 | 6 | 0 | 2 | 0 | 1 |
| 734 | 343 | 167 | 5:6:1:0:0 | 5 | 6 | 1 | 0 | 0 | 16 |
| 735 | 343 | 167 | 5:6:1:0:1 | 5 | 6 | 1 | 0 | 1 | 8 |
| 736 | 343 | 167 | 5:6:1:1:0 | 5 | 6 | 1 | 1 | 0 | 7 |
| 737 | 343 | 167 | 5:6:1:1:1 | 5 | 6 | 1 | 1 | 1 | 1 |
| 738 | 343 | 167 | 5:6:2:0:0 | 5 | 6 | 2 | 0 | 0 | 16 |
| 739 | 343 | 167 | 5:6:2:0:1 | 5 | 6 | 2 | 0 | 1 | 9 |
| 740 | 343 | 167 | 5:6:2:0:2 | 5 | 6 | 2 | 0 | 2 | 2 |
| 741 | 343 | 167 | 5:6:3:0:0 | 5 | 6 | 3 | 0 | 0 | 17 |
| 742 | 343 | 167 | 5:6:3:0:1 | 5 | 6 | 3 | 0 | 1 | 7 |
| 743 | 343 | 167 | 5:6:4:0:0 | 5 | 6 | 4 | 0 | 0 | 14 |
| 744 | 343 | 167 | 5:7:0:1:0 | 5 | 7 | 0 | 1 | 0 | 8 |
| 745 | 343 | 167 | 5:7:0:2:0 | 5 | 7 | 0 | 2 | 0 | 1 |
| 746 | 343 | 167 | 5:7:1:1:0 | 5 | 7 | 1 | 1 | 0 | 9 |
| 747 | 343 | 167 | 5:7:1:1:1 | 5 | 7 | 1 | 1 | 1 | 2 |
| 748 | 343 | 167 | 5:7:2:0:2 | 5 | 7 | 2 | 0 | 2 | 1 |
| 749 | 343 | 167 | 5:7:2:1:0 | 5 | 7 | 2 | 1 | 0 | 7 |
| 750 | 343 | 167 | 5:8:0:2:0 | 5 | 8 | 0 | 2 | 0 | 2 |
| 751 | 343 | 167 | 5:8:1:1:1 | 5 | 8 | 1 | 1 | 1 | 1 |
| 752 | 343 | 167 | 5:8:2:0:2 | 5 | 8 | 2 | 0 | 2 | 1 |
| 753 | 343 | 167 | 5:9:0:2:0 | 5 | 9 | 0 | 2 | 0 | 1 |
| 754 | 343 | 167 | 5:9:1:1:1 | 5 | 9 | 1 | 1 | 1 | 1 |
| 755 | 343 | 167 | 5:9:2:0:2 | 5 | 9 | 2 | 0 | 2 | 1 |
| 756 | 343 | 167 | 6:10:0:2:0 | 6 | 10 | 0 | 2 | 0 | 1 |
| 757 | 343 | 167 | 6:10:1:2:0 | 6 | 10 | 1 | 2 | 0 | 1 |
| 758 | 343 | 167 | 6:5:1:0:0 | 6 | 5 | 1 | 0 | 0 | 10 |
| 759 | 343 | 167 | 6:5:2:0:0 | 6 | 5 | 2 | 0 | 0 | 4 |
| 760 | 343 | 167 | 6:5:2:0:1 | 6 | 5 | 2 | 0 | 1 | 4 |
| 761 | 343 | 167 | 6:5:2:0:2 | 6 | 5 | 2 | 0 | 2 | 3 |
| 762 | 343 | 167 | 6:5:3:0:2 | 6 | 5 | 3 | 0 | 2 | 2 |

| | | | | | | | | | |
|---|---|---|---|---|---|---|---|---|---|
| 763 | 343 | 167 | 6:6:1:0:0 | 6 | 6 | 1 | 0 | 0 | 19 |
| 764 | 343 | 167 | 6:6:1:0:1 | 6 | 6 | 1 | 0 | 1 | 7 |
| 765 | 343 | 167 | 6:6:1:1:0 | 6 | 6 | 1 | 1 | 0 | 4 |
| 766 | 343 | 167 | 6:6:1:1:1 | 6 | 6 | 1 | 1 | 1 | 3 |
| 767 | 343 | 167 | 6:6:2:0:0 | 6 | 6 | 2 | 0 | 0 | 12 |
| 768 | 343 | 167 | 6:6:2:0:2 | 6 | 6 | 2 | 0 | 2 | 4 |
| 769 | 343 | 167 | 6:6:2:1:1 | 6 | 6 | 2 | 1 | 1 | 2 |
| 770 | 343 | 167 | 6:6:3:0:0 | 6 | 6 | 3 | 0 | 0 | 14 |
| 771 | 343 | 167 | 6:6:3:0:1 | 6 | 6 | 3 | 0 | 1 | 7 |
| 772 | 343 | 167 | 6:6:3:0:2 | 6 | 6 | 3 | 0 | 2 | 3 |
| 773 | 343 | 167 | 6:6:4:0:0 | 6 | 6 | 4 | 0 | 0 | 10 |
| 774 | 343 | 167 | 6:7:0:1:0 | 6 | 7 | 0 | 1 | 0 | 7 |
| 775 | 343 | 167 | 6:7:0:2:0 | 6 | 7 | 0 | 2 | 0 | 3 |
| 776 | 343 | 167 | 6:7:1:0:0 | 6 | 7 | 1 | 0 | 0 | 17 |
| 777 | 343 | 167 | 6:7:1:0:1 | 6 | 7 | 1 | 0 | 1 | 13 |
| 778 | 343 | 167 | 6:7:1:1:1 | 6 | 7 | 1 | 1 | 1 | 4 |
| 779 | 343 | 167 | 6:7:1:2:0 | 6 | 7 | 1 | 2 | 0 | 2 |
| 780 | 343 | 167 | 6:7:2:0:0 | 6 | 7 | 2 | 0 | 0 | 16 |
| 781 | 343 | 167 | 6:7:2:0:2 | 6 | 7 | 2 | 0 | 2 | 5 |
| 782 | 343 | 167 | 6:7:2:1:0 | 6 | 7 | 2 | 1 | 0 | 7 |
| 783 | 343 | 167 | 6:7:2:1:1 | 6 | 7 | 2 | 1 | 1 | 3 |
| 784 | 343 | 167 | 6:7:3:0:0 | 6 | 7 | 3 | 0 | 0 | 16 |
| 785 | 343 | 167 | 6:7:4:0:0 | 6 | 7 | 4 | 0 | 0 | 16 |
| 786 | 343 | 167 | 6:7:4:0:1 | 6 | 7 | 4 | 0 | 1 | 5 |
| 787 | 343 | 167 | 6:8:0:1:0 | 6 | 8 | 0 | 1 | 0 | 13 |
| 788 | 343 | 167 | 6:8:0:2:0 | 6 | 8 | 0 | 2 | 0 | 4 |
| 789 | 343 | 167 | 6:8:1:1:1 | 6 | 8 | 1 | 1 | 1 | 5 |
| 790 | 343 | 167 | 6:8:1:2:0 | 6 | 8 | 1 | 2 | 0 | 3 |
| 791 | 343 | 167 | 6:8:2:0:2 | 6 | 8 | 2 | 0 | 2 | 1 |
| 792 | 343 | 167 | 6:8:3:0:2 | 6 | 8 | 3 | 0 | 2 | 1 |
| 793 | 343 | 167 | 6:8:3:1:0 | 6 | 8 | 3 | 1 | 0 | 5 |
| 794 | 343 | 167 | 6:9:0:2:0 | 6 | 9 | 0 | 2 | 0 | 5 |
| 795 | 343 | 167 | 6:9:1:1:1 | 6 | 9 | 1 | 1 | 1 | 1 |
| 796 | 343 | 167 | 6:9:2:1:1 | 6 | 9 | 2 | 1 | 1 | 1 |
| 797 | 343 | 167 | 7:5:2:0:2 | 7 | 5 | 2 | 0 | 2 | 1 |
| 798 | 343 | 167 | 7:5:3:0:2 | 7 | 5 | 3 | 0 | 2 | 1 |
| 799 | 343 | 167 | 7:5:4:0:2 | 7 | 5 | 4 | 0 | 2 | 1 |
| 800 | 343 | 167 | 7:6:1:1:1 | 7 | 6 | 1 | 1 | 1 | 1 |
| 801 | 343 | 167 | 7:6:2:1:1 | 7 | 6 | 2 | 1 | 1 | 1 |
| 802 | 343 | 167 | 7:6:3:0:2 | 7 | 6 | 3 | 0 | 2 | 2 |
| 803 | 343 | 167 | 7:6:3:1:1 | 7 | 6 | 3 | 1 | 1 | 1 |
| 804 | 343 | 167 | 7:6:4:0:2 | 7 | 6 | 4 | 0 | 2 | 1 |
| 805 | 343 | 167 | 7:7:0:2:0 | 7 | 7 | 0 | 2 | 0 | 1 |
| 806 | 343 | 167 | 7:7:1:2:0 | 7 | 7 | 1 | 2 | 0 | 1 |
| 807 | 343 | 167 | 7:7:2:1:1 | 7 | 7 | 2 | 1 | 1 | 2 |
| 808 | 343 | 167 | 7:7:2:2:0 | 7 | 7 | 2 | 2 | 0 | 1 |
| 809 | 343 | 167 | 7:7:3:0:0 | 7 | 7 | 3 | 0 | 0 | 8 |
| 810 | 343 | 167 | 7:7:3:1:1 | 7 | 7 | 3 | 1 | 1 | 1 |
| 811 | 343 | 167 | 7:7:4:0:0 | 7 | 7 | 4 | 0 | 0 | 4 |
| 812 | 343 | 167 | 7:8:1:2:0 | 7 | 8 | 1 | 2 | 0 | 2 |
| 813 | 343 | 167 | 7:8:2:2:0 | 7 | 8 | 2 | 2 | 0 | 1 |
| 814 | 343 | 167 | 8:4:2:0:2 | 8 | 4 | 2 | 0 | 2 | 1 |
| 815 | 343 | 167 | 8:4:3:0:2 | 8 | 4 | 3 | 0 | 2 | 1 |
| 816 | 343 | 167 | 8:5:1:1:1 | 8 | 5 | 1 | 1 | 1 | 1 |
| 817 | 343 | 167 | 8:5:2:0:2 | 8 | 5 | 2 | 0 | 2 | 1 |
| 818 | 343 | 167 | 8:5:2:1:1 | 8 | 5 | 2 | 1 | 1 | 1 |
| 819 | 343 | 167 | 8:5:3:0:2 | 8 | 5 | 3 | 0 | 2 | 1 |
| 820 | 343 | 167 | 8:6:0:2:0 | 8 | 6 | 0 | 2 | 0 | 1 |
| 821 | 343 | 167 | 8:6:1:1:1 | 8 | 6 | 1 | 1 | 1 | 1 |

| | | | | | | | | | |
|---|---|---|---|---|---|---|---|---|---|
| 822 | 343 | 167 | 8:6:1:2:0 | 8 | 6 | 1 | 2 | 0 | 1 |
| 823 | 343 | 167 | 8:6:2:1:1 | 8 | 6 | 2 | 1 | 1 | 1 |
| 824 | 343 | 167 | 8:6:3:0:2 | 8 | 6 | 3 | 0 | 2 | 1 |
| 825 | 343 | 167 | 8:7:0:2:0 | 8 | 7 | 0 | 2 | 0 | 1 |
| 826 | 343 | 167 | 8:7:1:2:0 | 8 | 7 | 1 | 2 | 0 | 1 |
| 827 | 343 | 167 | 8:7:2:1:1 | 8 | 7 | 2 | 1 | 1 | 1 |
| 828 | 343 | 167 | 8:7:3:0:2 | 8 | 7 | 3 | 0 | 2 | 1 |
| 829 | 343 | 167 | 8:8:1:2:0 | 8 | 8 | 1 | 2 | 0 | 1 |
| 830 | 343 | 167 | 8:8:2:1:1 | 8 | 8 | 2 | 1 | 1 | 1 |
| 831 | 343 | 167 | 8:9:1:2:0 | 8 | 9 | 1 | 2 | 0 | 1 |
| 832 | 343 | 167 | 9:10:0:0:0 | 9 | 10 | 0 | 0 | 0 | 2 |
| 833 | 343 | 167 | 9:10:1:0:0 | 9 | 10 | 1 | 0 | 0 | 2 |
| 834 | 343 | 167 | 9:10:1:2:0 | 9 | 10 | 1 | 2 | 0 | 1 |
| 835 | 343 | 167 | 9:10:2:2:0 | 9 | 10 | 2 | 2 | 0 | 1 |
| 836 | 343 | 167 | 9:4:3:0:2 | 9 | 4 | 3 | 0 | 2 | 1 |
| 837 | 343 | 167 | 9:5:2:0:1 | 9 | 5 | 2 | 0 | 1 | 1 |
| 838 | 343 | 167 | 9:5:2:1:1 | 9 | 5 | 2 | 1 | 1 | 1 |
| 839 | 343 | 167 | 9:5:3:0:2 | 9 | 5 | 3 | 0 | 2 | 1 |
| 840 | 343 | 167 | 9:6:1:1:0 | 9 | 6 | 1 | 1 | 0 | 1 |
| 841 | 343 | 167 | 9:6:1:2:0 | 9 | 6 | 1 | 2 | 0 | 1 |
| 842 | 343 | 167 | 9:6:2:1:1 | 9 | 6 | 2 | 1 | 1 | 1 |
| 843 | 343 | 167 | 9:6:3:0:2 | 9 | 6 | 3 | 0 | 2 | 1 |
| 844 | 343 | 167 | 9:7:1:2:0 | 9 | 7 | 1 | 2 | 0 | 1 |
| 845 | 343 | 167 | 9:7:2:1:1 | 9 | 7 | 2 | 1 | 1 | 1 |
| 846 | 343 | 167 | 9:7:3:0:2 | 9 | 7 | 3 | 0 | 2 | 1 |
| 847 | 343 | 167 | 9:8:0:0:0 | 9 | 8 | 0 | 0 | 0 | 2 |
| 848 | 343 | 167 | 9:8:1:2:0 | 9 | 8 | 1 | 2 | 0 | 1 |
| 849 | 343 | 167 | 9:8:2:1:1 | 9 | 8 | 2 | 1 | 1 | 1 |
| 850 | 343 | 167 | 9:8:3:0:2 | 9 | 8 | 3 | 0 | 2 | 1 |
| 851 | 343 | 167 | 9:8:4:0:2 | 9 | 8 | 4 | 0 | 2 | 1 |
| 852 | 343 | 167 | 9:9:0:0:0 | 9 | 9 | 0 | 0 | 0 | 3 |
| 853 | 343 | 167 | 9:9:1:2:0 | 9 | 9 | 1 | 2 | 0 | 1 |
| 854 | 343 | 167 | 9:9:2:1:1 | 9 | 9 | 2 | 1 | 1 | 1 |
| 855 | 343 | 167 | 9:9:3:1:1 | 9 | 9 | 3 | 1 | 1 | 1 |
| 856 | 603 | 26 | 2:5:0:0:0 | 2 | 5 | 0 | 0 | 0 | 13 |
| 857 | 603 | 26 | 2:6:0:0:0 | 2 | 6 | 0 | 0 | 0 | 13 |
| 858 | 603 | 26 | 2:7:0:0:0 | 2 | 7 | 0 | 0 | 0 | 11 |
| 859 | 603 | 26 | 2:8:0:0:0 | 2 | 8 | 0 | 0 | 0 | 10 |
| 860 | 603 | 26 | 3:5:0:0:0 | 3 | 5 | 0 | 0 | 0 | 11 |
| 861 | 603 | 26 | 3:5:1:0:0 | 3 | 5 | 1 | 0 | 0 | 14 |
| 862 | 603 | 26 | 3:5:2:0:0 | 3 | 5 | 2 | 0 | 0 | 13 |
| 863 | 603 | 26 | 3:6:0:0:0 | 3 | 6 | 0 | 0 | 0 | 11 |
| 864 | 603 | 26 | 3:6:1:0:0 | 3 | 6 | 1 | 0 | 0 | 13 |
| 865 | 603 | 26 | 4:3:1:0:0 | 4 | 3 | 1 | 0 | 0 | 14 |
| 866 | 603 | 26 | 4:4:1:0:0 | 4 | 4 | 1 | 0 | 0 | 12 |
| 867 | 603 | 26 | 4:5:0:0:0 | 4 | 5 | 0 | 0 | 0 | 14 |
| 868 | 603 | 26 | 4:5:1:0:0 | 4 | 5 | 1 | 0 | 0 | 19 |
| 869 | 603 | 26 | 4:5:2:0:0 | 4 | 5 | 2 | 0 | 0 | 18 |
| 870 | 603 | 26 | 4:5:3:0:0 | 4 | 5 | 3 | 0 | 0 | 17 |
| 871 | 603 | 26 | 5:4:1:0:0 | 5 | 4 | 1 | 0 | 0 | 16 |
| 872 | 603 | 26 | 5:5:1:0:0 | 5 | 5 | 1 | 0 | 0 | 18 |
| 873 | 603 | 26 | 5:6:2:0:0 | 5 | 6 | 2 | 0 | 0 | 16 |
| 874 | 603 | 26 | 5:6:3:0:0 | 5 | 6 | 3 | 0 | 0 | 17 |
| 875 | 603 | 26 | 5:6:4:0:0 | 5 | 6 | 4 | 0 | 0 | 14 |
| 876 | 603 | 26 | 6:6:1:0:0 | 6 | 6 | 1 | 0 | 0 | 19 |
| 877 | 603 | 26 | 6:7:1:0:0 | 6 | 7 | 1 | 0 | 0 | 17 |
| 878 | 603 | 26 | 6:7:2:0:0 | 6 | 7 | 2 | 0 | 0 | 16 |
| 879 | 603 | 26 | 6:7:3:0:0 | 6 | 7 | 3 | 0 | 0 | 16 |
| 880 | 603 | 26 | 6:7:4:0:0 | 6 | 7 | 4 | 0 | 0 | 16 |

| | | | | | | | | | |
|---|---|---|---|---|---|---|---|---|---|
| 881 | 616 | 27 | 2:5:0:0:0 | 2 | 5 | 0 | 0 | 0 | 13 |
| 882 | 616 | 27 | 3:4:0:0:0 | 3 | 4 | 0 | 0 | 0 | 5 |
| 883 | 616 | 27 | 3:4:1:0:1 | 3 | 4 | 1 | 0 | 1 | 5 |
| 884 | 616 | 27 | 3:5:0:0:0 | 3 | 5 | 0 | 0 | 0 | 11 |
| 885 | 616 | 27 | 3:5:0:1:0 | 3 | 5 | 0 | 1 | 0 | 5 |
| 886 | 616 | 27 | 3:5:1:0:0 | 3 | 5 | 1 | 0 | 0 | 14 |
| 887 | 616 | 27 | 4:5:0:0:0 | 4 | 5 | 0 | 0 | 0 | 14 |
| 888 | 616 | 27 | 4:5:1:0:0 | 4 | 5 | 1 | 0 | 0 | 19 |
| 889 | 616 | 27 | 4:5:1:0:1 | 4 | 5 | 1 | 0 | 1 | 14 |
| 890 | 616 | 27 | 4:5:2:0:0 | 4 | 5 | 2 | 0 | 0 | 18 |
| 891 | 616 | 27 | 4:5:3:0:0 | 4 | 5 | 3 | 0 | 0 | 17 |
| 892 | 616 | 27 | 4:6:0:1:0 | 4 | 6 | 0 | 1 | 0 | 14 |
| 893 | 616 | 27 | 5:5:0:0:0 | 5 | 5 | 0 | 0 | 0 | 6 |
| 894 | 616 | 27 | 5:5:1:0:0 | 5 | 5 | 1 | 0 | 0 | 18 |
| 895 | 616 | 27 | 5:5:2:0:0 | 5 | 5 | 2 | 0 | 0 | 15 |
| 896 | 616 | 27 | 5:6:0:0:0 | 5 | 6 | 0 | 0 | 0 | 9 |
| 897 | 616 | 27 | 5:6:1:0:0 | 5 | 6 | 1 | 0 | 0 | 16 |
| 898 | 616 | 27 | 5:6:2:0:0 | 5 | 6 | 2 | 0 | 0 | 16 |
| 899 | 616 | 27 | 5:6:3:0:0 | 5 | 6 | 3 | 0 | 0 | 17 |
| 900 | 616 | 27 | 5:6:4:0:0 | 5 | 6 | 4 | 0 | 0 | 14 |
| 901 | 616 | 27 | 6:6:1:0:0 | 6 | 6 | 1 | 0 | 0 | 19 |
| 902 | 616 | 27 | 6:7:0:0:0 | 6 | 7 | 0 | 0 | 0 | 6 |
| 903 | 616 | 27 | 6:7:1:0:0 | 6 | 7 | 1 | 0 | 0 | 17 |
| 904 | 616 | 27 | 6:7:1:0:1 | 6 | 7 | 1 | 0 | 1 | 13 |
| 905 | 616 | 27 | 6:7:2:0:0 | 6 | 7 | 2 | 0 | 0 | 16 |
| 906 | 616 | 27 | 6:7:3:0:0 | 6 | 7 | 3 | 0 | 0 | 16 |
| 907 | 616 | 27 | 6:8:0:1:0 | 6 | 8 | 0 | 1 | 0 | 13 |
| 908 | 657 | 21 | 3:5:2:0:0 | 3 | 5 | 2 | 0 | 0 | 13 |
| 909 | 657 | 21 | 4:5:1:0:0 | 4 | 5 | 1 | 0 | 0 | 19 |
| 910 | 657 | 21 | 4:5:2:0:0 | 4 | 5 | 2 | 0 | 0 | 18 |
| 911 | 657 | 21 | 4:5:3:0:0 | 4 | 5 | 3 | 0 | 0 | 17 |
| 912 | 657 | 21 | 5:5:1:0:0 | 5 | 5 | 1 | 0 | 0 | 18 |
| 913 | 657 | 21 | 5:5:2:0:0 | 5 | 5 | 2 | 0 | 0 | 15 |
| 914 | 657 | 21 | 5:5:3:0:0 | 5 | 5 | 3 | 0 | 0 | 12 |
| 915 | 657 | 21 | 5:6:3:0:0 | 5 | 6 | 3 | 0 | 0 | 17 |
| 916 | 657 | 21 | 6:6:2:0:0 | 6 | 6 | 2 | 0 | 0 | 12 |
| 917 | 657 | 21 | 6:6:3:0:0 | 6 | 6 | 3 | 0 | 0 | 14 |
| 918 | 657 | 21 | 6:6:4:0:0 | 6 | 6 | 4 | 0 | 0 | 10 |
| 919 | 657 | 21 | 6:7:2:0:0 | 6 | 7 | 2 | 0 | 0 | 16 |
| 920 | 657 | 21 | 6:7:2:0:1 | 6 | 7 | 2 | 0 | 1 | 9 |
| 921 | 657 | 21 | 6:7:3:0:0 | 6 | 7 | 3 | 0 | 0 | 16 |
| 922 | 657 | 21 | 6:7:3:0:1 | 6 | 7 | 3 | 0 | 1 | 8 |
| 923 | 657 | 21 | 6:7:4:0:0 | 6 | 7 | 4 | 0 | 0 | 16 |
| 924 | 657 | 21 | 6:8:1:1:0 | 6 | 8 | 1 | 1 | 0 | 9 |
| 925 | 657 | 21 | 6:8:2:1:0 | 6 | 8 | 2 | 1 | 0 | 8 |
| 926 | 657 | 21 | 7:7:2:0:0 | 7 | 7 | 2 | 0 | 0 | 6 |
| 927 | 657 | 21 | 7:7:3:0:0 | 7 | 7 | 3 | 0 | 0 | 8 |
| 928 | 709 | 20 | 2:5:0:0:0 | 2 | 5 | 0 | 0 | 0 | 13 |
| 929 | 709 | 20 | 2:6:0:0:0 | 2 | 6 | 0 | 0 | 0 | 13 |
| 930 | 709 | 20 | 2:7:0:0:0 | 2 | 7 | 0 | 0 | 0 | 11 |
| 931 | 709 | 20 | 2:8:0:0:0 | 2 | 8 | 0 | 0 | 0 | 10 |
| 932 | 709 | 20 | 3:6:0:0:0 | 3 | 6 | 0 | 0 | 0 | 11 |
| 933 | 709 | 20 | 4:5:0:0:0 | 4 | 5 | 0 | 0 | 0 | 14 |
| 934 | 709 | 20 | 4:5:1:0:0 | 4 | 5 | 1 | 0 | 0 | 19 |
| 935 | 709 | 20 | 5:4:0:0:0 | 5 | 4 | 0 | 0 | 0 | 5 |
| 936 | 709 | 20 | 5:4:1:0:0 | 5 | 4 | 1 | 0 | 0 | 16 |
| 937 | 709 | 20 | 5:6:1:0:0 | 5 | 6 | 1 | 0 | 0 | 16 |
| 938 | 709 | 20 | 5:6:2:0:0 | 5 | 6 | 2 | 0 | 0 | 16 |
| 939 | 709 | 20 | 6:5:1:0:0 | 6 | 5 | 1 | 0 | 0 | 10 |

| | | | | | | | | | |
|---|---|---|---|---|---|---|---|---|---|
| 940 | 709 | 20 | 6:6:1:0:0 | 6 | 6 | 1 | 0 | 0 | 19 |
| 941 | 709 | 20 | 6:7:1:0:0 | 6 | 7 | 1 | 0 | 0 | 17 |
| 942 | 709 | 20 | 6:7:1:0:1 | 6 | 7 | 1 | 0 | 1 | 13 |
| 943 | 709 | 20 | 6:7:2:0:0 | 6 | 7 | 2 | 0 | 0 | 16 |
| 944 | 709 | 20 | 6:7:3:0:0 | 6 | 7 | 3 | 0 | 0 | 16 |
| 945 | 709 | 20 | 6:7:4:0:0 | 6 | 7 | 4 | 0 | 0 | 16 |
| 946 | 709 | 20 | 6:8:0:1:0 | 6 | 8 | 0 | 1 | 0 | 13 |
| 947 | 709 | 20 | 7:6:1:0:0 | 7 | 6 | 1 | 0 | 0 | 4 |
| 948 | 717 | 8 | 2:6:0:0:0 | 2 | 6 | 0 | 0 | 0 | 13 |
| 949 | 717 | 8 | 2:7:0:0:0 | 2 | 7 | 0 | 0 | 0 | 11 |
| 950 | 717 | 8 | 2:8:0:0:0 | 2 | 8 | 0 | 0 | 0 | 10 |
| 951 | 717 | 8 | 3:5:0:0:0 | 3 | 5 | 0 | 0 | 0 | 11 |
| 952 | 717 | 8 | 3:5:1:0:0 | 3 | 5 | 1 | 0 | 0 | 14 |
| 953 | 717 | 8 | 3:6:0:0:0 | 3 | 6 | 0 | 0 | 0 | 11 |
| 954 | 717 | 8 | 3:6:0:0:1 | 3 | 6 | 0 | 0 | 1 | 8 |
| 955 | 717 | 8 | 3:6:1:0:0 | 3 | 6 | 1 | 0 | 0 | 13 |
| 956 | 801 | 76 | 2:4:0:0:0 | 2 | 4 | 0 | 0 | 0 | 3 |
| 957 | 801 | 76 | 2:5:0:0:0 | 2 | 5 | 0 | 0 | 0 | 13 |
| 958 | 801 | 76 | 2:6:0:0:0 | 2 | 6 | 0 | 0 | 0 | 13 |
| 959 | 801 | 76 | 2:7:0:0:0 | 2 | 7 | 0 | 0 | 0 | 11 |
| 960 | 801 | 76 | 2:8:0:0:0 | 2 | 8 | 0 | 0 | 0 | 10 |
| 961 | 801 | 76 | 2:9:0:0:0 | 2 | 9 | 0 | 0 | 0 | 5 |
| 962 | 801 | 76 | 3:4:0:0:0 | 3 | 4 | 0 | 0 | 0 | 5 |
| 963 | 801 | 76 | 3:4:1:0:1 | 3 | 4 | 1 | 0 | 1 | 5 |
| 964 | 801 | 76 | 3:5:0:0:0 | 3 | 5 | 0 | 0 | 0 | 11 |
| 965 | 801 | 76 | 3:5:0:0:1 | 3 | 5 | 0 | 0 | 1 | 6 |
| 966 | 801 | 76 | 3:5:0:1:0 | 3 | 5 | 0 | 1 | 0 | 5 |
| 967 | 801 | 76 | 3:5:1:0:0 | 3 | 5 | 1 | 0 | 0 | 14 |
| 968 | 801 | 76 | 3:5:1:0:1 | 3 | 5 | 1 | 0 | 1 | 4 |
| 969 | 801 | 76 | 3:5:2:0:0 | 3 | 5 | 2 | 0 | 0 | 13 |
| 970 | 801 | 76 | 3:6:0:0:0 | 3 | 6 | 0 | 0 | 0 | 11 |
| 971 | 801 | 76 | 3:6:0:0:1 | 3 | 6 | 0 | 0 | 1 | 8 |
| 972 | 801 | 76 | 3:6:0:1:0 | 3 | 6 | 0 | 1 | 0 | 4 |
| 973 | 801 | 76 | 3:6:1:0:0 | 3 | 6 | 1 | 0 | 0 | 13 |
| 974 | 801 | 76 | 3:7:1:0:0 | 3 | 7 | 1 | 0 | 0 | 4 |
| 975 | 801 | 76 | 4:3:0:0:0 | 4 | 3 | 0 | 0 | 0 | 6 |
| 976 | 801 | 76 | 4:3:1:0:0 | 4 | 3 | 1 | 0 | 0 | 14 |
| 977 | 801 | 76 | 4:4:0:0:0 | 4 | 4 | 0 | 0 | 0 | 6 |
| 978 | 801 | 76 | 4:4:1:0:0 | 4 | 4 | 1 | 0 | 0 | 12 |
| 979 | 801 | 76 | 4:4:1:0:1 | 4 | 4 | 1 | 0 | 1 | 7 |
| 980 | 801 | 76 | 4:4:2:0:0 | 4 | 4 | 2 | 0 | 0 | 8 |
| 981 | 801 | 76 | 4:4:2:0:1 | 4 | 4 | 2 | 0 | 1 | 8 |
| 982 | 801 | 76 | 4:4:2:0:2 | 4 | 4 | 2 | 0 | 2 | 1 |
| 983 | 801 | 76 | 4:5:0:0:0 | 4 | 5 | 0 | 0 | 0 | 14 |
| 984 | 801 | 76 | 4:5:0:0:1 | 4 | 5 | 0 | 0 | 1 | 7 |
| 985 | 801 | 76 | 4:5:0:0:2 | 4 | 5 | 0 | 0 | 2 | 1 |
| 986 | 801 | 76 | 4:5:0:1:0 | 4 | 5 | 0 | 1 | 0 | 7 |
| 987 | 801 | 76 | 4:5:1:0:0 | 4 | 5 | 1 | 0 | 0 | 19 |
| 988 | 801 | 76 | 4:5:1:0:1 | 4 | 5 | 1 | 0 | 1 | 14 |
| 989 | 801 | 76 | 4:5:1:1:0 | 4 | 5 | 1 | 1 | 0 | 8 |
| 990 | 801 | 76 | 4:5:1:1:1 | 4 | 5 | 1 | 1 | 1 | 1 |
| 991 | 801 | 76 | 4:5:2:0:0 | 4 | 5 | 2 | 0 | 0 | 18 |
| 992 | 801 | 76 | 4:5:2:0:1 | 4 | 5 | 2 | 0 | 1 | 9 |
| 993 | 801 | 76 | 4:5:3:0:0 | 4 | 5 | 3 | 0 | 0 | 17 |
| 994 | 801 | 76 | 4:6:0:0:0 | 4 | 6 | 0 | 0 | 0 | 3 |
| 995 | 801 | 76 | 4:6:0:1:0 | 4 | 6 | 0 | 1 | 0 | 14 |
| 996 | 801 | 76 | 4:6:0:2:0 | 4 | 6 | 0 | 2 | 0 | 1 |
| 997 | 801 | 76 | 4:6:1:1:0 | 4 | 6 | 1 | 1 | 0 | 9 |
| 998 | 801 | 76 | 4:6:2:0:0 | 4 | 6 | 2 | 0 | 0 | 5 |

| | | | | | | | | | |
|---|---|---|---|---|---|---|---|---|---|
| 999 | 801 | 76 | 4:7:0:0:0 | 4 | 7 | 0 | 0 | 0 | 3 |
| 1000 | 801 | 76 | 4:7:1:0:0 | 4 | 7 | 1 | 0 | 0 | 5 |
| 1001 | 801 | 76 | 4:7:2:0:0 | 4 | 7 | 2 | 0 | 0 | 5 |
| 1002 | 801 | 76 | 5:3:0:0:0 | 5 | 3 | 0 | 0 | 0 | 6 |
| 1003 | 801 | 76 | 5:3:1:0:0 | 5 | 3 | 1 | 0 | 0 | 11 |
| 1004 | 801 | 76 | 5:4:0:0:0 | 5 | 4 | 0 | 0 | 0 | 5 |
| 1005 | 801 | 76 | 5:4:1:0:0 | 5 | 4 | 1 | 0 | 0 | 16 |
| 1006 | 801 | 76 | 5:4:1:0:1 | 5 | 4 | 1 | 0 | 1 | 4 |
| 1007 | 801 | 76 | 5:4:2:0:0 | 5 | 4 | 2 | 0 | 0 | 8 |
| 1008 | 801 | 76 | 5:5:0:0:0 | 5 | 5 | 0 | 0 | 0 | 6 |
| 1009 | 801 | 76 | 5:5:0:1:0 | 5 | 5 | 0 | 1 | 0 | 4 |
| 1010 | 801 | 76 | 5:5:1:0:0 | 5 | 5 | 1 | 0 | 0 | 18 |
| 1011 | 801 | 76 | 5:5:1:0:1 | 5 | 5 | 1 | 0 | 1 | 7 |
| 1012 | 801 | 76 | 5:5:2:0:0 | 5 | 5 | 2 | 0 | 0 | 15 |
| 1013 | 801 | 76 | 5:5:2:0:1 | 5 | 5 | 2 | 0 | 1 | 7 |
| 1014 | 801 | 76 | 5:5:3:0:0 | 5 | 5 | 3 | 0 | 0 | 12 |
| 1015 | 801 | 76 | 5:6:0:0:0 | 5 | 6 | 0 | 0 | 0 | 9 |
| 1016 | 801 | 76 | 5:6:0:1:0 | 5 | 6 | 0 | 1 | 0 | 7 |
| 1017 | 801 | 76 | 5:6:0:1:2 | 5 | 6 | 0 | 1 | 2 | 1 |
| 1018 | 801 | 76 | 5:6:1:0:0 | 5 | 6 | 1 | 0 | 0 | 16 |
| 1019 | 801 | 76 | 5:6:1:0:1 | 5 | 6 | 1 | 0 | 1 | 8 |
| 1020 | 801 | 76 | 5:6:1:1:0 | 5 | 6 | 1 | 1 | 0 | 7 |
| 1021 | 801 | 76 | 5:6:2:0:0 | 5 | 6 | 2 | 0 | 0 | 16 |
| 1022 | 801 | 76 | 5:6:3:0:0 | 5 | 6 | 3 | 0 | 0 | 17 |
| 1023 | 801 | 76 | 5:6:4:0:0 | 5 | 6 | 4 | 0 | 0 | 14 |
| 1024 | 801 | 76 | 5:7:0:1:0 | 5 | 7 | 0 | 1 | 0 | 8 |
| 1025 | 801 | 76 | 6:3:1:0:0 | 6 | 3 | 1 | 0 | 0 | 9 |
| 1026 | 801 | 76 | 6:5:1:0:0 | 6 | 5 | 1 | 0 | 0 | 10 |
| 1027 | 801 | 76 | 6:6:0:0:0 | 6 | 6 | 0 | 0 | 0 | 6 |
| 1028 | 801 | 76 | 6:6:1:0:0 | 6 | 6 | 1 | 0 | 0 | 19 |
| 1029 | 801 | 76 | 6:6:2:0:0 | 6 | 6 | 2 | 0 | 0 | 12 |
| 1030 | 801 | 76 | 6:6:3:0:0 | 6 | 6 | 3 | 0 | 0 | 14 |
| 1031 | 1074 | 155 | 2:3:0:0:0 | 2 | 3 | 0 | 0 | 0 | 1 |
| 1032 | 1074 | 155 | 2:4:0:0:0 | 2 | 4 | 0 | 0 | 0 | 3 |
| 1033 | 1074 | 155 | 2:5:0:0:0 | 2 | 5 | 0 | 0 | 0 | 13 |
| 1034 | 1074 | 155 | 2:6:0:0:0 | 2 | 6 | 0 | 0 | 0 | 13 |
| 1035 | 1074 | 155 | 2:7:0:0:0 | 2 | 7 | 0 | 0 | 0 | 11 |
| 1036 | 1074 | 155 | 2:8:0:0:0 | 2 | 8 | 0 | 0 | 0 | 10 |
| 1037 | 1074 | 155 | 2:9:0:0:0 | 2 | 9 | 0 | 0 | 0 | 5 |
| 1038 | 1074 | 155 | 3:3:0:0:0 | 3 | 3 | 0 | 0 | 0 | 3 |
| 1039 | 1074 | 155 | 3:3:1:0:0 | 3 | 3 | 1 | 0 | 0 | 4 |
| 1040 | 1074 | 155 | 3:4:0:0:0 | 3 | 4 | 0 | 0 | 0 | 5 |
| 1041 | 1074 | 155 | 3:4:0:0:1 | 3 | 4 | 0 | 0 | 1 | 3 |
| 1042 | 1074 | 155 | 3:4:1:0:0 | 3 | 4 | 1 | 0 | 0 | 8 |
| 1043 | 1074 | 155 | 3:4:1:0:1 | 3 | 4 | 1 | 0 | 1 | 5 |
| 1044 | 1074 | 155 | 3:4:2:0:0 | 3 | 4 | 2 | 0 | 0 | 5 |
| 1045 | 1074 | 155 | 3:4:2:0:1 | 3 | 4 | 2 | 0 | 1 | 4 |
| 1046 | 1074 | 155 | 3:5:0:0:0 | 3 | 5 | 0 | 0 | 0 | 11 |
| 1047 | 1074 | 155 | 3:5:0:0:1 | 3 | 5 | 0 | 0 | 1 | 6 |
| 1048 | 1074 | 155 | 3:5:0:1:0 | 3 | 5 | 0 | 1 | 0 | 5 |
| 1049 | 1074 | 155 | 3:5:1:0:0 | 3 | 5 | 1 | 0 | 0 | 14 |
| 1050 | 1074 | 155 | 3:5:1:0:1 | 3 | 5 | 1 | 0 | 1 | 4 |
| 1051 | 1074 | 155 | 3:5:1:1:0 | 3 | 5 | 1 | 1 | 0 | 4 |
| 1052 | 1074 | 155 | 3:5:2:0:0 | 3 | 5 | 2 | 0 | 0 | 13 |
| 1053 | 1074 | 155 | 3:6:0:0:0 | 3 | 6 | 0 | 0 | 0 | 11 |
| 1054 | 1074 | 155 | 3:6:0:0:1 | 3 | 6 | 0 | 0 | 1 | 8 |
| 1055 | 1074 | 155 | 3:6:0:1:0 | 3 | 6 | 0 | 1 | 0 | 4 |
| 1056 | 1074 | 155 | 3:6:1:0:0 | 3 | 6 | 1 | 0 | 0 | 13 |
| 1057 | 1074 | 155 | 3:6:1:0:1 | 3 | 6 | 1 | 0 | 1 | 2 |

| | | | | | | | | | |
|---|---|---|---|---|---|---|---|---|---|
| 1058 | 1074 | 155 | 3:6:2:0:0 | 3 | 6 | 2 | 0 | 0 | 6 |
| 1059 | 1074 | 155 | 3:7:0:0:0 | 3 | 7 | 0 | 0 | 0 | 2 |
| 1060 | 1074 | 155 | 3:7:1:0:0 | 3 | 7 | 1 | 0 | 0 | 4 |
| 1061 | 1074 | 155 | 4:3:0:0:0 | 4 | 3 | 0 | 0 | 0 | 6 |
| 1062 | 1074 | 155 | 4:3:1:0:0 | 4 | 3 | 1 | 0 | 0 | 14 |
| 1063 | 1074 | 155 | 4:3:2:0:1 | 4 | 3 | 2 | 0 | 1 | 2 |
| 1064 | 1074 | 155 | 4:4:0:0:0 | 4 | 4 | 0 | 0 | 0 | 6 |
| 1065 | 1074 | 155 | 4:4:1:0:0 | 4 | 4 | 1 | 0 | 0 | 12 |
| 1066 | 1074 | 155 | 4:4:1:0:1 | 4 | 4 | 1 | 0 | 1 | 7 |
| 1067 | 1074 | 155 | 4:4:1:1:0 | 4 | 4 | 1 | 1 | 0 | 2 |
| 1068 | 1074 | 155 | 4:4:2:0:0 | 4 | 4 | 2 | 0 | 0 | 8 |
| 1069 | 1074 | 155 | 4:4:2:0:1 | 4 | 4 | 2 | 0 | 1 | 8 |
| 1070 | 1074 | 155 | 4:4:3:0:1 | 4 | 4 | 3 | 0 | 1 | 4 |
| 1071 | 1074 | 155 | 4:5:0:0:0 | 4 | 5 | 0 | 0 | 0 | 14 |
| 1072 | 1074 | 155 | 4:5:0:0:1 | 4 | 5 | 0 | 0 | 1 | 7 |
| 1073 | 1074 | 155 | 4:5:0:1:0 | 4 | 5 | 0 | 1 | 0 | 7 |
| 1074 | 1074 | 155 | 4:5:1:0:0 | 4 | 5 | 1 | 0 | 0 | 19 |
| 1075 | 1074 | 155 | 4:5:1:0:1 | 4 | 5 | 1 | 0 | 1 | 14 |
| 1076 | 1074 | 155 | 4:5:1:1:0 | 4 | 5 | 1 | 1 | 0 | 8 |
| 1077 | 1074 | 155 | 4:5:2:0:0 | 4 | 5 | 2 | 0 | 0 | 18 |
| 1078 | 1074 | 155 | 4:5:2:0:1 | 4 | 5 | 2 | 0 | 1 | 9 |
| 1079 | 1074 | 155 | 4:5:2:1:0 | 4 | 5 | 2 | 1 | 0 | 4 |
| 1080 | 1074 | 155 | 4:5:3:0:0 | 4 | 5 | 3 | 0 | 0 | 17 |
| 1081 | 1074 | 155 | 4:6:0:0:0 | 4 | 6 | 0 | 0 | 0 | 3 |
| 1082 | 1074 | 155 | 4:6:0:1:0 | 4 | 6 | 0 | 1 | 0 | 14 |
| 1083 | 1074 | 155 | 4:6:1:0:0 | 4 | 6 | 1 | 0 | 0 | 3 |
| 1084 | 1074 | 155 | 4:6:1:0:1 | 4 | 6 | 1 | 0 | 1 | 2 |
| 1085 | 1074 | 155 | 4:6:1:1:0 | 4 | 6 | 1 | 1 | 0 | 9 |
| 1086 | 1074 | 155 | 4:6:2:0:0 | 4 | 6 | 2 | 0 | 0 | 5 |
| 1087 | 1074 | 155 | 4:6:2:0:1 | 4 | 6 | 2 | 0 | 1 | 1 |
| 1088 | 1074 | 155 | 4:6:3:0:0 | 4 | 6 | 3 | 0 | 0 | 3 |
| 1089 | 1074 | 155 | 4:7:0:0:0 | 4 | 7 | 0 | 0 | 0 | 3 |
| 1090 | 1074 | 155 | 4:7:0:1:0 | 4 | 7 | 0 | 1 | 0 | 2 |
| 1091 | 1074 | 155 | 4:7:1:0:0 | 4 | 7 | 1 | 0 | 0 | 5 |
| 1092 | 1074 | 155 | 4:7:1:1:0 | 4 | 7 | 1 | 1 | 0 | 1 |
| 1093 | 1074 | 155 | 4:7:2:0:0 | 4 | 7 | 2 | 0 | 0 | 5 |
| 1094 | 1074 | 155 | 5:3:0:0:0 | 5 | 3 | 0 | 0 | 0 | 6 |
| 1095 | 1074 | 155 | 5:3:1:0:0 | 5 | 3 | 1 | 0 | 0 | 11 |
| 1096 | 1074 | 155 | 5:4:0:0:0 | 5 | 4 | 0 | 0 | 0 | 5 |
| 1097 | 1074 | 155 | 5:4:1:0:0 | 5 | 4 | 1 | 0 | 0 | 16 |
| 1098 | 1074 | 155 | 5:4:2:0:0 | 5 | 4 | 2 | 0 | 0 | 8 |
| 1099 | 1074 | 155 | 5:5:0:0:0 | 5 | 5 | 0 | 0 | 0 | 6 |
| 1100 | 1074 | 155 | 5:5:1:0:0 | 5 | 5 | 1 | 0 | 0 | 18 |
| 1101 | 1074 | 155 | 5:5:1:0:1 | 5 | 5 | 1 | 0 | 1 | 7 |
| 1102 | 1074 | 155 | 5:5:2:0:0 | 5 | 5 | 2 | 0 | 0 | 15 |
| 1103 | 1074 | 155 | 5:5:2:0:1 | 5 | 5 | 2 | 0 | 1 | 7 |
| 1104 | 1074 | 155 | 5:5:3:0:0 | 5 | 5 | 3 | 0 | 0 | 12 |
| 1105 | 1074 | 155 | 5:5:3:0:1 | 5 | 5 | 3 | 0 | 1 | 2 |
| 1106 | 1074 | 155 | 5:5:4:0:1 | 5 | 5 | 4 | 0 | 1 | 3 |
| 1107 | 1074 | 155 | 5:6:0:0:0 | 5 | 6 | 0 | 0 | 0 | 9 |
| 1108 | 1074 | 155 | 5:6:0:1:0 | 5 | 6 | 0 | 1 | 0 | 7 |
| 1109 | 1074 | 155 | 5:6:1:0:0 | 5 | 6 | 1 | 0 | 0 | 16 |
| 1110 | 1074 | 155 | 5:6:1:0:1 | 5 | 6 | 1 | 0 | 1 | 8 |
| 1111 | 1074 | 155 | 5:6:1:0:2 | 5 | 6 | 1 | 0 | 2 | 4 |
| 1112 | 1074 | 155 | 5:6:1:1:0 | 5 | 6 | 1 | 1 | 0 | 7 |
| 1113 | 1074 | 155 | 5:6:2:0:0 | 5 | 6 | 2 | 0 | 0 | 16 |
| 1114 | 1074 | 155 | 5:6:2:0:1 | 5 | 6 | 2 | 0 | 1 | 9 |
| 1115 | 1074 | 155 | 5:6:2:1:0 | 5 | 6 | 2 | 1 | 0 | 2 |
| 1116 | 1074 | 155 | 5:6:3:0:0 | 5 | 6 | 3 | 0 | 0 | 17 |

| | | | | | | | | | |
|---|---|---|---|---|---|---|---|---|---|
| 1117 | 1074 | 155 | 5:6:3:0:1 | 5 | 6 | 3 | 0 | 1 | 7 |
| 1118 | 1074 | 155 | 5:6:3:1:0 | 5 | 6 | 3 | 1 | 0 | 3 |
| 1119 | 1074 | 155 | 5:6:4:0:0 | 5 | 6 | 4 | 0 | 0 | 14 |
| 1120 | 1074 | 155 | 5:7:0:1:0 | 5 | 7 | 0 | 1 | 0 | 8 |
| 1121 | 1074 | 155 | 5:7:0:1:1 | 5 | 7 | 0 | 1 | 1 | 4 |
| 1122 | 1074 | 155 | 5:7:1:0:0 | 5 | 7 | 1 | 0 | 0 | 2 |
| 1123 | 1074 | 155 | 5:7:1:0:1 | 5 | 7 | 1 | 0 | 1 | 2 |
| 1124 | 1074 | 155 | 5:7:1:1:0 | 5 | 7 | 1 | 1 | 0 | 9 |
| 1125 | 1074 | 155 | 5:7:2:0:0 | 5 | 7 | 2 | 0 | 0 | 3 |
| 1126 | 1074 | 155 | 5:7:2:1:0 | 5 | 7 | 2 | 1 | 0 | 7 |
| 1127 | 1074 | 155 | 5:8:0:1:0 | 5 | 8 | 0 | 1 | 0 | 2 |
| 1128 | 1074 | 155 | 6:3:1:0:0 | 6 | 3 | 1 | 0 | 0 | 9 |
| 1129 | 1074 | 155 | 6:4:1:0:0 | 6 | 4 | 1 | 0 | 0 | 8 |
| 1130 | 1074 | 155 | 6:5:0:0:0 | 6 | 5 | 0 | 0 | 0 | 3 |
| 1131 | 1074 | 155 | 6:5:1:0:0 | 6 | 5 | 1 | 0 | 0 | 10 |
| 1132 | 1074 | 155 | 6:5:1:0:1 | 6 | 5 | 1 | 0 | 1 | 2 |
| 1133 | 1074 | 155 | 6:5:2:0:0 | 6 | 5 | 2 | 0 | 0 | 4 |
| 1134 | 1074 | 155 | 6:5:2:0:1 | 6 | 5 | 2 | 0 | 1 | 4 |
| 1135 | 1074 | 155 | 6:5:4:0:1 | 6 | 5 | 4 | 0 | 1 | 3 |
| 1136 | 1074 | 155 | 6:6:0:0:0 | 6 | 6 | 0 | 0 | 0 | 6 |
| 1137 | 1074 | 155 | 6:6:0:0:1 | 6 | 6 | 0 | 0 | 1 | 2 |
| 1138 | 1074 | 155 | 6:6:0:1:0 | 6 | 6 | 0 | 1 | 0 | 2 |
| 1139 | 1074 | 155 | 6:6:1:0:0 | 6 | 6 | 1 | 0 | 0 | 19 |
| 1140 | 1074 | 155 | 6:6:1:0:1 | 6 | 6 | 1 | 0 | 1 | 7 |
| 1141 | 1074 | 155 | 6:6:1:1:0 | 6 | 6 | 1 | 1 | 0 | 4 |
| 1142 | 1074 | 155 | 6:6:2:0:0 | 6 | 6 | 2 | 0 | 0 | 12 |
| 1143 | 1074 | 155 | 6:6:2:0:1 | 6 | 6 | 2 | 0 | 1 | 8 |
| 1144 | 1074 | 155 | 6:6:3:0:0 | 6 | 6 | 3 | 0 | 0 | 14 |
| 1145 | 1074 | 155 | 6:6:3:0:1 | 6 | 6 | 3 | 0 | 1 | 7 |
| 1146 | 1074 | 155 | 6:6:3:1:0 | 6 | 6 | 3 | 1 | 0 | 3 |
| 1147 | 1074 | 155 | 6:6:4:0:0 | 6 | 6 | 4 | 0 | 0 | 10 |
| 1148 | 1074 | 155 | 6:6:4:0:1 | 6 | 6 | 4 | 0 | 1 | 3 |
| 1149 | 1074 | 155 | 6:7:0:0:0 | 6 | 7 | 0 | 0 | 0 | 6 |
| 1150 | 1074 | 155 | 6:7:0:0:1 | 6 | 7 | 0 | 0 | 1 | 2 |
| 1151 | 1074 | 155 | 6:7:0:1:0 | 6 | 7 | 0 | 1 | 0 | 7 |
| 1152 | 1074 | 155 | 6:7:1:0:0 | 6 | 7 | 1 | 0 | 0 | 17 |
| 1153 | 1074 | 155 | 6:7:1:0:1 | 6 | 7 | 1 | 0 | 1 | 13 |
| 1154 | 1074 | 155 | 6:7:1:0:2 | 6 | 7 | 1 | 0 | 2 | 5 |
| 1155 | 1074 | 155 | 6:7:1:1:0 | 6 | 7 | 1 | 1 | 0 | 8 |
| 1156 | 1074 | 155 | 6:7:2:0:0 | 6 | 7 | 2 | 0 | 0 | 16 |
| 1157 | 1074 | 155 | 6:7:2:0:1 | 6 | 7 | 2 | 0 | 1 | 9 |
| 1158 | 1074 | 155 | 6:7:2:1:0 | 6 | 7 | 2 | 1 | 0 | 7 |
| 1159 | 1074 | 155 | 6:7:3:0:0 | 6 | 7 | 3 | 0 | 0 | 16 |
| 1160 | 1074 | 155 | 6:7:3:0:1 | 6 | 7 | 3 | 0 | 1 | 8 |
| 1161 | 1074 | 155 | 6:7:3:1:0 | 6 | 7 | 3 | 1 | 0 | 3 |
| 1162 | 1074 | 155 | 6:7:4:0:0 | 6 | 7 | 4 | 0 | 0 | 16 |
| 1163 | 1074 | 155 | 6:7:4:0:1 | 6 | 7 | 4 | 0 | 1 | 5 |
| 1164 | 1074 | 155 | 6:8:0:0:0 | 6 | 8 | 0 | 0 | 0 | 2 |
| 1165 | 1074 | 155 | 6:8:0:1:0 | 6 | 8 | 0 | 1 | 0 | 13 |
| 1166 | 1074 | 155 | 6:8:0:1:1 | 6 | 8 | 0 | 1 | 1 | 5 |
| 1167 | 1074 | 155 | 6:8:1:1:0 | 6 | 8 | 1 | 1 | 0 | 9 |
| 1168 | 1074 | 155 | 6:8:2:1:0 | 6 | 8 | 2 | 1 | 0 | 8 |
| 1169 | 1074 | 155 | 6:8:3:1:0 | 6 | 8 | 3 | 1 | 0 | 5 |
| 1170 | 1074 | 155 | 7:3:1:0:0 | 7 | 3 | 1 | 0 | 0 | 2 |
| 1171 | 1074 | 155 | 7:4:1:0:0 | 7 | 4 | 1 | 0 | 0 | 2 |
| 1172 | 1074 | 155 | 7:5:1:0:0 | 7 | 5 | 1 | 0 | 0 | 1 |
| 1173 | 1074 | 155 | 7:6:1:0:0 | 7 | 6 | 1 | 0 | 0 | 4 |
| 1174 | 1074 | 155 | 7:6:2:0:0 | 7 | 6 | 2 | 0 | 0 | 3 |
| 1175 | 1074 | 155 | 7:6:3:0:0 | 7 | 6 | 3 | 0 | 0 | 2 |

| | | | | | | | | | |
|---|---|---|---|---|---|---|---|---|---|
| 1176 | 1074 | 155 | 7:7:1:0:0 | 7 | 7 | 1 | 0 | 0 | 6 |
| 1177 | 1074 | 155 | 7:7:1:0:1 | 7 | 7 | 1 | 0 | 1 | 3 |
| 1178 | 1074 | 155 | 7:7:2:0:0 | 7 | 7 | 2 | 0 | 0 | 6 |
| 1179 | 1074 | 155 | 7:7:3:0:0 | 7 | 7 | 3 | 0 | 0 | 8 |
| 1180 | 1074 | 155 | 7:8:0:0:0 | 7 | 8 | 0 | 0 | 0 | 2 |
| 1181 | 1074 | 155 | 7:8:0:1:0 | 7 | 8 | 0 | 1 | 0 | 3 |
| 1182 | 1074 | 155 | 7:8:1:0:0 | 7 | 8 | 1 | 0 | 0 | 4 |
| 1183 | 1074 | 155 | 7:8:2:0:0 | 7 | 8 | 2 | 0 | 0 | 2 |
| 1184 | 1074 | 155 | 8:9:0:0:0 | 8 | 9 | 0 | 0 | 0 | 5 |
| 1185 | 1098 | 146 | 2:4:0:0:0 | 2 | 4 | 0 | 0 | 0 | 3 |
| 1186 | 1098 | 146 | 2:5:0:0:0 | 2 | 5 | 0 | 0 | 0 | 13 |
| 1187 | 1098 | 146 | 2:6:0:0:0 | 2 | 6 | 0 | 0 | 0 | 13 |
| 1188 | 1098 | 146 | 2:7:0:0:0 | 2 | 7 | 0 | 0 | 0 | 11 |
| 1189 | 1098 | 146 | 2:8:0:0:0 | 2 | 8 | 0 | 0 | 0 | 10 |
| 1190 | 1098 | 146 | 2:9:0:0:0 | 2 | 9 | 0 | 0 | 0 | 5 |
| 1191 | 1098 | 146 | 3:3:0:0:0 | 3 | 3 | 0 | 0 | 0 | 3 |
| 1192 | 1098 | 146 | 3:3:1:0:0 | 3 | 3 | 1 | 0 | 0 | 4 |
| 1193 | 1098 | 146 | 3:4:0:0:0 | 3 | 4 | 0 | 0 | 0 | 5 |
| 1194 | 1098 | 146 | 3:4:0:0:1 | 3 | 4 | 0 | 0 | 1 | 3 |
| 1195 | 1098 | 146 | 3:4:1:0:0 | 3 | 4 | 1 | 0 | 0 | 8 |
| 1196 | 1098 | 146 | 3:4:1:0:1 | 3 | 4 | 1 | 0 | 1 | 5 |
| 1197 | 1098 | 146 | 3:4:2:0:1 | 3 | 4 | 2 | 0 | 1 | 4 |
| 1198 | 1098 | 146 | 3:5:0:0:0 | 3 | 5 | 0 | 0 | 0 | 11 |
| 1199 | 1098 | 146 | 3:5:0:0:1 | 3 | 5 | 0 | 0 | 1 | 6 |
| 1200 | 1098 | 146 | 3:5:0:1:0 | 3 | 5 | 0 | 1 | 0 | 5 |
| 1201 | 1098 | 146 | 3:5:1:0:0 | 3 | 5 | 1 | 0 | 0 | 14 |
| 1202 | 1098 | 146 | 3:5:1:0:1 | 3 | 5 | 1 | 0 | 1 | 4 |
| 1203 | 1098 | 146 | 3:5:1:0:2 | 3 | 5 | 1 | 0 | 2 | 1 |
| 1204 | 1098 | 146 | 3:5:1:1:0 | 3 | 5 | 1 | 1 | 0 | 4 |
| 1205 | 1098 | 146 | 3:5:1:1:1 | 3 | 5 | 1 | 1 | 1 | 1 |
| 1206 | 1098 | 146 | 3:5:2:0:0 | 3 | 5 | 2 | 0 | 0 | 13 |
| 1207 | 1098 | 146 | 3:6:0:0:0 | 3 | 6 | 0 | 0 | 0 | 11 |
| 1208 | 1098 | 146 | 3:6:0:0:1 | 3 | 6 | 0 | 0 | 1 | 8 |
| 1209 | 1098 | 146 | 3:6:0:1:0 | 3 | 6 | 0 | 1 | 0 | 4 |
| 1210 | 1098 | 146 | 3:6:1:0:0 | 3 | 6 | 1 | 0 | 0 | 13 |
| 1211 | 1098 | 146 | 3:6:1:0:1 | 3 | 6 | 1 | 0 | 1 | 2 |
| 1212 | 1098 | 146 | 3:6:2:0:0 | 3 | 6 | 2 | 0 | 0 | 6 |
| 1213 | 1098 | 146 | 3:7:0:0:0 | 3 | 7 | 0 | 0 | 0 | 2 |
| 1214 | 1098 | 146 | 3:7:1:0:0 | 3 | 7 | 1 | 0 | 0 | 4 |
| 1215 | 1098 | 146 | 4:3:0:0:0 | 4 | 3 | 0 | 0 | 0 | 6 |
| 1216 | 1098 | 146 | 4:3:1:0:0 | 4 | 3 | 1 | 0 | 0 | 14 |
| 1217 | 1098 | 146 | 4:4:0:0:0 | 4 | 4 | 0 | 0 | 0 | 6 |
| 1218 | 1098 | 146 | 4:4:1:0:0 | 4 | 4 | 1 | 0 | 0 | 12 |
| 1219 | 1098 | 146 | 4:4:1:0:1 | 4 | 4 | 1 | 0 | 1 | 7 |
| 1220 | 1098 | 146 | 4:4:2:0:1 | 4 | 4 | 2 | 0 | 1 | 8 |
| 1221 | 1098 | 146 | 4:5:0:0:0 | 4 | 5 | 0 | 0 | 0 | 14 |
| 1222 | 1098 | 146 | 4:5:0:0:1 | 4 | 5 | 0 | 0 | 1 | 7 |
| 1223 | 1098 | 146 | 4:5:0:1:0 | 4 | 5 | 0 | 1 | 0 | 7 |
| 1224 | 1098 | 146 | 4:5:1:0:0 | 4 | 5 | 1 | 0 | 0 | 19 |
| 1225 | 1098 | 146 | 4:5:1:0:1 | 4 | 5 | 1 | 0 | 1 | 14 |
| 1226 | 1098 | 146 | 4:5:1:1:0 | 4 | 5 | 1 | 1 | 0 | 8 |
| 1227 | 1098 | 146 | 4:5:2:0:0 | 4 | 5 | 2 | 0 | 0 | 18 |
| 1228 | 1098 | 146 | 4:5:2:0:1 | 4 | 5 | 2 | 0 | 1 | 9 |
| 1229 | 1098 | 146 | 4:5:3:0:0 | 4 | 5 | 3 | 0 | 0 | 17 |
| 1230 | 1098 | 146 | 4:6:0:0:0 | 4 | 6 | 0 | 0 | 0 | 3 |
| 1231 | 1098 | 146 | 4:6:0:0:1 | 4 | 6 | 0 | 0 | 1 | 1 |
| 1232 | 1098 | 146 | 4:6:0:1:0 | 4 | 6 | 0 | 1 | 0 | 14 |
| 1233 | 1098 | 146 | 4:6:1:0:0 | 4 | 6 | 1 | 0 | 0 | 3 |
| 1234 | 1098 | 146 | 4:6:1:0:1 | 4 | 6 | 1 | 0 | 1 | 2 |

| | | | | | | | | | |
|---|---|---|---|---|---|---|---|---|---|
| 1235 | 1098 | 146 | 4:6:1:1:0 | 4 | 6 | 1 | 1 | 0 | 9 |
| 1236 | 1098 | 146 | 4:6:2:0:0 | 4 | 6 | 2 | 0 | 0 | 5 |
| 1237 | 1098 | 146 | 4:6:3:0:0 | 4 | 6 | 3 | 0 | 0 | 3 |
| 1238 | 1098 | 146 | 4:6:4:2:0 | 4 | 6 | 4 | 2 | 0 | 1 |
| 1239 | 1098 | 146 | 4:7:0:0:0 | 4 | 7 | 0 | 0 | 0 | 3 |
| 1240 | 1098 | 146 | 4:7:0:0:1 | 4 | 7 | 0 | 0 | 1 | 1 |
| 1241 | 1098 | 146 | 4:7:0:1:0 | 4 | 7 | 0 | 1 | 0 | 2 |
| 1242 | 1098 | 146 | 4:7:1:0:0 | 4 | 7 | 1 | 0 | 0 | 5 |
| 1243 | 1098 | 146 | 4:7:2:0:0 | 4 | 7 | 2 | 0 | 0 | 5 |
| 1244 | 1098 | 146 | 4:8:0:0:0 | 4 | 8 | 0 | 0 | 0 | 1 |
| 1245 | 1098 | 146 | 4:8:1:0:0 | 4 | 8 | 1 | 0 | 0 | 1 |
| 1246 | 1098 | 146 | 5:3:0:0:0 | 5 | 3 | 0 | 0 | 0 | 6 |
| 1247 | 1098 | 146 | 5:3:1:0:0 | 5 | 3 | 1 | 0 | 0 | 11 |
| 1248 | 1098 | 146 | 5:4:0:0:0 | 5 | 4 | 0 | 0 | 0 | 5 |
| 1249 | 1098 | 146 | 5:4:1:0:0 | 5 | 4 | 1 | 0 | 0 | 16 |
| 1250 | 1098 | 146 | 5:5:0:0:0 | 5 | 5 | 0 | 0 | 0 | 6 |
| 1251 | 1098 | 146 | 5:5:1:0:0 | 5 | 5 | 1 | 0 | 0 | 18 |
| 1252 | 1098 | 146 | 5:5:1:0:1 | 5 | 5 | 1 | 0 | 1 | 7 |
| 1253 | 1098 | 146 | 5:5:1:0:2 | 5 | 5 | 1 | 0 | 2 | 2 |
| 1254 | 1098 | 146 | 5:5:2:0:0 | 5 | 5 | 2 | 0 | 0 | 15 |
| 1255 | 1098 | 146 | 5:5:2:0:1 | 5 | 5 | 2 | 0 | 1 | 7 |
| 1256 | 1098 | 146 | 5:5:3:0:1 | 5 | 5 | 3 | 0 | 1 | 2 |
| 1257 | 1098 | 146 | 5:6:0:0:0 | 5 | 6 | 0 | 0 | 0 | 9 |
| 1258 | 1098 | 146 | 5:6:0:0:1 | 5 | 6 | 0 | 0 | 1 | 2 |
| 1259 | 1098 | 146 | 5:6:0:0:2 | 5 | 6 | 0 | 0 | 2 | 1 |
| 1260 | 1098 | 146 | 5:6:0:1:0 | 5 | 6 | 0 | 1 | 0 | 7 |
| 1261 | 1098 | 146 | 5:6:0:1:1 | 5 | 6 | 0 | 1 | 1 | 2 |
| 1262 | 1098 | 146 | 5:6:1:0:0 | 5 | 6 | 1 | 0 | 0 | 16 |
| 1263 | 1098 | 146 | 5:6:1:0:1 | 5 | 6 | 1 | 0 | 1 | 8 |
| 1264 | 1098 | 146 | 5:6:1:1:0 | 5 | 6 | 1 | 1 | 0 | 7 |
| 1265 | 1098 | 146 | 5:6:2:0:0 | 5 | 6 | 2 | 0 | 0 | 16 |
| 1266 | 1098 | 146 | 5:6:2:0:1 | 5 | 6 | 2 | 0 | 1 | 9 |
| 1267 | 1098 | 146 | 5:6:2:1:0 | 5 | 6 | 2 | 1 | 0 | 2 |
| 1268 | 1098 | 146 | 5:6:3:0:0 | 5 | 6 | 3 | 0 | 0 | 17 |
| 1269 | 1098 | 146 | 5:6:3:0:1 | 5 | 6 | 3 | 0 | 1 | 7 |
| 1270 | 1098 | 146 | 5:6:4:0:0 | 5 | 6 | 4 | 0 | 0 | 14 |
| 1271 | 1098 | 146 | 5:7:0:0:0 | 5 | 7 | 0 | 0 | 0 | 1 |
| 1272 | 1098 | 146 | 5:7:0:1:0 | 5 | 7 | 0 | 1 | 0 | 8 |
| 1273 | 1098 | 146 | 5:7:1:0:0 | 5 | 7 | 1 | 0 | 0 | 2 |
| 1274 | 1098 | 146 | 5:7:1:0:1 | 5 | 7 | 1 | 0 | 1 | 2 |
| 1275 | 1098 | 146 | 5:7:1:1:0 | 5 | 7 | 1 | 1 | 0 | 9 |
| 1276 | 1098 | 146 | 5:7:2:0:0 | 5 | 7 | 2 | 0 | 0 | 3 |
| 1277 | 1098 | 146 | 5:7:2:1:0 | 5 | 7 | 2 | 1 | 0 | 7 |
| 1278 | 1098 | 146 | 5:8:0:1:0 | 5 | 8 | 0 | 1 | 0 | 2 |
| 1279 | 1098 | 146 | 6:3:0:0:0 | 6 | 3 | 0 | 0 | 0 | 1 |
| 1280 | 1098 | 146 | 6:3:1:0:0 | 6 | 3 | 1 | 0 | 0 | 9 |
| 1281 | 1098 | 146 | 6:4:0:0:0 | 6 | 4 | 0 | 0 | 0 | 1 |
| 1282 | 1098 | 146 | 6:4:1:0:0 | 6 | 4 | 1 | 0 | 0 | 8 |
| 1283 | 1098 | 146 | 6:5:0:0:0 | 6 | 5 | 0 | 0 | 0 | 3 |
| 1284 | 1098 | 146 | 6:5:1:0:0 | 6 | 5 | 1 | 0 | 0 | 10 |
| 1285 | 1098 | 146 | 6:5:2:0:0 | 6 | 5 | 2 | 0 | 0 | 4 |
| 1286 | 1098 | 146 | 6:6:0:0:0 | 6 | 6 | 0 | 0 | 0 | 6 |
| 1287 | 1098 | 146 | 6:6:0:0:1 | 6 | 6 | 0 | 0 | 1 | 2 |
| 1288 | 1098 | 146 | 6:6:1:0:0 | 6 | 6 | 1 | 0 | 0 | 19 |
| 1289 | 1098 | 146 | 6:6:1:0:1 | 6 | 6 | 1 | 0 | 1 | 7 |
| 1290 | 1098 | 146 | 6:6:1:0:2 | 6 | 6 | 1 | 0 | 2 | 2 |
| 1291 | 1098 | 146 | 6:6:2:0:0 | 6 | 6 | 2 | 0 | 0 | 12 |
| 1292 | 1098 | 146 | 6:6:2:0:1 | 6 | 6 | 2 | 0 | 1 | 8 |
| 1293 | 1098 | 146 | 6:6:3:0:0 | 6 | 6 | 3 | 0 | 0 | 14 |

| | | | | | | | | | |
|---|---|---|---|---|---|---|---|---|---|
| 1294 | 1098 | 146 | 6:6:3:0:1 | 6 | 6 | 3 | 0 | 1 | 7 |
| 1295 | 1098 | 146 | 6:7:0:0:0 | 6 | 7 | 0 | 0 | 0 | 6 |
| 1296 | 1098 | 146 | 6:7:0:0:1 | 6 | 7 | 0 | 0 | 1 | 2 |
| 1297 | 1098 | 146 | 6:7:0:0:2 | 6 | 7 | 0 | 0 | 2 | 1 |
| 1298 | 1098 | 146 | 6:7:0:1:0 | 6 | 7 | 0 | 1 | 0 | 7 |
| 1299 | 1098 | 146 | 6:7:0:1:1 | 6 | 7 | 0 | 1 | 1 | 2 |
| 1300 | 1098 | 146 | 6:7:1:0:0 | 6 | 7 | 1 | 0 | 0 | 17 |
| 1301 | 1098 | 146 | 6:7:1:0:1 | 6 | 7 | 1 | 0 | 1 | 13 |
| 1302 | 1098 | 146 | 6:7:1:0:2 | 6 | 7 | 1 | 0 | 2 | 5 |
| 1303 | 1098 | 146 | 6:7:1:1:0 | 6 | 7 | 1 | 1 | 0 | 8 |
| 1304 | 1098 | 146 | 6:7:2:0:0 | 6 | 7 | 2 | 0 | 0 | 16 |
| 1305 | 1098 | 146 | 6:7:2:0:1 | 6 | 7 | 2 | 0 | 1 | 9 |
| 1306 | 1098 | 146 | 6:7:2:1:0 | 6 | 7 | 2 | 1 | 0 | 7 |
| 1307 | 1098 | 146 | 6:7:3:0:0 | 6 | 7 | 3 | 0 | 0 | 16 |
| 1308 | 1098 | 146 | 6:7:3:0:1 | 6 | 7 | 3 | 0 | 1 | 8 |
| 1309 | 1098 | 146 | 6:7:4:0:0 | 6 | 7 | 4 | 0 | 0 | 16 |
| 1310 | 1098 | 146 | 6:8:0:0:0 | 6 | 8 | 0 | 0 | 0 | 2 |
| 1311 | 1098 | 146 | 6:8:0:1:0 | 6 | 8 | 0 | 1 | 0 | 13 |
| 1312 | 1098 | 146 | 6:8:0:1:1 | 6 | 8 | 0 | 1 | 1 | 5 |
| 1313 | 1098 | 146 | 6:8:1:1:0 | 6 | 8 | 1 | 1 | 0 | 9 |
| 1314 | 1098 | 146 | 6:8:2:1:0 | 6 | 8 | 2 | 1 | 0 | 8 |
| 1315 | 1098 | 146 | 7:6:0:0:0 | 7 | 6 | 0 | 0 | 0 | 1 |
| 1316 | 1098 | 146 | 7:6:1:0:0 | 7 | 6 | 1 | 0 | 0 | 4 |
| 1317 | 1098 | 146 | 7:7:0:0:0 | 7 | 7 | 0 | 0 | 0 | 1 |
| 1318 | 1098 | 146 | 7:7:1:0:0 | 7 | 7 | 1 | 0 | 0 | 6 |
| 1319 | 1098 | 146 | 7:7:2:0:1 | 7 | 7 | 2 | 0 | 1 | 1 |
| 1320 | 1098 | 146 | 7:7:2:0:2 | 7 | 7 | 2 | 0 | 2 | 1 |
| 1321 | 1098 | 146 | 7:8:0:0:0 | 7 | 8 | 0 | 0 | 0 | 2 |
| 1322 | 1098 | 146 | 7:8:0:0:1 | 7 | 8 | 0 | 0 | 1 | 1 |
| 1323 | 1098 | 146 | 7:8:1:0:0 | 7 | 8 | 1 | 0 | 0 | 4 |
| 1324 | 1098 | 146 | 7:8:1:0:1 | 7 | 8 | 1 | 0 | 1 | 2 |
| 1325 | 1098 | 146 | 7:8:1:1:0 | 7 | 8 | 1 | 1 | 0 | 1 |
| 1326 | 1098 | 146 | 7:8:1:1:1 | 7 | 8 | 1 | 1 | 1 | 1 |
| 1327 | 1098 | 146 | 7:8:2:0:0 | 7 | 8 | 2 | 0 | 0 | 2 |
| 1328 | 1098 | 146 | 7:9:0:1:0 | 7 | 9 | 0 | 1 | 0 | 2 |
| 1329 | 1098 | 146 | 7:9:0:2:0 | 7 | 9 | 0 | 2 | 0 | 1 |
| 1330 | 1134 | 38 | 2:5:0:0:0 | 2 | 5 | 0 | 0 | 0 | 13 |
| 1331 | 1134 | 38 | 2:6:0:0:0 | 2 | 6 | 0 | 0 | 0 | 13 |
| 1332 | 1134 | 38 | 2:7:0:0:0 | 2 | 7 | 0 | 0 | 0 | 11 |
| 1333 | 1134 | 38 | 3:4:1:0:0 | 3 | 4 | 1 | 0 | 0 | 8 |
| 1334 | 1134 | 38 | 3:5:0:0:0 | 3 | 5 | 0 | 0 | 0 | 11 |
| 1335 | 1134 | 38 | 3:5:0:0:1 | 3 | 5 | 0 | 0 | 1 | 6 |
| 1336 | 1134 | 38 | 3:5:1:0:0 | 3 | 5 | 1 | 0 | 0 | 14 |
| 1337 | 1134 | 38 | 3:5:2:0:0 | 3 | 5 | 2 | 0 | 0 | 13 |
| 1338 | 1134 | 38 | 3:6:0:0:0 | 3 | 6 | 0 | 0 | 0 | 11 |
| 1339 | 1134 | 38 | 3:6:1:0:0 | 3 | 6 | 1 | 0 | 0 | 13 |
| 1340 | 1134 | 38 | 4:3:1:0:0 | 4 | 3 | 1 | 0 | 0 | 14 |
| 1341 | 1134 | 38 | 4:4:1:0:0 | 4 | 4 | 1 | 0 | 0 | 12 |
| 1342 | 1134 | 38 | 4:5:0:0:0 | 4 | 5 | 0 | 0 | 0 | 14 |
| 1343 | 1134 | 38 | 4:5:1:0:0 | 4 | 5 | 1 | 0 | 0 | 19 |
| 1344 | 1134 | 38 | 4:5:1:0:1 | 4 | 5 | 1 | 0 | 1 | 14 |
| 1345 | 1134 | 38 | 4:5:2:0:0 | 4 | 5 | 2 | 0 | 0 | 18 |
| 1346 | 1134 | 38 | 4:5:3:0:0 | 4 | 5 | 3 | 0 | 0 | 17 |
| 1347 | 1134 | 38 | 4:6:0:1:0 | 4 | 6 | 0 | 1 | 0 | 14 |
| 1348 | 1134 | 38 | 5:3:1:0:0 | 5 | 3 | 1 | 0 | 0 | 11 |
| 1349 | 1134 | 38 | 5:4:1:0:0 | 5 | 4 | 1 | 0 | 0 | 16 |
| 1350 | 1134 | 38 | 5:5:1:0:0 | 5 | 5 | 1 | 0 | 0 | 18 |
| 1351 | 1134 | 38 | 5:5:2:0:0 | 5 | 5 | 2 | 0 | 0 | 15 |
| 1352 | 1134 | 38 | 5:6:1:0:0 | 5 | 6 | 1 | 0 | 0 | 16 |

| | | | | | | | | | |
|---|---|---|---|---|---|---|---|---|---|
| 1353 | 1134 | 38 | 5:6:2:0:0 | 5 | 6 | 2 | 0 | 0 | 16 |
| 1354 | 1134 | 38 | 5:6:2:0:1 | 5 | 6 | 2 | 0 | 1 | 9 |
| 1355 | 1134 | 38 | 5:6:3:0:0 | 5 | 6 | 3 | 0 | 0 | 17 |
| 1356 | 1134 | 38 | 5:7:1:1:0 | 5 | 7 | 1 | 1 | 0 | 9 |
| 1357 | 1134 | 38 | 6:3:1:0:0 | 6 | 3 | 1 | 0 | 0 | 9 |
| 1358 | 1134 | 38 | 6:4:1:0:0 | 6 | 4 | 1 | 0 | 0 | 8 |
| 1359 | 1134 | 38 | 6:6:1:0:0 | 6 | 6 | 1 | 0 | 0 | 19 |
| 1360 | 1134 | 38 | 6:6:3:0:0 | 6 | 6 | 3 | 0 | 0 | 14 |
| 1361 | 1134 | 38 | 6:6:3:0:1 | 6 | 6 | 3 | 0 | 1 | 7 |
| 1362 | 1134 | 38 | 6:7:1:0:0 | 6 | 7 | 1 | 0 | 0 | 17 |
| 1363 | 1134 | 38 | 6:7:1:0:1 | 6 | 7 | 1 | 0 | 1 | 13 |
| 1364 | 1134 | 38 | 6:7:2:0:0 | 6 | 7 | 2 | 0 | 0 | 16 |
| 1365 | 1134 | 38 | 6:7:2:1:0 | 6 | 7 | 2 | 1 | 0 | 7 |
| 1366 | 1134 | 38 | 6:7:3:0:0 | 6 | 7 | 3 | 0 | 0 | 16 |
| 1367 | 1134 | 38 | 6:8:0:1:0 | 6 | 8 | 0 | 1 | 0 | 13 |
| 1368 | 1158 | 25 | 4:7:4:2:0 | 4 | 7 | 4 | 2 | 0 | 1 |
| 1369 | 1158 | 25 | 5:6:1:0:0 | 5 | 6 | 1 | 0 | 0 | 16 |
| 1370 | 1158 | 25 | 6:6:1:0:0 | 6 | 6 | 1 | 0 | 0 | 19 |
| 1371 | 1158 | 25 | 6:6:2:0:1 | 6 | 6 | 2 | 0 | 1 | 8 |
| 1372 | 1158 | 25 | 6:7:1:0:0 | 6 | 7 | 1 | 0 | 0 | 17 |
| 1373 | 1158 | 25 | 6:7:1:0:1 | 6 | 7 | 1 | 0 | 1 | 13 |
| 1374 | 1158 | 25 | 6:7:1:0:2 | 6 | 7 | 1 | 0 | 2 | 5 |
| 1375 | 1158 | 25 | 6:7:1:1:0 | 6 | 7 | 1 | 1 | 0 | 8 |
| 1376 | 1158 | 25 | 6:7:2:0:0 | 6 | 7 | 2 | 0 | 0 | 16 |
| 1377 | 1158 | 25 | 6:7:2:0:1 | 6 | 7 | 2 | 0 | 1 | 9 |
| 1378 | 1158 | 25 | 6:7:3:0:0 | 6 | 7 | 3 | 0 | 0 | 16 |
| 1379 | 1158 | 25 | 6:7:3:0:1 | 6 | 7 | 3 | 0 | 1 | 8 |
| 1380 | 1158 | 25 | 6:7:3:0:2 | 6 | 7 | 3 | 0 | 2 | 4 |
| 1381 | 1158 | 25 | 6:7:4:0:0 | 6 | 7 | 4 | 0 | 0 | 16 |
| 1382 | 1158 | 25 | 6:8:0:1:0 | 6 | 8 | 0 | 1 | 0 | 13 |
| 1383 | 1158 | 25 | 6:8:0:1:1 | 6 | 8 | 0 | 1 | 1 | 5 |
| 1384 | 1158 | 25 | 6:8:1:1:0 | 6 | 8 | 1 | 1 | 0 | 9 |
| 1385 | 1158 | 25 | 6:8:2:1:0 | 6 | 8 | 2 | 1 | 0 | 8 |
| 1386 | 1158 | 25 | 6:8:2:1:1 | 6 | 8 | 2 | 1 | 1 | 4 |
| 1387 | 1158 | 25 | 6:9:1:2:0 | 6 | 9 | 1 | 2 | 0 | 4 |
| 1388 | 1158 | 25 | 7:8:1:0:0 | 7 | 8 | 1 | 0 | 0 | 4 |
| 1389 | 1158 | 25 | 7:8:1:0:1 | 7 | 8 | 1 | 0 | 1 | 2 |
| 1390 | 1158 | 25 | 7:8:3:0:0 | 7 | 8 | 3 | 0 | 0 | 1 |
| 1391 | 1158 | 25 | 7:9:0:1:0 | 7 | 9 | 0 | 1 | 0 | 2 |
| 1392 | 1173 | 14 | 5:6:1:0:0 | 5 | 6 | 1 | 0 | 0 | 16 |
| 1393 | 1173 | 14 | 6:6:2:0:2 | 6 | 6 | 2 | 0 | 2 | 4 |
| 1394 | 1173 | 14 | 6:6:3:0:1 | 6 | 6 | 3 | 0 | 1 | 7 |
| 1395 | 1173 | 14 | 6:7:1:0:0 | 6 | 7 | 1 | 0 | 0 | 17 |
| 1396 | 1173 | 14 | 6:7:1:0:1 | 6 | 7 | 1 | 0 | 1 | 13 |
| 1397 | 1173 | 14 | 6:7:1:0:2 | 6 | 7 | 1 | 0 | 2 | 5 |
| 1398 | 1173 | 14 | 6:7:1:1:1 | 6 | 7 | 1 | 1 | 1 | 4 |
| 1399 | 1173 | 14 | 6:7:2:0:0 | 6 | 7 | 2 | 0 | 0 | 16 |
| 1400 | 1173 | 14 | 6:7:2:1:0 | 6 | 7 | 2 | 1 | 0 | 7 |
| 1401 | 1173 | 14 | 6:7:3:0:0 | 6 | 7 | 3 | 0 | 0 | 16 |
| 1402 | 1173 | 14 | 6:7:4:0:0 | 6 | 7 | 4 | 0 | 0 | 16 |
| 1403 | 1173 | 14 | 6:8:0:1:0 | 6 | 8 | 0 | 1 | 0 | 13 |
| 1404 | 1173 | 14 | 6:8:0:1:1 | 6 | 8 | 0 | 1 | 1 | 5 |
| 1405 | 1173 | 14 | 6:8:0:2:0 | 6 | 8 | 0 | 2 | 0 | 4 |
| 1406 | 1194 | 44 | 4:3:1:0:0 | 4 | 3 | 1 | 0 | 0 | 14 |
| 1407 | 1194 | 44 | 4:3:1:0:1 | 4 | 3 | 1 | 0 | 1 | 1 |
| 1408 | 1194 | 44 | 4:4:0:0:0 | 4 | 4 | 0 | 0 | 0 | 6 |
| 1409 | 1194 | 44 | 4:4:0:1:0 | 4 | 4 | 0 | 1 | 0 | 1 |
| 1410 | 1194 | 44 | 4:5:0:0:0 | 4 | 5 | 0 | 0 | 0 | 14 |
| 1411 | 1194 | 44 | 4:5:1:0:0 | 4 | 5 | 1 | 0 | 0 | 19 |

| | | | | | | | | | |
|---|---|---|---|---|---|---|---|---|---|
| 1412 | 1194 | 44 | 4:5:2:0:0 | 4 | 5 | 2 | 0 | 0 | 18 |
| 1413 | 1194 | 44 | 4:5:3:0:0 | 4 | 5 | 3 | 0 | 0 | 17 |
| 1414 | 1194 | 44 | 5:3:1:0:0 | 5 | 3 | 1 | 0 | 0 | 11 |
| 1415 | 1194 | 44 | 5:4:1:0:0 | 5 | 4 | 1 | 0 | 0 | 16 |
| 1416 | 1194 | 44 | 5:5:1:0:0 | 5 | 5 | 1 | 0 | 0 | 18 |
| 1417 | 1194 | 44 | 5:6:0:0:0 | 5 | 6 | 0 | 0 | 0 | 9 |
| 1418 | 1194 | 44 | 5:6:1:0:0 | 5 | 6 | 1 | 0 | 0 | 16 |
| 1419 | 1194 | 44 | 5:6:1:0:1 | 5 | 6 | 1 | 0 | 1 | 8 |
| 1420 | 1194 | 44 | 5:6:2:0:0 | 5 | 6 | 2 | 0 | 0 | 16 |
| 1421 | 1194 | 44 | 5:6:2:0:1 | 5 | 6 | 2 | 0 | 1 | 9 |
| 1422 | 1194 | 44 | 5:6:3:0:0 | 5 | 6 | 3 | 0 | 0 | 17 |
| 1423 | 1194 | 44 | 5:6:4:0:0 | 5 | 6 | 4 | 0 | 0 | 14 |
| 1424 | 1194 | 44 | 5:7:0:1:0 | 5 | 7 | 0 | 1 | 0 | 8 |
| 1425 | 1194 | 44 | 5:7:1:1:0 | 5 | 7 | 1 | 1 | 0 | 9 |
| 1426 | 1194 | 44 | 6:3:1:0:0 | 6 | 3 | 1 | 0 | 0 | 9 |
| 1427 | 1194 | 44 | 6:4:1:0:0 | 6 | 4 | 1 | 0 | 0 | 8 |
| 1428 | 1194 | 44 | 6:5:1:0:0 | 6 | 5 | 1 | 0 | 0 | 10 |
| 1429 | 1194 | 44 | 6:6:1:0:0 | 6 | 6 | 1 | 0 | 0 | 19 |
| 1430 | 1194 | 44 | 6:6:2:0:1 | 6 | 6 | 2 | 0 | 1 | 8 |
| 1431 | 1194 | 44 | 6:7:0:0:0 | 6 | 7 | 0 | 0 | 0 | 6 |
| 1432 | 1194 | 44 | 6:7:1:0:0 | 6 | 7 | 1 | 0 | 0 | 17 |
| 1433 | 1194 | 44 | 6:7:1:0:1 | 6 | 7 | 1 | 0 | 1 | 13 |
| 1434 | 1194 | 44 | 6:7:1:1:0 | 6 | 7 | 1 | 1 | 0 | 8 |
| 1435 | 1194 | 44 | 6:7:2:0:0 | 6 | 7 | 2 | 0 | 0 | 16 |
| 1436 | 1194 | 44 | 6:7:2:0:1 | 6 | 7 | 2 | 0 | 1 | 9 |
| 1437 | 1194 | 44 | 6:7:2:0:2 | 6 | 7 | 2 | 0 | 2 | 5 |
| 1438 | 1194 | 44 | 6:7:3:0:0 | 6 | 7 | 3 | 0 | 0 | 16 |
| 1439 | 1194 | 44 | 6:7:3:0:1 | 6 | 7 | 3 | 0 | 1 | 8 |
| 1440 | 1194 | 44 | 6:7:3:0:2 | 6 | 7 | 3 | 0 | 2 | 4 |
| 1441 | 1194 | 44 | 6:7:4:0:0 | 6 | 7 | 4 | 0 | 0 | 16 |
| 1442 | 1194 | 44 | 6:8:0:1:0 | 6 | 8 | 0 | 1 | 0 | 13 |
| 1443 | 1194 | 44 | 6:8:1:1:0 | 6 | 8 | 1 | 1 | 0 | 9 |
| 1444 | 1194 | 44 | 6:8:1:1:1 | 6 | 8 | 1 | 1 | 1 | 5 |
| 1445 | 1194 | 44 | 6:8:2:1:0 | 6 | 8 | 2 | 1 | 0 | 8 |
| 1446 | 1194 | 44 | 6:8:2:1:1 | 6 | 8 | 2 | 1 | 1 | 4 |
| 1447 | 1194 | 44 | 6:9:0:2:0 | 6 | 9 | 0 | 2 | 0 | 5 |
| 1448 | 1194 | 44 | 6:9:1:2:0 | 6 | 9 | 1 | 2 | 0 | 4 |
| 1449 | 1194 | 44 | 7:8:1:0:0 | 7 | 8 | 1 | 0 | 0 | 4 |

Supplementary Table S12. Total site-specific glycan compositions of SARS CoV-2 Spike protein (by GR + Byonic)

| No. | Glycosylation site | Total No. of composition / site | Glycan composition | No. of HexNAc | No. of Hex | No. of Fuc | No. of NeuAc | No. of NeuGc | Assigned by GR | Assigned by Byonic |
|---|---|---|---|---|---|---|---|---|---|---|
| 1 | 17 | 54 | 2:7:0:0:0 | 2 | 7 | 0 | 0 | 0 | | 1 |
| 2 | 17 | 54 | 3:5:1:0:0 | 3 | 5 | 1 | 0 | 0 | 1 | |
| 3 | 17 | 54 | 3:5:2:0:0 | 3 | 5 | 2 | 0 | 0 | 1 | |
| 4 | 17 | 54 | 3:6:0:0:0 | 3 | 6 | 0 | 0 | 0 | 1 | |
| 5 | 17 | 54 | 3:6:1:0:0 | 3 | 6 | 1 | 0 | 0 | 1 | 1 |
| 6 | 17 | 54 | 3:6:2:0:0 | 3 | 6 | 2 | 0 | 0 | 1 | |
| 7 | 17 | 54 | 4:3:0:1:0 | 4 | 3 | 0 | 1 | 0 | | 1 |
| 8 | 17 | 54 | 4:3:1:0:0 | 4 | 3 | 1 | 0 | 0 | 1 | 1 |
| 9 | 17 | 54 | 4:4:1:0:0 | 4 | 4 | 1 | 0 | 0 | 1 | |
| 10 | 17 | 54 | 4:4:1:0:1 | 4 | 4 | 1 | 0 | 1 | 1 | |
| 11 | 17 | 54 | 4:4:2:0:0 | 4 | 4 | 2 | 0 | 0 | 1 | 1 |
| 12 | 17 | 54 | 4:5:0:1:0 | 4 | 5 | 0 | 1 | 0 | 1 | |
| 13 | 17 | 54 | 4:5:1:0:0 | 4 | 5 | 1 | 0 | 0 | 1 | 1 |
| 14 | 17 | 54 | 4:5:1:0:1 | 4 | 5 | 1 | 0 | 1 | 1 | |
| 15 | 17 | 54 | 4:5:2:0:0 | 4 | 5 | 2 | 0 | 0 | 1 | 1 |
| 16 | 17 | 54 | 4:5:2:0:1 | 4 | 5 | 2 | 0 | 1 | 1 | |
| 17 | 17 | 54 | 4:5:3:0:0 | 4 | 5 | 3 | 0 | 0 | 1 | 1 |
| 18 | 17 | 54 | 4:6:0:1:0 | 4 | 6 | 0 | 1 | 0 | 1 | |
| 19 | 17 | 54 | 4:6:1:1:0 | 4 | 6 | 1 | 1 | 0 | 1 | |
| 20 | 17 | 54 | 5:3:1:0:0 | 5 | 3 | 1 | 0 | 0 | 1 | |
| 21 | 17 | 54 | 5:4:1:0:0 | 5 | 4 | 1 | 0 | 0 | 1 | 1 |
| 22 | 17 | 54 | 5:4:1:0:1 | 5 | 4 | 1 | 0 | 1 | 1 | |
| 23 | 17 | 54 | 5:4:2:0:0 | 5 | 4 | 2 | 0 | 0 | 1 | 1 |
| 24 | 17 | 54 | 5:4:2:0:1 | 5 | 4 | 2 | 0 | 1 | 1 | |
| 25 | 17 | 54 | 5:4:3:0:0 | 5 | 4 | 3 | 0 | 0 | 1 | |
| 26 | 17 | 54 | 5:5:0:1:0 | 5 | 5 | 0 | 1 | 0 | 1 | |
| 27 | 17 | 54 | 5:5:1:0:0 | 5 | 5 | 1 | 0 | 0 | 1 | 1 |
| 28 | 17 | 54 | 5:5:1:1:0 | 5 | 5 | 1 | 1 | 0 | 1 | |
| 29 | 17 | 54 | 5:5:2:0:0 | 5 | 5 | 2 | 0 | 0 | 1 | 1 |
| 30 | 17 | 54 | 5:5:3:0:0 | 5 | 5 | 3 | 0 | 0 | 1 | 1 |
| 31 | 17 | 54 | 5:6:0:3:0 | 5 | 6 | 0 | 3 | 0 | | 1 |
| 32 | 17 | 54 | 5:6:1:0:0 | 5 | 6 | 1 | 0 | 0 | 1 | |
| 33 | 17 | 54 | 5:6:1:0:1 | 5 | 6 | 1 | 0 | 1 | 1 | |
| 34 | 17 | 54 | 5:6:2:0:0 | 5 | 6 | 2 | 0 | 0 | | 1 |
| 35 | 17 | 54 | 5:6:2:0:1 | 5 | 6 | 2 | 0 | 1 | 1 | |
| 36 | 17 | 54 | 5:6:3:0:0 | 5 | 6 | 3 | 0 | 0 | 1 | 1 |
| 37 | 17 | 54 | 5:6:4:0:0 | 5 | 6 | 4 | 0 | 0 | 1 | |
| 38 | 17 | 54 | 5:7:0:1:0 | 5 | 7 | 0 | 1 | 0 | 1 | |
| 39 | 17 | 54 | 5:7:1:1:0 | 5 | 7 | 1 | 1 | 0 | 1 | |
| 40 | 17 | 54 | 6:3:1:0:0 | 6 | 3 | 1 | 0 | 0 | 1 | |
| 41 | 17 | 54 | 6:4:2:0:0 | 6 | 4 | 2 | 0 | 0 | 1 | |
| 42 | 17 | 54 | 6:5:4:0:0 | 6 | 5 | 4 | 0 | 0 | 1 | |
| 43 | 17 | 54 | 6:6:1:0:0 | 6 | 6 | 1 | 0 | 0 | 1 | |
| 44 | 17 | 54 | 6:6:1:0:1 | 6 | 6 | 1 | 0 | 1 | 1 | |
| 45 | 17 | 54 | 6:6:2:0:0 | 6 | 6 | 2 | 0 | 0 | 1 | |
| 46 | 17 | 54 | 6:6:3:0:0 | 6 | 6 | 3 | 0 | 0 | 1 | 1 |
| 47 | 17 | 54 | 6:6:4:0:0 | 6 | 6 | 4 | 0 | 0 | 1 | |
| 48 | 17 | 54 | 6:7:0:1:0 | 6 | 7 | 0 | 1 | 0 | 1 | |
| 49 | 17 | 54 | 6:7:1:0:0 | 6 | 7 | 1 | 0 | 0 | 1 | |
| 50 | 17 | 54 | 6:7:4:0:0 | 6 | 7 | 4 | 0 | 0 | | 1 |
| 51 | 17 | 54 | 6:7:5:0:0 | 6 | 7 | 5 | 0 | 0 | | 1 |
| 52 | 17 | 54 | 7:6:2:0:0 | 7 | 6 | 2 | 0 | 0 | 1 | |
| 53 | 17 | 54 | 7:8:1:1:0 | 7 | 8 | 1 | 1 | 0 | | 1 |
| 54 | 17 | 54 | 8:7:0:0:0 | 8 | 7 | 0 | 0 | 0 | | 1 |
| 55 | 61 | 54 | 2:5:0:0:0 | 2 | 5 | 0 | 0 | 0 | 1 | 1 |
| 56 | 61 | 54 | 2:6:0:0:0 | 2 | 6 | 0 | 0 | 0 | 1 | 1 |
| 57 | 61 | 54 | 2:7:0:0:0 | 2 | 7 | 0 | 0 | 0 | 1 | 1 |
| 58 | 61 | 54 | 2:8:0:0:0 | 2 | 8 | 0 | 0 | 0 | 1 | 1 |
| 59 | 61 | 54 | 3:4:1:0:0 | 3 | 4 | 1 | 0 | 0 | 1 | 1 |
| 60 | 61 | 54 | 3:5:0:0:0 | 3 | 5 | 0 | 0 | 0 | 1 | 1 |
| 61 | 61 | 54 | 3:5:0:0:1 | 3 | 5 | 0 | 0 | 1 | 1 | |

| | | | | | | | | | | |
|---|---|---|---|---|---|---|---|---|---|---|
| 62 | 61 | 54 | 3:5:1:0:0 | 3 | 5 | 1 | 0 | 0 | 1 | 1 |
| 63 | 61 | 54 | 3:5:2:0:0 | 3 | 5 | 2 | 0 | 0 | 1 | |
| 64 | 61 | 54 | 3:6:0:0:0 | 3 | 6 | 0 | 0 | 0 | | 1 |
| 65 | 61 | 54 | 3:6:0:0:1 | 3 | 6 | 0 | 0 | 1 | 1 | 1 |
| 66 | 61 | 54 | 3:6:1:0:0 | 3 | 6 | 1 | 0 | 0 | 1 | 1 |
| 67 | 61 | 54 | 4:3:3:0:0 | 4 | 3 | 3 | 0 | 0 | | 1 |
| 68 | 61 | 54 | 4:4:1:0:0 | 4 | 4 | 1 | 0 | 0 | | 1 |
| 69 | 61 | 54 | 4:5:0:0:0 | 4 | 5 | 0 | 0 | 0 | 1 | 1 |
| 70 | 61 | 54 | 4:5:0:0:1 | 4 | 5 | 0 | 0 | 1 | 1 | |
| 71 | 61 | 54 | 4:5:1:0:0 | 4 | 5 | 1 | 0 | 0 | 1 | 1 |
| 72 | 61 | 54 | 4:5:1:0:1 | 4 | 5 | 1 | 0 | 1 | 1 | |
| 73 | 61 | 54 | 4:5:2:0:0 | 4 | 5 | 2 | 0 | 0 | 1 | 1 |
| 74 | 61 | 54 | 4:5:3:0:0 | 4 | 5 | 3 | 0 | 0 | 1 | 1 |
| 75 | 61 | 54 | 4:6:0:1:0 | 4 | 6 | 0 | 1 | 0 | 1 | 1 |
| 76 | 61 | 54 | 4:6:1:0:0 | 4 | 6 | 1 | 0 | 0 | 1 | 1 |
| 77 | 61 | 54 | 4:6:2:0:0 | 4 | 6 | 2 | 0 | 0 | 1 | 1 |
| 78 | 61 | 54 | 4:7:1:0:0 | 4 | 7 | 1 | 0 | 0 | 1 | 1 |
| 79 | 61 | 54 | 4:7:2:0:0 | 4 | 7 | 2 | 0 | 0 | 1 | 1 |
| 80 | 61 | 54 | 5:4:2:0:0 | 5 | 4 | 2 | 0 | 0 | | 1 |
| 81 | 61 | 54 | 5:5:0:0:0 | 5 | 5 | 0 | 0 | 0 | | 1 |
| 82 | 61 | 54 | 5:5:1:0:0 | 5 | 5 | 1 | 0 | 0 | 1 | 1 |
| 83 | 61 | 54 | 5:5:2:0:0 | 5 | 5 | 2 | 0 | 0 | 1 | 1 |
| 84 | 61 | 54 | 5:5:3:0:0 | 5 | 5 | 3 | 0 | 0 | 1 | 1 |
| 85 | 61 | 54 | 5:6:0:0:0 | 5 | 6 | 0 | 0 | 0 | 1 | 1 |
| 86 | 61 | 54 | 5:6:1:0:0 | 5 | 6 | 1 | 0 | 0 | | 1 |
| 87 | 61 | 54 | 5:6:2:0:0 | 5 | 6 | 2 | 0 | 0 | 1 | 1 |
| 88 | 61 | 54 | 5:6:2:0:1 | 5 | 6 | 2 | 0 | 1 | 1 | |
| 89 | 61 | 54 | 5:6:3:0:0 | 5 | 6 | 3 | 0 | 0 | 1 | 1 |
| 90 | 61 | 54 | 5:6:4:0:0 | 5 | 6 | 4 | 0 | 0 | 1 | |
| 91 | 61 | 54 | 5:7:1:1:0 | 5 | 7 | 1 | 1 | 0 | 1 | |
| 92 | 61 | 54 | 5:7:2:0:0 | 5 | 7 | 2 | 0 | 0 | 1 | |
| 93 | 61 | 54 | 5:7:3:0:0 | 5 | 7 | 3 | 0 | 0 | 1 | |
| 94 | 61 | 54 | 6:4:0:0:0 | 6 | 4 | 0 | 0 | 0 | | 1 |
| 95 | 61 | 54 | 6:6:0:0:0 | 6 | 6 | 0 | 0 | 0 | 1 | 1 |
| 96 | 61 | 54 | 6:6:1:0:0 | 6 | 6 | 1 | 0 | 0 | 1 | 1 |
| 97 | 61 | 54 | 6:6:1:1:0 | 6 | 6 | 1 | 1 | 0 | | 1 |
| 98 | 61 | 54 | 6:6:2:0:0 | 6 | 6 | 2 | 0 | 0 | 1 | 1 |
| 99 | 61 | 54 | 6:6:2:0:1 | 6 | 6 | 2 | 0 | 1 | 1 | |
| 100 | 61 | 54 | 6:6:3:0:0 | 6 | 6 | 3 | 0 | 0 | 1 | 1 |
| 101 | 61 | 54 | 6:6:4:0:0 | 6 | 6 | 4 | 0 | 0 | 1 | |
| 102 | 61 | 54 | 6:7:1:1:0 | 6 | 7 | 1 | 1 | 0 | 1 | |
| 103 | 61 | 54 | 6:7:4:0:0 | 6 | 7 | 4 | 0 | 0 | 1 | |
| 104 | 61 | 54 | 6:8:1:1:0 | 6 | 8 | 1 | 1 | 0 | | 1 |
| 105 | 61 | 54 | 8:3:0:0:0 | 8 | 3 | 0 | 0 | 0 | | 1 |
| 106 | 61 | 54 | 8:5:0:0:0 | 8 | 5 | 0 | 0 | 0 | | 1 |
| 107 | 61 | 54 | 8:9:0:0:0 | 8 | 9 | 0 | 0 | 0 | 1 | |
| 108 | 61 | 54 | 9:3:1:0:0 | 9 | 3 | 1 | 0 | 0 | | 1 |
| 109 | 74 | 237 | 2:8:0:0:0 | 2 | 8 | 0 | 0 | 0 | | 1 |
| 110 | 74 | 237 | 3:3:1:0:1 | 3 | 3 | 1 | 0 | 1 | 1 | |
| 111 | 74 | 237 | 3:3:1:1:0 | 3 | 3 | 1 | 1 | 0 | 1 | |
| 112 | 74 | 237 | 3:3:2:0:1 | 3 | 3 | 2 | 0 | 1 | 1 | |
| 113 | 74 | 237 | 3:4:0:1:0 | 3 | 4 | 0 | 1 | 0 | 1 | |
| 114 | 74 | 237 | 3:4:1:0:0 | 3 | 4 | 1 | 0 | 0 | 1 | |
| 115 | 74 | 237 | 3:4:1:1:0 | 3 | 4 | 1 | 1 | 0 | 1 | |
| 116 | 74 | 237 | 3:4:2:0:0 | 3 | 4 | 2 | 0 | 0 | 1 | |
| 117 | 74 | 237 | 3:4:2:0:1 | 3 | 4 | 2 | 0 | 1 | 1 | |
| 118 | 74 | 237 | 3:5:1:1:0 | 3 | 5 | 1 | 1 | 0 | 1 | |
| 119 | 74 | 237 | 3:5:2:0:0 | 3 | 5 | 2 | 0 | 0 | 1 | |
| 120 | 74 | 237 | 3:5:2:0:1 | 3 | 5 | 2 | 0 | 1 | 1 | |
| 121 | 74 | 237 | 3:6:1:1:0 | 3 | 6 | 1 | 1 | 0 | 1 | |
| 122 | 74 | 237 | 3:6:2:0:0 | 3 | 6 | 2 | 0 | 0 | 1 | |
| 123 | 74 | 237 | 3:6:2:0:1 | 3 | 6 | 2 | 0 | 1 | 1 | |
| 124 | 74 | 237 | 4:3:0:0:0 | 4 | 3 | 0 | 0 | 0 | 1 | |
| 125 | 74 | 237 | 4:3:1:0:0 | 4 | 3 | 1 | 0 | 0 | 1 | 1 |
| 126 | 74 | 237 | 4:3:1:1:0 | 4 | 3 | 1 | 1 | 0 | 1 | |

| | | | | | | | | | | |
|---|---|---|---|---|---|---|---|---|---|---|
| 127 | 74 | 237 | 4:3:2:0:0 | 4 | 3 | 2 | 0 | 0 | 1 | 1 |
| 128 | 74 | 237 | 4:3:2:0:1 | 4 | 3 | 2 | 0 | 1 | 1 | |
| 129 | 74 | 237 | 4:4:0:2:0 | 4 | 4 | 0 | 2 | 0 | 1 | |
| 130 | 74 | 237 | 4:4:1:0:0 | 4 | 4 | 1 | 0 | 0 | 1 | 1 |
| 131 | 74 | 237 | 4:4:1:0:1 | 4 | 4 | 1 | 0 | 1 | 1 | 1 |
| 132 | 74 | 237 | 4:4:1:1:0 | 4 | 4 | 1 | 1 | 0 | 1 | |
| 133 | 74 | 237 | 4:4:1:1:1 | 4 | 4 | 1 | 1 | 1 | 1 | |
| 134 | 74 | 237 | 4:4:1:2:0 | 4 | 4 | 1 | 2 | 0 | 1 | |
| 135 | 74 | 237 | 4:4:2:0:0 | 4 | 4 | 2 | 0 | 0 | 1 | 1 |
| 136 | 74 | 237 | 4:4:2:0:1 | 4 | 4 | 2 | 0 | 1 | 1 | |
| 137 | 74 | 237 | 4:4:3:0:1 | 4 | 4 | 3 | 0 | 1 | 1 | |
| 138 | 74 | 237 | 4:5:0:0:0 | 4 | 5 | 0 | 0 | 0 | 1 | |
| 139 | 74 | 237 | 4:5:0:1:0 | 4 | 5 | 0 | 1 | 0 | 1 | |
| 140 | 74 | 237 | 4:5:0:2:0 | 4 | 5 | 0 | 2 | 0 | 1 | |
| 141 | 74 | 237 | 4:5:1:0:0 | 4 | 5 | 1 | 0 | 0 | 1 | 1 |
| 142 | 74 | 237 | 4:5:1:0:1 | 4 | 5 | 1 | 0 | 1 | 1 | 1 |
| 143 | 74 | 237 | 4:5:1:0:2 | 4 | 5 | 1 | 0 | 2 | 1 | 1 |
| 144 | 74 | 237 | 4:5:1:1:0 | 4 | 5 | 1 | 1 | 0 | 1 | 1 |
| 145 | 74 | 237 | 4:5:2:0:0 | 4 | 5 | 2 | 0 | 0 | 1 | 1 |
| 146 | 74 | 237 | 4:5:2:0:1 | 4 | 5 | 2 | 0 | 1 | 1 | 1 |
| 147 | 74 | 237 | 4:5:2:1:0 | 4 | 5 | 2 | 1 | 0 | 1 | |
| 148 | 74 | 237 | 4:5:3:0:0 | 4 | 5 | 3 | 0 | 0 | 1 | 1 |
| 149 | 74 | 237 | 4:5:3:0:1 | 4 | 5 | 3 | 0 | 1 | 1 | |
| 150 | 74 | 237 | 4:5:3:2:0 | 4 | 5 | 3 | 2 | 0 | | 1 |
| 151 | 74 | 237 | 4:6:0:1:0 | 4 | 6 | 0 | 1 | 0 | 1 | |
| 152 | 74 | 237 | 4:6:0:1:1 | 4 | 6 | 0 | 1 | 1 | 1 | |
| 153 | 74 | 237 | 4:6:1:1:0 | 4 | 6 | 1 | 1 | 0 | 1 | 1 |
| 154 | 74 | 237 | 4:6:2:0:0 | 4 | 6 | 2 | 0 | 0 | 1 | |
| 155 | 74 | 237 | 4:6:2:1:0 | 4 | 6 | 2 | 1 | 0 | 1 | |
| 156 | 74 | 237 | 4:6:3:0:0 | 4 | 6 | 3 | 0 | 0 | 1 | 1 |
| 157 | 74 | 237 | 4:7:0:1:0 | 4 | 7 | 0 | 1 | 0 | | 1 |
| 158 | 74 | 237 | 5:3:0:0:0 | 5 | 3 | 0 | 0 | 0 | 1 | |
| 159 | 74 | 237 | 5:3:1:0:0 | 5 | 3 | 1 | 0 | 0 | 1 | 1 |
| 160 | 74 | 237 | 5:3:1:1:0 | 5 | 3 | 1 | 1 | 0 | 1 | |
| 161 | 74 | 237 | 5:3:2:0:0 | 5 | 3 | 2 | 0 | 0 | 1 | |
| 162 | 74 | 237 | 5:3:2:0:1 | 5 | 3 | 2 | 0 | 1 | 1 | |
| 163 | 74 | 237 | 5:3:3:0:1 | 5 | 3 | 3 | 0 | 1 | 1 | |
| 164 | 74 | 237 | 5:4:0:0:1 | 5 | 4 | 0 | 0 | 1 | 1 | |
| 165 | 74 | 237 | 5:4:0:2:0 | 5 | 4 | 0 | 2 | 0 | 1 | |
| 166 | 74 | 237 | 5:4:1:0:0 | 5 | 4 | 1 | 0 | 0 | 1 | 1 |
| 167 | 74 | 237 | 5:4:1:0:1 | 5 | 4 | 1 | 0 | 1 | 1 | 1 |
| 168 | 74 | 237 | 5:4:1:0:2 | 5 | 4 | 1 | 0 | 2 | 1 | |
| 169 | 74 | 237 | 5:4:1:1:0 | 5 | 4 | 1 | 1 | 0 | 1 | |
| 170 | 74 | 237 | 5:4:2:0:0 | 5 | 4 | 2 | 0 | 0 | 1 | 1 |
| 171 | 74 | 237 | 5:4:2:0:1 | 5 | 4 | 2 | 0 | 1 | 1 | 1 |
| 172 | 74 | 237 | 5:4:2:1:0 | 5 | 4 | 2 | 1 | 0 | 1 | 1 |
| 173 | 74 | 237 | 5:4:3:0:0 | 5 | 4 | 3 | 0 | 0 | 1 | |
| 174 | 74 | 237 | 5:5:0:1:0 | 5 | 5 | 0 | 1 | 0 | 1 | |
| 175 | 74 | 237 | 5:5:0:1:1 | 5 | 5 | 0 | 1 | 1 | 1 | |
| 176 | 74 | 237 | 5:5:1:0:0 | 5 | 5 | 1 | 0 | 0 | 1 | 1 |
| 177 | 74 | 237 | 5:5:1:0:1 | 5 | 5 | 1 | 0 | 1 | 1 | 1 |
| 178 | 74 | 237 | 5:5:1:0:2 | 5 | 5 | 1 | 0 | 2 | 1 | 1 |
| 179 | 74 | 237 | 5:5:1:1:0 | 5 | 5 | 1 | 1 | 0 | 1 | 1 |
| 180 | 74 | 237 | 5:5:2:0:0 | 5 | 5 | 2 | 0 | 0 | 1 | 1 |
| 181 | 74 | 237 | 5:5:2:0:1 | 5 | 5 | 2 | 0 | 1 | 1 | |
| 182 | 74 | 237 | 5:5:3:0:0 | 5 | 5 | 3 | 0 | 0 | 1 | 1 |
| 183 | 74 | 237 | 5:5:4:0:0 | 5 | 5 | 4 | 0 | 0 | 1 | |
| 184 | 74 | 237 | 5:5:4:0:1 | 5 | 5 | 4 | 0 | 1 | 1 | |
| 185 | 74 | 237 | 5:6:0:1:0 | 5 | 6 | 0 | 1 | 0 | 1 | |
| 186 | 74 | 237 | 5:6:0:1:1 | 5 | 6 | 0 | 1 | 1 | 1 | |
| 187 | 74 | 237 | 5:6:1:0:0 | 5 | 6 | 1 | 0 | 0 | 1 | 1 |
| 188 | 74 | 237 | 5:6:1:0:2 | 5 | 6 | 1 | 0 | 2 | 1 | |
| 189 | 74 | 237 | 5:6:1:0:3 | 5 | 6 | 1 | 0 | 3 | 1 | |
| 190 | 74 | 237 | 5:6:1:1:0 | 5 | 6 | 1 | 1 | 0 | 1 | 1 |
| 191 | 74 | 237 | 5:6:2:0:0 | 5 | 6 | 2 | 0 | 0 | 1 | 1 |

| | | | | | | | | | | |
|---|---|---|---|---|---|---|---|---|---|---|
| 192 | 74 | 237 | 5:6:2:0:1 | 5 | 6 | 2 | 0 | 1 | 1 | 1 |
| 193 | 74 | 237 | 5:6:2:1:0 | 5 | 6 | 2 | 1 | 0 | | 1 |
| 194 | 74 | 237 | 5:6:3:0:0 | 5 | 6 | 3 | 0 | 0 | 1 | 1 |
| 195 | 74 | 237 | 5:6:3:0:1 | 5 | 6 | 3 | 0 | 1 | 1 | 1 |
| 196 | 74 | 237 | 5:6:3:1:0 | 5 | 6 | 3 | 1 | 0 | 1 | |
| 197 | 74 | 237 | 5:6:4:0:0 | 5 | 6 | 4 | 0 | 0 | 1 | |
| 198 | 74 | 237 | 5:6:4:0:1 | 5 | 6 | 4 | 0 | 1 | 1 | |
| 199 | 74 | 237 | 5:7:0:1:1 | 5 | 7 | 0 | 1 | 1 | 1 | |
| 200 | 74 | 237 | 5:7:0:1:2 | 5 | 7 | 0 | 1 | 2 | 1 | |
| 201 | 74 | 237 | 5:7:1:1:0 | 5 | 7 | 1 | 1 | 0 | 1 | |
| 202 | 74 | 237 | 5:7:1:1:1 | 5 | 7 | 1 | 1 | 1 | | 1 |
| 203 | 74 | 237 | 5:7:2:1:0 | 5 | 7 | 2 | 1 | 0 | 1 | |
| 204 | 74 | 237 | 5:7:3:1:0 | 5 | 7 | 3 | 1 | 0 | 1 | |
| 205 | 74 | 237 | 6:3:1:0:0 | 6 | 3 | 1 | 0 | 0 | 1 | 1 |
| 206 | 74 | 237 | 6:3:1:0:1 | 6 | 3 | 1 | 0 | 1 | 1 | 1 |
| 207 | 74 | 237 | 6:3:2:0:0 | 6 | 3 | 2 | 0 | 0 | 1 | 1 |
| 208 | 74 | 237 | 6:3:2:0:1 | 6 | 3 | 2 | 0 | 1 | 1 | |
| 209 | 74 | 237 | 6:3:3:0:0 | 6 | 3 | 3 | 0 | 0 | 1 | 1 |
| 210 | 74 | 237 | 6:4:0:1:0 | 6 | 4 | 0 | 1 | 0 | 1 | |
| 211 | 74 | 237 | 6:4:1:0:0 | 6 | 4 | 1 | 0 | 0 | 1 | 1 |
| 212 | 74 | 237 | 6:4:1:1:0 | 6 | 4 | 1 | 1 | 0 | 1 | |
| 213 | 74 | 237 | 6:4:2:0:0 | 6 | 4 | 2 | 0 | 0 | 1 | 1 |
| 214 | 74 | 237 | 6:4:3:0:0 | 6 | 4 | 3 | 0 | 0 | 1 | |
| 215 | 74 | 237 | 6:5:1:0:0 | 6 | 5 | 1 | 0 | 0 | 1 | 1 |
| 216 | 74 | 237 | 6:5:1:1:0 | 6 | 5 | 1 | 1 | 0 | | 1 |
| 217 | 74 | 237 | 6:5:1:1:1 | 6 | 5 | 1 | 1 | 1 | | 1 |
| 218 | 74 | 237 | 6:5:2:0:0 | 6 | 5 | 2 | 0 | 0 | 1 | |
| 219 | 74 | 237 | 6:5:2:0:1 | 6 | 5 | 2 | 0 | 1 | 1 | |
| 220 | 74 | 237 | 6:5:2:0:2 | 6 | 5 | 2 | 0 | 2 | 1 | |
| 221 | 74 | 237 | 6:5:3:0:0 | 6 | 5 | 3 | 0 | 0 | 1 | |
| 222 | 74 | 237 | 6:5:3:0:1 | 6 | 5 | 3 | 0 | 1 | 1 | |
| 223 | 74 | 237 | 6:5:3:0:2 | 6 | 5 | 3 | 0 | 2 | 1 | |
| 224 | 74 | 237 | 6:5:4:0:0 | 6 | 5 | 4 | 0 | 0 | 1 | |
| 225 | 74 | 237 | 6:5:4:0:1 | 6 | 5 | 4 | 0 | 1 | 1 | |
| 226 | 74 | 237 | 6:6:0:1:1 | 6 | 6 | 0 | 1 | 1 | | 1 |
| 227 | 74 | 237 | 6:6:1:0:0 | 6 | 6 | 1 | 0 | 0 | 1 | 1 |
| 228 | 74 | 237 | 6:6:1:0:1 | 6 | 6 | 1 | 0 | 1 | 1 | |
| 229 | 74 | 237 | 6:6:1:0:2 | 6 | 6 | 1 | 0 | 2 | 1 | |
| 230 | 74 | 237 | 6:6:1:1:0 | 6 | 6 | 1 | 1 | 0 | 1 | |
| 231 | 74 | 237 | 6:6:1:1:1 | 6 | 6 | 1 | 1 | 1 | 1 | 1 |
| 232 | 74 | 237 | 6:6:2:0:0 | 6 | 6 | 2 | 0 | 0 | 1 | 1 |
| 233 | 74 | 237 | 6:6:2:0:1 | 6 | 6 | 2 | 0 | 1 | 1 | |
| 234 | 74 | 237 | 6:6:2:0:2 | 6 | 6 | 2 | 0 | 2 | 1 | |
| 235 | 74 | 237 | 6:6:2:1:0 | 6 | 6 | 2 | 1 | 0 | 1 | |
| 236 | 74 | 237 | 6:6:2:1:1 | 6 | 6 | 2 | 1 | 1 | 1 | |
| 237 | 74 | 237 | 6:6:3:0:0 | 6 | 6 | 3 | 0 | 0 | 1 | 1 |
| 238 | 74 | 237 | 6:6:3:0:1 | 6 | 6 | 3 | 0 | 1 | 1 | |
| 239 | 74 | 237 | 6:6:3:0:2 | 6 | 6 | 3 | 0 | 2 | 1 | |
| 240 | 74 | 237 | 6:6:3:1:0 | 6 | 6 | 3 | 1 | 0 | 1 | |
| 241 | 74 | 237 | 6:6:4:0:0 | 6 | 6 | 4 | 0 | 0 | 1 | |
| 242 | 74 | 237 | 6:6:4:0:1 | 6 | 6 | 4 | 0 | 1 | 1 | |
| 243 | 74 | 237 | 6:6:4:0:2 | 6 | 6 | 4 | 0 | 2 | 1 | |
| 244 | 74 | 237 | 6:7:0:1:0 | 6 | 7 | 0 | 1 | 0 | 1 | 1 |
| 245 | 74 | 237 | 6:7:0:1:1 | 6 | 7 | 0 | 1 | 1 | 1 | |
| 246 | 74 | 237 | 6:7:0:2:0 | 6 | 7 | 0 | 2 | 0 | 1 | |
| 247 | 74 | 237 | 6:7:1:0:0 | 6 | 7 | 1 | 0 | 0 | 1 | 1 |
| 248 | 74 | 237 | 6:7:1:0:1 | 6 | 7 | 1 | 0 | 1 | 1 | 1 |
| 249 | 74 | 237 | 6:7:1:0:2 | 6 | 7 | 1 | 0 | 2 | 1 | |
| 250 | 74 | 237 | 6:7:1:0:3 | 6 | 7 | 1 | 0 | 3 | 1 | |
| 251 | 74 | 237 | 6:7:1:1:0 | 6 | 7 | 1 | 1 | 0 | 1 | 1 |
| 252 | 74 | 237 | 6:7:1:1:1 | 6 | 7 | 1 | 1 | 1 | 1 | |
| 253 | 74 | 237 | 6:7:1:2:0 | 6 | 7 | 1 | 2 | 0 | 1 | |
| 254 | 74 | 237 | 6:7:2:0:0 | 6 | 7 | 2 | 0 | 0 | 1 | |
| 255 | 74 | 237 | 6:7:2:0:1 | 6 | 7 | 2 | 0 | 1 | 1 | |
| 256 | 74 | 237 | 6:7:2:0:2 | 6 | 7 | 2 | 0 | 2 | 1 | |

| | | | | | | | | | | |
|---|---|---|---|---|---|---|---|---|---|---|
| 257 | 74 | 237 | 6:7:2:0:3 | 6 | 7 | 2 | 0 | 3 | 1 | |
| 258 | 74 | 237 | 6:7:2:1:0 | 6 | 7 | 2 | 1 | 0 | 1 | |
| 259 | 74 | 237 | 6:7:2:1:1 | 6 | 7 | 2 | 1 | 1 | 1 | |
| 260 | 74 | 237 | 6:7:3:0:0 | 6 | 7 | 3 | 0 | 0 | 1 | |
| 261 | 74 | 237 | 6:7:3:0:1 | 6 | 7 | 3 | 0 | 1 | 1 | |
| 262 | 74 | 237 | 6:7:3:0:2 | 6 | 7 | 3 | 0 | 2 | 1 | |
| 263 | 74 | 237 | 6:7:3:1:0 | 6 | 7 | 3 | 1 | 0 | 1 | |
| 264 | 74 | 237 | 6:7:3:1:1 | 6 | 7 | 3 | 1 | 1 | 1 | |
| 265 | 74 | 237 | 6:7:4:0:0 | 6 | 7 | 4 | 0 | 0 | 1 | 1 |
| 266 | 74 | 237 | 6:7:4:0:1 | 6 | 7 | 4 | 0 | 1 | 1 | 1 |
| 267 | 74 | 237 | 6:7:4:1:0 | 6 | 7 | 4 | 1 | 0 | 1 | |
| 268 | 74 | 237 | 6:7:5:0:0 | 6 | 7 | 5 | 0 | 0 | | 1 |
| 269 | 74 | 237 | 6:8:0:1:0 | 6 | 8 | 0 | 1 | 0 | 1 | |
| 270 | 74 | 237 | 6:8:0:1:1 | 6 | 8 | 0 | 1 | 1 | 1 | |
| 271 | 74 | 237 | 6:8:0:1:2 | 6 | 8 | 0 | 1 | 2 | 1 | |
| 272 | 74 | 237 | 6:8:0:2:0 | 6 | 8 | 0 | 2 | 0 | 1 | |
| 273 | 74 | 237 | 6:8:1:1:0 | 6 | 8 | 1 | 1 | 0 | 1 | |
| 274 | 74 | 237 | 6:8:1:1:1 | 6 | 8 | 1 | 1 | 1 | 1 | |
| 275 | 74 | 237 | 6:8:1:1:2 | 6 | 8 | 1 | 1 | 2 | 1 | |
| 276 | 74 | 237 | 6:8:1:2:0 | 6 | 8 | 1 | 2 | 0 | 1 | |
| 277 | 74 | 237 | 6:8:2:1:0 | 6 | 8 | 2 | 1 | 0 | 1 | |
| 278 | 74 | 237 | 6:8:2:1:1 | 6 | 8 | 2 | 1 | 1 | 1 | |
| 279 | 74 | 237 | 6:8:2:2:0 | 6 | 8 | 2 | 2 | 0 | 1 | |
| 280 | 74 | 237 | 6:8:3:1:0 | 6 | 8 | 3 | 1 | 0 | 1 | |
| 281 | 74 | 237 | 6:8:4:1:0 | 6 | 8 | 4 | 1 | 0 | 1 | |
| 282 | 74 | 237 | 6:9:0:2:0 | 6 | 9 | 0 | 2 | 0 | 1 | |
| 283 | 74 | 237 | 6:9:0:2:1 | 6 | 9 | 0 | 2 | 1 | 1 | |
| 284 | 74 | 237 | 6:9:1:2:0 | 6 | 9 | 1 | 2 | 0 | 1 | |
| 285 | 74 | 237 | 7:3:1:0:0 | 7 | 3 | 1 | 0 | 0 | 1 | 1 |
| 286 | 74 | 237 | 7:4:1:0:0 | 7 | 4 | 1 | 0 | 0 | 1 | |
| 287 | 74 | 237 | 7:4:2:0:0 | 7 | 4 | 2 | 0 | 0 | 1 | 1 |
| 288 | 74 | 237 | 7:4:2:0:1 | 7 | 4 | 2 | 0 | 1 | 1 | |
| 289 | 74 | 237 | 7:4:3:0:0 | 7 | 4 | 3 | 0 | 0 | 1 | |
| 290 | 74 | 237 | 7:4:3:0:1 | 7 | 4 | 3 | 0 | 1 | 1 | |
| 291 | 74 | 237 | 7:4:4:0:0 | 7 | 4 | 4 | 0 | 0 | 1 | |
| 292 | 74 | 237 | 7:5:1:1:0 | 7 | 5 | 1 | 1 | 0 | 1 | |
| 293 | 74 | 237 | 7:5:2:1:0 | 7 | 5 | 2 | 1 | 0 | 1 | |
| 294 | 74 | 237 | 7:5:3:0:0 | 7 | 5 | 3 | 0 | 0 | 1 | |
| 295 | 74 | 237 | 7:5:3:0:1 | 7 | 5 | 3 | 0 | 1 | 1 | |
| 296 | 74 | 237 | 7:5:4:0:0 | 7 | 5 | 4 | 0 | 0 | 1 | |
| 297 | 74 | 237 | 7:6:1:0:0 | 7 | 6 | 1 | 0 | 0 | 1 | |
| 298 | 74 | 237 | 7:6:1:0:1 | 7 | 6 | 1 | 0 | 1 | 1 | |
| 299 | 74 | 237 | 7:6:2:0:0 | 7 | 6 | 2 | 0 | 0 | 1 | |
| 300 | 74 | 237 | 7:6:2:0:2 | 7 | 6 | 2 | 0 | 2 | 1 | |
| 301 | 74 | 237 | 7:6:2:1:0 | 7 | 6 | 2 | 1 | 0 | 1 | |
| 302 | 74 | 237 | 7:6:3:0:0 | 7 | 6 | 3 | 0 | 0 | 1 | |
| 303 | 74 | 237 | 7:6:3:0:1 | 7 | 6 | 3 | 0 | 1 | 1 | |
| 304 | 74 | 237 | 7:6:3:0:2 | 7 | 6 | 3 | 0 | 2 | 1 | |
| 305 | 74 | 237 | 7:6:4:0:0 | 7 | 6 | 4 | 0 | 0 | 1 | |
| 306 | 74 | 237 | 7:6:4:0:1 | 7 | 6 | 4 | 0 | 1 | 1 | |
| 307 | 74 | 237 | 7:7:0:1:0 | 7 | 7 | 0 | 1 | 0 | 1 | |
| 308 | 74 | 237 | 7:7:1:0:0 | 7 | 7 | 1 | 0 | 0 | 1 | |
| 309 | 74 | 237 | 7:7:1:0:1 | 7 | 7 | 1 | 0 | 1 | 1 | |
| 310 | 74 | 237 | 7:7:1:1:1 | 7 | 7 | 1 | 1 | 1 | 1 | |
| 311 | 74 | 237 | 7:7:2:0:0 | 7 | 7 | 2 | 0 | 0 | 1 | |
| 312 | 74 | 237 | 7:7:2:1:0 | 7 | 7 | 2 | 1 | 0 | 1 | |
| 313 | 74 | 237 | 7:7:2:1:1 | 7 | 7 | 2 | 1 | 1 | 1 | |
| 314 | 74 | 237 | 7:7:3:0:0 | 7 | 7 | 3 | 0 | 0 | 1 | |
| 315 | 74 | 237 | 7:7:3:1:0 | 7 | 7 | 3 | 1 | 0 | 1 | |
| 316 | 74 | 237 | 7:7:4:0:0 | 7 | 7 | 4 | 0 | 0 | 1 | |
| 317 | 74 | 237 | 7:7:4:0:1 | 7 | 7 | 4 | 0 | 1 | 1 | |
| 318 | 74 | 237 | 7:7:4:1:0 | 7 | 7 | 4 | 1 | 0 | 1 | |
| 319 | 74 | 237 | 7:8:0:1:0 | 7 | 8 | 0 | 1 | 0 | 1 | |
| 320 | 74 | 237 | 7:8:0:2:0 | 7 | 8 | 0 | 2 | 0 | 1 | |
| 321 | 74 | 237 | 7:8:1:2:0 | 7 | 8 | 1 | 2 | 0 | 1 | |

| | | | | | | | | | | |
|---|---|---|---|---|---|---|---|---|---|---|
| 322 | 74 | 237 | 7:8:3:1:0 | 7 | 8 | 3 | 1 | 0 | 1 | |
| 323 | 74 | 237 | 8:3:1:0:0 | 8 | 3 | 1 | 0 | 0 | 1 | |
| 324 | 74 | 237 | 8:3:1:1:0 | 8 | 3 | 1 | 1 | 0 | 1 | |
| 325 | 74 | 237 | 8:5:1:0:0 | 8 | 5 | 1 | 0 | 0 | 1 | |
| 326 | 74 | 237 | 8:5:1:0:1 | 8 | 5 | 1 | 0 | 1 | 1 | |
| 327 | 74 | 237 | 8:5:2:0:0 | 8 | 5 | 2 | 0 | 0 | 1 | |
| 328 | 74 | 237 | 8:5:3:0:0 | 8 | 5 | 3 | 0 | 0 | 1 | |
| 329 | 74 | 237 | 8:5:4:0:0 | 8 | 5 | 4 | 0 | 0 | 1 | |
| 330 | 74 | 237 | 8:6:0:1:0 | 8 | 6 | 0 | 1 | 0 | 1 | |
| 331 | 74 | 237 | 8:6:2:0:0 | 8 | 6 | 2 | 0 | 0 | 1 | |
| 332 | 74 | 237 | 8:6:3:0:0 | 8 | 6 | 3 | 0 | 0 | 1 | |
| 333 | 74 | 237 | 8:6:4:0:0 | 8 | 6 | 4 | 0 | 0 | 1 | |
| 334 | 74 | 237 | 8:8:0:0:0 | 8 | 8 | 0 | 0 | 0 | 1 | 1 |
| 335 | 74 | 237 | 8:9:0:0:1 | 8 | 9 | 0 | 0 | 1 | 1 | |
| 336 | 74 | 237 | 8:9:1:0:0 | 8 | 9 | 1 | 0 | 0 | | 1 |
| 337 | 74 | 237 | 9:8:0:0:1 | 9 | 8 | 0 | 0 | 1 | 1 | |
| 338 | 74 | 237 | 9:9:1:0:0 | 9 | 9 | 1 | 0 | 0 | 1 | 1 |
| 339 | 74 | 237 | 9:10:0:0:0 | 9 | 10 | 0 | 0 | 0 | 1 | 1 |
| 340 | 74 | 237 | 9:10:1:0:0 | 9 | 10 | 1 | 0 | 0 | 1 | |
| 341 | 74 | 237 | 10:3:0:3:0 | 10 | 3 | 0 | 3 | 0 | 1 | |
| 342 | 74 | 237 | 10:8:0:0:0 | 10 | 8 | 0 | 0 | 0 | 1 | |
| 343 | 74 | 237 | 10:9:0:0:0 | 10 | 9 | 0 | 0 | 0 | 1 | |
| 344 | 74 | 237 | 10:9:1:0:0 | 10 | 9 | 1 | 0 | 0 | 1 | |
| 345 | 74 | 237 | 10:10:1:0:0 | 10 | 10 | 1 | 0 | 0 | 1 | |
| 346 | 122 | 102 | 2:5:0:0:0 | 2 | 5 | 0 | 0 | 0 | 1 | 1 |
| 347 | 122 | 102 | 2:6:0:0:0 | 2 | 6 | 0 | 0 | 0 | 1 | 1 |
| 348 | 122 | 102 | 2:7:0:0:0 | 2 | 7 | 0 | 0 | 0 | 1 | 1 |
| 349 | 122 | 102 | 2:8:0:0:0 | 2 | 8 | 0 | 0 | 0 | 1 | 1 |
| 350 | 122 | 102 | 2:9:0:0:0 | 2 | 9 | 0 | 0 | 0 | 1 | |
| 351 | 122 | 102 | 3:3:0:0:0 | 3 | 3 | 0 | 0 | 0 | 1 | 1 |
| 352 | 122 | 102 | 3:3:1:0:0 | 3 | 3 | 1 | 0 | 0 | 1 | 1 |
| 353 | 122 | 102 | 3:4:0:0:0 | 3 | 4 | 0 | 0 | 0 | 1 | 1 |
| 354 | 122 | 102 | 3:4:0:0:1 | 3 | 4 | 0 | 0 | 1 | 1 | 1 |
| 355 | 122 | 102 | 3:4:1:0:0 | 3 | 4 | 1 | 0 | 0 | 1 | 1 |
| 356 | 122 | 102 | 3:4:1:0:1 | 3 | 4 | 1 | 0 | 1 | 1 | |
| 357 | 122 | 102 | 3:4:2:0:0 | 3 | 4 | 2 | 0 | 0 | 1 | 1 |
| 358 | 122 | 102 | 3:4:2:0:1 | 3 | 4 | 2 | 0 | 1 | 1 | |
| 359 | 122 | 102 | 3:5:0:0:0 | 3 | 5 | 0 | 0 | 0 | 1 | 1 |
| 360 | 122 | 102 | 3:5:0:0:1 | 3 | 5 | 0 | 0 | 1 | 1 | 1 |
| 361 | 122 | 102 | 3:5:0:1:0 | 3 | 5 | 0 | 1 | 0 | 1 | 1 |
| 362 | 122 | 102 | 3:5:1:0:0 | 3 | 5 | 1 | 0 | 0 | 1 | 1 |
| 363 | 122 | 102 | 3:5:1:0:1 | 3 | 5 | 1 | 0 | 1 | 1 | |
| 364 | 122 | 102 | 3:5:1:1:0 | 3 | 5 | 1 | 1 | 0 | 1 | |
| 365 | 122 | 102 | 3:5:2:0:0 | 3 | 5 | 2 | 0 | 0 | 1 | |
| 366 | 122 | 102 | 3:6:0:0:0 | 3 | 6 | 0 | 0 | 0 | 1 | 1 |
| 367 | 122 | 102 | 3:6:0:0:1 | 3 | 6 | 0 | 0 | 1 | 1 | 1 |
| 368 | 122 | 102 | 3:6:0:1:0 | 3 | 6 | 0 | 1 | 0 | 1 | 1 |
| 369 | 122 | 102 | 3:6:1:0:0 | 3 | 6 | 1 | 0 | 0 | 1 | 1 |
| 370 | 122 | 102 | 3:6:1:1:0 | 3 | 6 | 1 | 1 | 0 | | 1 |
| 371 | 122 | 102 | 3:7:1:0:0 | 3 | 7 | 1 | 0 | 0 | 1 | |
| 372 | 122 | 102 | 4:3:0:0:0 | 4 | 3 | 0 | 0 | 0 | 1 | 1 |
| 373 | 122 | 102 | 4:3:1:0:0 | 4 | 3 | 1 | 0 | 0 | 1 | 1 |
| 374 | 122 | 102 | 4:4:0:0:0 | 4 | 4 | 0 | 0 | 0 | 1 | 1 |
| 375 | 122 | 102 | 4:4:1:0:0 | 4 | 4 | 1 | 0 | 0 | 1 | 1 |
| 376 | 122 | 102 | 4:4:1:0:1 | 4 | 4 | 1 | 0 | 1 | 1 | |
| 377 | 122 | 102 | 4:4:2:0:0 | 4 | 4 | 2 | 0 | 0 | 1 | 1 |
| 378 | 122 | 102 | 4:4:2:0:1 | 4 | 4 | 2 | 0 | 1 | 1 | |
| 379 | 122 | 102 | 4:5:0:0:0 | 4 | 5 | 0 | 0 | 0 | 1 | 1 |
| 380 | 122 | 102 | 4:5:0:0:1 | 4 | 5 | 0 | 0 | 1 | 1 | 1 |
| 381 | 122 | 102 | 4:5:0:1:0 | 4 | 5 | 0 | 1 | 0 | 1 | 1 |
| 382 | 122 | 102 | 4:5:0:1:1 | 4 | 5 | 0 | 1 | 1 | | 1 |
| 383 | 122 | 102 | 4:5:1:0:0 | 4 | 5 | 1 | 0 | 0 | 1 | 1 |
| 384 | 122 | 102 | 4:5:1:0:1 | 4 | 5 | 1 | 0 | 1 | 1 | 1 |
| 385 | 122 | 102 | 4:5:1:1:0 | 4 | 5 | 1 | 1 | 0 | 1 | 1 |
| 386 | 122 | 102 | 4:5:2:0:0 | 4 | 5 | 2 | 0 | 0 | 1 | 1 |

| | | | | | | | | | | |
|---|---|---|---|---|---|---|---|---|---|---|
| 387 | 122 | 102 | 4:5:2:0:1 | 4 | 5 | 2 | 0 | 1 | 1 | 1 |
| 388 | 122 | 102 | 4:5:3:0:0 | 4 | 5 | 3 | 0 | 0 | 1 | 1 |
| 389 | 122 | 102 | 4:6:0:0:0 | 4 | 6 | 0 | 0 | 0 | | 1 |
| 390 | 122 | 102 | 4:6:0:1:0 | 4 | 6 | 0 | 1 | 0 | 1 | 1 |
| 391 | 122 | 102 | 4:6:1:0:0 | 4 | 6 | 1 | 0 | 0 | | 1 |
| 392 | 122 | 102 | 4:6:1:1:0 | 4 | 6 | 1 | 1 | 0 | 1 | |
| 393 | 122 | 102 | 4:6:2:0:0 | 4 | 6 | 2 | 0 | 0 | | 1 |
| 394 | 122 | 102 | 4:7:1:0:0 | 4 | 7 | 1 | 0 | 0 | 1 | 1 |
| 395 | 122 | 102 | 4:7:2:0:0 | 4 | 7 | 2 | 0 | 0 | 1 | 1 |
| 396 | 122 | 102 | 5:3:0:0:0 | 5 | 3 | 0 | 0 | 0 | 1 | |
| 397 | 122 | 102 | 5:3:1:0:0 | 5 | 3 | 1 | 0 | 0 | 1 | 1 |
| 398 | 122 | 102 | 5:3:1:0:1 | 5 | 3 | 1 | 0 | 1 | 1 | |
| 399 | 122 | 102 | 5:4:0:0:0 | 5 | 4 | 0 | 0 | 0 | 1 | 1 |
| 400 | 122 | 102 | 5:4:0:0:1 | 5 | 4 | 0 | 0 | 1 | | 1 |
| 401 | 122 | 102 | 5:4:0:1:0 | 5 | 4 | 0 | 1 | 0 | 1 | |
| 402 | 122 | 102 | 5:4:1:0:0 | 5 | 4 | 1 | 0 | 0 | 1 | 1 |
| 403 | 122 | 102 | 5:4:1:0:1 | 5 | 4 | 1 | 0 | 1 | 1 | |
| 404 | 122 | 102 | 5:4:2:0:0 | 5 | 4 | 2 | 0 | 0 | 1 | 1 |
| 405 | 122 | 102 | 5:4:2:1:0 | 5 | 4 | 2 | 1 | 0 | | 1 |
| 406 | 122 | 102 | 5:4:3:0:0 | 5 | 4 | 3 | 0 | 0 | 1 | |
| 407 | 122 | 102 | 5:5:0:0:0 | 5 | 5 | 0 | 0 | 0 | 1 | 1 |
| 408 | 122 | 102 | 5:5:0:1:0 | 5 | 5 | 0 | 1 | 0 | 1 | 1 |
| 409 | 122 | 102 | 5:5:1:0:0 | 5 | 5 | 1 | 0 | 0 | 1 | 1 |
| 410 | 122 | 102 | 5:5:1:0:1 | 5 | 5 | 1 | 0 | 1 | 1 | |
| 411 | 122 | 102 | 5:5:2:0:0 | 5 | 5 | 2 | 0 | 0 | 1 | 1 |
| 412 | 122 | 102 | 5:5:3:0:0 | 5 | 5 | 3 | 0 | 0 | 1 | 1 |
| 413 | 122 | 102 | 5:6:0:0:0 | 5 | 6 | 0 | 0 | 0 | 1 | 1 |
| 414 | 122 | 102 | 5:6:0:0:1 | 5 | 6 | 0 | 0 | 1 | 1 | 1 |
| 415 | 122 | 102 | 5:6:0:1:0 | 5 | 6 | 0 | 1 | 0 | 1 | 1 |
| 416 | 122 | 102 | 5:6:1:0:0 | 5 | 6 | 1 | 0 | 0 | | 1 |
| 417 | 122 | 102 | 5:6:1:0:1 | 5 | 6 | 1 | 0 | 1 | 1 | |
| 418 | 122 | 102 | 5:6:1:1:0 | 5 | 6 | 1 | 1 | 0 | | 1 |
| 419 | 122 | 102 | 5:6:2:0:0 | 5 | 6 | 2 | 0 | 0 | 1 | 1 |
| 420 | 122 | 102 | 5:6:3:0:0 | 5 | 6 | 3 | 0 | 0 | 1 | 1 |
| 421 | 122 | 102 | 5:6:3:0:1 | 5 | 6 | 3 | 0 | 1 | 1 | |
| 422 | 122 | 102 | 5:6:4:0:0 | 5 | 6 | 4 | 0 | 0 | 1 | |
| 423 | 122 | 102 | 5:7:0:1:0 | 5 | 7 | 0 | 1 | 0 | 1 | |
| 424 | 122 | 102 | 5:7:1:1:0 | 5 | 7 | 1 | 1 | 0 | | 1 |
| 425 | 122 | 102 | 5:7:2:1:0 | 5 | 7 | 2 | 1 | 0 | 1 | |
| 426 | 122 | 102 | 6:3:0:0:0 | 6 | 3 | 0 | 0 | 0 | | 1 |
| 427 | 122 | 102 | 6:5:0:0:0 | 6 | 5 | 0 | 0 | 0 | | 1 |
| 428 | 122 | 102 | 6:6:0:0:0 | 6 | 6 | 0 | 0 | 0 | 1 | 1 |
| 429 | 122 | 102 | 6:6:0:1:0 | 6 | 6 | 0 | 1 | 0 | | 1 |
| 430 | 122 | 102 | 6:6:0:1:1 | 6 | 6 | 0 | 1 | 1 | | 1 |
| 431 | 122 | 102 | 6:6:1:0:0 | 6 | 6 | 1 | 0 | 0 | 1 | 1 |
| 432 | 122 | 102 | 6:6:1:1:0 | 6 | 6 | 1 | 1 | 0 | | 1 |
| 433 | 122 | 102 | 6:6:2:0:0 | 6 | 6 | 2 | 0 | 0 | 1 | 1 |
| 434 | 122 | 102 | 6:6:2:0:1 | 6 | 6 | 2 | 0 | 1 | 1 | |
| 435 | 122 | 102 | 6:6:3:0:0 | 6 | 6 | 3 | 0 | 0 | 1 | |
| 436 | 122 | 102 | 6:6:4:0:0 | 6 | 6 | 4 | 0 | 0 | 1 | |
| 437 | 122 | 102 | 6:7:0:1:0 | 6 | 7 | 0 | 1 | 0 | | 1 |
| 438 | 122 | 102 | 6:7:1:0:0 | 6 | 7 | 1 | 0 | 0 | | 1 |
| 439 | 122 | 102 | 6:7:1:1:0 | 6 | 7 | 1 | 1 | 0 | 1 | 1 |
| 440 | 122 | 102 | 6:7:2:0:1 | 6 | 7 | 2 | 0 | 1 | 1 | |
| 441 | 122 | 102 | 6:7:2:0:2 | 6 | 7 | 2 | 0 | 2 | 1 | |
| 442 | 122 | 102 | 6:7:4:0:0 | 6 | 7 | 4 | 0 | 0 | 1 | 1 |
| 443 | 122 | 102 | 6:8:1:1:0 | 6 | 8 | 1 | 1 | 0 | 1 | |
| 444 | 122 | 102 | 6:8:1:1:1 | 6 | 8 | 1 | 1 | 1 | 1 | |
| 445 | 122 | 102 | 6:9:0:2:0 | 6 | 9 | 0 | 2 | 0 | 1 | |
| 446 | 122 | 102 | 7:7:1:0:0 | 7 | 7 | 1 | 0 | 0 | 1 | 1 |
| 447 | 122 | 102 | 7:7:3:0:0 | 7 | 7 | 3 | 0 | 0 | 1 | |
| 448 | 149 | 132 | 2:5:0:0:0 | 2 | 5 | 0 | 0 | 0 | 1 | 1 |
| 449 | 149 | 132 | 2:6:0:0:0 | 2 | 6 | 0 | 0 | 0 | 1 | |
| 450 | 149 | 132 | 2:7:0:0:0 | 2 | 7 | 0 | 0 | 0 | 1 | |
| 451 | 149 | 132 | 2:8:0:0:0 | 2 | 8 | 0 | 0 | 0 | 1 | |

| | | | | | | | | | | |
|---|---|---|---|---|---|---|---|---|---|---|
| 452 | 149 | 132 | 3:3:1:0:0 | 3 | 3 | 1 | 0 | 0 | 1 | |
| 453 | 149 | 132 | 3:4:2:0:0 | 3 | 4 | 2 | 0 | 0 | 1 | |
| 454 | 149 | 132 | 3:5:1:0:0 | 3 | 5 | 1 | 0 | 0 | 1 | |
| 455 | 149 | 132 | 3:5:2:0:0 | 3 | 5 | 2 | 0 | 0 | 1 | |
| 456 | 149 | 132 | 3:6:1:0:0 | 3 | 6 | 1 | 0 | 0 | 1 | |
| 457 | 149 | 132 | 3:6:2:0:0 | 3 | 6 | 2 | 0 | 0 | 1 | |
| 458 | 149 | 132 | 4:3:1:0:0 | 4 | 3 | 1 | 0 | 0 | 1 | |
| 459 | 149 | 132 | 4:4:1:0:0 | 4 | 4 | 1 | 0 | 0 | 1 | |
| 460 | 149 | 132 | 4:4:1:1:0 | 4 | 4 | 1 | 1 | 0 | | 1 |
| 461 | 149 | 132 | 4:4:2:0:0 | 4 | 4 | 2 | 0 | 0 | 1 | 1 |
| 462 | 149 | 132 | 4:5:0:0:0 | 4 | 5 | 0 | 0 | 0 | | 1 |
| 463 | 149 | 132 | 4:5:1:0:0 | 4 | 5 | 1 | 0 | 0 | 1 | 1 |
| 464 | 149 | 132 | 4:5:1:0:1 | 4 | 5 | 1 | 0 | 1 | 1 | |
| 465 | 149 | 132 | 4:5:1:0:2 | 4 | 5 | 1 | 0 | 2 | 1 | |
| 466 | 149 | 132 | 4:5:2:0:0 | 4 | 5 | 2 | 0 | 0 | 1 | |
| 467 | 149 | 132 | 4:5:2:0:1 | 4 | 5 | 2 | 0 | 1 | 1 | |
| 468 | 149 | 132 | 4:5:3:0:0 | 4 | 5 | 3 | 0 | 0 | 1 | 1 |
| 469 | 149 | 132 | 4:5:3:2:0 | 4 | 5 | 3 | 2 | 0 | | 1 |
| 470 | 149 | 132 | 4:5:4:0:2 | 4 | 5 | 4 | 0 | 2 | 1 | |
| 471 | 149 | 132 | 4:6:0:1:0 | 4 | 6 | 0 | 1 | 0 | 1 | 1 |
| 472 | 149 | 132 | 4:6:0:1:1 | 4 | 6 | 0 | 1 | 1 | 1 | |
| 473 | 149 | 132 | 4:6:1:1:0 | 4 | 6 | 1 | 1 | 0 | 1 | 1 |
| 474 | 149 | 132 | 4:6:2:0:0 | 4 | 6 | 2 | 0 | 0 | | 1 |
| 475 | 149 | 132 | 4:6:3:0:1 | 4 | 6 | 3 | 0 | 1 | 1 | |
| 476 | 149 | 132 | 4:6:3:1:1 | 4 | 6 | 3 | 1 | 1 | 1 | |
| 477 | 149 | 132 | 4:7:2:1:0 | 4 | 7 | 2 | 1 | 0 | 1 | |
| 478 | 149 | 132 | 4:7:2:2:0 | 4 | 7 | 2 | 2 | 0 | 1 | |
| 479 | 149 | 132 | 5:3:1:0:0 | 5 | 3 | 1 | 0 | 0 | 1 | 1 |
| 480 | 149 | 132 | 5:3:2:0:1 | 5 | 3 | 2 | 0 | 1 | 1 | |
| 481 | 149 | 132 | 5:4:1:0:0 | 5 | 4 | 1 | 0 | 0 | 1 | |
| 482 | 149 | 132 | 5:4:1:1:0 | 5 | 4 | 1 | 1 | 0 | 1 | |
| 483 | 149 | 132 | 5:4:2:0:0 | 5 | 4 | 2 | 0 | 0 | 1 | 1 |
| 484 | 149 | 132 | 5:4:2:0:1 | 5 | 4 | 2 | 0 | 1 | 1 | 1 |
| 485 | 149 | 132 | 5:4:3:0:0 | 5 | 4 | 3 | 0 | 0 | 1 | |
| 486 | 149 | 132 | 5:5:0:1:1 | 5 | 5 | 0 | 1 | 1 | | 1 |
| 487 | 149 | 132 | 5:5:1:0:0 | 5 | 5 | 1 | 0 | 0 | 1 | 1 |
| 488 | 149 | 132 | 5:5:1:1:0 | 5 | 5 | 1 | 1 | 0 | 1 | 1 |
| 489 | 149 | 132 | 5:5:2:0:0 | 5 | 5 | 2 | 0 | 0 | 1 | 1 |
| 490 | 149 | 132 | 5:5:2:0:1 | 5 | 5 | 2 | 0 | 1 | 1 | |
| 491 | 149 | 132 | 5:5:3:0:0 | 5 | 5 | 3 | 0 | 0 | 1 | 1 |
| 492 | 149 | 132 | 5:5:4:0:1 | 5 | 5 | 4 | 0 | 1 | 1 | |
| 493 | 149 | 132 | 5:6:0:3:0 | 5 | 6 | 0 | 3 | 0 | | 1 |
| 494 | 149 | 132 | 5:6:1:0:0 | 5 | 6 | 1 | 0 | 0 | 1 | 1 |
| 495 | 149 | 132 | 5:6:1:0:2 | 5 | 6 | 1 | 0 | 2 | 1 | |
| 496 | 149 | 132 | 5:6:1:1:0 | 5 | 6 | 1 | 1 | 0 | 1 | |
| 497 | 149 | 132 | 5:6:1:1:1 | 5 | 6 | 1 | 1 | 1 | | 1 |
| 498 | 149 | 132 | 5:6:2:0:0 | 5 | 6 | 2 | 0 | 0 | 1 | |
| 499 | 149 | 132 | 5:6:2:0:1 | 5 | 6 | 2 | 0 | 1 | 1 | |
| 500 | 149 | 132 | 5:6:2:0:2 | 5 | 6 | 2 | 0 | 2 | 1 | |
| 501 | 149 | 132 | 5:6:2:1:0 | 5 | 6 | 2 | 1 | 0 | | 1 |
| 502 | 149 | 132 | 5:6:3:0:0 | 5 | 6 | 3 | 0 | 0 | 1 | |
| 503 | 149 | 132 | 5:6:3:0:1 | 5 | 6 | 3 | 0 | 1 | 1 | |
| 504 | 149 | 132 | 5:6:3:1:0 | 5 | 6 | 3 | 1 | 0 | 1 | |
| 505 | 149 | 132 | 5:6:4:0:0 | 5 | 6 | 4 | 0 | 0 | 1 | |
| 506 | 149 | 132 | 5:7:0:1:1 | 5 | 7 | 0 | 1 | 1 | 1 | |
| 507 | 149 | 132 | 5:7:1:1:0 | 5 | 7 | 1 | 1 | 0 | 1 | |
| 508 | 149 | 132 | 5:7:1:1:1 | 5 | 7 | 1 | 1 | 1 | 1 | 1 |
| 509 | 149 | 132 | 5:7:2:1:0 | 5 | 7 | 2 | 1 | 0 | 1 | |
| 510 | 149 | 132 | 5:8:0:2:0 | 5 | 8 | 0 | 2 | 0 | 1 | |
| 511 | 149 | 132 | 6:3:1:0:0 | 6 | 3 | 1 | 0 | 0 | 1 | 1 |
| 512 | 149 | 132 | 6:4:1:0:0 | 6 | 4 | 1 | 0 | 0 | 1 | |
| 513 | 149 | 132 | 6:5:1:0:0 | 6 | 5 | 1 | 0 | 0 | 1 | |
| 514 | 149 | 132 | 6:5:2:0:1 | 6 | 5 | 2 | 0 | 1 | 1 | |
| 515 | 149 | 132 | 6:5:2:0:2 | 6 | 5 | 2 | 0 | 2 | 1 | |
| 516 | 149 | 132 | 6:5:3:0:0 | 6 | 5 | 3 | 0 | 0 | 1 | |

| | | | | | | | | | | |
|---|---|---|---|---|---|---|---|---|---|---|
| 517 | 149 | 132 | 6:5:4:0:0 | 6 | 5 | 4 | 0 | 0 | 1 | |
| 518 | 149 | 132 | 6:5:4:0:1 | 6 | 5 | 4 | 0 | 1 | 1 | |
| 519 | 149 | 132 | 6:6:1:0:0 | 6 | 6 | 1 | 0 | 0 | 1 | 1 |
| 520 | 149 | 132 | 6:6:1:0:1 | 6 | 6 | 1 | 0 | 1 | 1 | |
| 521 | 149 | 132 | 6:6:1:1:0 | 6 | 6 | 1 | 1 | 0 | 1 | |
| 522 | 149 | 132 | 6:6:1:1:1 | 6 | 6 | 1 | 1 | 1 | 1 | 1 |
| 523 | 149 | 132 | 6:6:2:0:0 | 6 | 6 | 2 | 0 | 0 | 1 | 1 |
| 524 | 149 | 132 | 6:6:2:0:1 | 6 | 6 | 2 | 0 | 1 | 1 | |
| 525 | 149 | 132 | 6:6:2:0:2 | 6 | 6 | 2 | 0 | 2 | 1 | |
| 526 | 149 | 132 | 6:6:3:0:0 | 6 | 6 | 3 | 0 | 0 | 1 | |
| 527 | 149 | 132 | 6:6:3:0:1 | 6 | 6 | 3 | 0 | 1 | 1 | |
| 528 | 149 | 132 | 6:6:3:0:2 | 6 | 6 | 3 | 0 | 2 | 1 | |
| 529 | 149 | 132 | 6:6:3:1:0 | 6 | 6 | 3 | 1 | 0 | 1 | |
| 530 | 149 | 132 | 6:6:4:0:0 | 6 | 6 | 4 | 0 | 0 | 1 | |
| 531 | 149 | 132 | 6:6:4:0:1 | 6 | 6 | 4 | 0 | 1 | 1 | |
| 532 | 149 | 132 | 6:7:0:1:0 | 6 | 7 | 0 | 1 | 0 | 1 | |
| 533 | 149 | 132 | 6:7:0:1:1 | 6 | 7 | 0 | 1 | 1 | | 1 |
| 534 | 149 | 132 | 6:7:0:2:0 | 6 | 7 | 0 | 2 | 0 | 1 | |
| 535 | 149 | 132 | 6:7:1:0:0 | 6 | 7 | 1 | 0 | 0 | 1 | 1 |
| 536 | 149 | 132 | 6:7:1:0:1 | 6 | 7 | 1 | 0 | 1 | 1 | |
| 537 | 149 | 132 | 6:7:1:1:0 | 6 | 7 | 1 | 1 | 0 | 1 | 1 |
| 538 | 149 | 132 | 6:7:1:1:1 | 6 | 7 | 1 | 1 | 1 | 1 | |
| 539 | 149 | 132 | 6:7:2:0:0 | 6 | 7 | 2 | 0 | 0 | 1 | |
| 540 | 149 | 132 | 6:7:2:0:1 | 6 | 7 | 2 | 0 | 1 | 1 | |
| 541 | 149 | 132 | 6:7:2:0:2 | 6 | 7 | 2 | 0 | 2 | 1 | |
| 542 | 149 | 132 | 6:7:2:1:0 | 6 | 7 | 2 | 1 | 0 | 1 | |
| 543 | 149 | 132 | 6:7:2:1:1 | 6 | 7 | 2 | 1 | 1 | 1 | |
| 544 | 149 | 132 | 6:7:3:0:0 | 6 | 7 | 3 | 0 | 0 | 1 | |
| 545 | 149 | 132 | 6:7:3:0:1 | 6 | 7 | 3 | 0 | 1 | 1 | |
| 546 | 149 | 132 | 6:7:3:0:2 | 6 | 7 | 3 | 0 | 2 | 1 | |
| 547 | 149 | 132 | 6:7:3:1:0 | 6 | 7 | 3 | 1 | 0 | 1 | |
| 548 | 149 | 132 | 6:7:4:0:0 | 6 | 7 | 4 | 0 | 0 | 1 | |
| 549 | 149 | 132 | 6:7:4:0:1 | 6 | 7 | 4 | 0 | 1 | 1 | |
| 550 | 149 | 132 | 6:7:4:1:0 | 6 | 7 | 4 | 1 | 0 | 1 | 1 |
| 551 | 149 | 132 | 6:7:5:0:0 | 6 | 7 | 5 | 0 | 0 | | 1 |
| 552 | 149 | 132 | 6:8:0:1:0 | 6 | 8 | 0 | 1 | 0 | 1 | |
| 553 | 149 | 132 | 6:8:0:2:0 | 6 | 8 | 0 | 2 | 0 | 1 | |
| 554 | 149 | 132 | 6:8:1:1:0 | 6 | 8 | 1 | 1 | 0 | 1 | |
| 555 | 149 | 132 | 6:8:1:1:1 | 6 | 8 | 1 | 1 | 1 | 1 | |
| 556 | 149 | 132 | 6:8:1:2:0 | 6 | 8 | 1 | 2 | 0 | 1 | |
| 557 | 149 | 132 | 6:8:2:1:0 | 6 | 8 | 2 | 1 | 0 | 1 | |
| 558 | 149 | 132 | 6:8:2:1:1 | 6 | 8 | 2 | 1 | 1 | 1 | |
| 559 | 149 | 132 | 6:8:3:1:0 | 6 | 8 | 3 | 1 | 0 | 1 | |
| 560 | 149 | 132 | 6:9:0:2:0 | 6 | 9 | 0 | 2 | 0 | 1 | |
| 561 | 149 | 132 | 6:9:1:2:0 | 6 | 9 | 1 | 2 | 0 | 1 | |
| 562 | 149 | 132 | 7:5:4:0:1 | 7 | 5 | 4 | 0 | 1 | 1 | |
| 563 | 149 | 132 | 7:6:3:0:1 | 7 | 6 | 3 | 0 | 1 | 1 | |
| 564 | 149 | 132 | 7:6:3:1:0 | 7 | 6 | 3 | 1 | 0 | 1 | |
| 565 | 149 | 132 | 7:6:4:0:0 | 7 | 6 | 4 | 0 | 0 | 1 | |
| 566 | 149 | 132 | 7:6:4:0:1 | 7 | 6 | 4 | 0 | 1 | 1 | |
| 567 | 149 | 132 | 7:7:1:0:1 | 7 | 7 | 1 | 0 | 1 | 1 | |
| 568 | 149 | 132 | 7:7:2:0:0 | 7 | 7 | 2 | 0 | 0 | 1 | |
| 569 | 149 | 132 | 7:7:2:1:0 | 7 | 7 | 2 | 1 | 0 | 1 | |
| 570 | 149 | 132 | 7:7:3:0:0 | 7 | 7 | 3 | 0 | 0 | 1 | |
| 571 | 149 | 132 | 7:7:3:0:1 | 7 | 7 | 3 | 0 | 1 | 1 | |
| 572 | 149 | 132 | 7:7:3:1:0 | 7 | 7 | 3 | 1 | 0 | 1 | |
| 573 | 149 | 132 | 7:7:4:0:0 | 7 | 7 | 4 | 0 | 0 | 1 | |
| 574 | 149 | 132 | 7:8:0:1:0 | 7 | 8 | 0 | 1 | 0 | 1 | |
| 575 | 149 | 132 | 7:8:2:1:0 | 7 | 8 | 2 | 1 | 0 | 1 | |
| 576 | 149 | 132 | 8:8:0:0:1 | 8 | 8 | 0 | 0 | 1 | 1 | |
| 577 | 149 | 132 | 8:9:0:0:0 | 8 | 9 | 0 | 0 | 0 | 1 | |
| 578 | 149 | 132 | 9:8:0:0:0 | 9 | 8 | 0 | 0 | 0 | 1 | |
| 579 | 149 | 132 | 9:9:0:0:0 | 9 | 9 | 0 | 0 | 0 | 1 | |
| 580 | 165 | 50 | 2:5:0:0:0 | 2 | 5 | 0 | 0 | 0 | 1 | 1 |
| 581 | 165 | 50 | 2:6:0:0:0 | 2 | 6 | 0 | 0 | 0 | 1 | 1 |

| | | | | | | | | | | |
|---|---|---|---|---|---|---|---|---|---|---|
| 582 | 165 | 50 | 3:4:1:0:0 | 3 | 4 | 1 | 0 | 0 | 1 | 1 |
| 583 | 165 | 50 | 3:5:1:0:0 | 3 | 5 | 1 | 0 | 0 | 1 | 1 |
| 584 | 165 | 50 | 3:6:0:0:0 | 3 | 6 | 0 | 0 | 0 | 1 | |
| 585 | 165 | 50 | 3:6:0:0:1 | 3 | 6 | 0 | 0 | 1 | 1 | 1 |
| 586 | 165 | 50 | 3:6:1:0:0 | 3 | 6 | 1 | 0 | 0 | 1 | 1 |
| 587 | 165 | 50 | 4:5:0:0:0 | 4 | 5 | 0 | 0 | 0 | 1 | 1 |
| 588 | 165 | 50 | 4:5:0:0:1 | 4 | 5 | 0 | 0 | 1 | 1 | |
| 589 | 165 | 50 | 4:5:0:1:0 | 4 | 5 | 0 | 1 | 0 | | 1 |
| 590 | 165 | 50 | 4:5:1:0:0 | 4 | 5 | 1 | 0 | 0 | 1 | 1 |
| 591 | 165 | 50 | 4:5:1:0:1 | 4 | 5 | 1 | 0 | 1 | 1 | 1 |
| 592 | 165 | 50 | 4:5:1:1:0 | 4 | 5 | 1 | 1 | 0 | | 1 |
| 593 | 165 | 50 | 4:5:2:0:0 | 4 | 5 | 2 | 0 | 0 | 1 | 1 |
| 594 | 165 | 50 | 4:5:3:0:0 | 4 | 5 | 3 | 0 | 0 | 1 | 1 |
| 595 | 165 | 50 | 4:6:0:1:0 | 4 | 6 | 0 | 1 | 0 | 1 | 1 |
| 596 | 165 | 50 | 4:6:1:1:0 | 4 | 6 | 1 | 1 | 0 | | 1 |
| 597 | 165 | 50 | 5:4:1:0:0 | 5 | 4 | 1 | 0 | 0 | 1 | |
| 598 | 165 | 50 | 5:4:3:0:0 | 5 | 4 | 3 | 0 | 0 | 1 | |
| 599 | 165 | 50 | 5:5:0:0:0 | 5 | 5 | 0 | 0 | 0 | 1 | 1 |
| 600 | 165 | 50 | 5:5:1:0:0 | 5 | 5 | 1 | 0 | 0 | 1 | 1 |
| 601 | 165 | 50 | 5:5:1:1:0 | 5 | 5 | 1 | 1 | 0 | | 1 |
| 602 | 165 | 50 | 5:5:2:0:0 | 5 | 5 | 2 | 0 | 0 | 1 | 1 |
| 603 | 165 | 50 | 5:5:3:0:0 | 5 | 5 | 3 | 0 | 0 | 1 | 1 |
| 604 | 165 | 50 | 5:6:0:0:0 | 5 | 6 | 0 | 0 | 0 | | 1 |
| 605 | 165 | 50 | 5:6:1:1:0 | 5 | 6 | 1 | 1 | 0 | | 1 |
| 606 | 165 | 50 | 5:6:2:0:0 | 5 | 6 | 2 | 0 | 0 | 1 | 1 |
| 607 | 165 | 50 | 5:6:2:1:0 | 5 | 6 | 2 | 1 | 0 | | 1 |
| 608 | 165 | 50 | 5:6:3:0:0 | 5 | 6 | 3 | 0 | 0 | 1 | 1 |
| 609 | 165 | 50 | 5:6:4:0:0 | 5 | 6 | 4 | 0 | 0 | 1 | |
| 610 | 165 | 50 | 6:3:1:1:0 | 6 | 3 | 1 | 1 | 0 | | 1 |
| 611 | 165 | 50 | 6:6:1:0:0 | 6 | 6 | 1 | 0 | 0 | 1 | 1 |
| 612 | 165 | 50 | 6:6:1:1:0 | 6 | 6 | 1 | 1 | 0 | | 1 |
| 613 | 165 | 50 | 6:6:3:0:0 | 6 | 6 | 3 | 0 | 0 | 1 | |
| 614 | 165 | 50 | 6:6:4:0:0 | 6 | 6 | 4 | 0 | 0 | 1 | |
| 615 | 165 | 50 | 6:7:1:0:0 | 6 | 7 | 1 | 0 | 0 | 1 | 1 |
| 616 | 165 | 50 | 6:7:1:1:0 | 6 | 7 | 1 | 1 | 0 | | 1 |
| 617 | 165 | 50 | 6:7:1:2:0 | 6 | 7 | 1 | 2 | 0 | | 1 |
| 618 | 165 | 50 | 6:7:2:0:0 | 6 | 7 | 2 | 0 | 0 | 1 | |
| 619 | 165 | 50 | 6:7:3:0:0 | 6 | 7 | 3 | 0 | 0 | 1 | |
| 620 | 165 | 50 | 6:7:3:0:1 | 6 | 7 | 3 | 0 | 1 | 1 | |
| 621 | 165 | 50 | 6:7:4:0:0 | 6 | 7 | 4 | 0 | 0 | 1 | 1 |
| 622 | 165 | 50 | 6:7:5:0:0 | 6 | 7 | 5 | 0 | 0 | | 1 |
| 623 | 165 | 50 | 6:8:2:1:0 | 6 | 8 | 2 | 1 | 0 | 1 | |
| 624 | 165 | 50 | 7:7:1:0:0 | 7 | 7 | 1 | 0 | 0 | 1 | |
| 625 | 165 | 50 | 7:7:2:0:0 | 7 | 7 | 2 | 0 | 0 | 1 | |
| 626 | 165 | 50 | 7:7:3:0:0 | 7 | 7 | 3 | 0 | 0 | 1 | |
| 627 | 165 | 50 | 7:7:4:0:0 | 7 | 7 | 4 | 0 | 0 | 1 | |
| 628 | 165 | 50 | 8:9:0:0:0 | 8 | 9 | 0 | 0 | 0 | 1 | 1 |
| 629 | 165 | 50 | 9:9:0:0:0 | 9 | 9 | 0 | 0 | 0 | 1 | |
| 630 | 234 | 46 | 2:5:0:0:0 | 2 | 5 | 0 | 0 | 0 | 1 | 1 |
| 631 | 234 | 46 | 2:6:0:0:0 | 2 | 6 | 0 | 0 | 0 | 1 | 1 |
| 632 | 234 | 46 | 2:7:0:0:0 | 2 | 7 | 0 | 0 | 0 | 1 | 1 |
| 633 | 234 | 46 | 2:8:0:0:0 | 2 | 8 | 0 | 0 | 0 | 1 | 1 |
| 634 | 234 | 46 | 2:9:0:0:0 | 2 | 9 | 0 | 0 | 0 | 1 | 1 |
| 635 | 234 | 46 | 3:4:0:0:1 | 3 | 4 | 0 | 0 | 1 | | 1 |
| 636 | 234 | 46 | 3:4:1:0:0 | 3 | 4 | 1 | 0 | 0 | | 1 |
| 637 | 234 | 46 | 3:5:0:0:0 | 3 | 5 | 0 | 0 | 0 | 1 | 1 |
| 638 | 234 | 46 | 3:5:0:0:1 | 3 | 5 | 0 | 0 | 1 | | 1 |
| 639 | 234 | 46 | 3:5:1:0:0 | 3 | 5 | 1 | 0 | 0 | 1 | 1 |
| 640 | 234 | 46 | 3:6:0:0:0 | 3 | 6 | 0 | 0 | 0 | 1 | 1 |
| 641 | 234 | 46 | 3:6:0:0:1 | 3 | 6 | 0 | 0 | 1 | 1 | |
| 642 | 234 | 46 | 4:3:1:0:0 | 4 | 3 | 1 | 0 | 0 | 1 | 1 |
| 643 | 234 | 46 | 4:4:0:0:0 | 4 | 4 | 0 | 0 | 0 | 1 | 1 |
| 644 | 234 | 46 | 4:4:1:0:0 | 4 | 4 | 1 | 0 | 0 | 1 | |
| 645 | 234 | 46 | 4:5:0:0:0 | 4 | 5 | 0 | 0 | 0 | 1 | 1 |
| 646 | 234 | 46 | 4:5:0:0:1 | 4 | 5 | 0 | 0 | 1 | 1 | |

| | | | | | | | | | | |
|---|---|---|---|---|---|---|---|---|---|---|
| 647 | 234 | 46 | 4:5:1:0:0 | 4 | 5 | 1 | 0 | 0 | 1 | 1 |
| 648 | 234 | 46 | 4:5:2:0:0 | 4 | 5 | 2 | 0 | 0 | 1 | |
| 649 | 234 | 46 | 4:5:2:1:0 | 4 | 5 | 2 | 1 | 0 | | 1 |
| 650 | 234 | 46 | 4:5:3:0:0 | 4 | 5 | 3 | 0 | 0 | | 1 |
| 651 | 234 | 46 | 4:5:3:1:0 | 4 | 5 | 3 | 1 | 0 | | 1 |
| 652 | 234 | 46 | 5:3:1:0:0 | 5 | 3 | 1 | 0 | 0 | | 1 |
| 653 | 234 | 46 | 5:4:1:0:0 | 5 | 4 | 1 | 0 | 0 | 1 | 1 |
| 654 | 234 | 46 | 5:5:1:0:0 | 5 | 5 | 1 | 0 | 0 | 1 | 1 |
| 655 | 234 | 46 | 5:6:0:0:0 | 5 | 6 | 0 | 0 | 0 | 1 | |
| 656 | 234 | 46 | 5:6:1:0:0 | 5 | 6 | 1 | 0 | 0 | 1 | 1 |
| 657 | 234 | 46 | 5:7:0:0:0 | 5 | 7 | 0 | 0 | 0 | | 1 |
| 658 | 234 | 46 | 6:3:0:0:0 | 6 | 3 | 0 | 0 | 0 | | 1 |
| 659 | 234 | 46 | 6:3:1:0:0 | 6 | 3 | 1 | 0 | 0 | | 1 |
| 660 | 234 | 46 | 6:3:1:1:0 | 6 | 3 | 1 | 1 | 0 | | 1 |
| 661 | 234 | 46 | 6:3:2:0:0 | 6 | 3 | 2 | 0 | 0 | | 1 |
| 662 | 234 | 46 | 6:3:3:0:0 | 6 | 3 | 3 | 0 | 0 | | 1 |
| 663 | 234 | 46 | 6:4:0:0:0 | 6 | 4 | 0 | 0 | 0 | | 1 |
| 664 | 234 | 46 | 6:4:1:0:0 | 6 | 4 | 1 | 0 | 0 | 1 | 1 |
| 665 | 234 | 46 | 6:5:0:0:0 | 6 | 5 | 0 | 0 | 0 | 1 | 1 |
| 666 | 234 | 46 | 6:5:1:0:0 | 6 | 5 | 1 | 0 | 0 | 1 | |
| 667 | 234 | 46 | 6:6:0:0:0 | 6 | 6 | 0 | 0 | 0 | 1 | |
| 668 | 234 | 46 | 6:6:1:0:0 | 6 | 6 | 1 | 0 | 0 | 1 | 1 |
| 669 | 234 | 46 | 6:6:1:2:0 | 6 | 6 | 1 | 2 | 0 | | 1 |
| 670 | 234 | 46 | 6:7:0:0:0 | 6 | 7 | 0 | 0 | 0 | 1 | 1 |
| 671 | 234 | 46 | 6:7:1:0:0 | 6 | 7 | 1 | 0 | 0 | 1 | |
| 672 | 234 | 46 | 7:3:0:0:0 | 7 | 3 | 0 | 0 | 0 | | 1 |
| 673 | 234 | 46 | 7:6:1:3:1 | 7 | 6 | 1 | 3 | 1 | | 1 |
| 674 | 234 | 46 | 7:7:1:4:0 | 7 | 7 | 1 | 4 | 0 | | 1 |
| 675 | 234 | 46 | 8:6:0:0:0 | 8 | 6 | 0 | 0 | 0 | | 1 |
| 676 | 282 | 77 | 2:5:0:0:0 | 2 | 5 | 0 | 0 | 0 | 1 | 1 |
| 677 | 282 | 77 | 2:6:0:0:0 | 2 | 6 | 0 | 0 | 0 | 1 | 1 |
| 678 | 282 | 77 | 2:7:0:0:0 | 2 | 7 | 0 | 0 | 0 | | 1 |
| 679 | 282 | 77 | 3:3:0:0:0 | 3 | 3 | 0 | 0 | 0 | | 1 |
| 680 | 282 | 77 | 3:3:1:0:1 | 3 | 3 | 1 | 0 | 1 | 1 | |
| 681 | 282 | 77 | 3:3:2:0:1 | 3 | 3 | 2 | 0 | 1 | 1 | |
| 682 | 282 | 77 | 3:4:0:1:0 | 3 | 4 | 0 | 1 | 0 | 1 | |
| 683 | 282 | 77 | 3:4:1:0:0 | 3 | 4 | 1 | 0 | 0 | 1 | |
| 684 | 282 | 77 | 3:4:1:1:0 | 3 | 4 | 1 | 1 | 0 | 1 | |
| 685 | 282 | 77 | 3:5:0:0:0 | 3 | 5 | 0 | 0 | 0 | 1 | 1 |
| 686 | 282 | 77 | 3:5:0:0:1 | 3 | 5 | 0 | 0 | 1 | | 1 |
| 687 | 282 | 77 | 3:5:1:0:0 | 3 | 5 | 1 | 0 | 0 | 1 | 1 |
| 688 | 282 | 77 | 3:6:0:0:0 | 3 | 6 | 0 | 0 | 0 | | 1 |
| 689 | 282 | 77 | 3:6:1:0:0 | 3 | 6 | 1 | 0 | 0 | 1 | |
| 690 | 282 | 77 | 4:3:0:0:0 | 4 | 3 | 0 | 0 | 0 | 1 | |
| 691 | 282 | 77 | 4:3:1:0:0 | 4 | 3 | 1 | 0 | 0 | 1 | 1 |
| 692 | 282 | 77 | 4:4:0:0:1 | 4 | 4 | 0 | 0 | 1 | | 1 |
| 693 | 282 | 77 | 4:4:1:0:0 | 4 | 4 | 1 | 0 | 0 | 1 | |
| 694 | 282 | 77 | 4:4:1:0:1 | 4 | 4 | 1 | 0 | 1 | 1 | |
| 695 | 282 | 77 | 4:4:2:0:1 | 4 | 4 | 2 | 0 | 1 | 1 | |
| 696 | 282 | 77 | 4:4:3:0:1 | 4 | 4 | 3 | 0 | 1 | 1 | |
| 697 | 282 | 77 | 4:5:0:0:0 | 4 | 5 | 0 | 0 | 0 | 1 | 1 |
| 698 | 282 | 77 | 4:5:0:1:0 | 4 | 5 | 0 | 1 | 0 | 1 | |
| 699 | 282 | 77 | 4:5:1:0:0 | 4 | 5 | 1 | 0 | 0 | 1 | 1 |
| 700 | 282 | 77 | 4:5:1:0:1 | 4 | 5 | 1 | 0 | 1 | 1 | |
| 701 | 282 | 77 | 4:5:1:1:0 | 4 | 5 | 1 | 1 | 0 | 1 | 1 |
| 702 | 282 | 77 | 4:5:2:0:0 | 4 | 5 | 2 | 0 | 0 | 1 | 1 |
| 703 | 282 | 77 | 4:5:2:1:0 | 4 | 5 | 2 | 1 | 0 | 1 | |
| 704 | 282 | 77 | 4:5:3:0:0 | 4 | 5 | 3 | 0 | 0 | 1 | 1 |
| 705 | 282 | 77 | 4:5:3:1:0 | 4 | 5 | 3 | 1 | 0 | | 1 |
| 706 | 282 | 77 | 4:6:0:0:0 | 4 | 6 | 0 | 0 | 0 | | 1 |
| 707 | 282 | 77 | 4:6:0:1:0 | 4 | 6 | 0 | 1 | 0 | 1 | 1 |
| 708 | 282 | 77 | 5:3:0:0:0 | 5 | 3 | 0 | 0 | 0 | 1 | |
| 709 | 282 | 77 | 5:4:1:0:0 | 5 | 4 | 1 | 0 | 0 | 1 | 1 |
| 710 | 282 | 77 | 5:4:1:1:1 | 5 | 4 | 1 | 1 | 1 | | 1 |
| 711 | 282 | 77 | 5:4:2:0:0 | 5 | 4 | 2 | 0 | 0 | 1 | |

| | | | | | | | | | | |
|---|---|---|---|---|---|---|---|---|---|---|
| 712 | 282 | 77 | 5:5:0:0:0 | 5 | 5 | 0 | 0 | 0 | | 1 |
| 713 | 282 | 77 | 5:5:0:1:0 | 5 | 5 | 0 | 1 | 0 | | 1 |
| 714 | 282 | 77 | 5:5:1:0:0 | 5 | 5 | 1 | 0 | 0 | 1 | 1 |
| 715 | 282 | 77 | 5:5:1:1:0 | 5 | 5 | 1 | 1 | 0 | | 1 |
| 716 | 282 | 77 | 5:5:2:0:0 | 5 | 5 | 2 | 0 | 0 | 1 | |
| 717 | 282 | 77 | 5:5:3:0:0 | 5 | 5 | 3 | 0 | 0 | 1 | |
| 718 | 282 | 77 | 5:6:0:0:0 | 5 | 6 | 0 | 0 | 0 | 1 | 1 |
| 719 | 282 | 77 | 5:6:0:1:0 | 5 | 6 | 0 | 1 | 0 | | 1 |
| 720 | 282 | 77 | 5:6:1:0:0 | 5 | 6 | 1 | 0 | 0 | 1 | 1 |
| 721 | 282 | 77 | 5:6:1:0:1 | 5 | 6 | 1 | 0 | 1 | | 1 |
| 722 | 282 | 77 | 5:6:1:1:0 | 5 | 6 | 1 | 1 | 0 | | 1 |
| 723 | 282 | 77 | 5:6:2:0:0 | 5 | 6 | 2 | 0 | 0 | 1 | 1 |
| 724 | 282 | 77 | 5:6:3:0:0 | 5 | 6 | 3 | 0 | 0 | 1 | 1 |
| 725 | 282 | 77 | 5:7:0:0:0 | 5 | 7 | 0 | 0 | 0 | | 1 |
| 726 | 282 | 77 | 5:7:1:0:0 | 5 | 7 | 1 | 0 | 0 | | 1 |
| 727 | 282 | 77 | 6:3:1:0:0 | 6 | 3 | 1 | 0 | 0 | 1 | 1 |
| 728 | 282 | 77 | 6:4:1:0:0 | 6 | 4 | 1 | 0 | 0 | 1 | 1 |
| 729 | 282 | 77 | 6:5:1:0:0 | 6 | 5 | 1 | 0 | 0 | 1 | 1 |
| 730 | 282 | 77 | 6:5:1:0:1 | 6 | 5 | 1 | 0 | 1 | 1 | |
| 731 | 282 | 77 | 6:6:0:0:0 | 6 | 6 | 0 | 0 | 0 | | 1 |
| 732 | 282 | 77 | 6:6:0:1:0 | 6 | 6 | 0 | 1 | 0 | 1 | 1 |
| 733 | 282 | 77 | 6:6:1:0:0 | 6 | 6 | 1 | 0 | 0 | 1 | 1 |
| 734 | 282 | 77 | 6:6:2:0:0 | 6 | 6 | 2 | 0 | 0 | 1 | 1 |
| 735 | 282 | 77 | 6:6:3:0:0 | 6 | 6 | 3 | 0 | 0 | 1 | 1 |
| 736 | 282 | 77 | 6:7:0:0:0 | 6 | 7 | 0 | 0 | 0 | 1 | 1 |
| 737 | 282 | 77 | 6:7:0:1:0 | 6 | 7 | 0 | 1 | 0 | | 1 |
| 738 | 282 | 77 | 6:7:0:3:1 | 6 | 7 | 0 | 3 | 1 | | 1 |
| 739 | 282 | 77 | 6:7:1:0:0 | 6 | 7 | 1 | 0 | 0 | 1 | 1 |
| 740 | 282 | 77 | 6:7:1:0:1 | 6 | 7 | 1 | 0 | 1 | 1 | 1 |
| 741 | 282 | 77 | 6:7:1:1:0 | 6 | 7 | 1 | 1 | 0 | | 1 |
| 742 | 282 | 77 | 6:7:2:0:0 | 6 | 7 | 2 | 0 | 0 | 1 | |
| 743 | 282 | 77 | 6:7:3:0:0 | 6 | 7 | 3 | 0 | 0 | 1 | |
| 744 | 282 | 77 | 6:7:4:0:0 | 6 | 7 | 4 | 0 | 0 | 1 | 1 |
| 745 | 282 | 77 | 6:7:5:0:0 | 6 | 7 | 5 | 0 | 0 | | 1 |
| 746 | 282 | 77 | 6:8:0:1:0 | 6 | 8 | 0 | 1 | 0 | 1 | 1 |
| 747 | 282 | 77 | 6:8:1:1:0 | 6 | 8 | 1 | 1 | 0 | | 1 |
| 748 | 282 | 77 | 7:6:1:0:0 | 7 | 6 | 1 | 0 | 0 | | 1 |
| 749 | 282 | 77 | 7:7:1:0:0 | 7 | 7 | 1 | 0 | 0 | 1 | 1 |
| 750 | 282 | 77 | 7:7:2:0:0 | 7 | 7 | 2 | 0 | 0 | 1 | |
| 751 | 282 | 77 | 7:7:3:0:0 | 7 | 7 | 3 | 0 | 0 | 1 | |
| 752 | 282 | 77 | 9:3:0:0:0 | 9 | 3 | 0 | 0 | 0 | | 1 |
| 753 | 331 | 56 | 2:5:0:0:0 | 2 | 5 | 0 | 0 | 0 | | 1 |
| 754 | 331 | 56 | 2:7:0:0:0 | 2 | 7 | 0 | 0 | 0 | | 1 |
| 755 | 331 | 56 | 3:4:1:0:1 | 3 | 4 | 1 | 0 | 1 | | 1 |
| 756 | 331 | 56 | 3:5:2:0:0 | 3 | 5 | 2 | 0 | 0 | 1 | |
| 757 | 331 | 56 | 4:3:1:0:0 | 4 | 3 | 1 | 0 | 0 | 1 | |
| 758 | 331 | 56 | 4:4:2:0:0 | 4 | 4 | 2 | 0 | 0 | 1 | |
| 759 | 331 | 56 | 4:4:2:0:1 | 4 | 4 | 2 | 0 | 1 | 1 | |
| 760 | 331 | 56 | 4:5:1:0:0 | 4 | 5 | 1 | 0 | 0 | 1 | 1 |
| 761 | 331 | 56 | 4:5:1:0:1 | 4 | 5 | 1 | 0 | 1 | 1 | 1 |
| 762 | 331 | 56 | 4:5:1:1:0 | 4 | 5 | 1 | 1 | 0 | 1 | 1 |
| 763 | 331 | 56 | 4:5:2:0:0 | 4 | 5 | 2 | 0 | 0 | 1 | 1 |
| 764 | 331 | 56 | 4:5:2:0:1 | 4 | 5 | 2 | 0 | 1 | 1 | |
| 765 | 331 | 56 | 4:5:3:0:0 | 4 | 5 | 3 | 0 | 0 | 1 | 1 |
| 766 | 331 | 56 | 4:6:0:1:0 | 4 | 6 | 0 | 1 | 0 | 1 | |
| 767 | 331 | 56 | 4:6:1:1:0 | 4 | 6 | 1 | 1 | 0 | 1 | 1 |
| 768 | 331 | 56 | 4:6:2:0:0 | 4 | 6 | 2 | 0 | 0 | | 1 |
| 769 | 331 | 56 | 5:3:1:0:0 | 5 | 3 | 1 | 0 | 0 | 1 | |
| 770 | 331 | 56 | 5:4:1:0:0 | 5 | 4 | 1 | 0 | 0 | 1 | 1 |
| 771 | 331 | 56 | 5:4:3:0:0 | 5 | 4 | 3 | 0 | 0 | 1 | |
| 772 | 331 | 56 | 5:5:1:0:0 | 5 | 5 | 1 | 0 | 0 | 1 | 1 |
| 773 | 331 | 56 | 5:5:1:0:1 | 5 | 5 | 1 | 0 | 1 | 1 | |
| 774 | 331 | 56 | 5:5:1:1:0 | 5 | 5 | 1 | 1 | 0 | | 1 |
| 775 | 331 | 56 | 5:5:2:0:0 | 5 | 5 | 2 | 0 | 0 | 1 | |
| 776 | 331 | 56 | 5:5:2:0:1 | 5 | 5 | 2 | 0 | 1 | 1 | |

| | | | | | | | | | | |
|---|---|---|---|---|---|---|---|---|---|---|
| 777 | 331 | 56 | 5:5:3:0:0 | 5 | 5 | 3 | 0 | 0 | 1 | 1 |
| 778 | 331 | 56 | 5:6:0:1:0 | 5 | 6 | 0 | 1 | 0 | 1 | |
| 779 | 331 | 56 | 5:6:1:0:0 | 5 | 6 | 1 | 0 | 0 | 1 | 1 |
| 780 | 331 | 56 | 5:6:1:0:1 | 5 | 6 | 1 | 0 | 1 | 1 | 1 |
| 781 | 331 | 56 | 5:6:1:0:2 | 5 | 6 | 1 | 0 | 2 | 1 | |
| 782 | 331 | 56 | 5:6:1:1:0 | 5 | 6 | 1 | 1 | 0 | 1 | 1 |
| 783 | 331 | 56 | 5:6:2:0:0 | 5 | 6 | 2 | 0 | 0 | 1 | 1 |
| 784 | 331 | 56 | 5:6:2:1:0 | 5 | 6 | 2 | 1 | 0 | | 1 |
| 785 | 331 | 56 | 5:6:3:0:0 | 5 | 6 | 3 | 0 | 0 | 1 | 1 |
| 786 | 331 | 56 | 5:6:3:0:1 | 5 | 6 | 3 | 0 | 1 | 1 | |
| 787 | 331 | 56 | 5:6:4:0:0 | 5 | 6 | 4 | 0 | 0 | 1 | |
| 788 | 331 | 56 | 5:7:0:1:0 | 5 | 7 | 0 | 1 | 0 | 1 | |
| 789 | 331 | 56 | 5:7:0:1:1 | 5 | 7 | 0 | 1 | 1 | 1 | |
| 790 | 331 | 56 | 5:7:2:1:0 | 5 | 7 | 2 | 1 | 0 | 1 | |
| 791 | 331 | 56 | 6:6:1:0:0 | 6 | 6 | 1 | 0 | 0 | 1 | 1 |
| 792 | 331 | 56 | 6:6:1:0:1 | 6 | 6 | 1 | 0 | 1 | 1 | |
| 793 | 331 | 56 | 6:6:2:0:0 | 6 | 6 | 2 | 0 | 0 | 1 | 1 |
| 794 | 331 | 56 | 6:6:3:0:0 | 6 | 6 | 3 | 0 | 0 | 1 | |
| 795 | 331 | 56 | 6:6:4:0:0 | 6 | 6 | 4 | 0 | 0 | 1 | |
| 796 | 331 | 56 | 6:7:0:1:0 | 6 | 7 | 0 | 1 | 0 | 1 | 1 |
| 797 | 331 | 56 | 6:7:1:0:0 | 6 | 7 | 1 | 0 | 0 | 1 | |
| 798 | 331 | 56 | 6:7:1:0:1 | 6 | 7 | 1 | 0 | 1 | 1 | |
| 799 | 331 | 56 | 6:7:1:1:0 | 6 | 7 | 1 | 1 | 0 | | 1 |
| 800 | 331 | 56 | 6:7:2:0:0 | 6 | 7 | 2 | 0 | 0 | 1 | |
| 801 | 331 | 56 | 6:7:2:0:1 | 6 | 7 | 2 | 0 | 1 | 1 | |
| 802 | 331 | 56 | 6:7:3:0:0 | 6 | 7 | 3 | 0 | 0 | 1 | |
| 803 | 331 | 56 | 6:7:4:0:0 | 6 | 7 | 4 | 0 | 0 | 1 | 1 |
| 804 | 331 | 56 | 6:7:4:0:1 | 6 | 7 | 4 | 0 | 1 | 1 | |
| 805 | 331 | 56 | 6:8:0:1:0 | 6 | 8 | 0 | 1 | 0 | 1 | |
| 806 | 331 | 56 | 6:8:1:1:0 | 6 | 8 | 1 | 1 | 0 | 1 | |
| 807 | 331 | 56 | 6:8:3:1:0 | 6 | 8 | 3 | 1 | 0 | 1 | |
| 808 | 331 | 56 | 8:9:0:0:0 | 8 | 9 | 0 | 0 | 0 | 1 | 1 |
| 809 | 343 | 190 | 2:5:0:0:0 | 2 | 5 | 0 | 0 | 0 | | 1 |
| 810 | 343 | 190 | 2:6:0:0:0 | 2 | 6 | 0 | 0 | 0 | | 1 |
| 811 | 343 | 190 | 3:4:2:0:0 | 3 | 4 | 2 | 0 | 0 | 1 | |
| 812 | 343 | 190 | 3:5:2:0:0 | 3 | 5 | 2 | 0 | 0 | 1 | |
| 813 | 343 | 190 | 3:6:0:1:0 | 3 | 6 | 0 | 1 | 0 | | 1 |
| 814 | 343 | 190 | 3:6:1:0:0 | 3 | 6 | 1 | 0 | 0 | 1 | |
| 815 | 343 | 190 | 3:6:2:0:0 | 3 | 6 | 2 | 0 | 0 | 1 | |
| 816 | 343 | 190 | 4:3:1:0:0 | 4 | 3 | 1 | 0 | 0 | 1 | |
| 817 | 343 | 190 | 4:4:1:0:0 | 4 | 4 | 1 | 0 | 0 | 1 | 1 |
| 818 | 343 | 190 | 4:4:2:0:0 | 4 | 4 | 2 | 0 | 0 | 1 | |
| 819 | 343 | 190 | 4:4:2:0:1 | 4 | 4 | 2 | 0 | 1 | 1 | |
| 820 | 343 | 190 | 4:4:3:0:1 | 4 | 4 | 3 | 0 | 1 | 1 | |
| 821 | 343 | 190 | 4:5:0:0:0 | 4 | 5 | 0 | 0 | 0 | | 1 |
| 822 | 343 | 190 | 4:5:1:0:0 | 4 | 5 | 1 | 0 | 0 | 1 | 1 |
| 823 | 343 | 190 | 4:5:1:0:1 | 4 | 5 | 1 | 0 | 1 | 1 | 1 |
| 824 | 343 | 190 | 4:5:1:0:2 | 4 | 5 | 1 | 0 | 2 | 1 | |
| 825 | 343 | 190 | 4:5:1:1:0 | 4 | 5 | 1 | 1 | 0 | 1 | 1 |
| 826 | 343 | 190 | 4:5:2:0:0 | 4 | 5 | 2 | 0 | 0 | 1 | 1 |
| 827 | 343 | 190 | 4:5:2:0:1 | 4 | 5 | 2 | 0 | 1 | 1 | 1 |
| 828 | 343 | 190 | 4:5:2:1:0 | 4 | 5 | 2 | 1 | 0 | 1 | 1 |
| 829 | 343 | 190 | 4:5:3:0:0 | 4 | 5 | 3 | 0 | 0 | 1 | 1 |
| 830 | 343 | 190 | 4:5:3:1:0 | 4 | 5 | 3 | 1 | 0 | | 1 |
| 831 | 343 | 190 | 4:5:3:2:0 | 4 | 5 | 3 | 2 | 0 | | 1 |
| 832 | 343 | 190 | 4:6:0:1:0 | 4 | 6 | 0 | 1 | 0 | 1 | |
| 833 | 343 | 190 | 4:6:0:1:1 | 4 | 6 | 0 | 1 | 1 | 1 | |
| 834 | 343 | 190 | 4:6:1:1:0 | 4 | 6 | 1 | 1 | 0 | 1 | |
| 835 | 343 | 190 | 4:7:2:0:0 | 4 | 7 | 2 | 0 | 0 | | 1 |
| 836 | 343 | 190 | 5:3:1:0:0 | 5 | 3 | 1 | 0 | 0 | 1 | 1 |
| 837 | 343 | 190 | 5:4:0:0:0 | 5 | 4 | 0 | 0 | 0 | | 1 |
| 838 | 343 | 190 | 5:4:1:0:0 | 5 | 4 | 1 | 0 | 0 | 1 | 1 |
| 839 | 343 | 190 | 5:4:2:0:0 | 5 | 4 | 2 | 0 | 0 | 1 | 1 |
| 840 | 343 | 190 | 5:4:2:0:2 | 5 | 4 | 2 | 0 | 2 | 1 | |
| 841 | 343 | 190 | 5:4:3:0:0 | 5 | 4 | 3 | 0 | 0 | 1 | |

| | | | | | | | | | | |
|---|---|---|---|---|---|---|---|---|---|---|
| 842 | 343 | 190 | 5:4:3:0:1 | 5 | 4 | 3 | 0 | 1 | 1 | |
| 843 | 343 | 190 | 5:5:0:0:0 | 5 | 5 | 0 | 0 | 0 | | 1 |
| 844 | 343 | 190 | 5:5:1:0:0 | 5 | 5 | 1 | 0 | 0 | 1 | 1 |
| 845 | 343 | 190 | 5:5:1:0:1 | 5 | 5 | 1 | 0 | 1 | 1 | 1 |
| 846 | 343 | 190 | 5:5:1:1:1 | 5 | 5 | 1 | 1 | 1 | 1 | |
| 847 | 343 | 190 | 5:5:2:0:0 | 5 | 5 | 2 | 0 | 0 | 1 | 1 |
| 848 | 343 | 190 | 5:5:2:0:1 | 5 | 5 | 2 | 0 | 1 | 1 | |
| 849 | 343 | 190 | 5:5:2:0:2 | 5 | 5 | 2 | 0 | 2 | 1 | |
| 850 | 343 | 190 | 5:5:2:1:0 | 5 | 5 | 2 | 1 | 0 | 1 | |
| 851 | 343 | 190 | 5:5:3:0:0 | 5 | 5 | 3 | 0 | 0 | 1 | 1 |
| 852 | 343 | 190 | 5:5:4:0:0 | 5 | 5 | 4 | 0 | 0 | 1 | |
| 853 | 343 | 190 | 5:6:0:0:0 | 5 | 6 | 0 | 0 | 0 | | 1 |
| 854 | 343 | 190 | 5:6:0:1:0 | 5 | 6 | 0 | 1 | 0 | 1 | |
| 855 | 343 | 190 | 5:6:0:2:0 | 5 | 6 | 0 | 2 | 0 | 1 | |
| 856 | 343 | 190 | 5:6:1:0:0 | 5 | 6 | 1 | 0 | 0 | 1 | |
| 857 | 343 | 190 | 5:6:1:0:1 | 5 | 6 | 1 | 0 | 1 | 1 | 1 |
| 858 | 343 | 190 | 5:6:1:1:0 | 5 | 6 | 1 | 1 | 0 | 1 | 1 |
| 859 | 343 | 190 | 5:6:1:1:1 | 5 | 6 | 1 | 1 | 1 | 1 | 1 |
| 860 | 343 | 190 | 5:6:1:2:1 | 5 | 6 | 1 | 2 | 1 | | 1 |
| 861 | 343 | 190 | 5:6:2:0:0 | 5 | 6 | 2 | 0 | 0 | 1 | 1 |
| 862 | 343 | 190 | 5:6:2:0:1 | 5 | 6 | 2 | 0 | 1 | 1 | 1 |
| 863 | 343 | 190 | 5:6:2:0:2 | 5 | 6 | 2 | 0 | 2 | 1 | |
| 864 | 343 | 190 | 5:6:2:1:0 | 5 | 6 | 2 | 1 | 0 | | 1 |
| 865 | 343 | 190 | 5:6:3:0:0 | 5 | 6 | 3 | 0 | 0 | 1 | 1 |
| 866 | 343 | 190 | 5:6:3:0:1 | 5 | 6 | 3 | 0 | 1 | 1 | 1 |
| 867 | 343 | 190 | 5:6:4:0:0 | 5 | 6 | 4 | 0 | 0 | 1 | |
| 868 | 343 | 190 | 5:7:0:1:0 | 5 | 7 | 0 | 1 | 0 | 1 | |
| 869 | 343 | 190 | 5:7:0:2:0 | 5 | 7 | 0 | 2 | 0 | 1 | |
| 870 | 343 | 190 | 5:7:1:1:0 | 5 | 7 | 1 | 1 | 0 | 1 | |
| 871 | 343 | 190 | 5:7:1:1:1 | 5 | 7 | 1 | 1 | 1 | 1 | |
| 872 | 343 | 190 | 5:7:1:2:0 | 5 | 7 | 1 | 2 | 0 | | 1 |
| 873 | 343 | 190 | 5:7:2:0:2 | 5 | 7 | 2 | 0 | 2 | 1 | |
| 874 | 343 | 190 | 5:7:2:1:0 | 5 | 7 | 2 | 1 | 0 | 1 | |
| 875 | 343 | 190 | 5:8:0:2:0 | 5 | 8 | 0 | 2 | 0 | 1 | |
| 876 | 343 | 190 | 5:8:1:1:1 | 5 | 8 | 1 | 1 | 1 | 1 | |
| 877 | 343 | 190 | 5:8:2:0:2 | 5 | 8 | 2 | 0 | 2 | 1 | |
| 878 | 343 | 190 | 5:9:0:2:0 | 5 | 9 | 0 | 2 | 0 | 1 | |
| 879 | 343 | 190 | 5:9:1:1:1 | 5 | 9 | 1 | 1 | 1 | 1 | |
| 880 | 343 | 190 | 5:9:2:0:2 | 5 | 9 | 2 | 0 | 2 | 1 | |
| 881 | 343 | 190 | 6:3:1:0:0 | 6 | 3 | 1 | 0 | 0 | | 1 |
| 882 | 343 | 190 | 6:5:1:0:0 | 6 | 5 | 1 | 0 | 0 | 1 | 1 |
| 883 | 343 | 190 | 6:5:2:0:0 | 6 | 5 | 2 | 0 | 0 | 1 | 1 |
| 884 | 343 | 190 | 6:5:2:0:1 | 6 | 5 | 2 | 0 | 1 | 1 | |
| 885 | 343 | 190 | 6:5:2:0:2 | 6 | 5 | 2 | 0 | 2 | 1 | |
| 886 | 343 | 190 | 6:5:3:0:2 | 6 | 5 | 3 | 0 | 2 | 1 | |
| 887 | 343 | 190 | 6:6:1:0:0 | 6 | 6 | 1 | 0 | 0 | 1 | 1 |
| 888 | 343 | 190 | 6:6:1:0:1 | 6 | 6 | 1 | 0 | 1 | 1 | |
| 889 | 343 | 190 | 6:6:1:1:0 | 6 | 6 | 1 | 1 | 0 | 1 | |
| 890 | 343 | 190 | 6:6:1:1:1 | 6 | 6 | 1 | 1 | 1 | 1 | |
| 891 | 343 | 190 | 6:6:2:0:0 | 6 | 6 | 2 | 0 | 0 | 1 | 1 |
| 892 | 343 | 190 | 6:6:2:0:2 | 6 | 6 | 2 | 0 | 2 | 1 | |
| 893 | 343 | 190 | 6:6:2:1:1 | 6 | 6 | 2 | 1 | 1 | 1 | |
| 894 | 343 | 190 | 6:6:3:0:0 | 6 | 6 | 3 | 0 | 0 | 1 | |
| 895 | 343 | 190 | 6:6:3:0:1 | 6 | 6 | 3 | 0 | 1 | 1 | |
| 896 | 343 | 190 | 6:6:3:0:2 | 6 | 6 | 3 | 0 | 2 | 1 | |
| 897 | 343 | 190 | 6:6:4:0:0 | 6 | 6 | 4 | 0 | 0 | 1 | |
| 898 | 343 | 190 | 6:7:0:1:0 | 6 | 7 | 0 | 1 | 0 | 1 | |
| 899 | 343 | 190 | 6:7:0:1:1 | 6 | 7 | 0 | 1 | 1 | | 1 |
| 900 | 343 | 190 | 6:7:0:2:0 | 6 | 7 | 0 | 2 | 0 | 1 | |
| 901 | 343 | 190 | 6:7:1:0:0 | 6 | 7 | 1 | 0 | 0 | 1 | 1 |
| 902 | 343 | 190 | 6:7:1:0:1 | 6 | 7 | 1 | 0 | 1 | 1 | |
| 903 | 343 | 190 | 6:7:1:1:1 | 6 | 7 | 1 | 1 | 1 | 1 | |
| 904 | 343 | 190 | 6:7:1:2:0 | 6 | 7 | 1 | 2 | 0 | 1 | |
| 905 | 343 | 190 | 6:7:2:0:0 | 6 | 7 | 2 | 0 | 0 | 1 | |
| 906 | 343 | 190 | 6:7:2:0:2 | 6 | 7 | 2 | 0 | 2 | 1 | |

| | | | | | | | | | | |
|---|---|---|---|---|---|---|---|---|---|---|
| 907 | 343 | 190 | 6:7:2:1:0 | 6 | 7 | 2 | 1 | 0 | 1 | |
| 908 | 343 | 190 | 6:7:2:1:1 | 6 | 7 | 2 | 1 | 1 | 1 | |
| 909 | 343 | 190 | 6:7:3:0:0 | 6 | 7 | 3 | 0 | 0 | 1 | |
| 910 | 343 | 190 | 6:7:4:0:0 | 6 | 7 | 4 | 0 | 0 | 1 | 1 |
| 911 | 343 | 190 | 6:7:4:0:1 | 6 | 7 | 4 | 0 | 1 | 1 | 1 |
| 912 | 343 | 190 | 6:7:5:0:0 | 6 | 7 | 5 | 0 | 0 | | 1 |
| 913 | 343 | 190 | 6:8:0:1:0 | 6 | 8 | 0 | 1 | 0 | 1 | |
| 914 | 343 | 190 | 6:8:0:2:0 | 6 | 8 | 0 | 2 | 0 | 1 | |
| 915 | 343 | 190 | 6:8:1:1:0 | 6 | 8 | 1 | 1 | 0 | | 1 |
| 916 | 343 | 190 | 6:8:1:1:1 | 6 | 8 | 1 | 1 | 1 | 1 | |
| 917 | 343 | 190 | 6:8:1:2:0 | 6 | 8 | 1 | 2 | 0 | 1 | |
| 918 | 343 | 190 | 6:8:2:0:2 | 6 | 8 | 2 | 0 | 2 | 1 | |
| 919 | 343 | 190 | 6:8:3:0:2 | 6 | 8 | 3 | 0 | 2 | 1 | |
| 920 | 343 | 190 | 6:8:3:1:0 | 6 | 8 | 3 | 1 | 0 | 1 | |
| 921 | 343 | 190 | 6:9:0:2:0 | 6 | 9 | 0 | 2 | 0 | 1 | |
| 922 | 343 | 190 | 6:9:1:1:1 | 6 | 9 | 1 | 1 | 1 | 1 | |
| 923 | 343 | 190 | 6:9:1:2:0 | 6 | 9 | 1 | 2 | 0 | | 1 |
| 924 | 343 | 190 | 6:9:2:1:1 | 6 | 9 | 2 | 1 | 1 | 1 | |
| 925 | 343 | 190 | 6:10:0:2:0 | 6 | 10 | 0 | 2 | 0 | 1 | |
| 926 | 343 | 190 | 6:10:1:2:0 | 6 | 10 | 1 | 2 | 0 | 1 | |
| 927 | 343 | 190 | 7:5:2:0:2 | 7 | 5 | 2 | 0 | 2 | 1 | |
| 928 | 343 | 190 | 7:5:3:0:2 | 7 | 5 | 3 | 0 | 2 | 1 | |
| 929 | 343 | 190 | 7:5:4:0:2 | 7 | 5 | 4 | 0 | 2 | 1 | |
| 930 | 343 | 190 | 7:6:1:1:1 | 7 | 6 | 1 | 1 | 1 | 1 | |
| 931 | 343 | 190 | 7:6:1:3:1 | 7 | 6 | 1 | 3 | 1 | | 1 |
| 932 | 343 | 190 | 7:6:2:1:1 | 7 | 6 | 2 | 1 | 1 | 1 | |
| 933 | 343 | 190 | 7:6:3:0:2 | 7 | 6 | 3 | 0 | 2 | 1 | |
| 934 | 343 | 190 | 7:6:3:1:1 | 7 | 6 | 3 | 1 | 1 | 1 | |
| 935 | 343 | 190 | 7:6:4:0:2 | 7 | 6 | 4 | 0 | 2 | 1 | |
| 936 | 343 | 190 | 7:7:0:2:0 | 7 | 7 | 0 | 2 | 0 | 1 | |
| 937 | 343 | 190 | 7:7:1:1:2 | 7 | 7 | 1 | 1 | 2 | | 1 |
| 938 | 343 | 190 | 7:7:1:2:0 | 7 | 7 | 1 | 2 | 0 | 1 | |
| 939 | 343 | 190 | 7:7:2:1:1 | 7 | 7 | 2 | 1 | 1 | 1 | |
| 940 | 343 | 190 | 7:7:2:2:0 | 7 | 7 | 2 | 2 | 0 | 1 | |
| 941 | 343 | 190 | 7:7:3:0:0 | 7 | 7 | 3 | 0 | 0 | 1 | |
| 942 | 343 | 190 | 7:7:3:1:1 | 7 | 7 | 3 | 1 | 1 | 1 | |
| 943 | 343 | 190 | 7:7:4:0:0 | 7 | 7 | 4 | 0 | 0 | 1 | |
| 944 | 343 | 190 | 7:8:1:1:2 | 7 | 8 | 1 | 1 | 2 | | 1 |
| 945 | 343 | 190 | 7:8:1:2:0 | 7 | 8 | 1 | 2 | 0 | 1 | 1 |
| 946 | 343 | 190 | 7:8:2:2:0 | 7 | 8 | 2 | 2 | 0 | 1 | |
| 947 | 343 | 190 | 8:4:2:0:2 | 8 | 4 | 2 | 0 | 2 | 1 | |
| 948 | 343 | 190 | 8:4:3:0:2 | 8 | 4 | 3 | 0 | 2 | 1 | |
| 949 | 343 | 190 | 8:5:0:0:0 | 8 | 5 | 0 | 0 | 0 | | 1 |
| 950 | 343 | 190 | 8:5:1:1:1 | 8 | 5 | 1 | 1 | 1 | 1 | |
| 951 | 343 | 190 | 8:5:2:0:2 | 8 | 5 | 2 | 0 | 2 | 1 | |
| 952 | 343 | 190 | 8:5:2:1:1 | 8 | 5 | 2 | 1 | 1 | 1 | |
| 953 | 343 | 190 | 8:5:3:0:2 | 8 | 5 | 3 | 0 | 2 | 1 | |
| 954 | 343 | 190 | 8:6:0:2:0 | 8 | 6 | 0 | 2 | 0 | 1 | |
| 955 | 343 | 190 | 8:6:1:1:1 | 8 | 6 | 1 | 1 | 1 | 1 | |
| 956 | 343 | 190 | 8:6:1:2:0 | 8 | 6 | 1 | 2 | 0 | 1 | |
| 957 | 343 | 190 | 8:6:2:1:1 | 8 | 6 | 2 | 1 | 1 | 1 | |
| 958 | 343 | 190 | 8:6:3:0:2 | 8 | 6 | 3 | 0 | 2 | 1 | |
| 959 | 343 | 190 | 8:7:0:2:0 | 8 | 7 | 0 | 2 | 0 | 1 | |
| 960 | 343 | 190 | 8:7:1:2:0 | 8 | 7 | 1 | 2 | 0 | 1 | |
| 961 | 343 | 190 | 8:7:2:1:1 | 8 | 7 | 2 | 1 | 1 | 1 | |
| 962 | 343 | 190 | 8:7:3:0:2 | 8 | 7 | 3 | 0 | 2 | 1 | |
| 963 | 343 | 190 | 8:8:1:2:0 | 8 | 8 | 1 | 2 | 0 | 1 | |
| 964 | 343 | 190 | 8:8:2:1:1 | 8 | 8 | 2 | 1 | 1 | 1 | |
| 965 | 343 | 190 | 8:9:1:1:2 | 8 | 9 | 1 | 1 | 2 | | 1 |
| 966 | 343 | 190 | 8:9:1:2:0 | 8 | 9 | 1 | 2 | 0 | 1 | |
| 967 | 343 | 190 | 8:9:1:2:1 | 8 | 9 | 1 | 2 | 1 | | 1 |
| 968 | 343 | 190 | 9:4:3:0:2 | 9 | 4 | 3 | 0 | 2 | 1 | |
| 969 | 343 | 190 | 9:5:2:0:1 | 9 | 5 | 2 | 0 | 1 | 1 | |
| 970 | 343 | 190 | 9:5:2:1:1 | 9 | 5 | 2 | 1 | 1 | 1 | |
| 971 | 343 | 190 | 9:5:3:0:2 | 9 | 5 | 3 | 0 | 2 | 1 | |

| | | | | | | | | | | |
|---|---|---|---|---|---|---|---|---|---|---|
| 972 | 343 | 190 | 9:6:1:1:0 | 9 | 6 | 1 | 1 | 0 | 1 | |
| 973 | 343 | 190 | 9:6:1:2:0 | 9 | 6 | 1 | 2 | 0 | 1 | |
| 974 | 343 | 190 | 9:6:2:1:1 | 9 | 6 | 2 | 1 | 1 | 1 | |
| 975 | 343 | 190 | 9:6:3:0:2 | 9 | 6 | 3 | 0 | 2 | 1 | |
| 976 | 343 | 190 | 9:7:1:2:0 | 9 | 7 | 1 | 2 | 0 | 1 | |
| 977 | 343 | 190 | 9:7:2:1:1 | 9 | 7 | 2 | 1 | 1 | 1 | |
| 978 | 343 | 190 | 9:7:3:0:2 | 9 | 7 | 3 | 0 | 2 | 1 | |
| 979 | 343 | 190 | 9:8:0:0:0 | 9 | 8 | 0 | 0 | 0 | 1 | |
| 980 | 343 | 190 | 9:8:1:2:0 | 9 | 8 | 1 | 2 | 0 | 1 | |
| 981 | 343 | 190 | 9:8:2:1:1 | 9 | 8 | 2 | 1 | 1 | 1 | |
| 982 | 343 | 190 | 9:8:3:0:2 | 9 | 8 | 3 | 0 | 2 | 1 | |
| 983 | 343 | 190 | 9:8:4:0:2 | 9 | 8 | 4 | 0 | 2 | 1 | |
| 984 | 343 | 190 | 9:9:0:0:0 | 9 | 9 | 0 | 0 | 0 | 1 | |
| 985 | 343 | 190 | 9:9:1:2:0 | 9 | 9 | 1 | 2 | 0 | 1 | |
| 986 | 343 | 190 | 9:9:2:1:1 | 9 | 9 | 2 | 1 | 1 | 1 | |
| 987 | 343 | 190 | 9:9:3:1:1 | 9 | 9 | 3 | 1 | 1 | 1 | |
| 988 | 343 | 190 | 9:10:0:0:0 | 9 | 10 | 0 | 0 | 0 | 1 | |
| 989 | 343 | 190 | 9:10:1:0:0 | 9 | 10 | 1 | 0 | 0 | 1 | |
| 990 | 343 | 190 | 9:10:1:2:0 | 9 | 10 | 1 | 2 | 0 | 1 | |
| 991 | 343 | 190 | 9:10:2:2:0 | 9 | 10 | 2 | 2 | 0 | 1 | |
| 992 | 343 | 190 | 10:6:3:0:2 | 10 | 6 | 3 | 0 | 2 | 1 | |
| 993 | 343 | 190 | 10:7:2:1:1 | 10 | 7 | 2 | 1 | 1 | 1 | |
| 994 | 343 | 190 | 10:8:1:2:0 | 10 | 8 | 1 | 2 | 0 | 1 | |
| 995 | 343 | 190 | 10:9:0:0:0 | 10 | 9 | 0 | 0 | 0 | 1 | |
| 996 | 343 | 190 | 10:9:1:0:0 | 10 | 9 | 1 | 0 | 0 | 1 | |
| 997 | 343 | 190 | 10:10:0:0:0 | 10 | 10 | 0 | 0 | 0 | 1 | |
| 998 | 343 | 190 | 10:10:1:0:0 | 10 | 10 | 1 | 0 | 0 | 1 | 1 |
| 999 | 603 | 36 | 2:5:0:0:0 | 2 | 5 | 0 | 0 | 0 | 1 | 1 |
| 1000 | 603 | 36 | 2:6:0:0:0 | 2 | 6 | 0 | 0 | 0 | 1 | 1 |
| 1001 | 603 | 36 | 2:7:0:0:0 | 2 | 7 | 0 | 0 | 0 | 1 | 1 |
| 1002 | 603 | 36 | 2:8:0:0:0 | 2 | 8 | 0 | 0 | 0 | 1 | 1 |
| 1003 | 603 | 36 | 3:4:0:1:0 | 3 | 4 | 0 | 1 | 0 | | 1 |
| 1004 | 603 | 36 | 3:5:0:0:0 | 3 | 5 | 0 | 0 | 0 | 1 | 1 |
| 1005 | 603 | 36 | 3:5:1:0:0 | 3 | 5 | 1 | 0 | 0 | 1 | 1 |
| 1006 | 603 | 36 | 3:5:2:0:0 | 3 | 5 | 2 | 0 | 0 | 1 | |
| 1007 | 603 | 36 | 3:6:0:0:0 | 3 | 6 | 0 | 0 | 0 | 1 | 1 |
| 1008 | 603 | 36 | 3:6:1:0:0 | 3 | 6 | 1 | 0 | 0 | 1 | 1 |
| 1009 | 603 | 36 | 4:3:0:1:0 | 4 | 3 | 0 | 1 | 0 | | 1 |
| 1010 | 603 | 36 | 4:3:1:0:0 | 4 | 3 | 1 | 0 | 0 | 1 | 1 |
| 1011 | 603 | 36 | 4:4:0:0:0 | 4 | 4 | 0 | 0 | 0 | | 1 |
| 1012 | 603 | 36 | 4:4:1:0:0 | 4 | 4 | 1 | 0 | 0 | 1 | 1 |
| 1013 | 603 | 36 | 4:5:0:0:0 | 4 | 5 | 0 | 0 | 0 | 1 | 1 |
| 1014 | 603 | 36 | 4:5:1:0:0 | 4 | 5 | 1 | 0 | 0 | 1 | 1 |
| 1015 | 603 | 36 | 4:5:1:1:0 | 4 | 5 | 1 | 1 | 0 | | 1 |
| 1016 | 603 | 36 | 4:5:2:0:0 | 4 | 5 | 2 | 0 | 0 | 1 | 1 |
| 1017 | 603 | 36 | 4:5:3:0:0 | 4 | 5 | 3 | 0 | 0 | 1 | |
| 1018 | 603 | 36 | 4:5:3:1:0 | 4 | 5 | 3 | 1 | 0 | | 1 |
| 1019 | 603 | 36 | 5:4:1:0:0 | 5 | 4 | 1 | 0 | 0 | 1 | 1 |
| 1020 | 603 | 36 | 5:5:1:0:0 | 5 | 5 | 1 | 0 | 0 | 1 | 1 |
| 1021 | 603 | 36 | 5:5:1:1:0 | 5 | 5 | 1 | 1 | 0 | | 1 |
| 1022 | 603 | 36 | 5:6:0:1:0 | 5 | 6 | 0 | 1 | 0 | | 1 |
| 1023 | 603 | 36 | 5:6:1:1:0 | 5 | 6 | 1 | 1 | 0 | | 1 |
| 1024 | 603 | 36 | 5:6:2:0:0 | 5 | 6 | 2 | 0 | 0 | 1 | 1 |
| 1025 | 603 | 36 | 5:6:2:1:0 | 5 | 6 | 2 | 1 | 0 | | 1 |
| 1026 | 603 | 36 | 5:6:3:0:0 | 5 | 6 | 3 | 0 | 0 | 1 | 1 |
| 1027 | 603 | 36 | 5:6:4:0:0 | 5 | 6 | 4 | 0 | 0 | 1 | |
| 1028 | 603 | 36 | 5:7:1:0:0 | 5 | 7 | 1 | 0 | 0 | | 1 |
| 1029 | 603 | 36 | 6:6:1:0:0 | 6 | 6 | 1 | 0 | 0 | 1 | |
| 1030 | 603 | 36 | 6:7:1:0:0 | 6 | 7 | 1 | 0 | 0 | 1 | 1 |
| 1031 | 603 | 36 | 6:7:2:0:0 | 6 | 7 | 2 | 0 | 0 | 1 | |
| 1032 | 603 | 36 | 6:7:3:0:0 | 6 | 7 | 3 | 0 | 0 | 1 | |
| 1033 | 603 | 36 | 6:7:4:0:0 | 6 | 7 | 4 | 0 | 0 | 1 | 1 |
| 1034 | 603 | 36 | 7:3:0:0:0 | 7 | 3 | 0 | 0 | 0 | | 1 |
| 1035 | 616 | 57 | 2:4:0:0:0 | 2 | 4 | 0 | 0 | 0 | | 1 |
| 1036 | 616 | 57 | 2:5:0:0:0 | 2 | 5 | 0 | 0 | 0 | 1 | 1 |

| | | | | | | | | | | |
|---|---|---|---|---|---|---|---|---|---|---|
| 1037 | 616 | 57 | 3:4:0:0:0 | 3 | 4 | 0 | 0 | 0 | 1 | |
| 1038 | 616 | 57 | 3:4:1:0:1 | 3 | 4 | 1 | 0 | 1 | 1 | |
| 1039 | 616 | 57 | 3:4:1:1:0 | 3 | 4 | 1 | 1 | 0 | | 1 |
| 1040 | 616 | 57 | 3:5:0:0:0 | 3 | 5 | 0 | 0 | 0 | 1 | 1 |
| 1041 | 616 | 57 | 3:5:0:1:0 | 3 | 5 | 0 | 1 | 0 | 1 | |
| 1042 | 616 | 57 | 3:5:1:0:0 | 3 | 5 | 1 | 0 | 0 | 1 | 1 |
| 1043 | 616 | 57 | 3:6:0:0:0 | 3 | 6 | 0 | 0 | 0 | | 1 |
| 1044 | 616 | 57 | 3:6:1:0:0 | 3 | 6 | 1 | 0 | 0 | | 1 |
| 1045 | 616 | 57 | 4:4:0:0:1 | 4 | 4 | 0 | 0 | 1 | | 1 |
| 1046 | 616 | 57 | 4:4:1:0:0 | 4 | 4 | 1 | 0 | 0 | | 1 |
| 1047 | 616 | 57 | 4:5:0:0:0 | 4 | 5 | 0 | 0 | 0 | 1 | 1 |
| 1048 | 616 | 57 | 4:5:1:0:0 | 4 | 5 | 1 | 0 | 0 | 1 | 1 |
| 1049 | 616 | 57 | 4:5:1:0:1 | 4 | 5 | 1 | 0 | 1 | 1 | |
| 1050 | 616 | 57 | 4:5:1:1:0 | 4 | 5 | 1 | 1 | 0 | | 1 |
| 1051 | 616 | 57 | 4:5:2:0:0 | 4 | 5 | 2 | 0 | 0 | 1 | 1 |
| 1052 | 616 | 57 | 4:5:3:0:0 | 4 | 5 | 3 | 0 | 0 | 1 | |
| 1053 | 616 | 57 | 4:5:3:1:0 | 4 | 5 | 3 | 1 | 0 | | 1 |
| 1054 | 616 | 57 | 4:6:0:0:0 | 4 | 6 | 0 | 0 | 0 | | 1 |
| 1055 | 616 | 57 | 4:6:0:1:0 | 4 | 6 | 0 | 1 | 0 | 1 | |
| 1056 | 616 | 57 | 4:6:1:0:0 | 4 | 6 | 1 | 0 | 0 | | 1 |
| 1057 | 616 | 57 | 4:6:1:1:0 | 4 | 6 | 1 | 1 | 0 | | 1 |
| 1058 | 616 | 57 | 5:4:0:0:0 | 5 | 4 | 0 | 0 | 0 | | 1 |
| 1059 | 616 | 57 | 5:4:0:0:1 | 5 | 4 | 0 | 0 | 1 | | 1 |
| 1060 | 616 | 57 | 5:4:0:1:1 | 5 | 4 | 0 | 1 | 1 | | 1 |
| 1061 | 616 | 57 | 5:4:1:0:0 | 5 | 4 | 1 | 0 | 0 | | 1 |
| 1062 | 616 | 57 | 5:5:0:0:0 | 5 | 5 | 0 | 0 | 0 | 1 | 1 |
| 1063 | 616 | 57 | 5:5:0:1:0 | 5 | 5 | 0 | 1 | 0 | | 1 |
| 1064 | 616 | 57 | 5:5:0:1:1 | 5 | 5 | 0 | 1 | 1 | | 1 |
| 1065 | 616 | 57 | 5:5:1:0:0 | 5 | 5 | 1 | 0 | 0 | 1 | 1 |
| 1066 | 616 | 57 | 5:5:2:0:0 | 5 | 5 | 2 | 0 | 0 | 1 | |
| 1067 | 616 | 57 | 5:5:3:0:0 | 5 | 5 | 3 | 0 | 0 | | 1 |
| 1068 | 616 | 57 | 5:6:0:0:0 | 5 | 6 | 0 | 0 | 0 | 1 | 1 |
| 1069 | 616 | 57 | 5:6:0:1:0 | 5 | 6 | 0 | 1 | 0 | | 1 |
| 1070 | 616 | 57 | 5:6:0:1:2 | 5 | 6 | 0 | 1 | 2 | | 1 |
| 1071 | 616 | 57 | 5:6:1:0:0 | 5 | 6 | 1 | 0 | 0 | 1 | 1 |
| 1072 | 616 | 57 | 5:6:1:0:1 | 5 | 6 | 1 | 0 | 1 | | 1 |
| 1073 | 616 | 57 | 5:6:1:1:0 | 5 | 6 | 1 | 1 | 0 | | 1 |
| 1074 | 616 | 57 | 5:6:2:0:0 | 5 | 6 | 2 | 0 | 0 | 1 | |
| 1075 | 616 | 57 | 5:6:3:0:0 | 5 | 6 | 3 | 0 | 0 | 1 | |
| 1076 | 616 | 57 | 5:6:4:0:0 | 5 | 6 | 4 | 0 | 0 | 1 | |
| 1077 | 616 | 57 | 5:7:0:0:0 | 5 | 7 | 0 | 0 | 0 | | 1 |
| 1078 | 616 | 57 | 6:3:1:0:0 | 6 | 3 | 1 | 0 | 0 | | 1 |
| 1079 | 616 | 57 | 6:5:1:1:1 | 6 | 5 | 1 | 1 | 1 | | 1 |
| 1080 | 616 | 57 | 6:6:0:1:0 | 6 | 6 | 0 | 1 | 0 | | 1 |
| 1081 | 616 | 57 | 6:6:1:0:0 | 6 | 6 | 1 | 0 | 0 | 1 | 1 |
| 1082 | 616 | 57 | 6:6:2:0:0 | 6 | 6 | 2 | 0 | 0 | | 1 |
| 1083 | 616 | 57 | 6:7:0:0:0 | 6 | 7 | 0 | 0 | 0 | 1 | 1 |
| 1084 | 616 | 57 | 6:7:0:1:0 | 6 | 7 | 0 | 1 | 0 | | 1 |
| 1085 | 616 | 57 | 6:7:1:0:0 | 6 | 7 | 1 | 0 | 0 | 1 | 1 |
| 1086 | 616 | 57 | 6:7:1:0:1 | 6 | 7 | 1 | 0 | 1 | 1 | |
| 1087 | 616 | 57 | 6:7:1:1:0 | 6 | 7 | 1 | 1 | 0 | | 1 |
| 1088 | 616 | 57 | 6:7:2:0:0 | 6 | 7 | 2 | 0 | 0 | 1 | |
| 1089 | 616 | 57 | 6:7:3:0:0 | 6 | 7 | 3 | 0 | 0 | 1 | |
| 1090 | 616 | 57 | 6:8:0:1:0 | 6 | 8 | 0 | 1 | 0 | 1 | |
| 1091 | 616 | 57 | 7:7:1:0:0 | 7 | 7 | 1 | 0 | 0 | | 1 |
| 1092 | 657 | 55 | 2:6:0:0:0 | 2 | 6 | 0 | 0 | 0 | | 1 |
| 1093 | 657 | 55 | 2:7:0:0:0 | 2 | 7 | 0 | 0 | 0 | | 1 |
| 1094 | 657 | 55 | 2:9:0:0:0 | 2 | 9 | 0 | 0 | 0 | | 1 |
| 1095 | 657 | 55 | 3:3:2:0:0 | 3 | 3 | 2 | 0 | 0 | | 1 |
| 1096 | 657 | 55 | 3:5:0:0:1 | 3 | 5 | 0 | 0 | 1 | | 1 |
| 1097 | 657 | 55 | 3:5:1:0:0 | 3 | 5 | 1 | 0 | 0 | | 1 |
| 1098 | 657 | 55 | 3:5:2:0:0 | 3 | 5 | 2 | 0 | 0 | 1 | |
| 1099 | 657 | 55 | 3:6:1:0:0 | 3 | 6 | 1 | 0 | 0 | | 1 |
| 1100 | 657 | 55 | 3:6:1:1:0 | 3 | 6 | 1 | 1 | 0 | | 1 |
| 1101 | 657 | 55 | 4:5:0:1:0 | 4 | 5 | 0 | 1 | 0 | | 1 |

| | | | | | | | | | | |
|---|---|---|---|---|---|---|---|---|---|---|
| 1102 | 657 | 55 | 4:5:1:0:0 | 4 | 5 | 1 | 0 | 0 | 1 | 1 |
| 1103 | 657 | 55 | 4:5:1:1:0 | 4 | 5 | 1 | 1 | 0 | | 1 |
| 1104 | 657 | 55 | 4:5:1:2:0 | 4 | 5 | 1 | 2 | 0 | | 1 |
| 1105 | 657 | 55 | 4:5:2:0:0 | 4 | 5 | 2 | 0 | 0 | 1 | |
| 1106 | 657 | 55 | 4:5:3:0:0 | 4 | 5 | 3 | 0 | 0 | 1 | 1 |
| 1107 | 657 | 55 | 4:6:0:1:0 | 4 | 6 | 0 | 1 | 0 | | 1 |
| 1108 | 657 | 55 | 4:6:2:0:0 | 4 | 6 | 2 | 0 | 0 | | 1 |
| 1109 | 657 | 55 | 5:4:1:2:0 | 5 | 4 | 1 | 2 | 0 | | 1 |
| 1110 | 657 | 55 | 5:4:2:1:0 | 5 | 4 | 2 | 1 | 0 | | 1 |
| 1111 | 657 | 55 | 5:5:0:0:1 | 5 | 5 | 0 | 0 | 1 | | 1 |
| 1112 | 657 | 55 | 5:5:0:1:0 | 5 | 5 | 0 | 1 | 0 | | 1 |
| 1113 | 657 | 55 | 5:5:1:0:0 | 5 | 5 | 1 | 0 | 0 | 1 | 1 |
| 1114 | 657 | 55 | 5:5:1:1:0 | 5 | 5 | 1 | 1 | 0 | | 1 |
| 1115 | 657 | 55 | 5:5:2:0:0 | 5 | 5 | 2 | 0 | 0 | 1 | 1 |
| 1116 | 657 | 55 | 5:5:3:0:0 | 5 | 5 | 3 | 0 | 0 | 1 | 1 |
| 1117 | 657 | 55 | 5:6:0:0:0 | 5 | 6 | 0 | 0 | 0 | | 1 |
| 1118 | 657 | 55 | 5:6:0:1:0 | 5 | 6 | 0 | 1 | 0 | | 1 |
| 1119 | 657 | 55 | 5:6:1:0:0 | 5 | 6 | 1 | 0 | 0 | | 1 |
| 1120 | 657 | 55 | 5:6:2:0:0 | 5 | 6 | 2 | 0 | 0 | | 1 |
| 1121 | 657 | 55 | 5:6:2:0:1 | 5 | 6 | 2 | 0 | 1 | | 1 |
| 1122 | 657 | 55 | 5:6:2:1:0 | 5 | 6 | 2 | 1 | 0 | | 1 |
| 1123 | 657 | 55 | 5:6:3:0:0 | 5 | 6 | 3 | 0 | 0 | 1 | |
| 1124 | 657 | 55 | 6:3:1:2:0 | 6 | 3 | 1 | 2 | 0 | | 1 |
| 1125 | 657 | 55 | 6:3:2:0:1 | 6 | 3 | 2 | 0 | 1 | | 1 |
| 1126 | 657 | 55 | 6:4:1:0:0 | 6 | 4 | 1 | 0 | 0 | | 1 |
| 1127 | 657 | 55 | 6:5:1:1:1 | 6 | 5 | 1 | 1 | 1 | | 1 |
| 1128 | 657 | 55 | 6:6:0:2:0 | 6 | 6 | 0 | 2 | 0 | | 1 |
| 1129 | 657 | 55 | 6:6:1:0:0 | 6 | 6 | 1 | 0 | 0 | | 1 |
| 1130 | 657 | 55 | 6:6:2:0:0 | 6 | 6 | 2 | 0 | 0 | 1 | 1 |
| 1131 | 657 | 55 | 6:6:3:0:0 | 6 | 6 | 3 | 0 | 0 | 1 | 1 |
| 1132 | 657 | 55 | 6:6:4:0:0 | 6 | 6 | 4 | 0 | 0 | 1 | |
| 1133 | 657 | 55 | 6:7:0:1:0 | 6 | 7 | 0 | 1 | 0 | | 1 |
| 1134 | 657 | 55 | 6:7:1:0:0 | 6 | 7 | 1 | 0 | 0 | | 1 |
| 1135 | 657 | 55 | 6:7:1:1:0 | 6 | 7 | 1 | 1 | 0 | | 1 |
| 1136 | 657 | 55 | 6:7:2:0:0 | 6 | 7 | 2 | 0 | 0 | 1 | |
| 1137 | 657 | 55 | 6:7:2:0:1 | 6 | 7 | 2 | 0 | 1 | 1 | |
| 1138 | 657 | 55 | 6:7:3:0:0 | 6 | 7 | 3 | 0 | 0 | 1 | |
| 1139 | 657 | 55 | 6:7:3:0:1 | 6 | 7 | 3 | 0 | 1 | 1 | |
| 1140 | 657 | 55 | 6:7:4:0:0 | 6 | 7 | 4 | 0 | 0 | 1 | 1 |
| 1141 | 657 | 55 | 6:7:4:1:0 | 6 | 7 | 4 | 1 | 0 | | 1 |
| 1142 | 657 | 55 | 6:8:1:1:0 | 6 | 8 | 1 | 1 | 0 | 1 | |
| 1143 | 657 | 55 | 6:8:2:1:0 | 6 | 8 | 2 | 1 | 0 | 1 | |
| 1144 | 657 | 55 | 7:7:2:0:0 | 7 | 7 | 2 | 0 | 0 | 1 | |
| 1145 | 657 | 55 | 7:7:3:0:0 | 7 | 7 | 3 | 0 | 0 | 1 | |
| 1146 | 657 | 55 | 7:8:1:0:0 | 7 | 8 | 1 | 0 | 0 | | 1 |
| 1147 | 709 | 28 | 2:4:0:0:0 | 2 | 4 | 0 | 0 | 0 | | 1 |
| 1148 | 709 | 28 | 2:5:0:0:0 | 2 | 5 | 0 | 0 | 0 | 1 | 1 |
| 1149 | 709 | 28 | 2:6:0:0:0 | 2 | 6 | 0 | 0 | 0 | 1 | 1 |
| 1150 | 709 | 28 | 2:7:0:0:0 | 2 | 7 | 0 | 0 | 0 | 1 | 1 |
| 1151 | 709 | 28 | 2:8:0:0:0 | 2 | 8 | 0 | 0 | 0 | 1 | 1 |
| 1152 | 709 | 28 | 3:5:1:0:0 | 3 | 5 | 1 | 0 | 0 | | 1 |
| 1153 | 709 | 28 | 3:6:0:0:0 | 3 | 6 | 0 | 0 | 0 | 1 | 1 |
| 1154 | 709 | 28 | 4:3:0:1:0 | 4 | 3 | 0 | 1 | 0 | | 1 |
| 1155 | 709 | 28 | 4:5:0:0:0 | 4 | 5 | 0 | 0 | 0 | 1 | 1 |
| 1156 | 709 | 28 | 4:5:1:0:0 | 4 | 5 | 1 | 0 | 0 | 1 | 1 |
| 1157 | 709 | 28 | 5:4:0:0:0 | 5 | 4 | 0 | 0 | 0 | 1 | |
| 1158 | 709 | 28 | 5:4:1:0:0 | 5 | 4 | 1 | 0 | 0 | 1 | |
| 1159 | 709 | 28 | 5:6:0:2:1 | 5 | 6 | 0 | 2 | 1 | | 1 |
| 1160 | 709 | 28 | 5:6:1:0:0 | 5 | 6 | 1 | 0 | 0 | 1 | 1 |
| 1161 | 709 | 28 | 5:6:2:0:0 | 5 | 6 | 2 | 0 | 0 | 1 | |
| 1162 | 709 | 28 | 6:5:1:0:0 | 6 | 5 | 1 | 0 | 0 | 1 | 1 |
| 1163 | 709 | 28 | 6:6:1:0:0 | 6 | 6 | 1 | 0 | 0 | 1 | |
| 1164 | 709 | 28 | 6:7:0:2:0 | 6 | 7 | 0 | 2 | 0 | | 1 |
| 1165 | 709 | 28 | 6:7:1:0:0 | 6 | 7 | 1 | 0 | 0 | 1 | |
| 1166 | 709 | 28 | 6:7:1:0:1 | 6 | 7 | 1 | 0 | 1 | 1 | |

| | | | | | | | | | | |
|---|---|---|---|---|---|---|---|---|---|---|
| 1167 | 709 | 28 | 6:7:2:0:0 | 6 | 7 | 2 | 0 | 0 | 1 | |
| 1168 | 709 | 28 | 6:7:3:0:0 | 6 | 7 | 3 | 0 | 0 | 1 | |
| 1169 | 709 | 28 | 6:7:4:0:0 | 6 | 7 | 4 | 0 | 0 | 1 | |
| 1170 | 709 | 28 | 6:7:4:1:0 | 6 | 7 | 4 | 1 | 0 | | 1 |
| 1171 | 709 | 28 | 6:8:0:1:0 | 6 | 8 | 0 | 1 | 0 | 1 | |
| 1172 | 709 | 28 | 6:9:1:1:1 | 6 | 9 | 1 | 1 | 1 | | 1 |
| 1173 | 709 | 28 | 6:10:1:1:0 | 6 | 10 | 1 | 1 | 0 | | 1 |
| 1174 | 709 | 28 | 7:6:1:0:0 | 7 | 6 | 1 | 0 | 0 | 1 | |
| 1175 | 717 | 18 | 2:5:0:0:0 | 2 | 5 | 0 | 0 | 0 | | 1 |
| 1176 | 717 | 18 | 2:6:0:0:0 | 2 | 6 | 0 | 0 | 0 | 1 | 1 |
| 1177 | 717 | 18 | 2:7:0:0:0 | 2 | 7 | 0 | 0 | 0 | 1 | 1 |
| 1178 | 717 | 18 | 2:8:0:0:0 | 2 | 8 | 0 | 0 | 0 | 1 | 1 |
| 1179 | 717 | 18 | 2:9:0:0:0 | 2 | 9 | 0 | 0 | 0 | | 1 |
| 1180 | 717 | 18 | 3:5:0:0:0 | 3 | 5 | 0 | 0 | 0 | 1 | 1 |
| 1181 | 717 | 18 | 3:5:1:0:0 | 3 | 5 | 1 | 0 | 0 | 1 | 1 |
| 1182 | 717 | 18 | 3:6:0:0:0 | 3 | 6 | 0 | 0 | 0 | 1 | 1 |
| 1183 | 717 | 18 | 3:6:0:0:1 | 3 | 6 | 0 | 0 | 1 | 1 | 1 |
| 1184 | 717 | 18 | 3:6:1:0:0 | 3 | 6 | 1 | 0 | 0 | 1 | 1 |
| 1185 | 717 | 18 | 4:5:0:0:0 | 4 | 5 | 0 | 0 | 0 | | 1 |
| 1186 | 717 | 18 | 6:5:1:1:2 | 6 | 5 | 1 | 1 | 2 | | 1 |
| 1187 | 717 | 18 | 6:6:0:1:0 | 6 | 6 | 0 | 1 | 0 | | 1 |
| 1188 | 717 | 18 | 6:7:1:2:0 | 6 | 7 | 1 | 2 | 0 | | 1 |
| 1189 | 717 | 18 | 6:8:1:1:0 | 6 | 8 | 1 | 1 | 0 | | 1 |
| 1190 | 717 | 18 | 7:8:1:1:0 | 7 | 8 | 1 | 1 | 0 | | 1 |
| 1191 | 717 | 18 | 7:8:1:1:1 | 7 | 8 | 1 | 1 | 1 | | 1 |
| 1192 | 717 | 18 | 8:9:1:2:1 | 8 | 9 | 1 | 2 | 1 | | 1 |
| 1193 | 801 | 92 | 2:4:0:0:0 | 2 | 4 | 0 | 0 | 0 | 1 | 1 |
| 1194 | 801 | 92 | 2:5:0:0:0 | 2 | 5 | 0 | 0 | 0 | 1 | 1 |
| 1195 | 801 | 92 | 2:5:1:0:0 | 2 | 5 | 1 | 0 | 0 | | 1 |
| 1196 | 801 | 92 | 2:6:0:0:0 | 2 | 6 | 0 | 0 | 0 | 1 | 1 |
| 1197 | 801 | 92 | 2:7:0:0:0 | 2 | 7 | 0 | 0 | 0 | 1 | 1 |
| 1198 | 801 | 92 | 2:8:0:0:0 | 2 | 8 | 0 | 0 | 0 | 1 | 1 |
| 1199 | 801 | 92 | 2:9:0:0:0 | 2 | 9 | 0 | 0 | 0 | 1 | 1 |
| 1200 | 801 | 92 | 3:4:0:0:0 | 3 | 4 | 0 | 0 | 0 | 1 | |
| 1201 | 801 | 92 | 3:4:0:0:1 | 3 | 4 | 0 | 0 | 1 | | 1 |
| 1202 | 801 | 92 | 3:4:1:0:0 | 3 | 4 | 1 | 0 | 0 | | 1 |
| 1203 | 801 | 92 | 3:4:1:0:1 | 3 | 4 | 1 | 0 | 1 | 1 | |
| 1204 | 801 | 92 | 3:4:1:1:0 | 3 | 4 | 1 | 1 | 0 | | 1 |
| 1205 | 801 | 92 | 3:4:2:0:0 | 3 | 4 | 2 | 0 | 0 | | 1 |
| 1206 | 801 | 92 | 3:5:0:0:0 | 3 | 5 | 0 | 0 | 0 | 1 | 1 |
| 1207 | 801 | 92 | 3:5:0:0:1 | 3 | 5 | 0 | 0 | 1 | 1 | 1 |
| 1208 | 801 | 92 | 3:5:0:1:0 | 3 | 5 | 0 | 1 | 0 | 1 | 1 |
| 1209 | 801 | 92 | 3:5:1:0:0 | 3 | 5 | 1 | 0 | 0 | 1 | 1 |
| 1210 | 801 | 92 | 3:5:1:0:1 | 3 | 5 | 1 | 0 | 1 | 1 | |
| 1211 | 801 | 92 | 3:5:2:0:0 | 3 | 5 | 2 | 0 | 0 | 1 | |
| 1212 | 801 | 92 | 3:6:0:0:0 | 3 | 6 | 0 | 0 | 0 | 1 | 1 |
| 1213 | 801 | 92 | 3:6:0:0:1 | 3 | 6 | 0 | 0 | 1 | 1 | 1 |
| 1214 | 801 | 92 | 3:6:0:1:0 | 3 | 6 | 0 | 1 | 0 | 1 | 1 |
| 1215 | 801 | 92 | 3:6:1:0:0 | 3 | 6 | 1 | 0 | 0 | 1 | 1 |
| 1216 | 801 | 92 | 3:7:1:0:0 | 3 | 7 | 1 | 0 | 0 | 1 | |
| 1217 | 801 | 92 | 4:3:0:0:0 | 4 | 3 | 0 | 0 | 0 | 1 | 1 |
| 1218 | 801 | 92 | 4:3:1:0:0 | 4 | 3 | 1 | 0 | 0 | 1 | 1 |
| 1219 | 801 | 92 | 4:4:0:0:0 | 4 | 4 | 0 | 0 | 0 | 1 | |
| 1220 | 801 | 92 | 4:4:1:0:0 | 4 | 4 | 1 | 0 | 0 | 1 | 1 |
| 1221 | 801 | 92 | 4:4:1:0:1 | 4 | 4 | 1 | 0 | 1 | 1 | |
| 1222 | 801 | 92 | 4:4:2:0:0 | 4 | 4 | 2 | 0 | 0 | 1 | 1 |
| 1223 | 801 | 92 | 4:4:2:0:1 | 4 | 4 | 2 | 0 | 1 | 1 | |
| 1224 | 801 | 92 | 4:4:2:0:2 | 4 | 4 | 2 | 0 | 2 | 1 | |
| 1225 | 801 | 92 | 4:5:0:0:0 | 4 | 5 | 0 | 0 | 0 | 1 | 1 |
| 1226 | 801 | 92 | 4:5:0:0:1 | 4 | 5 | 0 | 0 | 1 | 1 | 1 |
| 1227 | 801 | 92 | 4:5:0:0:2 | 4 | 5 | 0 | 0 | 2 | 1 | |
| 1228 | 801 | 92 | 4:5:0:1:0 | 4 | 5 | 0 | 1 | 0 | 1 | 1 |
| 1229 | 801 | 92 | 4:5:1:0:0 | 4 | 5 | 1 | 0 | 0 | 1 | 1 |
| 1230 | 801 | 92 | 4:5:1:0:1 | 4 | 5 | 1 | 0 | 1 | 1 | 1 |
| 1231 | 801 | 92 | 4:5:1:1:0 | 4 | 5 | 1 | 1 | 0 | 1 | 1 |

| | | | | | | | | | | |
|---|---|---|---|---|---|---|---|---|---|---|
| 1232 | 801 | 92 | 4:5:1:1:1 | 4 | 5 | 1 | 1 | 1 | 1 | |
| 1233 | 801 | 92 | 4:5:2:0:0 | 4 | 5 | 2 | 0 | 0 | 1 | 1 |
| 1234 | 801 | 92 | 4:5:2:0:1 | 4 | 5 | 2 | 0 | 1 | 1 | 1 |
| 1235 | 801 | 92 | 4:5:3:0:0 | 4 | 5 | 3 | 0 | 0 | 1 | 1 |
| 1236 | 801 | 92 | 4:6:0:0:0 | 4 | 6 | 0 | 0 | 0 | 1 | 1 |
| 1237 | 801 | 92 | 4:6:0:1:0 | 4 | 6 | 0 | 1 | 0 | 1 | 1 |
| 1238 | 801 | 92 | 4:6:0:2:0 | 4 | 6 | 0 | 2 | 0 | 1 | |
| 1239 | 801 | 92 | 4:6:1:0:0 | 4 | 6 | 1 | 0 | 0 | | 1 |
| 1240 | 801 | 92 | 4:6:1:1:0 | 4 | 6 | 1 | 1 | 0 | 1 | |
| 1241 | 801 | 92 | 4:6:2:0:0 | 4 | 6 | 2 | 0 | 0 | 1 | 1 |
| 1242 | 801 | 92 | 4:6:3:0:0 | 4 | 6 | 3 | 0 | 0 | | 1 |
| 1243 | 801 | 92 | 4:7:0:0:0 | 4 | 7 | 0 | 0 | 0 | 1 | 1 |
| 1244 | 801 | 92 | 4:7:1:0:0 | 4 | 7 | 1 | 0 | 0 | 1 | |
| 1245 | 801 | 92 | 4:7:2:0:0 | 4 | 7 | 2 | 0 | 0 | 1 | |
| 1246 | 801 | 92 | 5:3:0:0:0 | 5 | 3 | 0 | 0 | 0 | 1 | 1 |
| 1247 | 801 | 92 | 5:3:1:0:0 | 5 | 3 | 1 | 0 | 0 | 1 | 1 |
| 1248 | 801 | 92 | 5:3:1:1:0 | 5 | 3 | 1 | 1 | 0 | | 1 |
| 1249 | 801 | 92 | 5:4:0:0:0 | 5 | 4 | 0 | 0 | 0 | 1 | 1 |
| 1250 | 801 | 92 | 5:4:1:0:0 | 5 | 4 | 1 | 0 | 0 | 1 | 1 |
| 1251 | 801 | 92 | 5:4:1:0:1 | 5 | 4 | 1 | 0 | 1 | 1 | |
| 1252 | 801 | 92 | 5:4:2:0:0 | 5 | 4 | 2 | 0 | 0 | 1 | 1 |
| 1253 | 801 | 92 | 5:5:0:0:0 | 5 | 5 | 0 | 0 | 0 | 1 | 1 |
| 1254 | 801 | 92 | 5:5:0:1:0 | 5 | 5 | 0 | 1 | 0 | 1 | 1 |
| 1255 | 801 | 92 | 5:5:0:1:1 | 5 | 5 | 0 | 1 | 1 | | 1 |
| 1256 | 801 | 92 | 5:5:1:0:0 | 5 | 5 | 1 | 0 | 0 | 1 | 1 |
| 1257 | 801 | 92 | 5:5:1:0:1 | 5 | 5 | 1 | 0 | 1 | 1 | |
| 1258 | 801 | 92 | 5:5:2:0:0 | 5 | 5 | 2 | 0 | 0 | 1 | 1 |
| 1259 | 801 | 92 | 5:5:2:0:1 | 5 | 5 | 2 | 0 | 1 | 1 | |
| 1260 | 801 | 92 | 5:5:3:0:0 | 5 | 5 | 3 | 0 | 0 | 1 | 1 |
| 1261 | 801 | 92 | 5:6:0:0:0 | 5 | 6 | 0 | 0 | 0 | 1 | 1 |
| 1262 | 801 | 92 | 5:6:0:1:0 | 5 | 6 | 0 | 1 | 0 | 1 | 1 |
| 1263 | 801 | 92 | 5:6:0:1:2 | 5 | 6 | 0 | 1 | 2 | 1 | |
| 1264 | 801 | 92 | 5:6:1:0:0 | 5 | 6 | 1 | 0 | 0 | 1 | 1 |
| 1265 | 801 | 92 | 5:6:1:0:1 | 5 | 6 | 1 | 0 | 1 | 1 | 1 |
| 1266 | 801 | 92 | 5:6:1:1:0 | 5 | 6 | 1 | 1 | 0 | 1 | 1 |
| 1267 | 801 | 92 | 5:6:2:0:0 | 5 | 6 | 2 | 0 | 0 | 1 | 1 |
| 1268 | 801 | 92 | 5:6:2:1:0 | 5 | 6 | 2 | 1 | 0 | | 1 |
| 1269 | 801 | 92 | 5:6:3:0:0 | 5 | 6 | 3 | 0 | 0 | 1 | 1 |
| 1270 | 801 | 92 | 5:6:4:0:0 | 5 | 6 | 4 | 0 | 0 | 1 | |
| 1271 | 801 | 92 | 5:7:0:1:0 | 5 | 7 | 0 | 1 | 0 | 1 | |
| 1272 | 801 | 92 | 5:7:1:0:0 | 5 | 7 | 1 | 0 | 0 | | 1 |
| 1273 | 801 | 92 | 6:3:1:0:0 | 6 | 3 | 1 | 0 | 0 | 1 | 1 |
| 1274 | 801 | 92 | 6:4:1:0:0 | 6 | 4 | 1 | 0 | 0 | | 1 |
| 1275 | 801 | 92 | 6:5:1:0:0 | 6 | 5 | 1 | 0 | 0 | 1 | 1 |
| 1276 | 801 | 92 | 6:6:0:0:0 | 6 | 6 | 0 | 0 | 0 | 1 | 1 |
| 1277 | 801 | 92 | 6:6:0:1:0 | 6 | 6 | 0 | 1 | 0 | | 1 |
| 1278 | 801 | 92 | 6:6:1:0:0 | 6 | 6 | 1 | 0 | 0 | 1 | 1 |
| 1279 | 801 | 92 | 6:6:1:1:0 | 6 | 6 | 1 | 1 | 0 | | 1 |
| 1280 | 801 | 92 | 6:6:2:0:0 | 6 | 6 | 2 | 0 | 0 | 1 | |
| 1281 | 801 | 92 | 6:6:3:0:0 | 6 | 6 | 3 | 0 | 0 | 1 | 1 |
| 1282 | 801 | 92 | 6:7:1:3:1 | 6 | 7 | 1 | 3 | 1 | | 1 |
| 1283 | 801 | 92 | 6:9:1:1:1 | 6 | 9 | 1 | 1 | 1 | | 1 |
| 1284 | 801 | 92 | 7:3:0:0:0 | 7 | 3 | 0 | 0 | 0 | | 1 |
| 1285 | 1074 | 169 | 2:0:1:0:0 | 2 | 0 | 1 | 0 | 0 | | 1 |
| 1286 | 1074 | 169 | 2:3:0:0:0 | 2 | 3 | 0 | 0 | 0 | 1 | 1 |
| 1287 | 1074 | 169 | 2:4:0:0:0 | 2 | 4 | 0 | 0 | 0 | 1 | 1 |
| 1288 | 1074 | 169 | 2:5:0:0:0 | 2 | 5 | 0 | 0 | 0 | 1 | 1 |
| 1289 | 1074 | 169 | 2:6:0:0:0 | 2 | 6 | 0 | 0 | 0 | 1 | 1 |
| 1290 | 1074 | 169 | 2:7:0:0:0 | 2 | 7 | 0 | 0 | 0 | 1 | 1 |
| 1291 | 1074 | 169 | 2:8:0:0:0 | 2 | 8 | 0 | 0 | 0 | 1 | 1 |
| 1292 | 1074 | 169 | 2:9:0:0:0 | 2 | 9 | 0 | 0 | 0 | 1 | 1 |
| 1293 | 1074 | 169 | 2:11:0:0:0 | 2 | 11 | 0 | 0 | 0 | | 1 |
| 1294 | 1074 | 169 | 3:3:0:0:0 | 3 | 3 | 0 | 0 | 0 | 1 | 1 |
| 1295 | 1074 | 169 | 3:3:1:0:0 | 3 | 3 | 1 | 0 | 0 | 1 | 1 |
| 1296 | 1074 | 169 | 3:4:0:0:0 | 3 | 4 | 0 | 0 | 0 | 1 | 1 |

| | | | | | | | | | | |
|---|---|---|---|---|---|---|---|---|---|---|
| 1297 | 1074 | 169 | 3:4:0:0:1 | 3 | 4 | 0 | 0 | 1 | 1 | |
| 1298 | 1074 | 169 | 3:4:1:0:0 | 3 | 4 | 1 | 0 | 0 | 1 | 1 |
| 1299 | 1074 | 169 | 3:4:1:0:1 | 3 | 4 | 1 | 0 | 1 | 1 | 1 |
| 1300 | 1074 | 169 | 3:4:1:1:0 | 3 | 4 | 1 | 1 | 0 | | 1 |
| 1301 | 1074 | 169 | 3:4:2:0:0 | 3 | 4 | 2 | 0 | 0 | 1 | 1 |
| 1302 | 1074 | 169 | 3:4:2:0:1 | 3 | 4 | 2 | 0 | 1 | 1 | |
| 1303 | 1074 | 169 | 3:5:0:0:0 | 3 | 5 | 0 | 0 | 0 | 1 | 1 |
| 1304 | 1074 | 169 | 3:5:0:0:1 | 3 | 5 | 0 | 0 | 1 | 1 | 1 |
| 1305 | 1074 | 169 | 3:5:0:1:0 | 3 | 5 | 0 | 1 | 0 | 1 | 1 |
| 1306 | 1074 | 169 | 3:5:1:0:0 | 3 | 5 | 1 | 0 | 0 | 1 | 1 |
| 1307 | 1074 | 169 | 3:5:1:0:1 | 3 | 5 | 1 | 0 | 1 | 1 | |
| 1308 | 1074 | 169 | 3:5:1:1:0 | 3 | 5 | 1 | 1 | 0 | 1 | |
| 1309 | 1074 | 169 | 3:5:2:0:0 | 3 | 5 | 2 | 0 | 0 | 1 | |
| 1310 | 1074 | 169 | 3:6:0:0:0 | 3 | 6 | 0 | 0 | 0 | 1 | 1 |
| 1311 | 1074 | 169 | 3:6:0:0:1 | 3 | 6 | 0 | 0 | 1 | 1 | 1 |
| 1312 | 1074 | 169 | 3:6:0:1:0 | 3 | 6 | 0 | 1 | 0 | 1 | 1 |
| 1313 | 1074 | 169 | 3:6:1:0:0 | 3 | 6 | 1 | 0 | 0 | 1 | 1 |
| 1314 | 1074 | 169 | 3:6:1:0:1 | 3 | 6 | 1 | 0 | 1 | 1 | 1 |
| 1315 | 1074 | 169 | 3:6:2:0:0 | 3 | 6 | 2 | 0 | 0 | 1 | |
| 1316 | 1074 | 169 | 3:7:0:0:0 | 3 | 7 | 0 | 0 | 0 | 1 | |
| 1317 | 1074 | 169 | 3:7:1:0:0 | 3 | 7 | 1 | 0 | 0 | 1 | |
| 1318 | 1074 | 169 | 4:3:0:0:0 | 4 | 3 | 0 | 0 | 0 | 1 | 1 |
| 1319 | 1074 | 169 | 4:3:1:0:0 | 4 | 3 | 1 | 0 | 0 | 1 | 1 |
| 1320 | 1074 | 169 | 4:3:2:0:1 | 4 | 3 | 2 | 0 | 1 | 1 | |
| 1321 | 1074 | 169 | 4:4:0:0:0 | 4 | 4 | 0 | 0 | 0 | 1 | 1 |
| 1322 | 1074 | 169 | 4:4:1:0:0 | 4 | 4 | 1 | 0 | 0 | 1 | 1 |
| 1323 | 1074 | 169 | 4:4:1:0:1 | 4 | 4 | 1 | 0 | 1 | 1 | |
| 1324 | 1074 | 169 | 4:4:1:1:0 | 4 | 4 | 1 | 1 | 0 | 1 | |
| 1325 | 1074 | 169 | 4:4:2:0:0 | 4 | 4 | 2 | 0 | 0 | 1 | |
| 1326 | 1074 | 169 | 4:4:2:0:1 | 4 | 4 | 2 | 0 | 1 | 1 | |
| 1327 | 1074 | 169 | 4:4:3:0:1 | 4 | 4 | 3 | 0 | 1 | 1 | |
| 1328 | 1074 | 169 | 4:5:0:0:0 | 4 | 5 | 0 | 0 | 0 | 1 | 1 |
| 1329 | 1074 | 169 | 4:5:0:0:1 | 4 | 5 | 0 | 0 | 1 | 1 | 1 |
| 1330 | 1074 | 169 | 4:5:0:1:0 | 4 | 5 | 0 | 1 | 0 | 1 | |
| 1331 | 1074 | 169 | 4:5:0:1:1 | 4 | 5 | 0 | 1 | 1 | | 1 |
| 1332 | 1074 | 169 | 4:5:0:2:0 | 4 | 5 | 0 | 2 | 0 | | 1 |
| 1333 | 1074 | 169 | 4:5:1:0:0 | 4 | 5 | 1 | 0 | 0 | 1 | 1 |
| 1334 | 1074 | 169 | 4:5:1:0:1 | 4 | 5 | 1 | 0 | 1 | 1 | 1 |
| 1335 | 1074 | 169 | 4:5:1:1:0 | 4 | 5 | 1 | 1 | 0 | 1 | 1 |
| 1336 | 1074 | 169 | 4:5:1:1:1 | 4 | 5 | 1 | 1 | 1 | | 1 |
| 1337 | 1074 | 169 | 4:5:1:2:0 | 4 | 5 | 1 | 2 | 0 | | 1 |
| 1338 | 1074 | 169 | 4:5:2:0:0 | 4 | 5 | 2 | 0 | 0 | 1 | 1 |
| 1339 | 1074 | 169 | 4:5:2:0:1 | 4 | 5 | 2 | 0 | 1 | 1 | 1 |
| 1340 | 1074 | 169 | 4:5:2:1:0 | 4 | 5 | 2 | 1 | 0 | 1 | |
| 1341 | 1074 | 169 | 4:5:3:0:0 | 4 | 5 | 3 | 0 | 0 | 1 | 1 |
| 1342 | 1074 | 169 | 4:5:3:1:0 | 4 | 5 | 3 | 1 | 0 | | 1 |
| 1343 | 1074 | 169 | 4:5:3:2:0 | 4 | 5 | 3 | 2 | 0 | | 1 |
| 1344 | 1074 | 169 | 4:6:0:0:0 | 4 | 6 | 0 | 0 | 0 | 1 | 1 |
| 1345 | 1074 | 169 | 4:6:0:1:0 | 4 | 6 | 0 | 1 | 0 | 1 | 1 |
| 1346 | 1074 | 169 | 4:6:1:0:0 | 4 | 6 | 1 | 0 | 0 | 1 | 1 |
| 1347 | 1074 | 169 | 4:6:1:0:1 | 4 | 6 | 1 | 0 | 1 | 1 | |
| 1348 | 1074 | 169 | 4:6:1:1:0 | 4 | 6 | 1 | 1 | 0 | 1 | |
| 1349 | 1074 | 169 | 4:6:2:0:0 | 4 | 6 | 2 | 0 | 0 | 1 | 1 |
| 1350 | 1074 | 169 | 4:6:2:0:1 | 4 | 6 | 2 | 0 | 1 | 1 | |
| 1351 | 1074 | 169 | 4:6:3:0:0 | 4 | 6 | 3 | 0 | 0 | 1 | 1 |
| 1352 | 1074 | 169 | 4:7:0:0:0 | 4 | 7 | 0 | 0 | 0 | 1 | 1 |
| 1353 | 1074 | 169 | 4:7:0:1:0 | 4 | 7 | 0 | 1 | 0 | 1 | 1 |
| 1354 | 1074 | 169 | 4:7:1:0:0 | 4 | 7 | 1 | 0 | 0 | 1 | |
| 1355 | 1074 | 169 | 4:7:1:1:0 | 4 | 7 | 1 | 1 | 0 | 1 | |
| 1356 | 1074 | 169 | 4:7:2:0:0 | 4 | 7 | 2 | 0 | 0 | 1 | 1 |
| 1357 | 1074 | 169 | 5:3:0:0:0 | 5 | 3 | 0 | 0 | 0 | 1 | 1 |
| 1358 | 1074 | 169 | 5:3:1:0:0 | 5 | 3 | 1 | 0 | 0 | 1 | 1 |
| 1359 | 1074 | 169 | 5:4:0:0:0 | 5 | 4 | 0 | 0 | 0 | 1 | 1 |
| 1360 | 1074 | 169 | 5:4:0:0:2 | 5 | 4 | 0 | 0 | 2 | | 1 |
| 1361 | 1074 | 169 | 5:4:0:1:0 | 5 | 4 | 0 | 1 | 0 | | 1 |

| | | | | | | | | | | |
|---|---|---|---|---|---|---|---|---|---|---|
| 1362 | 1074 | 169 | 5:4:1:0:0 | 5 | 4 | 1 | 0 | 0 | 1 | 1 |
| 1363 | 1074 | 169 | 5:4:2:0:0 | 5 | 4 | 2 | 0 | 0 | 1 | 1 |
| 1364 | 1074 | 169 | 5:5:0:0:0 | 5 | 5 | 0 | 0 | 0 | 1 | 1 |
| 1365 | 1074 | 169 | 5:5:1:0:0 | 5 | 5 | 1 | 0 | 0 | 1 | 1 |
| 1366 | 1074 | 169 | 5:5:1:0:1 | 5 | 5 | 1 | 0 | 1 | 1 | |
| 1367 | 1074 | 169 | 5:5:1:1:0 | 5 | 5 | 1 | 1 | 0 | | 1 |
| 1368 | 1074 | 169 | 5:5:2:0:0 | 5 | 5 | 2 | 0 | 0 | 1 | 1 |
| 1369 | 1074 | 169 | 5:5:2:0:1 | 5 | 5 | 2 | 0 | 1 | 1 | |
| 1370 | 1074 | 169 | 5:5:3:0:0 | 5 | 5 | 3 | 0 | 0 | 1 | 1 |
| 1371 | 1074 | 169 | 5:5:3:0:1 | 5 | 5 | 3 | 0 | 1 | 1 | |
| 1372 | 1074 | 169 | 5:5:4:0:1 | 5 | 5 | 4 | 0 | 1 | 1 | |
| 1373 | 1074 | 169 | 5:6:0:0:0 | 5 | 6 | 0 | 0 | 0 | 1 | 1 |
| 1374 | 1074 | 169 | 5:6:0:0:1 | 5 | 6 | 0 | 0 | 1 | | 1 |
| 1375 | 1074 | 169 | 5:6:0:1:0 | 5 | 6 | 0 | 1 | 0 | 1 | 1 |
| 1376 | 1074 | 169 | 5:6:1:0:0 | 5 | 6 | 1 | 0 | 0 | 1 | 1 |
| 1377 | 1074 | 169 | 5:6:1:0:1 | 5 | 6 | 1 | 0 | 1 | 1 | 1 |
| 1378 | 1074 | 169 | 5:6:1:0:2 | 5 | 6 | 1 | 0 | 2 | 1 | 1 |
| 1379 | 1074 | 169 | 5:6:1:1:0 | 5 | 6 | 1 | 1 | 0 | 1 | 1 |
| 1380 | 1074 | 169 | 5:6:2:0:0 | 5 | 6 | 2 | 0 | 0 | 1 | 1 |
| 1381 | 1074 | 169 | 5:6:2:0:1 | 5 | 6 | 2 | 0 | 1 | 1 | 1 |
| 1382 | 1074 | 169 | 5:6:2:1:0 | 5 | 6 | 2 | 1 | 0 | 1 | 1 |
| 1383 | 1074 | 169 | 5:6:3:0:0 | 5 | 6 | 3 | 0 | 0 | 1 | 1 |
| 1384 | 1074 | 169 | 5:6:3:0:1 | 5 | 6 | 3 | 0 | 1 | 1 | |
| 1385 | 1074 | 169 | 5:6:3:1:0 | 5 | 6 | 3 | 1 | 0 | 1 | |
| 1386 | 1074 | 169 | 5:6:4:0:0 | 5 | 6 | 4 | 0 | 0 | 1 | |
| 1387 | 1074 | 169 | 5:7:0:1:0 | 5 | 7 | 0 | 1 | 0 | 1 | |
| 1388 | 1074 | 169 | 5:7:0:1:1 | 5 | 7 | 0 | 1 | 1 | 1 | |
| 1389 | 1074 | 169 | 5:7:1:0:0 | 5 | 7 | 1 | 0 | 0 | 1 | 1 |
| 1390 | 1074 | 169 | 5:7:1:0:1 | 5 | 7 | 1 | 0 | 1 | 1 | |
| 1391 | 1074 | 169 | 5:7:1:1:0 | 5 | 7 | 1 | 1 | 0 | 1 | |
| 1392 | 1074 | 169 | 5:7:2:0:0 | 5 | 7 | 2 | 0 | 0 | 1 | |
| 1393 | 1074 | 169 | 5:7:2:1:0 | 5 | 7 | 2 | 1 | 0 | 1 | |
| 1394 | 1074 | 169 | 5:8:0:1:0 | 5 | 8 | 0 | 1 | 0 | 1 | |
| 1395 | 1074 | 169 | 6:3:1:0:0 | 6 | 3 | 1 | 0 | 0 | 1 | 1 |
| 1396 | 1074 | 169 | 6:4:1:0:0 | 6 | 4 | 1 | 0 | 0 | 1 | 1 |
| 1397 | 1074 | 169 | 6:5:0:0:0 | 6 | 5 | 0 | 0 | 0 | 1 | |
| 1398 | 1074 | 169 | 6:5:1:0:0 | 6 | 5 | 1 | 0 | 0 | 1 | 1 |
| 1399 | 1074 | 169 | 6:5:1:0:1 | 6 | 5 | 1 | 0 | 1 | 1 | |
| 1400 | 1074 | 169 | 6:5:2:0:0 | 6 | 5 | 2 | 0 | 0 | 1 | 1 |
| 1401 | 1074 | 169 | 6:5:2:0:1 | 6 | 5 | 2 | 0 | 1 | 1 | |
| 1402 | 1074 | 169 | 6:5:4:0:1 | 6 | 5 | 4 | 0 | 1 | 1 | |
| 1403 | 1074 | 169 | 6:6:0:0:0 | 6 | 6 | 0 | 0 | 0 | 1 | 1 |
| 1404 | 1074 | 169 | 6:6:0:0:1 | 6 | 6 | 0 | 0 | 1 | 1 | |
| 1405 | 1074 | 169 | 6:6:0:1:0 | 6 | 6 | 0 | 1 | 0 | 1 | 1 |
| 1406 | 1074 | 169 | 6:6:1:0:0 | 6 | 6 | 1 | 0 | 0 | 1 | 1 |
| 1407 | 1074 | 169 | 6:6:1:0:1 | 6 | 6 | 1 | 0 | 1 | 1 | |
| 1408 | 1074 | 169 | 6:6:1:1:0 | 6 | 6 | 1 | 1 | 0 | 1 | |
| 1409 | 1074 | 169 | 6:6:2:0:0 | 6 | 6 | 2 | 0 | 0 | 1 | 1 |
| 1410 | 1074 | 169 | 6:6:2:0:1 | 6 | 6 | 2 | 0 | 1 | 1 | |
| 1411 | 1074 | 169 | 6:6:3:0:0 | 6 | 6 | 3 | 0 | 0 | 1 | 1 |
| 1412 | 1074 | 169 | 6:6:3:0:1 | 6 | 6 | 3 | 0 | 1 | 1 | |
| 1413 | 1074 | 169 | 6:6:3:1:0 | 6 | 6 | 3 | 1 | 0 | 1 | |
| 1414 | 1074 | 169 | 6:6:4:0:0 | 6 | 6 | 4 | 0 | 0 | 1 | |
| 1415 | 1074 | 169 | 6:6:4:0:1 | 6 | 6 | 4 | 0 | 1 | 1 | |
| 1416 | 1074 | 169 | 6:7:0:0:0 | 6 | 7 | 0 | 0 | 0 | 1 | 1 |
| 1417 | 1074 | 169 | 6:7:0:0:1 | 6 | 7 | 0 | 0 | 1 | 1 | 1 |
| 1418 | 1074 | 169 | 6:7:0:1:0 | 6 | 7 | 0 | 1 | 0 | 1 | 1 |
| 1419 | 1074 | 169 | 6:7:1:0:0 | 6 | 7 | 1 | 0 | 0 | 1 | 1 |
| 1420 | 1074 | 169 | 6:7:1:0:1 | 6 | 7 | 1 | 0 | 1 | 1 | 1 |
| 1421 | 1074 | 169 | 6:7:1:0:2 | 6 | 7 | 1 | 0 | 2 | 1 | |
| 1422 | 1074 | 169 | 6:7:1:1:0 | 6 | 7 | 1 | 1 | 0 | 1 | 1 |
| 1423 | 1074 | 169 | 6:7:2:0:0 | 6 | 7 | 2 | 0 | 0 | 1 | |
| 1424 | 1074 | 169 | 6:7:2:0:1 | 6 | 7 | 2 | 0 | 1 | 1 | |
| 1425 | 1074 | 169 | 6:7:2:1:0 | 6 | 7 | 2 | 1 | 0 | 1 | |
| 1426 | 1074 | 169 | 6:7:3:0:0 | 6 | 7 | 3 | 0 | 0 | 1 | |

| | | | | | | | | | | |
|---|---|---|---|---|---|---|---|---|---|---|
| 1427 | 1074 | 169 | 6:7:3:0:1 | 6 | 7 | 3 | 0 | 1 | 1 | |
| 1428 | 1074 | 169 | 6:7:3:1:0 | 6 | 7 | 3 | 1 | 0 | 1 | |
| 1429 | 1074 | 169 | 6:7:4:0:0 | 6 | 7 | 4 | 0 | 0 | 1 | 1 |
| 1430 | 1074 | 169 | 6:7:4:0:1 | 6 | 7 | 4 | 0 | 1 | 1 | 1 |
| 1431 | 1074 | 169 | 6:7:5:0:0 | 6 | 7 | 5 | 0 | 0 | | 1 |
| 1432 | 1074 | 169 | 6:8:0:0:0 | 6 | 8 | 0 | 0 | 0 | 1 | |
| 1433 | 1074 | 169 | 6:8:0:1:0 | 6 | 8 | 0 | 1 | 0 | 1 | |
| 1434 | 1074 | 169 | 6:8:0:1:1 | 6 | 8 | 0 | 1 | 1 | 1 | |
| 1435 | 1074 | 169 | 6:8:1:1:0 | 6 | 8 | 1 | 1 | 0 | 1 | 1 |
| 1436 | 1074 | 169 | 6:8:2:1:0 | 6 | 8 | 2 | 1 | 0 | 1 | |
| 1437 | 1074 | 169 | 6:8:3:1:0 | 6 | 8 | 3 | 1 | 0 | 1 | |
| 1438 | 1074 | 169 | 7:3:1:0:0 | 7 | 3 | 1 | 0 | 0 | 1 | 1 |
| 1439 | 1074 | 169 | 7:4:1:0:0 | 7 | 4 | 1 | 0 | 0 | 1 | |
| 1440 | 1074 | 169 | 7:5:1:0:0 | 7 | 5 | 1 | 0 | 0 | 1 | |
| 1441 | 1074 | 169 | 7:6:1:0:0 | 7 | 6 | 1 | 0 | 0 | 1 | 1 |
| 1442 | 1074 | 169 | 7:6:2:0:0 | 7 | 6 | 2 | 0 | 0 | 1 | |
| 1443 | 1074 | 169 | 7:6:3:0:0 | 7 | 6 | 3 | 0 | 0 | 1 | |
| 1444 | 1074 | 169 | 7:7:0:0:0 | 7 | 7 | 0 | 0 | 0 | | 1 |
| 1445 | 1074 | 169 | 7:7:1:0:0 | 7 | 7 | 1 | 0 | 0 | 1 | 1 |
| 1446 | 1074 | 169 | 7:7:1:0:1 | 7 | 7 | 1 | 0 | 1 | 1 | |
| 1447 | 1074 | 169 | 7:7:2:0:0 | 7 | 7 | 2 | 0 | 0 | 1 | |
| 1448 | 1074 | 169 | 7:7:3:0:0 | 7 | 7 | 3 | 0 | 0 | 1 | |
| 1449 | 1074 | 169 | 7:8:0:0:0 | 7 | 8 | 0 | 0 | 0 | 1 | |
| 1450 | 1074 | 169 | 7:8:0:1:0 | 7 | 8 | 0 | 1 | 0 | 1 | |
| 1451 | 1074 | 169 | 7:8:1:0:0 | 7 | 8 | 1 | 0 | 0 | 1 | |
| 1452 | 1074 | 169 | 7:8:2:0:0 | 7 | 8 | 2 | 0 | 0 | 1 | |
| 1453 | 1074 | 169 | 8:9:0:0:0 | 8 | 9 | 0 | 0 | 0 | 1 | |
| 1454 | 1098 | 157 | 2:4:0:0:0 | 2 | 4 | 0 | 0 | 0 | 1 | |
| 1455 | 1098 | 157 | 2:5:0:0:0 | 2 | 5 | 0 | 0 | 0 | 1 | 1 |
| 1456 | 1098 | 157 | 2:6:0:0:0 | 2 | 6 | 0 | 0 | 0 | 1 | 1 |
| 1457 | 1098 | 157 | 2:7:0:0:0 | 2 | 7 | 0 | 0 | 0 | 1 | 1 |
| 1458 | 1098 | 157 | 2:8:0:0:0 | 2 | 8 | 0 | 0 | 0 | 1 | 1 |
| 1459 | 1098 | 157 | 2:9:0:0:0 | 2 | 9 | 0 | 0 | 0 | 1 | 1 |
| 1460 | 1098 | 157 | 3:3:0:0:0 | 3 | 3 | 0 | 0 | 0 | 1 | 1 |
| 1461 | 1098 | 157 | 3:3:1:0:0 | 3 | 3 | 1 | 0 | 0 | 1 | |
| 1462 | 1098 | 157 | 3:4:0:0:0 | 3 | 4 | 0 | 0 | 0 | 1 | 1 |
| 1463 | 1098 | 157 | 3:4:0:0:1 | 3 | 4 | 0 | 0 | 1 | 1 | 1 |
| 1464 | 1098 | 157 | 3:4:1:0:0 | 3 | 4 | 1 | 0 | 0 | 1 | |
| 1465 | 1098 | 157 | 3:4:1:0:1 | 3 | 4 | 1 | 0 | 1 | 1 | |
| 1466 | 1098 | 157 | 3:4:2:0:1 | 3 | 4 | 2 | 0 | 1 | 1 | |
| 1467 | 1098 | 157 | 3:5:0:0:0 | 3 | 5 | 0 | 0 | 0 | 1 | 1 |
| 1468 | 1098 | 157 | 3:5:0:0:1 | 3 | 5 | 0 | 0 | 1 | 1 | 1 |
| 1469 | 1098 | 157 | 3:5:0:1:0 | 3 | 5 | 0 | 1 | 0 | 1 | 1 |
| 1470 | 1098 | 157 | 3:5:1:0:0 | 3 | 5 | 1 | 0 | 0 | 1 | 1 |
| 1471 | 1098 | 157 | 3:5:1:0:1 | 3 | 5 | 1 | 0 | 1 | 1 | |
| 1472 | 1098 | 157 | 3:5:1:0:2 | 3 | 5 | 1 | 0 | 2 | 1 | |
| 1473 | 1098 | 157 | 3:5:1:1:0 | 3 | 5 | 1 | 1 | 0 | 1 | |
| 1474 | 1098 | 157 | 3:5:1:1:1 | 3 | 5 | 1 | 1 | 1 | 1 | |
| 1475 | 1098 | 157 | 3:5:2:0:0 | 3 | 5 | 2 | 0 | 0 | 1 | |
| 1476 | 1098 | 157 | 3:6:0:0:0 | 3 | 6 | 0 | 0 | 0 | 1 | 1 |
| 1477 | 1098 | 157 | 3:6:0:0:1 | 3 | 6 | 0 | 0 | 1 | 1 | 1 |
| 1478 | 1098 | 157 | 3:6:0:1:0 | 3 | 6 | 0 | 1 | 0 | 1 | 1 |
| 1479 | 1098 | 157 | 3:6:1:0:0 | 3 | 6 | 1 | 0 | 0 | 1 | 1 |
| 1480 | 1098 | 157 | 3:6:1:0:1 | 3 | 6 | 1 | 0 | 1 | 1 | 1 |
| 1481 | 1098 | 157 | 3:6:2:0:0 | 3 | 6 | 2 | 0 | 0 | 1 | |
| 1482 | 1098 | 157 | 3:7:0:0:0 | 3 | 7 | 0 | 0 | 0 | 1 | |
| 1483 | 1098 | 157 | 3:7:1:0:0 | 3 | 7 | 1 | 0 | 0 | 1 | |
| 1484 | 1098 | 157 | 4:3:0:0:0 | 4 | 3 | 0 | 0 | 0 | 1 | 1 |
| 1485 | 1098 | 157 | 4:3:1:0:0 | 4 | 3 | 1 | 0 | 0 | 1 | |
| 1486 | 1098 | 157 | 4:4:0:0:0 | 4 | 4 | 0 | 0 | 0 | 1 | 1 |
| 1487 | 1098 | 157 | 4:4:1:0:0 | 4 | 4 | 1 | 0 | 0 | 1 | 1 |
| 1488 | 1098 | 157 | 4:4:1:0:1 | 4 | 4 | 1 | 0 | 1 | 1 | |
| 1489 | 1098 | 157 | 4:4:2:0:1 | 4 | 4 | 2 | 0 | 1 | 1 | |
| 1490 | 1098 | 157 | 4:5:0:0:0 | 4 | 5 | 0 | 0 | 0 | 1 | 1 |
| 1491 | 1098 | 157 | 4:5:0:0:1 | 4 | 5 | 0 | 0 | 1 | 1 | 1 |

| | | | | | | | | | | |
|---|---|---|---|---|---|---|---|---|---|---|
| 1492 | 1098 | 157 | 4:5:0:0:2 | 4 | 5 | 0 | 0 | 2 | | 1 |
| 1493 | 1098 | 157 | 4:5:0:1:0 | 4 | 5 | 0 | 1 | 0 | 1 | 1 |
| 1494 | 1098 | 157 | 4:5:1:0:0 | 4 | 5 | 1 | 0 | 0 | 1 | 1 |
| 1495 | 1098 | 157 | 4:5:1:0:1 | 4 | 5 | 1 | 0 | 1 | 1 | 1 |
| 1496 | 1098 | 157 | 4:5:1:1:0 | 4 | 5 | 1 | 1 | 0 | 1 | 1 |
| 1497 | 1098 | 157 | 4:5:2:0:0 | 4 | 5 | 2 | 0 | 0 | 1 | 1 |
| 1498 | 1098 | 157 | 4:5:2:0:1 | 4 | 5 | 2 | 0 | 1 | 1 | |
| 1499 | 1098 | 157 | 4:5:3:0:0 | 4 | 5 | 3 | 0 | 0 | 1 | 1 |
| 1500 | 1098 | 157 | 4:5:3:1:0 | 4 | 5 | 3 | 1 | 0 | | 1 |
| 1501 | 1098 | 157 | 4:5:3:1:1 | 4 | 5 | 3 | 1 | 1 | | 1 |
| 1502 | 1098 | 157 | 4:5:3:2:0 | 4 | 5 | 3 | 2 | 0 | | 1 |
| 1503 | 1098 | 157 | 4:6:0:0:0 | 4 | 6 | 0 | 0 | 0 | 1 | 1 |
| 1504 | 1098 | 157 | 4:6:0:0:1 | 4 | 6 | 0 | 0 | 1 | 1 | 1 |
| 1505 | 1098 | 157 | 4:6:0:1:0 | 4 | 6 | 0 | 1 | 0 | 1 | 1 |
| 1506 | 1098 | 157 | 4:6:1:0:0 | 4 | 6 | 1 | 0 | 0 | 1 | 1 |
| 1507 | 1098 | 157 | 4:6:1:0:1 | 4 | 6 | 1 | 0 | 1 | 1 | |
| 1508 | 1098 | 157 | 4:6:1:1:0 | 4 | 6 | 1 | 1 | 0 | 1 | |
| 1509 | 1098 | 157 | 4:6:2:0:0 | 4 | 6 | 2 | 0 | 0 | 1 | 1 |
| 1510 | 1098 | 157 | 4:6:3:0:0 | 4 | 6 | 3 | 0 | 0 | 1 | |
| 1511 | 1098 | 157 | 4:6:4:2:0 | 4 | 6 | 4 | 2 | 0 | 1 | |
| 1512 | 1098 | 157 | 4:7:0:0:0 | 4 | 7 | 0 | 0 | 0 | 1 | 1 |
| 1513 | 1098 | 157 | 4:7:0:0:1 | 4 | 7 | 0 | 0 | 1 | 1 | |
| 1514 | 1098 | 157 | 4:7:0:1:0 | 4 | 7 | 0 | 1 | 0 | 1 | 1 |
| 1515 | 1098 | 157 | 4:7:1:0:0 | 4 | 7 | 1 | 0 | 0 | 1 | 1 |
| 1516 | 1098 | 157 | 4:7:2:0:0 | 4 | 7 | 2 | 0 | 0 | 1 | |
| 1517 | 1098 | 157 | 4:8:0:0:0 | 4 | 8 | 0 | 0 | 0 | 1 | |
| 1518 | 1098 | 157 | 4:8:1:0:0 | 4 | 8 | 1 | 0 | 0 | 1 | |
| 1519 | 1098 | 157 | 5:3:0:0:0 | 5 | 3 | 0 | 0 | 0 | 1 | 1 |
| 1520 | 1098 | 157 | 5:3:1:0:0 | 5 | 3 | 1 | 0 | 0 | 1 | 1 |
| 1521 | 1098 | 157 | 5:4:0:0:0 | 5 | 4 | 0 | 0 | 0 | 1 | 1 |
| 1522 | 1098 | 157 | 5:4:1:0:0 | 5 | 4 | 1 | 0 | 0 | 1 | 1 |
| 1523 | 1098 | 157 | 5:5:0:0:0 | 5 | 5 | 0 | 0 | 0 | 1 | 1 |
| 1524 | 1098 | 157 | 5:5:0:0:1 | 5 | 5 | 0 | 0 | 1 | | 1 |
| 1525 | 1098 | 157 | 5:5:0:0:2 | 5 | 5 | 0 | 0 | 2 | | 1 |
| 1526 | 1098 | 157 | 5:5:1:0:0 | 5 | 5 | 1 | 0 | 0 | 1 | 1 |
| 1527 | 1098 | 157 | 5:5:1:0:1 | 5 | 5 | 1 | 0 | 1 | 1 | 1 |
| 1528 | 1098 | 157 | 5:5:1:0:2 | 5 | 5 | 1 | 0 | 2 | 1 | |
| 1529 | 1098 | 157 | 5:5:2:0:0 | 5 | 5 | 2 | 0 | 0 | 1 | 1 |
| 1530 | 1098 | 157 | 5:5:2:0:1 | 5 | 5 | 2 | 0 | 1 | 1 | |
| 1531 | 1098 | 157 | 5:5:3:0:1 | 5 | 5 | 3 | 0 | 1 | 1 | |
| 1532 | 1098 | 157 | 5:6:0:0:0 | 5 | 6 | 0 | 0 | 0 | 1 | 1 |
| 1533 | 1098 | 157 | 5:6:0:0:1 | 5 | 6 | 0 | 0 | 1 | 1 | 1 |
| 1534 | 1098 | 157 | 5:6:0:0:2 | 5 | 6 | 0 | 0 | 2 | 1 | 1 |
| 1535 | 1098 | 157 | 5:6:0:1:0 | 5 | 6 | 0 | 1 | 0 | 1 | 1 |
| 1536 | 1098 | 157 | 5:6:0:1:1 | 5 | 6 | 0 | 1 | 1 | 1 | 1 |
| 1537 | 1098 | 157 | 5:6:1:0:0 | 5 | 6 | 1 | 0 | 0 | 1 | 1 |
| 1538 | 1098 | 157 | 5:6:1:0:1 | 5 | 6 | 1 | 0 | 1 | 1 | 1 |
| 1539 | 1098 | 157 | 5:6:1:0:2 | 5 | 6 | 1 | 0 | 2 | | 1 |
| 1540 | 1098 | 157 | 5:6:1:1:0 | 5 | 6 | 1 | 1 | 0 | 1 | 1 |
| 1541 | 1098 | 157 | 5:6:2:0:0 | 5 | 6 | 2 | 0 | 0 | 1 | 1 |
| 1542 | 1098 | 157 | 5:6:2:0:1 | 5 | 6 | 2 | 0 | 1 | 1 | 1 |
| 1543 | 1098 | 157 | 5:6:2:1:0 | 5 | 6 | 2 | 1 | 0 | 1 | |
| 1544 | 1098 | 157 | 5:6:3:0:0 | 5 | 6 | 3 | 0 | 0 | 1 | 1 |
| 1545 | 1098 | 157 | 5:6:3:0:1 | 5 | 6 | 3 | 0 | 1 | 1 | |
| 1546 | 1098 | 157 | 5:6:4:0:0 | 5 | 6 | 4 | 0 | 0 | 1 | |
| 1547 | 1098 | 157 | 5:7:0:0:0 | 5 | 7 | 0 | 0 | 0 | 1 | 1 |
| 1548 | 1098 | 157 | 5:7:0:1:0 | 5 | 7 | 0 | 1 | 0 | 1 | |
| 1549 | 1098 | 157 | 5:7:1:0:0 | 5 | 7 | 1 | 0 | 0 | 1 | |
| 1550 | 1098 | 157 | 5:7:1:0:1 | 5 | 7 | 1 | 0 | 1 | 1 | |
| 1551 | 1098 | 157 | 5:7:1:1:0 | 5 | 7 | 1 | 1 | 0 | 1 | |
| 1552 | 1098 | 157 | 5:7:2:0:0 | 5 | 7 | 2 | 0 | 0 | 1 | |
| 1553 | 1098 | 157 | 5:7:2:1:0 | 5 | 7 | 2 | 1 | 0 | 1 | |
| 1554 | 1098 | 157 | 5:8:0:1:0 | 5 | 8 | 0 | 1 | 0 | 1 | |
| 1555 | 1098 | 157 | 6:3:0:0:0 | 6 | 3 | 0 | 0 | 0 | 1 | 1 |
| 1556 | 1098 | 157 | 6:3:1:0:0 | 6 | 3 | 1 | 0 | 0 | 1 | 1 |

| | | | | | | | | | | |
|---|---|---|---|---|---|---|---|---|---|---|
| 1557 | 1098 | 157 | 6:3:2:0:0 | 6 | 3 | 2 | 0 | 0 | | 1 |
| 1558 | 1098 | 157 | 6:4:0:0:0 | 6 | 4 | 0 | 0 | 0 | 1 | 1 |
| 1559 | 1098 | 157 | 6:4:1:0:0 | 6 | 4 | 1 | 0 | 0 | 1 | 1 |
| 1560 | 1098 | 157 | 6:5:0:0:0 | 6 | 5 | 0 | 0 | 0 | 1 | 1 |
| 1561 | 1098 | 157 | 6:5:1:0:0 | 6 | 5 | 1 | 0 | 0 | 1 | 1 |
| 1562 | 1098 | 157 | 6:5:2:0:0 | 6 | 5 | 2 | 0 | 0 | 1 | |
| 1563 | 1098 | 157 | 6:6:0:0:0 | 6 | 6 | 0 | 0 | 0 | 1 | 1 |
| 1564 | 1098 | 157 | 6:6:0:0:1 | 6 | 6 | 0 | 0 | 1 | 1 | |
| 1565 | 1098 | 157 | 6:6:0:1:1 | 6 | 6 | 0 | 1 | 1 | | 1 |
| 1566 | 1098 | 157 | 6:6:1:0:0 | 6 | 6 | 1 | 0 | 0 | 1 | 1 |
| 1567 | 1098 | 157 | 6:6:1:0:1 | 6 | 6 | 1 | 0 | 1 | 1 | |
| 1568 | 1098 | 157 | 6:6:1:0:2 | 6 | 6 | 1 | 0 | 2 | 1 | 1 |
| 1569 | 1098 | 157 | 6:6:1:1:2 | 6 | 6 | 1 | 1 | 2 | | 1 |
| 1570 | 1098 | 157 | 6:6:2:0:0 | 6 | 6 | 2 | 0 | 0 | 1 | |
| 1571 | 1098 | 157 | 6:6:2:0:1 | 6 | 6 | 2 | 0 | 1 | 1 | |
| 1572 | 1098 | 157 | 6:6:3:0:0 | 6 | 6 | 3 | 0 | 0 | 1 | |
| 1573 | 1098 | 157 | 6:6:3:0:1 | 6 | 6 | 3 | 0 | 1 | 1 | |
| 1574 | 1098 | 157 | 6:7:0:0:0 | 6 | 7 | 0 | 0 | 0 | 1 | 1 |
| 1575 | 1098 | 157 | 6:7:0:0:1 | 6 | 7 | 0 | 0 | 1 | 1 | 1 |
| 1576 | 1098 | 157 | 6:7:0:0:2 | 6 | 7 | 0 | 0 | 2 | 1 | 1 |
| 1577 | 1098 | 157 | 6:7:0:1:0 | 6 | 7 | 0 | 1 | 0 | 1 | 1 |
| 1578 | 1098 | 157 | 6:7:0:1:1 | 6 | 7 | 0 | 1 | 1 | 1 | 1 |
| 1579 | 1098 | 157 | 6:7:0:2:1 | 6 | 7 | 0 | 2 | 1 | | 1 |
| 1580 | 1098 | 157 | 6:7:1:0:0 | 6 | 7 | 1 | 0 | 0 | 1 | 1 |
| 1581 | 1098 | 157 | 6:7:1:0:1 | 6 | 7 | 1 | 0 | 1 | 1 | 1 |
| 1582 | 1098 | 157 | 6:7:1:0:2 | 6 | 7 | 1 | 0 | 2 | 1 | |
| 1583 | 1098 | 157 | 6:7:1:1:0 | 6 | 7 | 1 | 1 | 0 | 1 | 1 |
| 1584 | 1098 | 157 | 6:7:2:0:0 | 6 | 7 | 2 | 0 | 0 | 1 | |
| 1585 | 1098 | 157 | 6:7:2:0:1 | 6 | 7 | 2 | 0 | 1 | 1 | |
| 1586 | 1098 | 157 | 6:7:2:1:0 | 6 | 7 | 2 | 1 | 0 | 1 | |
| 1587 | 1098 | 157 | 6:7:3:0:0 | 6 | 7 | 3 | 0 | 0 | 1 | |
| 1588 | 1098 | 157 | 6:7:3:0:1 | 6 | 7 | 3 | 0 | 1 | 1 | |
| 1589 | 1098 | 157 | 6:7:4:0:0 | 6 | 7 | 4 | 0 | 0 | 1 | 1 |
| 1590 | 1098 | 157 | 6:8:0:0:0 | 6 | 8 | 0 | 0 | 0 | 1 | |
| 1591 | 1098 | 157 | 6:8:0:1:0 | 6 | 8 | 0 | 1 | 0 | 1 | 1 |
| 1592 | 1098 | 157 | 6:8:0:1:1 | 6 | 8 | 0 | 1 | 1 | 1 | |
| 1593 | 1098 | 157 | 6:8:1:1:0 | 6 | 8 | 1 | 1 | 0 | 1 | 1 |
| 1594 | 1098 | 157 | 6:8:2:1:0 | 6 | 8 | 2 | 1 | 0 | 1 | |
| 1595 | 1098 | 157 | 6:9:0:0:0 | 6 | 9 | 0 | 0 | 0 | | 1 |
| 1596 | 1098 | 157 | 7:6:0:0:0 | 7 | 6 | 0 | 0 | 0 | 1 | 1 |
| 1597 | 1098 | 157 | 7:6:1:0:0 | 7 | 6 | 1 | 0 | 0 | 1 | |
| 1598 | 1098 | 157 | 7:7:0:0:0 | 7 | 7 | 0 | 0 | 0 | 1 | 1 |
| 1599 | 1098 | 157 | 7:7:1:0:0 | 7 | 7 | 1 | 0 | 0 | 1 | 1 |
| 1600 | 1098 | 157 | 7:7:2:0:1 | 7 | 7 | 2 | 0 | 1 | 1 | |
| 1601 | 1098 | 157 | 7:7:2:0:2 | 7 | 7 | 2 | 0 | 2 | 1 | |
| 1602 | 1098 | 157 | 7:8:0:0:0 | 7 | 8 | 0 | 0 | 0 | 1 | |
| 1603 | 1098 | 157 | 7:8:0:0:1 | 7 | 8 | 0 | 0 | 1 | 1 | |
| 1604 | 1098 | 157 | 7:8:1:0:0 | 7 | 8 | 1 | 0 | 0 | 1 | |
| 1605 | 1098 | 157 | 7:8:1:0:1 | 7 | 8 | 1 | 0 | 1 | 1 | |
| 1606 | 1098 | 157 | 7:8:1:1:0 | 7 | 8 | 1 | 1 | 0 | 1 | |
| 1607 | 1098 | 157 | 7:8:1:1:1 | 7 | 8 | 1 | 1 | 1 | 1 | |
| 1608 | 1098 | 157 | 7:8:2:0:0 | 7 | 8 | 2 | 0 | 0 | 1 | |
| 1609 | 1098 | 157 | 7:9:0:1:0 | 7 | 9 | 0 | 1 | 0 | 1 | |
| 1610 | 1098 | 157 | 7:9:0:2:0 | 7 | 9 | 0 | 2 | 0 | 1 | |
| 1611 | 1134 | 60 | 2:5:0:0:0 | 2 | 5 | 0 | 0 | 0 | 1 | 1 |
| 1612 | 1134 | 60 | 2:6:0:0:0 | 2 | 6 | 0 | 0 | 0 | 1 | 1 |
| 1613 | 1134 | 60 | 2:7:0:0:0 | 2 | 7 | 0 | 0 | 0 | 1 | 1 |
| 1614 | 1134 | 60 | 2:8:0:0:0 | 2 | 8 | 0 | 0 | 0 | | 1 |
| 1615 | 1134 | 60 | 3:3:1:0:0 | 3 | 3 | 1 | 0 | 0 | | 1 |
| 1616 | 1134 | 60 | 3:4:1:0:0 | 3 | 4 | 1 | 0 | 0 | 1 | 1 |
| 1617 | 1134 | 60 | 3:5:0:0:0 | 3 | 5 | 0 | 0 | 0 | 1 | 1 |
| 1618 | 1134 | 60 | 3:5:0:0:1 | 3 | 5 | 0 | 0 | 1 | 1 | 1 |
| 1619 | 1134 | 60 | 3:5:1:0:0 | 3 | 5 | 1 | 0 | 0 | 1 | 1 |
| 1620 | 1134 | 60 | 3:5:2:0:0 | 3 | 5 | 2 | 0 | 0 | 1 | |
| 1621 | 1134 | 60 | 3:6:0:0:0 | 3 | 6 | 0 | 0 | 0 | 1 | 1 |

| | | | | | | | | | | |
|---|---|---|---|---|---|---|---|---|---|---|
| 1622 | 1134 | 60 | 3:6:0:1:0 | 3 | 6 | 0 | 1 | 0 | | 1 |
| 1623 | 1134 | 60 | 3:6:1:0:0 | 3 | 6 | 1 | 0 | 0 | 1 | 1 |
| 1624 | 1134 | 60 | 4:3:1:0:0 | 4 | 3 | 1 | 0 | 0 | 1 | 1 |
| 1625 | 1134 | 60 | 4:4:1:0:0 | 4 | 4 | 1 | 0 | 0 | 1 | 1 |
| 1626 | 1134 | 60 | 4:5:0:0:0 | 4 | 5 | 0 | 0 | 0 | 1 | |
| 1627 | 1134 | 60 | 4:5:1:0:0 | 4 | 5 | 1 | 0 | 0 | 1 | 1 |
| 1628 | 1134 | 60 | 4:5:1:0:1 | 4 | 5 | 1 | 0 | 1 | 1 | |
| 1629 | 1134 | 60 | 4:5:2:0:0 | 4 | 5 | 2 | 0 | 0 | 1 | 1 |
| 1630 | 1134 | 60 | 4:5:3:0:0 | 4 | 5 | 3 | 0 | 0 | 1 | |
| 1631 | 1134 | 60 | 4:6:0:1:0 | 4 | 6 | 0 | 1 | 0 | 1 | 1 |
| 1632 | 1134 | 60 | 4:6:1:0:0 | 4 | 6 | 1 | 0 | 0 | | 1 |
| 1633 | 1134 | 60 | 4:6:2:0:0 | 4 | 6 | 2 | 0 | 0 | | 1 |
| 1634 | 1134 | 60 | 5:3:0:0:0 | 5 | 3 | 0 | 0 | 0 | | 1 |
| 1635 | 1134 | 60 | 5:3:1:0:0 | 5 | 3 | 1 | 0 | 0 | 1 | |
| 1636 | 1134 | 60 | 5:3:1:1:0 | 5 | 3 | 1 | 1 | 0 | | 1 |
| 1637 | 1134 | 60 | 5:4:1:0:0 | 5 | 4 | 1 | 0 | 0 | 1 | 1 |
| 1638 | 1134 | 60 | 5:5:1:0:0 | 5 | 5 | 1 | 0 | 0 | 1 | |
| 1639 | 1134 | 60 | 5:5:1:1:0 | 5 | 5 | 1 | 1 | 0 | | 1 |
| 1640 | 1134 | 60 | 5:5:2:0:0 | 5 | 5 | 2 | 0 | 0 | 1 | |
| 1641 | 1134 | 60 | 5:6:0:0:0 | 5 | 6 | 0 | 0 | 0 | | 1 |
| 1642 | 1134 | 60 | 5:6:0:1:0 | 5 | 6 | 0 | 1 | 0 | | 1 |
| 1643 | 1134 | 60 | 5:6:1:0:0 | 5 | 6 | 1 | 0 | 0 | 1 | 1 |
| 1644 | 1134 | 60 | 5:6:1:0:1 | 5 | 6 | 1 | 0 | 1 | | 1 |
| 1645 | 1134 | 60 | 5:6:1:0:2 | 5 | 6 | 1 | 0 | 2 | | 1 |
| 1646 | 1134 | 60 | 5:6:1:1:0 | 5 | 6 | 1 | 1 | 0 | | 1 |
| 1647 | 1134 | 60 | 5:6:2:0:0 | 5 | 6 | 2 | 0 | 0 | 1 | |
| 1648 | 1134 | 60 | 5:6:2:0:1 | 5 | 6 | 2 | 0 | 1 | 1 | |
| 1649 | 1134 | 60 | 5:6:3:0:0 | 5 | 6 | 3 | 0 | 0 | 1 | |
| 1650 | 1134 | 60 | 5:6:3:0:1 | 5 | 6 | 3 | 0 | 1 | | 1 |
| 1651 | 1134 | 60 | 5:7:1:1:0 | 5 | 7 | 1 | 1 | 0 | 1 | |
| 1652 | 1134 | 60 | 5:8:1:0:0 | 5 | 8 | 1 | 0 | 0 | | 1 |
| 1653 | 1134 | 60 | 6:3:1:0:0 | 6 | 3 | 1 | 0 | 0 | 1 | |
| 1654 | 1134 | 60 | 6:4:1:0:0 | 6 | 4 | 1 | 0 | 0 | 1 | 1 |
| 1655 | 1134 | 60 | 6:5:1:0:0 | 6 | 5 | 1 | 0 | 0 | | 1 |
| 1656 | 1134 | 60 | 6:5:2:0:0 | 6 | 5 | 2 | 0 | 0 | | 1 |
| 1657 | 1134 | 60 | 6:6:0:1:0 | 6 | 6 | 0 | 1 | 0 | | 1 |
| 1658 | 1134 | 60 | 6:6:1:0:0 | 6 | 6 | 1 | 0 | 0 | 1 | 1 |
| 1659 | 1134 | 60 | 6:6:3:0:0 | 6 | 6 | 3 | 0 | 0 | 1 | |
| 1660 | 1134 | 60 | 6:6:3:0:1 | 6 | 6 | 3 | 0 | 1 | 1 | |
| 1661 | 1134 | 60 | 6:7:0:0:0 | 6 | 7 | 0 | 0 | 0 | | 1 |
| 1662 | 1134 | 60 | 6:7:1:0:0 | 6 | 7 | 1 | 0 | 0 | 1 | 1 |
| 1663 | 1134 | 60 | 6:7:1:0:1 | 6 | 7 | 1 | 0 | 1 | 1 | |
| 1664 | 1134 | 60 | 6:7:2:0:0 | 6 | 7 | 2 | 0 | 0 | 1 | |
| 1665 | 1134 | 60 | 6:7:2:1:0 | 6 | 7 | 2 | 1 | 0 | 1 | |
| 1666 | 1134 | 60 | 6:7:3:0:0 | 6 | 7 | 3 | 0 | 0 | 1 | |
| 1667 | 1134 | 60 | 6:8:0:1:0 | 6 | 8 | 0 | 1 | 0 | 1 | |
| 1668 | 1134 | 60 | 7:8:1:2:1 | 7 | 8 | 1 | 2 | 1 | | 1 |
| 1669 | 1134 | 60 | 8:5:0:0:0 | 8 | 5 | 0 | 0 | 0 | | 1 |
| 1670 | 1134 | 60 | 9:3:1:0:0 | 9 | 3 | 1 | 0 | 0 | | 1 |
| 1671 | 1158 | 24 | 4:7:4:2:0 | 4 | 7 | 4 | 2 | 0 | 1 | |
| 1672 | 1158 | 24 | 5:6:1:0:0 | 5 | 6 | 1 | 0 | 0 | 1 | |
| 1673 | 1158 | 24 | 6:6:1:0:0 | 6 | 6 | 1 | 0 | 0 | 1 | |
| 1674 | 1158 | 24 | 6:6:2:0:1 | 6 | 6 | 2 | 0 | 1 | 1 | |
| 1675 | 1158 | 24 | 6:7:1:0:0 | 6 | 7 | 1 | 0 | 0 | 1 | 1 |
| 1676 | 1158 | 24 | 6:7:1:0:1 | 6 | 7 | 1 | 0 | 1 | 1 | |
| 1677 | 1158 | 24 | 6:7:1:0:2 | 6 | 7 | 1 | 0 | 2 | 1 | |
| 1678 | 1158 | 24 | 6:7:1:1:0 | 6 | 7 | 1 | 1 | 0 | 1 | |
| 1679 | 1158 | 24 | 6:7:2:0:0 | 6 | 7 | 2 | 0 | 0 | 1 | |
| 1680 | 1158 | 24 | 6:7:2:0:1 | 6 | 7 | 2 | 0 | 1 | 1 | |
| 1681 | 1158 | 24 | 6:7:3:0:0 | 6 | 7 | 3 | 0 | 0 | 1 | |
| 1682 | 1158 | 24 | 6:7:3:0:1 | 6 | 7 | 3 | 0 | 1 | 1 | |
| 1683 | 1158 | 24 | 6:7:3:0:2 | 6 | 7 | 3 | 0 | 2 | 1 | |
| 1684 | 1158 | 24 | 6:7:4:0:0 | 6 | 7 | 4 | 0 | 0 | 1 | |
| 1685 | 1158 | 24 | 6:8:0:1:0 | 6 | 8 | 0 | 1 | 0 | 1 | |
| 1686 | 1158 | 24 | 6:8:0:1:1 | 6 | 8 | 0 | 1 | 1 | 1 | |

| | | | | | | | | | | |
|---|---|---|---|---|---|---|---|---|---|---|
| 1687 | 1158 | 24 | 6:8:1:1:0 | 6 | 8 | 1 | 1 | 0 | 1 | |
| 1688 | 1158 | 24 | 6:8:2:1:0 | 6 | 8 | 2 | 1 | 0 | 1 | |
| 1689 | 1158 | 24 | 6:8:2:1:1 | 6 | 8 | 2 | 1 | 1 | 1 | |
| 1690 | 1158 | 24 | 6:9:1:2:0 | 6 | 9 | 1 | 2 | 0 | 1 | |
| 1691 | 1158 | 24 | 7:8:1:0:0 | 7 | 8 | 1 | 0 | 0 | 1 | |
| 1692 | 1158 | 24 | 7:8:1:0:1 | 7 | 8 | 1 | 0 | 1 | 1 | |
| 1693 | 1158 | 24 | 7:8:3:0:0 | 7 | 8 | 3 | 0 | 0 | 1 | |
| 1694 | 1158 | 24 | 7:9:0:1:0 | 7 | 9 | 0 | 1 | 0 | 1 | |
| 1695 | 1173 | 16 | 5:6:1:0:0 | 5 | 6 | 1 | 0 | 0 | 1 | 1 |
| 1696 | 1173 | 16 | 6:3:1:0:0 | 6 | 3 | 1 | 0 | 0 | | 1 |
| 1697 | 1173 | 16 | 6:6:2:0:2 | 6 | 6 | 2 | 0 | 2 | 1 | |
| 1698 | 1173 | 16 | 6:6:3:0:1 | 6 | 6 | 3 | 0 | 1 | 1 | |
| 1699 | 1173 | 16 | 6:7:1:0:0 | 6 | 7 | 1 | 0 | 0 | 1 | 1 |
| 1700 | 1173 | 16 | 6:7:1:0:1 | 6 | 7 | 1 | 0 | 1 | 1 | 1 |
| 1701 | 1173 | 16 | 6:7:1:0:2 | 6 | 7 | 1 | 0 | 2 | 1 | |
| 1702 | 1173 | 16 | 6:7:1:1:1 | 6 | 7 | 1 | 1 | 1 | 1 | |
| 1703 | 1173 | 16 | 6:7:2:0:0 | 6 | 7 | 2 | 0 | 0 | 1 | |
| 1704 | 1173 | 16 | 6:7:2:1:0 | 6 | 7 | 2 | 1 | 0 | 1 | |
| 1705 | 1173 | 16 | 6:7:3:0:0 | 6 | 7 | 3 | 0 | 0 | 1 | |
| 1706 | 1173 | 16 | 6:7:4:0:0 | 6 | 7 | 4 | 0 | 0 | 1 | 1 |
| 1707 | 1173 | 16 | 6:7:5:0:0 | 6 | 7 | 5 | 0 | 0 | | 1 |
| 1708 | 1173 | 16 | 6:8:0:1:0 | 6 | 8 | 0 | 1 | 0 | 1 | |
| 1709 | 1173 | 16 | 6:8:0:1:1 | 6 | 8 | 0 | 1 | 1 | 1 | |
| 1710 | 1173 | 16 | 6:8:0:2:0 | 6 | 8 | 0 | 2 | 0 | 1 | |
| 1711 | 1194 | 49 | 4:3:1:0:0 | 4 | 3 | 1 | 0 | 0 | 1 | 1 |
| 1712 | 1194 | 49 | 4:3:1:0:1 | 4 | 3 | 1 | 0 | 1 | 1 | |
| 1713 | 1194 | 49 | 4:4:0:0:0 | 4 | 4 | 0 | 0 | 0 | 1 | |
| 1714 | 1194 | 49 | 4:4:0:1:0 | 4 | 4 | 0 | 1 | 0 | 1 | |
| 1715 | 1194 | 49 | 4:5:0:0:0 | 4 | 5 | 0 | 0 | 0 | 1 | 1 |
| 1716 | 1194 | 49 | 4:5:1:0:0 | 4 | 5 | 1 | 0 | 0 | 1 | 1 |
| 1717 | 1194 | 49 | 4:5:2:0:0 | 4 | 5 | 2 | 0 | 0 | 1 | |
| 1718 | 1194 | 49 | 4:5:3:0:0 | 4 | 5 | 3 | 0 | 0 | 1 | 1 |
| 1719 | 1194 | 49 | 5:3:1:0:0 | 5 | 3 | 1 | 0 | 0 | 1 | 1 |
| 1720 | 1194 | 49 | 5:4:1:0:0 | 5 | 4 | 1 | 0 | 0 | 1 | 1 |
| 1721 | 1194 | 49 | 5:5:1:0:0 | 5 | 5 | 1 | 0 | 0 | 1 | 1 |
| 1722 | 1194 | 49 | 5:6:0:0:0 | 5 | 6 | 0 | 0 | 0 | 1 | 1 |
| 1723 | 1194 | 49 | 5:6:0:1:1 | 5 | 6 | 0 | 1 | 1 | | 1 |
| 1724 | 1194 | 49 | 5:6:1:0:0 | 5 | 6 | 1 | 0 | 0 | 1 | 1 |
| 1725 | 1194 | 49 | 5:6:1:0:1 | 5 | 6 | 1 | 0 | 1 | 1 | 1 |
| 1726 | 1194 | 49 | 5:6:2:0:0 | 5 | 6 | 2 | 0 | 0 | 1 | 1 |
| 1727 | 1194 | 49 | 5:6:2:0:1 | 5 | 6 | 2 | 0 | 1 | 1 | |
| 1728 | 1194 | 49 | 5:6:2:1:0 | 5 | 6 | 2 | 1 | 0 | | 1 |
| 1729 | 1194 | 49 | 5:6:3:0:0 | 5 | 6 | 3 | 0 | 0 | 1 | |
| 1730 | 1194 | 49 | 5:6:4:0:0 | 5 | 6 | 4 | 0 | 0 | 1 | |
| 1731 | 1194 | 49 | 5:7:0:1:0 | 5 | 7 | 0 | 1 | 0 | 1 | |
| 1732 | 1194 | 49 | 5:7:1:1:0 | 5 | 7 | 1 | 1 | 0 | 1 | 1 |
| 1733 | 1194 | 49 | 6:3:1:0:0 | 6 | 3 | 1 | 0 | 0 | 1 | 1 |
| 1734 | 1194 | 49 | 6:4:1:0:0 | 6 | 4 | 1 | 0 | 0 | 1 | 1 |
| 1735 | 1194 | 49 | 6:5:1:0:0 | 6 | 5 | 1 | 0 | 0 | 1 | 1 |
| 1736 | 1194 | 49 | 6:6:1:0:0 | 6 | 6 | 1 | 0 | 0 | 1 | |
| 1737 | 1194 | 49 | 6:6:2:0:1 | 6 | 6 | 2 | 0 | 1 | 1 | |
| 1738 | 1194 | 49 | 6:7:0:0:0 | 6 | 7 | 0 | 0 | 0 | 1 | 1 |
| 1739 | 1194 | 49 | 6:7:0:1:0 | 6 | 7 | 0 | 1 | 0 | | 1 |
| 1740 | 1194 | 49 | 6:7:0:1:1 | 6 | 7 | 0 | 1 | 1 | | 1 |
| 1741 | 1194 | 49 | 6:7:1:0:0 | 6 | 7 | 1 | 0 | 0 | 1 | 1 |
| 1742 | 1194 | 49 | 6:7:1:0:1 | 6 | 7 | 1 | 0 | 1 | 1 | 1 |
| 1743 | 1194 | 49 | 6:7:1:1:0 | 6 | 7 | 1 | 1 | 0 | 1 | 1 |
| 1744 | 1194 | 49 | 6:7:2:0:0 | 6 | 7 | 2 | 0 | 0 | 1 | |
| 1745 | 1194 | 49 | 6:7:2:0:1 | 6 | 7 | 2 | 0 | 1 | 1 | |
| 1746 | 1194 | 49 | 6:7:2:0:2 | 6 | 7 | 2 | 0 | 2 | 1 | |
| 1747 | 1194 | 49 | 6:7:3:0:0 | 6 | 7 | 3 | 0 | 0 | 1 | |
| 1748 | 1194 | 49 | 6:7:3:0:1 | 6 | 7 | 3 | 0 | 1 | 1 | |
| 1749 | 1194 | 49 | 6:7:3:0:2 | 6 | 7 | 3 | 0 | 2 | 1 | |
| 1750 | 1194 | 49 | 6:7:4:0:0 | 6 | 7 | 4 | 0 | 0 | 1 | 1 |
| 1751 | 1194 | 49 | 6:7:4:0:1 | 6 | 7 | 4 | 0 | 1 | | 1 |

| | | | | | | | | | | |
|---|---|---|---|---|---|---|---|---|---|---|
| 1752 | 1194 | 49 | 6:8:0:1:0 | 6 | 8 | 0 | 1 | 0 | 1 | |
| 1753 | 1194 | 49 | 6:8:1:1:0 | 6 | 8 | 1 | 1 | 0 | 1 | |
| 1754 | 1194 | 49 | 6:8:1:1:1 | 6 | 8 | 1 | 1 | 1 | 1 | |
| 1755 | 1194 | 49 | 6:8:2:1:0 | 6 | 8 | 2 | 1 | 0 | 1 | |
| 1756 | 1194 | 49 | 6:8:2:1:1 | 6 | 8 | 2 | 1 | 1 | 1 | |
| 1757 | 1194 | 49 | 6:9:0:2:0 | 6 | 9 | 0 | 2 | 0 | 1 | |
| 1758 | 1194 | 49 | 6:9:1:2:0 | 6 | 9 | 1 | 2 | 0 | 1 | |
| 1759 | 1194 | 49 | 7:8:1:0:0 | 7 | 8 | 1 | 0 | 0 | 1 | |
| Total | | | | | | | | | 1,449 | 919 |

Supplementary Table S13. Total glycome assigned on SARS CoV-2 Spike protein by GR and Byonic

| No. | Glycan composition | No. of HexNAc | No. of Hex | No. of Fuc | No. of NeuAc | No. of NeuGc | Assigned by GR | Assigned by Byonic |
|---|---|---|---|---|---|---|---|---|
| 1 | 2:0:1:0:0 | 2 | 0 | 1 | 0 | 0 | | 1 |
| 2 | 2:3:0:0:0 | 2 | 3 | 0 | 0 | 0 | 1 | 1 |
| 3 | 2:4:0:0:0 | 2 | 4 | 0 | 0 | 0 | 1 | 1 |
| 4 | 2:5:0:0:0 | 2 | 5 | 0 | 0 | 0 | 1 | 1 |
| 5 | 2:5:1:0:0 | 2 | 5 | 1 | 0 | 0 | | 1 |
| 6 | 2:6:0:0:0 | 2 | 6 | 0 | 0 | 0 | 1 | 1 |
| 7 | 2:7:0:0:0 | 2 | 7 | 0 | 0 | 0 | 1 | 1 |
| 8 | 2:8:0:0:0 | 2 | 8 | 0 | 0 | 0 | 1 | 1 |
| 9 | 2:9:0:0:0 | 2 | 9 | 0 | 0 | 0 | 1 | 1 |
| 10 | 2:11:0:0:0 | 2 | 11 | 0 | 0 | 0 | | 1 |
| 11 | 3:3:0:0:0 | 3 | 3 | 0 | 0 | 0 | 1 | 1 |
| 12 | 3:3:1:0:0 | 3 | 3 | 1 | 0 | 0 | 1 | 1 |
| 13 | 3:3:1:0:1 | 3 | 3 | 1 | 0 | 1 | 1 | |
| 14 | 3:3:1:1:0 | 3 | 3 | 1 | 1 | 0 | 1 | |
| 15 | 3:3:2:0:0 | 3 | 3 | 2 | 0 | 0 | | 1 |
| 16 | 3:3:2:0:1 | 3 | 3 | 2 | 0 | 1 | 1 | |
| 17 | 3:4:0:0:0 | 3 | 4 | 0 | 0 | 0 | 1 | 1 |
| 18 | 3:4:0:0:1 | 3 | 4 | 0 | 0 | 1 | 1 | 1 |
| 19 | 3:4:0:1:0 | 3 | 4 | 0 | 1 | 0 | 1 | 1 |
| 20 | 3:4:1:0:0 | 3 | 4 | 1 | 0 | 0 | 1 | 1 |
| 21 | 3:4:1:0:1 | 3 | 4 | 1 | 0 | 1 | 1 | 1 |
| 22 | 3:4:1:1:0 | 3 | 4 | 1 | 1 | 0 | 1 | 1 |
| 23 | 3:4:2:0:0 | 3 | 4 | 2 | 0 | 0 | 1 | 1 |
| 24 | 3:4:2:0:1 | 3 | 4 | 2 | 0 | 1 | 1 | |
| 25 | 3:5:0:0:0 | 3 | 5 | 0 | 0 | 0 | 1 | 1 |
| 26 | 3:5:0:0:1 | 3 | 5 | 0 | 0 | 1 | 1 | 1 |
| 27 | 3:5:0:1:0 | 3 | 5 | 0 | 1 | 0 | 1 | 1 |
| 28 | 3:5:1:0:0 | 3 | 5 | 1 | 0 | 0 | 1 | 1 |
| 29 | 3:5:1:0:1 | 3 | 5 | 1 | 0 | 1 | 1 | |
| 30 | 3:5:1:0:2 | 3 | 5 | 1 | 0 | 2 | 1 | |
| 31 | 3:5:1:1:0 | 3 | 5 | 1 | 1 | 0 | 1 | |
| 32 | 3:5:1:1:1 | 3 | 5 | 1 | 1 | 1 | 1 | |
| 33 | 3:5:2:0:0 | 3 | 5 | 2 | 0 | 0 | 1 | |
| 34 | 3:5:2:0:1 | 3 | 5 | 2 | 0 | 1 | 1 | |
| 35 | 3:6:0:0:0 | 3 | 6 | 0 | 0 | 0 | 1 | 1 |
| 36 | 3:6:0:0:1 | 3 | 6 | 0 | 0 | 1 | 1 | 1 |
| 37 | 3:6:0:1:0 | 3 | 6 | 0 | 1 | 0 | 1 | 1 |
| 38 | 3:6:1:0:0 | 3 | 6 | 1 | 0 | 0 | 1 | 1 |
| 39 | 3:6:1:0:1 | 3 | 6 | 1 | 0 | 1 | 1 | 1 |
| 40 | 3:6:1:1:0 | 3 | 6 | 1 | 1 | 0 | 1 | 1 |
| 41 | 3:6:2:0:0 | 3 | 6 | 2 | 0 | 0 | 1 | |
| 42 | 3:6:2:0:1 | 3 | 6 | 2 | 0 | 1 | 1 | |
| 43 | 3:7:0:0:0 | 3 | 7 | 0 | 0 | 0 | 1 | |
| 44 | 3:7:1:0:0 | 3 | 7 | 1 | 0 | 0 | 1 | |
| 45 | 4:3:0:0:0 | 4 | 3 | 0 | 0 | 0 | 1 | 1 |
| 46 | 4:3:0:1:0 | 4 | 3 | 0 | 1 | 0 | | 1 |

| | | | | | | | | |
|---|---|---|---|---|---|---|---|---|
| 47 | 4:3:1:0:0 | 4 | 3 | 1 | 0 | 0 | 1 | 1 |
| 48 | 4:3:1:0:1 | 4 | 3 | 1 | 0 | 1 | 1 | |
| 49 | 4:3:1:1:0 | 4 | 3 | 1 | 1 | 0 | 1 | |
| 50 | 4:3:2:0:0 | 4 | 3 | 2 | 0 | 0 | 1 | 1 |
| 51 | 4:3:2:0:1 | 4 | 3 | 2 | 0 | 1 | 1 | |
| 52 | 4:3:3:0:0 | 4 | 3 | 3 | 0 | 0 | | 1 |
| 53 | 4:4:0:0:0 | 4 | 4 | 0 | 0 | 0 | 1 | 1 |
| 54 | 4:4:0:0:1 | 4 | 4 | 0 | 0 | 1 | | 1 |
| 55 | 4:4:0:1:0 | 4 | 4 | 0 | 1 | 0 | 1 | |
| 56 | 4:4:0:2:0 | 4 | 4 | 0 | 2 | 0 | 1 | |
| 57 | 4:4:1:0:0 | 4 | 4 | 1 | 0 | 0 | 1 | 1 |
| 58 | 4:4:1:0:1 | 4 | 4 | 1 | 0 | 1 | 1 | 1 |
| 59 | 4:4:1:1:0 | 4 | 4 | 1 | 1 | 0 | 1 | 1 |
| 60 | 4:4:1:1:1 | 4 | 4 | 1 | 1 | 1 | 1 | |
| 61 | 4:4:1:2:0 | 4 | 4 | 1 | 2 | 0 | 1 | |
| 62 | 4:4:2:0:0 | 4 | 4 | 2 | 0 | 0 | 1 | 1 |
| 63 | 4:4:2:0:1 | 4 | 4 | 2 | 0 | 1 | 1 | |
| 64 | 4:4:2:0:2 | 4 | 4 | 2 | 0 | 2 | 1 | |
| 65 | 4:4:3:0:1 | 4 | 4 | 3 | 0 | 1 | 1 | |
| 66 | 4:5:0:0:0 | 4 | 5 | 0 | 0 | 0 | 1 | 1 |
| 67 | 4:5:0:0:1 | 4 | 5 | 0 | 0 | 1 | 1 | 1 |
| 68 | 4:5:0:0:2 | 4 | 5 | 0 | 0 | 2 | 1 | 1 |
| 69 | 4:5:0:1:0 | 4 | 5 | 0 | 1 | 0 | 1 | 1 |
| 70 | 4:5:0:1:1 | 4 | 5 | 0 | 1 | 1 | | 1 |
| 71 | 4:5:0:2:0 | 4 | 5 | 0 | 2 | 0 | 1 | 1 |
| 72 | 4:5:1:0:0 | 4 | 5 | 1 | 0 | 0 | 1 | 1 |
| 73 | 4:5:1:0:1 | 4 | 5 | 1 | 0 | 1 | 1 | 1 |
| 74 | 4:5:1:0:2 | 4 | 5 | 1 | 0 | 2 | 1 | 1 |
| 75 | 4:5:1:1:0 | 4 | 5 | 1 | 1 | 0 | 1 | 1 |
| 76 | 4:5:1:1:1 | 4 | 5 | 1 | 1 | 1 | 1 | 1 |
| 77 | 4:5:1:2:0 | 4 | 5 | 1 | 2 | 0 | | 1 |
| 78 | 4:5:2:0:0 | 4 | 5 | 2 | 0 | 0 | 1 | 1 |
| 79 | 4:5:2:0:1 | 4 | 5 | 2 | 0 | 1 | 1 | 1 |
| 80 | 4:5:2:1:0 | 4 | 5 | 2 | 1 | 0 | 1 | 1 |
| 81 | 4:5:3:0:0 | 4 | 5 | 3 | 0 | 0 | 1 | 1 |
| 82 | 4:5:3:0:1 | 4 | 5 | 3 | 0 | 1 | 1 | |
| 83 | 4:5:3:1:0 | 4 | 5 | 3 | 1 | 0 | | 1 |
| 84 | 4:5:3:1:1 | 4 | 5 | 3 | 1 | 1 | | 1 |
| 85 | 4:5:3:2:0 | 4 | 5 | 3 | 2 | 0 | | 1 |
| 86 | 4:5:4:0:2 | 4 | 5 | 4 | 0 | 2 | 1 | |
| 87 | 4:6:0:0:0 | 4 | 6 | 0 | 0 | 0 | 1 | 1 |
| 88 | 4:6:0:0:1 | 4 | 6 | 0 | 0 | 1 | 1 | 1 |
| 89 | 4:6:0:1:0 | 4 | 6 | 0 | 1 | 0 | 1 | 1 |
| 90 | 4:6:0:1:1 | 4 | 6 | 0 | 1 | 1 | 1 | |
| 91 | 4:6:0:2:0 | 4 | 6 | 0 | 2 | 0 | 1 | |
| 92 | 4:6:1:0:0 | 4 | 6 | 1 | 0 | 0 | 1 | 1 |
| 93 | 4:6:1:0:1 | 4 | 6 | 1 | 0 | 1 | 1 | |
| 94 | 4:6:1:1:0 | 4 | 6 | 1 | 1 | 0 | 1 | 1 |
| 95 | 4:6:2:0:0 | 4 | 6 | 2 | 0 | 0 | 1 | 1 |

| | | | | | | | | |
|---|---|---|---|---|---|---|---|---|
| 96 | 4:6:2:0:1 | 4 | 6 | 2 | 0 | 1 | 1 | |
| 97 | 4:6:2:1:0 | 4 | 6 | 2 | 1 | 0 | 1 | |
| 98 | 4:6:3:0:0 | 4 | 6 | 3 | 0 | 0 | 1 | 1 |
| 99 | 4:6:3:0:1 | 4 | 6 | 3 | 0 | 1 | 1 | |
| 100 | 4:6:3:1:1 | 4 | 6 | 3 | 1 | 1 | 1 | |
| 101 | 4:6:4:2:0 | 4 | 6 | 4 | 2 | 0 | 1 | |
| 102 | 4:7:0:0:0 | 4 | 7 | 0 | 0 | 0 | 1 | 1 |
| 103 | 4:7:0:0:1 | 4 | 7 | 0 | 0 | 1 | 1 | |
| 104 | 4:7:0:1:0 | 4 | 7 | 0 | 1 | 0 | 1 | 1 |
| 105 | 4:7:1:0:0 | 4 | 7 | 1 | 0 | 0 | 1 | 1 |
| 106 | 4:7:1:1:0 | 4 | 7 | 1 | 1 | 0 | 1 | |
| 107 | 4:7:2:0:0 | 4 | 7 | 2 | 0 | 0 | 1 | 1 |
| 108 | 4:7:2:1:0 | 4 | 7 | 2 | 1 | 0 | 1 | |
| 109 | 4:7:2:2:0 | 4 | 7 | 2 | 2 | 0 | 1 | |
| 110 | 4:7:4:2:0 | 4 | 7 | 4 | 2 | 0 | 1 | |
| 111 | 4:8:0:0:0 | 4 | 8 | 0 | 0 | 0 | 1 | |
| 112 | 4:8:1:0:0 | 4 | 8 | 1 | 0 | 0 | 1 | |
| 113 | 5:3:0:0:0 | 5 | 3 | 0 | 0 | 0 | 1 | 1 |
| 114 | 5:3:1:0:0 | 5 | 3 | 1 | 0 | 0 | 1 | 1 |
| 115 | 5:3:1:0:1 | 5 | 3 | 1 | 0 | 1 | 1 | |
| 116 | 5:3:1:1:0 | 5 | 3 | 1 | 1 | 0 | 1 | 1 |
| 117 | 5:3:2:0:0 | 5 | 3 | 2 | 0 | 0 | 1 | |
| 118 | 5:3:2:0:1 | 5 | 3 | 2 | 0 | 1 | 1 | |
| 119 | 5:3:3:0:1 | 5 | 3 | 3 | 0 | 1 | 1 | |
| 120 | 5:4:0:0:0 | 5 | 4 | 0 | 0 | 0 | 1 | 1 |
| 121 | 5:4:0:0:1 | 5 | 4 | 0 | 0 | 1 | 1 | 1 |
| 122 | 5:4:0:0:2 | 5 | 4 | 0 | 0 | 2 | | 1 |
| 123 | 5:4:0:1:0 | 5 | 4 | 0 | 1 | 0 | 1 | 1 |
| 124 | 5:4:0:1:1 | 5 | 4 | 0 | 1 | 1 | | 1 |
| 125 | 5:4:0:2:0 | 5 | 4 | 0 | 2 | 0 | 1 | |
| 126 | 5:4:1:0:0 | 5 | 4 | 1 | 0 | 0 | 1 | 1 |
| 127 | 5:4:1:0:1 | 5 | 4 | 1 | 0 | 1 | 1 | 1 |
| 128 | 5:4:1:0:2 | 5 | 4 | 1 | 0 | 2 | 1 | |
| 129 | 5:4:1:1:0 | 5 | 4 | 1 | 1 | 0 | 1 | |
| 130 | 5:4:1:1:1 | 5 | 4 | 1 | 1 | 1 | | 1 |
| 131 | 5:4:1:2:0 | 5 | 4 | 1 | 2 | 0 | | 1 |
| 132 | 5:4:2:0:0 | 5 | 4 | 2 | 0 | 0 | 1 | 1 |
| 133 | 5:4:2:0:1 | 5 | 4 | 2 | 0 | 1 | 1 | 1 |
| 134 | 5:4:2:0:2 | 5 | 4 | 2 | 0 | 2 | 1 | |
| 135 | 5:4:2:1:0 | 5 | 4 | 2 | 1 | 0 | 1 | 1 |
| 136 | 5:4:3:0:0 | 5 | 4 | 3 | 0 | 0 | 1 | |
| 137 | 5:4:3:0:1 | 5 | 4 | 3 | 0 | 1 | 1 | |
| 138 | 5:5:0:0:0 | 5 | 5 | 0 | 0 | 0 | 1 | 1 |
| 139 | 5:5:0:0:1 | 5 | 5 | 0 | 0 | 1 | | 1 |
| 140 | 5:5:0:0:2 | 5 | 5 | 0 | 0 | 2 | | 1 |
| 141 | 5:5:0:1:0 | 5 | 5 | 0 | 1 | 0 | 1 | 1 |
| 142 | 5:5:0:1:1 | 5 | 5 | 0 | 1 | 1 | 1 | 1 |
| 143 | 5:5:1:0:0 | 5 | 5 | 1 | 0 | 0 | 1 | 1 |
| 144 | 5:5:1:0:1 | 5 | 5 | 1 | 0 | 1 | 1 | 1 |

| | | | | | | | | |
|---|---|---|---|---|---|---|---|---|
| 145 | 5:5:1:0:2 | 5 | 5 | 1 | 0 | 2 | 1 | 1 |
| 146 | 5:5:1:1:0 | 5 | 5 | 1 | 1 | 0 | 1 | 1 |
| 147 | 5:5:1:1:1 | 5 | 5 | 1 | 1 | 1 | 1 | |
| 148 | 5:5:2:0:0 | 5 | 5 | 2 | 0 | 0 | 1 | 1 |
| 149 | 5:5:2:0:1 | 5 | 5 | 2 | 0 | 1 | 1 | |
| 150 | 5:5:2:0:2 | 5 | 5 | 2 | 0 | 2 | 1 | |
| 151 | 5:5:2:1:0 | 5 | 5 | 2 | 1 | 0 | 1 | |
| 152 | 5:5:3:0:0 | 5 | 5 | 3 | 0 | 0 | 1 | 1 |
| 153 | 5:5:3:0:1 | 5 | 5 | 3 | 0 | 1 | 1 | |
| 154 | 5:5:4:0:0 | 5 | 5 | 4 | 0 | 0 | 1 | |
| 155 | 5:5:4:0:1 | 5 | 5 | 4 | 0 | 1 | 1 | |
| 156 | 5:6:0:0:0 | 5 | 6 | 0 | 0 | 0 | 1 | 1 |
| 157 | 5:6:0:0:1 | 5 | 6 | 0 | 0 | 1 | 1 | 1 |
| 158 | 5:6:0:0:2 | 5 | 6 | 0 | 0 | 2 | 1 | 1 |
| 159 | 5:6:0:1:0 | 5 | 6 | 0 | 1 | 0 | 1 | 1 |
| 160 | 5:6:0:1:1 | 5 | 6 | 0 | 1 | 1 | 1 | 1 |
| 161 | 5:6:0:1:2 | 5 | 6 | 0 | 1 | 2 | 1 | 1 |
| 162 | 5:6:0:2:0 | 5 | 6 | 0 | 2 | 0 | 1 | |
| 163 | 5:6:0:2:1 | 5 | 6 | 0 | 2 | 1 | | 1 |
| 164 | 5:6:0:3:0 | 5 | 6 | 0 | 3 | 0 | | 1 |
| 165 | 5:6:1:0:0 | 5 | 6 | 1 | 0 | 0 | 1 | 1 |
| 166 | 5:6:1:0:1 | 5 | 6 | 1 | 0 | 1 | 1 | 1 |
| 167 | 5:6:1:0:2 | 5 | 6 | 1 | 0 | 2 | 1 | 1 |
| 168 | 5:6:1:0:3 | 5 | 6 | 1 | 0 | 3 | 1 | |
| 169 | 5:6:1:1:0 | 5 | 6 | 1 | 1 | 0 | 1 | 1 |
| 170 | 5:6:1:1:1 | 5 | 6 | 1 | 1 | 1 | 1 | 1 |
| 171 | 5:6:1:2:1 | 5 | 6 | 1 | 2 | 1 | | 1 |
| 172 | 5:6:2:0:0 | 5 | 6 | 2 | 0 | 0 | 1 | 1 |
| 173 | 5:6:2:0:1 | 5 | 6 | 2 | 0 | 1 | 1 | 1 |
| 174 | 5:6:2:0:2 | 5 | 6 | 2 | 0 | 2 | 1 | |
| 175 | 5:6:2:1:0 | 5 | 6 | 2 | 1 | 0 | 1 | 1 |
| 176 | 5:6:3:0:0 | 5 | 6 | 3 | 0 | 0 | 1 | 1 |
| 177 | 5:6:3:0:1 | 5 | 6 | 3 | 0 | 1 | 1 | 1 |
| 178 | 5:6:3:1:0 | 5 | 6 | 3 | 1 | 0 | 1 | |
| 179 | 5:6:4:0:0 | 5 | 6 | 4 | 0 | 0 | 1 | |
| 180 | 5:6:4:0:1 | 5 | 6 | 4 | 0 | 1 | 1 | |
| 181 | 5:7:0:0:0 | 5 | 7 | 0 | 0 | 0 | 1 | 1 |
| 182 | 5:7:0:1:0 | 5 | 7 | 0 | 1 | 0 | 1 | |
| 183 | 5:7:0:1:1 | 5 | 7 | 0 | 1 | 1 | 1 | |
| 184 | 5:7:0:1:2 | 5 | 7 | 0 | 1 | 2 | 1 | |
| 185 | 5:7:0:2:0 | 5 | 7 | 0 | 2 | 0 | 1 | |
| 186 | 5:7:1:0:0 | 5 | 7 | 1 | 0 | 0 | 1 | 1 |
| 187 | 5:7:1:0:1 | 5 | 7 | 1 | 0 | 1 | 1 | |
| 188 | 5:7:1:1:0 | 5 | 7 | 1 | 1 | 0 | 1 | 1 |
| 189 | 5:7:1:1:1 | 5 | 7 | 1 | 1 | 1 | 1 | 1 |
| 190 | 5:7:1:2:0 | 5 | 7 | 1 | 2 | 0 | | 1 |
| 191 | 5:7:2:0:0 | 5 | 7 | 2 | 0 | 0 | 1 | |
| 192 | 5:7:2:0:2 | 5 | 7 | 2 | 0 | 2 | 1 | |
| 193 | 5:7:2:1:0 | 5 | 7 | 2 | 1 | 0 | 1 | |

| | | | | | | | | |
|---|---|---|---|---|---|---|---|---|
| 194 | 5:7:3:0:0 | 5 | 7 | 3 | 0 | 0 | 1 | |
| 195 | 5:7:3:1:0 | 5 | 7 | 3 | 1 | 0 | 1 | |
| 196 | 5:8:0:1:0 | 5 | 8 | 0 | 1 | 0 | 1 | |
| 197 | 5:8:0:2:0 | 5 | 8 | 0 | 2 | 0 | 1 | |
| 198 | 5:8:1:0:0 | 5 | 8 | 1 | 0 | 0 | | 1 |
| 199 | 5:8:1:1:1 | 5 | 8 | 1 | 1 | 1 | 1 | |
| 200 | 5:8:2:0:2 | 5 | 8 | 2 | 0 | 2 | 1 | |
| 201 | 5:9:0:2:0 | 5 | 9 | 0 | 2 | 0 | 1 | |
| 202 | 5:9:1:1:1 | 5 | 9 | 1 | 1 | 1 | 1 | |
| 203 | 5:9:2:0:2 | 5 | 9 | 2 | 0 | 2 | 1 | |
| 204 | 6:3:0:0:0 | 6 | 3 | 0 | 0 | 0 | 1 | 1 |
| 205 | 6:3:1:0:0 | 6 | 3 | 1 | 0 | 0 | 1 | 1 |
| 206 | 6:3:1:0:1 | 6 | 3 | 1 | 0 | 1 | 1 | 1 |
| 207 | 6:3:1:1:0 | 6 | 3 | 1 | 1 | 0 | | 1 |
| 208 | 6:3:1:2:0 | 6 | 3 | 1 | 2 | 0 | | 1 |
| 209 | 6:3:2:0:0 | 6 | 3 | 2 | 0 | 0 | 1 | 1 |
| 210 | 6:3:2:0:1 | 6 | 3 | 2 | 0 | 1 | 1 | 1 |
| 211 | 6:3:3:0:0 | 6 | 3 | 3 | 0 | 0 | 1 | 1 |
| 212 | 6:4:0:0:0 | 6 | 4 | 0 | 0 | 0 | 1 | 1 |
| 213 | 6:4:0:1:0 | 6 | 4 | 0 | 1 | 0 | 1 | |
| 214 | 6:4:1:0:0 | 6 | 4 | 1 | 0 | 0 | 1 | 1 |
| 215 | 6:4:1:1:0 | 6 | 4 | 1 | 1 | 0 | 1 | |
| 216 | 6:4:2:0:0 | 6 | 4 | 2 | 0 | 0 | 1 | 1 |
| 217 | 6:4:3:0:0 | 6 | 4 | 3 | 0 | 0 | 1 | |
| 218 | 6:5:0:0:0 | 6 | 5 | 0 | 0 | 0 | 1 | 1 |
| 219 | 6:5:1:0:0 | 6 | 5 | 1 | 0 | 0 | 1 | 1 |
| 220 | 6:5:1:0:1 | 6 | 5 | 1 | 0 | 1 | 1 | |
| 221 | 6:5:1:1:0 | 6 | 5 | 1 | 1 | 0 | | 1 |
| 222 | 6:5:1:1:1 | 6 | 5 | 1 | 1 | 1 | | 1 |
| 223 | 6:5:1:1:2 | 6 | 5 | 1 | 1 | 2 | | 1 |
| 224 | 6:5:2:0:0 | 6 | 5 | 2 | 0 | 0 | 1 | 1 |
| 225 | 6:5:2:0:1 | 6 | 5 | 2 | 0 | 1 | 1 | |
| 226 | 6:5:2:0:2 | 6 | 5 | 2 | 0 | 2 | 1 | |
| 227 | 6:5:3:0:0 | 6 | 5 | 3 | 0 | 0 | 1 | |
| 228 | 6:5:3:0:1 | 6 | 5 | 3 | 0 | 1 | 1 | |
| 229 | 6:5:3:0:2 | 6 | 5 | 3 | 0 | 2 | 1 | |
| 230 | 6:5:4:0:0 | 6 | 5 | 4 | 0 | 0 | 1 | |
| 231 | 6:5:4:0:1 | 6 | 5 | 4 | 0 | 1 | 1 | |
| 232 | 6:6:0:0:0 | 6 | 6 | 0 | 0 | 0 | 1 | 1 |
| 233 | 6:6:0:0:1 | 6 | 6 | 0 | 0 | 1 | 1 | |
| 234 | 6:6:0:1:0 | 6 | 6 | 0 | 1 | 0 | 1 | 1 |
| 235 | 6:6:0:1:1 | 6 | 6 | 0 | 1 | 1 | | 1 |
| 236 | 6:6:0:2:0 | 6 | 6 | 0 | 2 | 0 | | 1 |
| 237 | 6:6:1:0:0 | 6 | 6 | 1 | 0 | 0 | 1 | 1 |
| 238 | 6:6:1:0:1 | 6 | 6 | 1 | 0 | 1 | 1 | |
| 239 | 6:6:1:0:2 | 6 | 6 | 1 | 0 | 2 | 1 | 1 |
| 240 | 6:6:1:1:0 | 6 | 6 | 1 | 1 | 0 | 1 | 1 |
| 241 | 6:6:1:1:1 | 6 | 6 | 1 | 1 | 1 | 1 | 1 |
| 242 | 6:6:1:1:2 | 6 | 6 | 1 | 1 | 2 | | 1 |

| | | | | | | | | |
|---|---|---|---|---|---|---|---|---|
| 243 | 6:6:1:2:0 | 6 | 6 | 1 | 2 | 0 | | 1 |
| 244 | 6:6:2:0:0 | 6 | 6 | 2 | 0 | 0 | 1 | 1 |
| 245 | 6:6:2:0:1 | 6 | 6 | 2 | 0 | 1 | 1 | |
| 246 | 6:6:2:0:2 | 6 | 6 | 2 | 0 | 2 | 1 | |
| 247 | 6:6:2:1:0 | 6 | 6 | 2 | 1 | 0 | 1 | |
| 248 | 6:6:2:1:1 | 6 | 6 | 2 | 1 | 1 | 1 | |
| 249 | 6:6:3:0:0 | 6 | 6 | 3 | 0 | 0 | 1 | 1 |
| 250 | 6:6:3:0:1 | 6 | 6 | 3 | 0 | 1 | 1 | |
| 251 | 6:6:3:0:2 | 6 | 6 | 3 | 0 | 2 | 1 | |
| 252 | 6:6:3:1:0 | 6 | 6 | 3 | 1 | 0 | 1 | |
| 253 | 6:6:4:0:0 | 6 | 6 | 4 | 0 | 0 | 1 | |
| 254 | 6:6:4:0:1 | 6 | 6 | 4 | 0 | 1 | 1 | |
| 255 | 6:6:4:0:2 | 6 | 6 | 4 | 0 | 2 | 1 | |
| 256 | 6:7:0:0:0 | 6 | 7 | 0 | 0 | 0 | 1 | 1 |
| 257 | 6:7:0:0:1 | 6 | 7 | 0 | 0 | 1 | 1 | 1 |
| 258 | 6:7:0:0:2 | 6 | 7 | 0 | 0 | 2 | 1 | 1 |
| 259 | 6:7:0:1:0 | 6 | 7 | 0 | 1 | 0 | 1 | 1 |
| 260 | 6:7:0:1:1 | 6 | 7 | 0 | 1 | 1 | 1 | 1 |
| 261 | 6:7:0:2:0 | 6 | 7 | 0 | 2 | 0 | 1 | 1 |
| 262 | 6:7:0:2:1 | 6 | 7 | 0 | 2 | 1 | | 1 |
| 263 | 6:7:0:3:1 | 6 | 7 | 0 | 3 | 1 | | 1 |
| 264 | 6:7:1:0:0 | 6 | 7 | 1 | 0 | 0 | 1 | 1 |
| 265 | 6:7:1:0:1 | 6 | 7 | 1 | 0 | 1 | 1 | 1 |
| 266 | 6:7:1:0:2 | 6 | 7 | 1 | 0 | 2 | 1 | |
| 267 | 6:7:1:0:3 | 6 | 7 | 1 | 0 | 3 | 1 | |
| 268 | 6:7:1:1:0 | 6 | 7 | 1 | 1 | 0 | 1 | 1 |
| 269 | 6:7:1:1:1 | 6 | 7 | 1 | 1 | 1 | 1 | |
| 270 | 6:7:1:2:0 | 6 | 7 | 1 | 2 | 0 | 1 | 1 |
| 271 | 6:7:1:3:1 | 6 | 7 | 1 | 3 | 1 | | 1 |
| 272 | 6:7:2:0:0 | 6 | 7 | 2 | 0 | 0 | 1 | |
| 273 | 6:7:2:0:1 | 6 | 7 | 2 | 0 | 1 | 1 | |
| 274 | 6:7:2:0:2 | 6 | 7 | 2 | 0 | 2 | 1 | |
| 275 | 6:7:2:0:3 | 6 | 7 | 2 | 0 | 3 | 1 | |
| 276 | 6:7:2:1:0 | 6 | 7 | 2 | 1 | 0 | 1 | |
| 277 | 6:7:2:1:1 | 6 | 7 | 2 | 1 | 1 | 1 | |
| 278 | 6:7:3:0:0 | 6 | 7 | 3 | 0 | 0 | 1 | |
| 279 | 6:7:3:0:1 | 6 | 7 | 3 | 0 | 1 | 1 | |
| 280 | 6:7:3:0:2 | 6 | 7 | 3 | 0 | 2 | 1 | |
| 281 | 6:7:3:1:0 | 6 | 7 | 3 | 1 | 0 | 1 | |
| 282 | 6:7:3:1:1 | 6 | 7 | 3 | 1 | 1 | 1 | |
| 283 | 6:7:4:0:0 | 6 | 7 | 4 | 0 | 0 | 1 | 1 |
| 284 | 6:7:4:0:1 | 6 | 7 | 4 | 0 | 1 | 1 | 1 |
| 285 | 6:7:4:1:0 | 6 | 7 | 4 | 1 | 0 | 1 | 1 |
| 286 | 6:7:5:0:0 | 6 | 7 | 5 | 0 | 0 | | 1 |
| 287 | 6:8:0:0:0 | 6 | 8 | 0 | 0 | 0 | 1 | |
| 288 | 6:8:0:1:0 | 6 | 8 | 0 | 1 | 0 | 1 | 1 |
| 289 | 6:8:0:1:1 | 6 | 8 | 0 | 1 | 1 | 1 | |
| 290 | 6:8:0:1:2 | 6 | 8 | 0 | 1 | 2 | 1 | |
| 291 | 6:8:0:2:0 | 6 | 8 | 0 | 2 | 0 | 1 | |

| | | | | | | | | |
|---|---|---|---|---|---|---|---|---|
| 292 | 6:8:1:1:0 | 6 | 8 | 1 | 1 | 0 | 1 | 1 |
| 293 | 6:8:1:1:1 | 6 | 8 | 1 | 1 | 1 | 1 | |
| 294 | 6:8:1:1:2 | 6 | 8 | 1 | 1 | 2 | 1 | |
| 295 | 6:8:1:2:0 | 6 | 8 | 1 | 2 | 0 | 1 | |
| 296 | 6:8:2:0:2 | 6 | 8 | 2 | 0 | 2 | 1 | |
| 297 | 6:8:2:1:0 | 6 | 8 | 2 | 1 | 0 | 1 | |
| 298 | 6:8:2:1:1 | 6 | 8 | 2 | 1 | 1 | 1 | |
| 299 | 6:8:2:2:0 | 6 | 8 | 2 | 2 | 0 | 1 | |
| 300 | 6:8:3:0:2 | 6 | 8 | 3 | 0 | 2 | 1 | |
| 301 | 6:8:3:1:0 | 6 | 8 | 3 | 1 | 0 | 1 | |
| 302 | 6:8:4:1:0 | 6 | 8 | 4 | 1 | 0 | 1 | |
| 303 | 6:9:0:0:0 | 6 | 9 | 0 | 0 | 0 | | 1 |
| 304 | 6:9:0:2:0 | 6 | 9 | 0 | 2 | 0 | 1 | |
| 305 | 6:9:0:2:1 | 6 | 9 | 0 | 2 | 1 | 1 | |
| 306 | 6:9:1:1:1 | 6 | 9 | 1 | 1 | 1 | 1 | 1 |
| 307 | 6:9:1:2:0 | 6 | 9 | 1 | 2 | 0 | 1 | 1 |
| 308 | 6:9:2:1:1 | 6 | 9 | 2 | 1 | 1 | 1 | |
| 309 | 6:10:0:2:0 | 6 | 10 | 0 | 2 | 0 | 1 | |
| 310 | 6:10:1:1:0 | 6 | 10 | 1 | 1 | 0 | | 1 |
| 311 | 6:10:1:2:0 | 6 | 10 | 1 | 2 | 0 | 1 | |
| 312 | 7:3:0:0:0 | 7 | 3 | 0 | 0 | 0 | | 1 |
| 313 | 7:3:1:0:0 | 7 | 3 | 1 | 0 | 0 | 1 | 1 |
| 314 | 7:4:1:0:0 | 7 | 4 | 1 | 0 | 0 | 1 | |
| 315 | 7:4:2:0:0 | 7 | 4 | 2 | 0 | 0 | 1 | 1 |
| 316 | 7:4:2:0:1 | 7 | 4 | 2 | 0 | 1 | 1 | |
| 317 | 7:4:3:0:0 | 7 | 4 | 3 | 0 | 0 | 1 | |
| 318 | 7:4:3:0:1 | 7 | 4 | 3 | 0 | 1 | 1 | |
| 319 | 7:4:4:0:0 | 7 | 4 | 4 | 0 | 0 | 1 | |
| 320 | 7:5:1:0:0 | 7 | 5 | 1 | 0 | 0 | 1 | |
| 321 | 7:5:1:1:0 | 7 | 5 | 1 | 1 | 0 | 1 | |
| 322 | 7:5:2:0:2 | 7 | 5 | 2 | 0 | 2 | 1 | |
| 323 | 7:5:2:1:0 | 7 | 5 | 2 | 1 | 0 | 1 | |
| 324 | 7:5:3:0:0 | 7 | 5 | 3 | 0 | 0 | 1 | |
| 325 | 7:5:3:0:1 | 7 | 5 | 3 | 0 | 1 | 1 | |
| 326 | 7:5:3:0:2 | 7 | 5 | 3 | 0 | 2 | 1 | |
| 327 | 7:5:4:0:0 | 7 | 5 | 4 | 0 | 0 | 1 | |
| 328 | 7:5:4:0:1 | 7 | 5 | 4 | 0 | 1 | 1 | |
| 329 | 7:5:4:0:2 | 7 | 5 | 4 | 0 | 2 | 1 | |
| 330 | 7:6:0:0:0 | 7 | 6 | 0 | 0 | 0 | 1 | 1 |
| 331 | 7:6:1:0:0 | 7 | 6 | 1 | 0 | 0 | 1 | 1 |
| 332 | 7:6:1:0:1 | 7 | 6 | 1 | 0 | 1 | 1 | |
| 333 | 7:6:1:1:1 | 7 | 6 | 1 | 1 | 1 | 1 | |
| 334 | 7:6:1:3:1 | 7 | 6 | 1 | 3 | 1 | | 1 |
| 335 | 7:6:2:0:0 | 7 | 6 | 2 | 0 | 0 | 1 | |
| 336 | 7:6:2:0:2 | 7 | 6 | 2 | 0 | 2 | 1 | |
| 337 | 7:6:2:1:0 | 7 | 6 | 2 | 1 | 0 | 1 | |
| 338 | 7:6:2:1:1 | 7 | 6 | 2 | 1 | 1 | 1 | |
| 339 | 7:6:3:0:0 | 7 | 6 | 3 | 0 | 0 | 1 | |
| 340 | 7:6:3:0:1 | 7 | 6 | 3 | 0 | 1 | 1 | |

| | | | | | | | | |
|---|---|---|---|---|---|---|---|---|
| 341 | 7:6:3:0:2 | 7 | 6 | 3 | 0 | 2 | 1 | |
| 342 | 7:6:3:1:0 | 7 | 6 | 3 | 1 | 0 | 1 | |
| 343 | 7:6:3:1:1 | 7 | 6 | 3 | 1 | 1 | 1 | |
| 344 | 7:6:4:0:0 | 7 | 6 | 4 | 0 | 0 | 1 | |
| 345 | 7:6:4:0:1 | 7 | 6 | 4 | 0 | 1 | 1 | |
| 346 | 7:6:4:0:2 | 7 | 6 | 4 | 0 | 2 | 1 | |
| 347 | 7:7:0:0:0 | 7 | 7 | 0 | 0 | 0 | 1 | 1 |
| 348 | 7:7:0:1:0 | 7 | 7 | 0 | 1 | 0 | 1 | |
| 349 | 7:7:0:2:0 | 7 | 7 | 0 | 2 | 0 | 1 | |
| 350 | 7:7:1:0:0 | 7 | 7 | 1 | 0 | 0 | 1 | 1 |
| 351 | 7:7:1:0:1 | 7 | 7 | 1 | 0 | 1 | 1 | |
| 352 | 7:7:1:1:1 | 7 | 7 | 1 | 1 | 1 | 1 | |
| 353 | 7:7:1:1:2 | 7 | 7 | 1 | 1 | 2 | | 1 |
| 354 | 7:7:1:2:0 | 7 | 7 | 1 | 2 | 0 | 1 | |
| 355 | 7:7:1:4:0 | 7 | 7 | 1 | 4 | 0 | | 1 |
| 356 | 7:7:2:0:0 | 7 | 7 | 2 | 0 | 0 | 1 | |
| 357 | 7:7:2:0:1 | 7 | 7 | 2 | 0 | 1 | 1 | |
| 358 | 7:7:2:0:2 | 7 | 7 | 2 | 0 | 2 | 1 | |
| 359 | 7:7:2:1:0 | 7 | 7 | 2 | 1 | 0 | 1 | |
| 360 | 7:7:2:1:1 | 7 | 7 | 2 | 1 | 1 | 1 | |
| 361 | 7:7:2:2:0 | 7 | 7 | 2 | 2 | 0 | 1 | |
| 362 | 7:7:3:0:0 | 7 | 7 | 3 | 0 | 0 | 1 | |
| 363 | 7:7:3:0:1 | 7 | 7 | 3 | 0 | 1 | 1 | |
| 364 | 7:7:3:1:0 | 7 | 7 | 3 | 1 | 0 | 1 | |
| 365 | 7:7:3:1:1 | 7 | 7 | 3 | 1 | 1 | 1 | |
| 366 | 7:7:4:0:0 | 7 | 7 | 4 | 0 | 0 | 1 | |
| 367 | 7:7:4:0:1 | 7 | 7 | 4 | 0 | 1 | 1 | |
| 368 | 7:7:4:1:0 | 7 | 7 | 4 | 1 | 0 | 1 | |
| 369 | 7:8:0:0:0 | 7 | 8 | 0 | 0 | 0 | 1 | |
| 370 | 7:8:0:0:1 | 7 | 8 | 0 | 0 | 1 | 1 | |
| 371 | 7:8:0:1:0 | 7 | 8 | 0 | 1 | 0 | 1 | |
| 372 | 7:8:0:2:0 | 7 | 8 | 0 | 2 | 0 | 1 | |
| 373 | 7:8:1:0:0 | 7 | 8 | 1 | 0 | 0 | 1 | 1 |
| 374 | 7:8:1:0:1 | 7 | 8 | 1 | 0 | 1 | 1 | |
| 375 | 7:8:1:1:0 | 7 | 8 | 1 | 1 | 0 | 1 | 1 |
| 376 | 7:8:1:1:1 | 7 | 8 | 1 | 1 | 1 | 1 | 1 |
| 377 | 7:8:1:1:2 | 7 | 8 | 1 | 1 | 2 | | 1 |
| 378 | 7:8:1:2:0 | 7 | 8 | 1 | 2 | 0 | 1 | 1 |
| 379 | 7:8:1:2:1 | 7 | 8 | 1 | 2 | 1 | | 1 |
| 380 | 7:8:2:0:0 | 7 | 8 | 2 | 0 | 0 | 1 | |
| 381 | 7:8:2:1:0 | 7 | 8 | 2 | 1 | 0 | 1 | |
| 382 | 7:8:2:2:0 | 7 | 8 | 2 | 2 | 0 | 1 | |
| 383 | 7:8:3:0:0 | 7 | 8 | 3 | 0 | 0 | 1 | |
| 384 | 7:8:3:1:0 | 7 | 8 | 3 | 1 | 0 | 1 | |
| 385 | 7:9:0:1:0 | 7 | 9 | 0 | 1 | 0 | 1 | |
| 386 | 7:9:0:2:0 | 7 | 9 | 0 | 2 | 0 | 1 | |
| 387 | 8:3:0:0:0 | 8 | 3 | 0 | 0 | 0 | | 1 |
| 388 | 8:3:1:0:0 | 8 | 3 | 1 | 0 | 0 | 1 | |
| 389 | 8:3:1:1:0 | 8 | 3 | 1 | 1 | 0 | 1 | |

| | | | | | | | | |
|---|---|---|---|---|---|---|---|---|
| 390 | 8:4:2:0:2 | 8 | 4 | 2 | 0 | 2 | 1 | |
| 391 | 8:4:3:0:2 | 8 | 4 | 3 | 0 | 2 | 1 | |
| 392 | 8:5:0:0:0 | 8 | 5 | 0 | 0 | 0 | | 1 |
| 393 | 8:5:1:0:0 | 8 | 5 | 1 | 0 | 0 | 1 | |
| 394 | 8:5:1:0:1 | 8 | 5 | 1 | 0 | 1 | 1 | |
| 395 | 8:5:1:1:1 | 8 | 5 | 1 | 1 | 1 | 1 | |
| 396 | 8:5:2:0:0 | 8 | 5 | 2 | 0 | 0 | 1 | |
| 397 | 8:5:2:0:2 | 8 | 5 | 2 | 0 | 2 | 1 | |
| 398 | 8:5:2:1:1 | 8 | 5 | 2 | 1 | 1 | 1 | |
| 399 | 8:5:3:0:0 | 8 | 5 | 3 | 0 | 0 | 1 | |
| 400 | 8:5:3:0:2 | 8 | 5 | 3 | 0 | 2 | 1 | |
| 401 | 8:5:4:0:0 | 8 | 5 | 4 | 0 | 0 | 1 | |
| 402 | 8:6:0:0:0 | 8 | 6 | 0 | 0 | 0 | | 1 |
| 403 | 8:6:0:1:0 | 8 | 6 | 0 | 1 | 0 | 1 | |
| 404 | 8:6:0:2:0 | 8 | 6 | 0 | 2 | 0 | 1 | |
| 405 | 8:6:1:1:1 | 8 | 6 | 1 | 1 | 1 | 1 | |
| 406 | 8:6:1:2:0 | 8 | 6 | 1 | 2 | 0 | 1 | |
| 407 | 8:6:2:0:0 | 8 | 6 | 2 | 0 | 0 | 1 | |
| 408 | 8:6:2:1:1 | 8 | 6 | 2 | 1 | 1 | 1 | |
| 409 | 8:6:3:0:0 | 8 | 6 | 3 | 0 | 0 | 1 | |
| 410 | 8:6:3:0:2 | 8 | 6 | 3 | 0 | 2 | 1 | |
| 411 | 8:6:4:0:0 | 8 | 6 | 4 | 0 | 0 | 1 | |
| 412 | 8:7:0:0:0 | 8 | 7 | 0 | 0 | 0 | | 1 |
| 413 | 8:7:0:2:0 | 8 | 7 | 0 | 2 | 0 | 1 | |
| 414 | 8:7:1:2:0 | 8 | 7 | 1 | 2 | 0 | 1 | |
| 415 | 8:7:2:1:1 | 8 | 7 | 2 | 1 | 1 | 1 | |
| 416 | 8:7:3:0:2 | 8 | 7 | 3 | 0 | 2 | 1 | |
| 417 | 8:8:0:0:0 | 8 | 8 | 0 | 0 | 0 | 1 | 1 |
| 418 | 8:8:0:0:1 | 8 | 8 | 0 | 0 | 1 | 1 | |
| 419 | 8:8:1:2:0 | 8 | 8 | 1 | 2 | 0 | 1 | |
| 420 | 8:8:2:1:1 | 8 | 8 | 2 | 1 | 1 | 1 | |
| 421 | 8:9:0:0:0 | 8 | 9 | 0 | 0 | 0 | 1 | 1 |
| 422 | 8:9:0:0:1 | 8 | 9 | 0 | 0 | 1 | 1 | |
| 423 | 8:9:1:0:0 | 8 | 9 | 1 | 0 | 0 | | 1 |
| 424 | 8:9:1:1:2 | 8 | 9 | 1 | 1 | 2 | | 1 |
| 425 | 8:9:1:2:0 | 8 | 9 | 1 | 2 | 0 | 1 | |
| 426 | 8:9:1:2:1 | 8 | 9 | 1 | 2 | 1 | | 1 |
| 427 | 9:3:0:0:0 | 9 | 3 | 0 | 0 | 0 | | 1 |
| 428 | 9:3:1:0:0 | 9 | 3 | 1 | 0 | 0 | | 1 |
| 429 | 9:4:3:0:2 | 9 | 4 | 3 | 0 | 2 | 1 | |
| 430 | 9:5:2:0:1 | 9 | 5 | 2 | 0 | 1 | 1 | |
| 431 | 9:5:2:1:1 | 9 | 5 | 2 | 1 | 1 | 1 | |
| 432 | 9:5:3:0:2 | 9 | 5 | 3 | 0 | 2 | 1 | |
| 433 | 9:6:1:1:0 | 9 | 6 | 1 | 1 | 0 | 1 | |
| 434 | 9:6:1:2:0 | 9 | 6 | 1 | 2 | 0 | 1 | |
| 435 | 9:6:2:1:1 | 9 | 6 | 2 | 1 | 1 | 1 | |
| 436 | 9:6:3:0:2 | 9 | 6 | 3 | 0 | 2 | 1 | |
| 437 | 9:7:1:2:0 | 9 | 7 | 1 | 2 | 0 | 1 | |
| 438 | 9:7:2:1:1 | 9 | 7 | 2 | 1 | 1 | 1 | |

| | | | | | | | | |
|---|---|---|---|---|---|---|---|---|
| 439 | 9:7:3:0:2 | 9 | 7 | 3 | 0 | 2 | 1 | |
| 440 | 9:8:0:0:0 | 9 | 8 | 0 | 0 | 0 | 1 | |
| 441 | 9:8:0:0:1 | 9 | 8 | 0 | 0 | 1 | 1 | |
| 442 | 9:8:1:2:0 | 9 | 8 | 1 | 2 | 0 | 1 | |
| 443 | 9:8:2:1:1 | 9 | 8 | 2 | 1 | 1 | 1 | |
| 444 | 9:8:3:0:2 | 9 | 8 | 3 | 0 | 2 | 1 | |
| 445 | 9:8:4:0:2 | 9 | 8 | 4 | 0 | 2 | 1 | |
| 446 | 9:9:0:0:0 | 9 | 9 | 0 | 0 | 0 | 1 | |
| 447 | 9:9:1:0:0 | 9 | 9 | 1 | 0 | 0 | 1 | 1 |
| 448 | 9:9:1:2:0 | 9 | 9 | 1 | 2 | 0 | 1 | |
| 449 | 9:9:2:1:1 | 9 | 9 | 2 | 1 | 1 | 1 | |
| 450 | 9:9:3:1:1 | 9 | 9 | 3 | 1 | 1 | 1 | |
| 451 | 9:10:0:0:0 | 9 | 10 | 0 | 0 | 0 | 1 | 1 |
| 452 | 9:10:1:0:0 | 9 | 10 | 1 | 0 | 0 | 1 | |
| 453 | 9:10:1:2:0 | 9 | 10 | 1 | 2 | 0 | 1 | |
| 454 | 9:10:2:2:0 | 9 | 10 | 2 | 2 | 0 | 1 | |
| 455 | 10:3:0:3:0 | 10 | 3 | 0 | 3 | 0 | 1 | |
| 456 | 10:6:3:0:2 | 10 | 6 | 3 | 0 | 2 | 1 | |
| 457 | 10:7:2:1:1 | 10 | 7 | 2 | 1 | 1 | 1 | |
| 458 | 10:8:0:0:0 | 10 | 8 | 0 | 0 | 0 | 1 | |
| 459 | 10:8:1:2:0 | 10 | 8 | 1 | 2 | 0 | 1 | |
| 460 | 10:9:0:0:0 | 10 | 9 | 0 | 0 | 0 | 1 | |
| 461 | 10:9:1:0:0 | 10 | 9 | 1 | 0 | 0 | 1 | |
| 462 | 10:10:0:0:0 | 10 | 10 | 0 | 0 | 0 | 1 | |
| 463 | 10:10:1:0:0 | 10 | 10 | 1 | 0 | 0 | 1 | 1 |
| Total | | | | | | | 410 | 204 |

Supplementary Table S14. Number of glycan compositions on each site

| Site (sequon) | No. of composition / site (Total) | No. of composition / site (by Glyco-RIDGE) | No. of composition / site (by Byonic) |
|---|---|---|---|
| 17(NLT) | 54 | 46 | 21 |
| 61(NVT) | 54 | 42 | 40 |
| 74(NGT) | 237 | 227 | 64 |
| 122(NAT) | 102 | 85 | 72 |
| 149(NKS) | 132 | 122 | 32 |
| 165(NCT) | 50 | 38 | 34 |
| 234(NIT) | 46 | 27 | 38 |
| 282(NGT) | 77 | 53 | 54 |
| 331(NIT) | 56 | 49 | 26 |
| 343(NAT) | 190 | 166 | 55 |
| 603(NTS) | 36 | 25 | 30 |
| 616(NCT) | 57 | 27 | 43 |
| 657(NNS) | 55 | 20 | 43 |
| 709(NNS) | 28 | 20 | 17 |
| 717(NFT) | 18 | 8 | 18 |
| 801(NFS) | 92 | 75 | 70 |
| 1074(NFT) | 169 | 154 | 98 |
| 1098(NGT) | 157 | 145 | 91 |
| 1134(NNT) | 60 | 38 | 41 |
| 1158(NHT) | 24 | 24 | 1 |
| 1173(NAS) | 16 | 14 | 6 |
| 1194(NES) | 49 | 44 | 25 |
| Total | 1,759 | 1,449 | 919 |
